# Low additive genetic variation in a trait under selection in domesticated rice

**DOI:** 10.1101/637330

**Authors:** Nicholas G. Karavolias, Anthony J. Greenberg, Luz S. Barrero, Lyza G. Maron, Yuxin Shi, Eliana Monteverde, Miguel A. Piñeros, Susan R. McCouch

## Abstract

Quantitative traits are important targets of both natural and artificial selection. The genetic architecture of these traits and its change during the adaptive process is thus of fundamental interest. The fate of the additive effects of variants underlying a trait receives particular attention because they constitute the genetic variation component that is transferred from parents to offspring and thus governs the response to selection. While estimation of this component of phenotypic variation is challenging, the increasing availability of dense molecular markers puts it within reach. Inbred plant species offer an additional advantage because phenotypes of genetically identical individuals can be measured in replicate. This makes it possible to estimate marker effects separately from the contribution of the genetic background. We focused on root growth in domesticated rice, *Oryza sativa*, under normal and aluminum (Al) stress conditions, a trait under recent selection because it correlates with survival under drought. A dense single nucleotide polymorphism (SNP) map is available for all accessions studied. Taking advantage of this map and a set of Bayesian models, we assessed additive marker effects. While total genetic variation accounted for a large proportion of phenotypic variance, marker effects contributed little information, particularly in the Al-tolerant *tropical japonica* population of rice. We were unable to identify any loci associated with root growth in this population. Models estimating the aggregate effects of all measured genotypes likewise produced low estimates of marker heritability and were unable to predict total genetic values accurately. Our results support the long-standing conjecture that additive genetic variation is depleted in traits under selection. We further provide evidence that this depletion is due to the prevalence of low-frequency alleles that underlie the trait.

## 1 Introduction

Unraveling the mechanisms of phenotypic evolution under selection is a fundamental task in biology. Among-individual variation in most traits of interest results from a collective action of multiple genes in conjunction with environmental effects (Lynch and Walsh, 1998). To a first approximation, only the component of the genetic variation that is due to the additive contribution of DNA sequence polymorphisms is passed on to the following generations. The fate of this additive genetic variance is thus of particular interest since it is an important factor determining both the speed and direction of adaptation.

Fisher (1930) predicted that selection should deplete this component of phenotypic variance as beneficial alleles become fixed. This intuition has been extensively explored in theoretical studies. Most analyses focus on two scenarios: stabilizing selection, where the population is close to optimum fitness and extreme phenotypes are deleterious, and directional selection, where the optimal phenotype is shifted and the population is allowed to evolve towards the new state. While the studies involve different sets of assumptions and employ either analytical or simulation approaches, the general conclusion is that the predictions are very sensitive to model parameters. In some cases (Lande, 1975; Turelli, 1984) mutation-selection balance is expected to maintain a substantial amount of additive genetic variation. In other scenarios, the variance is either reduced or preserved due to segregation of a few large-effect alleles (Turelli, 1984; Barton, 1986; Johnson and Barton, 2005; de Valdar and Barton, 2014; Jain and Stephan, 2017; Stetter *et al.*, 2018). The assumed shape of the mutation effect size distribution appears to play a major role in determining whether substantial additive genetic variation is maintained in the face of selection.

Empirical observations can help us distinguish which theoretical models, and thus genetic mechanisms, are most consistent with data. The difficulty is that we cannot directly observe the components of phenotypic variance, but must rely on statistical models to estimate them. Assuming reliable estimates, the common approach is to compare additive genetic variances in fitness-related (often called life-history) traits to a control group (e.g., seemingly unimportant morphological characteristics). Since measurement scales differ among traits, some scale factor must be used to make comparisons meaningful. A popular scaling factor is the total variance. Its use is well-motivated by theory as the ratio of additive genetic to total variance yields estimates of narrow-sense heritability, a parameter important in predicting outcomes of single-generation selection on phenotypes (Lynch and Walsh, 1998). Most empirical studies (Mousseau and Roff, 1987; Gustafsson, 1986; Kruuk *et al.*, 2000; Johnson and Barton, 2005) find that narrow-sense heritability of traits correlated with fitness is indeed low. However, two traits can have equal levels of additive genetic variance but different heritabilities due to differing amounts of environmental variance (Price and Schluter, 1991). Fisher’s prediction applies to the total amounts of additive variation, so arguably estimates of heritability are misleading. There is in fact some evidence (Price and Schluter, 1991; Houle, 1992; Johnson and Barton, 2005) that scaling genetic variances by means reveals that life-history traits, if anything, possess a larger store of additive variation than other traits.

While proper scaling of variance estimates for among-trait comparisons is important, precise estimates of variance magnitudes are required for reliable inference. Observed (natural population pedigrees) or experimental (diallel) crosses have been necessary in the past to partition phenotypic variance into the various genetic and environmental components (Sprague and Tatum, 1942; Lynch and Walsh, 1998; Greenberg *et al.*, 2010). Wide availability of dense DNA sequence markers makes it possible to dispense with crossing and instead fit models that directly incorporate nucleotide polymorphism effects (Meuwissen *et al.*, 2001; Van Raden, 2008). Plant species, as well as some model animals, offer the additional possibility of assaying genetically identical individuals in replicate. This makes it possible to separate the overall genetic contribution into that explained by markers and the rest (we refer to this quasi-residual as the “background” effect). Interpretation of this partitioning is not always clear, however, and depends on the model used for the molecular markers.

Cultivated rice *Oryza sativa* is a selfing species with extensive genomic resources, making it a useful model system to test quantitative genetic theory. It is a major food crop that is mostly grown under irrigated or lowland rain-fed conditions, where standing water and anaerobic soil conditions are at least intermittently present (Khush, 1997; Bernier *et al.*, 2008). However, a minority of area under cultivation is occupied by rice grown under upland (aerobic) rainfed conditions, characterized by absence of standing water and frequent droughts (De Datta *et al.*, 1975; Khush, 1997; Bernier *et al.*, 2008). Since initial rice domestication is associated with paddy rice (Oka, 1988; Fuller *et al.*, 2010), this arid growth regime has likely imposed a recent and strong selection for survival in the face of water scarcity (De Datta, 1975). An additional difficulty for upland rice farmers is the prevalence of acidic soils (von Uexküll and Mutert, 1995) that lead to metal toxicity (e.g., aluminum) and nutrient deficiency (e.g., phosphorus). These problems, in turn, exacerbate drought stress in plants. Thus, rice accessions adapted to upland conditions have been under additional selection pressure for tolerance to aluminum toxicity and nutrient deficiency.

Root architecture is an important determinant of drought tolerance in rice. Accessions successfully grown under rain-fed (rather than irrigated) conditions tend to have relatively long and thick roots (De Datta *et al.*, 1975; Champoux *et al.*, 1995; Bernier *et al.*, 2008). There is evidence of a genetic correlation between root length, survival, and grain yield under arid conditions (Champoux *et al.*, 1995; Bernier *et al.*, 2009; Cairns *et al.*, 2009; Lyu *et al.*, 2014). Aluminum tolerance is also assessed by comparing root length of treated and control plants (von Uexküll and Mutert, 1995; Famoso *et al.*, 2010). While rice is relatively aluminum tolerant overall, *japonica*, particularly *tropical japonica*, cultivars are particularly resistant (Famoso *et al.*, 2010, 2011). Accessions from this subpopulation are often grown on acidic, upland, rain-fed soils in Africa, Latin America, and SouthEast Asia (Khush, 1997; Lyu *et al.*, 2014). Root length, including under aluminum treatment, is thus a good candidate for a character undergoing strong directional selection. Examining its genetic architecture thus may shed light on the evolution of quantitative traits. In addition, any information gained can be used to explore the adaptive potential of rice root length in water-limited environments.

Using a world-wide panel of *O. sativa* accessions and a dense set of single-nucleotide polymorphisms (SNP), Famoso *et al.* (2011) performed a genomewide association study (GWAS) to look for associations between genotyped loci and relative root growth under aluminum treatment. While they found associations in the *aus* and *indica* subpopulations, they did not detect any signal in *tropical japonica* despite significant overall variation in root growth. To determine if this result reflects a depletion of additive genetic variance under selection, we set out to build on this work. We first recapitulated the previous analyses using a new, ten-fold denser, genotyping platform (McCouch *et al.*, 2016). We also introduced a more robust approach to modeling replicated data. Despite these improvements, we still failed to find any GWA signal in *tropical japonica*. Adding more accessions, increasing the duration of aluminum treatment, and measurement of additional traits did not produce a higher power to detect loci associated with aluminum tolerance. Furthermore, while total genetic variation accounted for a large portion of overall phenotypic variance, additive genetic variation captured by SNP genotypes was consistently low. We provide evidence that this is due to a low frequency of alleles conferring susceptibility or tolerance. These results are consistent with a set of theoretical expectations and allow us to deduce a possible genetic architecture of root growth in *tropical japonica* varieties adapted to upland cultivation. Our inferences provide testable predictions for future experiments and point to breeding strategies that are more likely to improve resistance to drought and cultivation on acid soils in this population.

## 2 Materials and Methods

### 2.1 Root measurements and accession sets

We expanded the *tropical japonica* accession set (n=92) analyzed by Famoso *et al.* (2011) by adding 77 new accessions. These were treated and measured using the same protocol as before (Famoso *et al.*, 2010, 2011). We also chose a new set of 119 *tropical japonica* accessions (including some overlap with the original 169). The Al treatment in the latter experiment was conducted as before, but the duration was extended from three to six days. The root systems of seedlings were imaged and total and longest root length measured using RootReader 2D, a digital imaging and quantification system (Clark *et al.*, 2013).

### 2.2 Genotype data

The accessions we chose have been previously genotyped using the HDRA array platform (McCouch *et al.*, 2016). We filtered the original 700,000 SNPs for each accession set, including only variants with minor allele count greater than two and with fewer than 30% of the genotype calls missing. The minor allele count of two represents a genotype of a single diploid individual, but because our accessions are highly inbred (domesticated rice is a self-pollinating species) our threshold effectively eliminates singleton SNPs. We were left with 432,607 variants in the *aus* population, 434,753 SNPs in the *tropical japonica* set used by Famoso *et al.* (2011), 467,536 for the augmented *tropical japonica* set, and 461,298 for the new set of 119 accessions. Given that the rice genome is about 380 megabases in length (Kawahara *et al.*, 2013), the average between-SNP distance in our data is roughly 0.8 kilobases. This is well within the linkage disequilibrium decay window for all rice populations (McCouch *et al.*, 2016). Since average gene size is 3 kb http://rice.plantbiology.msu.edu/, each gene contains about three SNPs. Thus moderate to high-frequency causal variants should be tagged by at least several SNPs in our data set with high probability.

### 2.3 Modeling of replicated experiments

All experiments involved measuring root traits on replicates of genetically identical individuals (homozygous accessions). The plants were grown either in control or aluminum-containing media. This experimental structure informed our modeling approach. We sought to concurrently estimate contributions of genotyped SNPs, total genotypic effects, and random environmental uctuations to overall phenotypic variance. In addition, in the analyses of the original Famoso *et al.* (2011) experiments and our augmentation of their *tropical japonica* data set, we needed to take into account the year each plant was measured, as there were a priori obvious year effects (Supplemental Figure 2).

We fitted multi-trait Bayesian hierarchical models (Gelman *et al.*, 2004; Greenberg *et al.*, 2010, 2011) to estimate parameters of interest. We treated each data set (the original Famoso *et al.* (2011) data in *aus* and *tropical japonica* populations, the augmented *tropical japonica*, and the new single-year measurements) separately. The single-year experiment measured two root traits (total and longest root length). We included them together in a multi-trait model, but analyzed the Al treated and control sets separately The other experiments measured only total root length, but we modeled the treated and control measurements as separate traits. However, because the treatments were applied to different individual plants, the error covariance between the two pseudo-traits was set to exactly zero. Genetic covariances were estimated from data in all models, since the same accessions were subjected to both treatment and control conditions.

The portions of our hierarchical models that estimate location parameters (accession means, aggregate marker effects, and year covariates) take the following form:

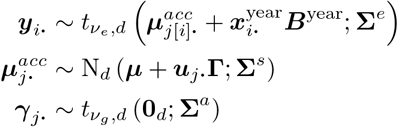

Location parameters of the form ***y***_***i***_ are row-vectors of the corresponding parameter matrices that have data points as rows and traits as columns (for example, as mentioned above, ***y***_***i***_ has two values for the Famoso *et al.* (2011) data: the treated and untreated total root length). The year effect is included via the 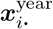 ***B***^year^ term. The correction is performed at the individual measurement level for the Famoso *et al.* (2011) data set. There is some risk of confounding between year and accession effects in the augmented data (see Supplemental File 1). Therefore, year effects were modeled at the level of accessions for that set. The third experiment was performed in one batch and therefore no year effect correction was required. The year effect coefficients (as well as the overall intercept ***μ***) were modeled with high-variance 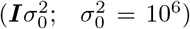 Gaussian priors. The errors were modeled using a multivariate Student-*t* distribution with three degrees of freedom to dampen the effects of outliers (Greenberg *et al.*, 2010, 2011) for the last experiment. Our set-up did not allow for Student-*t* models for block-diagonal error matrices. Therefore, the first two data sets were analyzed with Gaussian models for errors.

The marker effects enter through the eigenvectors *U* of the relationship matrix, weighted by square roots of their eigenvalues. We estimated the relationship matrix from all SNPs in each data set using the Van Raden method (Van Raden, 2008). We are using all the eigenvectors that correspond to nonzero eigenvalues. If this regression is performed using a Gaussian prior on the coefficients *γ*_*j*_, it is the Bayesian analog of the (matrix-variate) mixed effect model (Kang *et al.*, 2010; Hoffman, 2013). In this case, Σ^*a*^ is the additive marker covariance matrix. However, the first few principal components of the relationship matrix often correspond to large-scale population structure and may have relatively large regression coefficients (if the phenotypes are also stratified by subpopulation). A Gaussian prior may over-shrink these coefficients. Therefore, we used multivariate Student-*t* prior distribution on **Γ**. This is a multivariate version of the BayesA model (Meuwissen *et al.*, 2001). The results presented in the main text were obtained by setting the degrees of freedom to three (the minimal number that still results in defined moments of the Student-*t*). We repeated all analyses with the degrees of freedom set to 1000 (making the prior close to Gaussian) with the same results. This is likely because we work within populations, in the absence of deep population structure.

The covariance matrices Σ^*x*^ were modeled using Wishart distributions with weakly-informative Wishart priors with two degrees of freedom for Σ^*e*^ and Σ^*s*^ and four degrees of freedom for Σ^*a*^ (Gelman *et al.*, 2004; Greenberg *et al.*, 2011). This implies a uniform prior on narrow-sense (marker) heritability.

Detailed model descriptions are listed in the documentation provided in the Supplemental File 1. This supplemental file also includes the raw phenotypic data, scripts we used to manage the raw data, Markov chain convergence diagnostics, and post-processing and plotting of the results. The analyses were implemented in C++. The source code is included in the Supplemental File 1. The underlying library is also available from https://github.com/tonymugen/MuGen.

### 2.4 Heritability estimates

The hierarchical model allows us to estimate heritabilities. Narrow-sense heritability (Lynch and Walsh, 1998) is the fraction of total phenotypic varivariance that is explained by additive genetic effects. If we were to genotype all causal variants and accurately estimate their effects, the fraction of total variance explained by the sum of these polymorphisms’ contributions would be the narrow sense heritability. In practice, we only assay tag SNPs that may be in linkage disequilibrium with a fraction of causal variants. In addition, since linkage is likely imperfect, most linked genotyped SNPs do not capture the whole contribution of causal loci. We calculate marker heritability 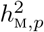 of the *p*-th trait from the total marker effects estimated in our models:

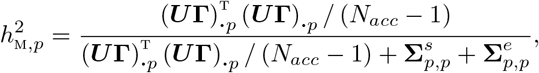

where *N*_*acc*_ is the number of accessions and the rest of the notation is as described above. We used the variance of GEBV estimates 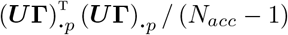 rather than values from the marker covariance matrix 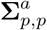 because the latter reects the additive covariance only when the prior on the principal component regression (see above) is Gaussian. We use a Student-*t* prior, thus 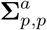 may underestimate the additive genetic variance captured by SNPs.

The 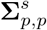 values estimate the portion of total genetic variance not explained by genotyped markers. They include the non-additive effects and additive effects not tagged by SNPs. We call these estimates “background effects” (e.g., Figure 1). The sum of the GEBV and background variance, divided by total phenotypic variance, is broad-sense heritability (Lynch and Walsh, 1998).

**Figure 1:**
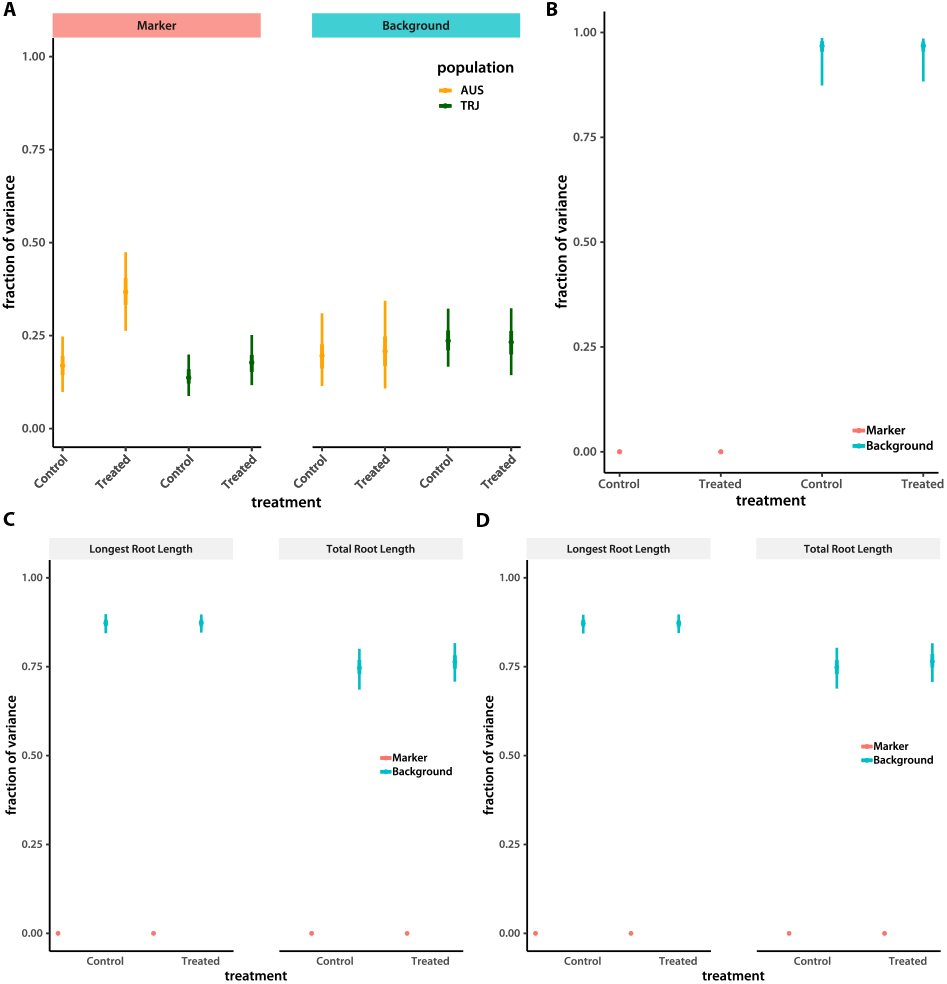
Fraction of variance explained by markers and genetic background. The plots depict regions of highest posterior density (HPD; thin lines are 95% and thick lines are 50% intervals, the points are modes) of separate estimates of variance fractions explained by markers and the rest of the genetic background. (A) Results of the re-analysis of the (Famoso *et al.*, 2011) data. (B) The larger data set of *tropical japonica* accessions. (C) Tropical *japonica* accessions subjected to new measurements and experimental conditions. (D) as in (C), but using a quadratic kernel to estimate pairwise epistatic interactions.

Our models were fitted using a Markov chain Monte Carlo approach, resulting in posterior samples of parameters. We used calculated heritabilities for each sample, yielding numerical approximations of posterior distributions of these statistics. Summaries of these distributions are plotted in the relevant figures.

### 2.5 GWA and genome prediction

Because full MCMC Bayesian model fitting is computationally demanding, we did not attempt to fit in dividual SNP effects within that framework. In addition, treated to control root length ratio (a measure of Al tolerance, Famoso *et al.*, 2010, 2011) was not modeled directly and could not be used as a response in our hierarchical models. We thus took a two-step approach, using the posterior modes of accession means as responses with mixed-model correction for population structure (Kang *et al.*, 2010) using our own R package (Greenberg, 2019), available from https://github.com/tonymugen/GWAlikeMeth. This is a maximum-likelihood method, first fitting the mixed model

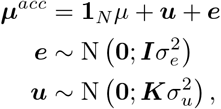

where ***y*** is the response vector of posterior accession modes, **1**_*N*_ is a column vector of ones of order, *μ* is the overall mean, *u* is the column vector of random genetic values, 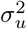 is the among-***u*** variance, ***K*** = ***XX***^*T*^=/*N*_*acc*_ is the among-genotype covariance matrix estimated from the *N*_*acc*_ × *N*_*SNP*_ genotype matrix ***X***, ***e*** is the column vector of random errors, and 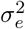 is the error variance. Residuals of this model are then used in individual SNP regressions. Each trait (including the logarithm of the treated to control root length ratio) is treated independently, although computations that are shared among traits are performed only once. We report −log_10_ *p* values as SNP effect scores.

To verify results obtained using the hierarchical Bayesian model outlined in the previous section, we also took the two-step approach to perform genome predictions. We again used modes of accession mean distributions, as described above for GWA. We started with a linear kernel ***K*** calculated as above. This is the genome best linear unbiased predictor (GBLUP) model of additive effects. To test whether modeling among-locus epistatic interactions improves prediction, we repeated the analyses with a Gaussian kernel for ***K***, setting its elements to 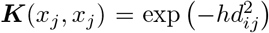. Here, *d*_*ij*_ is the Euclidian distance between genotypes *i* and *j*, *h* is the bandwidth parameter that controls the speed of decay of ***K*** elements with genetic distance. Following Crossa *et al.* (2010), we set 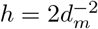, where *d*_*m*_ is the sample median among all *d*_*ij*_.

We used the R package Bayesian generalized linear regression (BGLR, Pérez and de los Campos, 2014) to fit the genome prediction models. The package implements a Gibbs sampler. We ran 55,000 iterations including 5,000 burn-in steps and 5-fold thinning.

We assessed prediction accuracy using 50 training-validation random partitions. Pearson’s correlations (*r*) between the predicted and the observed values were employed as accuracy measurements. To calculate summary statistics, we transformed raw correlation values using Fisher’s 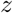 transformation 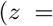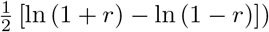. The mean and interval values presented in Table 1 were back-transformed to the original scale.

**Table 1:** Genome prediction accuracy. Mean (lower 95%, upper 95% confidence interval).

| Trait name | GBLUP | RKHS |
| --- | --- | --- |
| Longest root length, Al treated | 0.034 (-0.147, 0.214) | 0.130 (-0.052, 0.304) |
| Total root length, Al treated | 0.118 (-0.065, 0.292) | 0.102 (-0.081, 0.277) |
| Longest root length, control | 0.138 (-0.043, 0.311) | 0.025 (-0.156, 0.205) |
| Total root length, control | 0.036 (-0.146, 0.216) | -0.205 (-0.372, -0.025) |
| Longest root length, $\lg(RRG)$ | -0.223 (-0.388, -0.044) | -0.200 (-0.368, -0.020) |
| Total root length, $\lg(RRG)$ | -0.194 (-0.362, -0.014) | 0.0385 (-0.143, 0.218) |

### 2.6 Introgression mapping

We used RFMix (Maples *et al.*, 2013) to identify introgressions from *aus* and *indica* in the six admixed *tropical japonica* accessions. We used 10 training accessions from each of the three populations of interest (see the Supplemental File 2 for the full list). We ran Beagle v4.1 (Browning and Browning, 2016) with default parameters to phase genotypes, after filtering loci with minor allele frequencies below 0.08 and more than 30% missing genotypes. Our final data set included 304,488 SNPs. We then ran RFMix with the option “TrioPhased” on the phased genotypes from 36 lines: three sets of 10 training pure accessions plus the six admixed accessions. The results were plotted using an R script included in the Supplemental File 1.

### 2.7 Data and reagent availability statement

Supplemental files are available at FigShare. Supplemental File 1 contains detailed models, raw phenotypic data, scripts to manage the raw data, Markov chain convergence diagnostics, post-processing and plotting of the results, and source code. The raw phenotype data include those generated by Famoso *et al.* (2011). Genotype data are from McCouch *et al.* (2016) and available at ricediversity.org/data/index.cfm. Supplemental File 2 lists the training accessions we used for admixture mapping.

## 3 Results

The initial study (Famoso *et al.*, 2011) only examined relative root growth between Al-treated and control genotypes. To measure Al tolerance, separate individuals from the same accession are subjected to aluminum treatment and compared to plants grown under control conditions. The relative root growth is then estimated by calculating the ratio of the means among the treated and control plants within each genotype. Partitioning the phenotypic variance of this compound trait into genetic and environmental components using this experimental design is thus impossible. Therefore, we focused on estimating genetic parameters for root length itself under the two experimental regimes. This approach also fits better with results from previous studies that suggest a role for root length in water stress tolerance. The data are more sparse and ambiguous for relative root growth.

Since Famoso *et al.* (2011) did not report heritabilities for root length, we first re-analyzed their data, but using a denser SNP genotype map (HDRA, McCouch *et al.*, 2016). All accessions in the original study were genotyped using this newer platform. Of the 700,000 variants available, we retained about 450,000 SNPs after filtering out loci with singletons or more than 30% of the data missing in our accession sets (see Methods for details). We built a Bayesian hierarchical model (Gelman *et al.*, 2004, see Methods for details) that accounts for replicated measurements of accessions and the fact that experiments were performed across several years (Supplemental Fig. 2). We considered root length under control and Al treatment conditions as separate traits and used a multivariate hierarchical model (Greenberg *et al.*, 2011) that included a SNP-derived relationship matrix to account for marker effects (see Methods for model details).

Replication of identical genotypes, possible because rice is a selfing species with almost completely homozygous accessions (McCouch *et al.*, 2016), allowed us to separate additive marker-based variance from the total genetic variation and residual environmental effects. We call the fraction of total phenotypic variance explained by the sum of the additive effects of all SNPs “marker heritability.” If we estimate accurately all causal variant effects, this is narrow-sense heritability. However, most genotyped loci are likely only in linkage disequilibrium with causal variants and our estimates of even cumulative SNP effects are imperfect. Therefore, the residual genetic variance (total genetic variance, estimated from the hierarchical model as the accession effect, minus the marker variance) absorbs both additive effects of loci not tagged by our SNP panel and non-additive components. We call this portion of genetic variance the “background effect.”

We find that the heritability contributed by all markers considered together accounts for less than half of overall genetic variance in the *tropical japonica* population (Fig. 1A). This does not seem to be due to a lack of power resulting from a small sample size, since marker heritability is higher in *aus* under stress (Fig. 1A) despite a smaller number of accessions in that population (55 in *aus vs* 92 in *tropical japonica*). To see if any individual SNPs are associated with either trait, we used the posterior modes of accession effects (as well as log_10_ of the control to treated ratio, −lg *RRG*) as predictors in a standard single-trait mixed-model GWA (Kang *et al.*, 2010, see Methods for details). We were able to reproduce the GWA peak found by Famoso *et al.* (2011) for untransformed RRG in the *aus* subpopulation (Supplemental Fig. 1A), but did not detect any new associations. We did not unearth any associations in tropical *japonica* (Supplemental Fig. 1B), and no associations of SNPs with root length traits under Al treated or control conditions in either population.

Perhaps, as some theoretical results suggest, recent selection on root traits keeps adaptive alleles contributing to trait variation close to fixation. The resulting low minor allele frequency would diminish opportunities for linkage disequilibrium with our genotyped SNPs. In this case, increasing sample size should increase minor allele counts, boosting power to detect marker heritability. We therefore added more accessions to our *tropical japonica* sample, bringing the total to 169. These lines were already genotyped using HDRA (McCouch *et al.*, 2016). We administered the same treatment protocol as Famoso *et al.* (2011) and analyzed the data as before. The measurements of each accession were highly reproducible, with total genetic effects explaining almost all phenotypic variance. However, the proportion of this genetic variation accounted for by the sum of marker effects was much smaller than in the initial limited sample (Fig. 1B). Neither did we find any significant individual SNP associations (Supplemental Fig. 4A).

The estimates on a larger panel of *tropical japonica* accessions further supports the idea that the sum of all genotyped marker effects contribute little to phenotypic variation in root traits in this population. However, we wanted further confirmation that these estimates are not the result of experimental or modeling artifacts. Therefore, we selected a new panel of 119 *tropical japonica* accessions that represented the full diversity of the representatives of this population genotyped using HDRA (Supplemental Fig. 3). To increase confidence in our estimates of accession means (and thus the fraction of phenotypic variance explained by all genetic factors), we increased the number of replicates per line (eight on average). We performed all experiments at the same time, eliminating the year effect as a potential confounding variable. We measured two root traits: total and longest root length (see Methods for details) and included them together in a hierarchical Bayesian model (see Methods for details). However, we fit the model to the Al-treated and control data sets separately.

Aluminum treatment of individuals from the *aus* population increases the fraction of genetic variation explained by markers (Fig. 1A). This is not evident in *tropical japonica*. Furthermore, we see a high total genetic correlation (mode: 0.886; 95% highest posterior density (HPD) interval: [0.803, 0.945]) between control and treated root lengths in our initial experiment. Therefore, we increased the duration of aluminum treatment to put extra stress on the plants. The longer treatment did reduce the genetic relationships between treatment and control, although the correlations remained fairly high: 0.770 (0.733, 0.811) for total root length (the trait measured in the previous set of experiments) and 0.816 (0.777, 0.844) for longest root length, closer to the correlation observed in *aus* (0.597 [0.484, 0.704]).

Despite all these changes in experimental protocol and analysis, we see the same low marker heritability despite the high fraction of the phenotypic variance explained by total genetic effects (Fig. 1C). No single locus stood out when we performed GWAS on these new accession means, either (Supplemental Fig. 4B).

If our observation of extremely low marker heritability of root length in *tropical japonica* is correct, we expect that the genotyped SNPs would be poor predictors of phenotypic values. To test this, we implemented a two-step genome prediction scheme, using the posterior modes of accession means generated in the last experiment and used a separate implementation of a penalized regression on genotypes (Meuwissen *et al.*, 2001; Pérez and de los Campos, 2014) to model additive SNP effects (see Methods for details). Given the high total heritability, we are fairly confident in the robustness of accession mean estimates. We separated the data into two sets. The first set was used to train a model that used all markers to estimate model parameters. Given these training set estimates, we attempted to predict phenotypes of the remaining accessions. We repeated this process 50 times and generated empirical distributions of prediction accuracy. Consistent with the low additive marker heritability estimates presented above, genome-enabled prediction accuracy was low, with most distributions containing zero (Table 1).

All lines of evidence presented so far point to the conclusion that additive marker effects do not explain much of the total genetic variance of the root growth traits measured in *tropical japonica*. Given the high broad-sense heritability, we wondered if nonadditive SNP effects can explain our results. Given the very high homozygosity of individual lines (McCouch *et al.*, 2016), dominance is highly unlikely to play a measurable role. We are therefore left with epistatic interactions among loci as the remaining possibility. However, the parameter space of all interactions is prohibitively large, making it statistically and computationally unfeasible to estimate individual effects. However, models that rely on summaries of interactions are available. It has been shown that using the Hadamard (element-wise) product of the SNP-based relationship matrix (Van Raden, 2008) with itself in a genomic model gives approximate estimates of the sum of all pairwise epistatic interactions (Jiang and Reif, 2015). Similarly, use of a Gaussian kernel (see Methods) approximates all by all interactions (Jiang and Reif, 2015). We used the quadratic in place of the additive kernel in our hierarchical multi-trait model and the Gaussian kernel for genome prediction. We did not observe any measurable improvement in explanatory ability (Fig. 1D) or prediction accuracy (Table 1).

Since we cannot find any evidence that epistatic interactions account for much genetic variance, we are left with the possibility that low-frequency DNA sequence variants underlie our root traits. Because such alleles are not shared widely among accessions in our panel, and are not in high enough linkage disequilibrium with the SNPs we genotyped, we do not detect their effects in our models. If this conjecture is correct, we would expect many regions of the genome to make small contributions to the genetic variance. We see some evidence of this when we look at accessions that carry introgressions from the less Al tolerant *indica* and *aus* populations into the *tropical japonica* background (Fig. 2). We found six such lines in the Famoso *et al.* (2011) data set. All of them have shorter roots than the average *tropical japonica* accession in the sample, particularly under aluminum stress (Fig. 2A,B). When we looked at the genomic regions that come from *indica* or *aus* in these lines using RFMix (Maples *et al.*, 2013) we found an idiosyncratic collection of introgressed loci (Fig. 2C). This is expected if each *tropical japonica* accession carries a different set of rare causal variants.

**Figure 2:**
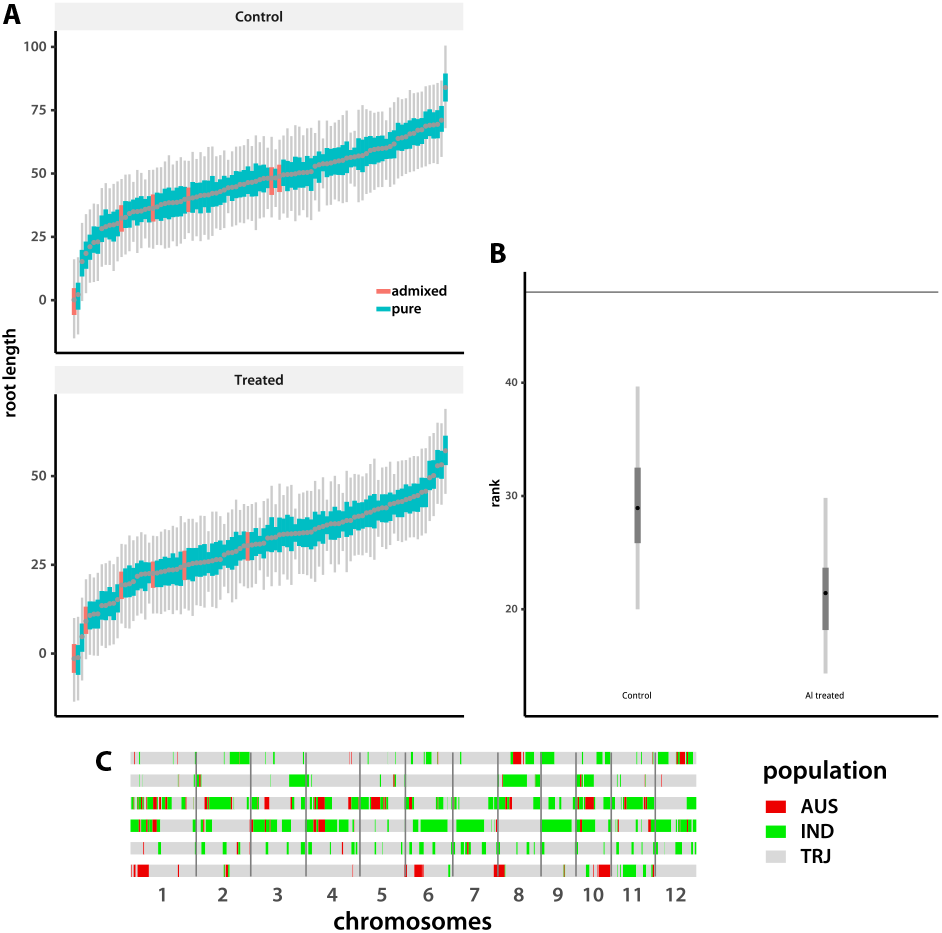
Rank of accessions with aus admixture. (A) Posterior highest-density intervals of accession means, sorted from smallest mode to largest. Tropical *japonica* accessions with *aus* admixture are marked with orange (see plot legend). (B) Posterior distributions of mean rank of admixed accessions under control and treatment conditions, with the expected value shown by the horizontal line. (C) Introgression locations in the genomes of the six admixed accessions. The six horizontal lines represent the genomes, with the accession with the shortest roots under aluminum stress on the bottom and the longest on top. Colored blocks mark the regions of introgression from *aus* (“AUS”) or *indica* (“IND”), as indicated on the legend. The *tropical japonica* (“TRJ”) background is in gray.

## 4 Discussion

Informative tests of the effects of selection on the genetic architecture of quantitative traits require partitioning of the total genetic variation into components. This can be achieved in selfing plants, *O. sativa* among them, because replicated measurements can be performed on genetically identical individuals. Relying on genotyping rather than diallel crosses saves on phenotyping costs. It also allows us to directly model additive and various non-additive effects of variants in our genotyping panel as well as loci that were not assayed but are in LD with those we assayed.

We employed this strategy to study the genetic architecture of root length in the *tropical japonica* population of rice. A significant fraction of accessions from this group are adapted to upland rain-fed conditions (Khush, 1997; Lyu *et al.*, 2014) where root length is under strong selection because it underlies drought and nutrient toxicity tolerance (De Datta, 1975; Champoux *et al.*, 1995; Bernier *et al.*, 2009; Cairns *et al.*, 2009; Lyu *et al.*, 2014). We find high broad-sense heritability using two experimental setups and three sets of accessions. However, we find that aggregate contributions of all genotyped loci, assessed using relationship matrices in and Bayesian hierarchical models, are very small in *tropical japonica*. Neither do we find any individual loci contributing significantly to phenotypic variation. This is in contrast to the situation in the *aus* population, where we see moderate marker heritability, particularly under aluminum stress, and a significant effect of the *Nrat1* locus despite a smaller sample of accessions. Alternative modeling approaches, including genome-enabled prediction, yielded the same results. Employing models that account for among-locus epistatic interactions, the only plausible source of non-additive genetic variation in this highly inbred panel, does not increase estimated marker heritability or phenotype prediction accuracy.

We are then left with the possibility that the alleles underlying these traits are at low frequency. The distribution of minor allele frequencies of the SNPs genotyped by our array is skewed towards rare alleles compared to the neutral expectation (Supplemental Fig. 5). These variants are thus unlikely to be in linkage disequilibrium with ungenotyped low-frequency causal polymorphisms and would not account for any phenotypic variance. A genetic architecture dominated by small-effect loci with low-frequency alleles is predicted by models of directional selection on quantitative traits (Jain and Stephan, 2017; Stetter *et al.*, 2018). The models envision a shift in the optimal phenotype value, followed by a change in the adaptive direction and finally stabilizing selection at the new optimum. This scenario fits with our under standing of the role of root architecture in the relatively recent adaptation of rice to cultivation under the arid upland rain-fed conditions (De Datta *et al.*, 1975; Oka, 1988; Khush, 1997; Fuller *et al.*, 2010). Genomic analyses of shifts in cultivation practices of domesticated plants may thus be a promising avenue for empirical studies of genetic architectures of traits under selection.

If our conjecture that small-effect low-frequency alleles underlie root traits in *tropical japonica* is correct, each accession should have a unique set of causal variants. Some would have a positive effect, some would be negative. The balance would then determine the overall genotypic value of the individual line. This, in turn, implies that most regions of the genome can contribute to phenotypic variance. We indeed see some evidence of this: introgressions from *indica* and *aus* populations reduce total root length of *tropical japonica* accessions, particularly under aluminum stress (Fig. 2), and the introgressions are evenly distributed throughout the genome. Conversely, Famoso *et al.* (2011) report that introgressions from *tropical japonica* into *indica* increase relative root length under aluminum stress, consistent with our hypothesis.

Although the low-frequency alleles may not individually contribute much to within-population variance, they should be detectable in biparental crosses. This is because allele frequencies of the causal variants present in these lines will be 50% and thus at high enough frequency to measure. Indeed, multiple quantitative trait locus mapping studies found loci contributing to root length under various conditions (Champoux *et al.*, 1995; Bernier *et al.*, 2009; Famoso *et al.*, 2011; Arbelaez *et al.*, 2017). However, subsequent attempts to introgress favorable alleles into a variety of genetic backgrounds have been largely unsuccessful (Bernier *et al.*, 2008). This is expected under our model, since every pair of parents should have a different set of causal loci. Furthermore, the predicted presence of at least some negative-effect alleles even in the high-value lines (Stetter *et al.*, 2018) should lead to transgressive segregation in the progeny. This is indeed often observed (Champoux *et al.*, 1995; Arbelaez *et al.*, 2017).

Our results shed some light on a fundamental question in quantitative genetics. They also should be taken into consideration when designing breeding strategies to improve productivity of rice grown in aerobic soils under water-limited conditions. The upland cultivation method has been historically low-yielding (De Datta *et al.*, 1975; De Datta, 1975; Bernier *et al.*, 2008) and has resisted efforts at genetic improvement (Bernier *et al.*, 2008). If our model is correct, progress is possible as long as the number of founding accessions is not too high. All causal alleles would be at moderate frequency and thus visible to selection. Total genetic, rather than breeding, values of the founders should be used to select them from the base population. GWAS, marker-assisted selection, and even multi-parental mapping populations (Yu *et al.*, 2008) are not expected to be effective to determine the best founders. However, genome prediction can still be used if the focus is within breeding populations. Finally, periodic addition of new founders can introduce additional beneficial alleles even if the new lines themselves are not top performers in the environments of interest.

Our study highlights the importance of detailed genome-enabled investigations of polygenic traits under selection to shed light on fundamental evolutionary questions and guide practical artificial selection decisions. We hope that it will stimulate further analyses of such traits in agriculturally important plants, taking advantage of sophisticated experimental designs and increasing availability of deep wholegenome DNA sequence data.

## Supporting information

Supplemental methods and scripts

Raw phenotype data

## 5 Acknowledgments

We thank Sandra Harrington for seed and accession maintenance, Eric Craft for technical assistance with Al treatment experiments, and Elsa de Becker for her assistance with setting up the GWAS pipeline and training NK on it. This work was supported by a grants from NSF-PGRP (award #1444511), USDANIFA (award #2013-67013-21379), Bill and Melinda Gates Foundation via AfricaRice (“Rapid Mobilization of Alleles for Rice Cultivar Improvement in SubSaharan Africa”), and salary support for LSB from Agrosavia in Colombia.

---

**Supplemental Figure 1:**
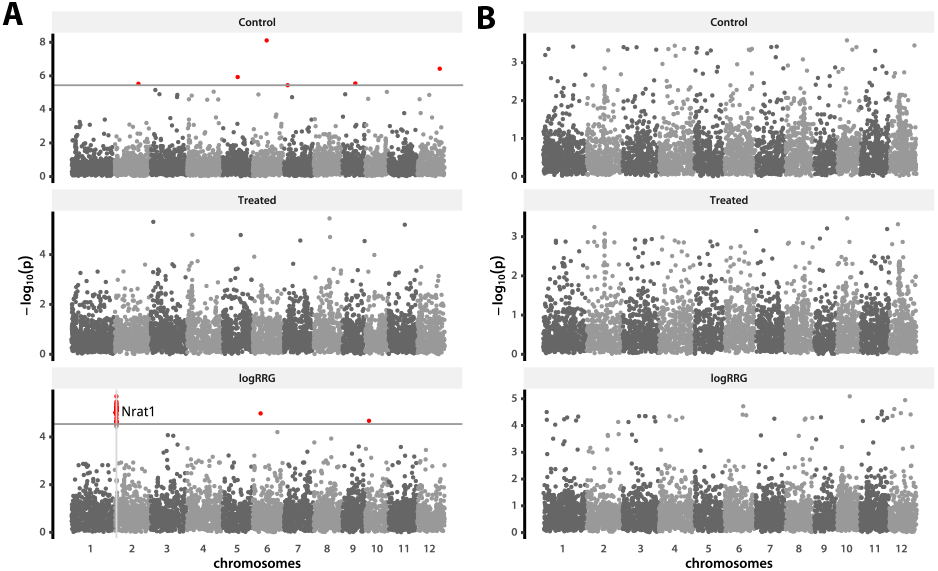
Genome-wide associations in the aus and tropical japonica populations. The Manhattan plots depict single-SNP associations (see Methods) of root length under control and Al treated conditions, as well as the logarithm of the treated to control ratio. The horizontal lines mark the −lg(*p*) values corresponding to the FDR cut-off at 0.1. (A) Associations within the *aus* population, with the Nrat1 locus marked on the lower panel; (B) associations within *tropical japonica*.

**Supplemental Figure 2:**
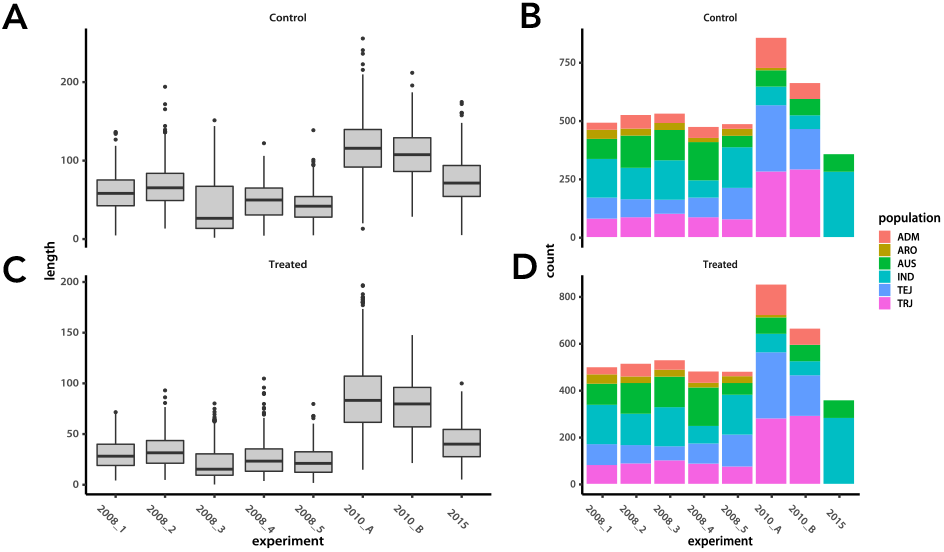
Experiment effect and accession distribution. The box plots (A, C) show distributions of raw replicate data from each experiment in Famoso *et al.* (2011) data. The bar plots (B, D) depict the total number of replicates coming from each subpopulation within each experiment.

**Supplemental Figure 3:**
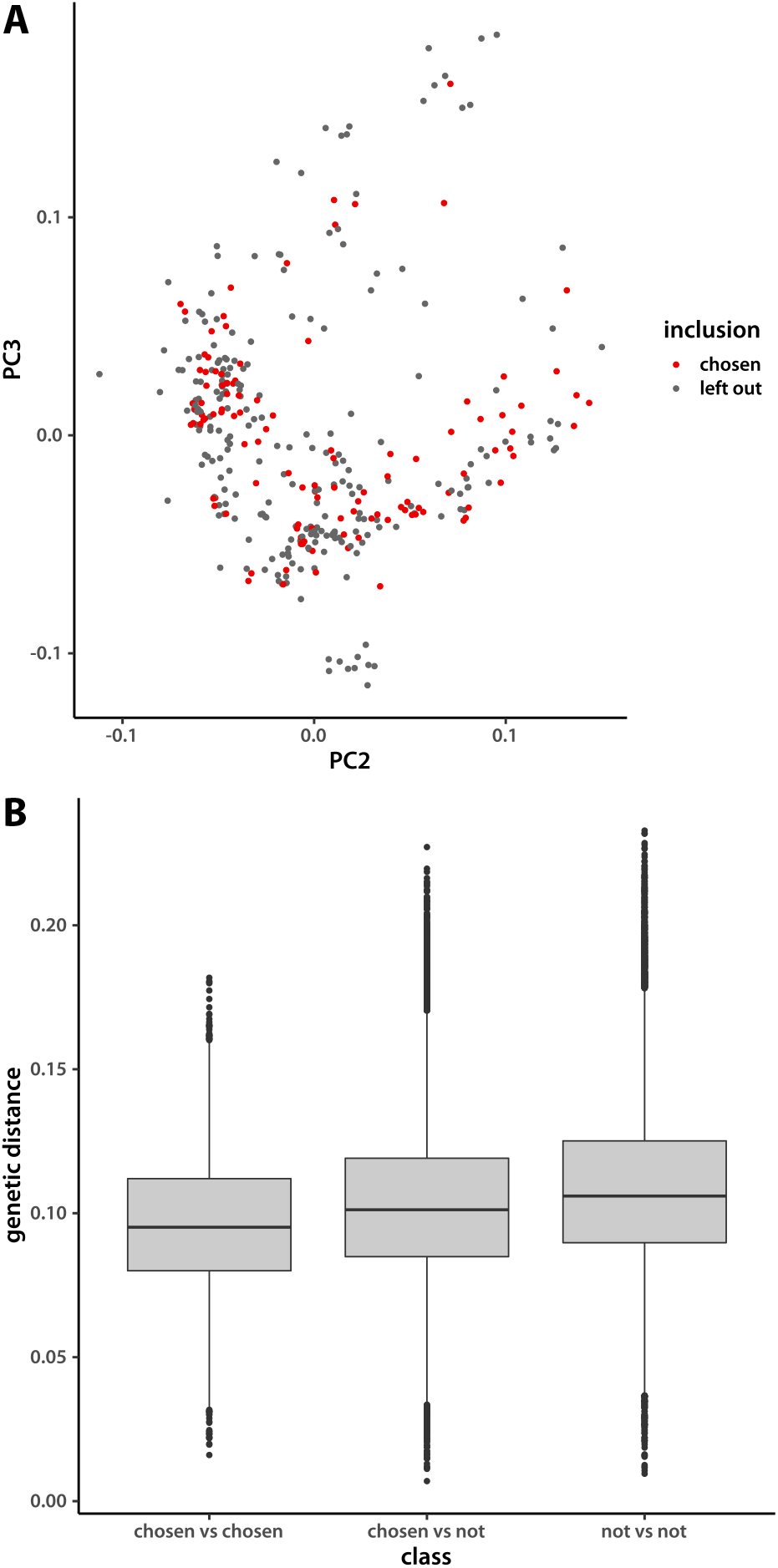
Genetic diversity of the chosen tropical japonica accessions. (A) Principal component plots of TRJ accessions with HDRA genotypes. Components 2 and 3 were chosen because outlier genotypes make PC1 less informative. (B) Distributions of genetic distances among chosen, chosen vs left out, and among left-out accessions.

**Supplemental Figure 4:**
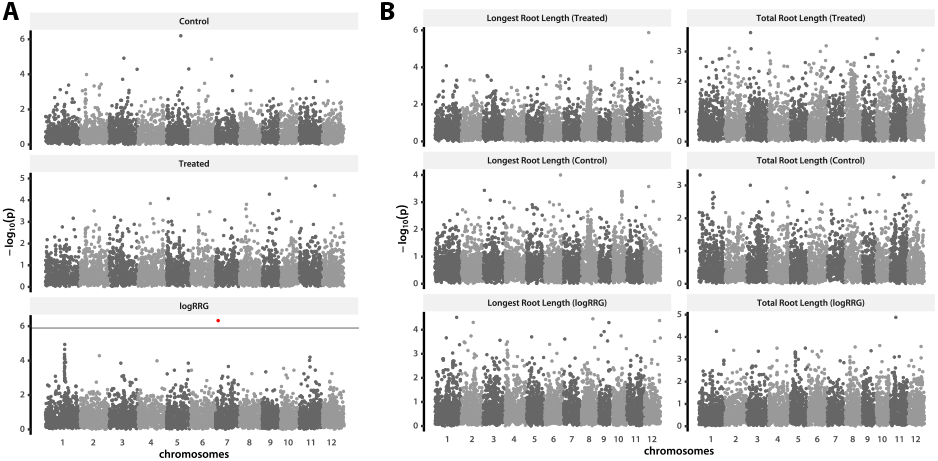
GWA in tropical japonica. (A) The population examined in Famoso *et al.* (2011) with additional accessions measured under the same conditions. (B) The new set of accessions measured under harsher aluminum treatment.

**Supplemental Figure 5:**
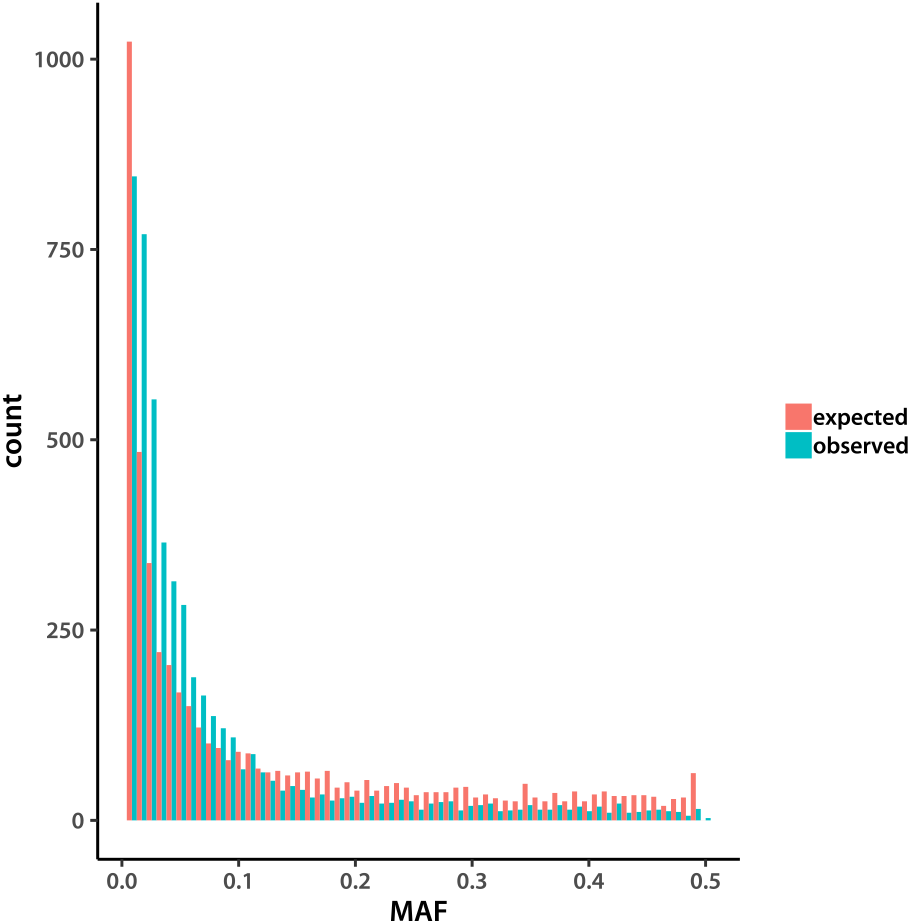
Allele frequency spectrum of tropical japonica SNPs. Folded site frequency spectrum of the 119 accessions genotyped with HDRA. Expected values are from simulations under an equilibrium neutral model. Minor allele frequency is on the *x* axis.

