## Supplemental methods and scripts for "Low additive genetic variation in a trait under selection in domesticated rice": altolMV.cpp

```
/*
 * Copyright (c) 2015 Anthony J. Greenberg
 *
 * Redistribution and use in source and binary forms, with or without modification, are permitted provided that the following conditions are met:
 *
 * 1. Redistributions of source code must retain the above copyright notice, this list of conditions and the following disclaimer.
 *
 * 2. Redistributions in binary form must reproduce the above copyright notice, this list of conditions and the following disclaimer in the documentation and/or other materials provided with the distribution.
 *
 * 3. Neither the name of the copyright holder nor the names of its contributors may be used to endorse or promote products derived from this software without specific prior written permission.
 *
 * THIS SOFTWARE IS PROVIDED BY THE COPYRIGHT HOLDERS AND CONTRIBUTORS "AS IS" AND ANY EXPRESS OR IMPLIED WARRANTIES, INCLUDING, BUT NOT LIMITED TO,
 * THE IMPLIED WARRANTIES OF MERCHANTABILITY AND FITNESS FOR A PARTICULAR PURPOSE ARE DISCLAIMED. IN NO EVENT SHALL THE COPYRIGHT HOLDER OR CONTRIBUTORS
 * BE LIABLE FOR ANY DIRECT, INDIRECT, INCIDENTAL, SPECIAL, EXEMPLARY, OR CONSEQUENTIAL DAMAGES (INCLUDING, BUT NOT LIMITED TO, PROCUREMENT OF
 * SUBSTITUTE GOODS OR SERVICES; LOSS OF USE, DATA, OR PROFITS; OR BUSINESS INTERRUPTION) HOWEVER CAUSED AND ON ANY THEORY OF LIABILITY, WHETHER
 * IN CONTRACT, STRICT LIABILITY, OR TORT (INCLUDING NEGLIGENCE OR OTHERWISE) ARISING IN ANY WAY OUT OF THE USE OF THIS SOFTWARE, EVEN IF ADVISED OF
 * THE POSSIBILITY OF SUCH DAMAGE.
 */

/*
 *	This is to analyze aluminum tolerance data. The data are root length with and without Al treatment. I am treating these as separate traits.
 *	The treated and untreated plants are different individuals, so the error covariance matrix is diagonal.
 */

#include <vector>
#include <string>
#include <sstream>
#include <iostream>
#include <cmath>
#include <cctype>
#include <cstdio>

#include <MuGen.h>

using std::vector;
using std::string;
using std::cout;
using std::cerr;
using std::endl;
using std::flush;
using std::stringstream;
using std::remove;


void finishFlNam(string &, const vector<string> &);
string db2str(const double &);

void finishFlNam(string &start, const vector<string> &add){
	for (vector<string>::const_iterator stIt = add.begin(); stIt != add.end(); ++stIt) {
		start += "_";
		start += *stIt;
	}
	start += ".gbin";
}

string db2str(const double &val){
	stringstream nuOS;
	nuOS << val;
	string vString = nuOS.str();
	size_t perPs = vString.find('.');
	if (perPs != string::npos) {
		//vString.erase(perPs, 1);
		vString.replace(perPs, 1, "-");
	}
	return vString;
}

int main(int argc, char *argv[]){
	
	// parsing command-line flags
	bool bOn = false; // burnin-in
	bool sOn = false; // sampling
	bool tOn = false; // thinning
	bool gOn = false; // nuG
	bool TOn = false; // # of threads
	bool cOn = false; // chain ID
	bool pOn = false; // population ID
	
	// main constants
	//double nuE   = 3.0;
	double nuG   = 3.0;
	int Nbnin    = 10;
	int Nsamp    = 10;
	int Nthin    = 1;
	int nThr     = 4;
	
	string popID;
	//char popTok;
	string cNam;
	
	string phenoFolder("phenoData/");
	string genoFolder("genoData/");
	string chainFolder("chains/");
	
	// number of "traits"
	const size_t d = 2;
	
	// processing command line flags
	for (int iArg = 1; iArg < argc; iArg++) {
		char *pchar = argv[iArg];
		switch (pchar[0]) {
			case '-': {
				switch (pchar[1]) {
					case 's':
						sOn = true;
						break;
						
					case 'b':
						bOn = true;
						break;
						
					case 't':
						tOn = true;
						break;
						
					case 'g':
						gOn = true;
						break;
						
					case 'p':
						pOn = true;
						break;
						
					case 'T':
						TOn = true;
						break;
						
					case 'c':
						cOn = true;
						break;
						
					default:
						cerr << "ERROR: unrecognized flag " << pchar[1] << endl;
						exit(-1);
						break;
					
				}
			}
			break;
				
			default: {
				if (sOn) {
					sOn   = false;
					Nsamp = atoi(pchar);
				}
				else if (bOn) {
					bOn   = false;
					Nbnin = atoi(pchar);
				}
				else if (tOn) {
					tOn   = false;
					Nthin = atoi(pchar);
				}
				else if (gOn) {
					gOn = false;
					nuG = atof(pchar);
				}
				else if (pOn){
					pOn   = false;
					popID = pchar;
				}
				else if (TOn){
					TOn = false;
					nThr = atoi(pchar);
				}
				else if (cOn){
					cOn = false;
					cNam = pchar;
				}
				
				break;
			}
		}
	}
	
	string popIDcaps;
	if (popID == "all") {
		popIDcaps = "All";
	}
	else if (popID == "ia"){
		popIDcaps = "IA";
	}
	else if (popID == "aus"){
		popIDcaps = "AUS";
	}
	else if (popID == "jap"){
		popIDcaps = "JAP";
	}
	else if (popID == "trj"){
		popIDcaps = "TRJ";
	}
	else if (popID == "trja"){
		popIDcaps = "TRJA";
	}
	else {
		cerr << "ERROR: Population name " << popID << " is not recognized" << endl;
		exit(1);
	}
	
	// Genotype file names
	string evecNam = genoFolder + "altl" + popIDcaps + "evc.gbin";
	string evalNam = genoFolder + "altl" + popIDcaps + "evl.gbin";
	// phenotype and index file names
	string phenInNam = phenoFolder + "Yat"   + popIDcaps + ".gbin"; // phenotype data
	string expMatNam = phenoFolder + "expMat"   + popIDcaps + ".gbin"; // experiment matrix
	string lnIndNam  = phenoFolder + "lnInd"    + popIDcaps + ".gbin"; // line index
	string mMatNam   = phenoFolder + "matNAind" + popIDcaps + ".gbin"; // missing value matrix
	string mVecNam   = phenoFolder + "totNAind" + popIDcaps+ ".gbin"; // missing value vector
	string trtBlkNam = phenoFolder + "trtBlk.gbin";                 // trait blocks for error covariance
	
	// parameters to save
	string LNout   = chainFolder + "LNout";  // line means
	string BVout   = chainFolder + "BVout";  // GEBVs
	string EXPout  = chainFolder + "EXPout"; // experiment effects
	string SgEout  = chainFolder + "SgEout"; // error covariance
	string SgSout  = chainFolder + "SgSout"; // sca covariance
	string SgAout  = chainFolder + "SgAout"; // gca covariance
	//string bSNPout = chainFolder + "abfSNPout"; // SNP ABFs
	
	vector<string> addOn;
	addOn.push_back(popID);
	addOn.push_back(db2str(nuG));
	addOn.push_back(cNam);
	
	finishFlNam(LNout, addOn);
	finishFlNam(BVout, addOn);
	finishFlNam(EXPout, addOn);
	finishFlNam(SgEout, addOn);
	finishFlNam(SgSout, addOn);
	finishFlNam(SgAout, addOn);
	//finishFlNam(bSNPout, addOn);
	
	size_t Nexp = 6; // number of levels in experiment
	// populating model constants
	size_t N;    // total number of data points
	size_t Nln;  // number of lines
	
	if (popID == "all") {
		N    = 4428;
		Nln  = 361;
	}
	else if (popID == "ia"){
		N    = 2009;
		Nln  = 141;
	}
	else if (popID == "aus"){
		N    = 794;
		Nln  = 55;
	}
	else if (popID == "jap"){
		N    = 2349;
		Nln  = 213;
	}
	else if (popID == "trj"){
		N    = 1007;
		Nln  = 92;
	}
	else if (popID == "trja"){
		N    = 1037;
		Nln  = 95;
	}
	else {
		cerr << "ERROR: Population name " << popID << " is not recognized" << endl;
		exit(1);
	}
	
	const size_t Npc = Nln - 1; // number of PCs in the relationship matrix
	
	RanIndex e2ln(N, Nln, lnIndNam);    // data to lines
	RanIndex e2e(N,N);                  // data to data prior
	RanIndex ln2mu(Nln);                // lines to intercept
	RanIndex ln2ln(Nln, Nln);           // line to genetic regression
	RanIndex exp2mu(Nexp);              // experiment to mean
	RanIndex gm2pr(Npc);                // PC regression coefficients to prior
	RanIndex mu2pr;                     // intercept to prior
	
	MuGrpMiss dataI(phenInNam, mMatNam, mVecNam, e2e, d);
	Grp &data = dataI;
	
	BetaGrpFt betaExpI(data, expMatNam, Nexp, exp2mu, EXPout, nThr);
	Grp &betaExp = betaExpI;
	
	MuGrp lnPredI = data - betaExp;
	Grp &lnPred   = lnPredI;
	
	MuGrp muLnI(lnPred, e2ln, ln2ln, LNout);
	Grp &muLn = muLnI;
	
	MuGrp betaExpPredI = data - muLn;
	Grp &betaExpPred = betaExpPredI;
	
	MuGrp muI(muLn, ln2mu, mu2pr);
	Grp &mu = muI;
	
	MuGrp gmPredI = muLn - mu;
	Grp &gmPred   = gmPredI;
	
	//BetaGrpPCpex gamI(gmPred, evecNam, evalNam, Npc, 1e-6, gm2pr, nThr);
	BetaGrpPC gamI(gmPred, evecNam, evalNam, Npc, gm2pr, nThr);
	Grp &gam = gamI;
	
	MuGrp gcaI = mu + gam;
	Grp &gca   = gcaI;
	
	MuGrp muPredI = muLn - gam;
	Grp &muPred   = muPredI;
	
	MuGrp dataPrI = muLn + betaExp;
	Grp &dataPr   = dataPrI;
	
	MuGrp eResdI = lnPred - muLn; // error residuals
	Grp &eResd   = eResdI;
	
	MuGrp sResdI = gmPred - gam; // genetic model residuals
	Grp &sResd   = sResdI;
	
	//SigmaI SigIe(eResd, SgEout, 1e-6, 2.0);
	SigmaIblk SigIeI = SigmaIblk(eResd, trtBlkNam, SgEout, 1.0, 2.0);
	SigmaI &SigIe = SigIeI;
	SigmaI SigIs(sResd, SgSout, 1.0, 2.0);
	SigmaI SigIa(gam, SgAout, 1.0, 4.0);
	SigmaI SigIpr(d, 1e-4);
	
	// For blocked residuals, no Student-t model for now
	//Qgrp qE(N, nuE, mVecNam);
	Qgrp qG(Npc, nuG);
	
	cout << "Burn-in..." << endl;
	for (int iBnin = 0; iBnin < Nbnin; iBnin++) {
		data.update(dataPr, SigIe);
		
		lnPredI = data - betaExp;
		muLn.update(lnPred, SigIe, gca, SigIs);
		
		betaExpPredI = data - muLn;
		betaExp.update(betaExpPred, SigIe, SigIpr);
		dataPrI = muLn + betaExp;
		eResdI  = data - dataPr;
		SigIe.update(eResd);
		
		gmPredI = muLn - mu;
		gam.update(gmPred, SigIs, qG, SigIa);
		SigIa.update(gam, qG);
		qG.update(gmPred, SigIa);
		
		muPredI = muLn - gam;
		mu.update(muPred, SigIs, SigIpr);
		sResdI = muPred - mu;
		gcaI   = mu + gam;
		SigIs.update(sResd);
		
		cout << "+" << flush;
	}
	cout << endl;
	cout << "Sampling..." << endl;
	for (int iSam = 0; iSam < Nsamp; iSam++) {
		data.update(dataPr, SigIe);
		
		lnPredI = data - betaExp;
		muLn.update(lnPred, SigIe, gca, SigIs);
		
		betaExpPredI = data - muLn;
		betaExp.update(betaExpPred, SigIe, SigIpr);
		dataPrI = muLn + betaExp;
		eResdI  = data - dataPr;
		SigIe.update(eResd);
		
		gmPredI = muLn - mu;
		gam.update(gmPred, SigIs, qG, SigIa);
		SigIa.update(gam, qG);
		qG.update(gmPred, SigIa);
		
		muPredI = muLn - gam;
		mu.update(muPred, SigIs, SigIpr);
		sResdI = muPred - mu;
		gcaI   = mu + gam;
		SigIs.update(sResd);

		if ((iSam + 1) % Nthin) {
			cout << "." << flush;
		}
		else {
			muLn.save();
			betaExp.save();
			gca.save(BVout);
			SigIe.save();
			SigIs.save();
			SigIa.save();
			
			cout << "|" << flush;
		}
		
	}
	cout << endl;

}
```
