## Supplemental methods and scripts for "Low additive genetic variation in a trait under selection in domesticated rice": altolMVnew.cpp

```
/*
 * Copyright (c) 2019 Anthony J. Greenberg
 *
 * Redistribution and use in source and binary forms, with or without modification, are permitted provided that the following conditions are met:
 *
 * 1. Redistributions of source code must retain the above copyright notice, this list of conditions and the following disclaimer.
 *
 * 2. Redistributions in binary form must reproduce the above copyright notice, this list of conditions and the following disclaimer in the documentation and/or other materials provided with the distribution.
 *
 * 3. Neither the name of the copyright holder nor the names of its contributors may be used to endorse or promote products derived from this software without specific prior written permission.
 *
 * THIS SOFTWARE IS PROVIDED BY THE COPYRIGHT HOLDERS AND CONTRIBUTORS "AS IS" AND ANY EXPRESS OR IMPLIED WARRANTIES, INCLUDING, BUT NOT LIMITED TO,
 * THE IMPLIED WARRANTIES OF MERCHANTABILITY AND FITNESS FOR A PARTICULAR PURPOSE ARE DISCLAIMED. IN NO EVENT SHALL THE COPYRIGHT HOLDER OR CONTRIBUTORS
 * BE LIABLE FOR ANY DIRECT, INDIRECT, INCIDENTAL, SPECIAL, EXEMPLARY, OR CONSEQUENTIAL DAMAGES (INCLUDING, BUT NOT LIMITED TO, PROCUREMENT OF
 * SUBSTITUTE GOODS OR SERVICES; LOSS OF USE, DATA, OR PROFITS; OR BUSINESS INTERRUPTION) HOWEVER CAUSED AND ON ANY THEORY OF LIABILITY, WHETHER
 * IN CONTRACT, STRICT LIABILITY, OR TORT (INCLUDING NEGLIGENCE OR OTHERWISE) ARISING IN ANY WAY OUT OF THE USE OF THIS SOFTWARE, EVEN IF ADVISED OF
 * THE POSSIBILITY OF SUCH DAMAGE.
 */

/*
 * Analyzing the new TRJ aluminum tolerance data. Several traits, one exeriment with no experimental covariates.
 */

#include <vector>
#include <string>
#include <unordered_map>
#include <sstream>
#include <iostream>
#include <cmath>
#include <cctype>
#include <cstdio>

#include "MuGen.h"

using std::vector;
using std::string;
using std::stod;
using std::stoi;
using std::cout;
using std::cerr;
using std::endl;
using std::flush;
using std::stringstream;
using std::remove;
using std::unordered_map;


void parseCL(int&, char**, unordered_map<char, string> &);
void finishFlNam(string &, const vector<string> &);
string db2str(const double &);

/*
 * Command line parser. Takes the argv array and produces a map of flags to values (as strings)
 */
void parseCL(int &argc, char **argv, unordered_map<char, string> &cli){
	// set to true after encountering a flag token (the character after the dash)
	bool val = false;
	// store the token value here
	char curFlag;

	for (int iArg = 1; iArg < argc; iArg++) {
		const char *pchar = argv[iArg];

		if (pchar[0] == '-') { // encountered the dash, look for the token after it
			if (!pchar[1]) {
				cerr << "ERROR: forgot character after dash" << endl;
				exit(-1);
			}
			// what follows the dash?
			val     = true;
			curFlag = pchar[1];

		} else {
			if (val) {
				val = false;
				cli[curFlag] = pchar;
			} else {
				cerr << "WARNING: command line value " << pchar << " ignored because it is not preceded by a flag" << endl;
			}

		}

	}
}

void finishFlNam(string &start, const vector<string> &add){
	for (vector<string>::const_iterator stIt = add.begin(); stIt != add.end(); ++stIt) {
		start += "_";
		start += *stIt;
	}
	start += ".gbin";
}

string db2str(const double &val){
	stringstream nuOS;
	nuOS << val;
	string vString = nuOS.str();
	size_t perPs = vString.find('.');
	if (perPs != string::npos) {
		vString.replace(perPs, 1, "-");
	}
	return vString;
}

int main(int argc, char *argv[]){

	unordered_map <char, string> clInfo;
	parseCL(argc, argv, clInfo);
	auto clIter = clInfo.begin(); // iterator of the command line flags

	// define constants from CL input. Have to use find() and not subscript or else missing flags will be inserted
	// burn-in
	int Nbnin;
	clIter = clInfo.find('b');
	if (clIter == clInfo.end()) { // if not there, set a default
		Nbnin = 10;
	} else {
		Nbnin = stoi(clIter->second);
	}

	// sampling
	int Nsamp;
	clIter = clInfo.find('s');
	if (clIter == clInfo.end()) { // if not there, set a default
		Nsamp = 10;
	} else {
		Nsamp = stoi(clIter->second);
	}

	// thinning
	int Nthin;
	clIter = clInfo.find('t');
	if (clIter == clInfo.end()) { // if not there, set a default
		Nthin = 10;
	} else {
		Nthin = stoi(clIter->second);
	}

	// # of threads
	int nThr;
	clIter = clInfo.find('T');
	if (clIter == clInfo.end()) { // if not there, set a default
		nThr = 1;
	} else {
		nThr = stoi(clIter->second);
	}

	// genetic nu
	double nuG;
	clIter = clInfo.find('g');
	if (clIter == clInfo.end()) { // if not there, set a default
		nuG = 1000.0;
	} else {
		nuG = stod(clIter->second);
	}

	// residual nu
	double nuE;
	clIter = clInfo.find('e');
	if (clIter == clInfo.end()) { // if not there, set a default
		nuE = 1000.0;
	} else {
		nuE = stod(clIter->second);
	}

	// chain ID
	string cNam;
	clIter = clInfo.find('c');
	if (clIter == clInfo.end()) { // if not there, set a default
		cNam = "1";
	} else {
		cNam = clIter->second;
	}

	// data set (treatment or control)
	string datID;
	clIter = clInfo.find('d');
	if (clIter == clInfo.end()) { // if not there, set a default
		datID = "expt";
	} else {
		datID = clIter->second;
	}

	// K matrix type
	string relID;
	clIter = clInfo.find('r');
	if (clIter == clInfo.end()) { // if not there, set a default
		relID = "IBS";
	} else {
		relID = clIter->second;
	}

	// invariant constants
	const size_t d    = 2;        // # of traits
	const size_t Nacc = 119;      // # of accessions
	const size_t Npc  = (relID == "IBD" ? Nacc - 4 : Nacc - 1); // # of genetic PCs; equal to Nacc - 1 if using KoK or IBS K
	size_t N;                     // # of data points; varies by data set
	if (datID == "expt") {
		N = 954;
	} else if (datID == "ctrl"){
		N = 951;
	} else {
		N = 0;
		cerr << "ERROR: unknown data set name " << datID << endl;
		exit(1);
	}

	// invariant names
	const string dataFolder("modelData/");
	const string chainFolder("chains/");
	const string evecNam   = dataFolder + relID + "evc.gbin";  // eigenvectors
	const string evalNam   = dataFolder + relID + "evl.gbin";  // eigenvalues
	const string phenInNam = dataFolder + datID + "Data.gbin"; // phenotype data
	const string accIndNam = dataFolder + datID + "Acc.gbin";  // accession index

	// parameters to save
	string LNout  = chainFolder + "LN";   // line means
	string BVout  = chainFolder + "BV";   // GEBV
	string SgEout = chainFolder + "SgE";  // error covariance
	string SgSout = chainFolder + "SgS";  // sca covariance
	string SgAout = chainFolder + "SgA";  // gca covariance

	vector<string> addOn;
	addOn.push_back(datID);
	addOn.push_back(relID);
	addOn.push_back(db2str(nuE));
	addOn.push_back(db2str(nuG));
	addOn.push_back(cNam);

	// finish deriving output file names
	finishFlNam(LNout, addOn);
	finishFlNam(BVout, addOn);
	finishFlNam(SgEout, addOn);
	finishFlNam(SgSout, addOn);
	finishFlNam(SgAout, addOn);

	//
	RanIndex e2acc(N, Nacc, accIndNam); // data to accessions
	RanIndex acc2mu(Nacc);              // accessions to intercept
	RanIndex acc2acc(Nacc, Nacc);       // accessions to genetic regression
	RanIndex gm2pr(Npc);                // PC regression coefficients to prior
	RanIndex mu2pr;                     // intercept to prior

	// data
	MuGrp yI(phenInNam, e2acc, d);
	Grp  &y = yI;

	// accession means
	MuGrp muAccI(y, e2acc, acc2acc);
	Grp  &muAcc = muAccI;

	// intercept
	MuGrp muI(muAcc, acc2mu, mu2pr);
	Grp  &mu = muI;

	// gamma response
	MuGrp gmRspI = muAcc - mu;
	Grp  &gmRsp  = gmRspI;

	// PC regression
	BetaGrpPC gamI(gmRsp, evecNam, evalNam, Npc, gm2pr, nThr); // regression on genetic PCs
	Grp      &gam = gamI;

	// intercept response
	MuGrp muRspI = muAcc - gam;
	Grp  &muRsp  = muRspI;

	// GCA
	MuGrp gcaI = mu + gam;
	Grp  &gca  = gcaI;

	// residual
	MuGrp eRsdI = y - muAcc;
	Grp  &eRsd  = eRsdI;

	// non-additive genetic residual
	MuGrp sRsdI = muAcc - gca;
	Grp  &sRsd  = sRsdI;

	// inverse-covariances
	SigmaI iSigE(eRsdI, SgEout, 1.0, 2.0); // residual
	SigmaI iSigS(sRsd, SgSout, 1.0, 2.0);  // SCA
	SigmaI iSigA(gam, SgAout, 1.0, 4.0);   // GCA
	SigmaI iSigPr(d, 1e-6);                // prior covariance for the intercept

	// Student-t weights
	Qgrp qE(N, nuE);
	Qgrp qG(Npc, nuG);

	cout << "Burn-in..." << endl;
	for (int iBnin = 0; iBnin < Nbnin; iBnin++) {
		muAcc.update(y, qE, iSigE, gca, iSigS);
		gam.update(gmRsp, iSigS, qG, iSigA);
		mu.update(muRsp, iSigS, iSigPr);

		eRsdI  = y - muAcc;
		gmRspI = muAcc - mu;
		muRspI = muAcc - gam;
		gcaI   = mu + gam;
		sRsdI  = muAcc - gca;

		iSigE.update(eRsd, qE);
		qE.update(eRsd, iSigE);
		iSigS.update(sRsd);
		iSigA.update(gam, qG);
		qG.update(gam, iSigA);

		cout << "+" << flush;
	}
	cout << endl;

	cout << "Sampling..." << endl;
	for (int iSam = 0; iSam < Nsamp; iSam++) {
		muAcc.update(y, qE, iSigE, gca, iSigS);
		gam.update(gmRsp, iSigS, qG, iSigA);
		mu.update(muRsp, iSigS, iSigPr);

		eRsdI  = y - muAcc;
		gmRspI = muAcc - mu;
		muRspI = muAcc - gam;
		gcaI   = mu + gam;
		sRsdI  = muAcc - gca;

		iSigE.update(eRsd, qE);
		qE.update(eRsd, iSigE);
		iSigS.update(sRsd);
		iSigA.update(gam, qG);
		qG.update(gam, iSigA);

		if ((iSam + 1) % Nthin) {
			cout << "." << flush;
		}
		else {
			muAcc.save(LNout);
			gca.save(BVout);
			iSigE.save();
			iSigS.save();
			iSigA.save();

			cout << "|" << flush;
		}

	}
	cout << endl;
}
```
