## Supplemental methods and scripts for "Low additive genetic variation in a trait under selection in domesticated rice": altolMVtrj.cpp

```
/*
 * Copyright (c) 2019 Anthony J. Greenberg
 *
 * Redistribution and use in source and binary forms, with or without modification, are permitted provided that the following conditions are met:
 *
 * 1. Redistributions of source code must retain the above copyright notice, this list of conditions and the following disclaimer.
 *
 * 2. Redistributions in binary form must reproduce the above copyright notice, this list of conditions and the following disclaimer in the documentation and/or other materials provided with the distribution.
 *
 * 3. Neither the name of the copyright holder nor the names of its contributors may be used to endorse or promote products derived from this software without specific prior written permission.
 *
 * THIS SOFTWARE IS PROVIDED BY THE COPYRIGHT HOLDERS AND CONTRIBUTORS "AS IS" AND ANY EXPRESS OR IMPLIED WARRANTIES, INCLUDING, BUT NOT LIMITED TO,
 * THE IMPLIED WARRANTIES OF MERCHANTABILITY AND FITNESS FOR A PARTICULAR PURPOSE ARE DISCLAIMED. IN NO EVENT SHALL THE COPYRIGHT HOLDER OR CONTRIBUTORS
 * BE LIABLE FOR ANY DIRECT, INDIRECT, INCIDENTAL, SPECIAL, EXEMPLARY, OR CONSEQUENTIAL DAMAGES (INCLUDING, BUT NOT LIMITED TO, PROCUREMENT OF
 * SUBSTITUTE GOODS OR SERVICES; LOSS OF USE, DATA, OR PROFITS; OR BUSINESS INTERRUPTION) HOWEVER CAUSED AND ON ANY THEORY OF LIABILITY, WHETHER
 * IN CONTRACT, STRICT LIABILITY, OR TORT (INCLUDING NEGLIGENCE OR OTHERWISE) ARISING IN ANY WAY OUT OF THE USE OF THIS SOFTWARE, EVEN IF ADVISED OF
 * THE POSSIBILITY OF SUCH DAMAGE.
 */

/*
 *	This is to analyze aluminum tolerance data. There are added data from Stella on extra TRJ accessions. Only TRJ are included from the previous experiment (analyzed with altolMV.cpp).
 *	The data are root length with and without Al treatment. I am treating these as separate traits.
 *	The treated and untreated plants are different individuals, so the error covariance matrix is diagonal. The errors have to be Gaussian for the block-diagonal model.
 */

#include <vector>
#include <string>
#include <unordered_map>
#include <sstream>
#include <iostream>
#include <cmath>
#include <cctype>
#include <cstdio>

#include "MuGen.h"

using std::vector;
using std::string;
using std::stod;
using std::stoi;
using std::cout;
using std::cerr;
using std::endl;
using std::flush;
using std::stringstream;
using std::remove;
using std::unordered_map;


void parseCL(int&, char**, unordered_map<char, string> &);
void finishFlNam(string &, const vector<string> &);
string db2str(const double &);

/*
 * Command line parser. Takes the argv array and produces a map of flags to values (as strings)
 */
void parseCL(int &argc, char **argv, unordered_map<char, string> &cli){
	// set to true after encountering a flag token (the character after the dash)
	bool val = false;
	// store the token value here
	char curFlag;

	for (int iArg = 1; iArg < argc; iArg++) {
		const char *pchar = argv[iArg];

		if (pchar[0] == '-') { // encountered the dash, look for the token after it
			if (!pchar[1]) {
				cerr << "ERROR: forgot character after dash" << endl;
				exit(-1);
			}
			// what follows the dash?
			val     = true;
			curFlag = pchar[1];

		} else {
			if (val) {
				val = false;
				cli[curFlag] = pchar;
			} else {
				cerr << "WARNING: command line value " << pchar << " ignored because it is not preceded by a flag" << endl;
			}

		}
		
	}
}

void finishFlNam(string &start, const vector<string> &add){
	for (vector<string>::const_iterator stIt = add.begin(); stIt != add.end(); ++stIt) {
		start += "_";
		start += *stIt;
	}
	start += ".gbin";
}

string db2str(const double &val){
	stringstream nuOS;
	nuOS << val;
	string vString = nuOS.str();
	size_t perPs = vString.find('.');
	if (perPs != string::npos) {
		vString.replace(perPs, 1, "-");
	}
	return vString;
}

int main(int argc, char *argv[]){
	
	unordered_map <char, string> clInfo;
	parseCL(argc, argv, clInfo);
	auto clIter = clInfo.begin(); // iterator of the command line flags
	
	
	// define constants from CL input. Have to use find() and not subscript or else missing flags will be inserted
	// burn-in
	int Nbnin;
	clIter = clInfo.find('b');
	if (clIter == clInfo.end()) { // if not there, set a default
		Nbnin = 10;
	} else {
		Nbnin = stoi(clIter->second);
	}
	
	// sampling
	int Nsamp;
	clIter = clInfo.find('s');
	if (clIter == clInfo.end()) { // if not there, set a default
		Nsamp = 10;
	} else {
		Nsamp = stoi(clIter->second);
	}
	
	// thinning
	int Nthin;
	clIter = clInfo.find('t');
	if (clIter == clInfo.end()) { // if not there, set a default
		Nthin = 10;
	} else {
		Nthin = stoi(clIter->second);
	}
	
	// # of threads
	int nThr;
	clIter = clInfo.find('T');
	if (clIter == clInfo.end()) { // if not there, set a default
		nThr = 1;
	} else {
		nThr = stoi(clIter->second);
	}
	
	// genetic nu
	double nuG;
	clIter = clInfo.find('g');
	if (clIter == clInfo.end()) { // if not there, set a default
		nuG = 1000.0;
	} else {
		nuG = stod(clIter->second);
	}
	
	string cNam;
	clIter = clInfo.find('c');
	if (clIter == clInfo.end()) { // if not there, set a default
		cNam = "1";
	} else {
		cNam = clIter->second;
	}
	
	// invariant constants
	const size_t d = 2;           // # of "traits"
	const size_t N = 2007;        // # of data points
	const size_t Nacc = 169;      // # of accessions
	const size_t Npc  = Nacc - 1; // # of genetic PCs
	const size_t Nyr = 3;         // # of years (plus the intercept)
	
	// invariant names
	const string phenoFolder("phenoData/");
	const string genoFolder("genoData/");
	const string chainFolder("chains/");
	const string evecNam = genoFolder + "altlTRJevc.gbin"; // eigenvectors
	const string evalNam = genoFolder + "altlTRJevl.gbin"; // eigenvalues
	string phenInNam = phenoFolder + "yMatS.gbin";         // phenotype data
	string yrPrdNam = phenoFolder + "Xyr.gbin";            // experiment contrast predictor
	string lnIndNam  = phenoFolder + "accIndS.gbin";       // accession index
	string mMatNam   = phenoFolder + "matNAindS.gbin";     // missing value matrix
	string mVecNam   = phenoFolder + "totNAindS.gbin";     // missing value vector
	string trtBlkNam = phenoFolder + "trtBlk.gbin";        // trait blocks for error covariance
	
	// parameters to save
	string LNout  = chainFolder + "LN";   // line means
	string LYout  = chainFolder + "LY";   // line means corrected by year effect
	string BVout  = chainFolder + "BV";   // GEBV
	string YRout  = chainFolder + "YR";   // experiment effects
	string SgEout = chainFolder + "SgE";  // error covariance
	string SgSout = chainFolder + "SgS";  // sca covariance
	string SgAout = chainFolder + "SgA";  // gca covariance
	
	vector<string> addOn;
	addOn.push_back(db2str(nuG));
	addOn.push_back(cNam);
	
	finishFlNam(LNout, addOn);
	finishFlNam(LYout, addOn);
	finishFlNam(BVout, addOn);
	finishFlNam(YRout, addOn);
	finishFlNam(SgEout, addOn);
	finishFlNam(SgSout, addOn);
	finishFlNam(SgAout, addOn);
	
	RanIndex e2acc(N, Nacc, lnIndNam);  // data to accessions
	RanIndex acc2mu(Nacc);              // accessions to intercept
	RanIndex acc2acc(Nacc, Nacc);       // accessions to genetic regression
	RanIndex yr2pr(Nyr);				// year to prior
	RanIndex gm2pr(Npc);                // PC regression coefficients to prior
	RanIndex mu2pr;                     // intercept to prior
	
	MuGrpMiss dataI(phenInNam, mMatNam, mVecNam, e2acc, d);
	Grp &data = dataI;
	
	MuGrp muAccI(data, e2acc, acc2acc, LNout); // accession means
	Grp &muAcc = muAccI;
	
	BetaGrpFt betaYrI(muAcc, yrPrdNam, Nyr, yr2pr, nThr);
	Grp &betaYr = betaYrI;
	
	MuGrp gmRspI = muAcc - betaYr; // response for the regression on genetic PCs
	Grp &gmRsp   = gmRspI;
	
	BetaGrpPC gamI(gmRsp, evecNam, evalNam, Npc, gm2pr, nThr); // regression on genetic PCs
	Grp &gam = gamI;
	
	MuGrp yrRspI = muAcc - gam; // response for intercept
	Grp &yrRsp   = yrRspI;

	MuGrp mu0I(acc2mu, d); // zero-vector to calculate fitted additive effects
	Grp &mu0 = mu0I;
	
	MuGrp gcaI = mu0 + gam;
	Grp &gca   = gcaI;
	
	MuGrp corrAccI = muAcc - betaYr;
	Grp &corrAcc   = corrAccI;
	
	MuGrp accPrI = betaYr + gam;
	Grp &accPr   = accPrI;
	
	MuGrp eResdI = data - muAcc; // error residuals
	Grp &eResd   = eResdI;

	MuGrp sResdI = gmRsp - betaYr; // genetic model residuals
	Grp &sResd   = sResdI;
	
	SigmaIblk SigIeI = SigmaIblk(eResd, trtBlkNam, SgEout, 1.0, 2.0);
	SigmaI &SigIe = SigIeI;                // error covariance
	SigmaI SigIs(sResd, SgSout, 1.0, 2.0); // non-additive genetic covariance
	SigmaI SigIa(gam, SgAout, 1.0, 4.0);   // additive genetic covariance
	SigmaI SigIpr(d, 1e-6);                // prior covariance for the intercept
	
	Qgrp qG(Npc, nuG);
	
	cout << "Burn-in..." << endl;
	for (int iBnin = 0; iBnin < Nbnin; iBnin++) {
		data.update(muAcc, SigIe);
		muAcc.update(data, SigIe, accPr, SigIs);
		betaYr.update(yrRsp, SigIs, SigIpr);
		gam.update(gmRsp, SigIs, qG, SigIa);
		
		yrRspI = muAcc - gam;
		gmRspI = muAcc - betaYr;
		accPrI = gam + betaYr;
		
		eResdI  = data - muAcc;
		sResdI  = gmRsp - betaYr;
		
		SigIe.update(eResd);
		SigIs.update(sResd);
		SigIa.update(gam, qG);
		qG.update(gam, SigIa);
		
		cout << "+" << flush;
	}
	cout << endl;
	
	cout << "Sampling..." << endl;
	for (int iSam = 0; iSam < Nsamp; iSam++) {
		data.update(muAcc, SigIe);
		muAcc.update(data, SigIe, accPr, SigIs);
		betaYr.update(yrRsp, SigIs, SigIpr);
		gam.update(gmRsp, SigIs, qG, SigIa);
		
		yrRspI = muAcc - gam;
		gmRspI = muAcc - betaYr;
		accPrI = gam + betaYr;
		
		eResdI  = data - muAcc;
		sResdI  = gmRsp - betaYr;
		
		SigIe.update(eResd);
		SigIs.update(sResd);
		SigIa.update(gam, qG);
		qG.update(gam, SigIa);
		
		if ((iSam + 1) % Nthin) {
			cout << "." << flush;
		}
		else {
			muAcc.save();
			gcaI = mu0 + gam;
			gca.save(BVout);
			corrAccI = muAcc - betaYr;
			corrAcc.save(LYout);
			betaYr.save(YRout);
			SigIe.save();
			SigIs.save();
			SigIa.save();
			
			cout << "|" << flush;
		}
		
	}
	cout << endl;

}
```
