## Supplemental methods and scripts for "Low additive genetic variation in a trait under selection in domesticated rice": GSLRIO.cpp

```
/*
 * Copyright (c) 2015 Anthony J. Greenberg
 *
 * Redistribution and use in source and binary forms, with or without modification, are permitted provided that the following conditions are met:
 *
 * 1. Redistributions of source code must retain the above copyright notice, this list of conditions and the following disclaimer.
 *
 * 2. Redistributions in binary form must reproduce the above copyright notice, this list of conditions and the following disclaimer in the documentation and/or other materials provided with the distribution.
 *
 * 3. Neither the name of the copyright holder nor the names of its contributors may be used to endorse or promote products derived from this software without specific prior written permission.
 *
 * THIS SOFTWARE IS PROVIDED BY THE COPYRIGHT HOLDERS AND CONTRIBUTORS "AS IS" AND ANY EXPRESS OR IMPLIED WARRANTIES, INCLUDING, BUT NOT LIMITED TO,
 * THE IMPLIED WARRANTIES OF MERCHANTABILITY AND FITNESS FOR A PARTICULAR PURPOSE ARE DISCLAIMED. IN NO EVENT SHALL THE COPYRIGHT HOLDER OR CONTRIBUTORS
 * BE LIABLE FOR ANY DIRECT, INDIRECT, INCIDENTAL, SPECIAL, EXEMPLARY, OR CONSEQUENTIAL DAMAGES (INCLUDING, BUT NOT LIMITED TO, PROCUREMENT OF
 * SUBSTITUTE GOODS OR SERVICES; LOSS OF USE, DATA, OR PROFITS; OR BUSINESS INTERRUPTION) HOWEVER CAUSED AND ON ANY THEORY OF LIABILITY, WHETHER
 * IN CONTRACT, STRICT LIABILITY, OR TORT (INCLUDING NEGLIGENCE OR OTHERWISE) ARISING IN ANY WAY OUT OF THE USE OF THIS SOFTWARE, EVEN IF ADVISED OF
 * THE POSSIBILITY OF SUCH DAMAGE.
 */

/// R interface for GSL binary file format
/** \file
 * \author Anthony J. Greenberg
 * \copyright Copyright (c) 2015 Anthony J. Greenberg
 * \version 0.9.1
 *
 * Functions to read and write binary files saved in GSL binary format from R. 
 * 
 * To compile, run on command line:
 * 
 *    R CMD SHLIB GSLRIO.cpp -lgsl -O3 -DHAVE_INLINE -DGSL_RANGE_CHECK_OFF
 *
 * and put the resulting GSLRIO.o where it can be found by `dyn.load()` in R.  The functions are then ready to use with the `.C()` command.
 *
 * \note To pass double-quoted strings from `.C()`, the corresponding arguments (here they are file names) must be in the form **fileNam.
 *
 */

#include <R.h>
#include <string>
#include <iostream>
#include <algorithm>
#include <gsl/gsl_matrix.h>
#include <gsl/gsl_vector.h>

using std::string;
using std::remove;
using std::cerr;
using std::endl;

/** \brief Save a double-precision matrix
 *
 * The fact that R matrices are column-major, while in C/C++ they are row-major is dealt with internally.  The R user does not have to worry about that.
 *
 * \param[in] char** output file name
 * \param[in] double* vectorized matrix to be saved
 * \param[in] int* pointer to the number of rows
 * \param[in] int* pointer to the number of columns
 */
extern "C"{
	void GSLmatSave(const char **fileNam, const double *vec, const int *nRows, const int *nCols){
		gsl_matrix_const_view mat = gsl_matrix_const_view_array(vec, *nCols, *nRows); // because the R matrix is vectorized row by row
		remove(*fileNam);
		
		for (size_t iRw = 0; iRw < *nRows; iRw++) { // the rows of the saved matrix are the columns of the passed matrix
			gsl_vector_const_view rw = gsl_matrix_const_column(&mat.matrix, iRw);
			FILE *fl = fopen(*fileNam, "a");
			gsl_vector_fwrite(fl, &rw.vector);
			fclose(fl);
		}
		
	}
	
}

/** \brief Save a vector of integers
 *
 * Typically used to save index (factor) vectors to read into RanIndex objects.
 *
 * \param[in] char** output file name
 * \param[in] int* array of integers
 * \param[in] int* pointer to the array length
 */
extern "C"{
	void GSLvecSaveInt(const char **fileNam, const int *vec, const int *len){
		gsl_vector_int_const_view svVec = gsl_vector_int_const_view_array(vec, *len);
		
		FILE *fl = fopen(*fileNam, "w");
		gsl_vector_int_fwrite(fl, &svVec.vector);
		fclose(fl);
		
	}
	
}
 
/** \brief Save a double-precision vector
 *
 * \param[in] char** output file name
 * \param[in] double* double-precision array
 * \param[in] int* pointer to the array size
 */
extern "C"{
	void GSLvecSave(const char **fileNam, const double *vec, const int *len){
		gsl_vector_const_view svVec = gsl_vector_const_view_array(vec, *len);
		
		FILE *fl = fopen(*fileNam, "w");
		gsl_vector_fwrite(fl, &svVec.vector);
		fclose(fl);
		
	}
	
}

/** \brief Appending a double-precision vector to an existing file
 *
 * \param[in] char** output file name
 * \param[in] double* double-precision array
 * \param[in] int* pointer to the array size
 */
extern "C"{
	void GSLvecAppend(const char **fileNam, const double *vec, const int *len){
		gsl_vector_const_view svVec = gsl_vector_const_view_array(vec, *len);
		
		FILE *fl = fopen(*fileNam, "a");
		gsl_vector_fwrite(fl, &svVec.vector);
		fclose(fl);
		
	}
	
}
/** \brief Read a vector of integers
 *
 * Reading a file into an array of integers.
 *
 * \warning While file existence is checked, there currently is no way to check file size.  If the file is too small to hold an array of integers of the required length, the function will crash and likely take the R session with it.
 *
 * \param[in] char** output file name
 * \param[in] int* pointer to array length
 * \param[out] int* array of integers to be passed to R
 */
extern "C" {
	void GSLiVecLoad(const char **fileNam, const int *len, int *arr){
		gsl_vector_int_view vec = gsl_vector_int_view_array(arr, *len);
		
		FILE *fl = fopen(*fileNam, "r");
		if (fl == NULL) {
			cerr << "ERROR: cannot open file" << endl;
			exit(-1);
		}
		gsl_vector_int_fread(fl, &vec.vector);
		fclose(fl);
		
	}
	
}

/** \brief Read a double-precision array or matrix
 *
 * Reads a double-precision array (vector) or matrix.  If a vector is desired, one of the dimensions should be set to one. The matrix is stored by row, so make sure to use matrix(..., byrow = T) in the R code.
 *
 * \warning While file existence is checked, there currently is no way to check file size.  If the file is too small to hold an array of doubles of the required length, the function will crash and likely take the R session with it.
 *
 * \param[in] char** output file name
 * \param[in] int* pointer to the number of rows
 * \param[in] int* pointer to the number of columns
 * \param[out] double* array to be passed to R and converted to a matrix if necessary
 */

extern "C" {
	void GSLmatLoad(const char **fileNam, const int *nRows, const int *nCols, double *arr){
		int len = (*nRows)*(*nCols);
		gsl_vector_view vec = gsl_vector_view_array(arr, len);
		
		FILE *fl = fopen(*fileNam, "r");
		if (fl == NULL) {
			cerr << "ERROR: cannot open file" << endl;
			exit(-1);
		}
		gsl_vector_fread(fl, &vec.vector);
		fclose(fl);
		
	}
	
}
```
