## Supplemental methods and scripts for "Low additive genetic variation in a trait under selection in domesticated rice": MuGen.cpp

```
/*
 * Copyright (c) 2015 Anthony J. Greenberg
 *
 * Redistribution and use in source and binary forms, with or without modification, are permitted provided that the following conditions are met:
 *
 * 1. Redistributions of source code must retain the above copyright notice, this list of conditions and the following disclaimer.
 *
 * 2. Redistributions in binary form must reproduce the above copyright notice, this list of conditions and the following disclaimer in the documentation and/or other materials provided with the distribution.
 *
 * 3. Neither the name of the copyright holder nor the names of its contributors may be used to endorse or promote products derived from this software without specific prior written permission.
 *
 * THIS SOFTWARE IS PROVIDED BY THE COPYRIGHT HOLDERS AND CONTRIBUTORS "AS IS" AND ANY EXPRESS OR IMPLIED WARRANTIES, INCLUDING, BUT NOT LIMITED TO,
 * THE IMPLIED WARRANTIES OF MERCHANTABILITY AND FITNESS FOR A PARTICULAR PURPOSE ARE DISCLAIMED. IN NO EVENT SHALL THE COPYRIGHT HOLDER OR CONTRIBUTORS
 * BE LIABLE FOR ANY DIRECT, INDIRECT, INCIDENTAL, SPECIAL, EXEMPLARY, OR CONSEQUENTIAL DAMAGES (INCLUDING, BUT NOT LIMITED TO, PROCUREMENT OF
 * SUBSTITUTE GOODS OR SERVICES; LOSS OF USE, DATA, OR PROFITS; OR BUSINESS INTERRUPTION) HOWEVER CAUSED AND ON ANY THEORY OF LIABILITY, WHETHER
 * IN CONTRACT, STRICT LIABILITY, OR TORT (INCLUDING NEGLIGENCE OR OTHERWISE) ARISING IN ANY WAY OUT OF THE USE OF THIS SOFTWARE, EVEN IF ADVISED OF
 * THE POSSIBILITY OF SUCH DAMAGE.
 */

/// Implementation of C++ classes for Hierarchical Bayesian Multi-trait quantitative-genetic models.
/** \file
 * \author Anthony J. Greenberg
 * \copyright Copyright (c) 2015 Anthony J. Greenberg
 * \version 0.9.1
 *
 * This file contains the implementation of the methods documented in the MuGen.h file.
 *
 */


#include <gsl/gsl_errno.h>
#include <gsl/gsl_matrix.h>
#include <gsl/gsl_vector.h>
#include <gsl/gsl_permutation.h>
#include <gsl/gsl_sort_vector.h>
#include <gsl/gsl_rng.h>
#include <gsl/gsl_randist.h>
#include <gsl/gsl_cdf.h>
#include <gsl/gsl_blas.h>
#include <gsl/gsl_linalg.h>
#include <gsl/gsl_statistics.h>
#include <gsl/gsl_math.h>
#include <gsl/gsl_machine.h>
#include <cmath>
#include <algorithm>
#include <vector>
#include <list>
#include <iostream>
#include <fstream>
#include <omp.h>

#include "MuGen.h"

using std::vector;
using std::list;
using std::isnan;
using std::cerr;
using std::cout;
using std::endl;
using std::flush;
using std::ofstream;
using std::ifstream;
using std::ios;
using std::max;
using std::max_element;
using std::min_element;
using std::remove;

/*
 *	Sampling functions
 *	IMPORTANT: Once started using CblasLower (required for MVgauss), have to use it everywhere.  Otherwise, matrix estimates diverge
 */

void MVgauss(const gsl_vector *mn, const gsl_matrix *SigChl, const gsl_rng *r, gsl_vector *samp){
	for (size_t iV = 0; iV < mn->size; iV++) {
		gsl_vector_set(samp, iV, gsl_ran_gaussian_ziggurat(r, 1.0));
	}
	gsl_blas_dtrmv(CblasLower, CblasNoTrans, CblasNonUnit, SigChl, samp);
	gsl_vector_add(samp, mn);
}

void Wishart(const gsl_matrix *SigC, const int &df, const gsl_rng *r, gsl_matrix *SigIout){
	gsl_matrix *tmp = gsl_matrix_calloc(SigC->size1, SigC->size2);
	for (size_t rwI = 0; rwI < SigC->size1; rwI++){
		gsl_matrix_set(tmp, rwI, rwI, sqrt(gsl_ran_chisq(r, df - rwI)));
		if (rwI == 0) {
			continue;
		}
		for (size_t clI = 0; clI < rwI - 1; clI++){
			gsl_matrix_set(tmp, rwI, clI, gsl_ran_gaussian_ziggurat(r, 1.0));
		}
	}
	gsl_blas_dtrmm(CblasLeft, CblasLower, CblasNoTrans, CblasNonUnit, 1.0, SigC, tmp);
	gsl_matrix_memcpy(SigIout, tmp); // necessary even though I am multiplying the matrix by itself.  Matrix can diverge otherwise
	gsl_blas_dtrmm(CblasRight, CblasLower, CblasTrans, CblasNonUnit, 1.0, tmp, SigIout);
	gsl_matrix_free(tmp);
}

void Wishart(const gsl_matrix *SigC, const size_t &df, const gsl_rng *r, gsl_matrix *SigIout){
	gsl_matrix *tmp = gsl_matrix_calloc(SigC->size1, SigC->size2);
	for (size_t rwI = 0; rwI < SigC->size1; rwI++){
		gsl_matrix_set(tmp, rwI, rwI, sqrt(gsl_ran_chisq(r, df - rwI)));
		if (rwI == 0) {
			continue;
		}
		for (size_t clI = 0; clI < rwI - 1; clI++){
			gsl_matrix_set(tmp, rwI, clI, gsl_ran_gaussian_ziggurat(r, 1.0));
		}
	}
	gsl_blas_dtrmm(CblasLeft, CblasLower, CblasNoTrans, CblasNonUnit, 1.0, SigC, tmp);
	gsl_matrix_memcpy(SigIout, tmp); // necessary even though I am multiplying the matrix by itself.  Matrix can diverge otherwise
	gsl_blas_dtrmm(CblasRight, CblasLower, CblasTrans, CblasNonUnit, 1.0, tmp, SigIout);
	gsl_matrix_free(tmp);
}

// Truncated geometric distribution; gives the unshifted (base-0) distribution
size_t rtgeom(const double &p, const size_t &limit, const gsl_rng *r){
	double u  = gsl_ran_flat(r, 0.0, 1.0);
	double up = gsl_cdf_geometric_P(static_cast<unsigned int>(limit), p);
	return static_cast<size_t>(floor(log(1.0 - u*up)/log(1.0 - p))); // inverse CDF of the geometric
}

/*
 *	function to calculate centered residuals to use for SNP regessions
 */

double mhl(const gsl_vector *beta, const gsl_matrix *SigI){ // distance to 0
	double m = 0.0;
	gsl_vector *tmp    = gsl_vector_alloc(SigI->size1);
	
	gsl_blas_dsymv(CblasLower, 1.0, SigI, beta, 0.0, tmp);
	gsl_blas_ddot(beta, tmp, &m);
	
	gsl_vector_free(tmp);
	
	return m;
}

/*
 *	Functions to mean-center columns of matrices
 */
void colCenter(gsl_matrix *inplace){
	gsl_vector *cl  = gsl_vector_alloc(inplace->size1);
	double clMn;
	for (size_t jCl = 0; jCl < inplace->size2; jCl++) {
		gsl_vector_view vCl = gsl_matrix_column(inplace, jCl);
		gsl_matrix_get_col(cl, inplace, jCl);
		clMn = gsl_stats_mean(cl->data, 1, cl->size);
		gsl_vector_add_constant(&vCl.vector, -clMn);
	}
	gsl_vector_free(cl);
}
void colCenter(const gsl_matrix *source, gsl_matrix *res){
	gsl_vector *cl  = gsl_vector_alloc(source->size1);
	double clMn;
	for (size_t jCl = 0; jCl < source->size2; jCl++) {
		gsl_matrix_get_col(cl, source, jCl);
		clMn = gsl_stats_mean(cl->data, 1, cl->size);
		gsl_vector_add_constant(cl, -clMn);
		gsl_matrix_set_col(res, jCl, cl);
	}
	gsl_vector_free(cl);

}
void colCenter(gsl_matrix *inplace, const double &absLab){
	double clMn;
	for (size_t jCl = 0; jCl < inplace->size2; jCl++) {
		vector<double> prNonMiss;
		vector<size_t> missInd;
		double val;
		for (size_t iRw = 0; iRw < inplace->size1; iRw++) {
			val = gsl_matrix_get(inplace, iRw, jCl);
			if (val > absLab) {
				prNonMiss.push_back(val);
			}
			else {
				missInd.push_back(iRw);
			}
		}
		
		gsl_vector_view vCl = gsl_matrix_column(inplace, jCl);
		clMn = gsl_stats_mean(prNonMiss.data(), 1, prNonMiss.size());
		gsl_vector_add_constant(&vCl.vector, -clMn);
		for (vector<size_t>::iterator missIt = missInd.begin(); missIt != missInd.end(); ++missIt) {
			gsl_matrix_set(inplace, *missIt, jCl, 0.0);
		}
	}
}
void colCenter(const gsl_matrix *source, gsl_matrix *res, const double &absLab){
	gsl_vector *cl  = gsl_vector_alloc(source->size1);
	double clMn;
	for (size_t jCl = 0; jCl < source->size2; jCl++) {
		vector<double> prNonMiss;
		vector<size_t> missInd;
		double val;
		for (size_t iRw = 0; iRw < source->size1; iRw++) {
			val = gsl_matrix_get(source, iRw, jCl);
			if (val > absLab) {
				prNonMiss.push_back(val);
			}
			else {
				missInd.push_back(iRw);
			}
		}
		
		gsl_matrix_get_col(cl, source, jCl);
		clMn = gsl_stats_mean(prNonMiss.data(), 1, prNonMiss.size());
		gsl_vector_add_constant(cl, -clMn);
		for (vector<size_t>::iterator missIt = missInd.begin(); missIt != missInd.end(); ++missIt) {
			gsl_vector_set(cl, *missIt, 0.0);
		}
		gsl_matrix_set_col(res, jCl, cl);
	}
	gsl_vector_free(cl);
	
}

void vecCenter(gsl_vector *inplace){
	double mn = gsl_stats_mean(inplace->data, 1, inplace->size);
	gsl_vector_add_constant(inplace, -mn);
}
void vecCenter(const gsl_vector *source, gsl_vector *res){
	gsl_vector_memcpy(res, source);
	double mn = gsl_stats_mean(res->data, 1, res->size);
	gsl_vector_add_constant(res, -mn);
}

void meanImpute(gsl_matrix *inplace, const double &absLab){
	double clMn;
	for (size_t jCl = 0; jCl < inplace->size2; jCl++) {
		vector<double> prNonMiss;
		vector<size_t> missInd;
		double val;
		for (size_t iRw = 0; iRw < inplace->size1; iRw++) {
			val = gsl_matrix_get(inplace, iRw, jCl);
			if (val > absLab) {
				prNonMiss.push_back(val);
			}
			else {
				missInd.push_back(iRw);
			}
		}
		
		clMn = gsl_stats_mean(prNonMiss.data(), 1, prNonMiss.size());
		for (vector<size_t>::iterator missIt = missInd.begin(); missIt != missInd.end(); ++missIt) {
			gsl_matrix_set(inplace, *missIt, jCl, 0.0);
		}
	}
}
// to print a GSL matrix to stdout
void printMat(const gsl_matrix *m){
	for (size_t i = 0; i < m->size1; i++) {
		for (size_t j = 0; j < m->size2; j++) {
			cout << gsl_matrix_get(m, i, j) << " " << flush;
		}
		cout << endl;
	}
}

// function that returns the number of CPU cycles since reset for use as a random number seed
unsigned long long rdtsc(){
	unsigned int lo,hi;
	__asm__ __volatile__ ("rdtsc" : "=a" (lo), "=d" (hi));
	return ((unsigned long long)hi << 32) | lo;
}

/*
 *	MVnorm methods
 */

MVnorm::MVnorm(gsl_vector *mn){
	_d = mn->size;
	_vec = gsl_vector_view_array(mn->data, 1);
}

MVnorm::MVnorm(gsl_vector *mn, const gsl_vector *sd, const gsl_rng *r){
	_d = mn->size;
	_vec = gsl_vector_view_array(mn->data, 1);
	
	gsl_vector *tmp = gsl_vector_alloc(_d);
	gsl_vector_memcpy(tmp, mn);
	for (int i = 0; i < _d; i++) {
		gsl_vector_set(&_vec.vector, i, gsl_vector_get(tmp, i) + gsl_ran_gaussian_ziggurat(r, gsl_vector_get(sd, i)));
	}
	gsl_vector_free(tmp);
}

MVnorm::MVnorm(gsl_vector *mn, const gsl_matrix *Sig, const gsl_rng *r){
	_d = mn->size;
	_vec = gsl_vector_view_array(mn->data, 1);
	gsl_matrix *chl = gsl_matrix_alloc(_d, _d);
	gsl_matrix_memcpy(chl, Sig);
	gsl_linalg_cholesky_decomp(chl);
	
	gsl_vector *tmp = gsl_vector_alloc(_d);
	gsl_vector_memcpy(tmp, mn);
	
	MVgauss(tmp, chl, r, &_vec.vector);
	gsl_matrix_free(chl);
	gsl_vector_free(tmp);
}

MVnorm::MVnorm(gsl_matrix *mn, const size_t &iRw){
	_d = mn->size2;
	_vec = gsl_matrix_row(mn, iRw);
}
MVnorm::MVnorm(gsl_matrix *mn, const size_t &iRw, const gsl_vector *sd, const gsl_rng *r){
	_d = mn->size2;
	_vec = gsl_matrix_row(mn, iRw);
	
	gsl_vector *tmp = gsl_vector_alloc(_d);
	gsl_vector_memcpy(tmp, &_vec.vector);
	for (int i = 0; i < _d; i++) {
		gsl_vector_set(&_vec.vector, i, gsl_vector_get(tmp, i) + gsl_ran_gaussian_ziggurat(r, gsl_vector_get(sd, i)));
	}
	gsl_vector_free(tmp);
}
MVnorm::MVnorm(gsl_matrix *mn, const size_t &iRw, const gsl_matrix *Sig, const gsl_rng *r){
	_d = mn->size2;
	_vec = gsl_matrix_row(mn, iRw);
	
	gsl_vector *tmp = gsl_vector_alloc(_d);
	gsl_vector_memcpy(tmp, &_vec.vector);
	
	gsl_matrix *chl = gsl_matrix_alloc(_d, _d);
	gsl_matrix_memcpy(chl, Sig);
	gsl_linalg_cholesky_decomp(chl);
	
	MVgauss(tmp, chl, r, &_vec.vector);
	
	gsl_matrix_free(chl);
	gsl_vector_free(tmp);
}

MVnorm::MVnorm(const MVnorm &mu){	//	Copy constructor
	_d = mu._d;
	_vec = mu._vec;
}

MVnorm & MVnorm::operator=(const MVnorm &mu){
	_d   = mu._d;
	_vec = mu._vec;
	
	return *this;
}

// Mahalanobis distance functions (in fact, square Mahalanobis distance)
double MVnorm::mhl(const gsl_vector *x, const SigmaI &SigI){
	double mhl = 0.0;
	gsl_vector *dif = gsl_vector_alloc(_d);
	gsl_vector *tmp = gsl_vector_alloc(_d);
	
	gsl_vector_memcpy(dif, x);
	gsl_vector_sub(dif, &_vec.vector);
	
	gsl_blas_dsymv(CblasLower, 1.0, SigI.getMat(), dif, 0.0, tmp);
	gsl_blas_ddot(dif, tmp, &mhl);
	
	gsl_vector_free(tmp);
	gsl_vector_free(dif);
	
	return mhl;
}
double MVnorm::mhl(const gsl_vector *x, const SigmaI &SigI) const{
	double mhl = 0.0;
	gsl_vector *dif = gsl_vector_alloc(_d);
	gsl_vector *tmp = gsl_vector_alloc(_d);
	
	gsl_vector_memcpy(dif, x);
	gsl_vector_sub(dif, &_vec.vector);
	
	gsl_blas_dsymv(CblasLower, 1.0, SigI.getMat(), dif, 0.0, tmp);
	gsl_blas_ddot(dif, tmp, &mhl);
	
	gsl_vector_free(tmp);
	gsl_vector_free(dif);
	
	return mhl;
}

double MVnorm::mhl(const MVnorm *x, const SigmaI &SigI){
	double mhl = 0.0;
	gsl_vector *dif = gsl_vector_alloc(_d);
	gsl_vector *tmp = gsl_vector_alloc(_d);
	
	gsl_vector_memcpy(dif, x->getVec());
	gsl_vector_sub(dif, &_vec.vector);
	
	gsl_blas_dsymv(CblasLower, 1.0, SigI.getMat(), dif, 0.0, tmp);
	gsl_blas_ddot(dif, tmp, &mhl);
	
	gsl_vector_free(tmp);
	gsl_vector_free(dif);
	
	return mhl;
	
}
double MVnorm::mhl(const MVnorm *x, const SigmaI &SigI) const{
	double mhl = 0.0;
	gsl_vector *dif = gsl_vector_alloc(_d);
	gsl_vector *tmp = gsl_vector_alloc(_d);
	
	gsl_vector_memcpy(dif, x->getVec());
	gsl_vector_sub(dif, &_vec.vector);
	
	gsl_blas_dsymv(CblasLower, 1.0, SigI.getMat(), dif, 0.0, tmp);
	gsl_blas_ddot(dif, tmp, &mhl);
	
	gsl_vector_free(tmp);
	gsl_vector_free(dif);
	
	return mhl;
	
}

double MVnorm::mhl(const SigmaI &SigI){ // distance to 0
	double mhl = 0.0;
	gsl_vector *tmp = gsl_vector_alloc(_d);
	
	gsl_blas_dsymv(CblasLower, 1.0, SigI.getMat(), &_vec.vector, 0.0, tmp);
	gsl_blas_ddot(&_vec.vector, tmp, &mhl);
	
	gsl_vector_free(tmp);
	
	return mhl;
	
}
double MVnorm::mhl(const SigmaI &SigI) const{ // distance to 0
	double mhl = 0.0;
	gsl_vector *tmp = gsl_vector_alloc(_d);
	
	gsl_blas_dsymv(CblasLower, 1.0, SigI.getMat(), &_vec.vector, 0.0, tmp);
	gsl_blas_ddot(&_vec.vector, tmp, &mhl);
	
	gsl_vector_free(tmp);
	
	return mhl;
	
}


// density function with various formats for theta.  Density calculated up to the constant (2pi)^{-d/2}, since mostly we will be dealing with likelihood rations where it cancels out
double MVnorm::density(const gsl_vector *theta, const SigmaI &SigI){
	double m = mhl(theta, SigI);
	double p = SigI.getSrDet() * exp(-0.5 * m);
	return p;
}
double MVnorm::density(const gsl_vector *theta, const SigmaI &SigI) const{
	double m = mhl(theta, SigI);
	double p = SigI.getSrDet() * exp(-0.5 * m);
	return p;
}
double MVnorm::density(const MVnorm *theta, const SigmaI &SigI){
	double m = mhl(theta, SigI);
	double p = SigI.getSrDet() * exp(-0.5 * m);
	return p;
	
}
double MVnorm::density(const MVnorm *theta, const SigmaI &SigI) const{
	double m = mhl(theta, SigI);
	double p = SigI.getSrDet() * exp(-0.5 * m);
	return p;
	
}


// save function, taking file name, appending by default
void MVnorm::save(const string &fileNam, const char *how){
	FILE *outFl = fopen(fileNam.c_str(), how);
	gsl_vector_fwrite(outFl, &_vec.vector);
	fclose(outFl);
	
}

void MVnorm::save(FILE *fileStr){
	gsl_vector_fwrite(fileStr, &_vec.vector);
	
}


/*
 *	MVnormMu methods
 */


/*
 *	various constructors to be used for initialization, in place of the .init() functions used in the R implementation
 */
MVnormMu::MVnormMu() : MVnorm(){
	_lowLevel = 0;
	_upLevel  = 0;
}

MVnormMu::MVnormMu(const size_t &d, const vector<size_t> &low, const size_t &up) : MVnorm(d){
	_lowLevel = &low;
	_upLevel  = &up;
}


MVnormMu::MVnormMu(gsl_vector *mn, const vector<size_t> &low, const size_t &up) : MVnorm(mn){
	_lowLevel = &low;
	_upLevel  = &up;
}

MVnormMu::MVnormMu(gsl_vector *mn, const size_t &up) : MVnorm(mn){
	_lowLevel = 0;
	_upLevel  = &up;
}

MVnormMu::MVnormMu(gsl_matrix *mn, const size_t &iRw){
	_vec      = gsl_matrix_row(mn, iRw);
	_d        = mn->size2;
	_lowLevel = 0;
	_upLevel  = 0;
}

MVnormMu::MVnormMu(gsl_matrix *mn, const size_t &iRw, const vector<size_t> &low){
	_vec      = gsl_matrix_row(mn, iRw);
	_d        = mn->size2;
	_lowLevel = &low;
	_upLevel  = 0;
}

MVnormMu::MVnormMu(gsl_vector *mn, const gsl_vector *sd, const gsl_rng *r, const vector<size_t> &low, const size_t &up) : MVnorm(mn, sd, r){
	_lowLevel = &low;
	_upLevel  = &up;
}

MVnormMu::MVnormMu(gsl_vector *mn, const gsl_matrix *Sig, const gsl_rng *r, const vector<size_t> &low, const size_t &up) : MVnorm(mn, Sig, r){
	_lowLevel = &low;
	_upLevel  = &up;
}

MVnormMu::MVnormMu(gsl_matrix *mn, const size_t &iRw, const vector<size_t> &low, const size_t &up) : MVnorm(mn, iRw){
	_lowLevel = &low;
	_upLevel  = &up;
}
MVnormMu::MVnormMu(gsl_matrix *mn, const size_t &iRw, const size_t &up) : MVnorm(mn, iRw){
	_lowLevel = 0;
	_upLevel  = &up;
}
MVnormMu::MVnormMu(gsl_matrix *mn, const size_t &iRw, const gsl_vector *sd, const gsl_rng *r, const vector<size_t> &low, const size_t &up) : MVnorm(mn, iRw, sd, r){
	_lowLevel = &low;
	_upLevel  = &up;
}
MVnormMu::MVnormMu(gsl_matrix *mn, const size_t &iRw, const gsl_matrix *Sig, const gsl_rng *r, const vector<size_t> &low, const size_t &up) : MVnorm(mn, iRw, Sig, r){
	_lowLevel = &low;
	_upLevel  = &up;
}

MVnormMu::MVnormMu(const MVnormMu &mu){	//	Copy constructor
	_d   = mu._d;
	_vec = mu._vec;
	
	_lowLevel = mu._lowLevel;
	_upLevel  = mu._upLevel;
}

MVnormMu & MVnormMu::operator=(const MVnormMu &mu){
	_d   = mu._d;
	_vec = mu._vec;
	_lowLevel = mu._lowLevel;
	_upLevel  = mu._upLevel;
	
	return *this;
}

MVnormMu::~MVnormMu(){
	_lowLevel = NULL;
	delete _lowLevel;
	_upLevel = NULL;
	delete _upLevel;
}

/*
 *	overloaded update() methods
 */

void MVnormMu::update(const Grp &dat, const SigmaI &SigIm, const gsl_rng *r){
	gsl_vector *smVec  = gsl_vector_calloc(_d);
	gsl_matrix *SigSum = gsl_matrix_alloc(_d, _d);
	
	for (vector<size_t>::const_iterator it = _lowLevel->begin(); it != _lowLevel->end(); ++it) {
		gsl_vector_add(smVec, dat[*it]->getVec());
	}
	gsl_vector_scale(smVec, 1.0/_lowLevel->size());
	// prepare the scale matrices
	gsl_matrix_memcpy(SigSum, SigIm.getMat());
	gsl_matrix_scale(SigSum, _lowLevel->size());
	gsl_linalg_cholesky_decomp(SigSum);
	gsl_linalg_cholesky_invert(SigSum);
	gsl_linalg_cholesky_decomp(SigSum);
	
	MVgauss(smVec, SigSum, r, &_vec.vector);
	
	gsl_vector_free(smVec);
	gsl_matrix_free(SigSum);
}
void MVnormMu::update(const Grp &dat, const Qgrp &q, const SigmaI &SigIm, const gsl_rng *r){
	gsl_vector *smVec  = gsl_vector_calloc(_d);
	gsl_matrix *SigSum = gsl_matrix_alloc(_d, _d);
	
	double qSum = 0.0;
	
	for (vector<size_t>::const_iterator it = _lowLevel->begin(); it != _lowLevel->end(); ++it) {
		gsl_blas_daxpy(q[*it], dat[*it]->getVec(), smVec);
		qSum += q[*it];
	}
	gsl_vector_scale(smVec, 1.0/qSum);
	
	// prepare the scale matrices
	gsl_matrix_memcpy(SigSum, SigIm.getMat());
	gsl_matrix_scale(SigSum, qSum);
	gsl_linalg_cholesky_decomp(SigSum);
	gsl_linalg_cholesky_invert(SigSum);
	gsl_linalg_cholesky_decomp(SigSum);
	
	MVgauss(smVec, SigSum, r, &_vec.vector);
	
	gsl_vector_free(smVec);
	gsl_matrix_free(SigSum);
}

void MVnormMu::update(const Grp &dat, const SigmaI &SigIm, const SigmaI &SigIp, const gsl_rng *r){
	gsl_vector *smVec  = gsl_vector_calloc(_d);
	gsl_vector *tmpV   = gsl_vector_alloc(_d);
	gsl_matrix *SigSum = gsl_matrix_alloc(_d, _d);
	
	for (vector<size_t>::const_iterator it = _lowLevel->begin(); it != _lowLevel->end(); ++it) {
		gsl_vector_add(smVec, dat[*it]->getVec());
	}
	
	// prepare the scale matrices
	gsl_matrix_memcpy(SigSum, SigIm.getMat());
	gsl_matrix_scale(SigSum, _lowLevel->size());
	gsl_matrix_add(SigSum, SigIp.getMat());
	gsl_linalg_cholesky_decomp(SigSum);
	gsl_linalg_cholesky_invert(SigSum);
	
	// prepare the scaled mean vector
	gsl_blas_dsymv(CblasLower, 1.0, SigIm.getMat(), smVec, 0.0, tmpV);
	gsl_blas_dsymv(CblasLower, 1.0, SigSum, tmpV, 0.0, smVec);
	
	gsl_linalg_cholesky_decomp(SigSum);
	MVgauss(smVec, SigSum, r, &_vec.vector);
	
	gsl_vector_free(tmpV);
	gsl_vector_free(smVec);
	gsl_matrix_free(SigSum);
}
void MVnormMu::update(const Grp &dat, const SigmaI &SigIm, const double &qPr, const SigmaI &SigIp, const gsl_rng *r){
	gsl_vector *smVec  = gsl_vector_calloc(_d);
	gsl_vector *tmpV   = gsl_vector_alloc(_d);
	gsl_matrix *SigSum = gsl_matrix_alloc(_d, _d);
	gsl_matrix *SigPr  = gsl_matrix_alloc(_d, _d);
	
	for (vector<size_t>::const_iterator it = _lowLevel->begin(); it != _lowLevel->end(); ++it) {
		gsl_vector_add(smVec, dat[*it]->getVec());
	}
	
	// prepare the scale matrices
	gsl_matrix_memcpy(SigSum, SigIm.getMat());
	gsl_matrix_scale(SigSum, _lowLevel->size());
	gsl_matrix_memcpy(SigPr, SigIp.getMat());
	gsl_matrix_scale(SigPr, qPr);
	gsl_matrix_add(SigSum, SigPr);
	gsl_linalg_cholesky_decomp(SigSum);
	gsl_linalg_cholesky_invert(SigSum);
	
	// prepare the scaled mean vector
	gsl_blas_dsymv(CblasLower, 1.0, SigIm.getMat(), smVec, 0.0, tmpV);
	gsl_blas_dsymv(CblasLower, 1.0, SigSum, tmpV, 0.0, smVec);
	
	gsl_linalg_cholesky_decomp(SigSum);
	MVgauss(smVec, SigSum, r, &_vec.vector);
	
	gsl_vector_free(tmpV);
	gsl_vector_free(smVec);
	gsl_matrix_free(SigSum);
	gsl_matrix_free(SigPr);
}
void MVnormMu::update(const Grp &dat, const Qgrp &q, const SigmaI &SigIm, const SigmaI &SigIp, const gsl_rng *r){
	gsl_vector *smVec  = gsl_vector_calloc(_d);
	gsl_vector *tmpV   = gsl_vector_alloc(_d);
	gsl_matrix *SigSum = gsl_matrix_alloc(_d, _d);
	
	double qSum = 0.0;
	
	for (vector<size_t>::const_iterator it = _lowLevel->begin(); it != _lowLevel->end(); ++it) {
		gsl_blas_daxpy(q[*it], dat[*it]->getVec(), smVec);
		qSum += q[*it];
	}
	
	// prepare the scale matrices
	gsl_matrix_memcpy(SigSum, SigIm.getMat());
	gsl_matrix_scale(SigSum, qSum);
	gsl_matrix_add(SigSum, SigIp.getMat());
	gsl_linalg_cholesky_decomp(SigSum);
	gsl_linalg_cholesky_invert(SigSum);
	
	
	// prepare the scaled mean vector
	gsl_blas_dsymv(CblasLower, 1.0, SigIm.getMat(), smVec, 0.0, tmpV);
	gsl_blas_dsymv(CblasLower, 1.0, SigSum, tmpV, 0.0, smVec);
	
	gsl_linalg_cholesky_decomp(SigSum);
	MVgauss(smVec, SigSum, r, &_vec.vector);
	
	gsl_vector_free(tmpV);
	gsl_vector_free(smVec);
	gsl_matrix_free(SigSum);
}
void MVnormMu::update(const Grp &dat, const Qgrp &q, const SigmaI &SigIm, const double &qPr, const SigmaI &SigIp, const gsl_rng *r){
	gsl_vector *smVec  = gsl_vector_calloc(_d);
	gsl_vector *tmpV   = gsl_vector_alloc(_d);
	gsl_matrix *SigSum = gsl_matrix_alloc(_d, _d);
	gsl_matrix *SigPr  = gsl_matrix_alloc(_d, _d);
	
	double qSum = 0.0;
	
	for (vector<size_t>::const_iterator it = _lowLevel->begin(); it != _lowLevel->end(); ++it) {
		gsl_blas_daxpy(q[*it], dat[*it]->getVec(), smVec);
		qSum += q[*it];
	}
	
	// prepare the scale matrices
	gsl_matrix_memcpy(SigSum, SigIm.getMat());
	gsl_matrix_scale(SigSum, qSum);
	gsl_matrix_memcpy(SigPr, SigIp.getMat());
	gsl_matrix_scale(SigPr, qPr);
	gsl_matrix_add(SigSum, SigPr);
	gsl_linalg_cholesky_decomp(SigSum);
	gsl_linalg_cholesky_invert(SigSum);
	
	
	// prepare the scaled mean vector
	gsl_blas_dsymv(CblasLower, 1.0, SigIm.getMat(), smVec, 0.0, tmpV);
	gsl_blas_dsymv(CblasLower, 1.0, SigSum, tmpV, 0.0, smVec);
	
	gsl_linalg_cholesky_decomp(SigSum);
	MVgauss(smVec, SigSum, r, &_vec.vector);
	
	gsl_vector_free(tmpV);
	gsl_vector_free(smVec);
	gsl_matrix_free(SigSum);
	gsl_matrix_free(SigPr);
}

void MVnormMu::update(const Grp &dat, const SigmaI &SigIm, const Grp &muPr, const SigmaI &SigIp, const gsl_rng *r){
	gsl_vector *smVec  = gsl_vector_calloc(_d);
	gsl_vector *tmpV   = gsl_vector_alloc(_d);
	gsl_matrix *SigSum = gsl_matrix_alloc(_d, _d);
	
	for (vector<size_t>::const_iterator it = _lowLevel->begin(); it != _lowLevel->end(); ++it) {
		gsl_vector_add(smVec, dat[*it]->getVec());
	}
	
	// prepare the scale matrices
	gsl_matrix_memcpy(SigSum, SigIm.getMat());
	gsl_matrix_scale(SigSum, _lowLevel->size());
	gsl_matrix_add(SigSum, SigIp.getMat());
	gsl_linalg_cholesky_decomp(SigSum);
	gsl_linalg_cholesky_invert(SigSum);
	
	// prepare the scaled mean vector
	gsl_blas_dsymv(CblasLower, 1.0, SigIm.getMat(), smVec, 0.0, tmpV);
	gsl_blas_dsymv(CblasLower, 1.0, SigIp.getMat(), muPr[*_upLevel]->getVec(), 1.0, tmpV);
	gsl_blas_dsymv(CblasLower, 1.0, SigSum, tmpV, 0.0, smVec);
	
	gsl_linalg_cholesky_decomp(SigSum);
	MVgauss(smVec, SigSum, r, &_vec.vector);
	
	gsl_vector_free(tmpV);
	gsl_vector_free(smVec);
	gsl_matrix_free(SigSum);
}
void MVnormMu::update(const Grp &dat, const SigmaI &SigIm, const Grp &muPr, const double &qPr, const SigmaI &SigIp, const gsl_rng *r){
	gsl_vector *smVec  = gsl_vector_calloc(_d);
	gsl_vector *tmpV   = gsl_vector_alloc(_d);
	gsl_matrix *SigSum = gsl_matrix_alloc(_d, _d);
	gsl_matrix *SigPr  = gsl_matrix_alloc(_d, _d);
	
	for (vector<size_t>::const_iterator it = _lowLevel->begin(); it != _lowLevel->end(); ++it) {
		gsl_vector_add(smVec, dat[*it]->getVec());
	}
	
	// prepare the scale matrices
	gsl_matrix_memcpy(SigSum, SigIm.getMat());
	gsl_matrix_scale(SigSum, _lowLevel->size());
	gsl_matrix_memcpy(SigPr, SigIp.getMat());
	gsl_matrix_scale(SigPr, qPr);
	gsl_matrix_add(SigSum, SigPr);
	gsl_linalg_cholesky_decomp(SigSum);
	gsl_linalg_cholesky_invert(SigSum);
	
	// prepare the scaled mean vector
	gsl_blas_dsymv(CblasLower, 1.0, SigIm.getMat(), smVec, 0.0, tmpV);
	gsl_blas_dsymv(CblasLower, 1.0, SigPr, muPr[*_upLevel]->getVec(), 1.0, tmpV);
	gsl_blas_dsymv(CblasLower, 1.0, SigSum, tmpV, 0.0, smVec);
	
	gsl_linalg_cholesky_decomp(SigSum);
	MVgauss(smVec, SigSum, r, &_vec.vector);
	
	gsl_vector_free(tmpV);
	gsl_vector_free(smVec);
	gsl_matrix_free(SigSum);
	gsl_matrix_free(SigPr);
}
void MVnormMu::update(const Grp &dat, const Qgrp &q, const SigmaI &SigIm, const Grp &muPr, const SigmaI &SigIp, const gsl_rng *r){
	gsl_vector *smVec  = gsl_vector_calloc(_d);
	gsl_vector *tmpV   = gsl_vector_alloc(_d);
	gsl_matrix *SigSum = gsl_matrix_alloc(_d, _d);
	
	double qSum = 0.0;
	
	for (vector<size_t>::const_iterator it = _lowLevel->begin(); it != _lowLevel->end(); ++it) {
		gsl_blas_daxpy(q[*it], dat[*it]->getVec(), smVec);
		qSum += q[*it];
	}
	
	// prepare the scale matrices
	gsl_matrix_memcpy(SigSum, SigIm.getMat());
	gsl_matrix_scale(SigSum, qSum);
	gsl_matrix_add(SigSum, SigIp.getMat());
	gsl_linalg_cholesky_decomp(SigSum);
	gsl_linalg_cholesky_invert(SigSum);
	
	
	// prepare the scaled mean vector
	gsl_blas_dsymv(CblasLower, 1.0, SigIm.getMat(), smVec, 0.0, tmpV);
	gsl_blas_dsymv(CblasLower, 1.0, SigIp.getMat(), muPr[*_upLevel]->getVec(), 1.0, tmpV);
	gsl_blas_dsymv(CblasLower, 1.0, SigSum, tmpV, 0.0, smVec);
	
	gsl_linalg_cholesky_decomp(SigSum);
	MVgauss(smVec, SigSum, r, &_vec.vector);
	
	gsl_vector_free(tmpV);
	gsl_vector_free(smVec);
	gsl_matrix_free(SigSum);
}

void MVnormMu::update(const Grp &dat, const Qgrp &q, const SigmaI &SigIm, const Grp &muPr, const double &qPr, const SigmaI &SigIp, const gsl_rng *r){
	gsl_vector *smVec  = gsl_vector_calloc(_d);
	gsl_vector *tmpV   = gsl_vector_alloc(_d);
	gsl_matrix *SigSum = gsl_matrix_alloc(_d, _d);
	gsl_matrix *SigPr  = gsl_matrix_alloc(_d, _d);
	
	double qSum = 0.0;
	
	for (vector<size_t>::const_iterator it = _lowLevel->begin(); it != _lowLevel->end(); ++it) {
		gsl_blas_daxpy(q[*it], dat[*it]->getVec(), smVec);
		qSum += q[*it];
	}
	
	// prepare the scale matrices
	gsl_matrix_memcpy(SigSum, SigIm.getMat());
	gsl_matrix_scale(SigSum, qSum);
	gsl_matrix_memcpy(SigPr, SigIp.getMat());
	gsl_matrix_scale(SigPr, qPr);
	gsl_matrix_add(SigSum, SigPr);
	gsl_linalg_cholesky_decomp(SigSum);
	gsl_linalg_cholesky_invert(SigSum);
	
	// prepare the scaled mean vector
	gsl_blas_dsymv(CblasLower, 1.0, SigIm.getMat(), smVec, 0.0, tmpV);
	gsl_blas_dsymv(CblasLower, 1.0, SigPr, muPr[*_upLevel]->getVec(), 1.0, tmpV);
	gsl_blas_dsymv(CblasLower, 1.0, SigSum, tmpV, 0.0, smVec);
	
	gsl_linalg_cholesky_decomp(SigSum);
	MVgauss(smVec, SigSum, r, &_vec.vector);
	
	gsl_vector_free(tmpV);
	gsl_vector_free(smVec);
	gsl_matrix_free(SigSum);
	gsl_matrix_free(SigPr);
}

/*
 *	MVnormMuPEX methods
 */
MVnormMuPEX::MVnormMuPEX(gsl_matrix *mn, const size_t &iRw, const vector<size_t> &low, const size_t &up, Apex &A, gsl_matrix *tSigIAt) : MVnormMu(mn, iRw, low, up) {
	_A = &A;
	_tSAprod = gsl_matrix_submatrix(tSigIAt, 0, 0, (_A->getMat())->size1, (_A->getMat())->size2);
}

MVnormMuPEX::MVnormMuPEX(gsl_matrix *mn, const size_t &iRw, const gsl_vector *sd, const gsl_rng *r, const vector<size_t> &low, const size_t &up, Apex &A, gsl_matrix *tSigIAt) : MVnormMu(mn, iRw, sd, r, low, up){
	_A = &A;
	_tSAprod = gsl_matrix_submatrix(tSigIAt, 0, 0, (_A->getMat())->size1, (_A->getMat())->size2);
}


MVnormMuPEX::MVnormMuPEX(const MVnormMuPEX &mu){
	_d        = mu._d;
	_vec      = mu._vec;
	_lowLevel = mu._lowLevel;
	_upLevel  = mu._upLevel;
	_A        = mu._A;
	_tSAprod  = mu._tSAprod;
}
MVnormMuPEX & MVnormMuPEX::operator=(const MVnormMuPEX &mu){
	_d        = mu._d;
	_vec      = mu._vec;
	_lowLevel = mu._lowLevel;
	_upLevel  = mu._upLevel;
	_A        = mu._A;
	_tSAprod  = mu._tSAprod;
	
	return *this;
}

void MVnormMuPEX::update(const Grp &dat, const SigmaI &SigIm, const SigmaI &SigIp, const gsl_rng *r){
	gsl_vector *smVec  = gsl_vector_calloc(_d);
	gsl_vector *tmpV   = gsl_vector_alloc(_d);
	gsl_matrix *SigSum = gsl_matrix_alloc(_d, _d);
	
	for (vector<size_t>::const_iterator it = _lowLevel->begin(); it != _lowLevel->end(); ++it) {
		gsl_vector_add(smVec, dat[*it]->getVec());
	}
	
	// prepare the scale matrices
	
	gsl_blas_dgemm(CblasNoTrans, CblasTrans, _lowLevel->size(), _A->getMat(), &_tSAprod.matrix, 0.0, SigSum); // getting n_j*_A%*%SigI%*%t(_A) by multiplying _A by t(_tSAprod)
	gsl_matrix_add(SigSum, SigIp.getMat());
	gsl_linalg_cholesky_decomp(SigSum);
	gsl_linalg_cholesky_invert(SigSum);
	
	// prepare the scaled mean vector
	gsl_blas_dgemv(CblasNoTrans, 1.0, &_tSAprod.matrix, smVec, 0.0, tmpV); // again, transposing _tSAprod, but because of the dgemv has the vector on the right, while I need it on the left, I use CblasNoTrans
	gsl_blas_dsymv(CblasLower, 1.0, SigSum, tmpV, 0.0, smVec);
	
	gsl_linalg_cholesky_decomp(SigSum);
	MVgauss(smVec, SigSum, r, &_vec.vector);
	
	gsl_vector_free(tmpV);
	gsl_vector_free(smVec);
	gsl_matrix_free(SigSum);
}

void MVnormMuPEX::update(const Grp &dat, const SigmaI &SigIm, const double &qPr, const SigmaI &SigIp, const gsl_rng *r){
	gsl_vector *smVec  = gsl_vector_calloc(_d);
	gsl_vector *tmpV   = gsl_vector_alloc(_d);
	gsl_matrix *SigSum = gsl_matrix_alloc(_d, _d);
	gsl_matrix *SigPr  = gsl_matrix_alloc(_d, _d);
	
	for (vector<size_t>::const_iterator it = _lowLevel->begin(); it != _lowLevel->end(); ++it) {
		gsl_vector_add(smVec, dat[*it]->getVec());
	}
	
	// prepare the scale matrices
	gsl_blas_dgemm(CblasNoTrans, CblasTrans, _lowLevel->size(), _A->getMat(), &_tSAprod.matrix, 0.0, SigSum); // getting n_j*_A%*%SigI%*%t(_A) by multiplying _A by t(_tSAprod)
	gsl_matrix_memcpy(SigPr, SigIp.getMat());
	gsl_matrix_scale(SigPr, qPr);
	gsl_matrix_add(SigSum, SigPr);
	gsl_linalg_cholesky_decomp(SigSum);
	gsl_linalg_cholesky_invert(SigSum);
	
	// prepare the scaled mean vector
	gsl_blas_dgemv(CblasNoTrans, 1.0, &_tSAprod.matrix, smVec, 0.0, tmpV); // again, transposing _tSAprod, but because of the dgemv has the vector on the right, while I need it on the left, I use CblasNoTrans
	gsl_blas_dsymv(CblasLower, 1.0, SigSum, tmpV, 0.0, smVec);
	
	gsl_linalg_cholesky_decomp(SigSum);
	MVgauss(smVec, SigSum, r, &_vec.vector);
	
	gsl_vector_free(tmpV);
	gsl_vector_free(smVec);
	gsl_matrix_free(SigSum);
	gsl_matrix_free(SigPr);
}
void MVnormMuPEX::update(const Grp &dat, const Qgrp &q, const SigmaI &SigIm, const SigmaI &SigIp, const gsl_rng *r){
	gsl_vector *smVec  = gsl_vector_calloc(_d);
	gsl_vector *tmpV   = gsl_vector_alloc(_d);
	gsl_matrix *SigSum = gsl_matrix_alloc(_d, _d);
	
	double qSum = 0.0;
	for (vector<size_t>::const_iterator it = _lowLevel->begin(); it != _lowLevel->end(); ++it) {
		gsl_blas_daxpy(q[*it], dat[*it]->getVec(), smVec);
		qSum += q[*it];
	}
	
	// prepare the scale matrices
	
	gsl_blas_dgemm(CblasNoTrans, CblasTrans, qSum, _A->getMat(), &_tSAprod.matrix, 0.0, SigSum); // getting sum(q)*_A%*%SigI%*%t(_A) by multiplying _A by t(_tSAprod)
	gsl_matrix_add(SigSum, SigIp.getMat());
	gsl_linalg_cholesky_decomp(SigSum);
	gsl_linalg_cholesky_invert(SigSum);
	
	// prepare the scaled mean vector
	gsl_blas_dgemv(CblasNoTrans, 1.0, &_tSAprod.matrix, smVec, 0.0, tmpV); // again, transposing _tSAprod, but because of the dgemv has the vector on the right, while I need it on the left, I use CblasNoTrans
	gsl_blas_dsymv(CblasLower, 1.0, SigSum, tmpV, 0.0, smVec);
	
	gsl_linalg_cholesky_decomp(SigSum);
	MVgauss(smVec, SigSum, r, &_vec.vector);
	
	gsl_vector_free(tmpV);
	gsl_vector_free(smVec);
	gsl_matrix_free(SigSum);
}
void MVnormMuPEX::update(const Grp &dat, const Qgrp &q, const SigmaI &SigIm, const double &qPr, const SigmaI &SigIp, const gsl_rng *r){
	gsl_vector *smVec  = gsl_vector_calloc(_d);
	gsl_vector *tmpV   = gsl_vector_alloc(_d);
	gsl_matrix *SigSum = gsl_matrix_alloc(_d, _d);
	gsl_matrix *SigPr  = gsl_matrix_alloc(_d, _d);
	
	double qSum = 0.0;
	for (vector<size_t>::const_iterator it = _lowLevel->begin(); it != _lowLevel->end(); ++it) {
		gsl_blas_daxpy(q[*it], dat[*it]->getVec(), smVec);
		qSum += q[*it];
	}
	
	// prepare the scale matrices
	gsl_blas_dgemm(CblasNoTrans, CblasTrans, qSum, _A->getMat(), &_tSAprod.matrix, 0.0, SigSum); // getting sum(q)*_A%*%SigI%*%t(_A) by multiplying _A by t(_tSAprod)
	gsl_matrix_memcpy(SigPr, SigIp.getMat());
	gsl_matrix_scale(SigPr, qPr);
	gsl_matrix_add(SigSum, SigPr);
	gsl_linalg_cholesky_decomp(SigSum);
	gsl_linalg_cholesky_invert(SigSum);
	
	// prepare the scaled mean vector
	gsl_blas_dgemv(CblasNoTrans, 1.0, &_tSAprod.matrix, smVec, 0.0, tmpV); // again, transposing _tSAprod, but because of the dgemv has the vector on the right, while I need it on the left, I use CblasNoTrans
	gsl_blas_dsymv(CblasLower, 1.0, SigSum, tmpV, 0.0, smVec);
	
	gsl_linalg_cholesky_decomp(SigSum);
	MVgauss(smVec, SigSum, r, &_vec.vector);
	
	gsl_vector_free(tmpV);
	gsl_vector_free(smVec);
	gsl_matrix_free(SigSum);
	gsl_matrix_free(SigPr);
}

void MVnormMuPEX::update(const Grp &dat, const SigmaI &SigIm, const Grp &muPr, const SigmaI &SigIp, const gsl_rng *r){
	gsl_vector *smVec  = gsl_vector_calloc(_d);
	gsl_vector *tmpV   = gsl_vector_alloc(_d);
	gsl_matrix *SigSum = gsl_matrix_alloc(_d, _d);
	
	for (vector<size_t>::const_iterator it = _lowLevel->begin(); it != _lowLevel->end(); ++it) {
		gsl_vector_add(smVec, dat[*it]->getVec());
	}
	
	// prepare the scale matrices
	
	gsl_blas_dgemm(CblasNoTrans, CblasTrans, _lowLevel->size(), _A->getMat(), &_tSAprod.matrix, 0.0, SigSum); // getting n_j*_A%*%SigI%*%t(_A) by multiplying _A by t(_tSAprod)
	gsl_matrix_add(SigSum, SigIp.getMat());
	gsl_linalg_cholesky_decomp(SigSum);
	gsl_linalg_cholesky_invert(SigSum);
	
	// prepare the scaled mean vector
	gsl_blas_dgemv(CblasNoTrans, 1.0, &_tSAprod.matrix, smVec, 0.0, tmpV); // again, transposing _tSAprod, but because of the dgemv has the vector on the right, while I need it on the left, I use CblasNoTrans
	gsl_blas_dsymv(CblasLower, 1.0, SigIp.getMat(), muPr[*_upLevel]->getVec(), 1.0, tmpV);
	gsl_blas_dsymv(CblasLower, 1.0, SigSum, tmpV, 0.0, smVec);
	
	gsl_linalg_cholesky_decomp(SigSum);
	MVgauss(smVec, SigSum, r, &_vec.vector);
	
	gsl_vector_free(tmpV);
	gsl_vector_free(smVec);
	gsl_matrix_free(SigSum);
	
}
void MVnormMuPEX::update(const Grp &dat, const SigmaI &SigIm, const Grp &muPr, const double &qPr, const SigmaI &SigIp, const gsl_rng *r){
	gsl_vector *smVec  = gsl_vector_calloc(_d);
	gsl_vector *tmpV   = gsl_vector_alloc(_d);
	gsl_matrix *SigSum = gsl_matrix_alloc(_d, _d);
	gsl_matrix *SigPr  = gsl_matrix_alloc(_d, _d);
	
	for (vector<size_t>::const_iterator it = _lowLevel->begin(); it != _lowLevel->end(); ++it) {
		gsl_vector_add(smVec, dat[*it]->getVec());
	}
	
	// prepare the scale matrices
	gsl_blas_dgemm(CblasNoTrans, CblasTrans, _lowLevel->size(), _A->getMat(), &_tSAprod.matrix, 0.0, SigSum); // getting n_j*_A%*%SigI%*%t(_A) by multiplying _A by t(_tSAprod)
	gsl_matrix_memcpy(SigPr, SigIp.getMat());
	gsl_matrix_scale(SigPr, qPr);
	gsl_matrix_add(SigSum, SigPr);
	gsl_linalg_cholesky_decomp(SigSum);
	gsl_linalg_cholesky_invert(SigSum);
	
	// prepare the scaled mean vector
	gsl_blas_dgemv(CblasNoTrans, 1.0, &_tSAprod.matrix, smVec, 0.0, tmpV); // again, transposing _tSAprod, but because of the dgemv has the vector on the right, while I need it on the left, I use CblasNoTrans
	gsl_blas_dsymv(CblasLower, 1.0, SigPr, muPr[*_upLevel]->getVec(), 1.0, tmpV);
	gsl_blas_dsymv(CblasLower, 1.0, SigSum, tmpV, 0.0, smVec);
	
	gsl_linalg_cholesky_decomp(SigSum);
	MVgauss(smVec, SigSum, r, &_vec.vector);
	
	gsl_vector_free(tmpV);
	gsl_vector_free(smVec);
	gsl_matrix_free(SigSum);
	gsl_matrix_free(SigPr);
}
void MVnormMuPEX::update(const Grp &dat, const Qgrp &q, const SigmaI &SigIm, const Grp &muPr, const SigmaI &SigIp, const gsl_rng *r){
	gsl_vector *smVec  = gsl_vector_calloc(_d);
	gsl_vector *tmpV   = gsl_vector_alloc(_d);
	gsl_matrix *SigSum = gsl_matrix_alloc(_d, _d);
	
	double qSum = 0.0;
	for (vector<size_t>::const_iterator it = _lowLevel->begin(); it != _lowLevel->end(); ++it) {
		gsl_blas_daxpy(q[*it], dat[*it]->getVec(), smVec);
		qSum += q[*it];
	}
	
	// prepare the scale matrices
	
	gsl_blas_dgemm(CblasNoTrans, CblasTrans, qSum, _A->getMat(), &_tSAprod.matrix, 0.0, SigSum); // getting sum(q)*_A%*%SigI%*%t(_A) by multiplying _A by t(_tSAprod)
	gsl_matrix_add(SigSum, SigIp.getMat());
	gsl_linalg_cholesky_decomp(SigSum);
	gsl_linalg_cholesky_invert(SigSum);
	
	// prepare the scaled mean vector
	gsl_blas_dgemv(CblasNoTrans, 1.0, &_tSAprod.matrix, smVec, 0.0, tmpV); // again, transposing _tSAprod, but because of the dgemv has the vector on the right, while I need it on the left, I use CblasNoTrans
	gsl_blas_dsymv(CblasLower, 1.0, SigIp.getMat(), muPr[*_upLevel]->getVec(), 1.0, tmpV);
	gsl_blas_dsymv(CblasLower, 1.0, SigSum, tmpV, 0.0, smVec);
	
	gsl_linalg_cholesky_decomp(SigSum);
	MVgauss(smVec, SigSum, r, &_vec.vector);
	
	gsl_vector_free(tmpV);
	gsl_vector_free(smVec);
	gsl_matrix_free(SigSum);
}
void MVnormMuPEX::update(const Grp &dat, const Qgrp &q, const SigmaI &SigIm, const Grp &muPr, const double &qPr, const SigmaI &SigIp, const gsl_rng *r){
	gsl_vector *smVec  = gsl_vector_calloc(_d);
	gsl_vector *tmpV   = gsl_vector_alloc(_d);
	gsl_matrix *SigSum = gsl_matrix_alloc(_d, _d);
	gsl_matrix *SigPr  = gsl_matrix_alloc(_d, _d);
	
	double qSum = 0.0;
	for (vector<size_t>::const_iterator it = _lowLevel->begin(); it != _lowLevel->end(); ++it) {
		gsl_blas_daxpy(q[*it], dat[*it]->getVec(), smVec);
		qSum += q[*it];
	}
	
	// prepare the scale matrices
	gsl_blas_dgemm(CblasNoTrans, CblasTrans, qSum, _A->getMat(), &_tSAprod.matrix, 0.0, SigSum); // getting sum(q)*_A%*%SigI%*%t(_A) by multiplying _A by t(_tSAprod)
	gsl_matrix_memcpy(SigPr, SigIp.getMat());
	gsl_matrix_scale(SigPr, qPr);
	gsl_matrix_add(SigSum, SigPr);
	gsl_linalg_cholesky_decomp(SigSum);
	gsl_linalg_cholesky_invert(SigSum);
	
	// prepare the scaled mean vector
	gsl_blas_dgemv(CblasNoTrans, 1.0, &_tSAprod.matrix, smVec, 0.0, tmpV); // again, transposing _tSAprod, but because of the dgemv has the vector on the right, while I need it on the left, I use CblasNoTrans
	gsl_blas_dsymv(CblasLower, 1.0, SigPr, muPr[*_upLevel]->getVec(), 1.0, tmpV);
	gsl_blas_dsymv(CblasLower, 1.0, SigSum, tmpV, 0.0, smVec);
	
	gsl_linalg_cholesky_decomp(SigSum);
	MVgauss(smVec, SigSum, r, &_vec.vector);
	
	gsl_vector_free(tmpV);
	gsl_vector_free(smVec);
	gsl_matrix_free(SigSum);
	gsl_matrix_free(SigPr);
}

/*
 *	MVnormBetaPEX methods
 */

MVnormBetaPEX::MVnormBetaPEX(const gsl_matrix *resp, gsl_matrix *pred, const size_t &iCl, vector<double> &eaFt, const gsl_matrix *Sig, const gsl_rng *r, const size_t &up, gsl_matrix *bet, const size_t &iRw, Apex &A, gsl_matrix *tSigIAt)  : MVnormMuPEX() {
	_A = &A;
	_tSAprod = gsl_matrix_submatrix(tSigIAt, 0, 0, (_A->getMat())->size1, (_A->getMat())->size2);
	_fitted  = gsl_matrix_view_array(eaFt.data(), pred->size1, _d);
	
	_upLevel = &up;
	_d       = resp->size2;
	_N       = pred->size1;
	
	_vec = gsl_matrix_row(bet, iRw);
	
	if (_N != resp->size1) {
		cerr << "ERROR: unequal number of rows in predictor (" << _N << ") and response (" << resp->size1 << ") when initializing MVnormBeta" << endl;
		exit(1);
	}
	else if (resp->size2 != Sig->size1){
		cerr << "ERROR: incompatible matrices *resp (d = " << resp->size2 << ") and *Sig (d = " << Sig->size1 << ") when initializing MVnormBeta" << endl;
		exit(1);
	}
	
	_X = gsl_matrix_column(pred, iCl);
	gsl_blas_ddot(&_X.vector, &_X.vector, &_scale);
	
	gsl_vector *bH    = gsl_vector_alloc(_d);
	gsl_matrix *chl   = gsl_matrix_alloc(_d, _d);
	
	gsl_blas_dgemv(CblasTrans, 1.0/_scale, resp, &_X.vector, 0.0, bH);
	
	gsl_matrix_memcpy(chl, Sig);
	gsl_matrix_scale(chl, 1.0/_scale);
	gsl_linalg_cholesky_decomp(chl);
	
	MVgauss(bH, chl, r, &_vec.vector);
	
	gsl_vector_free(bH);
	gsl_matrix_free(chl);
}
MVnormBetaPEX::MVnormBetaPEX(const size_t &d, gsl_matrix *pred, const size_t &iCl, vector<double> &eaFt, const size_t &up, gsl_matrix *bet, const size_t &iRw, Apex &A, gsl_matrix *tSigIAt) : MVnormMuPEX() {
	_A = &A;
	_tSAprod = gsl_matrix_submatrix(tSigIAt, 0, 0, (_A->getMat())->size1, (_A->getMat())->size2);
	_fitted  = gsl_matrix_view_array(eaFt.data(), pred->size1, d);
	
	_upLevel = &up;
	_d       = d;
	_N       = pred->size1;
	
	_vec = gsl_matrix_row(bet, iRw);
	
	_X = gsl_matrix_column(pred, iCl);
	gsl_blas_ddot(&_X.vector, &_X.vector, &_scale);
	
}

MVnormBetaPEX::MVnormBetaPEX(const MVnormBetaPEX &bet){
	_d        = bet._d;
	_vec      = bet._vec;
	_lowLevel = bet._lowLevel;
	_upLevel  = bet._upLevel;
	_A        = bet._A;
	_tSAprod  = bet._tSAprod;
	_X        = bet._X;
	_scale    = bet._scale;
	_N        = bet._N;
	_fitted   = bet._fitted;
	
}
MVnormBetaPEX & MVnormBetaPEX::operator=(const MVnormBetaPEX &bet){
	_d        = bet._d;
	_vec      = bet._vec;
	_lowLevel = bet._lowLevel;
	_upLevel  = bet._upLevel;
	_A        = bet._A;
	_tSAprod  = bet._tSAprod;
	_X        = bet._X;
	_scale    = bet._scale;
	_N        = bet._N;
	_fitted   = bet._fitted;
	
	return *this;
}

void MVnormBetaPEX::update(const Grp &dat, const SigmaI &SigIm, const SigmaI &SigIp, const gsl_rng *r){
	gsl_matrix *rsd    = gsl_matrix_alloc(_N, _d);
	gsl_matrix *SigSum = gsl_matrix_alloc(_d, _d);
	gsl_vector *tmpV   = gsl_vector_alloc(_d);
	gsl_vector *bH     = gsl_vector_alloc(_d);
	
	gsl_matrix_memcpy(rsd, dat.dMat());
	gsl_matrix_sub(rsd, &_fitted.matrix);
	
	gsl_blas_dgemm(CblasNoTrans, CblasTrans, _scale, _A->getMat(), &_tSAprod.matrix, 0.0, SigSum); // getting n_j*_A%*%SigI%*%t(_A) by multiplying _A by t(_tSAprod)
	gsl_matrix_add(SigSum, SigIp.getMat());
	gsl_linalg_cholesky_decomp(SigSum);
	gsl_linalg_cholesky_invert(SigSum);
	
	gsl_blas_dgemv(CblasTrans, 1.0, rsd, &_X.vector, 0.0, bH);
	gsl_blas_dgemv(CblasNoTrans, 1.0, &_tSAprod.matrix, bH, 0.0, tmpV); // again, transposing _tSAprod, but because of the dgemv has the vector on the right, while I need it on the left, I use CblasNoTrans
	gsl_blas_dsymv(CblasLower, 1.0, SigSum, tmpV, 0.0, bH);
	
	gsl_linalg_cholesky_decomp(SigSum);
	MVgauss(bH, SigSum, r, &_vec.vector);
	
	gsl_matrix_free(rsd);
	gsl_matrix_free(SigSum);
	gsl_vector_free(tmpV);
	gsl_vector_free(bH);
}
void MVnormBetaPEX::update(const Grp &dat, const SigmaI &SigIm, const double &qPr, const SigmaI &SigIp, const gsl_rng *r){
	gsl_matrix *rsd    = gsl_matrix_alloc(_N, _d);
	gsl_matrix *SigSum = gsl_matrix_alloc(_d, _d);
	gsl_matrix *SigPr  = gsl_matrix_alloc(_d, _d);
	gsl_vector *tmpV   = gsl_vector_alloc(_d);
	gsl_vector *bH     = gsl_vector_alloc(_d);
	
	gsl_matrix_memcpy(rsd, dat.dMat());
	gsl_matrix_sub(rsd, &_fitted.matrix);
	
	gsl_blas_dgemm(CblasNoTrans, CblasTrans, _scale, _A->getMat(), &_tSAprod.matrix, 0.0, SigSum); // getting n_j*_A%*%SigI%*%t(_A) by multiplying _A by t(_tSAprod)
	gsl_matrix_memcpy(SigPr, SigIp.getMat());
	gsl_matrix_scale(SigPr, qPr);
	gsl_matrix_add(SigSum, SigPr);
	gsl_linalg_cholesky_decomp(SigSum);
	gsl_linalg_cholesky_invert(SigSum);
	
	gsl_blas_dgemv(CblasTrans, 1.0, rsd, &_X.vector, 0.0, bH);
	gsl_blas_dgemv(CblasNoTrans, 1.0, &_tSAprod.matrix, bH, 0.0, tmpV); // again, transposing _tSAprod, but because of the dgemv has the vector on the right, while I need it on the left, I use CblasNoTrans
	gsl_blas_dsymv(CblasLower, 1.0, SigSum, tmpV, 0.0, bH);
	
	gsl_linalg_cholesky_decomp(SigSum);
	MVgauss(bH, SigSum, r, &_vec.vector);
	
	gsl_matrix_free(rsd);
	gsl_matrix_free(SigSum);
	gsl_matrix_free(SigPr);
	gsl_vector_free(tmpV);
	gsl_vector_free(bH);
}
void MVnormBetaPEX::update(const Grp &dat, const Qgrp &q, const SigmaI &SigIm, const SigmaI &SigIp, const gsl_rng *r){
	gsl_matrix *rsd    = gsl_matrix_alloc(_N, _d);
	gsl_matrix *SigSum = gsl_matrix_alloc(_d, _d);
	gsl_vector *tmpV   = gsl_vector_alloc(_d);
	gsl_vector *bH     = gsl_vector_alloc(_d);
	gsl_vector *XQ     = gsl_vector_alloc(_N);
	double xQx = 0.0;
	
	for (int iDat = 0; iDat < _N; iDat++) {
		gsl_vector_set(XQ, iDat, q[iDat]*gsl_vector_get(&_X.vector, iDat));
	}
	gsl_matrix_memcpy(rsd, dat.dMat());
	gsl_matrix_sub(rsd, &_fitted.matrix);
	
	gsl_blas_ddot(XQ, &_X.vector, &xQx);
	xQx = 1.0/xQx;
	
	gsl_blas_dgemm(CblasNoTrans, CblasTrans, xQx, _A->getMat(), &_tSAprod.matrix, 0.0, SigSum); // getting n_j*_A%*%SigI%*%t(_A) by multiplying _A by t(_tSAprod)
	gsl_matrix_add(SigSum, SigIp.getMat());
	gsl_linalg_cholesky_decomp(SigSum);
	gsl_linalg_cholesky_invert(SigSum);
	
	gsl_blas_dgemv(CblasTrans, 1.0, rsd, XQ, 0.0, bH);
	gsl_blas_dgemv(CblasNoTrans, 1.0, &_tSAprod.matrix, bH, 0.0, tmpV); // again, transposing _tSAprod, but because of the dgemv has the vector on the right, while I need it on the left, I use CblasNoTrans
	gsl_blas_dsymv(CblasLower, 1.0, SigSum, tmpV, 0.0, bH);
	
	gsl_linalg_cholesky_decomp(SigSum);
	MVgauss(bH, SigSum, r, &_vec.vector);
	
	gsl_matrix_free(rsd);
	gsl_matrix_free(SigSum);
	gsl_vector_free(tmpV);
	gsl_vector_free(bH);
	gsl_vector_free(XQ);
}
void MVnormBetaPEX::update(const Grp &dat, const Qgrp &q, const SigmaI &SigIm, const double &qPr, const SigmaI &SigIp, const gsl_rng *r){
	gsl_matrix *rsd    = gsl_matrix_alloc(_N, _d);
	gsl_matrix *SigSum = gsl_matrix_alloc(_d, _d);
	gsl_matrix *SigPr  = gsl_matrix_alloc(_d, _d);
	gsl_vector *tmpV   = gsl_vector_alloc(_d);
	gsl_vector *bH     = gsl_vector_alloc(_d);
	gsl_vector *XQ     = gsl_vector_alloc(_N);
	double xQx = 0.0;
	
	for (int iDat = 0; iDat < _N; iDat++) {
		gsl_vector_set(XQ, iDat, q[iDat]*gsl_vector_get(&_X.vector, iDat));
	}
	
	gsl_matrix_memcpy(rsd, dat.dMat());
	gsl_matrix_sub(rsd, &_fitted.matrix);
	
	gsl_blas_ddot(XQ, &_X.vector, &xQx);
	xQx = 1.0/xQx;
	
	gsl_blas_dgemm(CblasNoTrans, CblasTrans, xQx, _A->getMat(), &_tSAprod.matrix, 0.0, SigSum); // getting n_j*_A%*%SigI%*%t(_A) by multiplying _A by t(_tSAprod)
	gsl_matrix_memcpy(SigPr, SigIp.getMat());
	gsl_matrix_scale(SigPr, qPr);
	gsl_matrix_add(SigSum, SigPr);
	gsl_linalg_cholesky_decomp(SigSum);
	gsl_linalg_cholesky_invert(SigSum);
	
	gsl_blas_dgemv(CblasTrans, 1.0, rsd, XQ, 0.0, bH);
	gsl_blas_dgemv(CblasNoTrans, 1.0, &_tSAprod.matrix, bH, 0.0, tmpV); // again, transposing _tSAprod, but because of the dgemv has the vector on the right, while I need it on the left, I use CblasNoTrans
	gsl_blas_dsymv(CblasLower, 1.0, SigSum, tmpV, 0.0, bH);
	
	gsl_linalg_cholesky_decomp(SigSum);
	MVgauss(bH, SigSum, r, &_vec.vector);
	
	gsl_matrix_free(rsd);
	gsl_matrix_free(SigSum);
	gsl_matrix_free(SigPr);
	gsl_vector_free(tmpV);
	gsl_vector_free(bH);
	gsl_vector_free(XQ);
}

void MVnormBetaPEX::update(const Grp &dat, const SigmaI &SigIm, const Grp &muPr, const SigmaI &SigIp, const gsl_rng *r){
	gsl_matrix *rsd    = gsl_matrix_alloc(_N, _d);
	gsl_matrix *SigSum = gsl_matrix_alloc(_d, _d);
	gsl_vector *tmpV   = gsl_vector_alloc(_d);
	gsl_vector *bH     = gsl_vector_alloc(_d);
	
	gsl_matrix_memcpy(rsd, dat.dMat());
	gsl_matrix_sub(rsd, &_fitted.matrix);
	
	gsl_blas_dgemm(CblasNoTrans, CblasTrans, _scale, _A->getMat(), &_tSAprod.matrix, 0.0, SigSum); // getting n_j*_A%*%SigI%*%t(_A) by multiplying _A by t(_tSAprod)
	gsl_matrix_add(SigSum, SigIp.getMat());
	gsl_linalg_cholesky_decomp(SigSum);
	gsl_linalg_cholesky_invert(SigSum);
	
	gsl_blas_dgemv(CblasTrans, 1.0, rsd, &_X.vector, 0.0, bH);
	gsl_blas_dgemv(CblasNoTrans, 1.0, &_tSAprod.matrix, bH, 0.0, tmpV); // again, transposing _tSAprod, but because of the dgemv has the vector on the right, while I need it on the left, I use CblasNoTrans
	gsl_blas_dsymv(CblasLower, 1.0, SigIp.getMat(), muPr[*_upLevel]->getVec(), 1.0, tmpV);
	
	gsl_blas_dsymv(CblasLower, 1.0, SigSum, tmpV, 0.0, bH);
	
	gsl_linalg_cholesky_decomp(SigSum);
	MVgauss(bH, SigSum, r, &_vec.vector);
	
	gsl_matrix_free(rsd);
	gsl_matrix_free(SigSum);
	gsl_vector_free(tmpV);
	gsl_vector_free(bH);
}
void MVnormBetaPEX::update(const Grp &dat, const SigmaI &SigIm, const Grp &muPr, const double &qPr, const SigmaI &SigIp, const gsl_rng *r){
	gsl_matrix *rsd    = gsl_matrix_alloc(_N, _d);
	gsl_matrix *SigSum = gsl_matrix_alloc(_d, _d);
	gsl_matrix *SigPr  = gsl_matrix_alloc(_d, _d);
	gsl_vector *tmpV   = gsl_vector_alloc(_d);
	gsl_vector *bH     = gsl_vector_alloc(_d);
	
	gsl_matrix_memcpy(rsd, dat.dMat());
	gsl_matrix_sub(rsd, &_fitted.matrix);
	
	gsl_blas_dgemm(CblasNoTrans, CblasTrans, _scale, _A->getMat(), &_tSAprod.matrix, 0.0, SigSum); // getting n_j*_A%*%SigI%*%t(_A) by multiplying _A by t(_tSAprod)
	gsl_matrix_memcpy(SigPr, SigIp.getMat());
	gsl_matrix_scale(SigPr, qPr);
	gsl_matrix_add(SigSum, SigPr);
	gsl_linalg_cholesky_decomp(SigSum);
	gsl_linalg_cholesky_invert(SigSum);
	
	gsl_blas_dgemv(CblasTrans, 1.0, rsd, &_X.vector, 0.0, bH);
	gsl_blas_dgemv(CblasNoTrans, 1.0, &_tSAprod.matrix, bH, 0.0, tmpV); // again, transposing _tSAprod, but because of the dgemv has the vector on the right, while I need it on the left, I use CblasNoTrans
	gsl_blas_dsymv(CblasLower, 1.0, SigPr, muPr[*_upLevel]->getVec(), 1.0, tmpV);
	
	gsl_blas_dsymv(CblasLower, 1.0, SigSum, tmpV, 0.0, bH);
	
	gsl_linalg_cholesky_decomp(SigSum);
	MVgauss(bH, SigSum, r, &_vec.vector);
	
	gsl_matrix_free(rsd);
	gsl_matrix_free(SigSum);
	gsl_matrix_free(SigPr);
	gsl_vector_free(tmpV);
	gsl_vector_free(bH);
}
void MVnormBetaPEX::update(const Grp &dat, const Qgrp &q, const SigmaI &SigIm, const Grp &muPr, const SigmaI &SigIp, const gsl_rng *r){
	gsl_matrix *rsd    = gsl_matrix_alloc(_N, _d);
	gsl_matrix *SigSum = gsl_matrix_alloc(_d, _d);
	gsl_vector *tmpV   = gsl_vector_alloc(_d);
	gsl_vector *bH     = gsl_vector_alloc(_d);
	gsl_vector *XQ     = gsl_vector_alloc(_N);
	double xQx = 0.0;
	
	for (int iDat = 0; iDat < _N; iDat++) {
		gsl_vector_set(XQ, iDat, q[iDat]*gsl_vector_get(&_X.vector, iDat));
	}
	gsl_matrix_memcpy(rsd, dat.dMat());
	gsl_matrix_sub(rsd, &_fitted.matrix);
	
	gsl_blas_ddot(XQ, &_X.vector, &xQx);
	xQx = 1.0/xQx;
	
	gsl_matrix_memcpy(rsd, dat.dMat());
	gsl_matrix_sub(rsd, &_fitted.matrix);
	
	gsl_blas_dgemm(CblasNoTrans, CblasTrans, xQx, _A->getMat(), &_tSAprod.matrix, 0.0, SigSum); // getting n_j*_A%*%SigI%*%t(_A) by multiplying _A by t(_tSAprod)
	gsl_matrix_add(SigSum, SigIp.getMat());
	gsl_linalg_cholesky_decomp(SigSum);
	gsl_linalg_cholesky_invert(SigSum);
	
	gsl_blas_dgemv(CblasTrans, 1.0, rsd, XQ, 0.0, bH);
	gsl_blas_dgemv(CblasNoTrans, 1.0, &_tSAprod.matrix, bH, 0.0, tmpV); // again, transposing _tSAprod, but because of the dgemv has the vector on the right, while I need it on the left, I use CblasNoTrans
	gsl_blas_dsymv(CblasLower, 1.0, SigIp.getMat(), muPr[*_upLevel]->getVec(), 1.0, tmpV);
	
	gsl_blas_dsymv(CblasLower, 1.0, SigSum, tmpV, 0.0, bH);
	
	gsl_linalg_cholesky_decomp(SigSum);
	MVgauss(bH, SigSum, r, &_vec.vector);
	
	gsl_matrix_free(rsd);
	gsl_matrix_free(SigSum);
	gsl_vector_free(tmpV);
	gsl_vector_free(bH);
	gsl_vector_free(XQ);
}
void MVnormBetaPEX::update(const Grp &dat, const Qgrp &q, const SigmaI &SigIm, const Grp &muPr, const double &qPr, const SigmaI &SigIp, const gsl_rng *r){
	gsl_matrix *rsd    = gsl_matrix_alloc(_N, _d);
	gsl_matrix *SigSum = gsl_matrix_alloc(_d, _d);
	gsl_matrix *SigPr  = gsl_matrix_alloc(_d, _d);
	gsl_vector *tmpV   = gsl_vector_alloc(_d);
	gsl_vector *bH     = gsl_vector_alloc(_d);
	gsl_vector *XQ     = gsl_vector_alloc(_N);
	double xQx = 0.0;
	
	for (int iDat = 0; iDat < _N; iDat++) {
		gsl_vector_set(XQ, iDat, q[iDat]*gsl_vector_get(&_X.vector, iDat));
	}
	
	gsl_matrix_memcpy(rsd, dat.dMat());
	gsl_matrix_sub(rsd, &_fitted.matrix);
	
	gsl_blas_ddot(XQ, &_X.vector, &xQx);
	xQx = 1.0/xQx;
	
	gsl_blas_dgemm(CblasNoTrans, CblasTrans, xQx, _A->getMat(), &_tSAprod.matrix, 0.0, SigSum); // getting n_j*_A%*%SigI%*%t(_A) by multiplying _A by t(_tSAprod)
	gsl_matrix_memcpy(SigPr, SigIp.getMat());
	gsl_matrix_scale(SigPr, qPr);
	gsl_matrix_add(SigSum, SigPr);
	gsl_linalg_cholesky_decomp(SigSum);
	gsl_linalg_cholesky_invert(SigSum);
	
	gsl_blas_dgemv(CblasTrans, 1.0, rsd, XQ, 0.0, bH);
	gsl_blas_dgemv(CblasNoTrans, 1.0, &_tSAprod.matrix, bH, 0.0, tmpV); // again, transposing _tSAprod, but because of the dgemv has the vector on the right, while I need it on the left, I use CblasNoTrans
	gsl_blas_dsymv(CblasLower, 1.0, SigPr, muPr[*_upLevel]->getVec(), 1.0, tmpV);
	
	gsl_blas_dsymv(CblasLower, 1.0, SigSum, tmpV, 0.0, bH);
	
	gsl_linalg_cholesky_decomp(SigSum);
	MVgauss(bH, SigSum, r, &_vec.vector);
	
	gsl_matrix_free(rsd);
	gsl_matrix_free(SigSum);
	gsl_matrix_free(SigPr);
	gsl_vector_free(tmpV);
	gsl_vector_free(bH);
	gsl_vector_free(XQ);
}

/*
 *	MVnormMuMiss methods
 */

// Constructors

MVnormMuMiss::MVnormMuMiss(const size_t &d, const vector<size_t> &low, const size_t &up, const vector<size_t> &mis) : MVnormMu(d, low, up){
	_misPhenInd = mis;
}
MVnormMuMiss::MVnormMuMiss(gsl_vector *mn, const vector<size_t> &low, const size_t &up, const vector<size_t> &mis) : MVnormMu(mn, low, up){
	_misPhenInd = mis;
}
MVnormMuMiss::MVnormMuMiss(gsl_vector *mn, const size_t &up, const vector<size_t> &mis) : MVnormMu(mn, up){
	_misPhenInd = mis;
}
MVnormMuMiss::MVnormMuMiss(gsl_vector *mn, const gsl_vector *sd, const gsl_rng *r, const vector<size_t> &low, const size_t &up, const vector<size_t> &mis) : MVnormMu(mn, sd, r, low, up){
	_misPhenInd = mis;
}
MVnormMuMiss::MVnormMuMiss(gsl_vector *mn, const gsl_matrix *Sig, const gsl_rng *r, const vector<size_t> &low, const size_t &up, const vector<size_t> &mis) : MVnormMu(mn, Sig, r, low, up){
	_misPhenInd = mis;
}
MVnormMuMiss::MVnormMuMiss(gsl_matrix *mn, const size_t &iRw, const vector<size_t> &low, const size_t &up, const vector<size_t> &mis) : MVnormMu(mn, iRw, low, up){
	_myInd      = iRw;
	_misPhenInd = mis;
}
MVnormMuMiss::MVnormMuMiss(gsl_matrix *mn, const size_t &iRw, const size_t &up, const vector<size_t> &mis) : MVnormMu(mn, iRw, up){
	_myInd      = iRw;
	_misPhenInd = mis;
}
MVnormMuMiss::MVnormMuMiss(gsl_matrix *mn, const size_t &iRw, const gsl_vector *sd, const gsl_rng *r, const vector<size_t> &low, const size_t &up, const vector<size_t> &mis) : MVnormMu(mn, iRw, sd, r, low, up){
	_myInd      = iRw;
	_misPhenInd = mis;
}
MVnormMuMiss::MVnormMuMiss(gsl_matrix *mn, const size_t &iRw, const gsl_matrix *Sig, const gsl_rng *r, const vector<size_t> &low, const size_t &up, const vector<size_t> &mis) : MVnormMu(mn, iRw, Sig, r, low, up){
	_myInd      = iRw;
	_misPhenInd = mis;
}

// Phenotype imputation, improper prior
void MVnormMuMiss::update(const Grp &mu, const SigmaI &SigIm, const gsl_rng *r){
	// _misPhenInd is a vector that stores the positions of missing phenotypes.  Its size indicates the number of missing phenotypes
	
	// Previous versions treated cases of Nmis == 1 and _d-1 separately.  Here, I let GSL deal with these automatically with vectors and matrices that are actually scalars
	// Keep in mind, though, that each 1x1 matrix inversion via Cholesky is up to 10 times slower than a scalar inversion
	// However, the missing data imputation is not the rate-limiting step in the sampler, so I don't bother optimizing for the special cases
	if (_misPhenInd.size() == _d) {
		gsl_matrix *Sig = gsl_matrix_alloc(_d, _d);
		gsl_matrix_memcpy(Sig, SigIm.getMat());
		gsl_linalg_cholesky_decomp(Sig);
		gsl_linalg_cholesky_invert(Sig);
		gsl_linalg_cholesky_decomp(Sig);
		MVgauss(mu[*_upLevel]->getVec(), Sig, r, &_vec.vector);
		
		gsl_matrix_free(Sig);
	}
	else{
		gsl_vector *xUnkn   = gsl_vector_alloc(_misPhenInd.size());
		gsl_vector *xKnwn   = gsl_vector_alloc(_d - _misPhenInd.size());
		gsl_vector *muUnkn  = gsl_vector_alloc(_misPhenInd.size());
		gsl_vector *muKnwn  = gsl_vector_alloc(_d - _misPhenInd.size());
		gsl_matrix *Sig     = gsl_matrix_alloc(_d, _d);
		gsl_matrix *SigUnkn = gsl_matrix_alloc(_misPhenInd.size(), _misPhenInd.size());
		gsl_matrix *SigKnwn = gsl_matrix_alloc(_d - _misPhenInd.size(), _d - _misPhenInd.size());
		gsl_matrix *Sig12   = gsl_matrix_alloc(_misPhenInd.size(), _d - _misPhenInd.size());
		vector<size_t> prsInd; // to save the positions of non-missing phenotypes
		
		
		gsl_matrix_memcpy(Sig, SigIm.getMat());
		gsl_linalg_cholesky_decomp(Sig);
		gsl_linalg_cholesky_invert(Sig);
		
		/*
		 * when copying over elements of SigI etc, keep in mind that only the lower triangle is stably defined, so we need only assign to the lower triangles
		 */
		int trkMis = 0;
		int trkPrs = 0;
		for (int id = 0; id < _d; ++id) {
			if (id == _misPhenInd[trkMis]) {
				gsl_vector_set(muUnkn, trkMis, (*mu[*_upLevel])[id]);
				for (int iMs = 0; iMs <= trkMis; iMs++) {
					gsl_matrix_set(SigUnkn, trkMis, iMs, gsl_matrix_get(SigIm.getMat(), _misPhenInd[trkMis], _misPhenInd[iMs]));
				}
				if (trkMis < _misPhenInd.size() - 1) { // to insure we are not going past the end of the _misPhenInd vector
					trkMis++;
				}
			}
			else {
				gsl_vector_set(xKnwn, trkPrs, gsl_vector_get(&_vec.vector, id));
				gsl_vector_set(muKnwn, trkPrs, (*mu[*_upLevel])[id]);
				prsInd.push_back(id);
				for (int iPs = 0; iPs <= trkPrs; iPs++) {
					gsl_matrix_set(SigKnwn, trkPrs, iPs, gsl_matrix_get(Sig, prsInd[trkPrs], prsInd[iPs]));
				}
				for (int iRw = 0; iRw < _misPhenInd.size(); iRw++) {
					double elem;
					_misPhenInd[iRw] > id ? elem = gsl_matrix_get(Sig, _misPhenInd[iRw], id) : elem = gsl_matrix_get(Sig, id, _misPhenInd[iRw]); // making sure that we always copy from the lower triangle
					gsl_matrix_set(Sig12, iRw, trkPrs, elem);
				}
				
				trkPrs++;
			}
		}
		
		gsl_linalg_cholesky_decomp(SigUnkn);
		gsl_linalg_cholesky_invert(SigUnkn);
		gsl_linalg_cholesky_decomp(SigUnkn);
		
		gsl_linalg_cholesky_decomp(SigKnwn);
		gsl_linalg_cholesky_invert(SigKnwn);
		
		// comments track the notation in the Wikipedia MV normal page
		gsl_vector_sub(xKnwn, muKnwn); // a - mu2
		gsl_blas_dsymv(CblasLower, 1.0, SigKnwn, xKnwn, 0.0, muKnwn);  // V = Sig_{22}^{-1}(a - mu2)
		gsl_blas_dgemv(CblasNoTrans, 1.0, Sig12, muKnwn, 1.0, muUnkn); // mu1 + Sig12 %*% V
		
		MVgauss(muUnkn, SigUnkn, r, xUnkn);
		
		for (int iM = 0; iM < _misPhenInd.size(); iM++) {
			gsl_vector_set(&_vec.vector, _misPhenInd[iM], gsl_vector_get(xUnkn, iM));
		}
		
		gsl_vector_free(xUnkn);
		gsl_vector_free(xKnwn);
		gsl_vector_free(muUnkn);
		gsl_vector_free(muKnwn);
		gsl_matrix_free(SigUnkn);
		gsl_matrix_free(SigKnwn);
		gsl_matrix_free(Sig12);
		gsl_matrix_free(Sig);

	}
}

void MVnormMuMiss::update(const Grp &mu, const SigmaI &SigIm, const SigmaI &SigIp, const gsl_rng *r){
	// see comments for the normal imputation function
	if (_misPhenInd.size() == _d) {
		gsl_matrix *Sig = gsl_matrix_alloc(_d, _d);
		gsl_matrix_memcpy(Sig, SigIm.getMat());
		gsl_linalg_cholesky_decomp(Sig);
		gsl_linalg_cholesky_invert(Sig);
		gsl_linalg_cholesky_decomp(Sig);
		MVgauss(mu[*_upLevel]->getVec(), Sig, r, &_vec.vector);
		
		gsl_matrix_free(Sig);
	}
	else {
		gsl_vector *scPr     = gsl_vector_alloc(_d);
		gsl_vector *xUnkn    = gsl_vector_alloc(_misPhenInd.size());
		gsl_vector *xKnwn    = gsl_vector_alloc(_d - _misPhenInd.size());
		gsl_vector *muUnkn   = gsl_vector_alloc(_misPhenInd.size());
		gsl_vector *muKnwn   = gsl_vector_alloc(_d - _misPhenInd.size());
		gsl_matrix *Sig      = gsl_matrix_alloc(_d, _d);
		gsl_matrix *SigIunkn = gsl_matrix_alloc(_misPhenInd.size(), _misPhenInd.size());
		gsl_matrix *SigIpr   = gsl_matrix_alloc(_misPhenInd.size(), _misPhenInd.size());
		gsl_matrix *SigKnwn  = gsl_matrix_alloc(_d - _misPhenInd.size(), _d - _misPhenInd.size());
		gsl_matrix *Sig12    = gsl_matrix_alloc(_misPhenInd.size(), _d - _misPhenInd.size());
		vector<size_t> prsInd;
		
		
		gsl_matrix_memcpy(Sig, SigIm.getMat());
		gsl_linalg_cholesky_decomp(Sig);
		gsl_linalg_cholesky_invert(Sig);
		
		gsl_blas_dsymv(CblasLower, 1.0, SigIp.getMat(), mu[*_upLevel]->getVec(), 0.0, scPr);
		
		int trkMis = 0;
		int trkPrs = 0;
		for (int id = 0; id < _d; ++id) {
			if (id == _misPhenInd[trkMis]) {
				gsl_vector_set(muUnkn, trkMis, (*mu[*_upLevel])[id]);
				gsl_vector_set(xUnkn, trkMis, gsl_vector_get(scPr, id));
				for (int iMs = 0; iMs <= trkMis; iMs++) {
					gsl_matrix_set(SigIunkn, trkMis, iMs, gsl_matrix_get(SigIm.getMat(), _misPhenInd[trkMis], _misPhenInd[iMs]));
					gsl_matrix_set(SigIpr, trkMis, iMs, gsl_matrix_get(SigIp.getMat(), _misPhenInd[trkMis], _misPhenInd[iMs]));
				}
				if (trkMis < _misPhenInd.size() - 1) {
					trkMis++;
				}
			}
			else {
				gsl_vector_set(xKnwn, trkPrs, gsl_vector_get(&_vec.vector, id));
				gsl_vector_set(muKnwn, trkPrs, (*mu[*_upLevel])[id]);
				prsInd.push_back(id);
				for (int iPs = 0; iPs <= trkPrs; iPs++) {
					gsl_matrix_set(SigKnwn, trkPrs, iPs, gsl_matrix_get(Sig, prsInd[trkPrs], prsInd[iPs]));
				}
				for (int iRw = 0; iRw < _misPhenInd.size(); iRw++) {
					double elem;
					_misPhenInd[iRw] > id ? elem = gsl_matrix_get(Sig, _misPhenInd[iRw], id) : elem = gsl_matrix_get(Sig, id, _misPhenInd[iRw]); // making sure that we always copy from the lower triangle
					gsl_matrix_set(Sig12, iRw, trkPrs, elem);
				}
				
				trkPrs++;
			}
		}
		
		gsl_linalg_cholesky_decomp(SigKnwn);
		gsl_linalg_cholesky_invert(SigKnwn);
		
		// comments track the notation in the Wikipedia MV normal page
		gsl_vector_sub(xKnwn, muKnwn); // a - mu2
		gsl_blas_dsymv(CblasLower, 1.0, SigKnwn, xKnwn, 0.0, muKnwn);  // V = Sig_{22}^{-1}(a - mu2)
		gsl_blas_dgemv(CblasNoTrans, 1.0, Sig12, muKnwn, 1.0, muUnkn); // mu1 + Sig12 %*% V
		
		gsl_blas_dsymv(CblasLower, 1.0, SigIunkn, muUnkn, 1.0, xUnkn); // xUnkn has the scaled prior values
		
		gsl_matrix_add(SigIunkn, SigIpr);
		gsl_linalg_cholesky_decomp(SigIunkn);
		gsl_linalg_cholesky_invert(SigIunkn);
		
		gsl_blas_dsymv(CblasLower, 1.0, SigIunkn, xUnkn, 0.0, muUnkn);
		
		gsl_linalg_cholesky_decomp(SigIunkn);
		
		MVgauss(muUnkn, SigIunkn, r, xUnkn);
		
		for (int iM = 0; iM < _misPhenInd.size(); iM++) {
			gsl_vector_set(&_vec.vector, _misPhenInd[iM], gsl_vector_get(xUnkn, iM));
		}
		
		gsl_vector_free(scPr);
		gsl_vector_free(xUnkn);
		gsl_vector_free(xKnwn);
		gsl_vector_free(muUnkn);
		gsl_vector_free(muKnwn);
		gsl_matrix_free(SigIunkn);
		gsl_matrix_free(SigIpr);
		gsl_matrix_free(SigKnwn);
		gsl_matrix_free(Sig12);
		gsl_matrix_free(Sig);

	}
}


/*
 *	MVnormBeta methods
 */


/*
 *	various constructors to be used for initialization, in place of the .init() functions used in the R implementation
 */
MVnormBeta::MVnormBeta() : MVnorm(){
	_N = 0;
	_scale = 1.0;
	_upLevel = 0;
}

MVnormBeta::MVnormBeta(const size_t d)  : MVnorm(d){
	_N = 0;
	_scale = 1.0;
	_upLevel = 0;
}

MVnormBeta::MVnormBeta(gsl_vector *b, const gsl_vector *sd, gsl_matrix *pred, const size_t &iCl, const gsl_rng *r) : MVnorm(b, sd, r){
	_X = gsl_matrix_column(pred, iCl);
	_N = pred->size1;
	gsl_blas_ddot(&_X.vector, &_X.vector, &_scale);
	_upLevel = 0;
}
MVnormBeta::MVnormBeta(gsl_vector *b, const gsl_vector *sd, gsl_matrix *pred, const size_t &iCl, const gsl_rng *r, const size_t &up) : MVnorm(b, sd, r){
	_X = gsl_matrix_column(pred, iCl);
	_N = pred->size1;
	gsl_blas_ddot(&_X.vector, &_X.vector, &_scale);
	_upLevel = &up;
}

MVnormBeta::MVnormBeta(gsl_vector *b, const gsl_matrix *Sig, gsl_matrix *pred, const size_t &iCl, const gsl_rng *r) : MVnorm(b, Sig, r){
	_X = gsl_matrix_column(pred, iCl);
	_N = pred->size1;
	gsl_blas_ddot(&_X.vector, &_X.vector, &_scale);
	_upLevel = 0;
}
MVnormBeta::MVnormBeta(gsl_vector *b, const gsl_matrix *Sig, gsl_matrix *pred, const size_t &iCl, const gsl_rng *r, const size_t &up) : MVnorm(b, Sig, r){
	_X = gsl_matrix_column(pred, iCl);
	_N = pred->size1;
	gsl_blas_ddot(&_X.vector, &_X.vector, &_scale);
	_upLevel = &up;
}

MVnormBeta::MVnormBeta(const gsl_matrix *resp, gsl_matrix *pred, const size_t &iCl, const gsl_matrix *Sig, const gsl_rng *r, gsl_matrix *bet, const size_t &iRw){ // NOTE: no-intercept regression for initialization here
	_upLevel = 0;
	_d       = resp->size2;
	_N       = pred->size1;
	
	_vec = gsl_matrix_row(bet, iRw);
	
	if (_N != resp->size1) {
		cerr << "ERROR: unequal number of rows in predictor (" << _N << ") and response (" << resp->size1 << ") when initializing MVnormBeta" << endl;
		exit(1);
	}
	else if (resp->size2 != Sig->size1){
		cerr << "ERROR: incompatible matrices *resp (d = " << resp->size2 << ") and *Sig (d = " << Sig->size1 << ") when initializing MVnormBeta" << endl;
		exit(1);
	}
	
	_X = gsl_matrix_column(pred, iCl);
	gsl_blas_ddot(&_X.vector, &_X.vector, &_scale);
	
	gsl_vector *bH    = gsl_vector_alloc(_d);
	gsl_matrix *chl   = gsl_matrix_alloc(_d, _d);
	
	gsl_blas_dgemv(CblasTrans, 1.0/_scale, resp, &_X.vector, 0.0, bH);
	
	gsl_matrix_memcpy(chl, Sig);
	gsl_matrix_scale(chl, 1.0/_scale);
	gsl_linalg_cholesky_decomp(chl);
	
	MVgauss(bH, chl, r, &_vec.vector);
	
	gsl_vector_free(bH);
	gsl_matrix_free(chl);
}
MVnormBeta::MVnormBeta(const gsl_matrix *resp, gsl_matrix *pred, const size_t &iCl, const gsl_matrix *Sig, const gsl_rng *r, const size_t &up, gsl_matrix *bet, const size_t &iRw){ // NOTE: no-intercept regression for initialization here
	_upLevel = &up;
	_d       = resp->size2;
	_N       = pred->size1;
	
	_vec = gsl_matrix_row(bet, iRw);
	
	if (_N != resp->size1) {
		cerr << "ERROR: unequal number of rows in predictor (" << _N << ") and response (" << resp->size1 << ") when initializing MVnormBeta" << endl;
		exit(1);
	}
	else if (resp->size2 != Sig->size1){
		cerr << "ERROR: incompatible matrices *resp (d = " << resp->size2 << ") and *Sig (d = " << Sig->size1 << ") when initializing MVnormBeta" << endl;
		exit(1);
	}
	
	_X = gsl_matrix_column(pred, iCl);
	gsl_blas_ddot(&_X.vector, &_X.vector, &_scale);
	
	gsl_vector *bH    = gsl_vector_alloc(_d);
	gsl_matrix *chl   = gsl_matrix_alloc(_d, _d);
	
	gsl_blas_dgemv(CblasTrans, 1.0/_scale, resp, &_X.vector, 0.0, bH);
	
	gsl_matrix_memcpy(chl, Sig);
	gsl_matrix_scale(chl, 1.0/_scale);
	gsl_linalg_cholesky_decomp(chl);
	
	MVgauss(bH, chl, r, &_vec.vector);
	
	gsl_vector_free(bH);
	gsl_matrix_free(chl);
}

MVnormBeta::MVnormBeta(const Grp &resp, gsl_matrix *pred, const size_t &iCl, const gsl_matrix *Sig, const gsl_rng *r, gsl_matrix *bet, const size_t &iRw){
	_upLevel = 0;
	_N       = pred->size1;
	_d       = resp.phenD();
	
	_vec = gsl_matrix_row(bet, iRw);
	
	if (_N != resp.Ndata()) {
		cerr << "ERROR: unequal number of rows in predictor (" << _N << ") and response (" << resp.Ndata() << ") when initializing MVnormBeta" << endl;
		exit(1);
	}
	else if (resp.phenD() != Sig->size1){
		cerr << "ERROR: incompatible matrices *resp (d = " << resp.phenD() << ") and *Sig (d = " << Sig->size1 << ") when initializing MVnormBeta" << endl;
		exit(1);
	}
	
	_X = gsl_matrix_column(pred, iCl);
	gsl_blas_ddot(&_X.vector, &_X.vector, &_scale);
	
	gsl_vector *bH   = gsl_vector_alloc(_d);
	gsl_matrix *chl  = gsl_matrix_alloc(_d, _d);
	
	gsl_blas_dgemv(CblasTrans, 1.0/_scale, resp.dMat(), &_X.vector, 0.0, bH);
	
	gsl_matrix_memcpy(chl, Sig);
	gsl_matrix_scale(chl, 1.0/_scale);
	gsl_linalg_cholesky_decomp(chl);
	
	MVgauss(bH, chl, r, &_vec.vector);
	
	gsl_vector_free(bH);
	gsl_matrix_free(chl);
}
MVnormBeta::MVnormBeta(const Grp &resp, gsl_matrix *pred, const size_t &iCl, const gsl_matrix *Sig, const gsl_rng *r, const size_t &up, gsl_matrix *bet, const size_t &iRw){
	_upLevel = &up;
	_N       = pred->size1;
	_d       = resp.phenD();
	
	_vec = gsl_matrix_row(bet, iRw);
	
	if (_N != resp.Ndata()) {
		cerr << "ERROR: unequal number of rows in predictor (" << _N << ") and response (" << resp.Ndata() << ") when initializing MVnormBeta" << endl;
		exit(1);
	}
	else if (resp.phenD() != Sig->size1){
		cerr << "ERROR: incompatible matrices *resp (d = " << resp.phenD() << ") and *Sig (d = " << Sig->size1 << ") when initializing MVnormBeta" << endl;
		exit(1);
	}
	
	_X = gsl_matrix_column(pred, iCl);
	gsl_blas_ddot(&_X.vector, &_X.vector, &_scale);
	
	gsl_vector *bH   = gsl_vector_alloc(_d);
	gsl_matrix *chl  = gsl_matrix_alloc(_d, _d);
	
	gsl_blas_dgemv(CblasTrans, 1.0/_scale, resp.dMat(), &_X.vector, 0.0, bH);
	
	gsl_matrix_memcpy(chl, Sig);
	gsl_matrix_scale(chl, 1.0/_scale);
	gsl_linalg_cholesky_decomp(chl);
	
	MVgauss(bH, chl, r, &_vec.vector);
	
	gsl_vector_free(bH);
	gsl_matrix_free(chl);
}

MVnormBeta::MVnormBeta(gsl_matrix *pred, const size_t &iCl, gsl_matrix *bet, const size_t &iRw){
	_upLevel = 0;
	_N       = pred->size1;
	_d       = bet->size2;
	
	_vec = gsl_matrix_row(bet, iRw);
	
	_X = gsl_matrix_column(pred, iCl);
	gsl_blas_ddot(&_X.vector, &_X.vector, &_scale);
	
}
MVnormBeta::MVnormBeta(gsl_matrix *pred, const size_t &iCl, const size_t &up, gsl_matrix *bet, const size_t &iRw){
	_upLevel = &up;
	_N       = pred->size1;
	_d       = bet->size2;
	
	_vec = gsl_matrix_row(bet, iRw);
	
	_X = gsl_matrix_column(pred, iCl);
	gsl_blas_ddot(&_X.vector, &_X.vector, &_scale);
	
}


MVnormBeta::MVnormBeta(const MVnormBeta &b){	//	Copy constructor
	_d       = b._d;
	_N       = b._N;
	_scale   = b._scale;
	_vec     = b._vec;
	_X       = b._X;
	_upLevel = b._upLevel;
}

MVnormBeta & MVnormBeta::operator=(const MVnormBeta &b){
	_d       = b._d;
	_N       = b._N;
	_scale   = b._scale;
	_vec     = b._vec;
	_X       = b._X;
	_upLevel = b._upLevel;
	
	return *this;
}

MVnormBeta::~MVnormBeta(){
	_upLevel = NULL;
	delete _upLevel;
}

/*
 *	overloaded update() methods
 */

void MVnormBeta::update(const Grp &mu, const SigmaI &SigIb, const gsl_rng *r){
	gsl_matrix *SigSum = gsl_matrix_alloc(_d, _d);
	gsl_vector *tmpV   = gsl_vector_alloc(_d);
	
	gsl_matrix_memcpy(SigSum, SigIb.getMat());
	gsl_matrix_scale(SigSum, _scale);
	gsl_linalg_cholesky_decomp(SigSum);
	gsl_linalg_cholesky_invert(SigSum);
	gsl_linalg_cholesky_decomp(SigSum);
	
	gsl_blas_dgemv(CblasTrans, 1.0/_scale, mu.dMat(), &_X.vector, 0.0, tmpV);
	
	MVgauss(tmpV, SigSum, r, &_vec.vector);
	
	gsl_matrix_free(SigSum);
	gsl_vector_free(tmpV);
}
void MVnormBeta::update(const Grp &dat, const Qgrp &q, const SigmaI &SigIb, const gsl_rng *r){
	gsl_matrix *Sig = gsl_matrix_alloc(_d, _d);
	gsl_vector *XQ  = gsl_vector_alloc(_N);
	gsl_vector *tmp = gsl_vector_alloc(_d);
	double xQx = 0.0;
	
	for (int iDat = 0; iDat < _N; iDat++) {
		gsl_vector_set(XQ, iDat, q[iDat]*gsl_vector_get(&_X.vector, iDat));
	}
	
	gsl_blas_ddot(XQ, &_X.vector, &xQx);
	xQx = 1.0/xQx;
	
	gsl_blas_dgemv(CblasTrans, xQx, dat.dMat(), XQ, 0.0, tmp);
	
	gsl_matrix_memcpy(Sig, SigIb.getMat());
	gsl_linalg_cholesky_decomp(Sig);
	gsl_linalg_cholesky_invert(Sig);
	gsl_matrix_scale(Sig, xQx);
	gsl_linalg_cholesky_decomp(Sig);
	
	MVgauss(tmp, Sig, r, &_vec.vector);
	
	gsl_matrix_free(Sig);
	gsl_vector_free(XQ);
	gsl_vector_free(tmp);
}

void MVnormBeta::update(const Grp &dat, const SigmaI &SigIb, const SigmaI &SigIp, const gsl_rng *r){
	gsl_matrix *SigSum = gsl_matrix_alloc(_d, _d);
	gsl_vector *tmpV   = gsl_vector_alloc(_d);
	gsl_vector *bH     = gsl_vector_alloc(_d);
	
	gsl_matrix_memcpy(SigSum, SigIb.getMat());
	gsl_matrix_scale(SigSum, _scale);
	gsl_matrix_add(SigSum, SigIp.getMat());
	gsl_linalg_cholesky_decomp(SigSum);
	gsl_linalg_cholesky_invert(SigSum);
	
	gsl_blas_dgemv(CblasTrans, 1.0, dat.dMat(), &_X.vector, 0.0, bH);
	gsl_blas_dsymv(CblasLower, 1.0, SigIb.getMat(), bH, 0.0, tmpV);
	gsl_blas_dsymv(CblasLower, 1.0, SigSum, tmpV, 0.0, bH);
	
	gsl_linalg_cholesky_decomp(SigSum);
	MVgauss(bH, SigSum, r, &_vec.vector);
	
	gsl_matrix_free(SigSum);
	gsl_vector_free(tmpV);
	gsl_vector_free(bH);
}
void MVnormBeta::update(const Grp &dat, const SigmaI &SigIb, const double &qPr, const SigmaI &SigIp, const gsl_rng *r){
	gsl_matrix *SigSum = gsl_matrix_alloc(_d, _d);
	gsl_matrix *SigPr  = gsl_matrix_alloc(_d, _d);
	gsl_vector *tmpV   = gsl_vector_alloc(_d);
	gsl_vector *bH     = gsl_vector_alloc(_d);
	
	gsl_matrix_memcpy(SigSum, SigIb.getMat());
	gsl_matrix_scale(SigSum, _scale);
	gsl_matrix_memcpy(SigPr, SigIp.getMat());
	gsl_matrix_scale(SigPr, qPr);
	gsl_matrix_add(SigSum, SigPr);
	gsl_linalg_cholesky_decomp(SigSum);
	gsl_linalg_cholesky_invert(SigSum);
	
	gsl_blas_dgemv(CblasTrans, 1.0, dat.dMat(), &_X.vector, 0.0, bH);
	gsl_blas_dsymv(CblasLower, 1.0, SigIb.getMat(), bH, 0.0, tmpV);
	gsl_blas_dsymv(CblasLower, 1.0, SigSum, tmpV, 0.0, bH);
	
	gsl_linalg_cholesky_decomp(SigSum);
	MVgauss(bH, SigSum, r, &_vec.vector);
	
	gsl_matrix_free(SigSum);
	gsl_matrix_free(SigPr);
	gsl_vector_free(tmpV);
	gsl_vector_free(bH);
}
void MVnormBeta::update(const Grp &dat, const Qgrp &q, const SigmaI &SigIb, const SigmaI &SigIp, const gsl_rng *r){
	gsl_matrix *SigSum = gsl_matrix_alloc(_d, _d);
	gsl_vector *XQ     = gsl_vector_alloc(_N);
	gsl_vector *tmpV   = gsl_vector_alloc(_d);
	gsl_vector *bH     = gsl_vector_alloc(_d);
	double xQx = 0.0;
	
	for (int iDat = 0; iDat < _N; iDat++) {
		gsl_vector_set(XQ, iDat, q[iDat]*gsl_vector_get(&_X.vector, iDat));
	}
	
	gsl_blas_ddot(XQ, &_X.vector, &xQx);
	
	gsl_matrix_memcpy(SigSum, SigIb.getMat());
	gsl_matrix_scale(SigSum, xQx);
	gsl_matrix_add(SigSum, SigIp.getMat());
	gsl_linalg_cholesky_decomp(SigSum);
	gsl_linalg_cholesky_invert(SigSum);
	
	gsl_blas_dgemv(CblasTrans, 1.0, dat.dMat(), XQ, 0.0, bH);
	gsl_blas_dsymv(CblasLower, 1.0, SigIb.getMat(), bH, 0.0, tmpV);
	gsl_blas_dsymv(CblasLower, 1.0, SigSum, tmpV, 0.0, bH);
	
	gsl_linalg_cholesky_decomp(SigSum);
	MVgauss(bH, SigSum, r, &_vec.vector);
	
	gsl_matrix_free(SigSum);
	gsl_vector_free(XQ);
	gsl_vector_free(tmpV);
	gsl_vector_free(bH);
}
void MVnormBeta::update(const Grp &dat, const Qgrp &q, const SigmaI &SigIb, const double &qPr, const SigmaI &SigIp, const gsl_rng *r){
	gsl_matrix *SigSum = gsl_matrix_alloc(_d, _d);
	gsl_matrix *SigPr  = gsl_matrix_alloc(_d, _d);
	gsl_vector *XQ     = gsl_vector_alloc(_N);
	gsl_vector *tmpV   = gsl_vector_alloc(_d);
	gsl_vector *bH     = gsl_vector_alloc(_d);
	double xQx = 0.0;
	
	for (int iDat = 0; iDat < _N; iDat++) {
		gsl_vector_set(XQ, iDat, q[iDat]*gsl_vector_get(&_X.vector, iDat));
	}
	
	gsl_blas_ddot(XQ, &_X.vector, &xQx);
	
	gsl_matrix_memcpy(SigSum, SigIb.getMat());
	gsl_matrix_scale(SigSum, xQx);
	gsl_matrix_memcpy(SigPr, SigIp.getMat());
	gsl_matrix_scale(SigPr, qPr);
	gsl_matrix_add(SigSum, SigPr);
	gsl_linalg_cholesky_decomp(SigSum);
	gsl_linalg_cholesky_invert(SigSum);
	
	gsl_blas_dgemv(CblasTrans, 1.0, dat.dMat(), XQ, 0.0, bH);
	gsl_blas_dsymv(CblasLower, 1.0, SigIb.getMat(), bH, 0.0, tmpV);
	gsl_blas_dsymv(CblasLower, 1.0, SigSum, tmpV, 0.0, bH);
	
	gsl_linalg_cholesky_decomp(SigSum);
	MVgauss(bH, SigSum, r, &_vec.vector);
	
	gsl_matrix_free(SigSum);
	gsl_matrix_free(SigPr);
	gsl_vector_free(XQ);
	gsl_vector_free(tmpV);
	gsl_vector_free(bH);
}

void MVnormBeta::update(const Grp &dat, const SigmaI &SigIb, const Grp &muPr, const SigmaI &SigIp, const gsl_rng *r){
	gsl_matrix *SigSum = gsl_matrix_alloc(_d, _d);
	gsl_vector *tmpV   = gsl_vector_alloc(_d);
	gsl_vector *bH     = gsl_vector_alloc(_d);
	
	gsl_matrix_memcpy(SigSum, SigIb.getMat());
	gsl_matrix_scale(SigSum, _scale);
	gsl_matrix_add(SigSum, SigIp.getMat());
	gsl_linalg_cholesky_decomp(SigSum);
	gsl_linalg_cholesky_invert(SigSum);
	
	gsl_blas_dgemv(CblasTrans, 1.0, dat.dMat(), &_X.vector, 0.0, bH);
	gsl_blas_dsymv(CblasLower, 1.0, SigIb.getMat(), bH, 0.0, tmpV);
	gsl_blas_dsymv(CblasLower, 1.0, SigIp.getMat(), muPr[*_upLevel]->getVec(), 1.0, tmpV);
	
	gsl_blas_dsymv(CblasLower, 1.0, SigSum, tmpV, 0.0, bH);
	
	gsl_linalg_cholesky_decomp(SigSum);
	MVgauss(bH, SigSum, r, &_vec.vector);
	
	gsl_matrix_free(SigSum);
	gsl_vector_free(tmpV);
	gsl_vector_free(bH);
}
void MVnormBeta::update(const Grp &dat, const SigmaI &SigIb, const Grp &muPr, const double &qPr, const SigmaI &SigIp, const gsl_rng *r){
	gsl_matrix *SigSum = gsl_matrix_alloc(_d, _d);
	gsl_matrix *SigPr  = gsl_matrix_alloc(_d, _d);
	gsl_vector *tmpV   = gsl_vector_alloc(_d);
	gsl_vector *bH     = gsl_vector_alloc(_d);
	
	gsl_matrix_memcpy(SigSum, SigIb.getMat());
	gsl_matrix_scale(SigSum, _scale);
	gsl_matrix_memcpy(SigPr, SigIp.getMat());
	gsl_matrix_scale(SigPr, qPr);
	gsl_matrix_add(SigSum, SigPr);
	gsl_linalg_cholesky_decomp(SigSum);
	gsl_linalg_cholesky_invert(SigSum);
	
	gsl_blas_dgemv(CblasTrans, 1.0, dat.dMat(), &_X.vector, 0.0, bH);
	gsl_blas_dsymv(CblasLower, 1.0, SigIb.getMat(), bH, 0.0, tmpV);
	gsl_blas_dsymv(CblasLower, 1.0, SigPr, muPr[*_upLevel]->getVec(), 1.0, tmpV);
	
	gsl_blas_dsymv(CblasLower, 1.0, SigSum, tmpV, 0.0, bH);
	
	gsl_linalg_cholesky_decomp(SigSum);
	MVgauss(bH, SigSum, r, &_vec.vector);
	
	gsl_matrix_free(SigSum);
	gsl_matrix_free(SigPr);
	gsl_vector_free(tmpV);
	gsl_vector_free(bH);
}
void MVnormBeta::update(const Grp &dat, const Qgrp &q, const SigmaI &SigIb, const Grp &muPr, const SigmaI &SigIp, const gsl_rng *r){
	gsl_matrix *SigSum = gsl_matrix_alloc(_d, _d);
	gsl_vector *XQ     = gsl_vector_alloc(_N);
	gsl_vector *tmpV   = gsl_vector_alloc(_d);
	gsl_vector *bH     = gsl_vector_alloc(_d);
	double xQx = 0.0;
	
	for (int iDat = 0; iDat < _N; iDat++) {
		gsl_vector_set(XQ, iDat, q[iDat]*gsl_vector_get(&_X.vector, iDat));
	}
	
	gsl_blas_ddot(XQ, &_X.vector, &xQx);
	
	gsl_matrix_memcpy(SigSum, SigIb.getMat());
	gsl_matrix_scale(SigSum, xQx);
	gsl_matrix_add(SigSum, SigIp.getMat());
	gsl_linalg_cholesky_decomp(SigSum);
	gsl_linalg_cholesky_invert(SigSum);
	
	gsl_blas_dgemv(CblasTrans, 1.0, dat.dMat(), XQ, 0.0, bH);
	gsl_blas_dsymv(CblasLower, 1.0, SigIb.getMat(), bH, 0.0, tmpV);
	gsl_blas_dsymv(CblasLower, 1.0, SigIp.getMat(), muPr[*_upLevel]->getVec(), 1.0, tmpV);
	
	gsl_blas_dsymv(CblasLower, 1.0, SigSum, tmpV, 0.0, bH);
	
	gsl_linalg_cholesky_decomp(SigSum);
	MVgauss(bH, SigSum, r, &_vec.vector);
	
	gsl_matrix_free(SigSum);
	gsl_vector_free(XQ);
	gsl_vector_free(tmpV);
	gsl_vector_free(bH);
}
void MVnormBeta::update(const Grp &dat, const Qgrp &q, const SigmaI &SigIb, const Grp &muPr, const double &qPr, const SigmaI &SigIp, const gsl_rng *r){
	gsl_matrix *SigSum = gsl_matrix_alloc(_d, _d);
	gsl_matrix *SigPr  = gsl_matrix_alloc(_d, _d);
	gsl_vector *XQ     = gsl_vector_alloc(_N);
	gsl_vector *tmpV   = gsl_vector_alloc(_d);
	gsl_vector *bH     = gsl_vector_alloc(_d);
	double xQx = 0.0;
	
	for (size_t iDat = 0; iDat < _N; iDat++) {
		gsl_vector_set(XQ, iDat, q[iDat]*gsl_vector_get(&_X.vector, iDat));
	}
	
	gsl_blas_ddot(XQ, &_X.vector, &xQx);
	
	gsl_matrix_memcpy(SigSum, SigIb.getMat());
	gsl_matrix_scale(SigSum, xQx);
	gsl_matrix_memcpy(SigPr, SigIp.getMat());
	gsl_matrix_scale(SigPr, qPr);
	gsl_matrix_add(SigSum, SigPr);
	gsl_linalg_cholesky_decomp(SigSum);
	gsl_linalg_cholesky_invert(SigSum);
	
	gsl_blas_dgemv(CblasTrans, 1.0, dat.dMat(), XQ, 0.0, bH);
	gsl_blas_dsymv(CblasLower, 1.0, SigIb.getMat(), bH, 0.0, tmpV);
	gsl_blas_dsymv(CblasLower, 1.0, SigPr, muPr[*_upLevel]->getVec(), 1.0, tmpV);
	
	gsl_blas_dsymv(CblasLower, 1.0, SigSum, tmpV, 0.0, bH);
	
	gsl_linalg_cholesky_decomp(SigSum);
	MVgauss(bH, SigSum, r, &_vec.vector);
	
	gsl_matrix_free(SigSum);
	gsl_matrix_free(SigPr);
	gsl_vector_free(XQ);
	gsl_vector_free(tmpV);
	gsl_vector_free(bH);
}

/*
 *	MVnormBetaFt methods
 */


MVnormBetaFt::MVnormBetaFt(const MVnormBetaFt &b){
	_d       = b._d;
	_N       = b._N;
	_scale   = b._scale;
	_vec     = b._vec;
	_X       = b._X;
	_fitted  = b._fitted;
	_upLevel = b._upLevel;
}

MVnormBetaFt & MVnormBetaFt::operator=(const MVnormBetaFt &b){
	_d       = b._d;
	_N       = b._N;
	_scale   = b._scale;
	_vec     = b._vec;
	_X       = b._X;
	_fitted  = b._fitted;
	_upLevel = b._upLevel;
	
	return *this;
}

void MVnormBetaFt::update(const Grp &resp, const SigmaI &SigIb, const gsl_rng *r){
	gsl_matrix *rsd = gsl_matrix_alloc(_N, _d);
	gsl_matrix *Sig = gsl_matrix_alloc(_d, _d);
	gsl_vector *tmp = gsl_vector_alloc(_d);
	
	gsl_matrix_memcpy(rsd, resp.dMat());
	gsl_matrix_sub(rsd, &_fitted.matrix);
	
	gsl_blas_dgemv(CblasTrans, 1.0/_scale, rsd, &_X.vector, 0.0, tmp);
	
	gsl_matrix_memcpy(Sig, SigIb.getMat());
	gsl_linalg_cholesky_decomp(Sig);
	gsl_linalg_cholesky_invert(Sig);
	gsl_matrix_scale(Sig, 1.0/_scale);
	gsl_linalg_cholesky_decomp(Sig);
	
	MVgauss(tmp, Sig, r, &_vec.vector);
	
	gsl_matrix_free(rsd);
	gsl_matrix_free(Sig);
	gsl_vector_free(tmp);
}

void MVnormBetaFt::update(const Grp &resp, const Qgrp &q, const SigmaI &SigIb, const gsl_rng *r){
	gsl_matrix *rsd = gsl_matrix_alloc(_N, _d);
	gsl_matrix *Sig = gsl_matrix_alloc(_d, _d);
	gsl_vector *XQ  = gsl_vector_alloc(_N);
	gsl_vector *tmp = gsl_vector_alloc(_d);
	double xQx = 0.0;
	
	for (int iDat = 0; iDat < _N; iDat++) {
		gsl_vector_set(XQ, iDat, q[iDat]*gsl_vector_get(&_X.vector, iDat));
	}
	gsl_matrix_memcpy(rsd, resp.dMat());
	gsl_matrix_sub(rsd, &_fitted.matrix);
	
	gsl_blas_ddot(XQ, &_X.vector, &xQx);
	xQx = 1.0/xQx;
	
	gsl_blas_dgemv(CblasTrans, xQx, rsd, XQ, 0.0, tmp);
	
	gsl_matrix_memcpy(Sig, SigIb.getMat());
	gsl_linalg_cholesky_decomp(Sig);
	gsl_linalg_cholesky_invert(Sig);
	gsl_matrix_scale(Sig, xQx);
	gsl_linalg_cholesky_decomp(Sig);
	
	MVgauss(tmp, Sig, r, &_vec.vector);
	
	gsl_matrix_free(rsd);
	gsl_matrix_free(Sig);
	gsl_vector_free(XQ);
	gsl_vector_free(tmp);
}

void MVnormBetaFt::update(const Grp &resp, const SigmaI &SigIb, const SigmaI &SigIp, const gsl_rng *r){
	gsl_matrix *rsd    = gsl_matrix_alloc(_N, _d);
	gsl_matrix *SigSum = gsl_matrix_alloc(_d, _d);
	gsl_vector *tmpV   = gsl_vector_alloc(_d);
	gsl_vector *bH     = gsl_vector_alloc(_d);
	
	gsl_matrix_memcpy(rsd, resp.dMat());
	gsl_matrix_sub(rsd, &_fitted.matrix);
	
	gsl_matrix_memcpy(SigSum, SigIb.getMat());
	gsl_matrix_scale(SigSum, _scale);
	gsl_matrix_add(SigSum, SigIp.getMat());
	gsl_linalg_cholesky_decomp(SigSum);
	gsl_linalg_cholesky_invert(SigSum);
	
	gsl_blas_dgemv(CblasTrans, 1.0, rsd, &_X.vector, 0.0, bH);
	gsl_blas_dsymv(CblasLower, 1.0, SigIb.getMat(), bH, 0.0, tmpV);
	gsl_blas_dsymv(CblasLower, 1.0, SigSum, tmpV, 0.0, bH);
	
	gsl_linalg_cholesky_decomp(SigSum);
	MVgauss(bH, SigSum, r, &_vec.vector);
	
	gsl_matrix_free(rsd);
	gsl_matrix_free(SigSum);
	gsl_vector_free(tmpV);
	gsl_vector_free(bH);
}

void MVnormBetaFt::update(const Grp &resp, const SigmaI &SigIb, const double &qPr, const SigmaI &SigIp, const gsl_rng *r){
	gsl_matrix *rsd    = gsl_matrix_alloc(_N, _d);
	gsl_matrix *SigSum = gsl_matrix_alloc(_d, _d);
	gsl_matrix *SigPr  = gsl_matrix_alloc(_d, _d);
	gsl_vector *tmpV   = gsl_vector_alloc(_d);
	gsl_vector *bH     = gsl_vector_alloc(_d);
	
	gsl_matrix_memcpy(rsd, resp.dMat());
	gsl_matrix_sub(rsd, &_fitted.matrix);
	
	gsl_matrix_memcpy(SigSum, SigIb.getMat());
	gsl_matrix_scale(SigSum, _scale);
	gsl_matrix_memcpy(SigPr, SigIp.getMat());
	gsl_matrix_scale(SigPr, qPr);
	gsl_matrix_add(SigSum, SigPr);
	gsl_linalg_cholesky_decomp(SigSum);
	gsl_linalg_cholesky_invert(SigSum);
	
	gsl_blas_dgemv(CblasTrans, 1.0, rsd, &_X.vector, 0.0, bH);
	gsl_blas_dsymv(CblasLower, 1.0, SigIb.getMat(), bH, 0.0, tmpV);
	gsl_blas_dsymv(CblasLower, 1.0, SigSum, tmpV, 0.0, bH);
	
	gsl_linalg_cholesky_decomp(SigSum);
	MVgauss(bH, SigSum, r, &_vec.vector);
	
	gsl_matrix_free(rsd);
	gsl_matrix_free(SigSum);
	gsl_matrix_free(SigPr);
	gsl_vector_free(tmpV);
	gsl_vector_free(bH);
}

void MVnormBetaFt::update(const Grp &resp, const SigmaI &SigIb, const Grp &muPr, const SigmaI &SigIp, const gsl_rng *r){
	gsl_matrix *rsd    = gsl_matrix_alloc(_N, _d);
	gsl_matrix *SigSum = gsl_matrix_alloc(_d, _d);
	gsl_vector *tmpV   = gsl_vector_alloc(_d);
	gsl_vector *bH     = gsl_vector_alloc(_d);
	
	gsl_matrix_memcpy(rsd, resp.dMat());
	gsl_matrix_sub(rsd, &_fitted.matrix);
	
	gsl_matrix_memcpy(SigSum, SigIb.getMat());
	gsl_matrix_scale(SigSum, _scale);
	gsl_matrix_add(SigSum, SigIp.getMat());
	gsl_linalg_cholesky_decomp(SigSum);
	gsl_linalg_cholesky_invert(SigSum);
	
	gsl_blas_dgemv(CblasTrans, 1.0, rsd, &_X.vector, 0.0, bH);
	gsl_blas_dsymv(CblasLower, 1.0, SigIb.getMat(), bH, 0.0, tmpV);
	gsl_blas_dsymv(CblasLower, 1.0, SigIp.getMat(), muPr[*_upLevel]->getVec(), 1.0, tmpV);
	
	gsl_blas_dsymv(CblasLower, 1.0, SigSum, tmpV, 0.0, bH);
	
	gsl_linalg_cholesky_decomp(SigSum);
	MVgauss(bH, SigSum, r, &_vec.vector);
	
	gsl_matrix_free(rsd);
	gsl_matrix_free(SigSum);
	gsl_vector_free(tmpV);
	gsl_vector_free(bH);
}
void MVnormBetaFt::update(const Grp &resp, const SigmaI &SigIb, const Grp &muPr, const double &qPr, const SigmaI &SigIp, const gsl_rng *r){
	gsl_matrix *rsd    = gsl_matrix_alloc(_N, _d);
	gsl_matrix *SigSum = gsl_matrix_alloc(_d, _d);
	gsl_matrix *SigPr  = gsl_matrix_alloc(_d, _d);
	gsl_vector *tmpV   = gsl_vector_alloc(_d);
	gsl_vector *bH     = gsl_vector_alloc(_d);
	
	gsl_matrix_memcpy(rsd, resp.dMat());
	gsl_matrix_sub(rsd, &_fitted.matrix);
	
	gsl_matrix_memcpy(SigSum, SigIb.getMat());
	gsl_matrix_scale(SigSum, _scale);
	gsl_matrix_memcpy(SigPr, SigIp.getMat());
	gsl_matrix_scale(SigPr, qPr);
	gsl_matrix_add(SigSum, SigPr);
	gsl_linalg_cholesky_decomp(SigSum);
	gsl_linalg_cholesky_invert(SigSum);
	
	gsl_blas_dgemv(CblasTrans, 1.0, rsd, &_X.vector, 0.0, bH);
	gsl_blas_dsymv(CblasLower, 1.0, SigIb.getMat(), bH, 0.0, tmpV);
	gsl_blas_dsymv(CblasLower, 1.0, SigPr, muPr[*_upLevel]->getVec(), 1.0, tmpV);
	
	gsl_blas_dsymv(CblasLower, 1.0, SigSum, tmpV, 0.0, bH);
	
	gsl_linalg_cholesky_decomp(SigSum);
	MVgauss(bH, SigSum, r, &_vec.vector);
	
	gsl_matrix_free(rsd);
	gsl_matrix_free(SigSum);
	gsl_matrix_free(SigPr);
	gsl_vector_free(tmpV);
	gsl_vector_free(bH);
}

void MVnormBetaFt::update(const Grp &resp, const Qgrp &q, const SigmaI &SigIb, const SigmaI &SigIp, const gsl_rng *r){
	gsl_matrix *rsd    = gsl_matrix_alloc(_N, _d);
	gsl_matrix *SigSum = gsl_matrix_alloc(_d, _d);
	gsl_vector *XQ     = gsl_vector_alloc(_N);
	gsl_vector *tmpV   = gsl_vector_alloc(_d);
	gsl_vector *bH     = gsl_vector_alloc(_d);
	double xQx = 0.0;
	
	for (int iDat = 0; iDat < _N; iDat++) {
		gsl_vector_set(XQ, iDat, q[iDat]*gsl_vector_get(&_X.vector, iDat));
	}
	gsl_matrix_memcpy(rsd, resp.dMat());
	gsl_matrix_sub(rsd, &_fitted.matrix);
	
	gsl_blas_ddot(XQ, &_X.vector, &xQx);
	
	gsl_matrix_memcpy(SigSum, SigIb.getMat());
	gsl_matrix_scale(SigSum, xQx);
	gsl_matrix_add(SigSum, SigIp.getMat());
	gsl_linalg_cholesky_decomp(SigSum);
	gsl_linalg_cholesky_invert(SigSum);
	
	gsl_blas_dgemv(CblasTrans, 1.0, rsd, XQ, 0.0, bH);
	gsl_blas_dsymv(CblasLower, 1.0, SigIb.getMat(), bH, 0.0, tmpV);
	gsl_blas_dsymv(CblasLower, 1.0, SigSum, tmpV, 0.0, bH);
	
	gsl_linalg_cholesky_decomp(SigSum);
	MVgauss(bH, SigSum, r, &_vec.vector);
	
	gsl_matrix_free(rsd);
	gsl_matrix_free(SigSum);
	gsl_vector_free(XQ);
	gsl_vector_free(tmpV);
	gsl_vector_free(bH);
}

void MVnormBetaFt::update(const Grp &resp, const Qgrp &q, const SigmaI &SigIb, const double &qPr, const SigmaI &SigIp, const gsl_rng *r){
	gsl_matrix *rsd    = gsl_matrix_alloc(_N, _d);
	gsl_matrix *SigSum = gsl_matrix_alloc(_d, _d);
	gsl_matrix *SigPr  = gsl_matrix_alloc(_d, _d);
	gsl_vector *XQ     = gsl_vector_alloc(_N);
	gsl_vector *tmpV   = gsl_vector_alloc(_d);
	gsl_vector *bH     = gsl_vector_alloc(_d);
	double xQx = 0.0;
	
	for (int iDat = 0; iDat < _N; iDat++) {
		gsl_vector_set(XQ, iDat, q[iDat]*gsl_vector_get(&_X.vector, iDat));
	}
	gsl_matrix_memcpy(rsd, resp.dMat());
	gsl_matrix_sub(rsd, &_fitted.matrix);
	
	gsl_blas_ddot(XQ, &_X.vector, &xQx);
	
	gsl_matrix_memcpy(SigSum, SigIb.getMat());
	gsl_matrix_scale(SigSum, xQx);
	gsl_matrix_memcpy(SigPr, SigIp.getMat());
	gsl_matrix_scale(SigPr, qPr);
	gsl_matrix_add(SigSum, SigPr);
	gsl_linalg_cholesky_decomp(SigSum);
	gsl_linalg_cholesky_invert(SigSum);
	
	gsl_blas_dgemv(CblasTrans, 1.0, rsd, XQ, 0.0, bH);
	gsl_blas_dsymv(CblasLower, 1.0, SigIb.getMat(), bH, 0.0, tmpV);
	gsl_blas_dsymv(CblasLower, 1.0, SigSum, tmpV, 0.0, bH);
	
	gsl_linalg_cholesky_decomp(SigSum);
	MVgauss(bH, SigSum, r, &_vec.vector);
	
	gsl_matrix_free(rsd);
	gsl_matrix_free(SigSum);
	gsl_matrix_free(SigPr);
	gsl_vector_free(XQ);
	gsl_vector_free(tmpV);
	gsl_vector_free(bH);
}

void MVnormBetaFt::update(const Grp &resp, const Qgrp &q, const SigmaI &SigIb, const Grp &muPr, const SigmaI &SigIp, const gsl_rng *r){
	gsl_matrix *rsd    = gsl_matrix_alloc(_N, _d);
	gsl_matrix *SigSum = gsl_matrix_alloc(_d, _d);
	gsl_vector *XQ     = gsl_vector_alloc(_N);
	gsl_vector *tmpV   = gsl_vector_alloc(_d);
	gsl_vector *bH     = gsl_vector_alloc(_d);
	double xQx = 0.0;
	
	for (int iDat = 0; iDat < _N; iDat++) {
		gsl_vector_set(XQ, iDat, q[iDat]*gsl_vector_get(&_X.vector, iDat));
	}
	gsl_matrix_memcpy(rsd, resp.dMat());
	gsl_matrix_sub(rsd, &_fitted.matrix);
	
	gsl_blas_ddot(XQ, &_X.vector, &xQx);
	
	gsl_matrix_memcpy(SigSum, SigIb.getMat());
	gsl_matrix_scale(SigSum, xQx);
	gsl_matrix_add(SigSum, SigIp.getMat());
	gsl_linalg_cholesky_decomp(SigSum);
	gsl_linalg_cholesky_invert(SigSum);
	
	gsl_blas_dgemv(CblasTrans, 1.0, rsd, XQ, 0.0, bH);
	gsl_blas_dsymv(CblasLower, 1.0, SigIb.getMat(), bH, 0.0, tmpV);
	gsl_blas_dsymv(CblasLower, 1.0, SigIp.getMat(), muPr[*_upLevel]->getVec(), 1.0, tmpV);
	
	gsl_blas_dsymv(CblasLower, 1.0, SigSum, tmpV, 0.0, bH);
	
	gsl_linalg_cholesky_decomp(SigSum);
	MVgauss(bH, SigSum, r, &_vec.vector);
	
	gsl_matrix_free(rsd);
	gsl_matrix_free(SigSum);
	gsl_vector_free(XQ);
	gsl_vector_free(tmpV);
	gsl_vector_free(bH);
}
void MVnormBetaFt::update(const Grp &resp, const Qgrp &q, const SigmaI &SigIb, const Grp &muPr, const double &qPr, const SigmaI &SigIp, const gsl_rng *r){
	gsl_matrix *rsd    = gsl_matrix_alloc(_N, _d);
	gsl_matrix *SigSum = gsl_matrix_alloc(_d, _d);
	gsl_matrix *SigPr  = gsl_matrix_alloc(_d, _d);
	gsl_vector *XQ     = gsl_vector_alloc(_N);
	gsl_vector *tmpV   = gsl_vector_alloc(_d);
	gsl_vector *bH     = gsl_vector_alloc(_d);
	double xQx = 0.0;
	
	for (int iDat = 0; iDat < _N; iDat++) {
		gsl_vector_set(XQ, iDat, q[iDat]*gsl_vector_get(&_X.vector, iDat));
	}
	gsl_matrix_memcpy(rsd, resp.dMat());
	gsl_matrix_sub(rsd, &_fitted.matrix);
	
	gsl_blas_ddot(XQ, &_X.vector, &xQx);
	
	gsl_matrix_memcpy(SigSum, SigIb.getMat());
	gsl_matrix_scale(SigSum, xQx);
	gsl_matrix_memcpy(SigPr, SigIp.getMat());
	gsl_matrix_scale(SigPr, qPr);
	gsl_matrix_add(SigSum, SigPr);
	gsl_linalg_cholesky_decomp(SigSum);
	gsl_linalg_cholesky_invert(SigSum);
	
	gsl_blas_dgemv(CblasTrans, 1.0, rsd, XQ, 0.0, bH);
	gsl_blas_dsymv(CblasLower, 1.0, SigIb.getMat(), bH, 0.0, tmpV);
	gsl_blas_dsymv(CblasLower, 1.0, SigPr, muPr[*_upLevel]->getVec(), 1.0, tmpV);
	
	gsl_blas_dsymv(CblasLower, 1.0, SigSum, tmpV, 0.0, bH);
	
	gsl_linalg_cholesky_decomp(SigSum);
	MVgauss(bH, SigSum, r, &_vec.vector);
	
	gsl_matrix_free(rsd);
	gsl_matrix_free(SigSum);
	gsl_matrix_free(SigPr);
	gsl_vector_free(XQ);
	gsl_vector_free(tmpV);
	gsl_vector_free(bH);
}

/*
 *	MVnormBlk methods
 */

MVnormMuBlk::MVnormMuBlk(gsl_matrix *mn, const size_t &iRw, const vector<size_t> &blkStart, const size_t &up) : MVnorm(mn, iRw){
	_upLevel = &up;
	_blkStart = &blkStart;
	
	_eachVec.resize(_blkStart->size());
	
	vector<size_t> len;
	for (size_t iBlk = 1; iBlk <= (*_blkStart).size(); iBlk++) {
		iBlk == (*_blkStart).size() ? len.push_back(_d - (*_blkStart)[iBlk - 1]) : len.push_back((*_blkStart)[iBlk] - (*_blkStart)[iBlk - 1]);
	}
	
	for (size_t iBlk = 0; iBlk < _blkStart->size(); iBlk++) {
		_eachVec[iBlk] = gsl_vector_subvector(&_vec.vector, (*_blkStart)[iBlk], len[iBlk]);
	}
}
MVnormMuBlk::MVnormMuBlk(gsl_matrix *mn, const size_t &iRw, const gsl_vector *sd, const gsl_rng *r, const vector<size_t> &blkStart, const vector< vector<size_t> > &eachLL, const size_t &up) : MVnorm(mn, iRw, sd, r){
	_upLevel  = &up;
	_blkStart = &blkStart;
	_eachLL   = &eachLL;
	_eachVec.resize(_blkStart->size());
	
	vector<size_t> len;
	for (size_t iBlk = 1; iBlk <= (*_blkStart).size(); iBlk++) {
		iBlk == (*_blkStart).size() ? len.push_back(_d - (*_blkStart)[iBlk - 1]) : len.push_back((*_blkStart)[iBlk] - (*_blkStart)[iBlk - 1]);
	}
	
	for (size_t iBlk = 0; iBlk < _blkStart->size(); iBlk++) {
		_eachVec[iBlk] = gsl_vector_subvector(&_vec.vector, (*_blkStart)[iBlk], len[iBlk]);
	}
}

void MVnormMuBlk::update(const Grp &dat, const SigmaI &SigIm, const gsl_rng *r){
	for (size_t iElm = 0; iElm < _eachVec.size(); iElm++) {
		if ((*_eachLL)[iElm].empty()) { // this is for situations where the number of levels is not the same for all blocks
			continue;
		}
		gsl_vector *smVec  = gsl_vector_calloc(_eachVec[iElm].vector.size);
		gsl_matrix *SigSum = gsl_matrix_alloc(_eachVec[iElm].vector.size, _eachVec[iElm].vector.size);
		gsl_matrix_const_view SigIsub = gsl_matrix_const_submatrix(SigIm.getMat(), (*_blkStart)[iElm], (*_blkStart)[iElm], _eachVec[iElm].vector.size, _eachVec[iElm].vector.size);
		
		for (vector<size_t>::const_iterator it = (*_eachLL)[iElm].begin(); it != (*_eachLL)[iElm].end(); ++it) {
			gsl_vector_const_view datSub = gsl_vector_const_subvector(dat[*it]->getVec(), (*_blkStart)[iElm], _eachVec[iElm].vector.size);
			gsl_vector_add(smVec, &datSub.vector);
		}
		gsl_vector_scale(smVec, 1.0/(*_eachLL)[iElm].size());
		// prepare the scale matrices
		gsl_matrix_memcpy(SigSum, &SigIsub.matrix);
		gsl_matrix_scale(SigSum, (*_eachLL)[iElm].size());
		gsl_linalg_cholesky_decomp(SigSum);
		gsl_linalg_cholesky_invert(SigSum);
		gsl_linalg_cholesky_decomp(SigSum);
		
		MVgauss(smVec, SigSum, r, &(_eachVec[iElm]).vector);
		
		gsl_vector_free(smVec);
		gsl_matrix_free(SigSum);
		
	}
}
void MVnormMuBlk::update(const Grp &dat, const Qgrp &q, const SigmaI &SigIm, const gsl_rng *r){
	for (size_t iElm = 0; iElm < _eachVec.size(); iElm++) {
		if ((*_eachLL)[iElm].empty()) { // this is for situations where the number of levels is not the same for all blocks
			continue;
		}
		gsl_vector *smVec  = gsl_vector_calloc(_eachVec[iElm].vector.size);
		gsl_matrix *SigSum = gsl_matrix_alloc(_eachVec[iElm].vector.size, _eachVec[iElm].vector.size);
		gsl_matrix_const_view SigIsub = gsl_matrix_const_submatrix(SigIm.getMat(), (*_blkStart)[iElm], (*_blkStart)[iElm], _eachVec[iElm].vector.size, _eachVec[iElm].vector.size);
		
		double qSum = 0.0;
		
		for (vector<size_t>::const_iterator it = (*_eachLL)[iElm].begin(); it != (*_eachLL)[iElm].end(); ++it) {
			gsl_vector_const_view datSub = gsl_vector_const_subvector(dat[*it]->getVec(), (*_blkStart)[iElm], _eachVec[iElm].vector.size);
			gsl_blas_daxpy(q[*it], &datSub.vector, smVec);
			qSum += q[*it];
		}
		gsl_vector_scale(smVec, 1.0/qSum);
		
		// prepare the scale matrices
		gsl_matrix_memcpy(SigSum, &SigIsub.matrix);
		gsl_matrix_scale(SigSum, qSum);
		gsl_linalg_cholesky_decomp(SigSum);
		gsl_linalg_cholesky_invert(SigSum);
		gsl_linalg_cholesky_decomp(SigSum);
		
		MVgauss(smVec, SigSum, r, &(_eachVec[iElm]).vector);
		
		gsl_vector_free(smVec);
		gsl_matrix_free(SigSum);

	}
	
}

void MVnormMuBlk::update(const Grp &dat, const SigmaI &SigIm, const SigmaI &SigIp, const gsl_rng *r){
	for (size_t iElm = 0; iElm < _eachVec.size(); iElm++) {
		if ((*_eachLL)[iElm].empty()) { // this is for situations where the number of levels is not the same for all blocks
			continue;
		}
		gsl_vector *smVec  = gsl_vector_calloc(_eachVec[iElm].vector.size);
		gsl_vector *tmpV   = gsl_vector_alloc(_eachVec[iElm].vector.size);
		gsl_matrix *SigSum = gsl_matrix_alloc(_eachVec[iElm].vector.size, _eachVec[iElm].vector.size);
		gsl_matrix_const_view SigIms = gsl_matrix_const_submatrix(SigIm.getMat(), (*_blkStart)[iElm], (*_blkStart)[iElm], _eachVec[iElm].vector.size, _eachVec[iElm].vector.size);
		gsl_matrix_const_view SigIps = gsl_matrix_const_submatrix(SigIp.getMat(), (*_blkStart)[iElm], (*_blkStart)[iElm], _eachVec[iElm].vector.size, _eachVec[iElm].vector.size);
		
		for (vector<size_t>::const_iterator it = (*_eachLL)[iElm].begin(); it != (*_eachLL)[iElm].end(); ++it) {
			gsl_vector_const_view datSub = gsl_vector_const_subvector(dat[*it]->getVec(), (*_blkStart)[iElm], _eachVec[iElm].vector.size);
			gsl_vector_add(smVec, &datSub.vector);
		}
		
		// prepare the scale matrices
		gsl_matrix_memcpy(SigSum, &SigIms.matrix);
		gsl_matrix_scale(SigSum, (*_eachLL)[iElm].size());
		gsl_matrix_add(SigSum, &SigIps.matrix);
		gsl_linalg_cholesky_decomp(SigSum);
		gsl_linalg_cholesky_invert(SigSum);
		
		// prepare the scaled mean vector
		gsl_blas_dsymv(CblasLower, 1.0, &SigIms.matrix, smVec, 0.0, tmpV);
		gsl_blas_dsymv(CblasLower, 1.0, SigSum, tmpV, 0.0, smVec);
		
		gsl_linalg_cholesky_decomp(SigSum);
		MVgauss(smVec, SigSum, r, &(_eachVec[iElm]).vector);
		
		gsl_vector_free(tmpV);
		gsl_vector_free(smVec);
		gsl_matrix_free(SigSum);
		
		
	}
	
}
void MVnormMuBlk::update(const Grp &dat, const SigmaI &SigIm, const double &qPr, const SigmaI &SigIp, const gsl_rng *r){
	for (size_t iElm = 0; iElm < _eachVec.size(); iElm++) {
		if ((*_eachLL)[iElm].empty()) { // this is for situations where the number of levels is not the same for all blocks
			continue;
		}
		gsl_vector *smVec  = gsl_vector_calloc(_eachVec[iElm].vector.size);
		gsl_vector *tmpV   = gsl_vector_alloc(_eachVec[iElm].vector.size);
		gsl_matrix *SigSum = gsl_matrix_alloc(_eachVec[iElm].vector.size, _eachVec[iElm].vector.size);
		gsl_matrix *SigPr  = gsl_matrix_alloc(_eachVec[iElm].vector.size, _eachVec[iElm].vector.size);
		gsl_matrix_const_view SigIms = gsl_matrix_const_submatrix(SigIm.getMat(), (*_blkStart)[iElm], (*_blkStart)[iElm], _eachVec[iElm].vector.size, _eachVec[iElm].vector.size);
		gsl_matrix_const_view SigIps = gsl_matrix_const_submatrix(SigIp.getMat(), (*_blkStart)[iElm], (*_blkStart)[iElm], _eachVec[iElm].vector.size, _eachVec[iElm].vector.size);
		
		for (vector<size_t>::const_iterator it = (*_eachLL)[iElm].begin(); it != (*_eachLL)[iElm].end(); ++it) {
			gsl_vector_const_view datSub = gsl_vector_const_subvector(dat[*it]->getVec(), (*_blkStart)[iElm], _eachVec[iElm].vector.size);
			gsl_vector_add(smVec, &datSub.vector);
		}
		
		// prepare the scale matrices
		gsl_matrix_memcpy(SigSum, &SigIms.matrix);
		gsl_matrix_scale(SigSum, (*_eachLL)[iElm].size());
		gsl_matrix_memcpy(SigPr, &SigIps.matrix);
		gsl_matrix_scale(SigPr, qPr);
		gsl_matrix_add(SigSum, SigPr);
		gsl_linalg_cholesky_decomp(SigSum);
		gsl_linalg_cholesky_invert(SigSum);
		
		// prepare the scaled mean vector
		gsl_blas_dsymv(CblasLower, 1.0, &SigIms.matrix, smVec, 0.0, tmpV);
		gsl_blas_dsymv(CblasLower, 1.0, SigSum, tmpV, 0.0, smVec);
		
		gsl_linalg_cholesky_decomp(SigSum);
		MVgauss(smVec, SigSum, r, &(_eachVec[iElm]).vector);
		
		gsl_vector_free(tmpV);
		gsl_vector_free(smVec);
		gsl_matrix_free(SigSum);
		gsl_matrix_free(SigPr);
		

	}
}
void MVnormMuBlk::update(const Grp &dat, const Qgrp &q, const SigmaI &SigIm, const SigmaI &SigIp, const gsl_rng *r){
	for (size_t iElm = 0; iElm < _eachVec.size(); iElm++) {
		if ((*_eachLL)[iElm].empty()) { // this is for situations where the number of levels is not the same for all blocks
			continue;
		}
		gsl_vector *smVec  = gsl_vector_calloc(_eachVec[iElm].vector.size);
		gsl_vector *tmpV   = gsl_vector_alloc(_eachVec[iElm].vector.size);
		gsl_matrix *SigSum = gsl_matrix_alloc(_eachVec[iElm].vector.size, _eachVec[iElm].vector.size);
		gsl_matrix_const_view SigIms = gsl_matrix_const_submatrix(SigIm.getMat(), (*_blkStart)[iElm], (*_blkStart)[iElm], _eachVec[iElm].vector.size, _eachVec[iElm].vector.size);
		gsl_matrix_const_view SigIps = gsl_matrix_const_submatrix(SigIp.getMat(), (*_blkStart)[iElm], (*_blkStart)[iElm], _eachVec[iElm].vector.size, _eachVec[iElm].vector.size);
		
		double qSum = 0.0;
		
		for (vector<size_t>::const_iterator it = (*_eachLL)[iElm].begin(); it != (*_eachLL)[iElm].end(); ++it) {
			gsl_vector_const_view datSub = gsl_vector_const_subvector(dat[*it]->getVec(), (*_blkStart)[iElm], _eachVec[iElm].vector.size);
			gsl_blas_daxpy(q[*it], &datSub.vector, smVec);
			qSum += q[*it];
		}
		
		// prepare the scale matrices
		gsl_matrix_memcpy(SigSum, &SigIms.matrix);
		gsl_matrix_scale(SigSum, qSum);
		gsl_matrix_add(SigSum, &SigIps.matrix);
		gsl_linalg_cholesky_decomp(SigSum);
		gsl_linalg_cholesky_invert(SigSum);
		
		
		// prepare the scaled mean vector
		gsl_blas_dsymv(CblasLower, 1.0, &SigIms.matrix, smVec, 0.0, tmpV);
		gsl_blas_dsymv(CblasLower, 1.0, SigSum, tmpV, 0.0, smVec);
		
		gsl_linalg_cholesky_decomp(SigSum);
		MVgauss(smVec, SigSum, r, &(_eachVec[iElm]).vector);
		
		gsl_vector_free(tmpV);
		gsl_vector_free(smVec);
		gsl_matrix_free(SigSum);

	}

}
void MVnormMuBlk::update(const Grp &dat, const Qgrp &q, const SigmaI &SigIm, const double &qPr, const SigmaI &SigIp, const gsl_rng *r){
	for (size_t iElm = 0; iElm < _eachVec.size(); iElm++) {
		if ((*_eachLL)[iElm].empty()) { // this is for situations where the number of levels is not the same for all blocks
			continue;
		}
		gsl_vector *smVec  = gsl_vector_calloc(_eachVec[iElm].vector.size);
		gsl_vector *tmpV   = gsl_vector_alloc(_eachVec[iElm].vector.size);
		gsl_matrix *SigSum = gsl_matrix_alloc(_eachVec[iElm].vector.size, _eachVec[iElm].vector.size);
		gsl_matrix *SigPr  = gsl_matrix_alloc(_eachVec[iElm].vector.size, _eachVec[iElm].vector.size);
		gsl_matrix_const_view SigIms = gsl_matrix_const_submatrix(SigIm.getMat(), (*_blkStart)[iElm], (*_blkStart)[iElm], _eachVec[iElm].vector.size, _eachVec[iElm].vector.size);
		gsl_matrix_const_view SigIps = gsl_matrix_const_submatrix(SigIp.getMat(), (*_blkStart)[iElm], (*_blkStart)[iElm], _eachVec[iElm].vector.size, _eachVec[iElm].vector.size);
		
		double qSum = 0.0;
		
		for (vector<size_t>::const_iterator it = (*_eachLL)[iElm].begin(); it != (*_eachLL)[iElm].end(); ++it) {
			gsl_vector_const_view datSub = gsl_vector_const_subvector(dat[*it]->getVec(), (*_blkStart)[iElm], _eachVec[iElm].vector.size);
			gsl_blas_daxpy(q[*it], &datSub.vector, smVec);
			qSum += q[*it];
		}
		
		// prepare the scale matrices
		gsl_matrix_memcpy(SigSum, &SigIms.matrix);
		gsl_matrix_scale(SigSum, qSum);
		gsl_matrix_memcpy(SigPr, &SigIps.matrix);
		gsl_matrix_scale(SigPr, qPr);
		gsl_matrix_add(SigSum, SigPr);
		gsl_linalg_cholesky_decomp(SigSum);
		gsl_linalg_cholesky_invert(SigSum);
		
		
		// prepare the scaled mean vector
		gsl_blas_dsymv(CblasLower, 1.0, &SigIms.matrix, smVec, 0.0, tmpV);
		gsl_blas_dsymv(CblasLower, 1.0, SigSum, tmpV, 0.0, smVec);
		
		gsl_linalg_cholesky_decomp(SigSum);
		MVgauss(smVec, SigSum, r, &(_eachVec[iElm]).vector);
		
		gsl_vector_free(tmpV);
		gsl_vector_free(smVec);
		gsl_matrix_free(SigSum);
		gsl_matrix_free(SigPr);

	}
	
}

void MVnormMuBlk::update(const Grp &dat, const SigmaI &SigIm, const Grp &muPr, const SigmaI &SigIp, const gsl_rng *r){
	for (size_t iElm = 0; iElm < _eachVec.size(); iElm++) {
		if ((*_eachLL)[iElm].empty()) { // this is for situations where the number of levels is not the same for all blocks
			continue;
		}
		gsl_vector *smVec  = gsl_vector_calloc(_eachVec[iElm].vector.size);
		gsl_vector *tmpV   = gsl_vector_alloc(_eachVec[iElm].vector.size);
		gsl_matrix *SigSum = gsl_matrix_alloc(_eachVec[iElm].vector.size, _eachVec[iElm].vector.size);
		gsl_matrix_const_view SigIms = gsl_matrix_const_submatrix(SigIm.getMat(), (*_blkStart)[iElm], (*_blkStart)[iElm], _eachVec[iElm].vector.size, _eachVec[iElm].vector.size);
		gsl_matrix_const_view SigIps = gsl_matrix_const_submatrix(SigIp.getMat(), (*_blkStart)[iElm], (*_blkStart)[iElm], _eachVec[iElm].vector.size, _eachVec[iElm].vector.size);
		gsl_vector_const_view muSub  = gsl_vector_const_subvector(muPr[*_upLevel]->getVec(), (*_blkStart)[iElm], _eachVec[iElm].vector.size);
		
		for (vector<size_t>::const_iterator it = (*_eachLL)[iElm].begin(); it != (*_eachLL)[iElm].end(); ++it) {
			gsl_vector_const_view datSub = gsl_vector_const_subvector(dat[*it]->getVec(), (*_blkStart)[iElm], _eachVec[iElm].vector.size);
			gsl_vector_add(smVec, &datSub.vector);
		}
		
		// prepare the scale matrices
		gsl_matrix_memcpy(SigSum, &SigIms.matrix);
		gsl_matrix_scale(SigSum, (*_eachLL)[iElm].size());
		gsl_matrix_add(SigSum, &SigIps.matrix);
		gsl_linalg_cholesky_decomp(SigSum);
		gsl_linalg_cholesky_invert(SigSum);
		
		// prepare the scaled mean vector
		gsl_blas_dsymv(CblasLower, 1.0, &SigIms.matrix, smVec, 0.0, tmpV);
		gsl_blas_dsymv(CblasLower, 1.0, &SigIps.matrix, &muSub.vector, 1.0, tmpV);
		gsl_blas_dsymv(CblasLower, 1.0, SigSum, tmpV, 0.0, smVec);
		
		gsl_linalg_cholesky_decomp(SigSum);
		MVgauss(smVec, SigSum, r, &(_eachVec[iElm]).vector);
		
		gsl_vector_free(tmpV);
		gsl_vector_free(smVec);
		gsl_matrix_free(SigSum);
		
	}
}
void MVnormMuBlk::update(const Grp &dat, const SigmaI &SigIm, const Grp &muPr, const double &qPr, const SigmaI &SigIp, const gsl_rng *r){
	for (size_t iElm = 0; iElm < _eachVec.size(); iElm++) {
		if ((*_eachLL)[iElm].empty()) { // this is for situations where the number of levels is not the same for all blocks
			continue;
		}
		gsl_vector *smVec  = gsl_vector_calloc(_eachVec[iElm].vector.size);
		gsl_vector *tmpV   = gsl_vector_alloc(_eachVec[iElm].vector.size);
		gsl_matrix *SigSum = gsl_matrix_alloc(_eachVec[iElm].vector.size, _eachVec[iElm].vector.size);
		gsl_matrix *SigPr  = gsl_matrix_alloc(_eachVec[iElm].vector.size, _eachVec[iElm].vector.size);
		gsl_matrix_const_view SigIms = gsl_matrix_const_submatrix(SigIm.getMat(), (*_blkStart)[iElm], (*_blkStart)[iElm], _eachVec[iElm].vector.size, _eachVec[iElm].vector.size);
		gsl_matrix_const_view SigIps = gsl_matrix_const_submatrix(SigIp.getMat(), (*_blkStart)[iElm], (*_blkStart)[iElm], _eachVec[iElm].vector.size, _eachVec[iElm].vector.size);
		gsl_vector_const_view muSub  = gsl_vector_const_subvector(muPr[*_upLevel]->getVec(), (*_blkStart)[iElm], _eachVec[iElm].vector.size);
		
		for (vector<size_t>::const_iterator it = (*_eachLL)[iElm].begin(); it != (*_eachLL)[iElm].end(); ++it) {
			gsl_vector_const_view datSub = gsl_vector_const_subvector(dat[*it]->getVec(), (*_blkStart)[iElm], _eachVec[iElm].vector.size);
			gsl_vector_add(smVec, &datSub.vector);
		}
		
		// prepare the scale matrices
		gsl_matrix_memcpy(SigSum, &SigIms.matrix);
		gsl_matrix_scale(SigSum, (*_eachLL)[iElm].size());
		gsl_matrix_memcpy(SigPr, &SigIps.matrix);
		gsl_matrix_scale(SigPr, qPr);
		gsl_matrix_add(SigSum, SigPr);
		gsl_linalg_cholesky_decomp(SigSum);
		gsl_linalg_cholesky_invert(SigSum);
		
		// prepare the scaled mean vector
		gsl_blas_dsymv(CblasLower, 1.0, &SigIms.matrix, smVec, 0.0, tmpV);
		gsl_blas_dsymv(CblasLower, 1.0, SigPr, &muSub.vector, 1.0, tmpV);
		gsl_blas_dsymv(CblasLower, 1.0, SigSum, tmpV, 0.0, smVec);
		
		gsl_linalg_cholesky_decomp(SigSum);
		MVgauss(smVec, SigSum, r, &(_eachVec[iElm]).vector);
		
		gsl_vector_free(tmpV);
		gsl_vector_free(smVec);
		gsl_matrix_free(SigSum);
		gsl_matrix_free(SigPr);

	}

}
void MVnormMuBlk::update(const Grp &dat, const Qgrp &q, const SigmaI &SigIm, const Grp &muPr, const SigmaI &SigIp, const gsl_rng *r){
	for (size_t iElm = 0; iElm < _eachVec.size(); iElm++) {
		if ((*_eachLL)[iElm].empty()) { // this is for situations where the number of levels is not the same for all blocks
			continue;
		}
		gsl_vector *smVec  = gsl_vector_calloc(_eachVec[iElm].vector.size);
		gsl_vector *tmpV   = gsl_vector_alloc(_eachVec[iElm].vector.size);
		gsl_matrix *SigSum = gsl_matrix_alloc(_eachVec[iElm].vector.size, _eachVec[iElm].vector.size);
		gsl_matrix_const_view SigIms = gsl_matrix_const_submatrix(SigIm.getMat(), (*_blkStart)[iElm], (*_blkStart)[iElm], _eachVec[iElm].vector.size, _eachVec[iElm].vector.size);
		gsl_matrix_const_view SigIps = gsl_matrix_const_submatrix(SigIp.getMat(), (*_blkStart)[iElm], (*_blkStart)[iElm], _eachVec[iElm].vector.size, _eachVec[iElm].vector.size);
		gsl_vector_const_view muSub  = gsl_vector_const_subvector(muPr[*_upLevel]->getVec(), (*_blkStart)[iElm], _eachVec[iElm].vector.size);
		
		double qSum = 0.0;
		
		for (vector<size_t>::const_iterator it = (*_eachLL)[iElm].begin(); it != (*_eachLL)[iElm].end(); ++it) {
			gsl_vector_const_view datSub = gsl_vector_const_subvector(dat[*it]->getVec(), (*_blkStart)[iElm], _eachVec[iElm].vector.size);
			gsl_blas_daxpy(q[*it], &datSub.vector, smVec);
			qSum += q[*it];
		}
		
		// prepare the scale matrices
		gsl_matrix_memcpy(SigSum, &SigIms.matrix);
		gsl_matrix_scale(SigSum, qSum);
		gsl_matrix_add(SigSum, &SigIps.matrix);
		gsl_linalg_cholesky_decomp(SigSum);
		gsl_linalg_cholesky_invert(SigSum);
		
		
		// prepare the scaled mean vector
		gsl_blas_dsymv(CblasLower, 1.0, &SigIms.matrix, smVec, 0.0, tmpV);
		gsl_blas_dsymv(CblasLower, 1.0, &SigIps.matrix, &muSub.vector, 1.0, tmpV);
		gsl_blas_dsymv(CblasLower, 1.0, SigSum, tmpV, 0.0, smVec);
		
		gsl_linalg_cholesky_decomp(SigSum);
		MVgauss(smVec, SigSum, r, &(_eachVec[iElm]).vector);
		
		gsl_vector_free(tmpV);
		gsl_vector_free(smVec);
		gsl_matrix_free(SigSum);

	}

}
void MVnormMuBlk::update(const Grp &dat, const Qgrp &q, const SigmaI &SigIm, const Grp &muPr, const double &qPr, const SigmaI &SigIp, const gsl_rng *r){
	for (size_t iElm = 0; iElm < _eachVec.size(); iElm++) {
		if ((*_eachLL)[iElm].empty()) { // this is for situations where the number of levels is not the same for all blocks
			continue;
		}
		gsl_vector *smVec  = gsl_vector_calloc(_eachVec[iElm].vector.size);
		gsl_vector *tmpV   = gsl_vector_alloc(_eachVec[iElm].vector.size);
		gsl_matrix *SigSum = gsl_matrix_alloc(_eachVec[iElm].vector.size, _eachVec[iElm].vector.size);
		gsl_matrix *SigPr  = gsl_matrix_alloc(_eachVec[iElm].vector.size, _eachVec[iElm].vector.size);
		gsl_matrix_const_view SigIms = gsl_matrix_const_submatrix(SigIm.getMat(), (*_blkStart)[iElm], (*_blkStart)[iElm], _eachVec[iElm].vector.size, _eachVec[iElm].vector.size);
		gsl_matrix_const_view SigIps = gsl_matrix_const_submatrix(SigIp.getMat(), (*_blkStart)[iElm], (*_blkStart)[iElm], _eachVec[iElm].vector.size, _eachVec[iElm].vector.size);
		gsl_vector_const_view muSub  = gsl_vector_const_subvector(muPr[*_upLevel]->getVec(), (*_blkStart)[iElm], _eachVec[iElm].vector.size);
		
		double qSum = 0.0;
		
		for (vector<size_t>::const_iterator it = (*_eachLL)[iElm].begin(); it != (*_eachLL)[iElm].end(); ++it) {
			gsl_vector_const_view datSub = gsl_vector_const_subvector(dat[*it]->getVec(), (*_blkStart)[iElm], _eachVec[iElm].vector.size);
			gsl_blas_daxpy(q[*it], &datSub.vector, smVec);
			qSum += q[*it];
		}
		
		// prepare the scale matrices
		gsl_matrix_memcpy(SigSum, &SigIms.matrix);
		gsl_matrix_scale(SigSum, qSum);
		gsl_matrix_memcpy(SigPr, &SigIps.matrix);
		gsl_matrix_scale(SigPr, qPr);
		gsl_matrix_add(SigSum, SigPr);
		gsl_linalg_cholesky_decomp(SigSum);
		gsl_linalg_cholesky_invert(SigSum);
		
		// prepare the scaled mean vector
		gsl_blas_dsymv(CblasLower, 1.0, &SigIms.matrix, smVec, 0.0, tmpV);
		gsl_blas_dsymv(CblasLower, 1.0, SigPr, &muSub.vector, 1.0, tmpV);
		gsl_blas_dsymv(CblasLower, 1.0, SigSum, tmpV, 0.0, smVec);
		
		gsl_linalg_cholesky_decomp(SigSum);
		MVgauss(smVec, SigSum, r, &(_eachVec[iElm]).vector);
		
		gsl_vector_free(tmpV);
		gsl_vector_free(smVec);
		gsl_matrix_free(SigSum);
		gsl_matrix_free(SigPr);

	}
}

/*
 *	MVnormBetaBlk methods
 */

MVnormBetaBlk::MVnormBetaBlk(const Grp &resp, gsl_matrix *pred, const vector<size_t> &iCl, const gsl_matrix *Sig, const vector<size_t> &blkStart, const gsl_rng *r, gsl_matrix *bet, const size_t &iRw){
	_upLevel  = 0;
	_d        = resp.phenD();
	_vec      = gsl_matrix_row(bet, iRw);
	_blkStart = &blkStart;
	
	_eachVec.resize(_blkStart->size());
	
	vector<size_t> len;
	for (size_t iBlk = 1; iBlk <= (*_blkStart).size(); iBlk++) {
		iBlk == (*_blkStart).size() ? len.push_back(_d - (*_blkStart)[iBlk - 1]) : len.push_back((*_blkStart)[iBlk] - (*_blkStart)[iBlk - 1]);
	}
	
	for (size_t iBlk = 0; iBlk < _blkStart->size(); iBlk++) {
		_eachVec[iBlk] = gsl_vector_subvector(&_vec.vector, (*_blkStart)[iBlk], len[iBlk]);
	}
	
	if (pred->size1 != resp.Ndata()) {
		cerr << "ERROR: unequal number of rows in predictor (" << pred->size1 << ") and response (" << resp.Ndata() << ") when initializing MVnormBeta" << endl;
		exit(1);
	}
	else if (resp.phenD() != Sig->size1){
		cerr << "ERROR: incompatible matrices *resp (d = " << resp.phenD() << ") and *Sig (d = " << Sig->size1 << ") when initializing MVnormBeta" << endl;
		exit(1);
	}
	
	_X.resize(_blkStart->size());
	_scale.resize(_blkStart->size());
	for (size_t iEl = 0; iEl < _blkStart->size(); iEl++) {
		_X[iEl] = gsl_matrix_column(pred, iCl[iEl]);
		gsl_blas_ddot(&(_X[iEl]).vector, &(_X[iEl]).vector, &(_scale[iEl]));
		
		gsl_vector *bH  = gsl_vector_alloc(_eachVec[iEl].vector.size);
		gsl_matrix *chl = gsl_matrix_alloc(_eachVec[iEl].vector.size, _eachVec[iEl].vector.size);
		
		gsl_matrix_const_view sigSub  = gsl_matrix_const_submatrix(Sig, (*_blkStart)[iEl], (*_blkStart)[iEl], _eachVec[iEl].vector.size, _eachVec[iEl].vector.size);
		gsl_matrix_const_view respSub = gsl_matrix_const_submatrix(resp.dMat(), 0, (*_blkStart)[iEl], resp.Ndata(), _eachVec[iEl].vector.size);
		
		gsl_blas_dgemv(CblasTrans, 1.0/_scale[iEl], &respSub.matrix, &(_X[iEl]).vector, 0.0, bH);
		
		gsl_matrix_memcpy(chl, &sigSub.matrix);
		gsl_matrix_scale(chl, 1.0/_scale[iEl]);
		gsl_linalg_cholesky_decomp(chl);
		
		MVgauss(bH, chl, r, &(_eachVec[iEl]).vector);
		
		gsl_vector_free(bH);
		gsl_matrix_free(chl);
		
	}

}
MVnormBetaBlk::MVnormBetaBlk(const Grp &resp, gsl_matrix *pred, const vector<size_t> &iCl, const gsl_matrix *Sig, const vector<size_t> &blkStart, const gsl_rng *r, const size_t &up, gsl_matrix *bet, const size_t &iRw){
	_upLevel  = &up;
	_d        = resp.phenD();
	_vec      = gsl_matrix_row(bet, iRw);
	_blkStart = &blkStart;
	
	_eachVec.resize(_blkStart->size());
	
	vector<size_t> len;
	for (size_t iBlk = 1; iBlk <= (*_blkStart).size(); iBlk++) {
		iBlk == (*_blkStart).size() ? len.push_back(_d - (*_blkStart)[iBlk - 1]) : len.push_back((*_blkStart)[iBlk] - (*_blkStart)[iBlk - 1]);
	}
	
	for (size_t iBlk = 0; iBlk < _blkStart->size(); iBlk++) {
		_eachVec[iBlk] = gsl_vector_subvector(&_vec.vector, (*_blkStart)[iBlk], len[iBlk]);
	}

	if (pred->size1 != resp.Ndata()) {
		cerr << "ERROR: unequal number of rows in predictor (" << pred->size1 << ") and response (" << resp.Ndata() << ") when initializing MVnormBeta" << endl;
		exit(1);
	}
	else if (resp.phenD() != Sig->size1){
		cerr << "ERROR: incompatible matrices *resp (d = " << resp.phenD() << ") and *Sig (d = " << Sig->size1 << ") when initializing MVnormBeta" << endl;
		exit(1);
	}
	
	_X.resize(_blkStart->size());
	_scale.resize(_blkStart->size());
	for (size_t iEl = 0; iEl < _blkStart->size(); iEl++) {
		_X[iEl] = gsl_matrix_column(pred, iCl[iEl]);
		gsl_blas_ddot(&(_X[iEl]).vector, &(_X[iEl]).vector, &(_scale[iEl]));
		
		gsl_vector *bH  = gsl_vector_alloc(_eachVec[iEl].vector.size);
		gsl_matrix *chl = gsl_matrix_alloc(_eachVec[iEl].vector.size, _eachVec[iEl].vector.size);
		
		gsl_matrix_const_view sigSub  = gsl_matrix_const_submatrix(Sig, (*_blkStart)[iEl], (*_blkStart)[iEl], _eachVec[iEl].vector.size, _eachVec[iEl].vector.size);
		gsl_matrix_const_view respSub = gsl_matrix_const_submatrix(resp.dMat(), 0, (*_blkStart)[iEl], resp.Ndata(), _eachVec[iEl].vector.size);
		
		gsl_blas_dgemv(CblasTrans, 1.0/_scale[iEl], &respSub.matrix, &(_X[iEl]).vector, 0.0, bH);
		
		gsl_matrix_memcpy(chl, &sigSub.matrix);
		gsl_matrix_scale(chl, 1.0/_scale[iEl]);
		gsl_linalg_cholesky_decomp(chl);
		
		MVgauss(bH, chl, r, &(_eachVec[iEl]).vector);
		
		gsl_vector_free(bH);
		gsl_matrix_free(chl);

	}
}

MVnormBetaBlk::MVnormBetaBlk(const gsl_matrix *resp, gsl_matrix *pred, const vector<size_t> &iCl, const gsl_matrix *Sig, const vector<size_t> &blkStart, const gsl_rng *r, gsl_matrix *bet, const size_t &iRw){
	_upLevel  = 0;
	_d        = resp->size2;
	_vec      = gsl_matrix_row(bet, iRw);
	_blkStart = &blkStart;
	
	_eachVec.resize(_blkStart->size());
	
	vector<size_t> len;
	for (size_t iBlk = 1; iBlk <= (*_blkStart).size(); iBlk++) {
		iBlk == (*_blkStart).size() ? len.push_back(_d - (*_blkStart)[iBlk - 1]) : len.push_back((*_blkStart)[iBlk] - (*_blkStart)[iBlk - 1]);
	}
	
	for (size_t iBlk = 0; iBlk < _blkStart->size(); iBlk++) {
		_eachVec[iBlk] = gsl_vector_subvector(&_vec.vector, (*_blkStart)[iBlk], len[iBlk]);
	}
	
	if (pred->size1 != resp->size1) {
		cerr << "ERROR: unequal number of rows in predictor (" << pred->size1 << ") and response (" << resp->size1 << ") when initializing MVnormBeta" << endl;
		exit(1);
	}
	else if (resp->size2 != Sig->size1){
		cerr << "ERROR: incompatible matrices *resp (d = " << resp->size2 << ") and *Sig (d = " << Sig->size1 << ") when initializing MVnormBeta" << endl;
		exit(1);
	}
	
	_X.resize(_blkStart->size());
	_scale.resize(_blkStart->size());
	for (size_t iEl = 0; iEl < _blkStart->size(); iEl++) {
		_X[iEl] = gsl_matrix_column(pred, iCl[iEl]);
		gsl_blas_ddot(&(_X[iEl]).vector, &(_X[iEl]).vector, &(_scale[iEl]));
		
		gsl_vector *bH  = gsl_vector_alloc(_eachVec[iEl].vector.size);
		gsl_matrix *chl = gsl_matrix_alloc(_eachVec[iEl].vector.size, _eachVec[iEl].vector.size);
		
		gsl_matrix_const_view sigSub  = gsl_matrix_const_submatrix(Sig, (*_blkStart)[iEl], (*_blkStart)[iEl], _eachVec[iEl].vector.size, _eachVec[iEl].vector.size);
		gsl_matrix_const_view respSub = gsl_matrix_const_submatrix(resp, 0, (*_blkStart)[iEl], resp->size1, _eachVec[iEl].vector.size);
		
		gsl_blas_dgemv(CblasTrans, 1.0/_scale[iEl], &respSub.matrix, &(_X[iEl]).vector, 0.0, bH);
		
		gsl_matrix_memcpy(chl, &sigSub.matrix);
		gsl_matrix_scale(chl, 1.0/_scale[iEl]);
		gsl_linalg_cholesky_decomp(chl);
		
		MVgauss(bH, chl, r, &(_eachVec[iEl]).vector);
		
		gsl_vector_free(bH);
		gsl_matrix_free(chl);
		
	}
	
}
MVnormBetaBlk::MVnormBetaBlk(const gsl_matrix *resp, gsl_matrix *pred, const vector<size_t> &iCl, const gsl_matrix *Sig, const vector<size_t> &blkStart, const gsl_rng *r, const size_t &up, gsl_matrix *bet, const size_t &iRw){
	_upLevel  = &up;
	_d        = resp->size2;
	_vec      = gsl_matrix_row(bet, iRw);
	_blkStart = &blkStart;
	
	_eachVec.resize(_blkStart->size());
	
	vector<size_t> len;
	for (size_t iBlk = 1; iBlk <= (*_blkStart).size(); iBlk++) {
		iBlk == (*_blkStart).size() ? len.push_back(_d - (*_blkStart)[iBlk - 1]) : len.push_back((*_blkStart)[iBlk] - (*_blkStart)[iBlk - 1]);
	}
	
	for (size_t iBlk = 0; iBlk < _blkStart->size(); iBlk++) {
		_eachVec[iBlk] = gsl_vector_subvector(&_vec.vector, (*_blkStart)[iBlk], len[iBlk]);
	}
	
	if (pred->size1 != resp->size1) {
		cerr << "ERROR: unequal number of rows in predictor (" << pred->size1 << ") and response (" << resp->size1 << ") when initializing MVnormBeta" << endl;
		exit(1);
	}
	else if (resp->size2 != Sig->size1){
		cerr << "ERROR: incompatible matrices *resp (d = " << resp->size2 << ") and *Sig (d = " << Sig->size1 << ") when initializing MVnormBeta" << endl;
		exit(1);
	}
	
	_X.resize(_blkStart->size());
	_scale.resize(_blkStart->size());
	for (size_t iEl = 0; iEl < _blkStart->size(); iEl++) {
		_X[iEl] = gsl_matrix_column(pred, iCl[iEl]);
		gsl_blas_ddot(&(_X[iEl]).vector, &(_X[iEl]).vector, &(_scale[iEl]));
		
		gsl_vector *bH  = gsl_vector_alloc(_eachVec[iEl].vector.size);
		gsl_matrix *chl = gsl_matrix_alloc(_eachVec[iEl].vector.size, _eachVec[iEl].vector.size);
		
		gsl_matrix_const_view sigSub  = gsl_matrix_const_submatrix(Sig, (*_blkStart)[iEl], (*_blkStart)[iEl], _eachVec[iEl].vector.size, _eachVec[iEl].vector.size);
		gsl_matrix_const_view respSub = gsl_matrix_const_submatrix(resp, 0, (*_blkStart)[iEl], resp->size1, _eachVec[iEl].vector.size);
		
		gsl_blas_dgemv(CblasTrans, 1.0/_scale[iEl], &respSub.matrix, &(_X[iEl]).vector, 0.0, bH);
		
		gsl_matrix_memcpy(chl, &sigSub.matrix);
		gsl_matrix_scale(chl, 1.0/_scale[iEl]);
		gsl_linalg_cholesky_decomp(chl);
		
		MVgauss(bH, chl, r, &(_eachVec[iEl]).vector);
		
		gsl_vector_free(bH);
		gsl_matrix_free(chl);
		
	}
}

void MVnormBetaBlk::update(const Grp &dat, const SigmaI &SigIb, const gsl_rng *r){
	for (size_t iElm = 0; iElm < _eachVec.size(); iElm++) {
		gsl_matrix_const_view SigIsub = gsl_matrix_const_submatrix(SigIb.getMat(), (*_blkStart)[iElm], (*_blkStart)[iElm], _eachVec[iElm].vector.size, _eachVec[iElm].vector.size);
		gsl_matrix_const_view respSub = gsl_matrix_const_submatrix(dat.dMat(), 0, (*_blkStart)[iElm], dat.Ndata(), _eachVec[iElm].vector.size);
		gsl_matrix *SigSum = gsl_matrix_alloc(_eachVec[iElm].vector.size, _eachVec[iElm].vector.size);
		gsl_vector *tmpV   = gsl_vector_alloc(_eachVec[iElm].vector.size);
		
		gsl_matrix_memcpy(SigSum, &SigIsub.matrix);
		gsl_matrix_scale(SigSum, _scale[iElm]);
		gsl_linalg_cholesky_decomp(SigSum);
		gsl_linalg_cholesky_invert(SigSum);
		gsl_linalg_cholesky_decomp(SigSum);
		
		gsl_blas_dgemv(CblasTrans, 1.0/_scale[iElm], &respSub.matrix, &(_X[iElm]).vector, 0.0, tmpV);
		
		MVgauss(tmpV, SigSum, r, &(_eachVec[iElm]).vector);
		
		gsl_matrix_free(SigSum);
		gsl_vector_free(tmpV);

	}
}
void MVnormBetaBlk::update(const Grp &dat, const Qgrp &q, const SigmaI &SigIb, const gsl_rng *r){
	for (size_t iElm = 0; iElm < _eachVec.size(); iElm++) {
		gsl_matrix_const_view SigIsub = gsl_matrix_const_submatrix(SigIb.getMat(), (*_blkStart)[iElm], (*_blkStart)[iElm], _eachVec[iElm].vector.size, _eachVec[iElm].vector.size);
		gsl_matrix_const_view respSub = gsl_matrix_const_submatrix(dat.dMat(), 0, (*_blkStart)[iElm], dat.Ndata(), _eachVec[iElm].vector.size);
		gsl_matrix *SigSum = gsl_matrix_alloc(_eachVec[iElm].vector.size, _eachVec[iElm].vector.size);
		gsl_vector *tmpV   = gsl_vector_alloc(_eachVec[iElm].vector.size);
		gsl_vector *XQ     = gsl_vector_alloc(_X[iElm].vector.size);
		double xQx = 0.0;
		
		for (int iDat = 0; iDat < _X[iElm].vector.size; iDat++) {
			gsl_vector_set(XQ, iDat, q[iDat]*gsl_vector_get(&(_X[iElm]).vector, iDat));
		}
		
		gsl_blas_ddot(XQ, &(_X[iElm]).vector, &xQx);
		xQx = 1.0/xQx;
		
		gsl_blas_dgemv(CblasTrans, xQx, &respSub.matrix, XQ, 0.0, tmpV);
		
		gsl_matrix_memcpy(SigSum, &SigIsub.matrix);
		gsl_linalg_cholesky_decomp(SigSum);
		gsl_linalg_cholesky_invert(SigSum);
		gsl_matrix_scale(SigSum, xQx);
		gsl_linalg_cholesky_decomp(SigSum);
		
		MVgauss(tmpV, SigSum, r, &(_eachVec[iElm]).vector);
		
		gsl_matrix_free(SigSum);
		gsl_vector_free(XQ);
		gsl_vector_free(tmpV);

	}
}
// 0-mean prior methods
void MVnormBetaBlk::update(const Grp &dat, const SigmaI &SigIb, const SigmaI &SigIp, const gsl_rng *r){
	for (size_t iElm = 0; iElm < _eachVec.size(); iElm++) {
		gsl_matrix_const_view SigIsb = gsl_matrix_const_submatrix(SigIb.getMat(), (*_blkStart)[iElm], (*_blkStart)[iElm], _eachVec[iElm].vector.size, _eachVec[iElm].vector.size);
		gsl_matrix_const_view SigIsp = gsl_matrix_const_submatrix(SigIp.getMat(), (*_blkStart)[iElm], (*_blkStart)[iElm], _eachVec[iElm].vector.size, _eachVec[iElm].vector.size);
		gsl_matrix_const_view respSub = gsl_matrix_const_submatrix(dat.dMat(), 0, (*_blkStart)[iElm], dat.Ndata(), _eachVec[iElm].vector.size);
		gsl_matrix *SigSum = gsl_matrix_alloc(_eachVec[iElm].vector.size, _eachVec[iElm].vector.size);
		gsl_vector *tmpV   = gsl_vector_alloc(_eachVec[iElm].vector.size);
		gsl_vector *bH     = gsl_vector_alloc(_eachVec[iElm].vector.size);
		
		gsl_matrix_memcpy(SigSum, &SigIsb.matrix);
		gsl_matrix_scale(SigSum, _scale[iElm]);
		gsl_matrix_add(SigSum, &SigIsp.matrix);
		gsl_linalg_cholesky_decomp(SigSum);
		gsl_linalg_cholesky_invert(SigSum);
		
		gsl_blas_dgemv(CblasTrans, 1.0, &respSub.matrix, &(_X[iElm]).vector, 0.0, bH);
		gsl_blas_dsymv(CblasLower, 1.0, &SigIsb.matrix, bH, 0.0, tmpV);
		gsl_blas_dsymv(CblasLower, 1.0, SigSum, tmpV, 0.0, bH);
		
		gsl_linalg_cholesky_decomp(SigSum);
		MVgauss(bH, SigSum, r, &(_eachVec[iElm]).vector);
		
		gsl_matrix_free(SigSum);
		gsl_vector_free(tmpV);
		gsl_vector_free(bH);

	}
}
void MVnormBetaBlk::update(const Grp &dat, const SigmaI &SigIb, const double &qPr, const SigmaI &SigIp, const gsl_rng *r){
	for (size_t iElm = 0; iElm < _eachVec.size(); iElm++) {
		gsl_matrix_const_view SigIsb = gsl_matrix_const_submatrix(SigIb.getMat(), (*_blkStart)[iElm], (*_blkStart)[iElm], _eachVec[iElm].vector.size, _eachVec[iElm].vector.size);
		gsl_matrix_const_view SigIsp = gsl_matrix_const_submatrix(SigIp.getMat(), (*_blkStart)[iElm], (*_blkStart)[iElm], _eachVec[iElm].vector.size, _eachVec[iElm].vector.size);
		gsl_matrix_const_view respSub = gsl_matrix_const_submatrix(dat.dMat(), 0, (*_blkStart)[iElm], dat.Ndata(), _eachVec[iElm].vector.size);
		gsl_matrix *SigSum = gsl_matrix_alloc(_eachVec[iElm].vector.size, _eachVec[iElm].vector.size);
		gsl_matrix *SigPr  = gsl_matrix_alloc(_eachVec[iElm].vector.size, _eachVec[iElm].vector.size);
		gsl_vector *tmpV   = gsl_vector_alloc(_eachVec[iElm].vector.size);
		gsl_vector *bH     = gsl_vector_alloc(_eachVec[iElm].vector.size);
		
		gsl_matrix_memcpy(SigSum, &SigIsb.matrix);
		gsl_matrix_scale(SigSum, _scale[iElm]);
		gsl_matrix_memcpy(SigPr, &SigIsp.matrix);
		gsl_matrix_scale(SigPr, qPr);
		gsl_matrix_add(SigSum, SigPr);
		gsl_linalg_cholesky_decomp(SigSum);
		gsl_linalg_cholesky_invert(SigSum);
		
		gsl_blas_dgemv(CblasTrans, 1.0, &respSub.matrix, &(_X[iElm]).vector, 0.0, bH);
		gsl_blas_dsymv(CblasLower, 1.0, &SigIsb.matrix, bH, 0.0, tmpV);
		gsl_blas_dsymv(CblasLower, 1.0, SigSum, tmpV, 0.0, bH);
		
		gsl_linalg_cholesky_decomp(SigSum);
		MVgauss(bH, SigSum, r, &(_eachVec[iElm]).vector);
		
		gsl_matrix_free(SigSum);
		gsl_matrix_free(SigPr);
		gsl_vector_free(tmpV);
		gsl_vector_free(bH);

	}
}
void MVnormBetaBlk::update(const Grp &dat, const Qgrp &q, const SigmaI &SigIb, const SigmaI &SigIp, const gsl_rng *r){
	for (size_t iElm = 0; iElm < _eachVec.size(); iElm++) {
		gsl_matrix_const_view SigIsb = gsl_matrix_const_submatrix(SigIb.getMat(), (*_blkStart)[iElm], (*_blkStart)[iElm], _eachVec[iElm].vector.size, _eachVec[iElm].vector.size);
		gsl_matrix_const_view SigIsp = gsl_matrix_const_submatrix(SigIp.getMat(), (*_blkStart)[iElm], (*_blkStart)[iElm], _eachVec[iElm].vector.size, _eachVec[iElm].vector.size);
		gsl_matrix_const_view respSub = gsl_matrix_const_submatrix(dat.dMat(), 0, (*_blkStart)[iElm], dat.Ndata(), _eachVec[iElm].vector.size);
		gsl_matrix *SigSum = gsl_matrix_alloc(_eachVec[iElm].vector.size, _eachVec[iElm].vector.size);
		gsl_vector *tmpV   = gsl_vector_alloc(_eachVec[iElm].vector.size);
		gsl_vector *bH     = gsl_vector_alloc(_eachVec[iElm].vector.size);
		gsl_vector *XQ     = gsl_vector_alloc(_X[iElm].vector.size);
		double xQx = 0.0;
		
		for (int iDat = 0; iDat < _X[iElm].vector.size; iDat++) {
			gsl_vector_set(XQ, iDat, q[iDat]*gsl_vector_get(&(_X[iElm]).vector, iDat));
		}
		
		gsl_blas_ddot(XQ, &(_X[iElm]).vector, &xQx);
		
		gsl_matrix_memcpy(SigSum, &SigIsb.matrix);
		gsl_matrix_scale(SigSum, xQx);
		gsl_matrix_add(SigSum, &SigIsp.matrix);
		gsl_linalg_cholesky_decomp(SigSum);
		gsl_linalg_cholesky_invert(SigSum);
		
		gsl_blas_dgemv(CblasTrans, 1.0, &respSub.matrix, XQ, 0.0, bH);
		gsl_blas_dsymv(CblasLower, 1.0, &SigIsb.matrix, bH, 0.0, tmpV);
		gsl_blas_dsymv(CblasLower, 1.0, SigSum, tmpV, 0.0, bH);
		
		gsl_linalg_cholesky_decomp(SigSum);
		MVgauss(bH, SigSum, r, &(_eachVec[iElm]).vector);
		
		gsl_matrix_free(SigSum);
		gsl_vector_free(XQ);
		gsl_vector_free(tmpV);
		gsl_vector_free(bH);

	}
}
void MVnormBetaBlk::update(const Grp &dat, const Qgrp &q, const SigmaI &SigIb, const double &qPr, const SigmaI &SigIp, const gsl_rng *r){
	for (size_t iElm = 0; iElm < _eachVec.size(); iElm++) {
		gsl_matrix_const_view SigIsb = gsl_matrix_const_submatrix(SigIb.getMat(), (*_blkStart)[iElm], (*_blkStart)[iElm], _eachVec[iElm].vector.size, _eachVec[iElm].vector.size);
		gsl_matrix_const_view SigIsp = gsl_matrix_const_submatrix(SigIp.getMat(), (*_blkStart)[iElm], (*_blkStart)[iElm], _eachVec[iElm].vector.size, _eachVec[iElm].vector.size);
		gsl_matrix_const_view respSub = gsl_matrix_const_submatrix(dat.dMat(), 0, (*_blkStart)[iElm], dat.Ndata(), _eachVec[iElm].vector.size);
		gsl_matrix *SigSum = gsl_matrix_alloc(_eachVec[iElm].vector.size, _eachVec[iElm].vector.size);
		gsl_matrix *SigPr  = gsl_matrix_alloc(_eachVec[iElm].vector.size, _eachVec[iElm].vector.size);
		gsl_vector *tmpV   = gsl_vector_alloc(_eachVec[iElm].vector.size);
		gsl_vector *bH     = gsl_vector_alloc(_eachVec[iElm].vector.size);
		gsl_vector *XQ     = gsl_vector_alloc(_X[iElm].vector.size);
		double xQx = 0.0;
		
		for (int iDat = 0; iDat < _X[iElm].vector.size; iDat++) {
			gsl_vector_set(XQ, iDat, q[iDat]*gsl_vector_get(&(_X[iElm]).vector, iDat));
		}
		
		gsl_blas_ddot(XQ, &(_X[iElm]).vector, &xQx);
		
		gsl_matrix_memcpy(SigSum, &SigIsb.matrix);
		gsl_matrix_scale(SigSum, xQx);
		gsl_matrix_memcpy(SigPr, &SigIsp.matrix);
		gsl_matrix_scale(SigPr, qPr);
		gsl_matrix_add(SigSum, SigPr);
		gsl_linalg_cholesky_decomp(SigSum);
		gsl_linalg_cholesky_invert(SigSum);
		
		gsl_blas_dgemv(CblasTrans, 1.0, &respSub.matrix, XQ, 0.0, bH);
		gsl_blas_dsymv(CblasLower, 1.0, &SigIsb.matrix, bH, 0.0, tmpV);
		gsl_blas_dsymv(CblasLower, 1.0, SigSum, tmpV, 0.0, bH);
		
		gsl_linalg_cholesky_decomp(SigSum);
		MVgauss(bH, SigSum, r, &(_eachVec[iElm]).vector);
		
		gsl_matrix_free(SigSum);
		gsl_matrix_free(SigPr);
		gsl_vector_free(XQ);
		gsl_vector_free(tmpV);
		gsl_vector_free(bH);

	}
}
// non-zero mean prior methods
void MVnormBetaBlk::update(const Grp &dat, const SigmaI &SigIb, const Grp &muPr, const SigmaI &SigIp, const gsl_rng *r){
	for (size_t iElm = 0; iElm < _eachVec.size(); iElm++) {
		gsl_matrix_const_view SigIsb = gsl_matrix_const_submatrix(SigIb.getMat(), (*_blkStart)[iElm], (*_blkStart)[iElm], _eachVec[iElm].vector.size, _eachVec[iElm].vector.size);
		gsl_matrix_const_view SigIsp = gsl_matrix_const_submatrix(SigIp.getMat(), (*_blkStart)[iElm], (*_blkStart)[iElm], _eachVec[iElm].vector.size, _eachVec[iElm].vector.size);
		gsl_matrix_const_view respSub = gsl_matrix_const_submatrix(dat.dMat(), 0, (*_blkStart)[iElm], dat.Ndata(), _eachVec[iElm].vector.size);
		gsl_vector_const_view muSub  = gsl_vector_const_subvector(muPr[*_upLevel]->getVec(), (*_blkStart)[iElm], _eachVec[iElm].vector.size);
		gsl_matrix *SigSum = gsl_matrix_alloc(_eachVec[iElm].vector.size, _eachVec[iElm].vector.size);
		gsl_vector *tmpV   = gsl_vector_alloc(_eachVec[iElm].vector.size);
		gsl_vector *bH     = gsl_vector_alloc(_eachVec[iElm].vector.size);
		
		gsl_matrix_memcpy(SigSum, &SigIsb.matrix);
		gsl_matrix_scale(SigSum, _scale[iElm]);
		gsl_matrix_add(SigSum, &SigIsp.matrix);
		gsl_linalg_cholesky_decomp(SigSum);
		gsl_linalg_cholesky_invert(SigSum);
		
		gsl_blas_dgemv(CblasTrans, 1.0, &respSub.matrix, &(_X[iElm]).vector, 0.0, bH);
		gsl_blas_dsymv(CblasLower, 1.0, &SigIsb.matrix, bH, 0.0, tmpV);
		gsl_blas_dsymv(CblasLower, 1.0, &SigIsp.matrix, &muSub.vector, 1.0, tmpV);
		
		gsl_blas_dsymv(CblasLower, 1.0, SigSum, tmpV, 0.0, bH);
		
		gsl_linalg_cholesky_decomp(SigSum);
		MVgauss(bH, SigSum, r, &(_eachVec[iElm]).vector);
		
		gsl_matrix_free(SigSum);
		gsl_vector_free(tmpV);
		gsl_vector_free(bH);

	}
}
void MVnormBetaBlk::update(const Grp &dat, const SigmaI &SigIb, const Grp &muPr, const double &qPr, const SigmaI &SigIp, const gsl_rng *r){
	for (size_t iElm = 0; iElm < _eachVec.size(); iElm++) {
		gsl_matrix_const_view SigIsb = gsl_matrix_const_submatrix(SigIb.getMat(), (*_blkStart)[iElm], (*_blkStart)[iElm], _eachVec[iElm].vector.size, _eachVec[iElm].vector.size);
		gsl_matrix_const_view SigIsp = gsl_matrix_const_submatrix(SigIp.getMat(), (*_blkStart)[iElm], (*_blkStart)[iElm], _eachVec[iElm].vector.size, _eachVec[iElm].vector.size);
		gsl_matrix_const_view respSub = gsl_matrix_const_submatrix(dat.dMat(), 0, (*_blkStart)[iElm], dat.Ndata(), _eachVec[iElm].vector.size);
		gsl_vector_const_view muSub  = gsl_vector_const_subvector(muPr[*_upLevel]->getVec(), (*_blkStart)[iElm], _eachVec[iElm].vector.size);
		gsl_matrix *SigSum = gsl_matrix_alloc(_eachVec[iElm].vector.size, _eachVec[iElm].vector.size);
		gsl_matrix *SigPr  = gsl_matrix_alloc(_eachVec[iElm].vector.size, _eachVec[iElm].vector.size);
		gsl_vector *tmpV   = gsl_vector_alloc(_eachVec[iElm].vector.size);
		gsl_vector *bH     = gsl_vector_alloc(_eachVec[iElm].vector.size);
		
		gsl_matrix_memcpy(SigSum, &SigIsb.matrix);
		gsl_matrix_scale(SigSum, _scale[iElm]);
		gsl_matrix_memcpy(SigPr, &SigIsp.matrix);
		gsl_matrix_scale(SigPr, qPr);
		gsl_matrix_add(SigSum, SigPr);
		gsl_linalg_cholesky_decomp(SigSum);
		gsl_linalg_cholesky_invert(SigSum);
		
		gsl_blas_dgemv(CblasTrans, 1.0, &respSub.matrix, &(_X[iElm]).vector, 0.0, bH);
		gsl_blas_dsymv(CblasLower, 1.0, &SigIsb.matrix, bH, 0.0, tmpV);
		gsl_blas_dsymv(CblasLower, 1.0, SigPr, &muSub.vector, 1.0, tmpV);
		
		gsl_blas_dsymv(CblasLower, 1.0, SigSum, tmpV, 0.0, bH);
		
		gsl_linalg_cholesky_decomp(SigSum);
		MVgauss(bH, SigSum, r, &(_eachVec[iElm]).vector);
		
		gsl_matrix_free(SigSum);
		gsl_matrix_free(SigPr);
		gsl_vector_free(tmpV);
		gsl_vector_free(bH);

	}
}
void MVnormBetaBlk::update(const Grp &dat, const Qgrp &q, const SigmaI &SigIb, const Grp &muPr, const SigmaI &SigIp, const gsl_rng *r){
	for (size_t iElm = 0; iElm < _eachVec.size(); iElm++) {
		gsl_matrix_const_view SigIsb = gsl_matrix_const_submatrix(SigIb.getMat(), (*_blkStart)[iElm], (*_blkStart)[iElm], _eachVec[iElm].vector.size, _eachVec[iElm].vector.size);
		gsl_matrix_const_view SigIsp = gsl_matrix_const_submatrix(SigIp.getMat(), (*_blkStart)[iElm], (*_blkStart)[iElm], _eachVec[iElm].vector.size, _eachVec[iElm].vector.size);
		gsl_matrix_const_view respSub = gsl_matrix_const_submatrix(dat.dMat(), 0, (*_blkStart)[iElm], dat.Ndata(), _eachVec[iElm].vector.size);
		gsl_vector_const_view muSub  = gsl_vector_const_subvector(muPr[*_upLevel]->getVec(), (*_blkStart)[iElm], _eachVec[iElm].vector.size);
		gsl_matrix *SigSum = gsl_matrix_alloc(_eachVec[iElm].vector.size, _eachVec[iElm].vector.size);
		gsl_vector *tmpV   = gsl_vector_alloc(_eachVec[iElm].vector.size);
		gsl_vector *bH     = gsl_vector_alloc(_eachVec[iElm].vector.size);
		gsl_vector *XQ     = gsl_vector_alloc(_X[iElm].vector.size);
		double xQx = 0.0;
		
		for (int iDat = 0; iDat < _X[iElm].vector.size; iDat++) {
			gsl_vector_set(XQ, iDat, q[iDat]*gsl_vector_get(&(_X[iElm]).vector, iDat));
		}
		
		gsl_blas_ddot(XQ, &(_X[iElm]).vector, &xQx);
		
		gsl_matrix_memcpy(SigSum, &SigIsb.matrix);
		gsl_matrix_scale(SigSum, xQx);
		gsl_matrix_add(SigSum, &SigIsp.matrix);
		gsl_linalg_cholesky_decomp(SigSum);
		gsl_linalg_cholesky_invert(SigSum);
		
		gsl_blas_dgemv(CblasTrans, 1.0, &respSub.matrix, XQ, 0.0, bH);
		gsl_blas_dsymv(CblasLower, 1.0, &SigIsb.matrix, bH, 0.0, tmpV);
		gsl_blas_dsymv(CblasLower, 1.0, &SigIsp.matrix, &muSub.vector, 1.0, tmpV);
		
		gsl_blas_dsymv(CblasLower, 1.0, SigSum, tmpV, 0.0, bH);
		
		gsl_linalg_cholesky_decomp(SigSum);
		MVgauss(bH, SigSum, r, &(_eachVec[iElm]).vector);
		
		gsl_matrix_free(SigSum);
		gsl_vector_free(XQ);
		gsl_vector_free(tmpV);
		gsl_vector_free(bH);

	}
}
void MVnormBetaBlk::update(const Grp &dat, const Qgrp &q, const SigmaI &SigIb, const Grp &muPr, const double &qPr, const SigmaI &SigIp, const gsl_rng *r){
	for (size_t iElm = 0; iElm < _eachVec.size(); iElm++) {
		gsl_matrix_const_view SigIsb = gsl_matrix_const_submatrix(SigIb.getMat(), (*_blkStart)[iElm], (*_blkStart)[iElm], _eachVec[iElm].vector.size, _eachVec[iElm].vector.size);
		gsl_matrix_const_view SigIsp = gsl_matrix_const_submatrix(SigIp.getMat(), (*_blkStart)[iElm], (*_blkStart)[iElm], _eachVec[iElm].vector.size, _eachVec[iElm].vector.size);
		gsl_matrix_const_view respSub = gsl_matrix_const_submatrix(dat.dMat(), 0, (*_blkStart)[iElm], dat.Ndata(), _eachVec[iElm].vector.size);
		gsl_vector_const_view muSub  = gsl_vector_const_subvector(muPr[*_upLevel]->getVec(), (*_blkStart)[iElm], _eachVec[iElm].vector.size);
		gsl_matrix *SigSum = gsl_matrix_alloc(_eachVec[iElm].vector.size, _eachVec[iElm].vector.size);
		gsl_matrix *SigPr  = gsl_matrix_alloc(_eachVec[iElm].vector.size, _eachVec[iElm].vector.size);
		gsl_vector *tmpV   = gsl_vector_alloc(_eachVec[iElm].vector.size);
		gsl_vector *bH     = gsl_vector_alloc(_eachVec[iElm].vector.size);
		gsl_vector *XQ     = gsl_vector_alloc(_X[iElm].vector.size);
		double xQx = 0.0;
		
		for (int iDat = 0; iDat < _X[iElm].vector.size; iDat++) {
			gsl_vector_set(XQ, iDat, q[iDat]*gsl_vector_get(&(_X[iElm]).vector, iDat));
		}
		
		gsl_blas_ddot(XQ, &(_X[iElm]).vector, &xQx);
		
		gsl_matrix_memcpy(SigSum, &SigIsb.matrix);
		gsl_matrix_scale(SigSum, xQx);
		gsl_matrix_memcpy(SigPr, &SigIsp.matrix);
		gsl_matrix_scale(SigPr, qPr);
		gsl_matrix_add(SigSum, SigPr);
		gsl_linalg_cholesky_decomp(SigSum);
		gsl_linalg_cholesky_invert(SigSum);
		
		gsl_blas_dgemv(CblasTrans, 1.0, &respSub.matrix, XQ, 0.0, bH);
		gsl_blas_dsymv(CblasLower, 1.0, &SigIsb.matrix, bH, 0.0, tmpV);
		gsl_blas_dsymv(CblasLower, 1.0, SigPr, &muSub.vector, 1.0, tmpV);
		
		gsl_blas_dsymv(CblasLower, 1.0, SigSum, tmpV, 0.0, bH);
		
		gsl_linalg_cholesky_decomp(SigSum);
		MVgauss(bH, SigSum, r, &(_eachVec[iElm]).vector);
		
		gsl_matrix_free(SigSum);
		gsl_matrix_free(SigPr);
		gsl_vector_free(XQ);
		gsl_vector_free(tmpV);
		gsl_vector_free(bH);

	}
}

/*
 *	MVnormBetaFtBlk methods
 */

void MVnormBetaFtBlk::update(const Grp &dat, const SigmaI &SigIb, const gsl_rng *r){
	gsl_matrix *rsd = gsl_matrix_alloc(dat.Ndata(), _d);
	gsl_matrix_memcpy(rsd, dat.dMat());
	gsl_matrix_sub(rsd, &_fitted.matrix);
	
	for (size_t iElm = 0; iElm < _eachVec.size(); iElm++) {
		gsl_matrix_const_view SigIsub = gsl_matrix_const_submatrix(SigIb.getMat(), (*_blkStart)[iElm], (*_blkStart)[iElm], _eachVec[iElm].vector.size, _eachVec[iElm].vector.size);
		gsl_matrix_const_view respSub = gsl_matrix_const_submatrix(rsd, 0, (*_blkStart)[iElm], dat.Ndata(), _eachVec[iElm].vector.size);
		gsl_matrix *SigSum = gsl_matrix_alloc(_eachVec[iElm].vector.size, _eachVec[iElm].vector.size);
		gsl_vector *tmpV   = gsl_vector_alloc(_eachVec[iElm].vector.size);
		
		gsl_matrix_memcpy(SigSum, &SigIsub.matrix);
		gsl_matrix_scale(SigSum, _scale[iElm]);
		gsl_linalg_cholesky_decomp(SigSum);
		gsl_linalg_cholesky_invert(SigSum);
		gsl_linalg_cholesky_decomp(SigSum);
		
		gsl_blas_dgemv(CblasTrans, 1.0/_scale[iElm], &respSub.matrix, &(_X[iElm]).vector, 0.0, tmpV);
		
		MVgauss(tmpV, SigSum, r, &(_eachVec[iElm]).vector);
		
		gsl_matrix_free(SigSum);
		gsl_vector_free(tmpV);
		
	}
	gsl_matrix_free(rsd);
}
void MVnormBetaFtBlk::update(const Grp &dat, const Qgrp &q, const SigmaI &SigIb, const gsl_rng *r){
	gsl_matrix *rsd = gsl_matrix_alloc(dat.Ndata(), _d);
	gsl_matrix_memcpy(rsd, dat.dMat());
	gsl_matrix_sub(rsd, &_fitted.matrix);
	
	for (size_t iElm = 0; iElm < _eachVec.size(); iElm++) {
		gsl_matrix_const_view SigIsub = gsl_matrix_const_submatrix(SigIb.getMat(), (*_blkStart)[iElm], (*_blkStart)[iElm], _eachVec[iElm].vector.size, _eachVec[iElm].vector.size);
		gsl_matrix_const_view respSub = gsl_matrix_const_submatrix(rsd, 0, (*_blkStart)[iElm], dat.Ndata(), _eachVec[iElm].vector.size);
		gsl_matrix *SigSum = gsl_matrix_alloc(_eachVec[iElm].vector.size, _eachVec[iElm].vector.size);
		gsl_vector *tmpV   = gsl_vector_alloc(_eachVec[iElm].vector.size);
		gsl_vector *XQ     = gsl_vector_alloc(_X[iElm].vector.size);
		double xQx = 0.0;
		
		for (int iDat = 0; iDat < _X[iElm].vector.size; iDat++) {
			gsl_vector_set(XQ, iDat, q[iDat]*gsl_vector_get(&(_X[iElm]).vector, iDat));
		}
		
		gsl_blas_ddot(XQ, &(_X[iElm]).vector, &xQx);
		xQx = 1.0/xQx;
		
		gsl_blas_dgemv(CblasTrans, xQx, &respSub.matrix, XQ, 0.0, tmpV);
		
		gsl_matrix_memcpy(SigSum, &SigIsub.matrix);
		gsl_linalg_cholesky_decomp(SigSum);
		gsl_linalg_cholesky_invert(SigSum);
		gsl_matrix_scale(SigSum, xQx);
		gsl_linalg_cholesky_decomp(SigSum);
		
		MVgauss(tmpV, SigSum, r, &(_eachVec[iElm]).vector);
		
		gsl_matrix_free(SigSum);
		gsl_vector_free(XQ);
		gsl_vector_free(tmpV);
		
	}
	gsl_matrix_free(rsd);
}
// 0-mean prior methods
void MVnormBetaFtBlk::update(const Grp &dat, const SigmaI &SigIb, const SigmaI &SigIp, const gsl_rng *r){
	gsl_matrix *rsd = gsl_matrix_alloc(dat.Ndata(), _d);
	gsl_matrix_memcpy(rsd, dat.dMat());
	gsl_matrix_sub(rsd, &_fitted.matrix);
	
	for (size_t iElm = 0; iElm < _eachVec.size(); iElm++) {
		gsl_matrix_const_view SigIsb = gsl_matrix_const_submatrix(SigIb.getMat(), (*_blkStart)[iElm], (*_blkStart)[iElm], _eachVec[iElm].vector.size, _eachVec[iElm].vector.size);
		gsl_matrix_const_view SigIsp = gsl_matrix_const_submatrix(SigIp.getMat(), (*_blkStart)[iElm], (*_blkStart)[iElm], _eachVec[iElm].vector.size, _eachVec[iElm].vector.size);
		gsl_matrix_const_view respSub = gsl_matrix_const_submatrix(rsd, 0, (*_blkStart)[iElm], dat.Ndata(), _eachVec[iElm].vector.size);
		gsl_matrix *SigSum = gsl_matrix_alloc(_eachVec[iElm].vector.size, _eachVec[iElm].vector.size);
		gsl_vector *tmpV   = gsl_vector_alloc(_eachVec[iElm].vector.size);
		gsl_vector *bH     = gsl_vector_alloc(_eachVec[iElm].vector.size);
		
		gsl_matrix_memcpy(SigSum, &SigIsb.matrix);
		gsl_matrix_scale(SigSum, _scale[iElm]);
		gsl_matrix_add(SigSum, &SigIsp.matrix);
		gsl_linalg_cholesky_decomp(SigSum);
		gsl_linalg_cholesky_invert(SigSum);
		
		gsl_blas_dgemv(CblasTrans, 1.0, &respSub.matrix, &(_X[iElm]).vector, 0.0, bH);
		gsl_blas_dsymv(CblasLower, 1.0, &SigIsb.matrix, bH, 0.0, tmpV);
		gsl_blas_dsymv(CblasLower, 1.0, SigSum, tmpV, 0.0, bH);
		
		gsl_linalg_cholesky_decomp(SigSum);
		MVgauss(bH, SigSum, r, &(_eachVec[iElm]).vector);
		
		gsl_matrix_free(SigSum);
		gsl_vector_free(tmpV);
		gsl_vector_free(bH);
		
	}
	gsl_matrix_free(rsd);
}
void MVnormBetaFtBlk::update(const Grp &dat, const SigmaI &SigIb, const double &qPr, const SigmaI &SigIp, const gsl_rng *r){
	gsl_matrix *rsd = gsl_matrix_alloc(dat.Ndata(), _d);
	gsl_matrix_memcpy(rsd, dat.dMat());
	gsl_matrix_sub(rsd, &_fitted.matrix);
	
	for (size_t iElm = 0; iElm < _eachVec.size(); iElm++) {
		gsl_matrix_const_view SigIsb = gsl_matrix_const_submatrix(SigIb.getMat(), (*_blkStart)[iElm], (*_blkStart)[iElm], _eachVec[iElm].vector.size, _eachVec[iElm].vector.size);
		gsl_matrix_const_view SigIsp = gsl_matrix_const_submatrix(SigIp.getMat(), (*_blkStart)[iElm], (*_blkStart)[iElm], _eachVec[iElm].vector.size, _eachVec[iElm].vector.size);
		gsl_matrix_const_view respSub = gsl_matrix_const_submatrix(rsd, 0, (*_blkStart)[iElm], dat.Ndata(), _eachVec[iElm].vector.size);
		gsl_matrix *SigSum = gsl_matrix_alloc(_eachVec[iElm].vector.size, _eachVec[iElm].vector.size);
		gsl_matrix *SigPr  = gsl_matrix_alloc(_eachVec[iElm].vector.size, _eachVec[iElm].vector.size);
		gsl_vector *tmpV   = gsl_vector_alloc(_eachVec[iElm].vector.size);
		gsl_vector *bH     = gsl_vector_alloc(_eachVec[iElm].vector.size);
		
		gsl_matrix_memcpy(SigSum, &SigIsb.matrix);
		gsl_matrix_scale(SigSum, _scale[iElm]);
		gsl_matrix_memcpy(SigPr, &SigIsp.matrix);
		gsl_matrix_scale(SigPr, qPr);
		gsl_matrix_add(SigSum, SigPr);
		gsl_linalg_cholesky_decomp(SigSum);
		gsl_linalg_cholesky_invert(SigSum);
		
		gsl_blas_dgemv(CblasTrans, 1.0, &respSub.matrix, &(_X[iElm]).vector, 0.0, bH);
		gsl_blas_dsymv(CblasLower, 1.0, &SigIsb.matrix, bH, 0.0, tmpV);
		gsl_blas_dsymv(CblasLower, 1.0, SigSum, tmpV, 0.0, bH);
		
		gsl_linalg_cholesky_decomp(SigSum);
		MVgauss(bH, SigSum, r, &(_eachVec[iElm]).vector);
		
		gsl_matrix_free(SigSum);
		gsl_matrix_free(SigPr);
		gsl_vector_free(tmpV);
		gsl_vector_free(bH);
		
	}
	gsl_matrix_free(rsd);
}
void MVnormBetaFtBlk::update(const Grp &dat, const Qgrp &q, const SigmaI &SigIb, const SigmaI &SigIp, const gsl_rng *r){
	gsl_matrix *rsd = gsl_matrix_alloc(dat.Ndata(), _d);
	gsl_matrix_memcpy(rsd, dat.dMat());
	gsl_matrix_sub(rsd, &_fitted.matrix);
	
	for (size_t iElm = 0; iElm < _eachVec.size(); iElm++) {
		gsl_matrix_const_view SigIsb = gsl_matrix_const_submatrix(SigIb.getMat(), (*_blkStart)[iElm], (*_blkStart)[iElm], _eachVec[iElm].vector.size, _eachVec[iElm].vector.size);
		gsl_matrix_const_view SigIsp = gsl_matrix_const_submatrix(SigIp.getMat(), (*_blkStart)[iElm], (*_blkStart)[iElm], _eachVec[iElm].vector.size, _eachVec[iElm].vector.size);
		gsl_matrix_const_view respSub = gsl_matrix_const_submatrix(rsd, 0, (*_blkStart)[iElm], dat.Ndata(), _eachVec[iElm].vector.size);
		gsl_matrix *SigSum = gsl_matrix_alloc(_eachVec[iElm].vector.size, _eachVec[iElm].vector.size);
		gsl_vector *tmpV   = gsl_vector_alloc(_eachVec[iElm].vector.size);
		gsl_vector *bH     = gsl_vector_alloc(_eachVec[iElm].vector.size);
		gsl_vector *XQ     = gsl_vector_alloc(_X[iElm].vector.size);
		double xQx = 0.0;
		
		for (int iDat = 0; iDat < _X[iElm].vector.size; iDat++) {
			gsl_vector_set(XQ, iDat, q[iDat]*gsl_vector_get(&(_X[iElm]).vector, iDat));
		}
		
		gsl_blas_ddot(XQ, &(_X[iElm]).vector, &xQx);
		
		gsl_matrix_memcpy(SigSum, &SigIsb.matrix);
		gsl_matrix_scale(SigSum, xQx);
		gsl_matrix_add(SigSum, &SigIsp.matrix);
		gsl_linalg_cholesky_decomp(SigSum);
		gsl_linalg_cholesky_invert(SigSum);
		
		gsl_blas_dgemv(CblasTrans, 1.0, &respSub.matrix, XQ, 0.0, bH);
		gsl_blas_dsymv(CblasLower, 1.0, &SigIsb.matrix, bH, 0.0, tmpV);
		gsl_blas_dsymv(CblasLower, 1.0, SigSum, tmpV, 0.0, bH);
		
		gsl_linalg_cholesky_decomp(SigSum);
		MVgauss(bH, SigSum, r, &(_eachVec[iElm]).vector);
		
		gsl_matrix_free(SigSum);
		gsl_vector_free(XQ);
		gsl_vector_free(tmpV);
		gsl_vector_free(bH);
		
	}
	gsl_matrix_free(rsd);
}
void MVnormBetaFtBlk::update(const Grp &dat, const Qgrp &q, const SigmaI &SigIb, const double &qPr, const SigmaI &SigIp, const gsl_rng *r){
	gsl_matrix *rsd = gsl_matrix_alloc(dat.Ndata(), _d);
	gsl_matrix_memcpy(rsd, dat.dMat());
	gsl_matrix_sub(rsd, &_fitted.matrix);
	
	for (size_t iElm = 0; iElm < _eachVec.size(); iElm++) {
		gsl_matrix_const_view SigIsb = gsl_matrix_const_submatrix(SigIb.getMat(), (*_blkStart)[iElm], (*_blkStart)[iElm], _eachVec[iElm].vector.size, _eachVec[iElm].vector.size);
		gsl_matrix_const_view SigIsp = gsl_matrix_const_submatrix(SigIp.getMat(), (*_blkStart)[iElm], (*_blkStart)[iElm], _eachVec[iElm].vector.size, _eachVec[iElm].vector.size);
		gsl_matrix_const_view respSub = gsl_matrix_const_submatrix(rsd, 0, (*_blkStart)[iElm], dat.Ndata(), _eachVec[iElm].vector.size);
		gsl_matrix *SigSum = gsl_matrix_alloc(_eachVec[iElm].vector.size, _eachVec[iElm].vector.size);
		gsl_matrix *SigPr  = gsl_matrix_alloc(_eachVec[iElm].vector.size, _eachVec[iElm].vector.size);
		gsl_vector *tmpV   = gsl_vector_alloc(_eachVec[iElm].vector.size);
		gsl_vector *bH     = gsl_vector_alloc(_eachVec[iElm].vector.size);
		gsl_vector *XQ     = gsl_vector_alloc(_X[iElm].vector.size);
		double xQx = 0.0;
		
		for (int iDat = 0; iDat < _X[iElm].vector.size; iDat++) {
			gsl_vector_set(XQ, iDat, q[iDat]*gsl_vector_get(&(_X[iElm]).vector, iDat));
		}
		
		gsl_blas_ddot(XQ, &(_X[iElm]).vector, &xQx);
		
		gsl_matrix_memcpy(SigSum, &SigIsb.matrix);
		gsl_matrix_scale(SigSum, xQx);
		gsl_matrix_memcpy(SigPr, &SigIsp.matrix);
		gsl_matrix_scale(SigPr, qPr);
		gsl_matrix_add(SigSum, SigPr);
		gsl_linalg_cholesky_decomp(SigSum);
		gsl_linalg_cholesky_invert(SigSum);
		
		gsl_blas_dgemv(CblasTrans, 1.0, &respSub.matrix, XQ, 0.0, bH);
		gsl_blas_dsymv(CblasLower, 1.0, &SigIsb.matrix, bH, 0.0, tmpV);
		gsl_blas_dsymv(CblasLower, 1.0, SigSum, tmpV, 0.0, bH);
		
		gsl_linalg_cholesky_decomp(SigSum);
		MVgauss(bH, SigSum, r, &(_eachVec[iElm]).vector);
		
		gsl_matrix_free(SigSum);
		gsl_matrix_free(SigPr);
		gsl_vector_free(XQ);
		gsl_vector_free(tmpV);
		gsl_vector_free(bH);
		
	}
	gsl_matrix_free(rsd);
}
// non-zero mean prior methods
void MVnormBetaFtBlk::update(const Grp &dat, const SigmaI &SigIb, const Grp &muPr, const SigmaI &SigIp, const gsl_rng *r){
	gsl_matrix *rsd = gsl_matrix_alloc(dat.Ndata(), _d);
	gsl_matrix_memcpy(rsd, dat.dMat());
	gsl_matrix_sub(rsd, &_fitted.matrix);
	
	for (size_t iElm = 0; iElm < _eachVec.size(); iElm++) {
		gsl_matrix_const_view SigIsb = gsl_matrix_const_submatrix(SigIb.getMat(), (*_blkStart)[iElm], (*_blkStart)[iElm], _eachVec[iElm].vector.size, _eachVec[iElm].vector.size);
		gsl_matrix_const_view SigIsp = gsl_matrix_const_submatrix(SigIp.getMat(), (*_blkStart)[iElm], (*_blkStart)[iElm], _eachVec[iElm].vector.size, _eachVec[iElm].vector.size);
		gsl_matrix_const_view respSub = gsl_matrix_const_submatrix(rsd, 0, (*_blkStart)[iElm], dat.Ndata(), _eachVec[iElm].vector.size);
		gsl_vector_const_view muSub  = gsl_vector_const_subvector(muPr[*_upLevel]->getVec(), (*_blkStart)[iElm], _eachVec[iElm].vector.size);
		gsl_matrix *SigSum = gsl_matrix_alloc(_eachVec[iElm].vector.size, _eachVec[iElm].vector.size);
		gsl_vector *tmpV   = gsl_vector_alloc(_eachVec[iElm].vector.size);
		gsl_vector *bH     = gsl_vector_alloc(_eachVec[iElm].vector.size);
		
		gsl_matrix_memcpy(SigSum, &SigIsb.matrix);
		gsl_matrix_scale(SigSum, _scale[iElm]);
		gsl_matrix_add(SigSum, &SigIsp.matrix);
		gsl_linalg_cholesky_decomp(SigSum);
		gsl_linalg_cholesky_invert(SigSum);
		
		gsl_blas_dgemv(CblasTrans, 1.0, &respSub.matrix, &(_X[iElm]).vector, 0.0, bH);
		gsl_blas_dsymv(CblasLower, 1.0, &SigIsb.matrix, bH, 0.0, tmpV);
		gsl_blas_dsymv(CblasLower, 1.0, &SigIsp.matrix, &muSub.vector, 1.0, tmpV);
		
		gsl_blas_dsymv(CblasLower, 1.0, SigSum, tmpV, 0.0, bH);
		
		gsl_linalg_cholesky_decomp(SigSum);
		MVgauss(bH, SigSum, r, &(_eachVec[iElm]).vector);
		
		gsl_matrix_free(SigSum);
		gsl_vector_free(tmpV);
		gsl_vector_free(bH);
		
	}
	gsl_matrix_free(rsd);
}
void MVnormBetaFtBlk::update(const Grp &dat, const SigmaI &SigIb, const Grp &muPr, const double &qPr, const SigmaI &SigIp, const gsl_rng *r){
	gsl_matrix *rsd = gsl_matrix_alloc(dat.Ndata(), _d);
	gsl_matrix_memcpy(rsd, dat.dMat());
	gsl_matrix_sub(rsd, &_fitted.matrix);
	
	for (size_t iElm = 0; iElm < _eachVec.size(); iElm++) {
		gsl_matrix_const_view SigIsb = gsl_matrix_const_submatrix(SigIb.getMat(), (*_blkStart)[iElm], (*_blkStart)[iElm], _eachVec[iElm].vector.size, _eachVec[iElm].vector.size);
		gsl_matrix_const_view SigIsp = gsl_matrix_const_submatrix(SigIp.getMat(), (*_blkStart)[iElm], (*_blkStart)[iElm], _eachVec[iElm].vector.size, _eachVec[iElm].vector.size);
		gsl_matrix_const_view respSub = gsl_matrix_const_submatrix(rsd, 0, (*_blkStart)[iElm], dat.Ndata(), _eachVec[iElm].vector.size);
		gsl_vector_const_view muSub  = gsl_vector_const_subvector(muPr[*_upLevel]->getVec(), (*_blkStart)[iElm], _eachVec[iElm].vector.size);
		gsl_matrix *SigSum = gsl_matrix_alloc(_eachVec[iElm].vector.size, _eachVec[iElm].vector.size);
		gsl_matrix *SigPr  = gsl_matrix_alloc(_eachVec[iElm].vector.size, _eachVec[iElm].vector.size);
		gsl_vector *tmpV   = gsl_vector_alloc(_eachVec[iElm].vector.size);
		gsl_vector *bH     = gsl_vector_alloc(_eachVec[iElm].vector.size);
		
		gsl_matrix_memcpy(SigSum, &SigIsb.matrix);
		gsl_matrix_scale(SigSum, _scale[iElm]);
		gsl_matrix_memcpy(SigPr, &SigIsp.matrix);
		gsl_matrix_scale(SigPr, qPr);
		gsl_matrix_add(SigSum, SigPr);
		gsl_linalg_cholesky_decomp(SigSum);
		gsl_linalg_cholesky_invert(SigSum);
		
		gsl_blas_dgemv(CblasTrans, 1.0, &respSub.matrix, &(_X[iElm]).vector, 0.0, bH);
		gsl_blas_dsymv(CblasLower, 1.0, &SigIsb.matrix, bH, 0.0, tmpV);
		gsl_blas_dsymv(CblasLower, 1.0, SigPr, &muSub.vector, 1.0, tmpV);
		
		gsl_blas_dsymv(CblasLower, 1.0, SigSum, tmpV, 0.0, bH);
		
		gsl_linalg_cholesky_decomp(SigSum);
		MVgauss(bH, SigSum, r, &(_eachVec[iElm]).vector);
		
		gsl_matrix_free(SigSum);
		gsl_matrix_free(SigPr);
		gsl_vector_free(tmpV);
		gsl_vector_free(bH);
		
	}
	gsl_matrix_free(rsd);
}
void MVnormBetaFtBlk::update(const Grp &dat, const Qgrp &q, const SigmaI &SigIb, const Grp &muPr, const SigmaI &SigIp, const gsl_rng *r){
	gsl_matrix *rsd = gsl_matrix_alloc(dat.Ndata(), _d);
	gsl_matrix_memcpy(rsd, dat.dMat());
	gsl_matrix_sub(rsd, &_fitted.matrix);
	
	for (size_t iElm = 0; iElm < _eachVec.size(); iElm++) {
		gsl_matrix_const_view SigIsb = gsl_matrix_const_submatrix(SigIb.getMat(), (*_blkStart)[iElm], (*_blkStart)[iElm], _eachVec[iElm].vector.size, _eachVec[iElm].vector.size);
		gsl_matrix_const_view SigIsp = gsl_matrix_const_submatrix(SigIp.getMat(), (*_blkStart)[iElm], (*_blkStart)[iElm], _eachVec[iElm].vector.size, _eachVec[iElm].vector.size);
		gsl_matrix_const_view respSub = gsl_matrix_const_submatrix(rsd, 0, (*_blkStart)[iElm], dat.Ndata(), _eachVec[iElm].vector.size);
		gsl_vector_const_view muSub  = gsl_vector_const_subvector(muPr[*_upLevel]->getVec(), (*_blkStart)[iElm], _eachVec[iElm].vector.size);
		gsl_matrix *SigSum = gsl_matrix_alloc(_eachVec[iElm].vector.size, _eachVec[iElm].vector.size);
		gsl_vector *tmpV   = gsl_vector_alloc(_eachVec[iElm].vector.size);
		gsl_vector *bH     = gsl_vector_alloc(_eachVec[iElm].vector.size);
		gsl_vector *XQ     = gsl_vector_alloc(_X[iElm].vector.size);
		double xQx = 0.0;
		
		for (int iDat = 0; iDat < _X[iElm].vector.size; iDat++) {
			gsl_vector_set(XQ, iDat, q[iDat]*gsl_vector_get(&(_X[iElm]).vector, iDat));
		}
		
		gsl_blas_ddot(XQ, &(_X[iElm]).vector, &xQx);
		
		gsl_matrix_memcpy(SigSum, &SigIsb.matrix);
		gsl_matrix_scale(SigSum, xQx);
		gsl_matrix_add(SigSum, &SigIsp.matrix);
		gsl_linalg_cholesky_decomp(SigSum);
		gsl_linalg_cholesky_invert(SigSum);
		
		gsl_blas_dgemv(CblasTrans, 1.0, &respSub.matrix, XQ, 0.0, bH);
		gsl_blas_dsymv(CblasLower, 1.0, &SigIsb.matrix, bH, 0.0, tmpV);
		gsl_blas_dsymv(CblasLower, 1.0, &SigIsp.matrix, &muSub.vector, 1.0, tmpV);
		
		gsl_blas_dsymv(CblasLower, 1.0, SigSum, tmpV, 0.0, bH);
		
		gsl_linalg_cholesky_decomp(SigSum);
		MVgauss(bH, SigSum, r, &(_eachVec[iElm]).vector);
		
		gsl_matrix_free(SigSum);
		gsl_vector_free(XQ);
		gsl_vector_free(tmpV);
		gsl_vector_free(bH);
		
	}
	gsl_matrix_free(rsd);
}
void MVnormBetaFtBlk::update(const Grp &dat, const Qgrp &q, const SigmaI &SigIb, const Grp &muPr, const double &qPr, const SigmaI &SigIp, const gsl_rng *r){
	gsl_matrix *rsd = gsl_matrix_alloc(dat.Ndata(), _d);
	gsl_matrix_memcpy(rsd, dat.dMat());
	gsl_matrix_sub(rsd, &_fitted.matrix);
	
	for (size_t iElm = 0; iElm < _eachVec.size(); iElm++) {
		gsl_matrix_const_view SigIsb = gsl_matrix_const_submatrix(SigIb.getMat(), (*_blkStart)[iElm], (*_blkStart)[iElm], _eachVec[iElm].vector.size, _eachVec[iElm].vector.size);
		gsl_matrix_const_view SigIsp = gsl_matrix_const_submatrix(SigIp.getMat(), (*_blkStart)[iElm], (*_blkStart)[iElm], _eachVec[iElm].vector.size, _eachVec[iElm].vector.size);
		gsl_matrix_const_view respSub = gsl_matrix_const_submatrix(rsd, 0, (*_blkStart)[iElm], dat.Ndata(), _eachVec[iElm].vector.size);
		gsl_vector_const_view muSub  = gsl_vector_const_subvector(muPr[*_upLevel]->getVec(), (*_blkStart)[iElm], _eachVec[iElm].vector.size);
		gsl_matrix *SigSum = gsl_matrix_alloc(_eachVec[iElm].vector.size, _eachVec[iElm].vector.size);
		gsl_matrix *SigPr  = gsl_matrix_alloc(_eachVec[iElm].vector.size, _eachVec[iElm].vector.size);
		gsl_vector *tmpV   = gsl_vector_alloc(_eachVec[iElm].vector.size);
		gsl_vector *bH     = gsl_vector_alloc(_eachVec[iElm].vector.size);
		gsl_vector *XQ     = gsl_vector_alloc(_X[iElm].vector.size);
		double xQx = 0.0;
		
		for (int iDat = 0; iDat < _X[iElm].vector.size; iDat++) {
			gsl_vector_set(XQ, iDat, q[iDat]*gsl_vector_get(&(_X[iElm]).vector, iDat));
		}
		
		gsl_blas_ddot(XQ, &(_X[iElm]).vector, &xQx);
		
		gsl_matrix_memcpy(SigSum, &SigIsb.matrix);
		gsl_matrix_scale(SigSum, xQx);
		gsl_matrix_memcpy(SigPr, &SigIsp.matrix);
		gsl_matrix_scale(SigPr, qPr);
		gsl_matrix_add(SigSum, SigPr);
		gsl_linalg_cholesky_decomp(SigSum);
		gsl_linalg_cholesky_invert(SigSum);
		
		gsl_blas_dgemv(CblasTrans, 1.0, &respSub.matrix, XQ, 0.0, bH);
		gsl_blas_dsymv(CblasLower, 1.0, &SigIsb.matrix, bH, 0.0, tmpV);
		gsl_blas_dsymv(CblasLower, 1.0, SigPr, &muSub.vector, 1.0, tmpV);
		
		gsl_blas_dsymv(CblasLower, 1.0, SigSum, tmpV, 0.0, bH);
		
		gsl_linalg_cholesky_decomp(SigSum);
		MVgauss(bH, SigSum, r, &(_eachVec[iElm]).vector);
		
		gsl_matrix_free(SigSum);
		gsl_matrix_free(SigPr);
		gsl_vector_free(XQ);
		gsl_vector_free(tmpV);
		gsl_vector_free(bH);
		
	}
	gsl_matrix_free(rsd);
}

/*
 *	RanIndex methods
 */

// Constructors
RanIndex::RanIndex() : _idx(0), _vecInd(1, 0) {
	const gsl_rng_type *T = gsl_rng_mt19937;
	_r = gsl_rng_alloc(T);
	gsl_rng_set(_r, time(NULL)+rdtsc());

}
RanIndex::RanIndex(const gsl_vector_int *lInd, const size_t &Ntot, const size_t &Nup){
	_idx.resize(Nup);
	for (size_t iLN = 0; iLN < Ntot; iLN++) {
		_idx[gsl_vector_int_get(lInd, iLN)].push_back(iLN);
		_vecInd.push_back(gsl_vector_int_get(lInd, iLN));
	}
	const gsl_rng_type *T = gsl_rng_mt19937;
	_r = gsl_rng_alloc(T);
	gsl_rng_set(_r, time(NULL)+rdtsc());
}

RanIndex::RanIndex(const size_t &Ntot){
	_idx.resize(1);
	for (size_t iLN = 0; iLN < Ntot; iLN++) {
		_idx[0].push_back(iLN);
		_vecInd.push_back(0);
	}
	const gsl_rng_type *T = gsl_rng_mt19937;
	_r = gsl_rng_alloc(T);
	gsl_rng_set(_r, time(NULL)+rdtsc());
}

RanIndex::RanIndex(const size_t &Ntot, const size_t &Nup){
	if (Nup > Ntot) {
		cerr << "ERROR: more upper levels than lower levels when initializing a one-to-one RanIndex" << endl;
		exit(1);
	}
	else if (Nup < Ntot){
		cerr << "WARNING: number of upper elements " << Nup << " is smaller than the number of lower elements " << Ntot << "; " << Ntot - Nup << " elements ignored" << endl;
	}
	_idx.resize(Nup);
	for (size_t iLN = 0; iLN < Ntot; iLN++) {
		_idx[iLN].push_back(iLN);
		_vecInd.push_back(iLN);
	}
	const gsl_rng_type *T = gsl_rng_mt19937;
	_r = gsl_rng_alloc(T);
	gsl_rng_set(_r, time(NULL)+rdtsc());
}

RanIndex::RanIndex(const size_t &Ntot, const size_t &Nup, const string &fileNam){
	_idx.resize(Nup);
	gsl_vector_int *tmp = gsl_vector_int_alloc(Ntot);
	
	FILE *indIn = fopen(fileNam.c_str(), "r");
	if (indIn == NULL) {
		cerr << "ERROR: cannot open index file " << fileNam << " in RanIndex" << endl;
		exit(-1);
	}
	gsl_vector_int_fread(indIn, tmp);
	fclose(indIn);
	int mnVal = gsl_vector_int_min(tmp);
	int mxVal = gsl_vector_int_max(tmp);
	
	if ((mnVal == 1) && (mxVal == Nup) ) {
		cerr << "WARNING: Looks like the saved array is base-1.  Converting to base-0" << endl;
		gsl_vector_int_add_constant(tmp, -1); // looks like the saved array is base-1 (as in R)
	}
	else if (mxVal > (Nup - 1)){
		cerr << "ERROR: the upper-level index is out of range.  Max value " << mxVal << " is bigger than " << Nup - 1 << endl;
		exit(-1);
	}
	else if (mnVal != 0) { cerr << "WARNING: hierarchical index may not be base-1 or base-0.  Check for errors." << endl; }
	
	for (size_t iLN = 0; iLN < Ntot; iLN++) {
		_idx[gsl_vector_int_get(tmp, iLN)].push_back(iLN);
		_vecInd.push_back(gsl_vector_int_get(tmp, iLN));
	}
	
	vector<size_t> zeroLen;
	for (size_t idxI = 0; idxI < _idx.size(); idxI++) {
		if (_idx[idxI].size() == 0) {
			zeroLen.push_back(idxI);
		}
	}
	
	if (zeroLen.size()) {
		cerr << "ERROR: some higher-level index values are not present. They are (on base-0): " << endl;
		for (vector<size_t>::iterator zIt = zeroLen.begin(); zIt != zeroLen.end(); ++zIt) {
			cerr << *zIt << " " << flush;
		}
		cerr << endl;
		cerr << "Remove them and re-run" << endl;
		exit(1);
	}
	const gsl_rng_type *T = gsl_rng_mt19937;
	_r = gsl_rng_alloc(T);
	gsl_rng_set(_r, time(NULL)+rdtsc());
	
	gsl_vector_int_free(tmp);
}

RanIndex::RanIndex(const size_t &Ntot, const size_t &Nup, FILE *fileStr){
	_idx.resize(Nup);
	gsl_vector_int *tmp = gsl_vector_int_alloc(Ntot);
	
	gsl_vector_int_fread(fileStr, tmp);
	
	int mnVal = gsl_vector_int_min(tmp);
	int mxVal = gsl_vector_int_max(tmp);
	
	if ((mnVal == 1) && (mxVal == Nup) ) {
		cerr << "WARNING: Looks like the saved array is base-1.  Converting to base-0" << endl;
		gsl_vector_int_add_constant(tmp, -1); // looks like the saved array is base-1 (as in R)
	}
	else if (mxVal > (Nup -1)){
		cerr << "ERROR: the upper-level index is out of range.  Max value " << mxVal << " is bigger than " << Nup - 1 << endl;
		exit(-1);
	}
	else if (mnVal != 0) { cerr << "WARNING: hierarchical index may not be base-1 or base-0.  Check for errors." << endl; }
	
	for (size_t iLN = 0; iLN < Ntot; iLN++) {
		_idx[gsl_vector_int_get(tmp, iLN)].push_back(iLN);
		_vecInd.push_back(gsl_vector_int_get(tmp, iLN));
	}
	
	const gsl_rng_type *T = gsl_rng_mt19937;
	_r = gsl_rng_alloc(T);
	gsl_rng_set(_r, time(NULL)+rdtsc());
	
	gsl_vector_int_free(tmp);
}

const vector<double> RanIndex::props() const{
	vector<double> res(_idx.size());
	double sum = 0.0;
	for (size_t iV = 0; iV < _idx.size(); iV++) {
		sum += _idx[iV].size();
	}
	for (size_t iR = 0; iR < _idx.size(); iR++) {
		res[iR] = static_cast<double>(_idx[iR].size())/sum;
	}
	
	return res;
}

void RanIndex::init(const vector< vector<size_t> > &idx, const vector< vector<size_t> > &rLD) { // rLD ignored here
	_idx = idx;
	
	// now re-populate _locInd using the new _idx
	size_t totSz = 0;
	for (vector< vector<size_t> >::iterator idxIt = _idx.begin(); idxIt != _idx.end(); ++idxIt) {
		totSz += (*idxIt).size();
	}
	
	_vecInd.resize(totSz);
	for (size_t idxEl = 0; idxEl < _idx.size(); idxEl++) {
		for (vector<size_t>::iterator elIt = _idx[idxEl].begin(); elIt != _idx[idxEl].end(); ++elIt) {
			_vecInd[*elIt] = idxEl;
		}
	}
}
/*
 *	Update functions
 */

void RanIndex::update(const Grp &theta, const Grp &mu, const vector<SigmaI> &SigI, const MixP &p){
	_idx.clear();
	_idx.resize(mu.Ndata());
	_vecInd.clear();
	
	double *scDensity = new double[mu.Ndata()];
	unsigned int *z   = new unsigned int[mu.Ndata()];
	
	for (size_t jInd = 0; jInd < theta.Ndata(); jInd++) {
		double sumDns = 0.0;
		for (int iTrk = 0; iTrk < mu.Ndata(); iTrk++) {
			scDensity[iTrk] = p[iTrk] * mu[iTrk]->density(theta[jInd], SigI[iTrk]);
			sumDns += scDensity[iTrk];
		}
		
		for (int iTrk = 0; iTrk < mu.Ndata(); iTrk++) {
			scDensity[iTrk] /= sumDns;
		}
		gsl_ran_multinomial(_r, mu.Ndata(), 1, scDensity, z);
		
		for (int iTrk = 0; iTrk < mu.Ndata(); iTrk++) {
			if (z[iTrk]) {
				_idx[iTrk].push_back(jInd);
				_vecInd.push_back(iTrk);
				break;
			}
		}
	}
	delete [] scDensity;
	delete [] z;
}

void RanIndex::update(const Grp &theta, const Grp &mu, const SigmaI &SigI, const MixP &p){
	_idx.clear();
	_idx.resize(mu.Ndata());
	_vecInd.clear();
	
	double *scDensity = new double[mu.Ndata()];
	unsigned int *z   = new unsigned int[mu.Ndata()];
	
	for (size_t jInd = 0; jInd < theta.Ndata(); jInd++) {
		double sumDns = 0.0;
		for (int iTrk = 0; iTrk < mu.Ndata()	; iTrk++) {
			scDensity[iTrk] = p[iTrk] * mu[iTrk]->density(theta[jInd], SigI);
			sumDns += scDensity[iTrk];
		}
		
		for (size_t iTrk = 0; iTrk < mu.Ndata(); iTrk++) {
			scDensity[iTrk] /= sumDns;
		}
		gsl_ran_multinomial(_r, mu.Ndata(), 1, scDensity, z);
		
		for (size_t iTrk = 0; iTrk < mu.Ndata(); iTrk++) {
			if (z[iTrk]) {
				_idx[iTrk].push_back(jInd);
				_vecInd.push_back(iTrk);
				break;
			}
		}
	}
	delete [] scDensity;
	delete [] z;
}

/*
	RanIndexVS methods
 */
RanIndexVS::RanIndexVS() : RanIndex() {
	
	_acceptDrop    = vector<bool>(4, false);
	_acceptDrop[0] = true;
	_acceptDrop[2] = true;
	
	_rejectDrop    = vector<bool>(4, false);
	_rejectDrop[2] = true;
	
	_acceptAdd    = vector<bool>(4, false);
	_acceptAdd[0] = true;
	_acceptAdd[1] = true;
	
	_rejectAdd    = vector<bool>(4, false);
	_rejectAdd[1] = true;
	
	_acceptSwap    = vector<bool>(4, false);
	_acceptSwap[0] = true;
	_acceptSwap[3] = true;
	
	_rejectSwap    = vector<bool>(4, false);
	_rejectSwap[3] = true;
}

RanIndexVS::RanIndexVS(const size_t &Ntot, const string &outFlNam, MixP &pr) : RanIndex(), _Ntp(Ntot), _numSaves(0){
	_prior = &pr;
	
	_idx.resize(2); // [0] is the vector of selected indexes, [1] -- not selected
	_vecInd.resize(Ntot);
	
	_acceptDrop    = vector<bool>(4, false);
	_acceptDrop[0] = true;
	_acceptDrop[2] = true;
	
	_rejectDrop    = vector<bool>(4, false);
	_rejectDrop[2] = true;
	
	_acceptAdd    = vector<bool>(4, false);
	_acceptAdd[0] = true;
	_acceptAdd[1] = true;
	
	_rejectAdd    = vector<bool>(4, false);
	_rejectAdd[1] = true;
	
	_acceptSwap    = vector<bool>(4, false);
	_acceptSwap[0] = true;
	_acceptSwap[3] = true;
	
	_rejectSwap    = vector<bool>(4, false);
	_rejectSwap[3] = true;
	
	_pepOutFlnam  = outFlNam;
	_mcmcPutFlNam = outFlNam;
	size_t pos = _mcmcPutFlNam.find(".");
	_mcmcPutFlNam.erase(pos, 5);
	_mcmcPutFlNam += "MetroTrk.tsv";
	remove(_mcmcPutFlNam.c_str());

}

RanIndexVS::RanIndexVS(const size_t &Ntot, const string &outFlNam) : RanIndex(), _Ntp(Ntot), _numSaves(0), _prior(0) {
	_idx.resize(2); // [0] is the vector of selected indexes, [1] -- not selected
	_vecInd.resize(Ntot);
	
	_acceptDrop    = vector<bool>(4, false);
	_acceptDrop[0] = true;
	_acceptDrop[2] = true;
	
	_rejectDrop    = vector<bool>(4, false);
	_rejectDrop[2] = true;
	
	_acceptAdd    = vector<bool>(4, false);
	_acceptAdd[0] = true;
	_acceptAdd[1] = true;
	
	_rejectAdd    = vector<bool>(4, false);
	_rejectAdd[1] = true;
	
	_acceptSwap    = vector<bool>(4, false);
	_acceptSwap[0] = true;
	_acceptSwap[3] = true;
	
	_rejectSwap    = vector<bool>(4, false);
	_rejectSwap[3] = true;
	
	_pepOutFlnam  = outFlNam;
	_mcmcPutFlNam = outFlNam;
	size_t pos = _mcmcPutFlNam.find(".");
	_mcmcPutFlNam.erase(pos, 5);
	_mcmcPutFlNam += "MetroTrk.tsv";
	remove(_mcmcPutFlNam.c_str());
	
}

size_t RanIndexVS::_proposal(const size_t &Ntot, const int &Nmn, const gsl_rng *r){
	size_t res = 0;
	unsigned int tst = gsl_ran_bernoulli(r, 0.7);  // geometric or uniform?  0.3U + 0.7G
	if (tst) {
		double mn = 1.0/(Nmn + 1.0);
		res = rtgeom(mn, Ntot, r);
	}
	else {
		res = floor(gsl_ran_flat(r, 0.0, Ntot));
	}
	return res;
}

void RanIndexVS::props(vector<double> &prp) const{
	prp.resize(2);
	for (size_t iV = 0; iV < 2; iV++) {
		prp[iV] = static_cast<double>(_idx[iV].size())/(static_cast<double>(_idx[0].size() + _idx[1].size()));
	}
}

void RanIndexVS::init(const vector< vector<size_t> > &idx, const vector< vector<size_t> > &rLD){
	_relLD = rLD;
	_idx   = idx;
	// now re-populate _locInd using the new _idx
	size_t totSz = 0;
	for (vector< vector<size_t> >::iterator idxIt = _idx.begin(); idxIt != _idx.end(); ++idxIt) {
		totSz += (*idxIt).size();
	}
	
	_pep.resize(totSz, 0.0);
	_vecInd.resize(totSz);
	for (size_t idxEl = 0; idxEl < _idx.size(); idxEl++) {
		for (vector<size_t>::iterator elIt = _idx[idxEl].begin(); elIt != _idx[idxEl].end(); ++elIt) {
			_vecInd[*elIt] = idxEl;
		}
	}
}

void RanIndexVS::save(const Grp &y, const BetaGrpBVSR *theta, const SigmaI &SigIe){
	_numSaves += 1.0;
	
	ofstream Mout;
	Mout.open(_mcmcPutFlNam.c_str(), ios::app);
	
	vector<bool>::iterator trkIt = _mcmcTrack.begin();
	Mout << *trkIt << flush;
	++trkIt;
	for (; trkIt != _mcmcTrack.end(); ++trkIt) {
		Mout << " " << *trkIt << flush;
	}
	Mout << endl;
	Mout.close();
	
	vector<double> curPep(_pep.size(), 1.0);
	for (vector<size_t>::iterator slIt = _idx[0].begin(); slIt != _idx[0].end(); ++slIt) {
		double lam = theta->lnOddsRat(y, SigIe, *slIt) + log((*_prior)[0]) - log(1.0 - (*_prior)[0]);
		curPep[*slIt] = 1.0/(1.0 + exp(lam));
	}
	gsl_vector_view oP = gsl_vector_view_array(_pep.data(), _pep.size());
	gsl_vector_view nP = gsl_vector_view_array(curPep.data(), curPep.size());
	gsl_vector_add(&oP.vector, &nP.vector);
	
}

void RanIndexVS::save(const Grp &y, const Grp &theta, const SigmaI &SigIe){
	_numSaves += 1.0;
	
	vector<double> curPep(_pep.size(), 1.0);
	for (vector<size_t>::iterator slIt = _idx[0].begin(); slIt != _idx[0].end(); ++slIt) {
		double lam = theta.lnOddsRat(y, SigIe, *slIt);
		curPep[*slIt] = 1.0/(1.0 + exp(lam));
	}
	gsl_vector_view oP = gsl_vector_view_array(_pep.data(), _pep.size());
	gsl_vector_view nP = gsl_vector_view_array(curPep.data(), curPep.size());
	gsl_vector_add(&oP.vector, &nP.vector);
	
}

void RanIndexVS::dump(){
	if (_pep.size() && _relLD.size() && _numSaves) {
		gsl_vector_view pepV = gsl_vector_view_array(_pep.data(), _pep.size());
		gsl_vector_scale(&pepV.vector, 1.0/_numSaves);
		
		gsl_vector *svPep = gsl_vector_alloc(_Ntp);
		gsl_vector_set_all(svPep, 1.0);
		
		for (size_t iPck = 0; iPck < _relLD.size(); iPck++) {
			for (vector<size_t>::iterator ldIt = _relLD[iPck].begin(); ldIt != _relLD[iPck].end(); ++ldIt) {  // setting all the SNPs in LD with the focal one the same pep
				gsl_vector_set(svPep, *ldIt, _pep[iPck]);
			}
		}
		
		FILE *pepOut = fopen(_pepOutFlnam.c_str(), "w");
		gsl_vector_fwrite(pepOut, svPep);
		fclose(pepOut);
		gsl_vector_free(svPep);
		
		string relFlNam = _pepOutFlnam;
		size_t pos      = relFlNam.find(".");
		relFlNam.erase(pos, 5);
		relFlNam += "Picked.tsv";
		remove(relFlNam.c_str());
		
		ofstream Rout;
		Rout.open(relFlNam.c_str(), ios::app);
		for (vector< vector<size_t> >::iterator pIt = _relLD.begin(); pIt != _relLD.end(); ++pIt) {
			for (vector<size_t>::iterator iLD = (*pIt).begin(); iLD != (*pIt).end(); ++iLD) {
				Rout << (*pIt)[0] + 1 << "\t" << (*iLD) + 1 << endl;
			}
		}
		Rout.close();
		
	}

}

void RanIndexVS::update(const Grp &y, const SigmaI &SigIe, BetaGrpBVSR *theta, const SigmaI &SigIp){
	double u = gsl_ran_flat(_r, 0.0, 1.0);
	if (u <= 0.4) { // add predictor
		if (_idx[1].size() == 0) {
			_mcmcTrack = _rejectAdd;
		}
		else {
			size_t iAdd  = floor(gsl_ran_flat(_r, 0.0, static_cast<double>(_idx[1].size() - 1)));  // uniform proposal for the index of the element to be added
			size_t addID = _idx[1][iAdd];
			
#ifdef MYDEBUG
			if (_vecInd[addID] == 0) {
				cerr << "WARNING!!! Seem to be trying to add element " << addID << " that is already in the model" << endl;
			}
#endif
			double lnr = 0.5*(theta->_MGkernel(y, SigIe) - theta->_MGkernel(y, SigIe, SigIp, addID)) + log((*_prior)[0]) - log(1.0 - (*_prior)[0]);
			if (lnr >= 0.0) {
				_mcmcTrack = _acceptAdd;
				
				vector<size_t>::iterator selIt = _idx[1].begin();
				selIt += iAdd;
				_idx[1].erase(selIt);
				_idx[0].push_back(addID);
				_vecInd[addID] = 0;
				
				/*
					now modify the relevant theta components to account for the acceptance of the new element
				 */
				gsl_vector_view Xcol = gsl_matrix_column(theta->_Xmat, addID);
				gsl_vector_view bRow = gsl_matrix_row(theta->_valueMat, addID);
#ifdef MYDEBUG
				if (!gsl_matrix_isnull(theta->_tmpXb)) {
					cerr << "ERROR: _tmpXb matrix is non-null when proposing element addition on " << __LINE__ << " of " << __FILE__ << endl;
					exit(1);
				}
#endif
				gsl_blas_dger(1.0, &Xcol.vector, &bRow.vector, theta->_tmpXb); // this assumes that _tmpXb is zeroed out before-hand
				gsl_matrix_add(theta->_fittedAll, theta->_tmpXb);
				for (size_t iEl = 0; iEl < _idx[0].size() - 1; iEl++) { // since we know that the new element is last, and we don't need to change it's Xb
					gsl_matrix_view curXb = gsl_matrix_view_array((theta->_fittedEach[_idx[0][iEl]]).data(), (theta->_fittedAll)->size1, (theta->_fittedAll)->size2);
					gsl_matrix_add(&curXb.matrix, theta->_tmpXb);
				}
				for (vector<size_t>::iterator absIt = _idx[1].begin(); absIt != _idx[1].end(); ++absIt) {
					gsl_matrix_view curXb = gsl_matrix_view_array((theta->_fittedEach[*absIt]).data(), (theta->_fittedAll)->size1, (theta->_fittedAll)->size2);
					gsl_matrix_memcpy(&curXb.matrix, theta->_fittedAll);
				}
				gsl_matrix_set_zero(theta->_tmpXb);
				
			}
			else if (gsl_ran_bernoulli(_r, exp(lnr))){
				_mcmcTrack = _acceptAdd;
				
				vector<size_t>::iterator selIt = _idx[1].begin();
				selIt += iAdd;
				_idx[1].erase(selIt);
				_idx[0].push_back(addID);
				_vecInd[addID] = 0;
				
				/*
				    now modify the relevant theta components to account for the acceptance of the new element
				 */
				gsl_vector_view Xcol = gsl_matrix_column(theta->_Xmat, addID);
				gsl_vector_view bRow = gsl_matrix_row(theta->_valueMat, addID);
#ifdef MYDEBUG
				if (!gsl_matrix_isnull(theta->_tmpXb)) {
					cerr << "ERROR: _tmpXb matrix is non-null when proposing element addition on " << __LINE__ << " of " << __FILE__ << endl;
					exit(1);
				}
#endif
				gsl_blas_dger(1.0, &Xcol.vector, &bRow.vector, theta->_tmpXb); // this assumes that _tmpXb is zeroed out before-hand
				gsl_matrix_add(theta->_fittedAll, theta->_tmpXb);
				for (size_t iEl = 0; iEl < _idx[0].size() - 1; iEl++) { // since we know that the new element is last, and we don't need to change it's Xb
					gsl_matrix_view curXb = gsl_matrix_view_array((theta->_fittedEach[_idx[0][iEl]]).data(), (theta->_fittedAll)->size1, (theta->_fittedAll)->size2);
					gsl_matrix_add(&curXb.matrix, theta->_tmpXb);
				}
				for (vector<size_t>::iterator absIt = _idx[1].begin(); absIt != _idx[1].end(); ++absIt) {
					gsl_matrix_view curXb = gsl_matrix_view_array((theta->_fittedEach[*absIt]).data(), (theta->_fittedAll)->size1, (theta->_fittedAll)->size2);
					gsl_matrix_memcpy(&curXb.matrix, theta->_fittedAll);
				}
				gsl_matrix_set_zero(theta->_tmpXb);
			}
			else {
				_mcmcTrack = _rejectAdd;
				gsl_vector_view bRow = gsl_matrix_row(theta->_valueMat, addID); // revert the unsuccessfully proposed theta elements to zero
				gsl_vector_set_zero(&bRow.vector);
			}
		}
	}
	else if (u <= 0.8){ // toss predictor
		if (_idx[0].size() <= y.dMat()->size2) {
			_mcmcTrack = _rejectDrop;
		}
		else {
			size_t iSub  = floor(gsl_ran_flat(_r, 0.0, static_cast<double>(_idx[0].size() - 1)));  // uniform proposal for the index of the element to be dropped
			size_t subID = _idx[0][iSub];
#ifdef MYDEBUG
			if (_vecInd[subID] == 1) {
				cerr << "WARNING!!! Seem to be trying to drop element " << subID << " that is not in the model" << endl;
			}
#endif

			double lnr = 0.5*(theta->_MGkernel(y, SigIe, subID) - theta->_MGkernel(y, SigIe)) - log((*_prior)[0]) + log(1.0 - (*_prior)[0]);
			if (lnr >= 0.0) {
				_mcmcTrack = _acceptDrop;
				
				vector<size_t>::iterator selIt = _idx[0].begin();
				selIt += iSub;
				_idx[0].erase(selIt);
				_idx[1].push_back(subID);
				_vecInd[subID] = 1;
				
				/*
				    now modify the relevant theta components to account for the element deletion
				 */
				gsl_vector_view Xcol = gsl_matrix_column(theta->_Xmat, subID);
				gsl_vector_view bRow = gsl_matrix_row(theta->_valueMat, subID);
#ifdef MYDEBUG
				if (!gsl_matrix_isnull(theta->_tmpXb)) {
					cerr << "ERROR: _tmpXb matrix is non-null when proposing element deletion on " << __LINE__ << " of " << __FILE__ << endl;
					exit(1);
				}
#endif
				gsl_blas_dger(1.0, &Xcol.vector, &bRow.vector, theta->_tmpXb); // this assumes that _tmpXb is zeroed out before-hand
				gsl_matrix_view drpXb = gsl_matrix_view_array((theta->_fittedEach)[subID].data(), (theta->_fittedAll)->size1, (theta->_fittedAll)->size2);
				gsl_matrix_memcpy(theta->_fittedAll, &drpXb.matrix);  // the new _fittedAll is the partial Xb corresponding to subID
				for (size_t iEl = 0; iEl < _idx[0].size(); iEl++) {
					gsl_matrix_view curXb = gsl_matrix_view_array((theta->_fittedEach[_idx[0][iEl]]).data(), (theta->_fittedAll)->size1, (theta->_fittedAll)->size2);
					gsl_matrix_sub(&curXb.matrix, theta->_tmpXb);
				}
				for (vector<size_t>::iterator absIt = _idx[1].begin(); absIt != _idx[1].end() - 1; ++absIt) { // since we know that the new element is last, and we don't need to change it's Xb
					gsl_matrix_view curXb = gsl_matrix_view_array((theta->_fittedEach[*absIt]).data(), (theta->_fittedAll)->size1, (theta->_fittedAll)->size2);
					gsl_matrix_memcpy(&curXb.matrix, &drpXb.matrix);
				}
				gsl_vector_set_zero(&bRow.vector); // zero out the dropped element
				gsl_matrix_set_zero(theta->_tmpXb);

			}
			else if (gsl_ran_bernoulli(_r, exp(lnr))){
				_mcmcTrack = _acceptDrop;
				
				vector<size_t>::iterator selIt = _idx[0].begin();
				selIt += iSub;
				_idx[0].erase(selIt);
				_idx[1].push_back(subID);
				_vecInd[subID] = 1;
				
				/*
				    now modify the relevant theta components to account for the element deletion
				 */
				gsl_vector_view Xcol = gsl_matrix_column(theta->_Xmat, subID);
				gsl_vector_view bRow = gsl_matrix_row(theta->_valueMat, subID);
#ifdef MYDEBUG
				if (!gsl_matrix_isnull(theta->_tmpXb)) {
					cerr << "ERROR: _tmpXb matrix is non-null when proposing element deletion on " << __LINE__ << " of " << __FILE__ << endl;
					exit(1);
				}
#endif
				gsl_blas_dger(1.0, &Xcol.vector, &bRow.vector, theta->_tmpXb); // this assumes that _tmpXb is zeroed out before-hand
				gsl_matrix_view drpXb = gsl_matrix_view_array((theta->_fittedEach)[subID].data(), (theta->_fittedAll)->size1, (theta->_fittedAll)->size2);
				gsl_matrix_memcpy(theta->_fittedAll, &drpXb.matrix);  // the new _fittedAll is the partial Xb corresponding to subID
				for (size_t iEl = 0; iEl < _idx[0].size(); iEl++) {
					gsl_matrix_view curXb = gsl_matrix_view_array((theta->_fittedEach[_idx[0][iEl]]).data(), (theta->_fittedAll)->size1, (theta->_fittedAll)->size2);
					gsl_matrix_sub(&curXb.matrix, theta->_tmpXb);
				}
				for (vector<size_t>::iterator absIt = _idx[1].begin(); absIt != _idx[1].end() - 1; ++absIt) { // since we know that the new element is last, and we don't need to change it's Xb
					gsl_matrix_view curXb = gsl_matrix_view_array((theta->_fittedEach[*absIt]).data(), (theta->_fittedAll)->size1, (theta->_fittedAll)->size2);
					gsl_matrix_memcpy(&curXb.matrix, &drpXb.matrix);
				}
				gsl_vector_set_zero(&bRow.vector);
				gsl_matrix_set_zero(theta->_tmpXb);
			}
			else {
				_mcmcTrack = _rejectDrop;
			}
		}
	}
	else { // swap predictors
		size_t iAdd  = floor(gsl_ran_flat(_r, 0.0, static_cast<double>(_idx[1].size() - 1)));
		size_t iSub  = floor(gsl_ran_flat(_r, 0.0, static_cast<double>(_idx[0].size() - 1)));
		size_t addID = _idx[1][iAdd];
		size_t subID = _idx[0][iSub];
#ifdef MYDEBUG
		if ((_vecInd[subID] == 1) || (_vecInd[addID] == 0)) {
			cerr << "WARNING!!! Seem to be trying to swap element " << subID << "(state: " << _vecInd[subID] << ") for element " << addID << "(state: " << _vecInd[addID] << ")" << endl;
		}
#endif
		
		double lnr =  0.5*(theta->_MGkernel(y, SigIe, subID) - theta->_MGkernel(y, SigIe, SigIp, addID));
		if (lnr >= 0.0) {
			_mcmcTrack = _acceptSwap;
			
			vector<size_t>::iterator selIt = _idx[0].begin();
			selIt += iSub;
			_idx[0].erase(selIt);
			_idx[1].push_back(subID);
			_vecInd[subID] = 1;
			vector<size_t>::iterator usIt = _idx[1].begin();
			usIt += iAdd;
			_idx[1].erase(usIt);
			_idx[0].push_back(addID);
			_vecInd[addID] = 0;
			
			/*
				now modify the relevant theta components to account for the element swap
			 */
#ifdef MYDEBUG
			if (!gsl_matrix_isnull(theta->_tmpXb)) {
				cerr << "ERROR: _tmpXb matrix is non-null when proposing element swap on " << __LINE__ << " of " << __FILE__ << endl;
				exit(1);
			}
#endif
			gsl_vector_view XcolS = gsl_matrix_column(theta->_Xmat, subID);
			gsl_vector_view bRowS = gsl_matrix_row(theta->_valueMat, subID);
			gsl_vector_view XcolA = gsl_matrix_column(theta->_Xmat, addID);
			gsl_vector_view bRowA = gsl_matrix_row(theta->_valueMat, addID);
			gsl_matrix_view drpXb = gsl_matrix_view_array((theta->_fittedEach)[subID].data(), (theta->_fittedAll)->size1, (theta->_fittedAll)->size2);
			(theta->_fittedEach)[addID] = (theta->_fittedEach)[subID]; // since there's no need add the Xb for _fittedEach[addID]
			
			gsl_blas_dger(1.0, &XcolA.vector, &bRowA.vector, theta->_tmpXb);
			gsl_matrix_memcpy(theta->_fittedAll, &drpXb.matrix);
			gsl_matrix_add(theta->_fittedAll, theta->_tmpXb);
			
			for (vector<size_t>::iterator absIt = _idx[1].begin(); absIt != _idx[1].end(); ++absIt) {
				gsl_matrix_view curXb = gsl_matrix_view_array((theta->_fittedEach[*absIt]).data(), (theta->_fittedAll)->size1, (theta->_fittedAll)->size2);
				gsl_matrix_memcpy(&curXb.matrix, theta->_fittedAll);
			}
			gsl_blas_dger(-1.0, &XcolS.vector, &bRowS.vector, theta->_tmpXb);
			for (size_t iEl = 0; iEl < _idx[0].size() - 1; iEl++) { // excluding the newly-added, which is last
				gsl_matrix_view curXb = gsl_matrix_view_array((theta->_fittedEach[_idx[0][iEl]]).data(), (theta->_fittedAll)->size1, (theta->_fittedAll)->size2);
				gsl_matrix_add(&curXb.matrix, theta->_tmpXb);
			}
			gsl_vector_set_zero(&bRowS.vector);
			gsl_matrix_set_zero(theta->_tmpXb);
		}
		else if (gsl_ran_bernoulli(_r, exp(lnr))){
			_mcmcTrack = _acceptSwap;
			
			vector<size_t>::iterator selIt = _idx[0].begin();
			selIt += iSub;
			_idx[0].erase(selIt);
			_idx[1].push_back(subID);
			_vecInd[subID] = 1;
			vector<size_t>::iterator usIt = _idx[1].begin();
			usIt += iAdd;
			_idx[1].erase(usIt);
			_idx[0].push_back(addID);
			_vecInd[addID] = 0;
			
			/*
			    now modify the relevant theta components to account for the element swap
			 */
#ifdef MYDEBUG
			if (!gsl_matrix_isnull(theta->_tmpXb)) {
				cerr << "ERROR: _tmpXb matrix is non-null when proposing element swap on " << __LINE__ << " of " << __FILE__ << endl;
				exit(1);
			}
#endif
			gsl_vector_view XcolS = gsl_matrix_column(theta->_Xmat, subID);
			gsl_vector_view bRowS = gsl_matrix_row(theta->_valueMat, subID);
			gsl_vector_view XcolA = gsl_matrix_column(theta->_Xmat, addID);
			gsl_vector_view bRowA = gsl_matrix_row(theta->_valueMat, addID);
			gsl_matrix_view drpXb = gsl_matrix_view_array((theta->_fittedEach)[subID].data(), (theta->_fittedAll)->size1, (theta->_fittedAll)->size2);
			(theta->_fittedEach)[addID] = (theta->_fittedEach)[subID]; // since there's no need add the Xb for _fittedEach[addID]
			
			gsl_blas_dger(1.0, &XcolA.vector, &bRowA.vector, theta->_tmpXb);
			gsl_matrix_memcpy(theta->_fittedAll, &drpXb.matrix);
			gsl_matrix_add(theta->_fittedAll, theta->_tmpXb);
			
			for (vector<size_t>::iterator absIt = _idx[1].begin(); absIt != _idx[1].end(); ++absIt) {
				gsl_matrix_view curXb = gsl_matrix_view_array((theta->_fittedEach[*absIt]).data(), (theta->_fittedAll)->size1, (theta->_fittedAll)->size2);
				gsl_matrix_memcpy(&curXb.matrix, theta->_fittedAll);
			}
			gsl_blas_dger(-1.0, &XcolS.vector, &bRowS.vector, theta->_tmpXb);
			for (size_t iEl = 0; iEl < _idx[0].size() - 1; iEl++) { // excluding the newly-added, which is last
				gsl_matrix_view curXb = gsl_matrix_view_array((theta->_fittedEach[_idx[0][iEl]]).data(), (theta->_fittedAll)->size1, (theta->_fittedAll)->size2);
				gsl_matrix_add(&curXb.matrix, theta->_tmpXb);
			}
			gsl_vector_set_zero(&bRowS.vector);
			gsl_matrix_set_zero(theta->_tmpXb);
		}
		else {
			_mcmcTrack = _rejectSwap;
			gsl_vector_view bRow = gsl_matrix_row(theta->_valueMat, addID); // revert the unsuccessfully proposed theta elements to zero
			gsl_vector_set_zero(&bRow.vector);
		}
	}
}

/*
 *	Apex methods
 */
Apex::Apex(){
	_Amat = gsl_matrix_alloc(1, 1);
	gsl_matrix_set_all(_Amat, 1.0);
	_SigIpr = gsl_matrix_calloc(1, 1);
	gsl_vector_view diag = gsl_matrix_diagonal(_SigIpr);
	gsl_vector_set_all(&diag.vector, 1e-6);
	_fitEach = 0;
	
	const gsl_rng_type *T = gsl_rng_mt19937;
	_r = gsl_rng_alloc(T);
	gsl_rng_set(_r, time(NULL)+rdtsc());
}
Apex::Apex(const size_t &d, vector<vector <double> > &fE){
	_Amat = gsl_matrix_calloc(d, d);
	gsl_vector_view diag = gsl_matrix_diagonal(_Amat);
	gsl_vector_set_all(&diag.vector, 1.0);
	_SigIpr = gsl_matrix_calloc(d, d);
	diag = gsl_matrix_diagonal(_SigIpr);
	gsl_vector_set_all(&diag.vector, 1e-6);
	_fitEach = &fE;
	
}
Apex::Apex(const double &dV, const size_t &d, vector<vector <double> > &fE){
	_Amat = gsl_matrix_calloc(d, d);
	gsl_vector_view diag = gsl_matrix_diagonal(_Amat);
	gsl_vector_set_all(&diag.vector, 1.0);
	_SigIpr = gsl_matrix_calloc(d, d);
	diag = gsl_matrix_diagonal(_SigIpr);
	gsl_vector_set_all(&diag.vector, dV);
	_fitEach = &fE;
	
	const gsl_rng_type *T = gsl_rng_mt19937;
	_r = gsl_rng_alloc(T);
	gsl_rng_set(_r, time(NULL)+rdtsc());
}
Apex::Apex(const SigmaI &SigI, vector<vector <double> > &fE){
	_SigIpr = gsl_matrix_alloc(SigI.getMat()->size1, SigI.getMat()->size2);
	gsl_matrix_memcpy(_SigIpr, SigI.getMat());
	_Amat = gsl_matrix_calloc(_SigIpr->size1, _SigIpr->size2);
	gsl_vector_view diag = gsl_matrix_diagonal(_Amat);
	gsl_vector_set_all(&diag.vector, 1.0);
	_fitEach = &fE;
	
	const gsl_rng_type *T = gsl_rng_mt19937;
	_r = gsl_rng_alloc(T);
	gsl_rng_set(_r, time(NULL)+rdtsc());
}

Apex::~Apex(){
	gsl_matrix_free(_Amat);
	gsl_matrix_free(_SigIpr);
	gsl_rng_free(_r);
}

void Apex::setPr(const double &pr){
	gsl_vector_view diag = gsl_matrix_diagonal(_SigIpr);
	gsl_vector_set_all(&diag.vector, pr);
}

Apex::Apex(const Apex &A){
	gsl_matrix_free(_Amat);
	gsl_matrix_free(_SigIpr);
	
	_Amat = gsl_matrix_alloc(A._Amat->size1, A._Amat->size2);
	gsl_matrix_memcpy(_Amat, A._Amat);
	_SigIpr = gsl_matrix_alloc(A._SigIpr->size1, A._SigIpr->size2);
	gsl_matrix_memcpy(_SigIpr, A._SigIpr);
	_fitEach = A._fitEach;
}
Apex &Apex::operator=(const Apex &A){
	gsl_matrix_free(_Amat);
	gsl_matrix_free(_SigIpr);
	
	_Amat = gsl_matrix_alloc(A._Amat->size1, A._Amat->size2);
	gsl_matrix_memcpy(_Amat, A._Amat);
	_SigIpr = gsl_matrix_alloc(A._SigIpr->size1, A._SigIpr->size2);
	gsl_matrix_memcpy(_SigIpr, A._SigIpr);
	_fitEach = A._fitEach;
	
	return *this;
}

void Apex::update(const Grp &y, const gsl_matrix *xi, const SigmaI &SigIm){
	double xTx = 0.0;
	
	gsl_matrix *SigSum = gsl_matrix_alloc(_SigIpr->size1, _SigIpr->size2);
	gsl_vector *tmp    = gsl_vector_alloc(_Amat->size2);
	gsl_matrix *ft     = gsl_matrix_calloc(y.dMat()->size1, y.dMat()->size2);
	
	for (size_t iRw = 0; iRw < _Amat->size1; iRw++) {
		gsl_matrix_view ftMat = gsl_matrix_view_array((*_fitEach)[iRw].data(), y.dMat()->size1, y.dMat()->size2);
		gsl_matrix_memcpy(ft, y.dMat());
		gsl_matrix_sub(ft, &ftMat.matrix);
		
		gsl_vector_view Arw  = gsl_matrix_row(_Amat, iRw);
		gsl_vector_view muPr = gsl_matrix_row(_SigIpr, iRw); // muPr%*%SigI, and muPr is I
		gsl_vector_const_view xiCol = gsl_matrix_const_column(xi, iRw); // the iRw refers to a row if A, which corresponds to a column of xi
		
		gsl_blas_ddot(&xiCol.vector, &xiCol.vector, &xTx);
		
		gsl_matrix_memcpy(SigSum, SigIm.getMat());
		gsl_matrix_scale(SigSum, xTx);
		gsl_matrix_add(SigSum, _SigIpr);
		gsl_linalg_cholesky_decomp(SigSum);
		gsl_linalg_cholesky_invert(SigSum);
		
		gsl_blas_dgemv(CblasTrans, 1.0, ft, &xiCol.vector, 0.0, tmp);
		gsl_blas_dsymv(CblasLower, 1.0, SigIm.getMat(), tmp, 0.0, &Arw.vector);
		gsl_vector_add(&Arw.vector, &muPr.vector);
		gsl_blas_dsymv(CblasLower, 1.0, SigSum, &Arw.vector, 0.0, tmp);
		
		gsl_linalg_cholesky_decomp(SigSum);
		MVgauss(tmp, SigSum, _r, &Arw.vector);
		
	}
	
	gsl_matrix_free(SigSum);
	gsl_matrix_free(ft);
	gsl_vector_free(tmp);
}

void Apex::update(const Grp &y, const gsl_matrix *xi, const SigmaI &SigIm, const RanIndex &ind){
	double xTx = 0.0;
	gsl_matrix *SigSum = gsl_matrix_alloc(_SigIpr->size1, _SigIpr->size2);
	gsl_vector *tmp    = gsl_vector_alloc(_Amat->size2);
	gsl_matrix *xiLg   = gsl_matrix_alloc(y.dMat()->size1, y.dMat()->size2);
	
	for (size_t iUp = 0; iUp < xi->size1; iUp++) {
		gsl_vector_const_view xiRw = gsl_matrix_const_row(xi, iUp);
		for (vector<size_t>::const_iterator lwIt = ind[iUp].begin(); lwIt != ind[iUp].end(); ++lwIt) {
			gsl_matrix_set_row(xiLg, *lwIt, &xiRw.vector);
		}
	}
	
	gsl_matrix *ft = gsl_matrix_alloc(y.dMat()->size1, y.dMat()->size2);
	
	for (size_t iRw = 0; iRw < _Amat->size1; iRw++) {
		gsl_matrix_view ftMat = gsl_matrix_view_array((*_fitEach)[iRw].data(), y.dMat()->size1, y.dMat()->size2);
		gsl_matrix_memcpy(ft, y.dMat());
		gsl_matrix_sub(ft, &ftMat.matrix);
		
		gsl_vector_view Arw  = gsl_matrix_row(_Amat, iRw);
		gsl_vector_view muPr = gsl_matrix_row(_SigIpr, iRw);  // i_m%*%Sig_A is the same as the m-th row of Sig_A
		gsl_vector_const_view xiCol = gsl_matrix_const_column(xiLg, iRw); // the iRw refers to a row if A, which corresponds to a column of xi
		
		gsl_blas_ddot(&xiCol.vector, &xiCol.vector, &xTx);
		
		gsl_matrix_memcpy(SigSum, SigIm.getMat());
		gsl_matrix_scale(SigSum, xTx);
		gsl_matrix_add(SigSum, _SigIpr);
		gsl_linalg_cholesky_decomp(SigSum);
		gsl_linalg_cholesky_invert(SigSum);
		
		gsl_blas_dgemv(CblasTrans, 1.0, ft, &xiCol.vector, 0.0, tmp);
		gsl_blas_dsymv(CblasLower, 1.0, SigIm.getMat(), tmp, 0.0, &Arw.vector);
		gsl_vector_add(&Arw.vector, &muPr.vector);
		gsl_blas_dsymv(CblasLower, 1.0, SigSum, &Arw.vector, 0.0, tmp);
		
		gsl_linalg_cholesky_decomp(SigSum);
		MVgauss(tmp, SigSum, _r, &Arw.vector);
		
	}
	
	gsl_matrix_free(SigSum);
	gsl_matrix_free(xiLg);
	gsl_matrix_free(ft);
	gsl_vector_free(tmp);
}
void Apex::update(const Grp &y, const gsl_matrix *xi, const Qgrp &q, const SigmaI &SigIm, const RanIndex &ind){
	double xQx = 0.0;
	gsl_matrix *SigSum = gsl_matrix_alloc(_SigIpr->size1, _SigIpr->size2);
	gsl_vector *tmp    = gsl_vector_alloc(_Amat->size2);
	gsl_matrix *xiLg   = gsl_matrix_alloc(y.dMat()->size1, y.dMat()->size2);
	gsl_vector *xiQ    = gsl_vector_alloc(y.dMat()->size1);
	
	for (size_t iUp = 0; iUp < xi->size1; iUp++) {
		gsl_vector_const_view xiRw = gsl_matrix_const_row(xi, iUp);
		for (vector<size_t>::const_iterator lwIt = ind[iUp].begin(); lwIt != ind[iUp].end(); ++lwIt) {
			gsl_matrix_set_row(xiLg, *lwIt, &xiRw.vector);
		}
	}
	
	gsl_matrix *ft = gsl_matrix_alloc(y.dMat()->size1, y.dMat()->size2);
	
	for (size_t iRw = 0; iRw < _Amat->size1; iRw++) {
		gsl_matrix_view ftMat = gsl_matrix_view_array((*_fitEach)[iRw].data(), y.dMat()->size1, y.dMat()->size2);
		gsl_matrix_memcpy(ft, y.dMat());
		gsl_matrix_sub(ft, &ftMat.matrix);
		
		gsl_vector_view Arw  = gsl_matrix_row(_Amat, iRw);
		gsl_vector_view muPr = gsl_matrix_row(_SigIpr, iRw);  // i_m%*%Sig_A is the same as the m-th row of Sig_A
		gsl_vector_const_view xiCol = gsl_matrix_const_column(xiLg, iRw); // the iRw refers to a row if A, which corresponds to a column of xi
		
		for (size_t iXQ = 0; iXQ < xiQ->size; iXQ++) {
			gsl_vector_set(xiQ, iXQ, gsl_matrix_get(xiLg, iXQ, iRw)*q[iXQ]);
		}
		
		gsl_blas_ddot(xiQ, &xiCol.vector, &xQx);
		
		gsl_matrix_memcpy(SigSum, SigIm.getMat());
		gsl_matrix_scale(SigSum, xQx);
		gsl_matrix_add(SigSum, _SigIpr);
		gsl_linalg_cholesky_decomp(SigSum);
		gsl_linalg_cholesky_invert(SigSum);
		
		gsl_blas_dgemv(CblasTrans, 1.0, ft, xiQ, 0.0, tmp);
		gsl_blas_dsymv(CblasLower, 1.0, SigIm.getMat(), tmp, 0.0, &Arw.vector);
		gsl_vector_add(&Arw.vector, &muPr.vector);
		gsl_blas_dsymv(CblasLower, 1.0, SigSum, &Arw.vector, 0.0, tmp);
		
		gsl_linalg_cholesky_decomp(SigSum);
		MVgauss(tmp, SigSum, _r, &Arw.vector);
		
	}
	
	gsl_matrix_free(SigSum);
	gsl_matrix_free(xiLg);
	gsl_matrix_free(ft);
	gsl_vector_free(tmp);
}

/*
 *	arithmetic operators for Grp classes
 */

MuGrp operator+(const Grp &m1, const Grp &m2){
	if ((m1.fMat())->size1 == (m2.fMat())->size1) {
		MuGrp res(m1);
		gsl_matrix_add(res._valueMat, m2.fMat());
		return res;
	}
	else if ((m1.fMat())->size1 > (m2.fMat())->size1){
		
		if (m2.fMat()->size1 == 1) {
			MuGrp res(m1);
			gsl_vector_const_view m2Row = gsl_matrix_const_row(m2.fMat(), 0);
			for (size_t iRw = 0; iRw < res._valueMat->size1; iRw++) {
				gsl_vector_view resRow = gsl_matrix_row(res._valueMat, iRw);
				gsl_vector_add(&resRow.vector, &m2Row.vector);
			}
			return res;
		}
		else if ((m1.fMat())->size1 == (m2._lowLevel)->getNtot()) { // this is the case where the m2 index directly points to levels of m1, so figuring out the correspondence is straight-forward
			MuGrp res(m1);
			for (size_t iRw2 = 0; iRw2 < (m2.fMat())->size1; iRw2++) {
				gsl_vector_const_view m2Row = gsl_matrix_const_row(m2.fMat(), iRw2);
				for (vector<size_t>::const_iterator lwIt = (*(m2._lowLevel))[iRw2].begin(); lwIt != (*(m2._lowLevel))[iRw2].end(); ++lwIt) {
					gsl_vector_view resRow = gsl_matrix_row(res._valueMat, *lwIt);
					gsl_vector_add(&resRow.vector, &m2Row.vector);
				}
			}
			return res;
		}
		else if ((m1._lowLevel)->getNtot() == (m2._lowLevel)->getNtot()){ // this is the case where m1 and m2 point to the same underlying level.  We have to use that lower level to figure out which level of m2 corresponds to which m1 trough their mutual targets
			gsl_matrix *tmp = gsl_matrix_calloc((m1._lowLevel)->getNtot(), m1.fMat()->size2);
			MuGrp res(tmp);
			gsl_matrix_free(tmp);
			for (size_t iRw = 0; iRw < res._valueMat->size1; iRw++) {
				gsl_vector_view resRow = gsl_matrix_row(res._valueMat, iRw);
				gsl_matrix_get_row(&resRow.vector, m1.fMat(), (m1._lowLevel)->priorInd(iRw));
				gsl_vector_const_view m2Row = gsl_matrix_const_row(m2.fMat(), (m2._lowLevel)->priorInd(iRw));
				gsl_vector_add(&resRow.vector, &m2Row.vector);
			}
			return res;
		}
		else {
			cerr << "ADDITION OPERATOR ERROR: Cannot relate m1 rows to m2 in addition with m1 > m2" << endl;
			exit(1);
		}
		
	}
	else {
		
		if (m1.fMat()->size1 == 1) {
			MuGrp res(m2);
			gsl_vector_const_view m1Row = gsl_matrix_const_row(m1.fMat(), 0);
			for (size_t iRw = 0; iRw < res._valueMat->size1; iRw++) {
				gsl_vector_view resRow = gsl_matrix_row(res._valueMat, iRw);
				gsl_vector_add(&resRow.vector, &m1Row.vector);
			}
			return res;
		}
		else if ((m2.fMat())->size1 == (m1._lowLevel)->getNtot()) {
			MuGrp res(m2);
			for (size_t iRw1 = 0; iRw1 < (m1.fMat())->size1; iRw1++) {
				gsl_vector_const_view m1Row = gsl_matrix_const_row(m1.fMat(), iRw1);
				for (vector<size_t>::const_iterator lwIt = (*(m1._lowLevel))[iRw1].begin(); lwIt != (*(m1._lowLevel))[iRw1].end(); ++lwIt) {
					gsl_vector_view resRow = gsl_matrix_row(res._valueMat, *lwIt);
					gsl_vector_add(&resRow.vector, &m1Row.vector);
				}
			}
			return res;
		}
		else if ((m1._lowLevel)->getNtot() == (m2._lowLevel)->getNtot()){
			gsl_matrix *tmp = gsl_matrix_calloc((m1._lowLevel)->getNtot(), m1.fMat()->size2);
			MuGrp res(tmp);
			gsl_matrix_free(tmp);
			for (size_t iRw = 0; iRw < res._valueMat->size1; iRw++) {
				gsl_vector_view resRow = gsl_matrix_row(res._valueMat, iRw);
				gsl_matrix_get_row(&resRow.vector, m2.fMat(), (m2._lowLevel)->priorInd(iRw));
				gsl_vector_const_view m1Row = gsl_matrix_const_row(m1.fMat(), (m1._lowLevel)->priorInd(iRw));
				gsl_vector_add(&resRow.vector, &m1Row.vector);
			}
			return res;
		}
		else {
			cerr << "ADDITION OPERATOR ERROR: Cannot relate m1 rows to m2 in addition with m1 < m2" << endl;
			exit(1);
		}
		
	}
	
}
MuGrp operator+(const MuGrp &m1, const Grp &m2){
	if ((m1.fMat())->size1 == (m2.fMat())->size1) {
		MuGrp res(m1);
		gsl_matrix_add(res._valueMat, m2.fMat());
		return res;
	}
	else if ((m1.fMat())->size1 > (m2.fMat())->size1){
		
		if (m2.fMat()->size1 == 1) {
			MuGrp res(m1);
			gsl_vector_const_view m2Row = gsl_matrix_const_row(m2.fMat(), 0);
			for (size_t iRw = 0; iRw < res._valueMat->size1; iRw++) {
				gsl_vector_view resRow = gsl_matrix_row(res._valueMat, iRw);
				gsl_vector_add(&resRow.vector, &m2Row.vector);
			}
			return res;
		}
		else if ((m1.fMat())->size1 == (m2._lowLevel)->getNtot()) { // this is the case where the m2 index directly points to levels of m1, so figuring out the correspondence is straight-forward
			MuGrp res(m1);
			for (size_t iRw2 = 0; iRw2 < (m2.fMat())->size1; iRw2++) {
				gsl_vector_const_view m2Row = gsl_matrix_const_row(m2.fMat(), iRw2);
				for (vector<size_t>::const_iterator lwIt = (*(m2._lowLevel))[iRw2].begin(); lwIt != (*(m2._lowLevel))[iRw2].end(); ++lwIt) {
					gsl_vector_view resRow = gsl_matrix_row(res._valueMat, *lwIt);
					gsl_vector_add(&resRow.vector, &m2Row.vector);
				}
			}
			return res;
		}
		else if ((m1._lowLevel)->getNtot() == (m2._lowLevel)->getNtot()){ // this is the case where m1 and m2 point to the same underlying level.  We have to use that lower level to figure out which level of m2 corresponds to which m1 trough their mutual targets
			gsl_matrix *tmp = gsl_matrix_calloc((m1._lowLevel)->getNtot(), m1.fMat()->size2);
			MuGrp res(tmp);
			gsl_matrix_free(tmp);
			for (size_t iRw = 0; iRw < res._valueMat->size1; iRw++) {
				gsl_vector_view resRow = gsl_matrix_row(res._valueMat, iRw);
				gsl_matrix_get_row(&resRow.vector, m1.fMat(), (m1._lowLevel)->priorInd(iRw));
				gsl_vector_const_view m2Row = gsl_matrix_const_row(m2.fMat(), (m2._lowLevel)->priorInd(iRw));
				gsl_vector_add(&resRow.vector, &m2Row.vector);
			}
			return res;
		}
		else {
			cerr << "ADDITION OPERATOR ERROR: Cannot relate m1 rows to m2 in addition with m1 > m2" << endl;
			exit(1);
		}
		
	}
	else {
		
		if (m1.fMat()->size1 == 1) {
			MuGrp res(m2);
			gsl_vector_const_view m1Row = gsl_matrix_const_row(m1.fMat(), 0);
			for (size_t iRw = 0; iRw < res._valueMat->size1; iRw++) {
				gsl_vector_view resRow = gsl_matrix_row(res._valueMat, iRw);
				gsl_vector_add(&resRow.vector, &m1Row.vector);
			}
			return res;
		}
		else if ((m2.fMat())->size1 == (m1._lowLevel)->getNtot()) {
			MuGrp res(m2);
			for (size_t iRw1 = 0; iRw1 < (m1.fMat())->size1; iRw1++) {
				gsl_vector_const_view m1Row = gsl_matrix_const_row(m1.fMat(), iRw1);
				for (vector<size_t>::const_iterator lwIt = (*(m1._lowLevel))[iRw1].begin(); lwIt != (*(m1._lowLevel))[iRw1].end(); ++lwIt) {
					gsl_vector_view resRow = gsl_matrix_row(res._valueMat, *lwIt);
					gsl_vector_add(&resRow.vector, &m1Row.vector);
				}
			}
			return res;
		}
		else if ((m1._lowLevel)->getNtot() == (m2._lowLevel)->getNtot()){
			gsl_matrix *tmp = gsl_matrix_calloc((m1._lowLevel)->getNtot(), m1.fMat()->size2);
			MuGrp res(tmp);
			gsl_matrix_free(tmp);
			for (size_t iRw = 0; iRw < res._valueMat->size1; iRw++) {
				gsl_vector_view resRow = gsl_matrix_row(res._valueMat, iRw);
				gsl_matrix_get_row(&resRow.vector, m2.fMat(), (m2._lowLevel)->priorInd(iRw));
				gsl_vector_const_view m1Row = gsl_matrix_const_row(m1.fMat(), (m1._lowLevel)->priorInd(iRw));
				gsl_vector_add(&resRow.vector, &m1Row.vector);
			}
			return res;
		}
		else {
			cerr << "ADDITION OPERATOR ERROR: Cannot relate m1 rows to m2 in addition with m1 < m2" << endl;
			exit(1);
		}
		
	}
	
}
MuGrp operator+(const Grp &m1, const MuGrp &m2){
	if ((m1.fMat())->size1 == (m2.fMat())->size1) {
		MuGrp res(m1);
		gsl_matrix_add(res._valueMat, m2.fMat());
		return res;
	}
	else if ((m1.fMat())->size1 > (m2.fMat())->size1){
		
		if (m2.fMat()->size1 == 1) {
			MuGrp res(m1);
			gsl_vector_const_view m2Row = gsl_matrix_const_row(m2.fMat(), 0);
			for (size_t iRw = 0; iRw < res._valueMat->size1; iRw++) {
				gsl_vector_view resRow = gsl_matrix_row(res._valueMat, iRw);
				gsl_vector_add(&resRow.vector, &m2Row.vector);
			}
			return res;
		}
		else if ((m1.fMat())->size1 == (m2._lowLevel)->getNtot()) { // this is the case where the m2 index directly points to levels of m1, so figuring out the correspondence is straight-forward
			MuGrp res(m1);
			for (size_t iRw2 = 0; iRw2 < (m2.fMat())->size1; iRw2++) {
				gsl_vector_const_view m2Row = gsl_matrix_const_row(m2.fMat(), iRw2);
				for (vector<size_t>::const_iterator lwIt = (*(m2._lowLevel))[iRw2].begin(); lwIt != (*(m2._lowLevel))[iRw2].end(); ++lwIt) {
					gsl_vector_view resRow = gsl_matrix_row(res._valueMat, *lwIt);
					gsl_vector_add(&resRow.vector, &m2Row.vector);
				}
			}
			return res;
		}
		else if ((m1._lowLevel)->getNtot() == (m2._lowLevel)->getNtot()){ // this is the case where m1 and m2 point to the same underlying level.  We have to use that lower level to figure out which level of m2 corresponds to which m1 trough their mutual targets
			gsl_matrix *tmp = gsl_matrix_calloc((m1._lowLevel)->getNtot(), m1.fMat()->size2);
			MuGrp res(tmp);
			gsl_matrix_free(tmp);
			for (size_t iRw = 0; iRw < res._valueMat->size1; iRw++) {
				gsl_vector_view resRow = gsl_matrix_row(res._valueMat, iRw);
				gsl_matrix_get_row(&resRow.vector, m1.fMat(), (m1._lowLevel)->priorInd(iRw));
				gsl_vector_const_view m2Row = gsl_matrix_const_row(m2.fMat(), (m2._lowLevel)->priorInd(iRw));
				gsl_vector_add(&resRow.vector, &m2Row.vector);
			}
			return res;
		}
		else {
			cerr << "ADDITION OPERATOR ERROR: Cannot relate m1 rows to m2 in addition with m1 > m2" << endl;
			exit(1);
		}
		
	}
	else {
		
		if (m1.fMat()->size1 == 1) {
			MuGrp res(m2);
			gsl_vector_const_view m1Row = gsl_matrix_const_row(m1.fMat(), 0);
			for (size_t iRw = 0; iRw < res._valueMat->size1; iRw++) {
				gsl_vector_view resRow = gsl_matrix_row(res._valueMat, iRw);
				gsl_vector_add(&resRow.vector, &m1Row.vector);
			}
			return res;
		}
		else if ((m2.fMat())->size1 == (m1._lowLevel)->getNtot()) {
			MuGrp res(m2);
			for (size_t iRw1 = 0; iRw1 < (m1.fMat())->size1; iRw1++) {
				gsl_vector_const_view m1Row = gsl_matrix_const_row(m1.fMat(), iRw1);
				for (vector<size_t>::const_iterator lwIt = (*(m1._lowLevel))[iRw1].begin(); lwIt != (*(m1._lowLevel))[iRw1].end(); ++lwIt) {
					gsl_vector_view resRow = gsl_matrix_row(res._valueMat, *lwIt);
					gsl_vector_add(&resRow.vector, &m1Row.vector);
				}
			}
			return res;
		}
		else if ((m1._lowLevel)->getNtot() == (m2._lowLevel)->getNtot()){
			gsl_matrix *tmp = gsl_matrix_calloc((m1._lowLevel)->getNtot(), m1.fMat()->size2);
			MuGrp res(tmp);
			gsl_matrix_free(tmp);
			for (size_t iRw = 0; iRw < res._valueMat->size1; iRw++) {
				gsl_vector_view resRow = gsl_matrix_row(res._valueMat, iRw);
				gsl_matrix_get_row(&resRow.vector, m2.fMat(), (m2._lowLevel)->priorInd(iRw));
				gsl_vector_const_view m1Row = gsl_matrix_const_row(m1.fMat(), (m1._lowLevel)->priorInd(iRw));
				gsl_vector_add(&resRow.vector, &m1Row.vector);
			}
			return res;
		}
		else {
			cerr << "ADDITION OPERATOR ERROR: Cannot relate m1 rows to m2 in addition with m1 < m2" << endl;
			exit(1);
		}
		
	}
	
}

MuGrp operator-(const Grp &m1, const Grp &m2){
	if ((m1.fMat())->size1 == (m2.fMat())->size1) {
		MuGrp res(m1);
		gsl_matrix_sub(res._valueMat, m2.fMat());
		return res;
	}
	else if ((m1.fMat())->size1 > (m2.fMat())->size1){
		
		if (m2.fMat()->size1 == 1) {
			MuGrp res(m1);
			gsl_vector_const_view m2Row = gsl_matrix_const_row(m2.fMat(), 0);
			for (size_t iRw = 0; iRw < res._valueMat->size1; iRw++) {
				gsl_vector_view resRow = gsl_matrix_row(res._valueMat, iRw);
				gsl_vector_sub(&resRow.vector, &m2Row.vector);
			}
			return res;
		}
		else if ((m1.fMat())->size1 == (m2._lowLevel)->getNtot()) { // this is the case where the m2 index directly points to levels of m1, so figuring out the correspondence is straight-forward
			MuGrp res(m1);
			for (size_t iRw2 = 0; iRw2 < (m2.fMat())->size1; iRw2++) {
				gsl_vector_const_view m2Row = gsl_matrix_const_row(m2.fMat(), iRw2);
				for (vector<size_t>::const_iterator lwIt = (*(m2._lowLevel))[iRw2].begin(); lwIt != (*(m2._lowLevel))[iRw2].end(); ++lwIt) {
					gsl_vector_view resRow = gsl_matrix_row(res._valueMat, *lwIt);
					gsl_vector_sub(&resRow.vector, &m2Row.vector);
				}
			}
			return res;
		}
		else if ((m1._lowLevel)->getNtot() == (m2._lowLevel)->getNtot()){ // this is the case where m1 and m2 point to the same underlying level.  We have to use that lower level to figure out which level of m2 corresponds to which m1 trough their mutual targets
			gsl_matrix *tmp = gsl_matrix_calloc((m1._lowLevel)->getNtot(), m1.fMat()->size2);
			MuGrp res(tmp);
			gsl_matrix_free(tmp);
			for (size_t iRw = 0; iRw < res._valueMat->size1; iRw++) {
				gsl_vector_view resRow = gsl_matrix_row(res._valueMat, iRw);
				gsl_matrix_get_row(&resRow.vector, m1.fMat(), (m1._lowLevel)->priorInd(iRw));
				gsl_vector_const_view m2Row = gsl_matrix_const_row(m2.fMat(), (m2._lowLevel)->priorInd(iRw));
				gsl_vector_sub(&resRow.vector, &m2Row.vector);
			}
			return res;
		}
		else {
			cerr << "SUBTRACTION OPERATOR ERROR: Cannot relate m1 rows to m2 in subtraction with m1 > m2" << endl;
			exit(1);
		}
		
	}
	else {
		
		if (m1.fMat()->size1 == 1) {
			gsl_matrix *tmp = gsl_matrix_alloc(m2.fMat()->size1, m2.fMat()->size2);
			MuGrp res(tmp);
			gsl_matrix_free(tmp);
			gsl_vector_const_view m1Row = gsl_matrix_const_row(m1.fMat(), 0);
			for (size_t iRw = 0; iRw < res._valueMat->size1; iRw++) {
				gsl_matrix_set_row(res._valueMat, iRw, &m1Row.vector);
			}
			gsl_matrix_sub(res._valueMat, m2.fMat());
			return res;
		}
		else if ((m2.fMat())->size1 == (m1._lowLevel)->getNtot()) {
			gsl_matrix *tmp = gsl_matrix_alloc(m2.fMat()->size1, m2.fMat()->size2);
			MuGrp res(tmp);
			gsl_matrix_free(tmp);
			for (size_t iRw1 = 0; iRw1 < (m1.fMat())->size1; iRw1++) {
				gsl_vector_const_view m1Row = gsl_matrix_const_row(m1.fMat(), iRw1);
				for (vector<size_t>::const_iterator lwIt = (*(m1._lowLevel))[iRw1].begin(); lwIt != (*(m1._lowLevel))[iRw1].end(); ++lwIt) {
					gsl_matrix_set_row(res._valueMat, *lwIt, &m1Row.vector);
				}
			}
			gsl_matrix_sub(res._valueMat, m2.fMat());
			return res;
		}
		else if ((m1._lowLevel)->getNtot() == (m2._lowLevel)->getNtot()){
			gsl_matrix *tmp = gsl_matrix_calloc((m1._lowLevel)->getNtot(), m1.fMat()->size2);
			MuGrp res(tmp);
			gsl_matrix_free(tmp);
			for (size_t iRw = 0; iRw < res._valueMat->size1; iRw++) {
				gsl_vector_view resRow = gsl_matrix_row(res._valueMat, iRw);
				gsl_matrix_get_row(&resRow.vector, m1.fMat(), (m1._lowLevel)->priorInd(iRw));
				gsl_vector_const_view m2Row = gsl_matrix_const_row(m2.fMat(), (m2._lowLevel)->priorInd(iRw));
				gsl_vector_sub(&resRow.vector, &m2Row.vector);
			}
			return res;
		}
		else {
			cerr << "SUBTRACTION OPERATOR ERROR: Cannot relate m1 rows to m2 in subtraction with m1 < m2" << endl;
			exit(1);
		}
	}
}
MuGrp operator-(const MuGrp &m1, const Grp &m2){
	if ((m1.fMat())->size1 == (m2.fMat())->size1) {
		MuGrp res(m1);
		gsl_matrix_sub(res._valueMat, m2.fMat());
		return res;
	}
	else if ((m1.fMat())->size1 > (m2.fMat())->size1){
		
		if (m2.fMat()->size1 == 1) {
			MuGrp res(m1);
			gsl_vector_const_view m2Row = gsl_matrix_const_row(m2.fMat(), 0);
			for (size_t iRw = 0; iRw < res._valueMat->size1; iRw++) {
				gsl_vector_view resRow = gsl_matrix_row(res._valueMat, iRw);
				gsl_vector_sub(&resRow.vector, &m2Row.vector);
			}
			return res;
		}
		else if ((m1.fMat())->size1 == (m2._lowLevel)->getNtot()) { // this is the case where the m2 index directly points to levels of m1, so figuring out the correspondence is straight-forward
			MuGrp res(m1);
			for (size_t iRw2 = 0; iRw2 < (m2.fMat())->size1; iRw2++) {
				gsl_vector_const_view m2Row = gsl_matrix_const_row(m2.fMat(), iRw2);
				for (vector<size_t>::const_iterator lwIt = (*(m2._lowLevel))[iRw2].begin(); lwIt != (*(m2._lowLevel))[iRw2].end(); ++lwIt) {
					gsl_vector_view resRow = gsl_matrix_row(res._valueMat, *lwIt);
					gsl_vector_sub(&resRow.vector, &m2Row.vector);
				}
			}
			return res;
		}
		else if ((m1._lowLevel)->getNtot() == (m2._lowLevel)->getNtot()){ // this is the case where m1 and m2 point to the same underlying level.  We have to use that lower level to figure out which level of m2 corresponds to which m1 trough their mutual targets
			gsl_matrix *tmp = gsl_matrix_calloc((m1._lowLevel)->getNtot(), m1.fMat()->size2);
			MuGrp res(tmp);
			gsl_matrix_free(tmp);
			for (size_t iRw = 0; iRw < res._valueMat->size1; iRw++) {
				gsl_vector_view resRow = gsl_matrix_row(res._valueMat, iRw);
				gsl_matrix_get_row(&resRow.vector, m1.fMat(), (m1._lowLevel)->priorInd(iRw));
				gsl_vector_const_view m2Row = gsl_matrix_const_row(m2.fMat(), (m2._lowLevel)->priorInd(iRw));
				gsl_vector_sub(&resRow.vector, &m2Row.vector);
			}
			return res;
		}
		else {
			cerr << "SUBTRACTION OPERATOR ERROR: Cannot relate m1 rows to m2 in subtraction with m1 > m2" << endl;
			exit(1);
		}
		
	}
	else {
		
		if (m1.fMat()->size1 == 1) {
			gsl_matrix *tmp = gsl_matrix_alloc(m2.fMat()->size1, m2.fMat()->size2);
			MuGrp res(tmp);
			gsl_matrix_free(tmp);
			gsl_vector_const_view m1Row = gsl_matrix_const_row(m1.fMat(), 0);
			for (size_t iRw = 0; iRw < res._valueMat->size1; iRw++) {
				gsl_matrix_set_row(res._valueMat, iRw, &m1Row.vector);
			}
			gsl_matrix_sub(res._valueMat, m2.fMat());
			return res;
		}
		else if ((m2.fMat())->size1 == (m1._lowLevel)->getNtot()) {
			gsl_matrix *tmp = gsl_matrix_alloc(m2.fMat()->size1, m2.fMat()->size2);
			MuGrp res(tmp);
			gsl_matrix_free(tmp);
			for (size_t iRw1 = 0; iRw1 < (m1.fMat())->size1; iRw1++) {
				gsl_vector_const_view m1Row = gsl_matrix_const_row(m1.fMat(), iRw1);
				for (vector<size_t>::const_iterator lwIt = (*(m1._lowLevel))[iRw1].begin(); lwIt != (*(m1._lowLevel))[iRw1].end(); ++lwIt) {
					gsl_matrix_set_row(res._valueMat, *lwIt, &m1Row.vector);
				}
			}
			gsl_matrix_sub(res._valueMat, m2.fMat());
			return res;
		}
		else if ((m1._lowLevel)->getNtot() == (m2._lowLevel)->getNtot()){
			gsl_matrix *tmp = gsl_matrix_calloc((m1._lowLevel)->getNtot(), m1.fMat()->size2);
			MuGrp res(tmp);
			gsl_matrix_free(tmp);
			for (size_t iRw = 0; iRw < res._valueMat->size1; iRw++) {
				gsl_vector_view resRow = gsl_matrix_row(res._valueMat, iRw);
				gsl_matrix_get_row(&resRow.vector, m1.fMat(), (m1._lowLevel)->priorInd(iRw));
				gsl_vector_const_view m2Row = gsl_matrix_const_row(m2.fMat(), (m2._lowLevel)->priorInd(iRw));
				gsl_vector_sub(&resRow.vector, &m2Row.vector);
			}
			return res;
		}
		else {
			cerr << "SUBTRACTION OPERATOR ERROR: Cannot relate m1 rows to m2 in subtraction with m1 < m2" << endl;
			exit(1);
		}
	}
}
MuGrp operator-(const Grp &m1, const MuGrp &m2){
	if ((m1.fMat())->size1 == (m2.fMat())->size1) {
		MuGrp res(m1);
		gsl_matrix_sub(res._valueMat, m2.fMat());
		return res;
	}
	else if ((m1.fMat())->size1 > (m2.fMat())->size1){
		
		if (m2.fMat()->size1 == 1) {
			MuGrp res(m1);
			gsl_vector_const_view m2Row = gsl_matrix_const_row(m2.fMat(), 0);
			for (size_t iRw = 0; iRw < res._valueMat->size1; iRw++) {
				gsl_vector_view resRow = gsl_matrix_row(res._valueMat, iRw);
				gsl_vector_sub(&resRow.vector, &m2Row.vector);
			}
			return res;
		}
		else if ((m1.fMat())->size1 == (m2._lowLevel)->getNtot()) { // this is the case where the m2 index directly points to levels of m1, so figuring out the correspondence is straight-forward
			MuGrp res(m1);
			for (size_t iRw2 = 0; iRw2 < (m2.fMat())->size1; iRw2++) {
				gsl_vector_const_view m2Row = gsl_matrix_const_row(m2.fMat(), iRw2);
				for (vector<size_t>::const_iterator lwIt = (*(m2._lowLevel))[iRw2].begin(); lwIt != (*(m2._lowLevel))[iRw2].end(); ++lwIt) {
					gsl_vector_view resRow = gsl_matrix_row(res._valueMat, *lwIt);
					gsl_vector_sub(&resRow.vector, &m2Row.vector);
				}
			}
			return res;
		}
		else if ((m1._lowLevel)->getNtot() == (m2._lowLevel)->getNtot()){ // this is the case where m1 and m2 point to the same underlying level.  We have to use that lower level to figure out which level of m2 corresponds to which m1 trough their mutual targets
			gsl_matrix *tmp = gsl_matrix_calloc((m1._lowLevel)->getNtot(), m1.fMat()->size2);
			MuGrp res(tmp);
			gsl_matrix_free(tmp);
			for (size_t iRw = 0; iRw < res._valueMat->size1; iRw++) {
				gsl_vector_view resRow = gsl_matrix_row(res._valueMat, iRw);
				gsl_matrix_get_row(&resRow.vector, m1.fMat(), (m1._lowLevel)->priorInd(iRw));
				gsl_vector_const_view m2Row = gsl_matrix_const_row(m2.fMat(), (m2._lowLevel)->priorInd(iRw));
				gsl_vector_sub(&resRow.vector, &m2Row.vector);
			}
			return res;
		}
		else {
			cerr << "SUBTRACTION OPERATOR ERROR: Cannot relate m1 rows to m2 in subtraction with m1 > m2" << endl;
			exit(1);
		}
		
	}
	else {
		
		if (m1.fMat()->size1 == 1) {
			gsl_matrix *tmp = gsl_matrix_alloc(m2.fMat()->size1, m2.fMat()->size2);
			MuGrp res(tmp);
			gsl_matrix_free(tmp);
			gsl_vector_const_view m1Row = gsl_matrix_const_row(m1.fMat(), 0);
			for (size_t iRw = 0; iRw < res._valueMat->size1; iRw++) {
				gsl_matrix_set_row(res._valueMat, iRw, &m1Row.vector);
			}
			gsl_matrix_sub(res._valueMat, m2.fMat());
			return res;
		}
		else if ((m2.fMat())->size1 == (m1._lowLevel)->getNtot()) {
			gsl_matrix *tmp = gsl_matrix_alloc(m2.fMat()->size1, m2.fMat()->size2);
			MuGrp res(tmp);
			gsl_matrix_free(tmp);
			for (size_t iRw1 = 0; iRw1 < (m1.fMat())->size1; iRw1++) {
				gsl_vector_const_view m1Row = gsl_matrix_const_row(m1.fMat(), iRw1);
				for (vector<size_t>::const_iterator lwIt = (*(m1._lowLevel))[iRw1].begin(); lwIt != (*(m1._lowLevel))[iRw1].end(); ++lwIt) {
					gsl_matrix_set_row(res._valueMat, *lwIt, &m1Row.vector);
				}
			}
			gsl_matrix_sub(res._valueMat, m2.fMat());
			return res;
		}
		else if ((m1._lowLevel)->getNtot() == (m2._lowLevel)->getNtot()){
			gsl_matrix *tmp = gsl_matrix_calloc((m1._lowLevel)->getNtot(), m1.fMat()->size2);
			MuGrp res(tmp);
			gsl_matrix_free(tmp);
			for (size_t iRw = 0; iRw < res._valueMat->size1; iRw++) {
				gsl_vector_view resRow = gsl_matrix_row(res._valueMat, iRw);
				gsl_matrix_get_row(&resRow.vector, m1.fMat(), (m1._lowLevel)->priorInd(iRw));
				gsl_vector_const_view m2Row = gsl_matrix_const_row(m2.fMat(), (m2._lowLevel)->priorInd(iRw));
				gsl_vector_sub(&resRow.vector, &m2Row.vector);
			}
			return res;
		}
		else {
			cerr << "SUBTRACTION OPERATOR ERROR: Cannot relate m1 rows to m2 in subtraction with m1 < m2" << endl;
			exit(1);
		}
	}
}

/*
 *	Grp methods
 */
Grp::Grp() : _theta(0), _lowLevel(0), _upLevel(0), _outFlNam("LOCout.gbin") {
	_valueMat = gsl_matrix_calloc(1, 1);
	_rV.resize(1);
	
	
	const gsl_rng_type *T = gsl_rng_mt19937;
	_rV[0] = gsl_rng_alloc(T);
	gsl_rng_set(_rV[0], time(NULL)+rdtsc());

}
Grp::~Grp(){
	for (vector<MVnorm *>::iterator each = _theta.begin(); each != _theta.end(); ++each) {
		delete *each;
	}
	
	_theta.clear();
	gsl_matrix_free(_valueMat);
	for (vector<gsl_rng *>::iterator rIt = _rV.begin(); rIt != _rV.end(); ++rIt) {
		gsl_rng_free(*rIt);
	}
	_rV.clear();
};

void Grp::save(){
	FILE *outH = fopen(_outFlNam.c_str(), "a");
	gsl_matrix_fwrite(outH, _valueMat);
	fclose(outH);
}

void Grp::save(const string &outFlNam){
	if (_outFlNam == outFlNam) {
		FILE *outH = fopen(outFlNam.c_str(), "a");
		gsl_matrix_fwrite(outH, _valueMat);
		fclose(outH);

	}
	else {
		_outFlNam = outFlNam;
		remove(_outFlNam.c_str());
		FILE *outH = fopen(outFlNam.c_str(), "a");
		gsl_matrix_fwrite(outH, _valueMat);
		fclose(outH);
	}
}

void Grp::save(const string &outMuFlNam, const string &outSigFlNam, const SigmaI &SigI){
	gsl_matrix *Sig   = gsl_matrix_alloc(SigI.getMat()->size1, SigI.getMat()->size1);
	gsl_matrix_memcpy(Sig, SigI.getMat());
	gsl_linalg_cholesky_decomp(Sig);
	gsl_linalg_cholesky_invert(Sig);
	
	FILE *outS = fopen(outSigFlNam.c_str(), "a");
	gsl_matrix_fwrite(outS, Sig);
	fclose(outS);
	
	gsl_matrix_free(Sig);
	
	if (_outFlNam == outMuFlNam) {
		FILE *outM = fopen(outMuFlNam.c_str(), "a");
		gsl_matrix_fwrite(outM, _valueMat);
		fclose(outM);

	}
	else {
		_outFlNam = outMuFlNam;
		remove(_outFlNam.c_str());
		
		FILE *outM = fopen(outMuFlNam.c_str(), "a");
		gsl_matrix_fwrite(outM, _valueMat);
		fclose(outM);
		
	}

}

void Grp::mhlSave(const string &outFlNam, const SigmaI SigI){
	gsl_vector *mhlVec = gsl_vector_alloc(_theta.size());
	
	for (size_t iEl = 0; iEl < _theta.size(); iEl++) {
		gsl_vector_set(mhlVec, iEl, _theta[iEl]->mhl(SigI));
	}
	
	FILE *outM = fopen(outFlNam.c_str(), "a");
	gsl_vector_fwrite(outM, mhlVec);
	fclose(outM);
	
	gsl_vector_free(mhlVec);
}

MuGrp Grp::mean(RanIndex &grp){
	MuGrp res(*this, grp);
	return res;
}
const MuGrp Grp::mean(RanIndex &grp) const{
	MuGrp res(*this, grp);
	return res;
}

MuGrp Grp::mean(RanIndex &grp, const Qgrp &q){
	MuGrp res(*this, q, grp);
	return res;
}
const MuGrp Grp::mean(RanIndex &grp, const Qgrp &q) const{
	MuGrp res(*this, q, grp);
	return res;
}


/*
 *	MuGrp methods
 */

MuGrp::MuGrp(RanIndex &low, const size_t &d) : Grp(){
	gsl_matrix_free(_valueMat);
	_valueMat = gsl_matrix_calloc(1, d);
	_theta.resize(1);
	_theta[0] = new MVnormMu(_valueMat, 0, low.getIndVec());
	_lowLevel = &low;
	
}


MuGrp::MuGrp(const string &datFlNam, RanIndex &low, RanIndex &up, const size_t &d) : Grp(){
	if (low.getNgrp() != up.getNtot()) {
		cerr << "ERROR: # of elements in index to the upper level is wrong!!!" << endl;
		exit(1);
	}
	_lowLevel = &low;
	_upLevel  = &up;
	
	gsl_matrix_free(_valueMat);
	gsl_matrix *tmpIn = gsl_matrix_alloc(low.getNtot(), d);
	gsl_vector *tmpMn = gsl_vector_alloc(d);
	gsl_vector *tmpSd = gsl_vector_alloc(d);
	
	FILE *datIn = fopen(datFlNam.c_str(), "r");
	if (datIn == NULL) {
		cerr << "ERROR: cannot open data file " << datIn << " in MuGrp" << endl;
		exit(-1);
	}
	gsl_matrix_fread(datIn, tmpIn);
	fclose(datIn);
	
	_theta.resize(low.getNgrp());
	_valueMat = gsl_matrix_alloc(low.getNgrp(), d);
	
	for (size_t iEl = 0; iEl < low.getNgrp(); iEl++) {
		gsl_vector_set_zero(tmpMn);
		for (vector<size_t>::const_iterator el = low[iEl].begin(); el != low[iEl].end(); ++el) {
			gsl_vector_view ln = gsl_matrix_row(tmpIn, *el);
			gsl_vector_add(tmpMn, &ln.vector);
		}
		gsl_vector_scale(tmpMn, 1.0/low[iEl].size());
		gsl_matrix_set_row(_valueMat, iEl, tmpMn);
		
		for (size_t j = 0; j < d; j++) {
			gsl_vector_set(tmpSd, j, fabs(gsl_vector_get(tmpMn, j)));
		}
		_theta[iEl] = new MVnormMu(_valueMat, iEl, tmpSd, _rV[0], (*_lowLevel)[iEl], _upLevel->priorInd(iEl));
	}
		
	gsl_matrix_free(tmpIn);
	gsl_vector_free(tmpSd);
	gsl_vector_free(tmpMn);
}

MuGrp::MuGrp(const string &datFlNam, RanIndex &up, const size_t &d){
	gsl_matrix_free(_valueMat);
	_valueMat = gsl_matrix_alloc(up.getNtot(), d);
	_lowLevel = 0;
	_upLevel  = &up;
	
	FILE *datIn = fopen(datFlNam.c_str(), "r");
	if (datIn == NULL) {
		cerr << "ERROR: cannot open data file " << datIn << " in MuGrp" << endl;
		exit(-1);
	}
	gsl_matrix_fread(datIn, _valueMat);
	fclose(datIn);
	
	_theta.resize(up.getNtot());
	
	for (size_t iEl = 0; iEl < up.getNtot(); iEl++) {
		_theta[iEl] = new MVnormMu(_valueMat, iEl, _upLevel->priorInd(iEl));
	}
	
}
MuGrp::MuGrp(const vector<MVnorm *> &dat, RanIndex &low, RanIndex &up){
	if (low.getNgrp() != up.getNtot()) {
		cerr << "ERROR: # of elements in index to the upper level is wrong!!!" << endl;
		exit(1);
	}
	else if (low.getNtot() != dat.size()){
		cerr << "ERROR: the number of rows in data does not correspond to the number of points in the low index!!!" << endl;
		exit(1);
	}
	_lowLevel = &low;
	_upLevel  = &up;
	
	gsl_matrix_free(_valueMat);
	gsl_vector *tmpMn = gsl_vector_alloc(dat[0]->len());
	gsl_vector *tmpSd = gsl_vector_alloc(dat[0]->len());
	
	_theta.resize(low.getNgrp());
	_valueMat = gsl_matrix_alloc(low.getNgrp(), dat[0]->len());
	for (size_t iEl = 0; iEl < low.getNgrp(); iEl++) {
		gsl_vector_set_zero(tmpMn);
		for (vector<size_t>::const_iterator el = low[iEl].begin(); el != low[iEl].end(); ++el) {
			gsl_vector_add(tmpMn, dat[*el]->getVec());
		}
		gsl_vector_scale(tmpMn, 1.0/low[iEl].size());
		gsl_matrix_set_row(_valueMat, iEl, tmpMn);
		for (size_t j = 0; j < dat[0]->len(); j++) {
			gsl_vector_set(tmpSd, j, fabs(gsl_vector_get(tmpMn, j)));
		}
		_theta[iEl] = new MVnormMu(_valueMat, iEl, tmpSd, _rV[0], (*_lowLevel)[iEl], _upLevel->priorInd(iEl));
	}
	
	gsl_vector_free(tmpSd);
	gsl_vector_free(tmpMn);
}

MuGrp::MuGrp(const vector<MVnorm *> &dat, RanIndex &low, RanIndex &up, const string &outFlNam){
	if (low.getNgrp() != up.getNtot()) {
		cerr << "ERROR: # of elements in index to the upper level is wrong!!!" << endl;
		exit(1);
	}
	else if (low.getNtot() != dat.size()){
		cerr << "ERROR: the number of rows in data does not correspond to the number of points in the low index!!!" << endl;
		exit(1);
	}
	_lowLevel = &low;
	_upLevel  = &up;
	_outFlNam = outFlNam;
	remove(_outFlNam.c_str());
	
	gsl_matrix_free(_valueMat);
	gsl_vector *tmpMn = gsl_vector_alloc(dat[0]->len());
	gsl_vector *tmpSd = gsl_vector_alloc(dat[0]->len());
	
	_theta.resize(low.getNgrp());
	_valueMat = gsl_matrix_alloc(low.getNgrp(), dat[0]->len());
	for (size_t iEl = 0; iEl < low.getNgrp(); iEl++) {
		gsl_vector_set_zero(tmpMn);
		for (vector<size_t>::const_iterator el = low[iEl].begin(); el != low[iEl].end(); ++el) {
			gsl_vector_add(tmpMn, dat[*el]->getVec());
		}
		gsl_vector_scale(tmpMn, 1.0/low[iEl].size());
		gsl_matrix_set_row(_valueMat, iEl, tmpMn);
		for (size_t j = 0; j < dat[0]->len(); j++) {
			gsl_vector_set(tmpSd, j, fabs(gsl_vector_get(tmpMn, j)));
		}
		_theta[iEl] = new MVnormMu(_valueMat, iEl, tmpSd, _rV[0], (*_lowLevel)[iEl], _upLevel->priorInd(iEl));
	}
		
	gsl_vector_free(tmpSd);
	gsl_vector_free(tmpMn);
	
}

MuGrp::MuGrp(const Grp &dat, RanIndex &low, RanIndex &up){
	if (low.getNgrp() != up.getNtot()) {
		cerr << "ERROR: # of elements in index to the upper level is wrong!!!" << endl;
		exit(1);
	}
	else if (low.getNtot() != dat.Ndata()){
		cerr << "ERROR: the number of rows in data does not correspond to the number of points in the low index!!!" << endl;
		exit(1);
	}
	_lowLevel = &low;
	_upLevel  = &up;
	
	gsl_matrix_free(_valueMat);
	gsl_vector *tmpMn = gsl_vector_alloc(dat.phenD());
	gsl_vector *tmpSd = gsl_vector_alloc(dat.phenD());
	
	_theta.resize(low.getNgrp());
	_valueMat = gsl_matrix_alloc(low.getNgrp(), dat.phenD());
	
	for (size_t iEl = 0; iEl < low.getNgrp(); iEl++) {
		gsl_vector_set_zero(tmpMn);
		for (vector<size_t>::const_iterator el = low[iEl].begin(); el != low[iEl].end(); ++el) {
			gsl_vector_const_view tmpRow = gsl_matrix_const_row(dat.dMat(), *el);
			gsl_vector_add(tmpMn, &tmpRow.vector);
		}
		gsl_vector_scale(tmpMn, 1.0/low[iEl].size());
		gsl_matrix_set_row(_valueMat, iEl, tmpMn);
		
		for (size_t j = 0; j < dat[0]->len(); j++) {
			gsl_vector_set(tmpSd, j, fabs(gsl_vector_get(tmpMn, j)));
		}
		_theta[iEl] = new MVnormMu(_valueMat, iEl, tmpSd, _rV[0], (*_lowLevel)[iEl], _upLevel->priorInd(iEl));
	}
	
	
	gsl_vector_free(tmpSd);
	gsl_vector_free(tmpMn);
}

MuGrp::MuGrp(const Grp &dat, RanIndex &low, RanIndex &up, const string &outFlNam){
	if (low.getNgrp() != up.getNtot()) {
		cerr << "ERROR: # of elements in index to the upper level is wrong!!!" << endl;
		exit(1);
	}
	else if (low.getNtot() != dat.Ndata()){
		cerr << "ERROR: the number of rows in data does not correspond to the number of points in the low index!!!" << endl;
		exit(1);
	}
	_lowLevel = &low;
	_upLevel  = &up;
	_outFlNam = outFlNam;
	remove(_outFlNam.c_str());
	
	gsl_matrix_free(_valueMat);
	gsl_vector *tmpMn = gsl_vector_alloc(dat.phenD());
	gsl_vector *tmpSd = gsl_vector_alloc(dat.phenD());
	
	_theta.resize(low.getNgrp());
	_valueMat = gsl_matrix_alloc(low.getNgrp(), dat.phenD());
	
	for (size_t iEl = 0; iEl < low.getNgrp(); iEl++) {
		gsl_vector_set_zero(tmpMn);
		for (vector<size_t>::const_iterator el = low[iEl].begin(); el != low[iEl].end(); ++el) {
			gsl_vector_const_view tmpRow = gsl_matrix_const_row(dat.dMat(), *el);
			gsl_vector_add(tmpMn, &tmpRow.vector);
		}
		gsl_vector_scale(tmpMn, 1.0/low[iEl].size());
		gsl_matrix_set_row(_valueMat, iEl, tmpMn);
		
		for (size_t j = 0; j < dat[0]->len(); j++) {
			gsl_vector_set(tmpSd, j, fabs(gsl_vector_get(tmpMn, j)));
		}
		_theta[iEl] = new MVnormMu(_valueMat, iEl, tmpSd, _rV[0], (*_lowLevel)[iEl], _upLevel->priorInd(iEl));
	}
		
	gsl_vector_free(tmpSd);
	gsl_vector_free(tmpMn);
	
}

MuGrp::MuGrp(const Grp &dat, RanIndex &low) : Grp(){
	
	gsl_matrix_free(_valueMat);
	if (low.getNtot() != dat.dMat()->size1) {
		cerr << "RanIndex low has different number of elements (" << low.getNtot() << ") than Grp object dat has rows in _valueMat (" << dat.dMat()->size1 << ") in file " << __FILE__ << " on line " << __LINE__ << endl;
		exit(1);
	}
	
	_theta.resize(low.getNgrp());
	_valueMat = gsl_matrix_calloc(low.getNgrp(), dat.phenD());
	_lowLevel = &low;
	
 	for (size_t iEl = 0; iEl < low.getNgrp(); iEl++) {
		gsl_vector_view vlRw = gsl_matrix_row(_valueMat, iEl);
		for (vector<size_t>::const_iterator el = low[iEl].begin(); el != low[iEl].end(); ++el) {
			gsl_vector_const_view tmpRow = gsl_matrix_const_row(dat.dMat(), *el);
			gsl_vector_add(&vlRw.vector, &tmpRow.vector);
		}
		gsl_vector_scale(&vlRw.vector, 1.0/low[iEl].size());
	}
	
	for (size_t iMn = 0; iMn < low.getNgrp(); iMn++) {
		_theta[iMn] = new MVnormMu(_valueMat, iMn, (*_lowLevel)[iMn], 0);
		
	}
	
}

MuGrp::MuGrp(const Grp &dat, const Qgrp &q, RanIndex &low) : Grp(){
	
	gsl_matrix_free(_valueMat);
	gsl_vector *tmpVl = gsl_vector_alloc(dat.phenD());
	if (low.getNtot() != dat.dMat()->size1) {
		cerr << "RanIndex low has different number of elements (" << low.getNtot() << ") than Grp object dat has rows in _valueMat (" << dat.dMat()->size1 << ") in file " << __FILE__ << " on line " << __LINE__ << endl;
		exit(1);
	}
	else if (q.size() != dat.dMat()->size1) {
		cerr << "Weighting object Qgrp has different number of elements (" << q.size() << ") than Grp object dat has rows in _valueMat (" << dat.dMat()->size1 << ") in file " << __FILE__ << " on line " << __LINE__ << endl;
		exit(1);
	}
	
	_theta.resize(low.getNgrp());
	_valueMat = gsl_matrix_calloc(low.getNgrp(), dat.phenD());
	_lowLevel = &low;
	
	for (size_t iEl = 0; iEl < low.getNgrp(); iEl++) {
		gsl_vector_view vlRw = gsl_matrix_row(_valueMat, iEl);
		double qSum = 0.0;
		for (vector<size_t>::const_iterator el = low[iEl].begin(); el != low[iEl].end(); ++el) {
			qSum += q[*el];
			gsl_vector_const_view tmpRow = gsl_matrix_const_row(dat.dMat(), *el);
			gsl_vector_memcpy(tmpVl, &tmpRow.vector);
			gsl_vector_scale(tmpVl, q[*el]);
			gsl_vector_add(&vlRw.vector, tmpVl);
		}
		gsl_vector_scale(&vlRw.vector, 1.0/qSum);
		
		
	}
	
	for (size_t iMn = 0; iMn < low.getNgrp(); iMn++) {
		_theta[iMn] = new MVnormMu(_valueMat, iMn, (*_lowLevel)[iMn], 0);
		
	}
	
	gsl_vector_free(tmpVl);
}

MuGrp::MuGrp(const gsl_matrix *dat) : Grp(){
	
	gsl_matrix_free(_valueMat);
	
	_theta.resize(dat->size1);
	_valueMat = gsl_matrix_alloc(dat->size1, dat->size2);
	gsl_matrix_memcpy(_valueMat, dat);
	_lowLevel = new RanIndex(dat->size1);
	
	for (size_t iMn = 0; iMn < dat->size1; iMn++) {
		_theta[iMn] = new MVnormMu(_valueMat, iMn, (*_lowLevel)[iMn], 0);
		
	}
	
}

MuGrp::MuGrp(const gsl_matrix *dat, RanIndex &low) : Grp(){
	
	gsl_matrix_free(_valueMat);
	if (low.getNtot() != dat->size1) {
		cerr << "RanIndex low has different number of elements (" << low.getNtot() << ") than GSL matrix dat has rows (" << dat->size1 << ") in file " << __FILE__ << " on line " << __LINE__ << endl;
		exit(1);
	}
	
	_theta.resize(low.getNgrp());
	_valueMat = gsl_matrix_calloc(low.getNgrp(), dat->size2);
	_lowLevel = &low;
	
	for (size_t iEl = 0; iEl < low.getNgrp(); iEl++) {
		gsl_vector_view vlRw = gsl_matrix_row(_valueMat, iEl);
		for (vector<size_t>::const_iterator el = low[iEl].begin(); el != low[iEl].end(); ++el) {
			gsl_vector_const_view datRw = gsl_matrix_const_row(dat, *el);
			gsl_vector_add(&vlRw.vector, &datRw.vector);
		}
		gsl_vector_scale(&vlRw.vector, 1.0/low[iEl].size());
		
		
	}
	
	for (size_t iMn = 0; iMn < low.getNgrp(); iMn++) {
		_theta[iMn] = new MVnormMu(_valueMat, iMn, (*_lowLevel)[iMn], 0);
		
	}
	
}

MuGrp::MuGrp(const gsl_matrix *dat, const Qgrp &q, RanIndex &low) : Grp(){
	
	gsl_matrix_free(_valueMat);
	gsl_vector *tmpVl = gsl_vector_alloc(dat->size2);
	if (low.getNtot() != dat->size1) {
		cerr << "RanIndex low has different number of elements (" << low.getNtot() << ") than the GSL matrix dat has rows (" << dat->size1 << ") in file " << __FILE__ << " on line " << __LINE__ << endl;
		exit(1);
	}
	else if (q.size() != dat->size1) {
		cerr << "Weighting object Qgrp has different number of elements (" << q.size() << ") than the GSL matrix dat has rows (" << dat->size1 << ") in file " << __FILE__ << " on line " << __LINE__ << endl;
		exit(1);
	}
	
	_theta.resize(low.getNgrp());
	_valueMat = gsl_matrix_calloc(low.getNgrp(), dat->size2);
	_lowLevel = &low;
	
	for (size_t iEl = 0; iEl < low.getNgrp(); iEl++) {
		gsl_vector_view vlRw = gsl_matrix_row(_valueMat, iEl);
		double qSum = 0.0;
		for (vector<size_t>::const_iterator el = low[iEl].begin(); el != low[iEl].end(); ++el) {
			qSum += q[*el];
			gsl_matrix_get_col(tmpVl, dat, *el);
			gsl_vector_scale(tmpVl, q[*el]);
			gsl_vector_add(&vlRw.vector, tmpVl);
		}
		gsl_vector_scale(&vlRw.vector, 1.0/qSum);
		
		
	}
	
	for (size_t iMn = 0; iMn < low.getNgrp(); iMn++) {
		_theta[iMn] = new MVnormMu(_valueMat, iMn, (*_lowLevel)[iMn], 0);
		
	}
	
	gsl_vector_free(tmpVl);
}

MuGrp::MuGrp(const MuGrp &mG){
	gsl_matrix_free(_valueMat);
	_valueMat = gsl_matrix_alloc((mG._valueMat)->size1, (mG._valueMat)->size2);
	gsl_matrix_memcpy(_valueMat, mG._valueMat);
	_lowLevel = mG._lowLevel;
	_upLevel  = mG._upLevel;
	
	_theta.resize(_valueMat->size1);
	for (size_t iVrw = 0; iVrw < _valueMat->size1; iVrw++) {
		delete _theta[iVrw];
		_theta[iVrw] = new MVnormMu(_valueMat, iVrw, *(mG._theta[iVrw])->down(), *(mG._theta[iVrw])->up()); // can't simply copy _theta: not guaranteed that the elements will point to the right thing);
	}
}
MuGrp::MuGrp(const Grp &g){ // this copy constructor only copies the fMat (that points either to _valueMat or _fittedAll, depending on the inherited class)
	gsl_matrix_free(_valueMat);
	_valueMat = gsl_matrix_alloc((g.fMat())->size1, (g.fMat())->size2);
	gsl_matrix_memcpy(_valueMat, g.fMat());
	_lowLevel = g._lowLevel;
	_upLevel  = g._upLevel;
	
	_theta.resize(_valueMat->size1);
	for (size_t iVrw = 0; iVrw < _valueMat->size1; iVrw++) {
		delete _theta[iVrw];
		_theta[iVrw] = new MVnormMu(_valueMat, iVrw);
	}
}
MuGrp &MuGrp::operator=(const MuGrp &mG){
	gsl_matrix_free(_valueMat);
	_valueMat = gsl_matrix_alloc((mG._valueMat)->size1, (mG._valueMat)->size2);
	gsl_matrix_memcpy(_valueMat, mG._valueMat);
	_lowLevel = mG._lowLevel;
	_upLevel  = mG._upLevel;
	
	_theta.resize(_valueMat->size1);
	for (size_t iVrw = 0; iVrw < _valueMat->size1; iVrw++) {
		delete _theta[iVrw];
		_theta[iVrw] = new MVnormMu(_valueMat, iVrw, *(mG._theta[iVrw])->down(), *(mG._theta[iVrw])->up()); // can't simply copy _theta: not guaranteed that the elements will point to the right thing);
	}
	
	return *this;
}

// improper prior
void MuGrp::update(const Grp &dat, const SigmaI &SigIm){
	for (vector<MVnorm *>::const_iterator elm = _theta.begin(); elm != _theta.end(); ++elm) {
		(*elm)->update(dat, SigIm, _rV[0]);
	}
}
void MuGrp::update(const Grp &dat, const Qgrp &q, const SigmaI &SigIm){
	for (vector<MVnorm *>::const_iterator elm = _theta.begin(); elm != _theta.end(); ++elm) {
		(*elm)->update(dat, q, SigIm, _rV[0]);
	}
}

// 0-mean prior
void MuGrp::update(const Grp &dat, const SigmaI &SigIm, const SigmaI &SigIp){
	for (vector<MVnorm *>::const_iterator elm = _theta.begin(); elm != _theta.end(); ++elm) {
		(*elm)->update(dat, SigIm, SigIp, _rV[0]);
	}
}
void MuGrp::update(const Grp &dat, const Qgrp &q, const SigmaI &SigIm, const SigmaI &SigIp){
	for (vector<MVnorm *>::const_iterator elm = _theta.begin(); elm != _theta.end(); ++elm) {
		(*elm)->update(dat, q, SigIm, SigIp, _rV[0]);
	}
}
void MuGrp::update(const Grp &dat, const SigmaI &SigIm, const Qgrp &qPr, const SigmaI &SigIp){
	for (size_t iTh = 0; iTh < _theta.size(); iTh++) {
		_theta[iTh]->update(dat, SigIm, qPr[iTh], SigIp, _rV[0]);
	}
}
void MuGrp::update(const Grp &dat, const Qgrp &q, const SigmaI &SigIm, const Qgrp &qPr, const SigmaI &SigIp){
	for (size_t iTh = 0; iTh < _theta.size(); iTh++) {
		_theta[iTh]->update(dat, q, SigIm, qPr[iTh], SigIp, _rV[0]);
	}
}

// non-0-mean prior
void MuGrp::update(const Grp &dat, const SigmaI &SigIm, const Grp &muPr, const SigmaI &SigIp){
	for (vector<MVnorm *>::const_iterator elm = _theta.begin(); elm != _theta.end(); ++elm) {
		(*elm)->update(dat, SigIm, muPr, SigIp, _rV[0]);
	}
}
void MuGrp::update(const Grp &dat, const Qgrp &q, const SigmaI &SigIm, const Grp &muPr, const SigmaI &SigIp){
	for (vector<MVnorm *>::const_iterator elm = _theta.begin(); elm != _theta.end(); ++elm) {
		(*elm)->update(dat, q, SigIm, muPr, SigIp, _rV[0]);
	}
}
void MuGrp::update(const Grp &dat, const SigmaI &SigIm, const Grp &muPr, const Qgrp &qPr, const SigmaI &SigIp){
	for (size_t iTh = 0; iTh < _theta.size(); iTh++) {
		_theta[iTh]->update(dat, SigIm, muPr, qPr[iTh], SigIp, _rV[0]);
	}
}
void MuGrp::update(const Grp &dat, const Qgrp &q, const SigmaI &SigIm, const Grp &muPr, const Qgrp &qPr, const SigmaI &SigIp){
	for (size_t iTh = 0; iTh < _theta.size(); iTh++) {
		_theta[iTh]->update(dat, q, SigIm, muPr, qPr[iTh], SigIp, _rV[0]);
	}
}

/*
 *	MuGrpPEX methods
 */

MuGrpPEX::MuGrpPEX() : _outSigFlNam("MuPEXsig.gbin"), MuGrp(){
	_adjValMat = gsl_matrix_calloc(1, 1);
	_tSigIAt   = gsl_matrix_alloc(1, 1);
}

MuGrpPEX::MuGrpPEX(const Grp &dat, RanIndex &low, RanIndex &up, const double &sPr, const int &nThr) : _outSigFlNam("MuPEXsig.gbin"), MuGrp(){
	gsl_matrix_free(_valueMat);
	
	_ftA.resize(dat.phenD());
	_nThr = nThr;
	
	if (low.getNgrp() != up.getNtot()) {
		cerr << "ERROR: # of elements in index to the upper level is wrong!!!" << endl;
		exit(1);
	}
	else if (low.getNtot() != dat.Ndata()){
		cerr << "ERROR: the number of rows in data does not correspond to the number of points in the low index!!!" << endl;
		exit(1);
	}
	_lowLevel = &low;
	_upLevel  = &up;
	
	remove(_outFlNam.c_str());
	remove(_outSigFlNam.c_str());
	
	gsl_vector *tmpMn = gsl_vector_alloc(dat.phenD());
	gsl_vector *tmpSd = gsl_vector_alloc(dat.phenD());
	
	_theta.resize(low.getNgrp());
	_valueMat  = gsl_matrix_alloc(low.getNgrp(), dat.phenD());
	_adjValMat = gsl_matrix_calloc(low.getNgrp(), dat.phenD());
	_tSigIAt   = gsl_matrix_alloc(dat.phenD(), dat.phenD());
	_A         = Apex(sPr, dat.phenD(), _ftA);
	
	for (size_t iEl = 0; iEl < low.getNgrp(); iEl++) {
		gsl_vector_set_zero(tmpMn);
		for (vector<size_t>::const_iterator el = low[iEl].begin(); el != low[iEl].end(); ++el) {
			gsl_vector_add(tmpMn, dat[*el]->getVec());
		}
		gsl_vector_scale(tmpMn, 1.0/low[iEl].size());
		gsl_matrix_set_row(_valueMat, iEl, tmpMn);
		
		for (size_t j = 0; j < dat[0]->len(); j++) {
			gsl_vector_set(tmpSd, j, fabs(gsl_vector_get(tmpMn, j)));
		}
		
	}
	
	for (size_t jCl = 0; jCl < _valueMat->size2; jCl++) {
		_ftA[jCl].resize(dat.Ndata()*dat.phenD());
	}
	for (size_t iMn = 0; iMn < low.getNgrp(); iMn++) {
		_theta[iMn] = new MVnormMuPEX(_valueMat, iMn, tmpSd, _rV[0], (*_lowLevel)[iMn], _upLevel->priorInd(iMn), _A, _tSigIAt);
	}
	
	_updateFitted();
	gsl_vector_free(tmpSd);
	gsl_vector_free(tmpMn);
	
}

MuGrpPEX::MuGrpPEX(const Grp &dat, RanIndex &low, RanIndex &up, const string outMuFlNam, const double &sPr, const int &nThr) : _outSigFlNam("MuPEXsig.gbin"), MuGrp(){
	gsl_matrix_free(_valueMat);
	
	_ftA.resize(dat.phenD());
	_nThr = nThr;
	
	if (low.getNgrp() != up.getNtot()) {
		cerr << "ERROR: # of elements in index to the upper level is wrong!!!" << endl;
		exit(1);
	}
	else if (low.getNtot() != dat.Ndata()){
		cerr << "ERROR: the number of rows in data does not correspond to the number of points in the low index!!!" << endl;
		exit(1);
	}
	_lowLevel = &low;
	_upLevel  = &up;
	
	_outFlNam = outMuFlNam;
	remove(_outFlNam.c_str());
	remove(_outSigFlNam.c_str());
	
	gsl_vector *tmpMn = gsl_vector_alloc(dat.phenD());
	gsl_vector *tmpSd = gsl_vector_alloc(dat.phenD());
	
	_theta.resize(low.getNgrp());
	_valueMat  = gsl_matrix_alloc(low.getNgrp(), dat.phenD());
	_adjValMat = gsl_matrix_calloc(low.getNgrp(), dat.phenD());
	_tSigIAt   = gsl_matrix_alloc(dat.phenD(), dat.phenD());
	_A         = Apex(sPr, dat.phenD(), _ftA);
	
	for (size_t iEl = 0; iEl < low.getNgrp(); iEl++) {
		gsl_vector_set_zero(tmpMn);
		for (vector<size_t>::const_iterator el = low[iEl].begin(); el != low[iEl].end(); ++el) {
			gsl_vector_add(tmpMn, dat[*el]->getVec());
		}
		gsl_vector_scale(tmpMn, 1.0/low[iEl].size());
		gsl_matrix_set_row(_valueMat, iEl, tmpMn);
		
		for (size_t j = 0; j < dat[0]->len(); j++) {
			gsl_vector_set(tmpSd, j, fabs(gsl_vector_get(tmpMn, j)));
		}
		
	}
	
	for (size_t jCl = 0; jCl < _valueMat->size2; jCl++) {
		_ftA[jCl].resize(dat.Ndata()*dat.phenD());
	}
	for (size_t iMn = 0; iMn < low.getNgrp(); iMn++) {
		_theta[iMn] = new MVnormMuPEX(_valueMat, iMn, tmpSd, _rV[0], (*_lowLevel)[iMn], _upLevel->priorInd(iMn), _A, _tSigIAt);
	}
	
	_updateFitted();
	gsl_vector_free(tmpSd);
	gsl_vector_free(tmpMn);
	
}

MuGrpPEX::MuGrpPEX(const MuGrpPEX &mGp){
	gsl_matrix_free(_valueMat);
	_valueMat = gsl_matrix_alloc((mGp._valueMat)->size1, (mGp._valueMat)->size2);
	gsl_matrix_memcpy(_valueMat, mGp._valueMat);
	gsl_matrix_free(_adjValMat);
	_adjValMat = gsl_matrix_alloc((mGp._adjValMat)->size1, (mGp._adjValMat)->size2);
	gsl_matrix_memcpy(_adjValMat, mGp._adjValMat);
	_tSigIAt = gsl_matrix_alloc((mGp._tSigIAt)->size1, (mGp._tSigIAt)->size2);
	gsl_matrix_memcpy(_tSigIAt, mGp._tSigIAt);
	_lowLevel = mGp._lowLevel;
	_upLevel  = mGp._upLevel;
	_A        = mGp._A;
	_ftA      = mGp._ftA;
	_nThr     = mGp._nThr;
	
	_theta.resize(_valueMat->size1);
	for (size_t iVrw = 0; iVrw < _valueMat->size1; iVrw++) {
		delete _theta[iVrw];
		_theta[iVrw] = new MVnormMuPEX(_valueMat, iVrw, *(mGp._theta[iVrw])->down(), *(mGp._theta[iVrw])->up(), _A, _tSigIAt); // can't simply copy _theta: not guaranteed that the elements will point to the right thing);
	}
}

MuGrpPEX &MuGrpPEX::operator=(const MuGrpPEX &mGp){
	gsl_matrix_free(_valueMat);
	_valueMat = gsl_matrix_alloc((mGp._valueMat)->size1, (mGp._valueMat)->size2);
	gsl_matrix_memcpy(_valueMat, mGp._valueMat);
	gsl_matrix_free(_adjValMat);
	_adjValMat = gsl_matrix_alloc((mGp._adjValMat)->size1, (mGp._adjValMat)->size2);
	gsl_matrix_memcpy(_adjValMat, mGp._adjValMat);
	_tSigIAt = gsl_matrix_alloc((mGp._tSigIAt)->size1, (mGp._tSigIAt)->size2);
	gsl_matrix_memcpy(_tSigIAt, mGp._tSigIAt);
	_lowLevel = mGp._lowLevel;
	_upLevel  = mGp._upLevel;
	_A        = mGp._A;
	_ftA      = mGp._ftA;
	_nThr     = mGp._nThr;
	
	_theta.resize(_valueMat->size1);
	for (size_t iVrw = 0; iVrw < _valueMat->size1; iVrw++) {
		delete _theta[iVrw];
		_theta[iVrw] = new MVnormMuPEX(_valueMat, iVrw, *(mGp._theta[iVrw])->down(), *(mGp._theta[iVrw])->up(), _A, _tSigIAt); // can't simply copy _theta: not guaranteed that the elements will point to the right thing);
	}
	
	return *this;
}

MuGrpPEX::~MuGrpPEX() {
	gsl_matrix_free(_adjValMat);
	gsl_matrix_free(_tSigIAt);
}

void MuGrpPEX::save(){
	FILE *outH = fopen(_outFlNam.c_str(), "a");
	gsl_matrix_fwrite(outH, _adjValMat);
	fclose(outH);

}

void MuGrpPEX::save(const string &outFlNam){
	if (_outFlNam == outFlNam) {
		FILE *outH = fopen(outFlNam.c_str(), "a");
		gsl_matrix_fwrite(outH, _adjValMat);
		fclose(outH);
		
	}
	else {
		_outFlNam = outFlNam;
		remove(_outFlNam.c_str());
		FILE *outH = fopen(outFlNam.c_str(), "a");
		gsl_matrix_fwrite(outH, _adjValMat);
		fclose(outH);
	}
	
}

void MuGrpPEX::save(const SigmaI &SigI){
	gsl_matrix *Sig   = gsl_matrix_alloc(SigI.getMat()->size1, SigI.getMat()->size1);
	gsl_matrix *SigTr = gsl_matrix_alloc(SigI.getMat()->size1, SigI.getMat()->size1);
	gsl_matrix_memcpy(Sig, SigI.getMat());
	gsl_linalg_cholesky_decomp(Sig);
	gsl_linalg_cholesky_invert(Sig);
	
	gsl_blas_dsymm(CblasLeft, CblasLower, 1.0, Sig, _A.getMat(), 0.0, SigTr);
	gsl_blas_dgemm(CblasTrans, CblasNoTrans, 1.0, _A.getMat(), SigTr, 0.0, Sig);
	
	FILE *outS = fopen(SigI.getOutFile().c_str(), "a");
	gsl_matrix_fwrite(outS, Sig);
	fclose(outS);
	
	gsl_matrix_free(Sig);
	gsl_matrix_free(SigTr);
	
	FILE *outM = fopen(_outFlNam.c_str(), "a");
	gsl_matrix_fwrite(outM, _adjValMat);
	fclose(outM);
}

MuGrp MuGrpPEX::mean(RanIndex &grp){
	MuGrp res(_adjValMat, grp);
	return res;
}
const MuGrp MuGrpPEX::mean(RanIndex &grp) const{
	MuGrp res(_adjValMat, grp);
	return res;
}

MuGrp MuGrpPEX::mean(RanIndex &grp, const Qgrp &q){
	MuGrp res(_adjValMat, q, grp);
	return res;
}
const MuGrp MuGrpPEX::mean(RanIndex &grp, const Qgrp &q) const{
	MuGrp res(_adjValMat, q, grp);
	return res;
}

void MuGrpPEX::update(const Grp &dat, const SigmaI &SigIm, const SigmaI &SigIp){
	
	gsl_blas_dsymm(CblasRight, CblasLower, 1.0, SigIm.getMat(), _A.getMat(), 0.0, _tSigIAt);
	
	for (vector<MVnorm *>::const_iterator elm = _theta.begin(); elm != _theta.end(); ++elm) {
		(*elm)->update(dat, SigIm, SigIp, _rV[0]);
	}
	
	_A.update(dat, _valueMat, SigIm, *_lowLevel);
	_updateFitted();
}
void MuGrpPEX::update(const Grp &dat, const Qgrp &q, const SigmaI &SigIm, const SigmaI &SigIp){
	gsl_blas_dsymm(CblasRight, CblasLower, 1.0, SigIm.getMat(), _A.getMat(), 0.0, _tSigIAt);
	
	for (vector<MVnorm *>::const_iterator elm = _theta.begin(); elm != _theta.end(); ++elm) {
		(*elm)->update(dat, q, SigIm, SigIp, _rV[0]);
	}
	
	_A.update(dat, _valueMat, q, SigIm, *_lowLevel);
	_updateFitted();
}
void MuGrpPEX::update(const Grp &dat, const SigmaI &SigIm, const Qgrp &qPr, const SigmaI &SigIp){
	gsl_blas_dsymm(CblasRight, CblasLower, 1.0, SigIm.getMat(), _A.getMat(), 0.0, _tSigIAt);
	
	for (int iTh = 0; iTh < _theta.size(); iTh++) {
		_theta[iTh]->update(dat, SigIm, qPr[iTh], SigIp, _rV[0]);
	}
	
	_A.update(dat, _valueMat, SigIm, *_lowLevel);
	_updateFitted();
}
void MuGrpPEX::update(const Grp &dat, const Qgrp &q, const SigmaI &SigIm, const Qgrp &qPr, const SigmaI &SigIp){
	gsl_blas_dsymm(CblasRight, CblasLower, 1.0, SigIm.getMat(), _A.getMat(), 0.0, _tSigIAt);
	
	for (int iTh = 0; iTh < _theta.size(); iTh++) {
		_theta[iTh]->update(dat, q, SigIm, qPr[iTh], SigIp, _rV[0]);
	}
	
	_A.update(dat, _valueMat, q, SigIm, *_lowLevel);
	_updateFitted();
}

void MuGrpPEX::update(const Grp &dat, const SigmaI &SigIm, const Grp &muPr, const SigmaI &SigIp){
	
	gsl_blas_dsymm(CblasRight, CblasLower, 1.0, SigIm.getMat(), _A.getMat(), 0.0, _tSigIAt);
	
	for (vector<MVnorm *>::const_iterator elm = _theta.begin(); elm != _theta.end(); ++elm) {
		(*elm)->update(dat, SigIm, muPr, SigIp, _rV[0]);
	}
	
	_A.update(dat, _valueMat, SigIm, *_lowLevel);
	_updateFitted();
	
}
void MuGrpPEX::update(const Grp &dat, const Qgrp &q, const SigmaI &SigIm, const Grp &muPr, const SigmaI &SigIp){
	
	gsl_blas_dsymm(CblasRight, CblasLower, 1.0, SigIm.getMat(), _A.getMat(), 0.0, _tSigIAt);
	
	for (vector<MVnorm *>::const_iterator elm = _theta.begin(); elm != _theta.end(); ++elm) {
		(*elm)->update(dat, q, SigIm, muPr, SigIp, _rV[0]);
	}
	
	_A.update(dat, _valueMat, q, SigIm, *_lowLevel);
	_updateFitted();
	
}
void MuGrpPEX::update(const Grp &dat, const SigmaI &SigIm, const Grp &muPr, const Qgrp &qPr, const SigmaI &SigIp){
	gsl_blas_dsymm(CblasRight, CblasLower, 1.0, SigIm.getMat(), _A.getMat(), 0.0, _tSigIAt);
	
	for (int iTh = 0; iTh < _theta.size(); iTh++) {
		_theta[iTh]->update(dat, SigIm, muPr, qPr[iTh], SigIp, _rV[0]);
	}
	
	_A.update(dat, _valueMat, SigIm, *_lowLevel);
	_updateFitted();
	
}
void MuGrpPEX::update(const Grp &dat, const Qgrp &q, const SigmaI &SigIm, const Grp &muPr, const Qgrp &qPr, const SigmaI &SigIp){
	gsl_blas_dsymm(CblasRight, CblasLower, 1.0, SigIm.getMat(), _A.getMat(), 0.0, _tSigIAt);
	
	for (int iTh = 0; iTh < _theta.size(); iTh++) {
		_theta[iTh]->update(dat, q, SigIm, muPr, qPr[iTh], SigIp, _rV[0]);
	}
	
	_A.update(dat, _valueMat, q, SigIm, *_lowLevel);
	_updateFitted();
	
}

void MuGrpPEX::_updateFitted(){
	
#pragma omp parallel num_threads(_nThr)
	{
		gsl_vector *tmpVec = gsl_vector_alloc(_valueMat->size2); // have to make a OMP team copy of the temp array
		double pSum;
		
#pragma omp for
		for (int prdRow = 0; prdRow < (*_lowLevel).getNgrp(); prdRow++) {
			for (int bcol = 0; bcol < _A.getMat()->size2; bcol++) {
				pSum = 0.0;
				gsl_matrix_get_row(tmpVec, _valueMat, prdRow);
				gsl_vector_view evCol = gsl_matrix_column(_A.getMat(), bcol);
				gsl_vector_mul(tmpVec, &evCol.vector);
				for (int iBt = 0; iBt < _A.getMat()->size1; iBt++) {
					pSum += gsl_vector_get(tmpVec, iBt);
				}
				gsl_matrix_set(_adjValMat,  prdRow, bcol, pSum);
				for (int jBt = 0; jBt < _A.getMat()->size1; jBt++) {
					double pSumDif = pSum - gsl_vector_get(tmpVec, jBt);
					for (size_t eachLw = 0; eachLw < (*_lowLevel)[prdRow].size(); eachLw++) {
						_ftA[jBt][(*_lowLevel)[prdRow][eachLw]*(_valueMat->size2) + bcol] = pSumDif;
					}
				}
			}
		}
		
		gsl_vector_free(tmpVec);
	} // end parallel block
}


/*
 *	MuGrpMiss methods
 */

MuGrpMiss::MuGrpMiss(const string &datFlNam, const string &misMatFlNam, const string &misVecFlNam, RanIndex &up, const size_t &d) : MuGrp(){
	gsl_matrix_free(_valueMat);
	
	_upLevel                  = &up;
	_valueMat                 = gsl_matrix_alloc(up.getNtot(), d);
	gsl_matrix_int *tmpMisMat = gsl_matrix_int_alloc(up.getNtot(), d);
	gsl_vector_int *tmpMisVec = gsl_vector_int_alloc(up.getNtot());
	gsl_vector *tmpRow        = gsl_vector_calloc(d);
	gsl_vector *tmpSd         = gsl_vector_alloc(d);
	vector<vector<size_t> > misPhenID;
	
	FILE *datIn = fopen(datFlNam.c_str(), "r");
	if (datIn == NULL) {
		cerr << "ERROR: cannot open data file " << datIn << " in MuGrpMiss" << endl;
		exit(-1);
	}
	gsl_matrix_fread(datIn, _valueMat);
	fclose(datIn);
	
	FILE *mmIn = fopen(misMatFlNam.c_str(), "r");
	if (mmIn == NULL) {
		cerr << "ERROR: cannot missing data index file " << misMatFlNam << " in MuGrpMiss" << endl;
		exit(-1);
	}
	gsl_matrix_int_fread(mmIn, tmpMisMat);
	fclose(mmIn);
	
	FILE *mvIn = fopen(misVecFlNam.c_str(), "r"); // this has the number of missing phenotypes
	if (datIn == NULL) {
		cerr << "ERROR: cannot open number of missing values file " << misVecFlNam << " in MuGrpMiss" << endl;
		exit(-1);
	}
	gsl_vector_int_fread(mvIn, tmpMisVec);
	fclose(mvIn);
	
	_theta.resize(up.getNtot());
	
	for (size_t iDatLn = 0; iDatLn < up.getNtot(); iDatLn++) {
		size_t curNmis = gsl_vector_int_get(tmpMisVec, iDatLn);
		
		if (curNmis) {
			gsl_vector_int_view mRow = gsl_matrix_int_row(tmpMisMat, iDatLn);
			_misInd.push_back(iDatLn);
			misPhenID.push_back(vector<size_t>(curNmis));
			size_t mP = 0;
			for (size_t iPh = 0; iPh < d; iPh++) {
				if (gsl_vector_int_get(&mRow.vector, iPh) == 1) {
					(misPhenID.back())[mP] = iPh;
					mP++;
				}
			}
			
		}
		else{
			gsl_vector_view datRow = gsl_matrix_row(_valueMat, iDatLn);
			gsl_vector_add(tmpRow, &datRow.vector); // calculating the mean among all the rows with no missing data for use in initializiation
			_theta[iDatLn] = new MVnormMu(_valueMat, iDatLn, _upLevel->priorInd(iDatLn)); // nothing to be done for rows with no missing phenotypes
		}
	}
	
	cout << "Found " << _misInd.size() << " rows with missing phenotypes" << endl;
	gsl_vector_scale(tmpRow, 1.0/(up.getNtot() - _misInd.size()));
	for (size_t j = 0; j < d; j++) {
		gsl_vector_set(tmpSd, j, fabs(gsl_vector_get(tmpRow, j)));
	}
	
	for (size_t msI = 0; msI < _misInd.size(); msI++) {
		gsl_vector_view msRw = gsl_matrix_row(_valueMat, _misInd[msI]);
		for (vector<size_t>::iterator phIt = misPhenID[msI].begin(); phIt != misPhenID[msI].end(); ++phIt) {
			gsl_vector_set(&msRw.vector, *phIt, gsl_vector_get(tmpRow, *phIt) + gsl_ran_gaussian_ziggurat(_rV[0], gsl_vector_get(tmpSd, *phIt))); // replacing the missing phenotypes with initial values
		}
		_theta[_misInd[msI]] = new MVnormMuMiss(_valueMat, _misInd[msI], _upLevel->priorInd(_misInd[msI]), misPhenID[msI]); // will indirectly modify _valueMat, but should be safe
	}
	
	gsl_matrix_int_free(tmpMisMat);
	gsl_vector_int_free(tmpMisVec);
	gsl_vector_free(tmpRow);
	gsl_vector_free(tmpSd);
}

MuGrpMiss::MuGrpMiss(const MuGrpMiss &mG){
	_misInd = mG._misInd;
	gsl_matrix_free(_valueMat);
	_valueMat   = gsl_matrix_alloc((mG._valueMat)->size1, (mG._valueMat)->size2);
	gsl_matrix_memcpy(_valueMat, mG._valueMat);
	
	_upLevel  = mG._upLevel;
	
	_theta.resize(_valueMat->size1);
	size_t mI = 0;
	for (size_t iVrw = 0; iVrw < _valueMat->size1; iVrw++) {
		if (iVrw == _misInd[mI]) {
			if (mI < _misInd.size()) {
				mI++;
			}
			_theta[iVrw] = new MVnormMuMiss(_valueMat, iVrw, *(mG._theta[iVrw])->up(), (mG._theta[iVrw])->getMisPhen());
		}
		else {
			_theta[iVrw] = new MVnormMu(_valueMat, iVrw, *(mG._theta[iVrw])->up());
		}
		
	}
	
}
MuGrpMiss & MuGrpMiss::operator=(const MuGrpMiss &mG){
	_misInd = mG._misInd;
	gsl_matrix_free(_valueMat);
	_valueMat   = gsl_matrix_alloc((mG._valueMat)->size1, (mG._valueMat)->size2);
	gsl_matrix_memcpy(_valueMat, mG._valueMat);
	
	_upLevel  = mG._upLevel;
	
	_theta.resize(_valueMat->size1);
	size_t mI = 0;
	for (size_t iVrw = 0; iVrw < _valueMat->size1; iVrw++) {
		if (iVrw == _misInd[mI]) {
			if (mI < _misInd.size()) {
				mI++;
			}
			_theta[iVrw] = new MVnormMuMiss(_valueMat, iVrw, *(mG._theta[iVrw])->up(), (mG._theta[iVrw])->getMisPhen());
		}
		else {
			_theta[iVrw] = new MVnormMu(_valueMat, iVrw, *(mG._theta[iVrw])->up());
		}
		
	}
	
	return *this;
}

void MuGrpMiss::update(const Grp &mu, const SigmaI &SigIm){
	for (vector<size_t>::iterator msIt = _misInd.begin(); msIt != _misInd.end(); ++msIt) {
		_theta[*msIt]->update(mu, SigIm, _rV[0]);
	}
}

void MuGrpMiss::update(const Grp &mu, const SigmaI &SigIm, const SigmaI &SigIp){
	for (vector<size_t>::iterator msIt = _misInd.begin(); msIt != _misInd.end(); ++msIt) {
		_theta[*msIt]->update(mu, SigIm, SigIp, _rV[0]);
	}
}

/*
 * MuGrpEE methods
 */
MuGrpEE::MuGrpEE() : MuGrp() {
	_errorInvVar = gsl_matrix_calloc(1, 1);
	_meanVal = gsl_matrix_calloc(1, 1);
}
MuGrpEE::MuGrpEE(const string &datFlNam, const string &varFlNam, const string &indFlNam, RanIndex &up, const size_t &d) : MuGrp(datFlNam, up, d){
	
	ifstream varIn(indFlNam.c_str());
	
	if (!varIn) {
		cerr << "ERROR: cannot open file " << indFlNam << " for reading error variance index in MuGrpEE initialization" << endl;
		exit(-1);
	}
	int tmpIn;
	while (varIn >> tmpIn) {
		if (tmpIn < 0) {
			cerr << "ERROR: negative indexes not alllowed when reading error variance index from file " << varFlNam << " on initialization of MuGrpEE" << endl;
			exit(-1);
		} else if (tmpIn > d){
			cerr << "ERROR: value of the index (" << tmpIn << ") exceeds the number of variables (" << d << "). Check if you saved a base-0 index for MuGrpEE" << endl;
			exit(-1);
		}
		_errInd.push_back(tmpIn);
	}
	if (_errInd.size() > d) {
		cerr << "ERROR: length of index of error variances (" << _errInd.size() << ") exceeds the total number of variables (" << d << ") while initializing MuGrpEE" << endl;
		exit(-1);
	}
	
	varIn.close();
	_errorInvVar = gsl_matrix_alloc(_valueMat->size1, _errInd.size());
	FILE *errIn = fopen(varFlNam.c_str(), "r");
	if (errIn == NULL) {
		cerr << "ERROR: Failed to open file " << varFlNam << " when inititializing MuGrpEE" << endl;
		exit(-1);
	}
	gsl_matrix_fread(errIn, _errorInvVar);
	fclose(errIn);
	for (size_t iRw = 0; iRw < _errorInvVar->size1; iRw++) {
		for (size_t jCl = 0; jCl < _errorInvVar->size2; jCl++) {
			double var = gsl_matrix_get(_errorInvVar, iRw, jCl);
			if ( var >= GSL_MACH_EPS) {
				gsl_matrix_set(_errorInvVar, iRw, jCl, 1.0/var);
			}
			else {
				cerr << "ERROR: experimental error variance " << var << " in row " << iRw << " and column " << jCl << " is too small in MuGrpEE initialization." << endl;
				exit(-1);
			}
		}
	}
	
	_meanVal = gsl_matrix_alloc(_errorInvVar->size1, _errorInvVar->size2);
	for (size_t jCl = 0; jCl < _meanVal->size2; jCl++) {
		gsl_vector_view col = gsl_matrix_column(_valueMat, _errInd[jCl]);
		gsl_matrix_set_col(_meanVal, jCl, &col.vector);
	}
	
}

MuGrpEE::MuGrpEE(const string &datFlNam, const string &varFlNam, const vector<size_t> &varInd, RanIndex &up, const size_t &d) : MuGrp(datFlNam, up, d){
	if (varInd.size() > d) {
		cerr << "ERROR: more elements in the experimenatl error index (" << varInd.size() << ") than overall variables (" << d << ") in MuGrpEE initialization" << endl;
		exit(-1);
	}
	if (varInd[*max_element(varInd.begin(), varInd.end())] > d) {
		cerr << "ERROR: largest element in the experimenatl error index (" << varInd.size() << ") is larger than the number of overall variables (" << d << ") in MuGrpEE initialization" << endl;
		exit(-1);
	}
	_errorInvVar = gsl_matrix_alloc(_valueMat->size1, varInd.size());
	_errInd   = varInd;
	
	
	FILE *errIn = fopen(varFlNam.c_str(), "r");
	if (errIn == NULL) {
		cerr << "ERROR: Failed to open file " << varFlNam << " when inititializing MuGrpEE" << endl;
		exit(-1);
	}
	gsl_matrix_fread(errIn, _errorInvVar);
	fclose(errIn);
	
	for (size_t iRw = 0; iRw < _errorInvVar->size1; iRw++) {
		for (size_t jCl = 0; jCl < _errorInvVar->size2; jCl++) {
			double var = gsl_matrix_get(_errorInvVar, iRw, jCl);
			if ( var >= GSL_MACH_EPS) {
				gsl_matrix_set(_errorInvVar, iRw, jCl, 1.0/var);
			}
			else {
				cerr << "ERROR: experimental error variance " << var << " in row " << iRw << " and column " << jCl << " is too small in MuGrpEE initialization." << endl;
				exit(-1);
			}
		}
	}
	
	_meanVal = gsl_matrix_alloc(_errorInvVar->size1, _errorInvVar->size2);
	for (size_t jCl = 0; jCl < _meanVal->size2; jCl++) {
		gsl_vector_view col = gsl_matrix_column(_valueMat, _errInd[jCl]);
		gsl_matrix_set_col(_meanVal, jCl, &col.vector);
	}
}

MuGrpEE::~MuGrpEE() {
	gsl_matrix_free(_errorInvVar);
	gsl_matrix_free(_meanVal);
}

MuGrpEE::MuGrpEE(const MuGrpEE &mG){
	gsl_matrix_free(_valueMat);
	gsl_matrix_free(_errorInvVar);
	gsl_matrix_free(_meanVal);
	_valueMat = gsl_matrix_alloc((mG._valueMat)->size1, (mG._valueMat)->size2);
	gsl_matrix_memcpy(_valueMat, mG._valueMat);
	_errorInvVar = gsl_matrix_alloc((mG._errorInvVar)->size1, (mG._errorInvVar)->size2);
	gsl_matrix_memcpy(_errorInvVar, mG._errorInvVar);
	_meanVal = gsl_matrix_alloc((mG._meanVal)->size1, (mG._meanVal)->size2);
	gsl_matrix_memcpy(_meanVal, mG._meanVal);
	_upLevel  = mG._upLevel;
	
	_theta.resize(_valueMat->size1);
	for (size_t iVrw = 0; iVrw < _valueMat->size1; iVrw++) {
		delete _theta[iVrw];
		_theta[iVrw] = new MVnormMu(_valueMat, iVrw, *(mG._theta[iVrw])->down(), *(mG._theta[iVrw])->up()); // can't simply copy _theta: not guaranteed that the elements will point to the right thing);
	}
}
MuGrpEE & MuGrpEE::operator=(const MuGrpEE &mG){
	gsl_matrix_free(_valueMat);
	gsl_matrix_free(_errorInvVar);
	gsl_matrix_free(_meanVal);
	_valueMat = gsl_matrix_alloc((mG._valueMat)->size1, (mG._valueMat)->size2);
	gsl_matrix_memcpy(_valueMat, mG._valueMat);
	_errorInvVar = gsl_matrix_alloc((mG._errorInvVar)->size1, (mG._errorInvVar)->size2);
	gsl_matrix_memcpy(_errorInvVar, mG._errorInvVar);
	_meanVal = gsl_matrix_alloc((mG._meanVal)->size1, (mG._meanVal)->size2);
	gsl_matrix_memcpy(_meanVal, mG._meanVal);
	_upLevel  = mG._upLevel;
	
	_theta.resize(_valueMat->size1);
	for (size_t iVrw = 0; iVrw < _valueMat->size1; iVrw++) {
		delete _theta[iVrw];
		_theta[iVrw] = new MVnormMu(_valueMat, iVrw, *(mG._theta[iVrw])->down(), *(mG._theta[iVrw])->up()); // can't simply copy _theta: not guaranteed that the elements will point to the right thing);
	}
	
	return *this;
}

// all of these are actually priors
void MuGrpEE::update(const Grp &muPr, const SigmaI &SigIm){
	for (size_t iRw = 0; iRw < _valueMat->size1; iRw++) {
		for (size_t jECl = 0; jECl < _errInd.size(); jECl++) {
			double var = 1.0/(gsl_matrix_get(_errorInvVar, iRw, jECl) + gsl_matrix_get(SigIm.getMat(), _errInd[jECl], _errInd[jECl]));
			double mn  = var * ( gsl_matrix_get(_meanVal, iRw, jECl) * gsl_matrix_get(_errorInvVar, iRw, jECl) +  gsl_matrix_get(SigIm.getMat(), _errInd[jECl], _errInd[jECl]) * gsl_matrix_get(muPr.fMat(), _upLevel->priorInd(iRw), _errInd[jECl]) );
			gsl_matrix_set(_valueMat, iRw, _errInd[jECl], mn + gsl_ran_gaussian_ziggurat(_rV[0], sqrt(var)));
		}
	}
}
void MuGrpEE::update(const Grp &muPr, const Qgrp &q, const SigmaI &SigIm){
	for (size_t iRw = 0; iRw < _valueMat->size1; iRw++) {
		for (size_t jECl = 0; jECl < _errInd.size(); jECl++) {
			double var = 1.0/(gsl_matrix_get(_errorInvVar, iRw, jECl) + q[iRw] * gsl_matrix_get(SigIm.getMat(), _errInd[jECl], _errInd[jECl]));
			double mn  = var * ( gsl_matrix_get(_meanVal, iRw, jECl) * gsl_matrix_get(_errorInvVar, iRw, jECl) +  q[iRw] * gsl_matrix_get(SigIm.getMat(), _errInd[jECl], _errInd[jECl]) * gsl_matrix_get(muPr.fMat(), _upLevel->priorInd(iRw), _errInd[jECl]) );
			gsl_matrix_set(_valueMat, iRw, _errInd[jECl], mn + gsl_ran_gaussian_ziggurat(_rV[0], sqrt(var)));
		}
	}
}

/*
 * MuGrpEEmiss methods
 */

MuGrpEEmiss::MuGrpEEmiss(const string &datFlNam, const string &varFlNam, const string &indFlNam, const string &misMatFlNam, const string &misVecFlNam, RanIndex &up, const size_t &d) : MuGrpMiss(datFlNam, misMatFlNam, misVecFlNam, up, d) {
	
	gsl_matrix_int *tmpMisMat = gsl_matrix_int_alloc(_valueMat->size1, d);
	FILE *mmIn = fopen(misMatFlNam.c_str(), "r");
	if (mmIn == NULL) {
		cerr << "ERROR: cannot open missing data index file " << misMatFlNam << " in MuGrpMiss" << endl;
		exit(-1);
	}
	gsl_matrix_int_fread(mmIn, tmpMisMat);
	fclose(mmIn);
	
	list<size_t> tmpErrInd;
	ifstream varIn(indFlNam.c_str());
	
	if (!varIn) {
		cerr << "ERROR: cannot open file " << indFlNam << " for reading error variance index in MuGrpEEmiss initialization" << endl;
		exit(-1);
	}
	int tmpIn;
	while (varIn >> tmpIn) {
		if (tmpIn < 0) {
			cerr << "ERROR: negative indexes not allowed when reading error variance index from file " << indFlNam << " on initialization of MuGrpEEmiss" << endl;
			exit(-1);
		} else if (tmpIn > d){
			cerr << "ERROR: value of the index (" << tmpIn << ") exceeds the number of variables (" << d << "). Check if you saved a base-0 index for MuGrpEEmiss" << endl;
			exit(-1);
		}
		tmpErrInd.push_back(tmpIn);
	}
	if (tmpErrInd.size() > d) {
		cerr << "ERROR: length of index of error variances (" << tmpErrInd.size() << ") exceeds the total number of variables (" << d << ") while initializing MuGrpEEmiss" << endl;
		exit(-1);
	}
	varIn.close();
	
	for (size_t iDat = 0; iDat < _valueMat->size1; iDat++) {
		_missErrMat.push_back(tmpErrInd);
	}
	
	gsl_matrix *tmpEV = gsl_matrix_alloc(_valueMat->size1, tmpErrInd.size());
	FILE *errIn = fopen(varFlNam.c_str(), "r");
	if (errIn == NULL) {
		cerr << "ERROR: Failed to open file " << varFlNam << " when inititializing MuGrpEEmiss" << endl;
		exit(-1);
	}
	gsl_matrix_fread(errIn, tmpEV);
	fclose(errIn);
	
	_errorInvVar.resize(tmpEV->size1);
	_meanVal.resize(tmpEV->size1);
	
	for (size_t iRw = 0; iRw < tmpEV->size1; iRw++) {
		list<size_t>::iterator trtIt = tmpErrInd.begin();
		for (size_t jCl = 0; jCl < tmpEV->size2; jCl++) {
			_errorInvVar[iRw].push_back(gsl_matrix_get(tmpEV, iRw, jCl));
			_meanVal[iRw].push_back(gsl_matrix_get(_valueMat, iRw, *trtIt));
			++trtIt;
		}
	}
	gsl_matrix_free(tmpEV);
	
	// for those lines in the _valueMat that have missing data, eliminate the traits that are missing from the measurement error sampling
	for (vector<size_t>::iterator misIt = _misInd.begin(); misIt != _misInd.end(); ++misIt) {
		
		list<size_t>::iterator trtIt = _missErrMat[*misIt].begin();
		list<double>::iterator varIt = _errorInvVar[*misIt].begin();
		list<double>::iterator mnIt  = _meanVal[*misIt].begin();
		while (trtIt != _missErrMat[*misIt].end()) {
			if (gsl_matrix_int_get(tmpMisMat, *misIt, *trtIt) == 1) {
				trtIt = _missErrMat[*misIt].erase(trtIt);
				varIt = _errorInvVar[*misIt].erase(varIt);
				mnIt  = _meanVal[*misIt].erase(mnIt);
			}
			else {
				++trtIt;
				++varIt;
				++mnIt;
			}
			
		}
	}
	gsl_matrix_int_free(tmpMisMat);
	
	// now ivert all the present variances
	for (vector< list<double> >::iterator rowIt = _errorInvVar.begin(); rowIt != _errorInvVar.end(); ++rowIt) {
		for (list<double>::iterator itemIt = (*rowIt).begin(); itemIt != (*rowIt).end(); ++itemIt) {
			if (*itemIt >= GSL_MACH_EPS) {
				*itemIt = 1.0/(*itemIt);
			}
			else {
				cerr << "ERROR: experimental error variance " << *itemIt << " is too small in MuGrpEEmiss initialization." << endl;
				exit(-1);
			}
			
		}
	}
}
MuGrpEEmiss::MuGrpEEmiss(const string &datFlNam, const string &varFlNam, const vector<size_t> &varInd, const string &misMatFlNam, const string &misVecFlNam, RanIndex &up, const size_t &d) : MuGrpMiss(datFlNam, misMatFlNam, misVecFlNam, up, d) {
	
	gsl_matrix_int *tmpMisMat = gsl_matrix_int_alloc(_valueMat->size1, d);
	FILE *mmIn = fopen(misMatFlNam.c_str(), "r");
	if (mmIn == NULL) {
		cerr << "ERROR: cannot open missing data index file " << misMatFlNam << " in MuGrpMiss" << endl;
		exit(-1);
	}
	gsl_matrix_int_fread(mmIn, tmpMisMat);
	fclose(mmIn);
	
	list<size_t> tmpErrInd(varInd.begin(),varInd.end());
	
	
	for (size_t iDat = 0; iDat < _valueMat->size1; iDat++) {
		_missErrMat.push_back(tmpErrInd);
	}
	
	gsl_matrix *tmpEV = gsl_matrix_alloc(_valueMat->size1, tmpErrInd.size());
	FILE *errIn = fopen(varFlNam.c_str(), "r");
	if (errIn == NULL) {
		cerr << "ERROR: Failed to open file " << varFlNam << " when inititializing MuGrpEEmiss" << endl;
		exit(-1);
	}
	gsl_matrix_fread(errIn, tmpEV);
	fclose(errIn);
	
	_errorInvVar.resize(tmpEV->size1);
	_meanVal.resize(tmpEV->size1);
	
	for (size_t iRw = 0; iRw < tmpEV->size1; iRw++) {
		list<size_t>::iterator trtIt = tmpErrInd.begin();
		for (size_t jCl = 0; jCl < tmpEV->size2; jCl++) {
			_errorInvVar[iRw].push_back(gsl_matrix_get(tmpEV, iRw, jCl));
			_meanVal[iRw].push_back(gsl_matrix_get(_valueMat, iRw, *trtIt));
			++trtIt;
		}
	}
	gsl_matrix_free(tmpEV);
	
	// for those lines in the _valueMat that have missing data, eliminate the traits that are missing from the measurement error sampling
	for (vector<size_t>::iterator misIt = _misInd.begin(); misIt != _misInd.end(); ++misIt) {
		
		list<size_t>::iterator trtIt = _missErrMat[*misIt].begin();
		list<double>::iterator varIt = _errorInvVar[*misIt].begin();
		list<double>::iterator mnIt  = _meanVal[*misIt].begin();
		while (trtIt != _missErrMat[*misIt].end()) {
			if (gsl_matrix_int_get(tmpMisMat, *misIt, *trtIt) == 1) {
				trtIt = _missErrMat[*misIt].erase(trtIt);
				varIt = _errorInvVar[*misIt].erase(varIt);
				mnIt  = _meanVal[*misIt].erase(mnIt);
			}
			else {
				++trtIt;
				++varIt;
				++mnIt;
			}
			
		}
	}
	gsl_matrix_int_free(tmpMisMat);
	// now ivert all the present variances
	for (vector< list<double> >::iterator rowIt = _errorInvVar.begin(); rowIt != _errorInvVar.end(); ++rowIt) {
		for (list<double>::iterator itemIt = (*rowIt).begin(); itemIt != (*rowIt).end(); ++itemIt) {
			if (*itemIt >= GSL_MACH_EPS) {
				*itemIt = 1.0/(*itemIt);
			}
			else {
				cerr << "ERROR: experimental error variance " << *itemIt << " is too small in MuGrpEEmiss initialization." << endl;
				exit(-1);
			}

		}
	}
	
}

MuGrpEEmiss::MuGrpEEmiss(const MuGrpEEmiss &mG){
	_misInd = mG._misInd;
	gsl_matrix_free(_valueMat);
	_valueMat   = gsl_matrix_alloc((mG._valueMat)->size1, (mG._valueMat)->size2);
	gsl_matrix_memcpy(_valueMat, mG._valueMat);
	_errorInvVar = mG._errorInvVar;
	_meanVal     = mG._meanVal;
	
	_upLevel    = mG._upLevel;
	_missErrMat = mG._missErrMat;
	
	_theta.resize(_valueMat->size1);
	size_t mI = 0;
	for (size_t iVrw = 0; iVrw < _valueMat->size1; iVrw++) {
		if (iVrw == _misInd[mI]) {
			if (mI < _misInd.size()) {
				mI++;
			}
			_theta[iVrw] = new MVnormMuMiss(_valueMat, iVrw, *(mG._theta[iVrw])->up(), (mG._theta[iVrw])->getMisPhen());
		}
		else {
			_theta[iVrw] = new MVnormMu(_valueMat, iVrw, *(mG._theta[iVrw])->up());
		}
		
	}
}
MuGrpEEmiss & MuGrpEEmiss::operator=(const MuGrpEEmiss &mG){
	_misInd = mG._misInd;
	gsl_matrix_free(_valueMat);
	_valueMat   = gsl_matrix_alloc((mG._valueMat)->size1, (mG._valueMat)->size2);
	gsl_matrix_memcpy(_valueMat, mG._valueMat);
	_errorInvVar = mG._errorInvVar;
	_meanVal     = mG._meanVal;
	
	_upLevel    = mG._upLevel;
	_missErrMat = mG._missErrMat;
	
	_theta.resize(_valueMat->size1);
	size_t mI = 0;
	for (size_t iVrw = 0; iVrw < _valueMat->size1; iVrw++) {
		if (iVrw == _misInd[mI]) {
			if (mI < _misInd.size()) {
				mI++;
			}
			_theta[iVrw] = new MVnormMuMiss(_valueMat, iVrw, *(mG._theta[iVrw])->up(), (mG._theta[iVrw])->getMisPhen());
		}
		else {
			_theta[iVrw] = new MVnormMu(_valueMat, iVrw, *(mG._theta[iVrw])->up());
		}
		
	}
	
	return *this;
}

// all of these are actually priors
void MuGrpEEmiss::update(const Grp &muPr, const SigmaI &SigIm){
	for (size_t iRw = 0; iRw < _valueMat->size1; iRw++) {
		list<size_t>::iterator indIt = _missErrMat[iRw].begin();
		list<double>::iterator mnIt  = _meanVal[iRw].begin();
		for ( list<double>::iterator colIt = _errorInvVar[iRw].begin(); colIt != _errorInvVar[iRw].end(); ++colIt) {
			double var = 1.0/((*colIt) + gsl_matrix_get(SigIm.getMat(), *indIt, *indIt));
			double mn  = var * ((*mnIt) * (*colIt) +  gsl_matrix_get(SigIm.getMat(), *indIt, *indIt) * gsl_matrix_get(muPr.fMat(), _upLevel->priorInd(iRw), *indIt) );
			gsl_matrix_set(_valueMat, iRw, *indIt, mn + gsl_ran_gaussian_ziggurat(_rV[0], sqrt(var)));
			++indIt;
			++mnIt;
		}
	}
	
	for (vector<size_t>::iterator msIt = _misInd.begin(); msIt != _misInd.end(); ++msIt) {
		_theta[*msIt]->update(muPr, SigIm, _rV[0]);
	}
	
}
void MuGrpEEmiss::update(const Grp &muPr, const Qgrp &q, const SigmaI &SigIm){
	for (size_t iRw = 0; iRw < _valueMat->size1; iRw++) {
		list<size_t>::iterator indIt = _missErrMat[iRw].begin();
		list<double>::iterator mnIt  = _meanVal[iRw].begin();
		for ( list<double>::iterator colIt = _errorInvVar[iRw].begin(); colIt != _errorInvVar[iRw].end(); ++colIt) {
			double var = 1.0/((*colIt) + q[iRw] * gsl_matrix_get(SigIm.getMat(), *indIt, *indIt));
			double mn  = var * ((*mnIt) * (*colIt) +  q[iRw] * gsl_matrix_get(SigIm.getMat(), *indIt, *indIt) * gsl_matrix_get(muPr.fMat(), _upLevel->priorInd(iRw), *indIt) );
			gsl_matrix_set(_valueMat, iRw, *indIt, mn + gsl_ran_gaussian_ziggurat(_rV[0], sqrt(var)));
			++indIt;
			++mnIt;
		}
	}
	
	for (vector<size_t>::iterator msIt = _misInd.begin(); msIt != _misInd.end(); ++msIt) {
		_theta[*msIt]->update(muPr, SigIm, _rV[0]);
	}
	
}


/*
 *	BetaGrpFt methods
 */
BetaGrpFt::BetaGrpFt() : _numSaves(0.0), Grp() {
	_fittedAll  = gsl_matrix_calloc(1, 1);
	_Xmat       = gsl_matrix_calloc(1, 1);
}

BetaGrpFt::BetaGrpFt(const Grp &rsp, const string &predFlNam, const size_t &Npred, const int &nThr) : _numSaves(0.0), _nThr(nThr), Grp(){
	_fittedEach.resize(Npred);
	_fittedAll  = gsl_matrix_calloc(rsp.Ndata(), rsp.phenD());
	_Xmat       = gsl_matrix_alloc(rsp.Ndata(), Npred);
	gsl_matrix_free(_valueMat);
	_valueMat   = gsl_matrix_alloc(Npred, rsp.phenD());
	
	if (_nThr > 1) {
		const gsl_rng_type *T = gsl_rng_mt19937;
		_rV.resize(_nThr);
		vector<gsl_rng *>::iterator rIt = _rV.begin();
		++rIt;
		for (; rIt != _rV.end(); ++rIt) {
			unsigned long seed = gsl_rng_get(_rV[0]);
			*rIt = gsl_rng_alloc(T);
			gsl_rng_set(*rIt, seed);
		}

	}
	
	gsl_matrix *Sig  = gsl_matrix_alloc(rsp.phenD(), rsp.phenD());
	gsl_matrix_set_identity(Sig);
	
	_theta.resize(Npred);
	
	FILE *prdIn = fopen(predFlNam.c_str(), "r");
	if (prdIn == NULL) {
		cerr << "ERROR: cannot open file " << predFlNam << " in BetaGrpFt" << endl;
		exit(-1);
	}
	gsl_matrix_fread(prdIn, _Xmat);
	fclose(prdIn);
	
	gsl_matrix *y = gsl_matrix_alloc(rsp.dMat()->size1, rsp.dMat()->size2);
	colCenter(rsp.dMat(), y); // approximating a regression with an intercept for initial values (not quite right, but close enough for initialization)
	
	if (Npred == 1) {
		_theta[0] = new MVnormBeta(y, _Xmat, 0, Sig, _rV[0], _valueMat, 0);
	}
	else {
		for (size_t Xj = 0; Xj < Npred; Xj++) {
			_fittedEach[Xj].resize(rsp.Ndata()*rsp.phenD());
			_theta[Xj] = new MVnormBetaFt(y, _Xmat, Xj, _fittedEach[Xj], Sig, _rV[0], _valueMat, Xj);
		}
	}
	
	_updateFitted();
	gsl_matrix_free(Sig);
	gsl_matrix_free(y);
}

BetaGrpFt::BetaGrpFt(const Grp &rsp, const string &predFlNam, const size_t &Npred, RanIndex &up, const int &nThr) : _numSaves(0.0), _nThr(nThr), Grp(){
	if (Npred != up.getNtot()) {
		cerr << "ERROR: number of predictors " << Npred << " is not equal to the number of elements " << up.getNtot() << " in the prior index in BetaGrpFt initialization." << endl;
		exit(-1);
	}
	
	_fittedEach.resize(Npred);
	_fittedAll  = gsl_matrix_calloc(rsp.Ndata(), rsp.phenD());
	_Xmat       = gsl_matrix_alloc(rsp.Ndata(), Npred);
	gsl_matrix_free(_valueMat);
	_valueMat   = gsl_matrix_alloc(Npred, rsp.phenD());
	
	_upLevel = &up;
	

	if (_nThr > 1) {
		const gsl_rng_type *T = gsl_rng_mt19937;
		_rV.resize(_nThr);
		vector<gsl_rng *>::iterator rIt = _rV.begin();
		++rIt;
		for (; rIt != _rV.end(); ++rIt) {
			unsigned long seed = gsl_rng_get(_rV[0]);
			*rIt = gsl_rng_alloc(T);
			gsl_rng_set(*rIt, seed);
		}
		
	}
	
	gsl_matrix *Sig  = gsl_matrix_alloc(rsp.phenD(), rsp.phenD());
	gsl_matrix_set_identity(Sig);
	
	_theta.resize(Npred);
	
	FILE *prdIn = fopen(predFlNam.c_str(), "r");
	if (prdIn == NULL) {
		cerr << "ERROR: cannot open file " << predFlNam << " in BetaGrpFt" << endl;
		exit(-1);
	}
	gsl_matrix_fread(prdIn, _Xmat);
	fclose(prdIn);
	
	gsl_matrix *y = gsl_matrix_alloc(rsp.dMat()->size1, rsp.dMat()->size2);
	colCenter(rsp.dMat(), y); // approximating a regression with an intercept for initial values (not quite right, but close enough for initialization)
	
	if (Npred == 1) {
		_theta[0] = new MVnormBeta(y, _Xmat, 0, Sig, _rV[0], _upLevel->priorInd(0), _valueMat, 0);
	}
	else {
		for (size_t Xj = 0; Xj < Npred; Xj++) {
			_fittedEach[Xj].resize(rsp.Ndata()*rsp.phenD());
			_theta[Xj] = new MVnormBetaFt(y, _Xmat, Xj, _fittedEach[Xj], Sig, _rV[0], _upLevel->priorInd(Xj), _valueMat, Xj);
		}

	}
	
	_updateFitted();
	gsl_matrix_free(Sig);
	gsl_matrix_free(y);
}

BetaGrpFt::BetaGrpFt(const Grp &rsp, const string &predFlNam, const size_t &Npred, RanIndex &low, RanIndex &up, const int &nThr) : _numSaves(0.0), _nThr(nThr), Grp(){
	//	the _lowLevel RanIndex plays the role of the design matrix, and relates the unique rows of the predictor (low.getNgrp()xNpred) to the predictor for all the rsp rows
	//  the _fittedAll matrix still has only the unique rows; this is for efficiency when adding it to other Grp classes
	
	if (Npred != up.getNtot()) {
		cerr << "ERROR: number of predictors " << Npred << " is not equal to the number of elements " << up.getNtot() << " in the prior index in BetaGrpFt initialization." << endl;
		exit(-1);
	}
	else if (low.getNtot() != rsp.dMat()->size1){
		cerr << "ERROR: number of elements " << low.getNgrp() << " in the replication index is not equal to the number of rows " << rsp.dMat()->size1 << " in the response matrix in BetaGrpFt initialization." << endl;
		exit(-1);
	}
	
	_fittedEach.resize(Npred);
	_fittedAll  = gsl_matrix_calloc(low.getNgrp(), rsp.phenD());
	_Xmat       = gsl_matrix_alloc(low.getNtot(), Npred);
	gsl_matrix_free(_valueMat);
	_valueMat   = gsl_matrix_alloc(Npred, rsp.phenD());
	_lowLevel   = &low;
	_upLevel    = &up;
	
	if (_nThr > 1) {
		const gsl_rng_type *T = gsl_rng_mt19937;
		_rV.resize(_nThr);
		vector<gsl_rng *>::iterator rIt = _rV.begin();
		++rIt;
		for (; rIt != _rV.end(); ++rIt) {
			unsigned long seed = gsl_rng_get(_rV[0]);
			*rIt = gsl_rng_alloc(T);
			gsl_rng_set(*rIt, seed);
		}
		
	}
	
	gsl_matrix *Sig  = gsl_matrix_alloc(rsp.phenD(), rsp.phenD());
	gsl_matrix_set_identity(Sig);
	
	_theta.resize(Npred);
	
	/*
	 *  will read one line at a time so as not to have two potentially big matrices in memory at the same time; taking advantage of the row-major arrangement
	 */
	
	FILE *prdIn = fopen(predFlNam.c_str(), "r");
	if (prdIn == NULL) {
		cerr << "ERROR: cannot open file " << predFlNam << " in BetaGrpFt" << endl;
		exit(-1);
	}
	gsl_vector *row = gsl_vector_alloc(Npred);
	for (size_t iGrp = 0; iGrp < _lowLevel->getNgrp(); iGrp++) {
		gsl_vector_fread(prdIn, row);
		for (vector<size_t>::iterator grpIt = (*_lowLevel)[iGrp].begin(); grpIt != (*_lowLevel)[iGrp].end(); ++grpIt) {
			gsl_matrix_set_row(_Xmat, *grpIt, row);
		}
		
	}
	gsl_vector_free(row);
	fclose(prdIn);
	
	gsl_matrix *y = gsl_matrix_alloc(rsp.dMat()->size1, rsp.dMat()->size2);
	colCenter(rsp.dMat(), y); // approximating a regression with an intercept for initial values (not quite right, but close enough for initialization)
	
	if (Npred == 1) {
		_theta[0] = new MVnormBeta(y, _Xmat, 0, Sig, _rV[0], _upLevel->priorInd(0), _valueMat, 0);
	}
	else {
		for (size_t Xj = 0; Xj < Npred; Xj++) {
			_fittedEach[Xj].resize(low.getNtot()*rsp.phenD());
			_theta[Xj] = new MVnormBetaFt(y, _Xmat, Xj, _fittedEach[Xj], Sig, _rV[0], _upLevel->priorInd(Xj), _valueMat, Xj);
		}
	}
	
	_updateFitted();
	gsl_matrix_free(Sig);
	gsl_matrix_free(y);
}

BetaGrpFt::BetaGrpFt(const Grp &rsp, const string &predFlNam, const size_t &Npred, const string &outFlNam, const int &nThr) : _numSaves(0.0), _nThr(nThr), Grp(){
	_fittedEach.resize(Npred);
	_fittedAll  = gsl_matrix_calloc(rsp.Ndata(), rsp.phenD());
	_Xmat       = gsl_matrix_alloc(rsp.Ndata(), Npred);
	gsl_matrix_free(_valueMat);
	_valueMat   = gsl_matrix_alloc(Npred, rsp.phenD());
	_outFlNam   = outFlNam;
	remove(_outFlNam.c_str());
	
	if (_nThr > 1) {
		const gsl_rng_type *T = gsl_rng_mt19937;
		_rV.resize(_nThr);
		vector<gsl_rng *>::iterator rIt = _rV.begin();
		++rIt;
		for (; rIt != _rV.end(); ++rIt) {
			unsigned long seed = gsl_rng_get(_rV[0]);
			*rIt = gsl_rng_alloc(T);
			gsl_rng_set(*rIt, seed);
		}
		
	}
	
	gsl_matrix *Sig  = gsl_matrix_alloc(rsp.phenD(), rsp.phenD());
	gsl_matrix_set_identity(Sig);
	
	_theta.resize(Npred);
	
	FILE *prdIn = fopen(predFlNam.c_str(), "r");
	if (prdIn == NULL) {
		cerr << "ERROR: cannot open file " << predFlNam << " in BetaGrpFt" << endl;
		exit(-1);
	}
	gsl_matrix_fread(prdIn, _Xmat);
	fclose(prdIn);
	
	gsl_matrix *y = gsl_matrix_alloc(rsp.dMat()->size1, rsp.dMat()->size2);
	colCenter(rsp.dMat(), y); // approximating a regression with an intercept for initial values (not quite right, but close enough for initialization)
	
	if (Npred == 1) {
		_theta[0] = new MVnormBeta(y, _Xmat, 0, Sig, _rV[0], _valueMat, 0);
	}
	else {
		for (size_t Xj = 0; Xj < Npred; Xj++) {
			_fittedEach[Xj].resize(rsp.Ndata()*rsp.phenD());
			_theta[Xj] = new MVnormBetaFt(y, _Xmat, Xj, _fittedEach[Xj], Sig, _rV[0], _valueMat, Xj);
		}

	}
	
	_updateFitted();
	gsl_matrix_free(Sig);
	gsl_matrix_free(y);
}

BetaGrpFt::BetaGrpFt(const Grp &rsp, const string &predFlNam, const size_t &Npred, RanIndex &up, const string &outFlNam, const int &nThr) : _numSaves(0.0), _nThr(nThr), Grp(){
	if (Npred != up.getNtot()) {
		cerr << "ERROR: number of predictors " << Npred << " is not equal to the number of elements " << up.getNtot() << " in the prior index in BetaGrpFt initialization." << endl;
		exit(-1);
	}

	_fittedEach.resize(Npred);
	_fittedAll  = gsl_matrix_calloc(rsp.Ndata(), rsp.phenD());
	_Xmat       = gsl_matrix_alloc(rsp.Ndata(), Npred);
	gsl_matrix_free(_valueMat);
	_valueMat   = gsl_matrix_alloc(Npred, rsp.phenD());
	
	_upLevel  = &up;
	_outFlNam = outFlNam;
	remove(_outFlNam.c_str());
	
	if (_nThr > 1) {
		const gsl_rng_type *T = gsl_rng_mt19937;
		_rV.resize(_nThr);
		vector<gsl_rng *>::iterator rIt = _rV.begin();
		++rIt;
		for (; rIt != _rV.end(); ++rIt) {
			unsigned long seed = gsl_rng_get(_rV[0]);
			*rIt = gsl_rng_alloc(T);
			gsl_rng_set(*rIt, seed);
		}
		
	}
	
	gsl_matrix *Sig  = gsl_matrix_alloc(rsp.phenD(), rsp.phenD());
	gsl_matrix_set_identity(Sig);
	
	_theta.resize(Npred);
	
	FILE *prdIn = fopen(predFlNam.c_str(), "r");
	if (prdIn == NULL) {
		cerr << "ERROR: cannot open file " << predFlNam << " in BetaGrpFt" << endl;
		exit(-1);
	}
	gsl_matrix_fread(prdIn, _Xmat);
	fclose(prdIn);
	
	gsl_matrix *y = gsl_matrix_alloc(rsp.dMat()->size1, rsp.dMat()->size2);
	colCenter(rsp.dMat(), y); // approximating a regression with an intercept for initial values (not quite right, but close enough for initialization)
	
	if (Npred == 1) {
		_theta[0] = new MVnormBeta(y, _Xmat, 0, Sig, _rV[0], _upLevel->priorInd(0), _valueMat, 0);
	}
	else {
		for (size_t Xj = 0; Xj < Npred; Xj++) {
			_fittedEach[Xj].resize(rsp.Ndata()*rsp.phenD());
			_theta[Xj] = new MVnormBetaFt(y, _Xmat, Xj, _fittedEach[Xj], Sig, _rV[0], _upLevel->priorInd(Xj), _valueMat, Xj);
		}
	}
	
	_updateFitted();
	gsl_matrix_free(Sig);
	gsl_matrix_free(y);
}

BetaGrpFt::BetaGrpFt(const Grp &rsp, const string &predFlNam, const size_t &Npred, RanIndex &low, RanIndex &up, const string &outFlNam, const int &nThr) : _numSaves(0.0), _nThr(nThr), Grp(){
	//	the _lowLevel RanIndex plays the role of the design matrix, and relates the unique values of the predictor (low.getNgrp()x1) to the predictor for all the rsp rows
	if (Npred != up.getNtot()) {
		cerr << "ERROR: number of predictors " << Npred << " is not equal to the number of elements " << up.getNtot() << " in the prior index in BetaGrpFt initialization." << endl;
		exit(-1);
	}
	else if (low.getNtot() != rsp.dMat()->size1){
		cerr << "ERROR: number of elements " << low.getNgrp() << " in the replication index is not equal to the number of rows " << rsp.dMat()->size1 << " in the response matrix in BetaGrpFt initialization." << endl;
		exit(-1);
	}
	
	_fittedEach.resize(Npred);
	_fittedAll  = gsl_matrix_calloc(low.getNgrp(), rsp.phenD());
	_Xmat       = gsl_matrix_alloc(low.getNtot(), Npred);
	gsl_matrix_free(_valueMat);
	_valueMat   = gsl_matrix_alloc(Npred, rsp.phenD());
	_lowLevel   = &low;
	_upLevel    = &up;
	_outFlNam   = outFlNam;
	remove(_outFlNam.c_str());
	
	if (_nThr > 1) {
		const gsl_rng_type *T = gsl_rng_mt19937;
		_rV.resize(_nThr);
		vector<gsl_rng *>::iterator rIt = _rV.begin();
		++rIt;
		for (; rIt != _rV.end(); ++rIt) {
			unsigned long seed = gsl_rng_get(_rV[0]);
			*rIt = gsl_rng_alloc(T);
			gsl_rng_set(*rIt, seed);
		}
		
	}
	
	gsl_matrix *Sig  = gsl_matrix_alloc(rsp.phenD(), rsp.phenD());
	gsl_matrix_set_identity(Sig);
	
	_theta.resize(Npred);
	
	/*
	 *  will read on line at a time so as not to have two potentially big matrices in memory at the same time; taking advantage of the row-major arrangement
	 */
	
	FILE *prdIn = fopen(predFlNam.c_str(), "r");
	if (prdIn == NULL) {
		cerr << "ERROR: cannot open file " << predFlNam << " in BetaGrpFt" << endl;
		exit(-1);
	}
	gsl_vector *row = gsl_vector_alloc(Npred);
	for (size_t iGrp = 0; iGrp < _lowLevel->getNgrp(); iGrp++) {
		gsl_vector_fread(prdIn, row);
		for (vector<size_t>::iterator grpIt = (*_lowLevel)[iGrp].begin(); grpIt != (*_lowLevel)[iGrp].end(); ++grpIt) {
			gsl_matrix_set_row(_Xmat, *grpIt, row);
		}
		
	}
	gsl_vector_free(row);
	fclose(prdIn);
	
	gsl_matrix *y = gsl_matrix_alloc(rsp.dMat()->size1, rsp.dMat()->size2);
	colCenter(rsp.dMat(), y); // approximating a regression with an intercept for initial values (not quite right, but close enough for initialization)
	
	if (Npred == 1) {
		_theta[0] = new MVnormBeta(y, _Xmat, 0, Sig, _rV[0], _upLevel->priorInd(0), _valueMat, 0);
	}
	else {
		for (size_t Xj = 0; Xj < Npred; Xj++) {
			_fittedEach[Xj].resize(low.getNtot()*rsp.phenD());
			_theta[Xj] = new MVnormBetaFt(y, _Xmat, Xj, _fittedEach[Xj], Sig, _rV[0], _upLevel->priorInd(Xj), _valueMat, Xj);
		}
	}
	
	_updateFitted();
	gsl_matrix_free(Sig);
	gsl_matrix_free(y);
}

/*
 *	constructors with missing predictor data
 */
BetaGrpFt::BetaGrpFt(const Grp &rsp, const string &predFlNam, const size_t &Npred, const double &absLab, const int &nThr) :  _numSaves(0.0), _nThr(nThr), Grp() {
	_fittedEach.resize(Npred);
	_fittedAll  = gsl_matrix_calloc(rsp.Ndata(), rsp.phenD());
	_Xmat       = gsl_matrix_alloc(rsp.Ndata(), Npred);
	gsl_matrix_free(_valueMat);
	_valueMat   = gsl_matrix_alloc(Npred, rsp.phenD());
	
	if (_nThr > 1) {
		const gsl_rng_type *T = gsl_rng_mt19937;
		_rV.resize(_nThr);
		vector<gsl_rng *>::iterator rIt = _rV.begin();
		++rIt;
		for (; rIt != _rV.end(); ++rIt) {
			unsigned long seed = gsl_rng_get(_rV[0]);
			*rIt = gsl_rng_alloc(T);
			gsl_rng_set(*rIt, seed);
		}
		
	}
	
	gsl_matrix *Sig  = gsl_matrix_alloc(rsp.phenD(), rsp.phenD());
	gsl_matrix_set_identity(Sig);
	
	_theta.resize(Npred);
	
	FILE *prdIn = fopen(predFlNam.c_str(), "r");
	if (prdIn == NULL) {
		cerr << "ERROR: cannot open file " << predFlNam << " in BetaGrpFt" << endl;
		exit(-1);
	}
	gsl_matrix_fread(prdIn, _Xmat);
	fclose(prdIn);
	
	meanImpute(_Xmat, absLab);
	gsl_matrix *y = gsl_matrix_alloc(rsp.dMat()->size1, rsp.dMat()->size2);
	colCenter(rsp.dMat(), y); // approximating a regression with an intercept for initial values (not quite right, but close enough for initialization)
	
	if (Npred == 1) {
		_theta[0] = new MVnormBeta(y, _Xmat, 0, Sig, _rV[0], _valueMat, 0);
	}
	else {
		for (size_t Xj = 0; Xj < Npred; Xj++) {
			_fittedEach[Xj].resize(rsp.Ndata()*rsp.phenD());
			_theta[Xj] = new MVnormBetaFt(y, _Xmat, Xj, _fittedEach[Xj], Sig, _rV[0], _valueMat, Xj);
		}
	}
	
	_updateFitted();
	gsl_matrix_free(Sig);
	gsl_matrix_free(y);
}
BetaGrpFt::BetaGrpFt(const Grp &rsp, const string &predFlNam, const size_t &Npred, const double &absLab, RanIndex &up, const int &nThr) :  _numSaves(0.0), _nThr(nThr), Grp() {
	if (Npred != up.getNtot()) {
		cerr << "ERROR: number of predictors " << Npred << " is not equal to the number of elements " << up.getNtot() << " in the prior index in BetaGrpFt initialization." << endl;
		exit(-1);
	}
	
	_fittedEach.resize(Npred);
	_fittedAll  = gsl_matrix_calloc(rsp.Ndata(), rsp.phenD());
	_Xmat       = gsl_matrix_alloc(rsp.Ndata(), Npred);
	gsl_matrix_free(_valueMat);
	_valueMat   = gsl_matrix_alloc(Npred, rsp.phenD());
	
	_upLevel = &up;
	
	if (_nThr > 1) {
		const gsl_rng_type *T = gsl_rng_mt19937;
		_rV.resize(_nThr);
		vector<gsl_rng *>::iterator rIt = _rV.begin();
		++rIt;
		for (; rIt != _rV.end(); ++rIt) {
			unsigned long seed = gsl_rng_get(_rV[0]);
			*rIt = gsl_rng_alloc(T);
			gsl_rng_set(*rIt, seed);
		}
		
	}
	
	gsl_matrix *Sig  = gsl_matrix_alloc(rsp.phenD(), rsp.phenD());
	gsl_matrix_set_identity(Sig);
	
	_theta.resize(Npred);
	
	FILE *prdIn = fopen(predFlNam.c_str(), "r");
	if (prdIn == NULL) {
		cerr << "ERROR: cannot open file " << predFlNam << " in BetaGrpFt" << endl;
		exit(-1);
	}
	gsl_matrix_fread(prdIn, _Xmat);
	fclose(prdIn);
	
	meanImpute(_Xmat, absLab);
	gsl_matrix *y = gsl_matrix_alloc(rsp.dMat()->size1, rsp.dMat()->size2);
	colCenter(rsp.dMat(), y); // approximating a regression with an intercept for initial values (not quite right, but close enough for initialization)
	
	if (Npred == 1) {
		_theta[0] = new MVnormBeta(y, _Xmat, 0, Sig, _rV[0], _upLevel->priorInd(0), _valueMat, 0);
	}
	else {
		for (size_t Xj = 0; Xj < Npred; Xj++) {
			_fittedEach[Xj].resize(rsp.Ndata()*rsp.phenD());
			_theta[Xj] = new MVnormBetaFt(y, _Xmat, Xj, _fittedEach[Xj], Sig, _rV[0], _upLevel->priorInd(Xj), _valueMat, Xj);
		}
	}
	
	_updateFitted();
	gsl_matrix_free(Sig);
	gsl_matrix_free(y);
}
BetaGrpFt::BetaGrpFt(const Grp &rsp, const string &predFlNam, const size_t &Npred, const double &absLab, RanIndex &low, RanIndex &up, const int &nThr) :  _numSaves(0.0), _nThr(nThr), Grp() {
	if (Npred != up.getNtot()) {
		cerr << "ERROR: number of predictors " << Npred << " is not equal to the number of elements " << up.getNtot() << " in the prior index in BetaGrpFt initialization." << endl;
		exit(-1);
	}
	else if (low.getNtot() != rsp.dMat()->size1){
		cerr << "ERROR: number of elements " << low.getNgrp() << " in the replication index is not equal to the number of rows " << rsp.dMat()->size1 << " in the response matrix in BetaGrpFt initialization." << endl;
		exit(-1);
	}

	_fittedEach.resize(Npred);
	_fittedAll  = gsl_matrix_calloc(low.getNgrp(), rsp.phenD());
	_Xmat       = gsl_matrix_alloc(low.getNtot(), Npred);
	gsl_matrix_free(_valueMat);
	_valueMat   = gsl_matrix_alloc(Npred, rsp.phenD());
	_lowLevel   = &low;
	_upLevel    = &up;
	
	if (_nThr > 1) {
		const gsl_rng_type *T = gsl_rng_mt19937;
		_rV.resize(_nThr);
		vector<gsl_rng *>::iterator rIt = _rV.begin();
		++rIt;
		for (; rIt != _rV.end(); ++rIt) {
			unsigned long seed = gsl_rng_get(_rV[0]);
			*rIt = gsl_rng_alloc(T);
			gsl_rng_set(*rIt, seed);
		}
		
	}
	
	gsl_matrix *Sig  = gsl_matrix_alloc(rsp.phenD(), rsp.phenD());
	gsl_matrix_set_identity(Sig);
	
	_theta.resize(Npred);
	
	gsl_matrix *tmpX = gsl_matrix_alloc(_lowLevel->getNgrp(), Npred);
	FILE *prdIn = fopen(predFlNam.c_str(), "r");
	if (prdIn == NULL) {
		cerr << "ERROR: cannot open file " << predFlNam << " in BetaGrpFt" << endl;
		exit(-1);
	}
	gsl_matrix_fread(prdIn, tmpX);
	fclose(prdIn);
	meanImpute(tmpX, absLab);
	
	for (size_t iGrp = 0; iGrp < _lowLevel->getNgrp(); iGrp++) {
		gsl_vector_view tmpRow = gsl_matrix_row(tmpX, iGrp);
		for (vector<size_t>::iterator grpIt = (*_lowLevel)[iGrp].begin(); grpIt != (*_lowLevel)[iGrp].end(); ++grpIt) {
			gsl_matrix_set_row(_Xmat, *grpIt, &tmpRow.vector);
		}
		
	}
	gsl_matrix_free(tmpX);
	
	gsl_matrix *y = gsl_matrix_alloc(rsp.dMat()->size1, rsp.dMat()->size2);
	colCenter(rsp.dMat(), y); // approximating a regression with an intercept for initial values (not quite right, but close enough for initialization)
	
	if (Npred == 1) {
		_theta[0] = new MVnormBeta(y, _Xmat, 0, Sig, _rV[0], _upLevel->priorInd(0), _valueMat, 0);
	}
	else {
		for (size_t Xj = 0; Xj < Npred; Xj++) {
			_fittedEach[Xj].resize(low.getNtot()*rsp.phenD());
			_theta[Xj] = new MVnormBetaFt(y, _Xmat, Xj, _fittedEach[Xj], Sig, _rV[0], _upLevel->priorInd(Xj), _valueMat, Xj);
		}
	}
	
	_updateFitted();
	gsl_matrix_free(Sig);
	gsl_matrix_free(y);
}
BetaGrpFt::BetaGrpFt(const Grp &rsp, const string &predFlNam, const size_t &Npred, const double &absLab, const string &outFlNam, const int &nThr) :  _numSaves(0.0), _nThr(nThr), Grp(){
	_fittedEach.resize(Npred);
	_fittedAll  = gsl_matrix_calloc(rsp.Ndata(), rsp.phenD());
	_Xmat       = gsl_matrix_alloc(rsp.Ndata(), Npred);
	gsl_matrix_free(_valueMat);
	_valueMat   = gsl_matrix_alloc(Npred, rsp.phenD());
	_outFlNam   = outFlNam;
	remove(_outFlNam.c_str());
	
	if (_nThr > 1) {
		const gsl_rng_type *T = gsl_rng_mt19937;
		_rV.resize(_nThr);
		vector<gsl_rng *>::iterator rIt = _rV.begin();
		++rIt;
		for (; rIt != _rV.end(); ++rIt) {
			unsigned long seed = gsl_rng_get(_rV[0]);
			*rIt = gsl_rng_alloc(T);
			gsl_rng_set(*rIt, seed);
		}
		
	}
	
	gsl_matrix *Sig  = gsl_matrix_alloc(rsp.phenD(), rsp.phenD());
	gsl_matrix_set_identity(Sig);
	
	_theta.resize(Npred);
	
	FILE *prdIn = fopen(predFlNam.c_str(), "r");
	if (prdIn == NULL) {
		cerr << "ERROR: cannot open file " << predFlNam << " in BetaGrpFt" << endl;
		exit(-1);
	}
	gsl_matrix_fread(prdIn, _Xmat);
	fclose(prdIn);
	
	meanImpute(_Xmat, absLab);
	gsl_matrix *y = gsl_matrix_alloc(rsp.dMat()->size1, rsp.dMat()->size2);
	colCenter(rsp.dMat(), y); // approximating a regression with an intercept for initial values (not quite right, but close enough for initialization)
	
	if (Npred == 1) {
		_theta[0] = new MVnormBeta(y, _Xmat, 0, Sig, _rV[0], _valueMat, 0);
	}
	else {
		for (size_t Xj = 0; Xj < Npred; Xj++) {
			_fittedEach[Xj].resize(rsp.Ndata()*rsp.phenD());
			_theta[Xj] = new MVnormBetaFt(y, _Xmat, Xj, _fittedEach[Xj], Sig, _rV[0], _valueMat, Xj);
		}
	}
	
	_updateFitted();
	gsl_matrix_free(Sig);
	gsl_matrix_free(y);

}
BetaGrpFt::BetaGrpFt(const Grp &rsp, const string &predFlNam, const size_t &Npred, const double &absLab, RanIndex &up, const string &outFlNam, const int &nThr) :  _numSaves(0.0), _nThr(nThr), Grp() {
	if (Npred != up.getNtot()) {
		cerr << "ERROR: number of predictors " << Npred << " is not equal to the number of elements " << up.getNtot() << " in the prior index in BetaGrpFt initialization." << endl;
		exit(-1);
	}

	_fittedEach.resize(Npred);
	_fittedAll  = gsl_matrix_calloc(rsp.Ndata(), rsp.phenD());
	_Xmat       = gsl_matrix_alloc(rsp.Ndata(), Npred);
	gsl_matrix_free(_valueMat);
	_valueMat   = gsl_matrix_alloc(Npred, rsp.phenD());
	
	_upLevel  = &up;
	_outFlNam = outFlNam;
	remove(_outFlNam.c_str());
	
	if (_nThr > 1) {
		const gsl_rng_type *T = gsl_rng_mt19937;
		_rV.resize(_nThr);
		vector<gsl_rng *>::iterator rIt = _rV.begin();
		++rIt;
		for (; rIt != _rV.end(); ++rIt) {
			unsigned long seed = gsl_rng_get(_rV[0]);
			*rIt = gsl_rng_alloc(T);
			gsl_rng_set(*rIt, seed);
		}
		
	}
	
	gsl_matrix *Sig  = gsl_matrix_alloc(rsp.phenD(), rsp.phenD());
	gsl_matrix_set_identity(Sig);
	
	_theta.resize(Npred);
	
	FILE *prdIn = fopen(predFlNam.c_str(), "r");
	if (prdIn == NULL) {
		cerr << "ERROR: cannot open file " << predFlNam << " in BetaGrpFt" << endl;
		exit(-1);
	}
	gsl_matrix_fread(prdIn, _Xmat);
	fclose(prdIn);
	
	meanImpute(_Xmat, absLab);
	gsl_matrix *y = gsl_matrix_alloc(rsp.dMat()->size1, rsp.dMat()->size2);
	colCenter(rsp.dMat(), y); // approximating a regression with an intercept for initial values (not quite right, but close enough for initialization)
	
	if (Npred == 1) {
		_theta[0] = new MVnormBeta(y, _Xmat, 0, Sig, _rV[0], _upLevel->priorInd(0), _valueMat, 0);
	}
	else {
		for (size_t Xj = 0; Xj < Npred; Xj++) {
			_fittedEach[Xj].resize(rsp.Ndata()*rsp.phenD());
			_theta[Xj] = new MVnormBetaFt(y, _Xmat, Xj, _fittedEach[Xj], Sig, _rV[0], _upLevel->priorInd(Xj), _valueMat, Xj);
		}
	}
	
	_updateFitted();
	gsl_matrix_free(Sig);
	gsl_matrix_free(y);
}
BetaGrpFt::BetaGrpFt(const Grp &rsp, const string &predFlNam, const size_t &Npred, const double &absLab, RanIndex &low, RanIndex &up, const string &outFlNam, const int &nThr) :  _numSaves(0.0), _nThr(nThr), Grp() {
	if (Npred != up.getNtot()) {
		cerr << "ERROR: number of predictors " << Npred << " is not equal to the number of elements " << up.getNtot() << " in the prior index in BetaGrpFt initialization." << endl;
		exit(-1);
	}
	else if (low.getNtot() != rsp.dMat()->size1){
		cerr << "ERROR: number of elements " << low.getNgrp() << " in the replication index is not equal to the number of rows " << rsp.dMat()->size1 << " in the response matrix in BetaGrpFt initialization." << endl;
		exit(-1);
	}

	_fittedEach.resize(Npred);
	_fittedAll  = gsl_matrix_calloc(low.getNgrp(), rsp.phenD());
	_Xmat       = gsl_matrix_alloc(low.getNtot(), Npred);
	gsl_matrix_free(_valueMat);
	_valueMat   = gsl_matrix_alloc(Npred, rsp.phenD());
	_lowLevel   = &low;
	_upLevel    = &up;
	_outFlNam   = outFlNam;
	remove(_outFlNam.c_str());
	
	if (_nThr > 1) {
		const gsl_rng_type *T = gsl_rng_mt19937;
		_rV.resize(_nThr);
		vector<gsl_rng *>::iterator rIt = _rV.begin();
		++rIt;
		for (; rIt != _rV.end(); ++rIt) {
			unsigned long seed = gsl_rng_get(_rV[0]);
			*rIt = gsl_rng_alloc(T);
			gsl_rng_set(*rIt, seed);
		}
		
	}
	
	gsl_matrix *Sig  = gsl_matrix_alloc(rsp.phenD(), rsp.phenD());
	gsl_matrix_set_identity(Sig);
	
	_theta.resize(Npred);
	
	gsl_matrix *tmpX = gsl_matrix_alloc(_lowLevel->getNgrp(), Npred);
	FILE *prdIn = fopen(predFlNam.c_str(), "r");
	if (prdIn == NULL) {
		cerr << "ERROR: cannot open file " << predFlNam << " in BetaGrpFt" << endl;
		exit(-1);
	}
	gsl_matrix_fread(prdIn, tmpX);
	fclose(prdIn);
	meanImpute(tmpX, absLab);
	
	for (size_t iGrp = 0; iGrp < _lowLevel->getNgrp(); iGrp++) {
		gsl_vector_view tmpRow = gsl_matrix_row(tmpX, iGrp);
		for (vector<size_t>::iterator grpIt = (*_lowLevel)[iGrp].begin(); grpIt != (*_lowLevel)[iGrp].end(); ++grpIt) {
			gsl_matrix_set_row(_Xmat, *grpIt, &tmpRow.vector);
		}
		
	}
	gsl_matrix_free(tmpX);
	
	gsl_matrix *y = gsl_matrix_alloc(rsp.dMat()->size1, rsp.dMat()->size2);
	colCenter(rsp.dMat(), y);
	
	if (Npred == 1) {
		_theta[0] = new MVnormBeta(y, _Xmat, 0, Sig, _rV[0], _upLevel->priorInd(0), _valueMat, 0);
	}
	else {
		for (size_t Xj = 0; Xj < Npred; Xj++) {
			_fittedEach[Xj].resize(low.getNtot()*rsp.phenD());
			_theta[Xj] = new MVnormBetaFt(y, _Xmat, Xj, _fittedEach[Xj], Sig, _rV[0], _upLevel->priorInd(Xj), _valueMat, Xj);
		}
	}
	
	_updateFitted();
	gsl_matrix_free(Sig);
	gsl_matrix_free(y);
}

/*
 *	constructors with rank pre-selection
 */
BetaGrpFt::BetaGrpFt(const Grp &rsp, const SigmaI &SigI, const string &predFlNam, const size_t &Npred, const double &Nmul, const double &rSqMax, RanIndex &up, const string &outFlNam, const int &nThr) : _numSaves(0.0), _nThr(nThr), Grp(){
	if (Npred != up.getNtot()) {
		cerr << "ERROR: number of predictors " << Npred << " is not equal to the number of elements " << up.getNtot() << " in the prior index in BetaGrpFt initialization." << endl;
		exit(-1);
	}

	gsl_matrix_free(_valueMat);
	_fittedAll = gsl_matrix_calloc(rsp.Ndata(), rsp.phenD());
	_Xmat      = gsl_matrix_alloc(rsp.Ndata(), Npred);
	
	_upLevel  = &up;
	_outFlNam = outFlNam;
	remove(_outFlNam.c_str());
	
	if (_nThr > 1) {
		const gsl_rng_type *T = gsl_rng_mt19937;
		_rV.resize(_nThr);
		vector<gsl_rng *>::iterator rIt = _rV.begin();
		++rIt;
		for (; rIt != _rV.end(); ++rIt) {
			unsigned long seed = gsl_rng_get(_rV[0]);
			*rIt = gsl_rng_alloc(T);
			gsl_rng_set(*rIt, seed);
		}
		
	}
	
	FILE *prdIn = fopen(predFlNam.c_str(), "r");
	if (prdIn == NULL) {
		cerr << "ERROR: cannot open file " << predFlNam << " in BetaGrpFt" << endl;
		exit(-1);
	}
	gsl_matrix_fread(prdIn, _Xmat);
	fclose(prdIn);
	
	gsl_matrix *y = gsl_matrix_alloc(rsp.dMat()->size1, rsp.dMat()->size2);
	colCenter(rsp.dMat(), y); // approximating a regression with an intercept for initial values (not quite right, but close enough for initialization)
	
	gsl_vector *XtX = gsl_vector_alloc(_Xmat->size2);
	gsl_permutation *allP = gsl_permutation_alloc(_Xmat->size2);
	
	_rankPred(y, SigI, XtX, allP);
	
	size_t Npck = ceil(_Xmat->size1*Nmul);
	vector< vector<size_t> > tmpUp(2);
	vector< vector<size_t> > tmpRLD;
	gsl_matrix *tmpX = gsl_matrix_alloc(_Xmat->size1, Npck);
	
	_ldToss(XtX, allP, rSqMax, Npck, tmpUp, tmpRLD, tmpX);
	gsl_vector_free(XtX);
	gsl_permutation_free(allP);
	
	_upLevel->init(tmpUp, tmpRLD);
	
	// keeping only the selected predictors
	gsl_matrix_free(_Xmat);
	_Xmat = gsl_matrix_alloc(tmpX->size1, tmpX->size2);
	gsl_matrix_memcpy(_Xmat, tmpX);
	gsl_matrix_free(tmpX);
	
	_theta.resize(_Xmat->size2);
	_valueMat  = gsl_matrix_calloc(_Xmat->size2, rsp.phenD());
	_fittedEach.resize(_Xmat->size2);

	for (size_t Xj = 0; Xj < (*_upLevel)[0].size(); Xj++) {
		_fittedEach[Xj].resize(rsp.Ndata()*rsp.phenD());
		_theta[Xj] = new MVnormBetaFt(y, _Xmat, Xj, _fittedEach[Xj], SigI.getMat(), _rV[0], _upLevel->priorInd(Xj), _valueMat, Xj);
	}
	
	_updateFitted();
	gsl_matrix_free(y);
}

BetaGrpFt::BetaGrpFt(const Grp &rsp, const SigmaI &SigI, const string &predFlNam, const size_t &Npred, const double &Nmul, const double &rSqMax, RanIndex &low, RanIndex &up, const string &outFlNam, const int &nThr) : _numSaves(0.0), _nThr(nThr), Grp(){
	if (Npred != up.getNtot()) {
		cerr << "ERROR: number of predictors " << Npred << " is not equal to the number of elements " << up.getNtot() << " in the prior index in BetaGrpFt initialization." << endl;
		exit(-1);
	}
	else if (low.getNtot() != rsp.dMat()->size1){
		cerr << "ERROR: number of elements " << low.getNgrp() << " in the replication index is not equal to the number of rows " << rsp.dMat()->size1 << " in the response matrix in BetaGrpFt initialization." << endl;
		exit(-1);
	}

	gsl_matrix_free(_valueMat);
	_fittedAll  = gsl_matrix_calloc(low.getNgrp(), rsp.phenD());
	_Xmat       = gsl_matrix_alloc(low.getNgrp(), Npred);
	_lowLevel   = &low;
	_upLevel    = &up;
	_outFlNam   = outFlNam;
	remove(_outFlNam.c_str());
	
	if (_nThr > 1) {
		const gsl_rng_type *T = gsl_rng_mt19937;
		_rV.resize(_nThr);
		vector<gsl_rng *>::iterator rIt = _rV.begin();
		++rIt;
		for (; rIt != _rV.end(); ++rIt) {
			unsigned long seed = gsl_rng_get(_rV[0]);
			*rIt = gsl_rng_alloc(T);
			gsl_rng_set(*rIt, seed);
		}
		
	}
	
	FILE *prdIn = fopen(predFlNam.c_str(), "r");
	if (prdIn == NULL) {
		cerr << "ERROR: cannot open predictor file " << predFlNam << " in BetaGrpFt" << endl;
		exit(-1);
	}
	gsl_matrix_fread(prdIn, _Xmat);
	fclose(prdIn);
	
	MuGrp rspMn = rsp.mean(*_lowLevel);
	gsl_matrix *y = gsl_matrix_alloc(rspMn.dMat()->size1, rspMn.dMat()->size2);
	colCenter(rspMn.dMat(), y); // approximating a regression with an intercept for initial values (not quite right, but close enough for initialization)
	
	gsl_vector *XtX = gsl_vector_alloc(_Xmat->size2);
	gsl_permutation *allP = gsl_permutation_alloc(_Xmat->size2);
	
	_rankPred(y, SigI, XtX, allP); // will center _Xmat
	gsl_matrix_free(y);
	
	size_t Npck = ceil(_Xmat->size1*Nmul);
	vector< vector<size_t> > tmpUp(2);
	vector< vector<size_t> > tmpRLD;
	gsl_matrix *tmpX = gsl_matrix_alloc(_Xmat->size1, Npck);
	
	_ldToss(XtX, allP, rSqMax, Npck, tmpUp, tmpRLD, tmpX);
	gsl_vector_free(XtX);
	gsl_permutation_free(allP);
	
	_upLevel->init(tmpUp, tmpRLD);
	
	// keeping only the selected predictors
	gsl_matrix_free(_Xmat);
	_Xmat = gsl_matrix_alloc(_lowLevel->getNtot(), tmpX->size2);
	for (size_t iGrp = 0; iGrp < _lowLevel->getNgrp(); iGrp++) {
		gsl_vector_view row = gsl_matrix_row(tmpX, iGrp);
		for (vector<size_t>::iterator grpIt = (*_lowLevel)[iGrp].begin(); grpIt != (*_lowLevel)[iGrp].end(); ++grpIt) {
			gsl_matrix_set_row(_Xmat, *grpIt, &row.vector);
		}
	}
	gsl_matrix_free(tmpX);
	
	_theta.resize(_Xmat->size2);
	_valueMat  = gsl_matrix_calloc(_Xmat->size2, rsp.phenD());
	_fittedEach.resize(_Xmat->size2);
	
	gsl_matrix *Ylng = gsl_matrix_alloc(rsp.dMat()->size1, rsp.dMat()->size2);
	colCenter(rsp.dMat(), Ylng);
	for (size_t Xj = 0; Xj < (*_upLevel)[0].size(); Xj++) {
		_fittedEach[Xj].resize(low.getNtot()*rsp.phenD());
		_theta[Xj] = new MVnormBetaFt(Ylng, _Xmat, Xj, _fittedEach[Xj], SigI.getMat(), _rV[0], _upLevel->priorInd(Xj), _valueMat, Xj);
	}
	
	_updateFitted();
	gsl_matrix_free(Ylng);
}

BetaGrpFt::BetaGrpFt(const Grp &rsp, const SigmaI &SigI, const string &predFlNam, const size_t &Npred, const double &Nmul, const double &rSqMax, const double &absLab, RanIndex &up, const string &outFlNam, const int &nThr) :  _numSaves(0.0), _nThr(nThr), Grp() {
	if (Npred != up.getNtot()) {
		cerr << "ERROR: number of predictors " << Npred << " is not equal to the number of elements " << up.getNtot() << " in the prior index in BetaGrpFt initialization." << endl;
		exit(-1);
	}
	
	gsl_matrix_free(_valueMat);
	_fittedAll  = gsl_matrix_calloc(rsp.Ndata(), rsp.phenD());
	_Xmat       = gsl_matrix_alloc(rsp.Ndata(), Npred);
	
	_upLevel  = &up;
	_outFlNam = outFlNam;
	remove(_outFlNam.c_str());
	
	if (_nThr > 1) {
		const gsl_rng_type *T = gsl_rng_mt19937;
		_rV.resize(_nThr);
		vector<gsl_rng *>::iterator rIt = _rV.begin();
		++rIt;
		for (; rIt != _rV.end(); ++rIt) {
			unsigned long seed = gsl_rng_get(_rV[0]);
			*rIt = gsl_rng_alloc(T);
			gsl_rng_set(*rIt, seed);
		}
		
	}
	
	FILE *prdIn = fopen(predFlNam.c_str(), "r");
	if (prdIn == NULL) {
		cerr << "ERROR: cannot open file " << predFlNam << " in BetaGrpFt" << endl;
		exit(-1);
	}
	gsl_matrix_fread(prdIn, _Xmat);
	fclose(prdIn);
	
	gsl_matrix *y = gsl_matrix_alloc(rsp.dMat()->size1, rsp.dMat()->size2);
	colCenter(rsp.dMat(), y); // approximating a regression with an intercept for initial values (not quite right, but close enough for initialization)
	
	gsl_vector *XtX = gsl_vector_alloc(_Xmat->size2);
	gsl_permutation *allP = gsl_permutation_alloc(_Xmat->size2);
	
	_rankPred(y, SigI, absLab, XtX, allP); // will center _Xmat
	
	size_t Npck = ceil(_Xmat->size1*Nmul);
	vector< vector<size_t> > tmpUp(2);
	vector< vector<size_t> > tmpRLD;
	gsl_matrix *tmpX = gsl_matrix_alloc(_Xmat->size1, Npck);
	
	_ldToss(XtX, allP, rSqMax, Npck, tmpUp, tmpRLD, tmpX);
	gsl_vector_free(XtX);
	gsl_permutation_free(allP);
	
	_upLevel->init(tmpUp, tmpRLD);
	
	// keeping only the selected predictors
	gsl_matrix_free(_Xmat);
	_Xmat = gsl_matrix_alloc(tmpX->size1, tmpX->size2);
	gsl_matrix_memcpy(_Xmat, tmpX);
	gsl_matrix_free(tmpX);
	
	_theta.resize(_Xmat->size2);
	_valueMat  = gsl_matrix_calloc(_Xmat->size2, rsp.phenD());
	_fittedEach.resize(_Xmat->size2);
	
	for (size_t Xj = 0; Xj < (*_upLevel)[0].size(); Xj++) {
		_fittedEach[Xj].resize(rsp.Ndata()*rsp.phenD());
		_theta[Xj] = new MVnormBetaFt(y, _Xmat, Xj, _fittedEach[Xj], SigI.getMat(), _rV[0], _upLevel->priorInd(Xj), _valueMat, Xj);
	}
	
	_updateFitted();
	gsl_matrix_free(y);
}
BetaGrpFt::BetaGrpFt(const Grp &rsp, const SigmaI &SigI, const string &predFlNam, const size_t &Npred, const double &Nmul, const double &rSqMax, const double &absLab, RanIndex &low, RanIndex &up, const string &outFlNam, const int &nThr) :  _numSaves(0.0), _nThr(nThr), Grp() {
	if (Npred != up.getNtot()) {
		cerr << "ERROR: number of predictors " << Npred << " is not equal to the number of elements " << up.getNtot() << " in the prior index in BetaGrpFt initialization." << endl;
		exit(-1);
	}
	else if (low.getNtot() != rsp.dMat()->size1){
		cerr << "ERROR: number of elements " << low.getNgrp() << " in the replication index is not equal to the number of rows " << rsp.dMat()->size1 << " in the response matrix in BetaGrpFt initialization." << endl;
		exit(-1);
	}

	gsl_matrix_free(_valueMat);
	_fittedAll  = gsl_matrix_calloc(low.getNgrp(), rsp.phenD());
	_Xmat       = gsl_matrix_alloc(low.getNgrp(), Npred);
	_lowLevel   = &low;
	_upLevel    = &up;
	_outFlNam   = outFlNam;
	remove(_outFlNam.c_str());
	
	if (_nThr > 1) {
		const gsl_rng_type *T = gsl_rng_mt19937;
		_rV.resize(_nThr);
		vector<gsl_rng *>::iterator rIt = _rV.begin();
		++rIt;
		for (; rIt != _rV.end(); ++rIt) {
			unsigned long seed = gsl_rng_get(_rV[0]);
			*rIt = gsl_rng_alloc(T);
			gsl_rng_set(*rIt, seed);
		}
		
	}
	
	FILE *prdIn = fopen(predFlNam.c_str(), "r");
	if (prdIn == NULL) {
		cerr << "ERROR: cannot open file " << predFlNam << " in BetaGrpFt" << endl;
		exit(-1);
	}
	gsl_matrix_fread(prdIn, _Xmat);
	fclose(prdIn);
	
	MuGrp rspMn = rsp.mean(*_lowLevel);
	gsl_matrix *y = gsl_matrix_alloc(rspMn.dMat()->size1, rspMn.dMat()->size2);
	colCenter(rspMn.dMat(), y); // approximating a regression with an intercept for initial values (not quite right, but close enough for initialization)
	
	gsl_vector *XtX = gsl_vector_alloc(_Xmat->size2);
	gsl_permutation *allP = gsl_permutation_alloc(_Xmat->size2);
	
	_rankPred(y, SigI, XtX, allP); // will center _Xmat
	gsl_matrix_free(y);
	
	size_t Npck = ceil(_Xmat->size1*Nmul);
	vector< vector<size_t> > tmpUp(2);
	vector< vector<size_t> > tmpRLD;
	gsl_matrix *tmpX = gsl_matrix_alloc(_Xmat->size1, Npck);
	
	_ldToss(XtX, allP, rSqMax, Npck, tmpUp, tmpRLD, tmpX);
	gsl_vector_free(XtX);
	gsl_permutation_free(allP);
	
	_upLevel->init(tmpUp, tmpRLD);
	
	// keeping only the selected predictors
	gsl_matrix_free(_Xmat);
	_Xmat = gsl_matrix_alloc(_lowLevel->getNtot(), tmpX->size2);
	for (size_t iGrp = 0; iGrp < _lowLevel->getNgrp(); iGrp++) {
		gsl_vector_view row = gsl_matrix_row(tmpX, iGrp);
		for (vector<size_t>::iterator grpIt = (*_lowLevel)[iGrp].begin(); grpIt != (*_lowLevel)[iGrp].end(); ++grpIt) {
			gsl_matrix_set_row(_Xmat, *grpIt, &row.vector);
		}
	}
	gsl_matrix_free(tmpX);
	
	_theta.resize(_Xmat->size2);
	_valueMat  = gsl_matrix_calloc(_Xmat->size2, rsp.phenD());
	_fittedEach.resize(_Xmat->size2);
	
	gsl_matrix *Ylng = gsl_matrix_alloc(rsp.dMat()->size1, rsp.dMat()->size2);
	colCenter(rsp.dMat(), Ylng);
	for (size_t Xj = 0; Xj < (*_upLevel)[0].size(); Xj++) {
		_fittedEach[Xj].resize(low.getNtot()*rsp.phenD());
		_theta[Xj] = new MVnormBetaFt(Ylng, _Xmat, Xj, _fittedEach[Xj], SigI.getMat(), _rV[0], _upLevel->priorInd(Xj), _valueMat, Xj);
	}
	
	_updateFitted();
	gsl_matrix_free(Ylng);
}


BetaGrpFt::BetaGrpFt(const BetaGrpFt &bG){
	_fittedAll = gsl_matrix_alloc((bG._fittedAll)->size1, (bG._fittedAll)->size2);
	gsl_matrix_memcpy(_fittedAll, bG._fittedAll);
	
	_fittedEach = bG._fittedEach;
	_lowLevel   = bG._lowLevel;
	_upLevel    = bG._upLevel;
	_nThr       = bG._nThr;
	
	if (_nThr > 1) {
		const gsl_rng_type *T = gsl_rng_mt19937;
		_rV.resize(_nThr);
		vector<gsl_rng *>::iterator rIt = _rV.begin();
		++rIt;
		for (; rIt != _rV.end(); ++rIt) {
			unsigned long seed = gsl_rng_get(_rV[0]);
			*rIt = gsl_rng_alloc(T);
			gsl_rng_set(*rIt, seed);
		}
		
	}

	_Xmat = gsl_matrix_alloc((bG._Xmat)->size1, (bG._Xmat)->size2);
	gsl_matrix_memcpy(_Xmat, bG._Xmat);
	
	gsl_matrix_free(_valueMat);
	_valueMat = gsl_matrix_alloc((bG._valueMat)->size1, (bG._valueMat)->size2);
	gsl_matrix_memcpy(_valueMat, bG._valueMat);
	
	_theta.resize(_valueMat->size1);
	for (size_t iVrw = 0; iVrw < _valueMat->size1; iVrw++) {
		_theta[iVrw] = new MVnormMu(_valueMat, iVrw, *(bG._theta[iVrw])->down(), *(bG._theta[iVrw])->up());
	}
	
}
BetaGrpFt & BetaGrpFt::operator=(const BetaGrpFt &bG){
	_fittedAll = gsl_matrix_alloc((bG._fittedAll)->size1, (bG._fittedAll)->size2);
	gsl_matrix_memcpy(_fittedAll, bG._fittedAll);
	
	_fittedEach = bG._fittedEach;
	_lowLevel   = bG._lowLevel;
	_upLevel    = bG._upLevel;
	_nThr       = bG._nThr;
	
	if (_nThr > 1) {
		const gsl_rng_type *T = gsl_rng_mt19937;
		_rV.resize(_nThr);
		vector<gsl_rng *>::iterator rIt = _rV.begin();
		++rIt;
		for (; rIt != _rV.end(); ++rIt) {
			unsigned long seed = gsl_rng_get(_rV[0]);
			*rIt = gsl_rng_alloc(T);
			gsl_rng_set(*rIt, seed);
		}
		
	}
	
	_Xmat = gsl_matrix_alloc((bG._Xmat)->size1, (bG._Xmat)->size2);
	gsl_matrix_memcpy(_Xmat, bG._Xmat);
	
	gsl_matrix_free(_valueMat);
	_valueMat = gsl_matrix_alloc((bG._valueMat)->size1, (bG._valueMat)->size2);
	gsl_matrix_memcpy(_valueMat, bG._valueMat);
	
	_theta.resize(_valueMat->size1);
	for (size_t iVrw = 0; iVrw < _valueMat->size1; iVrw++) {
		_theta[iVrw] = new MVnormMu(_valueMat, iVrw, *(bG._theta[iVrw])->down(), *(bG._theta[iVrw])->up());
	}
	
	return *this;
}


BetaGrpFt::~BetaGrpFt(){
	gsl_matrix_free(_fittedAll);
	gsl_matrix_free(_Xmat);
}

void BetaGrpFt::_updateFitted(){
	if (_lowLevel) {
#pragma omp parallel num_threads(_nThr)
		{
			gsl_vector *tmpVec = gsl_vector_alloc(_theta.size()); // have to make a OMP team copy of the temp array
			double totSum;
			
#pragma omp for
			for (int prdRow = 0; prdRow < _lowLevel->getNgrp(); prdRow++) {   // going through pred, row by row, but only non-redundant rows
				for (int bcol = 0; bcol < _valueMat->size2; bcol++) { // going through columns of coefficients (i.e. through _valueMat)
					totSum = 0.0;
					gsl_matrix_get_row(tmpVec, _Xmat, (*_lowLevel)[prdRow][0]);
					gsl_vector_view evCol = gsl_matrix_column(_valueMat, bcol);
					gsl_vector_mul(tmpVec, &evCol.vector); // multiplying a column of _valueMat (estimated values) and row of _Xmat (predictors), fill the tmpVec with a[im]*b[mi] (m = 1, ..., Npred), to sum next
					for (int iBt = 0; iBt < _valueMat->size1; iBt++) {
						totSum += gsl_vector_get(tmpVec, iBt);
					}
					gsl_matrix_set(_fittedAll,  prdRow, bcol, totSum); // _fittedAll only has the unique elements
					if (_valueMat->size1 > 1) {
						for (int jBt = 0; jBt < _valueMat->size1; jBt++) {
							double pSum = totSum - gsl_vector_get(tmpVec, jBt);
							for (vector<size_t>::iterator grpIt = (*_lowLevel)[prdRow].begin(); grpIt != (*_lowLevel)[prdRow].end(); ++grpIt) {
								_fittedEach[jBt][(*grpIt)*(_valueMat->size2) + bcol] = pSum; // following the idiom set in the GSL description of the matrix element representation in the underlying array
							}
						}
					}
				}
			}
			
			gsl_vector_free(tmpVec);
		} // end parallel block
	}
	else {
#pragma omp parallel num_threads(_nThr)
		{
			gsl_vector *tmpVec = gsl_vector_alloc(_theta.size()); // have to make a OMP team copy of the temp array
			double pSum;
			
#pragma omp for
			for (int prdRow = 0; prdRow < _Xmat->size1; prdRow++) {   // going through pred, row by row
				for (int bcol = 0; bcol < _valueMat->size2; bcol++) { // going through columns of coefficients (i.e. through _valueMat)
					pSum = 0.0;
					gsl_matrix_get_row(tmpVec, _Xmat, prdRow);
					gsl_vector_view evCol = gsl_matrix_column(_valueMat, bcol);
					gsl_vector_mul(tmpVec, &evCol.vector); // multiplying a column of _valueMat (estimated values) and row of _Xmat (predictors), fill the tmpVec with a[im]*b[mi] (m = 1, ..., Npred), to sum next
					for (int iBt = 0; iBt < _valueMat->size1; iBt++) {
						pSum += gsl_vector_get(tmpVec, iBt);
					}
					gsl_matrix_set(_fittedAll,  prdRow, bcol, pSum);
					if (_valueMat->size1 > 1) {
						for (int jBt = 0; jBt < _valueMat->size1; jBt++) {
							_fittedEach[jBt][prdRow*(_valueMat->size2) + bcol] = pSum - gsl_vector_get(tmpVec, jBt); // following the idiom set in the GSL description of the matrix element representation in the underlying array
						}
					}
				}
			}
			
			gsl_vector_free(tmpVec);
		} // end parallel block
	}
}

void BetaGrpFt::_rankPred(const gsl_matrix *y, const SigmaI &SigI, gsl_vector *XtX, gsl_permutation *prm){
	gsl_vector *mhl = gsl_vector_alloc(_Xmat->size2);
#pragma omp parallel num_threads(_nThr)
	{
		gsl_vector *b   = gsl_vector_alloc(y->size2);
		gsl_vector *bM  = gsl_vector_alloc(y->size2);
		gsl_vector *xCl = gsl_vector_alloc(y->size1);
		double dotPr;
#pragma omp for
		for (size_t Xj = 0; Xj < _Xmat->size2; Xj++) {
			// mean-center the predictor
			gsl_vector_view Xcol = gsl_matrix_column(_Xmat, Xj);
			gsl_matrix_get_col(xCl, _Xmat, Xj);
			double mn = gsl_stats_mean(xCl->data, 1, xCl->size);
			gsl_vector_add_constant(&Xcol.vector, -mn);
			gsl_blas_ddot(&Xcol.vector, &Xcol.vector, &dotPr);
			dotPr = 1.0/dotPr;
			gsl_vector_set(XtX, Xj, dotPr);
			
			// now, the regression
			gsl_blas_dgemv(CblasTrans, dotPr, y, &Xcol.vector, 0.0, b);
			
			double m;
			gsl_blas_dsymv(CblasLower, 1.0, SigI.getMat(), b, 0.0, bM);
			gsl_blas_ddot(b, bM, &m);
			
			gsl_vector_set(mhl, Xj, m*dotPr);
		}
		gsl_vector_free(b);
		gsl_vector_free(bM);
		gsl_vector_free(xCl);
	}
	gsl_sort_vector_index(prm, mhl);
	gsl_permutation_reverse(prm); // sort is in the order of increase
	gsl_vector_free(mhl);
	
}
void BetaGrpFt::_rankPred(const gsl_matrix *y, const SigmaI &SigI, const double &absLab, gsl_vector *XtX, gsl_permutation *prm){
	gsl_vector *mhl = gsl_vector_alloc(_Xmat->size2);
#pragma omp parallel num_threads(_nThr)
	{
		gsl_vector *b  = gsl_vector_alloc(y->size2);
		gsl_vector *bM = gsl_vector_alloc(y->size2);
		vector<double> XnonMiss;
		double dotPr;
#pragma omp for
		for (size_t Xj = 0; Xj < _Xmat->size2; Xj++) {
			vector<size_t> missInd;
			for (size_t Xi = 0; Xi < _Xmat->size1; Xi++) {
				double cX = gsl_matrix_get(_Xmat, Xi, Xj);
				if (cX > absLab) {
					XnonMiss.push_back(cX);
				}
				else {
					missInd.push_back(Xi);
				}
			}
			
			// mean-center the predictor
			gsl_vector_view Xcol = gsl_matrix_column(_Xmat, Xj);
			double mn = gsl_stats_mean(XnonMiss.data(), 1, XnonMiss.size());
			XnonMiss.clear();
			gsl_vector_add_constant(&Xcol.vector, -mn);
			if (missInd.size()) {
				for (size_t iMs = 0; iMs < missInd.size(); iMs++) { // again, not using an iterator b/c of concerns with openmp
					gsl_matrix_set(_Xmat, missInd[iMs], Xj, 0.0);
				}
				
			}
			
			gsl_blas_ddot(&Xcol.vector, &Xcol.vector, &dotPr);
			dotPr = 1.0/dotPr;
			gsl_vector_set(XtX, Xj, dotPr);
			
			// now, the regression
			gsl_blas_dgemv(CblasTrans, dotPr, y, &Xcol.vector, 0.0, b);
			
			double m;
			gsl_blas_dsymv(CblasLower, 1.0, SigI.getMat(), b, 0.0, bM);
			gsl_blas_ddot(b, bM, &m);
			
			gsl_vector_set(mhl, Xj, m*dotPr);
		}
		gsl_vector_free(b);
		gsl_vector_free(bM);
	}
	gsl_sort_vector_index(prm, mhl);
	gsl_permutation_reverse(prm); // sort is in the order of increase
	gsl_vector_free(mhl);
	
}

void BetaGrpFt::_ldToss(const gsl_vector *var, const gsl_permutation *prm, const double &rSqMax, const size_t &Npck, vector< vector<size_t> > &idx, vector< vector<size_t> > &rLd, gsl_matrix *Xpck){
	rLd.resize(Npck);
	size_t nPicked = 0;
	size_t crEdge  = Npck; // index of the permutation that is one beyond the current "edge" of the candidate picks
	list<size_t> cand;
	
	for (size_t iEl = 0; iEl < Npck; iEl++) {
		cand.push_back(gsl_permutation_get(prm, iEl));
	}
	
	// go through all the candidates, tossing out (and recording in tmpLd) all the ones in LD > rSqMax with the top picks
	while ((nPicked < Npck) && (crEdge < _Xmat->size2)) {
		cout << "\r" << nPicked + 1 << flush;
		list<size_t>::iterator cndIt = cand.begin();
		rLd[nPicked].push_back(*cndIt);  // the first element of ld[i] is the index of the picked predictor
		gsl_vector_view curX = gsl_matrix_column(_Xmat, *cndIt);
		double curVr = gsl_vector_get(var, *cndIt);
		gsl_matrix_set_col(Xpck, nPicked, &curX.vector);
		
		idx[0].push_back(nPicked);
		
		cndIt = cand.erase(cndIt);
		int delCt = 0;
		while (cndIt != cand.end()) {
			gsl_vector_view tstX = gsl_matrix_column(_Xmat, *cndIt);
			double tstVr = gsl_vector_get(var, *cndIt);
			double rSq;
			gsl_blas_ddot(&curX.vector, &tstX.vector, &rSq);
			rSq = gsl_pow_2(rSq)*(curVr*tstVr); // sample sizes are all the same and cancel out, so crossproducts will do; also XtX is inverted, so multiply rather than divide
			if (rSq >= rSqMax) {
				rLd[nPicked].push_back(*cndIt);
				cndIt = cand.erase(cndIt);
				delCt++;
			}
			else cndIt++;
			
		}
		if (delCt) {
			while (delCt) {
				gsl_vector_view tstX = gsl_matrix_column(_Xmat, gsl_permutation_get(prm, crEdge));
				double tstVr = gsl_vector_get(var, crEdge);
				for (size_t iEl = 0; iEl < nPicked; iEl++) {
					gsl_vector_view pckX = gsl_matrix_column(_Xmat, rLd[iEl][0]);
					double pckVr = gsl_vector_get(var, rLd[iEl][0]);
					double rSq;
					gsl_blas_ddot(&tstX.vector, &pckX.vector, &rSq);
					rSq = gsl_pow_2(rSq)*(tstVr*pckVr);
					if (rSq >= rSqMax) { // have to check if ANY of the previously picked SNPs are in LD with the additional candidate
						if (crEdge == _Xmat->size2 - 1) {
							rLd.resize(nPicked+1);
							break;
						}
						rLd[iEl].push_back(gsl_permutation_get(prm, crEdge));
						crEdge++;
						continue;
					}
				}
				cand.push_back(gsl_permutation_get(prm, crEdge));
				delCt--;
				crEdge++;
			}
		}
		nPicked++;
	}
	cout << endl;
	
}

double BetaGrpFt::_MGkernel(const Grp &dat, const SigmaI &SigI) const{
	/*
	 *  MV Gaussian kernel, typically written as sum{(x[i]-mu)Sig^{-1}(x[i]-m)'}, can be re-written as Tr[Sig^{-1}(x-m)'(x-m)], which in turn is ddot(vec[Sig^{-1}]vec[(x-m)'(x-m)]) and can be calculated several-fold faster b/c of the use of level3 BLAS (especially for big x)
	 *  Have to test for the case where there is replication!!!
	 */
	gsl_matrix *dif = gsl_matrix_alloc(_fittedAll->size1, _fittedAll->size2);
	gsl_matrix *s   = gsl_matrix_alloc(_fittedAll->size2, _fittedAll->size2);
	double res;
	gsl_matrix_memcpy(dif, dat.dMat());
	gsl_matrix_sub(dif, _fittedAll);
	
	gsl_blas_dsyrk(CblasLower, CblasTrans, 1.0, dif, 0.0, s);      // (x-m)'(x-m)
	gsl_vector *lt1 = gsl_vector_alloc(s->size1*(s->size1 + 1)/2);
	gsl_vector *lt2 = gsl_vector_alloc(s->size1*(s->size1 + 1)/2);
	size_t iTrk = 0;
	for (size_t iCl = 0; iCl < s->size2; iCl++) {  // will copy over only the lower triangle of the symmetric matrices b/c the upper is not necessarily defined
		gsl_vector_set(lt1, iTrk, gsl_matrix_get(SigI.getMat(), iCl, iCl));
		gsl_vector_set(lt2, iTrk, gsl_matrix_get(s, iCl, iCl));
		iTrk++;
		for (size_t iRw = iCl + 1; iRw < s->size1; iRw++) {
			gsl_vector_set(lt1, iTrk, 2.0*gsl_matrix_get(SigI.getMat(), iRw, iCl));
			gsl_vector_set(lt2, iTrk, gsl_matrix_get(s, iRw, iCl));
			iTrk++;
		}
	}
	gsl_blas_ddot(lt1, lt2, &res);
	
	gsl_matrix_free(dif);
	gsl_matrix_free(s);
	
	return res;
	
}
double BetaGrpFt::_MGkernel(const Grp &dat, const SigmaI &SigI, const size_t &prInd) const{ // same, but for the partial fitted matrices
	gsl_matrix *dif = gsl_matrix_alloc(_fittedAll->size1, _fittedAll->size2);
	gsl_matrix *s   = gsl_matrix_alloc(_fittedAll->size2, _fittedAll->size2);
	double res;
	
	gsl_matrix_memcpy(dif, dat.dMat());
	gsl_matrix_const_view pFit = gsl_matrix_const_view_array(_fittedEach[prInd].data(), _fittedAll->size1, _fittedAll->size2);
	gsl_matrix_sub(dif, &pFit.matrix);
	
	gsl_blas_dsyrk(CblasLower, CblasTrans, 1.0, dif, 0.0, s);      // (x-m)'(x-m)
	gsl_vector *lt1 = gsl_vector_alloc(s->size1*(s->size1 + 1)/2);
	gsl_vector *lt2 = gsl_vector_alloc(s->size1*(s->size1 + 1)/2);
	size_t iTrk = 0;
	for (size_t iCl = 0; iCl < s->size2; iCl++) {  // will copy over only the lower triangle of the symmetric matrices b/c the upper is not necessarily defined
		gsl_vector_set(lt1, iTrk, gsl_matrix_get(SigI.getMat(), iCl, iCl));
		gsl_vector_set(lt2, iTrk, gsl_matrix_get(s, iCl, iCl));
		iTrk++;
		for (size_t iRw = iCl + 1; iRw < s->size1; iRw++) {
			gsl_vector_set(lt1, iTrk, 2.0*gsl_matrix_get(SigI.getMat(), iRw, iCl));
			gsl_vector_set(lt2, iTrk, gsl_matrix_get(s, iRw, iCl));
			iTrk++;
		}
	}
	gsl_blas_ddot(lt1, lt2, &res);
	
	gsl_matrix_free(dif);
	gsl_matrix_free(s);
	
	return res;
}

void BetaGrpFt::save(const SigmaI &SigI){
	gsl_vector_const_view sI = gsl_matrix_const_diagonal(SigI.getMat());
	if (_numSaves) {
		gsl_vector *tmpB = gsl_vector_alloc(_valueMat->size2);
		gsl_vector *tmpS = gsl_vector_alloc(_valueMat->size2 + 1);
		for (size_t vmRw = 0; vmRw < _valueMat->size1; vmRw++) {
			gsl_vector_view vsRw = gsl_matrix_row(_valueSum, vmRw);
			gsl_vector_view bS   = gsl_vector_subvector(tmpS, 0, _valueMat->size2);
			gsl_matrix_get_row(tmpB, _valueMat, vmRw);
			gsl_vector_memcpy(&bS.vector, tmpB);
			gsl_vector_mul(&bS.vector, tmpB);
			gsl_vector_mul(&bS.vector, &sI.vector);
			
			double md = _theta[vmRw]->mhl(tmpB, SigI);
			gsl_vector_set(tmpS, _valueMat->size2, md);
			gsl_vector_add(&vsRw.vector, tmpS);
		}
		_numSaves += 1.0;
		gsl_vector_free(tmpB);
		gsl_vector_free(tmpS);
	}
	else {
		_valueSum = gsl_matrix_alloc(_valueMat->size1, _valueMat->size2 + 1);
		gsl_vector *tmpB = gsl_vector_alloc(_valueMat->size2);
		gsl_vector *tmpS = gsl_vector_alloc(_valueMat->size2 + 1);
		for (size_t vmRw = 0; vmRw < _valueMat->size1; vmRw++) {
			gsl_vector_view bS   = gsl_vector_subvector(tmpS, 0, _valueMat->size2);
			gsl_matrix_get_row(tmpB, _valueMat, vmRw);
			gsl_vector_memcpy(&bS.vector, tmpB);
			gsl_vector_mul(&bS.vector, tmpB);
			gsl_vector_mul(&bS.vector, &sI.vector);
			
			double md = _theta[vmRw]->mhl(tmpB, SigI);
			gsl_vector_set(tmpS, _valueMat->size2, md);
			gsl_matrix_set_row(_valueSum, vmRw, tmpS);
		}
		_numSaves += 1.0;
		gsl_vector_free(tmpB);
		gsl_vector_free(tmpS);

	}
}

void BetaGrpFt::dump(){
	if (_numSaves) {
		gsl_matrix_scale(_valueSum, 1.0/_numSaves);
		FILE *btOut = fopen(_outFlNam.c_str(), "w");
		gsl_matrix_fwrite(btOut, _valueSum);
		fclose(btOut);
		gsl_matrix_free(_valueSum);
	}
	
}
double BetaGrpFt::lnOddsRat(const Grp &y, const SigmaI &SigI, const size_t i) const {
	if (y.fMat()->size1 != _fittedAll->size1) {
#ifdef MYDEBUG
		if (y.fMat()->size1 != _lowLevel->getNtot()) {
			cerr << "ERROR: y does not have the same # of rows (" << y.fMat()->size1 << ") as the number of elements in _lowLevel (" << _lowLevel->getNtot() << ") when calculating lnOddsRat on line " << __LINE__ << " of file " << __FILE__ << endl;
			exit(-1);
		}
#endif
		MuGrp yMnI = y.mean((*_lowLevel));
		Grp &yMn   = yMnI;
		return 0.5*(_MGkernel(yMn, SigI, i) - _MGkernel(yMn, SigI));
	}
	else {
		return 0.5*(_MGkernel(y, SigI, i) - _MGkernel(y, SigI));
	}
}


/*
 *	Update functions
 */
// improper prior
void BetaGrpFt::update(const Grp &dat, const SigmaI &SigIm){
	for (vector<MVnorm *>::const_iterator elm = _theta.begin(); elm != _theta.end(); ++elm) {
		(*elm)->update(dat, SigIm, _rV[0]);
	}
	_updateFitted();
}
void BetaGrpFt::update(const Grp &dat, const Qgrp &q, const SigmaI &SigIm){
#pragma omp parallel for num_threads(_nThr)
	for (int iTh = 0; iTh < _theta.size(); iTh++) {
		_theta[iTh]->update(dat, q, SigIm, _rV[omp_get_thread_num()]);
	}
	_updateFitted();
}

// 0-mean prior
void BetaGrpFt::update(const Grp &dat, const SigmaI &SigIm, const SigmaI &SigIp){
#pragma omp parallel for num_threads(_nThr)
	for (int iTh = 0; iTh < _theta.size(); iTh++) {
		_theta[iTh]->update(dat, SigIm, SigIp, _rV[omp_get_thread_num()]);
	}
	_updateFitted();
}
void BetaGrpFt::update(const Grp &dat, const Qgrp &q, const SigmaI &SigIm, const SigmaI &SigIp){
#pragma omp parallel for num_threads(_nThr)
	for (int iTh = 0; iTh < _theta.size(); iTh++) {
		_theta[iTh]->update(dat, q, SigIm, SigIp, _rV[omp_get_thread_num()]);
	}
	_updateFitted();
}

void BetaGrpFt::update(const Grp &dat, const SigmaI &SigIm, const Qgrp &qPr, const SigmaI &SigIp){
#pragma omp parallel for num_threads(_nThr)
	for (size_t iTh = 0; iTh < _theta.size(); iTh++) {
		_theta[iTh]->update(dat, SigIm, qPr[iTh], SigIp, _rV[omp_get_thread_num()]);
	}
	_updateFitted();
}
void BetaGrpFt::update(const Grp &dat, const Qgrp &q, const SigmaI &SigIm, const Qgrp &qPr, const SigmaI &SigIp){
#pragma omp parallel for num_threads(_nThr)
	for (int iTh = 0; iTh < _theta.size(); iTh++) {
		_theta[iTh]->update(dat, q, SigIm, qPr[iTh], SigIp, _rV[omp_get_thread_num()]);
	}
	_updateFitted();
}

// non-0-mean prior
void BetaGrpFt::update(const Grp &dat, const SigmaI &SigIm, const Grp &muPr, const SigmaI &SigIp){
#pragma omp parallel for num_threads(_nThr)
	for (int iTh = 0; iTh < _theta.size(); iTh++) {
		_theta[iTh]->update(dat, SigIm, muPr, SigIp, _rV[omp_get_thread_num()]);
	}
	_updateFitted();
}
void BetaGrpFt::update(const Grp &dat, const Qgrp &q, const SigmaI &SigIm, const Grp &muPr, const SigmaI &SigIp){
#pragma omp parallel for num_threads(_nThr)
	for (int iTh = 0; iTh < _theta.size(); iTh++) {
		_theta[iTh]->update(dat, q, SigIm, muPr, SigIp, _rV[omp_get_thread_num()]);
	}
	_updateFitted();
}
void BetaGrpFt::update(const Grp &dat, const SigmaI &SigIm, const Grp &muPr, const Qgrp &qPr, const SigmaI &SigIp){
#pragma omp parallel for num_threads(_nThr)
	for (int iTh = 0; iTh < _theta.size(); iTh++) {
		_theta[iTh]->update(dat, SigIm, muPr, qPr[iTh], SigIp, _rV[omp_get_thread_num()]);
	}
	_updateFitted();
}
void BetaGrpFt::update(const Grp &dat, const Qgrp &q, const SigmaI &SigIm, const Grp &muPr, const Qgrp &qPr, const SigmaI &SigIp){
#pragma omp parallel for num_threads(_nThr)
	for (int iTh = 0; iTh < _theta.size(); iTh++) {
		_theta[iTh]->update(dat, q, SigIm, muPr, qPr[iTh], SigIp, _rV[omp_get_thread_num()]);
	}
	_updateFitted();
}

/*
 *	BetaGrpPEX methods
 */

BetaGrpPEX::~BetaGrpPEX() {
	gsl_matrix_free(_tSigIAt);
	gsl_matrix_free(_fittedAllAdj);
}

void BetaGrpPEX::_finishConstruct(const double &Spr){
	_ftA.resize(_valueMat->size2);
	_tSigIAt = gsl_matrix_alloc(_valueMat->size2, _valueMat->size2);
	_A       = Apex(Spr, _valueMat->size2, _ftA);
	
	_fittedAllAdj = gsl_matrix_alloc(_fittedAll->size1, _fittedAll->size2);
	gsl_matrix_memcpy(_fittedAllAdj, _fittedAll);
	
	for (size_t jCl = 0; jCl < _valueMat->size2; jCl++) {
		_ftA[jCl].resize(_Xmat->size1*_valueMat->size2);
	}
	for (size_t iMn = 0; iMn < _theta.size(); iMn++) {
		delete _theta[iMn];
		_theta[iMn] = new MVnormBetaPEX(_valueMat->size2, _Xmat, iMn, _fittedEach[iMn], _upLevel->priorInd(iMn), _valueMat, iMn, _A, _tSigIAt);
	}
}

void BetaGrpPEX::_finishFitted(){
	if (_lowLevel) {
#pragma omp parallel num_threads(_nThr)
		{
			gsl_matrix *tmpFt = gsl_matrix_alloc(_lowLevel->getNtot(), _fittedAll->size2);
			
#pragma omp for
			for (size_t iEa = 0; iEa < _fittedEach.size(); iEa++) { // not using an iterator in case the OpenMP version in use balks
				gsl_matrix_view curFE = gsl_matrix_view_array(_fittedEach[iEa].data(), _lowLevel->getNtot(), _fittedAll->size2);
				gsl_blas_dgemm(CblasNoTrans, CblasNoTrans, 1.0, &curFE.matrix, _A.getMat(), 0.0, tmpFt);
				gsl_matrix_memcpy(&curFE.matrix, tmpFt);
			}
			
			gsl_matrix_free(tmpFt);
		}

	}
	else {
#pragma omp parallel num_threads(_nThr)
		{
			gsl_matrix *tmpFt = gsl_matrix_alloc(_fittedAll->size1, _fittedAll->size2);
			
#pragma omp for
			for (size_t iEa = 0; iEa < _fittedEach.size(); iEa++) { // not using an iterator in case the OpenMP version in use balks
				gsl_matrix_view curFE = gsl_matrix_view_array(_fittedEach[iEa].data(), _fittedAll->size1, _fittedAll->size2);
				gsl_blas_dgemm(CblasNoTrans, CblasNoTrans, 1.0, &curFE.matrix, _A.getMat(), 0.0, tmpFt);
				gsl_matrix_memcpy(&curFE.matrix, tmpFt);
			}
			
			gsl_matrix_free(tmpFt);
		}
	}
	
}

void BetaGrpPEX::_updateAfitted(){ 
	if (_lowLevel) {
#pragma omp parallel num_threads(_nThr)
		{
			gsl_vector *tmpVec = gsl_vector_alloc(_fittedAll->size2); // have to make an OMP team copy of the temp array
			double totSum;
			
#pragma omp for
			for (int prdRow = 0; prdRow < _lowLevel->getNgrp(); prdRow++) {
				for (int bcol = 0; bcol < _A.getMat()->size2; bcol++) {
					totSum = 0.0;
					gsl_matrix_get_row(tmpVec, _fittedAll, prdRow);
					gsl_vector_view evCol = gsl_matrix_column(_A.getMat(), bcol);
					gsl_vector_mul(tmpVec, &evCol.vector);
					for (int iBt = 0; iBt < _A.getMat()->size1; iBt++) {
						totSum += gsl_vector_get(tmpVec, iBt);
					}
					gsl_matrix_set(_fittedAllAdj,  prdRow, bcol, totSum);
					for (int jBt = 0; jBt < _A.getMat()->size1; jBt++) {
						double pSumDif = totSum - gsl_vector_get(tmpVec, jBt);
						for (vector<size_t>::iterator grpIt = (*_lowLevel)[prdRow].begin(); grpIt != (*_lowLevel)[prdRow].end(); ++grpIt) {
							_ftA[jBt][(*grpIt)*(_fittedAll->size2) + bcol] = pSumDif; // individual fitted matrices always match the _lowLevel->getNtot()
						}
					}
				}
			}
			
			gsl_vector_free(tmpVec);
		} // end parallel block

	}
	else {
#pragma omp parallel num_threads(_nThr)
		{
			gsl_vector *tmpVec = gsl_vector_alloc(_fittedAll->size2); // have to make an OMP team copy of the temp array
			double pSum;
			
#pragma omp for
			for (int prdRow = 0; prdRow < _fittedAll->size1; prdRow++) {
				for (int bcol = 0; bcol < _A.getMat()->size2; bcol++) {
					pSum = 0.0;
					gsl_matrix_get_row(tmpVec, _fittedAll, prdRow);
					gsl_vector_view evCol = gsl_matrix_column(_A.getMat(), bcol);
					gsl_vector_mul(tmpVec, &evCol.vector);
					for (int iBt = 0; iBt < _A.getMat()->size1; iBt++) {
						pSum += gsl_vector_get(tmpVec, iBt);
					}
					gsl_matrix_set(_fittedAllAdj,  prdRow, bcol, pSum);
					for (int jBt = 0; jBt < _A.getMat()->size1; jBt++) {
						double pSumDif = pSum - gsl_vector_get(tmpVec, jBt);
						_ftA[jBt][prdRow*(_fittedAll->size2) + bcol] = pSumDif;
					}
				}
			}
			
			gsl_vector_free(tmpVec);
		} // end parallel block
	}
}

void BetaGrpPEX::save(){
	gsl_matrix *adjMat = gsl_matrix_alloc(_valueMat->size1, _valueMat->size2);
	gsl_blas_dgemm(CblasNoTrans, CblasNoTrans, 1.0, _valueMat, _A.getMat(), 0.0, adjMat);
	
	FILE *outH = fopen(_outFlNam.c_str(), "a");
	gsl_matrix_fwrite(outH, adjMat);
	fclose(outH);
	
	gsl_matrix_free(adjMat);
}
void BetaGrpPEX::save(const string &outFlNam){
	gsl_matrix *adjMat = gsl_matrix_alloc(_valueMat->size1, _valueMat->size2);
	gsl_blas_dgemm(CblasNoTrans, CblasNoTrans, 1.0, _valueMat, _A.getMat(), 0.0, adjMat);
	
	if (_outFlNam == outFlNam) {
		FILE *outH = fopen(outFlNam.c_str(), "a");
		gsl_matrix_fwrite(outH, adjMat);
		fclose(outH);
		
	}
	else {
		_outFlNam = outFlNam;
		remove(_outFlNam.c_str());
		FILE *outH = fopen(outFlNam.c_str(), "a");
		gsl_matrix_fwrite(outH, adjMat);
		fclose(outH);
	}
	gsl_matrix_free(adjMat);
}
void BetaGrpPEX::save(const SigmaI &SigI){
	gsl_matrix *Sig   = gsl_matrix_alloc(SigI.getMat()->size1, SigI.getMat()->size1);
	gsl_matrix *SigTr = gsl_matrix_alloc(SigI.getMat()->size1, SigI.getMat()->size1);
	gsl_matrix_memcpy(Sig, SigI.getMat());
	gsl_linalg_cholesky_decomp(Sig);
	gsl_linalg_cholesky_invert(Sig);
	
	gsl_blas_dsymm(CblasLeft, CblasLower, 1.0, Sig, _A.getMat(), 0.0, SigTr);
	gsl_blas_dgemm(CblasTrans, CblasNoTrans, 1.0, _A.getMat(), SigTr, 0.0, Sig);
	
	FILE *outS = fopen(SigI.getOutFile().c_str(), "a");
	gsl_matrix_fwrite(outS, Sig);
	fclose(outS);
	
	gsl_matrix_free(Sig);
	gsl_matrix_free(SigTr);
	
	gsl_matrix *adjMat = gsl_matrix_alloc(_valueMat->size1, _valueMat->size2);
	gsl_blas_dgemm(CblasNoTrans, CblasNoTrans, 1.0, _valueMat, _A.getMat(), 0.0, adjMat);
	FILE *outM = fopen(_outFlNam.c_str(), "a");
	gsl_matrix_fwrite(outM, adjMat);
	fclose(outM);
	gsl_matrix_free(adjMat);
}


void BetaGrpPEX::update(const Grp &dat, const SigmaI &SigIm, const SigmaI &SigIp){
	gsl_blas_dsymm(CblasRight, CblasLower, 1.0, SigIm.getMat(), _A.getMat(), 0.0, _tSigIAt);
	
#pragma omp parallel for num_threads(_nThr)
	for (int iEl = 0; iEl < _theta.size(); iEl++) {
		_theta[iEl]->update(dat, SigIm, SigIp, _rV[omp_get_thread_num()]);
	}
	
	_updateFitted();
	_finishFitted();
	_updateAfitted();
	if (_lowLevel) {
		_A.update(dat, _fittedAll, SigIm, *_lowLevel);
	}
	else {
		_A.update(dat, _fittedAll, SigIm);
	}

}
void BetaGrpPEX::update(const Grp &dat, const Qgrp &q, const SigmaI &SigIm, const SigmaI &SigIp){
	gsl_blas_dsymm(CblasRight, CblasLower, 1.0, SigIm.getMat(), _A.getMat(), 0.0, _tSigIAt);
	
#pragma omp parallel for num_threads(_nThr)
	for (int iEl = 0; iEl < _theta.size(); iEl++) {
		_theta[iEl]->update(dat, q, SigIm, SigIp, _rV[omp_get_thread_num()]);
	}
	
	_updateFitted();
	_finishFitted();
	_updateAfitted();
	if (_lowLevel) {
		_A.update(dat, _fittedAll, q, SigIm, *_lowLevel);
	}
	else {
		_A.update(dat, _fittedAll, SigIm);
	}
	
}
void BetaGrpPEX::update(const Grp &dat, const SigmaI &SigIm, const Qgrp &qPr, const SigmaI &SigIp){
	gsl_blas_dsymm(CblasRight, CblasLower, 1.0, SigIm.getMat(), _A.getMat(), 0.0, _tSigIAt);
	
#pragma omp parallel for num_threads(_nThr)
	for (int iEl = 0; iEl < _theta.size(); iEl++) {
		_theta[iEl]->update(dat, SigIm, qPr[iEl], SigIp, _rV[omp_get_thread_num()]);
	}
	
	_updateFitted();
	_finishFitted();
	_updateAfitted();
	if (_lowLevel) {
		_A.update(dat, _fittedAll, SigIm, *_lowLevel);
	}
	else {
		_A.update(dat, _fittedAll, SigIm);
	}
}
void BetaGrpPEX::update(const Grp &dat, const Qgrp &q, const SigmaI &SigIm, const Qgrp &qPr, const SigmaI &SigIp){
	gsl_blas_dsymm(CblasRight, CblasLower, 1.0, SigIm.getMat(), _A.getMat(), 0.0, _tSigIAt);
	
#pragma omp parallel for num_threads(_nThr)
	for (int iEl = 0; iEl < _theta.size(); iEl++) {
		_theta[iEl]->update(dat, q, SigIm, qPr[iEl], SigIp, _rV[omp_get_thread_num()]);
	}
	
	_updateFitted();
	_finishFitted();
	_updateAfitted();
	if (_lowLevel) {
		_A.update(dat, _fittedAll, q, SigIm, *_lowLevel);
	}
	else {
		_A.update(dat, _fittedAll, SigIm);
	}
}

void BetaGrpPEX::update(const Grp &dat, const SigmaI &SigIm, const Grp &muPr, const SigmaI &SigIp){
	gsl_blas_dsymm(CblasRight, CblasLower, 1.0, SigIm.getMat(), _A.getMat(), 0.0, _tSigIAt);
	
#pragma omp parallel for num_threads(_nThr)
	for (int iEl = 0; iEl < _theta.size(); iEl++) {
		_theta[iEl]->update(dat, SigIm, muPr, SigIp, _rV[omp_get_thread_num()]);
	}
	
	_updateFitted();
	_finishFitted();
	_updateAfitted();
	if (_lowLevel) {
		_A.update(dat, _fittedAll, SigIm, *_lowLevel);
	}
	else {
		_A.update(dat, _fittedAll, SigIm);
	}
}
void BetaGrpPEX::update(const Grp &dat, const Qgrp &q, const SigmaI &SigIm, const Grp &muPr, const SigmaI &SigIp){
	gsl_blas_dsymm(CblasRight, CblasLower, 1.0, SigIm.getMat(), _A.getMat(), 0.0, _tSigIAt);
	
#pragma omp parallel for num_threads(_nThr)
	for (int iEl = 0; iEl < _theta.size(); iEl++) {
		_theta[iEl]->update(dat, q, SigIm, muPr, SigIp, _rV[omp_get_thread_num()]);
	}
	
	_updateFitted();
	_finishFitted();
	_updateAfitted();
	if (_lowLevel) {
		_A.update(dat, _fittedAll, q, SigIm, *_lowLevel);
	}
	else {
		_A.update(dat, _fittedAll, SigIm);
	}
}
void BetaGrpPEX::update(const Grp &dat, const SigmaI &SigIm, const Grp &muPr, const Qgrp &qPr, const SigmaI &SigIp){
	gsl_blas_dsymm(CblasRight, CblasLower, 1.0, SigIm.getMat(), _A.getMat(), 0.0, _tSigIAt);
	
#pragma omp parallel for num_threads(_nThr)
	for (int iEl = 0; iEl < _theta.size(); iEl++) {
		_theta[iEl]->update(dat, SigIm, muPr, qPr[iEl], SigIp, _rV[omp_get_thread_num()]);
	}
	
	_updateFitted();
	_finishFitted();
	_updateAfitted();
	if (_lowLevel) {
		_A.update(dat, _fittedAll, SigIm, *_lowLevel);
	}
	else {
		_A.update(dat, _fittedAll, SigIm);
	}
}
void BetaGrpPEX::update(const Grp &dat, const Qgrp &q, const SigmaI &SigIm, const Grp &muPr, const Qgrp &qPr, const SigmaI &SigIp){
	gsl_blas_dsymm(CblasRight, CblasLower, 1.0, SigIm.getMat(), _A.getMat(), 0.0, _tSigIAt);
	
#pragma omp parallel for num_threads(_nThr)
	for (int iEl = 0; iEl < _theta.size(); iEl++) {
		_theta[iEl]->update(dat, q, SigIm, muPr, qPr[iEl], SigIp, _rV[omp_get_thread_num()]);
	}
	
	_updateFitted();
	_finishFitted();
	_updateAfitted();
	if (_lowLevel) {
		_A.update(dat, _fittedAll, q, SigIm, *_lowLevel);
	}
	else {
		_A.update(dat, _fittedAll, SigIm);
	}
}

/*
 *	BetaGrpPC methods
 */


BetaGrpPC::BetaGrpPC(const Grp &rsp, const string &predFlNam, const string &evFlNam, const size_t &Npred, RanIndex &up, const int &nThr){
	if (Npred != up.getNtot()) {
		cerr << "ERROR: number of predictors " << Npred << " is not equal to the number of elements " << up.getNtot() << " in the prior index in BetaGrpFt initialization." << endl;
		exit(-1);
	}
	
	gsl_matrix_free(_fittedAll);
	gsl_matrix_free(_Xmat);
	_fittedEach.resize(Npred);
	_fittedAll  = gsl_matrix_calloc(rsp.Ndata(), rsp.phenD());
	_nThr       = nThr;
	_Xmat       = gsl_matrix_alloc(rsp.Ndata(), Npred);
	gsl_matrix_free(_valueMat);
	_valueMat   = gsl_matrix_alloc(Npred, rsp.phenD());
	_lowLevel   = 0;
	_upLevel    = &up;
	
	if (_nThr > 1) {
		const gsl_rng_type *T = gsl_rng_mt19937;
		_rV.resize(_nThr);
		vector<gsl_rng *>::iterator rIt = _rV.begin();
		++rIt;
		for (; rIt != _rV.end(); ++rIt) {
			unsigned long seed = gsl_rng_get(_rV[0]);
			*rIt = gsl_rng_alloc(T);
			gsl_rng_set(*rIt, seed);
		}
		
	}
	
	gsl_matrix *Sig = gsl_matrix_alloc(rsp.phenD(), rsp.phenD());
	gsl_matrix_set_identity(Sig);
	gsl_vector *ev = gsl_vector_alloc(Npred);
	
	_theta.resize(Npred);
	
	FILE *prdIn = fopen(predFlNam.c_str(), "r");
	if (prdIn == NULL) {
		cerr << "ERROR: cannot open file " << predFlNam << " in BetaGrpPC" << endl;
		exit(-1);
	}
	gsl_matrix_fread(prdIn, _Xmat);
	fclose(prdIn);
	// no centering, since eigenvectors are centered by definition
	
	FILE *evIn = fopen(evFlNam.c_str(), "r");
	if (evIn == NULL) {
		cerr << "ERROR: cannot open file " << evFlNam << " in BetaGrpPC" << endl;
		exit(-1);
	}
	gsl_vector_fread(evIn, ev);
	fclose(evIn);
	
	for (size_t IpcCol = 0; IpcCol < ev->size; IpcCol++) { // scale the columns of the PC vector matrix by square root of eigenvalues
		gsl_vector_view pcCol = gsl_matrix_column(_Xmat, IpcCol);
		gsl_vector_scale(&pcCol.vector, sqrt(gsl_vector_get(ev, IpcCol)));
	}
	
	gsl_matrix *yCtr = gsl_matrix_alloc(rsp.dMat()->size1, rsp.dMat()->size2);
	colCenter(rsp.dMat(), yCtr); // approximating a regression with an intercept for initial values (not quite right, but close enough for initialization)
	for (size_t Xj = 0; Xj < Npred; Xj++) {
		_fittedEach[Xj].resize(rsp.Ndata()*rsp.phenD());
		_theta[Xj] = new MVnormBetaFt(yCtr, _Xmat, Xj, _fittedEach[Xj], Sig, _rV[0], _upLevel->priorInd(Xj), _valueMat, Xj);
	}
	
	_updateFitted();
	
	gsl_matrix_free(Sig);
	gsl_matrix_free(yCtr);
	gsl_vector_free(ev);
}

BetaGrpPC::BetaGrpPC(const Grp &rsp, const string &predFlNam, const string &evFlNam, const size_t &Npred, RanIndex &low, RanIndex &up, const int &nThr){
	if (Npred != up.getNtot()) {
		cerr << "ERROR: number of predictors " << Npred << " is not equal to the number of elements " << up.getNtot() << " in the prior index in BetaGrpFt initialization." << endl;
		exit(-1);
	}
	else if (low.getNtot() != rsp.dMat()->size1){
		cerr << "ERROR: number of elements " << low.getNgrp() << " in the replication index is not equal to the number of rows " << rsp.dMat()->size1 << " in the response matrix in BetaGrpFt initialization." << endl;
		exit(-1);
	}
	
	gsl_matrix_free(_fittedAll);
	gsl_matrix_free(_Xmat);
	_fittedEach.resize(Npred);
	_fittedAll  = gsl_matrix_calloc(low.getNgrp(), rsp.phenD());
	_nThr       = nThr;
	_Xmat       = gsl_matrix_alloc(low.getNtot(), Npred);
	gsl_matrix_free(_valueMat);
	_valueMat   = gsl_matrix_alloc(Npred, rsp.phenD());
	_lowLevel   = &low;
	_upLevel    = &up;
	
	if (_nThr > 1) {
		const gsl_rng_type *T = gsl_rng_mt19937;
		_rV.resize(_nThr);
		vector<gsl_rng *>::iterator rIt = _rV.begin();
		++rIt;
		for (; rIt != _rV.end(); ++rIt) {
			unsigned long seed = gsl_rng_get(_rV[0]);
			*rIt = gsl_rng_alloc(T);
			gsl_rng_set(*rIt, seed);
		}
		
	}
	
	gsl_matrix *Sig = gsl_matrix_alloc(rsp.phenD(), rsp.phenD());
	gsl_matrix_set_identity(Sig);
	gsl_vector *ev = gsl_vector_alloc(Npred);
	
	_theta.resize(Npred);
	
	FILE *prdIn = fopen(predFlNam.c_str(), "r");
	if (prdIn == NULL) {
		cerr << "ERROR: cannot open file " << predFlNam << " in BetaGrpPC" << endl;
		exit(-1);
	}
	gsl_vector *tmpRow = gsl_vector_alloc(Npred);
	for (size_t iGrp = 0; iGrp < _lowLevel->getNgrp(); iGrp++) {
		gsl_vector_fread(prdIn, tmpRow);
		for (vector<size_t>::iterator grpIt = (*_lowLevel)[iGrp].begin(); grpIt != (*_lowLevel)[iGrp].end(); ++grpIt) {
			gsl_matrix_set_row(_Xmat, *grpIt, tmpRow);
		}
	}
	gsl_vector_free(tmpRow);
	fclose(prdIn);
	
	FILE *evIn = fopen(evFlNam.c_str(), "r");
	if (evIn == NULL) {
		cerr << "ERROR: cannot open file " << evFlNam << " in BetaGrpPC" << endl;
		exit(-1);
	}
	gsl_vector_fread(evIn, ev);
	fclose(evIn);
	
	for (size_t IpcCol = 0; IpcCol < ev->size; IpcCol++) { // scale the columns of the PC vector matrix by square root of eigenvalues
		gsl_vector_view pcCol = gsl_matrix_column(_Xmat, IpcCol);
		gsl_vector_scale(&pcCol.vector, sqrt(gsl_vector_get(ev, IpcCol)));
	}
	
	gsl_matrix *y = gsl_matrix_alloc(_Xmat->size1, rsp.dMat()->size2);
	colCenter(rsp.dMat(), y); // approximating a regression with an intercept for initial values (not quite right, but close enough for initialization)
	
	for (size_t Xj = 0; Xj < Npred; Xj++) {
		_fittedEach[Xj].resize(low.getNtot()*rsp.phenD());
		_theta[Xj] = new MVnormBetaFt(y, _Xmat, Xj, _fittedEach[Xj], Sig, _rV[0], _upLevel->priorInd(Xj), _valueMat, Xj);
	}
	
	_updateFitted();
	
	gsl_matrix_free(Sig);
	gsl_matrix_free(y);
	gsl_vector_free(ev);
}

BetaGrpPC::BetaGrpPC(const Grp &rsp, const string &predFlNam, const string &evFlNam, const size_t &Npred, RanIndex &up, const string &outFlNam, const int &nThr){
	if (Npred != up.getNtot()) {
		cerr << "ERROR: number of predictors " << Npred << " is not equal to the number of elements " << up.getNtot() << " in the prior index in BetaGrpFt initialization." << endl;
		exit(-1);
	}

	gsl_matrix_free(_fittedAll);
	gsl_matrix_free(_Xmat);
	_fittedEach.resize(Npred);
	_fittedAll  = gsl_matrix_calloc(rsp.Ndata(), rsp.phenD());
	_nThr       = nThr;
	_Xmat       = gsl_matrix_alloc(rsp.Ndata(), Npred);
	gsl_matrix_free(_valueMat);
	_valueMat   = gsl_matrix_alloc(Npred, rsp.phenD());
	_lowLevel   = 0;
	_upLevel    = &up;
	_outFlNam   = outFlNam;
	remove(_outFlNam.c_str());
	
	if (_nThr > 1) {
		const gsl_rng_type *T = gsl_rng_mt19937;
		_rV.resize(_nThr);
		vector<gsl_rng *>::iterator rIt = _rV.begin();
		++rIt;
		for (; rIt != _rV.end(); ++rIt) {
			unsigned long seed = gsl_rng_get(_rV[0]);
			*rIt = gsl_rng_alloc(T);
			gsl_rng_set(*rIt, seed);
		}
		
	}
	
	gsl_matrix *Sig = gsl_matrix_alloc(rsp.phenD(), rsp.phenD());
	gsl_matrix_set_identity(Sig);
	gsl_vector *ev = gsl_vector_alloc(Npred);
	
	_theta.resize(Npred);
	
	FILE *prdIn = fopen(predFlNam.c_str(), "r");
	if (prdIn == NULL) {
		cerr << "ERROR: cannot open file " << predFlNam << " in BetaGrpPC" << endl;
		exit(-1);
	}
	gsl_matrix_fread(prdIn, _Xmat);
	fclose(prdIn);
	// no centering, since eigenvectors are centered by definition
	
	FILE *evIn = fopen(evFlNam.c_str(), "r");
	if (evIn == NULL) {
		cerr << "ERROR: cannot open file " << evFlNam << " in BetaGrpPC" << endl;
		exit(-1);
	}
	gsl_vector_fread(evIn, ev);
	fclose(evIn);
	
	for (size_t IpcCol = 0; IpcCol < ev->size; IpcCol++) { // scale the columns of the PC vector matrix by square root of eigenvalues
		gsl_vector_view pcCol = gsl_matrix_column(_Xmat, IpcCol);
		gsl_vector_scale(&pcCol.vector, sqrt(gsl_vector_get(ev, IpcCol)));
	}
	
	gsl_matrix *yCtr = gsl_matrix_alloc(rsp.dMat()->size1, rsp.dMat()->size2);
	colCenter(rsp.dMat(), yCtr); // approximating a regression with an intercept for initial values (not quite right, but close enough for initialization)
	for (size_t Xj = 0; Xj < Npred; Xj++) {
		_fittedEach[Xj].resize(rsp.Ndata()*rsp.phenD());
		_theta[Xj] = new MVnormBetaFt(yCtr, _Xmat, Xj, _fittedEach[Xj], Sig, _rV[0], _upLevel->priorInd(Xj), _valueMat, Xj);
	}
	
	_updateFitted();
	
	gsl_matrix_free(Sig);
	gsl_matrix_free(yCtr);
	gsl_vector_free(ev);
}

BetaGrpPC::BetaGrpPC(const Grp &rsp, const string &predFlNam, const string &evFlNam, const size_t &Npred, RanIndex &low, RanIndex &up, const string &outFlNam, const int &nThr){
	if (Npred != up.getNtot()) {
		cerr << "ERROR: number of predictors " << Npred << " is not equal to the number of elements " << up.getNtot() << " in the prior index in BetaGrpFt initialization." << endl;
		exit(-1);
	}
	else if (low.getNtot() != rsp.dMat()->size1){
		cerr << "ERROR: number of elements " << low.getNgrp() << " in the replication index is not equal to the number of rows " << rsp.dMat()->size1 << " in the response matrix in BetaGrpFt initialization." << endl;
		exit(-1);
	}
	
	gsl_matrix_free(_fittedAll);
	gsl_matrix_free(_Xmat);
	_fittedEach.resize(Npred);
	_fittedAll  = gsl_matrix_calloc(low.getNgrp(), rsp.phenD());
	_nThr       = nThr;
	_Xmat       = gsl_matrix_alloc(low.getNtot(), Npred);
	gsl_matrix_free(_valueMat);
	_valueMat   = gsl_matrix_alloc(Npred, rsp.phenD());
	_lowLevel   = &low;
	_upLevel    = &up;
	_outFlNam   = outFlNam;
	remove(_outFlNam.c_str());
	
	if (_nThr > 1) {
		const gsl_rng_type *T = gsl_rng_mt19937;
		_rV.resize(_nThr);
		vector<gsl_rng *>::iterator rIt = _rV.begin();
		++rIt;
		for (; rIt != _rV.end(); ++rIt) {
			unsigned long seed = gsl_rng_get(_rV[0]);
			*rIt = gsl_rng_alloc(T);
			gsl_rng_set(*rIt, seed);
		}
		
	}
	
	gsl_matrix *Sig = gsl_matrix_alloc(rsp.phenD(), rsp.phenD());
	gsl_matrix_set_identity(Sig);
	gsl_vector *ev = gsl_vector_alloc(Npred);
	
	_theta.resize(Npred);
	
	FILE *prdIn = fopen(predFlNam.c_str(), "r");
	if (prdIn == NULL) {
		cerr << "ERROR: cannot open file " << predFlNam << " in BetaGrpPC" << endl;
		exit(-1);
	}
	gsl_vector *tmpRow = gsl_vector_alloc(Npred);
	for (size_t iGrp = 0; iGrp < _lowLevel->getNgrp(); iGrp++) {
		gsl_vector_fread(prdIn, tmpRow);
		for (vector<size_t>::iterator grpIt = (*_lowLevel)[iGrp].begin(); grpIt != (*_lowLevel)[iGrp].end(); ++grpIt) {
			gsl_matrix_set_row(_Xmat, *grpIt, tmpRow);
		}
	}
	gsl_vector_free(tmpRow);
	fclose(prdIn);
	
	FILE *evIn = fopen(evFlNam.c_str(), "r");
	if (evIn == NULL) {
		cerr << "ERROR: cannot open file " << evFlNam << " in BetaGrpPC" << endl;
		exit(-1);
	}
	gsl_vector_fread(evIn, ev);
	fclose(evIn);
	
	for (size_t IpcCol = 0; IpcCol < ev->size; IpcCol++) { // scale the columns of the PC vector matrix by square root of eigenvalues
		gsl_vector_view pcCol = gsl_matrix_column(_Xmat, IpcCol);
		gsl_vector_scale(&pcCol.vector, sqrt(gsl_vector_get(ev, IpcCol)));
	}
	
	gsl_matrix *y = gsl_matrix_alloc(_Xmat->size1, rsp.dMat()->size2);
	colCenter(rsp.dMat(), y); // approximating a regression with an intercept for initial values (not quite right, but close enough for initialization)
	
	for (size_t Xj = 0; Xj < Npred; Xj++) {
		_fittedEach[Xj].resize(low.getNtot()*rsp.phenD());
		_theta[Xj] = new MVnormBetaFt(y, _Xmat, Xj, _fittedEach[Xj], Sig, _rV[0], _upLevel->priorInd(Xj), _valueMat, Xj);
	}
	
	_updateFitted();
	
	gsl_matrix_free(Sig);
	gsl_matrix_free(y);
	gsl_vector_free(ev);
}

BetaGrpPC::BetaGrpPC(const BetaGrpPC &bG){
	gsl_matrix_free(_fittedAll);
	gsl_matrix_free(_Xmat);
	gsl_matrix_free(_valueMat);
	
	_fittedAll = gsl_matrix_alloc((bG._fittedAll)->size1, (bG._fittedAll)->size2);
	gsl_matrix_memcpy(_fittedAll, bG._fittedAll);
	
	_fittedEach = bG._fittedEach;
	_lowLevel   = bG._lowLevel;
	_upLevel    = bG._upLevel;
	_nThr       = bG._nThr;
	
	if (_nThr > 1) {
		const gsl_rng_type *T = gsl_rng_mt19937;
		_rV.resize(_nThr);
		vector<gsl_rng *>::iterator rIt = _rV.begin();
		++rIt;
		for (; rIt != _rV.end(); ++rIt) {
			unsigned long seed = gsl_rng_get(_rV[0]);
			*rIt = gsl_rng_alloc(T);
			gsl_rng_set(*rIt, seed);
		}
		
	}
	
	_Xmat = gsl_matrix_alloc((bG._Xmat)->size1, (bG._Xmat)->size2);
	gsl_matrix_memcpy(_Xmat, bG._Xmat);
	
	_valueMat = gsl_matrix_alloc((bG._valueMat)->size1, (bG._valueMat)->size2);
	gsl_matrix_memcpy(_valueMat, bG._valueMat);
	
	_theta.resize(_valueMat->size1);
	for (size_t iVrw = 0; iVrw < _valueMat->size1; iVrw++) {
		_theta[iVrw] = new MVnormMu(_valueMat, iVrw, *(bG._theta[iVrw])->down(), *(bG._theta[iVrw])->up());
	}
	
}
BetaGrpPC & BetaGrpPC::operator=(const BetaGrpPC &bG){
	gsl_matrix_free(_fittedAll);
	gsl_matrix_free(_Xmat);
	gsl_matrix_free(_valueMat);
	
	_fittedAll = gsl_matrix_alloc((bG._fittedAll)->size1, (bG._fittedAll)->size2);
	gsl_matrix_memcpy(_fittedAll, bG._fittedAll);
	
	_fittedEach = bG._fittedEach;
	_lowLevel   = bG._lowLevel;
	_upLevel    = bG._upLevel;
	_nThr       = bG._nThr;
	
	if (_nThr > 1) {
		const gsl_rng_type *T = gsl_rng_mt19937;
		_rV.resize(_nThr);
		vector<gsl_rng *>::iterator rIt = _rV.begin();
		++rIt;
		for (; rIt != _rV.end(); ++rIt) {
			unsigned long seed = gsl_rng_get(_rV[0]);
			*rIt = gsl_rng_alloc(T);
			gsl_rng_set(*rIt, seed);
		}
		
	}
	
	_Xmat = gsl_matrix_alloc((bG._Xmat)->size1, (bG._Xmat)->size2);
	gsl_matrix_memcpy(_Xmat, bG._Xmat);
	
	_valueMat = gsl_matrix_alloc((bG._valueMat)->size1, (bG._valueMat)->size2);
	gsl_matrix_memcpy(_valueMat, bG._valueMat);
	
	_theta.resize(_valueMat->size1);
	for (size_t iVrw = 0; iVrw < _valueMat->size1; iVrw++) {
		_theta[iVrw] = new MVnormMu(_valueMat, iVrw, *(bG._theta[iVrw])->down(), *(bG._theta[iVrw])->up());
	}
	
	return *this;
}

/*
 *	BetaGrpSnp methods
 */

BetaGrpSnp::BetaGrpSnp() : _numSaves(0.0), _nThr(1), _Npred(1), _priorVar(0.0), MuGrp(){
	_Xmat    = gsl_matrix_calloc(1, 1);
	_Ystore  = gsl_matrix_calloc(1, 1);
}
BetaGrpSnp::BetaGrpSnp(const string &predFlNam, const string &outFlNam, const size_t &Ndat, const size_t &Npred, const size_t &d, const int &Nthr) : _numSaves(0.0), _nThr(Nthr), _Npred(Npred), _priorVar(0.0), _inPredFl(predFlNam), MuGrp(){
	_fakeFmat = gsl_matrix_calloc(Ndat, d);       // will be all zero, so we can use it for addition and subtraction without any consequences other than loss of efficiency
	_Ystore   = gsl_matrix_calloc(Ndat, d);
	
	_outFlNam = outFlNam;
	remove(_outFlNam.c_str());
	
	// the rest of the set-up happens before the results are dumped into a file by the dump() function
}
BetaGrpSnp::BetaGrpSnp(const string &predFlNam, const string &outFlNam, RanIndex &low, const size_t &Npred, const size_t &d, const int &Nthr) : _numSaves(0.0), _nThr(Nthr), _Npred(Npred), _priorVar(0.0), _inPredFl(predFlNam), MuGrp(){
	
	_fakeFmat = gsl_matrix_calloc(low.getNgrp(), d);       // will be all zero, so we can use it for addition and subtraction without any consequences other than loss of efficiency
	_lowLevel = &low;
	
	_Ystore  = gsl_matrix_calloc(low.getNtot(), d);
	
	_outFlNam = outFlNam;
	remove(_outFlNam.c_str());
	
	// the rest of the set-up happens before the results are dumped into a file by the dump() function
	
}

BetaGrpSnp::BetaGrpSnp(const string &predFlNam, const string &outFlNam, const size_t &Ndat, const size_t &Npred, const size_t &d, const int &Nthr, const double &prVar) : _numSaves(0.0), _nThr(Nthr), _Npred(Npred), _priorVar(prVar), _inPredFl(predFlNam), MuGrp(){
	_fakeFmat = gsl_matrix_calloc(Ndat, d);       // will be all zero, so we can use it for addition and subtraction without any consequences other than loss of efficiency
	_Ystore   = gsl_matrix_calloc(Ndat, d);
	
	_outFlNam = outFlNam;
	remove(_outFlNam.c_str());
	
	// the rest of the set-up happens before the results are dumped into a file by the dump() function
}
BetaGrpSnp::BetaGrpSnp(const string &predFlNam, const string &outFlNam, RanIndex &low, const size_t &Npred, const size_t &d, const int &Nthr, const double &prVar) : _numSaves(0.0), _nThr(Nthr), _Npred(Npred), _priorVar(prVar), _inPredFl(predFlNam), MuGrp(){
	
	_fakeFmat = gsl_matrix_calloc(low.getNgrp(), d);       // will be all zero, so we can use it for addition and subtraction without any consequences other than loss of efficiency
	_lowLevel = &low;
	
	_Ystore  = gsl_matrix_calloc(low.getNtot(), d);
	
	_outFlNam = outFlNam;
	remove(_outFlNam.c_str());
	
	// the rest of the set-up happens before the results are dumped into a file by the dump() function
	
}

BetaGrpSnp::BetaGrpSnp(const BetaGrpSnp &mG){
	gsl_matrix_free(_valueMat);
	_Ystore  = gsl_matrix_calloc((mG._Ystore)->size1, (mG._Ystore)->size2);
	
	_lowLevel = mG._lowLevel;
	_upLevel  = mG._upLevel;
	_nThr     = mG._nThr;
	_Npred    = mG._Npred;
	_inPredFl = mG._inPredFl;
	_priorVar = mG._priorVar;
	
	_Xmat = gsl_matrix_alloc((mG._Xmat)->size1, (mG._Xmat)->size2);
	gsl_matrix_memcpy(_Xmat, mG._Xmat);
	
	_valueMat = gsl_matrix_alloc((mG._valueMat)->size1, (mG._valueMat)->size2);
	gsl_matrix_memcpy(_valueMat, mG._valueMat);
	
}
BetaGrpSnp & BetaGrpSnp::operator=(const BetaGrpSnp &mG){
	gsl_matrix_free(_valueMat);
	_Ystore  = gsl_matrix_calloc((mG._Ystore)->size1, (mG._Ystore)->size2);
	
	_lowLevel = mG._lowLevel;
	_upLevel  = mG._upLevel;
	_nThr     = mG._nThr;
	_Npred    = mG._Npred;
	_inPredFl = mG._inPredFl;
	_priorVar = mG._priorVar;
	
	_Xmat = gsl_matrix_alloc((mG._Xmat)->size1, (mG._Xmat)->size2);
	gsl_matrix_memcpy(_Xmat, mG._Xmat);
	
	_valueMat = gsl_matrix_alloc((mG._valueMat)->size1, (mG._valueMat)->size2);
	gsl_matrix_memcpy(_valueMat, mG._valueMat);
	
	return *this;
}

BetaGrpSnp::~BetaGrpSnp(){
	if (_numSaves) {
		gsl_matrix_free(_Xmat);
	}
	gsl_matrix_free(_fakeFmat);
	gsl_matrix_free(_Ystore);
	
}


void BetaGrpSnp::dump(){
	if (_numSaves) {
		gsl_matrix_free(_valueMat);
		if (_lowLevel) {
			_Xmat = gsl_matrix_alloc(_lowLevel->getNgrp(), _Npred);
		}
		else {
			_Xmat = gsl_matrix_alloc(_Ystore->size1, _Npred);
		}
		_valueMat = gsl_matrix_calloc(_Npred, _Ystore->size2 + 1);  // the extra column will have the Hotelling-type statistic for all the traits
		
		FILE *prdIn = fopen(_inPredFl.c_str(), "r");
		if (prdIn == NULL) {
			cerr << "ERROR: cannot open file " << _inPredFl << " in BetaGrpSnp" << endl;
			exit(-1);
		}
		gsl_matrix_fread(prdIn, _Xmat);
		fclose(prdIn);
		colCenter(_Xmat); // essential to mimic an intercept; predictor should be centered by construction of the sampler
		
		gsl_matrix_scale(_Ystore, 1.0/_numSaves);
		if (_lowLevel) {
			gsl_matrix *tmpY = gsl_matrix_calloc(_Xmat->size1, _Ystore->size2);
			for (size_t iHi = 0; iHi < _lowLevel->getNgrp(); iHi++) {
				double scale = 1.0/static_cast<double>((*_lowLevel)[iHi].size());
				gsl_vector_view cvRow = gsl_matrix_row(tmpY, iHi);
				for (vector<size_t>::iterator lowIt = (*_lowLevel)[iHi].begin(); lowIt != (*_lowLevel)[iHi].end(); ++lowIt) {
					gsl_vector_view yRow = gsl_matrix_row(_Ystore, *lowIt);
					gsl_blas_daxpy(scale, &yRow.vector, &cvRow.vector); // calculating the Y mean on the fly
				}
			}
			gsl_matrix_free(_Ystore);
			_Ystore = gsl_matrix_alloc(tmpY->size1, tmpY->size2);
			gsl_matrix_memcpy(_Ystore, tmpY);
			gsl_matrix_free(tmpY);
			
			colCenter(_Ystore); // only strictly necessary when there is an unbalanced design
		}
		
#pragma omp parallel num_threads(_nThr)
		{
			double XtX;
			gsl_vector *bTmp  = gsl_vector_alloc(_Ystore->size2);
			gsl_matrix *S     = gsl_matrix_alloc(_Ystore->size2, _Ystore->size2);
			gsl_matrix *VW    = gsl_matrix_alloc(_Ystore->size2, _Ystore->size2);
			gsl_matrix *resd  = gsl_matrix_alloc(_Ystore->size1, _Ystore->size2);
			if (_priorVar) {  // if we have a prior variance, then we are doing approximate Bayes factors as God intended
#pragma omp for
				for (int iSnp = 0; iSnp < _Npred; iSnp++) {
					gsl_vector_view Xtmp = gsl_matrix_column(_Xmat, iSnp);
					gsl_blas_ddot(&Xtmp.vector, &Xtmp.vector, &XtX);
					if (XtX <= 1e-12) {
						gsl_vector_view vmRw = gsl_matrix_row(_valueMat, iSnp);
						gsl_vector_set_zero(&vmRw.vector);
						continue;
					}
					XtX = 1.0/XtX;
					
					gsl_blas_dgemv(CblasTrans, XtX, _Ystore, &Xtmp.vector, 0.0, bTmp);
					
					gsl_matrix_memcpy(resd, _Ystore);
					gsl_blas_dger(-1.0, &Xtmp.vector, bTmp, resd);  // subtracting the Xb
					
					gsl_blas_dsyrk(CblasLower, CblasTrans, XtX/static_cast<double>(_Ystore->size1 - 1), resd, 0.0, S);
					for (int iPh = 0; iPh < _Ystore->size2; iPh++) {
						double V  = gsl_matrix_get(S, iPh, iPh);
						double r  = _priorVar/(V + _priorVar);  // W/(V+W) of Wakefield (2007)
						double pV = 0.5*(log(1.0 - r) + gsl_pow_2(gsl_vector_get(bTmp, iPh))*r/V) ;
						gsl_matrix_set(_valueMat, iSnp, iPh, pV);
						
					}
					gsl_matrix_memcpy(VW, S);
					gsl_vector_view VWdiag = gsl_matrix_diagonal(VW);
					gsl_vector_add_constant(&VWdiag.vector, _priorVar);
					gsl_linalg_cholesky_decomp(S);
					gsl_linalg_cholesky_decomp(VW);
					double lnSrSdet  = 0.0;  // ln of the square root of the determinant
					double lnSrVWdet = 0.0;
					for (size_t iPh = 0; iPh < _Ystore->size2; iPh++) {
						lnSrSdet  += log(gsl_matrix_get(S, iPh, iPh));
						lnSrVWdet += log(gsl_matrix_get(VW, iPh, iPh));
					}
					gsl_linalg_cholesky_invert(S);
					gsl_linalg_cholesky_invert(VW);
					gsl_matrix_sub(S, VW);
					double m = mhl(bTmp, S);
					double pV = lnSrSdet - lnSrVWdet + 0.5*m;
					gsl_matrix_set(_valueMat, iSnp, _Ystore->size2, pV);
					
				}

			}
			else {
#pragma omp for
				for (int iSnp = 0; iSnp < _Npred; iSnp++) {
					gsl_vector_view Xtmp = gsl_matrix_column(_Xmat, iSnp);
					gsl_blas_ddot(&Xtmp.vector, &Xtmp.vector, &XtX);
					if (XtX <= 1e-12) {
						gsl_vector_view vmRw = gsl_matrix_row(_valueMat, iSnp);
						gsl_vector_set_zero(&vmRw.vector);
						continue;
					}
					XtX = 1.0/XtX;
					
					gsl_blas_dgemv(CblasTrans, XtX, _Ystore, &Xtmp.vector, 0.0, bTmp);
					
					gsl_matrix_memcpy(resd, _Ystore);
					gsl_blas_dger(-1.0, &Xtmp.vector, bTmp, resd);  // subtracting the Xb
					
					gsl_blas_dsyrk(CblasLower, CblasTrans, XtX, resd, 0.0, S);
					
					for (int iPh = 0; iPh < _Ystore->size2; iPh++) {
						double seB  = sqrt(gsl_matrix_get(S, iPh, iPh)/static_cast<double>(_Ystore->size1 - 1));
						double tVal = fabs(gsl_vector_get(bTmp, iPh))/seB;
						double pV   = -log10(gsl_cdf_tdist_Q(tVal, _Ystore->size1 - 1)*2.0);
						gsl_matrix_set(_valueMat, iSnp, iPh, pV);
					
					}
					
					
					gsl_linalg_cholesky_decomp(S);
					gsl_linalg_cholesky_invert(S);
					double m = mhl(bTmp, S);
					m = (static_cast<double>(_Ystore->size1) - static_cast<double>(_Ystore->size2))/static_cast<double>(_Ystore->size2)*m;
					double pV = -log10(gsl_cdf_fdist_Q(m, static_cast<double>(_Ystore->size2), static_cast<double>(_Ystore->size1) - static_cast<double>(_Ystore->size2)));
					
					gsl_matrix_set(_valueMat, iSnp, _Ystore->size2, pV);
					
				}
				
			}
			gsl_vector_free(bTmp);
			gsl_matrix_free(S);
			gsl_matrix_free(resd);
		}//end omp block
		
		FILE *btOut = fopen(_outFlNam.c_str(), "w");
		gsl_matrix_fwrite(btOut, _valueMat);
		fclose(btOut);
		

	}
	else {
		cout << "BetaGrpSnp: nothing to dump" << endl;
	}
}

void BetaGrpSnp::update(const Grp &dat, const SigmaI &SigIm){
	// Sigma ignored
	gsl_matrix_add(_Ystore, dat.fMat());
	_numSaves += 1.0;
}

/*
 * BetaGrpSnpCV methods
 */

BetaGrpSnpCV::BetaGrpSnpCV(const BetaGrpSnpCV &mG) {
	gsl_matrix_free(_valueMat);
	_Ystore  = gsl_matrix_calloc((mG._Ystore)->size1, (mG._Ystore)->size2);
	
	_lowLevel = mG._lowLevel;
	_upLevel  = mG._upLevel;
	_nThr     = mG._nThr;
	_Npred    = mG._Npred;
	_inPredFl = mG._inPredFl;
	_priorVar = mG._priorVar;
	
	_Xmat = gsl_matrix_alloc((mG._Xmat)->size1, (mG._Xmat)->size2);
	gsl_matrix_memcpy(_Xmat, mG._Xmat);
	
	_valueMat = gsl_matrix_alloc((mG._valueMat)->size1, (mG._valueMat)->size2);
	gsl_matrix_memcpy(_valueMat, mG._valueMat);
}
BetaGrpSnpCV & BetaGrpSnpCV::operator=(const BetaGrpSnpCV &mG){
	gsl_matrix_free(_valueMat);
	_Ystore  = gsl_matrix_calloc((mG._Ystore)->size1, (mG._Ystore)->size2);
	
	_lowLevel = mG._lowLevel;
	_upLevel  = mG._upLevel;
	_nThr     = mG._nThr;
	_Npred    = mG._Npred;
	_inPredFl = mG._inPredFl;
	_priorVar = mG._priorVar;
	
	_Xmat = gsl_matrix_alloc((mG._Xmat)->size1, (mG._Xmat)->size2);
	gsl_matrix_memcpy(_Xmat, mG._Xmat);
	
	_valueMat = gsl_matrix_alloc((mG._valueMat)->size1, (mG._valueMat)->size2);
	gsl_matrix_memcpy(_valueMat, mG._valueMat);
	
	return *this;
}


void BetaGrpSnpCV::dump(){
	if (_numSaves) {
		gsl_matrix_free(_valueMat);
		if (_lowLevel) {
			_Xmat = gsl_matrix_alloc(_lowLevel->getNgrp(), _Npred);
		}
		else {
			_Xmat = gsl_matrix_alloc(_Ystore->size1, _Npred);
		}
		_valueMat = gsl_matrix_calloc(_Npred, _Ystore->size2 + 1);  // the extra column will have the Hotelling-type statistic for all the traits
		
		FILE *prdIn = fopen(_inPredFl.c_str(), "r");
		if (prdIn == NULL) {
			cerr << "ERROR: cannot open file " << _inPredFl << " in BetaGrpSnpCV" << endl;
			exit(-1);
		}
		gsl_matrix_fread(prdIn, _Xmat);
		fclose(prdIn);
		colCenter(_Xmat); // essential to mimic an intercept; predictor should be centered by construction of the sampler
		
		gsl_matrix_scale(_Ystore, 1.0/_numSaves);
		if (_lowLevel) {
			gsl_matrix *tmpY = gsl_matrix_calloc(_Xmat->size1, _Ystore->size2);
			for (size_t iHi = 0; iHi < _lowLevel->getNgrp(); iHi++) {
				double scale = 1.0/static_cast<double>((*_lowLevel)[iHi].size());
				gsl_vector_view cvRow = gsl_matrix_row(tmpY, iHi);
				for (vector<size_t>::iterator lowIt = (*_lowLevel)[iHi].begin(); lowIt != (*_lowLevel)[iHi].end(); ++lowIt) {
					gsl_vector_view yRow = gsl_matrix_row(_Ystore, *lowIt);
					gsl_blas_daxpy(scale, &yRow.vector, &cvRow.vector); // calculating the Y mean on the fly
				}
			}
			gsl_matrix_free(_Ystore);
			_Ystore = gsl_matrix_alloc(tmpY->size1, tmpY->size2);
			gsl_matrix_memcpy(_Ystore, tmpY);
			gsl_matrix_free(tmpY);
			
			colCenter(_Ystore); // only strictly necessary when there is an unbalanced design
		}
		
#pragma omp parallel num_threads(_nThr)
		{
			double XtX;
			gsl_vector *bTmp  = gsl_vector_alloc(_Ystore->size2);
			gsl_matrix *S     = gsl_matrix_alloc(_Ystore->size2, _Ystore->size2);
			gsl_matrix *VW    = gsl_matrix_alloc(_Ystore->size2, _Ystore->size2);
			gsl_matrix *resd  = gsl_matrix_alloc(_Ystore->size1, _Ystore->size2);
			if (_priorVar) {  // if we have a prior variance, then we are doing approximate Bayes factors as God intended
#pragma omp for
				for (int iSnp = 0; iSnp < _Npred; iSnp++) {
					gsl_vector_view Xtmp = gsl_matrix_column(_Xmat, iSnp);
					gsl_blas_ddot(&Xtmp.vector, &Xtmp.vector, &XtX);
					if (XtX <= 1e-12) {
						gsl_vector_view vmRw = gsl_matrix_row(_valueMat, iSnp);
						gsl_vector_set_zero(&vmRw.vector);
						continue;
					}
					XtX = 1.0/XtX;
					
					gsl_blas_dgemv(CblasTrans, XtX, _Ystore, &Xtmp.vector, 0.0, bTmp);
					
					gsl_matrix_memcpy(resd, _Ystore);
					gsl_blas_dger(-1.0, &Xtmp.vector, bTmp, resd);  // subtracting the Xb
					
					gsl_blas_dsyrk(CblasLower, CblasTrans, XtX/static_cast<double>(_Ystore->size1 - 1), resd, 0.0, S);
					gsl_matrix_memcpy(VW, S);
					gsl_vector_view VWdiag = gsl_matrix_diagonal(VW);
					gsl_vector_add_constant(&VWdiag.vector, _priorVar);
					gsl_linalg_cholesky_decomp(S);
					gsl_linalg_cholesky_decomp(VW);
					double lnSrSdet  = 0.0;  // ln of the square root of the determinant
					double lnSrVWdet = 0.0;
					for (size_t iPh = 0; iPh < _Ystore->size2; iPh++) {
						lnSrSdet  += log(gsl_matrix_get(S, iPh, iPh));
						lnSrVWdet += log(gsl_matrix_get(VW, iPh, iPh));
					}
					gsl_linalg_cholesky_invert(S);
					gsl_linalg_cholesky_invert(VW);
					
					for (int iPh = 0; iPh < _Ystore->size2; iPh++) {
						double invV  = gsl_matrix_get(S, iPh, iPh);
						double r  = _priorVar * gsl_matrix_get(VW, iPh, iPh);  // W/(V+W) of Wakefield (2007)
						double pV = 0.5*(log(1.0 - r) + gsl_pow_2(gsl_vector_get(bTmp, iPh))*r * invV) ;
						gsl_matrix_set(_valueMat, iSnp, iPh, pV);
						
					}
					gsl_matrix_sub(S, VW);
					double m = mhl(bTmp, S);
					double pV = lnSrSdet - lnSrVWdet + 0.5*m;
					gsl_matrix_set(_valueMat, iSnp, _Ystore->size2, pV);
					
				}
				
			}
			else {
#pragma omp for
				for (int iSnp = 0; iSnp < _Npred; iSnp++) {
					gsl_vector_view Xtmp = gsl_matrix_column(_Xmat, iSnp);
					gsl_blas_ddot(&Xtmp.vector, &Xtmp.vector, &XtX);
					if (XtX <= 1e-12) {
						gsl_vector_view vmRw = gsl_matrix_row(_valueMat, iSnp);
						gsl_vector_set_zero(&vmRw.vector);
						continue;
					}
					XtX = 1.0/XtX;
					
					gsl_blas_dgemv(CblasTrans, XtX, _Ystore, &Xtmp.vector, 0.0, bTmp);
					
					gsl_matrix_memcpy(resd, _Ystore);
					gsl_blas_dger(-1.0, &Xtmp.vector, bTmp, resd);  // subtracting the Xb
					
					gsl_blas_dsyrk(CblasLower, CblasTrans, XtX, resd, 0.0, S);
					gsl_linalg_cholesky_decomp(S);
					gsl_linalg_cholesky_invert(S);
					
					for (int iPh = 0; iPh < _Ystore->size2; iPh++) {
						double invSeB  = sqrt(gsl_matrix_get(S, iPh, iPh) * static_cast<double>(_Ystore->size1 - 1));
						double tVal = fabs(gsl_vector_get(bTmp, iPh)) * invSeB;
						double pV   = -log10(gsl_cdf_tdist_Q(tVal, _Ystore->size1 - 1)*2.0);
						gsl_matrix_set(_valueMat, iSnp, iPh, pV);
						
					}
					
					
					double m = mhl(bTmp, S);
					m = (static_cast<double>(_Ystore->size1) - static_cast<double>(_Ystore->size2))/static_cast<double>(_Ystore->size2)*m;
					double pV = -log10(gsl_cdf_fdist_Q(m, static_cast<double>(_Ystore->size2), static_cast<double>(_Ystore->size1) - static_cast<double>(_Ystore->size2)));
					
					gsl_matrix_set(_valueMat, iSnp, _Ystore->size2, pV);
					
				}
				
			}
			gsl_vector_free(bTmp);
			gsl_matrix_free(S);
			gsl_matrix_free(resd);
		}//end omp block
		
		FILE *btOut = fopen(_outFlNam.c_str(), "w");
		gsl_matrix_fwrite(btOut, _valueMat);
		fclose(btOut);
		
		
	}
	else {
		cout << "BetaGrpSnpCV: nothing to dump" << endl;
	}
}

/*
 * BetaGrpPSR methods
 */


BetaGrpPSR::BetaGrpPSR(const BetaGrpPSR &mG){
	gsl_matrix_free(_valueMat);
	_Ystore  = gsl_matrix_calloc((mG._Ystore)->size1, (mG._Ystore)->size2);
	
	_lowLevel = mG._lowLevel;
	_upLevel  = mG._upLevel;
	_nThr     = mG._nThr;
	_Npred    = mG._Npred;
	_inPredFl = mG._inPredFl;
	_priorVar = mG._priorVar;
	
	_Xmat = gsl_matrix_alloc((mG._Xmat)->size1, (mG._Xmat)->size2);
	gsl_matrix_memcpy(_Xmat, mG._Xmat);
	
	_valueMat = gsl_matrix_alloc((mG._valueMat)->size1, (mG._valueMat)->size2);
	gsl_matrix_memcpy(_valueMat, mG._valueMat);
	
	_theta.resize(_valueMat->size1);
	for (size_t iVrw = 0; iVrw < _valueMat->size1; iVrw++) {
		delete _theta[iVrw];
		_theta[iVrw] = new MVnormMu(_valueMat, iVrw, *(mG._theta[iVrw])->down(), *(mG._theta[iVrw])->up());
	}
}

BetaGrpPSR & BetaGrpPSR::operator=(const BetaGrpPSR &mG){
	gsl_matrix_free(_valueMat);
	_Ystore  = gsl_matrix_calloc((mG._Ystore)->size1, (mG._Ystore)->size2);
	
	_lowLevel = mG._lowLevel;
	_upLevel  = mG._upLevel;
	_nThr     = mG._nThr;
	_Npred    = mG._Npred;
	_inPredFl = mG._inPredFl;
	_priorVar = mG._priorVar;
	
	_Xmat = gsl_matrix_alloc((mG._Xmat)->size1, (mG._Xmat)->size2);
	gsl_matrix_memcpy(_Xmat, mG._Xmat);
	
	_valueMat = gsl_matrix_alloc((mG._valueMat)->size1, (mG._valueMat)->size2);
	gsl_matrix_memcpy(_valueMat, mG._valueMat);
	
	_theta.resize(_valueMat->size1);
	for (size_t iVrw = 0; iVrw < _valueMat->size1; iVrw++) {
		delete _theta[iVrw];
		_theta[iVrw] = new MVnormMu(_valueMat, iVrw, *(mG._theta[iVrw])->down(), *(mG._theta[iVrw])->up());
	}
		
	return *this;
}


void BetaGrpPSR::dump(){
	if (_numSaves) {
		if (_Ystore->size2 == 1) {
			cerr << "WARNING: only one trait. No conditional regression in single-SNP analysis." << endl;
		}
		
		gsl_matrix_free(_valueMat);
		
		if (_lowLevel) {
			_Xmat = gsl_matrix_alloc(_lowLevel->getNgrp(), _Npred);
		}
		else {
			_Xmat = gsl_matrix_alloc(_Ystore->size1, _Npred);
		}
		_valueMat = gsl_matrix_calloc(_Npred, _Ystore->size2);  // no extra column.  Get the full multi-trait test through the BetaGrpSnp.dump()
		
		FILE *prdIn = fopen(_inPredFl.c_str(), "r");
		if (prdIn == NULL) {
			cerr << "ERROR: cannot open file " << _inPredFl << " in BetaGrpPSR" << endl;
			exit(-1);
		}
		gsl_matrix_fread(prdIn, _Xmat);
		fclose(prdIn);
		colCenter(_Xmat); // essential to mimic an intercept
		
		gsl_matrix_scale(_Ystore, 1.0/_numSaves);
		
		if (_lowLevel) {
			gsl_matrix *tmpY = gsl_matrix_calloc(_Xmat->size1, _Ystore->size2);
			for (size_t iHi = 0; iHi < _lowLevel->getNgrp(); iHi++) {
				double scale = 1.0/static_cast<double>((*_lowLevel)[iHi].size());
				gsl_vector_view cvRow = gsl_matrix_row(tmpY, iHi);
				for (vector<size_t>::iterator lowIt = (*_lowLevel)[iHi].begin(); lowIt != (*_lowLevel)[iHi].end(); ++lowIt) {
					gsl_vector_view yRow = gsl_matrix_row(_Ystore, *lowIt);
					gsl_blas_daxpy(scale, &yRow.vector, &cvRow.vector); // calculating the Y mean on the fly
				}
			}
			gsl_matrix_free(_Ystore);
			_Ystore = gsl_matrix_alloc(tmpY->size1, tmpY->size2);
			gsl_matrix_memcpy(_Ystore, tmpY);
			gsl_matrix_free(tmpY);
			
			colCenter(_Ystore);
		}
		
#pragma omp parallel num_threads(_nThr)
		{
			double sig;
			gsl_matrix *Xplus = gsl_matrix_alloc(_Ystore->size1, _Ystore->size2);
			gsl_matrix *XtX   = gsl_matrix_alloc(_Ystore->size2, _Ystore->size2);
			gsl_vector *bHat  = gsl_vector_alloc(_Ystore->size2);
			gsl_vector *Xty   = gsl_vector_alloc(_Ystore->size2);
			gsl_vector *y     = gsl_vector_alloc(_Ystore->size1);
			if (_priorVar) {  // if we have a prior variance, then we are doing approximate Bayes factors as God intended
#pragma omp for
				for (int iSnp = 0; iSnp < _Npred; iSnp++) {
					
					gsl_vector_view Xtmp = gsl_matrix_column(_Xmat, iSnp);
					for (size_t iTr = 0; iTr < _Ystore->size2; iTr++) {
						size_t curCol = 0;
						for (size_t jTr = 0; jTr < _Ystore->size2; jTr++) {
							gsl_vector_view yCol = gsl_matrix_column(_Ystore, jTr);
							if (jTr != iTr) {
								gsl_matrix_set_col(Xplus, curCol, &yCol.vector);
								curCol++;
							}
							else {
								gsl_vector_memcpy(y, &yCol.vector);  // copying it over (as opposed to doing a vector view) may not be the most efficient, but do it to be safe
							}
						}
						gsl_matrix_set_col(Xplus, Xplus->size2 - 1, &Xtmp.vector);
						gsl_blas_dsyrk(CblasLower, CblasTrans, 1.0, Xplus, 0.0, XtX);  // XtX
						if (gsl_matrix_get(XtX, Xplus->size2 - 1, Xplus->size2 - 1) <= 1e-12) {
							gsl_matrix_set(_valueMat, iSnp, iTr, 0.0);
							continue;
						}
						gsl_linalg_cholesky_decomp(XtX);
						gsl_linalg_cholesky_invert(XtX); // XtX^-1
						
						gsl_blas_dgemv(CblasTrans, 1.0, Xplus, y, 0.0, Xty); // Xty
						gsl_blas_dsymv(CblasLower, 1.0, XtX, Xty, 0.0, bHat); // XtX^-1Xty = bHat
						
						gsl_blas_dgemv(CblasNoTrans, -1.0, Xplus, bHat, 1.0, y);  // y is now the residual; only interested in the last element of bHat (the regression on X)
						gsl_blas_ddot(y, y, &sig);
						sig = sig*gsl_matrix_get(XtX, XtX->size1 - 1, XtX->size2 - 1)/static_cast<double>(_Ystore->size1 - _Ystore->size2);
						double r = _priorVar/(sig + _priorVar);  // W/(V+W) of Wakefield (2007)
						double pV = 0.5*(log(1.0 - r) + gsl_pow_2(gsl_vector_get(bHat, bHat->size - 1))*r/sig);
						gsl_matrix_set(_valueMat, iSnp, iTr, pV);
					}
					
				}
				
			}
			else {
#pragma omp for
				for (int iSnp = 0; iSnp < _Npred; iSnp++) {
					
					gsl_vector_view Xtmp = gsl_matrix_column(_Xmat, iSnp);
					for (size_t iTr = 0; iTr < _Ystore->size2; iTr++) {
						size_t curCol = 0;
						for (size_t jTr = 0; jTr < _Ystore->size2; jTr++) {
							gsl_vector_view yCol = gsl_matrix_column(_Ystore, jTr);
							if (jTr != iTr) {
								gsl_matrix_set_col(Xplus, curCol, &yCol.vector);
								curCol++;
							}
							else {
								gsl_vector_memcpy(y, &yCol.vector);  // copying it over (as opposed to doing a vector view) may not be the most efficient, but do it to be safe
							}
						}
						gsl_matrix_set_col(Xplus, Xplus->size2 - 1, &Xtmp.vector);
						gsl_blas_dsyrk(CblasLower, CblasTrans, 1.0, Xplus, 0.0, XtX);  // XtX
						if (gsl_matrix_get(XtX, Xplus->size2 - 1, Xplus->size2 - 1) <= 1e-12) {
							gsl_matrix_set(_valueMat, iSnp, iTr, 0.0);
							continue;
						}
						gsl_linalg_cholesky_decomp(XtX);
						gsl_linalg_cholesky_invert(XtX); // XtX^-1
						
						gsl_blas_dgemv(CblasTrans, 1.0, Xplus, y, 0.0, Xty); // Xty
						gsl_blas_dsymv(CblasLower, 1.0, XtX, Xty, 0.0, bHat); // XtX^-1Xty = bHat
						
						gsl_blas_dgemv(CblasNoTrans, -1.0, Xplus, bHat, 1.0, y);  // y is now the residual; only interested in the last element of bHat (the regression on X)
						sig = gsl_blas_dnrm2(y);
						sig = sig*sqrt(gsl_matrix_get(XtX, XtX->size1 - 1, XtX->size2 - 1))/sqrt(static_cast<double>(_Ystore->size1 - _Ystore->size2));
						double tVal = fabs(gsl_vector_get(bHat, bHat->size - 1))/sig;
						double pV   = -log10(gsl_cdf_tdist_Q(tVal, _Ystore->size1 - 1)*2.0);
						gsl_matrix_set(_valueMat, iSnp, iTr, pV);
					}
					
				}
				
			}
			gsl_vector_free(bHat);
			gsl_vector_free(Xty);
			gsl_vector_free(y);
			gsl_matrix_free(XtX);
			gsl_matrix_free(Xplus);
		}//end omp block
		
		FILE *btOut = fopen(_outFlNam.c_str(), "w");
		gsl_matrix_fwrite(btOut, _valueMat);
		fclose(btOut);
		
		
	}
	else {
		cout << "BetaGrpPSR: nothing to dump" << endl;
	}
}

/*
 *	BetaGrpSnpMiss methods
 */
BetaGrpSnpMiss::BetaGrpSnpMiss() : _numSaves(0.0), _nThr(1), _Npred(1), _priorVar(0.0), _absLab(-9.0), MuGrp() {
	_Ystore  = gsl_matrix_calloc(1, 1);
}

BetaGrpSnpMiss::BetaGrpSnpMiss(const string &predFlNam, const string &outFlNam, const size_t &Ndat, const size_t &Npred, const size_t &d, const int &Nthr, const double &absLab) : _numSaves(0.0), _nThr(Nthr), _Npred(Npred), _inPredFl(predFlNam), _priorVar(0.0), _absLab(absLab), MuGrp(){
	_Ystore  = gsl_matrix_calloc(Ndat, d);
	
	_fakeFmat = gsl_matrix_calloc(Ndat, d);     // will be all zero, so we can use it for addition and subtraction without any consequences other than loss of efficiency
	_lowLevel = 0;
	_outFlNam = outFlNam;
	remove(_outFlNam.c_str());
	
}
BetaGrpSnpMiss::BetaGrpSnpMiss(const string &predFlNam, const string &outFlNam, RanIndex &low, const size_t &Npred, const size_t &d, const int &Nthr, const double &absLab) : _numSaves(0.0), _nThr(Nthr), _Npred(Npred), _inPredFl(predFlNam), _priorVar(0.0), _absLab(absLab), MuGrp() {
	_Ystore  = gsl_matrix_calloc(low.getNtot(), d);
	
	_fakeFmat = gsl_matrix_calloc(low.getNgrp(), d);     // will be all zero, so we can use it for addition and subtraction without any consequences other than loss of efficiency
	_lowLevel = &low;
	_outFlNam = outFlNam;
	remove(_outFlNam.c_str());
}
BetaGrpSnpMiss::BetaGrpSnpMiss(const string &predFlNam, const string &outFlNam, const size_t &Ndat, const size_t &Npred, const size_t &d, const int &Nthr, const double &prVar, const double &absLab) : _numSaves(0.0), _nThr(Nthr), _Npred(Npred), _inPredFl(predFlNam), _priorVar(prVar), _absLab(absLab), MuGrp(){
	_Ystore  = gsl_matrix_calloc(Ndat, d);
	
	_fakeFmat = gsl_matrix_calloc(Ndat, d);     // will be all zero, so we can use it for addition and subtraction without any consequences other than loss of efficiency
	_lowLevel = 0;
	_outFlNam = outFlNam;
	remove(_outFlNam.c_str());
	
}
BetaGrpSnpMiss::BetaGrpSnpMiss(const string &predFlNam, const string &outFlNam, RanIndex &low, const size_t &Npred, const size_t &d, const int &Nthr, const double &prVar, const double &absLab) : _numSaves(0.0), _nThr(Nthr), _Npred(Npred), _inPredFl(predFlNam), _priorVar(prVar), _absLab(absLab), MuGrp() {
	_Ystore  = gsl_matrix_calloc(low.getNtot(), d);
	
	_fakeFmat = gsl_matrix_calloc(low.getNgrp(), d);     // will be all zero, so we can use it for addition and subtraction without any consequences other than loss of efficiency
	_lowLevel = &low;
	_outFlNam = outFlNam;
	remove(_outFlNam.c_str());
}

BetaGrpSnpMiss::BetaGrpSnpMiss(const BetaGrpSnpMiss &mG){
	gsl_matrix_free(_valueMat);
	_Ystore  = gsl_matrix_calloc((mG._Ystore)->size1, (mG._Ystore)->size2);
	
	_lowLevel = mG._lowLevel;
	_upLevel  = mG._upLevel;
	_nThr     = mG._nThr;
	_Npred    = mG._Npred;
	_absLab   = mG._absLab;
	_priorVar = mG._priorVar;
	_inPredFl = mG._inPredFl;
	
	_valueMat = gsl_matrix_alloc((mG._valueMat)->size1, (mG._valueMat)->size2);
	gsl_matrix_memcpy(_valueMat, mG._valueMat);
	
	
}
BetaGrpSnpMiss & BetaGrpSnpMiss::operator=(const BetaGrpSnpMiss &mG){
	gsl_matrix_free(_valueMat);
	_Ystore  = gsl_matrix_calloc((mG._Ystore)->size1, (mG._Ystore)->size2);
	
	_lowLevel = mG._lowLevel;
	_upLevel  = mG._upLevel;
	_nThr     = mG._nThr;
	_Npred    = mG._Npred;
	_absLab   = mG._absLab;
	_priorVar = mG._priorVar;
	_inPredFl = mG._inPredFl;
	
	_valueMat = gsl_matrix_alloc((mG._valueMat)->size1, (mG._valueMat)->size2);
	gsl_matrix_memcpy(_valueMat, mG._valueMat);
	
	
	return *this;
}

BetaGrpSnpMiss::~BetaGrpSnpMiss(){
	gsl_matrix_free(_fakeFmat);
	gsl_matrix_free(_Ystore);
}

void BetaGrpSnpMiss::dump(){
	if (_numSaves) {
		_Xmat.resize(_Npred);
		
		gsl_matrix_free(_valueMat);
		_valueMat = gsl_matrix_calloc(_Npred, _Ystore->size2 + 1);
		
		gsl_vector *tmpX = gsl_vector_alloc(_Npred);
		FILE *prdIn = fopen(_inPredFl.c_str(), "r");
		if (prdIn == NULL) {
			cerr << "ERROR: cannot open file " << _inPredFl << " in BetaGrpSnpMiss" << endl;
			exit(-1);
		}
		vector< vector<size_t> > presInd(_Npred);
		
		size_t Nrw;
		_lowLevel ? Nrw = _lowLevel->getNgrp() : Nrw = _Ystore->size1;
		
		for (size_t XiRw = 0; XiRw < Nrw; XiRw++) { // reading in one row (line of predictor) at a time so that two whole SNP matrix worth of stuff is not in memory at once;  remember that SNPs are saved in row-major
			gsl_vector_fread(prdIn, tmpX);
			for (size_t XjCl = 0; XjCl < _Npred; XjCl++) {
				if (gsl_vector_get(tmpX, XjCl) > _absLab) { // absLab labels the missing data, has to be smaller than the smallest real value
					presInd[XjCl].push_back(XiRw);
					_Xmat[XjCl].push_back(gsl_vector_get(tmpX, XjCl));
				}
			}
		}
		fclose(prdIn);
		gsl_vector_free(tmpX);
		
		gsl_matrix_scale(_Ystore, 1.0/_numSaves);
		if (_lowLevel) {
			gsl_matrix *tmpY = gsl_matrix_calloc(_lowLevel->getNgrp(), _Ystore->size2);
			for (size_t iHi = 0; iHi < _lowLevel->getNgrp(); iHi++) {
				double scale = 1.0/static_cast<double>((*_lowLevel)[iHi].size());
				gsl_vector_view cvRow = gsl_matrix_row(tmpY, iHi);
				for (vector<size_t>::iterator lowIt = (*_lowLevel)[iHi].begin(); lowIt != (*_lowLevel)[iHi].end(); ++lowIt) {
					gsl_vector_view yRow = gsl_matrix_row(_Ystore, *lowIt);
					gsl_blas_daxpy(scale, &yRow.vector, &cvRow.vector); // calculating the Y mean on the fly
				}
			}
			gsl_matrix_free(_Ystore);
			_Ystore = gsl_matrix_alloc(tmpY->size1, tmpY->size2);
			gsl_matrix_memcpy(_Ystore, tmpY);
			gsl_matrix_free(tmpY);
		}
		
		colCenter(_Ystore); // necessary for predictors with no missing data
#pragma omp parallel num_threads(_nThr)
		{
			double XtX;
			gsl_vector *bTmp  = gsl_vector_alloc(_Ystore->size2);
			gsl_matrix *S     = gsl_matrix_alloc(_Ystore->size2, _Ystore->size2);
			gsl_matrix *VW    = gsl_matrix_alloc(_Ystore->size2, _Ystore->size2);
			if (_priorVar) {  // if we have a prior variance, then we are doing approximate Bayes factors as God intended
#pragma omp for
				for (int iSnp = 0; iSnp < _Npred; iSnp++) {
					gsl_matrix *resd;
					if (presInd[iSnp].size() == 0) { // i.e., no data at all.  Shouldn't happen, but who knows
						resd = gsl_matrix_alloc(1, 1); // just so the freeing of the matrix later works
						gsl_vector_view pVrow = gsl_matrix_row(_valueMat, iSnp);
						gsl_vector_set_zero(&pVrow.vector);
						continue;
					}
					else if (presInd[iSnp].size() < _Ystore->size1) {  // i.e., some missing SNP data
						resd = gsl_matrix_alloc(presInd[iSnp].size(), _Ystore->size2);
						int ind = 0;
						for (vector<size_t>::const_iterator it = presInd[iSnp].begin(); it != presInd[iSnp].end(); ++it) {
							gsl_vector_view rspRw = gsl_matrix_row(_Ystore, *it);
							gsl_matrix_set_row(resd, ind, &rspRw.vector);
							ind++;
						}
						colCenter(resd);
						
					}
					else {
						resd = gsl_matrix_alloc(_Ystore->size1, _Ystore->size2);
						gsl_matrix_memcpy(resd, _Ystore);
					}
					gsl_vector_view Xtmp = gsl_vector_view_array(_Xmat[iSnp].data(), _Xmat[iSnp].size());
					gsl_blas_ddot(&Xtmp.vector, &Xtmp.vector, &XtX);
					if (XtX <= 1e-12) {
						gsl_vector_view vmRw = gsl_matrix_row(_valueMat, iSnp);
						gsl_vector_set_zero(&vmRw.vector);
						continue;
					}
					XtX = 1.0/XtX;
					
					gsl_blas_dgemv(CblasTrans, XtX, resd, &Xtmp.vector, 0.0, bTmp);
					
					gsl_blas_dger(-1.0, &Xtmp.vector, bTmp, resd);  // subtracting the Xb
					
					gsl_blas_dsyrk(CblasLower, CblasTrans, XtX/static_cast<double>(presInd[iSnp].size() - 1), resd, 0.0, S);
					
					for (int iPh = 0; iPh < _Ystore->size2; iPh++) {
						double V  = gsl_matrix_get(S, iPh, iPh);
						double r  = _priorVar/(V + _priorVar);  // W/(V+W) of Wakefield (2007)
						double pV = 0.5*(log(1.0 - r) + gsl_pow_2(gsl_vector_get(bTmp, iPh))*r/V) ;
						gsl_matrix_set(_valueMat, iSnp, iPh, pV);
						
					}
					gsl_matrix_memcpy(VW, S);
					gsl_vector_view VWdiag = gsl_matrix_diagonal(VW);
					gsl_vector_add_constant(&VWdiag.vector, _priorVar);
					gsl_linalg_cholesky_decomp(S);
					gsl_linalg_cholesky_decomp(VW);
					double lnSrSdet = 0.0;  // ln of the square root of the determinant
					double lnSrVWdet = 0.0;
					for (size_t iPh = 0; iPh < _Ystore->size2; iPh++) {
						lnSrSdet  += log(gsl_matrix_get(S, iPh, iPh));
						lnSrVWdet += log(gsl_matrix_get(VW, iPh, iPh));
					}
					gsl_linalg_cholesky_invert(S);
					gsl_linalg_cholesky_invert(VW);
					gsl_matrix_sub(S, VW);
					double m = mhl(bTmp, S);
					double pV = lnSrSdet - lnSrVWdet + 0.5*m;
					gsl_matrix_set(_valueMat, iSnp, _Ystore->size2, pV);
					
					gsl_matrix_free(resd);
				}
				
			}
			else {
#pragma omp for
				for (int iSnp = 0; iSnp < _Npred; iSnp++) {
					gsl_matrix *resd;
					if (presInd[iSnp].size() == 0) { // i.e., no data at all.  Shouldn't happen, but who knows
						resd = gsl_matrix_alloc(1, 1); // just so the freeing of the matrix later works
						gsl_vector_view pVrow = gsl_matrix_row(_valueMat, iSnp);
						gsl_vector_set_zero(&pVrow.vector);
						continue;
					}
					else if (presInd[iSnp].size() < _Ystore->size1) {  // i.e., some missing data for this SNP
						resd = gsl_matrix_alloc(presInd[iSnp].size(), _Ystore->size2);
						int ind = 0;
						for (vector<size_t>::const_iterator it = presInd[iSnp].begin(); it != presInd[iSnp].end(); ++it) {
							gsl_vector_view rspRw = gsl_matrix_row(_Ystore, *it);
							gsl_matrix_set_row(resd, ind, &rspRw.vector);
							ind++;
						}
						colCenter(resd);
						
					}
					else {
						resd = gsl_matrix_alloc(_Ystore->size1, _Ystore->size2);
						gsl_matrix_memcpy(resd, _Ystore);
					}
					gsl_vector_view Xtmp = gsl_vector_view_array(_Xmat[iSnp].data(), _Xmat[iSnp].size());
					gsl_blas_ddot(&Xtmp.vector, &Xtmp.vector, &XtX);
					if (XtX <= 1e-12) {
						gsl_vector_view vmRw = gsl_matrix_row(_valueMat, iSnp);
						gsl_vector_set_zero(&vmRw.vector);
						continue;
					}
					XtX = 1.0/XtX;
					
					gsl_blas_dgemv(CblasTrans, XtX, resd, &Xtmp.vector, 0.0, bTmp);
					gsl_blas_dger(-1.0, &Xtmp.vector, bTmp, resd);  // subtracting the Xb
					
					gsl_blas_dsyrk(CblasLower, CblasTrans, XtX, resd, 0.0, S);
					
					for (int iPh = 0; iPh < _Ystore->size2; iPh++) {
						double seB  = sqrt(gsl_matrix_get(S, iPh, iPh)/static_cast<double>(presInd[iSnp].size() - 1));
						double tVal = fabs(gsl_vector_get(bTmp, iPh))/seB;
						double pV   = -log10(gsl_cdf_tdist_Q(tVal, _Ystore->size1 - 1)*2.0);
						gsl_matrix_set(_valueMat, iSnp, iPh, pV);
						
					}
					
					
					gsl_linalg_cholesky_decomp(S);
					gsl_linalg_cholesky_invert(S);
					double m = mhl(bTmp, S);
					m = (static_cast<double>(presInd[iSnp].size()) - static_cast<double>(_Ystore->size2))/static_cast<double>(_Ystore->size2)*m;
					double pV = -log10(gsl_cdf_fdist_Q(m, static_cast<double>(_Ystore->size2), static_cast<double>(presInd[iSnp].size()) - static_cast<double>(_Ystore->size2)));
					
					gsl_matrix_set(_valueMat, iSnp, _Ystore->size2, pV);
					gsl_matrix_free(resd);
				}
				
			}
			gsl_vector_free(bTmp);
			gsl_matrix_free(S);
			
		}//end omp block
		
		FILE *btOut = fopen(_outFlNam.c_str(), "w");
		gsl_matrix_fwrite(btOut, _valueMat);
		fclose(btOut);
	}
	else {
		cout << "BetaGrpSnpMiss: nothing to dump" << endl;
	}
}

void BetaGrpSnpMiss::update(const Grp &dat, const SigmaI &SigIm){
	gsl_matrix_add(_Ystore, dat.fMat());
	_numSaves += 1.0;
}

/*
 * BetaGrpSnpMissCV methods
 */

BetaGrpSnpMissCV::BetaGrpSnpMissCV(const BetaGrpSnpMissCV &mG){
	gsl_matrix_free(_valueMat);
	_Ystore  = gsl_matrix_calloc((mG._Ystore)->size1, (mG._Ystore)->size2);
	
	_lowLevel = mG._lowLevel;
	_upLevel  = mG._upLevel;
	_nThr     = mG._nThr;
	_Npred    = mG._Npred;
	_absLab   = mG._absLab;
	_priorVar = mG._priorVar;
	_inPredFl = mG._inPredFl;
	
	_valueMat = gsl_matrix_alloc((mG._valueMat)->size1, (mG._valueMat)->size2);
	gsl_matrix_memcpy(_valueMat, mG._valueMat);
	
}
BetaGrpSnpMissCV & BetaGrpSnpMissCV::operator=(const BetaGrpSnpMissCV &mG){
	gsl_matrix_free(_valueMat);
	_Ystore  = gsl_matrix_calloc((mG._Ystore)->size1, (mG._Ystore)->size2);
	
	_lowLevel = mG._lowLevel;
	_upLevel  = mG._upLevel;
	_nThr     = mG._nThr;
	_Npred    = mG._Npred;
	_absLab   = mG._absLab;
	_priorVar = mG._priorVar;
	_inPredFl = mG._inPredFl;
	
	_valueMat = gsl_matrix_alloc((mG._valueMat)->size1, (mG._valueMat)->size2);
	gsl_matrix_memcpy(_valueMat, mG._valueMat);
	
	
	return *this;
}

void BetaGrpSnpMissCV::dump(){
	if (_numSaves) {
		_Xmat.resize(_Npred);
		
		gsl_matrix_free(_valueMat);
		_valueMat = gsl_matrix_calloc(_Npred, _Ystore->size2 + 1);
		
		gsl_vector *tmpX = gsl_vector_alloc(_Npred);
		FILE *prdIn = fopen(_inPredFl.c_str(), "r");
		if (prdIn == NULL) {
			cerr << "ERROR: cannot open file " << _inPredFl << " in BetaGrpMissCV" << endl;
			exit(-1);
		}
		vector< vector<size_t> > presInd(_Npred);
		
		size_t Nrw;
		_lowLevel ? Nrw = _lowLevel->getNgrp() : Nrw = _Ystore->size1;
		
		for (size_t XiRw = 0; XiRw < Nrw; XiRw++) { // reading in one row (line of predictor) at a time so that two whole SNP matrix worth of stuff is not in memory at once;  remember that SNPs are saved in row-major
			gsl_vector_fread(prdIn, tmpX);
			for (size_t XjCl = 0; XjCl < _Npred; XjCl++) {
				if (gsl_vector_get(tmpX, XjCl) > _absLab) { // absLab labels the missing data, has to be smaller than the smallest real value
					presInd[XjCl].push_back(XiRw);
					_Xmat[XjCl].push_back(gsl_vector_get(tmpX, XjCl));
				}
			}
		}
		fclose(prdIn);
		gsl_vector_free(tmpX);
		
		gsl_matrix_scale(_Ystore, 1.0/_numSaves);
		if (_lowLevel) {
			gsl_matrix *tmpY = gsl_matrix_calloc(_lowLevel->getNgrp(), _Ystore->size2);
			for (size_t iHi = 0; iHi < _lowLevel->getNgrp(); iHi++) {
				double scale = 1.0/static_cast<double>((*_lowLevel)[iHi].size());
				gsl_vector_view cvRow = gsl_matrix_row(tmpY, iHi);
				for (vector<size_t>::iterator lowIt = (*_lowLevel)[iHi].begin(); lowIt != (*_lowLevel)[iHi].end(); ++lowIt) {
					gsl_vector_view yRow = gsl_matrix_row(_Ystore, *lowIt);
					gsl_blas_daxpy(scale, &yRow.vector, &cvRow.vector); // calculating the Y mean on the fly
				}
			}
			gsl_matrix_free(_Ystore);
			_Ystore = gsl_matrix_alloc(tmpY->size1, tmpY->size2);
			gsl_matrix_memcpy(_Ystore, tmpY);
			gsl_matrix_free(tmpY);
		}
		
		colCenter(_Ystore); // necessary for predictors with no missing data
#pragma omp parallel num_threads(_nThr)
		{
			double XtX;
			gsl_vector *bTmp  = gsl_vector_alloc(_Ystore->size2);
			gsl_matrix *S     = gsl_matrix_alloc(_Ystore->size2, _Ystore->size2);
			gsl_matrix *VW    = gsl_matrix_alloc(_Ystore->size2, _Ystore->size2);
			if (_priorVar) {  // if we have a prior variance, then we are doing approximate Bayes factors as God intended
#pragma omp for
				for (int iSnp = 0; iSnp < _Npred; iSnp++) {
					gsl_matrix *resd;
					if (presInd[iSnp].size() == 0) { // i.e., no data at all.  Shouldn't happen, but who knows
						resd = gsl_matrix_alloc(1, 1); // just so the freeing of the matrix later works
						gsl_vector_view pVrow = gsl_matrix_row(_valueMat, iSnp);
						gsl_vector_set_zero(&pVrow.vector);
						continue;
					}
					else if (presInd[iSnp].size() < _Ystore->size1) {  // i.e., some missing SNP data
						resd = gsl_matrix_alloc(presInd[iSnp].size(), _Ystore->size2);
						int ind = 0;
						for (vector<size_t>::const_iterator it = presInd[iSnp].begin(); it != presInd[iSnp].end(); ++it) {
							gsl_vector_view rspRw = gsl_matrix_row(_Ystore, *it);
							gsl_matrix_set_row(resd, ind, &rspRw.vector);
							ind++;
						}
						colCenter(resd);
						
					}
					else {
						resd = gsl_matrix_alloc(_Ystore->size1, _Ystore->size2);
						gsl_matrix_memcpy(resd, _Ystore);
					}
					gsl_vector_view Xtmp = gsl_vector_view_array(_Xmat[iSnp].data(), _Xmat[iSnp].size());
					gsl_blas_ddot(&Xtmp.vector, &Xtmp.vector, &XtX);
					if (XtX <= 1e-12) {
						gsl_vector_view vmRw = gsl_matrix_row(_valueMat, iSnp);
						gsl_vector_set_zero(&vmRw.vector);
						continue;
					}
					XtX = 1.0/XtX;
					
					gsl_blas_dgemv(CblasTrans, XtX, resd, &Xtmp.vector, 0.0, bTmp);
					
					gsl_blas_dger(-1.0, &Xtmp.vector, bTmp, resd);  // subtracting the Xb
					
					gsl_blas_dsyrk(CblasLower, CblasTrans, XtX/static_cast<double>(presInd[iSnp].size() - 1), resd, 0.0, S);
					gsl_matrix_memcpy(VW, S);
					gsl_vector_view VWdiag = gsl_matrix_diagonal(VW);
					gsl_vector_add_constant(&VWdiag.vector, _priorVar);
					gsl_linalg_cholesky_decomp(S);
					gsl_linalg_cholesky_decomp(VW);
					
					double lnSrSdet = 0.0;  // ln of the square root of the determinant
					double lnSrVWdet = 0.0;
					for (size_t iPh = 0; iPh < _Ystore->size2; iPh++) {
						lnSrSdet  += log(gsl_matrix_get(S, iPh, iPh));
						lnSrVWdet += log(gsl_matrix_get(VW, iPh, iPh));
					}
					gsl_linalg_cholesky_invert(S);
					gsl_linalg_cholesky_invert(VW);
					
					for (int iPh = 0; iPh < _Ystore->size2; iPh++) {
						double invV  = gsl_matrix_get(S, iPh, iPh);
						double r  = _priorVar * gsl_matrix_get(VW, iPh, iPh);  // W/(V+W) of Wakefield (2007)
						double pV = 0.5*(log(1.0 - r) + gsl_pow_2(gsl_vector_get(bTmp, iPh))*r*invV) ;
						gsl_matrix_set(_valueMat, iSnp, iPh, pV);
						
					}
					gsl_matrix_sub(S, VW);
					double m = mhl(bTmp, S);
					double pV = lnSrSdet - lnSrVWdet + 0.5*m;
					gsl_matrix_set(_valueMat, iSnp, _Ystore->size2, pV);
					
					gsl_matrix_free(resd);
				}
				
			}
			else {
#pragma omp for
				for (int iSnp = 0; iSnp < _Npred; iSnp++) {
					gsl_matrix *resd;
					if (presInd[iSnp].size() == 0) { // i.e., no data at all.  Shouldn't happen, but who knows
						resd = gsl_matrix_alloc(1, 1); // just so the freeing of the matrix later works
						gsl_vector_view pVrow = gsl_matrix_row(_valueMat, iSnp);
						gsl_vector_set_zero(&pVrow.vector);
						continue;
					}
					else if (presInd[iSnp].size() < _Ystore->size1) {  // i.e., some missing data for this SNP
						resd = gsl_matrix_alloc(presInd[iSnp].size(), _Ystore->size2);
						int ind = 0;
						for (vector<size_t>::const_iterator it = presInd[iSnp].begin(); it != presInd[iSnp].end(); ++it) {
							gsl_vector_view rspRw = gsl_matrix_row(_Ystore, *it);
							gsl_matrix_set_row(resd, ind, &rspRw.vector);
							ind++;
						}
						colCenter(resd);
						
					}
					else {
						resd = gsl_matrix_alloc(_Ystore->size1, _Ystore->size2);
						gsl_matrix_memcpy(resd, _Ystore);
					}
					gsl_vector_view Xtmp = gsl_vector_view_array(_Xmat[iSnp].data(), _Xmat[iSnp].size());
					gsl_blas_ddot(&Xtmp.vector, &Xtmp.vector, &XtX);
					if (XtX <= 1e-12) {
						gsl_vector_view vmRw = gsl_matrix_row(_valueMat, iSnp);
						gsl_vector_set_zero(&vmRw.vector);
						continue;
					}
					XtX = 1.0/XtX;
					
					gsl_blas_dgemv(CblasTrans, XtX, resd, &Xtmp.vector, 0.0, bTmp);
					gsl_blas_dger(-1.0, &Xtmp.vector, bTmp, resd);  // subtracting the Xb
					
					gsl_blas_dsyrk(CblasLower, CblasTrans, XtX, resd, 0.0, S);
					gsl_linalg_cholesky_decomp(S);
					gsl_linalg_cholesky_invert(S);
					
					for (int iPh = 0; iPh < _Ystore->size2; iPh++) {
						double invSeB  = sqrt(gsl_matrix_get(S, iPh, iPh) * static_cast<double>(presInd[iSnp].size() - 1));
						double tVal = fabs(gsl_vector_get(bTmp, iPh)) * invSeB;
						double pV   = -log10(gsl_cdf_tdist_Q(tVal, _Ystore->size1 - 1)*2.0);
						gsl_matrix_set(_valueMat, iSnp, iPh, pV);
						
					}
					
					
					double m = mhl(bTmp, S);
					m = (static_cast<double>(presInd[iSnp].size()) - static_cast<double>(_Ystore->size2))/static_cast<double>(_Ystore->size2)*m;
					double pV = -log10(gsl_cdf_fdist_Q(m, static_cast<double>(_Ystore->size2), static_cast<double>(presInd[iSnp].size()) - static_cast<double>(_Ystore->size2)));
					
					gsl_matrix_set(_valueMat, iSnp, _Ystore->size2, pV);
					gsl_matrix_free(resd);
				}
				
			}
			gsl_vector_free(bTmp);
			gsl_matrix_free(S);
			
		}//end omp block
		
		FILE *btOut = fopen(_outFlNam.c_str(), "w");
		gsl_matrix_fwrite(btOut, _valueMat);
		fclose(btOut);
	}
	else {
		cout << "BetaGrpSnpMissCV: nothing to dump" << endl;
	}
}


/*
 *  BetaGrpPSRmiss methods
 */

BetaGrpPSRmiss::BetaGrpPSRmiss(const BetaGrpPSRmiss &mG){
	gsl_matrix_free(_valueMat);
	_Ystore  = gsl_matrix_calloc((mG._Ystore)->size1, (mG._Ystore)->size2);
	
	_lowLevel = mG._lowLevel;
	_upLevel  = mG._upLevel;
	_nThr     = mG._nThr;
	_Npred    = mG._Npred;
	_absLab   = mG._absLab;
	_priorVar = mG._priorVar;
	_inPredFl = mG._inPredFl;
	
	_valueMat = gsl_matrix_alloc((mG._valueMat)->size1, (mG._valueMat)->size2);
	gsl_matrix_memcpy(_valueMat, mG._valueMat);
	
	_theta.resize(_valueMat->size1);
	for (size_t iVrw = 0; iVrw < _valueMat->size1; iVrw++) {
		_theta[iVrw] = new MVnormMu(_valueMat, iVrw, *(mG._theta[iVrw])->down(), *(mG._theta[iVrw])->up());
	}
		
}
BetaGrpPSRmiss & BetaGrpPSRmiss::operator=(const BetaGrpPSRmiss &mG){
	gsl_matrix_free(_valueMat);
	_Ystore  = gsl_matrix_calloc((mG._Ystore)->size1, (mG._Ystore)->size2);
	
	_lowLevel = mG._lowLevel;
	_upLevel  = mG._upLevel;
	_nThr     = mG._nThr;
	_Npred    = mG._Npred;
	_absLab   = mG._absLab;
	_priorVar = mG._priorVar;
	_inPredFl = mG._inPredFl;
	
	_valueMat = gsl_matrix_alloc((mG._valueMat)->size1, (mG._valueMat)->size2);
	gsl_matrix_memcpy(_valueMat, mG._valueMat);
	
	_theta.resize(_valueMat->size1);
	for (size_t iVrw = 0; iVrw < _valueMat->size1; iVrw++) {
		_theta[iVrw] = new MVnormMu(_valueMat, iVrw, *(mG._theta[iVrw])->down(), *(mG._theta[iVrw])->up());
	}
		
	return *this;
}

void BetaGrpPSRmiss::dump(){
	if (_numSaves) {
		if (_Ystore->size2 == 1) {
			cerr << "WARNING: only one trait. No conditional regression in single-SNP analysis." << endl;
		}
		
		_Xmat.resize(_Npred);
		
		gsl_matrix_free(_valueMat);
		_valueMat = gsl_matrix_calloc(_Npred, _Ystore->size2 + 1);
		
		gsl_vector *tmpX = gsl_vector_alloc(_Npred);
		FILE *prdIn = fopen(_inPredFl.c_str(), "r");
		if (prdIn == NULL) {
			cerr << "ERROR: cannot open file " << _inPredFl << " in BetaGrpPSRmiss" << endl;
			exit(-1);
		}
		vector< vector<size_t> > presInd(_Npred);
		
		size_t Nrw;
		_lowLevel ? Nrw = _lowLevel->getNgrp() : Nrw = _Ystore->size1;
		
		for (size_t XiRw = 0; XiRw < Nrw; XiRw++) { // reading in one row (line of predictor) at a time so that two whole SNP matrix worth of stuff is not in memory at once;  remember that SNPs are saved in row-major
			gsl_vector_fread(prdIn, tmpX);
			for (size_t XjCl = 0; XjCl < _Npred; XjCl++) {
				if (gsl_vector_get(tmpX, XjCl) > _absLab) { // absLab labels the missing data, has to be smaller than the smallest real value
					presInd[XjCl].push_back(XiRw);
					_Xmat[XjCl].push_back(gsl_vector_get(tmpX, XjCl));
				}
			}
		}
		fclose(prdIn);
		gsl_vector_free(tmpX);
		
		gsl_matrix_scale(_Ystore, 1.0/_numSaves);
		if (_lowLevel) {
			gsl_matrix *tmpY = gsl_matrix_calloc(_lowLevel->getNgrp(), _Ystore->size2);
			for (size_t iHi = 0; iHi < _lowLevel->getNgrp(); iHi++) {
				double scale = 1.0/static_cast<double>((*_lowLevel)[iHi].size());
				gsl_vector_view cvRow = gsl_matrix_row(tmpY, iHi);
				for (vector<size_t>::iterator lowIt = (*_lowLevel)[iHi].begin(); lowIt != (*_lowLevel)[iHi].end(); ++lowIt) {
					gsl_vector_view yRow = gsl_matrix_row(_Ystore, *lowIt);
					gsl_blas_daxpy(scale, &yRow.vector, &cvRow.vector); // calculating the Y mean on the fly
				}
			}
			gsl_matrix_free(_Ystore);
			_Ystore = gsl_matrix_alloc(tmpY->size1, tmpY->size2);
			gsl_matrix_memcpy(_Ystore, tmpY);
			gsl_matrix_free(tmpY);
		}
		
		colCenter(_Ystore); // necessary for presidictors with no missing data
#pragma omp parallel num_threads(_nThr)
		{
			double sig;
			gsl_matrix *XtX   = gsl_matrix_alloc(_Ystore->size2, _Ystore->size2);
			gsl_vector *bHat  = gsl_vector_alloc(_Ystore->size2);
			gsl_vector *Xty   = gsl_vector_alloc(_Ystore->size2);
			if (_priorVar) {  // if we have a prior variance, then we are doing approximate Bayes factors as God intended
#pragma omp for
				for (int iSnp = 0; iSnp < _Npred; iSnp++) {
					
					gsl_vector_view Xtmp = gsl_vector_view_array(_Xmat[iSnp].data(), _Xmat[iSnp].size());
					if (presInd[iSnp].size() == 0) {    // i.e., no data at all.  Shouldn't happen, but who knows
						gsl_vector_view pVrow = gsl_matrix_row(_valueMat, iSnp);
						gsl_vector_set_zero(&pVrow.vector);
						continue;
					}
					else if (presInd[iSnp].size() < _Ystore->size1) { // i.e., some missing data
						gsl_matrix *Xplus = gsl_matrix_alloc(presInd[iSnp].size(), _Ystore->size2);
						gsl_vector *y     = gsl_vector_alloc(presInd[iSnp].size());
						
						for (size_t iTr = 0; iTr < _Ystore->size2; iTr++) {
							size_t curCol = 0;
							for (size_t jTr = 0; jTr < _Ystore->size2; jTr++) {
								if (jTr != iTr) {
									int ind = 0;
									for (vector<size_t>::const_iterator it = presInd[iSnp].begin(); it != presInd[iSnp].end(); ++it) {
										gsl_matrix_set(Xplus, ind, curCol, gsl_matrix_get(_Ystore, *it, jTr)) ;
										ind++;
									}
									curCol++;
								}
								else {
									int ind = 0;
									for (vector<size_t>::const_iterator it = presInd[iSnp].begin(); it != presInd[iSnp].end(); ++it) {
										gsl_vector_set(y, ind, gsl_matrix_get(_Ystore, *it, jTr)) ;
										ind++;
									}
								}
							}
							gsl_matrix_set_col(Xplus, Xplus->size2 - 1, &Xtmp.vector);
							colCenter(Xplus);
							vecCenter(y);
							gsl_blas_dsyrk(CblasLower, CblasTrans, 1.0, Xplus, 0.0, XtX);  // XtX
							if (gsl_matrix_get(XtX, Xplus->size2 - 1, Xplus->size2 - 1) <= 1e-12) {
								gsl_matrix_set(_valueMat, iSnp, iTr, 0.0);
								continue;
							}
							
							gsl_linalg_cholesky_decomp(XtX);
							gsl_linalg_cholesky_invert(XtX); // XtX^-1
							
							gsl_blas_dgemv(CblasTrans, 1.0, Xplus, y, 0.0, Xty); // Xty
							gsl_blas_dsymv(CblasLower, 1.0, XtX, Xty, 0.0, bHat); // XtX^-1Xty = bHat
							
							gsl_blas_dgemv(CblasNoTrans, -1.0, Xplus, bHat, 1.0, y);  // y is now the residual; only interested in the last element of bHat (the regression on X)
							gsl_blas_ddot(y, y, &sig);
							sig = sig*gsl_matrix_get(XtX, XtX->size1 - 1, XtX->size2 - 1)/static_cast<double>(presInd[iSnp].size() - _Ystore->size2);
							double r = _priorVar/(sig + _priorVar);  // W/(V+W) of Wakefield (2007)
							double pV = 0.5*(log(1.0 - r) + gsl_pow_2(gsl_vector_get(bHat, bHat->size - 1))*r/sig);
							gsl_matrix_set(_valueMat, iSnp, iTr, pV);
							
						}
						gsl_vector_free(y);
						gsl_matrix_free(Xplus);
					}
					else {
						gsl_matrix *Xplus = gsl_matrix_alloc(_Ystore->size1, _Ystore->size2);
						gsl_vector *y     = gsl_vector_alloc(_Ystore->size1);
						for (size_t iTr = 0; iTr < _Ystore->size2; iTr++) {
							size_t curCol = 0;
							for (size_t jTr = 0; jTr < _Ystore->size2; jTr++) {
								gsl_vector_view yCol = gsl_matrix_column(_Ystore, jTr);
								if (jTr != iTr) {
									gsl_matrix_set_col(Xplus, curCol, &yCol.vector);
									curCol++;
								}
								else {
									gsl_vector_memcpy(y, &yCol.vector);  // copying it over (as opposed to doing a vector view) may not be the most efficient, but do it to be safe
								}
							}
							gsl_matrix_set_col(Xplus, Xplus->size2 - 1, &Xtmp.vector);
							gsl_blas_dsyrk(CblasLower, CblasTrans, 1.0, Xplus, 0.0, XtX);  // XtX
							if (gsl_matrix_get(XtX, Xplus->size2 - 1, Xplus->size2 - 1) <= 1e-12) {
								gsl_matrix_set(_valueMat, iSnp, iTr, 0.0);
								continue;
							}
							gsl_linalg_cholesky_decomp(XtX);
							gsl_linalg_cholesky_invert(XtX); // XtX^-1
							
							gsl_blas_dgemv(CblasTrans, 1.0, Xplus, y, 0.0, Xty); // Xty
							gsl_blas_dsymv(CblasLower, 1.0, XtX, Xty, 0.0, bHat); // XtX^-1Xty = bHat
							
							gsl_blas_dgemv(CblasNoTrans, -1.0, Xplus, bHat, 1.0, y);  // y is now the residual; only interested in the last element of bHat (the regression on X)
							gsl_blas_ddot(y, y, &sig);
							sig = sig*gsl_matrix_get(XtX, XtX->size1 - 1, XtX->size2 - 1)/static_cast<double>(_Ystore->size1 - _Ystore->size2);
							double r = _priorVar/(sig + _priorVar);  // W/(V+W) of Wakefield (2007)
							double pV = 0.5*(log(1.0 - r) + gsl_pow_2(gsl_vector_get(bHat, bHat->size - 1))*r/sig);
							gsl_matrix_set(_valueMat, iSnp, iTr, pV);
							
						}
						gsl_vector_free(y);
						gsl_matrix_free(Xplus);
					}
					
				}
				
			}
			else {
#pragma omp for
				for (int iSnp = 0; iSnp < _Npred; iSnp++) {
					
					gsl_vector_view Xtmp = gsl_vector_view_array(_Xmat[iSnp].data(), _Xmat[iSnp].size());
					if (presInd[iSnp].size() == 0) {    // i.e., no data at all.  Shouldn't happen, but who knows
						gsl_vector_view pVrow = gsl_matrix_row(_valueMat, iSnp);
						gsl_vector_set_zero(&pVrow.vector);
						continue;
					}
					else if (presInd[iSnp].size() < _Ystore->size1) { // i.e., some missing data
						gsl_matrix *Xplus = gsl_matrix_alloc(presInd[iSnp].size(), _Ystore->size2);
						gsl_vector *y     = gsl_vector_alloc(presInd[iSnp].size());
						
						for (size_t iTr = 0; iTr < _Ystore->size2; iTr++) {
							size_t curCol = 0;
							for (size_t jTr = 0; jTr < _Ystore->size2; jTr++) {
								if (jTr != iTr) {
									int ind = 0;
									for (vector<size_t>::const_iterator it = presInd[iSnp].begin(); it != presInd[iSnp].end(); ++it) {
										gsl_matrix_set(Xplus, ind, curCol, gsl_matrix_get(_Ystore, *it, jTr)) ;
										ind++;
									}
									curCol++;
								}
								else {
									int ind = 0;
									for (vector<size_t>::const_iterator it = presInd[iSnp].begin(); it != presInd[iSnp].end(); ++it) {
										gsl_vector_set(y, ind, gsl_matrix_get(_Ystore, *it, jTr)) ;
										ind++;
									}
								}
							}
							gsl_matrix_set_col(Xplus, Xplus->size2 - 1, &Xtmp.vector);
							colCenter(Xplus);
							vecCenter(y);
							gsl_blas_dsyrk(CblasLower, CblasTrans, 1.0, Xplus, 0.0, XtX);  // XtX
							if (gsl_matrix_get(XtX, Xplus->size2 - 1, Xplus->size2 - 1) <= 1e-12) {
								gsl_matrix_set(_valueMat, iSnp, iTr, 0.0);
								continue;
							}
							gsl_linalg_cholesky_decomp(XtX);
							gsl_linalg_cholesky_invert(XtX); // XtX^-1
							
							gsl_blas_dgemv(CblasTrans, 1.0, Xplus, y, 0.0, Xty); // Xty
							gsl_blas_dsymv(CblasLower, 1.0, XtX, Xty, 0.0, bHat); // XtX^-1Xty = bHat
							
							gsl_blas_dgemv(CblasNoTrans, -1.0, Xplus, bHat, 1.0, y);  // y is now the residual; only interested in the last element of bHat (the regression on X)
							sig = gsl_blas_dnrm2(y);
							sig = sig*sqrt(gsl_matrix_get(XtX, XtX->size1 - 1, XtX->size2 - 1))/sqrt(static_cast<double>(_Ystore->size1 - _Ystore->size2));
							double tVal = fabs(gsl_vector_get(bHat, bHat->size - 1))/sig;
							double pV   = -log10(gsl_cdf_tdist_Q(tVal, _Ystore->size1 - 1)*2.0);
							gsl_matrix_set(_valueMat, iSnp, iTr, pV);
							
						}
						gsl_vector_free(y);
						gsl_matrix_free(Xplus);
					}
					else {
						gsl_matrix *Xplus = gsl_matrix_alloc(presInd[iSnp].size(), _Ystore->size2);
						gsl_vector *y     = gsl_vector_alloc(presInd[iSnp].size());
						for (size_t iTr = 0; iTr < _Ystore->size2; iTr++) {
							size_t curCol = 0;
							for (size_t jTr = 0; jTr < _Ystore->size2; jTr++) {
								gsl_vector_view yCol = gsl_matrix_column(_Ystore, jTr);
								if (jTr != iTr) {
									gsl_matrix_set_col(Xplus, curCol, &yCol.vector);
									curCol++;
								}
								else {
									gsl_vector_memcpy(y, &yCol.vector);  // copying it over (as opposed to doing a vector view) may not be the most efficient, but do it to be safe
								}
							}
							gsl_matrix_set_col(Xplus, Xplus->size2 - 1, &Xtmp.vector);
							gsl_blas_dsyrk(CblasLower, CblasTrans, 1.0, Xplus, 0.0, XtX);  // XtX
							if (gsl_matrix_get(XtX, Xplus->size2 - 1, Xplus->size2 - 1) <= 1e-12) {
								gsl_matrix_set(_valueMat, iSnp, iTr, 0.0);
								continue;
							}
							gsl_linalg_cholesky_decomp(XtX);
							gsl_linalg_cholesky_invert(XtX); // XtX^-1
							
							gsl_blas_dgemv(CblasTrans, 1.0, Xplus, y, 0.0, Xty); // Xty
							gsl_blas_dsymv(CblasLower, 1.0, XtX, Xty, 0.0, bHat); // XtX^-1Xty = bHat
							
							gsl_blas_dgemv(CblasNoTrans, -1.0, Xplus, bHat, 1.0, y);  // y is now the residual; only interested in the last element of bHat (the regression on X)
							sig = gsl_blas_dnrm2(y);
							sig = sig*sqrt(gsl_matrix_get(XtX, XtX->size1 - 1, XtX->size2 - 1))/sqrt(static_cast<double>(_Ystore->size1 - _Ystore->size2));
							double tVal = fabs(gsl_vector_get(bHat, bHat->size - 1))/sig;
							double pV   = -log10(gsl_cdf_tdist_Q(tVal, _Ystore->size1 - 1)*2.0);
							gsl_matrix_set(_valueMat, iSnp, iTr, pV);
						}
						gsl_vector_free(y);
						gsl_matrix_free(Xplus);
					}
					
				}
				
			}
			gsl_vector_free(bHat);
			gsl_vector_free(Xty);
			gsl_matrix_free(XtX);
		}//end omp block
		
		FILE *btOut = fopen(_outFlNam.c_str(), "w");
		gsl_matrix_fwrite(btOut, _valueMat);
		fclose(btOut);
		
		
	}
	else {
		cout << "BetaGrpPSRmiss: nothing to dump" << endl;
	}
}

/*
 *	BetaGrpBVSR methods
 */

BetaGrpBVSR::BetaGrpBVSR(const Grp &y, const SigmaI &SigI, const string &predFlNam, const double &Nmul, const double &rSqMax, RanIndex &up, const string &outFlNam, const int &nThr) : BetaGrpFt(){
	if (up.getNgrp() != 2) {
		cerr << "ERROR: variable selection index &up has " << up.getNgrp() << " elements instead of 2!" << endl;
		exit(1);
	}
	
	gsl_matrix_free(_fittedAll);
	gsl_matrix_free(_valueMat);
	_nThr  = nThr;
	_Xmat  = gsl_matrix_alloc(y.Ndata(), up.getNtot());
	_tmpXb = gsl_matrix_calloc(y.Ndata(), y.phenD());
	
	if (_nThr > 1) {
		const gsl_rng_type *T = gsl_rng_mt19937;
		_rV.resize(_nThr);
		vector<gsl_rng *>::iterator rIt = _rV.begin();
		++rIt;
		for (; rIt != _rV.end(); ++rIt) {
			unsigned long seed = gsl_rng_get(_rV[0]);
			*rIt = gsl_rng_alloc(T);
			gsl_rng_set(*rIt, seed);
		}
		
	}
	
	_upLevel = &up;
	_outFlNam = outFlNam;
	remove(_outFlNam.c_str());
	
	FILE *prdIn = fopen(predFlNam.c_str(), "r");
	if (prdIn == NULL) {
		cerr << "ERROR: cannot open file " << predFlNam << " in BetaGrpBVSR" << endl;
		exit(-1);
	}
	gsl_matrix_fread(prdIn, _Xmat);
	fclose(prdIn);
	
	gsl_matrix *yCtr = gsl_matrix_alloc(y.Ndata(), y.phenD());
	colCenter(y.dMat(), yCtr);
	
	gsl_vector *XtX = gsl_vector_alloc(_Xmat->size2);
	gsl_permutation *allP = gsl_permutation_alloc(_Xmat->size2);
	
	_rankPred(yCtr, SigI, XtX, allP);
	
	size_t Npck = ceil(_Xmat->size1*Nmul);
	size_t Nsel;
	if (Nmul < 1.0) {
		Nsel = floor(gsl_ran_flat(_rV[0], 1.0, static_cast<double>(Npck)));
	}
	else {
```
