## Supplemental methods and scripts for "Low additive genetic variation in a trait under selection in domesticated rice": MuGen.h

```
/*
 * Copyright (c) 2015 Anthony J. Greenberg
 *
 * Redistribution and use in source and binary forms, with or without modification, are permitted provided that the following conditions are met:
 *
 * 1. Redistributions of source code must retain the above copyright notice, this list of conditions and the following disclaimer.
 *
 * 2. Redistributions in binary form must reproduce the above copyright notice, this list of conditions and the following disclaimer in the documentation and/or other materials provided with the distribution.
 *
 * 3. Neither the name of the copyright holder nor the names of its contributors may be used to endorse or promote products derived from this software without specific prior written permission.
 *
 * THIS SOFTWARE IS PROVIDED BY THE COPYRIGHT HOLDERS AND CONTRIBUTORS "AS IS" AND ANY EXPRESS OR IMPLIED WARRANTIES, INCLUDING, BUT NOT LIMITED TO,
 * THE IMPLIED WARRANTIES OF MERCHANTABILITY AND FITNESS FOR A PARTICULAR PURPOSE ARE DISCLAIMED. IN NO EVENT SHALL THE COPYRIGHT HOLDER OR CONTRIBUTORS
 * BE LIABLE FOR ANY DIRECT, INDIRECT, INCIDENTAL, SPECIAL, EXEMPLARY, OR CONSEQUENTIAL DAMAGES (INCLUDING, BUT NOT LIMITED TO, PROCUREMENT OF
 * SUBSTITUTE GOODS OR SERVICES; LOSS OF USE, DATA, OR PROFITS; OR BUSINESS INTERRUPTION) HOWEVER CAUSED AND ON ANY THEORY OF LIABILITY, WHETHER
 * IN CONTRACT, STRICT LIABILITY, OR TORT (INCLUDING NEGLIGENCE OR OTHERWISE) ARISING IN ANY WAY OUT OF THE USE OF THIS SOFTWARE, EVEN IF ADVISED OF
 * THE POSSIBILITY OF SUCH DAMAGE.
 */

/// C++ classes for hierarchical Bayesian Multi-trait quantitative-genetic models.
/** \file
 * \author Anthony J. Greenberg
 * \copyright Copyright (c) 2015 Anthony J. Greenberg
 * \version 0.9.1
 * 
 * This is the project header file containing class definitions and interface documentation.
 *
 */

#ifndef LIBMUGEN_H
#define LIBMUGEN_H
#include <gsl/gsl_matrix.h>
#include <gsl/gsl_vector.h>
#include <gsl/gsl_rng.h>
#include <gsl/gsl_permutation.h>
#include <vector>
#include <string>
#include <list>

using std::vector;
using std::list;
using std::string;

class MVnorm;
class MVnormMu;
class MVnormMuPEX;
class MVnormBetaPEX;
class MVnormMuMiss;
class MVnormBeta;
class MVnormBetaMiss;
class MVnormBetaFt;
class MVnormBetaSc;
class MVnormMuBlk;
class MVnormBetaBlk;
class MVnormBetaFtBlk;
class RanIndex;
class RanIndexVS;
class Apex;
class Grp;
class MuGrp;
class MuGrpPEX;
class BetaGrpPEX;
class MuGrpMiss;
class MuGrpEE;
class MuGrpEEmiss;
class BetaGrpSnp;
class BetaGrpSnpCV;
class BetaGrpPSR;
class BetaGrpSnpMiss;
class BetaGrpSnpMissCV;
class BetaGrpPSRmiss;
class BetaGrpFt;
class BetaGrpSc;
class BetaGrpPC;
class BetaGrpPCpex;
class BetaGrpBVSR;
class MuBlk;
class BetaBlk;
class SigmaI;
class SigmaIblk;
class SigmaIpex;
class Qgrp;
class QgrpPEX;
class MixP;

/** \defgroup distributions Various distribution functions
 *
 * Sampling functions for distributions not already in GSL.
 *
 * \warning These are internal functions that are invoked from within classes. Because they need to be as efficient as possible, no range-checking is performed (especially if GSL range checking is turned off at compilation).  They rely on the user to keep track of dimensions.
 *
 *
 * @{
 */

/** \brief Multivariate Gaussian sampling function
 * 
 * This function gives a sample from the MV Gaussian distribution with a given mean and covariance matrix.
 *
 * \param[in] gsl_vector* pointer to a vector of means
 * \param[in] gsl_matrix* pointer to a matrix with the Cholesky-decomposed covariance matrix
 * \param[in] gsl_rng* pointer to a pseudo-random number generator (PNG)
 * \param[out] gsl_vector* pointer to a destination vector for the sample
 *
 */
void MVgauss(const gsl_vector*, const gsl_matrix*, const gsl_rng*, gsl_vector*);

/** \defgroup Overloaded Wishart sampling functions
 *
 * These functions give samples from the Wishart distribution, the only difference between them is the type of the degrees of freedom paramter.
 *
 * \param[in] gsl_matrix* pointer to a matrix with the Cholesky-transformed covariance matrix parameter
 * \param[in] int/size_t& degrees of freedom parameter
 * \param[in] gsl_rng* pointer to a PNG
 * \param[out] gsl_matrix* pointer to a matrix with the sample
 * 
 * @{
 */
void Wishart(const gsl_matrix*, const int&, const gsl_rng*, gsl_matrix*);
void Wishart(const gsl_matrix*, const size_t&, const gsl_rng*, gsl_matrix*);
/** @} */

/** \brief Truncated geometric distribution
 *
 * Sampling from the truncated geometric distribution, returning a size_t type index value.
 *
 * \param[in] double& probability of success
 * \param[in] size_t& maximum value
 * \param[in] gsl_rng* pointer to a PNG
 * \return a value of the type size_t that is a sample from the distribution.
 *
 */
size_t rtgeom(const double&, const size_t&, const gsl_rng*);
/** @} */

/** \defgroup auxFun Auxiliary functions
 *	 
 * Various functions that are needed internally.
 *
 * @{
 */
/** \brief Mahalanobis distance
 *
 * Calculates the Mahalanobis distance of a vector from zero.
 *
 * \param[in] gsl_vector* the vector
 * \param[in] gsl_matrix* the corresponding inverse-covariance matrix
 * \return a scalar value of type double.
 *
 */
double mhl(const gsl_vector *beta, const gsl_matrix *SigI);
/** \defgroup centerFun Centering functions
 *
 * Functions that center matrix columns and vectors.
 *
 * @{
 */
/** \brief Matrix centering in-place
 *
 * \param[in,out] gsl_matrix* the matrix to be modified in-place
 *
 */
void colCenter(gsl_matrix *inplace);
/** \brief Matrix centering with copy
 *
 * \param[in] gsl_matrix* the matrix to be centered, left unmodified
 * \param[out] gsl_matrix* the modified matrix
 *
 */
void colCenter(const gsl_matrix *source, gsl_matrix *res);
/** \brief Matrix centering in-place with missing values
 *
 * \param[in,out] gsl_matrix* matrix to be modified in-place
 * \param[in] double& label for missing values
 *
 */
void colCenter(gsl_matrix *inplace, const double &absLab);
/** \brief Matrix centering with copy and missing values
 *
 *	In addition to centering does mean-imputation of missing values.
 *
 * \param[in] gsl_matrix* matrix to be centered, but not changed
 * \param[out] gsl_matrix* centered matrix
 * \param[in] double& label for missing values
 *
 */
void colCenter(const gsl_matrix *source, gsl_matrix *res, const double &absLab);
/** \brief Vector centering in-place
 *
 * \param[in,out] gsl_vector* vector to be modified
 * \param[in] double& label for missing values
 *
 */
void vecCenter(gsl_vector *inplace);
/** \brief Vector centering with copy
 *
 * \param[in] gsl_vector* vector to be centered, but not changed
 * \param[out] gsl_vector* modified vector
 *
 */
void vecCenter(const gsl_vector *source, gsl_vector *res);
/** \brief Mean imputation without centering
 *
 * \param[in,out] gsl_matrix* matrix to be modified in-place
 * \param[in] double& label for missing values
 *
 */
void meanImpute(gsl_matrix *inplace, const double &absLab);
/** @} */
/** \brief Print matrix to screen
 *
 * Printing a GSL matrix to screen, with each row on a line, values seprated by spaces.
 *
 * \param[in] gsl_matrix* matrix to be displayed
 * \param[in] double& label for missing data
 *
 */
void printMat(const gsl_matrix *m);
/** \brief Accessing the processor RTDSC instruction
 *
 * \return a value of type unsigned long long.
 *
 * This function outputs the processor RTDSC instruction for use in seeding random number generators using assembly code.
 * \warning this function is unlikely to work under Windows.
 *
 */
unsigned long long rdtsc();
/** @} */

/** \defgroup rowLocParam Individual location parameters
 *
 * A hierarchy of classes that refer to rows of location parameter matrices. These classes are internal to Grp classes and are not directly declared by the user of the library.  
 * They are typically updated with samples from the multivariate normal distribution.  Because the methods are used for much of the computation, they are implemented for speed and almost no chacking for errors is done.
 * The assumption is that the encapsualting classes will do all of that correctly. Constructors are used to set initial values.
 *
 * @{
 */
/** \brief The abstract base class for location parameter rows
 *
 * This is the generic location parameter class and cannot be envoked directly.  All the constructors are protected.
 */
class MVnorm {
protected:
	/** \brief Data vector
	 * 
	 * This is actually a pointer (implemented as a GSL *vector_view*) to a row in the corresponding matrix of location parameters.
	 *
	 */
	gsl_vector_view _vec;
	
	/** \brief Length of the data vector */
	size_t _d;
	
	/** \brief Default constructor
	 *
	 * This is a deterministic constructor that results in a pointer to nowhere and the dimension member set to zero.
	 */
	MVnorm() : _d(0) {};
	/** \brief Dimension-only constructor
	 *
	 * Sets the dimension to a value, but the *vector_view* is not assigned a target.
	 * 
	 * \param[in] size_t& dimension value
	 */
	MVnorm(const size_t &d) : _d(d) {};
	/** \brief Dimension and vector value constructor
	 *
	 * A deterministic constructor that sets the _vec member to point to the provided *gsl_vector* and the dimension member to the size of the provided *gsl_vector*.
	 *
	 * \param[in] gsl_vector* target vector
	 */
	MVnorm(gsl_vector *mn);
	/** \brief Univariate Gaussian constructor
	 *
	 * Sets the _vec member to point to the provided *gsl_vector* and ads univariate Gaussian noise independtly to each element, modifying the target.
	 *
	 * \param[in,out] gsl_vector* vector of means
	 * \param[in] gsl_vector* vector of standard deviations for the univariate Gaussian, has to be no shorter than the mean vector.  If it is longer, the extra values are ignored.
	 * \param[in] gsl_rng* pointer to a PNG
	 *
	 */
	MVnorm(gsl_vector *mn, const gsl_vector *sd, const gsl_rng *r);
	/** \brief Multivariate Gaussian constructor
	 *
	 * Sets the _vec member to point to the provided *gsl_vector* and ads multivariate Gaussian noise, modifying the target.
	 *
	 * \param[in,out] gsl_vector* vector of means
	 * \param[in] gsl_matrix* covariance matrix for the multivariate Gaussian, has to match the mean vector in dimensions and has to be symmetric positive-definite
	 * \param[in] gsl_rng* pointer to a PNG
	 *
	 */
	MVnorm(gsl_vector *mn, const gsl_matrix *Sig, const gsl_rng *r);
	/** \brief Dimension and vector value constructor
	 *
	 * A deterministic constructor that sets the _vec member to point to a row (indicated by the row index) of the provided *gsl_matrix* and the dimension member to the number of columns in the provided *gsl_matrix*.
	 *
	 * \param[in] gsl_matrix* target matrix
	 * \param[in] size_t& row index of the matrix
	 */
	MVnorm(gsl_matrix *mn, const size_t &iRw);
	/** \brief Univariate Gaussian constructor with a matrix
	 *
	 * Sets the _vec member to point to a row (indicated by the row index) of the provided *gsl_matrix* and ads univariate Gaussian noise independtly to each element, modifying the target.
	 *
	 * \param[in,out] gsl_matrix* target matrix
	 * \param[in] size_t& row index of the target matrix
	 * \param[in] gsl_vector* vector of standard deviations for the univariate Gaussian, has to be no shorter than a row of the mean matrix.  If it is longer, the extra values are ignored.
	 * \param[in] gsl_rng* pointer to a PNG
	 *
	 */
	MVnorm(gsl_matrix *mn, const size_t &iRw, const gsl_vector *sd, const gsl_rng *r);
	/** \brief Multivariate Gaussian constructor with a matrix
	 *
	 * Sets the _vec member to point to a row (indicated by the row index) of the provided *gsl_matrix* and ads multivariate Gaussian noise, modifying the target.
	 *
	 * \param[in,out] gsl_matrix* target matrix
	 * \param[in] size_t& row index of the target matrix
	 * \param[in] gsl_matrix* covariance matrix for the multivariate Gaussian, has to be symmetric positive-definite, with diemsions equal to the number of columns in the target matrix
	 * \param[in] gsl_rng* pointer to a PNG
	 *
	 */
	MVnorm(gsl_matrix *mn, const size_t &iRw, const gsl_matrix *Sig, const gsl_rng *r);
	
public:
	/** \brief Copy constructor
	 *
	 * Provided for completeness and has not been extensively tested.  The constructor is deterministic, i.e.\ the values are copied exactly, with no stochastic perturbation.
	 *
	 * \param[in] MVnorm& object to be copied
	 * \return object of class _MVnorm_.
	 */
	MVnorm(const MVnorm &);
	/** \brief Assignement operator
	 *
	 * Provided for completeness and has not been extensively tested.  The operator is deterministic, i.e.\ the values are copied exactly, with no stochastic perturbation.
	 *
	 * \param[in] MVnorm& object to be assigned
	 * \return reference to an object of class _MVnorm_.
	 */
	MVnorm & operator=(const MVnorm &);
	
	/** \brief Virtual destructor
	 *
	 * The destructor has nothing to do since members of this class cannot be de-allocated.
	 */
	virtual ~MVnorm() {};
	
	/** \defgroup updateFun Overloaded update methods
	 *
	 * Update functions generate new stochastic values of a given parameter or set of parameters as we step through the Markov chain iteration.  These are mostly Gibbs updates, but occasional Metropolis steps are used in special cases.
	 *
	 * @{
	 */
	/** \defgroup locIP Improper prior methods
	 *
	 * @{
	 */
	/** \brief Gaussian likelihood
	 *
	 * \param[in] Grp& data
	 * \param[in] SigmaI& inverse-covariance matrix for the likelihood
	 * \param[in] gsl_rng* pointer to a PNG
	 */
	virtual void update(const Grp &, const SigmaI &, const gsl_rng *) = 0;
	/** \brief Sudent-\f$t\f$ likelihood
	 *
	 * \param[in] Grp& data
	 * \param[in] Qgrp& vector of Student-\f$t\f$ covariance scale parameters for the likelihood covariance
	 * \param[in] SigmaI& inverse-covariance matrix for the likelihood
	 * \param[in] gsl_rng* pointer to a PNG
	 */
	virtual void update(const Grp &, const Qgrp &, const SigmaI &, const gsl_rng *) = 0;
	/** @} */
	/** \addtogroup locZMn 0-mean prior methods
	 *
	 * @{
	 */
	/** \brief Gaussian likelihood, Gaussian prior
	 *
	 * \param[in] Grp& data for the likelihood
	 * \param[in] SigmaI& inverse-covariance matrix for the likelihood
	 * \param[in] SigmaI& prior inverse-covariance matrix
	 * \param[in] gsl_rng* pointer to a PNG
	 */
	virtual void update(const Grp &, const SigmaI &, const SigmaI &, const gsl_rng *) = 0;
	/** \brief Gaussian likelihood, Student-\f$t\f$ prior
	 *
	 * \param[in] Grp& data for the likelihood
	 * \param[in] SigmaI& inverse-covariance matrix for the likelihood
	 * \param[in] double& Student-\f$t\f$ scale parameter for the prior covariance
	 * \param[in] SigmaI& prior inverse-covariance matrix
	 * \param[in] gsl_rng* pointer to a PNG
	 */
	virtual void update(const Grp &, const SigmaI &, const double &, const SigmaI &, const gsl_rng *) = 0;
	/** \brief Student-\f$t\f$ likelihood, Gaussian prior
	 *
	 * \param[in] Grp& data for the likelihood
	 * \param[in] Qgrp& vector of Student-\f$t\f$ covariance scale parameters for the likelihood covariance
	 * \param[in] SigmaI& inverse-covariance matrix for the likelihood
	 * \param[in] SigmaI& prior inverse-covariance matrix
	 * \param[in] gsl_rng* pointer to a PNG
	 */
	virtual void update(const Grp &, const Qgrp &, const SigmaI &, const SigmaI &, const gsl_rng *) = 0;
	/** \brief Student-\f$t\f$ likelihood, Student-\f$t\f$ prior
	 *
	 * \param[in] Grp& data for the likelihood
	 * \param[in] Qgrp& vector Student-\f$t\f$ covariance scale parameter for the likelihood covariance
	 * \param[in] SigmaI& inverse-covariance matrix for the likelihood
	 * \param[in] double& Student-\f$t\f$ scale parameter for the prior covariance
	 * \param[in] SigmaI& prior inverse-covariance matrix
	 * \param[in] gsl_rng* pointer to a PNG
	 */
	virtual void update(const Grp &, const Qgrp &, const SigmaI &, const double &, const SigmaI &, const gsl_rng *) = 0;
	/** @} */
	/** \addtogroup locNZMn non-0-mean prior methods
	 *
	 * @{
	 */
	/** \brief Gaussian likelihood, Gaussian prior
	 *
	 * \param[in] Grp& data for the likelihood
	 * \param[in] SigmaI& inverse-covariance matrix for the likelihood
	 * \param[in] Grp& prior mean
	 * \param[in] SigmaI& prior inverse-covariance matrix
	 * \param[in] gsl_rng* pointer to a PNG
	 */
	virtual void update(const Grp &, const SigmaI &, const Grp &, const SigmaI &, const gsl_rng *) = 0;
	/** \brief Gaussian likelihood, Student-\f$t\f$ prior
	 *
	 * \param[in] Grp& data for the likelihood
	 * \param[in] SigmaI& inverse-covariance matrix for the likelihood
	 * \param[in] Grp& prior mean
	 * \param[in] double& Student-\f$t\f$ scale parameter for the prior covariance
	 * \param[in] SigmaI& prior inverse-covariance matrix
	 * \param[in] gsl_rng* pointer to a PNG
	 */
	virtual void update(const Grp &, const SigmaI &, const Grp &, const double &, const SigmaI &, const gsl_rng *) = 0;
	/** \brief Student-\f$t\f$ likelihood, Gaussian prior
	 *
	 * \param[in] Grp& data for the likelihood
	 * \param[in] Qgrp& vector Student-\f$t\f$ covariance scale parameter for the likelihood covariance
	 * \param[in] SigmaI& inverse-covariance matrix for the likelihood
	 * \param[in] Grp& prior mean
	 * \param[in] SigmaI& prior inverse-covariance matrix
	 * \param[in] gsl_rng* pointer to a PNG
	 */
	virtual void update(const Grp &, const Qgrp &, const SigmaI &, const Grp &, const SigmaI &, const gsl_rng *) = 0;
	/** \brief Student-\f$t\f$ likelihood, Student-\f$t\f$ prior
	 *
	 * \param[in] Grp& data for the likelihood
	 * \param[in] Qgrp& vector Student-\f$t\f$ covariance scale parameter for the likelihood covariance
	 * \param[in] SigmaI& inverse-covariance matrix for the likelihood
	 * \param[in] Grp& prior mean
	 * \param[in] double& Student-\f$t\f$ scale parameter for the prior covariance
	 * \param[in] SigmaI& prior inverse-covariance matrix
	 * \param[in] gsl_rng* pointer to a PNG
	 */
	virtual void update(const Grp &, const Qgrp &, const SigmaI &, const Grp &, const double &, const SigmaI &, const gsl_rng *) = 0;
	/** @} */
	/** @} */
	// end of the update group
	
	/** \defgroup mahal Mahalanobis distance functions
	 *
	 * Overloaded functions to calculate the Mahalanobis distance of the vector of location parameters stored in the class to another vector or to zero.
	 *
	 * @{
	 */
	/** \brief Mahalanobis distance to a vector
	 *
	 * Calculates the Mahalanobis distance between the vector stored in the class and the provided vector of class MVnorm.
	 *
	 * \param[in] MVnorm& vector to calculate the distance from
	 * \param[in] SigmaI& inverse covariance matrix for scaling
	 * \return the distance value of type _double_.
	 *
	 */
	virtual double mhl(const MVnorm *x, const SigmaI &SigI);
	/** \brief Mahalanobis distance to a vector
	 *
	 * Calculates the Mahalanobis distance between the vector stored in the class and the provided vector of class MVnorm.  The const version of the above function.
	 *
	 * \param[in] MVnorm& vector to calculate the distance from
	 * \param[in] SigmaI& inverse covariance matrix for scaling
	 * \return the distance value of type _double_.
	 *
	 */
	virtual double mhl(const MVnorm *x, const SigmaI &SigI) const;
	/** \brief Mahalanobis distance to a vector
	 *
	 * Calculates the Mahalanobis distance between the vector stored in the class and the provided GSL vector.
	 *
	 * \param[in] MVnorm& vector to calculate the distance from
	 * \param[in] SigmaI& inverse covariance matrix for scaling
	 * \return the distance value of type _double_.
	 *
	 */
	virtual double mhl(const gsl_vector *x, const SigmaI &SigI);
	/** \brief Mahalanobis distance to a vector
	 *
	 * Calculates the Mahalanobis distance between the vector stored in the class and the provided GSL vector. The const version of the above function.
	 *
	 * \param[in] MVnorm& vector to calculate the distance from
	 * \param[in] SigmaI& inverse covariance matrix for scaling
	 * \return the distance value of type _double_.
	 *
	 */
	virtual double mhl(const gsl_vector *x, const SigmaI &SigI) const;
	/** \brief Mahalanobis distance to zero
	 *
	 * Calculates the Mahalanobis distance between the vector stored in the class zero.
	 *
	 * \param[in] SigmaI& inverse covariance matrix for scaling
	 * \return the distance value of type _double_.
	 *
	 */
	virtual double mhl(const SigmaI &SigI); // distance from 0
	/** \brief Mahalanobis distance to zero
	 *
	 * Calculates the Mahalanobis distance between the vector stored in the class zero.  The const version of the above function.
	 *
	 * \param[in] SigmaI& inverse covariance matrix for scaling
	 * \return the distance value of type _double_.
	 *
	 */
	virtual double mhl(const SigmaI &SigI) const; // distance from 0
	/** @} */
	/** \defgroup MVnormDens Multivariate Gaussian density functions
	 *
	 * These functions calculate multivariate Gaussian density given the mean and inverse-covariance provided and the vector of values stored in the class as data.
	 *
	 * @{
	 */
	/** \brief Multivariate Gaussian density
	 *
	 * \param[in] gsl_vector* vector of means
	 * \param[in] SigmaI& inverse-covariance matrix
	 * \return density value of type _double_.
	 *
	 */
	double density(const gsl_vector *theta, const SigmaI &SigI);
	/** \brief Multivariate Gaussian density
	 *
	 * \note A const version of the above function.
	 *
	 * \param[in] gsl_vector* vector of means
	 * \param[in] SigmaI& inverse-covariance matrix
	 * \return density value of type _double_.
	 *
	 */
	double density(const gsl_vector *theta, const SigmaI &SigI) const;
	/** \brief Multivariate Gaussian density
	 *
	 * \param[in] MVnorm* vector of means
	 * \param[in] SigmaI& inverse-covariance matrix
	 * \return density value of type _double_
	 *
	 */
	double density(const MVnorm *theta, const SigmaI &SigI);
	/** \brief Multivariate Gaussian density
	 *
	 * \note A const version of the above function.
	 *
	 * \param[in] MVnorm* vector of means
	 * \param[in] SigmaI& inverse-covariance matrix
	 * \return density value of type _double_.
	 *
	 */
	double density(const MVnorm *theta, const SigmaI &SigI) const;
	/** @} */
	
	/** \brief Save function
	 *
	 * Saves the current value of the location parameter to a file, appending by default.
	 *
	 * \param[in] string& file name
	 * \param[in] char* save mode, "a" for append by default. The allowable values are as for a standard C file stream
	 *
	 */
	void save(const string &fileNam, const char *how = "a");
	/** \brief Save function
	 *
	 * Saves the current value of the location parameter to a file.
	 *
	 * \param[in] FILE* C file stream
	 *
	 */
	void save(FILE *fileStr);
	
	/** \brief Subscript operator
	 *
	 * Overloaded subscript operator for the vector of location parameter values.
	 *
	 * \param[in] size_t index value
	 * \return value of the member vector corresponding to the index value, of type _double_.
	 *
	 */
	double operator[](const size_t i) const{return gsl_vector_get(&_vec.vector, i); };
	/** \brief Setting an element to a value
	 *
	 * Sets the _i_-th element of the location vector to a value deterministically
	 *
	 * \param[in] size_t index
	 * \param[in] double new value of the element
	 *
	 */
	void valSet(const size_t i, const double x){gsl_vector_set(&_vec.vector, i, x); };
	/** \brief Access the location vector
	 *
	 * Provides access to the internal parameter vector.
	 *
	 * \return GSL vector pointing to the corresponding location parameter.
	 */
	const gsl_vector *getVec() const{return &_vec.vector;};
	/** \brief Length of the location vector
	 *
	 * \return Length of the parameter vector, of type _size_t_. Corresponds to the number of traits.
	 *
	 */
	size_t len() const{return _d; };
	/** \brief Number of missing values
	 *
	 * Accesses the  number of missing phenotype values.  Non-zero only for classes that implement treatment of missing values.
	 *
	 * \return Number of missing phenotypes, of type _size_t_.
	 */
	virtual size_t nMissP() const{return 0; };
	/** \brief Indexes of missing values
	 *
	 * Accesses a vector with indexes of missing phenotypes.
	 *
	 * \return Vector of _size_t_ that contains indexes that correspond to missing values. For classes that do not allow missing values it is empty.
	 *
	 */
	virtual const vector<size_t> getMisPhen() const{ return vector<size_t>(0); };
	/** \brief Points to the corresponding data
	 *
	 * Access to the pointer to a vector that indexes the data used to update an instance of the class.  Is non-zero only for classes which can be part of a hierarchical model.
	 *
	 * \return Pointer to a vector of _size_t_.
	 *
	 */
	virtual const vector<size_t> *down() const{return 0; };
	/** \brief Points to the prior
	 *
	 * Access to the pointer to the correspoding vector of priors.  Is non-zero only for classes where a prior is implemented.
	 *
	 * \return Pointer to _size_t_.
	 *
	 */
	virtual const size_t *up() const{return 0; };
	/** \brief Scale parameter
	 *
	 * Access to a scale parameter member, defined only for some derived regression-type classes.  If not defined, returns 1.0.
	 *
	 * \return Scale paramter value of type _double_.
	 *
	 */
	virtual double scalePar() const {return 1.0; };
};

/** \brief Individual vector of means
 *
 * Refers to a row of a matrix of location parameters that are updated with a likelihood that is an inverse-covariance weighted mean of the data. Variables of this class are typically imbedded in a model hierarchy, and in that case are updated with non-0-mean priors. 
 * This class allows us to keep track of which rows in data and prior matrices to use through pointers to index vectors (for data, where there may be more than one row) and to a single index (for the prior mean, since there is only one).  
 * We use pointers because they allow us to modify the indexes in other parts of the model, thus allowing for things like mixture models and model selection schemes.
 *
 */
class MVnormMu : public MVnorm {
protected:
	/** \brief Data indexes
	 *
	 * A pointer to a vector of _size_t_. The elements are row indexes of the data matrix.  Means among these rows are used in the likelihood of the _udpate_ functions.
	 * The pointer is _const_ to make sure we are not modifying it from this class.
	 *
	 */
	const vector<size_t> *_lowLevel;
	
	/** \brief Prior index
	 *
	 * Pointer to a value of type _size_t_ that is a row index of the prior location matrix. The pointer is _const_ to make sure we are not modifying it from this class.
	 *
	 */
	const size_t *_upLevel;
	
public:
	/** \brief Default constructor
	 *
	 * The _vec member is unassigned and the low-level and upper-level pointers are set to zero.
	 *
	 */
	MVnormMu();
	/** \brief Zero vector constructor
	 *
	 * Sets _vec to point to a vector of zeros, of length *d*.
	 *
	 * \param[in] size_t& size of the vector
	 */
	MVnormMu(const size_t &d) : MVnorm(d) {};
	/** \brief Zero vector with pointers
	 *
	 * Sets _vec to point to a vector of zeros, of length *d*.  Pointers to the lower and upper level of the model hierarchy are set.
	 *
	 * \param[in] size_t& size of the vector
	 * \param[in] vector<size_t>& vector of indexes of rows in the data matrix
	 * \param[in] size_t& index in the matrix of priors
	 *
	 */
	MVnormMu(const size_t &d, const vector<size_t> &low, const size_t &up);
	/** \brief Deterministic constructor
	 *
	 * Sets _vec to point to a GSL vector of values, without modifying the target during the invocation of the constructor. The pointers to the data and the prior indexes are set.
	 *
	 * \param[in] gsl_vector* vector of values
	 * \param[in] vector<size_t>& vector of indexes of rows in the data matrix
	 * \param[in] size_t& index in the matrix of priors
	 *
	 */
	MVnormMu(gsl_vector *mn, const vector<size_t> &low, const size_t &up);
	/** \brief Deterministic constructor, prior index only
	 *
	 * Sets _vec to point to a GSL vector of values, without modifying the target during the invocation of the constructor. Only the pointer to the prior index is set. This constructor can be used for the lowest hierarchy level.
	 *
	 * \param[in] gsl_vector* vector of values
	 * \param[in] size_t& index in the matrix of priors
	 *
	 */
	MVnormMu(gsl_vector *mn, const size_t &up);
	/** \brief Deterministic constructor with a matrix
	 *
	 * Sets _vec to point to a row of the GSL matrix of values, without modifying the target during the invocation of the constructor.
	 *
	 * \param[in] gsl_matrix* matrix of values
	 * \param[in] size_t& row index of the matrix of values, has to be no smaller than 0 and strictly smaller than the number of rows in _mn_
	 *
	 */
	MVnormMu(gsl_matrix *mn, const size_t &iRw);
	/** \brief Deterministic constructor with a matrix and an index to data
	 *
	 * Sets _vec to point to a row of the GSL matrix of values, without modifying the target during the invocation of the constructor.  None of the index pointers are set.
	 *
	 * \param[in] gsl_matrix* matrix of values
	 * \param[in] size_t& row index of the matrix of values, has to be no smaller than 0 and strictly smaller than the number of rows in _mn_
	 * \param[in] vector<size_t>& vector of indexes of rows in the data matrix
	 *
	 */
	MVnormMu(gsl_matrix *mn, const size_t &iRw, const vector<size_t> &low);
	/** \brief Deterministic constructor with a matrix and indexes to data and a prior
	 *
	 * Sets _vec to point to a row of the GSL matrix of values, without modifying the target during the invocation of the constructor.
	 *
	 * \param[in] gsl_matrix* vector of values
	 * \param[in] size_t& row index of the matrix of values, has to be no smaller than 0 and strictly smaller than the number of rows in _mn_
	 * \param[in] vector<size_t>& vector of indexes of rows in the data matrix
	 * \param[in] size_t& index in the matrix of priors
	 *
	 */
	MVnormMu(gsl_matrix *mn, const size_t &iRw, const vector<size_t> &low, const size_t &up);
	/** \brief Deterministic constructor with a matrix and an index to a prior
	 *
	 * Sets _vec to point to a row of the GSL matrix of values, without modifying the target during the invocation of the constructor.
	 *
	 * \param[in] gsl_matrix* vector of values
	 * \param[in] size_t& row index of the matrix of values, has to be no smaller than 0 and strictly smaller than the number of rows in _mn_
	 * \param[in] size_t& index in the matrix of priors
	 *
	 */
	MVnormMu(gsl_matrix *mn, const size_t &iRw, const size_t &up);
	/** \brief Univariate random constructor with a vector and indexes to data and a prior
	 *
	 * Sets _vec to point to a GSL vector of values, modifying the target during the invocation of the constructor. Modification is by adding independent samples from a univariate normal distribution with the provided standard deviations.
	 *
	 * \param[in] gsl_vector* vector of values
	 * \param[in] gsl_vector* vector of standard deviations
	 * \param[in] gsl_rng* pointer to a PNG
	 * \param[in] vector<size_t>& vector of indexes of rows in the data matrix
	 * \param[in] size_t& row index of the matrix of values, has to be no smaller than 0 and strictly smaller than the number of rows in _mn_
	 *
	 */
	MVnormMu(gsl_vector *mn, const gsl_vector *sd, const gsl_rng *r, const vector<size_t> &low, const size_t &up);
	/** \brief Multivariate random constructor with a vector and indexes to data and a prior
	 *
	 * Sets _vec to point to a GSL vector of values, modifying the target during the invocation of the constructor. Modification is by adding samples from a multivariate normal distribution with the provided covariance matrix.
	 *
	 * \param[in] gsl_vector* vector of values
	 * \param[in] gsl_matrix* covariance matrix
	 * \param[in] gsl_rng* pointer to a PNG
	 * \param[in] vector<size_t>& vector of indexes of rows in the data matrix
	 * \param[in] size_t& index in the matrix of priors
	 *
	 */
	MVnormMu(gsl_vector *mn, const gsl_matrix *Sig, const gsl_rng *r, const vector<size_t> &low, const size_t &up);
	/** \brief Univariate random constructor with a matrix and indexes to data and a prior
	 *
	 * Sets _vec to point to a row of the GSL matrix of values, modifying the target during the invocation of the constructor. Modification is by adding independent samples from a univariate normal distribution with the provided standard deviations.
	 *
	 * \param[in] gsl_matrix* matrix of values
	 * \param[in] size_t& row index of the matrix of values, has to be no smaller than 0 and strictly smaller than the number of rows in _mn_
	 * \param[in] gsl_vector* vector of standard deviations
	 * \param[in] gsl_rng* pointer to a PNG
	 * \param[in] vector<size_t>& vector of indexes of rows in the data matrix
	 * \param[in] size_t& index in the matrix of priors
	 *
	 */
	MVnormMu(gsl_matrix *mn, const size_t &iRw, const gsl_vector *sd, const gsl_rng *r, const vector<size_t> &low, const size_t &up);
	/** \brief Multivariate random constructor with a matrix and indexes to data and a prior
	 *
	 * Sets _vec to point to a row of the GSL matrix of values, modifying the target during the invocation of the constructor. Modification is by adding samples from a multivariate normal distribution with the provided covariance matrix.
	 *
	 * \param[in] gsl_matrix* matrix of values
	 * \param[in] size_t& row index of the matrix of values, has to be no smaller than 0 and strictly smaller than the number of rows in _mn_
	 * \param[in] gsl_matrix* covariance matrix
	 * \param[in] gsl_rng* pointer to a PNG
	 * \param[in] vector<size_t>& vector of indexes of rows in the data matrix
	 * \param[in] size_t& index in the matrix of priors
	 *
	 */
	MVnormMu(gsl_matrix *mn, const size_t &iRw, const gsl_matrix *Sig, const gsl_rng *r, const vector<size_t> &low, const size_t &up);
	
	/** \brief Copy constructor
	 *
	 * Deterministic copy constructor
	 *
	 * \param[in] MVnormMu& object of type MVnormMu
	 *
	 */
	MVnormMu(const MVnormMu &);
	/** \brief Assignment operator
	 *
	 * Deterministic assignement operator
	 *
	 * \param[in] MVnormMu& object of type MVnormMu
	 *
	 * \return A reference to an object of type MVnormMu
	 *
	 */
	MVnormMu & operator=(const MVnormMu &);
	
	/** \brief Destructor
	 *
	 */
	virtual ~MVnormMu();
	
	// improper prior methods
	virtual void update(const Grp &dat, const SigmaI &SigIm, const gsl_rng *r);
	virtual void update(const Grp &dat, const Qgrp &q, const SigmaI &SigIm, const gsl_rng *r);
	// 0-mean prior methods
	virtual void update(const Grp &dat, const SigmaI &SigIm, const SigmaI &SigIp, const gsl_rng *r);
	virtual void update(const Grp &dat, const SigmaI &SigIm, const double &qPr, const SigmaI &SigIp, const gsl_rng *r);
	virtual void update(const Grp &dat, const Qgrp &q, const SigmaI &SigIm, const SigmaI &SigIp, const gsl_rng *r);
	virtual void update(const Grp &dat, const Qgrp &q, const SigmaI &SigIm, const double &qPr, const SigmaI &SigIp, const gsl_rng *r);
	// non-zero mean prior methods
	virtual void update(const Grp &dat, const SigmaI &SigIm, const Grp &muPr, const SigmaI &SigIp, const gsl_rng *r);
	virtual void update(const Grp &dat, const SigmaI &SigIm, const Grp &muPr, const double &qPr, const SigmaI &SigIp, const gsl_rng *r);
	virtual void update(const Grp &dat, const Qgrp &q, const SigmaI &SigIm, const Grp &muPr, const SigmaI &SigIp, const gsl_rng *r);
	virtual void update(const Grp &dat, const Qgrp &q, const SigmaI &SigIm, const Grp &muPr, const double &qPr, const SigmaI &SigIp, const gsl_rng *r);
	
	virtual size_t nMissP() const{return 0; };
	virtual const vector<size_t> getMisPhen() const{ return vector<size_t>(0); };
	/** \brief Points to the corresponding data
	 *
	 * Access to the pointer to a vector that indexes the data used to update an instance of the class.  Is implemented for this class.
	 *
	 * \return Pointer to a vector of _size_t_.
	 *
	 */
	const vector<size_t> *down() const{return _lowLevel; };
	/** \brief Points to the prior
	 *
	 * Access to the pointer to the correspoding vector of priors.  Is implemented for this class.
	 *
	 * \return Pointer to _size_t_.
	 *
	 */
	const size_t *up() const{return _upLevel; };
};

/** \brief Individual vector of means with parameter expansion
 *
 * This class is the same as MVnormMu, but implements multivariate parameter expansion for updating (Greenberg, in prep.)
 *
 */
class MVnormMuPEX : public MVnormMu {
protected:
	/** \brief Pointer to the redundant parameter
	 *
	 * A pointer to an object of class Apex, that stores the matrix of redundant parameters.
	 *
	 */
	Apex *_A;
	/** \brief Pre-computed auxiliary matrix
	 *
	 * Vector view of a matrix that stores a pre-computed \f$ (SA)^{\textsc{t}} \f$ matrix that is common for all members of the encapsulating Grp class.
	 *
	 */
	gsl_matrix_view _tSAprod;
	
public:
	/** \brief Default constructor
	 *
	 */
	MVnormMuPEX() : MVnormMu() {_A = 0; };
	/** \brief Deterministic constructor
	 *
	 * \param[in] gsl_matrix* matrix of values
	 * \param[in] size_t& row index of the matrix of values
	 * \param[in] vector<size_t>& vector of indexes of rows in the data matrix
	 * \param[in] size_t& index in the matrix of priors
	 * \param[in] Apex& object storing the redundant parameter matrix
	 * \param[in] gsl_matrix* pre-computed auxiliary matrix
	 *
	 */
	MVnormMuPEX(gsl_matrix *mn, const size_t &iRw, const vector<size_t> &low, const size_t &up, Apex &A, gsl_matrix *tSigIAt);
	/** \brief Univariate random constructor
	 *
	 * \param[in] gsl_matrix* matrix of values
	 * \param[in] size_t& row index of the matrix of values
	 * \param[in] gsl_vector* vector of standard deviations for the univariate Gaussian
	 * \param[in] gsl_rng* pointer to a PNG
	 * \param[in] vector<size_t>& vector of indexes of rows in the data matrix
	 * \param[in] size_t& index in the matrix of priors
	 * \param[in] Apex& object storing the redundant parameter matrix
	 * \param[in] gsl_matrix* pre-computed auxiliary matrix
	 *
	 */
	MVnormMuPEX(gsl_matrix *mn, const size_t &iRw, const gsl_vector *sd, const gsl_rng *r, const vector<size_t> &low, const size_t &up, Apex &A, gsl_matrix *tSigIAt);
	
	/** \brief Deterministic copy constructor
	 *
	 * \param[in] MVnormMuPEX& object of type MVnormMuPEX
	 *
	 */
	MVnormMuPEX(const MVnormMuPEX &mu);
	/** \brief Assignment operator
	 *
	 * Deterministic assignement operator
	 *
	 * \param[in] MVnormMuPEX& object of type MVnormMuPEX
	 *
	 * \return A reference to an object of type MVnormMuPEX
	 *
	 */
	MVnormMuPEX & operator=(const MVnormMuPEX &mu);
	
	/** \brief Destructor */
	virtual ~MVnormMuPEX() {};
	
	virtual void update(const Grp &dat, const SigmaI &SigIm, const SigmaI &SigIp, const gsl_rng *r);
	virtual void update(const Grp &dat, const SigmaI &SigIm, const double &qPr, const SigmaI &SigIp, const gsl_rng *r);
	virtual void update(const Grp &dat, const Qgrp &q, const SigmaI &SigIm, const SigmaI &SigIp, const gsl_rng *r);
	virtual void update(const Grp &dat, const Qgrp &q, const SigmaI &SigIm, const double &qPr, const SigmaI &SigIp, const gsl_rng *r);
	
	virtual void update(const Grp &dat, const SigmaI &SigIm, const Grp &muPr, const SigmaI &SigIp, const gsl_rng *r);
	virtual void update(const Grp &dat, const SigmaI &SigIm, const Grp &muPr, const double &qPr, const SigmaI &SigIp, const gsl_rng *r);
	virtual void update(const Grp &dat, const Qgrp &q, const SigmaI &SigIm, const Grp &muPr, const SigmaI &SigIp, const gsl_rng *r);
	virtual void update(const Grp &dat, const Qgrp &q, const SigmaI &SigIm, const Grp &muPr, const double &qPr, const SigmaI &SigIp, const gsl_rng *r);
};

/** \brief Individual regression with parameter expansion
 *
 * Implements handling of one-at-a-time regressions, as in MVnormBetaFt but with parameter expansion.
 *
 */
class MVnormBetaPEX : public MVnormMuPEX {
protected:
	/** \brief Predictor
	 *
	 * Points to a vector of predictor values, typically a column of the predictor matrix that is in the encapsulating Grp class.
	 *
	 */
	gsl_vector_view _X;
	/** \brief Scale parameter
	 *
	 * Typically \f$ x^{\textsc{t}}x \f$, but some exceptions exist in special cases.
	 *
	 */
	double _scale;
	/** \brief Length of the predictor */
	size_t _N;
	/** \brief Fitted values
	 *
	 * Points to a matrix of partial fitted values that exclude the current regression (\f$ \boldsymbol{X}_{\cdot -i}\boldsymbol{B}_{-i \cdot} \f$).
	 *
	 */
	gsl_matrix_view _fitted;
	
public:
	/** \brief Default constructor */
	MVnormBetaPEX() : MVnormMuPEX() {};
	/** \brief Random constructor
	 *
	 * Calculates initial values for the regression using the provided responce and covariance matrix.  The initial values modify the corresponding row of the value matrix, stored in the encapsulating class.
	 *
	 * \param[in] gsl_matrix* response matrix
	 * \param[in] gsl_matrix* predictor matrix
	 * \param[in] size_t& column index of the current predictor in the predictor matrix
	 * \param[in] vector<double>& vectorized partial fitted matrix
	 * \param[in] gsl_matrix* covariance matrix
	 * \param[in] gsl_rng* pointer to a PNG
	 * \param[in] size_t& row index for the prior matrix
	 * \param[in] gsl_matrix* value matrix
	 * \param[in] size_t& row index of the value matrix
	 * \param[in] Apex& object storing the redundant parameter matrix
	 * \param[in] gsl_matrix* pre-computed auxiliary matrix
	 *
	 */
	MVnormBetaPEX(const gsl_matrix *resp, gsl_matrix *pred, const size_t &iCl, vector<double> &eaFt, const gsl_matrix *Sig, const gsl_rng *r, const size_t &up, gsl_matrix *bet, const size_t &iRw, Apex &A, gsl_matrix *tSigIAt);
	/** \brief Deterministic constructor
	 *
	 * Sets up the proper pointers to parameters of the regression that have already been calculated in the encapsulating class.  The corresponding value matrix is not modified.
	 *
	 * \param[in] gsl_matrix* response matrix
	 * \param[in] gsl_matrix* predictor matrix
	 * \param[in] size_t& column index of the current predictor in the predictor matrix
	 * \param[in] vector<double>& vectorized partial fitted matrix
	 * \param[in] size_t& row index for the prior matrix
	 * \param[in] gsl_matrix* value matrix
	 * \param[in] size_t& row index of the value matrix
	 * \param[in] Apex& object storing the redundant parameter matrix
	 * \param[in] gsl_matrix* pre-computed auxiliary matrix
	 *
	 */
	MVnormBetaPEX(const size_t &d, gsl_matrix *pred, const size_t &iCl, vector<double> &eaFt, const size_t &up, gsl_matrix *bet, const size_t &iRw, Apex &A, gsl_matrix *tSigIAt);
	
	/** \brief Deterministic copy constructor
	 *
	 * \param[in] MVnormBetaPEX& object of type MVnormBetaPEX
	 *
	 */
	MVnormBetaPEX(const MVnormBetaPEX &bet);
	/** \brief Assignment operator
	 *
	 * Deterministic assignement operator
	 *
	 * \param[in] MVnormBetaPEX& object of type MVnormBetaPEX
	 *
	 * \return A reference to an object of type MVnormBetaPEX
	 *
	 */
	MVnormBetaPEX & operator=(const MVnormBetaPEX &bet);
	
	/** \brief Destructor */
	~MVnormBetaPEX() {};
	
	void update(const Grp &dat, const SigmaI &SigIm, const SigmaI &SigIp, const gsl_rng *r);
	void update(const Grp &dat, const SigmaI &SigIm, const double &qPr, const SigmaI &SigIp, const gsl_rng *r);
	void update(const Grp &dat, const Qgrp &q, const SigmaI &SigIm, const SigmaI &SigIp, const gsl_rng *r);
	void update(const Grp &dat, const Qgrp &q, const SigmaI &SigIm, const double &qPr, const SigmaI &SigIp, const gsl_rng *r);
	
	void update(const Grp &dat, const SigmaI &SigIm, const Grp &muPr, const SigmaI &SigIp, const gsl_rng *r);
	void update(const Grp &dat, const SigmaI &SigIm, const Grp &muPr, const double &qPr, const SigmaI &SigIp, const gsl_rng *r);
	void update(const Grp &dat, const Qgrp &q, const SigmaI &SigIm, const Grp &muPr, const SigmaI &SigIp, const gsl_rng *r);
	void update(const Grp &dat, const Qgrp &q, const SigmaI &SigIm, const Grp &muPr, const double &qPr, const SigmaI &SigIp, const gsl_rng *r);
};

/** \brief Individual vector of means with missing data
 *
 * Implements missing phenotype data imputation.  Some parameters for update methods have a different meaning than for other MVnorm classes.
 *
 */
class MVnormMuMiss : public MVnormMu {
protected:
	/** \brief Missing data index
	 *
	 * Vector of indexes tagging missing data.
	 *
	 */
	vector<size_t> _misPhenInd;
	/** \brief own index
	 *
	 * Index of the row the current instance of this class is pointing to.  Only defined with constructors that take _iRw_.
	 *
	 */
	size_t _myInd;
	
public:
	/** \brief Default constructor */
	MVnormMuMiss() : MVnormMu(){};
	/** \brief 0-mean deterministic constructor
	 *
	 * Sets up the instance to point to a vector of zeros
	 *
	 * \param[in] size_t& dimension of the vector
	 *
	 */
	MVnormMuMiss(const size_t &d) : MVnormMu(d) {};
	/** \brief Deterministic constructor
	 *
	 * \param[in] size_t& dimension of the vector
	 * \param[in] vector<size_t>& vector of row indexes for data
	 * \param[in] size_t& row index for the prior matrix
	 * \param[in] vector<size_t>& indexes of missing data
	 *
	 */
	MVnormMuMiss(const size_t &d, const vector<size_t> &low, const size_t &up, const vector<size_t> &mis);
	/** \brief Deterministic constructor with a vector
	 *
	 * \param[in] gsl_vector* vector of values
	 * \param[in] size_t& dimension of the vector
	 * \param[in] vector<size_t>& vector of row indexes for data
	 * \param[in] size_t& row index for the prior matrix
	 * \param[in] vector<size_t>& indexes of missing data
	 *
	 */
	MVnormMuMiss(gsl_vector *mn, const vector<size_t> &low, const size_t &up, const vector<size_t> &mis);
	/** \brief Deterministic constructor with a vector
	 *
	 * \param[in] gsl_vector* vector of values
	 * \param[in] size_t& row index for the prior matrix
	 * \param[in] vector<size_t>& indexes of missing data
	 *
	 */
	MVnormMuMiss(gsl_vector *mn, const size_t &up, const vector<size_t> &mis);
	/** \brief Univariate random constructor with a vector
	 *
	 * \param[in] gsl_vector* vector of values
	 * \param[in] gsl_vector* vector of standard deviations
	 * \param[in] gsl_rng* pointer to a PNG
	 * \param[in] size_t& dimension of the vector
	 * \param[in] vector<size_t>& vector of row indexes for data
	 * \param[in] size_t& row index for the prior matrix
	 * \param[in] vector<size_t>& indexes of missing data
	 *
	 */
	MVnormMuMiss(gsl_vector *mn, const gsl_vector *sd, const gsl_rng *r, const vector<size_t> &low, const size_t &up, const vector<size_t> &mis);
	/** \brief Multivariate random constructor with a vector
	 *
	 * \param[in] gsl_vector* vector of values
	 * \param[in] gsl_matrix* covariance matrix
	 * \param[in] gsl_rng* pointer to a PNG
	 * \param[in] size_t& dimension of the vector
	 * \param[in] vector<size_t>& vector of row indexes for data
	 * \param[in] size_t& row index for the prior matrix
	 * \param[in] vector<size_t>& indexes of missing data
	 *
	 */
	MVnormMuMiss(gsl_vector *mn, const gsl_matrix *Sig, const gsl_rng *r, const vector<size_t> &low, const size_t &up, const vector<size_t> &mis);
	/** \brief Deterministic constructor with a matrix
	 *
	 * \param[in] gsl_matrix* vector of values
	 * \param[in] size_t& row index of the matrix of values
	 * \param[in] vector<size_t>& vector of row indexes for data
	 * \param[in] size_t& row index for the prior matrix
	 * \param[in] vector<size_t>& indexes of missing data
	 *
	 */
	MVnormMuMiss(gsl_matrix *mn, const size_t &iRw, const vector<size_t> &low, const size_t &up, const vector<size_t> &mis);
	/** \brief Deterministic constructor with a matrix
	 *
	 * \param[in] gsl_matrix* vector of values
	 * \param[in] size_t& row index of the matrix of values
	 * \param[in] size_t& row index for the prior matrix
	 * \param[in] vector<size_t>& indexes of missing data
	 *
	 */
	MVnormMuMiss(gsl_matrix *mn, const size_t &iRw, const size_t &up, const vector<size_t> &mis);
	/** \brief Univariate random constructor with a matrix
	 *
	 * \param[in] gsl_matrix* matrix of values
	 * \param[in] size_t& row index of the matrix of values
	 * \param[in] gsl_vector* vector of standard deviations
	 * \param[in] gsl_rng* pointer to a PNG
	 * \param[in] size_t& dimension of the vector
	 * \param[in] vector<size_t>& vector of row indexes for data
	 * \param[in] size_t& row index for the prior matrix
	 * \param[in] vector<size_t>& indexes of missing data
	 *
	 */
	MVnormMuMiss(gsl_matrix *mn, const size_t &iRw, const gsl_vector *sd, const gsl_rng *r, const vector<size_t> &low, const size_t &up, const vector<size_t> &mis);
	/** \brief Multivariate random constructor with a matrix
	 *
	 * \param[in] gsl_matrix* matrix of values
	 * \param[in] size_t& row index of the matrix of values
	 * \param[in] gsl_matrix* covariance matrix
	 * \param[in] gsl_rng* pointer to a PNG
	 * \param[in] size_t& dimension of the vector
	 * \param[in] vector<size_t>& vector of row indexes for data
	 * \param[in] size_t& row index for the prior matrix
	 * \param[in] vector<size_t>& indexes of missing data
	 *
	 */
	MVnormMuMiss(gsl_matrix *mn, const size_t &iRw, const gsl_matrix *Sig, const gsl_rng *r, const vector<size_t> &low, const size_t &up, const vector<size_t> &mis);
	
	/** \brief Deterministic copy constructor
	 *
	 * \param[in] MVnormMuMiss& object of type MVnormMuMiss
	 *
	 */
	MVnormMuMiss(const MVnormMuMiss &); // copy constructor
	/** \brief Assignment operator
	 *
	 * Deterministic assignement operator
	 *
	 * \param[in] MVnormMuMiss& object of type MVnormMuMiss
	 *
	 * \return A reference to an object of type MVnormMuMiss
	 *
	 */
	MVnormMuMiss & operator=(const MVnormMuMiss &);
	
	/** \brief Destructor */
	~MVnormMuMiss() {};
	
	/** \brief Gaussian likelihood
	 *
	 * Here, the Grp variable contains the means for imputation rather than the data
	 *
	 * \param[in] Grp& data
	 * \param[in] SigmaI& inverse-covariance matrix for the likelihood
	 * \param[in] gsl_rng* pointer to a PNG
	 */
	void update(const Grp &mu, const SigmaI &SigIm, const gsl_rng *r);
	/** \brief Gaussian likelihood, Gaussian 0-mean prior
	 *
	 * Here, the Grp variable contains the means for imputation rather than the data
	 *
	 * \param[in] Grp& data
	 * \param[in] SigmaI& inverse-covariance matrix for the likelihood
	 * \param[in] SigmaI& inverse-covariance matrix for the prior
	 * \param[in] gsl_rng* pointer to a PNG
	 */
	void update(const Grp &mu, const SigmaI &SigIm, const SigmaI &SigIp, const gsl_rng *r);
	
	/** \brief Number of missing values
	 *
	 * Accesses the  number of missing phenotype values.  Implemented in this class.
	 *
	 * \return Number of missing phenotypes, of type _size_t_.
	 */
	size_t nMissP() const{return _misPhenInd.size(); };
	/** \brief Indexes of missing values
	 *
	 * Accesses a vector with indexes of missing phenotypes.  Implemented for this class.
	 *
	 * \return Vector of _size_t_ that contains indexes that correspond to missing values.
	 *
	 */
	const vector<size_t> getMisPhen() const{ return _misPhenInd; };
};

/** \brief Generic regression
 *
 * Generic implementation of a regression. By itself to be used strictly for regressions with a single predictor because it ignores the effects of other predictors.
 *
 */
class MVnormBeta : public MVnorm {
protected:
	/** \brief Predictor
	 *
	 * Points to a vector of predictor values, typically a column of the predictor matrix that is in the encapsulating Grp class.
	 *
	 */
	gsl_vector_view _X;
	/** \brief Scale parameter
	 *
	 * Typically \f$ x^{\textsc{t}}x \f$, but some exceptions exist in special cases.
	 *
	 */
	double _scale;
	/** \brief Length of the predictor */
	size_t _N;
	
	/** \brief Row index of the prior
	 *
	 * Points to the index of a row in the prior matrix
	 *
	 */
	const size_t *_upLevel; // index into the vector of priors
	
public:
	/** \brief Default constructor */
	MVnormBeta();
	/** \brief Dimension-only constructor
	 *
	 * Sets the dimension to a value, but the _vec is not assigned a target.
	 *
	 */
	MVnormBeta(const size_t d);
	
	/** \brief Univariate random constructor with vector
	 *
	 * Sets up _vec to point to the provided vector of values, and the predictor to a column in the provided matrix of predictors.
	 *
	 * \param[in] gsl_vector* vector of values
	 * \param[in] gsl_vector* vector of standard deviations
	 * \param[in] gsl_matrix* matrix of predictors
	 * \param[in] size_t& column index for the predictor matrix
	 * \param[in] gsl_rng* pointer to a PNG
	 *
	 */
	MVnormBeta(gsl_vector *b, const gsl_vector *sd, gsl_matrix *pred, const size_t &iCl, const gsl_rng *r);
	/** \brief Univariate random constructor with vector and pointer to prior
	 *
	 * Sets up _vec to point to the provided vector of values, the predictor to a column in the provided matrix of predictors, and the pointer to the corresponding row in the prior matrix.
	 *
	 * \param[in] gsl_vector* vector of values
	 * \param[in] gsl_vector* vector of standard deviations
	 * \param[in] gsl_matrix* matrix of predictors
	 * \param[in] size_t& column index for the predictor matrix
	 * \param[in] gsl_rng* pointer to a PNG
	 * \param[in] size_t& index of the appropriate row in the prior matrix
	 *
	 */
	MVnormBeta(gsl_vector *b, const gsl_vector *sd, gsl_matrix *pred, const size_t &iCl, const gsl_rng *r, const size_t &up);
	
	/** \brief Multivariate random constructor with vector
	 *
	 * Sets up _vec to point to the provided vector of values, and the predictor to a column in the provided matrix of predictors.
	 *
	 * \param[in] gsl_vector* vector of values
	 * \param[in] gsl_matrix* covariance matrix
	 * \param[in] gsl_matrix* matrix of predictors
	 * \param[in] size_t& column index for the predictor matrix
	 * \param[in] gsl_rng* pointer to a PNG
	 *
	 */
	MVnormBeta(gsl_vector *b, const gsl_matrix *Sig, gsl_matrix *pred, const size_t &iCl, const gsl_rng *r);
	/** \brief Multivariate random constructor with vector and pointer to prior
	 *
	 * Sets up _vec to point to the provided vector of values, the predictor to a column in the provided matrix of predictors, and the pointer to the corresponding row in the prior matrix.
	 *
	 * \param[in] gsl_vector* vector of values
	 * \param[in] gsl_matrix* covariance matrix
	 * \param[in] gsl_matrix* matrix of predictors
	 * \param[in] size_t& column index for the predictor matrix
	 * \param[in] gsl_rng* pointer to a PNG
	 * \param[in] size_t& index of the appropriate row in the prior matrix
	 *
	 */
	MVnormBeta(gsl_vector *b, const gsl_matrix *Sig, gsl_matrix *pred, const size_t &iCl, const gsl_rng *r, const size_t &up);
	
	/** \brief Multivariate constructor with response matrix
	 *
	 * Sets up _vec to point to a row in the provided matrix of regression coefficients, and the predictor to a column of the matrix of predictors.  Initializes regression coefficients using the provided response matrix.
	 *
	 * \param[in] gsl_matrix* response matrix
	 * \param[in] gsl_matrix* predictor martix
	 * \param[in] size_t& column index of the predictor matrix
	 * \param[in] gsl_matrix* covariance matrix
	 * \param[in] gsl_rng* pointer to a PNG
	 * \param[in] gsl_matrix* matrix of regression coefficients
	 * \param[in] size_t& row index of the regression coefficient matrix
	 *
	 */
	MVnormBeta(const gsl_matrix *resp, gsl_matrix *pred, const size_t &iCl, const gsl_matrix *Sig, const gsl_rng *r, gsl_matrix *bet, const size_t &iRw);
	/** \brief Multivariate constructor with response matrix and index for a prior
	 *
	 * Sets up _vec to point to a row in the provided matrix of regression coefficients, and the predictor to a column of the matrix of predictors.  Initializes regression coefficients using the provided response matrix.  
	 * Initializes _upLevel to point to the appropriate row in the matrix of priors.
	 *
	 * \param[in] gsl_matrix* response matrix
	 * \param[in] gsl_matrix* predictor martix
	 * \param[in] size_t& column index of the predictor matrix
	 * \param[in] gsl_matrix* covariance matrix
	 * \param[in] gsl_rng* pointer to a PNG
	 * \param[in] size_t& row index of the prior matrix
	 * \param[in] gsl_matrix* matrix of regression coefficients
	 * \param[in] size_t& row index of the regression coefficient matrix
	 *
	 */
	MVnormBeta(const gsl_matrix *resp, gsl_matrix *pred, const size_t &iCl, const gsl_matrix *Sig, const gsl_rng *r, const size_t &up, gsl_matrix *bet, const size_t &iRw);
	
	/** \brief Multivariate constructor with a Grp type response
	 *
	 * Sets up _vec to point to a row in the provided matrix of regression coefficients, and the predictor to a column of the matrix of predictors.  Initializes regression coefficients using the provided response matrix.
	 *
	 * \param[in] Grp& response
	 * \param[in] gsl_matrix* predictor martix
	 * \param[in] size_t& column index of the predictor matrix
	 * \param[in] gsl_matrix* covariance matrix
	 * \param[in] gsl_rng* pointer to a PNG
	 * \param[in] gsl_matrix* matrix of regression coefficients
	 * \param[in] size_t& row index of the regression coefficient matrix
	 *
	 */
	MVnormBeta(const Grp &resp, gsl_matrix *pred, const size_t &iCl, const gsl_matrix *Sig, const gsl_rng *r, gsl_matrix *bet, const size_t &iRw);
	/** \brief Multivariate constructor with a Grp type response and index for a prior
	 *
	 * Sets up _vec to point to a row in the provided matrix of regression coefficients, and the predictor to a column of the matrix of predictors.  Initializes regression coefficients using the provided response matrix.
	 * Initializes _upLevel to point to the appropriate row in the matrix of priors.
	 *
	 * \param[in] Grp& response
	 * \param[in] gsl_matrix* predictor martix
	 * \param[in] size_t& column index of the predictor matrix
	 * \param[in] gsl_matrix* covariance matrix
	 * \param[in] gsl_rng* pointer to a PNG
	 * \param[in] size_t& row index of the prior matrix
	 * \param[in] gsl_matrix* matrix of regression coefficients
	 * \param[in] size_t& row index of the regression coefficient matrix
	 *
	 */
	MVnormBeta(const Grp &resp, gsl_matrix *pred, const size_t &iCl, const gsl_matrix *Sig, const gsl_rng *r, const size_t &up, gsl_matrix *bet, const size_t &iRw);
	
	/** \brief Deterministic constructor
	 *
	 * Does not initialize the regression coefficients, but simply points to the already-initialized matrix of values.
	 *
	 * \param[in] gsl_matrix* predictor matrix
	 * \param[in] size_t& column index of the predictor matrix
	 * \param[in] gsl_matrix* regression coefficient matrix
	 * \param[in] size_t& row index of the regression coefficient matrix
	 *
	 */
	MVnormBeta(gsl_matrix *pred, const size_t &iCl, gsl_matrix *bet, const size_t &iRw);
	/** \brief Deterministic constructor with prior index
	 *
	 * Does not initialize the regression coefficients, but simply points to the already-initialized matrix of values.
	 * Sets the _upLevel to point to the corresponding row of the prior matrix
	 *
	 * \param[in] gsl_matrix* predictor matrix
	 * \param[in] size_t& column index of the predictor matrix
	 * \param[in] size_t& row index of the prior matrix
	 * \param[in] gsl_matrix* regression coefficient matrix
	 * \param[in] size_t& row index of the regression coefficient matrix
	 *
	 */
	MVnormBeta(gsl_matrix *pred, const size_t &iCl, const size_t &up, gsl_matrix *bet, const size_t &iRw);
	
	/** \brief Deterministic copy constructor
	 *
	 * \param[in] MVnormBeta& object to be copied
	 *
	 */
	MVnormBeta(const MVnormBeta &);
	/** \brief Deterministic assignement operator
	 *
	 * \param[in] MVnormBeta& object to be assigned
	 *
	 * \return A reference to a MVnormBeta object
	 *
	 */
	MVnormBeta & operator=(const MVnormBeta &);
	
	/** \brief Destructor */
	virtual ~MVnormBeta(); 	// destructor
	
	// improper prior methods
	virtual void update(const Grp &dat, const SigmaI &SigIb, const gsl_rng *r);
	virtual void update(const Grp &dat, const Qgrp &q, const SigmaI &SigIb, const gsl_rng *r);
	// 0-mean prior methods
	virtual void update(const Grp &dat, const SigmaI &SigIb, const SigmaI &SigIp, const gsl_rng *r); // G data, G prior
	virtual void update(const Grp &dat, const SigmaI &SigIb, const double &qPr, const SigmaI &SigIp, const gsl_rng *r); // G data, t prior
	virtual void update(const Grp &dat, const Qgrp &q, const SigmaI &SigIb, const SigmaI &SigIp, const gsl_rng *r); // t data, G prior
	virtual void update(const Grp &dat, const Qgrp &q, const SigmaI &SigIb, const double &qPr, const SigmaI &SigIp, const gsl_rng *r); // t data, t prior
	// non-zero mean prrior methods
	virtual void update(const Grp &dat, const SigmaI &SigIb, const Grp &muPr, const SigmaI &SigIp, const gsl_rng *r); // G data, G prior
	virtual void update(const Grp &dat, const SigmaI &SigIb, const Grp &muPr, const double &qPr, const SigmaI &SigIp, const gsl_rng *r); // G data, t prior
	virtual void update(const Grp &dat, const Qgrp &q, const SigmaI &SigIb, const Grp &muPr, const SigmaI &SigIp, const gsl_rng *r); // t data, G prior
	virtual void update(const Grp &dat, const Qgrp &q, const SigmaI &SigIb, const Grp &muPr, const double &qPr, const SigmaI &SigIp, const gsl_rng *r); // t data, t prior
	
	const size_t *up() const{return _upLevel; };
	double scalePar() const {return _scale; };
};

/** \brief Regression with multiple predictors
 *
 * Full implementation of a multiple regression using one-at-a-time updating. Updates regression coefficients for the current predictor, while accounting for the effects of other predictors
 *
 */
class MVnormBetaFt: public MVnormBeta {
protected:
	/** \brief Matrix of already fitted values
	 *
	 * \f$ \boldsymbol{XB} \f$ matrix for all predictors but the current one (i.e., \f$ \boldsymbol{X}_{\cdot -j}\boldsymbol{B}_{-j\cdot} \f$).  It is subtracted from the response and the regression is performed on the residual.
	 *
	 */
	gsl_matrix_view _fitted;
	
public:
	/** \brief Default constructor */
	MVnormBetaFt(): MVnormBeta() {};
	
	/** \brief Univariate random constructor with vector
	 *
	 * Sets up _vec to point to the provided vector of values, and the predictor to a column in the provided matrix of predictors.  
	 * The _fitted member points to the indicated submatrix of a matrix that contains all the partial fitted matrices.
	 *
	 * \param[in] gsl_vector* vector of values
	 * \param[in] gsl_vector* vector of standard deviations
	 * \param[in] gsl_matrix* matrix of predictors
	 * \param[in] size_t& column index for the predictor matrix
	 * \param[in] gsl_matrix* matrix with all the partial fitted matrices stacked
	 * \param[in] size_t& index of the row where the relevant fitted matrix begins
	 * \param[in] gsl_rng* pointer to a PNG
	 *
	 */
	MVnormBetaFt(gsl_vector *b, const gsl_vector *sd, gsl_matrix *pred, const size_t &iCl, gsl_matrix *allFt, const size_t &begRw, const gsl_rng *r) : MVnormBeta(b, sd, pred, iCl, r) {_fitted = gsl_matrix_submatrix(allFt, begRw, 0, pred->size1, _d); };
	/** \brief Univariate random constructor with vector and prior index
	 *
	 * Sets up _vec to point to the provided vector of values, and the predictor to a column in the provided matrix of predictors.
	 * The _fitted member points to the indicated submatrix of a matrix that contains all the partial fitted matrices. The _upLevel member is set to point to a row in the prior matrix.
	 *
	 * \param[in] gsl_vector* vector of values
	 * \param[in] gsl_vector* vector of standard deviations
	 * \param[in] gsl_matrix* matrix of predictors
	 * \param[in] size_t& column index for the predictor matrix
	 * \param[in] gsl_matrix* matrix with all the partial fitted matrices stacked
	 * \param[in] size_t& index of the row where the relevant fitted matrix begins
	 * \param[in] gsl_rng* pointer to a PNG
	 * \param[in] size_t& row index of the prior matrix
	 *
	 */
	MVnormBetaFt(gsl_vector *b, const gsl_vector *sd, gsl_matrix *pred, const size_t &iCl, gsl_matrix *allFt, const size_t &begRw, const gsl_rng *r, const size_t &up) : MVnormBeta(b, sd, pred, iCl, r, up) {_fitted = gsl_matrix_submatrix(allFt, begRw, 0, pred->size1, _d); };
	
	/** \brief Multivariate random constructor with vector
	 *
	 * Sets up _vec to point to the provided vector of values, and the predictor to a column in the provided matrix of predictors.
	 * The _fitted member points to the indicated submatrix of a matrix that contains all the partial fitted matrices.
	 *
	 * \param[in] gsl_vector* vector of values
	 * \param[in] gsl_matrix* covariance matrix
	 * \param[in] gsl_matrix* matrix of predictors
	 * \param[in] size_t& column index for the predictor matrix
	 * \param[in] gsl_matrix* matrix with all the partial fitted matrices stacked
	 * \param[in] size_t& index of the row where the relevant fitted matrix begins
	 * \param[in] gsl_rng* pointer to a PNG
	 *
	 */
	MVnormBetaFt(gsl_vector *b, const gsl_matrix *Sig, gsl_matrix *pred, const size_t &iCl, gsl_matrix *allFt, const size_t &begRw, const gsl_rng *r) : MVnormBeta(b, Sig, pred, iCl, r) {_fitted = gsl_matrix_submatrix(allFt, begRw, 0, pred->size1, _d); };
	/** \brief Multivariate random constructor with vector and prior index
	 *
	 * Sets up _vec to point to the provided vector of values, and the predictor to a column in the provided matrix of predictors.
	 * The _fitted member points to the indicated submatrix of a matrix that contains all the partial fitted matrices. The _upLevel member is set to point to a row in the prior matrix.
	 *
	 * \param[in] gsl_vector* vector of values
	 * \param[in] gsl_matrix* covariance matrix
	 * \param[in] gsl_matrix* matrix of predictors
	 * \param[in] size_t& column index for the predictor matrix
	 * \param[in] gsl_matrix* matrix with all the partial fitted matrices stacked
	 * \param[in] size_t& index of the row where the relevant fitted matrix begins
	 * \param[in] gsl_rng* pointer to a PNG
	 * \param[in] size_t& row index of the prior matrix
	 *
	 */
	MVnormBetaFt(gsl_vector *b, const gsl_matrix *Sig, gsl_matrix *pred, const size_t &iCl, gsl_matrix *allFt, const size_t &begRw, const gsl_rng *r, const size_t &up) : MVnormBeta(b, Sig, pred, iCl, r, up) {_fitted = gsl_matrix_submatrix(allFt, begRw, 0, pred->size1, _d); };
	
	/** \brief Multivariate random constructor with response matrix
	 *
	 * Sets up _vec to point to the provided vector of values, and the predictor to a column in the provided matrix of predictors.   Initializes regression coefficients using the provided response matrix.
	 * The _fitted member points to the indicated submatrix of a matrix that contains all the partial fitted matrices.
	 *
	 * \param[in] gsl_matrix* response matrix
	 * \param[in] gsl_matrix* predictor matrix
	 * \param[in] size_t& column index for the predictor matrix
	 * \param[in] gsl_matrix* matrix with all the partial fitted matrices stacked
	 * \param[in] size_t& index of the row where the relevant fitted matrix begins
	 * \param[in] gsl_matrix* covariance matrix
	 * \param[in] gsl_rng* pointer to a PNG
	 * \param[in] gsl_matrix* matrix of regression coefficients
	 * \param[in] size_t& row index of the regression coefficient matrix
	 *
	 */
	MVnormBetaFt(const gsl_matrix *resp, gsl_matrix *pred, const size_t &iCl, gsl_matrix *allFt, const size_t &begRw, const gsl_matrix *Sig, const gsl_rng *r, gsl_matrix *bet, const size_t &iRw) : MVnormBeta(resp, pred, iCl, Sig, r, bet, iRw) {_fitted = gsl_matrix_submatrix(allFt, begRw, 0, pred->size1, _d); };
	/** \brief Multivariate random constructor with response matrix and prior index
	 *
	 * Sets up _vec to point to the provided vector of values, and the predictor to a column in the provided matrix of predictors. Initializes regression coefficients using the provided response matrix.
	 * The _fitted member points to the indicated submatrix of a matrix that contains all the partial fitted matrices.  The _upLevel member is set to point to a row in the prior matrix.
	 *
	 * \param[in] gsl_matrix* response matrix
	 * \param[in] gsl_matrix* predictor matrix
	 * \param[in] size_t& column index for the predictor matrix
	 * \param[in] gsl_matrix* matrix with all the partial fitted matrices stacked
	 * \param[in] size_t& index of the row where the relevant fitted matrix begins
	 * \param[in] gsl_matrix* covariance matrix
	 * \param[in] gsl_rng* pointer to a PNG
	 * \param[in] size_t& row index of the prior matrix
	 * \param[in] gsl_matrix* matrix of regression coefficients
	 * \param[in] size_t& row index of the regression coefficient matrix
	 *
	 */
	MVnormBetaFt(const gsl_matrix *resp, gsl_matrix *pred, const size_t &iCl, gsl_matrix *allFt, const size_t &begRw, const gsl_matrix *Sig, const gsl_rng *r, const size_t &up, gsl_matrix *bet, const size_t &iRw) : MVnormBeta(resp, pred, iCl, Sig, r, up, bet, iRw) {_fitted = gsl_matrix_submatrix(allFt, begRw, 0, pred->size1, _d); };
	
	/** \brief Multivariate random constructor with response matrix
	 *
	 * Sets up _vec to point to the provided vector of values, and the predictor to a column in the provided matrix of predictors. Initializes regression coefficients using the provided response matrix.
	 * The _fitted member points to the indicated vector that has the appropriate partial fitted values.
	 *
	 * \param[in] gsl_matrix* response matrix
	 * \param[in] gsl_matrix* predictor matrix
	 * \param[in] size_t& column index for the predictor matrix
	 * \param[in] vector<double>& vectorized partial fitted matrix
	 * \param[in] gsl_matrix* covariance matrix
	 * \param[in] gsl_rng* pointer to a PNG
	 * \param[in] gsl_matrix* matrix of regression coefficients
	 * \param[in] size_t& row index of the regression coefficient matrix
	 *
	 */
	MVnormBetaFt(const gsl_matrix *resp, gsl_matrix *pred, const size_t &iCl, vector<double> &eaFt, const gsl_matrix *Sig, const gsl_rng *r, gsl_matrix *bet, const size_t &iRw) : MVnormBeta(resp, pred, iCl, Sig, r, bet, iRw) {_fitted = gsl_matrix_view_array(eaFt.data(), pred->size1, _d); };
	/** \brief Multivariate random constructor with response matrix and prior index
	 *
	 * Sets up _vec to point to the provided vector of values, and the predictor to a column in the provided matrix of predictors. Initializes regression coefficients using the provided response matrix.
	 * The _fitted member points to the indicated vector that has the appropriate partial fitted values. The _upLevel member is set to point to a row in the prior matrix.
	 *
	 * \param[in] gsl_matrix* response matrix
	 * \param[in] gsl_matrix* predictor matrix
	 * \param[in] size_t& column index for the predictor matrix
	 * \param[in] vector<double>& vectorized partial fitted matrix
	 * \param[in] gsl_matrix* covariance matrix
	 * \param[in] gsl_rng* pointer to a PNG
	 * \param[in] size_t& row index of the prior matrix
	 * \param[in] gsl_matrix* matrix of regression coefficients
	 * \param[in] size_t& row index of the regression coefficient matrix
	 *
	 */
	MVnormBetaFt(const gsl_matrix *resp, gsl_matrix *pred, const size_t &iCl, vector<double> &eaFt, const gsl_matrix *Sig, const gsl_rng *r, const size_t &up, gsl_matrix *bet, const size_t &iRw) : MVnormBeta(resp, pred, iCl, Sig, r, up, bet, iRw) {_fitted = gsl_matrix_view_array(eaFt.data(), pred->size1, _d); };
	
	/** \brief Multivariate random constructor with Grp type response
	 *
	 * Sets up _vec to point to the provided vector of values, and the predictor to a column in the provided matrix of predictors.   Initializes regression coefficients using the provided response matrix.
	 * The _fitted member points to the indicated submatrix of a matrix that contains all the partial fitted matrices.
	 *
	 * \param[in] Grp& response
	 * \param[in] gsl_matrix* predictor matrix
	 * \param[in] size_t& column index for the predictor matrix
	 * \param[in] gsl_matrix* matrix with all the partial fitted matrices stacked
	 * \param[in] size_t& index of the row where the relevant fitted matrix begins
	 * \param[in] gsl_matrix* covariance matrix
	 * \param[in] gsl_rng* pointer to a PNG
	 * \param[in] gsl_matrix* matrix of regression coefficients
	 * \param[in] size_t& row index of the regression coefficient matrix
	 *
	 */
	MVnormBetaFt(const Grp &resp, gsl_matrix *pred, const size_t &iCl, gsl_matrix *allFt, const size_t &begRw, const gsl_matrix *Sig, const gsl_rng *r, gsl_matrix *bet, const size_t &iRw) : MVnormBeta(resp, pred, iCl, Sig, r, bet, iRw) {_fitted = gsl_matrix_submatrix(allFt, begRw, 0, pred->size1, _d); }; // deprecated
	/** \brief Multivariate random constructor with Grp type response and prior index
	 *
	 * Sets up _vec to point to the provided vector of values, and the predictor to a column in the provided matrix of predictors. Initializes regression coefficients using the provided response matrix.
	 * The _fitted member points to the indicated submatrix of a matrix that contains all the partial fitted matrices.  The _upLevel member is set to point to a row in the prior matrix.
	 *
	 * \param[in] Grp& response
	 * \param[in] gsl_matrix* predictor matrix
	 * \param[in] size_t& column index for the predictor matrix
	 * \param[in] gsl_matrix* matrix with all the partial fitted matrices stacked
	 * \param[in] size_t& index of the row where the relevant fitted matrix begins
	 * \param[in] gsl_matrix* covariance matrix
	 * \param[in] gsl_rng* pointer to a PNG
	 * \param[in] size_t& row index of the prior matrix
	 * \param[in] gsl_matrix* matrix of regression coefficients
	 * \param[in] size_t& row index of the regression coefficient matrix
	 *
	 */
	MVnormBetaFt(const Grp &resp, gsl_matrix *pred, const size_t &iCl, gsl_matrix *allFt, const size_t &begRw, const gsl_matrix *Sig, const gsl_rng *r, const size_t &up, gsl_matrix *bet, const size_t &iRw) : MVnormBeta(resp, pred, iCl, Sig, r, up, bet, iRw) {_fitted = gsl_matrix_submatrix(allFt, begRw, 0, pred->size1, _d); };
	
	/** \brief Multivariate random constructor with Grp type response
	 *
	 * Sets up _vec to point to the provided vector of values, and the predictor to a column in the provided matrix of predictors. Initializes regression coefficients using the provided response matrix.
	 * The _fitted member points to the indicated vector that has the appropriate partial fitted values.
	 *
	 * \param[in] Grp& response
	 * \param[in] gsl_matrix* predictor matrix
	 * \param[in] size_t& column index for the predictor matrix
	 * \param[in] vector<double>& vectorized partial fitted matrix
	 * \param[in] gsl_matrix* covariance matrix
	 * \param[in] gsl_rng* pointer to a PNG
	 * \param[in] gsl_matrix* matrix of regression coefficients
	 * \param[in] size_t& row index of the regression coefficient matrix
	 *
	 */
	MVnormBetaFt(const Grp &resp, gsl_matrix *pred, const size_t &iCl, vector<double> &eaFt, const gsl_matrix *Sig, const gsl_rng *r, gsl_matrix *bet, const size_t &iRw) : MVnormBeta(resp, pred, iCl, Sig, r, bet, iRw) {_fitted = gsl_matrix_view_array(eaFt.data(), pred->size1, _d); };
	/** \brief Multivariate random constructor with Grp type response and prior index
	 *
	 * Sets up _vec to point to the provided vector of values, and the predictor to a column in the provided matrix of predictors. Initializes regression coefficients using the provided response matrix.
	 * The _fitted member points to the indicated vector that has the appropriate partial fitted values. The _upLevel member is set to point to a row in the prior matrix.
	 *
	 * \param[in] Grp& response
	 * \param[in] gsl_matrix* predictor matrix
	 * \param[in] size_t& column index for the predictor matrix
	 * \param[in] vector<double>& vectorized partial fitted matrix
	 * \param[in] gsl_matrix* covariance matrix
	 * \param[in] gsl_rng* pointer to a PNG
	 * \param[in] size_t& row index of the prior matrix
	 * \param[in] gsl_matrix* matrix of regression coefficients
	 * \param[in] size_t& row index of the regression coefficient matrix
	 *
	 */
	MVnormBetaFt(const Grp &resp, gsl_matrix *pred, const size_t &iCl, vector<double> &eaFt, const gsl_matrix *Sig, const gsl_rng *r, const size_t &up, gsl_matrix *bet, const size_t &iRw) : MVnormBeta(resp, pred, iCl, Sig, r, up, bet, iRw) {_fitted = gsl_matrix_view_array(eaFt.data(), pred->size1, _d); };
	
	/** \brief Deterministic constructor
	 *
	 * Does not initialize the regression coefficients, but simply points to the already-initialized matrix of values.
	 * The _fitted member points to the indicated vector that has the appropriate partial fitted values.
	 *
	 * \param[in] gsl_matrix* predictor matrix
	 * \param[in] size_t& column index of the predictor matrix
	 * \param[in] vector<double>& vectorized partial fitted matrix
	 * \param[in] gsl_matrix* regression coefficient matrix
	 * \param[in] size_t& row index of the regression coefficient matrix
	 *
	 */
	MVnormBetaFt(gsl_matrix *pred, const size_t &iCl, vector<double> &eaFt, gsl_matrix *bet, const size_t &iRw) : MVnormBeta(pred, iCl, bet, iRw) {_fitted = gsl_matrix_view_array(eaFt.data(), pred->size1, _d); };
	/** \brief Deterministic constructor with a prior index
	 *
	 * Does not initialize the regression coefficients, but simply points to the already-initialized matrix of values.
	 * The _fitted member points to the indicated vector that has the appropriate partial fitted values.  The _upLevel member is set to point to a row in the prior matrix.
	 *
	 * \param[in] gsl_matrix* predictor matrix
	 * \param[in] size_t& column index of the predictor matrix
	 * \param[in] vector<double>& vectorized partial fitted matrix
	 * \param[in] size_t& row index of the prior matrix
	 * \param[in] gsl_matrix* regression coefficient matrix
	 * \param[in] size_t& row index of the regression coefficient matrix
	 *
	 */
	MVnormBetaFt(gsl_matrix *pred, const size_t &iCl, vector<double> &eaFt, const size_t &up, gsl_matrix *bet, const size_t &iRw) : MVnormBeta(pred, iCl, up, bet, iRw) {_fitted = gsl_matrix_view_array(eaFt.data(), pred->size1, _d); };
	
	/** \brief Deterministic copy constructor
	 *
	 * \param[in] MVnormBetaFt& object to be copied
	 *
	 */
	MVnormBetaFt(const MVnormBetaFt &);
	/** \brief Deterministic assignment operator
	 *
	 * \param[in] MVnormBetaFt& object to be copied
	 *
	 * \return Refernce to an object of class MVnormBetaFt
	 *
	 */
	MVnormBetaFt & operator=(const MVnormBetaFt &);
	
	/** \brief Destructor */
	virtual ~MVnormBetaFt() {};
	
	// update() methods
	// Gaussian model, improper prior (for "fixed effect" regression)
	void update(const Grp &resp, const SigmaI &SigIb, const gsl_rng *r);
	// Student-t model, improper prior (for "fixed effect" regression).  To use for replicate-level covariate regression
	void update(const Grp &resp, const Qgrp &q, const SigmaI &SigIb, const gsl_rng *r);
	
	// Gaussian model, 0-mean Gaussian prior
	void update(const Grp &resp, const SigmaI &SigIb, const SigmaI &SigIp, const gsl_rng *r);
	// Gaussian model, 0-mean Student-t prior
	void update(const Grp &resp, const SigmaI &SigIb, const double &qPr, const SigmaI &SigIp, const gsl_rng *r);
	// Student-t model, 0-mean Gaussian prior
	void update(const Grp &resp, const Qgrp &q, const SigmaI &SigIb, const SigmaI &SigIp, const gsl_rng *r);
	// Student-t model, 0-mean Student-t prior
	void update(const Grp &resp, const Qgrp &q, const SigmaI &SigIb, const double &qPr, const SigmaI &SigIp, const gsl_rng *r);
	
	// Gaussian model, Gaussian prior
	void update(const Grp &resp, const SigmaI &SigIb, const Grp &muPr, const SigmaI &SigIp, const gsl_rng *r);
	// Gaussian model, Student-t prior
	void update(const Grp &resp, const SigmaI &SigIb, const Grp &muPr, const double &qPr, const SigmaI &SigIp, const gsl_rng *r);
	// Student-t model, Gaussian prior
	void update(const Grp &resp, const Qgrp &q, const SigmaI &SigIb, const Grp &muPr, const SigmaI &SigIp, const gsl_rng *r);
	// Student-t model, Student-t prior
	void update(const Grp &resp, const Qgrp &q, const SigmaI &SigIb, const Grp &muPr, const double &qPr, const SigmaI &SigIp, const gsl_rng *r);
	
};
/** \brief Individual vector of means with blocks of traits
 *
 * Implements separate models for blocks of traits.  The likelihood covariance matrix is block-diagonal.  Traits of the same block have to be contiguous within the vector.
 *
 */
class MVnormMuBlk : public MVnorm {
protected:
	/** \brief Pointer to a row index of the prior
	 *
	 * This has to be the same for all the blocks.
	 *
	 */
	const size_t *_upLevel;
	/** \brief Start positions of blocks
	 *
	 * Pointer to the vector that's in the corresponding Grp class.
	 *
	 */
	const vector< size_t > *_blkStart;
	/** \brief Trait blocks
	 *
	 * Vector views of vector pieces, each corresponding to a block of variables. Pointer to the vector that's in the corresponding Grp class.
	 *
	 */
	vector< gsl_vector_view > _eachVec;
	/** \brief Data matrix row indexes
	 *
	 * Each block of traits can have a different number and identity of rows in the data matrix it refers to.  Each vector of *size_t* in this vector corresponds to a block, so the size of this should be the same as the size of *_eachVec*.
	 * This member is a pointer to the vector that's in the corresponding Grp class.
	 *
	 */
	const vector< vector<size_t> > *_eachLL;
	
public:
	/** \brief Default constructor */
	MVnormMuBlk() : MVnorm() {};
	/** \brief Deterministic constructor
	 *
	 * Sets up the member variables to point to blocks of traits in the matrix _mn_, but does not perform stochastic intitialization.
	 *
	 * \param[in] gsl_matrix* mean values matrix
	 * \param[in] size_t& row index of the mean values matrix
	 * \param[in] vector<size_t>& vector of start indexes for each block
	 * \param[in] size_t& row index of the prior matrix
	 *
	 */
	MVnormMuBlk(gsl_matrix *mn, const size_t &iRw, const vector<size_t> &blkStart, const size_t &up);
	/** \brief Univariate stochastic constructor
	 *
	 * Sets initial values independently for each trait, modifying the matrix row that corresponds to this vector.
	 *
	 * \param[in] gsl_matrix* mean values matrix
	 * \param[in] size_t& row index of the mean values matrix
	 * \param[in] gsl_vector* vector of standard deviations
	 * \param[in] gsl_rng* pointer to a PNG
	 * \param[in] vector<size_t>& vector of start indexes for each block
	 * \param[in] vector< vector<size_t> >& vector of lower-level index vectors
	 * \param[in] size_t& row index of the prior matrix
	 *
	 */
	MVnormMuBlk(gsl_matrix *mn, const size_t &iRw, const gsl_vector *sd, const gsl_rng *r, const vector<size_t> &blkStart, const vector< vector<size_t> > &eachLL, const size_t &up);
	
	/** \brief Destructor */
	virtual ~MVnormMuBlk() {};
	
	const size_t *up() const{return _upLevel; };
	
	void update(const Grp &dat, const SigmaI &SigIm, const gsl_rng *r);
	void update(const Grp &dat, const Qgrp &q, const SigmaI &SigIm, const gsl_rng *r);
	// 0-mean prior methods
	void update(const Grp &dat, const SigmaI &SigIm, const SigmaI &SigIp, const gsl_rng *r);
	void update(const Grp &dat, const SigmaI &SigIm, const double &qPr, const SigmaI &SigIp, const gsl_rng *r);
	void update(const Grp &dat, const Qgrp &q, const SigmaI &SigIm, const SigmaI &SigIp, const gsl_rng *r);
	void update(const Grp &dat, const Qgrp &q, const SigmaI &SigIm, const double &qPr, const SigmaI &SigIp, const gsl_rng *r);
	// non-zero mean prior methods
	void update(const Grp &dat, const SigmaI &SigIm, const Grp &muPr, const SigmaI &SigIp, const gsl_rng *r);
	void update(const Grp &dat, const SigmaI &SigIm, const Grp &muPr, const double &qPr, const SigmaI &SigIp, const gsl_rng *r);
	void update(const Grp &dat, const Qgrp &q, const SigmaI &SigIm, const Grp &muPr, const SigmaI &SigIp, const gsl_rng *r);
	void update(const Grp &dat, const Qgrp &q, const SigmaI &SigIm, const Grp &muPr, const double &qPr, const SigmaI &SigIp, const gsl_rng *r);
};

/** \brief Individual vector of regression coefficients with blocks of traits
 *
 * Implements separate regression models for blocks of traits.  The likelihood covariance matrix is block-diagonal.  Traits of the same block have to be contiguous within the vector.  
 *
 */
class MVnormBetaBlk : public MVnorm {
protected:
	/** \brief Start positions of blocks
	 *
	 * Pointer to the vector that's in the corresponding Grp class.
	 *
	 */
	const vector< size_t > *_blkStart;
	/** \brief Trait blocks
	 *
	 * Vector views of vector pieces, each corresponding to a block of variables. Pointer to the vector that's in the corresponding Grp class.
	 *
	 */
	vector< gsl_vector_view > _eachVec;
	/** \brief Separate predictors
	 *
	 * Vector views of predictor vectors, each corresponding to a block of variables. The predictor matrix is in the encompassing Grp class.
	 *
	 */
	vector< gsl_vector_view > _X;
	/** \brief Vector of scales
	 *
	 * Typically \f$X^{T}X\f$, but can bet something else in special cases.  Each _X has a correspodning _scale.
	 *
	 */
	vector<double> _scale;
	
	/** \brief Pointer to a row index of the prior
	 *
	 * This has to be the same for all the blocks.
	 *
	 */
	const size_t *_upLevel;

	
public:
	/** \brief Default constructor 
	 *
	 * Simply calls the MVnorm default constructor.
	 *
	 */
	MVnormBetaBlk() : MVnorm () {};
	/** \brief Constructor with no prior index
	 *
	 * Stochastic constructor for a regression with an improper prior.
	 *
	 * \param[in] Grp& response variable
	 * \param[in] gsl_matrix* predictor matrix
	 * \param[in] vector<size_t>& column indexes of the predictor matrix
	 * \param[in] gsl_matrix* covariance marix for stochastic initiation
	 * \param[in] vector<size_t>& vector of start indixes of each block
	 * \param[in] gsl_rng* pointer to a PNG
	 * \param[in] gsl_matrix* matrix of regression coefficients
	 * \param[in] size_t& row index of the regression coefficient matrix
	 *
	 */
	MVnormBetaBlk(const Grp &resp, gsl_matrix *pred, const vector<size_t> &iCl, const gsl_matrix *Sig, const vector<size_t> &blkStart, const gsl_rng *r, gsl_matrix *bet, const size_t &iRw);
	/** \brief Constructor with a prior index
	 *
	 * Stochastic constructor for a regression with a proper prior.
	 *
	 * \param[in] Grp& response variable
	 * \param[in] gsl_matrix* predictor matrix
	 * \param[in] vector<size_t>& column indexes of the predictor matrix
	 * \param[in] gsl_matrix* covariance marix for stochastic initiation
	 * \param[in] vector<size_t>& vector of start indixes of each block
	 * \param[in] gsl_rng* pointer to a PNG
	 * \param[in] size_t& row index of the prior matrix
	 * \param[in] gsl_matrix* matrix of regression coefficients
	 * \param[in] size_t& row index of the regression coefficient matrix
	 *
	 */
	MVnormBetaBlk(const Grp &resp, gsl_matrix *pred, const vector<size_t> &iCl, const gsl_matrix *Sig, const vector<size_t> &blkStart, const gsl_rng *r, const size_t &up, gsl_matrix *bet, const size_t &iRw);
	/** \brief Constructor with no prior index
	 *
	 * Stochastic constructor for a regression with an improper prior.
	 *
	 * \param[in] gsl_matrix* response matrix
	 * \param[in] gsl_matrix* predictor matrix
	 * \param[in] vector<size_t>& column indexes of the predictor matrix
	 * \param[in] gsl_matrix* covariance marix for stochastic initiation
	 * \param[in] vector<size_t>& vector of start indixes of each block
	 * \param[in] gsl_rng* pointer to a PNG
	 * \param[in] gsl_matrix* matrix of regression coefficients
	 * \param[in] size_t& row index of the regression coefficient matrix
	 *
	 */
	MVnormBetaBlk(const gsl_matrix *resp, gsl_matrix *pred, const vector<size_t> &iCl, const gsl_matrix *Sig, const vector<size_t> &blkStart, const gsl_rng *r, gsl_matrix *bet, const size_t &iRw);
	/** \brief Constructor with a prior index
	 *
	 * Stochastic constructor for a regression with a proper prior.
	 *
	 * \param[in] gsl_matrix* response matrix
	 * \param[in] gsl_matrix* predictor matrix
	 * \param[in] vector<size_t>& column indexes of the predictor matrix
	 * \param[in] gsl_matrix* covariance marix for stochastic initiation
	 * \param[in] vector<size_t>& vector of start indixes of each block
	 * \param[in] gsl_rng* pointer to a PNG
	 * \param[in] size_t& row index of the prior matrix
	 * \param[in] gsl_matrix* matrix of regression coefficients
	 * \param[in] size_t& row index of the regression coefficient matrix
	 *
	 */
	MVnormBetaBlk(const gsl_matrix *resp, gsl_matrix *pred, const vector<size_t> &iCl, const gsl_matrix *Sig, const vector<size_t> &blkStart, const gsl_rng *r, const size_t &up, gsl_matrix *bet, const size_t &iRw);
	
	/** \brief Destructor */
	virtual ~MVnormBetaBlk() {};
	
	const size_t *up() const{return _upLevel; };
	
	virtual void update(const Grp &dat, const SigmaI &SigIb, const gsl_rng *r);
	virtual void update(const Grp &dat, const Qgrp &q, const SigmaI &SigIb, const gsl_rng *r);
	// 0-mean prior methods
	virtual void update(const Grp &dat, const SigmaI &SigIb, const SigmaI &SigIp, const gsl_rng *r);
	virtual void update(const Grp &dat, const SigmaI &SigIb, const double &qPr, const SigmaI &SigIp, const gsl_rng *r);
	virtual void update(const Grp &dat, const Qgrp &q, const SigmaI &SigIb, const SigmaI &SigIp, const gsl_rng *r);
	virtual void update(const Grp &dat, const Qgrp &q, const SigmaI &SigIb, const double &qPr, const SigmaI &SigIp, const gsl_rng *r);
	// non-zero mean prior methods
	virtual void update(const Grp &dat, const SigmaI &SigIb, const Grp &muPr, const SigmaI &SigIp, const gsl_rng *r);
	virtual void update(const Grp &dat, const SigmaI &SigIb, const Grp &muPr, const double &qPr, const SigmaI &SigIp, const gsl_rng *r);
	virtual void update(const Grp &dat, const Qgrp &q, const SigmaI &SigIb, const Grp &muPr, const SigmaI &SigIp, const gsl_rng *r);
	virtual void update(const Grp &dat, const Qgrp &q, const SigmaI &SigIb, const Grp &muPr, const double &qPr, const SigmaI &SigIp, const gsl_rng *r);
};

/** \brief Individual vector of conditional regression coefficients with blocks of traits
 *
 * Implements separate regression models for blocks of traits.  The likelihood covariance matrix is block-diagonal.  Traits of the same block have to be contiguous within the vector.
 * The regression is conditional on other regression coefficient estimates within a group (i.e., one-at-a-time updating).
 *
 */
class MVnormBetaFtBlk : public MVnormBetaBlk {
protected:
	/** \brief Matrix of fitted values
	 *
	 * \f$ \boldsymbol{XB} \f$ matrix for all predictors but the current one (i.e., \f$ \boldsymbol{X}_{\cdot -j}\boldsymbol{B}_{-j\cdot} \f$).  It is subtracted from the response and the regression is performed on the residual.
	 *
	 */
	gsl_matrix_view _fitted;
	
public:
	/** \brief Default constructor
	 *
	 * Simply calls the default constructor of the MVnormBetaBlk class.  Pointer to the fitted matrix is not set to anything.
	 *
	 */
	MVnormBetaFtBlk() : MVnormBetaBlk() {};
	/** \brief Constructor with no prior index
	 *
	 * Stochastic constructor for a regression with an improper prior.
	 *
	 * \param[in] Grp& response variable
	 * \param[in] gsl_matrix* predictor matrix
	 * \param[in] vector<double>& vectorized fitted matrix
	 * \param[in] vector<size_t>& column indexes of the predictor matrix
	 * \param[in] gsl_matrix* covariance marix for stochastic initiation
	 * \param[in] vector<size_t>& vector of start indixes of each block
	 * \param[in] gsl_rng* pointer to a PNG
	 * \param[in] gsl_matrix* matrix of regression coefficients
	 * \param[in] size_t& row index of the regression coefficient matrix
	 *
	 */
	MVnormBetaFtBlk(const Grp &resp, gsl_matrix *pred, vector<double> &eaFt, const vector<size_t> &iCl, const gsl_matrix *Sig, const vector<size_t> &blkStart, const gsl_rng *r, gsl_matrix *bet, const size_t &iRw) : MVnormBetaBlk(resp, pred, iCl, Sig, blkStart, r, bet, iRw) {_fitted = gsl_matrix_view_array(eaFt.data(), pred->size1, _d);};
	/** \brief Constructor with a prior index
	 *
	 * Stochastic constructor for a regression with a proper prior.
	 *
	 * \param[in] Grp& response variable
	 * \param[in] gsl_matrix* predictor matrix
	 * \param[in] vector<double>& vectorized fitted matrix
	 * \param[in] vector<size_t>& column indexes of the predictor matrix
	 * \param[in] gsl_matrix* covariance marix for stochastic initiation
	 * \param[in] vector<size_t>& vector of start indixes of each block
	 * \param[in] gsl_rng* pointer to a PNG
	 * \param[in] size_t& row index of the prior matrix
	 * \param[in] gsl_matrix* matrix of regression coefficients
	 * \param[in] size_t& row index of the regression coefficient matrix
	 *
	 */
	MVnormBetaFtBlk(const Grp &resp, gsl_matrix *pred, vector<double> &eaFt, const vector<size_t> &iCl, const gsl_matrix *Sig, const vector<size_t> &blkStart, const gsl_rng *r, const size_t &up, gsl_matrix *bet, const size_t &iRw) : MVnormBetaBlk(resp, pred, iCl, Sig, blkStart, r, up, bet, iRw) {_fitted = gsl_matrix_view_array(eaFt.data(), pred->size1, _d);};
	/** \brief Constructor with no prior index
	 *
	 * Stochastic constructor for a regression with an improper prior.
	 *
	 * \param[in] gsl_matrix* response matrix
	 * \param[in] gsl_matrix* predictor matrix
	 * \param[in] vector<double>& vectorized fitted matrix
	 * \param[in] vector<size_t>& column indexes of the predictor matrix
	 * \param[in] gsl_matrix* covariance marix for stochastic initiation
	 * \param[in] vector<size_t>& vector of start indixes of each block
	 * \param[in] gsl_rng* pointer to a PNG
	 * \param[in] gsl_matrix* matrix of regression coefficients
	 * \param[in] size_t& row index of the regression coefficient matrix
	 *
	 */
	MVnormBetaFtBlk(const gsl_matrix *resp, gsl_matrix *pred, vector<double> &eaFt, const vector<size_t> &iCl, const gsl_matrix *Sig, const vector<size_t> &blkStart, const gsl_rng *r, gsl_matrix *bet, const size_t &iRw) : MVnormBetaBlk(resp, pred, iCl, Sig, blkStart, r, bet, iRw) {_fitted = gsl_matrix_view_array(eaFt.data(), pred->size1, _d);};
	/** \brief Constructor with a prior index
	 *
	 * Stochastic constructor for a regression with a proper prior.
	 *
	 * \param[in] gsl_matrix* response matrix
	 * \param[in] gsl_matrix* predictor matrix
	 * \param[in] vector<double>& vectorized fitted matrix
	 * \param[in] vector<size_t>& column indexes of the predictor matrix
	 * \param[in] gsl_matrix* covariance marix for stochastic initiation
	 * \param[in] vector<size_t>& vector of start indixes of each block
	 * \param[in] gsl_rng* pointer to a PNG
	 * \param[in] size_t& row index of the prior matrix
	 * \param[in] gsl_matrix* matrix of regression coefficients
	 * \param[in] size_t& row index of the regression coefficient matrix
	 *
	 */
	MVnormBetaFtBlk(const gsl_matrix *resp, gsl_matrix *pred, vector<double> &eaFt, const vector<size_t> &iCl, const gsl_matrix *Sig, const vector<size_t> &blkStart, const gsl_rng *r, const size_t &up, gsl_matrix *bet, const size_t &iRw) : MVnormBetaBlk(resp, pred, iCl, Sig, blkStart, r, up, bet, iRw) {_fitted = gsl_matrix_view_array(eaFt.data(), pred->size1, _d);};
	
	/** \brief Destructor */
	~MVnormBetaFtBlk() {};
	
	void update(const Grp &dat, const SigmaI &SigIb, const gsl_rng *r);
	void update(const Grp &dat, const Qgrp &q, const SigmaI &SigIb, const gsl_rng *r);
	// 0-mean prior methods
	void update(const Grp &dat, const SigmaI &SigIb, const SigmaI &SigIp, const gsl_rng *r);
	void update(const Grp &dat, const SigmaI &SigIb, const double &qPr, const SigmaI &SigIp, const gsl_rng *r);
	void update(const Grp &dat, const Qgrp &q, const SigmaI &SigIb, const SigmaI &SigIp, const gsl_rng *r);
	void update(const Grp &dat, const Qgrp &q, const SigmaI &SigIb, const double &qPr, const SigmaI &SigIp, const gsl_rng *r);
	// non-zero mean prior methods
	void update(const Grp &dat, const SigmaI &SigIb, const Grp &muPr, const SigmaI &SigIp, const gsl_rng *r);
	void update(const Grp &dat, const SigmaI &SigIb, const Grp &muPr, const double &qPr, const SigmaI &SigIp, const gsl_rng *r);
	void update(const Grp &dat, const Qgrp &q, const SigmaI &SigIb, const Grp &muPr, const SigmaI &SigIp, const gsl_rng *r);
	void update(const Grp &dat, const Qgrp &q, const SigmaI &SigIb, const Grp &muPr, const double &qPr, const SigmaI &SigIp, const gsl_rng *r);

};
/** @} */
// end of lineLoc group
/** \defgroup Index Index classes
 *
 * Index classes relate model hierarchy levels to each other through indexes.  The relationships can be kept the same throughout the Gibbs sampling process.  Alternatively, they can be updated according to various models, implemented in the base and derived class.
 *
 * \note The updating of indexes has not yet been extensively tested.
 *
 *@{
 */

/** \brief Generic index class.
 *
 * Establishes relationships of location parameter levels in a hierarchy.  If the constructor is deterministic and there is no updating, this class is used simply to relate levels.  Updating methods are provided that implement Gaussian mixture models.
 *
 */
class RanIndex {

protected:
	/** \brief Indexes of lower levels
	 *
	 * This vector of vectors relates a top level of a hierarchy to the corresponding lower level.  The total number of elements is the number of elements in the top level.  Each element is a vector of row indexes into the lower level location parameter matrix.
	 * Constituent vectors may not be of the same length (corresponding to an unbalanced design).
	 *
	 */
	vector< vector<size_t> > _idx;
	/** \brief Indexes of upper levels
	 *
	 * Stores the row indexes of the upper-level location parameter matrix corresponding to each lower-level index.  The length of this vector is the same as the sum of all the vectors in *_idx*, and the number of unique values is equal to the length of *_idx*.
	 *
	 */
	vector<size_t> _vecInd;        // vector that has the upper-level element IDs for each lower-level instance
	/** \brief Pointer to a PNG
	 *
	 * The PNG is initialized whether or not updating is going to be performed. We use the sum of _time(NULL)_ and RTDSC for seeding.
	 *
	 */
	gsl_rng *_r;
	
public:
	/** \brief Default constructor
	 *
	 * The PNG is initialized, _idx is set to length 0, and _vecInd has only one element set to 0.
	 *
	 */
	RanIndex();
	/** \brief Vector-based constructor
	 *
	 * The GSL vector of integers has the indexes of the upper level (number of unique values is _Nup_), and is the same length as the number of rows in the lower-level location parameter matrix (_Ntot_).
	 *
	 * \param[in] gsl_vector_int* GSL vector of index information
	 * \param[in] size_t& number of elements in the lower level
	 * \param[in] size_t& number of elements in the upper level
	 *
	 */
	RanIndex(const gsl_vector_int *lInd, const size_t &Ntot, const size_t &Nup);
	/** \brief Constructor with one upper level
	 *
	 * For cases where all the lower-level elemants have the same prior.
	 *
	 * \param[in] size_t& number of lower-level elements
	 *
	 */
	RanIndex(const size_t &Ntot);
	/** \brief Constructor for single-element lower levels
	 *
	 * Each element of the lower level has a separate corresponding upper level (prior).  Typically the two numbers passed will be the same, but the one-argument constructor is already used for setting up the single-prior index.
	 * If the number of elements in the upper level is bigger than that in the lower level, an error is issued.  If the number of upper elements is smaller than the _Ntot_, only the first _Nup_ elements of the lower level are used and the rest are ignored with a warning.
	 *
	 */
	RanIndex(const size_t &Ntot, const size_t &Nup);
	/** \brief Constructor with file input
	 *
	 * Reads the index information from a file. Base-0 indexing is generally assumed, but checks are attempted.  Since there are cases when indexes that look non-base-0 are needed (such as when only a subset of lower-level rows are accessed), these generate warnings rather than errors, unless there are upper-level index values outside the set range.
	 *
	 * \param[in] size_t& number of lower-level elements
	 * \param[in] size_t& number of upper-level elements
	 * \param[in] string& file name
	 *
	 */
	RanIndex(const size_t &Ntot, const size_t &Nup, const string &fileNam);
	/** \brief Constructor with file-stream input
	 *
	 * Reads the index information from a file. Base-0 indexing is generally assumed, but checks are attempted.  Since there are cases when indexes that look non-base-0 are needed (such as when only a subset of lower-level rows are accessed), these generate warnings rather than errors, unless there are upper-level index values outside the set range.
	 *
	 * \param[in] size_t& number of lower-level elements
	 * \param[in] size_t& number of upper-level elements
	 * \param[in] FILE* file stream name
	 *
	 */
	RanIndex(const size_t &Ntot, const size_t &Nup, FILE *fileStr);
	
	/** \brief Destructor */
	virtual ~RanIndex() {gsl_rng_free(_r);};
	
	/** \brief Subscript operator
	 *
	 * Returns a vector of lower-level indexes for a given upper-level index. This is the _const_ version.
	 *
	 * \param[in] size_t upper-level index value
	 * \return const vector<size_t>& vector of lower-level indexes
	 *
	 */
	const vector<size_t> &operator[](const size_t i) const{return _idx[i];};
	/** \brief Subscript operator
	 *
	 * Returns a vector of lower-level indexes for a given upper-level index.
	 *
	 * \param[in] size_t upper-level index value
	 * \return vector<size_t>& vector of lower-level indexes
	 *
	 */
	vector<size_t> &operator[](const size_t i) {return _idx[i];};
	
	/** \brief Retrieve the index vector
	 *
	 * Provides access to the vector of indexes into the upper level.  This is the _const_ version.
	 *
	 * \return const vector<size_t> vector of upper-level indexes
	 *
	 */
	const vector<size_t> &getIndVec() const{return _vecInd; };
	/** \brief Retrieve the index vector
	 *
	 * Provides access to the vector of indexes into the upper level.
	 *
	 * \return const vector<size_t> vector of upper-level indexes
	 *
	 */
	vector<size_t> &getIndVec() {return _vecInd; };
	
	/** \brief Upper-level index
	 *
	 * Returns the upper-level index that corresponds to the given lower-level index.  This is the _const_ version.
	 *
	 * \param[in] size_t lower-level index
	 * \return const size_t upper-level index
	 *
	 */
	const size_t &priorInd(const size_t i) const{return _vecInd[i]; };
	/** \brief Upper-level index
	 *
	 * Returns the upper-level index that corresponds to the given lower-level index.
	 *
	 * \param[in] size_t lower-level index
	 * \return const size_t upper-level index
	 *
	 */
	size_t &priorInd(const size_t i) {return _vecInd[i]; };
	
	/** \brief Number of lower-level elements
	 *
	 * Returns the number of rows in the lower-level location matrix.  This is the _const_ version.
	 *
	 * \return size_t number of lower-level elements
	 *
	 */
	size_t getNtot() const{return _vecInd.size(); };
	/** \brief Number of lower-level elements
	 *
	 * Returns the number of rows in the lower-level location matrix.
	 *
	 * \return size_t number of lower-level elements
	 *
	 */
	size_t getNtot(){return _vecInd.size(); };
	
	/** \brief Number of upper-level elements
	 *
	 * Returns the number of rows in the upper-level (prior) matrix. This is the _const_ version.
	 *
	 * \return size_t number of upper-level elements
	 *
	 */
	size_t getNgrp() const{return _idx.size(); };
	/** \brief Number of upper-level elements
	 *
	 * Returns the number of rows in the upper-level (prior) matrix.
	 *
	 * \return size_t number of upper-level elements
	 *
	 */
	size_t getNgrp(){return _idx.size(); };
	
	/** \brief Proportions of each group
	 *
	 * Returns the fraction of the total number of lower-level elements that fall into each of the upper-level groups.
	 *
	 * \return vector<double> a vector of proportions
	 *
	 */
	virtual const vector<double> props() const;
	
	/** \brief Initialization function
	 *
	 * Deterministically re-initializes the index with a new vector of vectors. Used for some mixture models.
	 *
	 * \param[in] vector< vector<size_t> >& vector to replace the current _idx
	 * \param[in] vector< vector<size_t> >& a relationship vector; ignored in this class
	 *
	 */
	virtual void init(const vector< vector<size_t> > &idx, const vector< vector<size_t> > &rLD);
	/** \brief Variable selection save
	 *
	 * Saves probablities of belonging to a group. For now, only implemented in the derived class.
	 *
	 * \param[in] Grp& data
	 * \param[in] BetaGrpBVSR* location parameters
	 * \param[in] SigmaI& inverse-covariance matrix
	 *
	 */
	virtual void save(const Grp &y, const BetaGrpBVSR *theta, const SigmaI &SigIe) {};
	/** \brief Probability save
	 *
	 * Saves probablities of belonging to a group. For now, only implemented in the derived class.
	 *
	 * \param[in] Grp& data
	 * \param[in] Grp& location parameters
	 * \param[in] SigmaI& inverse-covariance matrix
	 *
	 */
	virtual void save(const Grp &y, const Grp &theta, const SigmaI &SigIe) {};
	/** \brief Dump results to a file
	 *
	 * Dumping the mean values for group probabilites to a file. For now, only implemented in the derived class.
	 *
	 */
	virtual void dump(){};
	
	/** \brief Update mixture model with multiple covariances.
	 *
	 * \ingroup updateFun
	 *
	 * Updates the group relationships for Gaussian mixture models when each group has a different mean and covariance.
	 *
	 * \param[in] Grp& data
	 * \param[in] Grp& group means
	 * \param[in] vector<SigmaI>& vector of inverse-covariances
	 * \param[in] MixP& prior proportions
	 *
	 */
	void update(const Grp &theta, const Grp &mu, const vector<SigmaI> &SigI, const MixP &p);
	/** \brief Update mixture model with a single covariance.
	 *
	 * \ingroup updateFun
	 *
	 * Updates the group relationships for Gaussian mixture models when each group has a different mean but all covrariances are the same.
	 *
	 * \param[in] Grp& data
	 * \param[in] Grp& group means
	 * \param[in] SigmaI& common inverse-covariance
	 * \param[in] MixP& prior proportions
	 *
	 */
	void update(const Grp &theta, const Grp &mu, const SigmaI &SigI, const MixP &p);
	/** \brief Variable selection update
	 *
	 * \ingroup updateFun
	 *
	 * Update for a mixture that includes point-mass at 0.  Only implemented in the derived class.
	 *
	 * \param[in] Grp& data
	 * \param[in] SigmaI& likelihood inverse-covariance
	 * \param[in] BetaGrpBVSR* location parameter values
	 * \param[in] SigmaI& prior inverse-covariance
	 *
	 */
	virtual void update(const Grp &y, const SigmaI &SigIe, BetaGrpBVSR *theta, const SigmaI &SigIp) {};
	
};

/** \brief BVSR index class
 *
 * Keeps track of the variable selection process for a multivariate version of the Bayesian Variable Selection Regression (BVSR) multi-marker regression method of Guan and Stephens \cite guan11 (Greenberg, unpublished).   This clas is a friend of BetaGrpBVSR.
 *
 * \warning This class is experimental and has not been extensively tested.
 *
 */
class RanIndexVS : public RanIndex {
	
protected:
	/** \brief Posterior exclusion probabilities
	 *
	 * Posterior _exclusion_ probabilities.  This is the opposite if the implementation in Guan and Stephens, where the inclusion probabilities are exported.  My implementation results in output that is analogous to \f$ -\log_{10}p \f$, making it more intuitive to people used to frequentist approaches.
	 * The probabilities stored in this variable are in the order of the SNPs picked for initial inclusion in the BetaGrpBVSR class.  We keep adding PEP values to this at each save, then dump the mean into a file in the end, giving a Rao-Blackwellized point estimate of PEP.
	 *
	 */
	vector<double> _pep;
	/** \brief Transition tracking variable
	 *
	 * Used to track the Metropolis moves made after the current MCMC step. There are four elements, three last ones being mutually exclusive: [0]: accepted? [1]: proposed adding a predictor? [2]: proposed dropping a predictor? [3]: proposed swapping predictors?
	 *
	 */
	vector<bool> _mcmcTrack;
	/** \brief Mapping selected to original predictors
	 *
	 * Relates the new positions of selected predictors to the original positions.  Its length is the same as the selected number of predictors.  _relLD[i][0] is the original position of the selected predictor _i_; the rest of _relLD[i] (if any) are positions of linked SNPs (or, in general, correlated predictors)
	 *
	 */
	vector< vector<size_t> > _relLD;
	/** \brief Original number of predictors */
	size_t _Ntp;
	
	/** \brief Prior proportions
	 *
	 * Prior proportion of predictors in selected to be in the model.
	 */
	MixP *_prior;
	
	/** \brief Accept a drop
	 *
	 * The state of _mcmcTrack that corresponds to accepting a drop of a predictor.  Pre-set so that it is easy to modify _mcmcTrack.
	 *
	 */
	vector<bool> _acceptDrop;
	/** \brief Reject a drop
	 *
	 * The state of _mcmcTrack that corresponds to rejecting a drop of a predictor.  Pre-set so that it is easy to modify _mcmcTrack.
	 *
	 */
	vector<bool> _rejectDrop;
	/** \brief Accept an addition
	 *
	 * The state of _mcmcTrack that corresponds to accepting an addition of a predictor.  Pre-set so that it is easy to modify _mcmcTrack.
	 *
	 */
	vector<bool> _acceptAdd;
	/** \brief Rject an addition
	 *
	 * The state of _mcmcTrack that corresponds to rejecting an addition of a predictor.  Pre-set so that it is easy to modify _mcmcTrack.
	 *
	 */
	vector<bool> _rejectAdd;
	/** \brief Accept a swap
	 *
	 * The state of _mcmcTrack that corresponds to accepting a swap of predictors.  Pre-set so that it is easy to modify _mcmcTrack.
	 *
	 */
	vector<bool> _acceptSwap;
	/** \brief Reject a swap
	 *
	 * The state of _mcmcTrack that corresponds to rejecting a swap of predictors.  Pre-set so that it is easy to modify _mcmcTrack.
	 *
	 */
	vector<bool> _rejectSwap;
	
	/** \brief Number of calls to save()
	 *
	 * Tracks the number of times a save function is called.  Before dumping the PEP results, divide each element of _pep by this number.
	 *
	 */
	double _numSaves;
	/** \brief PEP output file name */
	string _pepOutFlnam;
	/** \brief Accept/reject file name
	 *
	 * Name of the file where the accept/reject and proposal kind statistics are stored.  This file is appended at each save.  These statistics are useful for checking and trouble-shooting Metropolis chain behavior.
	 */
	string _mcmcPutFlNam;
	
	/** \brief Rank proposal-generating function
	 *
	 * The rank-based Metropolis proposal-generating function proposed in Guan and Stephens \cite guan11 .
	 *
	 * \warning This function is not presently used in my implementation of BVSR because I am not sure it is a symmetric proposal.  If it is not, then we would need to come up with a Metropolis-Hastings updating scheme.
	 */
	size_t _proposal(const size_t &Ntot, const int &Nmn, const gsl_rng *r); // rank proposal-generating function for Metropolis updating; not clear if it is symmetric so don't use for now

public:
	/** \brief Default constructor */
	RanIndexVS();
	/** \brief Constructor with a prior
	 *
	 * This is a preliminary set-up.  The initiation is finished with the init() member function, after pre-screening of predictors is performed.
	 *
	 */
	RanIndexVS(const size_t &Ntot, const string &outFlNam, MixP &pr);
	/** \brief Constructor without a prior
	 *
	 * This is a preliminary set-up.  The initiation is finished with the init() member function, after pre-screening of predictors is performed.
	 *
	 */
	RanIndexVS(const size_t &Ntot, const string &outFlNam);
	
	/** \brief Destructor */
	~RanIndexVS(){};
	
	void props(vector<double> &prp) const;
	
	/** \brief Save PEP
	 *
	 * Stores current PEP values and saves Metropolis proposal statistics to a file.
	 *
	 * \param[in] Grp& data
	 * \param[in] BetaGrpBVSR* location parameters
	 * \param[in] SigmaI& inverse-covariance matrix
	 *
	 */
	void save(const Grp &y, const BetaGrpBVSR *theta, const SigmaI &SigIe);
	/** \brief Save PEP
	 *
	 * Stores current PEP values and saves Metropolis proposal statistics to a file.
	 *
	 * \param[in] Grp& data
	 * \param[in] Grp& location parameters
	 * \param[in] SigmaI& inverse-covariance matrix
	 *
	 */
	void save(const Grp &y, const Grp &theta, const SigmaI &SigIe);
	/** \brief Save PEP to a file
	 *
	 * Saves posterior exclusion probabilities (PEP) to a file. Name of the file is stored in the _pepOutFlnam variable.
	 *
	 */
	void dump();
	/** \brief Complete initialization
	 *
	 * Initializes the index instance after preliminary screening of predictors in performed by the corresponding BetaGrpBVSR variable.
	 *
	 * \param[in] vector< vector<size_t> >& vector to replace the current _idx
	 * \param[in] vector< vector<size_t> >& a relationship vector; ignored in this class
	 *
	 */
	void init(const vector< vector<size_t> > &idx, const vector< vector<size_t> > &rLD); // idx is _idx, rLD is _relLD
	
	void update(const Grp &y, const SigmaI &SigIe, BetaGrpBVSR *theta, const SigmaI &SigIp);
};

/** @} */

/** \brief Multiplicative redundant parameter
 *
 * Implements multiplicative redundant parametrization for analogs of frequentist random-effects models.  It is a multivariate extension of well-established multiplicative data augmentation schemes \cite gelman07 \cite gelman08 (Greenberg, unpublished).
 * Here, instead of a scalar redundant parameter, I am using a redundant parameter matrix.  This approach introduces additional per-iteration computational burden, but results in much faster model convergence in some cases (Greenberg, unpublished).
 * Therefore, it should only be used when convergence problems are severe and cannot be eliminated with simpler re-parametrizations.
 *
 */
class Apex {
private:
	/** \brief Redundant prameter matrix
	 *
	 * A square matrix with dimensions equal to the number of traits.
	 */
	gsl_matrix *_Amat;
	/** \brief Inverse-prior matrix
	 *
	 * Diagonal matrix with the same dimensions as _Amat. Diagonal values are set to a small value if a vague prior is desired.
	 */
	gsl_matrix *_SigIpr;
	/** \brief Each fitted value
	 *
	 * Each row of _Amat is a vector of regression coefficients updated given all other rows. This pointer addresses a vector of the same length as the dimensions of _Amat contains vectorized fitted matrices of all the other coefficients, \f$ \boldsymbol{\Xi}_{\cdot -m}\boldsymbol{A}_{-m\cdot} \f$.
	 */
	vector<vector<double> > *_fitEach;
	/** \brief Pointer to a PNG
	 *
	 * Seeded on construction with the sum of _time(NULL)_ and RTDSC.
	 */
	gsl_rng *_r;
	
public:
	/** \brief Default constructor 
	 *
	 * All matrices are set to be \f$ 1 \times 1 \f$ with the element equal to 1.
	 */
	Apex();
	/** \brief Dimension-only constructor
	 *
	 * Initializes a vague prior with diagonal elements set to \f$ 10^{-6} \f$.
	 *
	 * \param[in] size_t& dimension (i.e. number of traits)
	 * \param[in] vector<vector <double> >& fitted values
	 */
	Apex(const size_t &d, vector<vector <double> > &fE);
	/** \brief Diagonal prior constructor
	 *
	 * Initializes a prior with diagonal elements set to the given value.
	 *
	 * \param[in] double& inverse-prior value
	 * \param[in] size_t& dimension (i.e. number of traits)
	 * \param[in] vector<vector <double> >& fitted values
	 */
	Apex(const double &dV, const size_t &d, vector<vector <double> > &fE);
	/** \brief Covariance prior constructor
	 *
	 * Initializes a prior with the given inverse-covariance matrix. Dimension is set to the dimension of the prior matrix.
	 *
	 * \param[in] SigmaI& inverse-prior matrix
	 * \param[in] vector<vector <double> >& fitted values
	 */
	Apex(const SigmaI &SigI, vector<vector <double> > &fE);
	
	/** \brief Destructor */
	~Apex();
	
	/** \brief Copy constructor
	 *
	 * \param[in] Apex& object to be copied
	 */
	Apex(const Apex &A);
	/** \brief Assignement constructor
	 *
	 * \param[in] Apex& object to be copied
	 * \return Apex object
	 */
	Apex &operator=(const Apex &A);
	
	/** \brief Get redundant parameter matrix */
	gsl_matrix *getMat(){return _Amat; };
	/** \brief Get redundant parameter matrix */
	const gsl_matrix *getMat() const{return _Amat; };
	/** \brief Set prior value
	 *
	 * Sets the diagonal of the inverse-prior matrix to the given value
	 *
	 * \param[in] double& inverse-prior value
	 *
	 */
	void setPr(const double &pr);
	
	/** \brief Unreplicated Gaussian update
	 *
	 * \ingroup updateFun
	 *
	 * Gaussian data with no replication. 
	 *
	 * \param[in] Grp& data
	 * \param[in] gsl_matrix* "raw" location parameter matrix
	 * \param[in] SigmaI& data inverse-covariance matrix
	 *
	 */
	void update(const Grp &y, const gsl_matrix *xi, const SigmaI &SigIm);
	/** \brief Replicated Gaussian update
	 *
	 * \ingroup updateFun
	 *
	 * Gaussian data with replication, i.e., the number of rows in \f$ \boldsymbol{Y} \f$ is larger than the number of rows in \f$ \boldsymbol{\Xi} \f$.
	 *
	 * \param[in] Grp& data
	 * \param[in] gsl_matrix* "raw" location parameter matrix
	 * \param[in] SigmaI& data inverse-covariance matrix
	 * \param[in] RanIndex& index relating data rows to rows in \f$ \boldsymbol{\Xi} \f$
	 *
	 */
	void update(const Grp &y, const gsl_matrix *xi, const SigmaI &SigIm, const RanIndex &ind);
	/** \brief Replicated Student-\f$t\f$ update
	 *
	 * \ingroup updateFun
	 *
	 * Student-\f$t\f$ data with replication, i.e., the number of rows in \f$ \boldsymbol{Y} \f$ is larger than the number of rows in \f$ \boldsymbol{\Xi} \f$.
	 *
	 * \param[in] Grp& data
	 * \param[in] gsl_matrix* "raw" location parameter matrix
	 * \param[in] Qgrp& Student-\f$t\f$ weights
	 * \param[in] SigmaI& data inverse-covariance matrix
	 * \param[in] RanIndex& index relating data rows to rows in \f$ \boldsymbol{\Xi} \f$
	 *
	 */
	void update(const Grp &y, const gsl_matrix *xi, const Qgrp &q, const SigmaI &SigIm, const RanIndex &ind);
};

/** \defgroup arithmetic Arithmetic operators
 *
 * Arithmetic operators for Grp type location parameters.  The matrices addressed by the fMat() member functions are added.  These are essential to build Gibbs samplers with multiple parameters at the same level of hierarchy, such as variable-intercept regressions \cite gelman07 . However, care must be taken because only certain combinations of objects are compatible. These are
 *
 * 1. fMat() matrices have the same number of rows.  In this case, they are simply added or subtracted as is.
 * 2. One of the matrices is a single-row matrix.  In this case, the row is added to each row of the bigger matrix (or subtracted in the appropriate direction).  The resulting MuGrp object will have the same size value matrix as the bigger fMat().
 * 3. The RanIndex to the lower level of the Grp (or MuGrp) object with the smaller fMat() points to the same number of rows as in the fMat() of the larger object.  In this case, the RanIndex is used to relate the rows of the larger matrix to those of the smaller matrix, the rows of the smaller matrix are re-used as necessary. The resulting MuGrp object has a value matrix of the same size as the larger fMat().
 * 4. None of the above apply, but the RanIndex to the lower level of each object points to the same number of rows.  In this case, a MuGrp object with a value matrix with that number of rows is created, and the addition/subtraction is performed with rows corresponing to the same lower level of the RanIndex.
 *
 * In all other cases, an error is issued and execution is aborted. The number of traits (columns) has to always be the same. Addition is commutative, subtraction is anti-commutative.
 *
 * @{
 */
/** \brief Addition operator
 *
 * \param[in] Grp& first summand
 * \param[in] Grp& second summand
 * \return MuGrp object that is the sum of the two input objects
 *
 */
MuGrp operator+(const Grp &m1, const Grp &m2);
/** \brief Addition operator
 *
 * \param[in] MuGrp& first summand
 * \param[in] Grp& second summand
 * \return MuGrp object that is the sum of the two input objects
 *
 */
MuGrp operator+(const MuGrp &m1, const Grp &m2);
/** \brief Addition operator
 *
 * \param[in] Grp& first summand
 * \param[in] MuGrp& second summand
 * \return MuGrp object that is the sum of the two input objects
 *
 */
MuGrp operator+(const Grp &m1, const MuGrp &m2);
/** \brief Subtraction operator
 *
 * \param[in] Grp& minuend
 * \param[in] Grp& subtrahend
 * \return MuGrp object that is the difference between the two input objects
 *
 */
MuGrp operator-(const Grp &m1, const Grp &m2);
/** \brief Subtraction operator
 *
 * \param[in] MuGrp& minuend
 * \param[in] Grp& subtrahend
 * \return MuGrp object that is the difference between the two input objects
 *
 */
MuGrp operator-(const MuGrp &m1, const Grp &m2);
/** \brief Subtraction operator
 *
 * \param[in] Grp& minuend
 * \param[in] MuGrp& subtrahend
 * \return MuGrp object that is the difference between the two input objects
 *
 */
MuGrp operator-(const Grp &m1, const MuGrp &m2);

/**@} */

/** \defgroup locGrp Groups of location parameters
 *
 * Location parameters grouped by common modeling features. They have common priors, common data and prior covariances, and if they are regression coefficients a common set of predictors. 
 * Values are stored in a matrix with the columns representing traits (or analogous parameters) and rows -- individual location parameters.  The latter are targets of MVnorm classes and are typically modified in Markov chains through update functions implemented in these individual-row objects.
 *
 * @{
 */
/** \brief Base location parameter group class
 *
 * The abstract base location parameter group class.  Cannot be initialized directly.
 *
 */
class Grp {
	friend MuGrp operator+(const Grp &m1, const Grp &m2);
	friend MuGrp operator+(const MuGrp &m1, const Grp &m2);
	friend MuGrp operator+(const Grp &m1, const MuGrp &m2);
	friend MuGrp operator-(const Grp &m1, const Grp &m2);
	friend MuGrp operator-(const MuGrp &m1, const Grp &m2);
	friend MuGrp operator-(const Grp &m1, const MuGrp &m2);
	friend class MuGrp;
	
protected:
	/** \brief Vector of pointers to value rows
	 *
	 * Each individual MVnorm pointer corresponds to a row in _valueMat, which can be modified by invoking update functions that MVnorm class members.
	 *
	 */
	vector<MVnorm *> _theta;
	/** \brief Value matrix
	 *
	 * Matrix of location parameter values.  The columns are traits, and rows are individual replicates or regression coefficients corresponding to a given predictor.  Each row is addressed by an individual element of _theta for most classes.  Exceptions are noted in descriptions of those derived classes.
	 */
	gsl_matrix *_valueMat;
	/** \brief Lower level index
	 *
	 * If initialized, points to the lower (i.e., data) level in the hierarchy. Otherwise, set to 0.  For regressions, plays the role of the design matrix \f$ \boldsymbol{Z} \f$.
	 *
	 */
	RanIndex *_lowLevel;
	/** \brief Upper level index
	 *
	 * If initialized, points to the upper (i.e., prior) level in the hierarchy. Otherwise, set to 0.
	 *
	 */
	RanIndex *_upLevel;
	/** \brief Vector of PNG pointers
	 *
	 * Typically is of unit length.  However, for derived classes that use multi-threading its length is equal to the number of threads, each thread having its own PNG. PNGS are seeded on construction with the sum of _time(NULL)_ and RTDSC.
	 */
	vector<gsl_rng *> _rV;
	
	/** \brief Name of the output file */
	string _outFlNam;
	
	/** Default constructor */
	Grp();
	
public:
	/** \brief Destructor */
	virtual ~Grp();
	
	/** \brief Gaussian likelihood, improper prior
	 *
	 * \ingroup updateFun
	 * \ingroup locIP
	 *
	 * \param[in] Grp& data
	 * \param[in] SigmaI& data inverse-covariance
	 *
	 */
	virtual void update(const Grp &, const SigmaI &) = 0;
	/** \brief Student-\f$t\f$ likelihood, improper prior
	 *
	 * \ingroup updateFun
	 * \ingroup locIP
	 *
	 * \param[in] Grp& data
	 * \param[in] Qgrp& Student-\f$t\f$ weight parameter for data
	 * \param[in] SigmaI& data inverse-covariance
	 *
	 */
	virtual void update(const Grp &, const Qgrp &, const SigmaI &) = 0;
	/** \brief Gaussian likelihood, 0-mean Gaussian prior
	 *
	 * \ingroup updateFun
	 * \ingroup locZMn
	 *
	 * \param[in] Grp& data
	 * \param[in] SigmaI& data inverse-covariance
	 * \param[in] SigmaI& prior inverse-covariance
	 *
	 */
	virtual void update(const Grp &, const SigmaI &, const SigmaI &) = 0;
	/** \brief Student-\f$t\f$ likelihood, 0-mean Gaussian prior
	 *
	 * \ingroup updateFun
	 * \ingroup locZMn
	 *
	 * \param[in] Grp& data
	 * \param[in] Qgrp& Student-\f$t\f$ weight parameter for data
	 * \param[in] SigmaI& data inverse-covariance
	 * \param[in] SigmaI& prior inverse-covariance
	 *
	 */
	virtual void update(const Grp &, const Qgrp &, const SigmaI &, const SigmaI &) = 0;
	/** \brief Gaussian likelihood, 0-mean Student-\f$t\f$ prior
	 *
	 * \ingroup updateFun
	 * \ingroup locZMn
	 *
	 * \param[in] Grp& data
	 * \param[in] SigmaI& data inverse-covariance
	 * \param[in] Qgrp& Student-\f$t\f$ weight parameter for the prior
	 * \param[in] SigmaI& prior inverse-covariance
	 *
	 */
	virtual void update(const Grp &, const SigmaI &, const Qgrp &, const SigmaI &) = 0;
	/** \brief Student-\f$t\f$ likelihood, 0-mean Student-\f$t\f$ prior
	 *
	 * \ingroup updateFun
	 * \ingroup locZMn
	 *
	 * \param[in] Grp& data
	 * \param[in] Qgrp& Student-\f$t\f$ weight parameter for the data
	 * \param[in] SigmaI& data inverse-covariance
	 * \param[in] Qgrp& Student-\f$t\f$ weight parameter for the prior
	 * \param[in] SigmaI& prior inverse-covariance
	 *
	 */
	virtual void update(const Grp &, const Qgrp &, const SigmaI &, const Qgrp &, const SigmaI &) = 0;
	/** \brief Gaussian likelihood, non-zero mean Gaussian prior
	 *
	 * For the relationship to work properly, the _upLevel index of the focal object has to point to the rows of the prior mean matrix.  This is the matrix addressed by dMat().  The relationship to the data is dependent on the derived class.
	 *
	 * \ingroup updateFun
	 * \ingroup locNZMn
	 *
	 * \param[in] Grp& data
	 * \param[in] SigmaI& data inverse-covariance
	 * \param[in] Grp& prior mean
	 * \param[in] SigmaI& prior inverse-covariance
	 *
	 */
	virtual void update(const Grp &, const SigmaI &, const Grp &, const SigmaI &) = 0;
	/** \brief Student-\f$t\f$ likelihood, non-zero mean Gaussian prior
	 *
	 * For the relationship to work properly, the _upLevel index of the focal object has to point to the rows of the prior mean matrix.  This is the matrix addressed by dMat().  The relationship to the data is dependent on the derived class.
	 *
	 * \ingroup updateFun
	 * \ingroup locNZMn
	 *
	 * \param[in] Grp& data
	 * \param[in] Qgrp& Student-\f$t\f$ weight parameter for the data
	 * \param[in] SigmaI& data inverse-covariance
	 * \param[in] Grp& prior mean
	 * \param[in] SigmaI& prior inverse-covariance
	 *
	 */
	virtual void update(const Grp &, const Qgrp &, const SigmaI &, const Grp &, const SigmaI &) = 0;
	/** \brief Gaussian likelihood, non-zero mean Student-\f$t\f$ prior
	 *
	 * For the relationship to work properly, the _upLevel index of the focal object has to point to the rows of the prior mean matrix.  This is the matrix addressed by dMat().  The relationship to the data is dependent on the derived class.
	 *
	 * \ingroup updateFun
	 * \ingroup locNZMn
	 *
	 * \param[in] Grp& data
	 * \param[in] SigmaI& data inverse-covariance
	 * \param[in] Grp& prior mean
	 * \param[in] Qgrp& Student-\f$t\f$ weight parameter for the prior
	 * \param[in] SigmaI& prior inverse-covariance
	 *
	 */
	virtual void update(const Grp &, const SigmaI &, const Grp &, const Qgrp &, const SigmaI &) = 0;
	/** \brief Student-\f$t\f$ likelihood, non-zero mean Student-\f$t\f$ prior
	 *
	 * For the relationship to work properly, the _upLevel index of the focal object has to point to the rows of the prior mean matrix.  This is the matrix addressed by dMat().  The relationship to the data is dependent on the derived class.
	 *
	 * \ingroup updateFun
	 * \ingroup locNZMn
	 *
	 * \param[in] Grp& data
	 * \param[in] Qgrp& Student-\f$t\f$ weight parameter for the data
	 * \param[in] SigmaI& data inverse-covariance
	 * \param[in] Grp& prior mean
	 * \param[in] Qgrp& Student-\f$t\f$ weight parameter for the prior
	 * \param[in] SigmaI& prior inverse-covariance
	 *
	 */
	virtual void update(const Grp &, const Qgrp &, const SigmaI &, const Grp &, const Qgrp &, const SigmaI &) = 0;
	
	/** \brief Save to pre-specified file
	 *
	 * Saves a sample from the Markov chain for the object by appending to a file.  Unless re-implemented, the current contents of _valueMat are saved. File name is either generic or pre-specified at construction.
	 */
	virtual void save();
	/** \brief Save to file
	 *
	 * Saves a sample from the Markov chain for the object by appending to the named file.  Unless re-implemented, the current contents of _valueMat are saved.
	 *
	 * \param[in] string& file name
	 */
	virtual void save(const string &outFlNam);
	/** \brief Joint save
	 *
	 * Saves the current value of _valueMat to one file and the covariance matrix corresponding to the given inverse-covariance to the other.  Both files are appended.  Has more practical importance in some derived classes.
	 *
	 * \param[in] string& mean file name
	 * \param[in] string& covariance file name
	 * \param[in] SigmaI& inverse-covariance
	 *
	 */
	virtual void save(const string &outMuFlNam, const string &outSigFlNam, const SigmaI &SigI);
	/** \brief Save with inverse-covariance
	 *
	 * Reserved for saving to a variable rather than a file.  Has an effect only in classes where it is re-implemented.
	 *
	 * \param[in] SigmaI& inverse-covariance
	 *
	 */
	virtual void save(const SigmaI &SigI) {};
	/** \brief Save with data and inverse-covariance
	 *
	 * Reserved for saving to a variable rather than a file.  Has an effect only in classes where it is re-implemented.
	 *
	 * \param[in] Grp& data
	 * \param[in] SigmaI& inverse-covariance
	 *
	 */
	virtual void save(const Grp &y, const SigmaI &SigI) {};
	/** \brief Save Mahalanobis distance
	 *
	 * Saves Mahalanobis distances of each row to a vector of zeros, given the provided inverse-covariance.
	 *
	 * \param[in] string& file name
	 * \param[in] SigmaI& inverse-covariance
	 *
	 */
	void mhlSave(const string &outFlNam, const SigmaI SigI);
	
	/** \brief Dump to a file
	 *
	 * Dumps paramter values averaged over the MCMC run to a file in the end of the run.  Has an effect only in derived classes where it is implemented.  In these classes, the name of the file is pre-specified at initialization.
	 *
	 */
	virtual void dump(){}; // dump to a file in cases where save() saves to a matrix
	
	/** \brief Get vector of row pointers
	 *
	 * \return vector<MVnorm *> vector of objects addressing rows of the value matrix
	 */
	const vector<MVnorm *> &dataVec() const{return _theta; };
	/** \brief Access the value matrix
	 *
	 * Gives access to the value matrix.  The internal structure this points to depends on the derived class.
	 *
	 * \return gsl_matrix* pointer to a GSL matrix
	 */
	virtual const gsl_matrix *dMat() const{return _valueMat; };
	/** \brief Access the value matrix
	 *
	 * Gives access to the value matrix. The internal structure this points to depends on the derived class, and may be different from dMat().
	 *
	 * \return gsl_matrix* pointer to a GSL matrix
	 */
	virtual const gsl_matrix *fMat() const{return _valueMat; };
	
	/** \brief Get number of rows
	 *
	 * \return size_t number of rows in the value matrix, corresponds to the number of mean values or regression coefficients
	 */
	const size_t Ndata() const{return _valueMat->size1; };
	/** \brief Get number of traits
	 *
	 * \return size_t number of columns in the value matrix, corresponds to the number of traits
	 */
	const size_t phenD() const{return _valueMat->size2; };
	
	/** \brief Log-odds ratio
	 *
	 * Log-odds ration between models with and without the i-th regression coefficient (row).  Makes sense only in regression classes, everywhere else returns zero.
	 *
	 * \param[in] Grp& data
	 * \param[in] SigmaI& data inverse-covariance
	 * \param[in] size_t index of the element (row)
	 *
	 * \return double log-odds ratio
	 */
	virtual double lnOddsRat(const Grp &y, const SigmaI &SigI, const size_t i) const {return 0.0; };
	
	/** \brief Subscript operator
	 *
	 * Get a MVnorm object pointer to a row of the value matrix.
	 *
	 * \param[in] size_t row index
	 *
	 * \return MVnorm* pointer to an element in the value matrix
	 */
	const MVnorm * operator[](const size_t i) const{return _theta[i]; };
	/** \brief Subscript operator
	 *
	 * Get a MVnorm object pointer to a row of the value matrix.
	 *
	 * \param[in] size_t row index
	 *
	 * \return MVnorm* pointer to an element in the value matrix
	 */
	MVnorm * operator[](const size_t i) {return _theta[i]; };
	
	/** \brief Group mean
	 *
	 * Calculates within-group mean values of the value matrix (by row).  The groups are defined by the upper-level of the given RanIndex variable.  The lower-level number of elements of the index has to be the same as the number of rows in the current object's value matrix.
	 *
	 * \param[in] RanIndex& grouping index
	 * \return MuGrp object with the matrix of means
	 */
	virtual MuGrp mean(RanIndex &grp);
	/** \brief Group mean
	 *
	 * Calculates within-group mean values of the value matrix (by row).  The groups are defined by the upper-level of the given RanIndex variable.  The lower-level number of elements of the index has to be the same as the number of rows in the current object's value matrix.
	 *
	 * \param[in] RanIndex& grouping index
	 * \return MuGrp object with the matrix of means
	 */
	virtual const MuGrp mean(RanIndex &grp) const;
	
	/** \brief Group weighted mean
	 *
	 * Calculates within-group weighted mean values of the value matrix (by row).  The groups are defined by the upper-level of the given RanIndex variable.  The lower-level number of elements of the index has to be the same as the number of rows in the current object's value matrix.
	 * The weights are in the provided Qgrp object.  Typically, they are from a Student-\f$t\f$ model.
	 *
	 * \param[in] RanIndex& grouping index
	 * \param[in] Qgrp& weights
	 * \return MuGrp object with the matrix of means
	 */
	virtual MuGrp mean(RanIndex &grp, const Qgrp &q);
	/** \brief Group weighted mean
	 *
	 * Calculates within-group weighted mean values of the value matrix (by row).  The groups are defined by the upper-level of the given RanIndex variable.  The lower-level number of elements of the index has to be the same as the number of rows in the current object's value matrix.
	 * The weights are in the provided Qgrp object.  Typically, they are from a Student-\f$t\f$ model.
	 *
	 * \param[in] RanIndex& grouping index
	 * \param[in] Qgrp& weights
	 * \return MuGrp object with the matrix of means
	 */
	virtual const MuGrp mean(RanIndex &grp, const Qgrp &q) const;
	
	/** \brief Center the value matrix 
	 *
	 * Centering is performed by subtracting the mean for each column from each column in place.
	 */
	void center(){ colCenter(_valueMat); };
	
};

/** \brief Hierarchical mean
 *
 * Standard hierarchical mean parameter of Bayesian hierarchical models \cite gelman04 \cite gelman07 .  Can also be used to store raw data (i.e., the lowest level in the hierarchy or the repsonse in a mixed model; in which case it is not updated), and to implement a Bayesian analog of a frequentist random effect (in which case the prior is zero).
 *
 */
class MuGrp : public Grp {
	friend MuGrp operator+(const Grp &m1, const Grp &m2);
	friend MuGrp operator+(const MuGrp &m1, const Grp &m2);
	friend MuGrp operator+(const Grp &m1, const MuGrp &m2);
	friend MuGrp operator-(const Grp &m1, const Grp &m2);
	friend MuGrp operator-(const MuGrp &m1, const Grp &m2);
	friend MuGrp operator-(const Grp &m1, const MuGrp &m2);
protected:
	
public:
	/** \brief Default constructor */
	MuGrp() : Grp(){};
	/** \brief Deterministic zero-value constructor
	 *
	 * Sets all elements of the value matrix to zero.  Can be used to construct a 0-mean prior. No upper level is set and thus this is not typically updated.
	 *
	 * \param[in] RanIndex& index to the lower level
	 * \param[in] size_t& number of traits
	 */
	MuGrp(RanIndex &low, const size_t &d);
	/** \brief Constructor with data from file
	 *
	 * Reading in data from a file.  The resulting value matrix has the number of rows equal to the number of groups in the low index and the number of elements in the upper index.  
	 * Index consistency is required, otherwise an error is issued and the program aborted.  The constructor is deterministic.
	 *
	 * \param[in] string& data file name
	 * \param[in] RanIndex& index to the lower (data) level
	 * \param[in] RanIndex& index to the upper (prior) level
	 * \param[in] size_t& number of traits
	 */
	MuGrp(const string &datFlNam, RanIndex &low, RanIndex &up, const size_t &d); // RanIndex's are not const because I have a pointer to it as a member
	/** \brief Constructor with data from file and no lower level
	 *
	 * Reading in data from a file.  The resulting value matrix has the number of rows equal to the number of elements in the upper index.  This constructor is deterministic and can be used to set up the lowest (i.e. data) level of the hierarchy, and not update it.
	 *
	 * \param[in] string& data file name
	 * \param[in] RanIndex& index to the upper (prior) level
	 * \param[in] size_t& number of traits
	 */
	MuGrp(const string &datFlNam, RanIndex &up, const size_t &d);
	/** \brief Constructor with a vector of MVnorm pointers
	 *
	 * Initializes a value matrix with means among the elements of the vector of MVnorm pointers, according to the low index (i.e., the given vector contains the data). Values for the matrix are sampled around the mean values.  The number returned by low.getNgrp() has to be equal to the number returned by up.getNtot(), and is equal to the number of rows in the value matrix.  Violation of any of these conditions results in an error.
	 *
	 * \param[in] vector<MVnorm *>& vector of pointers to rows of the value matrix
	 * \param[in] RanIndex& index to the lower (data) level
	 * \param[in] RanIndex& index to the upper (prior) level
	 *
	 */
	MuGrp(const vector<MVnorm *> &dat, RanIndex &low, RanIndex &up);
	/** \brief Constructor with a Grp object
	 *
	 * Initializes a value matrix with means among the rows of the matrix in Grp addressed by the dMat() member, according to the low index (i.e., the Grp object contains the data). Values for the matrix are sampled around the mean values.  The number returned by low.getNgrp() has to be equal to the number returned by up.getNtot(), and is equal to the number of rows in the value matrix.  Violation of any of these conditions results in an error.
	 *
	 * \param[in] Grp& vector of pointers to rows of the value matrix
	 * \param[in] RanIndex& index to the lower (data) level
	 * \param[in] RanIndex& index to the upper (prior) level
	 *
	 */
	MuGrp(const Grp &dat, RanIndex &low, RanIndex &up);
	/** \brief Constructor with a vector of MVnorm pointers and output file name
	 *
	 * Initializes a value matrix with means among the elements of the vector of MVnorm pointers, according to the low index (i.e., the given vector contains the data). Values for the matrix are sampled around the mean values.  The number returned by low.getNgrp() has to be equal to the number returned by up.getNtot(), and is equal to the number of rows in the value matrix.  Violation of any of these conditions results in an error.
	 *
	 * \param[in] vector<MVnorm *>& vector of pointers to rows of the value matrix
	 * \param[in] RanIndex& index to the lower (data) level
	 * \param[in] RanIndex& index to the upper (prior) level
	 * \param[in] string& name of the output file
	 *
	 */
	MuGrp(const vector<MVnorm *> &dat, RanIndex &low, RanIndex &up, const string &outFlNam);
	/** \brief Constructor with a Grp object and output file name
	 *
	 * Initializes a value matrix with means among the rows of the matrix in Grp addressed by the dMat() member, according to the low index (i.e., the Grp object contains the data). Values for the matrix are sampled around the mean values.  The number returned by low.getNgrp() has to be equal to the number returned by up.getNtot(), and is equal to the number of rows in the value matrix.  Violation of any of these conditions results in an error.
	 *
	 * \param[in] Grp& vector of pointers to rows of the value matrix
	 * \param[in] RanIndex& index to the lower (data) level
	 * \param[in] RanIndex& index to the upper (prior) level
	 * \param[in] string& name of the output file
	 *
	 */
	MuGrp(const Grp &dat, RanIndex &low, RanIndex &up, const string &outFlNam);
	/** \brief Deterministic mean constructor
	 *
	 * The value matrix of the resulting object has low.getNgrp() rows and is deterministically set to within-group means.
	 *
	 * \param[in] Grp& data object
	 * \param[in] RanIndex& index that defines groups
	 *
	 */
	MuGrp(const Grp &dat, RanIndex &low);
	/** \brief Deterministic weighted mean constructor
	 *
	 * The value matrix of the resulting object has low.getNgrp() rows and is deterministically set to within-group weighted means. Weghts are in the Qgrp object.  They are typically Student-\f$t\f$ model weights.
	 *
	 * \param[in] Grp& data object
	 * \param[in] Qgrp& weights
	 * \param[in] RanIndex& index that defines groups
	 *
	 */
	MuGrp(const Grp &dat, const Qgrp &q, RanIndex &low);
	/** \brief Deterministic constructor with a GSL matrix
	 *
	 * Indexes not set.  The given matrix is copied to the value matrix.
	 *
	 * \param[in] gsl_matrix* data matrix
	 */
	MuGrp(const gsl_matrix *dat);
	/** \brief Deterministic GSL matrix mean constructor
	 *
	 * The value matrix of the resulting object has low.getNgrp() rows and is deterministically set to within-group means.  Mostly used internally in other derived classes.
	 *
	 * \param[in] gsl_matrix* data matrix
	 * \param[in] RanIndex& index that defines groups
	 */
	MuGrp(const gsl_matrix *dat, RanIndex &low);
	/** \brief Deterministic GSL matrix weighted mean constructor
	 *
	 * The value matrix of the resulting object has low.getNgrp() rows and is deterministically set to within-group weighted means.  Mostly used internally in other derived classes.
	 *
	 * \param[in] gsl_matrix* data matrix
	 * \param[in] Qgrp& weights
	 * \param[in] RanIndex& index that defines groups
	 */
	MuGrp(const gsl_matrix *dat, const Qgrp &q, RanIndex &low);
	
	/** \brief Destructor */
	virtual ~MuGrp() {};
	
	/** \brief Copy constructor
	 *
	 * \param[in] MuGrp& object to be copied
	 * \return MuGrp& copy of the object
	 */
	MuGrp(const MuGrp &mG);
	/** \brief Copy constructor
	 *
	 * \param[in] Grp& object to be copied
	 * \return MuGrp& copy of the object
	 */
	MuGrp(const Grp &g);
	/** \brief Assignemnt operator
	 *
	 * \param[in] MuGrp& object to be copied
	 * \return MuGrp& target object
	 */
	MuGrp & operator=(const MuGrp &mG);
	
	// improper prior
	virtual void update(const Grp &dat, const SigmaI &SigIm);
	virtual void update(const Grp &dat, const Qgrp &q, const SigmaI &SigIm);
	// 0-mean prior
	virtual void update(const Grp &dat, const SigmaI &SigIm, const SigmaI &SigIp);
	virtual void update(const Grp &dat, const Qgrp &q, const SigmaI &SigIm, const SigmaI &SigIp);
	virtual void update(const Grp &dat, const SigmaI &SigIm, const Qgrp &qPr, const SigmaI &SigIp);
	virtual void update(const Grp &dat, const Qgrp &q, const SigmaI &SigIm, const Qgrp &qPr, const SigmaI &SigIp);
	// non-0-mean prior
	virtual void update(const Grp &dat, const SigmaI &SigIm, const Grp &muPr, const SigmaI &SigIp);
	virtual void update(const Grp &dat, const Qgrp &q, const SigmaI &SigIm, const Grp &muPr, const SigmaI &SigIp);
	virtual void update(const Grp &dat, const SigmaI &SigIm, const Grp &muPr, const Qgrp &qPr, const SigmaI &SigIp);
	virtual void update(const Grp &dat, const Qgrp &q, const SigmaI &SigIm, const Grp &muPr, const Qgrp &qPr, const SigmaI &SigIp);
	
};

/** \brief "Random effect" with parameter expansion
 *
 * An implementation of a multivariate extension (Greenberg, unpublished) of the multiplicative PEX scheme for "random effects" model \cite gelman07 \cite gelman08 .  This object has to be outside the model hierarchy, i.e. have the same prior for all rows.
 * Variables are divided into "raw" and "adjusted" as in \cite gelman07 . "Adjusted" variables are on the correct scale for interpretation.  The "raw" value matrix is denoted \f$ \boldsymbol{\Xi} \f$ and the redundant parameter matrix (implemented in the Apex class) is \f$ \boldsymbol{A} \f$.
 *
 */
class MuGrpPEX : public MuGrp {
protected:
	/** \brief Adjusted value matrix
	 *
	 * The location paramter of interest, i.e. \f$ \boldsymbol{\Xi A} \f$.
	 */
	gsl_matrix *_adjValMat;
	/** \brief Scaled inverse-covariance
	 *
	 * The matrix \f$ \left( \boldsymbol{\Sigma}^{-1}\boldsymbol{A}^T \right)^T \f$ that is common for sampling each row of the value matrix.
	 */
	gsl_matrix *_tSigIAt;
	/** \brief Redundant parameter matrix */
	Apex _A;
	/** \brief Fitted values
	 *
	 * Vector of vectorized individual fitted matrices \f$ \boldsymbol{\Xi}_{\cdot -m}\boldsymbol{A}_{-m\cdot} \f$.
	 *
	 */
	vector<vector<double> > _ftA;
	/** \brief number of threads */
	int _nThr;
	
	/** \brief Covariance output file name
	 *
	 * This is to save the adjusted prior covariance matrix associated with this object.  Saving from the SigmaI object gives the raw matrix.
	 */
	string _outSigFlNam;
	
	/** \defgroup fitFun Functions to update fitted values
	 *
	 * Protected member functions of the derived Grp classes that implement multiple regressions in some form. In Gibbs sampling for multiple regression, it is necessary to calculate partial fitted matrices of the form \f$ \boldsymbol{X}_{\cdot -k}\boldsymbol{B}_{-k \cdot} \f$ for each \f$k\f$-th predictor (i.e., product for all but the \f$k\f$-th predictor),
	 * as well as the complete fitted matrix \f$ \boldsymbol{C} = \boldsymbol{XB} \f$.  Here, \f$ \boldsymbol{X} \f$ is the predictor matrix and \f$ \boldsymbol{B} \f$ is the matrix of regression coefficients.  This process is typically the bottleneck of the Gibbs sampler when even a moderate number of predictors (\f$ \geq 100 \f$) are in the model.
	 * The key idea in implementing these functions is that the multiplications required to calculate all the partial matrix products
	 * are also necessary for computing the complete fitted matrix.  Thus, elements of the complete matrix
	 * \f[
	 *     c_{i,j} = \sum_k x_{i,k}\beta_{k,j}
	 * \f]
	 * are calculated first, then the individual \f$ x_{i,k}\beta_{k,j} \f$ are subtracted to get the required partial fitted matrices. Further speed-ups are achieved by parallelizing the calculations by row of the regression coefficient matrix (i.e. by individual predictor).
	 *
	 * @{
	 */
	/** \brief Update adjusted values
	 *
	 * Creates the adjusted value matrix and the individual \f$ \boldsymbol{\Xi}_{\cdot -m}\boldsymbol{A}_{-m\cdot} \f$.
	 */
	void _updateFitted();
	/** @} */
	
public:
	/** \brief Default constructor */
	MuGrpPEX();
	/** \brief Full constructor
	 *
	 * Initializing constructor that provides starting values for the location parameter and redundant parameter matrices.
	 *
	 * \param[in] Grp& data
	 * \param[in] RanIndex& index to the lower (data) level
	 * \param[in] RanIndex& index to the upper (prior) level
	 * \param[in] double& inverse-prior for the redundant parameter matrix
	 * \param[in] int& number of threads
	 *
	 */
	MuGrpPEX(const Grp &dat, RanIndex &low, RanIndex &up, const double &Spr, const int &nThr);
	/** \brief Full constructor with location parameter file name
	 *
	 * Initializing constructor that provides starting values for the location parameter and redundant parameter matrices.  Sets the name of the file where the  adjusted location parameter chains will be saved.
	 *
	 * \param[in] Grp& data
	 * \param[in] RanIndex& index to the lower (data) level
	 * \param[in] RanIndex& index to the upper (prior) level
	 * \param[in] string& output file name
	 * \param[in] double& inverse-prior for the redundant parameter matrix
	 * \param[in] int& number of threads
	 *
	 */
	MuGrpPEX(const Grp &dat, RanIndex &low, RanIndex &up, const string outMuFlNam, const double &Spr, const int &nThr);
	
	/** \brief Destructor */
	~MuGrpPEX();
	
	/** \brief Copy constructor
	 *
	 * \param[in] MuGrpPEX& object to be copied
	 * \return MuGrpPEX object
	 */
	MuGrpPEX(const MuGrpPEX &mGp);
	/** \brief Assignment operator
	 *
	 * \param[in] MuGrpPEX& object to be copied
	 * \return MuGrpPEX& target object
	 */
	MuGrpPEX &operator=(const MuGrpPEX &mGp);
	
	/** \brief Access the adjusted value matrix
	 *
	 * \return gsl_matrix* pointer to the adjusted value matrix
	 */
	const gsl_matrix *fMat() const{return _adjValMat; };
	/** \brief Save the adjusted values
	 *
	 * Appends the current adjusted mean values to the file whose name was set during construction.
	 */
	void save();
	/** \brief Save adjusted values to named file
	 *
	 * Appends the current adjusted mean values to the named file.
	 *
	 * \param[in] string& file name
	 */
	void save(const string &outFlNam);
	/** \brief Save with the covariance matrix
	 *
	 * Appends the current adjusted mean values and prior covariance matrix to the files whose names were set during construction.
	 */
	void save(const SigmaI &SigI);
	/** \brief Access the redundant parameter
	 *
	 * \return Apex& the redundant parameter matrix
	 */
	Apex &getA(){return _A; };
	/** \brief Set the inverse-prior for the redundant parameter matrix
	 *
	 * \param[in] double& inverse-prior value
	 */
	void setApr(const double &pr){_A.setPr(pr); };
	
	MuGrp mean(RanIndex &grp);
	const MuGrp mean(RanIndex &grp) const;
	MuGrp mean(RanIndex &grp, const Qgrp &q);
	const MuGrp mean(RanIndex &grp, const Qgrp &q) const;
	
	void update(const Grp &dat, const SigmaI &SigIm, const SigmaI &SigIp);
	void update(const Grp &dat, const Qgrp &q, const SigmaI &SigIm, const SigmaI &SigIp);
	void update(const Grp &dat, const SigmaI &SigIm, const Qgrp &qPr, const SigmaI &SigIp);
	void update(const Grp &dat, const Qgrp &q, const SigmaI &SigIm, const Qgrp &qPr, const SigmaI &SigIp);
	
	void update(const Grp &dat, const SigmaI &SigIm, const Grp &muPr, const SigmaI &SigIp);
	void update(const Grp &dat, const Qgrp &q, const SigmaI &SigIm, const Grp &muPr, const SigmaI &SigIp);
	void update(const Grp &dat, const SigmaI &SigIm, const Grp &muPr, const Qgrp &qPr, const SigmaI &SigIp);
	void update(const Grp &dat, const Qgrp &q, const SigmaI &SigIm, const Grp &muPr, const Qgrp &qPr, const SigmaI &SigIp);
};

/** \brief Data with missing phenotype values
 *
 * Implements missing phenotypic (responce) data imputation.  This object has to be at the bottom of the hierarchy, so there is no index to the lower level.
 */
class MuGrpMiss : public MuGrp {
protected:
	/** \brief Index of the rows with missing data */
	vector<size_t> _misInd;
	
public:
	/** \brief Default constructor */
	MuGrpMiss() : MuGrp() {};
	/** \brief Full constructor
	 *
	 * Reads data and indexes that show which data are missing from files.  Matrix index has the same dimensions as the value matrix and has "1" in places where the data are missing and "0" otherwise.
	 * The vector missing data index indicates the number of data points missing in each row (zero for rows with all data present).  The data file can have any double-precision value in place of missing data.
	 * These values are ignored and replaced by the overall mean at construction, but subsequently updated.
	 *
	 * \param[in] string& data file name
	 * \param[in] string& matrix missing data index file name
	 * \param[in] string& vector missing data index file name
	 * \param[in] RanIndex& index pointing to the prior (next level in the hierarchy)
	 * \param[in] size_t& number of traits
	 */
	MuGrpMiss(const string &datFlNam, const string &misMatFlNam, const string &misVecFlNam, RanIndex &up, const size_t &d);
	
	/** \brief Destructor */
	~MuGrpMiss() {};
	
	/** \brief Copy constructor
	 *
	 * \param[in] MuGrpMiss& object to be copied
	 * \return MuGrpMiss object
	 */
	MuGrpMiss(const MuGrpMiss &mG); // copy constructor
	/** \brief Assignment operator
	 *
	 * \param[in] MuGrpMiss& object to be copied
	 * \return MuGrpMiss& target object
	 */
	MuGrpMiss & operator=(const MuGrpMiss &mG);
	
	/** \brief Standard Gaussian imputation
	 *
	 * If some data are present, performs standard Gaussian marginal imputation (desribed in, e.g., \cite chatfield80 ) with the provided Grp object as a mean and the SigmaI object as the inverse-covariance.
	 * If no data for a row are present, simply replaces the row values by a Gaussian sample with mean and inverse-covariance provided.  Rows with no missing data are ignored.
	 *
	 * \ingroup updateFun
	 *
	 * \param[in] Grp& mean
	 * \param[in] SigmaI& inverse-covariance
	 */
	virtual void update(const Grp &mu, const SigmaI &SigIm);
	/** \brief Gaussian imputation with a prior
	 *
	 * If some data are present, performs Gaussian marginal imputation (desribed in, e.g., \cite chatfield80 ) with the provided Grp object as a mean and the SigmaI object as the inverse-covariance, but with a 0-mean prior.
	 * If no data for a row are present, simply replaces the row values by a Gaussian sample with mean and inverse-covariance provided.  Rows with no missing data are ignored.
	 *
	 * \ingroup updateFun
	 *
	 * \warning Has not been extensively tested
	 *
	 * \param[in] Grp& mean
	 * \param[in] SigmaI& inverse-covariance
	 * \param[in] SigmaI& prior inverse-covariance
	 */
	virtual void update(const Grp &mu, const SigmaI &SigIm, const SigmaI &SigIp);
	
	size_t nMis() const{return _misInd.size();};
	size_t nMis(){return _misInd.size();};
};

/** \brief Data with measurement error
 *
 * If some phenotypes are measured with known error (e.g., technical replicates that are not idividually available), these can be sampled rather than used as point estimates. A mix of traits with and without measurement error is allowed.
 * This class has to be on the bottom of the model hierarchy.
 *
 */
class MuGrpEE : public MuGrp {
protected:
	/** \brief Matrix of error inverse-variances
	 *
	 * The matrix has the same number of rows as the value matrix, and the number of columns no larger than the value matrix.  Variances are read from a file and inverted on initialization.
	 */
	gsl_matrix *_errorInvVar;
	/** \brief Matrix of means
	 *
	 * Has the same dimensions as the _errorVar matrix, and stores mean values for the variables that will be sampled.
	 */
	gsl_matrix *_meanVal;
	/** \brief Trait index
	 *
	 * Contains indexes of the traits that have measurment errors.  Only one index for all rows, i.e. once a trait has measurment error all samples must have a non-zero error variance.
	 */
	vector<size_t> _errInd;
	
public:
	/** \brief Default constructor */
	MuGrpEE();
	/** \brief Constructor with index read from a file
	 *
	 * The data, error variances and the index identifying traits with errors are read from files.  The index file must be a white-space separated text file.  Variances are inverted after reading and are checked for sanity (i.e. not smaller than machine epsilon).
	 *
	 * \param[in] string& data file name
	 * \param[in] string& variance file name
	 * \param[in] string& index values file name
	 * \param[in] RainIndex& upper level (prior) index
	 * \param[in] size_t& number of traits
	 */
	MuGrpEE(const string &datFlNam, const string &varFlNam, const string &indFlNam, RanIndex &up, const size_t &d);
	/** \brief Constructor with index vector
	 *
	 * The data and error variances are read from files, but the index identifying traits with errors is provided in a vector.  Variances are inverted after reading and are checked for sanity (i.e. not smaller than machine epsilon).
	 *
	 * \param[in] string& data file name
	 * \param[in] string& variance file name
	 * \param[in] vector<size_t>& index values file name
	 * \param[in] RainIndex& upper level (prior) index
	 * \param[in] size_t& number of traits
	 */
	MuGrpEE(const string &datFlNam, const string &varFlNam, const vector<size_t> &varInd, RanIndex &up, const size_t &d);
	
	/** \brief Destructor */
	~MuGrpEE();
	
	/** \brief Copy constructor
	 *
	 * \param[in] MuGrpEE& object to be copied
	 * \return MuGrpEE object
	 */
	MuGrpEE(const MuGrpEE &mG); // copy constructor
	/** \brief Assignment operator
	 *
	 * \param[in] MuGrpEE& object to be copied
	 * \return MuGrpEE& target object
	 */
	MuGrpEE & operator=(const MuGrpEE &mG);
	
	/** \brief Gaussian prior
	 *
	 * The Grp object contains the prior means and the SigmaI object -- the prior inverse-covariance for the sampling of data values.  The upper index of the object must have the same number of groups as the number of rows in the prior matrix addressed by fMat().
	 * While the sampling is independent, the prior inverse variances for each trait are taken from the diagonal of the inverse-covariance matrix (in the SigmaI object), and are thus influenced by any correlated traits.
	 *
	 * \ingroup updateFun
	 *
	 * \param[in] Grp& prior mean
	 * \param[in] SigmaI& prior inverse-covariance
	 */
	virtual void update(const Grp &muPr, const SigmaI &SigIm);
	/** \brief Student-\f$t\f$ prior
	 *
	 * The Grp object contains the prior means and the SigmaI object -- the prior inverse-covariance for the sampling of data values.  The upper index of the object must have the same number of groups as the number of rows in the prior matrix addressed by fMat().
	 * While the sampling is independent, the prior inverse variances for each trait are taken from the diagonal of the inverse-covariance matrix (in the SigmaI object), and are thus influenced by any correlated traits.
	 *
	 * \ingroup updateFun
	 *
	 * \param[in] Grp& prior mean
	 * \param[in] Qgrp& Student-\f$t\f$ weights
	 * \param[in] SigmaI& prior inverse-covariance
	 */
	virtual void update(const Grp &muPr, const Qgrp &q, const SigmaI &SigIm);
	
};

/** \brief Data with measurement error and missing phenotypes
 *
 * Some traits have measurment errors, some (possibly an overlapping set) are missing.  Sampling with error is done only for the present phenotypes, thus not all rows have the same number of traits with error variances.
 */
class MuGrpEEmiss : public MuGrpMiss {
protected:
	/** \brief Measurement error inverse-variances
	 *
	 * The same configuration as the variance vector of lists.  This is the subset of _valueMat that corresponds to the values to be updated with experimental error.
	 */
	vector< list<double> > _errorInvVar;
	/** \brief Means for values with errors
	 *
	 * The length of the vector is equal to the number of rows of the value matrix.  For each row, the error variances are in a list that has as many members as there are non-missing phenotypes with errors.
	 */
	vector< list<double> > _meanVal;
	/** \brief Error index
	 *
	 * The same configuration as the variance vector of lists, but containing indexes of the elements of the value matrix that have measurement errors.
	 */
	vector< list<size_t> > _missErrMat;
	
public:
	/** \brief Default constructor */
	MuGrpEEmiss() : MuGrpMiss() {};
	/** \brief Constructor with index read from a file
	 *
	 * The data, error variances and the index identifying traits with errors are read from files.  The index file must be a white-space separated text file.  The index is initially the same for each row, but then the values corresponding to missing phenotypes are dropped for relevant rows.
	 *
	 * \param[in] string& data file name
	 * \param[in] string& variance file name
	 * \param[in] string& index values file name
	 * \param[in] string& matrix missing data index file name
	 * \param[in] string& vector missing data index file name
	 * \param[in] RainIndex& upper level (prior) index
	 * \param[in] size_t& number of traits
	 */
	MuGrpEEmiss(const string &datFlNam, const string &varFlNam, const string &indFlNam, const string &misMatFlNam, const string &misVecFlNam, RanIndex &up, const size_t &d);
	/** \brief Constructor with index vector
	 *
	 * The data and error variances are read from files, but the index identifying traits with errors is provided in a vector.  The index is initially the same for each row, but then the values corresponding to missing phenotypes are dropped for relevant rows.
	 *
	 * \param[in] string& data file name
	 * \param[in] string& variance file name
	 * \param[in] vector<size_t>& index values
	 * \param[in] string& matrix missing data index file name
	 * \param[in] string& vector missing data index file name
	 * \param[in] RainIndex& upper level (prior) index
	 * \param[in] size_t& number of traits
	 */
	MuGrpEEmiss(const string &datFlNam, const string &varFlNam, const vector<size_t> &varInd, const string &misMatFlNam, const string &misVecFlNam, RanIndex &up, const size_t &d);
	
	/** \brief Destructor */
	~MuGrpEEmiss() { };
	
	/** \brief Copy constructor
	 *
	 * \param[in] MuGrpEEmiss& object to be copied
	 * \return MuGrpEEmiss object
	 */
	MuGrpEEmiss(const MuGrpEEmiss &mG); // copy constructor
	/** \brief Assignment operator
	 *
	 * \param[in] MuGrpEEmiss& object to be copied
	 * \return MuGrpEEmiss& target object
	 */
	MuGrpEEmiss & operator=(const MuGrpEEmiss &mG);
	
	void update(const Grp &muPr, const SigmaI &SigIm);
	void update(const Grp &muPr, const Qgrp &q, const SigmaI &SigIm);
	
};

/** \brief Multivariate multiple regression
 *
 * Implements multivariate (multitrait) regression with multiple predictors.  A single predictor is treated as a special case internally, the user does not have to do anything different.  A variety for penalized regression methods is available, implemented through priors and pre-selection of variables, although the latter is still experimental.
 * As the number of predictors grows, the computational burden increases and updates involving these objects become bottlenecks in the Markov chain computation.  Therefore, great care has been taken to optimize computation for this class, sometimes at the expense of error checking.
 *
 */
class BetaGrpFt : public Grp {
protected:
	/** \brief Partial fitted value matrices
	 *
	 * Each member of the vector stores the element-specific fitted matrix (\f$ \boldsymbol{X}_{\cdot -k}\boldsymbol{B}_{-k \cdot} \f$) as a vector in the row-major format, to be accessed as a matrix_view of an array. If there is only one predictor, this is empty.
	 * If there is replication (i.e. the index to the lower level is initialized), these matrices have the same number of rows as the response (equivalent to \f$ \boldsymbol{Z}\boldsymbol{X}_{\cdot -k}\boldsymbol{B}_{-k \cdot} \f$, where \f$ \boldsymbol{Z} \f$ is the design matrix).
	 */
	vector< vector<double> > _fittedEach;
	/** \brief Matrix of fitted values
	 *
	 * The \f$ \boldsymbol{XB} matrix \f$. In cases where the index to the lower level is initialized, i.e. there is replication in the response, this matrix has the same number of rows as the number of unique values of the predictor.
	 */
	gsl_matrix *_fittedAll;
	/** \brief Sample storage matrix
	 *
	 * Stores Markov chain samples of the value matrix if we want only the point estimates in the end. Stores the sum of all the estimates saved up to now; allocated only if needed (under the condition that _numSaves != 0.0)
	 */
	gsl_matrix *_valueSum;
	/** \brief Predictor matrix
	 *
	 * Individual predictors are in columns.  If there is replication, the number of rows is expanded to fit the number of rows in the response matrix (\f$ \boldsymbol{Z}\boldsymbol{X}\boldsymbol{B} \f$).
	 */
	gsl_matrix *_Xmat;
	/** \brief Number of threads */
	int _nThr;
	
	/** \brief Number of saves
	 *
	 * Number of saves made, to calculate the mean at the end.
	 */
	double _numSaves;
	
	/** \brief Update fitted values 
	 * 
	 * \ingroup fitFun
	 */
	virtual void _updateFitted();
	/** \brief Rank predictors
	 *
	 * Ranking the predictors by the size of their effects in preparation for eliminating the ones below a certain rank.
	 *
	 * \param[in] gsl_matrix* data (response)
	 * \param[in] SigmaI& data inverse-covariance
	 * \param[in] gsl_vector* vector of regression scales \f$ \left( \boldsymbol{x}^T\boldsymbol{x} \right)^{-1} \f$
	 * \param[out] gsl_permutation* predictor ranks
	 */
	void _rankPred(const gsl_matrix *y, const SigmaI &SigI, gsl_vector *XtX, gsl_permutation *prm);
	/** \brief Rank predictors with missing data.
	 *
	 * Ranking the predictors by the size of their effects in preparation for eliminating the ones below a certain rank.  Missing predictor values are labeled by a given value.
	 *
	 * \param[in] gsl_matrix* data (response)
	 * \param[in] SigmaI& data inverse-covariance
	 * \param[in] double& missing data label
	 * \param[in] gsl_vector* vector of regression scales \f$ \left( \boldsymbol{x}^T\boldsymbol{x} \right)^{-1} \f$
	 * \param[out] gsl_permutation* predictor ranks
	 */
	void _rankPred(const gsl_matrix *y, const SigmaI &SigI, const double &absLab, gsl_vector *XtX, gsl_permutation *prm);
	/** \brief Testing candidates for correlation
	 *
	 * Goes through the list of "top" predictors and eliminates lower-ranked candidates correlated with them.  If any candidates are eliminated, the list of top predictors is augmented with predictors previously discarded.
	 * The identity of the predictors eliminated for correlation is saved.  The correlation among predictors arises, for example, when estimating SNP effect in genetics (GWAS).
	 *
	 * \warning Tested only superficially
	 *
	 * \param[in] gsl_vector* \f$ \left( \boldsymbol{x}^T\boldsymbol{x} \right)^{-1} \f$
	 * \param[in] gsl_permutation* predictor ranks
	 * \param[in] double& \f$ r^2 \f$ cut-off
	 * \param[in] size_t& number of predictors to pick
	 * \param[out] vector< vector<size_t> >& index of picked predictors
	 * \param[out] vector< vector<size_t> >& index relating dropped correlated predictors to their group-defining predictor
	 * \param[out] gsl_matrix* matrix of picked predictor values
	 */
	void _ldToss(const gsl_vector *var, const gsl_permutation *prm, const double &rSqMax, const size_t &Npck, vector< vector<size_t> > &idx, vector< vector<size_t> > &rLd, gsl_matrix *Xpck);
	/** \brief Gaussian kernel
	 *
	 * Calculates the multivariate Gaussian kernel value for all regression coefficients in the model.
	 *
	 * \param[in] Grp& data
	 * \param[in] SigmaI& inverse-covariance
	 *
	 * \return double kernel value
	 */
	virtual double _MGkernel(const Grp &dat, const SigmaI &SigI) const;
	/** \brief Gaussian kernel dropping one predictor
	 *
	 * Calculates the multivariate Gaussian kernel value for all regression coefficients in the model, except for the one indicated.
	 *
	 * \param[in] Grp& data
	 * \param[in] SigmaI& inverse-covariance
	 * \param[in] size_t& index of the dropped predictor
	 *
	 * \return double kernel value
	 */
	virtual double _MGkernel(const Grp &dat, const SigmaI &SigI, const size_t &prInd) const;
public:
	/** \brief Default constructor */
	BetaGrpFt();
	/** \brief Simple constructor
	 *
	 * Reads the predictor from a file and initiates regression coefficients using the provided response data.  Number of rows of the predictor is equal to the number of rows in the response (no replication).
	 * No upper index is specified, so this object will implement a regression with an improper flat prior.  Caution is advised if the number of predictors approaches the number of rows in the response, since inference will not be well-conditioned.
	 *
	 * \param[in] Grp& response
	 * \param[in] srting& predictor file name
	 * \param[in] size_t& number of predictors
	 * \param[in] int& number of threads
	 */
	BetaGrpFt(const Grp &rsp, const string &predFlNam, const size_t &Npred, const int &nThr);
	/** \brief Simple constructor with a prior index
	 *
	 * Reads the predictor from a file and initiates regression coefficients using the provided response data.  Number of rows of the predictor is equal to the number of rows in the response (no replication).
	 *
	 * \param[in] Grp& response
	 * \param[in] srting& predictor file name
	 * \param[in] size_t& number of predictors
	 * \param[in] RanIndex& index to the prior
	 * \param[in] int& number of threads
	 */
	BetaGrpFt(const Grp &rsp, const string &predFlNam, const size_t &Npred, RanIndex &up, const int &nThr);
	/** \brief Simple constructor with a prior index and replication
	 *
	 * Reads the predictor from a file and initiates regression coefficients using the provided response data.  Number of rows of the predictor is equal to the number of groups in the lower (data) index.
	 * The number of rows in the response is equal to the number of elements in the low index.
	 *
	 * \param[in] Grp& response
	 * \param[in] srting& predictor file name
	 * \param[in] size_t& number of predictors
	 * \param[in] RanIndex& replication index
	 * \param[in] RanIndex& index to the prior
	 * \param[in] int& number of threads
	 */
	BetaGrpFt(const Grp &rsp, const string &predFlNam, const size_t &Npred, RanIndex &low, RanIndex &up, const int &nThr);
	/** \brief Simple constructor with output file name
	 *
	 * Reads the predictor from a file and initiates regression coefficients using the provided response data.  Number of rows of the predictor is equal to the number of rows in the response (no replication).
	 * No upper index is specified, so this object will implement a regression with an improper flat prior.  Caution is advised if the number of predictors approaches the number of rows in the response, since inference will not be well-conditioned.
	 *
	 * \param[in] Grp& response
	 * \param[in] srting& predictor file name
	 * \param[in] size_t& number of predictors
	 * \param[in] string& output file name
	 * \param[in] int& number of threads
	 */
	BetaGrpFt(const Grp &rsp, const string &predFlNam, const size_t &Npred, const string &outFlNam, const int &nThr);
	/** \brief Simple constructor with a prior index and output file name
	 *
	 * Reads the predictor from a file and initiates regression coefficients using the provided response data.  Number of rows of the predictor is equal to the number of rows in the response (no replication).
	 *
	 * \param[in] Grp& response
	 * \param[in] srting& predictor file name
	 * \param[in] size_t& number of predictors
	 * \param[in] RanIndex& index to the prior
	 * \param[in] string& output file name
	 * \param[in] int& number of threads
	 */
	BetaGrpFt(const Grp &rsp, const string &predFlNam, const size_t &Npred, RanIndex &up, const string &outFlNam, const int &nThr);
	/** \brief Simple constructor with a prior index, replication and output file name
	 *
	 * Reads the predictor from a file and initiates regression coefficients using the provided response data.  Number of rows of the predictor is equal to the number of groups in the lower (data) index.
	 * The number of rows in the response is equal to the number of elements in the low index
	 *
	 * \param[in] Grp& response
	 * \param[in] srting& predictor file name
	 * \param[in] size_t& number of predictors
	 * \param[in] RanIndex& replication index
	 * \param[in] RanIndex& index to the prior
	 * \param[in] string& output file name
	 * \param[in] int& number of threads
	 */
	BetaGrpFt(const Grp &rsp, const string &predFlNam, const size_t &Npred, RanIndex &low, RanIndex &up, const string &outFlNam, const int &nThr);
	
	/** \brief Missing data constructor
	 *
	 * Reads the predictor from a file and initiates regression coefficients using the provided response data. Predictor has missing data labeled by the provided value.  Number of rows of the predictor is equal to the number of rows in the response (no replication).
	 * No upper index is specified, so this object will implement a regression with an improper flat prior.  Caution is advised if the number of predictors approaches the number of rows in the response, since inference will not be well-conditioned.
	 *
	 * \param[in] Grp& response
	 * \param[in] srting& predictor file name
	 * \param[in] size_t& number of predictors
	 * \param[in] double& missing data label
	 * \param[in] int& number of threads
	 */
	BetaGrpFt(const Grp &rsp, const string &predFlNam, const size_t &Npred, const double &absLab, const int &nThr);
	/** \brief Missing data constructor with a prior index
	 *
	 * Reads the predictor from a file and initiates regression coefficients using the provided response data. Predictor has missing data labeled by the provided value.  Number of rows of the predictor is equal to the number of rows in the response (no replication).
	 *
	 * \param[in] Grp& response
	 * \param[in] srting& predictor file name
	 * \param[in] size_t& number of predictors
	 * \param[in] double& missing data label
	 * \param[in] RanIndex& index to the prior
	 * \param[in] int& number of threads
	 */
	BetaGrpFt(const Grp &rsp, const string &predFlNam, const size_t &Npred, const double &absLab, RanIndex &up, const int &nThr);
	/** \brief Missing data constructor with a prior index and replication
	 *
	 * Reads the predictor from a file and initiates regression coefficients using the provided response data. Predictor has missing data labeled by the provided value.  Number of rows of the predictor is equal to the number of groups in the lower (data) index.
	 * The number of rows in the response is equal to the number of elements in the low index
	 *
	 * \param[in] Grp& response
	 * \param[in] srting& predictor file name
	 * \param[in] size_t& number of predictors
	 * \param[in] double& missing data label
	 * \param[in] RanIndex& replication index
	 * \param[in] RanIndex& index to the prior
	 * \param[in] int& number of threads
	 */
	BetaGrpFt(const Grp &rsp, const string &predFlNam, const size_t &Npred, const double &absLab, RanIndex &low, RanIndex &up, const int &nThr);
	/** \brief Missing data constructor with output file name
	 *
	 * Reads the predictor from a file and initiates regression coefficients using the provided response data. Predictor has missing data labeled by the provided value.  Number of rows of the predictor is equal to the number of rows in the response (no replication).
	 * No upper index is specified, so this object will implement a regression with an improper flat prior.  Caution is advised if the number of predictors approaches the number of rows in the response, since inference will not be well-conditioned.
	 *
	 * \param[in] Grp& response
	 * \param[in] srting& predictor file name
	 * \param[in] size_t& number of predictors
	 * \param[in] double& missing data label
	 * \param[in] string& output file name
	 * \param[in] int& number of threads
	 */
	BetaGrpFt(const Grp &rsp, const string &predFlNam, const size_t &Npred, const double &absLab, const string &outFlNam, const int &nThr);
	/** \brief Missing data constructor with a prior index and output file name
	 *
	 * Reads the predictor from a file and initiates regression coefficients using the provided response data. Predictor has missing data labeled by the provided value.  Number of rows of the predictor is equal to the number of rows in the response (no replication).
	 *
	 * \param[in] Grp& response
	 * \param[in] srting& predictor file name
	 * \param[in] size_t& number of predictors
	 * \param[in] double& missing data label
	 * \param[in] RanIndex& index to the prior
	 * \param[in] string& output file name
	 * \param[in] int& number of threads
	 */
	BetaGrpFt(const Grp &rsp, const string &predFlNam, const size_t &Npred, const double &absLab, RanIndex &up, const string &outFlNam, const int &nThr);
	/** \brief Missing data constructor with a prior index, replication and output file name
	 *
	 * Reads the predictor from a file and initiates regression coefficients using the provided response data. Predictor has missing data labeled by the provided value.  Number of rows of the predictor is equal to the number of groups in the lower (data) index.
	 * The number of rows in the response is equal to the number of elements in the low index
	 *
	 * \param[in] Grp& response
	 * \param[in] srting& predictor file name
	 * \param[in] size_t& number of predictors
	 * \param[in] double& missing data label
	 * \param[in] RanIndex& replication index
	 * \param[in] RanIndex& index to the prior
	 * \param[in] string& output file name
	 * \param[in] int& number of threads
	 */
	BetaGrpFt(const Grp &rsp, const string &predFlNam, const size_t &Npred, const double &absLab, RanIndex &low, RanIndex &up, const string &outFlNam, const int &nThr);
	
	/** \brief Selection constructor
	 *
	 * Unlike the previous constructors, selection-type constructors pre-screen predictors for association with any of the traits (using Hoteling-like multi-trait statistics), and keeps only the specified proportion in the model.  
	 * The candidate predictors are further screened for between-predictor correlation and only ones with \f$ r^2 \f$ below the provided threshold are kept in the model.  This among-predictor correlation often arises in SNP data, where there is likage disequilbrium (LD) among markers.
	 * Once the predictors are picked, the updating proceeds like for other BetaGrpFt objects, i.e. the set remains the same (unlike variable selection).
	 *
	 * \warning This set of models is still experimental and has not been extenisively tested.  For example, it seems clear that the rest of the model has to be pre-run to convergence before initializing this kind of object.  Furthermore, the initializeing predictor probably has to be a mean of several (\f$ \sim 50 \f$) MCMC samples.
	 *
	 * \param[in] Grp& response
	 * \param[in] SigmaI& data inverse-covariance
	 * \param[in] srting& predictor file name
	 * \param[in] size_t& number of predictors
	 * \param[in] double& fraction of predictors to retain, as a fraction of the number of data points
	 * \param[in] double& \f$ r^2 \f$ cut-off for among-predictor correlation
	 * \param[in] RanIndex& index to the prior
	 * \param[in] string& output file name
	 * \param[in] int& number of threads
	 *
	 */
	BetaGrpFt(const Grp &rsp, const SigmaI &SigI, const string &predFlNam, const size_t &Npred, const double &Nmul, const double &rSqMax, RanIndex &up, const string &outFlNam, const int &nThr);
	/** \brief Selection constructor with replication
	 *
	 * Unlike the previous constructors, selection-type constructors pre-screen predictors for association with any of the traits (using Hoteling-like multi-trait statistics), and keeps only the specified proportion in the model.
	 * The candidate predictors are further screened for between-predictor correlation and only ones with \f$ r^2 \f$ below the provided threshold are kept in the model.  This among-predictor correlation often arises in SNP data, where there is likage disequilbrium (LD) among markers.
	 * Once the predictors are picked, the updating proceeds like for other BetaGrpFt objects, i.e. the set remains the same (unlike variable selection).
	 *
	 * \warning This set of models is still experimental and has not been extenisively tested.  For example, it seems clear that the rest of the model has to be pre-run to convergence before initializing this kind of object.  Furthermore, the initializeing predictor probably has to be a mean of several (\f$ \sim 50 \f$) MCMC samples.
	 *
	 * \param[in] Grp& response
	 * \param[in] SigmaI& data inverse-covariance
	 * \param[in] srting& predictor file name
	 * \param[in] size_t& number of predictors
	 * \param[in] double& fraction of predictors to retain, as a fraction of the number of data points
	 * \param[in] double& \f$ r^2 \f$ cut-off for among-predictor correlation
	 * \param[in] RanIndex& replication index
	 * \param[in] RanIndex& index to the prior
	 * \param[in] string& output file name
	 * \param[in] int& number of threads
	 *
	 */
	BetaGrpFt(const Grp &rsp, const SigmaI &SigI, const string &predFlNam, const size_t &Npred, const double &Nmul, const double &rSqMax, RanIndex &low, RanIndex &up, const string &outFlNam, const int &nThr);
	/** \brief Selection constructor with missing predictor data
	 *
	 * Unlike the previous constructors, selection-type constructors pre-screen predictors for association with any of the traits (using Hoteling-like multi-trait statistics), and keeps only the specified proportion in the model.
	 * The candidate predictors are further screened for between-predictor correlation and only ones with \f$ r^2 \f$ below the provided threshold are kept in the model.  This among-predictor correlation often arises in SNP data, where there is likage disequilbrium (LD) among markers.
	 * Once the predictors are picked, the updating proceeds like for other BetaGrpFt objects, i.e. the set remains the same (unlike variable selection).  Missing predictor data are filled in by mean imputation.
	 *
	 * \warning This set of models is still experimental and has not been extenisively tested.  For example, it seems clear that the rest of the model has to be pre-run to convergence before initializing this kind of object.  Furthermore, the initializeing predictor probably has to be a mean of several (\f$ \sim 50 \f$) MCMC samples.
	 *
	 * \param[in] Grp& response
	 * \param[in] SigmaI& data inverse-covariance
	 * \param[in] srting& predictor file name
	 * \param[in] size_t& number of predictors
	 * \param[in] double& fraction of predictors to retain, as a fraction of the number of data points
	 * \param[in] double& \f$ r^2 \f$ cut-off for among-predictor correlation
	 * \param[in] double& missing data label
	 * \param[in] RanIndex& index to the prior
	 * \param[in] string& output file name
	 * \param[in] int& number of threads
	 *
	 */
	BetaGrpFt(const Grp &rsp, const SigmaI &SigI, const string &predFlNam, const size_t &Npred, const double &Nmul, const double &rSqMax, const double &absLab, RanIndex &up, const string &outFlNam, const int &nThr);
	/** \brief Selection constructor with missing predictor data and replication
	 *
	 * Unlike the previous constructors, selection-type constructors pre-screen predictors for association with any of the traits (using Hoteling-like multi-trait statistics), and keeps only the specified proportion in the model.
	 * The candidate predictors are further screened for between-predictor correlation and only ones with \f$ r^2 \f$ below the provided threshold are kept in the model.  This among-predictor correlation often arises in SNP data, where there is likage disequilbrium (LD) among markers.
	 * Once the predictors are picked, the updating proceeds like for other BetaGrpFt objects, i.e. the set remains the same (unlike variable selection).  Missing predictor data are filled in by mean imputation.
	 *
	 * \warning This set of models is still experimental and has not been extenisively tested.  For example, it seems clear that the rest of the model has to be pre-run to convergence before initializing this kind of object.  Furthermore, the initializeing predictor probably has to be a mean of several (\f$ \sim 50 \f$) MCMC samples.
	 *
	 * \param[in] Grp& response
	 * \param[in] SigmaI& data inverse-covariance
	 * \param[in] srting& predictor file name
	 * \param[in] size_t& number of predictors
	 * \param[in] double& fraction of predictors to retain, as a fraction of the number of data points
	 * \param[in] double& \f$ r^2 \f$ cut-off for among-predictor correlation
	 * \param[in] double& missing data label
	 * \param[in] RanIndex& replication index
	 * \param[in] RanIndex& index to the prior
	 * \param[in] string& output file name
	 * \param[in] int& number of threads
	 *
	 */
	BetaGrpFt(const Grp &rsp, const SigmaI &SigI, const string &predFlNam, const size_t &Npred, const double &Nmul, const double &rSqMax, const double &absLab, RanIndex &low, RanIndex &up, const string &outFlNam, const int &nThr);
	
	/** \brief Destructor */
	virtual ~BetaGrpFt();
	
	/** \brief Copy constructor
	 *
	 * \param[in] BetaGrpFt& object to be copied
	 * \return BetaGrpFt object
	 */
	BetaGrpFt(const BetaGrpFt &mG);
	/** \brief Assignment operator
	 *
	 * \param[in] BetaGrpFt& object to be copied
	 * \return BetaGrpFt& target object
	 */
	BetaGrpFt & operator=(const BetaGrpFt &mG);
	
	/** \brief Access to the fitted value matrix
	 *
	 * Pointer to the \f$ \boldsymbol{XB} \f$ matrix, while dMat() accesses the regression coefficient matrix.
	 *
	 * \return gsl_matrix* pointer to the matrix of fitted values
	 *
	 */
	virtual const gsl_matrix *fMat() const{return _fittedAll; };
	/** \brief Store samples
	 *
	 * Stores samples of predictor effect scores in a matrix to be dumped at the end of the MCMC run.
	 *
	 * \param[in] SigmaI& data inverse-covariance
	 *
	 */
	void save(const SigmaI &SigI);
	void dump();
	
	double lnOddsRat(const Grp &y, const SigmaI &SigI, const size_t i) const;
	
	// improper prior
	void update(const Grp &dat, const SigmaI &SigIm);
	void update(const Grp &dat, const Qgrp &q, const SigmaI &SigIm);
	// 0-mean prior
	virtual void update(const Grp &dat, const SigmaI &SigIm, const SigmaI &SigIp);
	virtual void update(const Grp &dat, const Qgrp &q, const SigmaI &SigIm, const SigmaI &SigIp);
	virtual void update(const Grp &dat, const SigmaI &SigIm, const Qgrp &qPr, const SigmaI &SigIp);
	virtual void update(const Grp &dat, const Qgrp &q, const SigmaI &SigIm, const Qgrp &qPr, const SigmaI &SigIp);
	// non-0-mean prior
	virtual void update(const Grp &dat, const SigmaI &SigIm, const Grp &muPr, const SigmaI &SigIp);
	virtual void update(const Grp &dat, const Qgrp &q, const SigmaI &SigIm, const Grp &muPr, const SigmaI &SigIp);
	virtual void update(const Grp &dat, const SigmaI &SigIm, const Grp &muPr, const Qgrp &qPr, const SigmaI &SigIp);
	virtual void update(const Grp &dat, const Qgrp &q, const SigmaI &SigIm, const Grp &muPr, const Qgrp &qPr, const SigmaI &SigIp);
	
};
/** \brief Multivariate multiple regression with parameter expansion
 *
 * Implements multiplicative prameter expansion as in MuGrpPEX, but for regression coefficients. The "raw" value matrix \f$ \boldsymbol{\Xi} \f$ contains regression coefficients on the unadjusted scale (see MuGrpPEX for explanation of these terms).
 * The adjusted regression coefficient matrix is not calculated, but fMat() point to the adjusted fitted value matrix \f$ \boldsymbol{X \Xi A} \f$.
 * Regression models with this method must have a prior, typically one with mean zero.
 *
 */
class BetaGrpPEX : virtual public BetaGrpFt {
protected:
	/** \brief Scaled inverse-covariance
	 *
	 * The matrix \f$ \left( \boldsymbol{\Sigma}^{-1}\boldsymbol{A}^T \right)^T \f$ that is common for sampling each row of the value matrix.
	 */
	gsl_matrix *_tSigIAt;
	/** \brief Multiplicative redundant parameter */
	Apex _A;
	/** \brief Fitted values
	 *
	 * Vector of vectorized individual fitted matrices \f$ \boldsymbol{\Xi}_{\cdot -m}\boldsymbol{A}_{-m\cdot} \f$.
	 *
	 */
	vector<vector<double> > _ftA;
	/** \brief Adjusted fitted matrix
	 *
	 * The \f$ \boldsymbol{X \Xi A} \f$ matrix.  The inherited _fittedAll matrix is \f$ \boldsymbol{X \Xi} \f$, i.e. the "raw" version.
	 *
	 */
	gsl_matrix *_fittedAllAdj;
	
	/** \brief Finish construction
	 *
	 * Most of the construction tasks are handled by the parent class.  This function sets up the PEX portion of the class.
	 *
	 * \param[in] double& prior inverse variance for \f$ \boldsymbol{A} \f$
	 *
	 */
	void _finishConstruct(const double &Spr);
	/** \brief Adjusted matrix calculation
	 *
	 * Calculates adjusted fitted matrix and all sub-matrices.
	 *
	 */
	void _finishFitted();
	/** \brief Calculate redundant parameter fitted values
	 *
	 * \ingroup fitFun
	 *
	 * Calculates separate \f$ \left(\boldsymbol{X \Xi A}\right)_{\cdot -p} \f$ for each trait \f$ p \f$.
	 */
	void _updateAfitted();
	
	/** \brief Finishing constructor
	 *
	 * For use in BetaGrpPCpex.
	 *
	 * \param[in] double& prior inverse variance for \f$ \boldsymbol{A} \f$
	 */
	BetaGrpPEX(const double &Spr) {_finishConstruct(Spr); };
public:
	/** \brief Default constructor */
	BetaGrpPEX() : BetaGrpFt() {};
	/** \brief Simple constructor with a prior index
	 *
	 * Reads the predictor from a file and initiates regression coefficients using the provided response data.  Number of rows of the predictor is equal to the number of rows in the response (no replication).
	 *
	 * \param[in] Grp& response
	 * \param[in] srting& predictor file name
	 * \param[in] size_t& number of predictors
	 * \param[in] double& prior inverse variance for \f$ \boldsymbol{A} \f$
	 * \param[in] RanIndex& index to the prior
	 * \param[in] int& number of threads
	 */
	BetaGrpPEX(const Grp &rsp, const string &predFlNam, const size_t &Npred, const double &Spr, RanIndex &up, const int &nThr) : BetaGrpFt(rsp, predFlNam, Npred, up, nThr) {_finishConstruct(Spr); };
	/** \brief Simple constructor with a prior index and replication
	 *
	 * Reads the predictor from a file and initiates regression coefficients using the provided response data.  Number of rows of the predictor is equal to the number of groups in the lower (data) index.
	 * The number of rows in the response is equal to the number of elements in the low index
	 *
	 * \param[in] Grp& response
	 * \param[in] srting& predictor file name
	 * \param[in] size_t& number of predictors
	 * \param[in] double& prior inverse variance for \f$ \boldsymbol{A} \f$
	 * \param[in] RanIndex& replication index
	 * \param[in] RanIndex& index to the prior
	 * \param[in] int& number of threads
	 */
	BetaGrpPEX(const Grp &rsp, const string &predFlNam, const size_t &Npred, const double &Spr, RanIndex &low, RanIndex &up, const int &nThr) : BetaGrpFt(rsp, predFlNam, Npred, low, up, nThr) {_finishConstruct(Spr); };
	/** \brief Simple constructor with a prior index and output file name
	 *
	 * Reads the predictor from a file and initiates regression coefficients using the provided response data.  Number of rows of the predictor is equal to the number of rows in the response (no replication).
	 *
	 * \param[in] Grp& response
	 * \param[in] srting& predictor file name
	 * \param[in] size_t& number of predictors
	 * \param[in] double& prior inverse variance for \f$ \boldsymbol{A} \f$
	 * \param[in] RanIndex& index to the prior
	 * \param[in] string& output file name
	 * \param[in] int& number of threads
	 */
	BetaGrpPEX(const Grp &rsp, const string &predFlNam, const size_t &Npred, const double &Spr, RanIndex &up, const string &outFlNam, const int &nThr) : BetaGrpFt(rsp, predFlNam, Npred, up, outFlNam, nThr) {_finishConstruct(Spr); };
	/** \brief Simple constructor with a prior index, replication and output file name
	 *
	 * Reads the predictor from a file and initiates regression coefficients using the provided response data.  Number of rows of the predictor is equal to the number of groups in the lower (data) index.
	 * The number of rows in the response is equal to the number of elements in the low index
	 *
	 * \param[in] Grp& response
	 * \param[in] srting& predictor file name
	 * \param[in] size_t& number of predictors
	 * \param[in] double& prior inverse variance for \f$ \boldsymbol{A} \f$
	 * \param[in] RanIndex& replication index
	 * \param[in] RanIndex& index to the prior
	 * \param[in] string& output file name
	 * \param[in] int& number of threads
	 */
	BetaGrpPEX(const Grp &rsp, const string &predFlNam, const size_t &Npred, const double &Spr, RanIndex &low, RanIndex &up, const string &outFlNam, const int &nThr) : BetaGrpFt(rsp, predFlNam, Npred, low, up, outFlNam, nThr) {_finishConstruct(Spr); };
	
	/** \brief Missing data constructor with a prior index
	 *
	 * Reads the predictor from a file and initiates regression coefficients using the provided response data. Predictor has missing data labeled by the provided value.  Number of rows of the predictor is equal to the number of rows in the response (no replication).
	 *
	 * \param[in] Grp& response
	 * \param[in] srting& predictor file name
	 * \param[in] size_t& number of predictors
	 * \param[in] double& prior inverse variance for \f$ \boldsymbol{A} \f$
	 * \param[in] double& missing data label
	 * \param[in] RanIndex& index to the prior
	 * \param[in] int& number of threads
	 */
	BetaGrpPEX(const Grp &rsp, const string &predFlNam, const size_t &Npred, const double &Spr, const double &absLab, RanIndex &up, const int &nThr) : BetaGrpFt(rsp, predFlNam, Npred, absLab, up, nThr) {_finishConstruct(Spr); };
	/** \brief Missing data constructor with a prior index and replication
	 *
	 * Reads the predictor from a file and initiates regression coefficients using the provided response data. Predictor has missing data labeled by the provided value.  Number of rows of the predictor is equal to the number of groups in the lower (data) index.
	 * The number of rows in the response is equal to the number of elements in the low index
	 *
	 * \param[in] Grp& response
	 * \param[in] srting& predictor file name
	 * \param[in] size_t& number of predictors
	 * \param[in] double& prior inverse variance for \f$ \boldsymbol{A} \f$
	 * \param[in] double& missing data label
	 * \param[in] RanIndex& replication index
	 * \param[in] RanIndex& index to the prior
	 * \param[in] int& number of threads
	 */
	BetaGrpPEX(const Grp &rsp, const string &predFlNam, const size_t &Npred, const double &Spr, const double &absLab, RanIndex &low, RanIndex &up, const int &nThr) : BetaGrpFt(rsp, predFlNam, Npred, absLab, low, up, nThr) {_finishConstruct(Spr); };
	/** \brief Missing data constructor with a prior index and output file name
	 *
	 * Reads the predictor from a file and initiates regression coefficients using the provided response data. Predictor has missing data labeled by the provided value.  Number of rows of the predictor is equal to the number of rows in the response (no replication).
	 *
	 * \param[in] Grp& response
	 * \param[in] srting& predictor file name
	 * \param[in] size_t& number of predictors
	 * \param[in] double& prior inverse variance for \f$ \boldsymbol{A} \f$
	 * \param[in] double& missing data label
	 * \param[in] RanIndex& index to the prior
	 * \param[in] string& output file name
	 * \param[in] int& number of threads
	 */
	BetaGrpPEX(const Grp &rsp, const string &predFlNam, const size_t &Npred, const double &Spr, const double &absLab, RanIndex &up, const string &outFlNam, const int &nThr) : BetaGrpFt(rsp, predFlNam, Npred, absLab, up, outFlNam, nThr) {_finishConstruct(Spr); };
	/** \brief Missing data constructor with a prior index, replication and output file name
	 *
	 * Reads the predictor from a file and initiates regression coefficients using the provided response data. Predictor has missing data labeled by the provided value.  Number of rows of the predictor is equal to the number of groups in the lower (data) index.
	 * The number of rows in the response is equal to the number of elements in the low index
	 *
	 * \param[in] Grp& response
	 * \param[in] srting& predictor file name
	 * \param[in] size_t& number of predictors
	 * \param[in] double& prior inverse variance for \f$ \boldsymbol{A} \f$
	 * \param[in] double& missing data label
	 * \param[in] RanIndex& replication index
	 * \param[in] RanIndex& index to the prior
	 * \param[in] string& output file name
	 * \param[in] int& number of threads
	 */
	BetaGrpPEX(const Grp &rsp, const string &predFlNam, const size_t &Npred, const double &Spr, const double &absLab, RanIndex &low, RanIndex &up, const string &outFlNam, const int &nThr) : BetaGrpFt(rsp, predFlNam, Npred, absLab, low, up, outFlNam, nThr) {_finishConstruct(Spr); };
	
	/** \brief Selection constructor
	 *
	 * Unlike the previous constructors, selection-type constructors pre-screen predictors for association with any of the traits (using Hoteling-like multi-trait statistics), and keeps only the specified proportion in the model.
	 * The candidate predictors are further screened for between-predictor correlation and only ones with \f$ r^2 \f$ below the provided threshold are kept in the model.  This among-predictor correlation often arises in SNP data, where there is likage disequilbrium (LD) among markers.
	 * Once the predictors are picked, the updating proceeds like for other BetaGrpFt objects, i.e. the set remains the same (unlike variable selection).
	 *
	 * \warning This set of models is still experimental and has not been extenisively tested.  For example, it seems clear that the rest of the model has to be pre-run to convergence before initializing this kind of object.  Furthermore, the initializeing predictor probably has to be a mean of several (\f$ \sim 50 \f$) MCMC samples.
	 *
	 * \param[in] Grp& response
	 * \param[in] SigmaI& data inverse-covariance
	 * \param[in] srting& predictor file name
	 * \param[in] size_t& number of predictors
	 * \param[in] double& prior inverse variance for \f$ \boldsymbol{A} \f$
	 * \param[in] double& fraction of predictors to retain, as a fraction of the number of data points
	 * \param[in] double& \f$ r^2 \f$ cut-off for among-predictor correlation
	 * \param[in] RanIndex& index to the prior
	 * \param[in] string& output file name
	 * \param[in] int& number of threads
	 *
	 */
	BetaGrpPEX(const Grp &rsp, const SigmaI &SigI, const string &predFlNam, const size_t &Npred, const double &Spr, const double &Nmul, const double &rSqMax, RanIndex &up, const string &outFlNam, const int &nThr) : BetaGrpFt(rsp, SigI, predFlNam, Npred, Nmul, rSqMax, up, outFlNam, nThr) {_finishConstruct(Spr); };
	/** \brief Selection constructor with replication
	 *
	 * Unlike the previous constructors, selection-type constructors pre-screen predictors for association with any of the traits (using Hoteling-like multi-trait statistics), and keeps only the specified proportion in the model.
	 * The candidate predictors are further screened for between-predictor correlation and only ones with \f$ r^2 \f$ below the provided threshold are kept in the model.  This among-predictor correlation often arises in SNP data, where there is likage disequilbrium (LD) among markers.
	 * Once the predictors are picked, the updating proceeds like for other BetaGrpFt objects, i.e. the set remains the same (unlike variable selection).
	 *
	 * \warning This set of models is still experimental and has not been extenisively tested.  For example, it seems clear that the rest of the model has to be pre-run to convergence before initializing this kind of object.  Furthermore, the initializeing predictor probably has to be a mean of several (\f$ \sim 50 \f$) MCMC samples.
	 *
	 * \param[in] Grp& response
	 * \param[in] SigmaI& data inverse-covariance
	 * \param[in] srting& predictor file name
	 * \param[in] size_t& number of predictors
	 * \param[in] double& prior inverse variance for \f$ \boldsymbol{A} \f$
	 * \param[in] double& fraction of predictors to retain, as a fraction of the number of data points
	 * \param[in] double& \f$ r^2 \f$ cut-off for among-predictor correlation
	 * \param[in] RanIndex& replication index
	 * \param[in] RanIndex& index to the prior
	 * \param[in] string& output file name
	 * \param[in] int& number of threads
	 *
	 */
	BetaGrpPEX(const Grp &rsp, const SigmaI &SigI, const string &predFlNam, const size_t &Npred, const double &Spr, const double &Nmul, const double &rSqMax, RanIndex &low, RanIndex &up, const string &outFlNam, const int &nThr) : BetaGrpFt(rsp, SigI, predFlNam, Npred, Nmul, rSqMax, low, up, outFlNam, nThr) {_finishConstruct(Spr); };
	/** \brief Selection constructor with missing predictor data
	 *
	 * Unlike the previous constructors, selection-type constructors pre-screen predictors for association with any of the traits (using Hoteling-like multi-trait statistics), and keeps only the specified proportion in the model.
	 * The candidate predictors are further screened for between-predictor correlation and only ones with \f$ r^2 \f$ below the provided threshold are kept in the model.  This among-predictor correlation often arises in SNP data, where there is likage disequilbrium (LD) among markers.
	 * Once the predictors are picked, the updating proceeds like for other BetaGrpFt objects, i.e. the set remains the same (unlike variable selection).  Missing predictor data are filled in by mean imputation.
	 *
	 * \warning This set of models is still experimental and has not been extenisively tested.  For example, it seems clear that the rest of the model has to be pre-run to convergence before initializing this kind of object.  Furthermore, the initializeing predictor probably has to be a mean of several (\f$ \sim 50 \f$) MCMC samples.
	 *
	 * \param[in] Grp& response
	 * \param[in] SigmaI& data inverse-covariance
	 * \param[in] srting& predictor file name
	 * \param[in] size_t& number of predictors
	 * \param[in] double& prior inverse variance for \f$ \boldsymbol{A} \f$
	 * \param[in] double& fraction of predictors to retain, as a fraction of the number of data points
	 * \param[in] double& \f$ r^2 \f$ cut-off for among-predictor correlation
	 * \param[in] double& missing data label
	 * \param[in] RanIndex& index to the prior
	 * \param[in] string& output file name
	 * \param[in] int& number of threads
	 *
	 */
	BetaGrpPEX(const Grp &rsp, const SigmaI &SigI, const string &predFlNam, const size_t &Npred, const double &Spr, const double &Nmul, const double &rSqMax, const double &absLab, RanIndex &up, const string &outFlNam, const int &nThr) : BetaGrpFt(rsp, SigI, predFlNam, Npred, Nmul, rSqMax, absLab, up, outFlNam, nThr) {_finishConstruct(Spr); };
	/** \brief Selection constructor with missing predictor data and replication
	 *
	 * Unlike the previous constructors, selection-type constructors pre-screen predictors for association with any of the traits (using Hoteling-like multi-trait statistics), and keeps only the specified proportion in the model.
	 * The candidate predictors are further screened for between-predictor correlation and only ones with \f$ r^2 \f$ below the provided threshold are kept in the model.  This among-predictor correlation often arises in SNP data, where there is likage disequilbrium (LD) among markers.
	 * Once the predictors are picked, the updating proceeds like for other BetaGrpFt objects, i.e. the set remains the same (unlike variable selection).  Missing predictor data are filled in by mean imputation.
	 *
	 * \warning This set of models is still experimental and has not been extenisively tested.  For example, it seems clear that the rest of the model has to be pre-run to convergence before initializing this kind of object.  Furthermore, the initializeing predictor probably has to be a mean of several (\f$ \sim 50 \f$) MCMC samples.
	 *
	 * \param[in] Grp& response
	 * \param[in] SigmaI& data inverse-covariance
	 * \param[in] srting& predictor file name
	 * \param[in] size_t& number of predictors
	 * \param[in] double& prior inverse variance for \f$ \boldsymbol{A} \f$
	 * \param[in] double& fraction of predictors to retain, as a fraction of the number of data points
	 * \param[in] double& \f$ r^2 \f$ cut-off for among-predictor correlation
	 * \param[in] double& missing data label
	 * \param[in] RanIndex& replication index
	 * \param[in] RanIndex& index to the prior
	 * \param[in] string& output file name
	 * \param[in] int& number of threads
	 *
	 */
	BetaGrpPEX(const Grp &rsp, const SigmaI &SigI, const string &predFlNam, const size_t &Npred, const double &Spr, const double &Nmul, const double &rSqMax, const double &absLab, RanIndex &low, RanIndex &up, const string &outFlNam, const int &nThr) : BetaGrpFt(rsp, SigI, predFlNam, Npred, Nmul, rSqMax, absLab, low, up, outFlNam, nThr) {_finishConstruct(Spr); };
	
	/** \brief Destructor */
	virtual ~BetaGrpPEX();
	
	/** \brief Access adjusted fitted value matrix 
	 *
	 * \return gsl_matrix* pointer to the adjusted-value fitted matrix
	 */
	virtual const gsl_matrix *fMat() const{return _fittedAllAdj; };
	/** \brief Save adjusted values
	 *
	 * Appends the current adjusted mean values to the file whose name was set during construction.
	 */
	virtual void save();
	/** \brief Save adjusted values to named file
	 *
	 * Appends the current adjusted mean values to the named file.
	 *
	 * \param[in] string& file name
	 */
	virtual void save(const string &outFlNam);
	/** \brief Save adjusted values with the adjusted covariance matrix
	 *
	 * Appends the current adjusted mean values and prior covariance matrix to the files whose names were set during construction.
	 */
	virtual void save(const SigmaI &SigI);
	
	virtual void update(const Grp &dat, const SigmaI &SigIm, const SigmaI &SigIp);
	virtual void update(const Grp &dat, const Qgrp &q, const SigmaI &SigIm, const SigmaI &SigIp);
	virtual void update(const Grp &dat, const SigmaI &SigIm, const Qgrp &qPr, const SigmaI &SigIp);
	virtual void update(const Grp &dat, const Qgrp &q, const SigmaI &SigIm, const Qgrp &qPr, const SigmaI &SigIp);
	
	virtual void update(const Grp &dat, const SigmaI &SigIm, const Grp &muPr, const SigmaI &SigIp);
	virtual void update(const Grp &dat, const Qgrp &q, const SigmaI &SigIm, const Grp &muPr, const SigmaI &SigIp);
	virtual void update(const Grp &dat, const SigmaI &SigIm, const Grp &muPr, const Qgrp &qPr, const SigmaI &SigIp);
	virtual void update(const Grp &dat, const Qgrp &q, const SigmaI &SigIm, const Grp &muPr, const Qgrp &qPr, const SigmaI &SigIp);
};

/** \brief Relationship matrix regression
 *
 * Implements regression models with principal components of a relationship matrix as predictors.  Only principal vectors with non-zero eigenvalues are used.  If the likelihood and the prior are Gaussian, and the prior mean is zero, this is equivalent to the "mixed model" in packages like EMMA and TASSEL.
 * Because the PC vectors are orthonormal there are some speed-ups in initialization.  The updates are handled by the same internal row classes as in BetaGrpFt.
 *
 */
class BetaGrpPC : virtual public BetaGrpFt {
protected:
	
public:
	/** \brief Default constructor */
	BetaGrpPC() : BetaGrpFt() {};
	/** \brief Constructor with a prior index
	 *
	 * Reads principal vectors and eigen-values from files. Number of rows of the predictor is equal to the number of rows in the response (no replication).
	 *
	 * \param[in] Grp& data for initialization
	 * \param[in] string& PC vector file name
	 * \param[in] string& eigen-value file name
	 * \param[in] size_t& number of predictors
	 * \param[in] RanIndex& prior index
	 * \param[in] int& number of threads
	 */
	BetaGrpPC(const Grp &rsp, const string &predFlNam, const string &evFlNam, const size_t &Npred, RanIndex &up, const int &nThr);
	/** \brief Constructor with a prior index and replication
	 *
	 * Reads principal vectors and eigen-values from files.   Number of rows of the predictor is equal to the number of groups in the lower (data) index.
	 * The number of rows in the response is equal to the number of elements in the low index.
	 *
	 * \param[in] Grp& data for initialization
	 * \param[in] string& PC vector file name
	 * \param[in] string& eigen-value file name
	 * \param[in] size_t& number of predictors
	 * \param[in] RanIndex& lower-level (replicate) index
	 * \param[in] RanIndex& prior index
	 * \param[in] int& number of threads
	 */
	BetaGrpPC(const Grp &rsp, const string &predFlNam, const string &evFlNam, const size_t &Npred, RanIndex &low, RanIndex &up, const int &nThr);
	/** \brief Constructor with a prior index, replication and output file name
	 *
	 * Reads principal vectors and eigen-values from files.   Number of rows of the predictor is equal to the number of groups in the lower (data) index.
	 * The number of rows in the response is equal to the number of elements in the low index.
	 *
	 * \param[in] Grp& data for initialization
	 * \param[in] string& PC vector file name
	 * \param[in] string& eigen-value file name
	 * \param[in] size_t& number of predictors
	 * \param[in] RanIndex& lower-level (replicate) index
	 * \param[in] RanIndex& prior index
	 * \param[in] string& output file name
	 * \param[in] int& number of threads
	 */
	BetaGrpPC(const Grp &rsp, const string &predFlNam, const string &evFlNam, const size_t &Npred, RanIndex &up, const string &outFlNam, const int &nThr);
	/** \brief Constructor with a prior index and replication
	 *
	 * Reads principal vectors and eigen-values from files.   Number of rows of the predictor is equal to the number of groups in the lower (data) index.
	 * The number of rows in the response is equal to the number of elements in the low index.
	 *
	 * \param[in] Grp& data for initialization
	 * \param[in] string& PC vector file name
	 * \param[in] string& eigen-value file name
	 * \param[in] size_t& number of predictors
	 * \param[in] RanIndex& lower-level (replicate) index
	 * \param[in] RanIndex& prior index
	 * \param[in] string& output file name
	 * \param[in] int& number of threads
	 */
	BetaGrpPC(const Grp &rsp, const string &predFlNam, const string &evFlNam, const size_t &Npred, RanIndex &low, RanIndex &up, const string &outFlNam, const int &nThr);
	
	/** \brief Destructor */
	virtual ~BetaGrpPC() {};
	
	/** \brief Copy constructor
	 *
	 * \param[in] BetaGrpPC& object to be copied
	 */
	BetaGrpPC(const BetaGrpPC &mG);
	/** \brief Assignment operator
	 *
	 * \param[in] BetaGrpPC& object to be copied
	 * \return BetaGrpPC& target object
	 */
	BetaGrpPC & operator=(const BetaGrpPC &mG);
	
	//update methods taken care of by BetaGrpFt
};

/** \brief Multiplicative parameter expansion for PC regression
 *
 * Brings together implementations from BetaGrpPC and BetaGrpPEX.
 *
 */
class BetaGrpPCpex : public BetaGrpPC, public BetaGrpPEX {
protected:
	
public:
	/** \brief Default constructor */
	BetaGrpPCpex(){};
	/** \brief Constructor with a prior index
	 *
	 * Reads principal vectors and eigen-values from files. Number of rows of the predictor is equal to the number of rows in the response (no replication).
	 *
	 * \param[in] Grp& data for initialization
	 * \param[in] string& PC vector file name
	 * \param[in] string& eigen-value file name
	 * \param[in] size_t& number of predictors
	 * \param[in] double& prior inverse variance for \f$ \boldsymbol{A} \f$
	 * \param[in] RanIndex& prior index
	 * \param[in] int& number of threads
	 */
	BetaGrpPCpex(const Grp &rsp, const string &predFlNam, const string &evFlNam, const size_t &Npred, const double &Spr, RanIndex &up, const int &nThr) : BetaGrpFt(), BetaGrpPC(rsp, predFlNam, evFlNam, Npred, up, nThr), BetaGrpPEX(Spr) {};
	/** \brief Constructor with a prior index and replication
	 *
	 * Reads principal vectors and eigen-values from files.   Number of rows of the predictor is equal to the number of groups in the lower (data) index.
	 * The number of rows in the response is equal to the number of elements in the low index.
	 *
	 * \param[in] Grp& data for initialization
	 * \param[in] string& PC vector file name
	 * \param[in] string& eigen-value file name
	 * \param[in] size_t& number of predictors
	 * \param[in] double& prior inverse variance for \f$ \boldsymbol{A} \f$
	 * \param[in] RanIndex& lower-level (replicate) index
	 * \param[in] RanIndex& prior index
	 * \param[in] int& number of threads
	 */
	BetaGrpPCpex(const Grp &rsp, const string &predFlNam, const string &evFlNam, const size_t &Npred, const double &Spr, RanIndex &low, RanIndex &up, const int &nThr) : BetaGrpFt(), BetaGrpPC(rsp, predFlNam, evFlNam, Npred, low, up, nThr), BetaGrpPEX(Spr) {};
	/** \brief Constructor with a prior index, replication and output file name
	 *
	 * Reads principal vectors and eigen-values from files.   Number of rows of the predictor is equal to the number of groups in the lower (data) index.
	 * The number of rows in the response is equal to the number of elements in the low index.
	 *
	 * \param[in] Grp& data for initialization
	 * \param[in] string& PC vector file name
	 * \param[in] string& eigen-value file name
	 * \param[in] size_t& number of predictors
	 * \param[in] double& prior inverse variance for \f$ \boldsymbol{A} \f$
	 * \param[in] RanIndex& lower-level (replicate) index
	 * \param[in] RanIndex& prior index
	 * \param[in] string& output file name
	 * \param[in] int& number of threads
	 */
	BetaGrpPCpex(const Grp &rsp, const string &predFlNam, const string &evFlNam, const size_t &Npred, const double &Spr, RanIndex &up, const string &outFlNam, const int &nThr) : BetaGrpFt(), BetaGrpPC(rsp, predFlNam, evFlNam, Npred, up, outFlNam, nThr), BetaGrpPEX(Spr) {};
	/** \brief Constructor with a prior index and replication
	 *
	 * Reads principal vectors and eigen-values from files.   Number of rows of the predictor is equal to the number of groups in the lower (data) index.
	 * The number of rows in the response is equal to the number of elements in the low index.
	 *
	 * \param[in] Grp& data for initialization
	 * \param[in] string& PC vector file name
	 * \param[in] string& eigen-value file name
	 * \param[in] size_t& number of predictors
	 * \param[in] double& prior inverse variance for \f$ \boldsymbol{A} \f$
	 * \param[in] RanIndex& lower-level (replicate) index
	 * \param[in] RanIndex& prior index
	 * \param[in] string& output file name
	 * \param[in] int& number of threads
	 */
	BetaGrpPCpex(const Grp &rsp, const string &predFlNam, const string &evFlNam, const size_t &Npred, const double &Spr, RanIndex &low, RanIndex &up, const string &outFlNam, const int &nThr) : BetaGrpFt(), BetaGrpPC(rsp, predFlNam, evFlNam, Npred, low, up, outFlNam, nThr), BetaGrpPEX(Spr) {};
	
	/** \brief Destructor */
	~BetaGrpPCpex() {};
	
	/** \brief Access adjusted fitted value matrix
	 *
	 * \return gsl_matrix* pointer to the adjusted-value fitted matrix
	 */
	const gsl_matrix *fMat() const{return _fittedAllAdj; };
	
	void save(){ BetaGrpPEX::save(); };
	void save(const string &outFlNam){ BetaGrpPEX::save(outFlNam); };
	void save(const SigmaI &SigI){ BetaGrpPEX::save(SigI); };
	
	void update(const Grp &dat, const SigmaI &SigIm, const SigmaI &SigIp) { BetaGrpPEX::update(dat, SigIm, SigIp); };
	void update(const Grp &dat, const Qgrp &q, const SigmaI &SigIm, const SigmaI &SigIp){ BetaGrpPEX::update(dat, q, SigIm, SigIp); };
	void update(const Grp &dat, const SigmaI &SigIm, const Qgrp &qPr, const SigmaI &SigIp) { BetaGrpPEX::update(dat, SigIm, qPr, SigIp); };
	void update(const Grp &dat, const Qgrp &q, const SigmaI &SigIm, const Qgrp &qPr, const SigmaI &SigIp){ BetaGrpPEX::update(dat, q, SigIm, qPr, SigIp); };
	
	void update(const Grp &dat, const SigmaI &SigIm, const Grp &muPr, const SigmaI &SigIp) { BetaGrpPEX::update(dat, SigIm, muPr, SigIp); };
	void update(const Grp &dat, const Qgrp &q, const SigmaI &SigIm, const Grp &muPr, const SigmaI &SigIp){ BetaGrpPEX::update(dat, q, SigIm, muPr, SigIp); };
	void update(const Grp &dat, const SigmaI &SigIm, const Grp &muPr, const Qgrp &qPr, const SigmaI &SigIp) { BetaGrpPEX::update(dat, SigIm, muPr, qPr, SigIp); };
	void update(const Grp &dat, const Qgrp &q, const SigmaI &SigIm, const Grp &muPr, const Qgrp &qPr, const SigmaI &SigIp){ BetaGrpPEX::update(dat, q, SigIm, muPr, qPr, SigIp); };
};

/** \brief Simple single-SNP regression class
 *
 * Implements single-marker GWAS.  Each trait is treated separately, including the variances (i.e., \f$ \widehat{\sigma}^2_p = \widehat{\boldsymbol{\Sigma}}_{p,p} \f$).  However, in addition to the single-trait tests an extra Hotelling-type mutivariate association test is performed.
 * This multivariate test reflects the distance of the whole vector of trait effects to zero, potentially increasing the power to detect pleiotropic SNPs with moderate effects on mutiple traits.
 * The saved value matrix thus has one extra column for this test.  The output is either \f$ -\log_{10}p \f$, (although this statistic is not strictly a frequentist \f$p\f$-value, it performs very similarly in simulations), or Wakefield's \cite wakefield07 approximation of \f$ -ln BF \f$ (log-Bayes factor ratio).  In the latter case, the user can set the prior variance manually.
 * The SNP regression is performed on the point estimate of a response, as described in documentation of the update() and dump() functions.  The latter can be, say, a residual of a mixed-model type GEBV estimate done to control population structure.
 *
 */
class BetaGrpSnp : public MuGrp {
protected:
	/** \brief SNP predictor matrix
	 *
	 * Read in only at the end of the run in the dump() function.
	 */
	gsl_matrix *_Xmat;
	/** \brief Matrix for addition operators
	 *
	 * This matrix is addressed by fMat() and set to all zero values, so that if this object is in an addition/subtraction operator things don't break.  Because this object operates outside of the regular Markov chains, on the residuals of the model, it ordinarily makes no sense to add or subtract it from anything.
	 */
	gsl_matrix *_fakeFmat;
	
	/** \brief Response storage
	 *
	 * MCMC samples from a response variable (representing a residual of the model) are stored here and divided by the total number of saves at the end before the regression is performed.
	 */
	gsl_matrix *_Ystore;
	
	/** \brief Number of predictors (SNPs) */
	size_t _Npred;
	/** \brief Number of threads */
	int _nThr;
	/** \brief Number of saves
	 * 
	 * Number of times the response samples have been saved.
	 */
	double _numSaves;
	/** \brief Prior variance
	 *
	 * Prior variance for Wakefield's \cite wakefield07 ABF calculation (Wakefield's notation for it is \f$W\f$).  If it zero, \f$ -\log_{10}p \f$ are calculated.
	 */
	double _priorVar;  // W in Wakefield's (2007) ABF paper
	
	/** \brief Predictor (SNP) file name */
	string _inPredFl;
	
public:
	/** \brief Default constructor */
	BetaGrpSnp();
	/** \brief Constructor with no replication and \f$p\f$-values
	 *
	 * Initializes the object with no replication (i.e., the number of rows in the predictor is the same as in the response) and leaves the prior variance for SNP regressions at zero, thereby resulting in a dump of \f$ -\log_{10}p \f$ values.
	 *
	 * \param[in] string& predictor file name
	 * \param[in] string& output file name
	 * \param[in] size_t& number of rows in the response matrix
	 * \param[in] size_t& number of predictors
	 * \param[in] size_t& number of traits
	 * \param[in] int& number of threads
	 *
	 */
	BetaGrpSnp(const string &predFlNam, const string &outFlNam, const size_t &Ndat, const size_t &Npred, const size_t &d, const int &Nthr);
	/** \brief Constructor with replication and \f$p\f$-values
	 *
	 * Initializes the object with replication, encoded with the given index, and leaves the prior variance for SNP regressions at zero, thereby resulting in a dump of \f$ -\log_{10}p \f$ values.
	 *
	 * \param[in] string& predictor file name
	 * \param[in] string& output file name
	 * \param[in] RanIndex& replicate index
	 * \param[in] size_t& number of predictors
	 * \param[in] size_t& number of traits
	 * \param[in] int& number of threads
	 *
	 */
	BetaGrpSnp(const string &predFlNam, const string &outFlNam, RanIndex &low, const size_t &Npred, const size_t &d, const int &Nthr);
	/** \brief Constructor with no replication and ABF
	 *
	 * Initializes the object with no replication (i.e., the number of rows in the predictor is the same as in the response) and sets the prior variance for SNP regressions to the given value, therefore dumping Wakefield's log of approximate Bayes factor (ABF) ratios to the output file.
	 *
	 * \param[in] string& predictor file name
	 * \param[in] string& output file name
	 * \param[in] size_t& number of rows in the response matrix
	 * \param[in] size_t& number of predictors
	 * \param[in] size_t& number of traits
	 * \param[in] int& number of threads
	 * \param[in] double& prior variance
	 *
	 */
	BetaGrpSnp(const string &predFlNam, const string &outFlNam, const size_t &Ndat, const size_t &Npred, const size_t &d, const int &Nthr, const double &prVar);
	/** \brief Constructor with replication and ABF
	 *
	 * Initializes the object with replication, encoded with the given index, and sets the prior variance for SNP regressions to the given value, therefore dumping Wakefield's log of approximate Bayes factor (ABF) ratios to the output file.
	 *
	 * \param[in] string& predictor file name
	 * \param[in] string& output file name
	 * \param[in] RanIndex& replicate index
	 * \param[in] size_t& number of predictors
	 * \param[in] size_t& number of traits
	 * \param[in] int& number of threads
	 * \param[in] double& prior variance
	 *
	 */
	BetaGrpSnp(const string &predFlNam, const string &outFlNam, RanIndex &low, const size_t &Npred, const size_t &d, const int &Nthr, const double &prVar);
	
	/** \brief Destructor */
	~BetaGrpSnp();
	
	/** \brief Dump results to the output file
	 *
	 * The predictor (SNP) file is read by this function, and the single-predictor regression (i.e., each predictor is tested separately, ignoring all others) is performed.  The results are saved to the pre-specified output file.
	 */
	virtual void dump();
	
	/** \brief Copy constructor
	 *
	 * \param[in] BetaGrpSnp& object to be copied
	 */
	BetaGrpSnp(const BetaGrpSnp &mG);
	/** \brief Assignment operator
	 *
	 * \param[in] BetaGrpSnp& object to be copied
	 * \return BetaGrpSnp& target object
	 */
	BetaGrpSnp & operator=(const BetaGrpSnp &mG);
	
	/** \brief Access adjusted fitted value matrix
	 *
	 * \return gsl_matrix* pointer to a zero-valued matrix
	 */
	const gsl_matrix *fMat() const{return _fakeFmat; };
	
	/** \brief Response update function
	 *
	 * Unlike standard update functions, this one only saves MCMC samples of the response.  The covariance provided is ignored.  
	 * The actual regression is performed on the point estimates of response values calculated as means of the stored MCMC values, and is done at the end by envoking the dump() function.
	 *
	 * \ingroup updateFun
	 *
	 * \param[in] Grp& response variable to be saved
	 * \param[in] SigmaI& inverse-covariance (ignored)
	 */
	void update(const Grp &dat, const SigmaI &SigIm);
};

/** \brief Single-SNP regression with conditional variance
 *
 * Implements single-marker GWAS.  Each trait is treated separately for coefficient calculation.  In contrast, the variance estimates are taken from the inverse-covariance: \f$ \widehat{\sigma}^2_p = \left[ \widehat{\boldsymbol{\Sigma}}^{-1}_{p,p} \right]^{-1} \le \widehat{\boldsymbol{\Sigma}}_{p,p} \f$.
 * In addition to the single-trait tests an extra Hotelling-type mutivariate association test is performed.
 * This multivariate test reflects the distance of the whole vector of trait effects to zero, potentially increasing the power to detect pleiotropic SNPs with moderate effects on mutiple traits.
 * The saved value matrix thus has one extra column for this test.  The output is either \f$ -\log_{10}p \f$, (although this statistic is not strictly a frequentist \f$p\f$-value, it performs very similarly in simulations), or Wakefield's \cite wakefield07 approximation of \f$ -ln BF \f$ (log-Bayes factor ratio).  In the latter case, the user can set the prior variance manually.
 * The SNP regression is performed on the point estimate of a response, as described in documentation of the update() and dump() functions.  The latter can be, say, a residual of a mixed-model type GEBV estimate done to control population structure.
 */
class BetaGrpSnpCV : public BetaGrpSnp {
protected:
	
public:
	/** \brief Default constructor */
	BetaGrpSnpCV() : BetaGrpSnp() {};
	/** \brief Constructor with no replication and \f$p\f$-values
	 *
	 * Initializes the object with no replication (i.e., the number of rows in the predictor is the same as in the response) and leaves the prior variance for SNP regressions at zero, thereby resulting in a dump of \f$ -\log_{10}p \f$ values.
	 *
	 * \param[in] string& predictor file name
	 * \param[in] string& output file name
	 * \param[in] size_t& number of rows in the response matrix
	 * \param[in] size_t& number of predictors
	 * \param[in] size_t& number of traits
	 * \param[in] int& number of threads
	 *
	 */
	BetaGrpSnpCV(const string &predFlNam, const string &outFlNam, const size_t &Ndat, const size_t &Npred, const size_t &d, const int &Nthr) : BetaGrpSnp(predFlNam, outFlNam, Ndat, Npred, d, Nthr) {};
	/** \brief Constructor with replication and \f$p\f$-values
	 *
	 * Initializes the object with replication, encoded with the given index, and leaves the prior variance for SNP regressions at zero, thereby resulting in a dump of \f$ -\log_{10}p \f$ values.
	 *
	 * \param[in] string& predictor file name
	 * \param[in] string& output file name
	 * \param[in] RanIndex& replicate index
	 * \param[in] size_t& number of predictors
	 * \param[in] size_t& number of traits
	 * \param[in] int& number of threads
	 *
	 */
	BetaGrpSnpCV(const string &predFlNam, const string &outFlNam, RanIndex &low, const size_t &Npred, const size_t &d, const int &Nthr) : BetaGrpSnp(predFlNam, outFlNam, low, Npred, d, Nthr) {};
	/** \brief Constructor with no replication and ABF
	 *
	 * Initializes the object with no replication (i.e., the number of rows in the predictor is the same as in the response) and sets the prior variance for SNP regressions to the given value, therefore dumping Wakefield's log of approximate Bayes factor (ABF) ratios to the output file.
	 *
	 * \param[in] string& predictor file name
	 * \param[in] string& output file name
	 * \param[in] size_t& number of rows in the response matrix
	 * \param[in] size_t& number of predictors
	 * \param[in] size_t& number of traits
	 * \param[in] int& number of threads
	 * \param[in] double& prior variance
	 *
	 */
	BetaGrpSnpCV(const string &predFlNam, const string &outFlNam, const size_t &Ndat, const size_t &Npred, const size_t &d, const int &Nthr, const double &prVar) : BetaGrpSnp(predFlNam, outFlNam, Ndat, Npred, d, Nthr, prVar) {};
	/** \brief Constructor with replication and ABF
	 *
	 * Initializes the object with replication, encoded with the given index, and sets the prior variance for SNP regressions to the given value, therefore dumping Wakefield's log of approximate Bayes factor (ABF) ratios to the output file.
	 *
	 * \param[in] string& predictor file name
	 * \param[in] string& output file name
	 * \param[in] RanIndex& replicate index
	 * \param[in] size_t& number of predictors
	 * \param[in] size_t& number of traits
	 * \param[in] int& number of threads
	 * \param[in] double& prior variance
	 *
	 */
	BetaGrpSnpCV(const string &predFlNam, const string &outFlNam, RanIndex &low, const size_t &Npred, const size_t &d, const int &Nthr, const double &prVar) : BetaGrpSnp(predFlNam, outFlNam, low, Npred, d, Nthr, prVar) {};
	
	/** \brief Destructor */
	~BetaGrpSnpCV() {};
	
	/** \brief Copy constructor
	 *
	 * \param[in] BetaGrpSnpCV& object to be copied
	 */
	BetaGrpSnpCV(const BetaGrpSnpCV &mG);
	/** \brief Assignment operator
	 *
	 * \param[in] BetaGrpSnp& object to be copied
	 * \return BetaGrpSnp& target object
	 */
	BetaGrpSnpCV & operator=(const BetaGrpSnpCV &mG);
	
	void dump();
	
};

/** \brief Single-SNP regression with partial effects
 *
 * Implements single-marker GWAS.  Each trait is treated as a single response, and the other traits are added to the SNP as predictors.  Only the SNP test is reported.
 * No Hotelling-type mutivariate association test is performed, so the number of columns in the results matrix is the same as the number of traits.
 * The output is either \f$ -\log_{10}p \f$, (although this statistic is not strictly a frequentist \f$p\f$-value, it performs very similarly in simulations), or Wakefield's \cite wakefield07 approximation of \f$ -ln BF \f$ (log-Bayes factor ratio).  In the latter case, the user can set the prior variance manually.
 * The SNP regression is performed on the point estimate of a response, as described in documentation of the update() and dump() functions.  The latter can be, say, a residual of a mixed-model type GEBV estimate done to control population structure.
 */
class BetaGrpPSR : public BetaGrpSnp {
protected:
	
public:
	/** \brief Default constructor */
	BetaGrpPSR() : BetaGrpSnp() {};
	/** \brief Constructor with no replication and \f$p\f$-values
	 *
	 * Initializes the object with no replication (i.e., the number of rows in the predictor is the same as in the response) and leaves the prior variance for SNP regressions at zero, thereby resulting in a dump of \f$ -\log_{10}p \f$ values.
	 *
	 * \param[in] string& predictor file name
	 * \param[in] string& output file name
	 * \param[in] size_t& number of rows in the response matrix
	 * \param[in] size_t& number of predictors
	 * \param[in] size_t& number of traits
	 * \param[in] int& number of threads
	 *
	 */
	BetaGrpPSR(const string &predFlNam, const string &outFlNam, const size_t &Ndat, const size_t &Npred, const size_t &d, const int &Nthr) : BetaGrpSnp(predFlNam, outFlNam, Ndat, Npred, d, Nthr) {};
	/** \brief Constructor with replication and \f$p\f$-values
	 *
	 * Initializes the object with replication, encoded with the given index, and leaves the prior variance for SNP regressions at zero, thereby resulting in a dump of \f$ -\log_{10}p \f$ values.
	 *
	 * \param[in] string& predictor file name
	 * \param[in] string& output file name
	 * \param[in] RanIndex& replicate index
	 * \param[in] size_t& number of predictors
	 * \param[in] size_t& number of traits
	 * \param[in] int& number of threads
	 *
	 */
	BetaGrpPSR(const string &predFlNam, const string &outFlNam, RanIndex &low, const size_t &Npred, const size_t &d, const int &Nthr) : BetaGrpSnp(predFlNam, outFlNam, low, Npred, d, Nthr) {};
	/** \brief Constructor with no replication and ABF
	 *
	 * Initializes the object with no replication (i.e., the number of rows in the predictor is the same as in the response) and sets the prior variance for SNP regressions to the given value, therefore dumping Wakefield's log of approximate Bayes factor (ABF) ratios to the output file.
	 *
	 * \param[in] string& predictor file name
	 * \param[in] string& output file name
	 * \param[in] size_t& number of rows in the response matrix
	 * \param[in] size_t& number of predictors
	 * \param[in] size_t& number of traits
	 * \param[in] int& number of threads
	 * \param[in] double& prior variance
	 *
	 */
	BetaGrpPSR(const string &predFlNam, const string &outFlNam, const size_t &Ndat, const size_t &Npred, const size_t &d, const int &Nthr, const double &prVar) : BetaGrpSnp(predFlNam, outFlNam, Ndat, Npred, d, Nthr, prVar) {};
	/** \brief Constructor with replication and ABF
	 *
	 * Initializes the object with replication, encoded with the given index, and sets the prior variance for SNP regressions to the given value, therefore dumping Wakefield's log of approximate Bayes factor (ABF) ratios to the output file.
	 *
	 * \param[in] string& predictor file name
	 * \param[in] string& output file name
	 * \param[in] RanIndex& replicate index
	 * \param[in] size_t& number of predictors
	 * \param[in] size_t& number of traits
	 * \param[in] int& number of threads
	 * \param[in] double& prior variance
	 *
	 */
	BetaGrpPSR(const string &predFlNam, const string &outFlNam, RanIndex &low, const size_t &Npred, const size_t &d, const int &Nthr, const double &prVar) : BetaGrpSnp(predFlNam, outFlNam, low, Npred, d, Nthr, prVar) {};
	
	/** \brief Destructor */
	~BetaGrpPSR() {};
	
	/** \brief Copy constructor
	 *
	 * \param[in] BetaGrpPSR& object to be copied
	 */
	BetaGrpPSR(const BetaGrpPSR &mG);
	/** \brief Assignment operator
	 *
	 * \param[in] BetaGrpPSR& object to be copied
	 * \return BetaGrpPSR& target object
	 */
	BetaGrpPSR & operator=(const BetaGrpPSR &mG);
	
	void dump();
	
};

/** \brief Simple single-SNP regression class with missing data
 *
 * Implements single-marker GWAS with missing genotypes.  Rows with missing genotypes are dropped.  Each trait is treated separately, including the variances (i.e., \f$ \widehat{\sigma}^2_p = \widehat{\boldsymbol{\Sigma}}_{p,p} \f$).  However, in addition to the single-trait tests an extra Hotelling-type mutivariate association test is performed.
 * This multivariate test reflects the distance of the whole vector of trait effects to zero, potentially increasing the power to detect pleiotropic SNPs with moderate effects on mutiple traits.
 * The saved value matrix thus has one extra column for this test.  The output is either \f$ -\log_{10}p \f$, (although this statistic is not strictly a frequentist \f$p\f$-value, it performs very similarly in simulations), or Wakefield's \cite wakefield07 approximation of \f$ -ln BF \f$ (log-Bayes factor ratio).  In the latter case, the user can set the prior variance manually.
 * The SNP regression is performed on the point estimate of a response, as described in documentation of the update() and dump() functions.  The latter can be, say, a residual of a mixed-model type GEBV estimate done to control population structure.
 *
 */
class BetaGrpSnpMiss : public MuGrp {
protected:
	/** \brief Predictor array
	 *
	 * The outer vector is the same length as the number of predictors.  Each constituent vector stores only the non-missing genotypes.
	 */
	vector< vector<double> > _Xmat;
	
	/** \brief Matrix for addition operators
	 *
	 * This matrix is addressed by fMat() and set to all zero values, so that if this object is in an addition/subtraction operator things don't break.  Because this object operates outside of the regular Markov chains, on the residuals of the model, it ordinarily makes no sense to add or subtract it from anything.
	 */
	gsl_matrix *_fakeFmat;
	
	/** \brief Response storage
	 *
	 * MCMC samples from a response variable (representing a residual of the model) are stored here and divided by the total number of saves at the end before the regression is performed.
	 */
	gsl_matrix *_Ystore;
	
	/** \brief Number of predictors (SNPs) */
	size_t _Npred;
	/** \brief Number of threads to use */
	int _nThr;
	/** \brief Number of saves
	 *
	 * Number of times the response samples have been saved.
	 */
	double _numSaves;
	/** \brief Missing value label
	 * 
	 * Value that labels the missing genotypes (predictors). Provided by the user.
	 */
	double _absLab;
	/** \brief Prior variance
	 *
	 * Prior variance for Wakefield's \cite wakefield07 ABF calculation (Wakefield's notation for it is \f$W\f$).  If it zero, \f$ -\log_{10}p \f$ are calculated.
	 */
	double _priorVar;
	
	/** \brief Predictor (SNP) file name */
	string _inPredFl;
	
public:
	/** \brief Default constructor */
	BetaGrpSnpMiss();
	/** \brief Constructor with no replication and \f$p\f$-values
	 *
	 * Initializes the object with no replication (i.e., the number of rows in the predictor is the same as in the response) and leaves the prior variance for SNP regressions at zero, thereby resulting in a dump of \f$ -\log_{10}p \f$ values.
	 *
	 * \param[in] string& predictor file name
	 * \param[in] string& output file name
	 * \param[in] size_t& number of rows in the response matrix
	 * \param[in] size_t& number of predictors
	 * \param[in] size_t& number of traits
	 * \param[in] int& number of threads
	 * \param[in] double& missing value label
	 *
	 */
	BetaGrpSnpMiss(const string &predFlNam, const string &outFlNam, const size_t &Ndat, const size_t &Npred, const size_t &d, const int &Nthr, const double &absLab);
	/** \brief Constructor with replication and \f$p\f$-values
	 *
	 * Initializes the object with replication, encoded with the given index, and leaves the prior variance for SNP regressions at zero, thereby resulting in a dump of \f$ -\log_{10}p \f$ values.
	 *
	 * \param[in] string& predictor file name
	 * \param[in] string& output file name
	 * \param[in] RanIndex& replicate index
	 * \param[in] size_t& number of predictors
	 * \param[in] size_t& number of traits
	 * \param[in] int& number of threads
	 * \param[in] double& missing value label
	 *
	 */
	BetaGrpSnpMiss(const string &predFlNam, const string &outFlNam, RanIndex &low, const size_t &Npred, const size_t &d, const int &Nthr, const double &absLab);
	/** \brief Constructor with no replication and ABF
	 *
	 * Initializes the object with no replication (i.e., the number of rows in the predictor is the same as in the response) and sets the prior variance for SNP regressions to the given value, therefore dumping Wakefield's log of approximate Bayes factor (ABF) ratios to the output file.
	 *
	 * \param[in] string& predictor file name
	 * \param[in] string& output file name
	 * \param[in] size_t& number of rows in the response matrix
	 * \param[in] size_t& number of predictors
	 * \param[in] size_t& number of traits
	 * \param[in] int& number of threads
	 * \param[in] double& prior variance
	 * \param[in] double& missing value label
	 *
	 */
	BetaGrpSnpMiss(const string &predFlNam, const string &outFlNam, const size_t &Ndat, const size_t &Npred, const size_t &d, const int &Nthr, const double &prVar, const double &absLab);
	/** \brief Constructor with replication and ABF
	 *
	 * Initializes the object with replication, encoded with the given index, and sets the prior variance for SNP regressions to the given value, therefore dumping Wakefield's log of approximate Bayes factor (ABF) ratios to the output file.
	 *
	 * \param[in] string& predictor file name
	 * \param[in] string& output file name
	 * \param[in] RanIndex& replicate index
	 * \param[in] size_t& number of predictors
	 * \param[in] size_t& number of traits
	 * \param[in] int& number of threads
	 * \param[in] double& prior variance
	 * \param[in] double& missing value label
	 *
	 */
	BetaGrpSnpMiss(const string &predFlNam, const string &outFlNam, RanIndex &low, const size_t &Npred, const size_t &d, const int &Nthr, const double &prVar, const double &absLab);
	
	/** \brief Destructor */
	~BetaGrpSnpMiss();
	
	/** \brief Copy constructor
	 *
	 * \param[in] BetaGrpSnpMiss& object to be copied
	 */
	BetaGrpSnpMiss(const BetaGrpSnpMiss &mG); // copy constructor
	/** \brief Assignment operator
	 *
	 * \param[in] BetaGrpSnpMiss& object to be copied
	 * \return BetaGrpSnpMiss& target object
	 */
	BetaGrpSnpMiss & operator=(const BetaGrpSnpMiss &mG);
	
	/** \brief Dump results to the output file
	 *
	 * The predictor (SNP) file is read by this function, and the single-predictor regression (i.e., each predictor is tested separately, ignoring all others) is performed.  The results are saved to the pre-specified output file.
	 */
	virtual void dump();
	
	/** \brief Access adjusted fitted value matrix
	 *
	 * \return gsl_matrix* pointer to a zero-valued matrix
	 */
	const gsl_matrix *fMat() const{return _fakeFmat; };
	
	/** \brief Response update function
	 *
	 * Unlike standard update functions, this one only saves MCMC samples of the response.  The covariance provided is ignored.
	 * The actual regression is performed on the point estimates of response values calculated as means of the stored MCMC values, and is done at the end by envoking the dump() function.
	 *
	 * \ingroup updateFun
	 *
	 * \param[in] Grp& response variable to be saved
	 * \param[in] SigmaI& inverse-covariance (ignored)
	 */
	void update(const Grp &dat, const SigmaI &SigIm);
	
};

/** \brief Single-SNP regression with conditional variance and missing data
 *
 * Implements single-marker GWAS with missing genotype data.  Rows with missing genotypes are dropped.  Each trait is treated separately for coefficient calculation.  In contrast, the variance estimates are taken from the inverse-covariance: \f$ \widehat{\sigma}^2_p = \left[ \widehat{\boldsymbol{\Sigma}}^{-1}_{p,p} \right]^{-1} \le \widehat{\boldsymbol{\Sigma}}_{p,p} \f$.
 * In addition to the single-trait tests an extra Hotelling-type mutivariate association test is performed.
 * This multivariate test reflects the distance of the whole vector of trait effects to zero, potentially increasing the power to detect pleiotropic SNPs with moderate effects on mutiple traits.
 * The saved value matrix thus has one extra column for this test.  The output is either \f$ -\log_{10}p \f$, (although this statistic is not strictly a frequentist \f$p\f$-value, it performs very similarly in simulations), or Wakefield's \cite wakefield07 approximation of \f$ -ln BF \f$ (log-Bayes factor ratio).  In the latter case, the user can set the prior variance manually.
 * The SNP regression is performed on the point estimate of a response, as described in documentation of the update() and dump() functions.  The latter can be, say, a residual of a mixed-model type GEBV estimate done to control population structure.
 */
class BetaGrpSnpMissCV : public BetaGrpSnpMiss {
protected:
	
public:
	/** \brief Default constructor */
	BetaGrpSnpMissCV() : BetaGrpSnpMiss() {};
	/** \brief Constructor with no replication and \f$p\f$-values
	 *
	 * Initializes the object with no replication (i.e., the number of rows in the predictor is the same as in the response) and leaves the prior variance for SNP regressions at zero, thereby resulting in a dump of \f$ -\log_{10}p \f$ values.
	 *
	 * \param[in] string& predictor file name
	 * \param[in] string& output file name
	 * \param[in] size_t& number of rows in the response matrix
	 * \param[in] size_t& number of predictors
	 * \param[in] size_t& number of traits
	 * \param[in] int& number of threads
	 * \param[in] double& missing value label
	 *
	 */
	BetaGrpSnpMissCV(const string &predFlNam, const string &outFlNam, const size_t &Ndat, const size_t &Npred, const size_t &d, const int &Nthr, const double &absLab) : BetaGrpSnpMiss(predFlNam, outFlNam, Ndat, Npred, d, Nthr, absLab) {};
	/** \brief Constructor with replication and \f$p\f$-values
	 *
	 * Initializes the object with replication, encoded with the given index, and leaves the prior variance for SNP regressions at zero, thereby resulting in a dump of \f$ -\log_{10}p \f$ values.
	 *
	 * \param[in] string& predictor file name
	 * \param[in] string& output file name
	 * \param[in] RanIndex& replicate index
	 * \param[in] size_t& number of predictors
	 * \param[in] size_t& number of traits
	 * \param[in] int& number of threads
	 * \param[in] double& missing value label
	 *
	 */
	BetaGrpSnpMissCV(const string &predFlNam, const string &outFlNam, RanIndex &low, const size_t &Npred, const size_t &d, const int &Nthr, const double &absLab) : BetaGrpSnpMiss(predFlNam, outFlNam, low, Npred, d, Nthr, absLab) {};
	/** \brief Constructor with no replication and ABF
	 *
	 * Initializes the object with no replication (i.e., the number of rows in the predictor is the same as in the response) and sets the prior variance for SNP regressions to the given value, therefore dumping Wakefield's log of approximate Bayes factor (ABF) ratios to the output file.
	 *
	 * \param[in] string& predictor file name
	 * \param[in] string& output file name
	 * \param[in] size_t& number of rows in the response matrix
	 * \param[in] size_t& number of predictors
	 * \param[in] size_t& number of traits
	 * \param[in] int& number of threads
	 * \param[in] double& prior variance
	 * \param[in] double& missing value label
	 *
	 */
	BetaGrpSnpMissCV(const string &predFlNam, const string &outFlNam, const size_t &Ndat, const size_t &Npred, const size_t &d, const int &Nthr, const double &prVar, const double &absLab) : BetaGrpSnpMiss(predFlNam, outFlNam, Ndat, Npred, d, Nthr, prVar, absLab) {};
	/** \brief Constructor with replication and ABF
	 *
	 * Initializes the object with replication, encoded with the given index, and sets the prior variance for SNP regressions to the given value, therefore dumping Wakefield's log of approximate Bayes factor (ABF) ratios to the output file.
	 *
	 * \param[in] string& predictor file name
	 * \param[in] string& output file name
	 * \param[in] RanIndex& replicate index
	 * \param[in] size_t& number of predictors
	 * \param[in] size_t& number of traits
	 * \param[in] int& number of threads
	 * \param[in] double& prior variance
	 * \param[in] double& missing value label
	 *
	 */
	BetaGrpSnpMissCV(const string &predFlNam, const string &outFlNam, RanIndex &low, const size_t &Npred, const size_t &d, const int &Nthr, const double &prVar, const double &absLab) : BetaGrpSnpMiss(predFlNam, outFlNam, low, Npred, d, Nthr, prVar, absLab) {};
	
	/** \brief Destructor */
	~BetaGrpSnpMissCV(){};
	
	/** \brief Copy constructor
	 *
	 * \param[in] BetaGrpSnpMissCV& object to be copied
	 */
	BetaGrpSnpMissCV(const BetaGrpSnpMissCV &mG);
	/** \brief Assignment operator
	 *
	 * \param[in] BetaGrpSnpMissCV& object to be copied
	 * \return BetaGrpSnpMissCV& target object
	 */
	BetaGrpSnpMissCV & operator=(const BetaGrpSnpMissCV &mG);
	
	void dump();
	
};

/** \brief Single-SNP regression with partial effects and missing data
 *
 * Implements single-marker GWAS with missing genotype data.  Rows with missing genotypes are dropped.  Each trait is treated as a single response, and the other traits are added to the SNP as predictors.  Only the SNP test is reported.
 * No Hotelling-type mutivariate association test is performed, so the number of columns in the results matrix is the same as the number of traits.
 * The output is either \f$ -\log_{10}p \f$, (although this statistic is not strictly a frequentist \f$p\f$-value, it performs very similarly in simulations), or Wakefield's \cite wakefield07 approximation of \f$ -ln BF \f$ (log-Bayes factor ratio).  In the latter case, the user can set the prior variance manually.
 * The SNP regression is performed on the point estimate of a response, as described in documentation of the update() and dump() functions.  The latter can be, say, a residual of a mixed-model type GEBV estimate done to control population structure.
 */
class BetaGrpPSRmiss : public BetaGrpSnpMiss {
protected:
	
public:
	/** \brief Default constructor */
	BetaGrpPSRmiss() : BetaGrpSnpMiss() {};
	/** \brief Constructor with no replication and \f$p\f$-values
	 *
	 * Initializes the object with no replication (i.e., the number of rows in the predictor is the same as in the response) and leaves the prior variance for SNP regressions at zero, thereby resulting in a dump of \f$ -\log_{10}p \f$ values.
	 *
	 * \param[in] string& predictor file name
	 * \param[in] string& output file name
	 * \param[in] size_t& number of rows in the response matrix
	 * \param[in] size_t& number of predictors
	 * \param[in] size_t& number of traits
	 * \param[in] int& number of threads
	 * \param[in] double& missing value label
	 *
	 */
	BetaGrpPSRmiss(const string &predFlNam, const string &outFlNam, const size_t &Ndat, const size_t &Npred, const size_t &d, const int &Nthr, const double &absLab) : BetaGrpSnpMiss(predFlNam, outFlNam, Ndat, Npred, d, Nthr, absLab) {};
	/** \brief Constructor with replication and \f$p\f$-values
	 *
	 * Initializes the object with replication, encoded with the given index, and leaves the prior variance for SNP regressions at zero, thereby resulting in a dump of \f$ -\log_{10}p \f$ values.
	 *
	 * \param[in] string& predictor file name
	 * \param[in] string& output file name
	 * \param[in] RanIndex& replicate index
	 * \param[in] size_t& number of predictors
	 * \param[in] size_t& number of traits
	 * \param[in] int& number of threads
	 * \param[in] double& missing value label
	 *
	 */
	BetaGrpPSRmiss(const string &predFlNam, const string &outFlNam, RanIndex &low, const size_t &Npred, const size_t &d, const int &Nthr, const double &absLab) : BetaGrpSnpMiss(predFlNam, outFlNam, low, Npred, d, Nthr, absLab) {};
	/** \brief Constructor with no replication and ABF
	 *
	 * Initializes the object with no replication (i.e., the number of rows in the predictor is the same as in the response) and sets the prior variance for SNP regressions to the given value, therefore dumping Wakefield's log of approximate Bayes factor (ABF) ratios to the output file.
	 *
	 * \param[in] string& predictor file name
	 * \param[in] string& output file name
	 * \param[in] size_t& number of rows in the response matrix
	 * \param[in] size_t& number of predictors
	 * \param[in] size_t& number of traits
	 * \param[in] int& number of threads
	 * \param[in] double& prior variance
	 * \param[in] double& missing value label
	 *
	 */
	BetaGrpPSRmiss(const string &predFlNam, const string &outFlNam, const size_t &Ndat, const size_t &Npred, const size_t &d, const int &Nthr, const double &prVar, const double &absLab) : BetaGrpSnpMiss(predFlNam, outFlNam, Ndat, Npred, d, Nthr, prVar, absLab) {};
	/** \brief Constructor with replication and ABF
	 *
	 * Initializes the object with replication, encoded with the given index, and sets the prior variance for SNP regressions to the given value, therefore dumping Wakefield's log of approximate Bayes factor (ABF) ratios to the output file.
	 *
	 * \param[in] string& predictor file name
	 * \param[in] string& output file name
	 * \param[in] RanIndex& replicate index
	 * \param[in] size_t& number of predictors
	 * \param[in] size_t& number of traits
	 * \param[in] int& number of threads
	 * \param[in] double& prior variance
	 * \param[in] double& missing value label
	 *
	 */
	BetaGrpPSRmiss(const string &predFlNam, const string &outFlNam, RanIndex &low, const size_t &Npred, const size_t &d, const int &Nthr, const double &prVar, const double &absLab) : BetaGrpSnpMiss(predFlNam, outFlNam, low, Npred, d, Nthr, prVar, absLab) {};
	
	/** \brief Destructor */
	~BetaGrpPSRmiss() {};
	
	/** \brief Copy constructor
	 *
	 * \param[in] BetaGrpPSRmiss& object to be copied
	 */
	BetaGrpPSRmiss(const BetaGrpPSRmiss &mG);
	/** \brief Assignment operator
	 *
	 * \param[in] BetaGrpPSRmiss& object to be copied
	 * \return BetaGrpPSRmiss& target object
	 */
	BetaGrpPSRmiss & operator=(const BetaGrpPSRmiss &mG);
	
	void dump();

};

/** \brief Bayesian variable selection regression
 *
 * A multivariate implementation of Bayesian variable selection regression (BVSR), together with the RanIndexVS class.  The model is similar to that described in \cite guan11 , but unlike their method a pre-selection of predictors based on a first-pass single-predictor regression is made.
 * The user has control over the number of predictors to choose, and the cut-off for correlation among predictors (which arises as linkage disequilibrium in SNP regressions).  This numebr is expressed as a multiple of genetic sample size. 
 * Once the pre-selected group of predictors is formed, variable selection is performed within the smaller group.  Predictors discarded for correlation with picked variables are assigned the same probability of retention in the model as their corresponding picked predictors.
 * Rao-Blackwellised values of regression coefficients are saved by the dump() function.
 *
 * \warning This class is experimental and has not been extensively tested
 */
class BetaGrpBVSR : public BetaGrpFt {
	friend void RanIndexVS::update(const Grp &, const SigmaI &, BetaGrpBVSR *, const SigmaI &);
protected:
	/** \brief Matrix of selected predictors
	 *
	 * Contains only the predictors currently in the model.
	 */
	gsl_matrix *_selB;
	/** \brief \f$ \boldsymbol{\mathrm{x}}_{\cdot l}\boldsymbol{\beta}_{l \cdot}\f$ matrix for the candidate dropped/added element  */
	gsl_matrix *_tmpXb;
	
	/** \brief Update fitted values 
	 *
	 * \ingroup fitFun
	 */
	void _updateFitted();
	/** \brief Testing candidates for correlation
	 *
	 * Goes through the list of "top" predictors and eliminates lower-ranked candidates correlated with them.  If any candidates are eliminated, the list of top predictors is augmented with predictors previously discarded.
	 * The identity of the predictors eliminated for correlation is saved.  The correlation among predictors arises, for example, when estimating SNP effect in genetics (GWAS).
	 *
	 * \warning Tested only superficially
	 *
	 * \param[in] gsl_vector* \f$ \left( \boldsymbol{x}^T\boldsymbol{x} \right)^{-1} \f$
	 * \param[in] gsl_permutation* predictor ranks
	 * \param[in] double& \f$ r^2 \f$ cut-off
	 * \param[in] size_t& number of predictors to pick
	 * \param[out] vector< vector<size_t> >& index of picked predictors
	 * \param[out] vector< vector<size_t> >& index relating dropped correlated predictors to their group-defining predictor
	 * \param[out] gsl_matrix* matrix of picked predictor values
	 */
	void _ldToss(const gsl_vector *var, const gsl_permutation *prm, const double &rSqMax, const size_t &Nsel, const size_t &Npck, vector< vector<size_t> > &idx, vector< vector<size_t> > &rLd, gsl_matrix *Xpck);
	/** \brief Gaussian kernel
	 *
	 * Calculates the multivariate Gaussian kernel value for all regression coefficients in the model.
	 *
	 * \param[in] Grp& data
	 * \param[in] SigmaI& inverse-covariance
	 *
	 * \return double kernel value
	 */
	double _MGkernel(const Grp &dat, const SigmaI &SigI) const;
	/** \brief Gaussian kernel dropping one predictor
	 *
	 * Calculates the multivariate Gaussian kernel value for all regression coefficients in the model, except for the one indicated.
	 *
	 * \param[in] Grp& data
	 * \param[in] SigmaI& inverse-covariance
	 * \param[in] size_t& index of the dropped predictor
	 *
	 * \return double kernel value
	 */
	double _MGkernel(const Grp &dat, const SigmaI &SigI, const size_t &prInd) const;  // MV Gaussian kernel, dropping predictor prInd
	/** \brief Gaussian kernel adding one predictor
	 *
	 * Calculates the multivariate Gaussian kernel value for all regression coefficients in the model, plus the indicated extra predictor with a Gaussian zero-mean prior.
	 *
	 * \param[in] Grp& data
	 * \param[in] SigmaI& data inverse-covariance
	 * \param[in] SigmaI& prior inverse-covariance
	 * \param[in] size_t& index of the dropped predictor
	 *
	 * \return double kernel value
	 */
	double _MGkernel(const Grp &dat, const SigmaI &SigIe, const SigmaI &SigIpr, const size_t &prInd);
public:
	/** \brief Default constructor */
	BetaGrpBVSR() : BetaGrpFt() {};
	/** \brief Basic constructor
	 *
	 * Constructor with no missing data in predictors and no replication.  Performs the initial pre-selection of predictors.
	 *
	 * \param[in] Grp& data
	 * \param[in] SigmaI& data inverse-covariance
	 * \param[in] string& preddictor file name
	 * \param[in] double& number of predictors to keep, in multiples of sample size
	 * \param[in] double& \f$ r^2 \f$ cut-off for predictor correlation
	 * \param[in] RanIndex& prior index
	 * \param[in] string& output file name
	 * \param[in] int& number of threads
	 *
	 */
	BetaGrpBVSR(const Grp &y, const SigmaI &SigI, const string &predFlNam, const double &Nmul, const double &rSqMax, RanIndex &up, const string &outFlNam, const int &nThr);
	/** \brief Basic constructor with replication
	 *
	 * Constructor with no missing data in predictors and replication.  A RanIndex index is used instead of a design matrix.  Performs the initial pre-selection of predictors.
	 *
	 * \param[in] Grp& data
	 * \param[in] SigmaI& data inverse-covariance
	 * \param[in] string& preddictor file name
	 * \param[in] double& number of predictors to keep, in multiples of sample size
	 * \param[in] double& \f$ r^2 \f$ cut-off for predictor correlation
	 * \param[in] RanIndex& replicate (data) index
	 * \param[in] RanIndex& prior index
	 * \param[in] string& output file name
	 * \param[in] int& number of threads
	 *
	 */
	BetaGrpBVSR(const Grp &y, const SigmaI &SigI, const string &predFlNam, const double &Nmul, const double &rSqMax, RanIndex &low, RanIndex &up, const string &outFlNam, const int &nThr);
	/** \brief Basic constructor with missing data
	 *
	 * Constructor with missing data in predictors and no replication. Missing data in the selected predictors are filled in using mean imputation. Performs the initial pre-selection of predictors.
	 *
	 * \param[in] Grp& data
	 * \param[in] SigmaI& data inverse-covariance
	 * \param[in] string& preddictor file name
	 * \param[in] double& number of predictors to keep, in multiples of sample size
	 * \param[in] double& \f$ r^2 \f$ cut-off for predictor correlation
	 * \param[in] double& missing data label
	 * \param[in] RanIndex& prior index
	 * \param[in] string& output file name
	 * \param[in] int& number of threads
	 *
	 */
	BetaGrpBVSR(const Grp &y, const SigmaI &SigI, const string &predFlNam, const double &Nmul, const double &rSqMax, const double &absLab, RanIndex &up, const string &outFlNam, const int &nThr); // absLab is the token that signifies missing data
	/** \brief Basic constructor with replication and missing data
	 *
	 * Constructor with missing data in predictors and replication.  Missing data in the selected predictors are filled in using mean imputation. A RanIndex index is used instead of a design matrix.  Performs the initial pre-selection of predictors.
	 *
	 * \param[in] Grp& data
	 * \param[in] SigmaI& data inverse-covariance
	 * \param[in] string& preddictor file name
	 * \param[in] double& number of predictors to keep, in multiples of sample size
	 * \param[in] double& \f$ r^2 \f$ cut-off for predictor correlation
	 * \param[in] double& missing data label
	 * \param[in] RanIndex& replicate (data) index
	 * \param[in] RanIndex& prior index
	 * \param[in] string& output file name
	 * \param[in] int& number of threads
	 *
	 */
	BetaGrpBVSR(const Grp &y, const SigmaI &SigI, const string &predFlNam, const double &Nmul, const double &rSqMax, const double &absLab, RanIndex &low, RanIndex &up, const string &outFlNam, const int &nThr);
	
	/** \brief Destructor */
	~BetaGrpBVSR();
	
	/** \brief Regression value store
	 *
	 * Stores \f$t\f$-statistic values for each trait and pre-selected (i.e. whether or not currently in the model) regression coefficient in a variable to subsequently save to a file by dump().
	 *
	 * \param[in] SigmaI& data inverse-covariance
	 */
	void save(const SigmaI &SigI);
	/** \brief Regression value and probability store
	 *
	 * Stores \f$t\f$-statistic values for each trait and pre-selected (i.e. whether or not currently in the model) regression coefficient in a variable to subsequently save to a file by dump().
	 * Additionally, (\f$ -\log_{10}) Rao-Blackwellised probabilities that each pre-selected predictor is not in the model (PEP of RanIndexVS) is also stored.
	 *
	 * \param[in] Grp& data
	 * \param[in] SigmaI& data inverse-covariance
	 */
	void save(const Grp &y, const SigmaI &SigI);
	void dump();
	/** \brief Access to the values currently in the model */
	const gsl_matrix *dMat() const{return _selB; };
	
	void update(const Grp &dat, const SigmaI &SigIm, const SigmaI &SigIp);
};

/** \brief Hierarchical mean with independent blocks of traits
 *
 * Blocks of inter-correlated traits with correlations among blocks set to exactly zero.  These models arise, for example, when traits measured in different environments are modeled together as different traits.
 * Replication structure and the number of replicates and levels (rows in the would-be _valueMat) can be different among blocks.  The only limitation is that the traits belonging to the same block have to be in adjacent columns.
 */
class MuBlk : public MuGrp {
protected:
	/** \brief Block start indexes
	 *
	 * Vector of indexes that indicate the column that starts each block.
	 */
	vector< size_t > _blkStart;
	/** \brief Vector of low level indexes
	 *
	 * Indexes to data, similar to _lowLevel.  These are separate for each block, not necessarily the same number of levels or replicates in each.
	 */
	vector< vector< vector<size_t> > > _blkLow;
	/** \brief Expanded _valueMat
	 *
	 * Expanded value matrix, with the rows repeated according to _blkLow, separately for each block.  To be used for arithmetic operators.
	 */
	gsl_matrix *_expandedVM;
	/** \brief Blocks with few levels
	 *
	 * For cases where the number of levels of _blkLow differ among blocks, this vector stores the index of the first element past the available one.  
	 * For blocks that have the same number of levels as the largest among blocks, the value is zero.  Each object has to have at least one zero in this vector, which is for the block with the largest number of levels.
	 */
	vector<size_t> _shortLevels;
	
	/** \brief Update the _expandedVM matrix */
	void _updateExp();
	
	/** \brief Fill in blocks with few levels
	 *
	 * For blocks with the number of levels smaller than the maximum, fill in the value matrix with the overall mean among levels.  This insures that the overall mean is equal to what it would be if only the available rows were used.
	 */
	void _fillIn();
	/** \brief Fill in blocks with few levels and a prior grouping
	 *
	 * The same as _fillIn(), but the means are calculated according to the grouping in the prior (upper) level in the hierarchy.
	 */
	void _fillInUp();
	
public:
	/** \brief Default constructor */
	MuBlk() : MuGrp() {};
	/** \brief Basic constructor
	 *
	 * The provided number of values is the number of rows in the value matrix, i.e. the maximum number of levels of the lower level (data) index.
	 * The file that has the lower level index information should have a matrix of _int_ with the number of rows the same as the number of rows in the data, and the number of columns equal to the number of blocks.
	 *
	 * \param[in] Grp& data for initialization
	 * \param[in] string& name of the file with the low-level index
	 * \param[in] size_t& maximum number of levels
	 * \param[in] RanIndex& prior index
	 * \param[in] string& name of the file with the block indexes
	 */
	MuBlk(const Grp &dat, const string &lowIndFlName, const size_t &Nval, RanIndex &up, const string &blkIndFileNam);
	/** \brief Constructor with output file name
	 *
	 * The provided number of values is the number of rows in the value matrix, i.e. the maximum number of levels of the lower level (data) index.
	 * The file that has the lower level index information should have a matrix of _int_ with the number of rows the same as the number of rows in the data, and the number of columns equal to the number of blocks.
	 *
	 * \param[in] Grp& data for initialization
	 * \param[in] string& name of the file with the low-level index
	 * \param[in] size_t& maximum number of levels
	 * \param[in] RanIndex& prior index
	 * \param[in] string& output file name
	 * \param[in] string& name of the file with the block indexes
	 */
	MuBlk(const Grp &dat, const string &lowIndFlName, const size_t &Nval, RanIndex &up, const string &outFlNam, const string &blkIndFileNam);
	
	/** \brief Destructor */
	~MuBlk() {gsl_matrix_free(_expandedVM); };
	
	/** \brief Access to the expanded value matrix */
	const gsl_matrix *fMat() const{ return _expandedVM; };
	
	// improper prior
	void update(const Grp &dat, const SigmaI &SigIm);
	void update(const Grp &dat, const Qgrp &q, const SigmaI &SigIm);
	// 0-mean prior
	void update(const Grp &dat, const SigmaI &SigIm, const SigmaI &SigIp);
	void update(const Grp &dat, const Qgrp &q, const SigmaI &SigIm, const SigmaI &SigIp);
	void update(const Grp &dat, const SigmaI &SigIm, const Qgrp &qPr, const SigmaI &SigIp);
	void update(const Grp &dat, const Qgrp &q, const SigmaI &SigIm, const Qgrp &qPr, const SigmaI &SigIp);
	// non-0-mean prior
	void update(const Grp &dat, const SigmaI &SigIm, const Grp &muPr, const SigmaI &SigIp);
	void update(const Grp &dat, const Qgrp &q, const SigmaI &SigIm, const Grp &muPr, const SigmaI &SigIp);
	void update(const Grp &dat, const SigmaI &SigIm, const Grp &muPr, const Qgrp &qPr, const SigmaI &SigIp);
	void update(const Grp &dat, const Qgrp &q, const SigmaI &SigIm, const Grp &muPr, const Qgrp &qPr, const SigmaI &SigIp);
};


/** \brief Regression with independent blocks of traits
 *
 * Blocks of inter-correlated traits with correlations among blocks set to exactly zero.  These models arise, for example, when traits measured in different environments are modeled together as different traits.
 * The traits belonging to the same block have to be in adjacent columns.  The number of predictors per block must be the same (future devolpment will drop this requirement), and the predictor columns corresponding to the same trait block have to be adjacent in the _Xmat matrix.
 */
class BetaBlk : public BetaGrpFt {

protected:
	/** \brief Block start indexes
	 *
	 * Vector of indexes that indicate the column that starts each block.
	 */
	vector< size_t > _blkStart;
	/** \brief Predictor submatrices
	 *
	 * A vector of GSL matrix views of the submatrices of the overall predictor matrix that correspond to each trait block.
	 */
	vector< gsl_matrix_view > _eachX;
	/** \brief Regression coefficient submatrices
	 *
	 * A vector of GSL matrix views of the overall value matrix.
	 */
	vector< gsl_matrix_view > _eachB;
	
	/** \brief Update fitted values
	 *
	 * \ingroup fitFun
	 */
	void _updateFitted();
public:
	/** \brief Default constructor */
	BetaBlk() : BetaGrpFt() {};
	/** \brief Basic constructor
	 *
	 * The total number of predictors (columns in _Xmat) is the provided number of predictors times the number of blocks.
	 * The file that has the block start indexes should be a white-space delimited text file.  The file that has the lower level index information should have a matrix of _int_ with the number of rows the same as the number of rows in the data, and the number of columns equal to the number of blocks.
	 *
	 * \param[in] Grp& data for initialization
	 * \param[in] string& name of the predictor file
	 * \param[in] size_t& number of predictors per block
	 * \param[in] string& name of the file with the block indexes
	 * \param[in] int& number of threads
	 */
	BetaBlk(const Grp &dat, const string &predFlName, const size_t &Npred, const string &blkIndFileNam, const int &nThr);
	/** \brief Constructor with output file name
	 *
	 * The total number of predictors (columns in _Xmat) is the provided number of predictors times the number of blocks.
	 * The file that has the block start indexes should be a white-space delimited text file.  The file that has the lower level index information should have a matrix of _int_ with the number of rows the same as the number of rows in the data, and the number of columns equal to the number of blocks.
	 *
	 * \param[in] Grp& data for initialization
	 * \param[in] string& name of the predictor file
	 * \param[in] size_t& number of predictors per block
	 * \param[in] string& output file name
	 * \param[in] string& name of the file with the block indexes
	 * \param[in] int& number of threads
	 */
	BetaBlk(const Grp &dat, const string &predFlName, const size_t &Npred, const string &outFlNam, const string &blkIndFileNam, const int &nThr);
	/** \brief Constructor with a prior index
	 *
	 * The total number of predictors (columns in _Xmat) is the provided number of predictors times the number of blocks.
	 * The file that has the block start indexes should be a white-space delimited text file.  The file that has the lower level index information should have a matrix of _int_ with the number of rows the same as the number of rows in the data, and the number of columns equal to the number of blocks.
	 *
	 * \param[in] Grp& data for initialization
	 * \param[in] string& name of the predictor file
	 * \param[in] size_t& number of predictors per block
	 * \param[in] RanIndex& prior (upper-level) index
	 * \param[in] string& name of the file with the block indexes
	 * \param[in] int& number of threads
	 */
	BetaBlk(const Grp &dat, const string &predFlName, const size_t &Npred, RanIndex &up, const string &blkIndFileNam, const int &nThr);
	/** \brief Constructor with a prior index and output file
	 *
	 * The total number of predictors (columns in _Xmat) is the provided number of predictors times the number of blocks.
	 * The file that has the block start indexes should be a white-space delimited text file.  The file that has the lower level index information should have a matrix of _int_ with the number of rows the same as the number of rows in the data, and the number of columns equal to the number of blocks.
	 *
	 * \param[in] Grp& data for initialization
	 * \param[in] string& name of the predictor file
	 * \param[in] size_t& number of predictors per block
	 * \param[in] RanIndex& prior (upper-level) index
	 * \param[in] string& output file name
	 * \param[in] string& name of the file with the block indexes
	 * \param[in] int& number of threads
	 */
	BetaBlk(const Grp &dat, const string &predFlName, const size_t &Npred, RanIndex &up, const string &outFlNam, const string &blkIndFileNam, const int &nThr);
	
	/** \brief Constructor with missing predictor data
	 *
	 * The total number of predictors (columns in _Xmat) is the provided number of predictors times the number of blocks.
	 * The file that has the block start indexes should be a white-space delimited text file.  The file that has the lower level index information should have a matrix of _int_ with the number of rows the same as the number of rows in the data, and the number of columns equal to the number of blocks.
	 *
	 * \param[in] Grp& data for initialization
	 * \param[in] string& name of the predictor file
	 * \param[in] size_t& number of predictors per block
	 * \param[in] double& missing value label
	 * \param[in] string& name of the file with the block indexes
	 * \param[in] int& number of threads
	 */
	BetaBlk(const Grp &dat, const string &predFlName, const size_t &Npred, const double &absLab, const string &blkIndFileNam, const int &nThr);
	/** \brief Constructor with missing predictor values and output file name
	 *
	 * The total number of predictors (columns in _Xmat) is the provided number of predictors times the number of blocks.
	 * The file that has the block start indexes should be a white-space delimited text file.  The file that has the lower level index information should have a matrix of _int_ with the number of rows the same as the number of rows in the data, and the number of columns equal to the number of blocks.
	 *
	 * \param[in] Grp& data for initialization
	 * \param[in] string& name of the predictor file
	 * \param[in] size_t& number of predictors per block
	 * \param[in] double& missing value label
	 * \param[in] string& output file name
	 * \param[in] string& name of the file with the block indexes
	 * \param[in] int& number of threads
	 */
	BetaBlk(const Grp &dat, const string &predFlName, const size_t &Npred, const double &absLab, const string &outFlNam, const string &blkIndFileNam, const int &nThr);
	/** \brief Constructor with a prior index and missing predictor data
	 *
	 * The total number of predictors (columns in _Xmat) is the provided number of predictors times the number of blocks.
	 * The file that has the block start indexes should be a white-space delimited text file.  The file that has the lower level index information should have a matrix of _int_ with the number of rows the same as the number of rows in the data, and the number of columns equal to the number of blocks.
	 *
	 * \param[in] Grp& data for initialization
	 * \param[in] string& name of the predictor file
	 * \param[in] size_t& number of predictors per block
	 * \param[in] double& missing data label
	 * \param[in] RanIndex& prior (upper-level) index
	 * \param[in] string& name of the file with the block indexes
	 * \param[in] int& number of threads
	 */
	BetaBlk(const Grp &dat, const string &predFlName, const size_t &Npred, const double &absLab, RanIndex &up, const string &blkIndFileNam, const int &nThr);
	/** \brief Constructor with a prior index, missing predictor data, and output file
	 *
	 * The total number of predictors (columns in _Xmat) is the provided number of predictors times the number of blocks.
	 * The file that has the block start indexes should be a white-space delimited text file.  The file that has the lower level index information should have a matrix of _int_ with the number of rows the same as the number of rows in the data, and the number of columns equal to the number of blocks.
	 *
	 * \param[in] Grp& data for initialization
	 * \param[in] string& name of the predictor file
	 * \param[in] size_t& number of predictors per block
	 * \param[in] double& missing data label
	 * \param[in] RanIndex& prior (upper-level) index
	 * \param[in] string& output file name
	 * \param[in] string& name of the file with the block indexes
	 * \param[in] int& number of threads
	 */
	BetaBlk(const Grp &dat, const string &predFlName, const size_t &Npred, const double &absLab, RanIndex &up, const string &outFlNam, const string &blkIndFileNam, const int &nThr);
	
	/** \brief Destructor */
	~BetaBlk() {};
	
	// updates handled by BetaGrpFt
};

// end of the locGrp group
/** @} */

/** \defgroup sigInv Inverse covariance matrices
 *
 * Group of classes that implement inverse-covariance matrix updating.  We are dealing with inverse-covarince (a.k.a. concentration) matrices because most algorithms in hierarchical models use these rather than covariance matrices. 
 * Thus, we save on matrix inversion operations by keeping track only of inverse-covariances.  The save functions, however, output covariance matrices.  All the internal algorithms for all the classes use the lower triangles of the underlying symmetric matrix.  On saving, the matrices are symmetric and it does not matter for downstream applications.
 * However, if a user wants to manipulate or interact with these in a sampler, addressing the lower triangle ensures stability of the algorithm.
 * @{
 */
/** \brief Basic inverse-covariance
 *
 * Standard model for inverse-covariance matrices with a diagonal Wishart prior (i.e., pre-supposing no correlations among traits).  The values on the diagonal of the prior matrix and the prior degrees of freedom can be varied to reflect the vagueness of prior information.
 * These conjugate priors allow for Gibbs sampling even when the number of traits exceeds the number of data points.  However, in the latter case the estimation of correlations will likely be inaccurate.  Algorithms involving direct penalties on correlations should perform better and are inder development.
 */
class SigmaI {
protected:
	/** \brief Inverse-covariance matrix 
	 *
	 * Square symmetric matrix.
	 */
	gsl_matrix *_mat;
	/** \brief Dimension of the matrix 
	 *
	 * The number of rows and columns of _mat.
	 */
	size_t _d;
	/** \brief Square root of the determinant
	 *
	 * Square root of the determinant of the corrent value of the inverse-covariance matrix.  Is calculated only if necessary, otherwise is set to -1.0 to provide an easy way to find out if it had been calculated at least once.
	 */
	double _srDet;
	
	/** \brief Prior inverse-covariance
	 *
	 * Diagonal matrix with zeros on the off-diagonal and inverse variance scaled by degrees of freedom on the diagonal.
	 */
	gsl_matrix *_LamSc;
	/** \brief Prior degrees of freedom */
	double _n0;
	
	/** \brief Output file name */
	string _outFlNam;
	
	/** \brief Pseudo-random number generator
	 *
	 * Initialized the same way as Grp type PNGs, with a sum of time and RTDSC.
	 */
	gsl_rng *_r;
	
public:
	/** \brief Default constructor 
	 *
	 * Creates a \f$ 2\times2 \f$ identity matrix.
	 */
	SigmaI();
	/** \brief Deterministic diagonal matrix constructor
	 *
	 * Creates a diagonal matrix with the provided inverse-variance and the degrees of freedom parameter equal to one.  Principally used to create vague large-variance Gaussian priors for location parameters.
	 *
	 * \param[in] size_t& number of rows and columns
	 * \param[in] double& inverse variance
	 */
	SigmaI(const size_t &d, const double &invVar);
	/** \brief Deterministic diagonal matrix constructor
	 *
	 * Creates a diagonal matrix with the provided inverse-variance and degrees of freedom parameter.
	 *
	 * \param[in] size_t& number of rows and columns
	 * \param[in] double& inverse variance
	 * \param[in] double& degrees of freedom
	 */
	SigmaI(const size_t &d, const double &invVar, const double &df);
	/** \brief Deterministic diagonal matrix constructor with output file name
	 *
	 * Creates a diagonal matrix with the provided inverse-variance and degrees of freedom parameter.
	 *
	 * \param[in] size_t& number of rows and columns
	 * \param[in] double& inverse variance
	 * \param[in] double& degrees of freedom
	 * \param[in] string& output file name
	 */
	SigmaI(const size_t &d, const double &invVar, const double &df, const string &outFlNam);
	/** \brief Deterministic matrix constructor
	 *
	 * Used to initialize the inverse-covariance with a given matrix deterministically, i.e. the initial values are not sampled. Only the lower-triangular part of the matrix is used, so if it is not symmetric no error is produced.
	 *
	 * \param[in] gsl_matrix* value matrix
	 */
	SigmaI(const gsl_matrix *mat);
	
	/** \brief Constructor with a matrix prior
	 *
	 * Initializes the matrix with a Wishart sample with the provided matrix as data and another matrix as a prior.  
	 *
	 * \param[in] gsl_matrix* data matrix
	 * \param[in] size_t& dimension parameter
	 * \param[in] size_t& Wishart sample degrees of freedom
	 * \param[in] gsl_matrix* prior matrix
	 * \param[in] double& prior degrees of freedom
	 */
	SigmaI(const gsl_matrix *S, const size_t &d, const size_t &df, const gsl_matrix *LamPr, const double &nu0);
	/** \brief Constructor with a diagonal prior
	 *
	 * Initializes the matrix with a Wishart sample with the provided matrix as data and a diagonal prior with all elements set to the provided value.
	 *
	 * \param[in] gsl_matrix* data matrix
	 * \param[in] size_t& dimension parameter
	 * \param[in] size_t& Wishart sample degrees of freedom
	 * \param[in] double* prior inverse-variance
	 * \param[in] double& prior degrees of freedom
	 */
	SigmaI(const gsl_matrix *S, const size_t &d, const size_t &df, const double &diagPr, const double &nu0);
	
	/** \brief Location data-based constructor
	 *
	 * Initializes the object with an inverse-covariance of the provided location data.  The dimension of the object os equal to the number of columns in the data.
	 *
	 * \param[in] Grp& data
	 * \param[in] double& prior inverse-variance
	 * \param[in] double& prior degrees of freedom
	 */
	SigmaI(const Grp &dat, const double &prDiag, const double &nu0);
	/** \brief Location data-based constructor with output file name
	 *
	 * Initializes the object with an inverse-covariance of the provided location data.  The dimension of the object os equal to the number of columns in the data.
	 *
	 * \param[in] Grp& data
	 * \param[in] string& output file name
	 * \param[in] double& prior inverse-variance
	 * \param[in] double& prior degrees of freedom
	 */
	SigmaI(const Grp &dat, const string &outFlNam, const double &prDiag, const double &nu0);
	
	/** \brief Copy constructor
	 *
	 * \param[in] SigmaI& oject to be copied
	 */
	SigmaI(const SigmaI &);
	/** \brief Assignment operator
	 *
	 * \param[in] SigmaI& object to be copied
	 * \return SigmaI& target object
	 */
	SigmaI & operator=(const SigmaI &);
	
	/** \brief Destructor */
	virtual ~SigmaI();
	
	/** \brief Basic Gaussian update
	 *
	 * \ingroup updateFun
	 *
	 * The data are assumed already centered, so the update is based on the simple cross-product of the data.
	 *
	 * \param[in] Grp& data
	 */
	virtual void update(const Grp &dat);
	/** \brief Gaussian update with a mean
	 *
	 * \ingroup updateFun
	 *
	 * The mean value is subtracted from the data according to the up-pointing RanIndex in the data object.
	 *
	 * \param[in] Grp& data
	 * \param[in] Grp& mean
	 */
	virtual void update(const Grp &dat, const Grp &mu);
	/** \brief Basic Student-\f$t\f$ update
	 *
	 * \ingroup updateFun
	 *
	 * The data are assumed already centered, so the update is based on the simple cross-product of the data and the current value of the scale parameter.
	 *
	 * \param[in] Grp& data
	 * \param[in] Qgrp& scale parameter
	 */
	virtual void update(const Grp &dat, const Qgrp &q);
	/** \brief Student-\f$t\f$ update with a mean
	 *
	 * \ingroup updateFun
	 *
	 * The mean value is subtracted from the data according to the up-pointing RanIndex in the data object.
	 *
	 * \param[in] Grp& data
	 * \param[in] Grp& mean
	 * \param[in] Qgrp& scale parameter
	 */
	virtual void update(const Grp &dat, const Grp &mu, const Qgrp &q);
	
	/** \brief Save to a stored file name
	 *
	 * \param[in] char* saving mode, appending by default
	 */
	virtual void save(const char *how = "a");
	/** \brief Save to a given file name
	 *
	 * \param[in] string& output file name
	 * \param[in] char* saving mode, appending by default
	 */
	virtual void save(const string &fileNam, const char *how = "a");
	
	/** \brief Save adjusted matrix
	 *
	 * Saving the adjusted matrix for the multiplicative parameter expansion models.
	 *
	 * \param[in] string& output file name
	 * \param[in] Apex& redundant parameter matrix
	 * \param[in] char* saving mode, appending by default
	 */
	void save(const string &fileNam, const Apex &A, const char *how = "a");
	/** \brief Save to file stream
	 *
	 * \param[in] FILE* file stream name
	 */
	void save(FILE *fileStr) {gsl_matrix_fwrite(fileStr, _mat); };
	/** \brief Access the output file
	 *
	 * \return string file name
	 */
	string getOutFile() const {return _outFlNam; };
	
	/** \brief Access to the inverse-covariance matrix
	 *
	 * \return gsl_matrix* pointer to the inverse-covariance matrix
	 */
	const gsl_matrix *getMat() const{return _mat; };
	
	/** \brief Update square-root of the determinant */
	void   srDetUpdate();
	/** \brief Access the square-root of the determinant
	 *
	 * \return double square root of the determinant of the inverse-covariance
	 */
	double getSrDet() const {return _srDet; };
	
};

/** \brief Block-diagonal inverse-covariance
 *
 * Implements block-diagonal inverse-covariance, to go with MuBlk and BetaBlk classes. All covariances among blocks are set to exactly zero and are never modified. All sampling is done block by block.
 */
class SigmaIblk : public SigmaI {
protected:
	/** \brief Block start indexes
	 *
	 * Vector of indexes that indicate the column that starts each block.
	 */
	vector< size_t > _blkStart;
	/** \brief Inverse-covariance matrix submatrices
	 *
	 * Vector of submatrices of the current sample of the inverse-covariance matrix (_mat).  Is the same length as the number of blocks.
	 */
	vector< gsl_matrix_view > _eachBlk;
	/** \brief Prior matrix submatrices
	 *
	 * Vector of submatrices of the prior inverse-covariance matrix (_LamSc).  Is the same length as the number of blocks.
	 */
	vector< gsl_matrix_view > _eachLS;   // matrix-view submatrices of the prior
	
public:
	/** \brief Default constructor */
	SigmaIblk();
	/** \brief Deterministic diagonal matrix constructor
	 *
	 * Creates a diagonal matrix with the provided inverse-variance and degrees of freedom parameter.
	 *
	 * \param[in] size_t& number of rows and columns
	 * \param[in] double& inverse variance
	 * \param[in] double& degrees of freedom
	 * \param[in] string& name of the file storing block indexes
	 */
	SigmaIblk(const size_t &d, const double &invVar, const double &df, const string &blkIndFileNam);
	/** \brief Deterministic diagonal matrix constructor with output file
	 *
	 * Creates a diagonal matrix with the provided inverse-variance and degrees of freedom parameter.
	 *
	 * \param[in] size_t& number of rows and columns
	 * \param[in] double& inverse variance
	 * \param[in] double& degrees of freedom
	 * \param[in] string& name of the file storing block indexes
	 * \param[in] string& output file name
	 */
	SigmaIblk(const size_t &d, const double &invVar, const double &df, const string &blkIndFileNam, const string &outFlNam);
	/** \brief Location data-based constructor
	 *
	 * Initializes the object with an inverse-covariance of the provided location data.  The dimension of the object os equal to the number of columns in the data.
	 *
	 * \param[in] Grp& data
	 * \param[in] string& name of the file storing block indexes
	 * \param[in] double& prior inverse-variance
	 * \param[in] double& prior degrees of freedom
	 */
	SigmaIblk(const Grp &dat, const string &blkIndFileNam, const double &prDiag, const double &nu0);
	/** \brief Location data-based constructor with output file
	 *
	 * Initializes the object with an inverse-covariance of the provided location data.  The dimension of the object os equal to the number of columns in the data.
	 *
	 * \param[in] Grp& data
	 * \param[in] string& name of the file storing block indexes
	 * \param[in] string& output file name
	 * \param[in] double& prior inverse-variance
	 * \param[in] double& prior degrees of freedom
	 */
	SigmaIblk(const Grp &dat, const string &blkIndFileNam, const string &outFlNam, const double &prDiag, const double &nu0);
	
	/** \brief Destructor */
	~SigmaIblk() {};
	
	// Gaussian only
	void update(const Grp &dat);
	void update(const Grp &dat, const Grp &mu);
	
};

/** \brief PEX inverse-covariance
 *
 * Implementation of the van Dyk and Meng's \cite dyk01 parameter expansion scheme from multivariate Student-\f$t\f$ sampling.  Note that it only works when all the scale values in Qgrp are sampled.  So, for example, when we have missing data with a Student-\f$t\f$ model for the rest (see Qgrp documentation for details), this class should not be used, it will cause divergence and crashing.
 * 
 * \note Not to be confused with location parameter PEX such as MuGrpPEX
 */
class SigmaIpex : public SigmaI {
	
protected:
	/** \brief The PEX \f$ \alpha \f$ parameter */
	double _alpha;
	
public:
	/** \brief Default constructor */
	SigmaIpex() : _alpha(1.0), SigmaI() {};
	/** \brief Deterministic diagonal matrix constructor
	 *
	 * Creates a diagonal matrix with the provided inverse-variance and degrees of freedom parameter.
	 *
	 * \param[in] size_t& number of rows and columns
	 * \param[in] double& inverse variance
	 * \param[in] double& degrees of freedom
	 */
	SigmaIpex(const size_t &d, const double &invVar, const double &df) : _alpha(1.0), SigmaI(d, invVar, df) {};
	/** \brief Deterministic diagonal matrix constructor with output file name
	 *
	 * Creates a diagonal matrix with the provided inverse-variance and degrees of freedom parameter.
	 *
	 * \param[in] size_t& number of rows and columns
	 * \param[in] double& inverse variance
	 * \param[in] double& degrees of freedom
	 * \param[in] string& output file name
	 */
	SigmaIpex(const size_t &d, const double &invVar, const double &df, const string &outFlNam) : _alpha(1.0), SigmaI(d, invVar, df, outFlNam) {};
	/** \brief Location data-based constructor
	 *
	 * Initializes the object with an inverse-covariance of the provided location data.  The dimension of the object os equal to the number of columns in the data.
	 *
	 * \param[in] Grp& data
	 * \param[in] double& prior inverse-variance
	 * \param[in] double& prior degrees of freedom
	 */
	SigmaIpex(const Grp &dat, const double &prDiag, const double &nu0) : _alpha(1.0), SigmaI(dat, prDiag, nu0) {};
	/** \brief Location data-based constructor with output file name
	 *
	 * Initializes the object with an inverse-covariance of the provided location data.  The dimension of the object os equal to the number of columns in the data.
	 *
	 * \param[in] Grp& data
	 * \param[in] string& output file name
	 * \param[in] double& prior inverse-variance
	 * \param[in] double& prior degrees of freedom
	 */
	SigmaIpex(const Grp &dat, const string &outFlNam, const double &prDiag, const double &nu0) : _alpha(1.0), SigmaI(dat, outFlNam, prDiag, nu0) {};
	
	/** \brief Destructor */
	~SigmaIpex() {};
	
	/** \brief Save to a stored file name
	 *
	 * Saves the "adjusted" value, i.e. scaled by the multiplicative parameter.
	 *
	 * \param[in] char* saving mode, appending by default
	 */
	void save(const char *how = "a");
	/** \brief Save to a given file name
	 *
	 * Saves the "adjusted" value, i.e. scaled by the multiplicative parameter.
	 *
	 * \param[in] string& output file name
	 * \param[in] char* saving mode, appending by default
	 */
	void save(const string &fileNam, const char *how = "a");
	
	void update(const Grp &dat, const Qgrp &q);
	void update(const Grp &dat, const Grp &mu, const Qgrp &q);
};
// end of the sigInv group
/** @} */

/** \defgroup qGrp Student-t weight parameters
 *
 * A popular way to build a Gibbs sampler for a Student-\f$t\f$ model \cite dyk01 \cite gelman04  is to treat it like mixture of Gaussians, with the covariance of each data point weighted proportionally to its distabce from the mean.
 * The weights are modeled as \f$\chi^2_{\nu}\f$ scalars, with the degrees of freedom \f$\nu\f$ equal to the Student-\f$t\f$ degrees of freedom.  It is possible to further construct Metropolis updating steps for the degrees of freedom, but in our experience it complicates computation without adding much to inference.
 * We therefore leave degrees of freedom as constant, to be varied manually by the user.  We encourage trying a few degree of freedom values to see how the values affect inference.
 * All constructors for these classes intitialize all weights deterministically to 1.0.  Empirically, this seems the most numerically stable approach, and still results in reasonably well-spread starting points as long the location parameters and inverse-covariances are initiated stochastically.
 *
 * @{
 */
/** \brief Standard Student-\f$t\f$ weights
 *
 * Mostly a standard implementation of the weight parameter sampling for Student-\f$t\f$ modeling.  The only deviation of the standard approach concerns the situation where some values in the data are missing.
 * If we impute values for these elements, the weights become confounded with means.  So the weights are kept strictly equal to 1.0 and not updated.
 *
 * \note The weights are relatively small for values far from the mean, i.e. outliers have small \f$q\f$.
 */
class Qgrp {
protected:
	/** \brief Vector of weights */
	vector<double> _qVec;
	/** \brief Vector of indexes of present data
	 *
	 * Only _qVec elements corresponding to rows of data with no missing values are updated.
	 */
	vector<size_t> _presInd;
	/** \brief Student-\f$t\f$ degrees of freedom */
	double _nu;
	/** \brief Output file name */
	string _outFile;
	
	/** \brief Pseudo-random number generator
	 *
	 * Initialized the same way as Grp type PNGs, with a sum of time and RTDSC.
	 */
	gsl_rng *_r;
	
public:
	/** \brief Default constructor 
	 *
	 * Results in an empty vector of weights.
	 */
	Qgrp() : _nu(3.0), _outFile("notSet") {};
	/** \brief Deterministic size-only constructor
	 *
	 * Creates a vector of weights that all equal to 1.0, and three degrees of freedom (the smallest possible that still gives a destribution with defined mean and variance).
	 *
	 * \param[in] size_t& number of weights
	 */
	Qgrp(const size_t &N);
	/** \brief Deterministic constructor with degrees of freedom
	 *
	 * Creates a vector of weights that all equal to 1.0, and degrees of freedom set to the given value.
	 *
	 * \param[in] size_t& number of weights
	 * \param[in] double& degrees of freedom
	 */
	Qgrp(const size_t &N, const double &nu);
	/** \brief Deterministic constructor with degrees of freedom and missing values
	 *
	 * Creates a vector of weights that all equal to 1.0, and degrees of freedom set to the given value.  The given file is read to determine which rows of data have missing values.
	 *
	 * \param[in] size_t& number of weights
	 * \param[in] double& degrees of freedom
	 * \param[in] string& missing value file name
	 */
	Qgrp(const size_t &N, const double &nu, const string &misVecFlNam);
	/** \brief Deterministic size-only constructor with output file name
	 *
	 * Creates a vector of weights that all equal to 1.0, and three degrees of freedom (the smallest possible that still gives a destribution with defined mean and variance).
	 *
	 * \param[in] size_t& number of weights
	 * \param[in] string& output file name
	 */
	Qgrp(const size_t &N, const string &outFileNam);
	/** \brief Deterministic constructor with degrees of freedom and output file name
	 *
	 * Creates a vector of weights that all equal to 1.0, and degrees of freedom set to the given value.
	 *
	 * \param[in] size_t& number of weights
	 * \param[in] string& output file name
	 * \param[in] double& degrees of freedom
	 */
	Qgrp(const size_t &N, const string &outFileNam, const double &nu);
	/** \brief Deterministic constructor with degrees of freedom, missing values and output file name
	 *
	 * Creates a vector of weights that all equal to 1.0, and degrees of freedom set to the given value.  The given file is read to determine which rows of data have missing values.
	 *
	 * \param[in] size_t& number of weights
	 * \param[in] string& output file name
	 * \param[in] double& degrees of freedom
	 * \param[in] string& missing value file name
	 */
	Qgrp(const size_t &N, const string &outFileNam, const double &nu, const string &misVecFlNam);
	
	/** \brief Destructor */
	virtual ~Qgrp() {};
	
	/** \brief Const subscript operator
	 *
	 * \param[in] size_t& index
	 * \return double weight value
	 */
	double operator[](const size_t i) const{return _qVec[i]; };
	/** \brief Subscript operator
	 *
	 * \param[in] size_t& index
	 * \return double weight value
	 */
	double operator[](const size_t i) {return _qVec[i]; };
	
	/** \brief Const access to \f$\alpha\f$
	 *
	 * Access to the van Dyk and Meng \f$\alpha\f$ multiplicative redundant parameter.  Is equal to 1.0 in this class.
	 *
	 * \return double 1.0
	 */
	virtual double alpha() const {return 1.0; };
	/** \brief Access to \f$\alpha\f$
	 *
	 * Access to the van Dyk and Meng \f$\alpha\f$ multiplicative redundant parameter.  Is equal to 1.0 in this class.
	 *
	 * \return double 1.0
	 */
	virtual double alpha() {return 1.0; };
	
	/** \brief Const length of the weight vector
	 *
	 * \return size_t& vector length
	 */
	size_t size() const {return _qVec.size(); };
	/** \brief Length of the weight vector
	 *
	 * \return size_t& vector length
	 */
	size_t size() {return _qVec.size(); };
	
	/** \brief Save to a stored file name
	 *
	 * \param[in] char* saving mode, appending by default
	 */
	void save(const char *how = "a");
	/** \brief Save to a given file name
	 *
	 * \param[in] string& output file name
	 * \param[in] char* saving mode, appending by default
	 */
	void save(const string &fileNam, const char *how = "a");
	
	/** \brief Update with a mean
	 *
	 * \ingroup updateFun
	 *
	 * The mean value is subtracted from the data according to the up-pointing RanIndex in the data object.
	 *
	 * \param[in] Grp& data
	 * \param[in] Grp& mean
	 * \param[in] SigmaI& inverse-covariance
	 */
	virtual void update(const Grp &dat, const Grp &mu, const SigmaI &SigI);
	/** \brief Basic update
	 *
	 * \ingroup updateFun
	 *
	 * The data are assumed already centered, so the update is based on the simple cross-product of the data and the current value of the inverse-covariance.
	 *
	 * \param[in] Grp& data
	 * \param[in] SigmaI& inverse-covariance
	 */
	virtual void update(const Grp &dat, const SigmaI &SigI);
	
};

/** \brief Student-\f$t\f$ weights for PEX
 *
 * Implements the weight updating for van Dyk and Meng's \cite dyk01 PEX scheme.  The multiplicative redundant parameter \f$\alpha\f$ is a member of this class and is updated internally.
 * \f$\alpha\f$ is initialized deterministically to 1.0.
 */
class QgrpPEX : public Qgrp {
protected:
	/** \brief Redundant parameter \f$\alpha\f$ */
	double _alpha;
	
public:
	/** \brief Default constructor */
	QgrpPEX() : _alpha(1.0), Qgrp() {};
	/** \brief Deterministic size-only constructor
	 *
	 * Creates a vector of weights that all equal to 1.0, and three degrees of freedom (the smallest possible that still gives a destribution with defined mean and variance).
	 *
	 * \param[in] size_t& number of weights
	 */
	QgrpPEX(const size_t &N) : _alpha(1.0), Qgrp(N) {};
	/** \brief Deterministic constructor with degrees of freedom
	 *
	 * Creates a vector of weights that all equal to 1.0, and degrees of freedom set to the given value.
	 *
	 * \param[in] size_t& number of weights
	 * \param[in] double& degrees of freedom
	 */
	QgrpPEX(const size_t &N, const double &nu) : _alpha(1.0), Qgrp(N, nu) {};
	/** \brief Deterministic constructor with degrees of freedom and missing values
	 *
	 * Creates a vector of weights that all equal to 1.0, and degrees of freedom set to the given value.  The given file is read to determine which rows of data have missing values.
	 *
	 * \param[in] size_t& number of weights
	 * \param[in] double& degrees of freedom
	 * \param[in] string& missing value file name
	 */
	QgrpPEX(const size_t &N, const double &nu, const string &misVecFlNam) : _alpha(1.0), Qgrp(N, nu, misVecFlNam) {};
	/** \brief Deterministic size-only constructor with output file name
	 *
	 * Creates a vector of weights that all equal to 1.0, and three degrees of freedom (the smallest possible that still gives a destribution with defined mean and variance).
	 *
	 * \param[in] size_t& number of weights
	 * \param[in] string& output file name
	 */
	QgrpPEX(const size_t &N, const string &outFileNam) : _alpha(1.0), Qgrp(N, outFileNam) {};
	/** \brief Deterministic constructor with degrees of freedom and output file name
	 *
	 * Creates a vector of weights that all equal to 1.0, and degrees of freedom set to the given value.
	 *
	 * \param[in] size_t& number of weights
	 * \param[in] string& output file name
	 * \param[in] double& degrees of freedom
	 */
	QgrpPEX(const size_t &N, const string &outFileNam, const double &nu) : _alpha(1.0), Qgrp(N, outFileNam, nu) {};
	/** \brief Deterministic constructor with degrees of freedom, missing values and output file name
	 *
	 * Creates a vector of weights that all equal to 1.0, and degrees of freedom set to the given value.  The given file is read to determine which rows of data have missing values.
	 *
	 * \param[in] size_t& number of weights
	 * \param[in] string& output file name
	 * \param[in] double& degrees of freedom
	 * \param[in] string& missing value file name
	 */
	QgrpPEX(const size_t &N, const string &outFileNam, const double &nu, const string &misVecFlNam) : _alpha(1.0), Qgrp(N, outFileNam, nu, misVecFlNam) {};
	
	/** \brief Const access to \f$\alpha\f$
	 *
	 * Access to the van Dyk and Meng \f$\alpha\f$ multiplicative redundant parameter.  Is equal to 1.0 in this class.
	 *
	 * \return double \f$\alpha\f$
	 */
	double alpha() const {return _alpha; };
	/** \brief Access to \f$\alpha\f$
	 *
	 * Access to the van Dyk and Meng \f$\alpha\f$ multiplicative redundant parameter.  Is equal to 1.0 in this class.
	 *
	 * \return double \f$\alpha\f$
	 */
	double alpha() {return _alpha; };
	
	void update(const Grp &dat, const Grp &mu, const SigmaI &SigI);
	void update(const Grp &dat, const SigmaI &SigI);
};
// end of qGrp
/** @} */

/** \brief Dirichlet-multinomial mixture prior
 *
 * Implements the Dirichlet-multinomial prior on proportions in a mixture of Gaussians.
 *
 * \warning This has not been extenisively tested
 */
class MixP {
private:
	/** \brief Vector of mixture proportions */
	vector<double> _p;
	/** \brief Vector of prior sample sizes */
	vector<double> _alpha;
	/** \brief Pseudo-random number generator
	 *
	 * Initialized the same way as Grp type PNGs, with a sum of time and RTDSC.
	 */
	gsl_rng *_r;
	
public:
	/** \brief Default constructor */
	MixP();
	/** \brief Constructor with a vector of proportions
	 *
	 * The prior is set to all ones, i.e. the vaguest possible.
	 *
	 * \param[in] vector<double>& vector of initial proportions
	 */
	MixP(const vector<double> &initP) : _p(initP), _alpha(initP.size(), 1.0) {};
	/** \brief Constructor with a GSL vector of proportions
	 *
	 * The prior is set to all ones, i.e. the vaguest possible.
	 *
	 * \param[in] gsl_vector* vector of initial proportions
	 */
	MixP(const gsl_vector *initP);
	/** \brief Constructor with a vector of proportions and a single prior class size
	 *
	 * The prior for each mixture category is set to the same provided value.
	 *
	 * \param[in] vector<double>& vector of initial proportions
	 * \param[in] double& prior category size
	 */
	MixP(const vector<double> &initP, const double &a);
	/** \brief Constructor with a GSL vector of proportions and a single prior class size
	 *
	 * The prior for each mixture category is set to the same provided value.
	 *
	 * \param[in] gsl_vector* vector of initial proportions
	 * \param[in] double& prior category size
	 */
	MixP(const gsl_vector *initP, const double &a);
	/** \brief Constructor with a vector of proportions and an array of prior class sizes
	 *
	 * The prior for each mixture category is set to the corresponding value in the array.
	 *
	 * \param[in] vector<double>& vector of initial proportions
	 * \param[in] double* prior category size
	 * \param[in] size_t& prior array length
	 */
	MixP(const vector<double> &initP, const double *a, const size_t &len);
	/** \brief Constructor with a GSL vector of proportions and an array of prior class sizes
	 *
	 * The prior for each mixture category is set to the corresponding value in the array.
	 *
	 * \param[in] gsl_vector* vector of initial proportions
	 * \param[in] double* prior category size
	 */
	MixP(const gsl_vector *initP, const double *a);
	/** \brief Constructor with a vector of proportions and a vector of prior class sizes
	 *
	 * The prior for each mixture category is set to the corresponding value in the vector.
	 *
	 * \param[in] vector<double>& vector of initial proportions
	 * \param[in] vector<double>& prior category size
	 */
	MixP(const vector<double> &initP, const vector<double> &a);
	/** \brief Constructor with a GSL vector of proportions and a vector of prior class sizes
	 *
	 * The prior for each mixture category is set to the corresponding value in the vector.
	 *
	 * \param[in] gsl_vector* vector of initial proportions
	 * \param[in] vector<double> prior category size
	 */
	MixP(const gsl_vector *initP, const vector<double> &a);
	/** \brief Constructor with a RanIndex of group IDs and a vector of prior class sizes
	 *
	 * The prior for each mixture category is set to the same provided value.  The proportions are calculated using the data in the RanIndex object.
	 *
	 * \param[in] gsl_vector* vector of initial proportions
	 * \param[in] double& prior category size
	 */
	MixP(const RanIndex &ind, const double &a);
	
	/** \brief Destructor */
	~MixP() {};
	
	/** \brief Copy constructor
	 *
	 * \param[in] MixP& object to be copied
	 */
	MixP(const MixP &);
	/** \brief Assignement operator
	 *
	 * \param[in] Mixp& object to be copied
	 * \return MixP target object
	 */
	MixP & operator=(const MixP &);
	/** \brief Subscript operator
	 *
	 * \param[in] size_t& index of the category
	 * \return double& proprtion of the elements in the given category
	 */
	const double &operator[](const size_t i) const{return _p[i]; };
	
	/** \brief Gibbs update function
	 *
	 * \ingroup updateFun
	 *
	 * Performs standard Gibbs mixture updating (e.g., \cite gelman04 ) with samples from a Dirichlet distribution.
	 *
	 * \param[in] RanIndex& data index of which elements belong to which category
	 */
	void update(const RanIndex &Nvec);
	
};


#endif
```
