## Supplemental methods and scripts for "Low additive genetic variation in a trait under selection in domesticated rice": README.md

This directory contains the C++ source code for the Gibbs sampler programs we used to analyze phenotypic data. The _GSLRIO.cpp_ file is necessary to read and write the binary data format that these programs read and write. To compile it, run

	R CMD SHLIB GSLRIO.cpp -lgsl -O3 -DHAVE_INLINE -DGSL_RANGE_CHECK_OFF

and move the resulting _GSLRIO.so_ object file to the directory where the R scripts included in the *R_scripts* directory are to be run.

The two analysis programs included are _altolMV.cpp_, _altolMVnew.cpp_ and _altolMVtrj.cpp_. They depend on [BLAS](http://www.netlib.org/blas/) and the [GNU Scientific library](http://www.gnu.org/software/gsl/), as well as the [MuGen](https://github.com/tonymugen/MuGen) library included in this directory. To compile, run

	g++ altolMV.cpp MuGen.cpp -o altolMV -lz -lgslcblas -lgsl -lpthread -lm -fopenmp \\
		-DHAVE_INLINE -DGSL_RANGE_CHECK_OFF -O3 -march=native

To compile _altolMVnew.cpp_ and _altolMVtrj.cpp_, substitute them in the above.

The following flags are needed to run these programs:

	./altolMV -b <number_of_burnin_iterations> -s <number_of_sampling_iterations> -t <thinning_interval> -T <number_of_threads> \\
		-g <degrees_of_freedom_for_PCs> -p <population_ID[trj/trja/aus]> -c <chain_ID>

	./altolMVtrj -b <number_of_burnin_iterations> -s <number_of_sampling_iterations> -t <thinning_interval> -T <number_of_threads> \\
		-g <degrees_of_freedom_for_PCs> -c <chain_ID>

	./altolMVnew -b <number_of_burnin_iterations> -s <number_of_sampling_iterations> -t <thinning_interval> -T <number_of_threads> \\
		-e <degrees_of_freedom_for_errors> -g <degrees_of_freedom_for_PCs> -c <chain_ID> \\
		-d <data_set[expt/ctrl] -r <relationship_matrix[IBS/IBSS]
