## Supplemental methods and scripts for "Low additive genetic variation in a trait under selection in domesticated rice": dataPrepADtrj.pdf

### Preparing TRJ Al tolerance data for analysis

Tony Greenberg

March 24, 2017

```
               -
platform       x86_64-apple-darwin16.0.0
version.string R version 3.3.1 (2016-06-21)
```

In this document I prepare the new data set that includes additional measurements on the tropical *japonica* accessions. Start by reading the phenotype data. The file has been modified from the original Excel spreadsheet to add ".0" to two accession names that did not have it.

#### 1 Phenotype data

```
> rawData <- read.table(file = "rawData.tsv", sep = "\t", header = T)
> summary(rawData)
```

```
       assay_id      accession_id                Original_sample_ID   day_3_TRL
 08de34ee.0: 201   NSFTV174: 201   10_Tub1A_0uM_01.RR2Dat:   2     Min.   :  1.78
 45d3c920.0:  44   NSFTV397:  44   10_Tub1A_0uM_02.RR2Dat:   2     1st Qu.: 44.59
 06334433.0:  39   NSFTV251:  39   10_Tub1A_0uM_03.RR2Dat:   2     Median : 66.88
 f25ad349.0:  39   NSFTV54 :  39   10_Tub1A_0uM_04.RR2Dat:   2     Mean   : 69.90
 68c2ecf8.0:  38   NSFTV139:  38   10_Tub1A_0uM_05.RR2Dat:   2     3rd Qu.: 92.14
 (Other)   :3739   (Other) :1767   10_Tub1A_0uM_06.RR2Dat:   2     Max.   :211.91
 NA's      :  90   NA's    :2062   (Other)               :4178     NA's   :1
         Experiment
 2017-January :1438
 2016-December: 750
 2010-panelB  : 580
 2010-panelA  : 570
 2008-screen3 : 200
 2008-screen2 : 172
 (Other)      : 480
```

There are a few accessions that do not have IDs. There are not genotyped, so I am eliminating them.

```
> dim(rawData)
```

```
[1] 4190    5
```

```
> rawData <- rawData[!is.na(rawData$assay_id),]
> dim(rawData)
```

```
[1] 4100    5
```

The control individuals are marked by "0uM" in the `Original_sample_ID` column and the treated plants are "160uM." I am making them into two separate traits. Here is the function that does that within each accession.

```
> buildY <- cmpfun(function(accNam){
+     accMat <- rawData[rawData$assay_id == accNam,]
+     ctrlID <- grep("_0uM_", as.character(accMat$Original_sample_ID))
+     ctrl   <- accMat$day_3_TRL[ctrlID]
+     cExp   <- as.character(accMat$Experiment)[ctrlID]
+
+     trtID  <- grep("_160uM_", as.character(accMat$Original_sample_ID))
+     trt    <- accMat$day_3_TRL[trtID]
+     tExp   <- as.character(accMat$Experiment)[trtID]
+     res    <- matrix(NA, nrow = max(length(trt), length(ctrl)), ncol = 2)
+     res[1:length(ctrl),1] <- ctrl
+     res[1:length(trt),2]  <- trt
+
+     acc  <- rep(accNam, nrow(res))
+     if(length(cExp) >= length(tExp)) {
+         expr <- cExp
+     } else {
+         expr <- tExp
+     }
+
+     Ymat  <<- rbind(Ymat, res)
+     accID <<- c(accID, acc)
+     expID <<- c(expID, expr)
+     return(NULL)
+ })
```

Now run the function to build the trait matrix and the accession and experiment ID factors.

```
> Ymat  <- NULL
> accID <- NULL
> expID <- NULL
> trash <- sapply(levels(rawData$assay_id), buildY)
> colnames(Ymat) <- c("CTRL", "TRT")
> dim(Ymat)
```

```
[1] 2067    2
```

```
> length(unique(accID))
```

```
[1] 169
```

```
> range(apply(Ymat, 1, function(vec){sum(is.na(vec))}))
```

```
[1] 0 1
```

Prepare the missing data matrix and save the phenotype and meta-data.

```
> matNA <- apply(Ymat, 2, function(vec){ifelse(is.na(vec), 1, 0)})
> colSums(matNA)
```

```
CTRL  TRT
  24   11
```

```
> totNA <- rowSums(matNA)
> trash   <- .C("GSLvecSaveInt", "phenoData/matNAind.gbin",
+     as.integer(t(matNA)), as.integer(prod(dim(matNA))))
> trash <- .C("GSLvecSaveInt", "phenoData/totNAind.gbin",
+     as.integer(totNA), length(totNA))
> Ymat[is.na(Ymat)] <- 0.0
> trash <- .C("GSLmatSave", "phenoData/yMat.gbin",
+     as.double(Ymat), nrow(Ymat), ncol(Ymat))
> accID <- factor(accID)
> trash <- .C("GSLvecSaveInt", "phenoData/accInd.gbin",
+     as.integer(accID), length(accID))
> accIDmat <- matrix(levels(accID), nrow = nlevels(accID), ncol = 2)
> cat(apply(accIDmat, 1, paste, collapse = "\t"), file = "accID.tsv", sep = "\n")
```

Save the trait-block index for errors. In our case each of the two “traits” are in their own block so that the errors are not correlated (the data for treated and untreated come from different individuals).

```
> blkIdx <- 1:2
> trash  <- .C("GSLvecSaveInt", "phenoData/trtBlk.gbin",
+     as.integer(blkIdx), length(blkIdx))
```

I will use a regression on contrasts to account for the experiment effects. To pick which level to set as the base, I plot effect distributions.

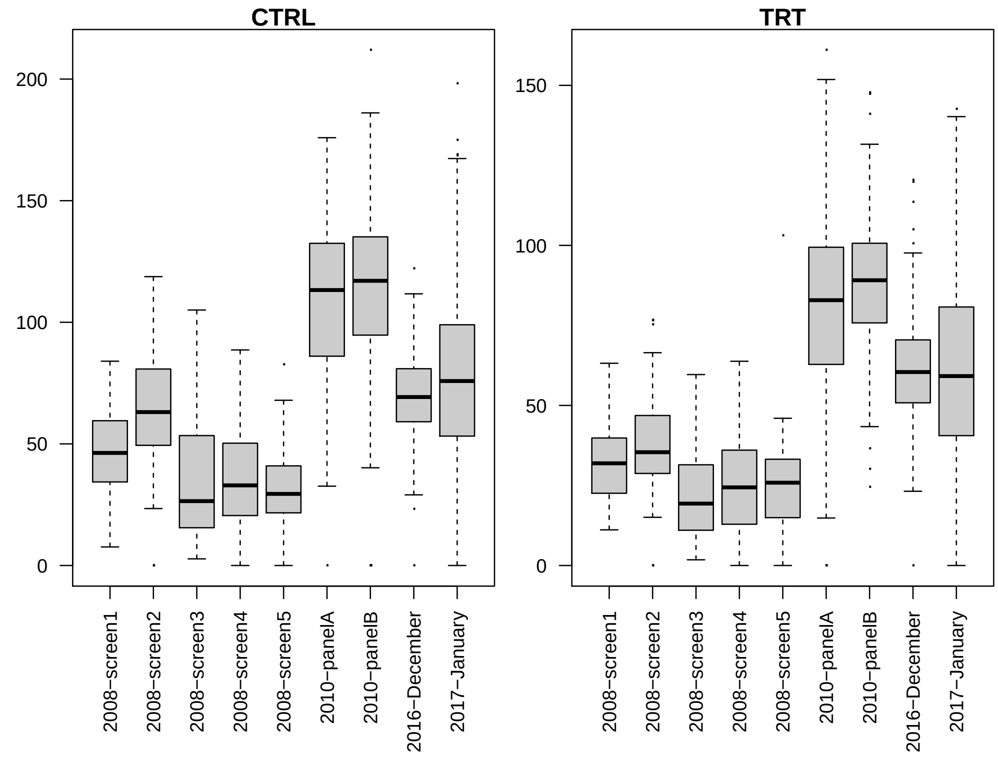


`2016-December` seems to be close to the overall mean, so I will use that. Furthermore, differences within a year are minor compared to among-year divergence. Trying to model each experiment resulted in shrinkage to zero and no correction. I will just estimate year effects, contrasting to 2016/2017 (those were done by the same person using the same reagents and there is no evident difference in mean).

```
> nYr    <- length(unique(sapply(strsplit(expID, "-"), function(vec){vec[1]}))) - 2
> Xexp   <- matrix(0, nrow = nrow(Ymat), ncol = nYr)
> Xexp[grep("2008-", expID),1] <- 1
> Xexp[grep("2010-", expID),2] <- 1
> dim(Xexp)
```

```
[1] 2067    2
```

```
> trash <- .C("GSLmatSave", "phenoData/Xexp.gbin",
+     as.double(Xexp), nrow(Xexp), ncol(Xexp))
```

I also want to construct an experiment predictor that corrects at the level of accessions (i.e. after accession mean estimation). I first test how many accessions were measured in more than one year.

```
> multiExpAcc <- levels(accID)[
+     sapply(
+         tapply(
+             sapply(strsplit(expID, "-"), function(vec){vec[1]}),
```

```
+             accID, function(vec){unique(vec)}),
+         length) > 1]
> multiExpAcc
```

```
[1] "08de34ee.0" "45d3c920.0"
```

There are two accessions. I will keep only the data for the 2016/2017 for these.

```
> dropInd <- which((rowSums(Xexp) == 1) & (accID %in% multiExpAcc))
> YmatS   <- Ymat[-dropInd,]
> accIDs  <- accID[-dropInd]
> matNAs  <- matNA[-dropInd,]
> totNAs  <- totNA[-dropInd]
> trash   <- .C("GSLvecSaveInt", "phenoData/matNAindS.gbin",
+     as.integer(t(matNAs)), as.integer(prod(dim(matNAs))))
> trash <- .C("GSLvecSaveInt", "phenoData/totNAindS.gbin",
+     as.integer(totNAs), length(totNAs))
> trash <- .C("GSLmatSave", "phenoData/yMatS.gbin",
+     as.double(YmatS), nrow(YmatS), ncol(YmatS))
> trash <- .C("GSLvecSaveInt", "phenoData/accIndS.gbin",
+     as.integer(accIDs), length(accIDs))
```

I determine the year ID for each accession and build the contrast matrix. I will include the intercept.

```
> yrID <- tapply(sapply(strsplit(expID[-dropInd], "-"), function(vec){vec[1]}),
+             accIDs, function(vec){unique(vec)[1]})
> Xyr <- cbind(rep(1, nlevels(accIDs)),
+     sapply(c("2008", "2010"), function(yr){yrID == yr}))
> trash <- .C("GSLmatSave", "phenoData/Xyr.gbin",
+     as.double(Xyr), nrow(Xyr), ncol(Xyr))
```

#### 2 Genotype data

Next I prepare genotype data: the SNP table and the relationship matrix eigenvectors and -values. I first run `plink` on the HDRA SNP master file to extract the accessions I need and trim out missing data and singleton SNPs. To do that, the MAF cut-off is $2/N_{acc} = 0.012$.

```
> if(!file.exists("genoData/snpHDRAaltlTRJ.gbin")){
+     plinkComm <- paste(
+             "plink --noweb --silent",
+             "--tfile HDRA-G6-4-RDP1-RDP2-sativa-only",
+             "--keep accID.tsv --maf 0.012 --geno 0.3",
+             "--recode12 --transpose --out alTolTRJ"
+         )
+     system(plinkComm)
+ }
```

Next I read in the genotype file and format the data.

```
> if(!file.exists("genoData/snpHDRAaltlTRJ.gbin")){
+     dimPipe <- pipe("wc -l alTolTRJ.tped")
+     nSNP  <- as.integer(scan(dimPipe, what = character())[1])
+     close(dimPipe)
+     snpFl <- pipe("cut -d ' ' -f5- alTolTRJ.tped")
+     SNPtab <- matrix(scan(snpFl, what = integer()), nrow = nSNP, byrow = T)
+     close(snpFl)
+ }
```

Extract SNP ID, chromosome position and chromosome number from the `.tped` file. This information is in the first four columns. I re-arrange the columns so that I have SNP ID then chromosome number then position. This is done only if the SNP data have not been saved yet.

```
> if(!file.exists("genoData/snpHDRAaltlTRJ.gbin")){
+     snpFl <- pipe("cut -d ' ' -f-4 alTolTRJ.tped")
+     snpID <- matrix(scan(snpFl, what = character()), nrow = nSNP, byrow = T)
+     close(snpFl)
+     system("rm alTolTRJ.*")
+     snpID <- snpID[,c(2, 1, 4)]
+     cat(
+         apply(rbind(c("SNP_ID", "CHROM", "POS"), snpID),
+             1, paste, collapse = "\t"),
+         file = "genoData/snpID.tsv", sep = "\n"
+         )
+     SNPtab[SNPtab == 0] <- NA
+     SNPtab <- SNPtab - 1
+     dim(SNPtab)
+ }
```

Convert the `.tped` two rows per individual to one row per individual.

```
> if(!file.exists("genoData/snpHDRAaltlTRJ.gbin")){
+     SNPtab <- (SNPtab[,seq(from = 1, to = ncol(SNPtab) - 1, by = 2)] +
+         SNPtab[,seq(from = 2, to = ncol(SNPtab), by = 2)])
+     SNPtab <- t(SNPtab)
+     dim(SNPtab)
+ }
```

Calculate the relationship matrix and its eigendecomposition unless that has been done before.

```
> if(!file.exists("genoData/altlTRJevc.gbin") | !file.exists("genoData/altlTRJevl.gbin")){
+     library(rrBLUP)
+     SNPtab <- SNPtab - 1
+     K      <- A.mat(SNPtab, impute.method="EM")
+     Keig   <- eigen(K, symmetric = T)
+     Keig$values
+
+     eVal  <- Keig$val[Keig$val > 1e-8]
```

```
+     eVec  <- Keig$vec[,Keig$val > 1e-8]
+     trash <- .C("GSLvecSave", "genoData/altlTRJevl.gbin",
+         as.double(eVal), length(eVal))
+     trash <- .C("GSLmatSave", "genoData/altlTRJevc.gbin",
+         as.double(eVec), nrow(eVec), ncol(eVec))
+
+     SNPtab[is.na(SNPtab)] <- -9
+     trash <- .C("GSLmatSave", "genoData/snpHDRAaltlTRJ.gbin",
+         as.double(SNPtab), nrow(SNPtab), ncol(SNPtab))
+     dim(SNPtab)
+     dim(eVec)
+ }
```

Print dimensions for future reference.

```
> dim(Ymat)

[1] 2067    2

> dim(YmatS)

[1] 2007    2

> nlevels(accID)

[1] 169

> dim(Xexp)

[1] 2067    2

> dim(Xyr)

[1] 169   3
```
