## Supplemental methods and scripts for "Low additive genetic variation in a trait under selection in domesticated rice": dataPrepNDtrj.pdf

### Preparing and checking raw data

Tony Greenberg

December 22, 2018

```
R version 3.5.1 (2018-07-02)
Platform: x86_64-pc-linux-gnu (64-bit)
Running under: Pop!_OS 18.10

Matrix products: default
BLAS/LAPACK: /opt/intel/compilers_and_libraries_2019.1.144/linux/mkl/lib/intel64_lin/libmkl_gf_lp64.so

locale:
 [1] LC_CTYPE=en_US.UTF-8       LC_NUMERIC=C               LC_TIME=en_US.UTF-8
 [4] LC_COLLATE=en_US.UTF-8     LC_MONETARY=en_US.UTF-8    LC_MESSAGES=en_US.UTF-8
 [7] LC_PAPER=en_US.UTF-8       LC_NAME=C                  LC_ADDRESS=C
[10] LC_TELEPHONE=C             LC_MEASUREMENT=en_US.UTF-8 LC_IDENTIFICATION=C

attached base packages:
[1] compiler  stats     graphics  grDevices utils     datasets  methods   base

other attached packages:
[1] showtext_0.5-1 showtextdb_2.0 sysfonts_0.8   gridExtra_2.3  ggplot2_3.1.0

loaded via a namespace (and not attached):
 [1] Rcpp_1.0.0       crayon_1.3.4     withr_2.1.2      grid_3.5.1       plyr_1.8.4
 [6] gtable_0.2.0     scales_1.0.0     pillar_1.3.0     rlang_0.3.0.1    lazyeval_0.2.1
[11] tools_3.5.1      munsell_0.5.0    colorspace_1.3-2 tibble_1.4.2
```

In this document I check and save raw data from the new Al tolerance experiment. The data files were first modified by replacing commas with tabs, fixing line endings, and inserting `NA` where data point were left blank. I also changed all instances of `196` sample ID to `196_RDP` and added `NA` where a field was blank. I also had to fix some of the HDRA IDs and remove underscores at ends of some accession RPD2 IDs. I begin by reading the data.

#### 1 Phenotype data

```
> ctrl <- read.table("longest+total_control.tsv", header = T, sep = "\t")
> summary(ctrl)
```

```
       ID            RDP2_names           HDRA      Total.Selected.Root.Length
 Min.   :   1.0   118_RDP: 16   5c60aacf.0: 16   Min.   : 6.75
 1st Qu.: 253.8   169_RDP: 16   7c86efa6.0: 16   1st Qu.:15.97
```

```
 Median : 506.5   171_RDP: 16   edc86429.0: 16   Median :18.80
 Mean   : 506.5   150_RDP: 14   f3d9c979.0: 16   Mean   :19.27
 3rd Qu.: 759.2   019_RDP:  8   a51b2a39.0: 14   3rd Qu.:22.59
 Max.   :1012.0   07     :  8   (Other)   :910   Max.   :36.47
                  (Other):934   NA's      : 24   NA's   :38
 Total.Root.Length
 Min.   : 25.37
 1st Qu.: 95.70
 Median :122.73
 Mean   :123.14
 3rd Qu.:149.75
 Max.   :265.13
 NA's   :36
```

```
> dim(ctrl)
```

```
[1] 1012    5
```

```
> expt <- read.table("longest+total_treatment.tsv", header = T, sep = "\t")
> summary(expt)
```

```
       ID          RDP2_names          HDRA      Total.Selected.Root.Length
 Min.   :   1   118_RDP: 16   5c60aacf.0: 16   Min.   : 6.31
 1st Qu.: 254   150_RDP: 16   a51b2a39.0: 16   1st Qu.:13.39
 Median : 507   169_RDP: 16   edc86429.0: 16   Median :16.15
 Mean   : 507   171_RDP: 14   f3d9c979.0: 16   Mean   :16.56
 3rd Qu.: 760   019_RDP:  8   7c86efa6.0: 14   3rd Qu.:19.08
 Max.   :1013   07     :  8   (Other)   :911   Max.   :30.89
                (Other):935   NA's      : 24   NA's   :36
 Total.Root.Length
 Min.   : 26.23
 1st Qu.: 80.15
 Median :101.86
 Mean   :105.12
 3rd Qu.:127.86
 Max.   :218.88
 NA's   :35
```

```
> dim(expt)
```

```
[1] 1013    5
```

I eliminate rows where both traits are missing, or that correspond to ungenotyped accessions.

```
> ctrl <- subset(ctrl, !is.na(HDRA))
> ctrl <- subset(ctrl,
+     (!is.na(Total.Selected.Root.Length)) & (!is.na(Total.Root.Length)))
> summary(ctrl)
```

```
       ID           RDP2_names          HDRA     Total.Selected.Root.Length
 Min.   :   9.0   169_RDP: 16   5c60aacf.0: 16   Min.   : 6.75
 1st Qu.: 265.5   171_RDP: 16   7c86efa6.0: 16   1st Qu.:15.95
 Median : 517.0   118_RDP: 14   f3d9c979.0: 15   Median :18.75
 Mean   : 513.8   150_RDP: 10   edc86429.0: 14   Mean   :19.25
 3rd Qu.: 763.5   019_RDP:  8   a51b2a39.0: 10   3rd Qu.:22.47
 Max.   :1012.0   07     :  8   128dd425.0:  9   Max.   :36.47
                  (Other):879   (Other)   :871
 Total.Root.Length
 Min.   : 33.61
 1st Qu.: 96.27
 Median :122.92
 Mean   :123.67
 3rd Qu.:150.21
 Max.   :265.13
```

```
> dim(ctrl)
```

```
[1] 951   5
```

```
> expt <- subset(expt, !is.na(HDRA))
> expt <- subset(expt,
+     (!is.na(Total.Selected.Root.Length)) & (!is.na(Total.Root.Length)))
> summary(expt)
```

```
       ID           RDP2_names          HDRA     Total.Selected.Root.Length
 Min.   :   1.0   118_RDP: 16   5c60aacf.0: 16   Min.   : 6.31
 1st Qu.: 263.5   150_RDP: 16   a51b2a39.0: 16   1st Qu.:13.30
 Median : 513.5   169_RDP: 16   edc86429.0: 16   Median :16.11
 Mean   : 512.7   171_RDP: 14   7c86efa6.0: 14   Mean   :16.52
 3rd Qu.: 762.8   019_RDP:  8   f3d9c979.0: 14   3rd Qu.:18.98
 Max.   :1013.0   071_RDP:  8   02275e26.0:  8   Max.   :30.89
                  (Other):876   (Other)   :870
 Total.Root.Length
 Min.   : 26.23
 1st Qu.: 80.38
 Median :102.51
 Mean   :105.45
 3rd Qu.:128.43
 Max.   :218.88
```

```
> dim(expt)
```

```
[1] 954   5
```

No more missing data, so the modeling will not involve imputation. To plot the distributions, I first put everything in a single data frame.

```
> alTol <- data.frame(value=c(unlist(ctrl[,4:5]), unlist(expt[,4:5])),
+             RDP2.ID=c(rep(as.character(ctrl[,2]), 2), rep(as.character(expt[,2]), 2)),
+             HDRA.ID=c(rep(as.character(ctrl[,3]), 2), rep(as.character(expt[,3]), 2)),
+             trait=c(rep(c("Selected.Root.Length", "Total.Root.Length"), each=nrow(ctrl)),
+                     rep(c("Selected.Root.Length", "Total.Root.Length"), each=nrow(expt))),
+             treatment=rep(c("control", "treatment"), times = c(2*nrow(ctrl), 2*nrow(expt)))
+             )
```

Plot histograms of the two traits.

```
> showtext_auto()
> ggplot(data=alTol,
+         aes(x=value, ..density.., fill=treatment)) +
+     geom_histogram(bins=100) +
+     facet_wrap(~trait, scale="free") +
+     theme_classic(base_size=18, base_family="myriad")
```

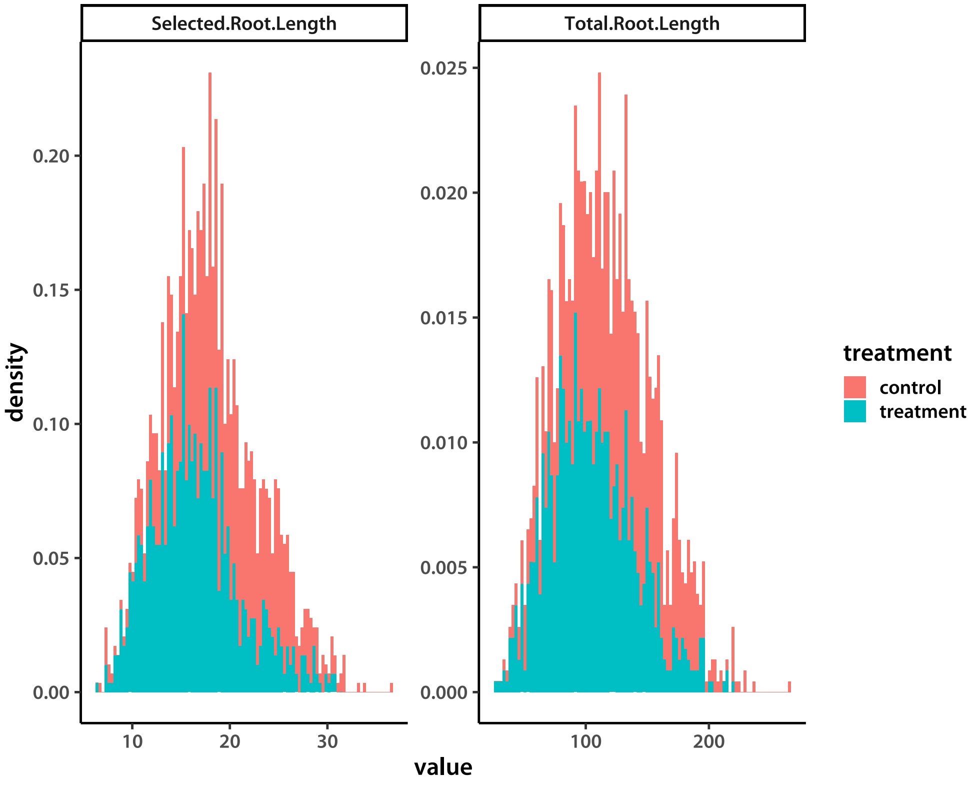


Save the raw data to a file.

```
> write.table(alTol, file="karavolias_RAW_TRJ.tsv", sep = "\t", quote=F, row.names=F)
```

I also look at total between-trait correlations.

```
> signif(cor(ctrl[,4:5])[1,2], 4)
```

```
[1] 0.5288
```

```
> signif(cor(expt[,4:5])[1,2], 4)
```

```
[1] 0.5486
```

Correlations are moderate, so will be good for multi-trait treatment.

I proceed to save the phenotypic data and the single factors that will relate the accessions to individual replicates. The model will be simple, with no experiment-level covariates. I first test if the control and experiment data frames have the same accessions, and if their order is the same.

```
> dim(unique(cbind(levels(ctrl$HDRA), levels(expt$HDRA)), MARGIN = 2))
```

```
[1] 119   1
```

It is. Proceed to make the factors and save.

```
> accID   <- levels(ctrl$HDRA)
> ctrlFac <- as.integer(ctrl$HDRA)
> exptFac <- as.integer(expt$HDRA)
> trash   <- .C("GSLvecSaveInt", "modelData/ctrlAcc.gbin", ctrlFac, length(ctrlFac))
> trash   <- .C("GSLvecSaveInt", "modelData/exptAcc.gbin", exptFac, length(exptFac))
> cat(accID, "\n", file = "accList.tsv", sep = "\t")
```

Now save the phenotype data matrices.

```
> ctrlDat <- as.matrix(ctrl[,4:5])
> trash <- .C("GSLmatSave", "modelData/ctrlData.gbin",
+          as.double(ctrlDat), nrow(ctrlDat), ncol(ctrlDat))
> exptDat <- as.matrix(expt[,4:5])
> trash <- .C("GSLmatSave", "modelData/exptData.gbin",
+          as.double(exptDat), nrow(exptDat), ncol(exptDat))
```

List all relevant dimensions for future reference.

```
> dim(ctrlDat)
```

```
[1] 951   2
```

```
> dim(exptDat)
```

```
[1] 954   2
```

```
> nlevels(ctrl$HDRA)
```

```
[1] 119
```

#### 2 Genotype data

Now I move to extracting genotype data to construct a relationship matrices. First, format the HDRA IDs to be usable by `plink`.

```
> cat(apply(cbind(accID, accID), 1, paste, collapse = "\t"),
+     file = "accPlink.tsv", sep = "\n")
```

Run `plink` to extract the accessions.

```
> plinkComm <- paste(
+     "plink1.9 --silent",
+     "--bfile ../../SNPdata/HDRA-G6-4-RDP1-RDP2-sativa-only",
+     "--keep accPlink.tsv --mac 2 --geno 0.3",
+     "--make-bed --out AlTolGeno"
+ )
> system(plinkComm)
```

Now make the IBS and IBD matrices with `plink`.

```
> system("plink1.9 --silent --bfile AlTolGeno --distance square ibs --out AlTolGeno")
> system("plink1.9 --silent --bfile AlTolGeno --make-rel square ibc2 --out AlTolGeno")
```

I read the relationship matrices and calculate eigenvectors and -values. `plink` does not automatically order individuals according to the list I provide in the `accPlink.tsv` file. I therefore have to re-order the rows and columns of the $K$ matrices.

```
> K     <- matrix(scan(file = "AlTolGeno.mibs", what = double()), ncol = length(accID))
> Knam  <- matrix(scan(file="AlTolGeno.fam", what=character()), ncol=6, byrow=T)[,1]
> rownames(K) <- Knam
> colnames(K) <- Knam
> K     <- K[accID, accID]
> Keig  <- eigen(K)
> signif(Keig$values, 4)
```

```
  [1] 104.30000   1.38400   1.13800   0.58800   0.46620   0.44920   0.40810   0.35210
  [9]   0.29120   0.27210   0.25680   0.24580   0.22260   0.21950   0.20440   0.19990
 [17]   0.19300   0.18810   0.17900   0.17590   0.17010   0.16220   0.15960   0.15900
 [25]   0.15290   0.14910   0.14680   0.13810   0.13630   0.13490   0.12940   0.12870
 [33]   0.12570   0.12300   0.12200   0.11790   0.11690   0.11450   0.11120   0.10990
 [41]   0.10810   0.10490   0.10460   0.10350   0.10140   0.10040   0.09955   0.09677
 [49]   0.09531   0.09442   0.09262   0.09149   0.08965   0.08730   0.08548   0.08447
 [57]   0.08332   0.08123   0.07966   0.07878   0.07720   0.07586   0.07501   0.07229
 [65]   0.07187   0.07091   0.06826   0.06574   0.06544   0.06421   0.06324   0.06188
 [73]   0.06054   0.06006   0.05902   0.05886   0.05654   0.05567   0.05507   0.05397
 [81]   0.05278   0.05202   0.05025   0.05001   0.04911   0.04760   0.04630   0.04568
 [89]   0.04460   0.04330   0.04261   0.04207   0.04095   0.03975   0.03853   0.03823
 [97]   0.03801   0.03586   0.03568   0.03484   0.03337   0.03292   0.03254   0.03168
[105]   0.03125   0.03050   0.02986   0.02956   0.02782   0.02641   0.02579   0.02366
[113]   0.02230   0.02182   0.02018   0.01980   0.01926   0.01677   0.01597
```

```
> eVal  <- Keig$val[Keig$val > 1e-8]
> eVec  <- Keig$vec[,Keig$val > 1e-8]
> trash <- .C("GSLvecSave", "modelData/IBSevl.gbin", as.double(eVal), length(eVal))
> trash <- .C("GSLmatSave", "modelData/IBSevc.gbin",
+     as.double(eVec), nrow(eVec), ncol(eVec))
> dim(eVec)
```

```
[1] 119 119
```

```
> K     <- matrix(scan(file = "AlTolGeno.rel", what = double()), ncol = length(accID))
> rownames(K) <- Knam
> colnames(K) <- Knam
> K     <- K[accID, accID]
> Keig  <- eigen(K)
> signif(Keig$values, 4)
```

```
  [1] 18.9200  7.4740  4.8950  4.4090  4.0020  3.7860  3.3590  3.0860  2.9980  2.9060
 [11]  2.7610  2.6360  2.5520  2.4190  2.3620  2.3410  2.2910  2.2460  2.1880  2.1540
 [21]  2.1390  2.1110  2.0940  2.0710  2.0400  1.9950  1.9750  1.9640  1.9470  1.9350
 [31]  1.9040  1.9040  1.8980  1.8690  1.8670  1.8510  1.8470  1.8360  1.8340  1.8220
 [41]  1.8100  1.7990  1.7930  1.7800  1.7770  1.7710  1.7610  1.7530  1.7490  1.7360
 [51]  1.7350  1.7330  1.7180  1.7110  1.7090  1.7030  1.6970  1.6930  1.6810  1.6790
 [61]  1.6760  1.6700  1.6670  1.6610  1.6540  1.6520  1.6450  1.6350  1.6260  1.6250
 [71]  1.6200  1.6070  1.6030  1.5940  1.5890  1.5810  1.5740  1.5660  1.5630  1.5600
 [81]  1.5570  1.5410  1.5350  1.5210  1.5180  1.5070  1.4960  1.4870  1.4760  1.4680
 [91]  1.4580  1.4480  1.4340  1.4170  1.4100  1.4050  1.3980  1.3650  1.3500  1.3340
[101]  1.3090  1.2900  1.2590  1.2470  1.1960  1.1740  1.1570  1.1240  1.0530  1.0230
[111]  0.9863  0.9260  0.8120  0.7447  0.6761 -0.5490 -2.7760 -3.3190 -3.9550
```

```
> eVal  <- Keig$val[Keig$val > 1e-8]
> eVec  <- Keig$vec[,Keig$val > 1e-8]
> trash <- .C("GSLvecSave", "modelData/IBDevl.gbin", as.double(eVal), length(eVal))
> trash <- .C("GSLmatSave", "modelData/IBDevc.gbin",
+     as.double(eVec), nrow(eVec), ncol(eVec))
> dim(eVec)
```

```
[1] 119 115
```

I also save the eigenvectors and -values of the square of the IBS matrix in an attempt to capture non-additive relationships.

```
> K     <- matrix(scan(file = "AlTolGeno.mibs", what = double()), ncol = length(accID))
> rownames(K) <- Knam
> colnames(K) <- Knam
> K     <- K[accID, accID]
> K     <- K*K
> Keig  <- eigen(K)
> signif(Keig$values, 4)
```

```
  [1] 92.15000  2.28200  1.74500  1.05800  0.81790  0.81160  0.74480  0.59810  0.52960
 [10]  0.49760  0.46880  0.44570  0.41170  0.40250  0.37910  0.36580  0.35490  0.34810
 [19]  0.33210  0.32580  0.31600  0.30320  0.30010  0.29640  0.28630  0.27870  0.27750
 [28]  0.25970  0.25600  0.25340  0.24390  0.24290  0.23790  0.23340  0.23120  0.22360
 [37]  0.22230  0.21780  0.21230  0.20950  0.20720  0.20100  0.19970  0.19800  0.19400
 [46]  0.19350  0.19180  0.18540  0.18330  0.18200  0.17860  0.17630  0.17270  0.16840
 [55]  0.16570  0.16390  0.16090  0.15800  0.15410  0.15340  0.15030  0.14790  0.14600
 [64]  0.14070  0.14000  0.13870  0.13340  0.12870  0.12840  0.12570  0.12440  0.12180
 [73]  0.11880  0.11840  0.11630  0.11560  0.11120  0.10960  0.10860  0.10670  0.10380
 [82]  0.10290  0.09961  0.09926  0.09737  0.09450  0.09197  0.09016  0.08847  0.08592
 [91]  0.08455  0.08367  0.08132  0.07908  0.07686  0.07651  0.07565  0.07174  0.07111
[100]  0.06966  0.06677  0.06561  0.06488  0.06326  0.06218  0.06082  0.05977  0.05920
[109]  0.05588  0.05278  0.05167  0.04749  0.04474  0.04400  0.04065  0.03985  0.03890
[118]  0.03376  0.03222
```

```
> eVal  <- Keig$val[Keig$val > 1e-8]
> eVec  <- Keig$vec[,Keig$val > 1e-8]
> trash <- .C("GSLvecSave", "modelData/IBSSevl.gbin", as.double(eVal), length(eVal))
> trash <- .C("GSLmatSave", "modelData/IBSSevc.gbin",
+     as.double(eVec), nrow(eVec), ncol(eVec))
> dim(eVec)
```

```
[1] 119 119
```
