## Supplemental methods and scripts for "Low additive genetic variation in a trait under selection in domesticated rice": mcmcAnalysisADtrj.pdf

### Results from the extended TRJ panel (Stella + Famoso)

Tony Greenberg

March 4, 2019

```
R version 3.5.2 (2018-12-20)
Platform: x86_64-pc-linux-gnu (64-bit)
Running under: Arch Linux

Matrix products: default
BLAS/LAPACK: /opt/intel/compilers_and_libraries_2019.1.144/linux/mkl/lib/intel64_lin/libmkl_gf_lp64.so

locale:
 [1] LC_CTYPE=en_US.UTF-8       LC_NUMERIC=C               LC_TIME=en_US.UTF-8
 [4] LC_COLLATE=en_US.UTF-8     LC_MONETARY=en_US.UTF-8    LC_MESSAGES=en_US.UTF-8
 [7] LC_PAPER=en_US.UTF-8       LC_NAME=C                  LC_ADDRESS=C
[10] LC_TELEPHONE=C             LC_MEASUREMENT=en_US.UTF-8 LC_IDENTIFICATION=C

attached base packages:
[1] compiler  stats     graphics  grDevices utils     datasets  methods   base

other attached packages:
[1] showtext_0.6   showtextdb_2.0 sysfonts_0.8   gridExtra_2.3  ggplot2_3.1.0

loaded via a namespace (and not attached):
 [1] Rcpp_1.0.0       crayon_1.3.4     withr_2.1.2      grid_3.5.2       plyr_1.8.4
 [6] gtable_0.2.0     scales_1.0.0     pillar_1.3.1     rlang_0.3.1      lazyeval_0.2.1
[11] tools_3.5.2      munsell_0.5.0    pkgconfig_2.0.2  colorspace_1.4-0 tibble_2.0.1
```

In this document I summarize the results of modeling the combined data from Famoso *et al.* and Stella. I start with accession mean generation.

#### 1 Accession means

Accession means were derived using the following hierarchical model.

$$
\begin{aligned}
\boldsymbol{y}_{i\cdot} &\sim \mathrm{N}_d\left(\boldsymbol{\mu}^{acc}_{j[i]\cdot}; \boldsymbol{\Sigma}_e\right) \\
\boldsymbol{\mu}^{acc}_{j\cdot} &\sim \mathrm{N}_d\left(\boldsymbol{\mu} + \boldsymbol{x}_{i\cdot}\boldsymbol{B}^{yr} + \boldsymbol{u}_{j\cdot}\boldsymbol{\Gamma}; \boldsymbol{\Sigma}_s\right) \\
\gamma_{j\cdot} &\sim t_{\nu_g,d}\left(\mathbf{0}_d; \boldsymbol{\Sigma}_a\right) \\
\boldsymbol{\Sigma}_e^{-1} &\sim \mathrm{W}_{d,\nu_0}\left(\left[\sum_i \left(\boldsymbol{y}_{i\cdot} - \boldsymbol{\mu}^{acc}_{j[i]\cdot}\right)\left(\boldsymbol{y}_{i\cdot} - \boldsymbol{\mu}^{acc}_{j[i]\cdot}\right)^{\mathrm{T}}\right]^{-1}\right) \\
\boldsymbol{\Sigma}_s^{-1} &\sim \mathrm{W}_{d,\nu_0}\left(\left[\sum_j \left(\boldsymbol{\mu}^{acc}_{j\cdot} - \boldsymbol{\mu} - \boldsymbol{x}_{i\cdot}\boldsymbol{B}^{yr} - \boldsymbol{u}_{j\cdot}\boldsymbol{\Gamma}\right)\left(\boldsymbol{\mu}^{acc}_{j\cdot} - \boldsymbol{\mu} - \boldsymbol{x}_{i\cdot}\boldsymbol{B}^{yr} - \boldsymbol{u}_{j\cdot}\boldsymbol{\Gamma}\right)^{\mathrm{T}}\right]^{-1}\right) \\
\boldsymbol{\Sigma}_a^{-1} &\sim \mathrm{W}_{d,\nu_0}\left(\left[\boldsymbol{\Gamma}^{\mathrm{T}}\boldsymbol{\Gamma}\right]^{-1}\right),
\end{aligned}
$$

where bold lower-case symbols refer to row vectors and bold upper-case symbols are matrices. All vectors are of length $d$ and all matrices have $d$ columns, where $d$ is the number of traits ($d = 2$ in our case). $\boldsymbol{B}^{yr}$ is the matrix of year effects. There are four years, but the 2016 and 2017 are essentially the same experiment. Therefore, I constructed two contrasts with 2016+2017 set as the base (see `dataPrep` documents). The degrees of freedom parameter $\nu_g$ was set to 1000, leading to essentially a Gaussian model for the genetic component. Notation of the $i\cdot$ type refers to rows in matrices. Subscripts of the $j[i]$ type refer to a row $j$ in a upper level (in the model hierarchy) that corresponds to a replicate row $i$.

I define functions that calculate summaries of Markov chains.

```
> pmode <- cmpfun(function(vec){
+     dst <- density(vec, adjust = 2)
+     mxi <- which(dst$y == max(dst$y))
+     if(length(mxi) > 1){
+         warning("More than one mode in call to pmode(); picking randomly")
+         mxi <- sample(mxi, 1)
+     }
+     dst$x[mxi]
+ })
> HPDint <- cmpfun(function(vec, prob = 0.95){
+     nsamp <- length(vec)
+     if (nsamp <= 2) stop("vector must have length > 2")
+
+     vals <- sort(vec)
+     gap  <- max(1, min(nsamp - 1, round(nsamp * prob)))
+     init <- 1:(nsamp - gap)
+     mInd <- which.min(vals[init + gap] - vals[init])
+     res  <- c(vals[mInd], vals[mInd + gap])
+     names(res) <- c("lower", "upper")
+     return(res)
+ })
```

Next, I write a function that is similar to `quantile()`, but outputs the mode, lower and upper limits of the 95% and 50% HPD. The `outr` parameter is the probability for the outer margins, default is 95%.

```
> quantileLike <- cmpfun(function(vec, outr = 0.95){
+     if(outr <= 0.5) stop("Outer margin has to be >= 50%")
+     md    <- pmode(vec)
+     hpd95 <- HPDint(vec, outr)
+     hpd50 <- HPDint(vec, 0.5)
+     res   <- c(hpd95[1], hpd50[1], md, hpd50[2], hpd95[2])
+
+     outNm <- paste(c("lower", "upper"), outr*100, sep = "")
+     names(res) <- c(outNm[1], "lower50", "mode", "upper50", outNm[2])
+     return(res)
+ })
```

Write a function that will read in specified chains.

```
> addSamp <- cmpfun(function(i, vrNam, ncl){
+     inFlNam <- paste("../chains/", vrNam, "_3_", i, ".gbin", sep = "")
+     chn <<- rbind(chn,
+                 matrix(.C("GSLmatLoad",
+                     inFlNam, as.integer(chnLen), as.integer(ncl),
+                     out = double(chnLen*ncl))$out,
+                 nrow = chnLen, byrow = T)
+         )
+     return(NULL)
+ })
```

Define constants.

```
> trtNam <- c("Control", "Treated")
> d      <- length(trtNam)
> nChn   <- 5
> chnLen <- 2000
```

Read year effect chains to check how well the correction worked.

```
> yrNam  <- c("2016/2017", "2008", "2010")
> Nyr      <- length(yrNam)
> yrDim <- d*Nyr
> chn <- NULL
> trash <- sapply(1:nChn, addSamp, "YR", yrDim)
> yrMn       <- as.data.frame(t(apply(chn, 2, quantileLike)))
> yrMn$trait <- rep(trtNam, times = Nyr)
```

Plot the results.

```
> pdfFlNam <- "yearEffectADD.pdf"
> showtext_auto()
> ggplot(data=yrMn, aes(x=1:nrow(yrMn),y=mode)) +
+     geom_segment(aes(x=1:nrow(yrMn), y=lower95, xend=1:nrow(yrMn), yend=upper95),
+         color="grey80", size=0.75) +
+     geom_segment(aes(x=1:nrow(yrMn), y=lower50, xend=1:nrow(yrMn), yend=upper50),
+         color="grey50", size=1) +
+     geom_point() +
+     facet_wrap(~trait, scales="free_y", nrow=2) +
+     theme_classic(base_size=18, base_family="myriad") +
+     theme(strip.background=element_rect(fill="grey95", linetype="blank")) +
+     #theme(axis.title.x=element_blank(), axis.text.x=element_blank(),
+     #    axis.ticks.x=element_blank()) +
+     scale_x_continuous(name="year", breaks=c(1.5, 3.5, 5.5), labels=yrNam) +
+     labs(y="root length")
> ggsave(pdfFlNam, width=8, height=10, units="in", device="pdf", useDingbats=F)
> cat("\\includegraphics{", pdfFlNam, "}\n\n", sep="")
```

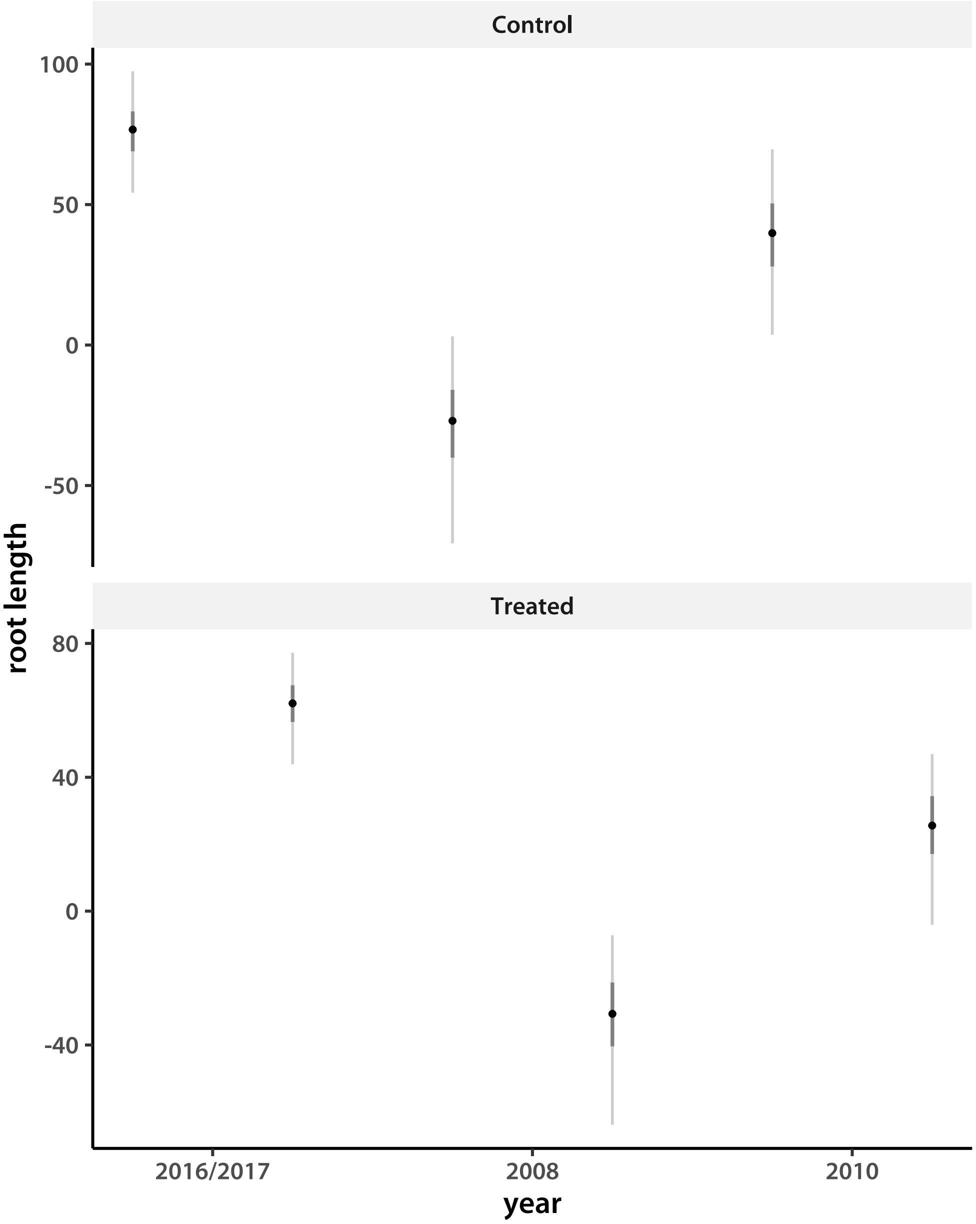


The year effects are in line with expectation from looking at raw data plots.

I move on to read accession mean chains. The accession means corrected for the year effect are in files marked LY.

```
> accNam <- matrix(scan("../accID.tsv", what = character()), ncol = 2, byrow = T)[,1]
> Nacc   <- length(accNam)
> accDim <- d*Nacc
> nChn   <- 5
> chnLen <- 2000
> chn   <- NULL
```

```
> trash <- sapply(1:nChn, addSamp, "LY", accDim)
> accMn       <- as.data.frame(t(apply(chn, 2, quantileLike)))
> accMn$trait <- rep(trtNam, times = Nacc)
> accMnS      <- accMn[order(accMn[,6], accMn[,3]),]
>
```

I plot sorted accession means.

```
> pdfFlNam <- "lineMeansADD.pdf"
> showtext_auto()
> ggplot(data=accMnS, aes(x=1:nrow(accMnS),y=mode)) +
+     geom_segment(aes(x=1:nrow(accMnS), y=lower95, xend=1:nrow(accMnS), yend=upper95),
+         color="grey80", size=0.75) +
+     geom_segment(aes(x=1:nrow(accMnS), y=lower50, xend=1:nrow(accMnS), yend=upper50),
+         color="grey50", size=1) +
+     geom_point() +
+     facet_wrap(~trait, scales="free", nrow=2) +
+     theme_classic(base_size=18, base_family="myriad") +
+     theme(axis.title.x=element_blank(), axis.text.x=element_blank(),
+         strip.background=element_rect(fill="grey95", linetype="blank"),
+         axis.ticks.x=element_blank()) +
+     labs(y="root length")
> ggsave(pdfFlNam, width=8, height=10, units="in", device="pdf", useDingbats=F)
> cat("\\includegraphics{", pdfFlNam, "}\n\n", sep="")
```

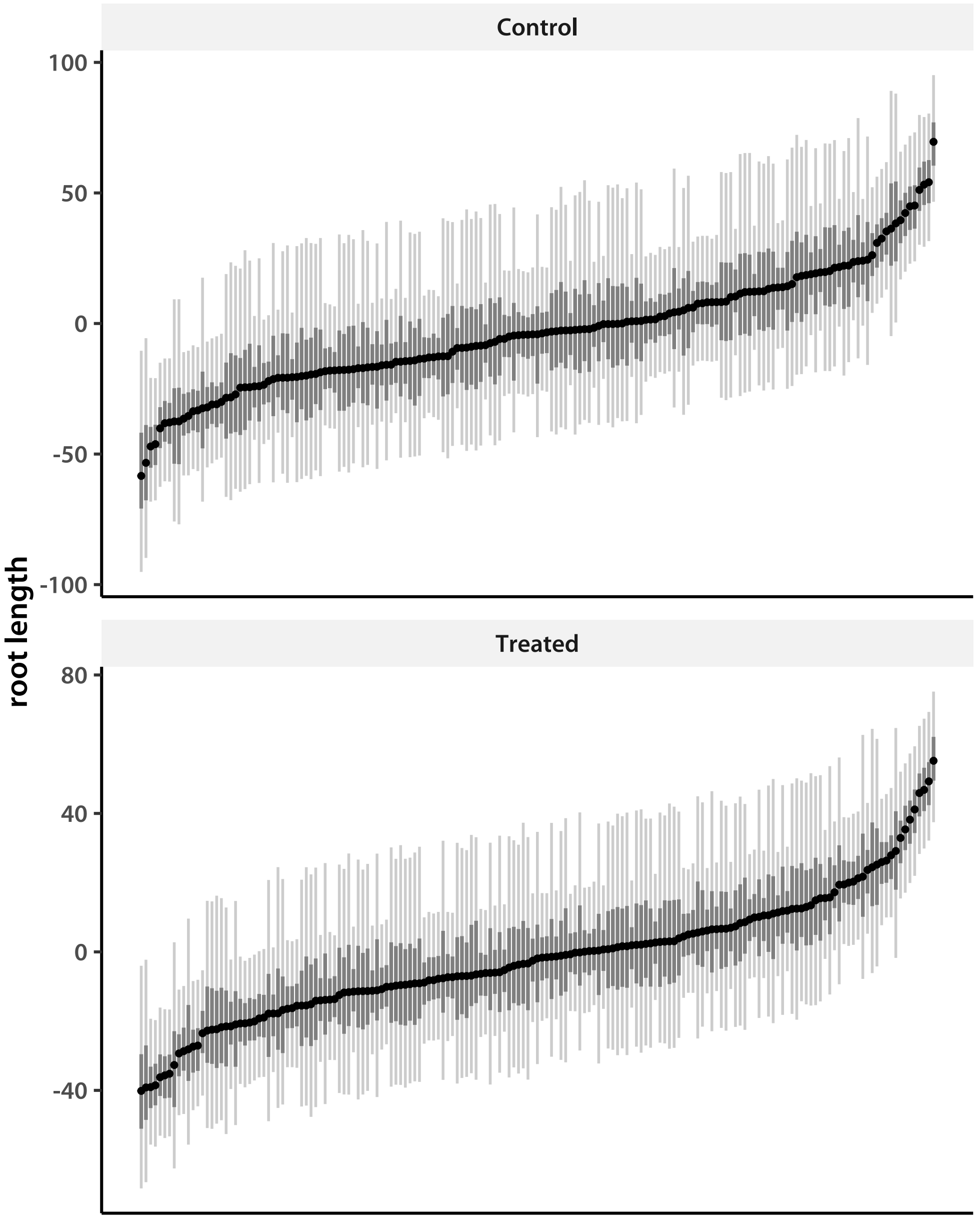


I write a function to check genetic correlations between the two "traits."

```
> trtCor <- cmpfun(function(vec){
+     cor(matrix(vec, ncol=d, byrow=T))[1,2]
+ })
> lnCor <- apply(chn, 1, trtCor)
> round(quantileLike(lnCor), 3)

lower95 lower50    mode upper50 upper95
  0.803   0.858   0.886   0.906   0.945
```

There does not seem to be much interaction between accession ID and treatment (the genetic correlation between treated and control is very high). Next I look at GEBV distributions.

```
> chn     <- NULL
> trash   <- sapply(1:nChn, addSamp, "BV", accDim)
> gebvCHN <- chn
> gebv       <- as.data.frame(t(apply(chn, 2, quantileLike)))
> gebv$trait <- rep(trtNam, times = Nacc)
> gebvS      <- gebv[order(gebv[,6], gebv[,3]),]
```

I plot sorted accession means.

```
> pdfFlNam <- "gebvADD.pdf"
> showtext_auto()
> ggplot(data=gebvS, aes(x=1:nrow(gebvS),y=mode)) +
+     geom_segment(aes(x=1:nrow(gebvS), y=lower95, xend=1:nrow(gebvS), yend=upper95),
+         color="grey80", size=0.75) +
+     geom_segment(aes(x=1:nrow(gebvS), y=lower50, xend=1:nrow(gebvS), yend=upper50),
+         color="grey50", size=1) +
+     geom_point() +
+     facet_wrap(~trait, scales="free", nrow=2) +
+     theme_classic(base_size=18, base_family="myriad") +
+     theme(axis.title.x=element_blank(), axis.text.x=element_blank(),
+         strip.background=element_rect(fill="grey95", linetype="blank"),
+         axis.ticks.x=element_blank()) +
+     labs(y="root length")
> ggsave(pdfFlNam, width=8, height=10, units="in", device="pdf", useDingbats=F)
> cat("\\includegraphics{", pdfFlNam, "}\n\n", sep="")
```

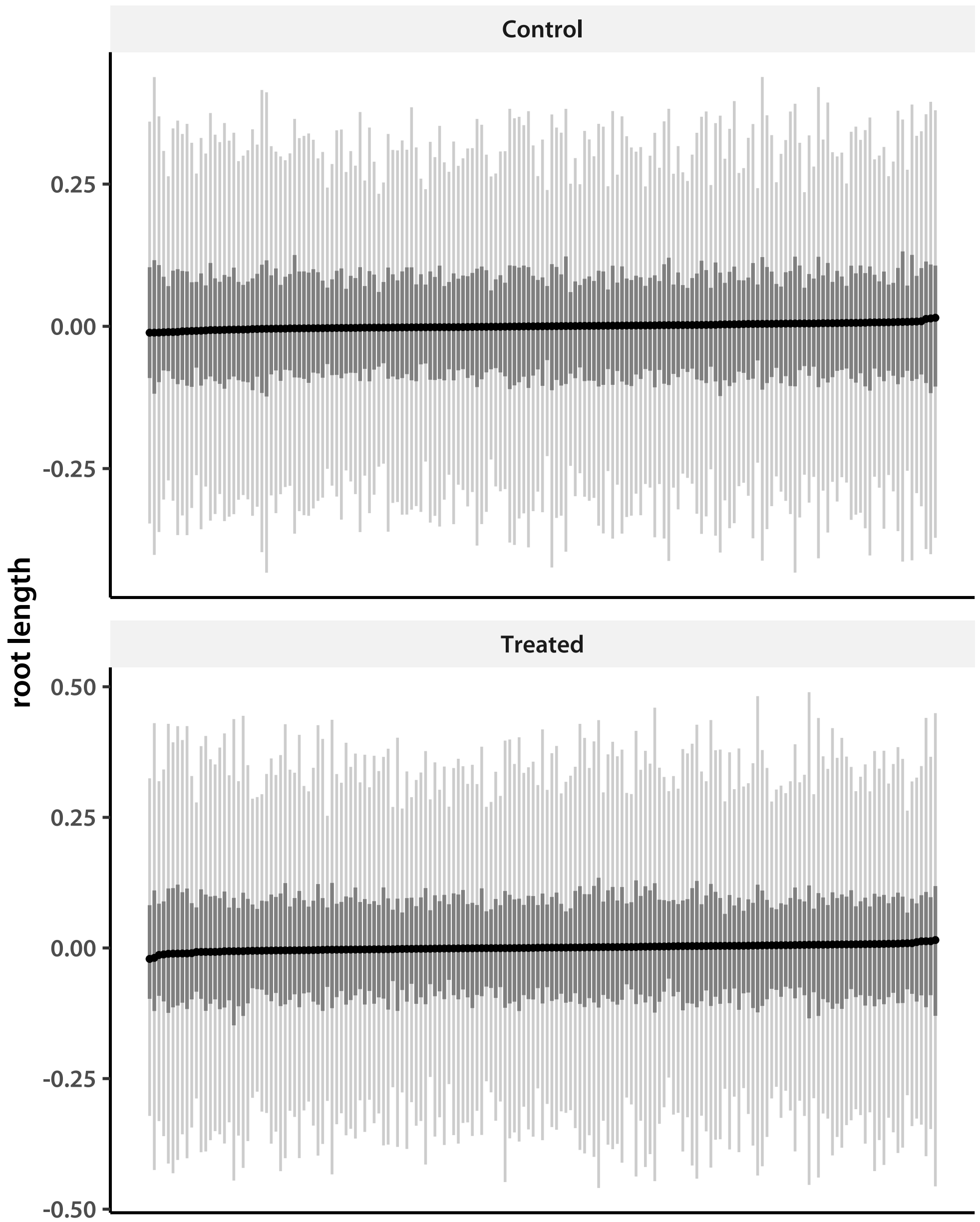


#### 2 Heritability

I next estimate heritability, both broad-sense and marker-based narrow sense. Because I am using a Student-$t$ model for marker effects, I cannot use $\mathbf{\Sigma}_a$ directly, but have to calculate the variances from the GEBV estimates. In addition to broad-sense heritability, I am interested in the separate contribution of background effects. I first define the functions I need.

```
> makeVar <- cmpfun(function(vec){
+     apply(matrix(vec, ncol=d, byrow=T), 2, var)
+ })
```

```
> get.hsq <- cmpfun(function(vec){
+     vec[1:d]/rowSums(matrix(vec, nrow = d))
+ })
> get.Hsq <- cmpfun(function(vec){
+     (vec[1:d] + vec[(d+1):(2*d)])/rowSums(matrix(vec, nrow = d))
+ })
> get.nad <- cmpfun(function(vec){
+     (vec[(d+1):(2*d)])/rowSums(matrix(vec, nrow = d))
+ })
```

The function `get.hsq` calculates the marker heritability, which is a kind of narrow-sense heritability. It is

$$h^2 = \frac{\sigma^2_{\text{GEBV}}}{\sigma^2_{\text{GEBV}} + \sigma^2_s + \sigma^2_e}$$

for each treatment. The `get.Hsq` function calculates the broad-sense (among-accession) heritability:

$$H^2 = \frac{\sigma^2_{\text{GEBV}} + \sigma^2_s}{\sigma^2_{\text{GEBV}} + \sigma^2_s + \sigma^2_e}$$

The `get.nad` function calculates the fraction of total variance contributed by the non-additive effects alone:

$$FVE_s = \frac{\sigma^2_s}{\sigma^2_{\text{GEBV}} + \sigma^2_s + \sigma^2_e}$$

I read in covariance matrix chains.

```
> diagInd <- diag(matrix(1:(d^2), ncol = d, byrow = T))
> chn1 <- matrix(.C("GSLmatLoad",
+     "../chains/SgS_3_1.gbin",
+     as.integer(chnLen), as.integer(d^2), out = double(chnLen*d^2))$out,
+         nrow = chnLen, byrow = T)[,diagInd]
> chn1 <- cbind(chn1,
+     matrix(.C("GSLmatLoad",
+     "../chains/SgE_3_1.gbin",
+     as.integer(chnLen), as.integer(d^2), out = double(chnLen*d^2))$out,
+         nrow = chnLen, byrow = T)[,diagInd])
> chn2 <- matrix(.C("GSLmatLoad",
+     "../chains/SgS_3_2.gbin",
+     as.integer(chnLen), as.integer(d^2), out = double(chnLen*d^2))$out,
+         nrow = chnLen, byrow = T)[,diagInd]
> chn2 <- cbind(chn2,
+     matrix(.C("GSLmatLoad",
+     "../chains/SgE_3_2.gbin",
+     as.integer(chnLen), as.integer(d^2), out = double(chnLen*d^2))$out,
```

```
+         nrow = chnLen, byrow = T)[,diagInd])
> chn3 <- matrix(.C("GSLmatLoad",
+     "../chains/SgS_3_3.gbin",
+     as.integer(chnLen), as.integer(d^2), out = double(chnLen*d^2))$out,
+         nrow = chnLen, byrow = T)[,diagInd]
> chn3 <- cbind(chn3,
+     matrix(.C("GSLmatLoad",
+     "../chains/SgE_3_3.gbin",
+     as.integer(chnLen), as.integer(d^2), out = double(chnLen*d^2))$out,
+         nrow = chnLen, byrow = T)[,diagInd])
> chn4 <- matrix(.C("GSLmatLoad",
+     "../chains/SgS_3_4.gbin",
+     as.integer(chnLen), as.integer(d^2), out = double(chnLen*d^2))$out,
+         nrow = chnLen, byrow = T)[,diagInd]
> chn4 <- cbind(chn4,
+     matrix(.C("GSLmatLoad",
+     "../chains/SgE_3_4.gbin",
+     as.integer(chnLen), as.integer(d^2), out = double(chnLen*d^2))$out,
+         nrow = chnLen, byrow = T)[,diagInd])
> chn5 <- matrix(.C("GSLmatLoad",
+     "../chains/SgS_3_5.gbin",
+     as.integer(chnLen), as.integer(d^2), out = double(chnLen*d^2))$out,
+         nrow = chnLen, byrow = T)[,diagInd]
> chn5 <- cbind(chn5,
+     matrix(.C("GSLmatLoad",
+     "../chains/SgE_3_5.gbin",
+     as.integer(chnLen), as.integer(d^2), out = double(chnLen*d^2))$out,
+         nrow = chnLen, byrow = T)[,diagInd])
> sigChn <- rbind(chn1, chn2, chn3, chn4, chn5)
> sigChn <- cbind(t(apply(gebvCHN, 1, makeVar)), sigChn)
> chnhsq <- t(apply(sigChn, 1, get.hsq))
> chnHsq <- t(apply(sigChn, 1, get.Hsq))
> chnNad <- t(apply(sigChn, 1, get.nad))
> hsqHPD <- as.data.frame(rbind(t(apply(chnhsq, 2, quantileLike)),
+                           t(apply(chnHsq, 2, quantileLike))))
> hsqHPD$treatment    <- rep(trtNam, times=2)
> hsqHPD$heritability <- rep(c("GEBV", "Broad"), each=d)
> fveHPD <- as.data.frame(rbind(t(apply(chnhsq, 2, quantileLike)),
+                           t(apply(chnNad, 2, quantileLike))))
> fveHPD$treatment <- rep(trtNam, times=2)
> fveHPD$variance  <- factor(rep(c("Marker", "Background"), each=d),
+                           levels=c("Marker", "Background"))
```

Now plot both kinds of heritabilities on the same panel.

```
> pdfFlNam <- "heritabilityADD.pdf"
```

```
> showtext_auto()
> ggplot(data=hsqHPD, aes(x=1:nrow(hsqHPD), y=mode, color=heritability)) +
+     geom_segment(aes(x=1:nrow(hsqHPD), y=lower95, xend=1:nrow(hsqHPD),
+         yend=upper95, color=heritability),
+         size=1.2) +
+     geom_segment(aes(x=1:nrow(hsqHPD), y=lower50, xend=1:nrow(hsqHPD),
+         yend=upper50, color=heritability),
+         size=1.75) +
+     geom_point(size=2) +
+     theme_classic(base_size=18, base_family="myriad") +
+     theme(legend.title=element_blank(),
+         strip.background=element_rect(fill="grey95", linetype="blank")) +
+     scale_x_continuous(breaks=1:nrow(hsqHPD), labels=rep(trtNam, 2)) +
+     ylim(c(0,1)) +
+     labs(y="heritability", x="treatment")
> ggsave(pdfFlNam, width=8, height=8, units="in", device="pdf", useDingbats=F)
> cat("\\includegraphics{", pdfFlNam, "}\n\n", sep="")
```

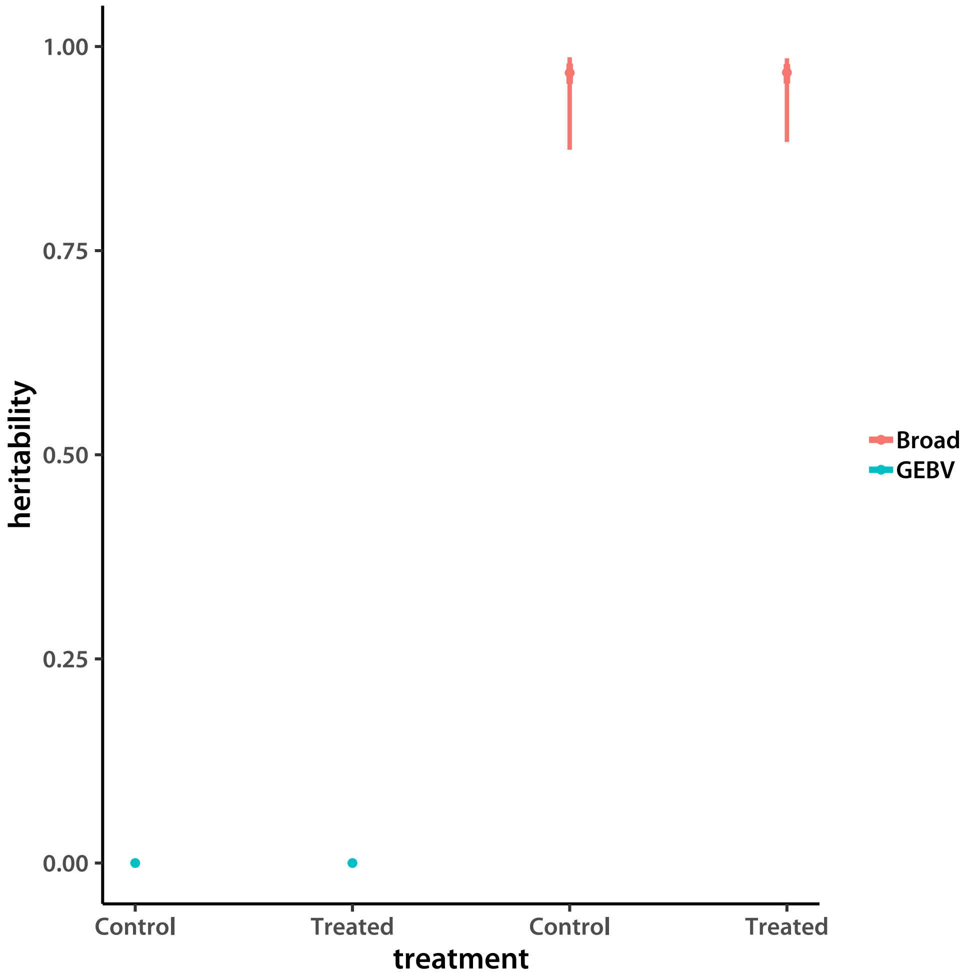

Finally plot fractions of variance explained on the same panel.

```
> pdfFlNam <- "fveADD.pdf"
> showtext_auto()
> ggplot(data=fveHPD, aes(x=1:nrow(fveHPD), y=mode, color=variance)) +
+     geom_segment(aes(x=1:nrow(fveHPD), y=lower95, xend=1:nrow(fveHPD),
+         yend=upper95, color=variance),
+         size=1.2) +
+     geom_segment(aes(x=1:nrow(fveHPD), y=lower50, xend=1:nrow(fveHPD),
+         yend=upper50, color=variance),
+         size=1.75) +
+     geom_point(size=2) +
+     theme_classic(base_size=18, base_family="myriad") +
+     theme(legend.title=element_blank(),
+         strip.background=element_rect(fill="grey95", linetype="blank")) +
+     scale_x_continuous(breaks=1:nrow(fveHPD), labels=rep(trtNam, 2)) +
+     ylim(c(0,1)) +
+     labs(y="fraction of variance", x="treatment")
> ggsave(pdfFlNam, width=8, height=8, units="in", device="pdf", useDingbats=F)
> cat("\\includegraphics{", pdfFlNam, "}\n\n", sep="")
```

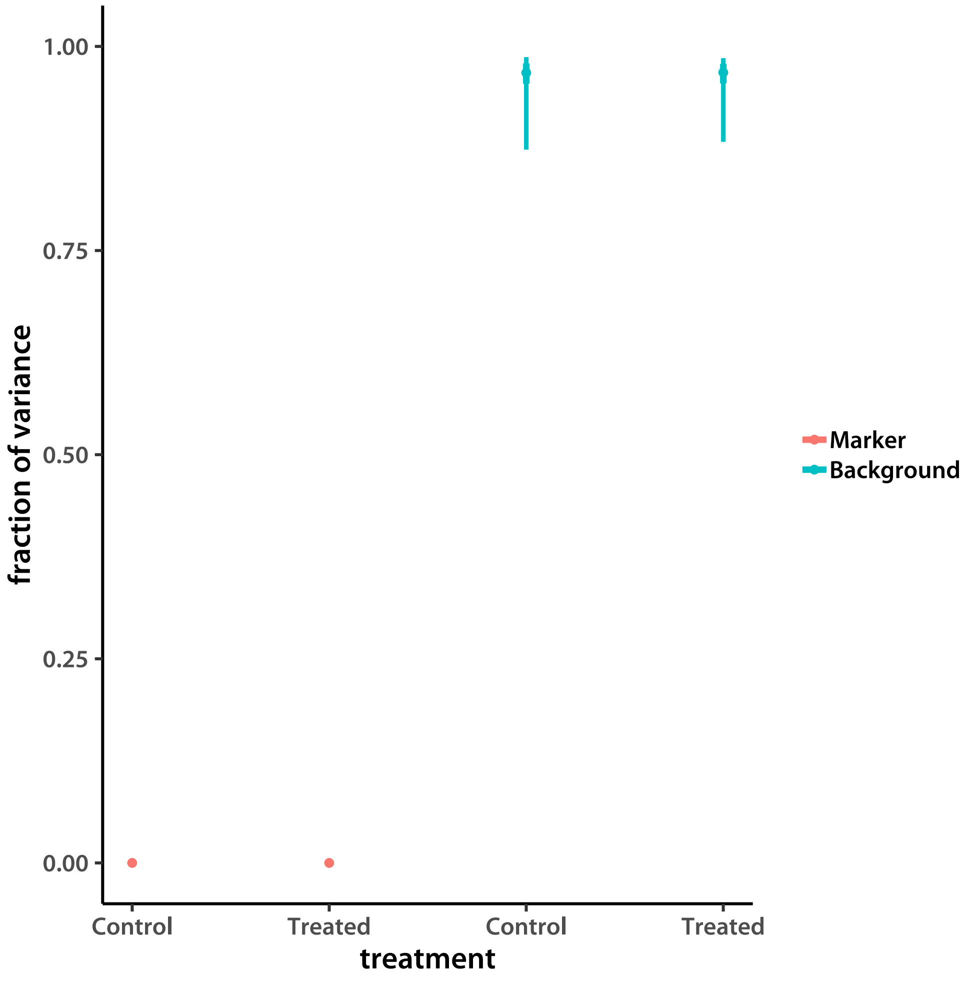
1.00
0.75
0.50
0.25
0.00
fraction of variance
Control
Treated
Control
Treated
treatment
Marker
Background
