## Supplemental methods and scripts for "Low additive genetic variation in a trait under selection in domesticated rice": mcmcExamineADtrj.pdf

### Examining MCMC chains for convergence and mixing

Tony Greenberg

December 10, 2018

```
R version 3.5.1 (2018-07-02)
Platform: x86_64-pc-linux-gnu (64-bit)
Running under: Pop!_OS 18.10

Matrix products: default
BLAS/LAPACK: /opt/intel/compilers_and_libraries_2019.1.144/linux/mkl/lib/intel64_lin/libmkl_gf_lp64.so

locale:
 [1] LC_CTYPE=en_US.UTF-8       LC_NUMERIC=C               LC_TIME=en_US.UTF-8
 [4] LC_COLLATE=en_US.UTF-8     LC_MONETARY=en_US.UTF-8    LC_MESSAGES=en_US.UTF-8
 [7] LC_PAPER=en_US.UTF-8       LC_NAME=C                  LC_ADDRESS=C
[10] LC_TELEPHONE=C             LC_MEASUREMENT=en_US.UTF-8 LC_IDENTIFICATION=C

attached base packages:
[1] compiler  stats     graphics  grDevices utils     datasets  methods   base

loaded via a namespace (and not attached):
[1] tools_3.5.1
```

Examining the Gibbs sampler chains generated for the Al tolerance experiment on TRJ. Starting with the bivariate model that generates accession means for further analyses relative root growth.

Start by defining some useful functions.

I first define a function that calculates the Gelman-Rubin convergence statistic for multiple MCMC chains.

```
> gelRub <- cmpfun(function(vrNam, nChn, chnLen){
+     mat <- NULL
+     for (i in 1:nChn){
+         mat <- rbind(mat, eval(as.symbol(paste(vrNam, i, sep=""))))
+     }
+     chnFac <- factor(rep(1:nChn, each=chnLen))
+     W <- colMeans(apply(mat, 2, tapply, chnFac, var))
+     B <- apply(apply(mat, 2, tapply, chnFac, mean), 2, var)
+     return( sqrt((((chnLen - 1)/chnLen)*W + B)/W) )
+ })
>
```

I am also interested in the quality of chain mixing. I use autocorrelations to test for that. The following function examines autocorrelations for each chain for a given variable and reports the chain with the highest mean value. The autocorrelations are calculated using default parameters of the `acf()` function from the `stats` R package.

```
> eachAcf <- cmpfun(function(i, nChn, vrNam){
+     tmp <- NULL
+     for (j in 1:nChn){
+         tmp <- cbind(tmp,
+                 acf(eval(as.symbol(paste(vrNam, j, sep="")))[,i], plot = F)$acf
+             )
+     }
+     return(tmp[,which.max(colMeans(tmp[-1,]))])
+ })
> mcmcAcf <- cmpfun(function(vrNam, nChn){
+     N <- ncol(eval(as.symbol(paste(vrNam, 1, sep="")))) # all chains should have the same di
+     t(sapply(1:N, eachAcf, nChn, vrNam))
+ })
```

### 1 Raw data modeling

The first task is to model raw data and extract accession means. I ran a 5000 iteration burn-in and 10 000 samples with, keeping every fifth value. I first look at the experiment effect.

```
> trtNam <- c("CTRL", "TRT")
> d      <- 2
> Nyr    <- 3
> yrDim  <- d*Nyr
> nChn   <- 5
> chnLen <- 2000
> chn1 <- matrix(.C("GSLmatLoad", "chains/YR_3_1.gbin",
+     as.integer(chnLen), as.integer(yrDim), out = double(chnLen*yrDim))$out,
+         nrow = chnLen, byrow = T)
> chn2 <- matrix(.C("GSLmatLoad", "chains/YR_3_2.gbin",
+     as.integer(chnLen), as.integer(yrDim), out = double(chnLen*yrDim))$out,
+         nrow = chnLen, byrow = T)
> chn3 <- matrix(.C("GSLmatLoad", "chains/YR_3_3.gbin",
+     as.integer(chnLen), as.integer(yrDim), out = double(chnLen*yrDim))$out,
+         nrow = chnLen, byrow = T)
> chn4 <- matrix(.C("GSLmatLoad", "chains/YR_3_4.gbin",
+     as.integer(chnLen), as.integer(yrDim), out = double(chnLen*yrDim))$out,
+         nrow = chnLen, byrow = T)
> chn5 <- matrix(.C("GSLmatLoad", "chains/YR_3_5.gbin",
+     as.integer(chnLen), as.integer(yrDim), out = double(chnLen*yrDim))$out,
+         nrow = chnLen, byrow = T)
> grMat    <- matrix(gelRub("chn",nChn,chnLen), ncol = d, byrow = T)
> grMaxInd <- apply(grMat, 2, function(vec){which(vec == max(vec))[1]})
```

```
> acMat    <- mcmcAcf("chn", 5)
>
```

Now I plot the Gelman-Rubin for each trait.

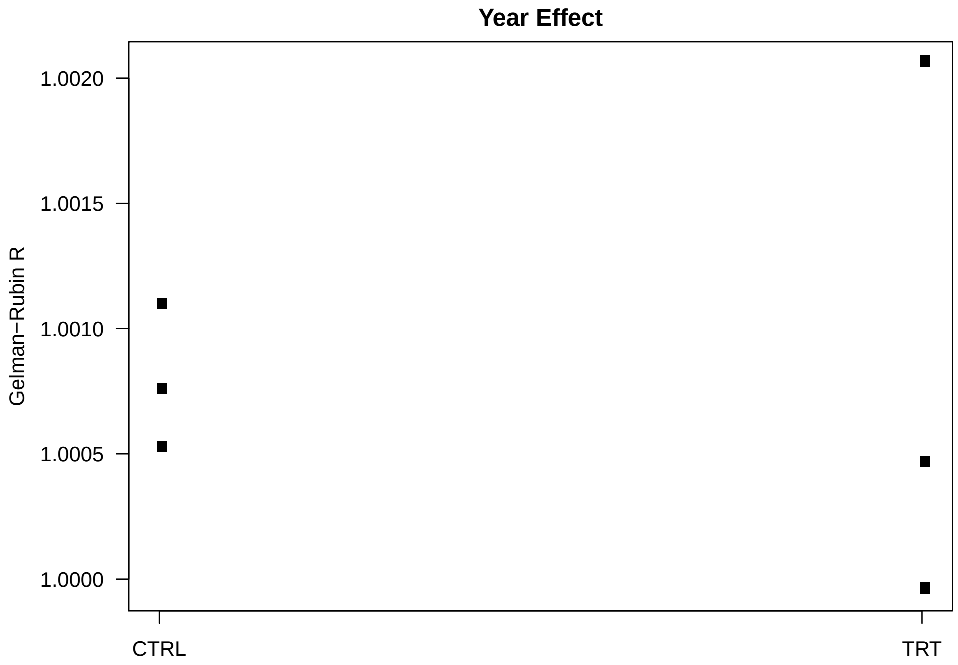


Values smaller than ~1.2 indicate good convergence. I check by plotting an the chains with the worst G-R statistic for each trait.

```
> plotLocEx <- cmpfun(function(trt.i, samp.j, N){
+     i <- ( d*(samp.j-1) ) + trt.i
+     plot( chn1[,i], type = "l",
+         ylim = range(c(chn1[,i], chn2[,i], chn3[,i], chn4[,i], chn5[,i])),
+         col = "orange", ylab = trtNam[trt.i], xaxt = "n", las = 1)
+     lines(chn2[,i], col = "red")
+     lines(chn3[,i], col = "blue")
+     lines(chn4[,i], col = "green")
+     lines(chn5[,i], col = "magenta")
+ })
```

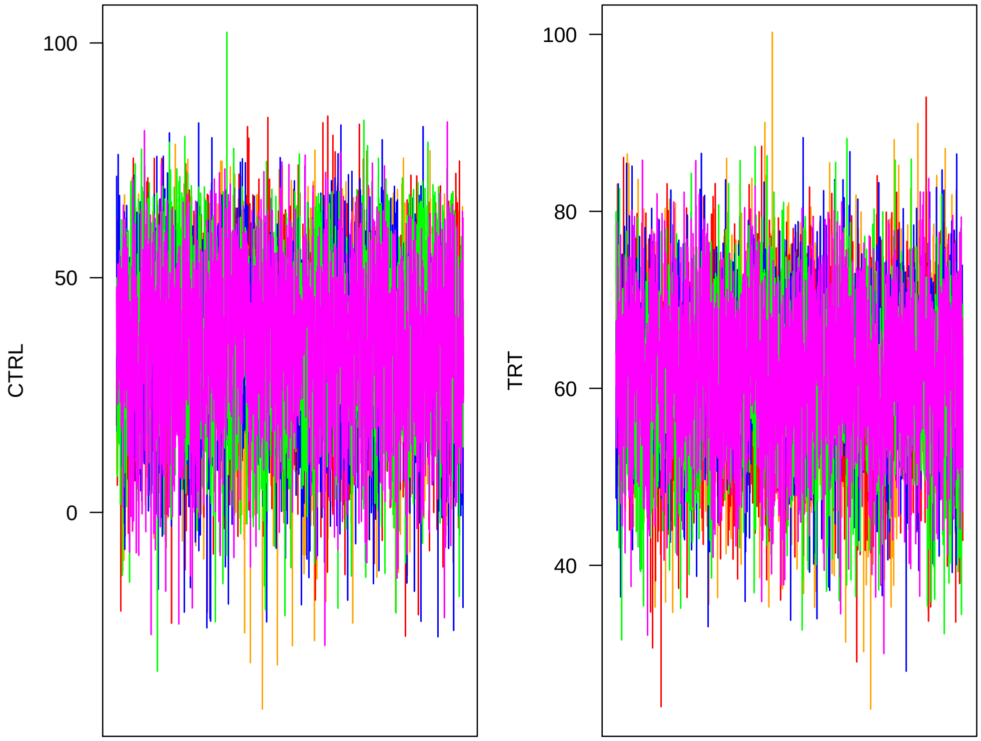


Now I look at mixing. Lag is on the $x$ axis.

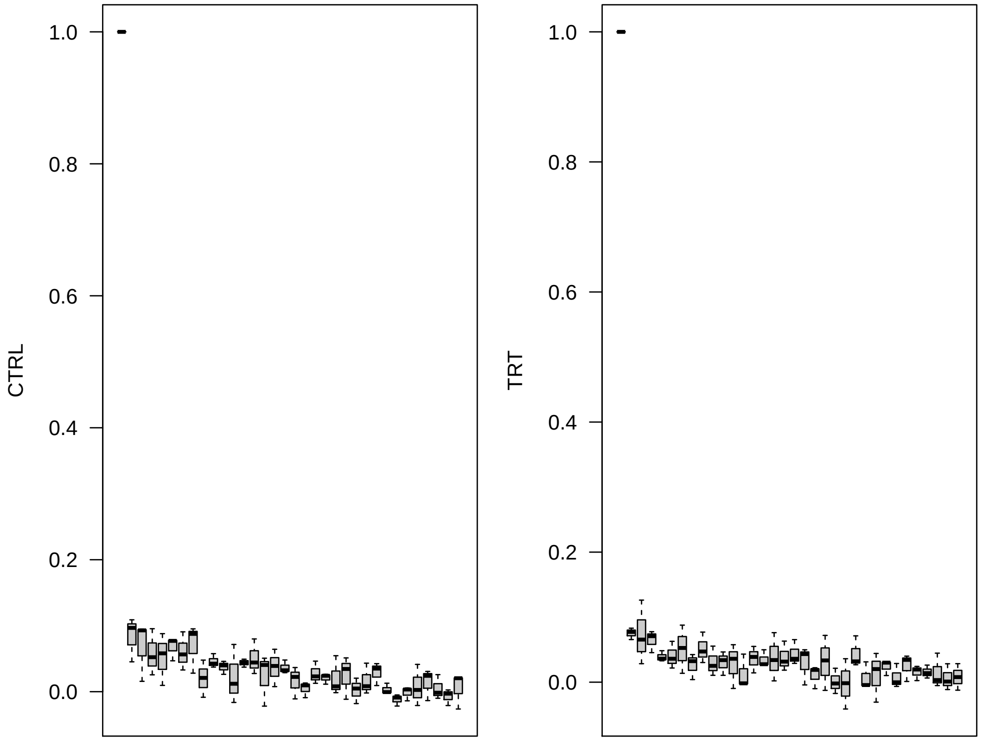


Next I look at accession means.

```
> Nacc   <- 169
> accDim <- d*Nacc
> chn1 <- matrix(.C("GSLmatLoad", "chains/LN_3_1.gbin",
+     as.integer(chnLen), as.integer(accDim), out = double(chnLen*accDim))$out,
+         nrow = chnLen, byrow = T)
> chn2 <- matrix(.C("GSLmatLoad", "chains/LN_3_2.gbin",
+     as.integer(chnLen), as.integer(accDim), out = double(chnLen*accDim))$out,
+         nrow = chnLen, byrow = T)
> chn3 <- matrix(.C("GSLmatLoad", "chains/LN_3_3.gbin",
+     as.integer(chnLen), as.integer(accDim), out = double(chnLen*accDim))$out,
+         nrow = chnLen, byrow = T)
> chn4 <- matrix(.C("GSLmatLoad", "chains/LN_3_4.gbin",
+     as.integer(chnLen), as.integer(accDim), out = double(chnLen*accDim))$out,
+         nrow = chnLen, byrow = T)
> chn5 <- matrix(.C("GSLmatLoad", "chains/LN_3_5.gbin",
+     as.integer(chnLen), as.integer(accDim), out = double(chnLen*accDim))$out,
+         nrow = chnLen, byrow = T)
> grMat    <- matrix(gelRub("chn",nChn,chnLen), ncol = d, byrow = T)
> grMaxInd <- apply(grMat, 2, function(vec){which(vec == max(vec))[1]})
> acMat    <- mcmcAcf("chn", 5)
>
```

Now I plot the Gelman-Rubin for each trait.

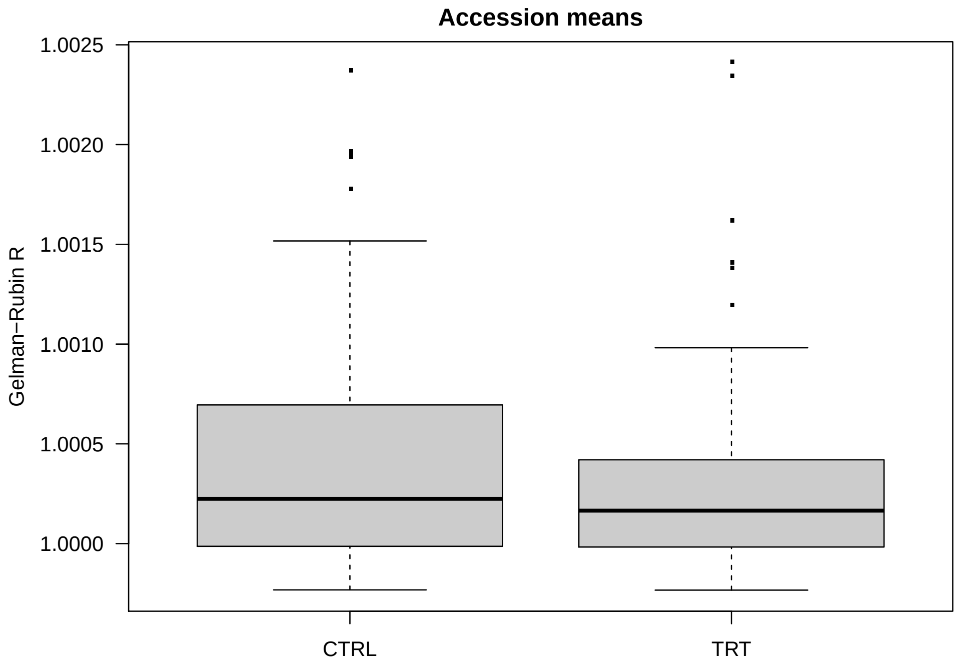


Values smaller than ∼1.2 indicate good convergence. I check by plotting an the chains with the worst G-R statistic for each trait.

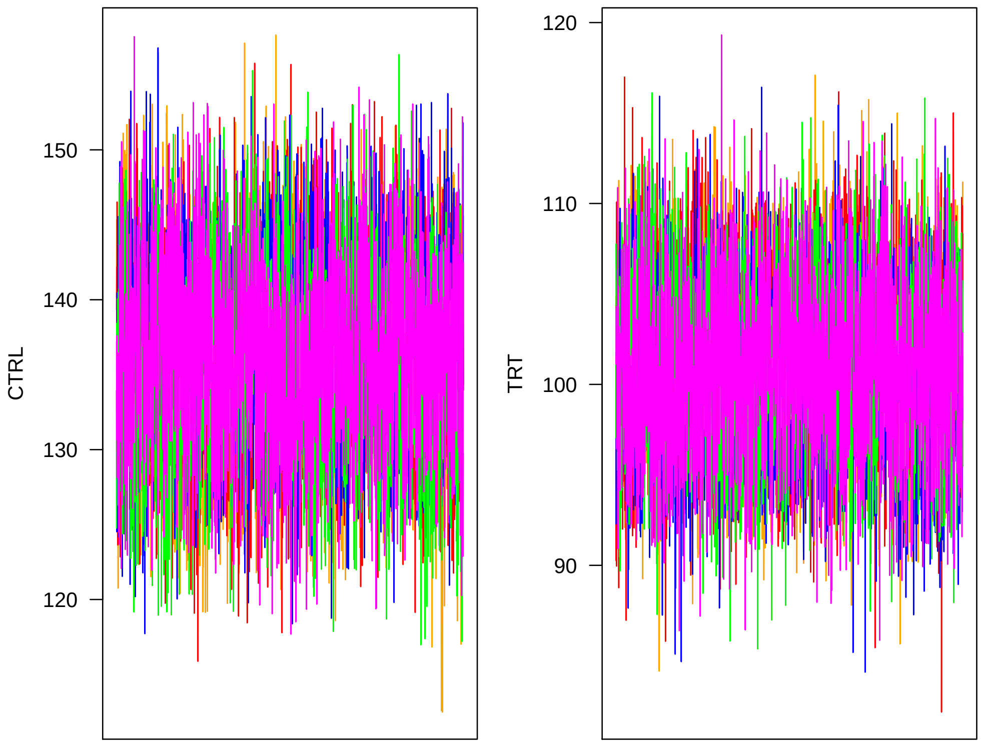


Now I look at mixing. Lag is on the $x$ axis.

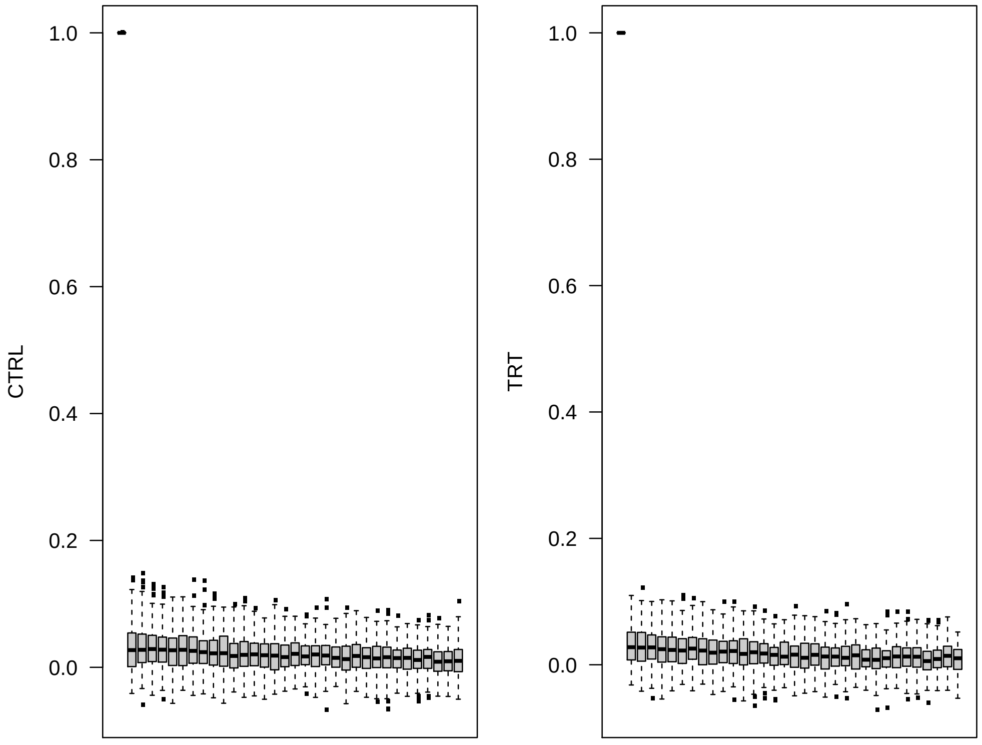

Next I examine the GEBV chains.

```
> chn1 <- matrix(.C("GSLmatLoad", "chains/BV_3_1.gbin",
+     as.integer(chnLen), as.integer(accDim), out = double(chnLen*accDim))$out,
+         nrow = chnLen, byrow = T)
> chn2 <- matrix(.C("GSLmatLoad", "chains/BV_3_2.gbin",
+     as.integer(chnLen), as.integer(accDim), out = double(chnLen*accDim))$out,
+         nrow = chnLen, byrow = T)
> chn3 <- matrix(.C("GSLmatLoad", "chains/BV_3_3.gbin",
+     as.integer(chnLen), as.integer(accDim), out = double(chnLen*accDim))$out,
+         nrow = chnLen, byrow = T)
> chn4 <- matrix(.C("GSLmatLoad", "chains/BV_3_4.gbin",
+     as.integer(chnLen), as.integer(accDim), out = double(chnLen*accDim))$out,
+         nrow = chnLen, byrow = T)
> chn5 <- matrix(.C("GSLmatLoad", "chains/BV_3_5.gbin",
+     as.integer(chnLen), as.integer(accDim), out = double(chnLen*accDim))$out,
+         nrow = chnLen, byrow = T)
> grMat    <- matrix(gelRub("chn",nChn,chnLen), ncol = d, byrow = T)
> grMaxInd <- apply(grMat, 2, function(vec){which(vec == max(vec))[1]})
> acMat    <- mcmcAcf("chn", 5)
```

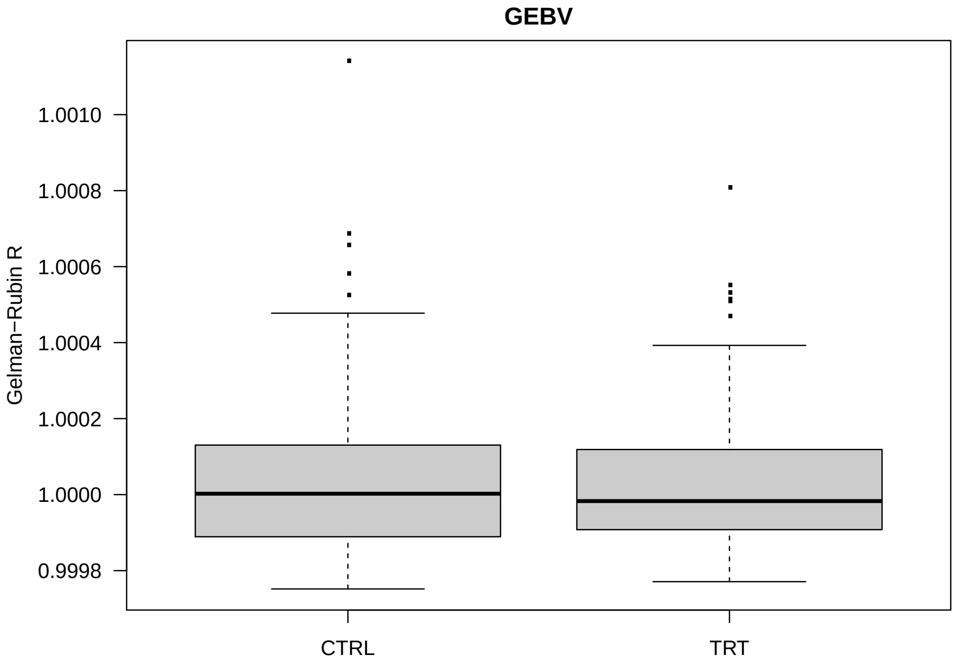


Plot the worst chains.

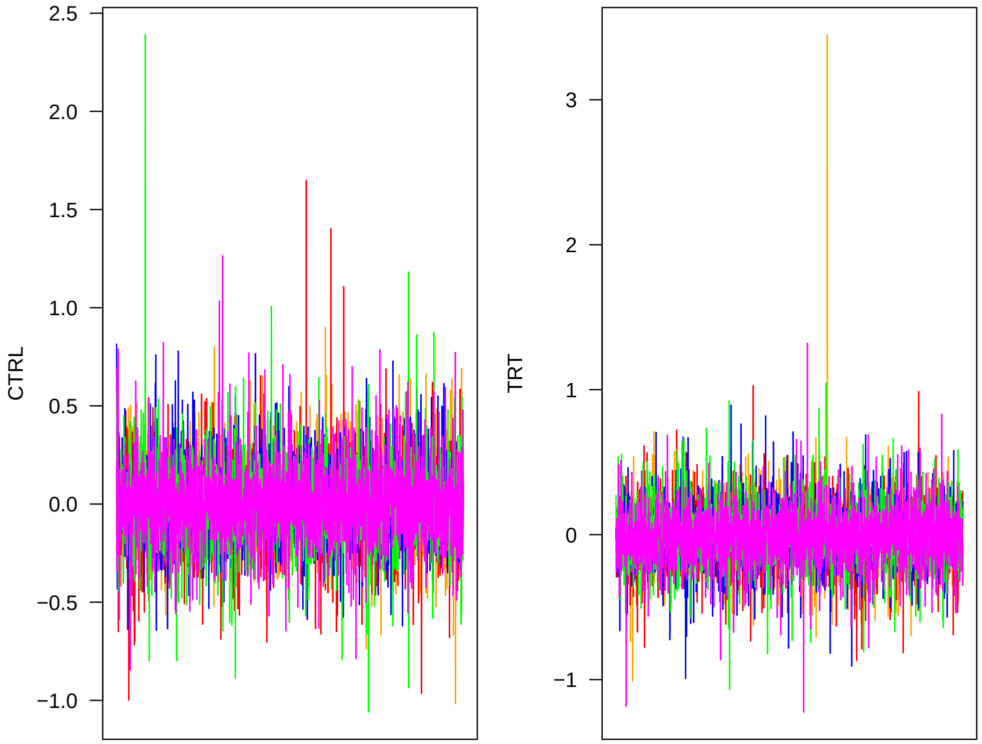


Now I look at mixing.

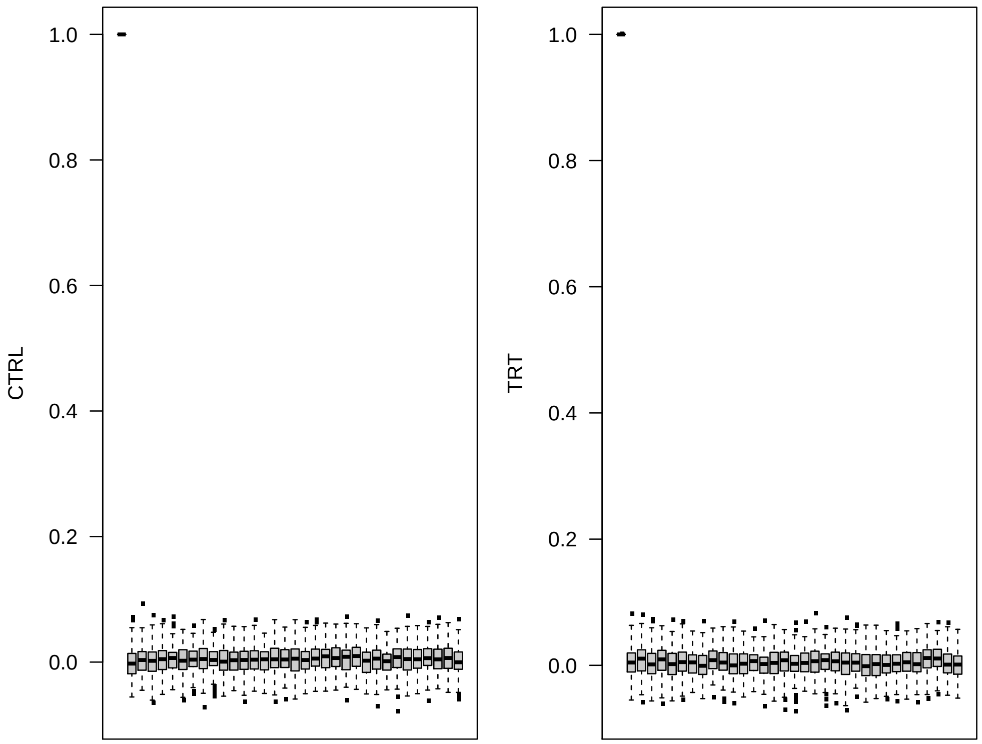


Next I look at covariance matrices. I create an index to isolate just the upper triangles.

```
> upInd   <- matrix(1:(d^2), ncol = d, byrow = T)
> upInd   <- upInd[row(upInd) <= col(upInd)]
> diagInd <- diag(matrix(1:(d^2), ncol = d, byrow = T))
```

Read in covariance matrix chains. Error correlations are set to zero, I exclude them.

```
> chn1 <- matrix(.C("GSLmatLoad",
+     "chains/SgE_3_1.gbin",
+     as.integer(chnLen), as.integer(d^2), out = double(chnLen*d^2))$out,
+         nrow = chnLen, byrow = T)[,diagInd]
> chn1 <- cbind(chn1,
+     matrix(.C("GSLmatLoad",
+     "chains/SgS_3_1.gbin",
+     as.integer(chnLen), as.integer(d^2), out = double(chnLen*d^2))$out,
+         nrow = chnLen, byrow = T)[,upInd])
> chn1 <- cbind(chn1,
+     matrix(.C("GSLmatLoad",
+     "chains/SgA_3_1.gbin",
+     as.integer(chnLen), as.integer(d^2), out = double(chnLen*d^2))$out,
+         nrow = chnLen, byrow = T)[,upInd])
> chn2 <- matrix(.C("GSLmatLoad",
+     "chains/SgE_3_2.gbin",
+     as.integer(chnLen), as.integer(d^2), out = double(chnLen*d^2))$out,
+         nrow = chnLen, byrow = T)[,diagInd]
> chn2 <- cbind(chn2,
+     matrix(.C("GSLmatLoad",
+     "chains/SgS_3_2.gbin",
+     as.integer(chnLen), as.integer(d^2), out = double(chnLen*d^2))$out,
+         nrow = chnLen, byrow = T)[,upInd])
> chn2 <- cbind(chn2,
+     matrix(.C("GSLmatLoad",
+     "chains/SgA_3_2.gbin",
+     as.integer(chnLen), as.integer(d^2), out = double(chnLen*d^2))$out,
+         nrow = chnLen, byrow = T)[,upInd])
> chn3 <- matrix(.C("GSLmatLoad",
+     "chains/SgE_3_3.gbin",
+     as.integer(chnLen), as.integer(d^2), out = double(chnLen*d^2))$out,
+         nrow = chnLen, byrow = T)[,diagInd]
> chn3 <- cbind(chn3,
+     matrix(.C("GSLmatLoad",
+     "chains/SgS_3_3.gbin",
+     as.integer(chnLen), as.integer(d^2), out = double(chnLen*d^2))$out,
+         nrow = chnLen, byrow = T)[,upInd])
> chn3 <- cbind(chn3,
+     matrix(.C("GSLmatLoad",
+     "chains/SgA_3_3.gbin",
```

```
+     as.integer(chnLen), as.integer(d^2), out = double(chnLen*d^2))$out,
+         nrow = chnLen, byrow = T)[,upInd])
> chn4 <- matrix(.C("GSLmatLoad",
+     "chains/SgE_3_4.gbin",
+     as.integer(chnLen), as.integer(d^2), out = double(chnLen*d^2))$out,
+         nrow = chnLen, byrow = T)[,diagInd]
> chn4 <- cbind(chn4,
+     matrix(.C("GSLmatLoad",
+     "chains/SgS_3_4.gbin",
+     as.integer(chnLen), as.integer(d^2), out = double(chnLen*d^2))$out,
+         nrow = chnLen, byrow = T)[,upInd])
> chn4 <- cbind(chn4,
+     matrix(.C("GSLmatLoad",
+     "chains/SgA_3_4.gbin",
+     as.integer(chnLen), as.integer(d^2), out = double(chnLen*d^2))$out,
+         nrow = chnLen, byrow = T)[,upInd])
> chn5 <- matrix(.C("GSLmatLoad",
+     "chains/SgE_3_5.gbin",
+     as.integer(chnLen), as.integer(d^2), out = double(chnLen*d^2))$out,
+         nrow = chnLen, byrow = T)[,diagInd]
> chn5 <- cbind(chn5,
+     matrix(.C("GSLmatLoad",
+     "chains/SgS_3_5.gbin",
+     as.integer(chnLen), as.integer(d^2), out = double(chnLen*d^2))$out,
+         nrow = chnLen, byrow = T)[,upInd])
> chn5 <- cbind(chn5,
+     matrix(.C("GSLmatLoad",
+     "chains/SgA_3_5.gbin",
+     as.integer(chnLen), as.integer(d^2), out = double(chnLen*d^2))$out,
+         nrow = chnLen, byrow = T)[,upInd])
> acMat <- mcmcAcf("chn", 5)
>
```

I have to make an index that marks each covariance matrix elements.

```
> cvrID <- c(rep("e", length(diagInd)), rep(c("s", "a"), each = length(upInd)))
> cvrInd <- 1:ncol(chn1)
```

Now plot the diagnostics.

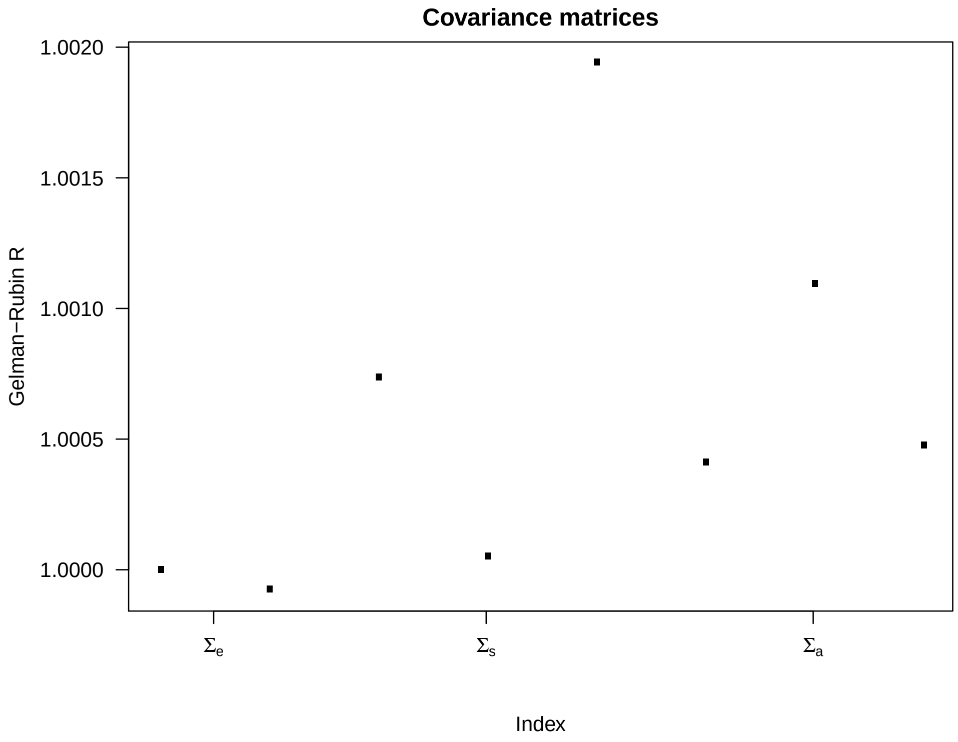


Now I look at mixing.

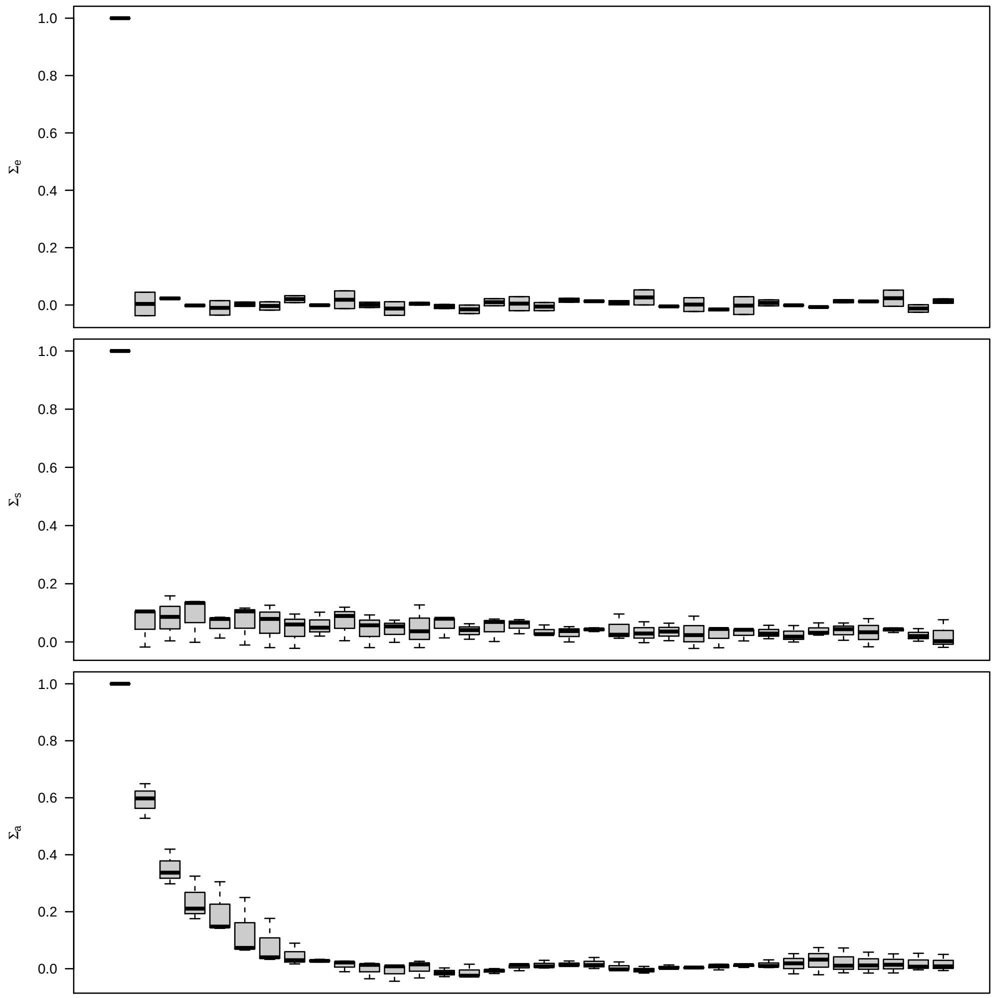


I plot the MCMC chains of variances (diagonal elements) for each covariance matrix. First, I get the diagonal index from the upper triangle index.

```
> diagInd <- list(1:2, c(3,5), c(6,8))
```

First, I plot the error variances.

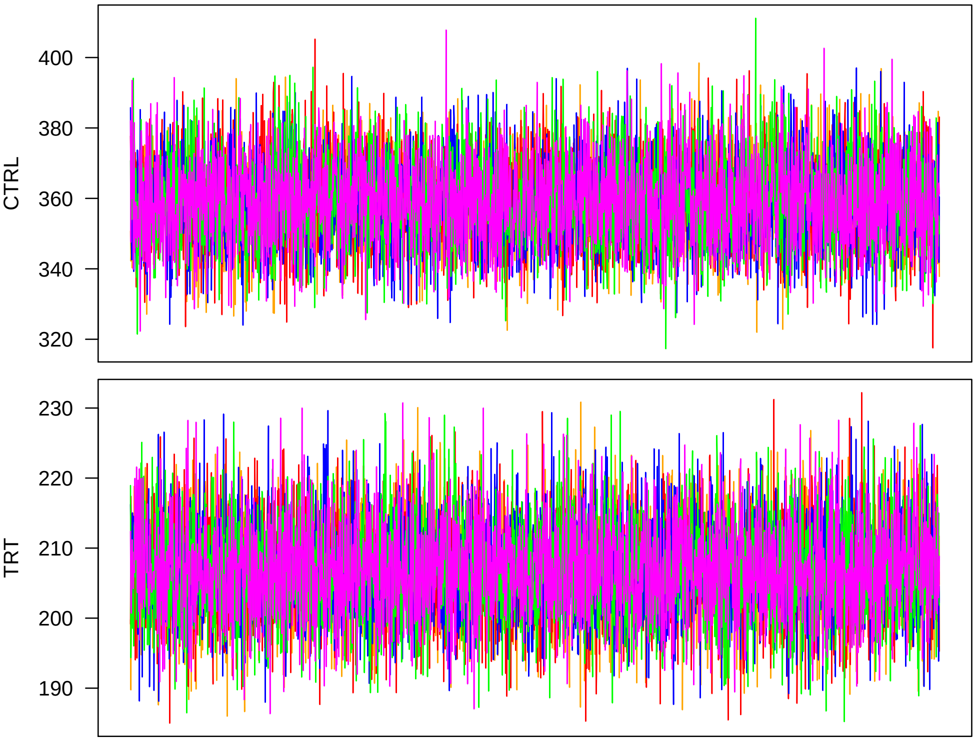


Next, non-additive genetic variances.

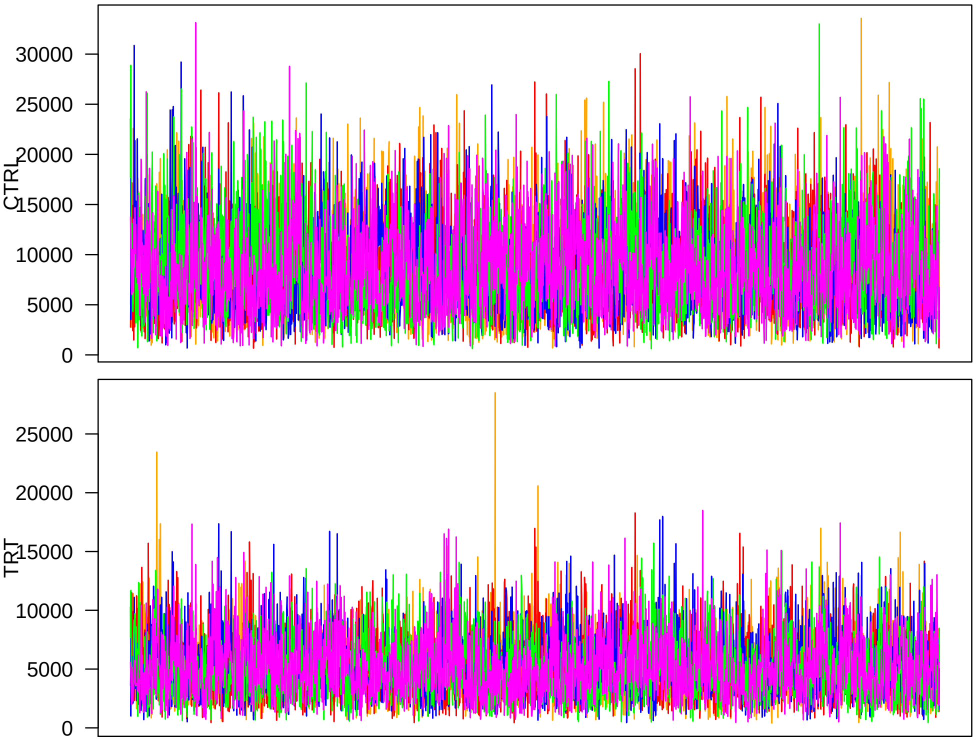

Finally, $\mathbf{\Sigma}_a$ which can be interpreted as the additive genetic covariance matrix in this case, where $\nu_g = 1000$ and the model is essentially Gaussian.

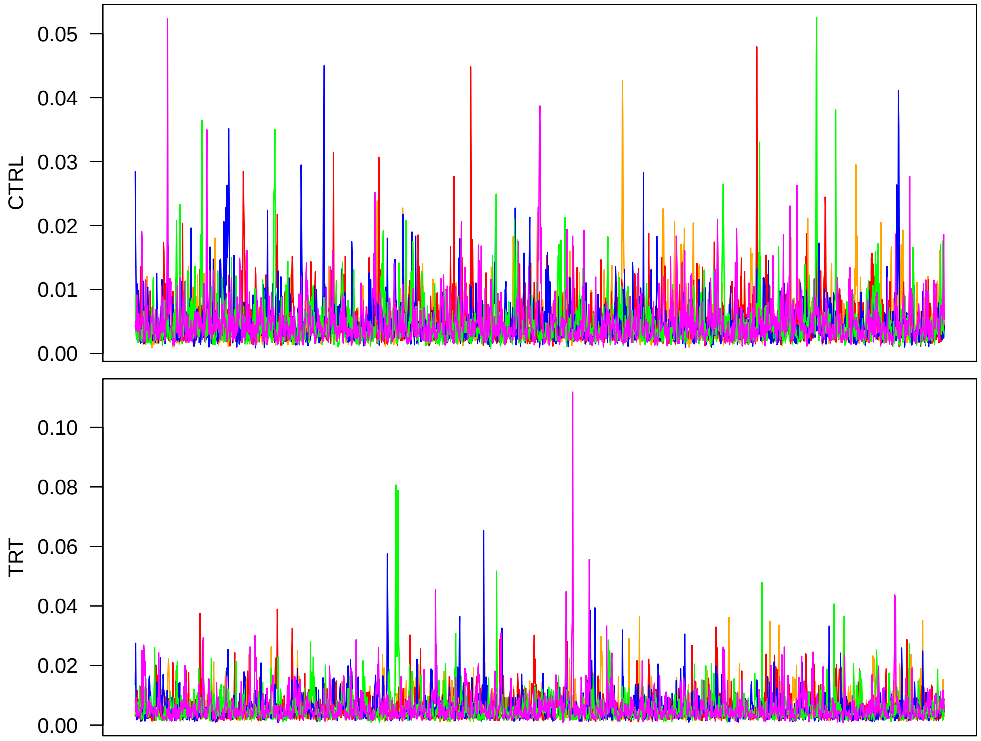


It is OK to proceed with summarizing the MCMC output for further analyses.

#### 2 Population structure control for GWA

Having extracted accession means from the model above, I collected three traits (the original two plus $\log_{10}$ RRG) and ran a model for genotypes with the accession means as responce. Here I check the MCMC output of this model.

I start with the genetic PC regression coefficients.

```
> trtNam <- c("CTRL", "TRT", "RRG")
> d      <- 3
> Npc    <- 168
> pcDim  <- d*Npc
> chn1 <- matrix(.C("GSLmatLoad", "chainsRRG/GM_3_1.gbin",
+     as.integer(chnLen), as.integer(pcDim), out = double(chnLen*pcDim))$out,
+         nrow = chnLen, byrow = T)
> chn2 <- matrix(.C("GSLmatLoad", "chainsRRG/GM_3_2.gbin",
+     as.integer(chnLen), as.integer(pcDim), out = double(chnLen*pcDim))$out,
+         nrow = chnLen, byrow = T)
> chn3 <- matrix(.C("GSLmatLoad", "chainsRRG/GM_3_3.gbin",
+     as.integer(chnLen), as.integer(pcDim), out = double(chnLen*pcDim))$out,
```

```
+         nrow = chnLen, byrow = T)
> chn4 <- matrix(.C("GSLmatLoad", "chainsRRG/GM_3_4.gbin",
+     as.integer(chnLen), as.integer(pcDim), out = double(chnLen*pcDim))$out,
+         nrow = chnLen, byrow = T)
> chn5 <- matrix(.C("GSLmatLoad", "chainsRRG/GM_3_5.gbin",
+     as.integer(chnLen), as.integer(pcDim), out = double(chnLen*pcDim))$out,
+         nrow = chnLen, byrow = T)
> grMat    <- matrix(gelRub("chn",nChn,chnLen), ncol = d, byrow = T)
> grMaxInd <- apply(grMat, 2, function(vec){which(vec == max(vec))[1]})
> acMat    <- mcmcAcf("chn", 5)
>
```

Plot the results.

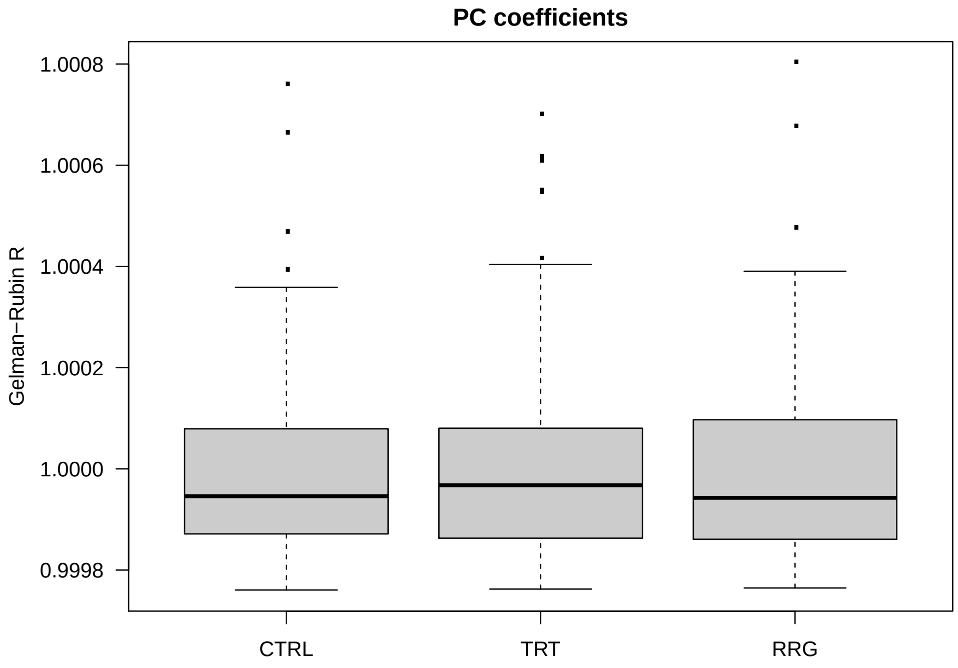


Plot the worst chains.

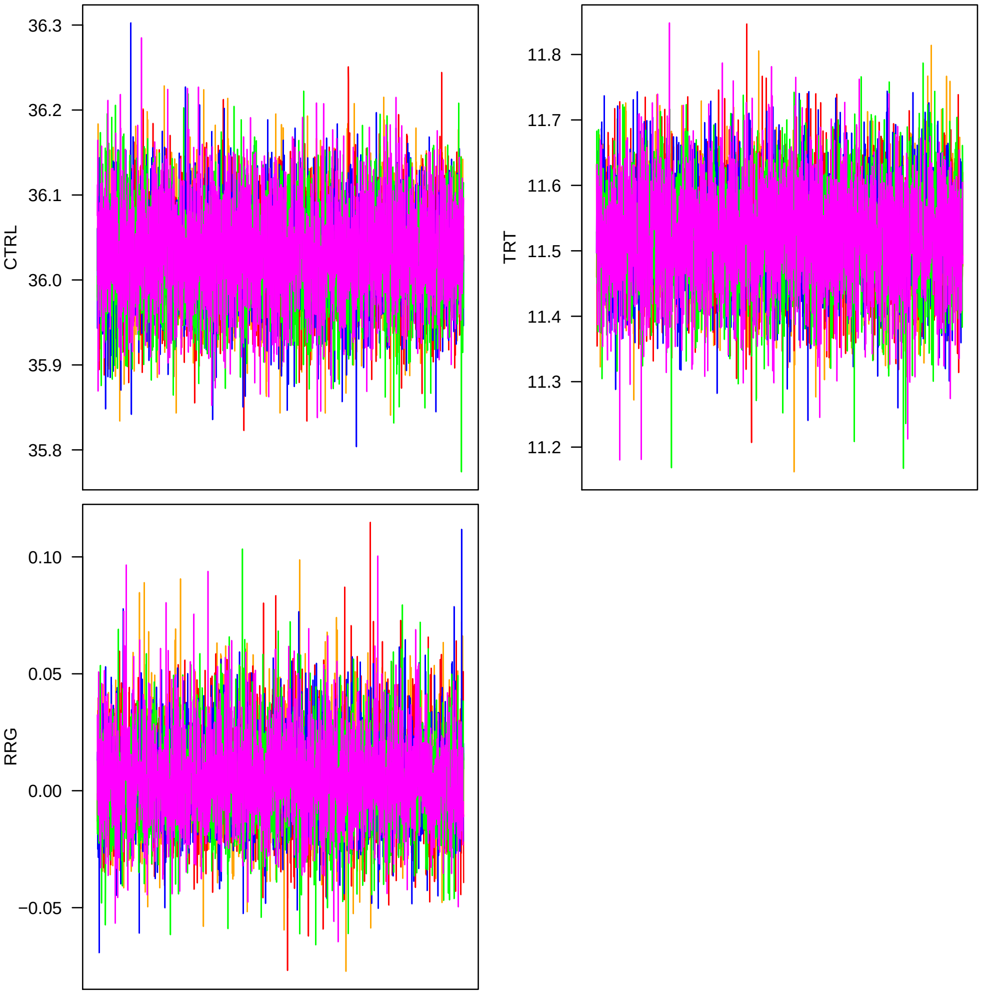


Now I look at mixing.

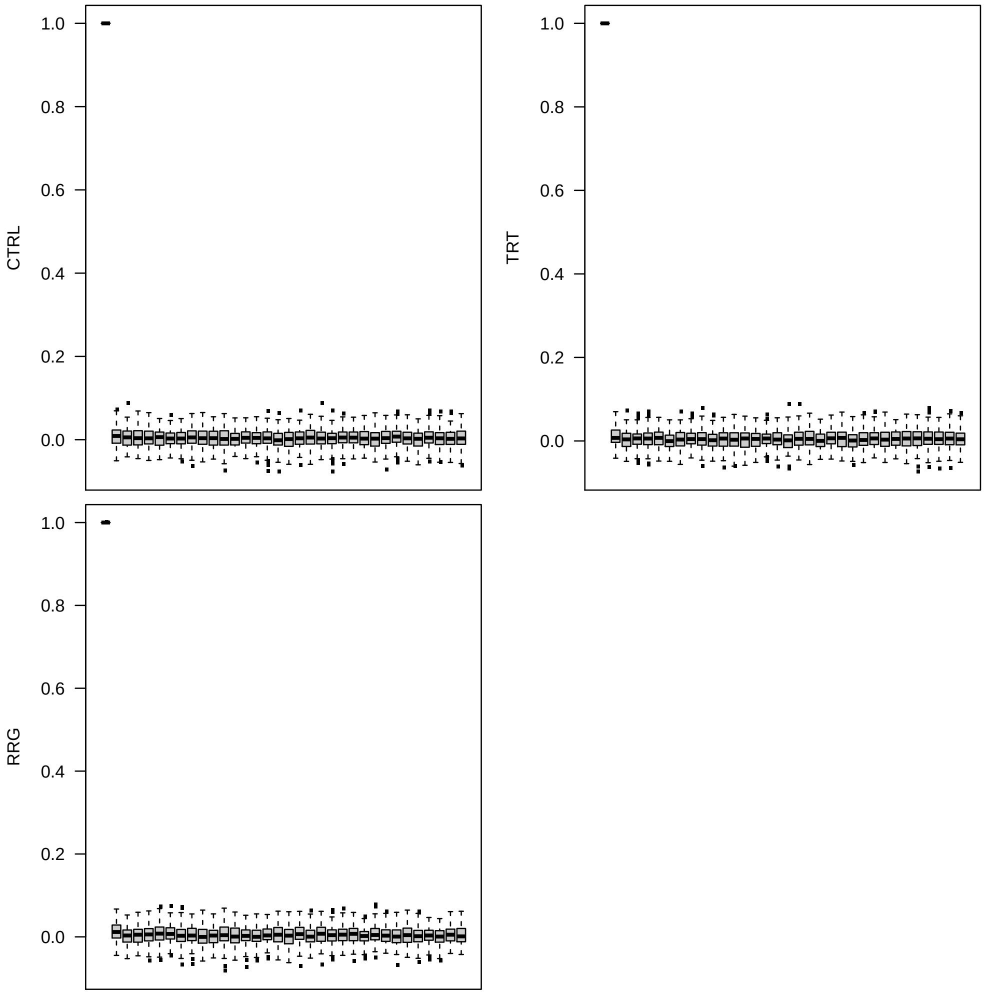


Next, look at GEBVs.

```
> accDim <- d*Nacc
> chn1 <- matrix(.C("GSLmatLoad", "chainsRRG/BV_3_1.gbin",
+     as.integer(chnLen), as.integer(accDim), out = double(chnLen*accDim))$out,
+         nrow = chnLen, byrow = T)
> chn2 <- matrix(.C("GSLmatLoad", "chainsRRG/BV_3_2.gbin",
+     as.integer(chnLen), as.integer(accDim), out = double(chnLen*accDim))$out,
+         nrow = chnLen, byrow = T)
> chn3 <- matrix(.C("GSLmatLoad", "chainsRRG/BV_3_3.gbin",
+     as.integer(chnLen), as.integer(accDim), out = double(chnLen*accDim))$out,
+         nrow = chnLen, byrow = T)
> chn4 <- matrix(.C("GSLmatLoad", "chainsRRG/BV_3_4.gbin",
+     as.integer(chnLen), as.integer(accDim), out = double(chnLen*accDim))$out,
+         nrow = chnLen, byrow = T)
> chn5 <- matrix(.C("GSLmatLoad", "chainsRRG/BV_3_5.gbin",
+     as.integer(chnLen), as.integer(accDim), out = double(chnLen*accDim))$out,
```

```
+          nrow = chnLen, byrow = T)
> grMat    <- matrix(gelRub("chn",nChn,chnLen), ncol = d, byrow = T)
> grMaxInd <- apply(grMat, 2, function(vec){which(vec == max(vec))[1]})
> acMat    <- mcmcAcf("chn", 5)
>
```

Plot the results.

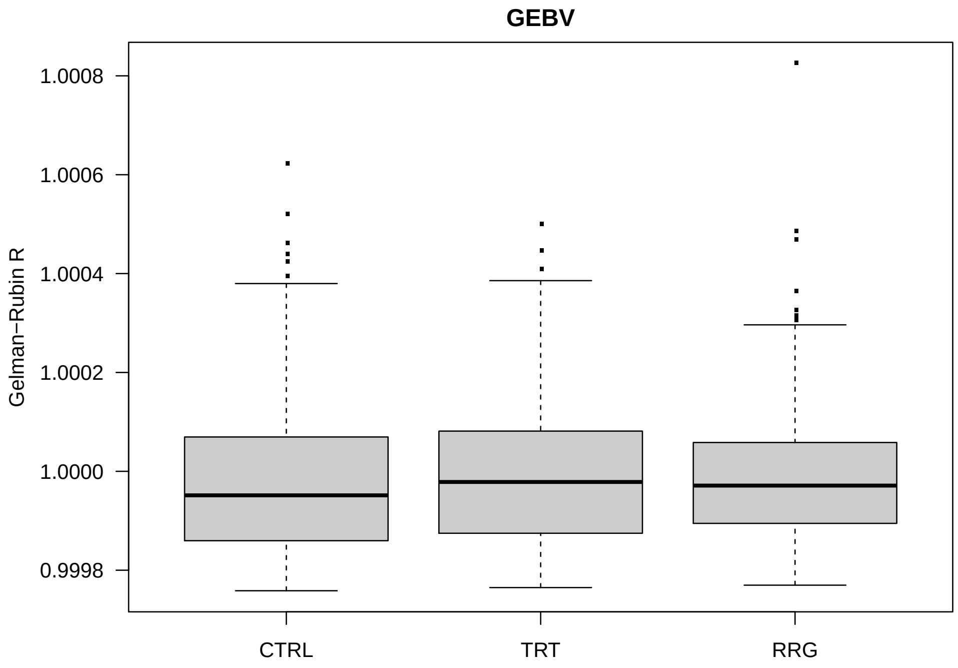


Plot the worst chains.

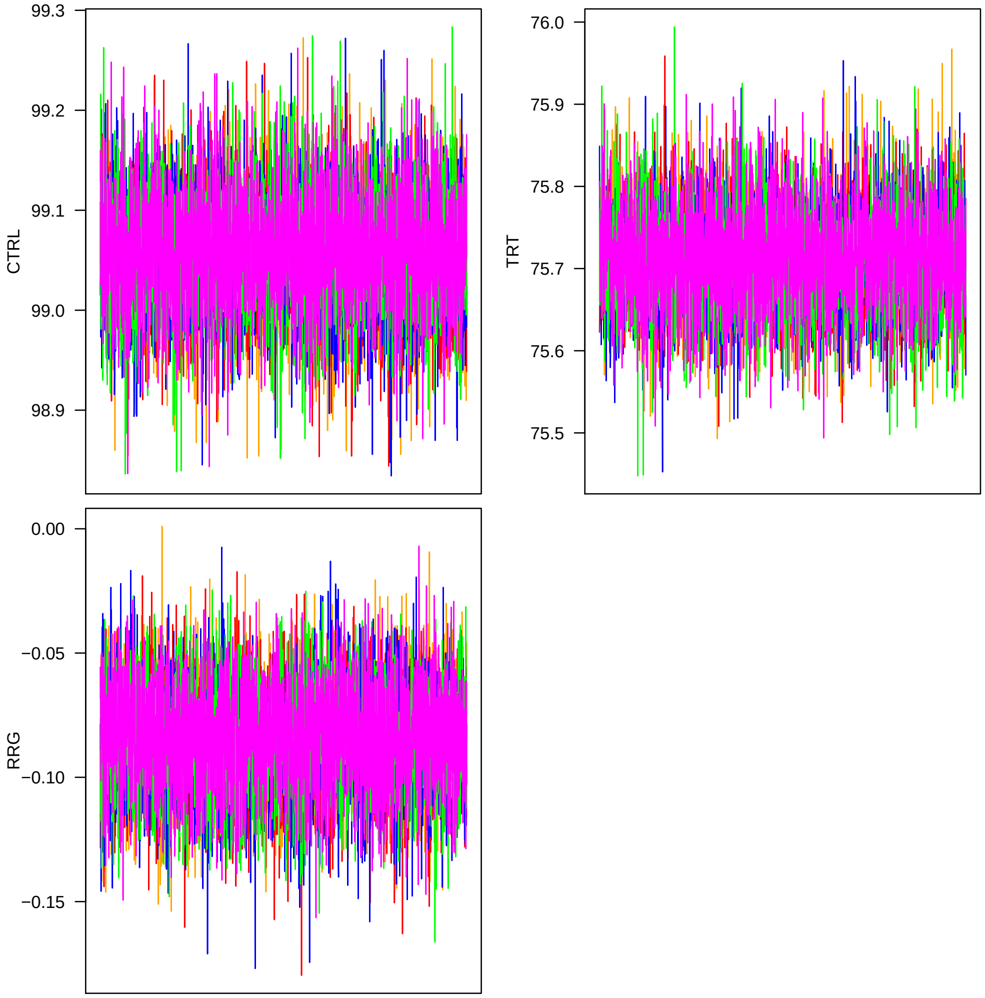


Now I look at mixing.

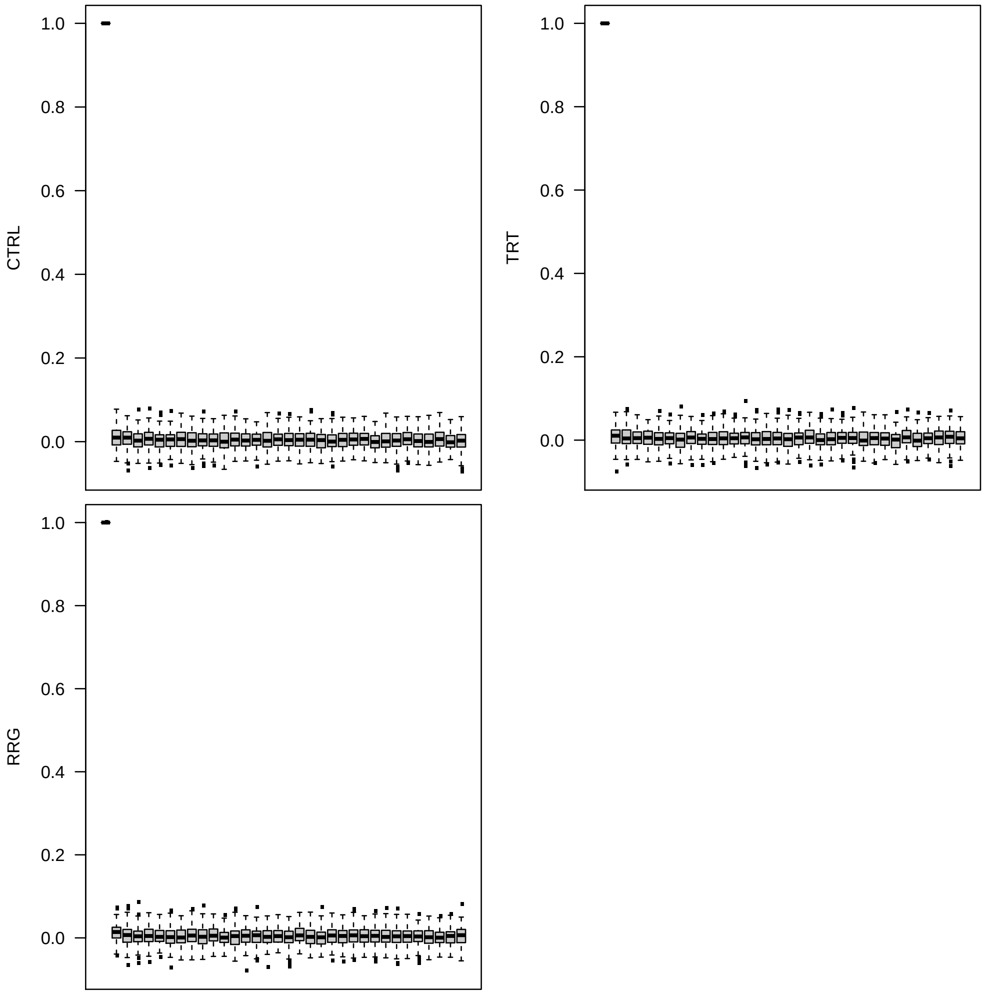


Next I look at covariance matrices. I create an index to isolate just the upper triangles.

```
> upInd   <- matrix(1:(d^2), ncol = d, byrow = T)
> upInd   <- upInd[row(upInd) <= col(upInd)]
```

Read in covariance matrix chains. Error correlations are set to zero, I exclude them.

```
> chn1 <- matrix(.C("GSLmatLoad",
+     "chainsRRG/SgS_3_1.gbin",
+     as.integer(chnLen), as.integer(d^2), out = double(chnLen*d^2))$out,
+         nrow = chnLen, byrow = T)[,upInd]
> chn1 <- cbind(chn1,
+     matrix(.C("GSLmatLoad",
+     "chainsRRG/SgA_3_1.gbin",
+     as.integer(chnLen), as.integer(d^2), out = double(chnLen*d^2))$out,
+         nrow = chnLen, byrow = T)[,upInd])
> chn2 <- matrix(.C("GSLmatLoad",
```

```
+     "chainsRRG/SgS_3_2.gbin",
+     as.integer(chnLen), as.integer(d^2), out = double(chnLen*d^2))$out,
+         nrow = chnLen, byrow = T)[,upInd]
> chn2 <- cbind(chn2,
+     matrix(.C("GSLmatLoad",
+     "chainsRRG/SgA_3_2.gbin",
+     as.integer(chnLen), as.integer(d^2), out = double(chnLen*d^2))$out,
+         nrow = chnLen, byrow = T)[,upInd])
> chn3 <- matrix(.C("GSLmatLoad",
+     "chainsRRG/SgS_3_3.gbin",
+     as.integer(chnLen), as.integer(d^2), out = double(chnLen*d^2))$out,
+         nrow = chnLen, byrow = T)[,upInd]
> chn3 <- cbind(chn3,
+     matrix(.C("GSLmatLoad",
+     "chainsRRG/SgA_3_3.gbin",
+     as.integer(chnLen), as.integer(d^2), out = double(chnLen*d^2))$out,
+         nrow = chnLen, byrow = T)[,upInd])
> chn4 <- matrix(.C("GSLmatLoad",
+     "chainsRRG/SgS_3_4.gbin",
+     as.integer(chnLen), as.integer(d^2), out = double(chnLen*d^2))$out,
+         nrow = chnLen, byrow = T)[,upInd]
> chn4 <- cbind(chn4,
+     matrix(.C("GSLmatLoad",
+     "chainsRRG/SgA_3_4.gbin",
+     as.integer(chnLen), as.integer(d^2), out = double(chnLen*d^2))$out,
+         nrow = chnLen, byrow = T)[,upInd])
> chn5 <- matrix(.C("GSLmatLoad",
+     "chainsRRG/SgS_3_5.gbin",
+     as.integer(chnLen), as.integer(d^2), out = double(chnLen*d^2))$out,
+         nrow = chnLen, byrow = T)[,upInd]
> chn5 <- cbind(chn5,
+     matrix(.C("GSLmatLoad",
+     "chainsRRG/SgA_3_5.gbin",
+     as.integer(chnLen), as.integer(d^2), out = double(chnLen*d^2))$out,
+         nrow = chnLen, byrow = T)[,upInd])
> acMat <- mcmcAcf("chn", 5)
>
```

Plot the results.

Gelman-Rubin:

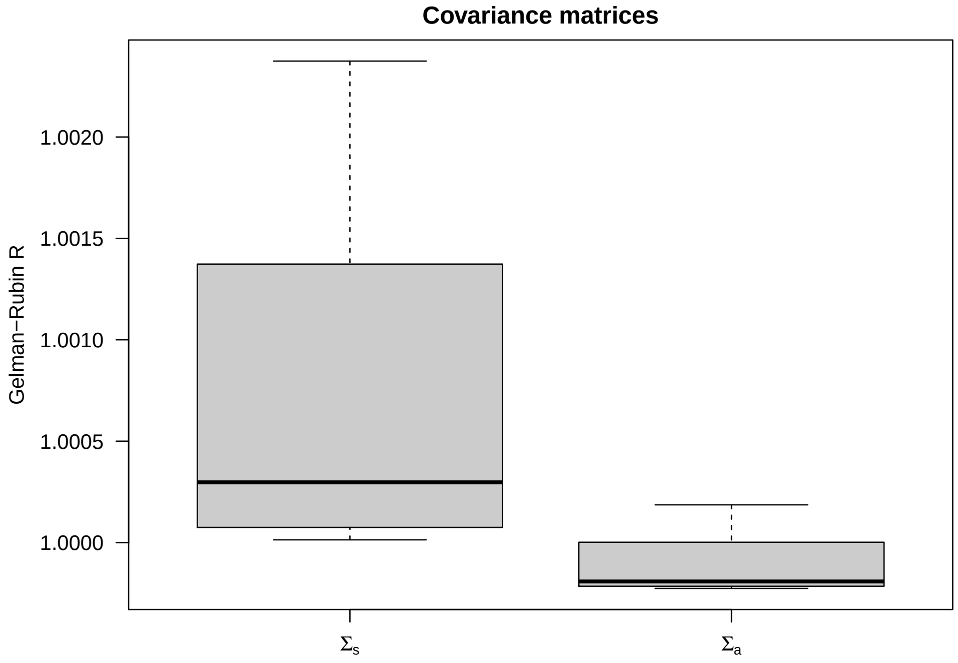


Now I look at mixing.

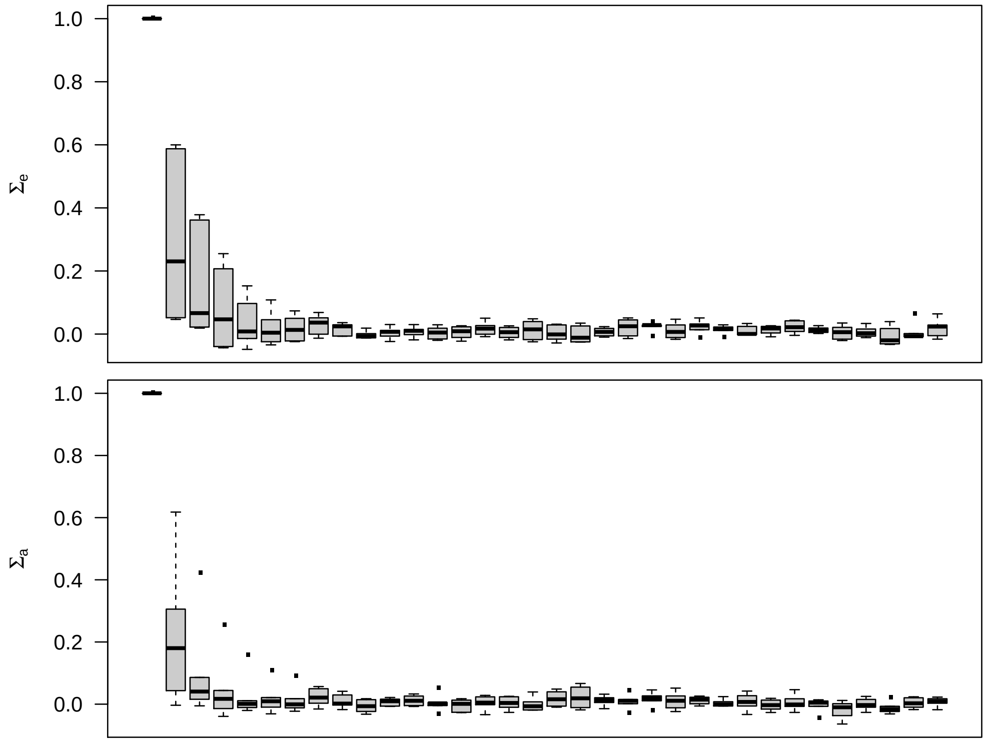


```
> tmpInd  <- matrix(1:(d^2), ncol = d, byrow = T)
```

```
> tmpInd  <- tmpInd[row(tmpInd) < col(tmpInd)]
> diagInd <- which(!(upInd %in% tmpInd))
> rm(tmpInd)
```

First, I plot the error (SCA) variances.

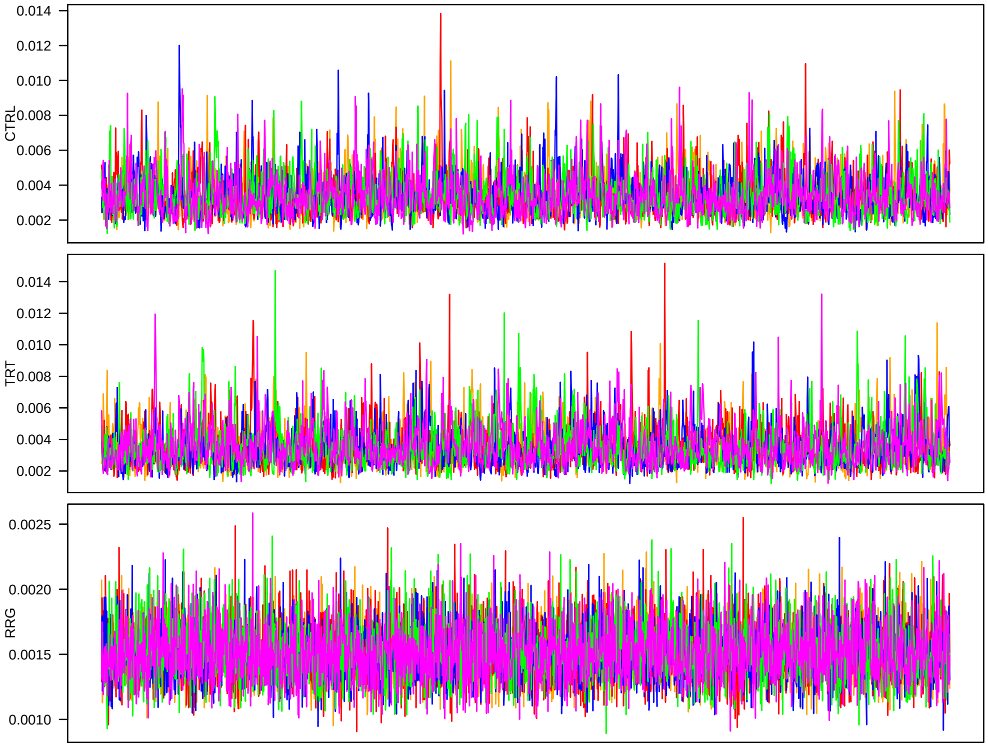


Next, the $\sigma_a^2$.

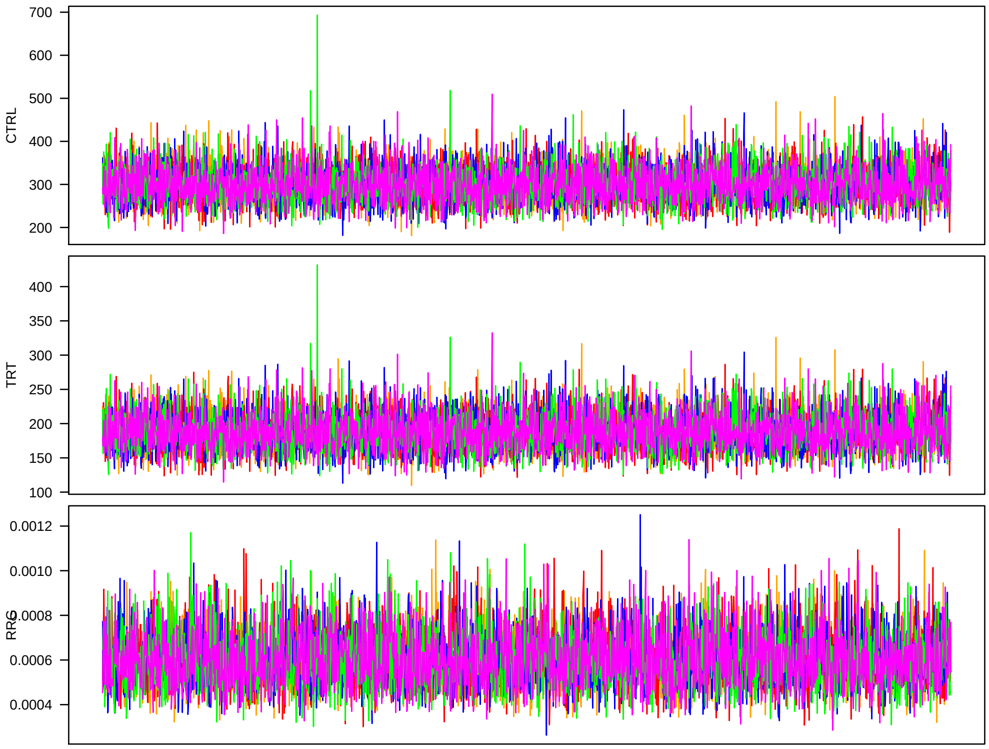


Everything looks reasonable. Proceeding with MCMC summaries for use in GWA.

Finally, I try a model with essentially a Gaussian prior on the PC coefficients (analogous to a regular mixed model).

```
> chn1 <- matrix(.C("GSLmatLoad", "chainsRRG/GM_1000_1.gbin",
+     as.integer(chnLen), as.integer(pcDim), out = double(chnLen*pcDim))$out,
+         nrow = chnLen, byrow = T)
> chn2 <- matrix(.C("GSLmatLoad", "chainsRRG/GM_1000_2.gbin",
+     as.integer(chnLen), as.integer(pcDim), out = double(chnLen*pcDim))$out,
+         nrow = chnLen, byrow = T)
> chn3 <- matrix(.C("GSLmatLoad", "chainsRRG/GM_1000_3.gbin",
+     as.integer(chnLen), as.integer(pcDim), out = double(chnLen*pcDim))$out,
+         nrow = chnLen, byrow = T)
> chn4 <- matrix(.C("GSLmatLoad", "chainsRRG/GM_1000_4.gbin",
+     as.integer(chnLen), as.integer(pcDim), out = double(chnLen*pcDim))$out,
+         nrow = chnLen, byrow = T)
> chn5 <- matrix(.C("GSLmatLoad", "chainsRRG/GM_1000_5.gbin",
+     as.integer(chnLen), as.integer(pcDim), out = double(chnLen*pcDim))$out,
+         nrow = chnLen, byrow = T)
> grMat    <- matrix(gelRub("chn",nChn,chnLen), ncol = d, byrow = T)
> grMaxInd <- apply(grMat, 2, function(vec){which(vec == max(vec))[1]})
> acMat    <- mcmcAcf("chn", 5)
>
```

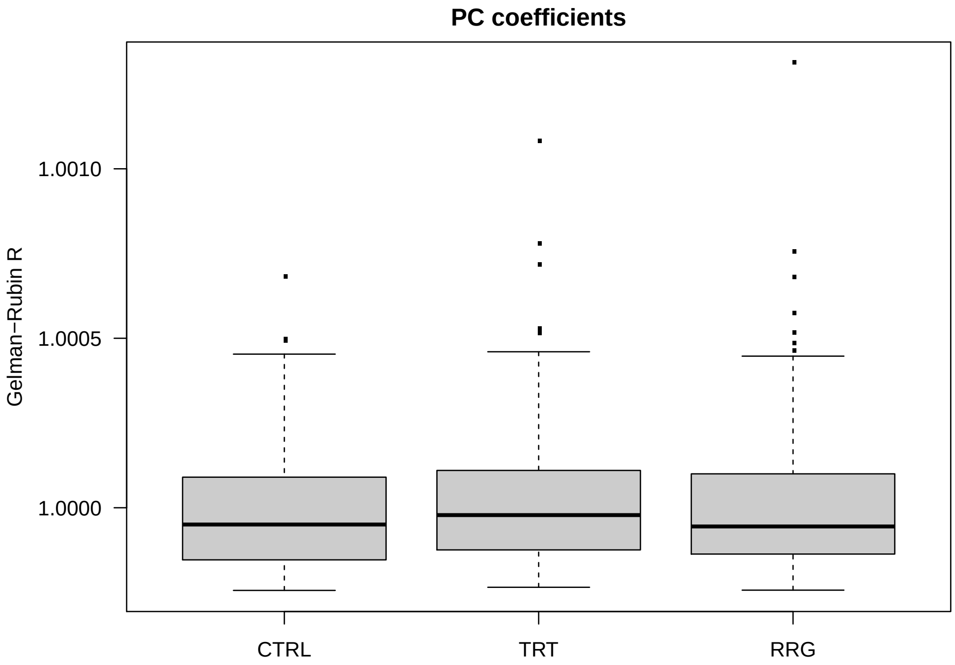


Plot the worst chains.

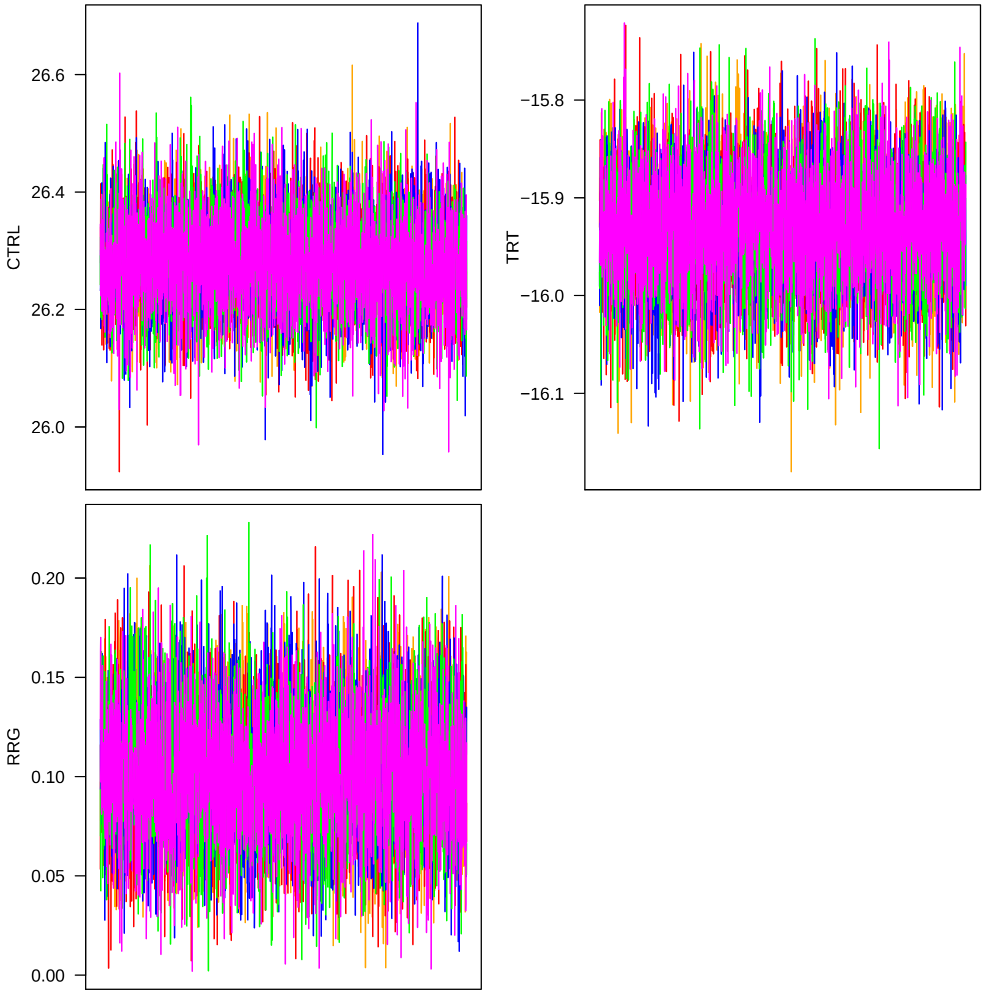


Now I look at mixing.

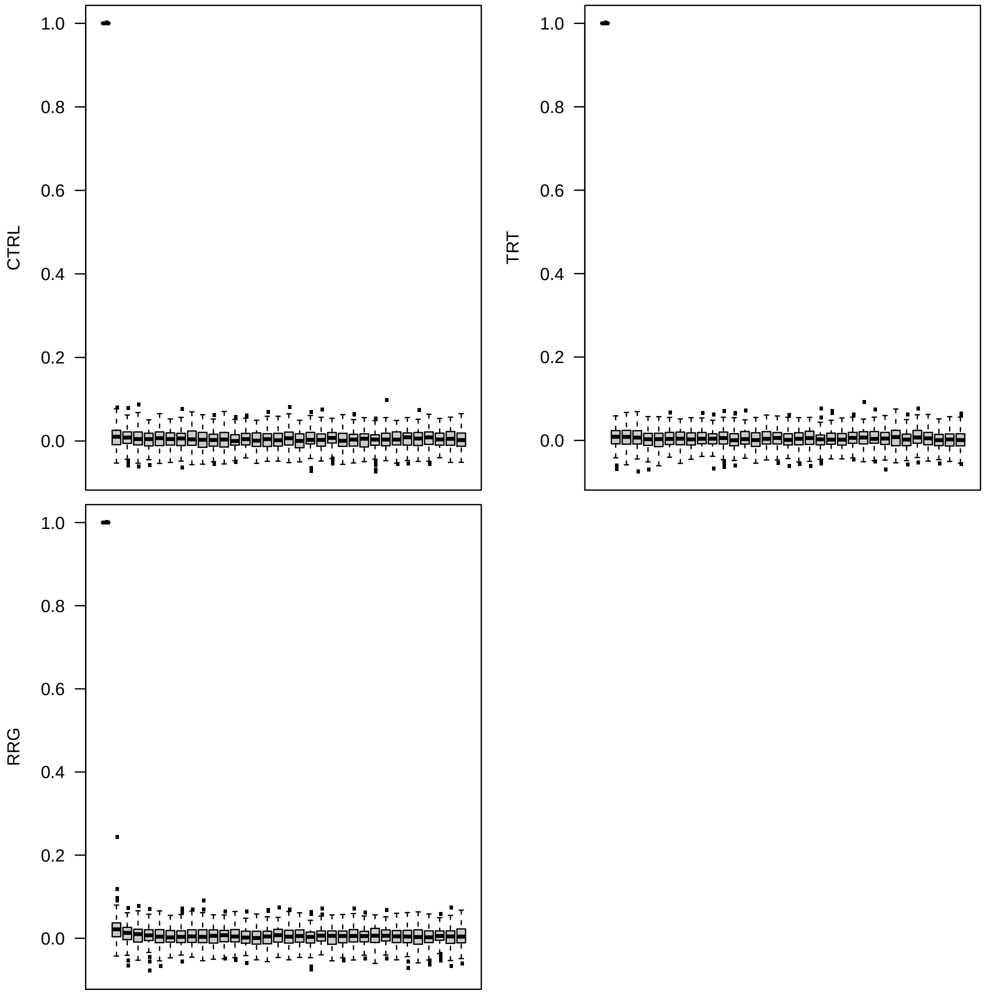


Next, look at GEBVs.

```
> chn1 <- matrix(.C("GSLmatLoad", "chainsRRG/BV_1000_1.gbin",
+     as.integer(chnLen), as.integer(accDim), out = double(chnLen*accDim))$out,
+         nrow = chnLen, byrow = T)
> chn2 <- matrix(.C("GSLmatLoad", "chainsRRG/BV_1000_2.gbin",
+     as.integer(chnLen), as.integer(accDim), out = double(chnLen*accDim))$out,
+         nrow = chnLen, byrow = T)
> chn3 <- matrix(.C("GSLmatLoad", "chainsRRG/BV_1000_3.gbin",
+     as.integer(chnLen), as.integer(accDim), out = double(chnLen*accDim))$out,
+         nrow = chnLen, byrow = T)
> chn4 <- matrix(.C("GSLmatLoad", "chainsRRG/BV_1000_4.gbin",
+     as.integer(chnLen), as.integer(accDim), out = double(chnLen*accDim))$out,
+         nrow = chnLen, byrow = T)
> chn5 <- matrix(.C("GSLmatLoad", "chainsRRG/BV_1000_5.gbin",
+     as.integer(chnLen), as.integer(accDim), out = double(chnLen*accDim))$out,
+         nrow = chnLen, byrow = T)
```

```
> grMat    <- matrix(gelRub("chn",nChn,chnLen), ncol = d, byrow = T)
> grMaxInd <- apply(grMat, 2, function(vec){which(vec == max(vec))[1]})
> acMat    <- mcmcAcf("chn", 5)
```

Plot the results.

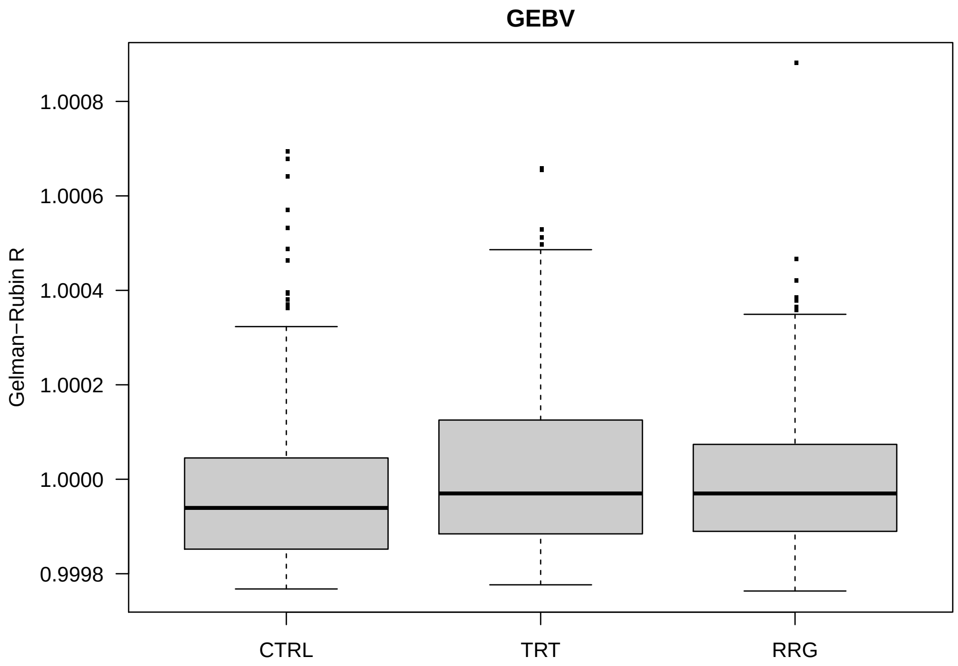


Plot the worst chains.

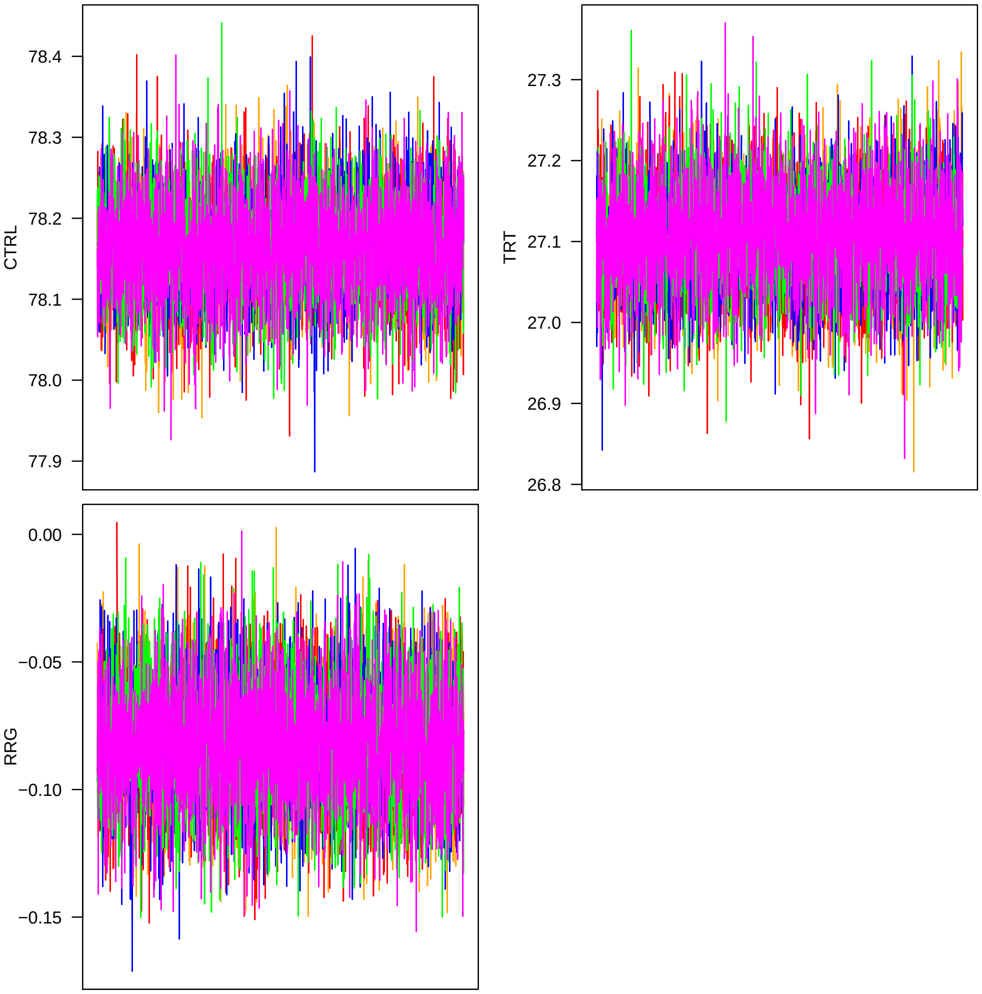


Now I look at mixing.

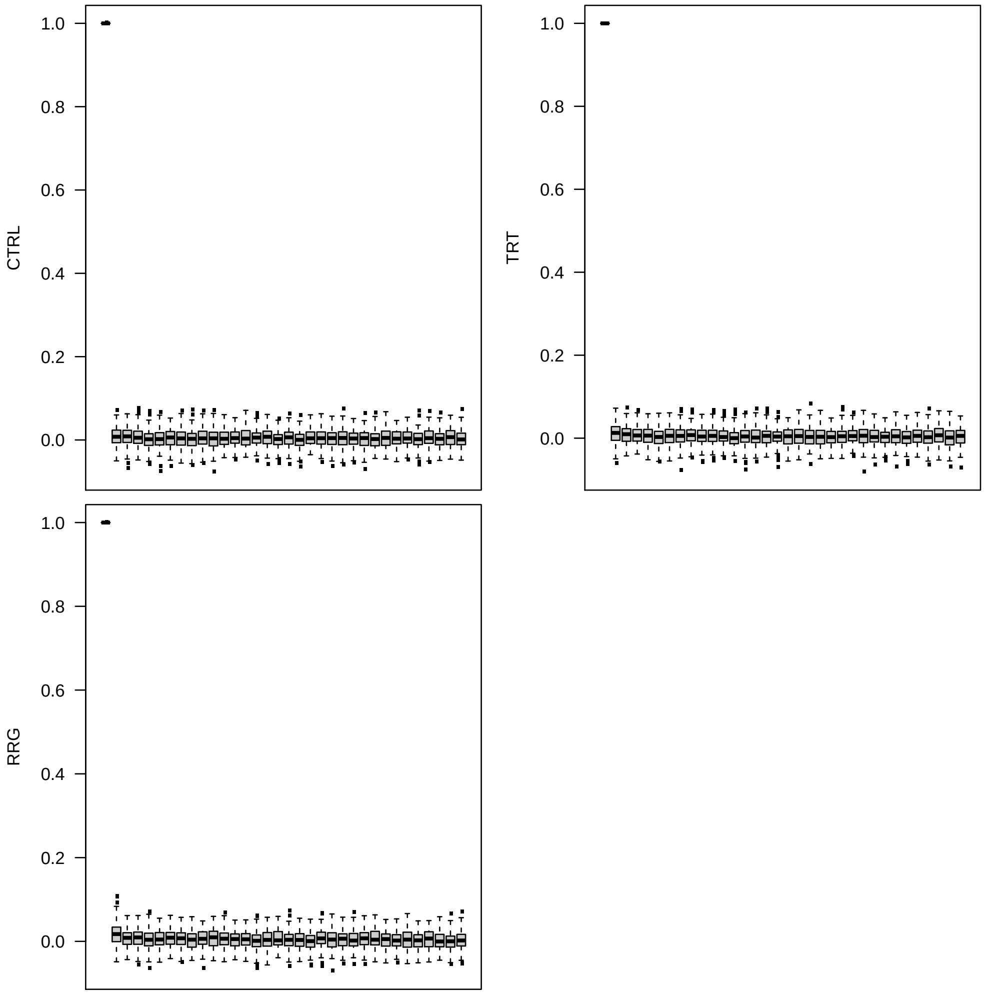


Next I look at covariance matrices. I create an index to isolate just the upper triangles.

```
> chn1 <- matrix(.C("GSLmatLoad",
+     "chainsRRG/SgS_1000_1.gbin",
+     as.integer(chnLen), as.integer(d^2), out = double(chnLen*d^2))$out,
+         nrow = chnLen, byrow = T)[,upInd]
> chn1 <- cbind(chn1,
+     matrix(.C("GSLmatLoad",
+     "chainsRRG/SgA_1000_1.gbin",
+     as.integer(chnLen), as.integer(d^2), out = double(chnLen*d^2))$out,
+         nrow = chnLen, byrow = T)[,upInd])
> chn2 <- matrix(.C("GSLmatLoad",
+     "chainsRRG/SgS_1000_2.gbin",
+     as.integer(chnLen), as.integer(d^2), out = double(chnLen*d^2))$out,
+         nrow = chnLen, byrow = T)[,upInd]
> chn2 <- cbind(chn2,
+     matrix(.C("GSLmatLoad",
```

```
+     "chainsRRG/SgA_1000_2.gbin",
+     as.integer(chnLen), as.integer(d^2), out = double(chnLen*d^2))$out,
+         nrow = chnLen, byrow = T)[,upInd])
> chn3 <- matrix(.C("GSLmatLoad",
+     "chainsRRG/SgS_1000_3.gbin",
+     as.integer(chnLen), as.integer(d^2), out = double(chnLen*d^2))$out,
+         nrow = chnLen, byrow = T)[,upInd]
> chn3 <- cbind(chn3,
+     matrix(.C("GSLmatLoad",
+     "chainsRRG/SgA_1000_3.gbin",
+     as.integer(chnLen), as.integer(d^2), out = double(chnLen*d^2))$out,
+         nrow = chnLen, byrow = T)[,upInd])
> chn4 <- matrix(.C("GSLmatLoad",
+     "chainsRRG/SgS_1000_4.gbin",
+     as.integer(chnLen), as.integer(d^2), out = double(chnLen*d^2))$out,
+         nrow = chnLen, byrow = T)[,upInd]
> chn4 <- cbind(chn4,
+     matrix(.C("GSLmatLoad",
+     "chainsRRG/SgA_1000_4.gbin",
+     as.integer(chnLen), as.integer(d^2), out = double(chnLen*d^2))$out,
+         nrow = chnLen, byrow = T)[,upInd])
> chn5 <- matrix(.C("GSLmatLoad",
+     "chainsRRG/SgS_1000_5.gbin",
+     as.integer(chnLen), as.integer(d^2), out = double(chnLen*d^2))$out,
+         nrow = chnLen, byrow = T)[,upInd]
> chn5 <- cbind(chn5,
+     matrix(.C("GSLmatLoad",
+     "chainsRRG/SgA_1000_5.gbin",
+     as.integer(chnLen), as.integer(d^2), out = double(chnLen*d^2))$out,
+         nrow = chnLen, byrow = T)[,upInd])
> acMat <- mcmcAcf("chn", 5)
>
```

Plot the results.

Gelman-Rubin:

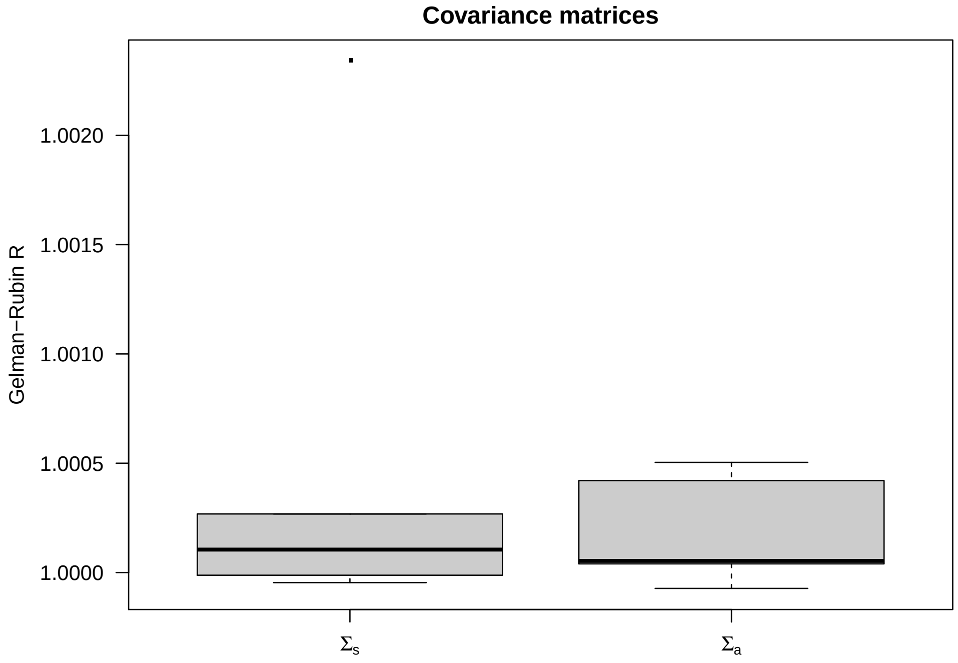


Now I look at mixing.

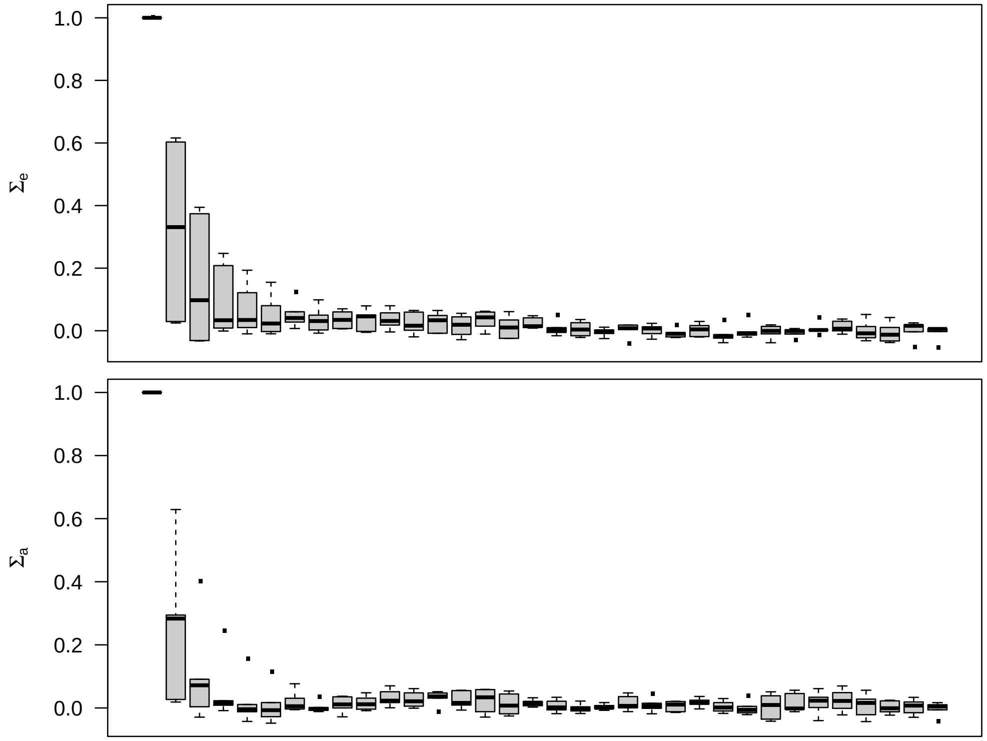

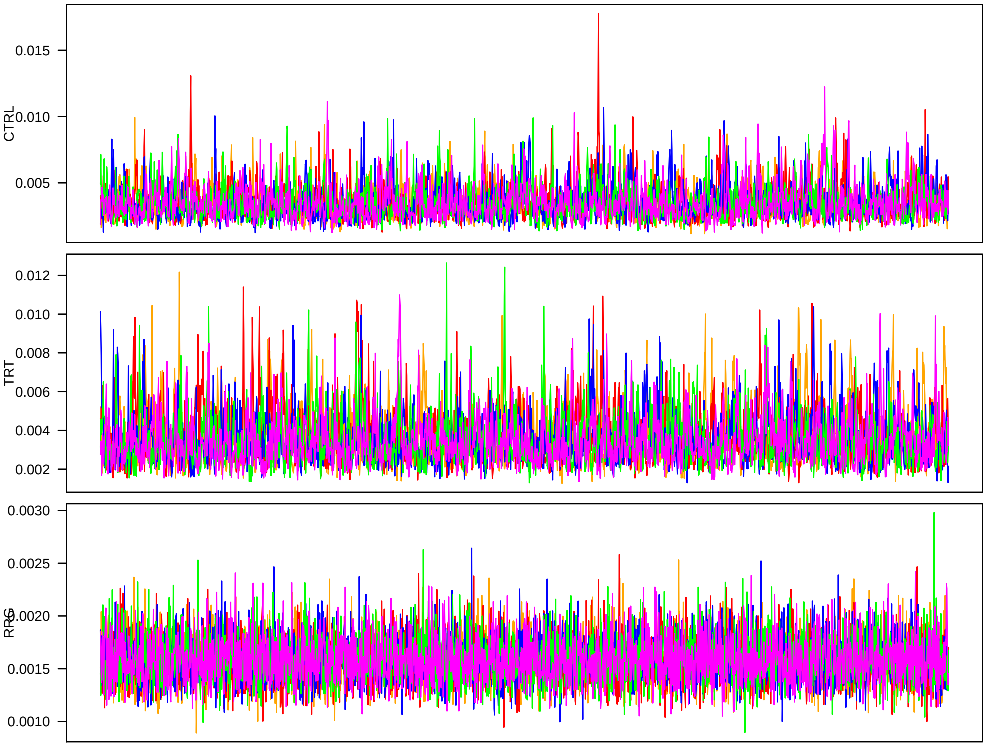


Next, the $\sigma_a^2$.

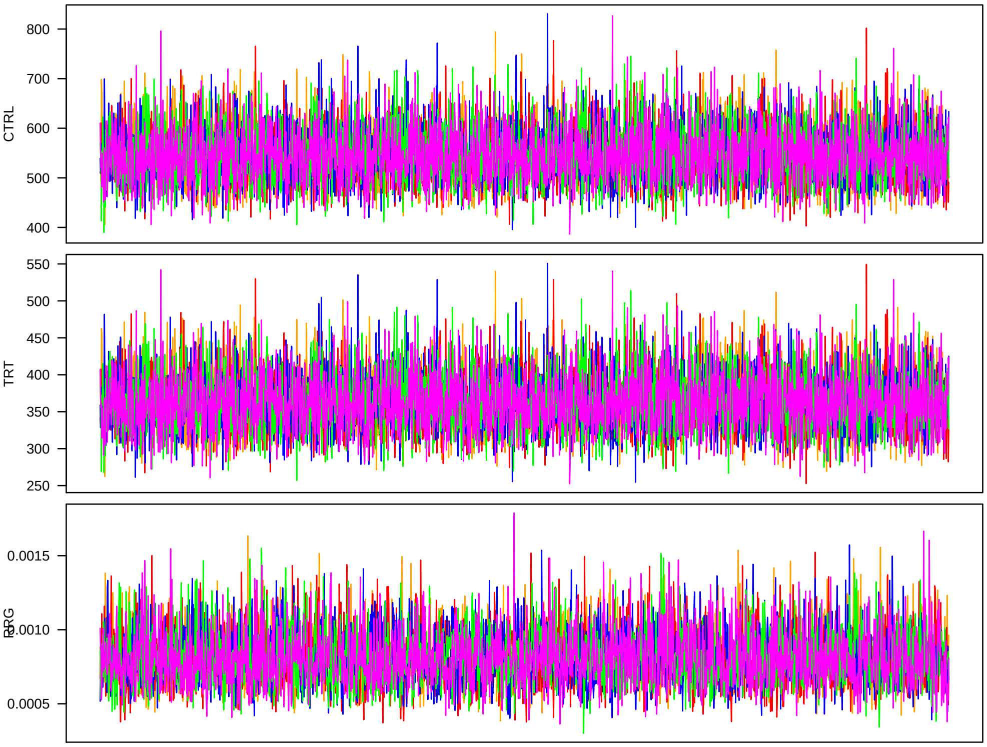


Everything is ready to proceed.
