## Supplemental methods and scripts for "Low additive genetic variation in a trait under selection in domesticated rice": mcmcExamineFAM.pdf

### MCMC convergence analyses

Tony Greenberg

May 4, 2018

```
R version 3.4.2 (2017-09-28)
Platform: x86_64-apple-darwin17.2.0 (64-bit)
Running under: macOS High Sierra 10.13.4

Matrix products: default
BLAS: /System/Library/Frameworks/Accelerate.framework/Versions/A/Frameworks/vecLib.framework/Versions/A/
LAPACK: /Library/Frameworks/R.framework/Versions/3.4/Resources/lib/libRlapack.dylib

locale:
[1] en_US.UTF-8/en_US.UTF-8/en_US.UTF-8/C/en_US.UTF-8/en_US.UTF-8

attached base packages:
[1] compiler  stats     graphics  grDevices utils     methods   base

other attached packages:
[1] showtext_0.5    showtextdb_2.0  sysfonts_0.7.1  gridExtra_2.3   ggplot2_2.2.1
[6] lattice_0.20-35

loaded via a namespace (and not attached):
 [1] Rcpp_0.12.13     digest_0.6.12    grid_3.4.2       plyr_1.8.4       gtable_0.2.0
 [6] scales_0.5.0     rlang_0.1.4      lazyeval_0.2.1   labeling_0.3     tools_3.4.2
[11] munsell_0.4.3    colorspace_1.3-2 tibble_1.3.4
```

In this document I examine convergence of Markov chains generated for the Al tolerance project on the old multipopulation data set. I start by looking at the experiment ID regression coefficients, first in analyses using *aus* accessions, then using tropical *japonica* lines.

The chain files are in the binary format generated by saving GSL matrices. R functions to read these files are available from the MuGen GitHub page.

I first define a function that calculates the Gelman-Rubin convergence statistic for multiple MCMC chains.

```
> gelRub <- cmpfun(function(vrNam, nChn, chnLen){
+     mat <- NULL
+     for (i in 1:nChn){
+         mat <- rbind(mat, eval(as.symbol(paste(vrNam, i, sep=""))))
+     }
+     chnFac <- factor(rep(1:nChn, each=chnLen))
+     W <- colMeans(apply(mat, 2, tapply, chnFac, var))
```

```
+     B <- apply(apply(mat, 2, tapply, chnFac, mean), 2, var)
+     return( sqrt((((chnLen - 1)/chnLen)*W + B)/W) )
+ })
>
```

I am also interested in the quality of chain mixing. I use autocorrelations to test for that. The following function examines autocorrelations for each chain for a given variable and reports the chain with the highest mean value. The autocorrelations are calculated using default parameters of the `acf()` function from the `stats` R package.

```
> eachAcf <- cmpfun(function(i, nChn, vrNam){
+     tmp <- NULL
+     for (j in 1:nChn){
+         tmp <- cbind(tmp,
+                 acf(eval(as.symbol(paste(vrNam, j, sep=""))))[,i], plot = F)$acf
+             )
+     }
+     return(tmp[,which.max(colMeans(tmp[-1,]))])
+ })
> mcmcAcf <- cmpfun(function(vrNam, nChn){
+      # all chains should have the same dimensions, so take the first one
+     N <- ncol(eval(as.symbol(paste(vrNam, 1, sep=""))))
+     t(sapply(1:N, eachAcf, nChn, vrNam))
+ })
```

Read the chains.

```
> trtNam <- c("Control", "160uM_Al")
> d      <- 2
> Nexp   <- 6
> expDim <- d*Nexp
> nChn   <- 5
> chnLen <- 2000
> chn1 <- matrix(.C("GSLmatLoad",
+     "chains/EXPout_aus_3_1.gbin",
+     as.integer(chnLen), as.integer(expDim), out = double(chnLen*expDim))$out,
+         nrow = chnLen, byrow = T)
> chn2 <- matrix(.C("GSLmatLoad",
+     "chains/EXPout_aus_3_2.gbin",
+     as.integer(chnLen), as.integer(expDim), out = double(chnLen*expDim))$out,
+         nrow = chnLen, byrow = T)
> chn3 <- matrix(.C("GSLmatLoad",
+     "chains/EXPout_aus_3_3.gbin",
+     as.integer(chnLen), as.integer(expDim), out = double(chnLen*expDim))$out,
+         nrow = chnLen, byrow = T)
> chn4 <- matrix(.C("GSLmatLoad",
+     "chains/EXPout_aus_3_4.gbin",
```

```
+     as.integer(chnLen), as.integer(expDim), out = double(chnLen*expDim))$out,
+         nrow = chnLen, byrow = T)
> chn5 <- matrix(.C("GSLmatLoad",
+     "chains/EXPout_aus_3_5.gbin",
+     as.integer(chnLen), as.integer(expDim), out = double(chnLen*expDim))$out,
+         nrow = chnLen, byrow = T)
> grMat    <- matrix(gelRub("chn", nChn, chnLen), ncol = d, byrow = T)
> grFrame  <- data.frame(Gelman.Rubin=c(grMat), trait=rep(trtNam, each = Nexp))
> grMaxInd <- apply(grMat, 2, function(vec){which(vec == max(vec))[1]})
> acMat    <- mcmcAcf("chn", 5)
> acFrame  <- data.frame(autocorrelation=c(acMat), trait=rep(trtNam, prod(dim(acMat))/2),
+     lag=rep(0:33, each = nrow(acMat)))
>
```

Plot the *aus* results.

```
> showtext_auto()
> ggplot(data=grFrame,
+         aes(x=trait, y=Gelman.Rubin)) +
+     geom_boxplot(fill="grey80") +
+     theme_classic(base_size=18, base_family="myriad")
>
```

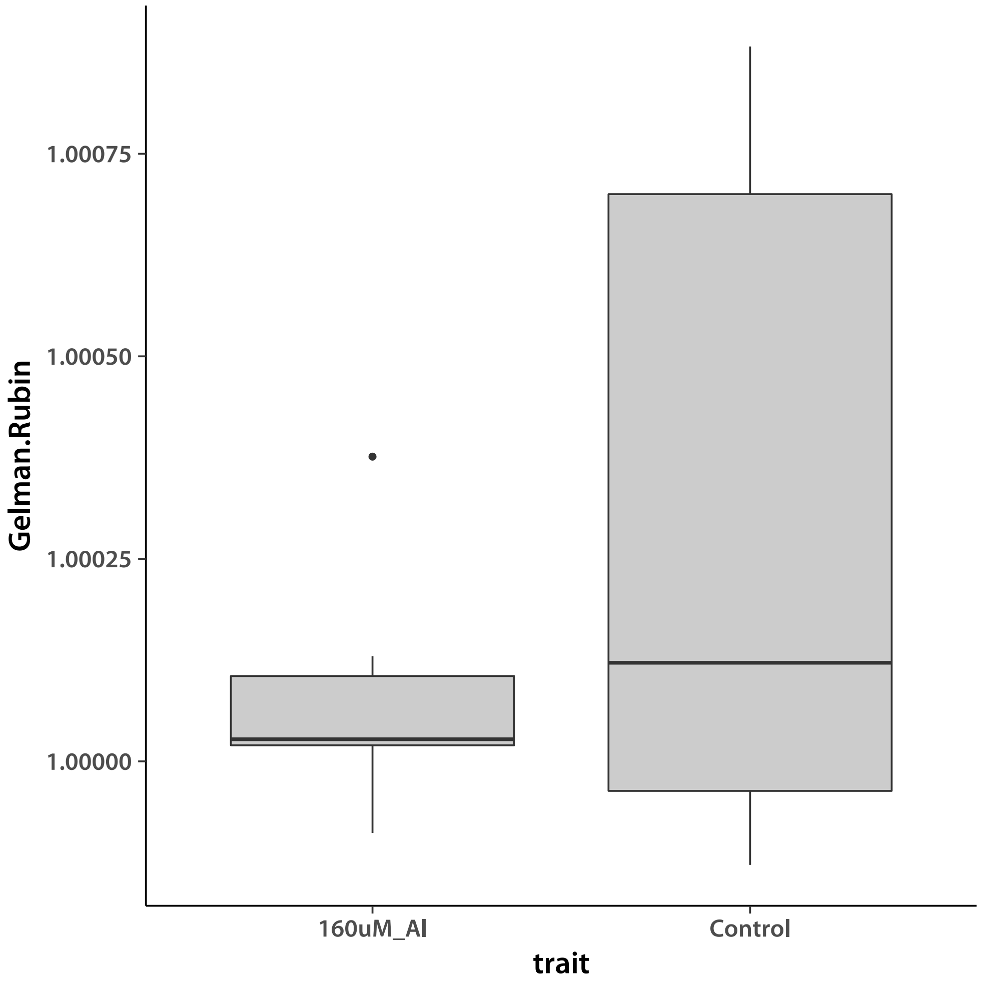


Values smaller than ∼1.2 indicate good convergence. I check by plotting an the chains with the worst G-R statistic for each trait.

```
> plotLocEx <- cmpfun(function(trt.i, samp.j, N){
+     i <- ( d*(samp.j-1) ) + trt.i
+     plot( chn1[,i], type = "l",
+         ylim = range(c(chn1[,i], chn2[,i], chn3[,i], chn4[,i], chn5[,i])),
+         col = "orange", ylab = trtNam[trt.i], xaxt = "n", las = 1)
+     lines(chn2[,i], col = "red")
+     lines(chn3[,i], col = "blue")
+     lines(chn4[,i], col = "green")
+     lines(chn5[,i], col = "magenta")
+ })
```

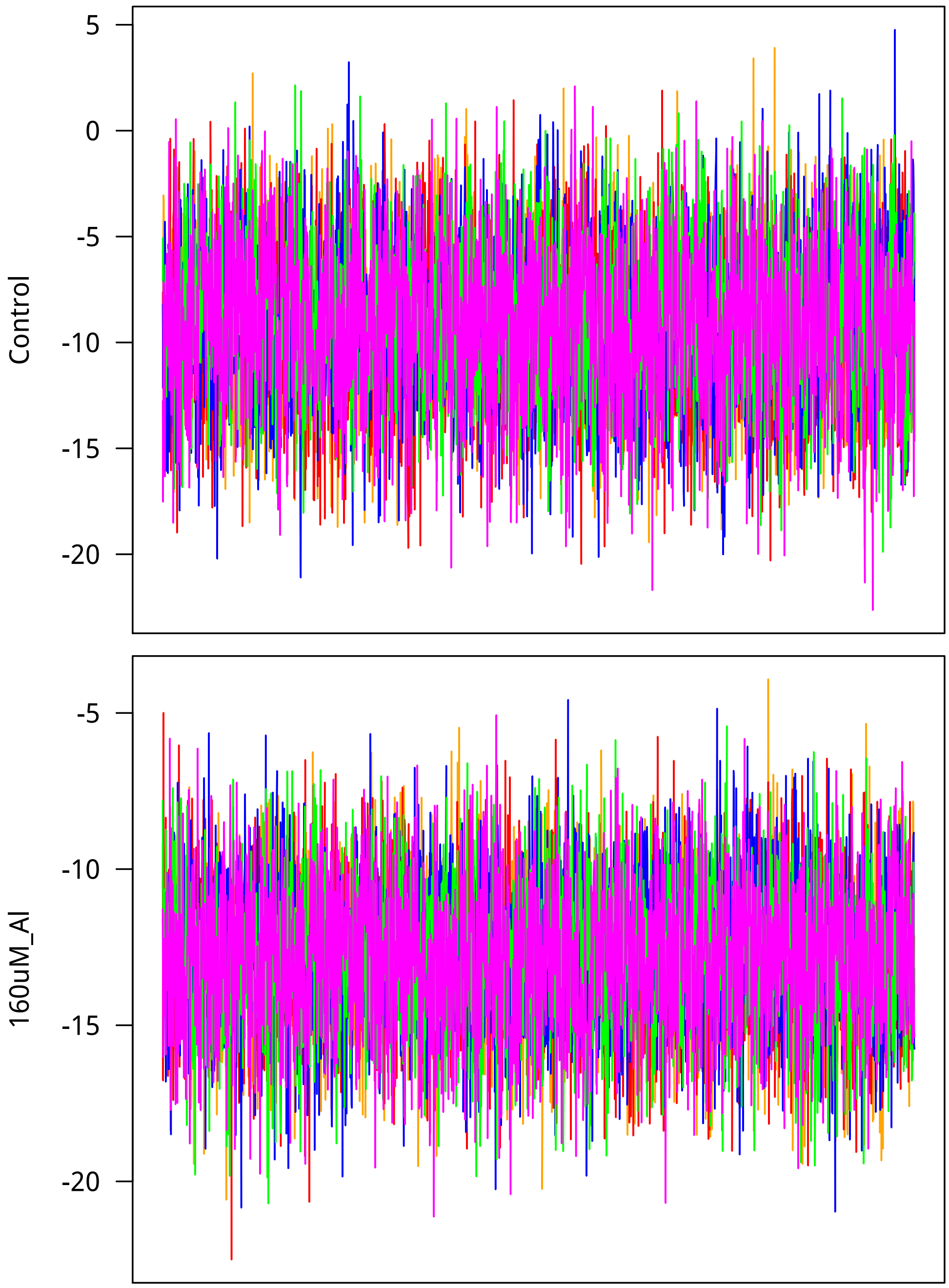


Now I look at mixing. Lag is on the $x$ axis.

```
> ggTrt1 <- ggplot(data=subset(acFrame, trait==trtNam[1]),
+         aes(x=lag, y=autocorrelation)) +
+     geom_boxplot(fill="grey80", aes(group=cut_interval(lag, n=34))) +
+     theme_classic(base_size=18, base_family="myriad") +
+     labs(title=trtNam[1])
> ggTrt2 <- ggplot(data=subset(acFrame, trait==trtNam[2]),
```

```
+         aes(x=lag, y=autocorrelation)) +
+     geom_boxplot(fill="grey80", aes(group=cut_interval(lag, n=34))) +
+     theme_classic(base_size=18, base_family="myriad") +
+     labs(title=trtNam[2])
> showtext_auto()
> grid.arrange(ggTrt1, ggTrt2, nrow = 2)
>
```

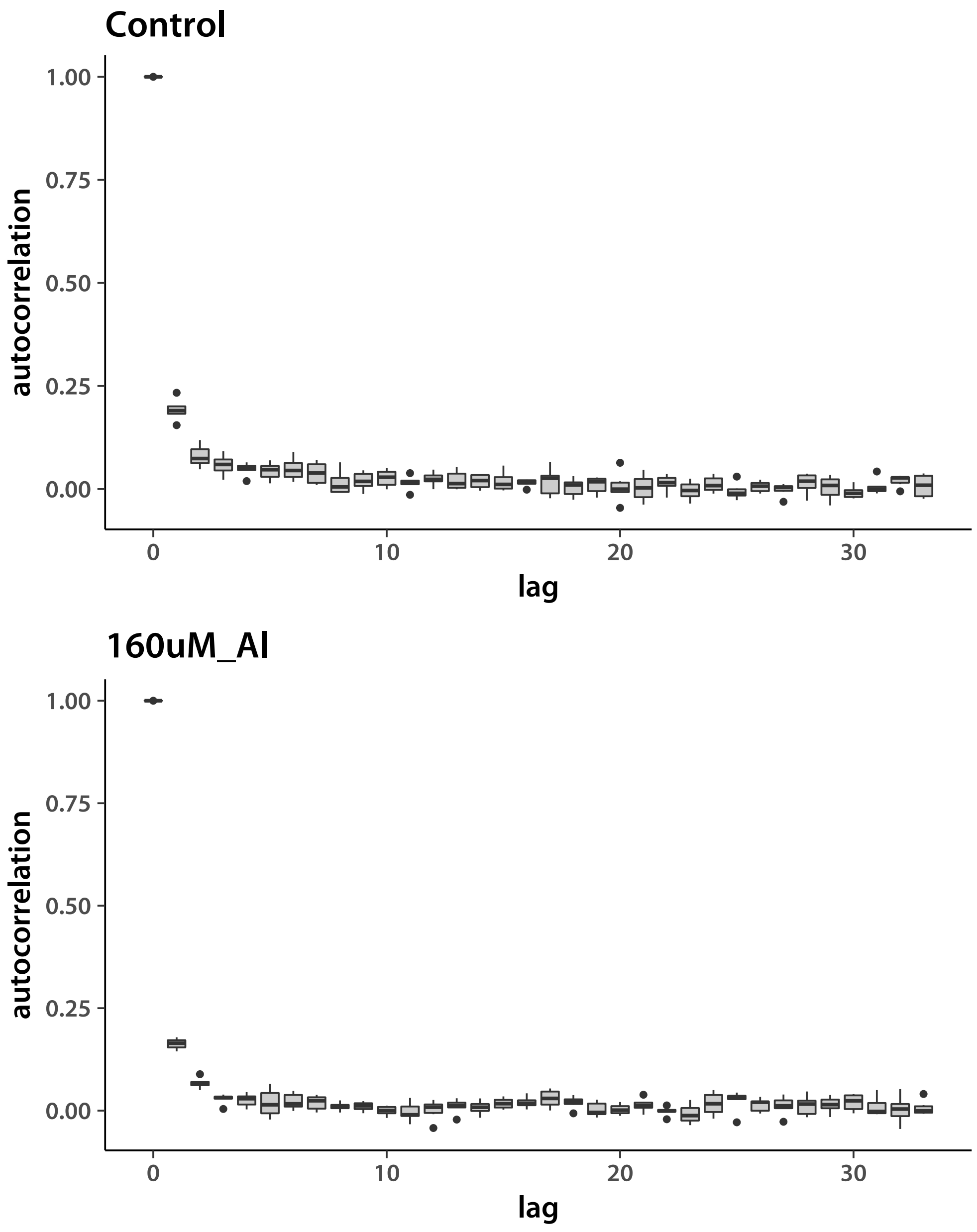


Now do the tropical *japonica* results.

```
> chn1 <- matrix(.C("GSLmatLoad",
```

```
+     "chains/EXPout_trj_3_1.gbin",
+     as.integer(chnLen), as.integer(expDim), out = double(chnLen*expDim))$out,
+         nrow = chnLen, byrow = T)
> chn2 <- matrix(.C("GSLmatLoad",
+     "chains/EXPout_trj_3_2.gbin",
+     as.integer(chnLen), as.integer(expDim), out = double(chnLen*expDim))$out,
+         nrow = chnLen, byrow = T)
> chn3 <- matrix(.C("GSLmatLoad",
+     "chains/EXPout_trj_3_3.gbin",
+     as.integer(chnLen), as.integer(expDim), out = double(chnLen*expDim))$out,
+         nrow = chnLen, byrow = T)
> chn4 <- matrix(.C("GSLmatLoad",
+     "chains/EXPout_trj_3_4.gbin",
+     as.integer(chnLen), as.integer(expDim), out = double(chnLen*expDim))$out,
+         nrow = chnLen, byrow = T)
> chn5 <- matrix(.C("GSLmatLoad",
+     "chains/EXPout_trj_3_5.gbin",
+     as.integer(chnLen), as.integer(expDim), out = double(chnLen*expDim))$out,
+         nrow = chnLen, byrow = T)
> grMat    <- matrix(gelRub("chn", nChn, chnLen), ncol = d, byrow = T)
> grFrame  <- data.frame(Gelman.Rubin=c(grMat), trait=rep(trtNam, each = Nexp))
> grMaxInd <- apply(grMat, 2, function(vec){which(vec == max(vec))[1]})
> acMat    <- mcmcAcf("chn", 5)
> acFrame  <- data.frame(autocorrelation=c(acMat), trait=rep(trtNam, prod(dim(acMat))/2),
+     lag=rep(0:33, each = nrow(acMat)))
>

> showtext_auto()
> ggplot(data=grFrame,
+         aes(x=trait, y=Gelman.Rubin)) +
+     geom_boxplot(fill="grey80") +
+     theme_classic(base_size=18, base_family="myriad")
>
```

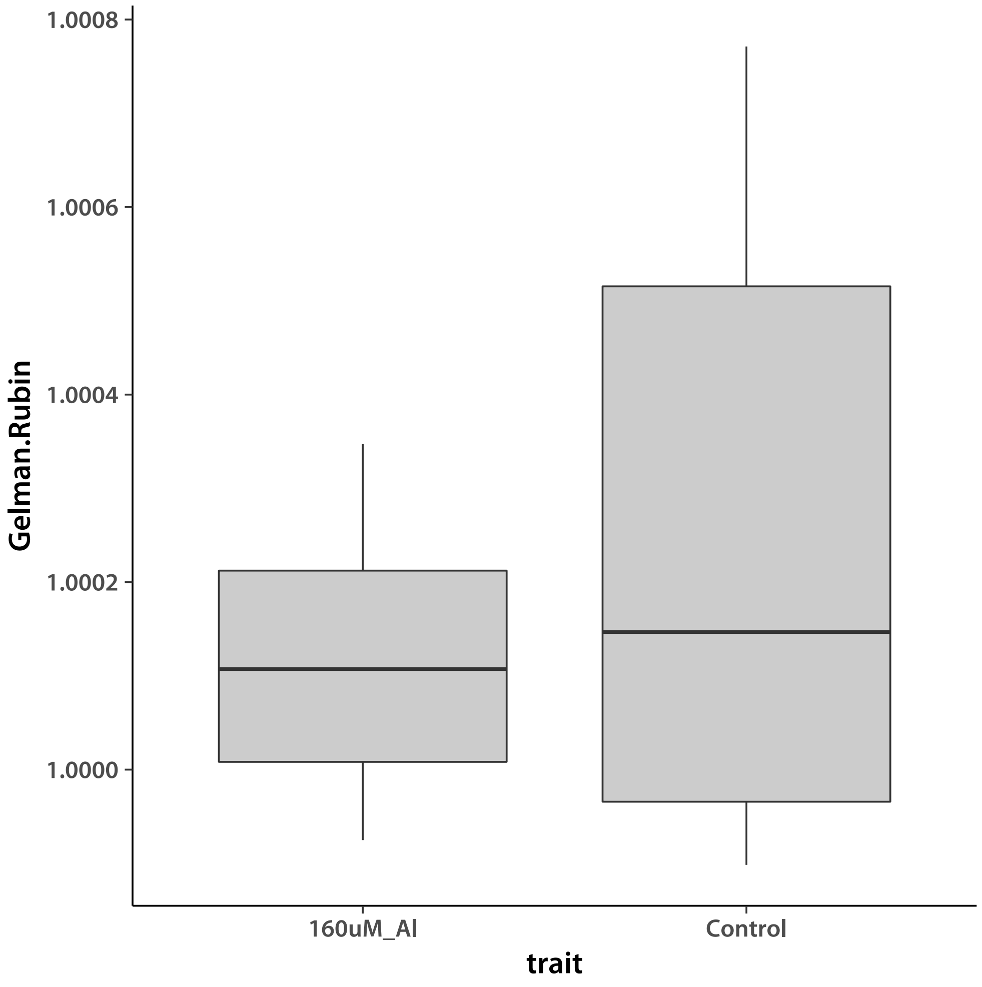


Values smaller than ~1.2 indicate good convergence. I check by plotting an the chains with the worst G-R statistic for each trait.

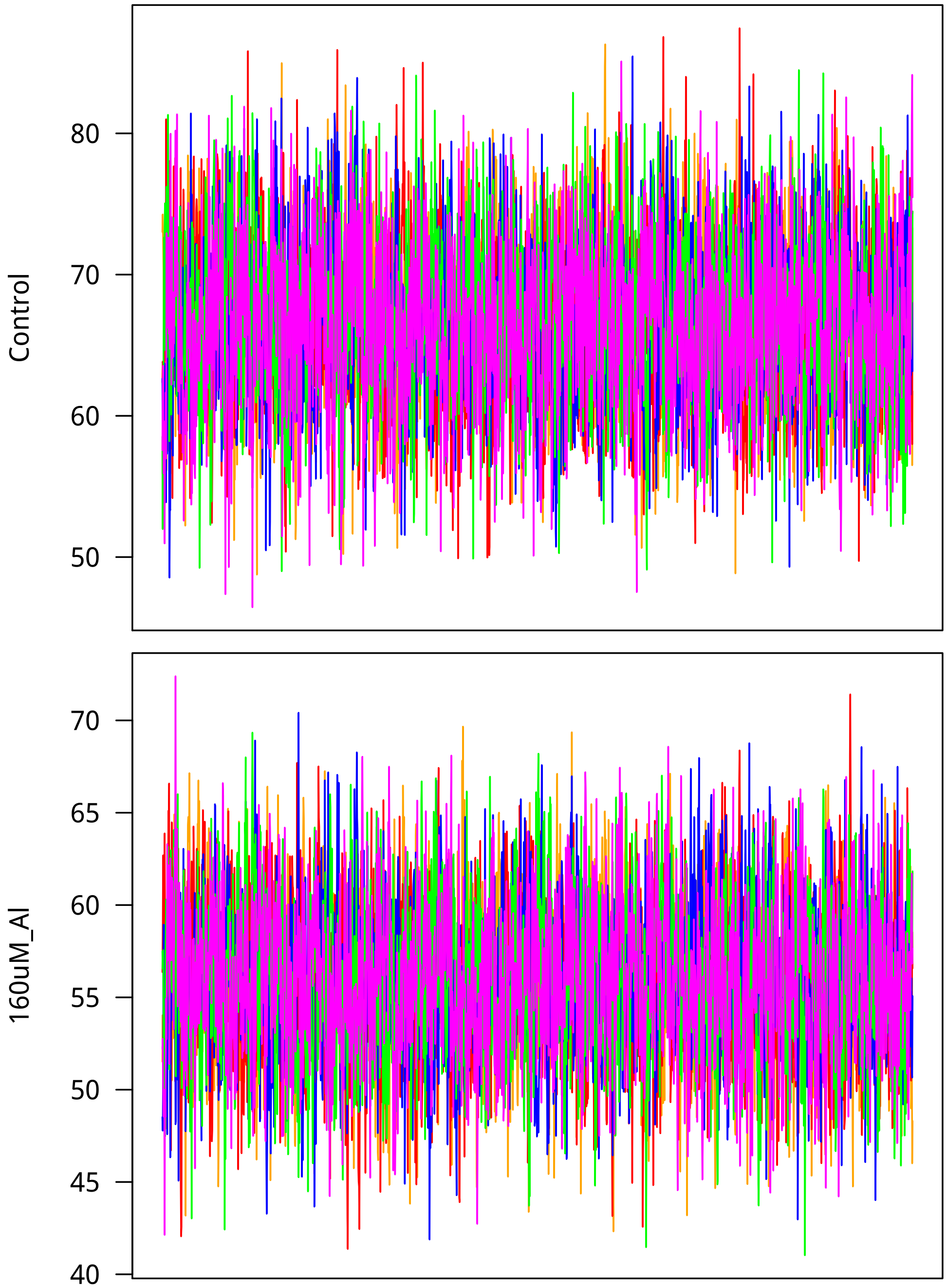


Now I look at mixing. Lag is on the $x$ axis.

```
> ggTrt1 <- ggplot(data=subset(acFrame, trait==trtNam[1]),
+         aes(x=lag, y=autocorrelation)) +
+     geom_boxplot(fill="grey80", aes(group=cut_interval(lag, n=34))) +
+     theme_classic(base_size=18, base_family="myriad") +
+     labs(title=trtNam[1])
> ggTrt2 <- ggplot(data=subset(acFrame, trait==trtNam[2]),
```

```
+         aes(x=lag, y=autocorrelation)) +
+     geom_boxplot(fill="grey80", aes(group=cut_interval(lag, n=34))) +
+     theme_classic(base_size=18, base_family="myriad") +
+     labs(title=trtNam[2])
> showtext_auto()
> grid.arrange(ggTrt1, ggTrt2, nrow = 2)
>
```

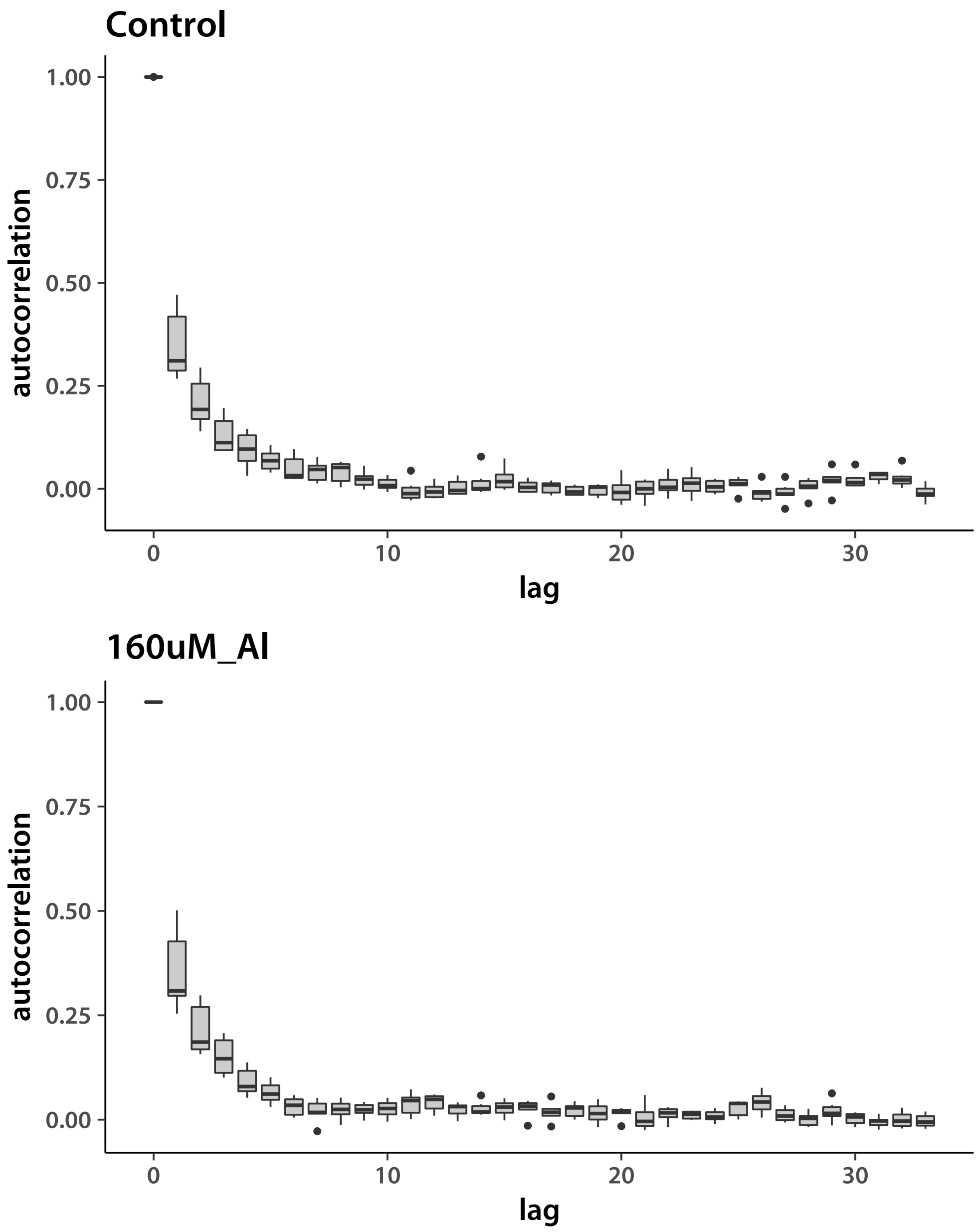


Next, tropical *japonica* with admixture.

```
> chn1 <- matrix(.C("GSLmatLoad",
```

```
+     "chains/EXPout_trja_3_1.gbin",
+     as.integer(chnLen), as.integer(expDim), out = double(chnLen*expDim))$out,
+         nrow = chnLen, byrow = T)
> chn2 <- matrix(.C("GSLmatLoad",
+     "chains/EXPout_trja_3_2.gbin",
+     as.integer(chnLen), as.integer(expDim), out = double(chnLen*expDim))$out,
+         nrow = chnLen, byrow = T)
> chn3 <- matrix(.C("GSLmatLoad",
+     "chains/EXPout_trja_3_3.gbin",
+     as.integer(chnLen), as.integer(expDim), out = double(chnLen*expDim))$out,
+         nrow = chnLen, byrow = T)
> chn4 <- matrix(.C("GSLmatLoad",
+     "chains/EXPout_trja_3_4.gbin",
+     as.integer(chnLen), as.integer(expDim), out = double(chnLen*expDim))$out,
+         nrow = chnLen, byrow = T)
> chn5 <- matrix(.C("GSLmatLoad",
+     "chains/EXPout_trja_3_5.gbin",
+     as.integer(chnLen), as.integer(expDim), out = double(chnLen*expDim))$out,
+         nrow = chnLen, byrow = T)
> grMat    <- matrix(gelRub("chn", nChn, chnLen), ncol = d, byrow = T)
> grFrame  <- data.frame(Gelman.Rubin=c(grMat), trait=rep(trtNam, each = Nexp))
> grMaxInd <- apply(grMat, 2, function(vec){which(vec == max(vec))[1]})
> acMat    <- mcmcAcf("chn", 5)
> acFrame  <- data.frame(autocorrelation=c(acMat), trait=rep(trtNam, prod(dim(acMat))/2),
+     lag=rep(0:33, each = nrow(acMat)))
>

> showtext_auto()
> ggplot(data=grFrame,
+         aes(x=trait, y=Gelman.Rubin)) +
+     geom_boxplot(fill="grey80") +
+     theme_classic(base_size=18, base_family="myriad")
>
```

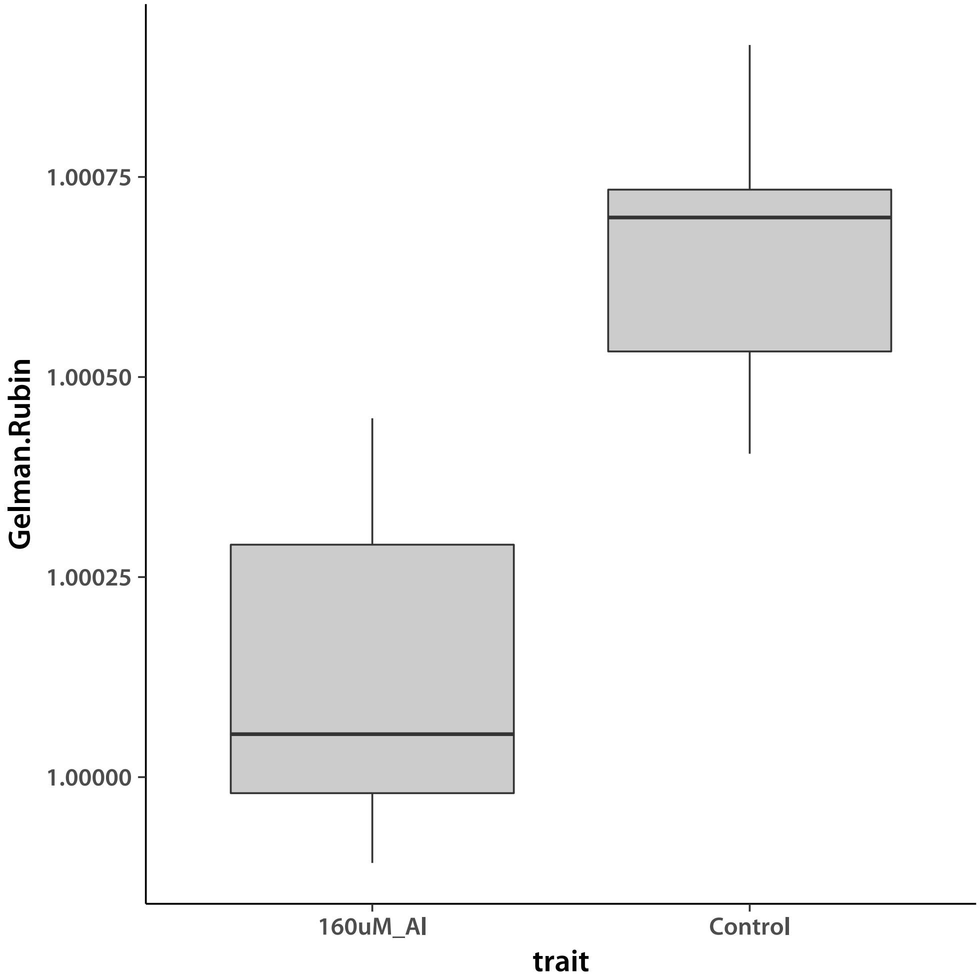


Values smaller than ∼1.2 indicate good convergence. I check by plotting an the chains with the worst G-R statistic for each trait.

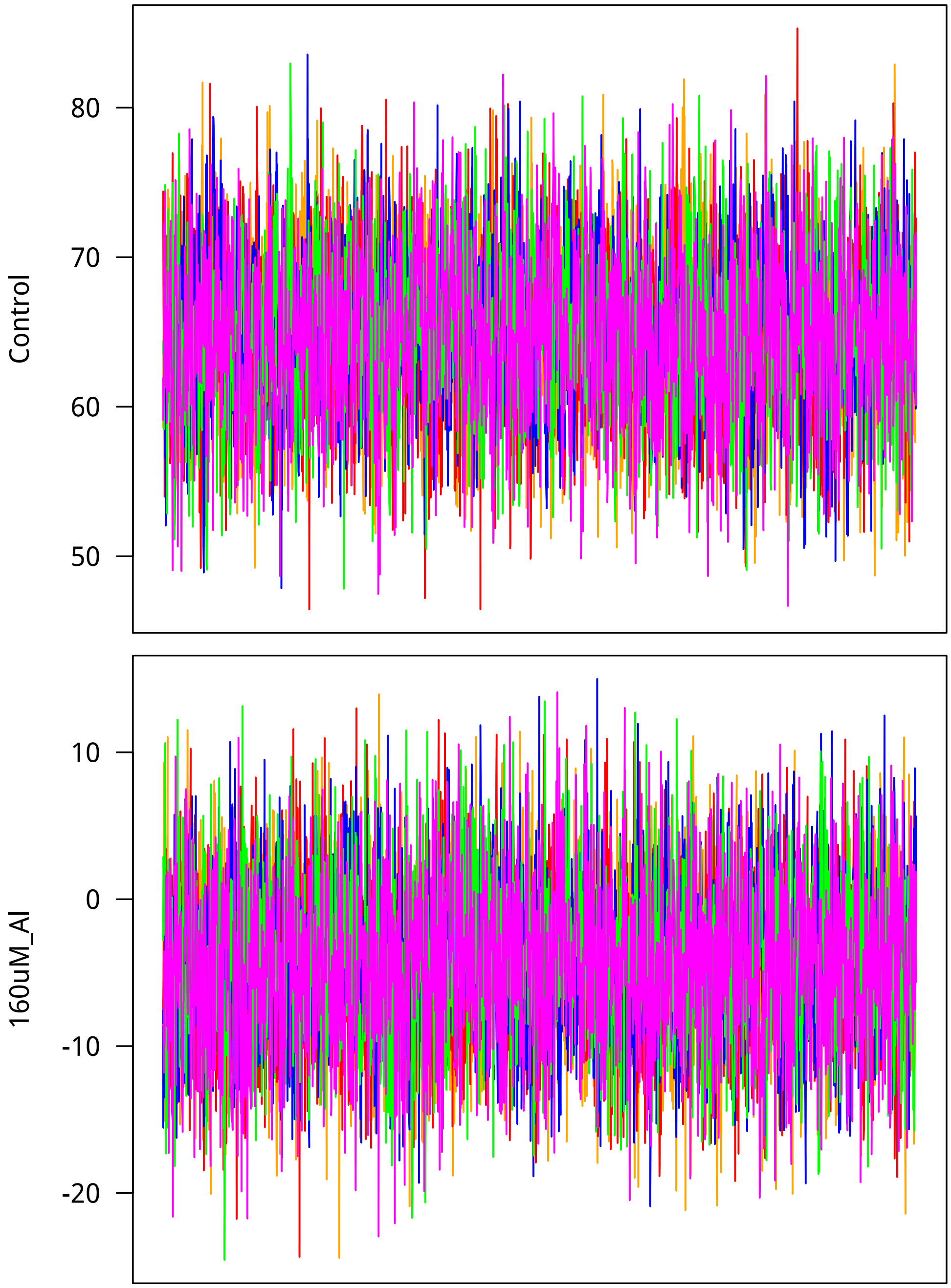


Now I look at mixing. Lag is on the $x$ axis.

```
> ggTrt1 <- ggplot(data=subset(acFrame, trait==trtNam[1]),
+          aes(x=lag, y=autocorrelation)) +
+     geom_boxplot(fill="grey80", aes(group=cut_interval(lag, n=34))) +
+     theme_classic(base_size=18, base_family="myriad") +
+     labs(title=trtNam[1])
> ggTrt2 <- ggplot(data=subset(acFrame, trait==trtNam[2]),
```

```
+         aes(x=lag, y=autocorrelation)) +
+     geom_boxplot(fill="grey80", aes(group=cut_interval(lag, n=34))) +
+     theme_classic(base_size=18, base_family="myriad") +
+     labs(title=trtNam[2])
> showtext_auto()
> grid.arrange(ggTrt1, ggTrt2, nrow = 2)
>
```

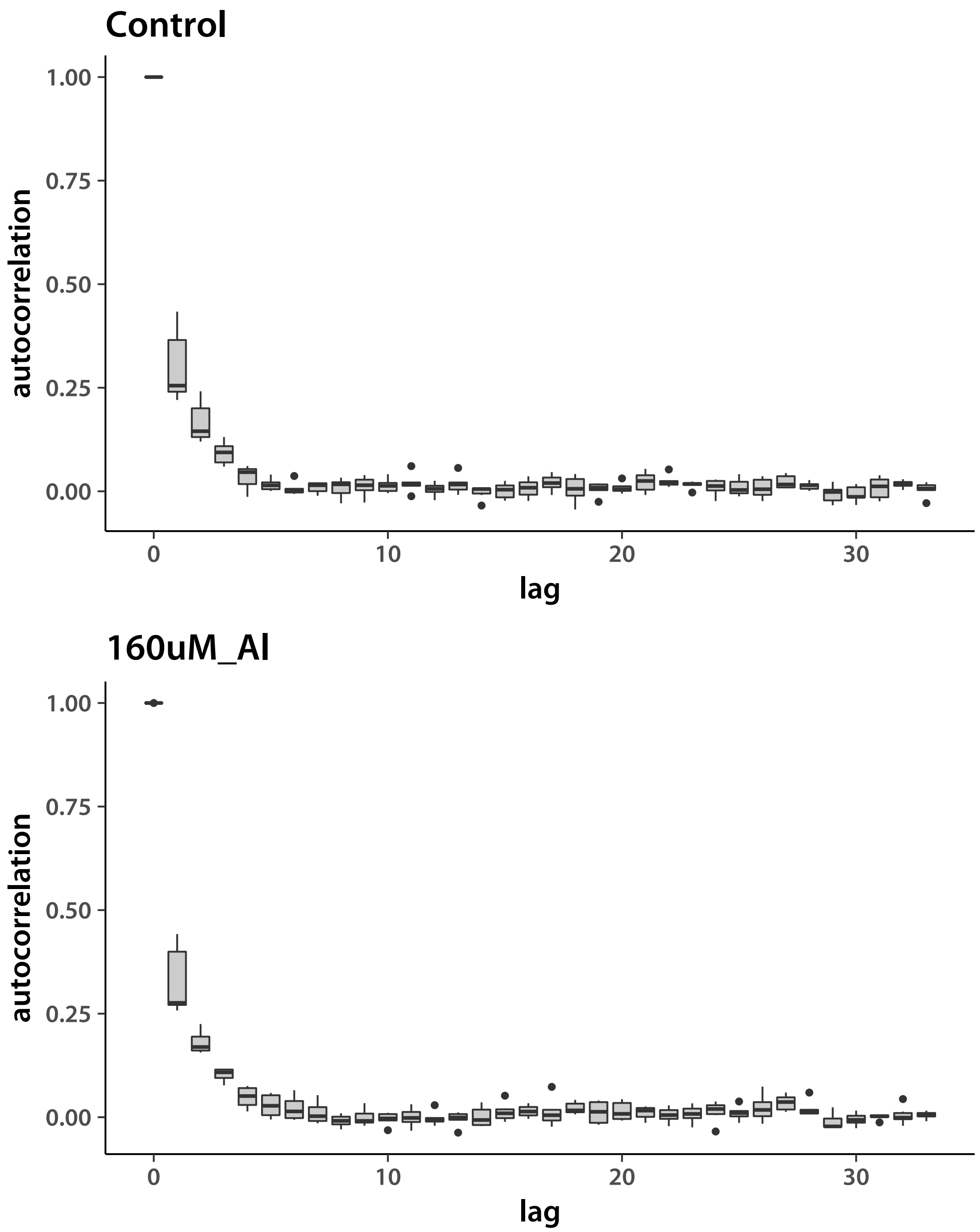


Everything looks good. I next look at the accession means, *aus* first.

```
> Nacc   <- 55
```

```
> locDim <- d*Nacc
> chn1 <- matrix(.C("GSLmatLoad",
+     "chains/LNout_aus_3_1.gbin",
+     as.integer(chnLen), as.integer(locDim), out = double(chnLen*locDim))$out,
+         nrow = chnLen, byrow = T)
> chn2 <- matrix(.C("GSLmatLoad",
+     "chains/LNout_aus_3_2.gbin",
+     as.integer(chnLen), as.integer(locDim), out = double(chnLen*locDim))$out,
+         nrow = chnLen, byrow = T)
> chn3 <- matrix(.C("GSLmatLoad",
+     "chains/LNout_aus_3_3.gbin",
+     as.integer(chnLen), as.integer(locDim), out = double(chnLen*locDim))$out,
+         nrow = chnLen, byrow = T)
> chn4 <- matrix(.C("GSLmatLoad",
+     "chains/LNout_aus_3_4.gbin",
+     as.integer(chnLen), as.integer(locDim), out = double(chnLen*locDim))$out,
+         nrow = chnLen, byrow = T)
> chn5 <- matrix(.C("GSLmatLoad",
+     "chains/LNout_aus_3_5.gbin",
+     as.integer(chnLen), as.integer(locDim), out = double(chnLen*locDim))$out,
+         nrow = chnLen, byrow = T)
> grMat    <- matrix(gelRub("chn", nChn, chnLen), ncol = d, byrow = T)
> grFrame  <- data.frame(Gelman.Rubin=c(grMat), trait=rep(trtNam, each = Nacc))
> grMaxInd <- apply(grMat, 2, function(vec){which(vec == max(vec))[1]})
> acMat    <- mcmcAcf("chn", 5)
> acFrame  <- data.frame(autocorrelation=c(acMat), trait=rep(trtNam, prod(dim(acMat))/2),
+     lag=rep(0:33, each = nrow(acMat)))
>
```

Plot the *aus* results.

```
> showtext_auto()
> ggplot(data=grFrame,
+         aes(x=trait, y=Gelman.Rubin)) +
+     geom_boxplot(fill="grey80") +
+     theme_classic(base_size=18, base_family="myriad")
>
```

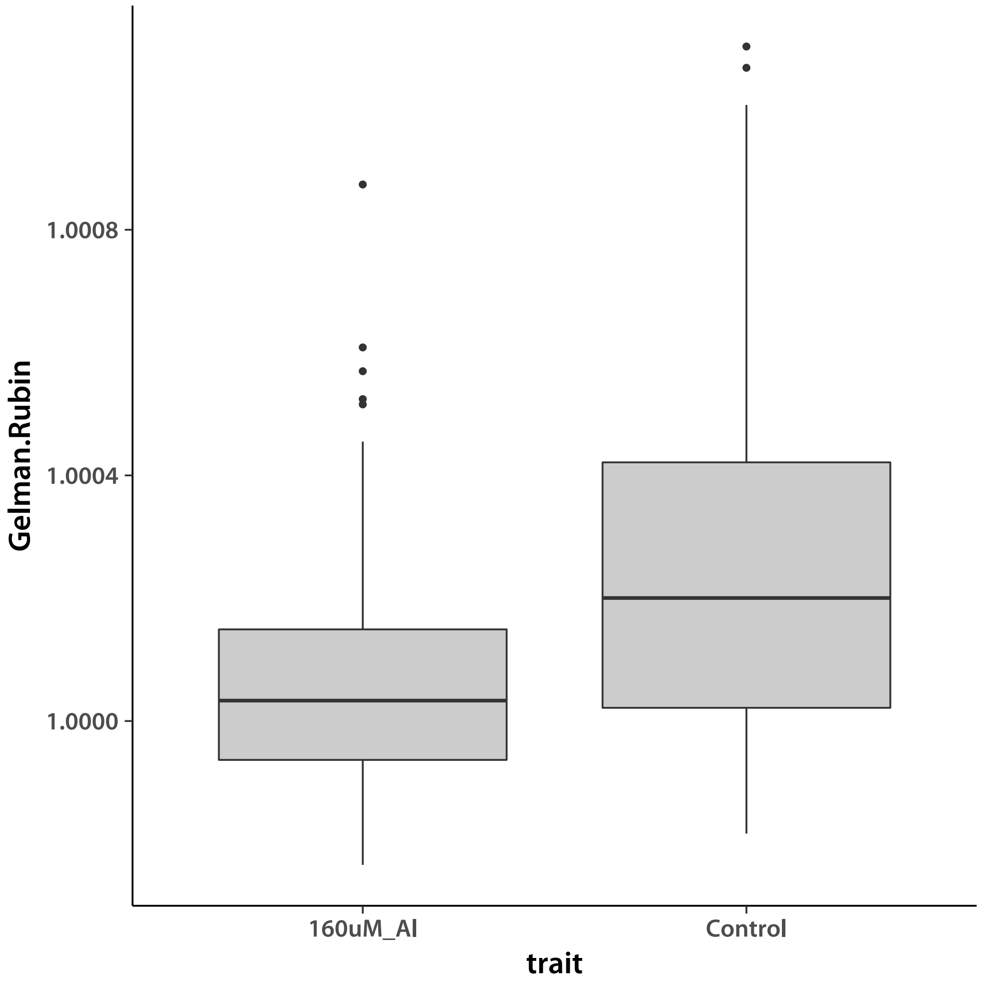


Values smaller than ∼1.2 indicate good convergence. I check by plotting an the chains with the worst G-R statistic for each trait.

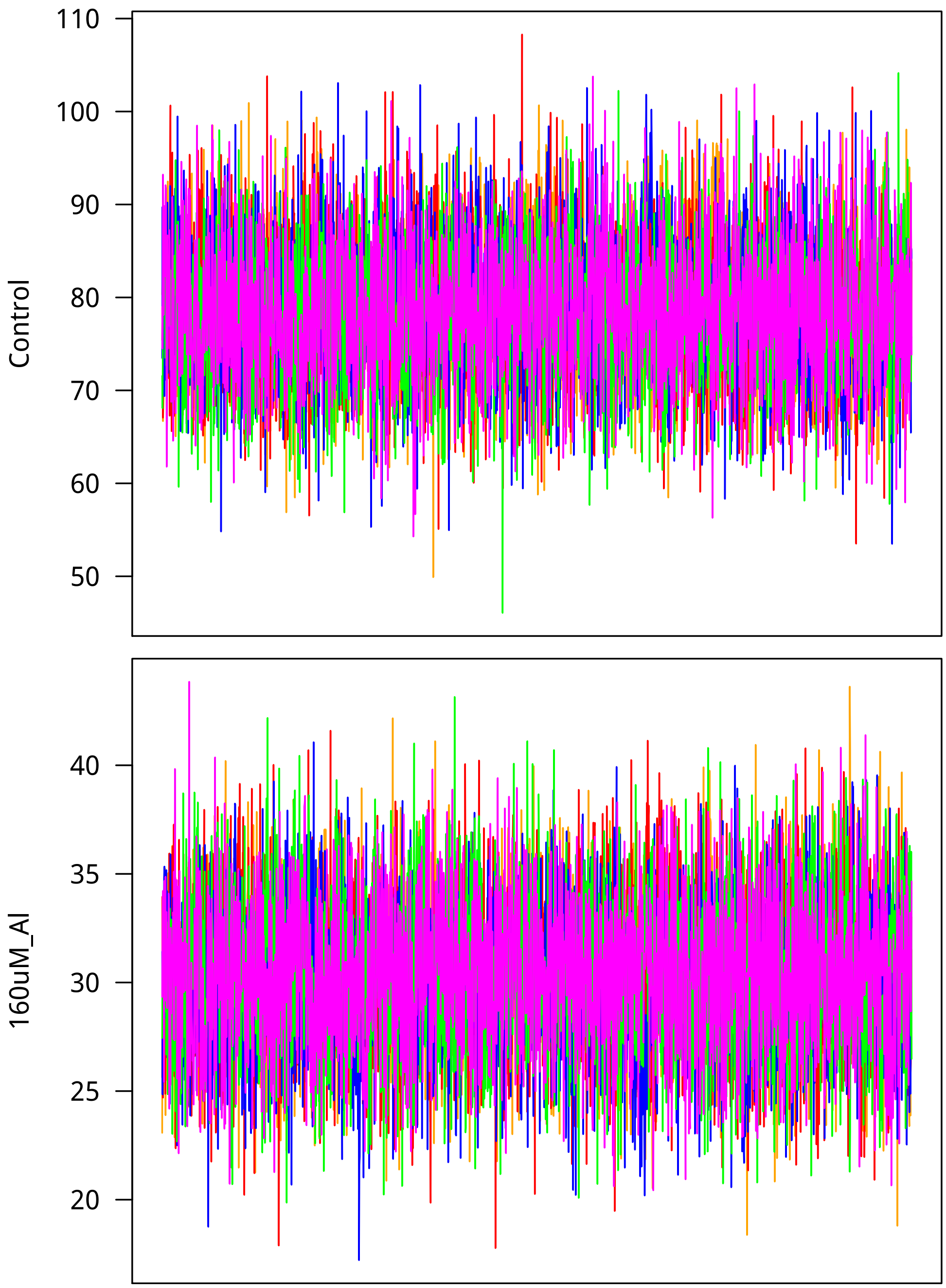


Now I look at mixing. Lag is on the $x$ axis.

```
> ggTrt1 <- ggplot(data=subset(acFrame, trait==trtNam[1]),
+         aes(x=lag, y=autocorrelation)) +
+     geom_boxplot(fill="grey80", aes(group=cut_interval(lag, n=34))) +
+     theme_classic(base_size=18, base_family="myriad") +
+     labs(title=trtNam[1])
> ggTrt2 <- ggplot(data=subset(acFrame, trait==trtNam[2]),
```

```
+         aes(x=lag, y=autocorrelation)) +
+     geom_boxplot(fill="grey80", aes(group=cut_interval(lag, n=34))) +
+     theme_classic(base_size=18, base_family="myriad") +
+     labs(title=trtNam[2])
> showtext_auto()
> grid.arrange(ggTrt1, ggTrt2, nrow = 2)
>
```

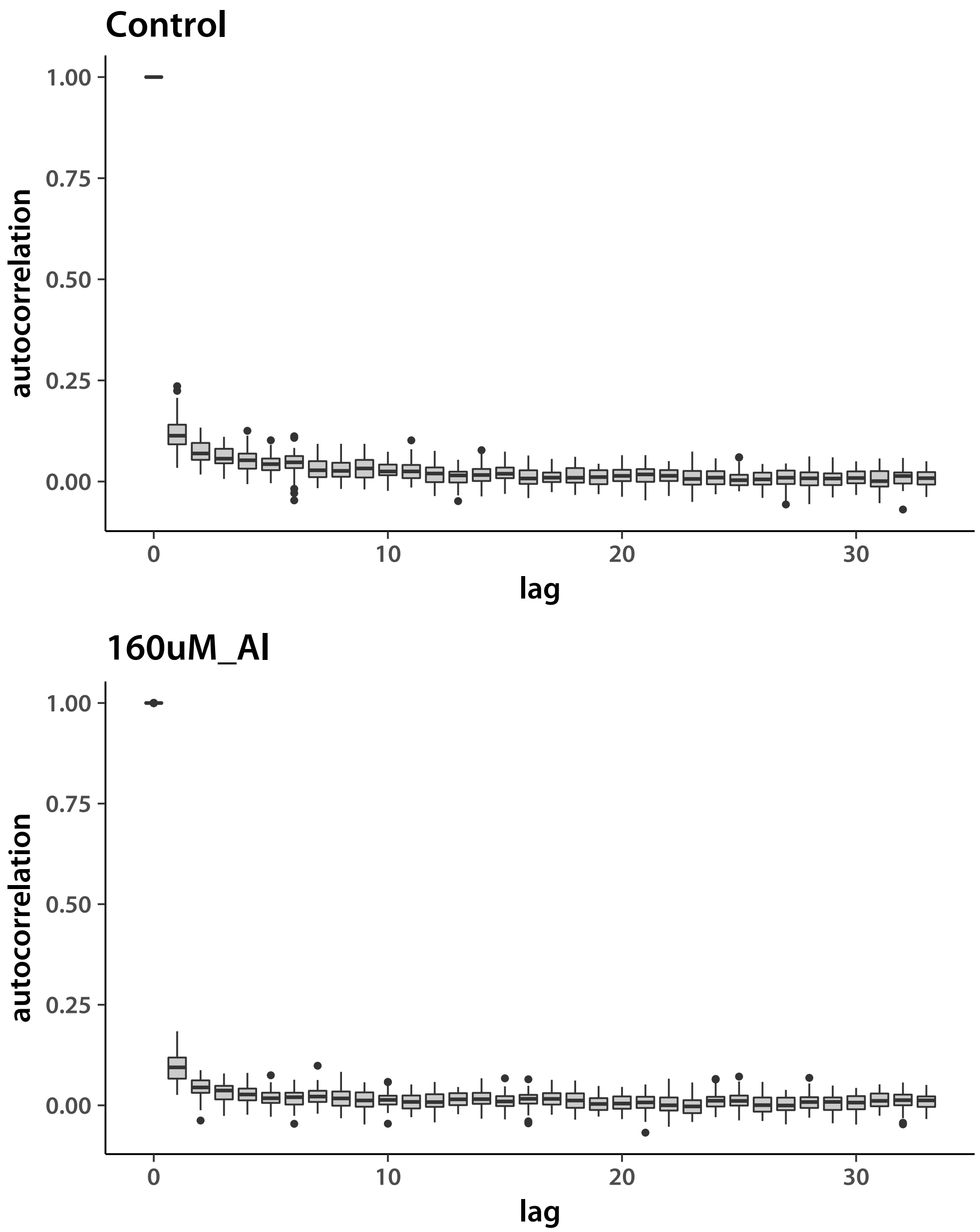


Looks good. Now do the tropical *japonica*.

```
> Nacc   <- 92
```

```
> locDim <- d*Nacc
> chn1 <- matrix(.C("GSLmatLoad",
+     "chains/LNout_trj_3_1.gbin",
+     as.integer(chnLen), as.integer(locDim), out = double(chnLen*locDim))$out,
+         nrow = chnLen, byrow = T)
> chn2 <- matrix(.C("GSLmatLoad",
+     "chains/LNout_trj_3_2.gbin",
+     as.integer(chnLen), as.integer(locDim), out = double(chnLen*locDim))$out,
+         nrow = chnLen, byrow = T)
> chn3 <- matrix(.C("GSLmatLoad",
+     "chains/LNout_trj_3_3.gbin",
+     as.integer(chnLen), as.integer(locDim), out = double(chnLen*locDim))$out,
+         nrow = chnLen, byrow = T)
> chn4 <- matrix(.C("GSLmatLoad",
+     "chains/LNout_trj_3_4.gbin",
+     as.integer(chnLen), as.integer(locDim), out = double(chnLen*locDim))$out,
+         nrow = chnLen, byrow = T)
> chn5 <- matrix(.C("GSLmatLoad",
+     "chains/LNout_trj_3_5.gbin",
+     as.integer(chnLen), as.integer(locDim), out = double(chnLen*locDim))$out,
+         nrow = chnLen, byrow = T)
> grMat    <- matrix(gelRub("chn", nChn, chnLen), ncol = d, byrow = T)
> grFrame  <- data.frame(Gelman.Rubin=c(grMat), trait=rep(trtNam, each = Nacc))
> grMaxInd <- apply(grMat, 2, function(vec){which(vec == max(vec))[1]})
> acMat    <- mcmcAcf("chn", 5)
> acFrame  <- data.frame(autocorrelation=c(acMat), trait=rep(trtNam, prod(dim(acMat))/2),
+     lag=rep(0:33, each = nrow(acMat)))
>
```

Plot the *aus* results.

```
> showtext_auto()
> ggplot(data=grFrame,
+         aes(x=trait, y=Gelman.Rubin)) +
+     geom_boxplot(fill="grey80") +
+     theme_classic(base_size=18, base_family="myriad")
>
```

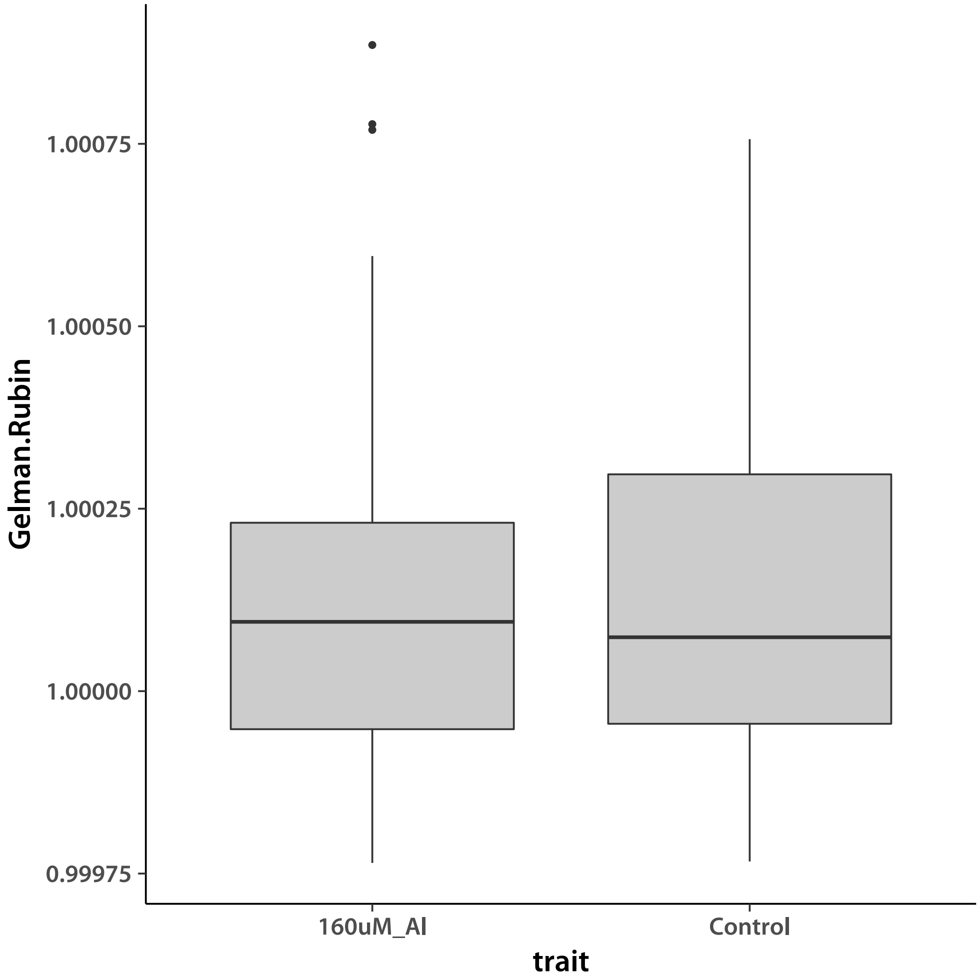


Values smaller than ∼1.2 indicate good convergence. I check by plotting an the chains with the worst G-R statistic for each trait.

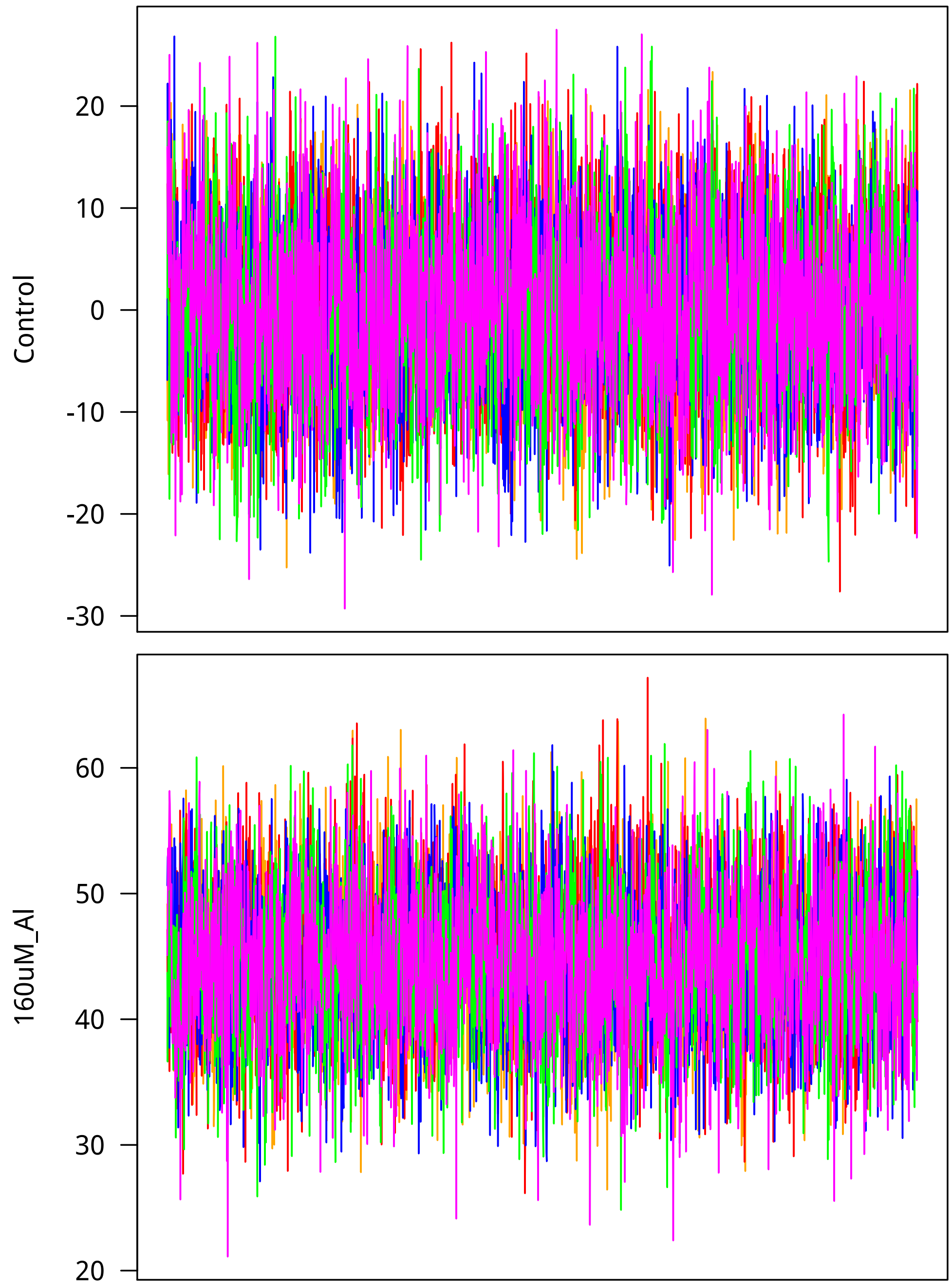


Now I look at mixing. Lag is on the $x$ axis.

```
> ggTrt1 <- ggplot(data=subset(acFrame, trait==trtNam[1]),
+         aes(x=lag, y=autocorrelation)) +
+     geom_boxplot(fill="grey80", aes(group=cut_interval(lag, n=34))) +
+     theme_classic(base_size=18, base_family="myriad") +
+     labs(title=trtNam[1])
> ggTrt2 <- ggplot(data=subset(acFrame, trait==trtNam[2]),
```

```
+         aes(x=lag, y=autocorrelation)) +
+     geom_boxplot(fill="grey80", aes(group=cut_interval(lag, n=34))) +
+     theme_classic(base_size=18, base_family="myriad") +
+     labs(title=trtNam[2])
> showtext_auto()
> grid.arrange(ggTrt1, ggTrt2, nrow = 2)
>
```

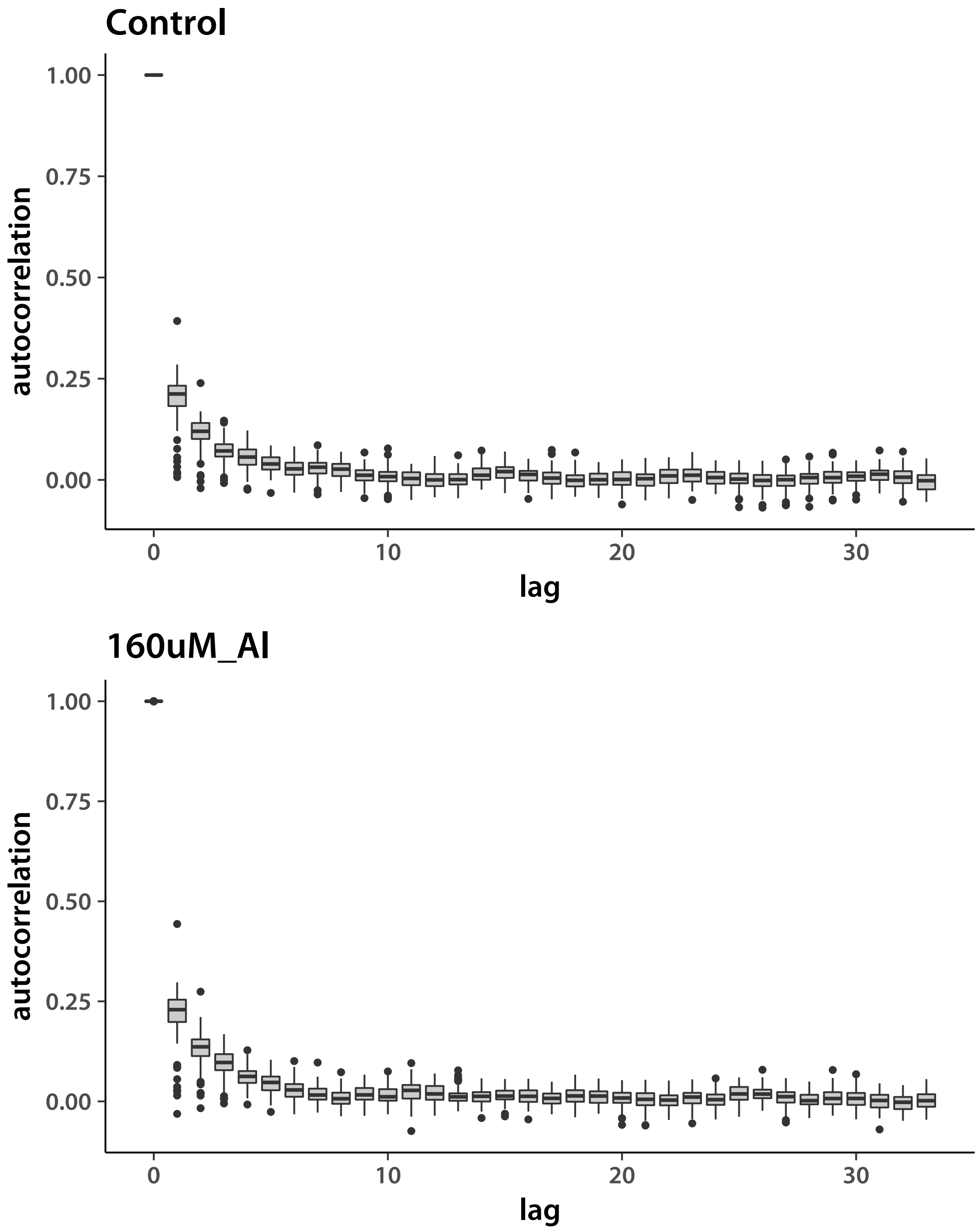


Next, admixed tropical *japonica*.

```
> Nacc   <- 95
```

```
> locDim <- d*Nacc
> chn1 <- matrix(.C("GSLmatLoad",
+     "chains/LNout_trja_3_1.gbin",
+     as.integer(chnLen), as.integer(locDim), out = double(chnLen*locDim))$out,
+         nrow = chnLen, byrow = T)
> chn2 <- matrix(.C("GSLmatLoad",
+     "chains/LNout_trja_3_2.gbin",
+     as.integer(chnLen), as.integer(locDim), out = double(chnLen*locDim))$out,
+         nrow = chnLen, byrow = T)
> chn3 <- matrix(.C("GSLmatLoad",
+     "chains/LNout_trja_3_3.gbin",
+     as.integer(chnLen), as.integer(locDim), out = double(chnLen*locDim))$out,
+         nrow = chnLen, byrow = T)
> chn4 <- matrix(.C("GSLmatLoad",
+     "chains/LNout_trja_3_4.gbin",
+     as.integer(chnLen), as.integer(locDim), out = double(chnLen*locDim))$out,
+         nrow = chnLen, byrow = T)
> chn5 <- matrix(.C("GSLmatLoad",
+     "chains/LNout_trja_3_5.gbin",
+     as.integer(chnLen), as.integer(locDim), out = double(chnLen*locDim))$out,
+         nrow = chnLen, byrow = T)
> grMat    <- matrix(gelRub("chn", nChn, chnLen), ncol = d, byrow = T)
> grFrame  <- data.frame(Gelman.Rubin=c(grMat), trait=rep(trtNam, each = Nacc))
> grMaxInd <- apply(grMat, 2, function(vec){which(vec == max(vec))[1]})
> acMat    <- mcmcAcf("chn", 5)
> acFrame  <- data.frame(autocorrelation=c(acMat), trait=rep(trtNam, prod(dim(acMat))/2),
+     lag=rep(0:33, each = nrow(acMat)))
>
```

Plot the *aus* results.

```
> showtext_auto()
> ggplot(data=grFrame,
+         aes(x=trait, y=Gelman.Rubin)) +
+     geom_boxplot(fill="grey80") +
+     theme_classic(base_size=18, base_family="myriad")
>
```

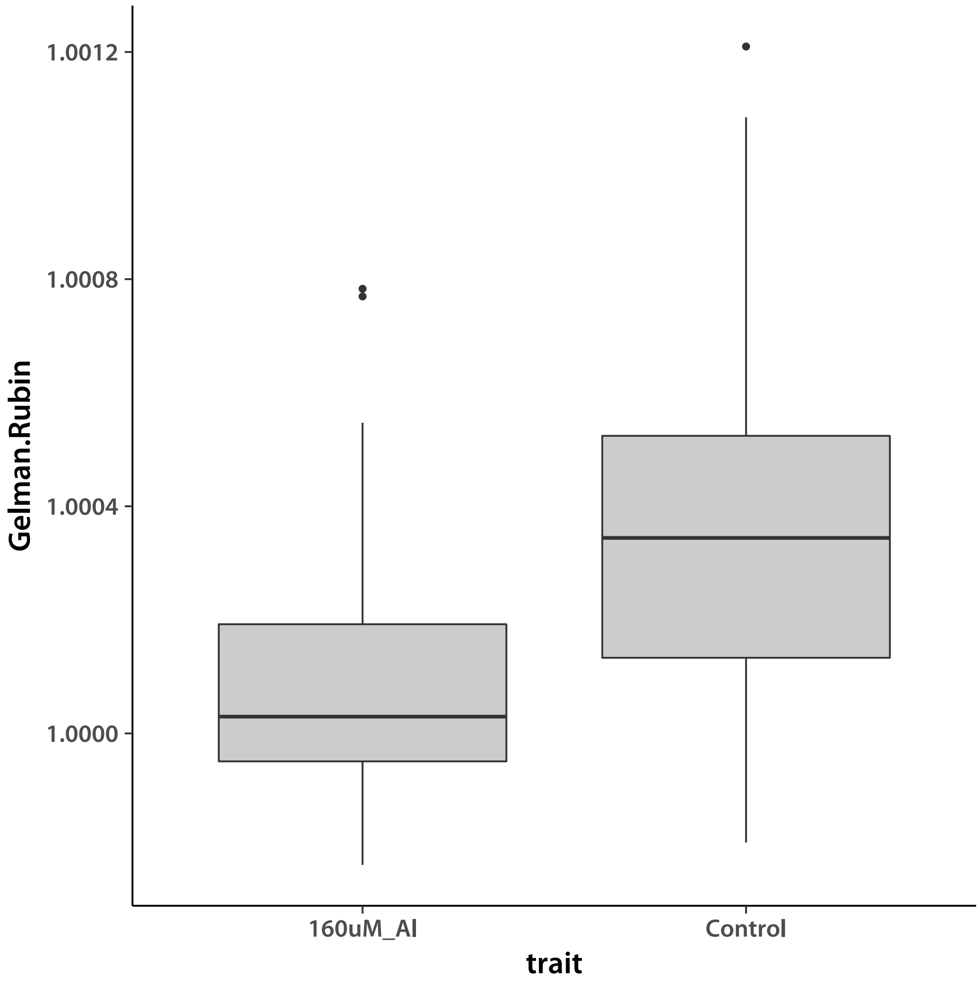


Values smaller than ∼1.2 indicate good convergence. I check by plotting an the chains with the worst G-R statistic for each trait.

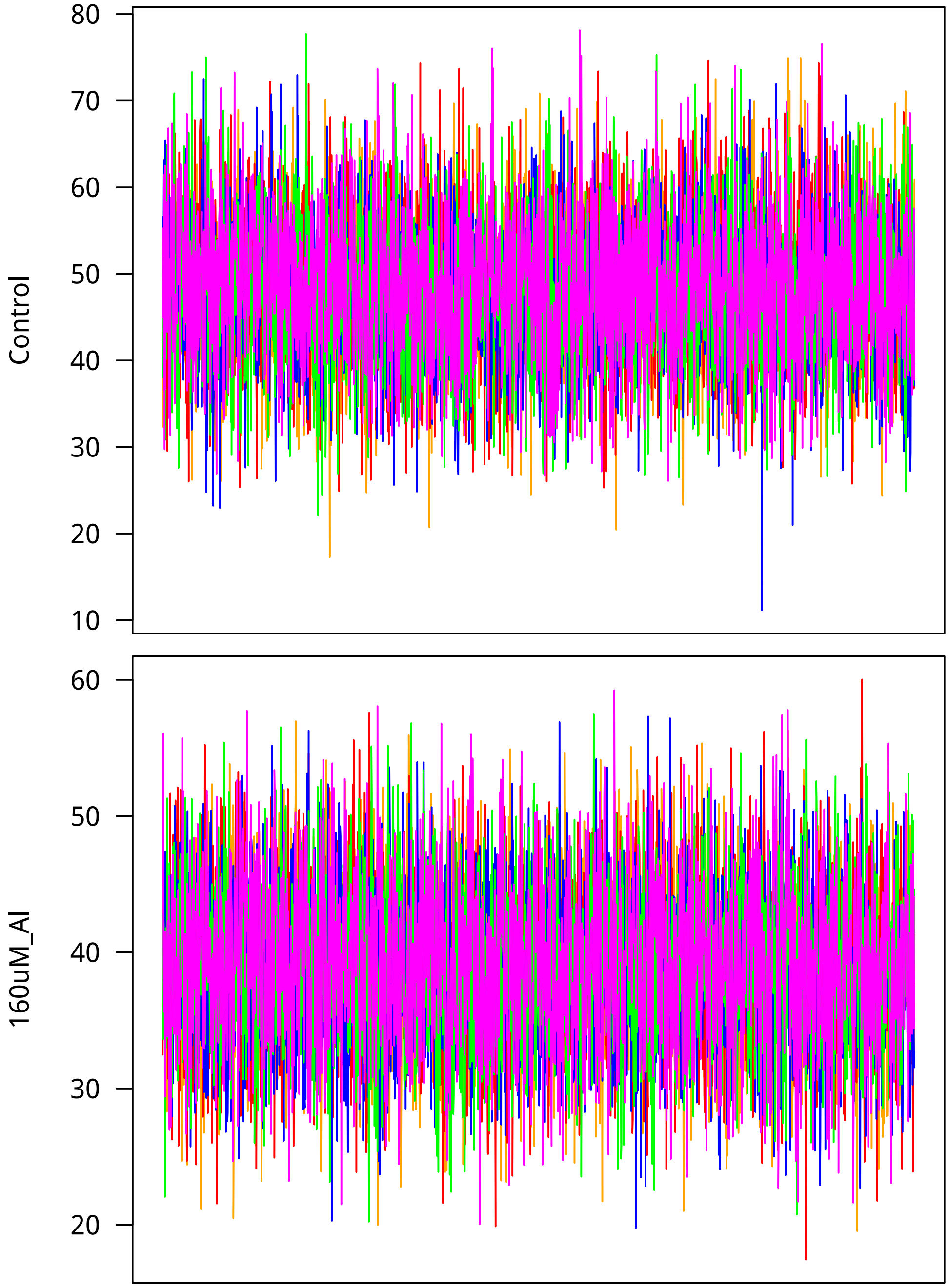


Now I look at mixing. Lag is on the $x$ axis.

```
> ggTrt1 <- ggplot(data=subset(acFrame, trait==trtNam[1]),
+         aes(x=lag, y=autocorrelation)) +
+     geom_boxplot(fill="grey80", aes(group=cut_interval(lag, n=34))) +
+     theme_classic(base_size=18, base_family="myriad") +
+     labs(title=trtNam[1])
> ggTrt2 <- ggplot(data=subset(acFrame, trait==trtNam[2]),
```

```
+         aes(x=lag, y=autocorrelation)) +
+     geom_boxplot(fill="grey80", aes(group=cut_interval(lag, n=34))) +
+     theme_classic(base_size=18, base_family="myriad") +
+     labs(title=trtNam[2])
> showtext_auto()
> grid.arrange(ggTrt1, ggTrt2, nrow = 2)
>
```

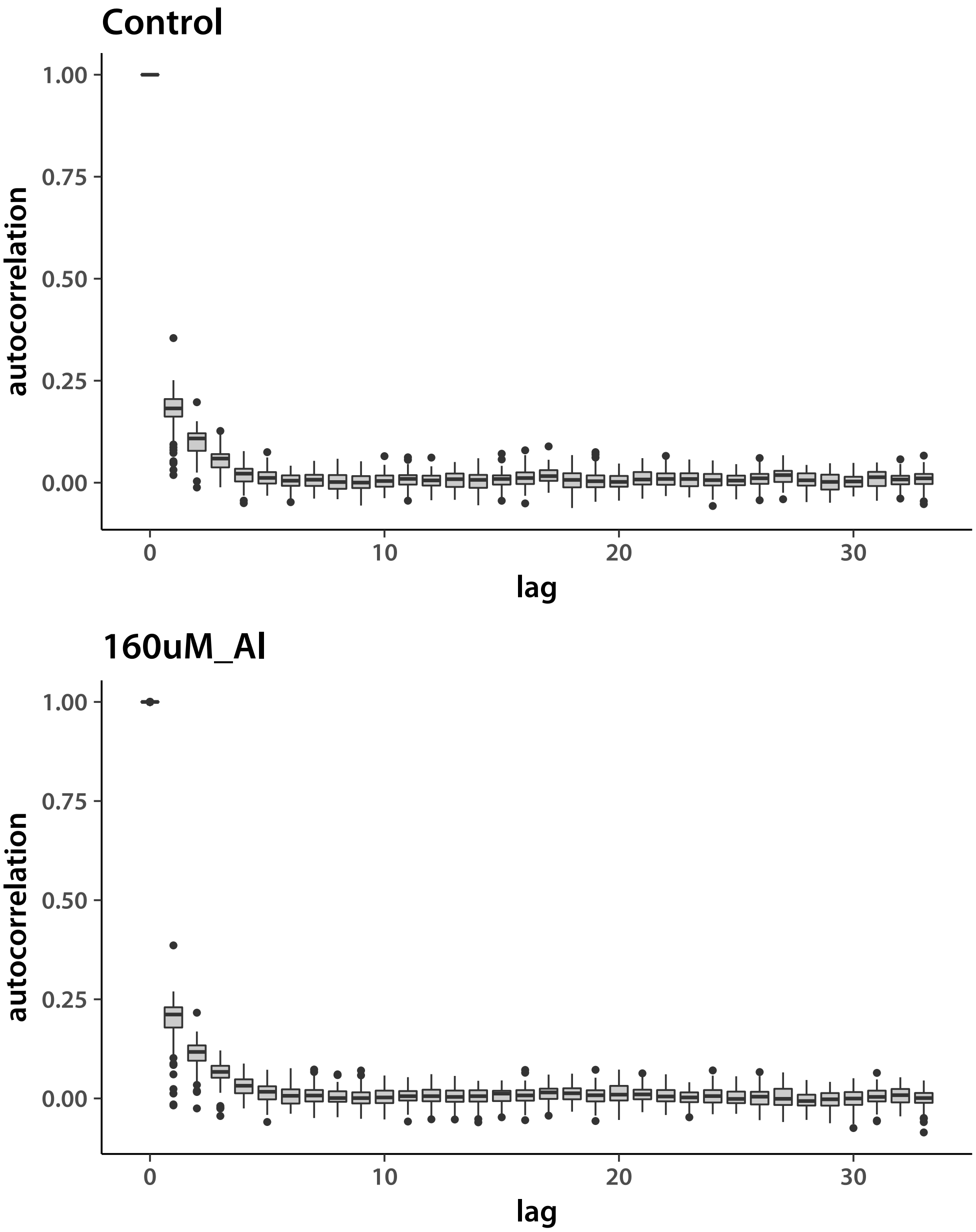


Everything looks good. Now examine GEBV chains, again starting with *aus*.

```
> Nacc   <- 55
```

```
> locDim <- d*Nacc
> chn1 <- matrix(.C("GSLmatLoad",
+     "chains/BVout_aus_3_1.gbin",
+     as.integer(chnLen), as.integer(locDim), out = double(chnLen*locDim))$out,
+         nrow = chnLen, byrow = T)
> chn2 <- matrix(.C("GSLmatLoad",
+     "chains/BVout_aus_3_2.gbin",
+     as.integer(chnLen), as.integer(locDim), out = double(chnLen*locDim))$out,
+         nrow = chnLen, byrow = T)
> chn3 <- matrix(.C("GSLmatLoad",
+     "chains/BVout_aus_3_3.gbin",
+     as.integer(chnLen), as.integer(locDim), out = double(chnLen*locDim))$out,
+         nrow = chnLen, byrow = T)
> chn4 <- matrix(.C("GSLmatLoad",
+     "chains/BVout_aus_3_4.gbin",
+     as.integer(chnLen), as.integer(locDim), out = double(chnLen*locDim))$out,
+         nrow = chnLen, byrow = T)
> chn5 <- matrix(.C("GSLmatLoad",
+     "chains/BVout_aus_3_5.gbin",
+     as.integer(chnLen), as.integer(locDim), out = double(chnLen*locDim))$out,
+         nrow = chnLen, byrow = T)
> grMat    <- matrix(gelRub("chn", nChn, chnLen), ncol = d, byrow = T)
> grFrame  <- data.frame(Gelman.Rubin=c(grMat), trait=rep(trtNam, each = Nacc))
> grMaxInd <- apply(grMat, 2, function(vec){which(vec == max(vec))[1]})
> acMat    <- mcmcAcf("chn", 5)
> acFrame  <- data.frame(autocorrelation=c(acMat), trait=rep(trtNam, prod(dim(acMat))/2),
+     lag=rep(0:33, each = nrow(acMat)))
>
```

Plot the *aus* results.

```
> showtext_auto()
> ggplot(data=grFrame,
+         aes(x=trait, y=Gelman.Rubin)) +
+     geom_boxplot(fill="grey80") +
+     theme_classic(base_size=18, base_family="myriad")
>
```

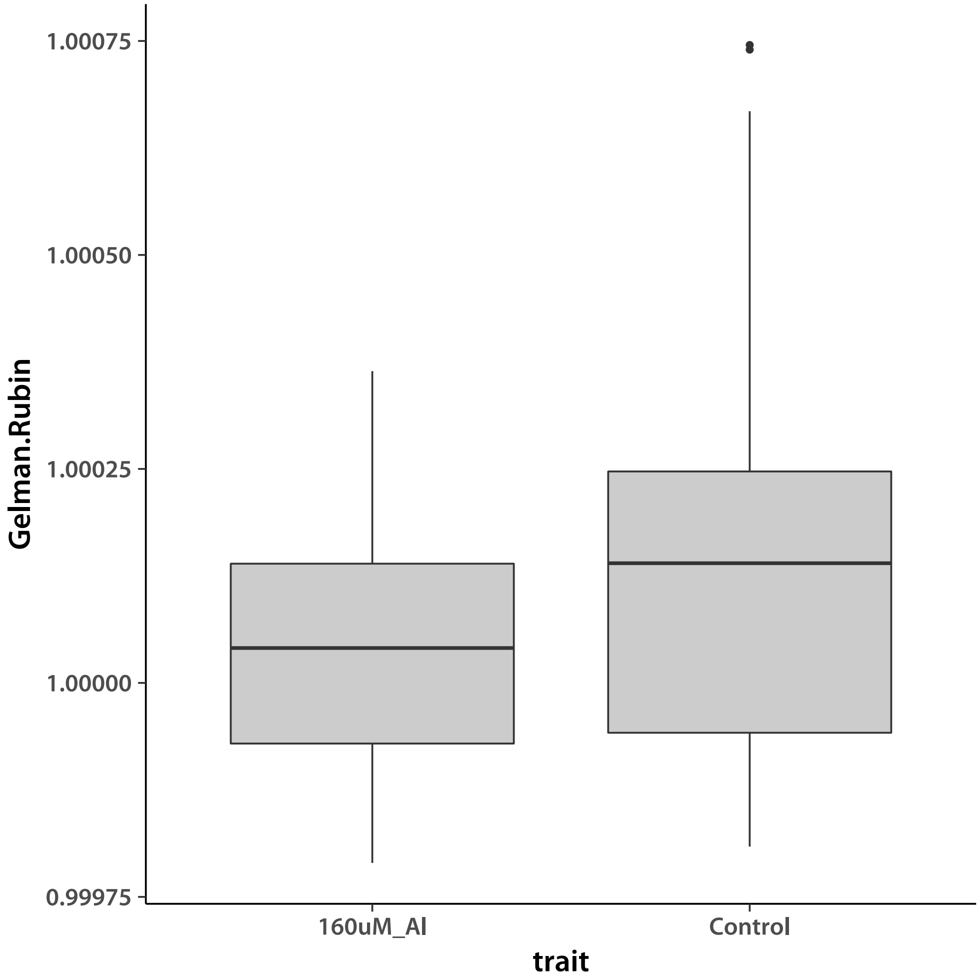


Values smaller than ∼1.2 indicate good convergence. I check by plotting an the chains with the worst G-R statistic for each trait.

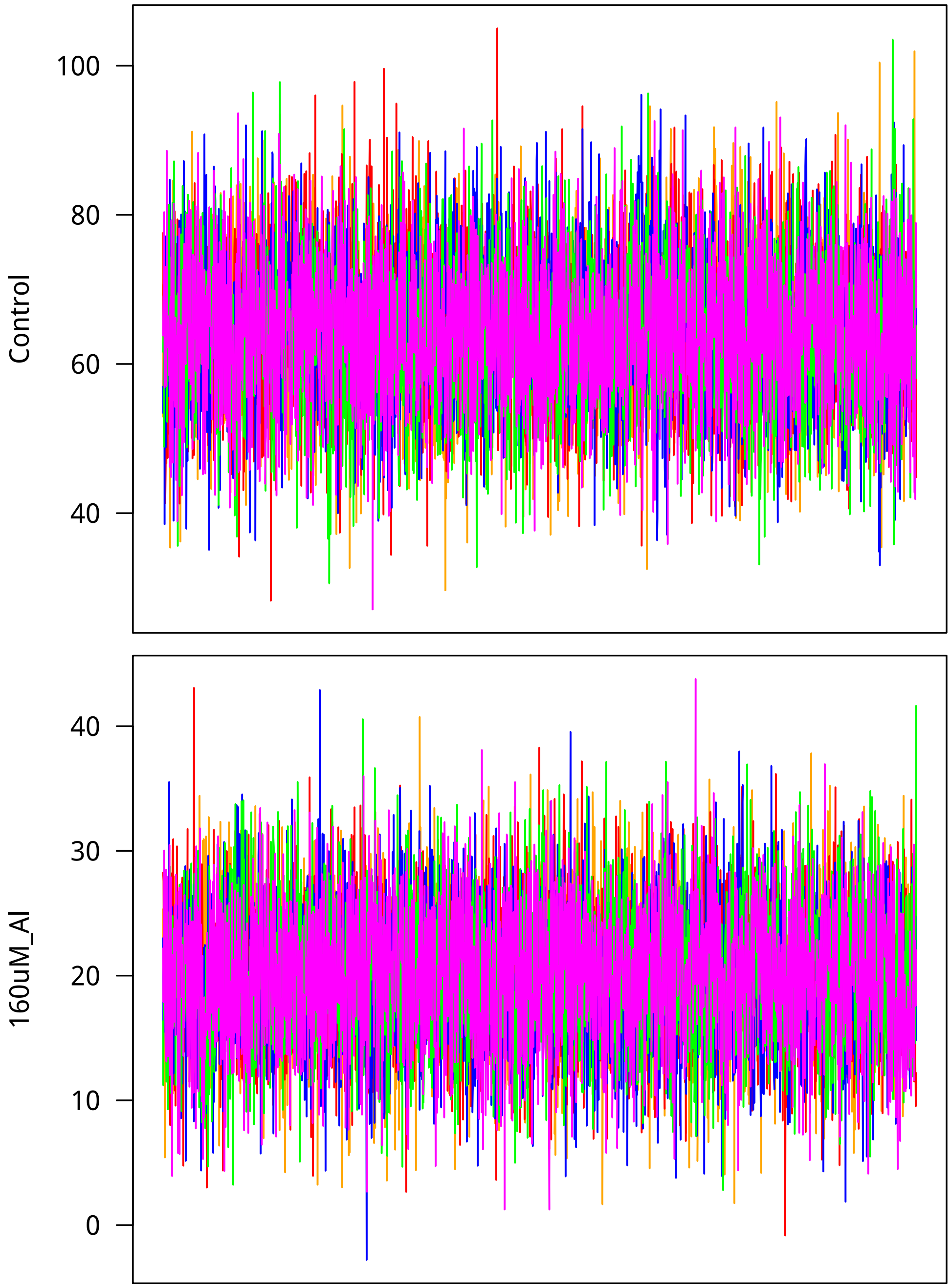


Now I look at mixing. Lag is on the $x$ axis.

```
> ggTrt1 <- ggplot(data=subset(acFrame, trait==trtNam[1]),
+         aes(x=lag, y=autocorrelation)) +
+     geom_boxplot(fill="grey80", aes(group=cut_interval(lag, n=34))) +
+     theme_classic(base_size=18, base_family="myriad") +
+     labs(title=trtNam[1])
> ggTrt2 <- ggplot(data=subset(acFrame, trait==trtNam[2]),
```

```
+         aes(x=lag, y=autocorrelation)) +
+     geom_boxplot(fill="grey80", aes(group=cut_interval(lag, n=34))) +
+     theme_classic(base_size=18, base_family="myriad") +
+     labs(title=trtNam[2])
> showtext_auto()
> grid.arrange(ggTrt1, ggTrt2, nrow = 2)
>
```

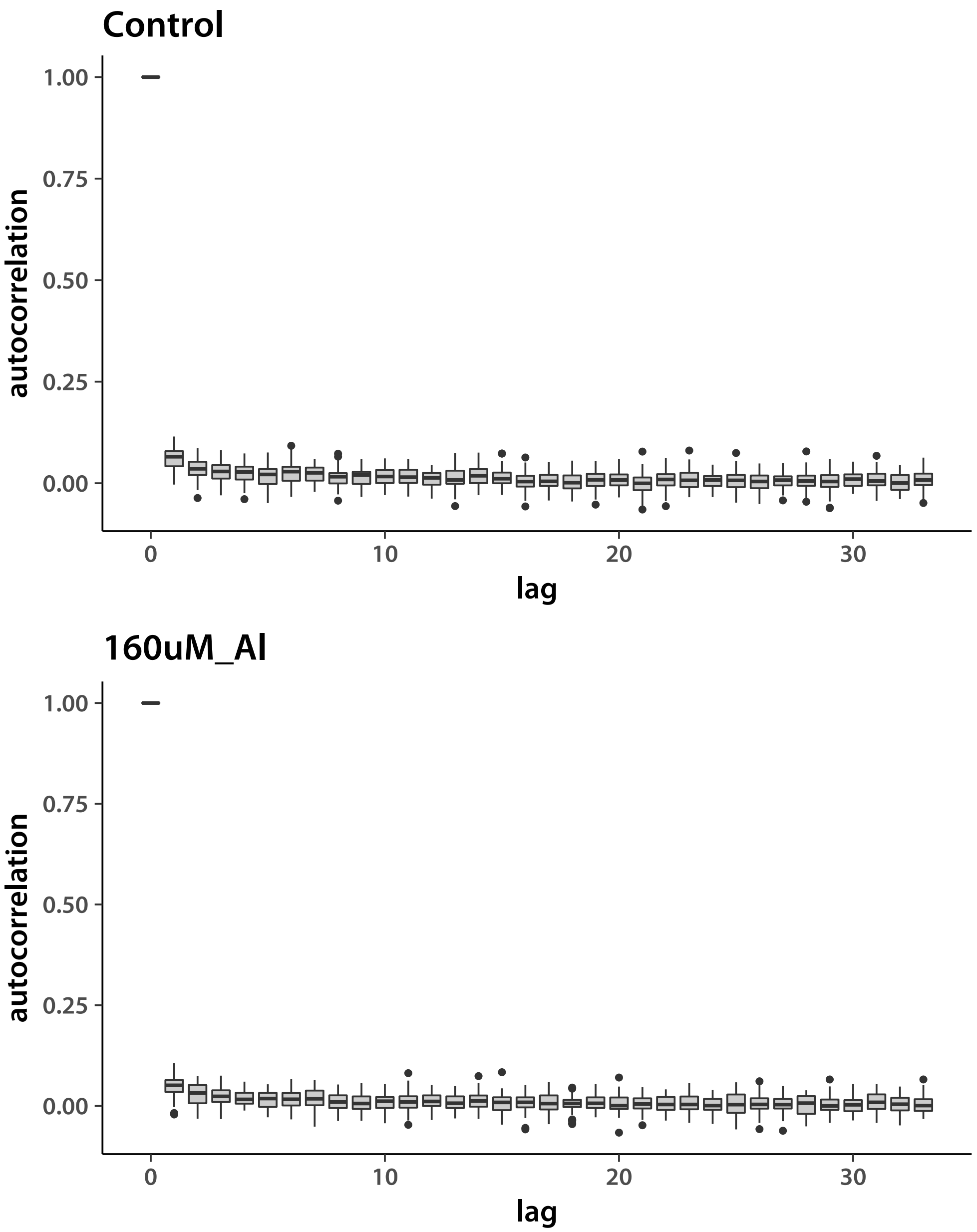


Looks good. Now do the tropical *japonica*.

```
> Nacc   <- 92
```

```
> locDim <- d*Nacc
> chn1 <- matrix(.C("GSLmatLoad",
+     "chains/BVout_trj_3_1.gbin",
+     as.integer(chnLen), as.integer(locDim), out = double(chnLen*locDim))$out,
+         nrow = chnLen, byrow = T)
> chn2 <- matrix(.C("GSLmatLoad",
+     "chains/BVout_trj_3_2.gbin",
+     as.integer(chnLen), as.integer(locDim), out = double(chnLen*locDim))$out,
+         nrow = chnLen, byrow = T)
> chn3 <- matrix(.C("GSLmatLoad",
+     "chains/BVout_trj_3_3.gbin",
+     as.integer(chnLen), as.integer(locDim), out = double(chnLen*locDim))$out,
+         nrow = chnLen, byrow = T)
> chn4 <- matrix(.C("GSLmatLoad",
+     "chains/BVout_trj_3_4.gbin",
+     as.integer(chnLen), as.integer(locDim), out = double(chnLen*locDim))$out,
+         nrow = chnLen, byrow = T)
> chn5 <- matrix(.C("GSLmatLoad",
+     "chains/BVout_trj_3_5.gbin",
+     as.integer(chnLen), as.integer(locDim), out = double(chnLen*locDim))$out,
+         nrow = chnLen, byrow = T)
> grMat    <- matrix(gelRub("chn", nChn, chnLen), ncol = d, byrow = T)
> grFrame  <- data.frame(Gelman.Rubin=c(grMat), trait=rep(trtNam, each = Nacc))
> grMaxInd <- apply(grMat, 2, function(vec){which(vec == max(vec))[1]})
> acMat    <- mcmcAcf("chn", 5)
> acFrame  <- data.frame(autocorrelation=c(acMat), trait=rep(trtNam, prod(dim(acMat))/2),
+     lag=rep(0:33, each = nrow(acMat)))
>
```

Plot the *aus* results.

```
> showtext_auto()
> ggplot(data=grFrame,
+         aes(x=trait, y=Gelman.Rubin)) +
+     geom_boxplot(fill="grey80") +
+     theme_classic(base_size=18, base_family="myriad")
>
```

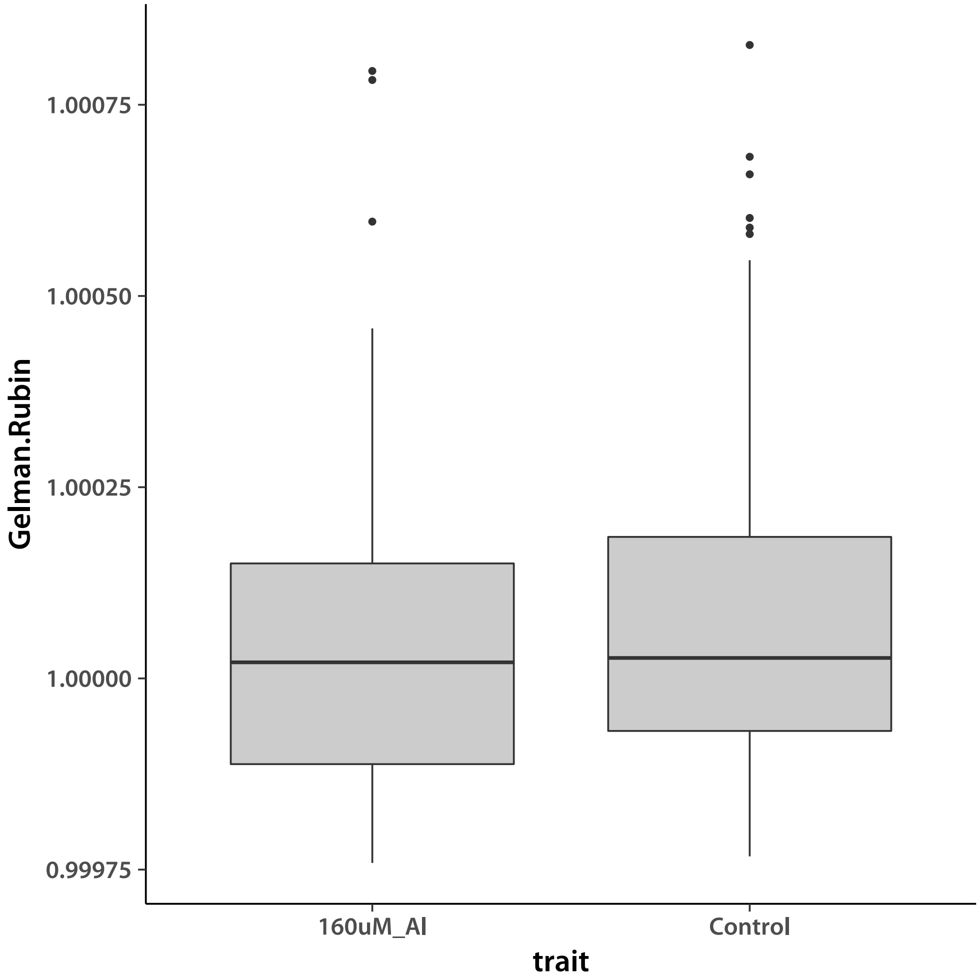


Values smaller than ~1.2 indicate good convergence. I check by plotting an the chains with the worst G-R statistic for each trait.

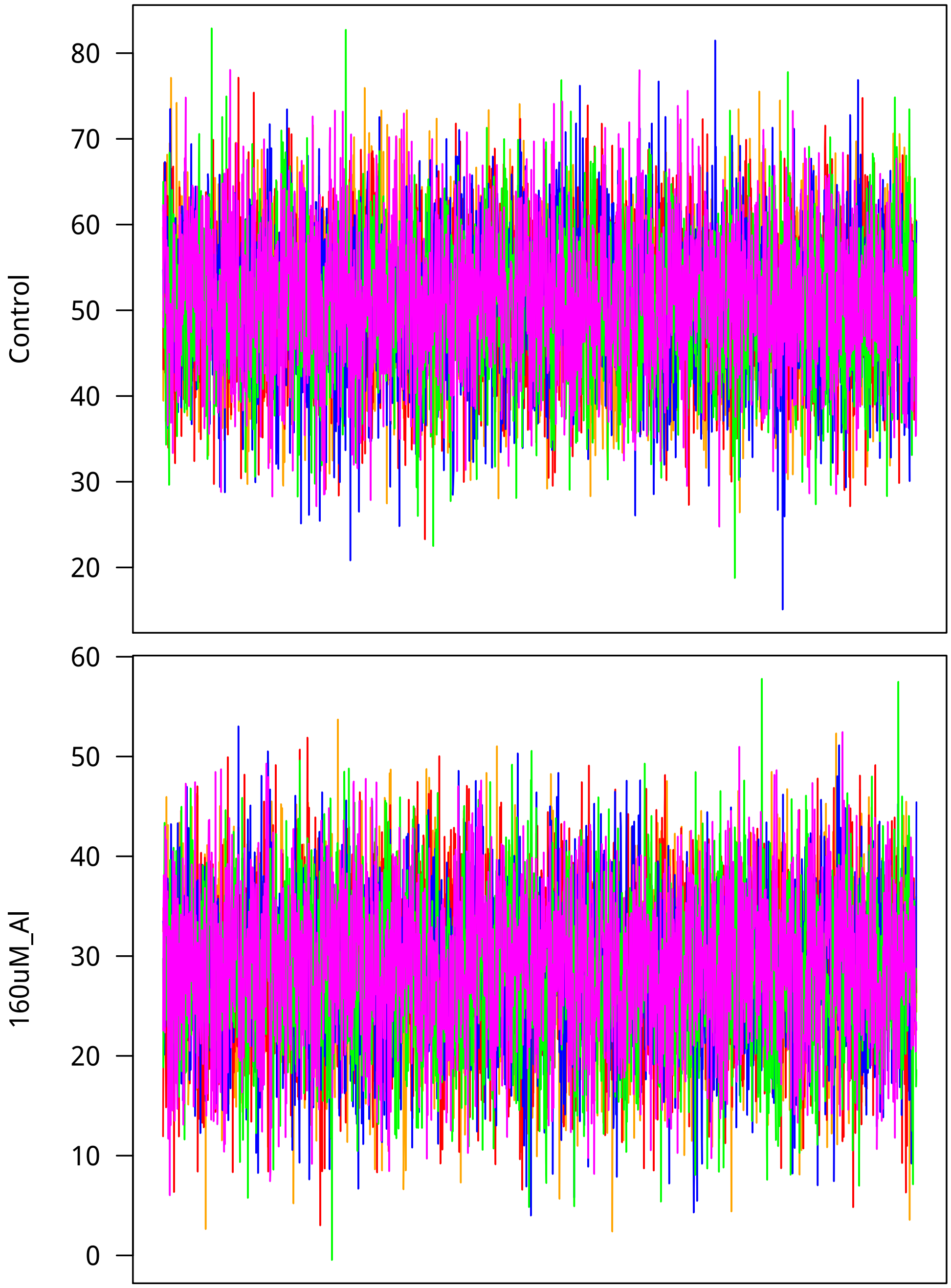


Now I look at mixing. Lag is on the $x$ axis.

```
> ggTrt1 <- ggplot(data=subset(acFrame, trait==trtNam[1]),
+          aes(x=lag, y=autocorrelation)) +
+      geom_boxplot(fill="grey80", aes(group=cut_interval(lag, n=34))) +
+      theme_classic(base_size=18, base_family="myriad") +
+      labs(title=trtNam[1])
> ggTrt2 <- ggplot(data=subset(acFrame, trait==trtNam[2]),
```

```
+         aes(x=lag, y=autocorrelation)) +
+     geom_boxplot(fill="grey80", aes(group=cut_interval(lag, n=34))) +
+     theme_classic(base_size=18, base_family="myriad") +
+     labs(title=trtNam[2])
> showtext_auto()
> grid.arrange(ggTrt1, ggTrt2, nrow = 2)
>
```

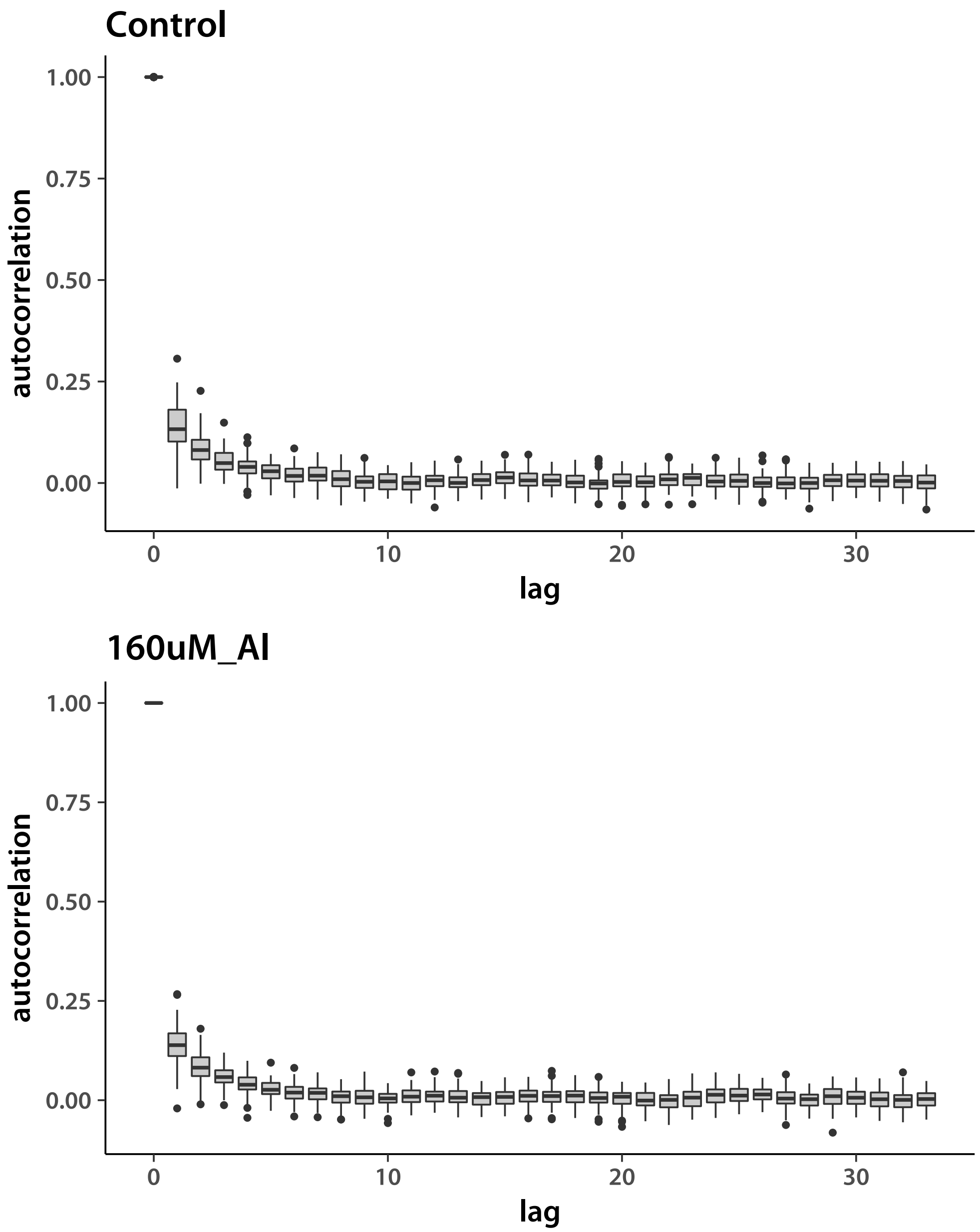


Next, admixed tropical *japonica*.

```
> Nacc   <- 95
```

```
> locDim <- d*Nacc
> chn1 <- matrix(.C("GSLmatLoad",
+     "chains/BVout_trja_3_1.gbin",
+     as.integer(chnLen), as.integer(locDim), out = double(chnLen*locDim))$out,
+         nrow = chnLen, byrow = T)
> chn2 <- matrix(.C("GSLmatLoad",
+     "chains/BVout_trja_3_2.gbin",
+     as.integer(chnLen), as.integer(locDim), out = double(chnLen*locDim))$out,
+         nrow = chnLen, byrow = T)
> chn3 <- matrix(.C("GSLmatLoad",
+     "chains/BVout_trja_3_3.gbin",
+     as.integer(chnLen), as.integer(locDim), out = double(chnLen*locDim))$out,
+         nrow = chnLen, byrow = T)
> chn4 <- matrix(.C("GSLmatLoad",
+     "chains/BVout_trja_3_4.gbin",
+     as.integer(chnLen), as.integer(locDim), out = double(chnLen*locDim))$out,
+         nrow = chnLen, byrow = T)
> chn5 <- matrix(.C("GSLmatLoad",
+     "chains/BVout_trja_3_5.gbin",
+     as.integer(chnLen), as.integer(locDim), out = double(chnLen*locDim))$out,
+         nrow = chnLen, byrow = T)
> grMat    <- matrix(gelRub("chn", nChn, chnLen), ncol = d, byrow = T)
> grFrame  <- data.frame(Gelman.Rubin=c(grMat), trait=rep(trtNam, each = Nacc))
> grMaxInd <- apply(grMat, 2, function(vec){which(vec == max(vec))[1]})
> acMat    <- mcmcAcf("chn", 5)
> acFrame  <- data.frame(autocorrelation=c(acMat), trait=rep(trtNam, prod(dim(acMat))/2),
+     lag=rep(0:33, each = nrow(acMat)))
>
```

Plot the *aus* results.

```
> showtext_auto()
> ggplot(data=grFrame,
+         aes(x=trait, y=Gelman.Rubin)) +
+     geom_boxplot(fill="grey80") +
+     theme_classic(base_size=18, base_family="myriad")
>
```

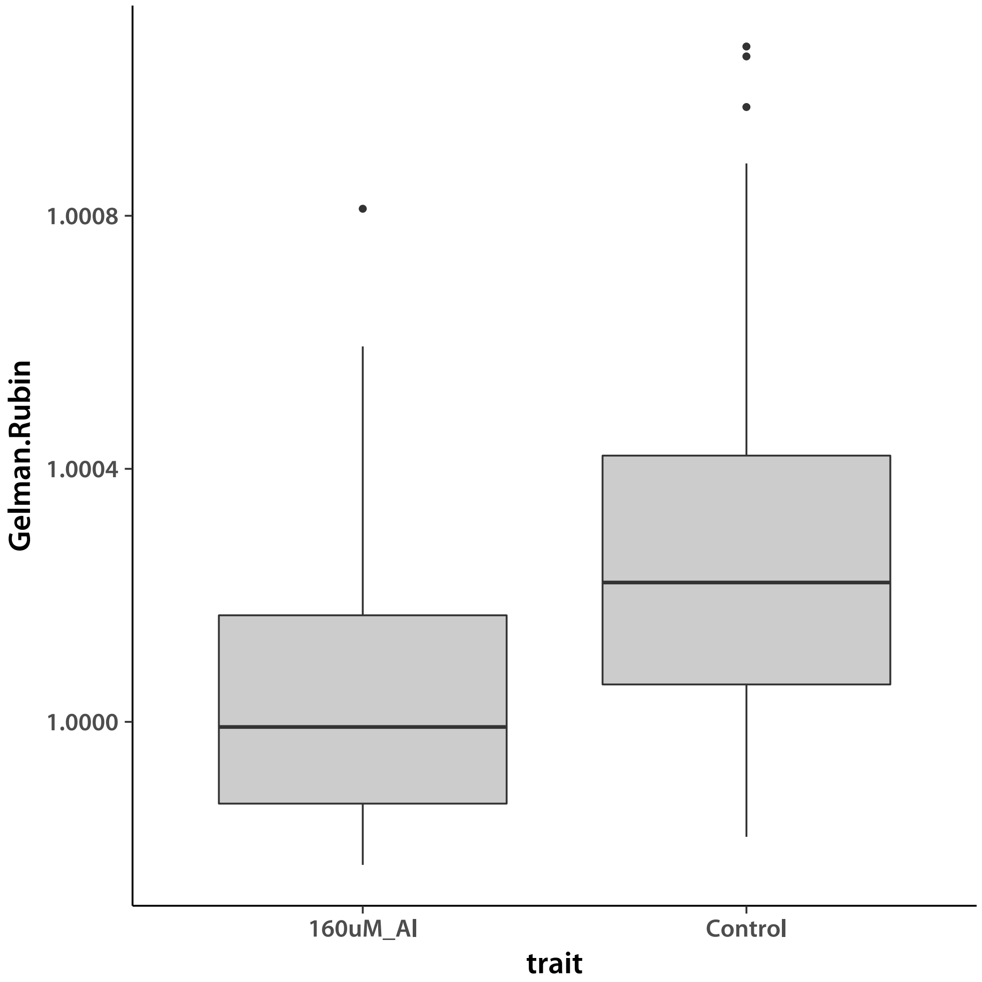


Values smaller than ~1.2 indicate good convergence. I check by plotting an the chains with the worst G-R statistic for each trait.

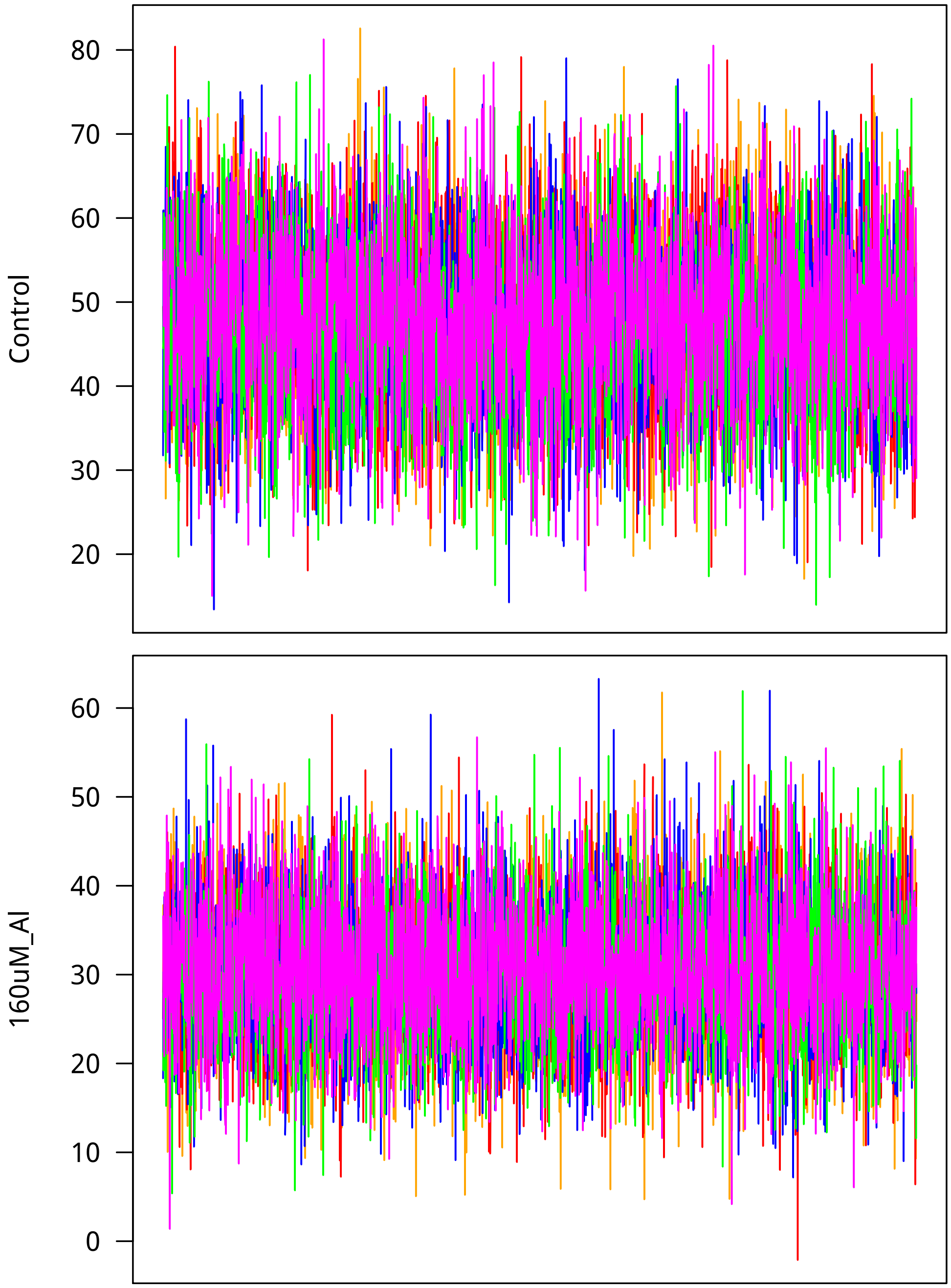


Now I look at mixing. Lag is on the $x$ axis.

```
> ggTrt1 <- ggplot(data=subset(acFrame, trait==trtNam[1]),
+         aes(x=lag, y=autocorrelation)) +
+     geom_boxplot(fill="grey80", aes(group=cut_interval(lag, n=34))) +
+     theme_classic(base_size=18, base_family="myriad") +
+     labs(title=trtNam[1])
> ggTrt2 <- ggplot(data=subset(acFrame, trait==trtNam[2]),
```

```
+         aes(x=lag, y=autocorrelation)) +
+     geom_boxplot(fill="grey80", aes(group=cut_interval(lag, n=34))) +
+     theme_classic(base_size=18, base_family="myriad") +
+     labs(title=trtNam[2])
> showtext_auto()
> grid.arrange(ggTrt1, ggTrt2, nrow = 2)
>
```

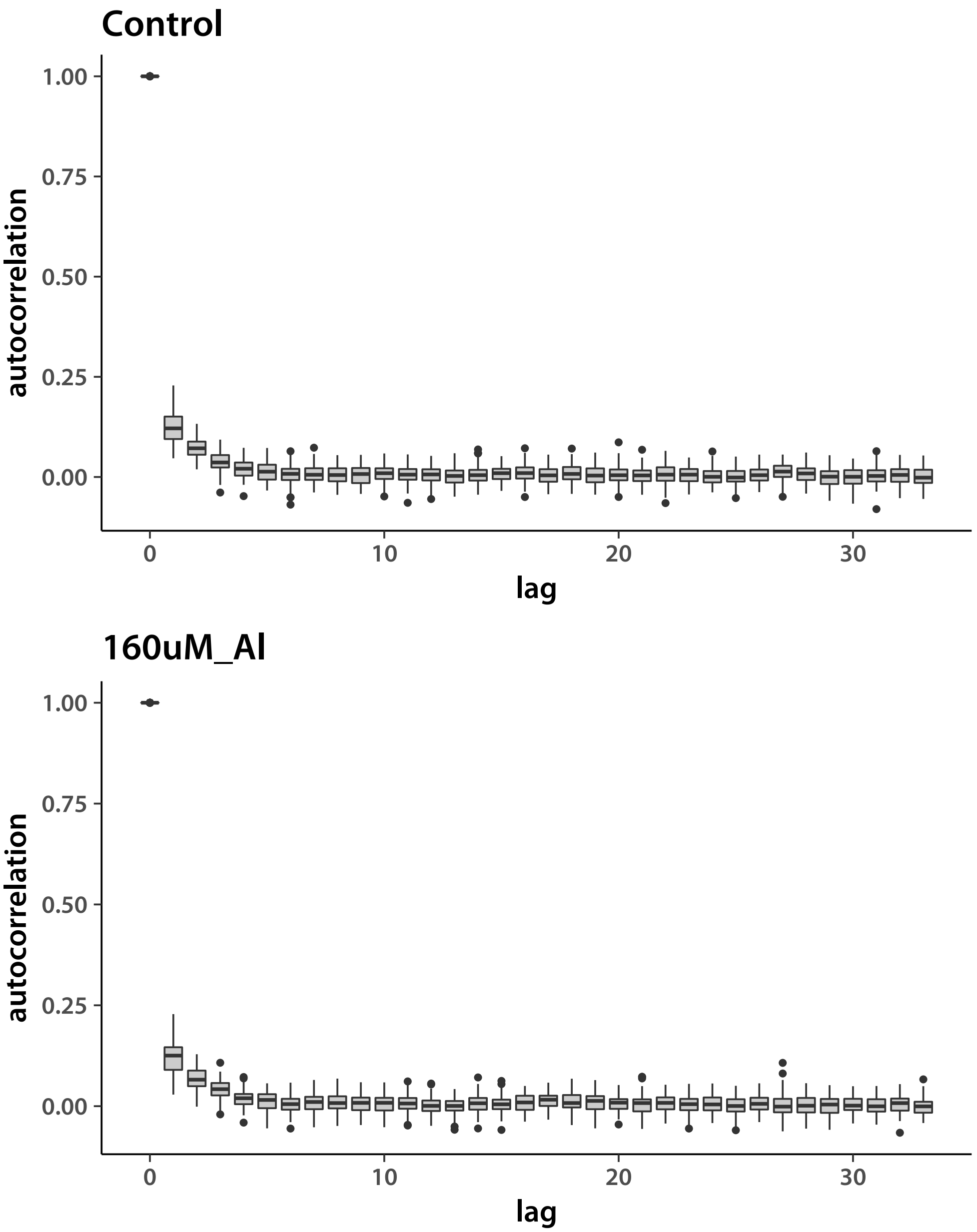


Finally, I look at covariance matrices. I create an index to isolate just the upper triangles. I again start with *aus*.

```
> upInd   <- matrix(1:(d^2), ncol = d, byrow = T)
> upInd   <- upInd[row(upInd) <= col(upInd)]
> diagInd <- diag(matrix(1:(d^2), ncol = d, byrow = T))
```

Read in covariance matrix chains. Error correlations are set to zero, I exclude them.

```
> chn1 <- matrix(.C("GSLmatLoad",
+     "chains/SgEout_aus_3_1.gbin",
+     as.integer(chnLen), as.integer(d^2), out = double(chnLen*d^2))$out,
+         nrow = chnLen, byrow = T)[,diagInd]
> chn1 <- cbind(chn1,
+     matrix(.C("GSLmatLoad",
+     "chains/SgSout_aus_3_1.gbin",
+     as.integer(chnLen), as.integer(d^2), out = double(chnLen*d^2))$out,
+         nrow = chnLen, byrow = T)[,upInd])
> chn1 <- cbind(chn1,
+     matrix(.C("GSLmatLoad",
+     "chains/SgAout_aus_3_1.gbin",
+     as.integer(chnLen), as.integer(d^2), out = double(chnLen*d^2))$out,
+         nrow = chnLen, byrow = T)[,upInd])
> chn2 <- matrix(.C("GSLmatLoad",
+     "chains/SgEout_aus_3_2.gbin",
+     as.integer(chnLen), as.integer(d^2), out = double(chnLen*d^2))$out,
+         nrow = chnLen, byrow = T)[,diagInd]
> chn2 <- cbind(chn2,
+     matrix(.C("GSLmatLoad",
+     "chains/SgSout_aus_3_2.gbin",
+     as.integer(chnLen), as.integer(d^2), out = double(chnLen*d^2))$out,
+         nrow = chnLen, byrow = T)[,upInd])
> chn2 <- cbind(chn2,
+     matrix(.C("GSLmatLoad",
+     "chains/SgAout_aus_3_2.gbin",
+     as.integer(chnLen), as.integer(d^2), out = double(chnLen*d^2))$out,
+         nrow = chnLen, byrow = T)[,upInd])
> chn3 <- matrix(.C("GSLmatLoad",
+     "chains/SgEout_aus_3_3.gbin",
+     as.integer(chnLen), as.integer(d^2), out = double(chnLen*d^2))$out,
+         nrow = chnLen, byrow = T)[,diagInd]
> chn3 <- cbind(chn3,
+     matrix(.C("GSLmatLoad",
+     "chains/SgSout_aus_3_3.gbin",
+     as.integer(chnLen), as.integer(d^2), out = double(chnLen*d^2))$out,
+         nrow = chnLen, byrow = T)[,upInd])
> chn3 <- cbind(chn3,
+     matrix(.C("GSLmatLoad",
+     "chains/SgAout_aus_3_3.gbin",
```

```
+     as.integer(chnLen), as.integer(d^2), out = double(chnLen*d^2))$out,
+         nrow = chnLen, byrow = T)[,upInd])
> chn4 <- matrix(.C("GSLmatLoad",
+     "chains/SgEout_aus_3_4.gbin",
+     as.integer(chnLen), as.integer(d^2), out = double(chnLen*d^2))$out,
+         nrow = chnLen, byrow = T)[,diagInd]
> chn4 <- cbind(chn4,
+     matrix(.C("GSLmatLoad",
+     "chains/SgSout_aus_3_4.gbin",
+     as.integer(chnLen), as.integer(d^2), out = double(chnLen*d^2))$out,
+         nrow = chnLen, byrow = T)[,upInd])
> chn4 <- cbind(chn4,
+     matrix(.C("GSLmatLoad",
+     "chains/SgAout_aus_3_4.gbin",
+     as.integer(chnLen), as.integer(d^2), out = double(chnLen*d^2))$out,
+         nrow = chnLen, byrow = T)[,upInd])
> chn5 <- matrix(.C("GSLmatLoad",
+     "chains/SgEout_aus_3_5.gbin",
+     as.integer(chnLen), as.integer(d^2), out = double(chnLen*d^2))$out,
+         nrow = chnLen, byrow = T)[,diagInd]
> chn5 <- cbind(chn5,
+     matrix(.C("GSLmatLoad",
+     "chains/SgSout_aus_3_5.gbin",
+     as.integer(chnLen), as.integer(d^2), out = double(chnLen*d^2))$out,
+         nrow = chnLen, byrow = T)[,upInd])
> chn5 <- cbind(chn5,
+     matrix(.C("GSLmatLoad",
+     "chains/SgAout_aus_3_5.gbin",
+     as.integer(chnLen), as.integer(d^2), out = double(chnLen*d^2))$out,
+         nrow = chnLen, byrow = T)[,upInd])
> acMat   <- mcmcAcf("chn", 5)
> acFrame <- data.frame(autocorrelation=c(acMat),
+     Sigma=rep(c(rep("SgE", length(diagInd)),
+     rep(c("SgS", "SgA"), each=length(upInd))), ncol(acMat)),
+     lag=rep(0:33, each = nrow(acMat)))
> gelRub("chn",nChn,chnLen)
```

```
[1] 0.9999770 0.9999225 1.0004899 1.0001130 0.9998007 1.0001715 1.0005032 0.9999878
```

```
> ggSgE <- ggplot(data=subset(acFrame, Sigma=="SgE"),
+         aes(x=lag, y=autocorrelation)) +
+     geom_point() +
+     theme_classic(base_size=18, base_family="myriad") +
+     labs(title=expression(Sigma[e]))
> ggSgS <- ggplot(data=subset(acFrame, Sigma=="SgS"),
```

```
+         aes(x=lag, y=autocorrelation)) +
+     geom_point() +
+     theme_classic(base_size=18, base_family="myriad") +
+     labs(title=expression(Sigma[s]))
> ggSgA <- ggplot(data=subset(acFrame, Sigma=="SgA"),
+         aes(x=lag, y=autocorrelation)) +
+     geom_point() +
+     theme_classic(base_size=18, base_family="myriad") +
+     labs(title=expression(Sigma[a]))
> showtext_auto()
> grid.arrange(ggSgE, ggSgS, ggSgA, nrow = 3)
>
```

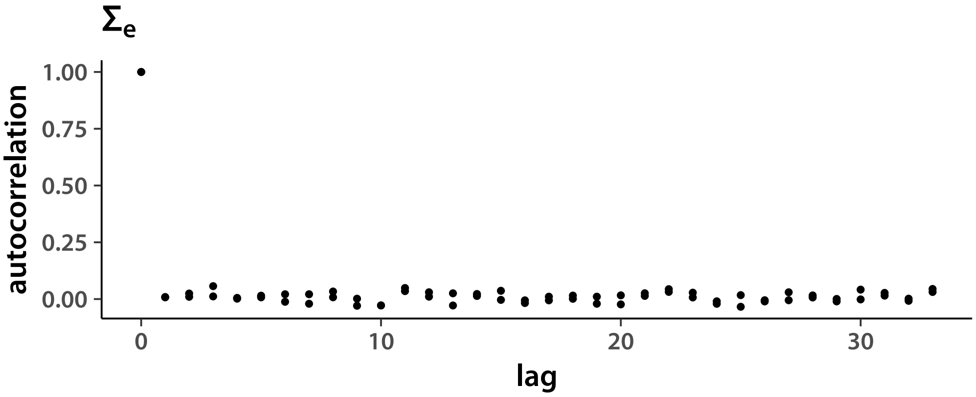


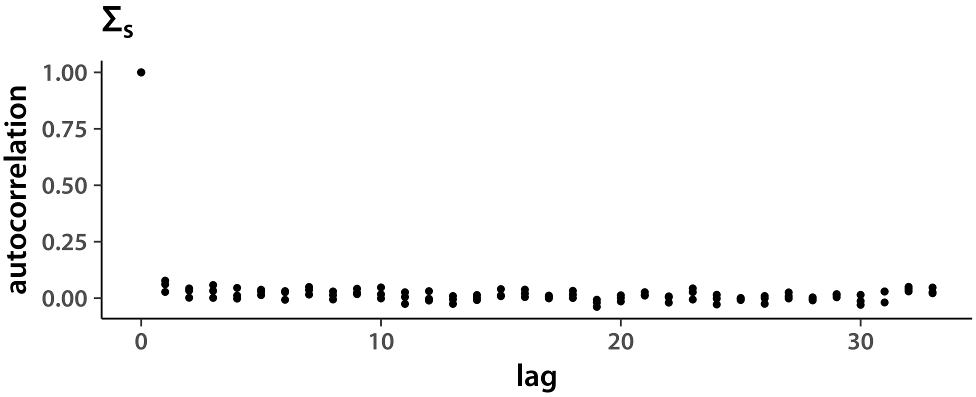


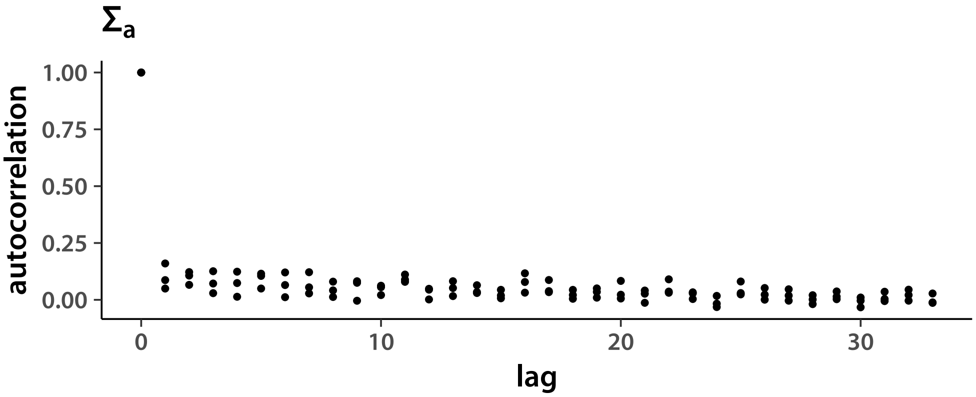


Plot variance chains. First, I get the diagonal index from the upper triangle index.

```
> diagList <- list(1:d, c(3,5), c(6,8))
```

$\Sigma_e$:

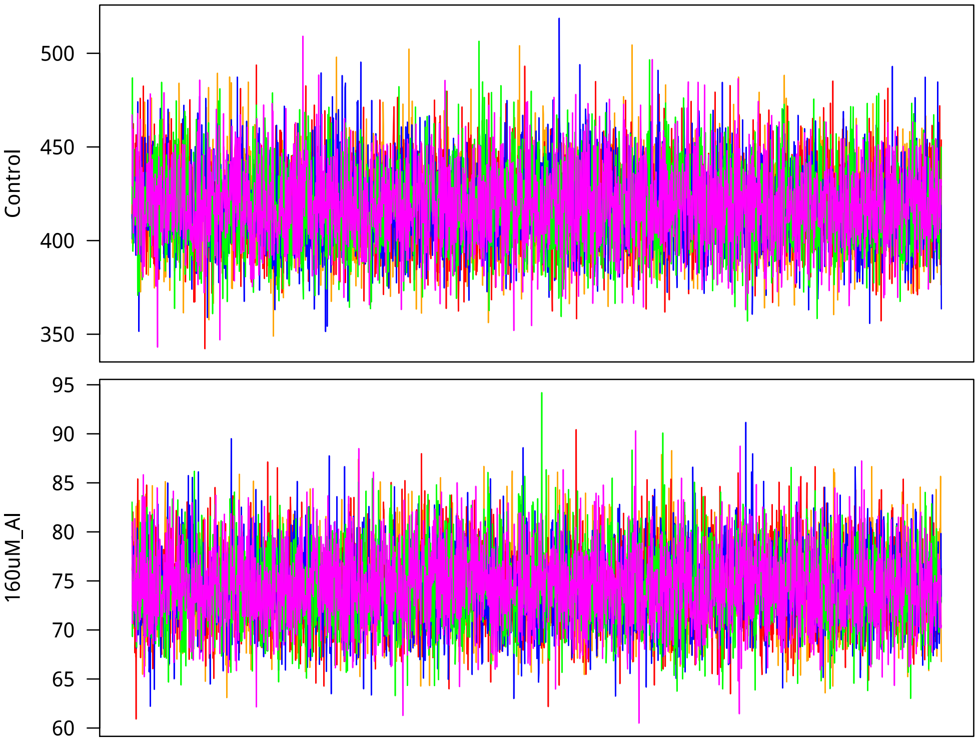


Next, non-additive genetic variances.

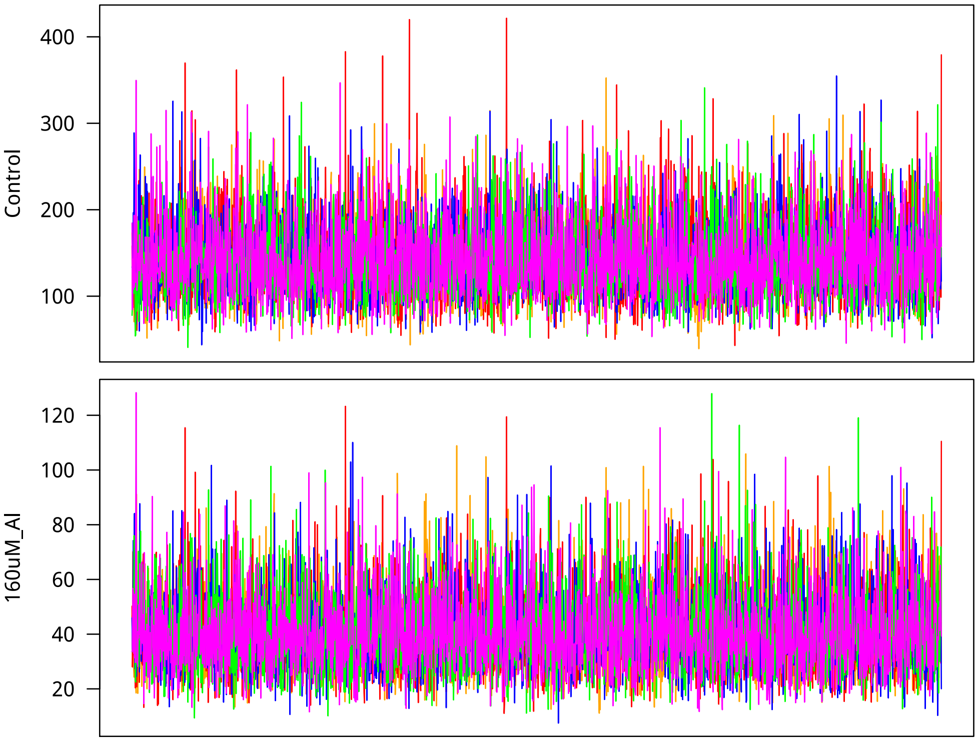

Finally, $\mathbf{\Sigma}_a$ which can not be interpreted as the additive genetic covariance matrix in this case, where $\nu_g = 3$ and the model is Student-$t$.

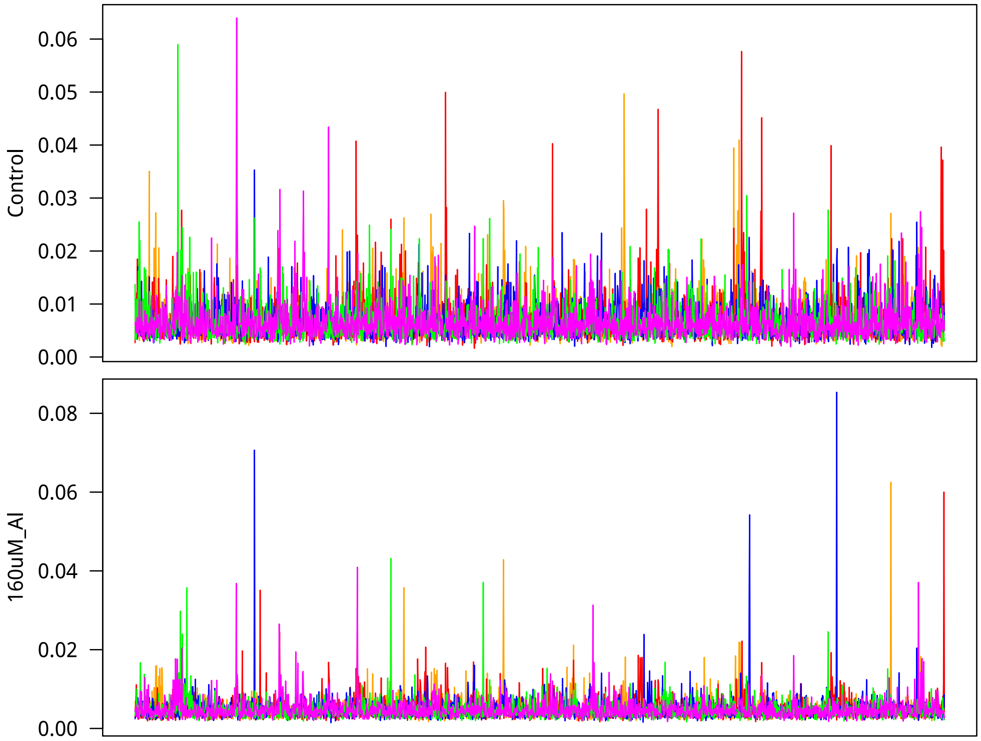


Everything looks good and I move on to the tropical *japonica* covariances.

```
> chn1 <- matrix(.C("GSLmatLoad",
+     "chains/SgEout_trj_3_1.gbin",
+     as.integer(chnLen), as.integer(d^2), out = double(chnLen*d^2))$out,
+         nrow = chnLen, byrow = T)[,diagInd]
> chn1 <- cbind(chn1,
+     matrix(.C("GSLmatLoad",
+     "chains/SgSout_trj_3_1.gbin",
+     as.integer(chnLen), as.integer(d^2), out = double(chnLen*d^2))$out,
+         nrow = chnLen, byrow = T)[,upInd])
> chn1 <- cbind(chn1,
+     matrix(.C("GSLmatLoad",
+     "chains/SgAout_trj_3_1.gbin",
+     as.integer(chnLen), as.integer(d^2), out = double(chnLen*d^2))$out,
+         nrow = chnLen, byrow = T)[,upInd])
> chn2 <- matrix(.C("GSLmatLoad",
+     "chains/SgEout_trj_3_2.gbin",
+     as.integer(chnLen), as.integer(d^2), out = double(chnLen*d^2))$out,
+         nrow = chnLen, byrow = T)[,diagInd]
> chn2 <- cbind(chn2,
```

```
+     matrix(.C("GSLmatLoad",
+     "chains/SgSout_trj_3_2.gbin",
+     as.integer(chnLen), as.integer(d^2), out = double(chnLen*d^2))$out,
+         nrow = chnLen, byrow = T)[,upInd])
> chn2 <- cbind(chn2,
+     matrix(.C("GSLmatLoad",
+     "chains/SgAout_trj_3_2.gbin",
+     as.integer(chnLen), as.integer(d^2), out = double(chnLen*d^2))$out,
+         nrow = chnLen, byrow = T)[,upInd])
> chn3 <- matrix(.C("GSLmatLoad",
+     "chains/SgEout_trj_3_3.gbin",
+     as.integer(chnLen), as.integer(d^2), out = double(chnLen*d^2))$out,
+         nrow = chnLen, byrow = T)[,diagInd]
> chn3 <- cbind(chn3,
+     matrix(.C("GSLmatLoad",
+     "chains/SgSout_trj_3_3.gbin",
+     as.integer(chnLen), as.integer(d^2), out = double(chnLen*d^2))$out,
+         nrow = chnLen, byrow = T)[,upInd])
> chn3 <- cbind(chn3,
+     matrix(.C("GSLmatLoad",
+     "chains/SgAout_trj_3_3.gbin",
+     as.integer(chnLen), as.integer(d^2), out = double(chnLen*d^2))$out,
+         nrow = chnLen, byrow = T)[,upInd])
> chn4 <- matrix(.C("GSLmatLoad",
+     "chains/SgEout_trj_3_4.gbin",
+     as.integer(chnLen), as.integer(d^2), out = double(chnLen*d^2))$out,
+         nrow = chnLen, byrow = T)[,diagInd]
> chn4 <- cbind(chn4,
+     matrix(.C("GSLmatLoad",
+     "chains/SgSout_trj_3_4.gbin",
+     as.integer(chnLen), as.integer(d^2), out = double(chnLen*d^2))$out,
+         nrow = chnLen, byrow = T)[,upInd])
> chn4 <- cbind(chn4,
+     matrix(.C("GSLmatLoad",
+     "chains/SgAout_trj_3_4.gbin",
+     as.integer(chnLen), as.integer(d^2), out = double(chnLen*d^2))$out,
+         nrow = chnLen, byrow = T)[,upInd])
> chn5 <- matrix(.C("GSLmatLoad",
+     "chains/SgEout_trj_3_5.gbin",
+     as.integer(chnLen), as.integer(d^2), out = double(chnLen*d^2))$out,
+         nrow = chnLen, byrow = T)[,diagInd]
> chn5 <- cbind(chn5,
+     matrix(.C("GSLmatLoad",
+     "chains/SgSout_trj_3_5.gbin",
+     as.integer(chnLen), as.integer(d^2), out = double(chnLen*d^2))$out,
```

```
+         nrow = chnLen, byrow = T)[,upInd])
> chn5 <- cbind(chn5,
+     matrix(.C("GSLmatLoad",
+     "chains/SgAout_trj_3_5.gbin",
+     as.integer(chnLen), as.integer(d^2), out = double(chnLen*d^2))$out,
+         nrow = chnLen, byrow = T)[,upInd])
> acMat   <- mcmcAcf("chn", 5)
> acFrame <- data.frame(autocorrelation=c(acMat),
+     Sigma=rep(c(rep("SgE", length(diagInd)),
+     rep(c("SgS", "SgA"), each=length(upInd))), ncol(acMat)),
+     lag=rep(0:33, each = nrow(acMat)))
> gelRub("chn",nChn,chnLen)

[1] 0.9999211 1.0000121 1.0005728 1.0005353 1.0004180 0.9998886 0.9997742 0.9997981


> ggSgE <- ggplot(data=subset(acFrame, Sigma=="SgE"),
+         aes(x=lag, y=autocorrelation)) +
+     geom_point() +
+     theme_classic(base_size=18, base_family="myriad") +
+     labs(title=expression(Sigma[e]))
> ggSgS <- ggplot(data=subset(acFrame, Sigma=="SgS"),
+         aes(x=lag, y=autocorrelation)) +
+     geom_point() +
+     theme_classic(base_size=18, base_family="myriad") +
+     labs(title=expression(Sigma[s]))
> ggSgA <- ggplot(data=subset(acFrame, Sigma=="SgA"),
+         aes(x=lag, y=autocorrelation)) +
+     geom_point() +
+     theme_classic(base_size=18, base_family="myriad") +
+     labs(title=expression(Sigma[a]))
> showtext_auto()
> grid.arrange(ggSgE, ggSgS, ggSgA, nrow = 3)
>
```

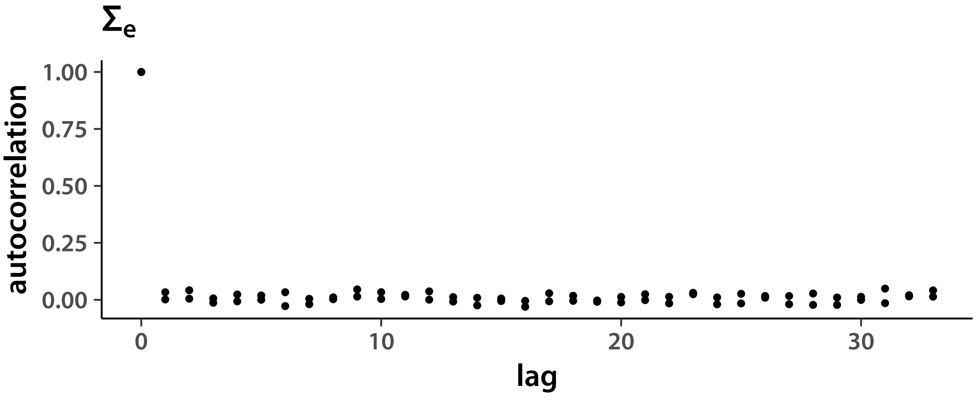


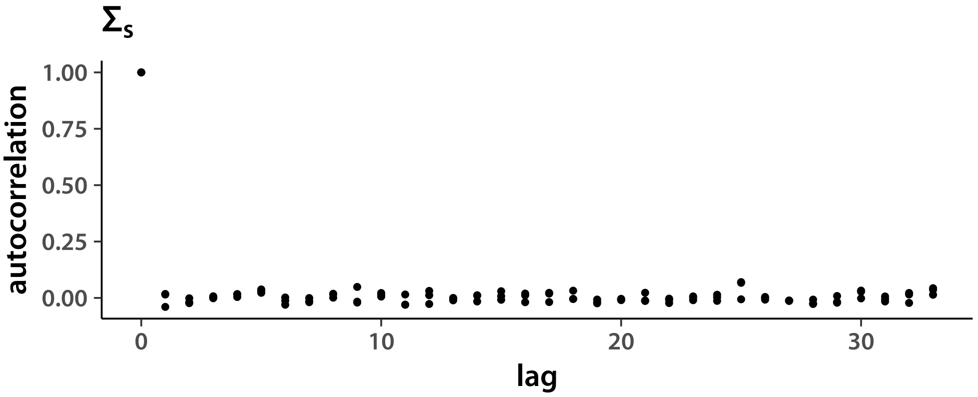


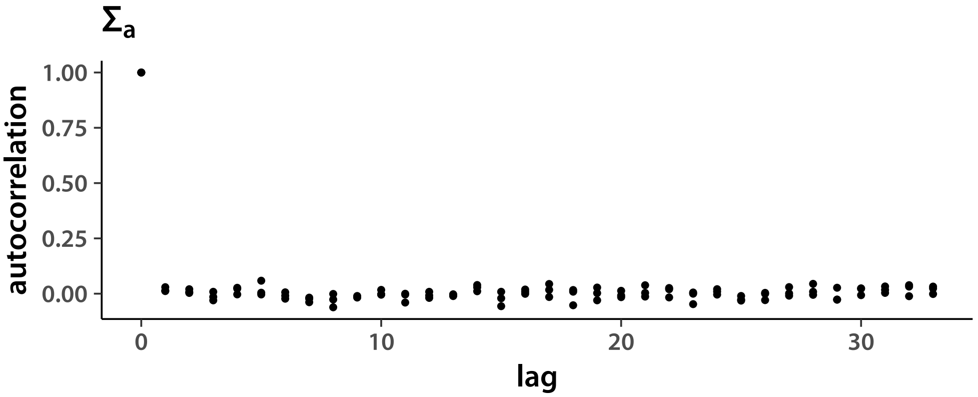


Plot variance chains. First, I get the diagonal index from the upper triangle index.

```
> diagList <- list(1:d, c(3,5), c(6,8))
```

$\Sigma_e$:

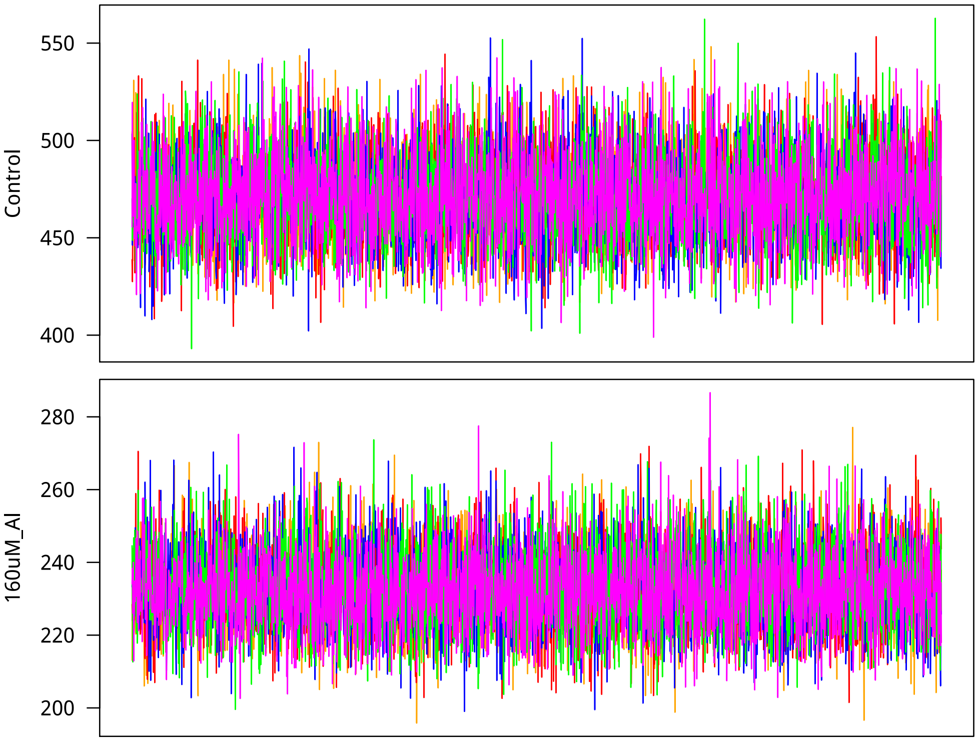


Next, non-additive genetic variances.

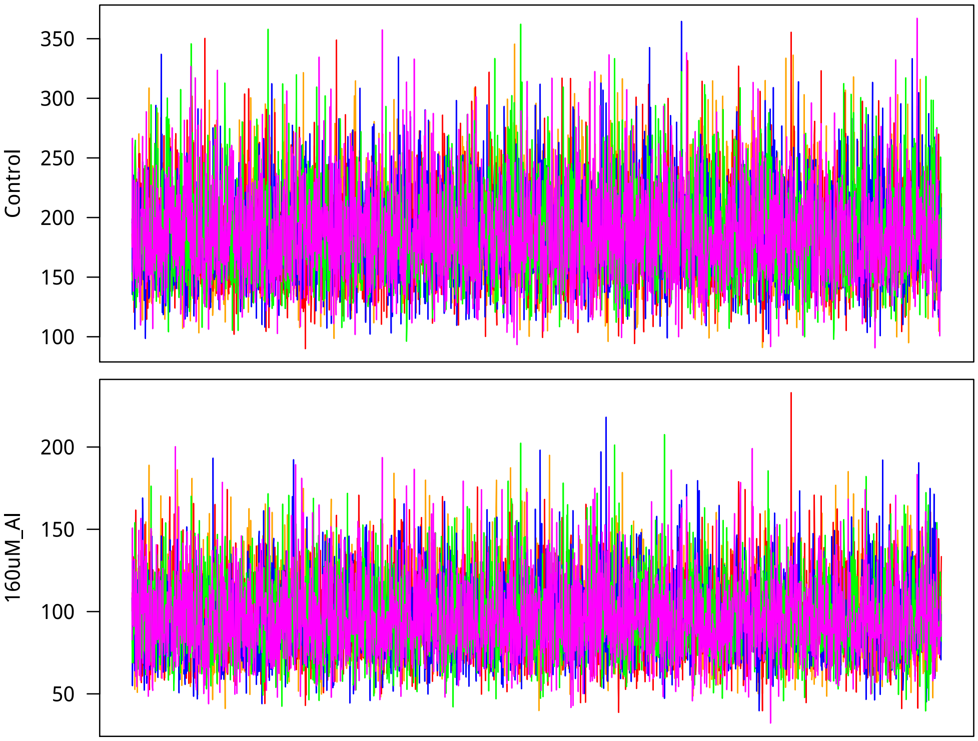

Finally, $\mathbf{\Sigma}_a$ which can not be interpreted as the additive genetic covariance matrix in this case, where $\nu_g = 3$ and the model is Student-$t$.

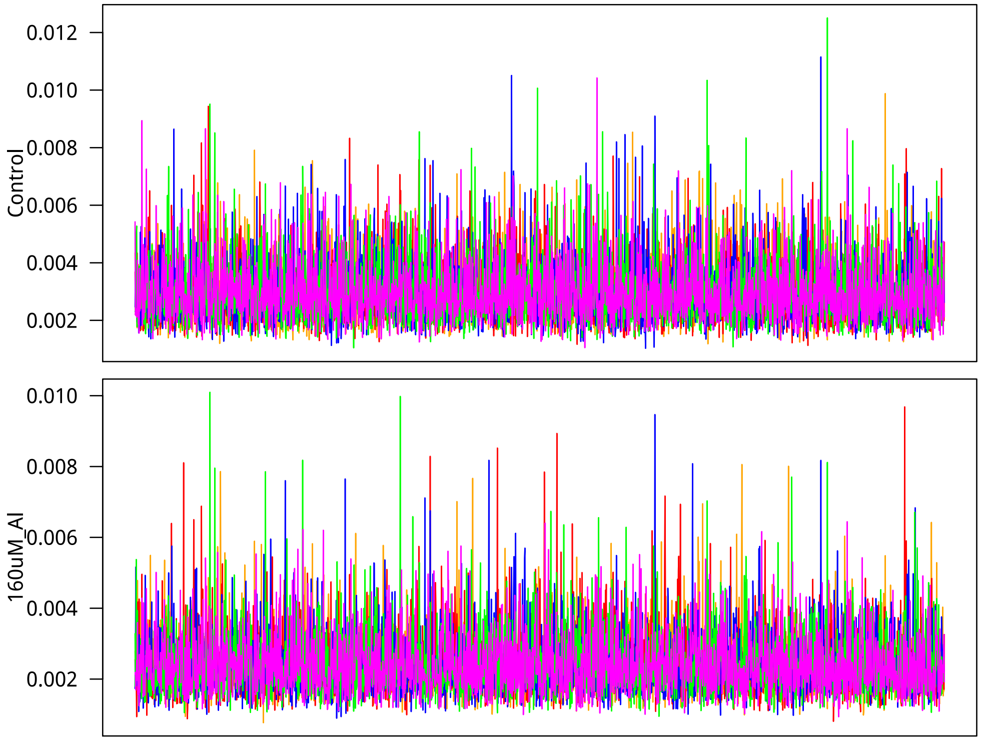


Finally, admixed tropical *japonica* covariances.

```
> chn1 <- matrix(.C("GSLmatLoad",
+     "chains/SgEout_trja_3_1.gbin",
+     as.integer(chnLen), as.integer(d^2), out = double(chnLen*d^2))$out,
+         nrow = chnLen, byrow = T)[,diagInd]
> chn1 <- cbind(chn1,
+     matrix(.C("GSLmatLoad",
+     "chains/SgSout_trja_3_1.gbin",
+     as.integer(chnLen), as.integer(d^2), out = double(chnLen*d^2))$out,
+         nrow = chnLen, byrow = T)[,upInd])
> chn1 <- cbind(chn1,
+     matrix(.C("GSLmatLoad",
+     "chains/SgAout_trja_3_1.gbin",
+     as.integer(chnLen), as.integer(d^2), out = double(chnLen*d^2))$out,
+         nrow = chnLen, byrow = T)[,upInd])
> chn2 <- matrix(.C("GSLmatLoad",
+     "chains/SgEout_trja_3_2.gbin",
+     as.integer(chnLen), as.integer(d^2), out = double(chnLen*d^2))$out,
+         nrow = chnLen, byrow = T)[,diagInd]
> chn2 <- cbind(chn2,
```

```
+     matrix(.C("GSLmatLoad",
+     "chains/SgSout_trja_3_2.gbin",
+     as.integer(chnLen), as.integer(d^2), out = double(chnLen*d^2))$out,
+         nrow = chnLen, byrow = T)[,upInd])
> chn2 <- cbind(chn2,
+     matrix(.C("GSLmatLoad",
+     "chains/SgAout_trja_3_2.gbin",
+     as.integer(chnLen), as.integer(d^2), out = double(chnLen*d^2))$out,
+         nrow = chnLen, byrow = T)[,upInd])
> chn3 <- matrix(.C("GSLmatLoad",
+     "chains/SgEout_trja_3_3.gbin",
+     as.integer(chnLen), as.integer(d^2), out = double(chnLen*d^2))$out,
+         nrow = chnLen, byrow = T)[,diagInd]
> chn3 <- cbind(chn3,
+     matrix(.C("GSLmatLoad",
+     "chains/SgSout_trja_3_3.gbin",
+     as.integer(chnLen), as.integer(d^2), out = double(chnLen*d^2))$out,
+         nrow = chnLen, byrow = T)[,upInd])
> chn3 <- cbind(chn3,
+     matrix(.C("GSLmatLoad",
+     "chains/SgAout_trja_3_3.gbin",
+     as.integer(chnLen), as.integer(d^2), out = double(chnLen*d^2))$out,
+         nrow = chnLen, byrow = T)[,upInd])
> chn4 <- matrix(.C("GSLmatLoad",
+     "chains/SgEout_trja_3_4.gbin",
+     as.integer(chnLen), as.integer(d^2), out = double(chnLen*d^2))$out,
+         nrow = chnLen, byrow = T)[,diagInd]
> chn4 <- cbind(chn4,
+     matrix(.C("GSLmatLoad",
+     "chains/SgSout_trja_3_4.gbin",
+     as.integer(chnLen), as.integer(d^2), out = double(chnLen*d^2))$out,
+         nrow = chnLen, byrow = T)[,upInd])
> chn4 <- cbind(chn4,
+     matrix(.C("GSLmatLoad",
+     "chains/SgAout_trja_3_4.gbin",
+     as.integer(chnLen), as.integer(d^2), out = double(chnLen*d^2))$out,
+         nrow = chnLen, byrow = T)[,upInd])
> chn5 <- matrix(.C("GSLmatLoad",
+     "chains/SgEout_trja_3_5.gbin",
+     as.integer(chnLen), as.integer(d^2), out = double(chnLen*d^2))$out,
+         nrow = chnLen, byrow = T)[,diagInd]
> chn5 <- cbind(chn5,
+     matrix(.C("GSLmatLoad",
+     "chains/SgSout_trja_3_5.gbin",
+     as.integer(chnLen), as.integer(d^2), out = double(chnLen*d^2))$out,
```

```
+         nrow = chnLen, byrow = T)[,upInd])
> chn5 <- cbind(chn5,
+     matrix(.C("GSLmatLoad",
+     "chains/SgAout_trja_3_5.gbin",
+     as.integer(chnLen), as.integer(d^2), out = double(chnLen*d^2))$out,
+         nrow = chnLen, byrow = T)[,upInd])
> acMat   <- mcmcAcf("chn", 5)
> acFrame <- data.frame(autocorrelation=c(acMat),
+     Sigma=rep(c(rep("SgE", length(diagInd)),
+     rep(c("SgS", "SgA"), each=length(upInd))), ncol(acMat)),
+     lag=rep(0:33, each = nrow(acMat)))
> gelRub("chn",nChn,chnLen)
```

```
[1] 0.9999800 0.9998059 0.9999920 1.0001017 1.0002182 0.9998683 0.9997681 0.9998172
```

```
> ggSgE <- ggplot(data=subset(acFrame, Sigma=="SgE"),
+         aes(x=lag, y=autocorrelation)) +
+     geom_point() +
+     theme_classic(base_size=18, base_family="myriad") +
+     labs(title=expression(Sigma[e]))
> ggSgS <- ggplot(data=subset(acFrame, Sigma=="SgS"),
+         aes(x=lag, y=autocorrelation)) +
+     geom_point() +
+     theme_classic(base_size=18, base_family="myriad") +
+     labs(title=expression(Sigma[s]))
> ggSgA <- ggplot(data=subset(acFrame, Sigma=="SgA"),
+         aes(x=lag, y=autocorrelation)) +
+     geom_point() +
+     theme_classic(base_size=18, base_family="myriad") +
+     labs(title=expression(Sigma[a]))
> showtext_auto()
> grid.arrange(ggSgE, ggSgS, ggSgA, nrow = 3)
>
```

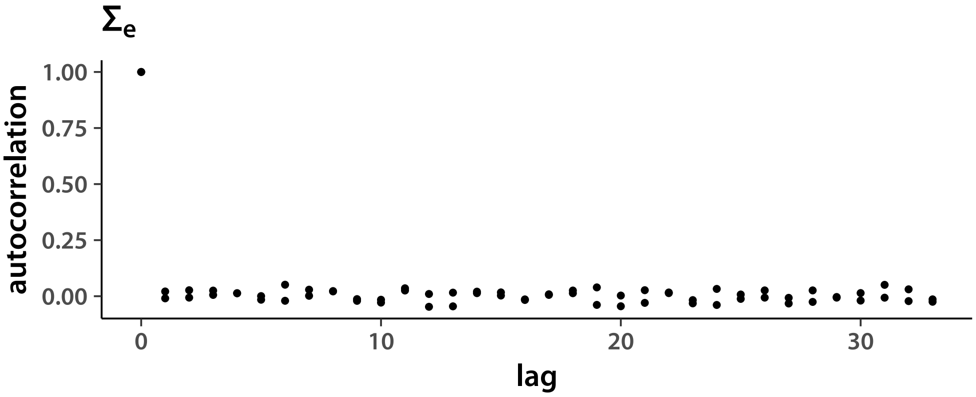


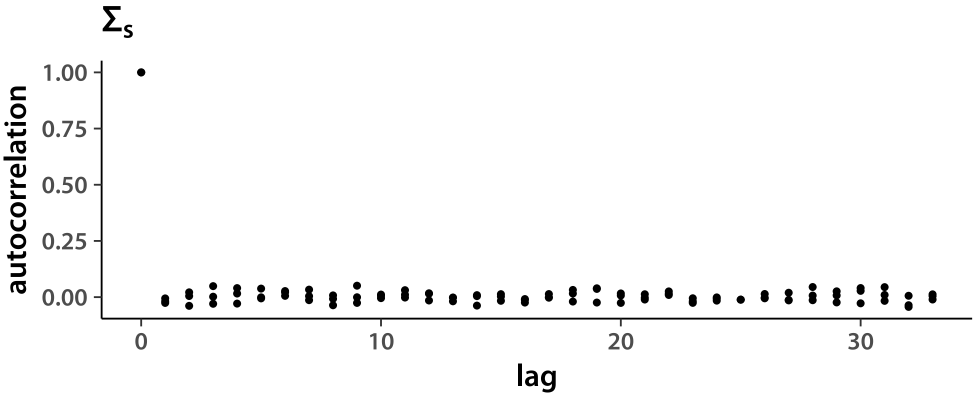


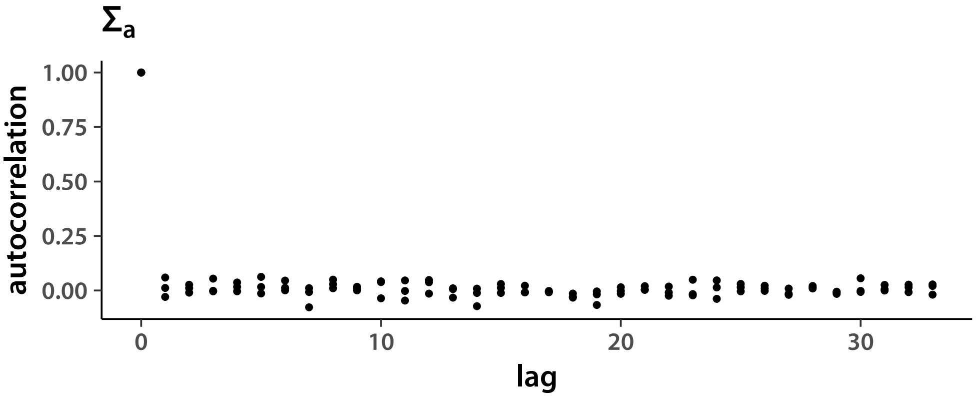


Plot variance chains. First, I get the diagonal index from the upper triangle index.

```
> diagList <- list(1:d, c(3,5), c(6,8))
```

$\Sigma_e$:

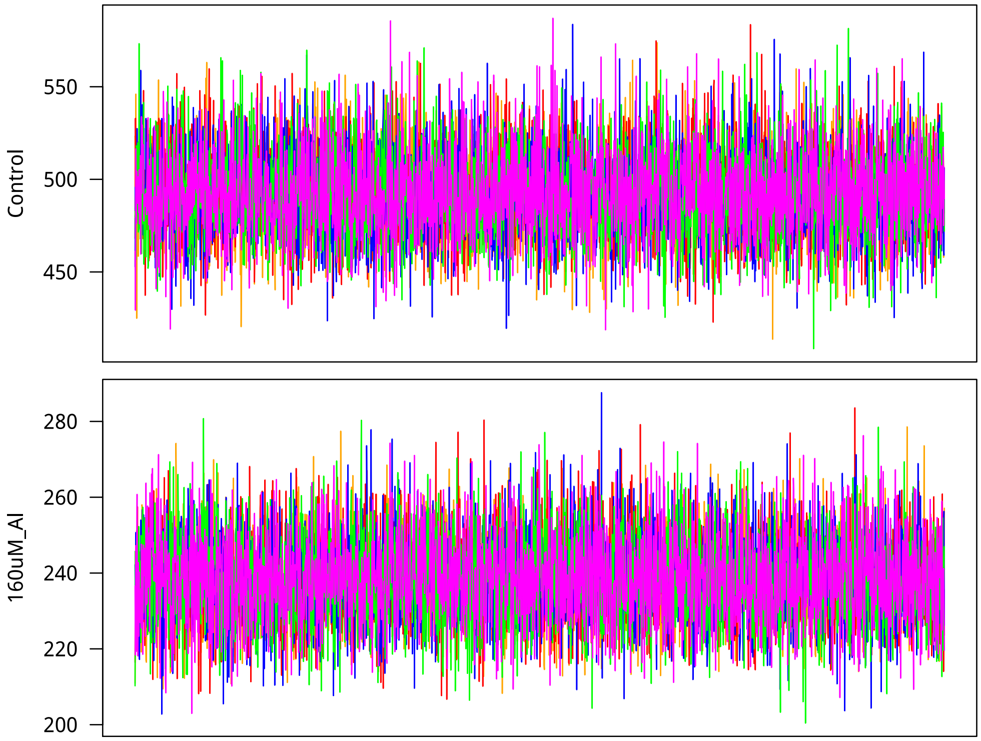


Next, non-additive genetic variances.

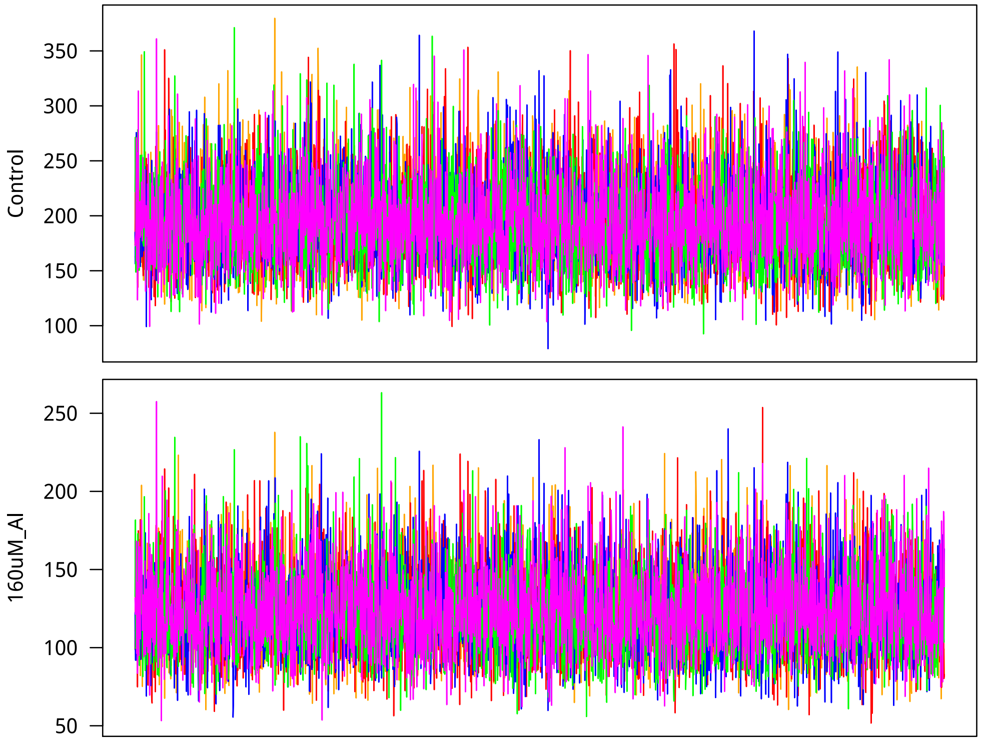

Finally, $\boldsymbol{\Sigma}_a$ which can not be interpreted as the additive genetic covariance matrix in this case, where $\nu_g = 3$ and the model is Student-$t$.

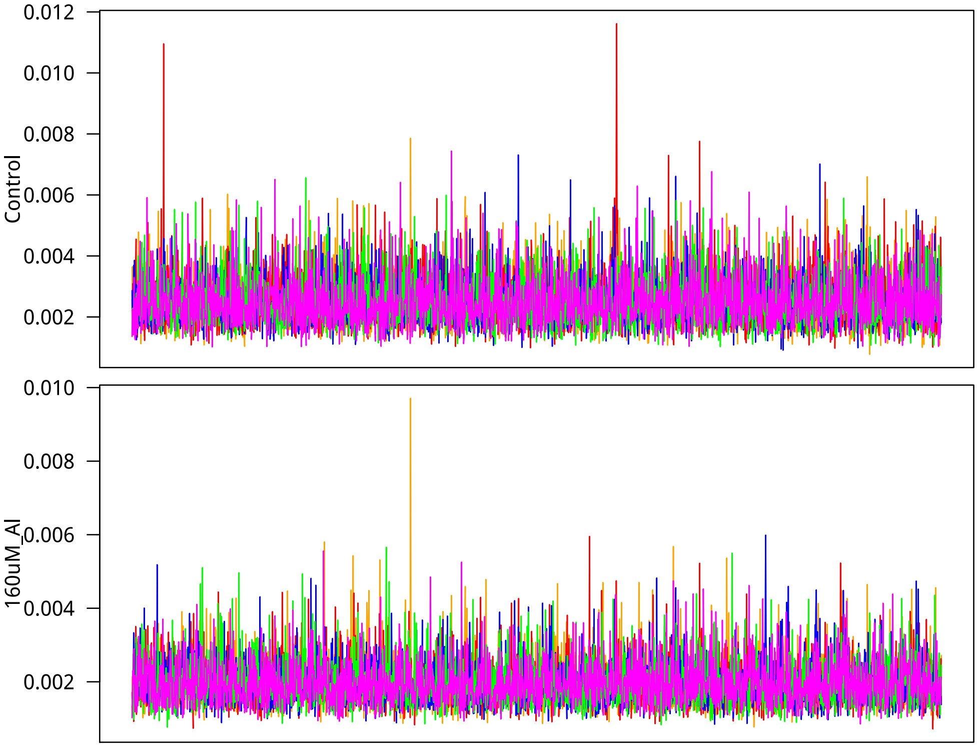
