## Supplemental methods and scripts for "Low additive genetic variation in a trait under selection in domesticated rice": mcmcResultsFAM.pdf

### Markov chain results

Tony Greenberg

March 25, 2019

```
R version 3.5.2 (2018-12-20)
Platform: x86_64-pc-linux-gnu (64-bit)
Running under: Arch Linux

Matrix products: default
BLAS/LAPACK: /opt/intel/compilers_and_libraries_2019.1.144/linux/mkl/lib/intel64_lin/libmkl_gf_lp64.so

locale:
 [1] LC_CTYPE=en_US.UTF-8       LC_NUMERIC=C               LC_TIME=en_US.UTF-8
 [4] LC_COLLATE=en_US.UTF-8     LC_MONETARY=en_US.UTF-8    LC_MESSAGES=en_US.UTF-8
 [7] LC_PAPER=en_US.UTF-8       LC_NAME=C                  LC_ADDRESS=C
[10] LC_TELEPHONE=C             LC_MEASUREMENT=en_US.UTF-8 LC_IDENTIFICATION=C

attached base packages:
[1] compiler  stats     graphics  grDevices utils     datasets  methods   base

other attached packages:
[1] showtext_0.6   showtextdb_2.0 sysfonts_0.8   gridExtra_2.3  ggplot2_3.1.0

loaded via a namespace (and not attached):
 [1] Rcpp_1.0.0       crayon_1.3.4     withr_2.1.2      grid_3.5.2       plyr_1.8.4
 [6] gtable_0.2.0     scales_1.0.0     pillar_1.3.1     rlang_0.3.1      lazyeval_0.2.1
[11] tools_3.5.2      munsell_0.5.0    pkgconfig_2.0.2  colorspace_1.4-0 tibble_2.0.1
```

In this document I summarize the results of modeling the old Al tolerance data using Bayesian hierarchical modeling.

#### 1 Model

I used the following hierarchical model to account for experiments from a number of years.

$$
\begin{aligned}
\boldsymbol{y}_{i\cdot} &\sim \mathrm{N}_d\left(\boldsymbol{\mu}^{acc}_{j[i]\cdot} + \boldsymbol{x}^{\mathrm{EXP}}_{i\cdot}\boldsymbol{B}^{\mathrm{EXP}}; \boldsymbol{\Sigma}_e\right)\\
\boldsymbol{\mu}^{acc}_{j\cdot} &\sim \mathrm{N}_d\left(\boldsymbol{\mu} + \boldsymbol{u}_{j\cdot}\boldsymbol{\Gamma}; \boldsymbol{\Sigma}_s\right)\\
\gamma_{j\cdot} &\sim t_{\nu_g,d}\left(\boldsymbol{0}_d; \boldsymbol{\Sigma}_a\right)\\
\boldsymbol{\Sigma}^{-1}_e &\sim \mathrm{W}_{d,\nu_0}\left(\left[\sum_i \left(\boldsymbol{y}_{i\cdot} - \boldsymbol{\mu}^{acc}_{j[i]\cdot} - \boldsymbol{x}^{\mathrm{EXP}}_{i\cdot}\boldsymbol{B}^{\mathrm{EXP}}\right)\left(\boldsymbol{y}_{i\cdot} - \boldsymbol{\mu}^{acc}_{j[i]\cdot} - \boldsymbol{x}^{\mathrm{EXP}}_{i\cdot}\boldsymbol{B}^{\mathrm{EXP}}\right)^{\mathrm{T}}\right]^{-1}\right)\\
\boldsymbol{\Sigma}^{-1}_s &\sim \mathrm{W}_{d,\nu_0}\left(\left[\sum_j \left(\boldsymbol{\mu}^{acc}_{j\cdot} - \boldsymbol{\mu} - \boldsymbol{u}_{j\cdot}\boldsymbol{\Gamma}\right)\left(\boldsymbol{\mu}^{acc}_{j\cdot} - \boldsymbol{\mu} - \boldsymbol{u}_{j\cdot}\boldsymbol{\Gamma}\right)^{\mathrm{T}}\right]^{-1}\right)\\
\boldsymbol{\Sigma}^{-1}_a &\sim \mathrm{W}_{d,\nu_0}\left(\left[\boldsymbol{\Gamma}^{\mathrm{T}}\boldsymbol{\Gamma}\right]^{-1}\right),
\end{aligned}
$$

where bold lower-case symbols refer to row vectors and bold upper-case symbols are matrices. All vectors are of length $d$ and all matrices have $d$ columns, where $d$ is the number of traits ($d = 2$ in our case). $\boldsymbol{X}^{\mathrm{EXP}}$ is the matrix of experiment IDs and $\boldsymbol{B}^{\mathrm{EXP}}$ contains experimental effect coefficients. These are essentially years, but see the `phenoDataPrep` documents for details. The $\boldsymbol{U}$ matrix contains principal components of the genetic relationship matrix, scaled by square roots of their eigenvalues. The $\boldsymbol{U\Gamma}$ matrix is the matrix of GEBVs (marker breeding values). The degrees of freedom parameter $\nu_g$ was set to 3, the smallest value that results in a distribution with all moments defined. Notation of the $i\cdot$ type refers to rows in matrices. Subscripts of the $j[i]$ type refer to a row $j$ in a upper level (in the model hierarchy) that corresponds to a replicate row $i$. Note also that the residual covariance matrix $\boldsymbol{\Sigma}^{-1}_e$ is diagonal, since the experiment and control phenotypes were measured on different plants and thus the residual correlation is undefined. Location parameters without priors were modeled with very high-variance Gaussian priors.

I define functions that calculate summaries of Markov chains.

```
> pmode <- cmpfun(function(vec){
+     dst <- density(vec, adjust = 2)
+     mxi <- which(dst$y == max(dst$y))
+     if(length(mxi) > 1){
+         warning("More than one mode in call to pmode(); picking randomly")
+         mxi <- sample(mxi, 1)
+     }
+     dst$x[mxi]
+ })
> HPDint <- cmpfun(function(vec, prob = 0.95){
+     nsamp <- length(vec)
+     if (nsamp <= 2) stop("vector must have length > 2")
+
+     vals <- sort(vec)
+     gap  <- max(1, min(nsamp - 1, round(nsamp * prob)))
+     init <- 1:(nsamp - gap)
+     mInd <- which.min(vals[init + gap] - vals[init])
```

```
+     res  <- c(vals[mInd], vals[mInd + gap])
+     names(res) <- c("lower", "upper")
+     return(res)
+ })
```

Next, I write a function that is similar to `quantile()`, but outputs the mode, lower and upper limits of the 95% and 50% HPD. The `outr` parameter is the probability for the outer margins, default is 95%.

```
> quantileLike <- cmpfun(function(vec, outr = 0.95){
+     if(outr <= 0.5) stop("Outer margin has to be >= 50%")
+     md    <- pmode(vec)
+     hpd95 <- HPDint(vec, outr)
+     hpd50 <- HPDint(vec, 0.5)
+     res   <- c(hpd95[1], hpd50[1], md, hpd50[2], hpd95[2])
+
+     outNm <- paste(c("lower", "upper"), outr*100, sep = "")
+     names(res) <- c(outNm[1], "lower50", "mode", "upper50", outNm[2])
+     return(res)
+ })
```

#### 2 Experiment effect

I start by checking the experiment effect. Start with the *aus* results.

I start with total accession means.

```
> trtNam <- c("Control", "Treated")
> d      <- 2
> Nexp   <- 6
> expDim <- d*Nexp
> nChn   <- 5
> chnLen <- 2000
> chn <- NULL
> addSamp <- cmpfun(function(i, vrNam, popNam, ncl){
+     inFlNam <- paste("chains/", vrNam, "out_", popNam, "_3_", i, ".gbin", sep = "")
+     chn <<- rbind(chn,
+                 matrix(.C("GSLmatLoad",
+                     inFlNam, as.integer(chnLen), as.integer(ncl),
+                     out = double(chnLen*ncl))$out,
+                 nrow = chnLen, byrow = T)
+         )
+     return(NULL)
+ })
> trash <- sapply(1:nChn, addSamp, "EXP", "aus", expDim)
> expID               <- scan(file = "expIDs.txt", what=character())
> expMnAUS            <- as.data.frame(t(apply(chn, 2, quantileLike)))
```

```
> expMnAUS$trait      <- rep(trtNam, times = Nexp)
> expMnAUS$experiment <- rep(expID, each=d)

> ggTrt1 <- ggplot(data=subset(expMnAUS, trait==trtNam[1]), aes(x=experiment,y=mode)) +
+     geom_segment(aes(x=experiment, y=lower95, xend=experiment, yend=upper95),
+         color="grey80", size=0.75) +
+     geom_segment(aes(x=experiment, y=lower50, xend=experiment, yend=upper50),
+         color="grey50", size=1) +
+     geom_point() +
+     theme_classic(base_size=18, base_family="myriad") +
+     theme(axis.title.x=element_blank(), axis.ticks.x=element_blank()) +
+     labs(y=trtNam[1])
> ggTrt2 <- ggplot(data=subset(expMnAUS, trait==trtNam[2]), aes(x=experiment,y=mode)) +
+     geom_segment(aes(x=experiment, y=lower95, xend=experiment, yend=upper95),
+         color="grey80", size=0.75) +
+     geom_segment(aes(x=experiment, y=lower50, xend=experiment, yend=upper50),
+         color="grey50", size=1) +
+     geom_point() +
+     theme_classic(base_size=18, base_family="myriad") +
+     theme(axis.ticks.x=element_blank()) +
+     labs(y=trtNam[2])
> showtext_auto()
> grid.arrange(ggTrt1, ggTrt2, nrow = 2)
```

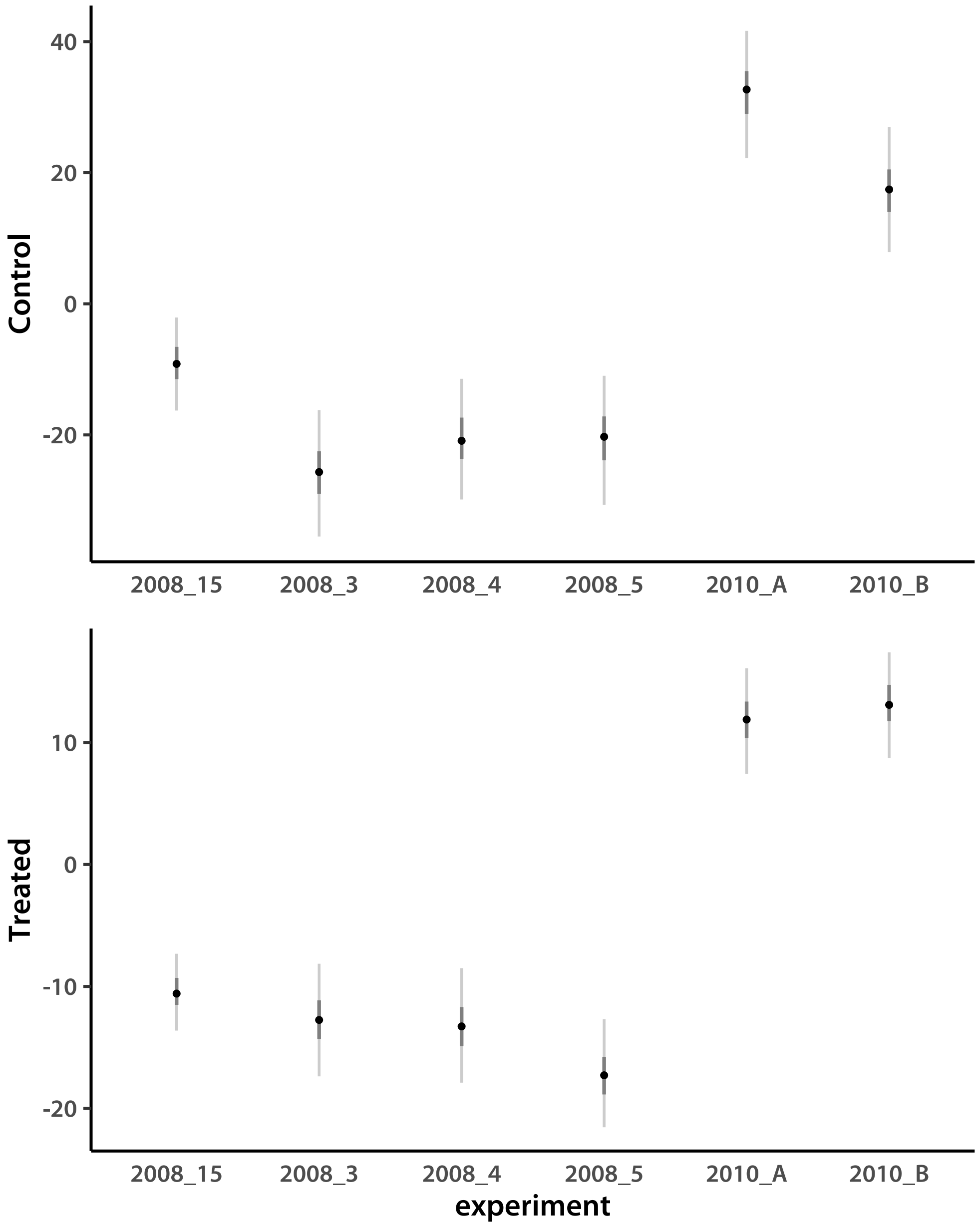


The estimates look reasonable compared to plots of raw data. I look at the tropical *japonica* results.

```
> chn <- NULL
> trash <- sapply(1:nChn, addSamp, "EXP", "trj", expDim)
> expMnTRJ            <- as.data.frame(t(apply(chn, 2, quantileLike)))
> expMnTRJ$trait      <- rep(trtNam, times = Nexp)
> expMnTRJ$experiment <- rep(expID, each=d)

> ggTrt1 <- ggplot(data=subset(expMnTRJ, trait==trtNam[1]), aes(x=experiment,y=mode)) +
+     geom_segment(aes(x=experiment, y=lower95, xend=experiment, yend=upper95),
```

```
+         color="grey80", size=0.75) +
+     geom_segment(aes(x=experiment, y=lower50, xend=experiment, yend=upper50),
+         color="grey50", size=1) +
+     geom_point() +
+     theme_classic(base_size=18, base_family="myriad") +
+     theme(axis.title.x=element_blank(), axis.ticks.x=element_blank()) +
+     labs(y=trtNam[1])
> ggTrt2 <- ggplot(data=subset(expMnTRJ, trait==trtNam[2]), aes(x=experiment,y=mode)) +
+     geom_segment(aes(x=experiment, y=lower95, xend=experiment, yend=upper95),
+         color="grey80", size=0.75) +
+     geom_segment(aes(x=experiment, y=lower50, xend=experiment, yend=upper50),
+         color="grey50", size=1) +
+     geom_point() +
+     theme_classic(base_size=18, base_family="myriad") +
+     theme(axis.ticks.x=element_blank()) +
+     labs(y=trtNam[2])
> showtext_auto()
> grid.arrange(ggTrt1, ggTrt2, nrow = 2)
```

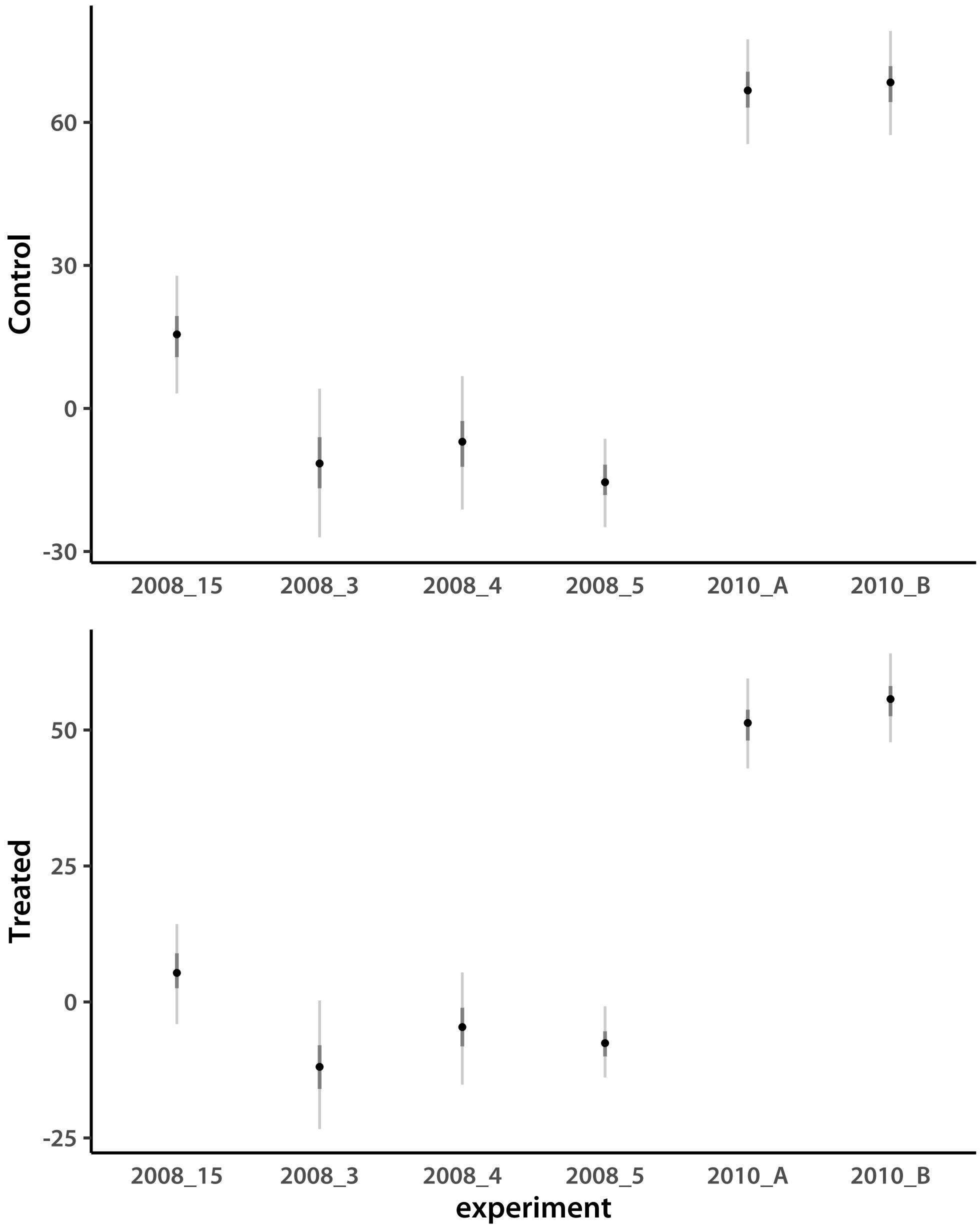


Results are very similar to *aus*. Finally, I look at the admixed tropical *japonica* results.

```
> chn <- NULL
> trash <- sapply(1:nChn, addSamp, "EXP", "trja", expDim)
> expMnTRJA            <- as.data.frame(t(apply(chn, 2, quantileLike)))
> expMnTRJA$trait      <- rep(trtNam, times = Nexp)
> expMnTRJA$experiment <- rep(expID, each=d)

> ggTrt1 <- ggplot(data=subset(expMnTRJA, trait==trtNam[1]), aes(x=experiment,y=mode)) +
+     geom_segment(aes(x=experiment, y=lower95, xend=experiment, yend=upper95),
```

```
+         color="grey80", size=0.75) +
+     geom_segment(aes(x=experiment, y=lower50, xend=experiment, yend=upper50),
+         color="grey50", size=1) +
+     geom_point() +
+     theme_classic(base_size=18, base_family="myriad") +
+     theme(axis.title.x=element_blank(), axis.ticks.x=element_blank()) +
+     labs(y=trtNam[1])
> ggTrt2 <- ggplot(data=subset(expMnTRJA, trait==trtNam[2]), aes(x=experiment,y=mode)) +
+     geom_segment(aes(x=experiment, y=lower95, xend=experiment, yend=upper95),
+         color="grey80", size=0.75) +
+     geom_segment(aes(x=experiment, y=lower50, xend=experiment, yend=upper50),
+         color="grey50", size=1) +
+     geom_point() +
+     theme_classic(base_size=18, base_family="myriad") +
+     theme(axis.ticks.x=element_blank()) +
+     labs(y=trtNam[2])
> showtext_auto()
> grid.arrange(ggTrt1, ggTrt2, nrow = 2)
```

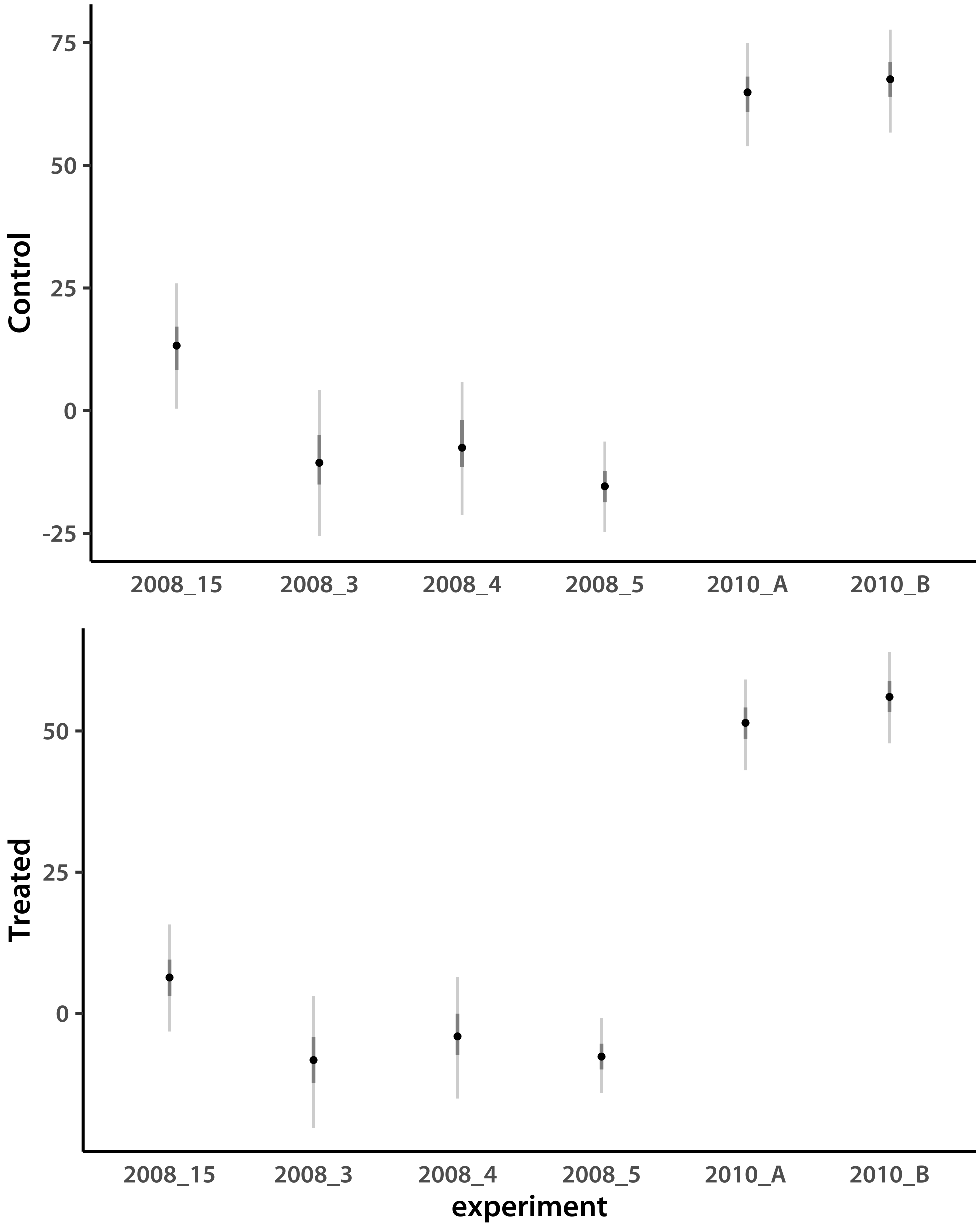


#### 3 Accession means

Next, I look at accession means. Their variance reflects broad-sense heritability. Starting with *aus*.

```
> Nacc   <- 55
> accDim <- d*Nacc
> chn <- NULL
> trash <- sapply(1:nChn, addSamp, "LN", "aus", accDim)
> accMnAUS       <- as.data.frame(t(apply(chn, 2, quantileLike)))
```

```
> accMnAUS$trait <- rep(trtNam, times = Nacc)
> accMnAUSs      <- accMnAUS[order(accMnAUS[,6], accMnAUS[,3]),]
```

I calculate the relative root growth and check if all values are positive.

```
> accNamAUS     <- scan(file="lineIDsAUS.txt", what=character())
> colnames(chn) <- paste(rep(trtNam, Nacc), rep(accNamAUS, each=d), sep=".")
> chnRRG        <- chn[,grep(trtNam[2], colnames(chn))]/chn[,grep(trtNam[1], colnames(chn))]
> sum(chnRRG<=0)

[1] 0
```

They are. I calculate the $-\log_{10}(RRG)$ and save the modes of accession effect estimates for further analyses.

```
> rrgModeAUS    <- apply(-log10(chnRRG), 2, pmode)
> accModeTabAUS <- cbind(matrix(round(accMnAUS$mode, 4), ncol=d, byrow=T),
+     round(rrgModeAUS, 4))
> cat(c("NSFTVID\tControl\t160uM_Al\tneglgRRG",
+         apply(cbind(accNamAUS, accModeTabAUS), 1, paste, collapse="\t")),
+     file="accModeAUS.tsv",sep="\n"
+     )
> trash <- .C("GSLmatSave", "phenoData/accMdAUS.gbin",
+     as.double(accModeTabAUS), nrow(accModeTabAUS), ncol(accModeTabAUS))
```

I plot sorted accession means.

```
> pdfFlNam <- "lineMeansFAMaus.pdf"
> showtext_auto()
> ggplot(data=accMnAUSs, aes(x=1:nrow(accMnAUSs),y=mode)) +
+     geom_segment(aes(x=1:nrow(accMnAUSs), y=lower95, xend=1:nrow(accMnAUSs), yend=upper95),
+         color="grey80", size=0.75) +
+     geom_segment(aes(x=1:nrow(accMnAUSs), y=lower50, xend=1:nrow(accMnAUSs), yend=upper50),
+         color="grey50", size=1) +
+     geom_point() +
+     facet_wrap(~trait, scales="free", nrow=2) +
+     theme_classic(base_size=18, base_family="myriad") +
+     theme(axis.title.x=element_blank(), axis.text.x=element_blank(),
+         strip.background=element_rect(fill="grey95", linetype="blank"),
+         axis.ticks.x=element_blank()) +
+     labs(y="root length")
> ggsave(pdfFlNam, width=8, height=10, units="in", device="pdf", useDingbats=F)
> cat("\\includegraphics{", pdfFlNam, "}\n\n", sep="")
```

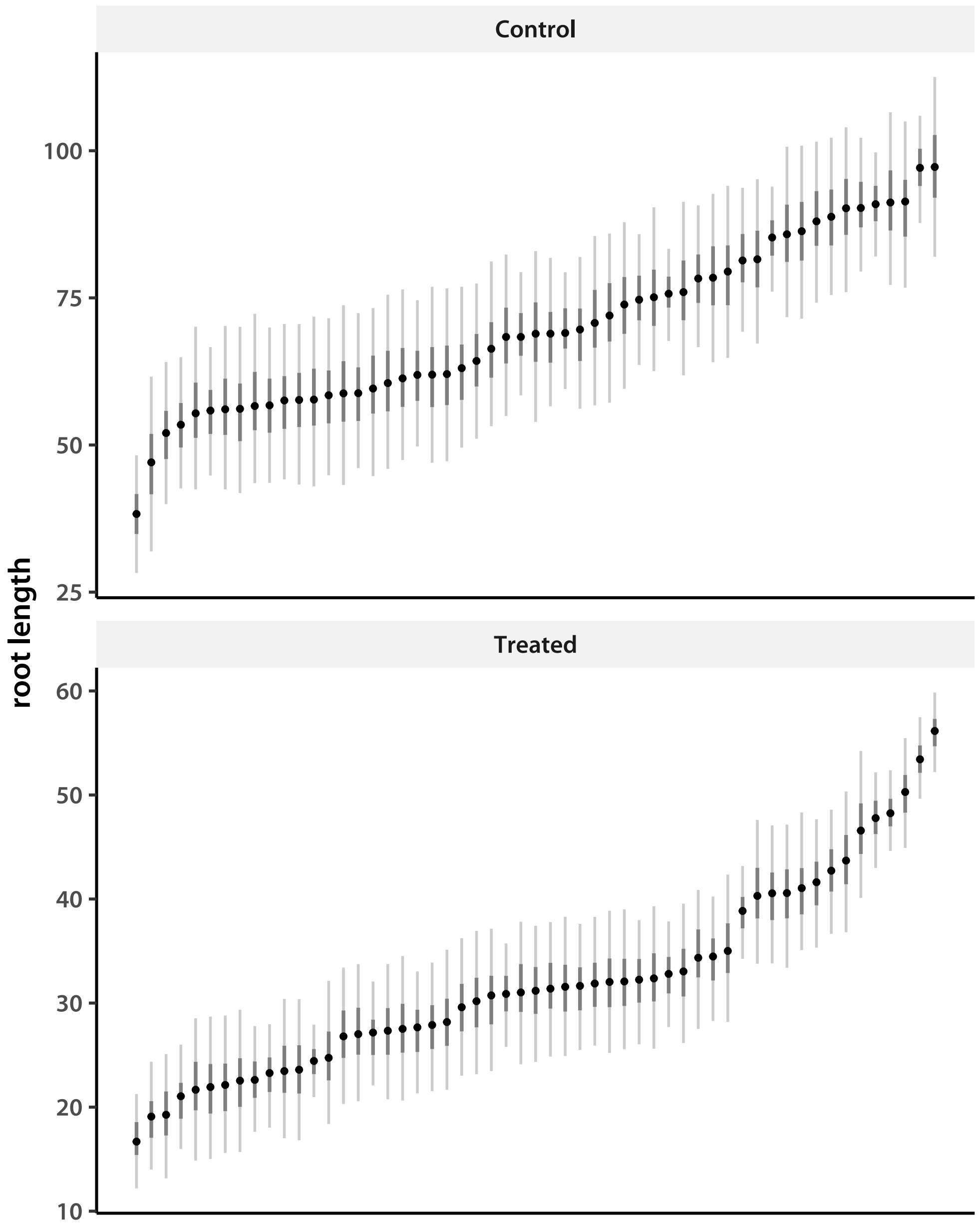


Seems like there are some particularly resistant accessions, which is expected when treatment has the desired stress effect. Now look at the tropical *japonica*.

```
> Nacc   <- 92
> accDim <- d*Nacc
> chn <- NULL
> trash <- sapply(1:nChn, addSamp, "LN", "trj", accDim)
> accMnTRJ       <- as.data.frame(t(apply(chn, 2, quantileLike)))
> accMnTRJ$trait <- rep(trtNam, times = Nacc)
> accMnTRJs      <- accMnTRJ[order(accMnTRJ[,6], accMnTRJ[,3]),]
```

I calculate the relative root growth and check if all values are positive.

```
> accNamTRJ     <- scan(file="lineIDsTRJ.txt", what=character())
> colnames(chn) <- paste(rep(trtNam, Nacc), rep(accNamTRJ, each=d), sep=".")
> chnRRG        <- chn[,grep(trtNam[2], colnames(chn))]/chn[,grep(trtNam[1], colnames(chn))]
> sum(chnRRG<=0)
```

```
[1] 11652
```

There are negative values, due to correction for the experiment effect. I subtract the smallest value to make everything positive.

```
> chnRRG <- (chn[,grep(trtNam[2], colnames(chn))]-min(chn)+1e-4)/
+     (chn[,grep(trtNam[1], colnames(chn))]-min(chn)+1e-4)
> sum(chnRRG<=0)
```

```
[1] 0
```

I calculate the $-\log_{10}(RRG)$ and save the modes of accession effect estimates for further analyses.

```
> rrgModeTRJ    <- apply(-log10(chnRRG), 2, pmode)
> accModeTabTRJ <- cbind(matrix(round(accMnTRJ$mode, 4), ncol=d, byrow=T),
+     round(rrgModeTRJ, 4))
> cat(c("NSFTVID\tControl\t160uM_Al\tneglgRRG",
+         apply(cbind(accNamTRJ, accModeTabTRJ), 1, paste, collapse="\t")),
+     file="accModeTRJ.tsv",sep="\n"
+     )
> trash <- .C("GSLmatSave", "phenoData/accMdTRJ.gbin",
+     as.double(accModeTabTRJ), nrow(accModeTabTRJ), ncol(accModeTabTRJ))
```

I plot sorted accession means.

```
> pdfFlNam <- "lineMeansFAMtrj.pdf"
> showtext_auto()
> ggplot(data=accMnTRJs, aes(x=1:nrow(accMnTRJs),y=mode)) +
+     geom_segment(aes(x=1:nrow(accMnTRJs), y=lower95, xend=1:nrow(accMnTRJs), yend=upper95),
+         color="grey80", size=0.75) +
+     geom_segment(aes(x=1:nrow(accMnTRJs), y=lower50, xend=1:nrow(accMnTRJs), yend=upper50),
+         color="grey50", size=1) +
+     geom_point() +
+     facet_wrap(~trait, scales="free", nrow=2) +
+     theme_classic(base_size=18, base_family="myriad") +
+     theme(axis.title.x=element_blank(), axis.text.x=element_blank(),
+         strip.background=element_rect(fill="grey95", linetype="blank"),
+         axis.ticks.x=element_blank()) +
+     labs(y="root length")
> ggsave(pdfFlNam, width=8, height=10, units="in", device="pdf", useDingbats=F)
> cat("\\includegraphics{", pdfFlNam, "}\n\n", sep="")
```

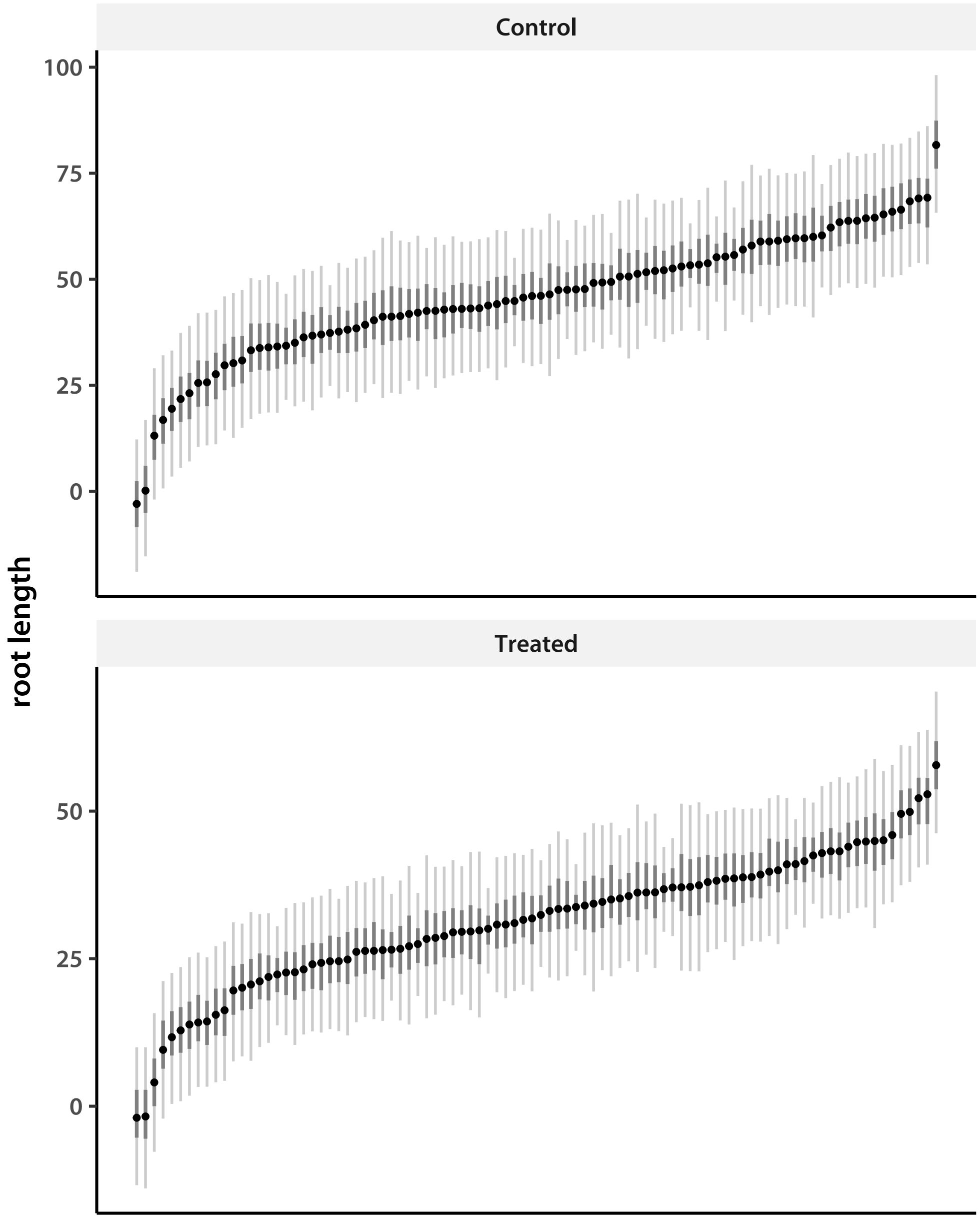


The distributions are very similar. Finally, look at the admixed tropical *japonica*.

```
> Nacc   <- 95
> accDim <- d*Nacc
> chn <- NULL
> trash <- sapply(1:nChn, addSamp, "LN", "trja", accDim)
> accNamTRJA      <- scan(file="lineIDsTRJA.txt", what=character())
> accNamADM       <- scan(file="lineIDsADM.txt", what=character())
> accMnTRJA       <- as.data.frame(t(apply(chn, 2, quantileLike)))
> accMnTRJA$trait <- rep(trtNam, times = Nacc)
```

```
> accMnTRJA$admix <- rep(ifelse(accNamTRJA %in% accNamADM, "admixed", "pure"), each=d)
> accMnTRJAs      <- accMnTRJA[order(accMnTRJA[,6], accMnTRJA[,3]),]
```

Save the modes to a text file.

```
> accModeTabTRJA <- matrix(round(accMnTRJA$mode, 4), ncol=d, byrow=T)
> cat(c("NSFTVID\tControl\t160uM_Al",
+         apply(cbind(accNamTRJA, accModeTabTRJA), 1, paste, collapse="\t")),
+     file="accModeTRJA.tsv",sep="\n"
+     )
>
```

I plot sorted accession means.

```
> pdfFlNam <- "lineMeansFAMtrja.pdf"
> showtext_auto()
> ggplot(data=accMnTRJAs, aes(x=1:nrow(accMnTRJAs),y=mode)) +
+     geom_segment(aes(x=1:nrow(accMnTRJAs), y=lower95, xend=1:nrow(accMnTRJAs), yend=upper95)
+         color="grey80", size=0.75) +
+     geom_segment(aes(x=1:nrow(accMnTRJAs), y=lower50,
+                 xend=1:nrow(accMnTRJAs), yend=upper50, color=admix),size=1.5) +
+     geom_point(size=1.5, color="grey60") +
+     facet_wrap(~trait, scales="free", nrow=2) +
+     theme_classic(base_size=18, base_family="myriad") +
+     theme(axis.title.x=element_blank(), axis.text.x=element_blank(),
+         strip.background=element_rect(fill="grey95", linetype="blank"),
+         axis.ticks.x=element_blank(), legend.title=element_blank()) +
+     labs(y="root length")
> ggsave(pdfFlNam, width=8, height=10, units="in", device="pdf", useDingbats=F)
> cat("\\includegraphics{", pdfFlNam, "}\n\n", sep="")
```

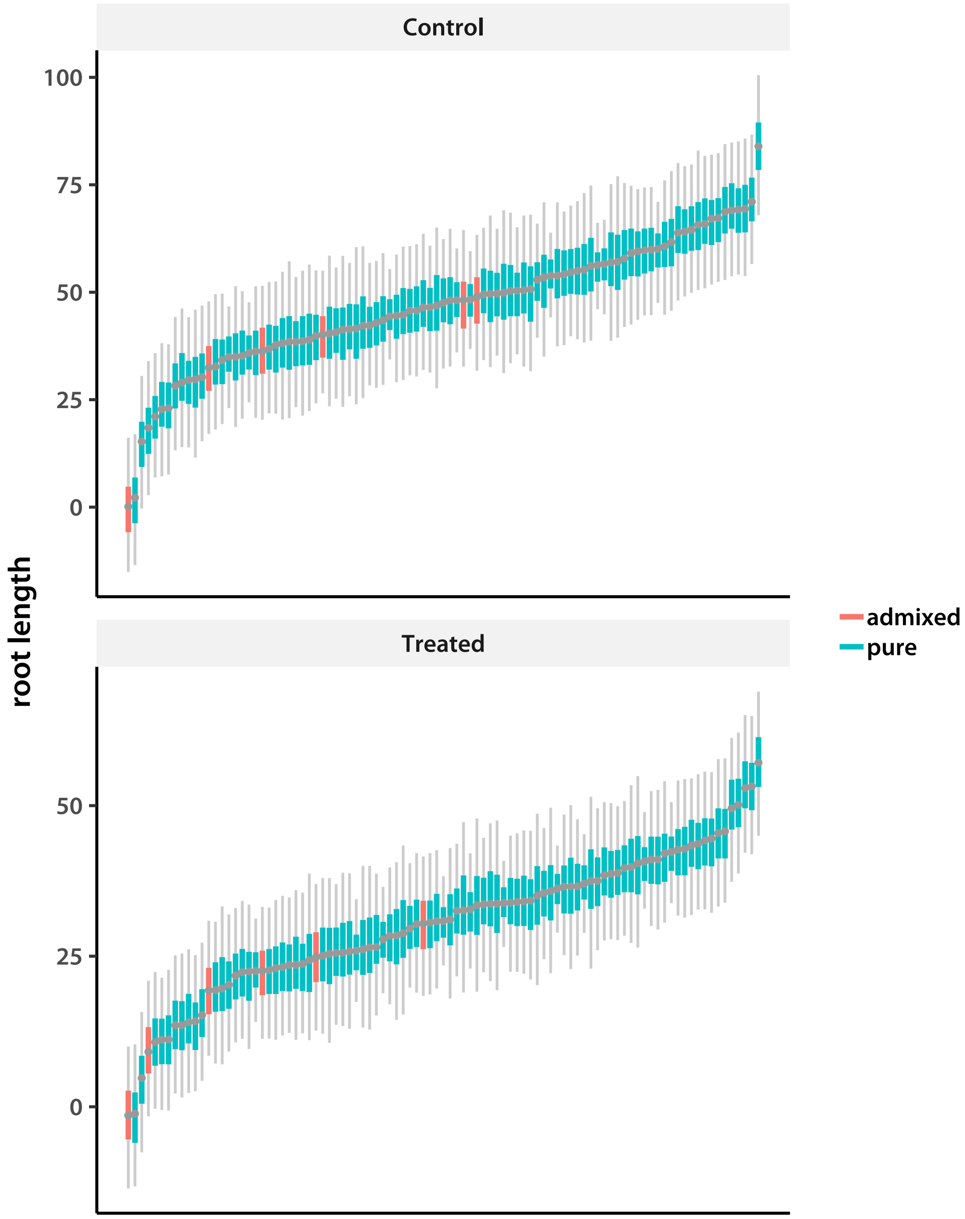


The admixed lines have smaller roots even in control, and even smaller in treatment (compared to the average tropical *japonica*). I test this more formally by estimating mean rank of the admixed accessions, compared to the overall mean (which is $(1 + N_{acc})/2$).

```
> admInd <- accNamTRJA %in% accNamADM
> mnRank <- cmpfun(function(vec, ind){
+     colMeans(apply(matrix(vec, ncol=2, byrow=T), 2, rank)[ind,])
+ })
> rankChn         <- t(apply(chn, 1, mnRank, admInd))
> rankMnTRJ       <- as.data.frame(t(apply(rankChn, 2, quantileLike)))
```

```
> rankMnTRJ$trait <- trtNam
> rankMnTRJ
```

```
   lower95  lower50     mode  upper50  upper95   trait
1 20.00000 25.83333 28.93652 32.50000 39.66667 Control
2 14.33333 18.16667 21.42600 23.66667 29.83333 Treated
```

Plot the ranks.

```
> pdfFlNam <- "rankAdmLineMeans.pdf"
> showtext_auto()
> ggplot(data=rankMnTRJ, aes(x=1:d,y=mode)) +
+     geom_segment(aes(x=1:d, y=lower95, xend=1:d, yend=upper95),
+         color="grey80", size=1.25) +
+     geom_segment(aes(x=1:d, y=lower50, xend=1:d, yend=upper50),
+         color="grey50", size=2) +
+     geom_point() +
+     geom_hline(yintercept=(mean(1:Nacc)), color="grey50") +
+     theme_classic(base_size=18, base_family="myriad") +
+     theme(axis.title.x=element_blank(), axis.text.x=element_blank(),
+         strip.background=element_rect(fill="grey95", linetype="blank"),
+         axis.ticks.x=element_blank()) +
+     labs(y="rank") + xlim(0.5, d+0.5) +
+     annotate("text", label="Control", x=1, y=min(rankMnTRJ$lower95)-2) +
+     annotate("text", label="Al treated", x=2, y=min(rankMnTRJ$lower95)-2)
> ggsave(pdfFlNam, width=8, height=8, units="in", device="pdf", useDingbats=F)
> cat("\\includegraphics{", pdfFlNam, "}\n\n", sep="")
```

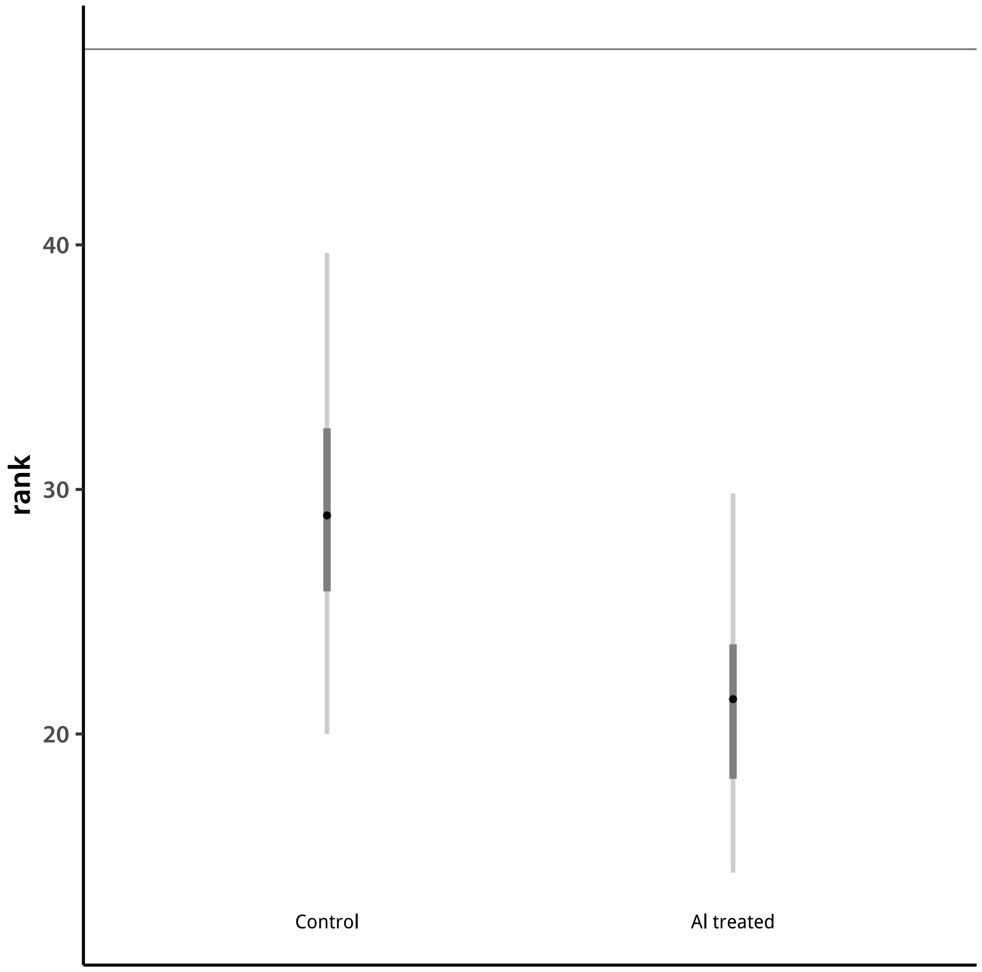


### 4 GEBV

I next examine the distributions of genome (or SNP) estimated breeding values (GEBV).

```
> Nacc   <- 55
> accDim <- d*Nacc
> chn        <- NULL
> trash      <- sapply(1:nChn, addSamp, "BV", "aus", accDim)
> gebvChnAUS <- chn
> gebvMnAUS       <- as.data.frame(t(apply(chn, 2, quantileLike)))
> gebvMnAUS$trait <- rep(trtNam, times = Nacc)
> gebvMnAUSs      <- gebvMnAUS[order(gebvMnAUS[,6], gebvMnAUS[,3]),]
```

I plot sorted GEBVs.

```
> pdfFlNam <- "gebvFAMaus.pdf"
> showtext_auto()
```

```
> ggplot(data=gebvMnAUSs, aes(x=1:nrow(gebvMnAUSs),y=mode)) +
+     geom_segment(aes(x=1:nrow(gebvMnAUSs), y=lower95, xend=1:nrow(gebvMnAUSs), yend=upper95)
+         color="grey80", size=0.75) +
+     geom_segment(aes(x=1:nrow(gebvMnAUSs), y=lower50, xend=1:nrow(gebvMnAUSs), yend=upper50)
+         color="grey50", size=1) +
+     geom_point() +
+     facet_wrap(~trait, scales="free", nrow=2) +
+     theme_classic(base_size=18, base_family="myriad") +
+     theme(axis.title.x=element_blank(), axis.text.x=element_blank(),
+         strip.background=element_rect(fill="grey95", linetype="blank"),
+         axis.ticks.x=element_blank()) +
+     labs(y="root length")
> ggsave(pdfFlNam, width=8, height=10, units="in", device="pdf", useDingbats=F)
> cat("\\includegraphics{", pdfFlNam, "}\n\n", sep="")
```

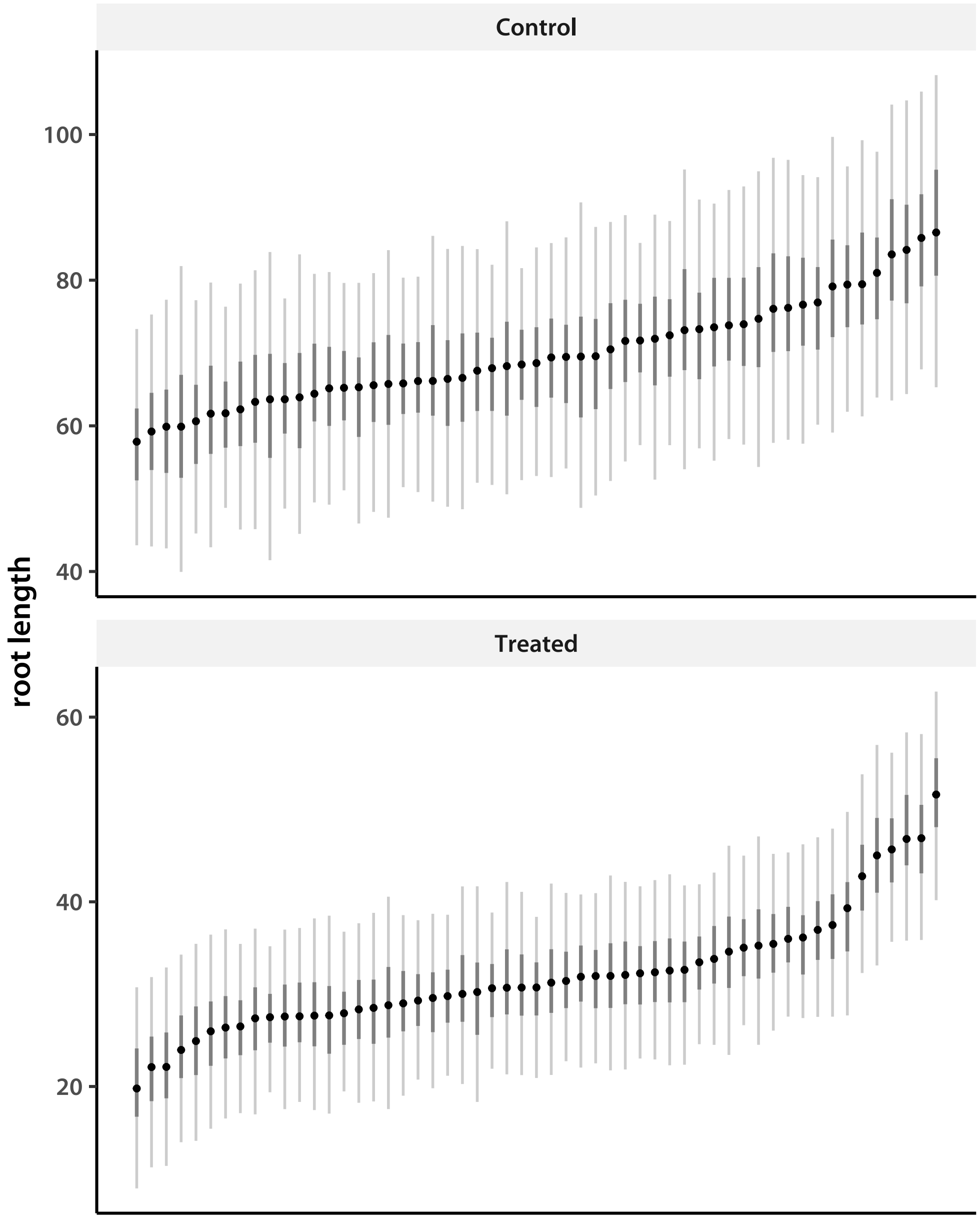


Again, seems like some breeding values stand out as high in the treated group. Now look at the tropical *japonica*.

```
> Nacc   <- 92
> accDim <- d*Nacc
> chn <- NULL
> trash <- sapply(1:nChn, addSamp, "BV", "trj", accDim)
> gebvChnTRJ <- chn
> gebvMnTRJ       <- as.data.frame(t(apply(chn, 2, quantileLike)))
> gebvMnTRJ$trait <- rep(trtNam, times = Nacc)
```

```
> gebvMnTRJs      <- gebvMnTRJ[order(gebvMnTRJ[,6], gebvMnTRJ[,3]),]
```

I plot sorted accession means.

```
> pdfFlNam <- "gebvFAMtrj.pdf"
> showtext_auto()
> ggplot(data=gebvMnTRJs, aes(x=1:nrow(gebvMnTRJs),y=mode)) +
+     geom_segment(aes(x=1:nrow(gebvMnTRJs), y=lower95, xend=1:nrow(gebvMnTRJs), yend=upper95)
+         color="grey80", size=0.75) +
+     geom_segment(aes(x=1:nrow(gebvMnTRJs), y=lower50, xend=1:nrow(gebvMnTRJs), yend=upper50)
+         color="grey50", size=1) +
+     geom_point() +
+     facet_wrap(~trait, scales="free", nrow=2) +
+     theme_classic(base_size=18, base_family="myriad") +
+     theme(axis.title.x=element_blank(), axis.text.x=element_blank(),
+         strip.background=element_rect(fill="grey95", linetype="blank"),
+         axis.ticks.x=element_blank()) +
+     labs(y="root length")
> ggsave(pdfFlNam, width=8, height=10, units="in", device="pdf", useDingbats=F)
> cat("\\includegraphics{", pdfFlNam, "}\n\n", sep="")
```

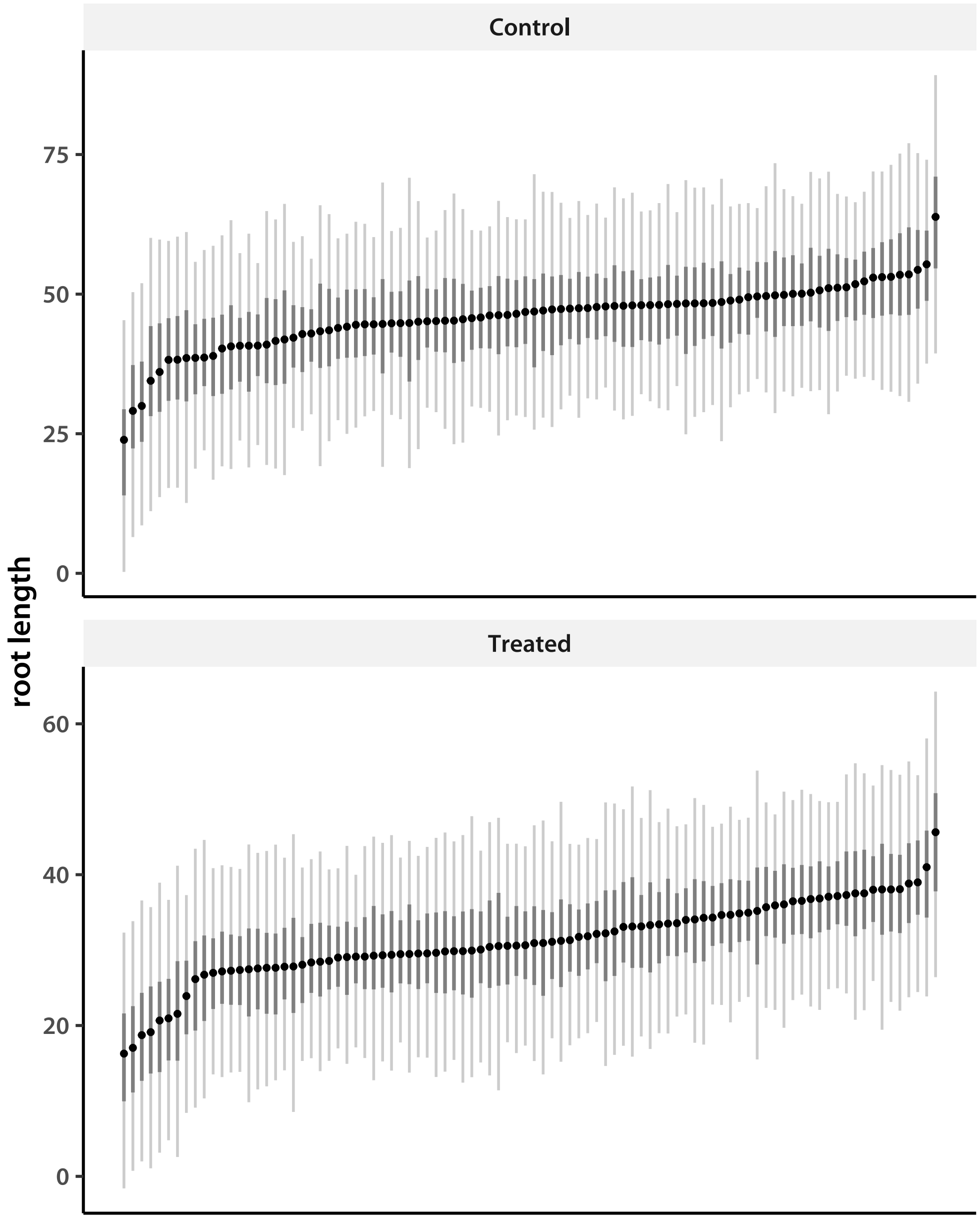


Finally, look at the admixed tropical *japonica*.

```
> Nacc   <- 95
> accDim <- d*Nacc
> chn <- NULL
> trash <- sapply(1:nChn, addSamp, "BV", "trja", accDim)
> gebvChnTRJA <- chn
> gebvMnTRJA       <- as.data.frame(t(apply(chn, 2, quantileLike)))
> gebvMnTRJA$trait <- rep(trtNam, times = Nacc)
> gebvMnTRJA$admix <- rep(ifelse(accNamTRJA %in% accNamADM, "admixed", "pure"), each=d)
```

```
> gebvMnTRJAs      <- gebvMnTRJA[order(gebvMnTRJA[,6], gebvMnTRJA[,3]),]
```

I plot sorted accession means.

```
> pdfFlNam <- "gebvFAMtrja.pdf"
> showtext_auto()
> ggplot(data=gebvMnTRJAs, aes(x=1:nrow(gebvMnTRJAs),y=mode,color=admix)) +
+     geom_segment(aes(x=1:nrow(gebvMnTRJAs), y=lower95, xend=1:nrow(gebvMnTRJAs), yend=upper9
+         color="grey80", size=0.75) +
+     geom_segment(aes(x=1:nrow(gebvMnTRJAs), y=lower50, xend=1:nrow(gebvMnTRJAs), yend=upper5
+         color="grey50", size=1) +
+     geom_point() +
+     facet_wrap(~trait, scales="free", nrow=2) +
+     theme_classic(base_size=18, base_family="myriad") +
+     theme(axis.title.x=element_blank(), axis.text.x=element_blank(),
+         strip.background=element_rect(fill="grey95", linetype="blank"),
+         axis.ticks.x=element_blank()) +
+     labs(y=trtNam[1])
> ggsave(pdfFlNam, width=8, height=10, units="in", device="pdf", useDingbats=F)
> cat("\\includegraphics{", pdfFlNam, "}\n\n", sep="")
```

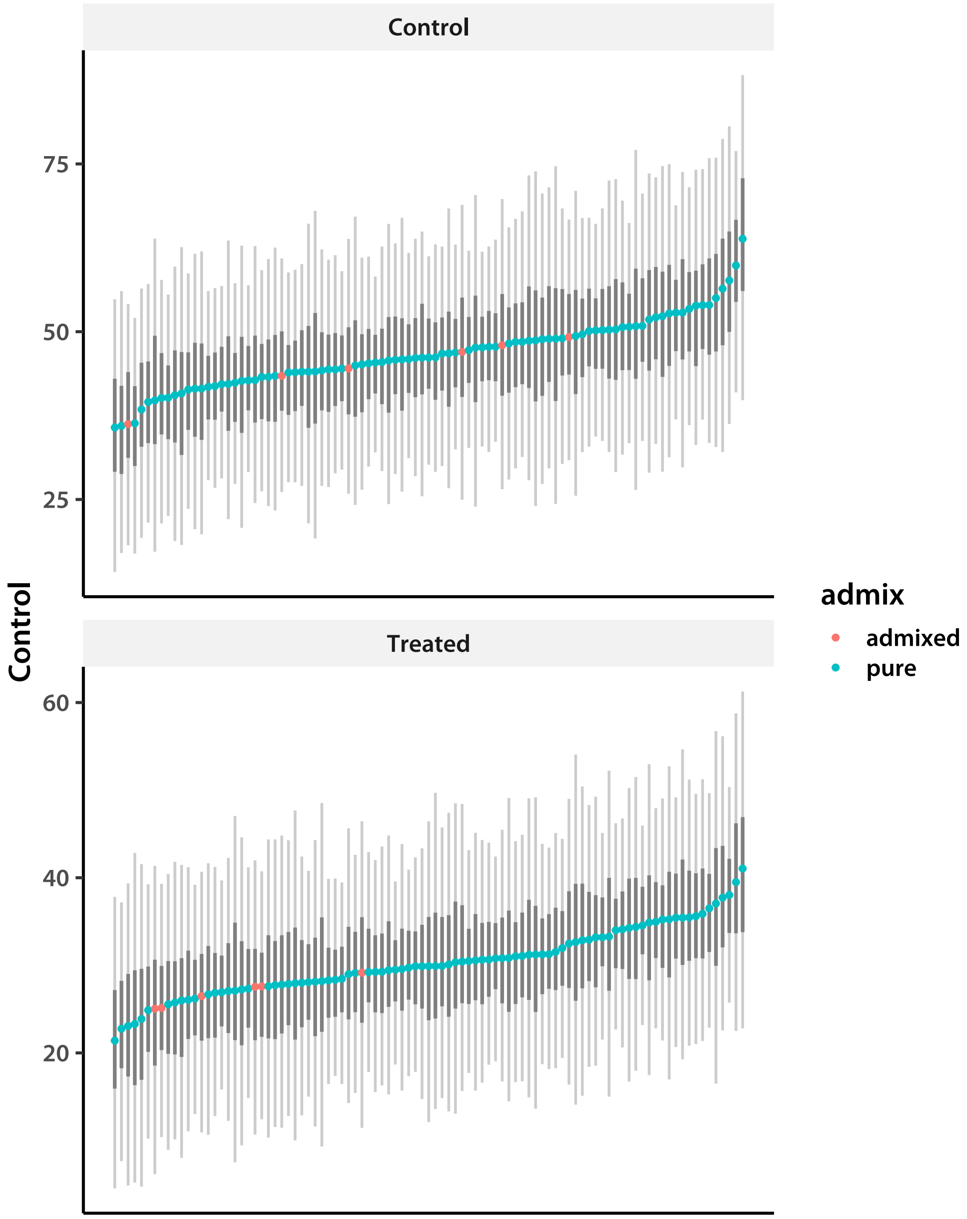


The effect is similar to the accession means, but seems more pronounced. Plot the ranks as before.

```
> pdfFlNam        <- "rankAdmGEBV.pdf"
> rankChn         <- t(apply(chn, 1, mnRank, admInd))
> rankBVTRJ       <- as.data.frame(t(apply(rankChn, 2, quantileLike)))
> rankBVTRJ$trait <- trtNam
> showtext_auto()
> ggplot(data=rankBVTRJ, aes(x=1:d,y=mode)) +
+     geom_segment(aes(x=1:d, y=lower95, xend=1:d, yend=upper95),
+         color="grey80", size=1.25) +
```

```
+     geom_segment(aes(x=1:d, y=lower50, xend=1:d, yend=upper50),
+         color="grey50", size=2) +
+     geom_point() +
+     geom_hline(yintercept=(mean(1:Nacc)), color="grey50") +
+     theme_classic(base_size=18, base_family="myriad") +
+     theme(axis.title.x=element_blank(), axis.text.x=element_blank(),
+         strip.background=element_rect(fill="grey95", linetype="blank"),
+         axis.ticks.x=element_blank()) +
+     labs(y="rank") + xlim(0.5, d+0.5) +
+     annotate("text", label="Control", x=1, y=min(rankMnTRJ$lower95)-2) +
+     annotate("text", label="Al treated", x=2, y=min(rankMnTRJ$lower95)-2)
> ggsave(pdfFlNam, width=8, height=8, units="in", device="pdf", useDingbats=F)
> cat("\\includegraphics{", pdfFlNam, "}\n\n", sep="")
```

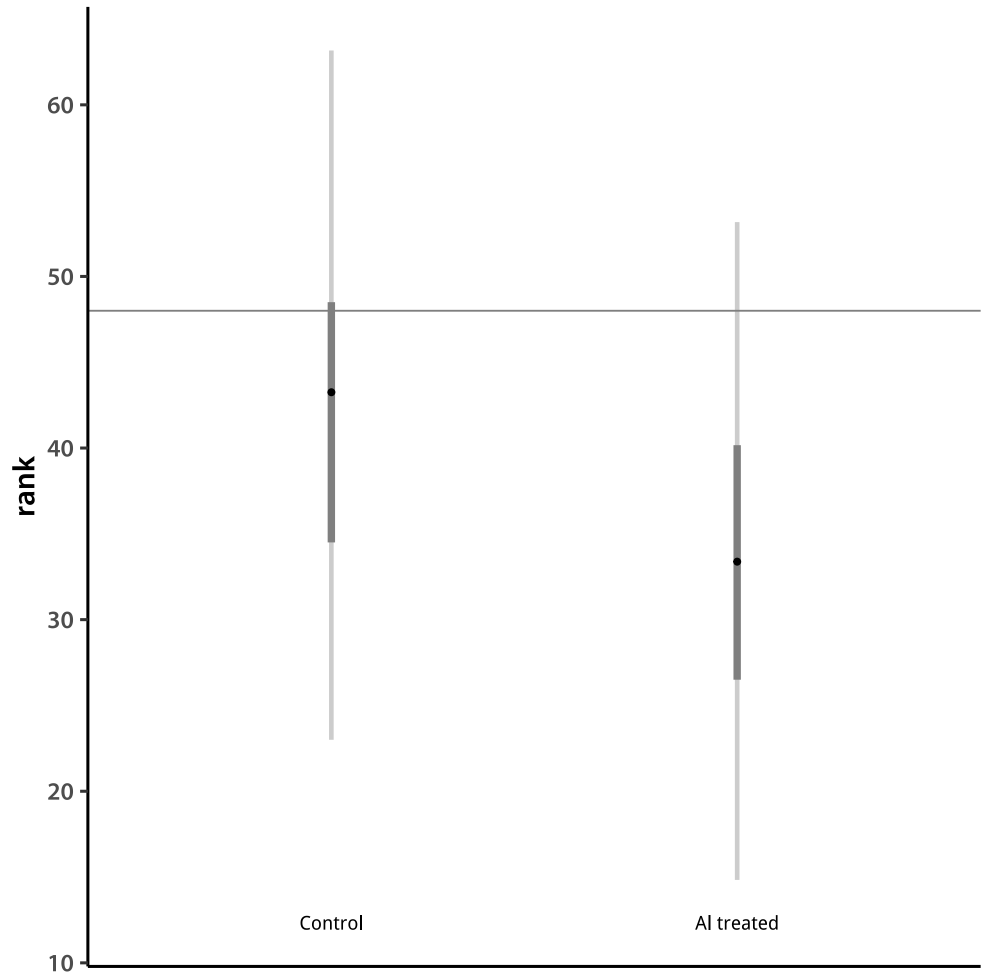


Numerical values:

```
> rankBVTRJ
```

```
   lower95 lower50     mode  upper50  upper95   trait
1 23.00000    34.5 43.25091 48.50000 63.16667 Control
2 14.83333    26.5 33.37917 40.16667 53.16667 Treated
```

### 5 Heritabilities

Finally, I directly compare heritabilities between control and treated, as well as between populations. The background genetic and residual variances are taken from the respective covariance matrices. However, because we used a Student-$t$ genetic model, the additive variance is estimated from the GEBV values rather $\mathbf{\Sigma}_a$.

I first define the functions I need.

```
> makeVar <- cmpfun(function(vec){
+     apply(matrix(vec, ncol=d, byrow=T), 2, var)
+ })
> get.hsq <- cmpfun(function(vec){
+     vec[1:d]/rowSums(matrix(vec, nrow = d))
+ })
```

The function `get.hsq` calculates the marker heritability, which is a kind of narrow-sense heritability. It is

$$h^2 = \frac{\sigma^2_{\text{GEBV}}}{\sigma^2_{\text{GEBV}} + \sigma^2_s + \sigma^2_e}$$

for each treatment.

I read in the covariance matrices and extract variances (diagonal elements).

```
> diagInd <- diag(matrix(1:(d^2), ncol = d, byrow = T))
> chn1 <- matrix(.C("GSLmatLoad",
+     "chains/SgEout_aus_3_1.gbin",
+     as.integer(chnLen), as.integer(d^2), out = double(chnLen*d^2))$out,
+         nrow = chnLen, byrow = T)[,diagInd]
> chn1 <- cbind(chn1,
+     matrix(.C("GSLmatLoad",
+     "chains/SgSout_aus_3_1.gbin",
+     as.integer(chnLen), as.integer(d^2), out = double(chnLen*d^2))$out,
+         nrow = chnLen, byrow = T)[,diagInd])
> chn2 <- matrix(.C("GSLmatLoad",
+     "chains/SgEout_aus_3_2.gbin",
+     as.integer(chnLen), as.integer(d^2), out = double(chnLen*d^2))$out,
+         nrow = chnLen, byrow = T)[,diagInd]
> chn2 <- cbind(chn2,
+     matrix(.C("GSLmatLoad",
+     "chains/SgSout_aus_3_2.gbin",
+     as.integer(chnLen), as.integer(d^2), out = double(chnLen*d^2))$out,
```

```
+         nrow = chnLen, byrow = T)[,diagInd])
> chn3 <- matrix(.C("GSLmatLoad",
+     "chains/SgEout_aus_3_3.gbin",
+     as.integer(chnLen), as.integer(d^2), out = double(chnLen*d^2))$out,
+         nrow = chnLen, byrow = T)[,diagInd]
> chn3 <- cbind(chn3,
+     matrix(.C("GSLmatLoad",
+     "chains/SgSout_aus_3_3.gbin",
+     as.integer(chnLen), as.integer(d^2), out = double(chnLen*d^2))$out,
+         nrow = chnLen, byrow = T)[,diagInd])
> chn4 <- matrix(.C("GSLmatLoad",
+     "chains/SgEout_aus_3_4.gbin",
+     as.integer(chnLen), as.integer(d^2), out = double(chnLen*d^2))$out,
+         nrow = chnLen, byrow = T)[,diagInd]
> chn4 <- cbind(chn4,
+     matrix(.C("GSLmatLoad",
+     "chains/SgSout_aus_3_4.gbin",
+     as.integer(chnLen), as.integer(d^2), out = double(chnLen*d^2))$out,
+         nrow = chnLen, byrow = T)[,diagInd])
> chn5 <- matrix(.C("GSLmatLoad",
+     "chains/SgEout_aus_3_5.gbin",
+     as.integer(chnLen), as.integer(d^2), out = double(chnLen*d^2))$out,
+         nrow = chnLen, byrow = T)[,diagInd]
> chn5 <- cbind(chn5,
+     matrix(.C("GSLmatLoad",
+     "chains/SgSout_aus_3_5.gbin",
+     as.integer(chnLen), as.integer(d^2), out = double(chnLen*d^2))$out,
+         nrow = chnLen, byrow = T)[,diagInd])
> sigChnAUS <- rbind(chn1, chn2, chn3, chn4, chn5)
> sigChnAUS <- cbind(t(apply(gebvChnAUS, 1, makeVar)), sigChnAUS)
> chnhsqAUS <- t(apply(sigChnAUS, 1, get.hsq))
> chn1 <- matrix(.C("GSLmatLoad",
+     "chains/SgEout_trj_3_1.gbin",
+     as.integer(chnLen), as.integer(d^2), out = double(chnLen*d^2))$out,
+         nrow = chnLen, byrow = T)[,diagInd]
> chn1 <- cbind(chn1,
+     matrix(.C("GSLmatLoad",
+     "chains/SgSout_trj_3_1.gbin",
+     as.integer(chnLen), as.integer(d^2), out = double(chnLen*d^2))$out,
+         nrow = chnLen, byrow = T)[,diagInd])
> chn2 <- matrix(.C("GSLmatLoad",
+     "chains/SgEout_trj_3_2.gbin",
+     as.integer(chnLen), as.integer(d^2), out = double(chnLen*d^2))$out,
+         nrow = chnLen, byrow = T)[,diagInd]
> chn2 <- cbind(chn2,
```

```
+     matrix(.C("GSLmatLoad",
+     "chains/SgSout_trj_3_2.gbin",
+     as.integer(chnLen), as.integer(d^2), out = double(chnLen*d^2))$out,
+         nrow = chnLen, byrow = T)[,diagInd])
> chn3 <- matrix(.C("GSLmatLoad",
+     "chains/SgEout_trj_3_3.gbin",
+     as.integer(chnLen), as.integer(d^2), out = double(chnLen*d^2))$out,
+         nrow = chnLen, byrow = T)[,diagInd]
> chn3 <- cbind(chn3,
+     matrix(.C("GSLmatLoad",
+     "chains/SgSout_trj_3_3.gbin",
+     as.integer(chnLen), as.integer(d^2), out = double(chnLen*d^2))$out,
+         nrow = chnLen, byrow = T)[,diagInd])
> chn4 <- matrix(.C("GSLmatLoad",
+     "chains/SgEout_trj_3_4.gbin",
+     as.integer(chnLen), as.integer(d^2), out = double(chnLen*d^2))$out,
+         nrow = chnLen, byrow = T)[,diagInd]
> chn4 <- cbind(chn4,
+     matrix(.C("GSLmatLoad",
+     "chains/SgSout_trj_3_4.gbin",
+     as.integer(chnLen), as.integer(d^2), out = double(chnLen*d^2))$out,
+         nrow = chnLen, byrow = T)[,diagInd])
> chn5 <- matrix(.C("GSLmatLoad",
+     "chains/SgEout_trj_3_5.gbin",
+     as.integer(chnLen), as.integer(d^2), out = double(chnLen*d^2))$out,
+         nrow = chnLen, byrow = T)[,diagInd]
> chn5 <- cbind(chn5,
+     matrix(.C("GSLmatLoad",
+     "chains/SgSout_trj_3_5.gbin",
+     as.integer(chnLen), as.integer(d^2), out = double(chnLen*d^2))$out,
+         nrow = chnLen, byrow = T)[,diagInd])
> sigChnTRJ <- rbind(chn1, chn2, chn3, chn4, chn5)
> sigChnTRJ <- cbind(t(apply(gebvChnTRJ, 1, makeVar)), sigChnTRJ)
> chnhsqTRJ <- t(apply(sigChnTRJ, 1, get.hsq))
> chn1 <- matrix(.C("GSLmatLoad",
+     "chains/SgEout_trja_3_1.gbin",
+     as.integer(chnLen), as.integer(d^2), out = double(chnLen*d^2))$out,
+         nrow = chnLen, byrow = T)[,diagInd]
> chn1 <- cbind(chn1,
+     matrix(.C("GSLmatLoad",
+     "chains/SgSout_trja_3_1.gbin",
+     as.integer(chnLen), as.integer(d^2), out = double(chnLen*d^2))$out,
+         nrow = chnLen, byrow = T)[,diagInd])
> chn2 <- matrix(.C("GSLmatLoad",
+     "chains/SgEout_trja_3_2.gbin",
```

```
+     as.integer(chnLen), as.integer(d^2), out = double(chnLen*d^2))$out,
+         nrow = chnLen, byrow = T)[,diagInd]
> chn2 <- cbind(chn2,
+     matrix(.C("GSLmatLoad",
+     "chains/SgSout_trja_3_2.gbin",
+     as.integer(chnLen), as.integer(d^2), out = double(chnLen*d^2))$out,
+         nrow = chnLen, byrow = T)[,diagInd])
> chn3 <- matrix(.C("GSLmatLoad",
+     "chains/SgEout_trja_3_3.gbin",
+     as.integer(chnLen), as.integer(d^2), out = double(chnLen*d^2))$out,
+         nrow = chnLen, byrow = T)[,diagInd]
> chn3 <- cbind(chn3,
+     matrix(.C("GSLmatLoad",
+     "chains/SgSout_trja_3_3.gbin",
+     as.integer(chnLen), as.integer(d^2), out = double(chnLen*d^2))$out,
+         nrow = chnLen, byrow = T)[,diagInd])
> chn4 <- matrix(.C("GSLmatLoad",
+     "chains/SgEout_trja_3_4.gbin",
+     as.integer(chnLen), as.integer(d^2), out = double(chnLen*d^2))$out,
+         nrow = chnLen, byrow = T)[,diagInd]
> chn4 <- cbind(chn4,
+     matrix(.C("GSLmatLoad",
+     "chains/SgSout_trja_3_4.gbin",
+     as.integer(chnLen), as.integer(d^2), out = double(chnLen*d^2))$out,
+         nrow = chnLen, byrow = T)[,diagInd])
> chn5 <- matrix(.C("GSLmatLoad",
+     "chains/SgEout_trja_3_5.gbin",
+     as.integer(chnLen), as.integer(d^2), out = double(chnLen*d^2))$out,
+         nrow = chnLen, byrow = T)[,diagInd]
> chn5 <- cbind(chn5,
+     matrix(.C("GSLmatLoad",
+     "chains/SgSout_trja_3_5.gbin",
+     as.integer(chnLen), as.integer(d^2), out = double(chnLen*d^2))$out,
+         nrow = chnLen, byrow = T)[,diagInd])
> sigChnTRJA <- rbind(chn1, chn2, chn3, chn4, chn5)
> sigChnTRJA <- cbind(t(apply(gebvChnTRJA, 1, makeVar)), sigChnTRJA)
> chnhsqTRJA <- t(apply(sigChnTRJA, 1, get.hsq))
> Npop   <- 3
> hsqHPD <- as.data.frame(rbind(t(apply(chnhsqAUS, 2, quantileLike)),
+     t(apply(chnhsqTRJ, 2, quantileLike)),
+     t(apply(chnhsqTRJA, 2, quantileLike))))
> hsqHPD$treatment  <- rep(trtNam, Npop)
> hsqHPD$population <- rep(c("AUS", "TRJ", "TRJA"), each=d)
>
```

Plot the results.

```
> pdfFlNam <- "hSqFAMall.pdf"
> showtext_auto()
> ggplot(data=hsqHPD, aes(x=c(1,2,4,5,7,8), y=mode, color=population)) +
+     geom_segment(aes(x=c(1,2,4,5,7,8), y=lower95, xend=c(1,2,4,5,7,8),
+         yend=upper95, color=population),
+         size=1.2) +
+     geom_segment(aes(x=c(1,2,4,5,7,8), y=lower50, xend=c(1,2,4,5,7,8),
+         yend=upper50, color=population),
+         size=1.75) +
+     geom_point(size=2) +
+     theme_classic(base_size=18, base_family="myriad") +
+     scale_x_continuous(breaks=c(1,2,4,5,7,8), labels=rep(trtNam, Npop)) +
+     ylim(c(0,1)) +
+     labs(y=expression(italic(h)^2), x="treatment")
> ggsave(pdfFlNam, width=8, height=8, units="in", device="pdf", useDingbats=F)
> cat("\\includegraphics{", pdfFlNam, "}\n\n", sep="")
```

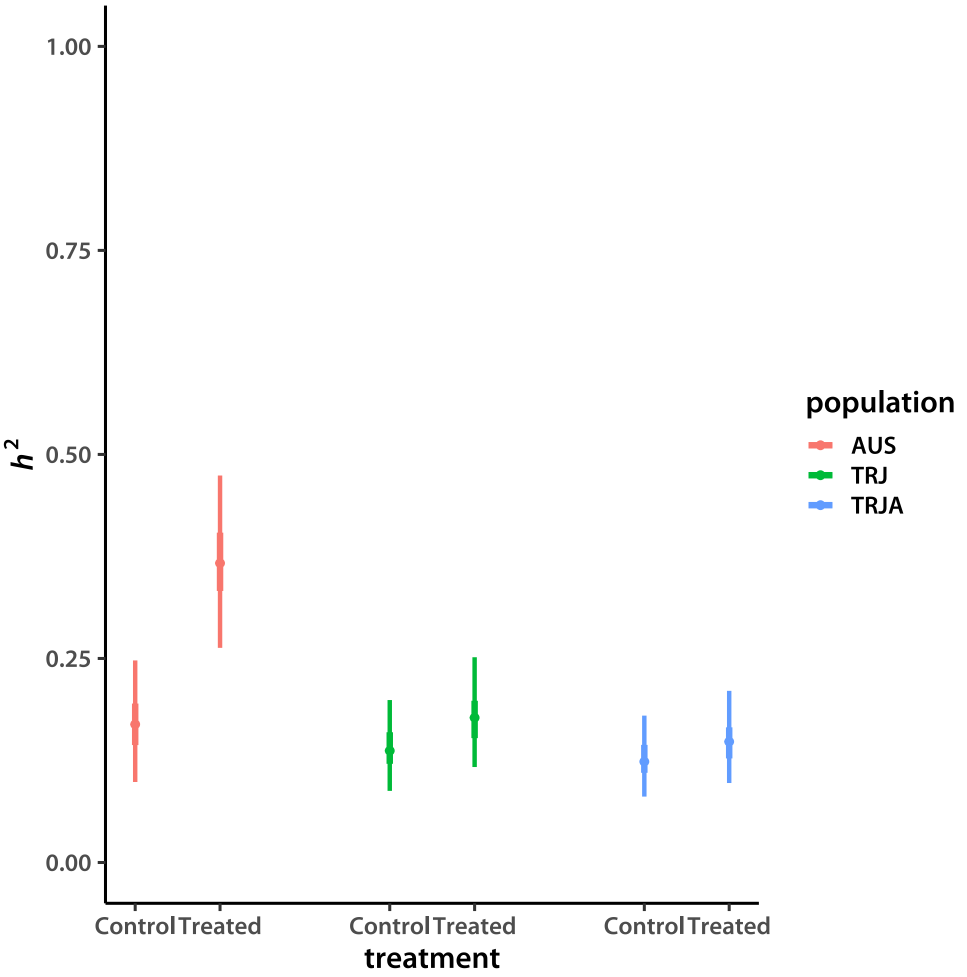


Repeat, including only the pure accessions.

```
> pdfFlNam <- "hSqFAMpure.pdf"
> showtext_auto()
> ggplot(data=subset(hsqHPD, population != "TRJA"), aes(x=c(1,2,4,5), y=mode, color=population
+     geom_segment(aes(x=c(1,2,4,5), y=lower95, xend=c(1,2,4,5),
+         yend=upper95, color=population),
+         size=1.2) +
+     geom_segment(aes(x=c(1,2,4,5), y=lower50, xend=c(1,2,4,5),
+         yend=upper50, color=population),
+         size=1.75) +
+     geom_point(size=2) +
+     theme_classic(base_size=18, base_family="myriad") +
+     scale_x_continuous(breaks=c(1,2,4,5), labels=rep(trtNam, Npop-1)) +
+     ylim(c(0,1)) +
+     labs(y=expression(italic(h)^2), x="treatment")
> ggsave(pdfFlNam, width=8, height=8, units="in", device="pdf", useDingbats=F)
```

```
> cat("\\includegraphics{", pdfFlNam, "}\n\n", sep="")
```

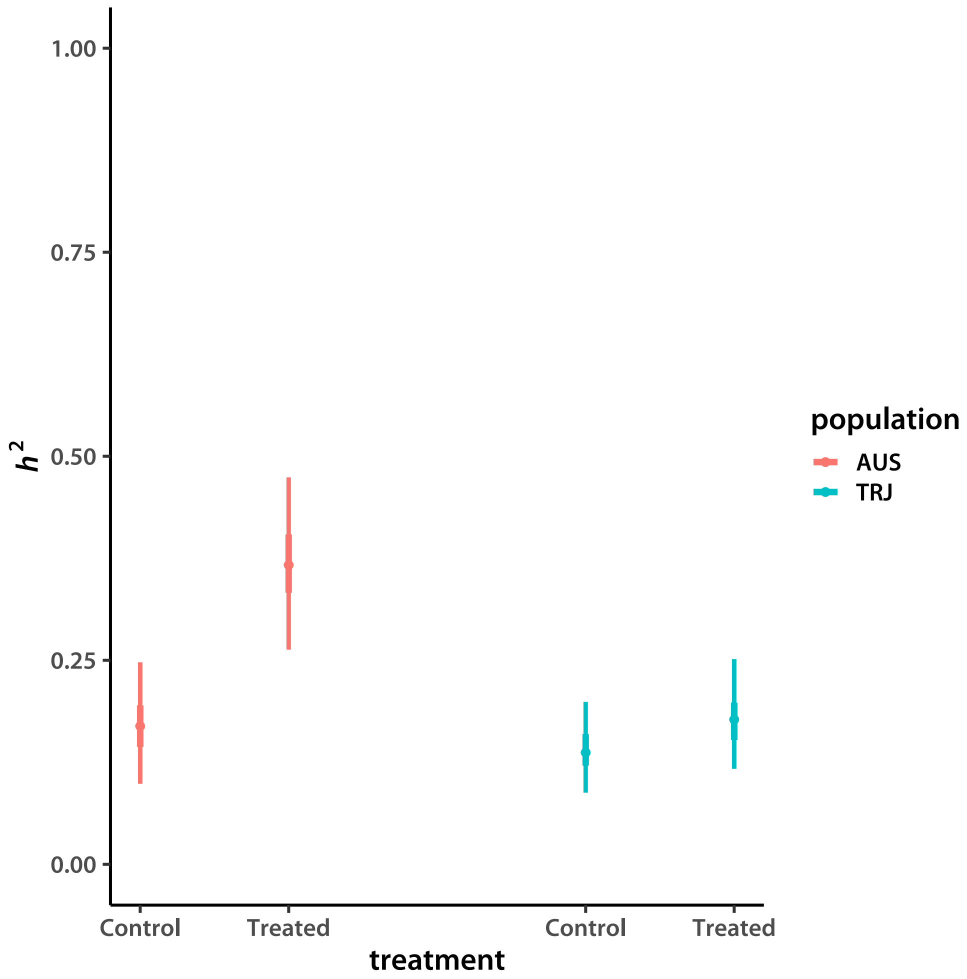


Now I plot broad-sense (among-accession) heritability:

$$H^2 = \frac{\sigma^2_{\text{GEBV}} + \sigma^2_s}{\sigma^2_{\text{GEBV}} + \sigma^2_s + \sigma^2_e}$$

I am also interested in the contribution of the non-additive effects alone, excluding the marker influence:

$$FVE_s = \frac{\sigma^2_s}{\sigma^2_{\text{GEBV}} + \sigma^2_s + \sigma^2_e}$$

```
> get.Hsq <- cmpfun(function(vec){
+     (vec[1:d] + vec[(2*d+1):(3*d)])/rowSums(matrix(vec, nrow = d))
+ })
> get.fve <- cmpfun(function(vec){
+     (vec[(2*d+1):(3*d)])/rowSums(matrix(vec, nrow = d))
```

```
+ })
> chnHsqAUS  <- t(apply(sigChnAUS,  1, get.Hsq))
> chnHsqTRJ  <- t(apply(sigChnTRJ,  1, get.Hsq))
> chnHsqTRJA <- t(apply(sigChnTRJA, 1, get.Hsq))
> chnFveAUS  <- t(apply(sigChnAUS,  1, get.fve))
> chnFveTRJ  <- t(apply(sigChnTRJ,  1, get.fve))
> chnFveTRJA <- t(apply(sigChnTRJA, 1, get.fve))
> HsqHPD <- as.data.frame(rbind(t(apply(chnHsqAUS, 2, quantileLike)),
+     t(apply(chnHsqTRJ, 2, quantileLike)),
+     t(apply(chnHsqTRJA, 2, quantileLike))))
> HsqHPD$treatment <- rep(trtNam, Npop)
> HsqHPD$population <- rep(c("AUS", "TRJ", "TRJA"), each=d)
> fveHPD <- as.data.frame(rbind(t(apply(chnhsqAUS, 2, quantileLike)),
+     t(apply(chnhsqTRJ, 2, quantileLike)),
+     t(apply(chnFveAUS, 2, quantileLike)),
+     t(apply(chnFveTRJ, 2, quantileLike))))
> fveHPD$treatment  <- rep(trtNam, 2*(Npop-1))
> fveHPD$population <- rep(rep(c("AUS", "TRJ"), each=2), d)
> fveHPD$kind       <- factor(rep(c("Marker", "Background"), each=d*(Npop-1)),
+                              levels=c("Marker", "Background"))
```

Plot the results.

```
> pdfFlNam <- "HsqFAMall.pdf"
> showtext_auto()
> ggplot(data=HsqHPD, aes(x=c(1,2,4,5,7,8), y=mode, color=population)) +
+     geom_segment(aes(x=c(1,2,4,5,7,8), y=lower95, xend=c(1,2,4,5,7,8),
+         yend=upper95, color=population),
+         size=1.2) +
+     geom_segment(aes(x=c(1,2,4,5,7,8), y=lower50, xend=c(1,2,4,5,7,8),
+         yend=upper50, color=population),
+         size=1.75) +
+     geom_point(size=2) +
+     theme_classic(base_size=18, base_family="myriad") +
+     scale_x_continuous(breaks=c(1,2,4,5,7,8), labels=rep(trtNam, Npop)) +
+     ylim(c(0,1)) +
+     labs(y=expression(italic(H)^2), x="treatment")
> ggsave(pdfFlNam, width=8, height=8, units="in", device="pdf", useDingbats=F)
> cat("\\includegraphics{", pdfFlNam, "}\n\n", sep="")
```

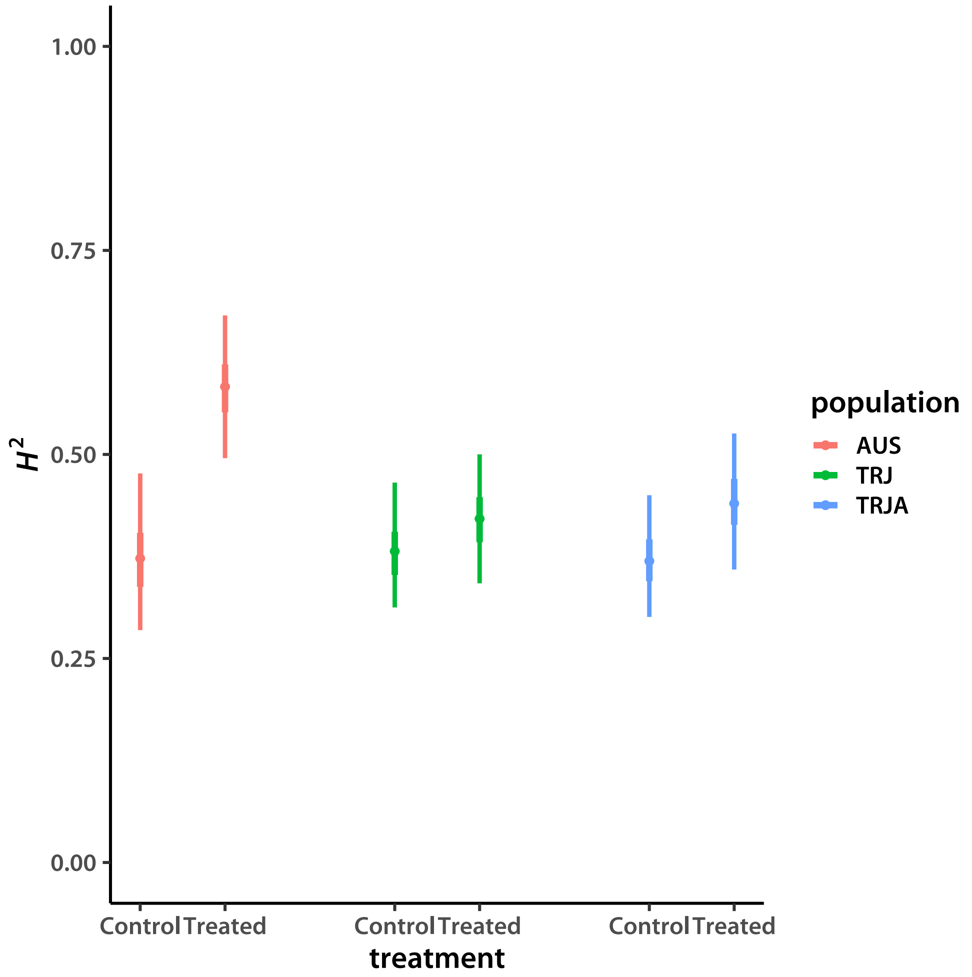


Repeat, including only the pure accessions.

```
> pdfFlNam <- "HsqFAMpure.pdf"
> showtext_auto()
> ggplot(data=subset(HsqHPD, population != "TRJA"), aes(x=c(1,2,4,5), y=mode, color=population)
+     geom_segment(aes(x=c(1,2,4,5), y=lower95, xend=c(1,2,4,5),
+         yend=upper95, color=population),
+         size=1.2) +
+     geom_segment(aes(x=c(1,2,4,5), y=lower50, xend=c(1,2,4,5),
+         yend=upper50, color=population),
+         size=1.75) +
+     geom_point(size=2) +
+     theme_classic(base_size=18, base_family="myriad") +
+     scale_x_continuous(breaks=c(1,2,4,5), labels=rep(trtNam, Npop-1)) +
+     ylim(c(0,1)) +
+     labs(y=expression(italic(H)^2), x="treatment")
> ggsave(pdfFlNam, width=8, height=8, units="in", device="pdf", useDingbats=F)
```

```
> cat("\\includegraphics{", pdfFlNam, "}\n\n", sep="")
```

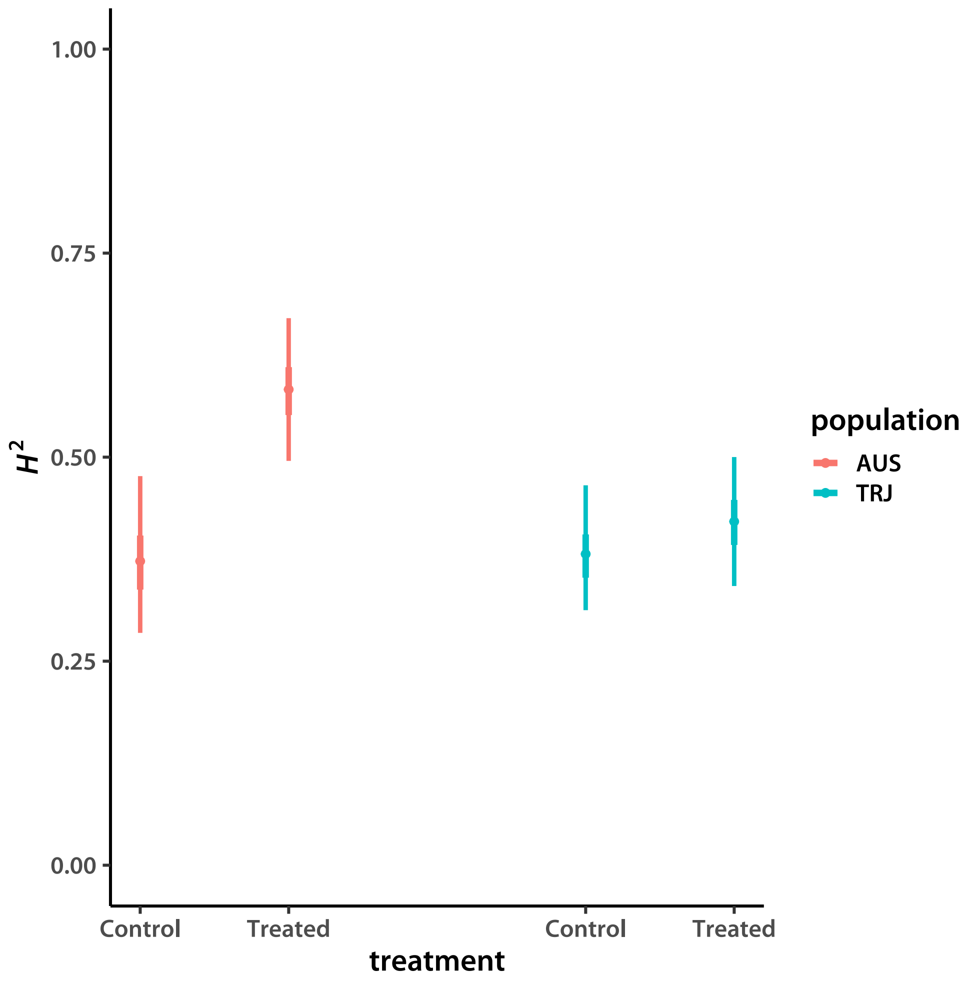


I plot the fractions of variance explained side by side.

```
> pdfFlNam <- "fveFAMpure.pdf"
> showtext_auto()
> ggplot(data=fveHPD, aes(x=rep(1:4,2), y=mode, color=population)) +
+     geom_segment(aes(x=rep(1:4,2), y=lower95, xend=rep(1:4,2),
+         yend=upper95, color=population),
+         size=1.2) +
+     geom_segment(aes(x=rep(1:4,2), y=lower50, xend=rep(1:4,2),
+         yend=upper50, color=population),
+         size=1.75) +
+     geom_point(size=2) +
+     facet_wrap(~kind, scales="free_x", ncol=2) +
+     theme_classic(base_size=18, base_family="myriad") +
+     scale_x_continuous(breaks=rep(1:4,2), labels=rep(trtNam, d*(Npop-1))) +
```

```
+     theme(axis.text.x=element_text(angle=45, hjust=1.0, vjust=1.0),
+         strip.background=element_rect(fill="grey95", linetype="blank"),
+         panel.spacing.x=unit(4.0, "lines")) +
+     ylim(c(0,1)) +
+     labs(y="fraction of variance", x="treatment")
> ggsave(pdfFlNam, width=10, height=8, units="in", device="pdf", useDingbats=F)
> cat("\\includegraphics{", pdfFlNam, "}\n\n", sep="")
```

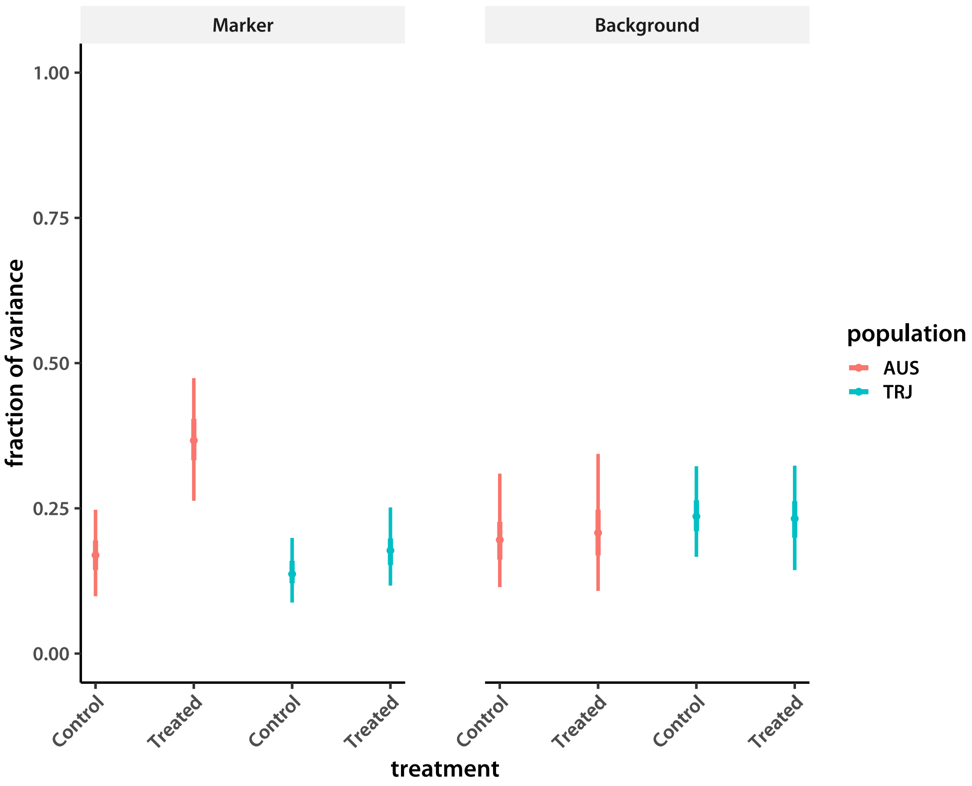


Finally, put the plots together for direct comparison.

```
> ggGEBV <- ggplot(data=hsqHPD, aes(x=c(1,2,4,5,7,8), y=mode, color=population)) +
+     geom_segment(aes(x=c(1,2,4,5,7,8), y=lower95, xend=c(1,2,4,5,7,8),
+         yend=upper95, color=population),
+         size=1.2) +
+     geom_segment(aes(x=c(1,2,4,5,7,8), y=lower50, xend=c(1,2,4,5,7,8),
+         yend=upper50, color=population),
+         size=1.75) +
+     geom_point(size=2) +
+     theme_classic(base_size=18, base_family="myriad") +
+     scale_x_continuous(breaks=c(1,2,4,5,7,8), labels=rep(trtNam, Npop)) +
+     theme(axis.text.x=element_text(angle=45, hjust=1.0, vjust=1.0)) +
+     ylim(c(0,1)) +
+     labs(y=expression(italic(h)^2), x="treatment")
```

```
> ggHsq <- ggplot(data=HsqHPD, aes(x=c(1,2,4,5,7,8), y=mode, color=population)) +
+     geom_segment(aes(x=c(1,2,4,5,7,8), y=lower95, xend=c(1,2,4,5,7,8),
+         yend=upper95, color=population),
+         size=1.2) +
+     geom_segment(aes(x=c(1,2,4,5,7,8), y=lower50, xend=c(1,2,4,5,7,8),
+         yend=upper50, color=population),
+         size=1.75) +
+     geom_point(size=2) +
+     theme_classic(base_size=18, base_family="myriad") +
+     scale_x_continuous(breaks=c(1,2,4,5,7,8), labels=rep(trtNam, Npop)) +
+     theme(axis.text.x=element_text(angle=45, hjust=1.0, vjust=1.0)) +
+     ylim(c(0,1)) +
+     labs(y=expression(italic(H)^2), x="treatment")
> showtext_auto()
> grid.arrange(ggGEBV, ggHsq, ncol = 2)
>
```

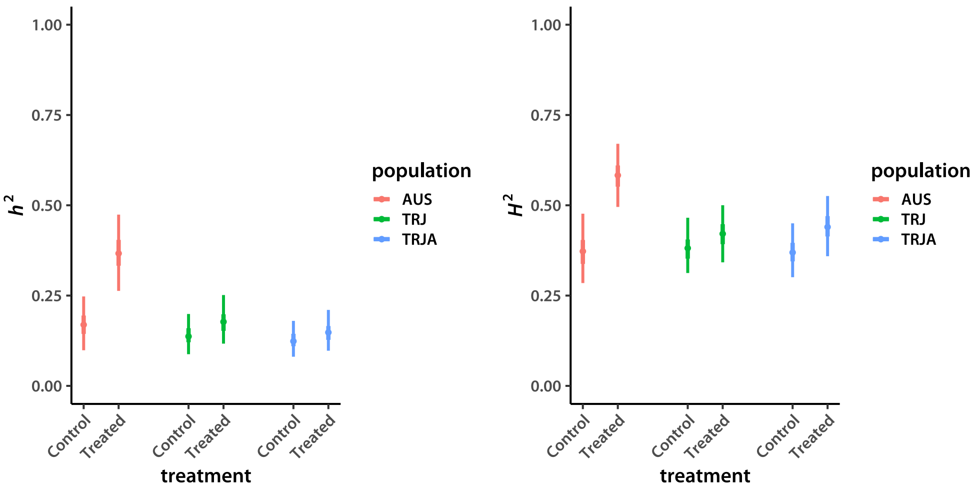
