## Supplemental methods and scripts for "Low additive genetic variation in a trait under selection in domesticated rice": mcmcResultsNDtrjIBSS.pdf

### Parameter estimates from model Markov chains with the quadratic kernel

Tony Greenberg

April 30, 2019

```
R version 3.5.3 (2019-03-11)
Platform: x86_64-pc-linux-gnu (64-bit)
Running under: Arch Linux

Matrix products: default
BLAS/LAPACK: /opt/intel/compilers_and_libraries_2019.1.144/linux/mkl/lib/intel64_lin/libmkl_gf_lp64.so

locale:
 [1] LC_CTYPE=en_US.UTF-8       LC_NUMERIC=C               LC_TIME=en_US.UTF-8
 [4] LC_COLLATE=en_US.UTF-8     LC_MONETARY=en_US.UTF-8    LC_MESSAGES=en_US.UTF-8
 [7] LC_PAPER=en_US.UTF-8       LC_NAME=C                  LC_ADDRESS=C
[10] LC_TELEPHONE=C             LC_MEASUREMENT=en_US.UTF-8 LC_IDENTIFICATION=C

attached base packages:
[1] compiler  stats     graphics  grDevices utils     datasets  methods   base

other attached packages:
[1] showtext_0.6   showtextdb_2.0 sysfonts_0.8   gridExtra_2.3  ggplot2_3.1.0

loaded via a namespace (and not attached):
 [1] Rcpp_1.0.0       crayon_1.3.4     withr_2.1.2      grid_3.5.3       plyr_1.8.4
 [6] gtable_0.2.0     scales_1.0.0     pillar_1.3.1     rlang_0.3.1      lazyeval_0.2.1
[11] tools_3.5.3      munsell_0.5.0    pkgconfig_2.0.2  colorspace_1.4-0 tibble_2.0.1
```

In this document I calculate parameter estimates from the model Gibbs sampler Markov chains. Control and Al-treatment data sets were modeled separately, with a simple replication scheme without experiment-level covariates. I used the quadratic kernel ($K \circ K$) to model all-pairwise epistasis.

```
> trtNam <- c("Longest Root Length", "Total Root Length")
> d      <- 2
> Nacc   <- 119
> locDim <- d*Nacc
> nChn   <- 5
> chnLen <- 2000
> chn <- NULL
> addSamp <- cmpfun(function(i, vrNam, exNam, ncl){
+     inFlNam <- paste("chains/", vrNam, "_", exNam, "_IBSS_3_1000_", i, ".gbin", sep = "")
+     chn <<- rbind(chn,
+                 matrix(.C("GSLmatLoad",
+                     inFlNam, as.integer(chnLen), as.integer(ncl),
+                     out = double(chnLen*ncl))$out,
+                 nrow = chnLen, byrow = T)
+         )
+     return(NULL)
+ })
> trash <- sapply(1:nChn, addSamp, "LN", "ctrl", locDim)
> accMnCtrlCHN    <- chn
> accMnCtrl       <- as.data.frame(t(apply(chn, 2, quantileLike)))
> accMnCtrl$trait <- rep(trtNam, times = Nacc)
> accMnCtrlS      <- accMnCtrl[order(accMnCtrl[,6], accMnCtrl[,3]),]

> pdfFlNam <- "lineMeansCtrlNEWqk.pdf"
> showtext_auto()
> ggplot(data=accMnCtrlS, aes(x=1:nrow(accMnCtrlS),y=mode)) +
+     geom_segment(aes(x=1:nrow(accMnCtrlS), y=lower95,
+         xend=1:nrow(accMnCtrlS), yend=upper95),
+         color="grey80", size=0.75) +
+     geom_segment(aes(x=1:nrow(accMnCtrlS), y=lower50,
+         xend=1:nrow(accMnCtrlS), yend=upper50),
+         color="grey50", size=1) +
+     geom_point() +
+     facet_wrap(~trait, scales="free", nrow=2) +
+     theme_classic(base_size=18, base_family="myriad") +
```

```
+     theme(axis.title.x=element_blank(), axis.text.x=element_blank(),
+         axis.ticks.x=element_blank(),
+         strip.background=element_rect(fill="grey95", linetype="blank")) +
+     labs(y="length")
> ggsave(pdfFlNam, width=8, height=10, units="in", device="pdf", useDingbats=F)
> cat("\\includegraphics{", pdfFlNam, "}\n\n", sep="")
```

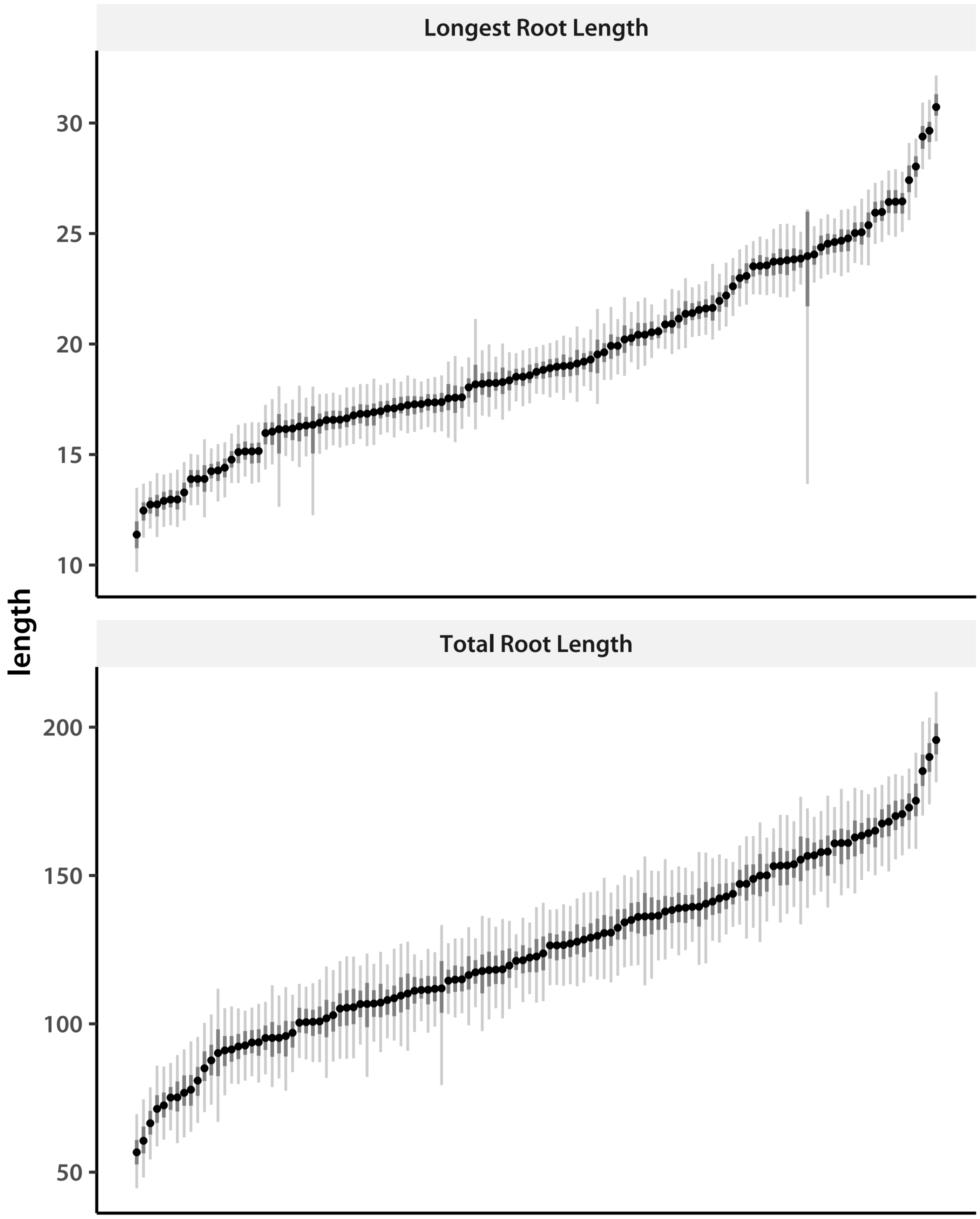


Process the GEBV estimates the same.

```
> chn             <- NULL
> trash           <- sapply(1:nChn, addSamp, "BV", "ctrl", locDim)
```

```
> chnGEBVctrl      <- chn
> accBVCtrl        <- as.data.frame(t(apply(chn, 2, quantileLike)))
> accBVCtrl$trait <- rep(trtNam, times = Nacc)
> accBVCtrlS       <- accBVCtrl[order(accBVCtrl[,6], accBVCtrl[,3]),]

> pdfFlNam <- "gebvCtrlNEWqk.pdf"
> showtext_auto()
> ggplot(data=accBVCtrlS, aes(x=1:nrow(accBVCtrlS),y=mode)) +
+     geom_segment(aes(x=1:nrow(accBVCtrlS), y=lower95,
+         xend=1:nrow(accBVCtrlS), yend=upper95),
+         color="grey80", size=0.75) +
+     geom_segment(aes(x=1:nrow(accBVCtrlS), y=lower50,
+         xend=1:nrow(accBVCtrlS), yend=upper50),
+         color="grey50", size=1) +
+     geom_point() +
+     facet_wrap(~trait, scales="free", nrow=2) +
+     theme_classic(base_size=18, base_family="myriad") +
+     theme(axis.title.x=element_blank(), axis.text.x=element_blank(),
+         axis.ticks.x=element_blank(),
+         strip.background=element_rect(fill="grey95", linetype="blank")) +
+     labs(y="length")
> ggsave(pdfFlNam, width=8, height=10, units="in", device="pdf", useDingbats=F)
> cat("\\includegraphics{", pdfFlNam, "}\n\n", sep="")
```

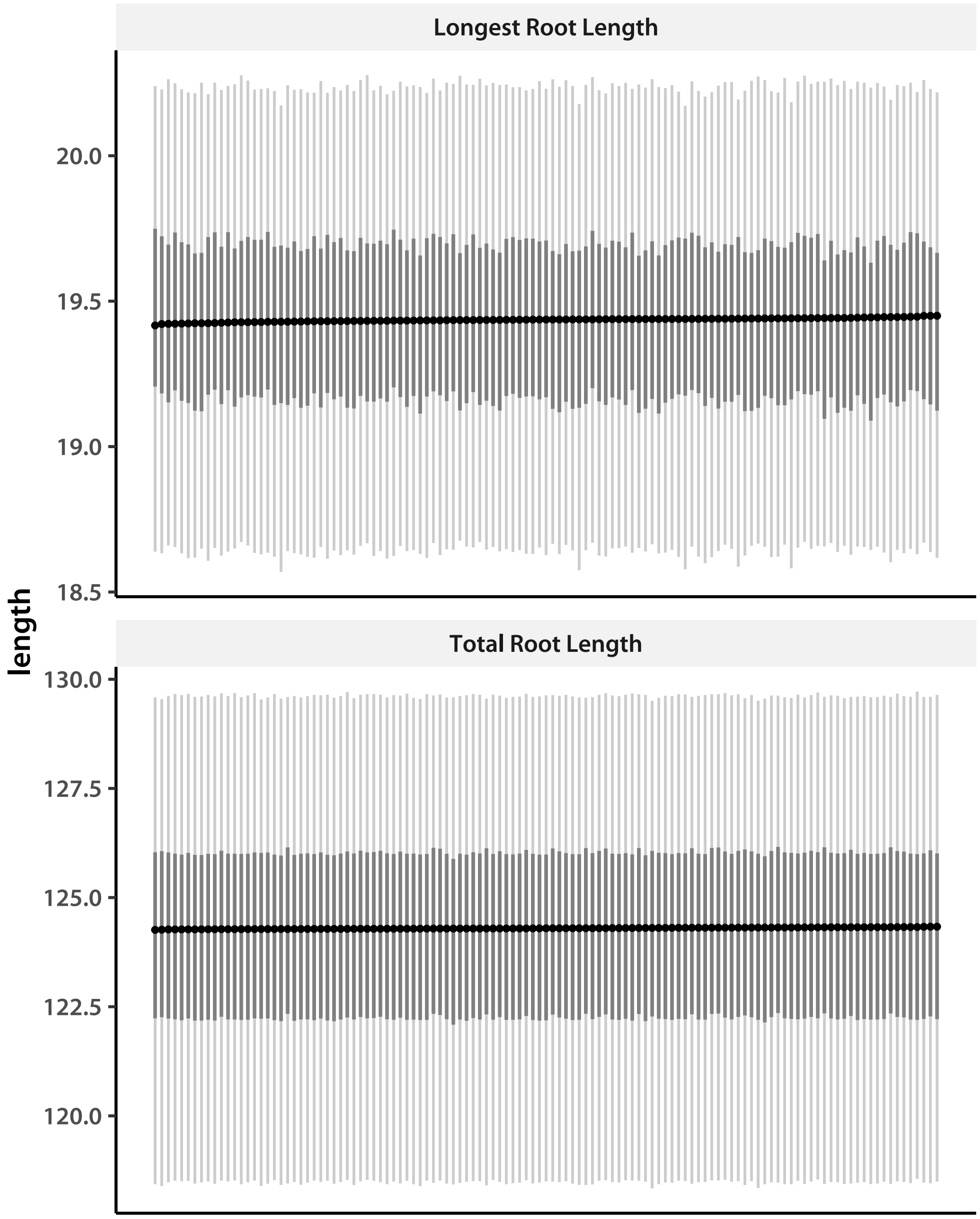


Clearly not much GEBV variation.

Now estimate the parameters from Al treatment data.

```
> chn   <- NULL
> trash <- sapply(1:nChn, addSamp, "LN", "expt", locDim)
> accMnExpt       <- as.data.frame(t(apply(chn, 2, quantileLike)))
> accMnExpt$trait <- rep(trtNam, times = Nacc)
> accMnExptS      <- accMnExpt[order(accMnExpt[,6], accMnExpt[,3]),]
```

```
> pdfFlNam <- "lineMeansExptNEWqk.pdf"
> showtext_auto()
```

```
> ggplot(data=accMnExptS, aes(x=1:nrow(accMnExptS),y=mode)) +
+     geom_segment(aes(x=1:nrow(accMnExptS), y=lower95, xend=1:nrow(accMnExptS), yend=upper95)
+         color="grey80", size=0.75) +
+     geom_segment(aes(x=1:nrow(accMnExptS), y=lower50, xend=1:nrow(accMnExptS), yend=upper50)
+         color="grey50", size=1) +
+     geom_point() +
+     facet_wrap(~trait, scales="free", nrow=2) +
+     theme_classic(base_size=18, base_family="myriad") +
+     theme(axis.title.x=element_blank(), axis.text.x=element_blank(),
+         axis.ticks.x=element_blank(),
+         strip.background=element_rect(fill="grey95", linetype="blank")) +
+     labs(y="length")
> ggsave(pdfFlNam, width=8, height=10, units="in", device="pdf", useDingbats=F)
> cat("\\includegraphics{", pdfFlNam, "}\n\n", sep="")
```

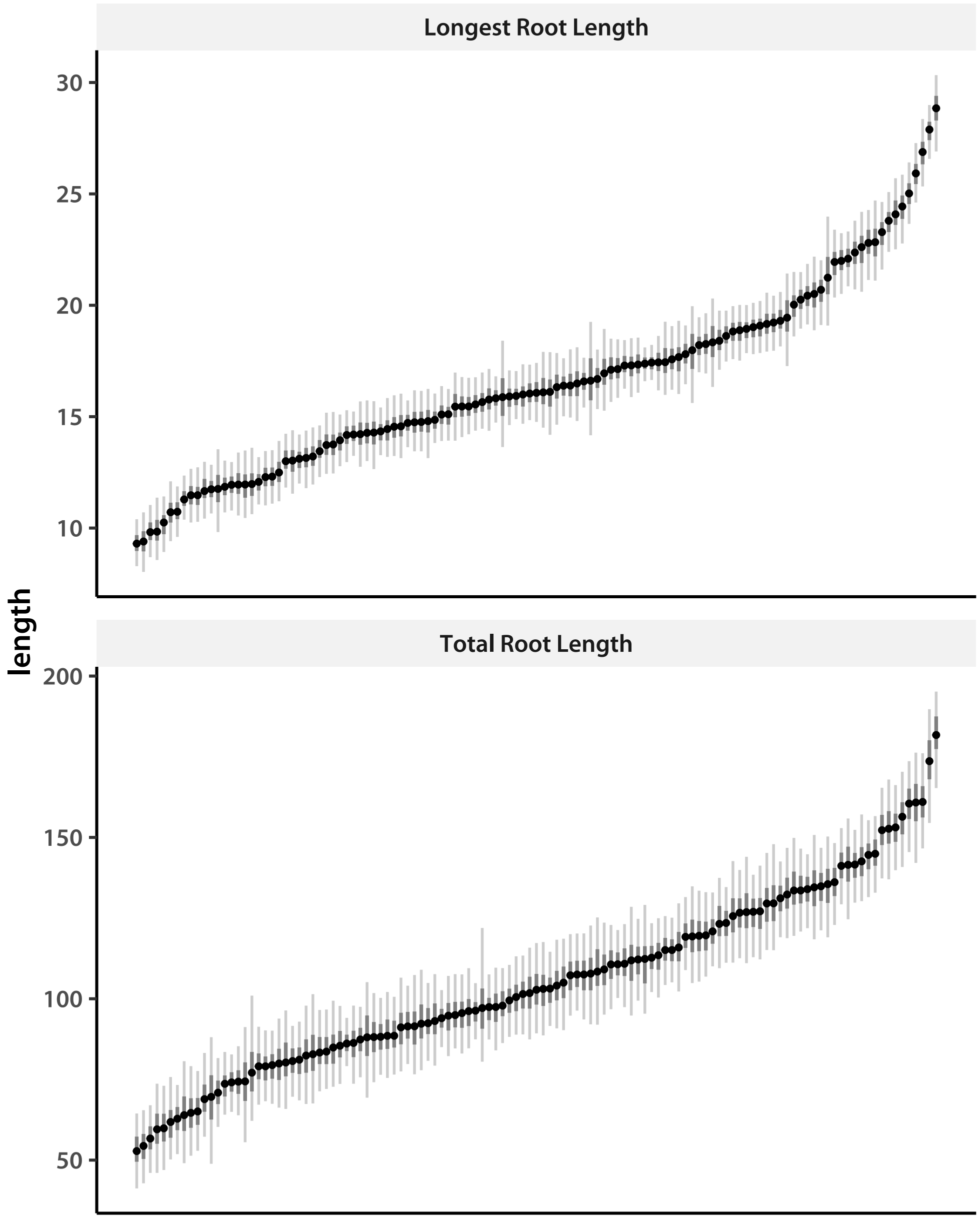


Now look at the genetic correlation of treatment *vs* control.

```
> gCor <- cmpfun(function(i){
+     res <- cor(cbind(matrix(accMnCtrlCHN[i,], ncol=d, byrow=T),
+                  matrix(chn[i,], ncol=d, byrow=T)))
+     return(res[col(res)>row(res)]) # upper triangle
+ })
> corCHN   <- sapply(1:nrow(chn), gCor)
> corHPD   <- apply(corCHN, 1, quantileLike)
> corNames <- paste(rep(trtNam, 2), rep(c("CTL", "EXP"), each=d), sep=":")
```

```
> corInd   <- col(matrix(1, 2*d, 2*d))>row(matrix(1, 2*d, 2*d))
> colnames(corHPD) <- paste(matrix(corNames, 2*d, 2*d)[corInd],
+                           matrix(corNames, 2*d, 2*d, byrow=T)[corInd], sep="|")
> round(t(corHPD), 4)
```

```
                                                 lower95 lower50   mode upper50 upper95
Longest Root Length:CTL|Total Root Length:CTL     0.5498  0.5777 0.5912  0.6062  0.6334
Longest Root Length:CTL|Longest Root Length:EXP   0.7760  0.8022 0.8159  0.8254  0.8429
Total Root Length:CTL|Longest Root Length:EXP     0.4513  0.4853 0.5039  0.5196  0.5519
Longest Root Length:CTL|Total Root Length:EXP     0.5973  0.6228 0.6368  0.6512  0.6797
Total Root Length:CTL|Total Root Length:EXP       0.7307  0.7596 0.7713  0.7866  0.8085
Longest Root Length:EXP|Total Root Length:EXP     0.6429  0.6702 0.6832  0.6956  0.7179
```

And, finally, GEBV.

```
> chn             <- NULL
> trash           <- sapply(1:nChn, addSamp, "BV", "expt", locDim)
> chnGEBVexpt     <- chn
> accBVExpt       <- as.data.frame(t(apply(chn, 2, quantileLike)))
> accBVExpt$trait <- rep(trtNam, times = Nacc)
> accBVExptS      <- accBVExpt[order(accBVExpt[,6], accBVExpt[,3]),]

> pdfFlNam <- "gebvExptNEWqk.pdf"
> showtext_auto()
> ggplot(data=accBVExptS, aes(x=1:nrow(accBVExptS),y=mode)) +
+     geom_segment(aes(x=1:nrow(accBVExptS), y=lower95,
+                      xend=1:nrow(accBVExptS), yend=upper95),
+         color="grey80", size=0.75) +
+     geom_segment(aes(x=1:nrow(accBVExptS), y=lower50,
+                      xend=1:nrow(accBVExptS), yend=upper50),
+         color="grey50", size=1) +
+     geom_point() +
+     facet_wrap(~trait, scales="free", nrow=2) +
+     theme_classic(base_size=18, base_family="myriad") +
+     theme(axis.title.x=element_blank(), axis.text.x=element_blank(),
+         axis.ticks.x=element_blank(),
+         strip.background=element_rect(fill="grey95", linetype="blank")) +
+     labs(y="length")
> ggsave(pdfFlNam, width=8, height=10, units="in", device="pdf", useDingbats=F)
> cat("\\includegraphics{", pdfFlNam, "}\n\n", sep="")
```

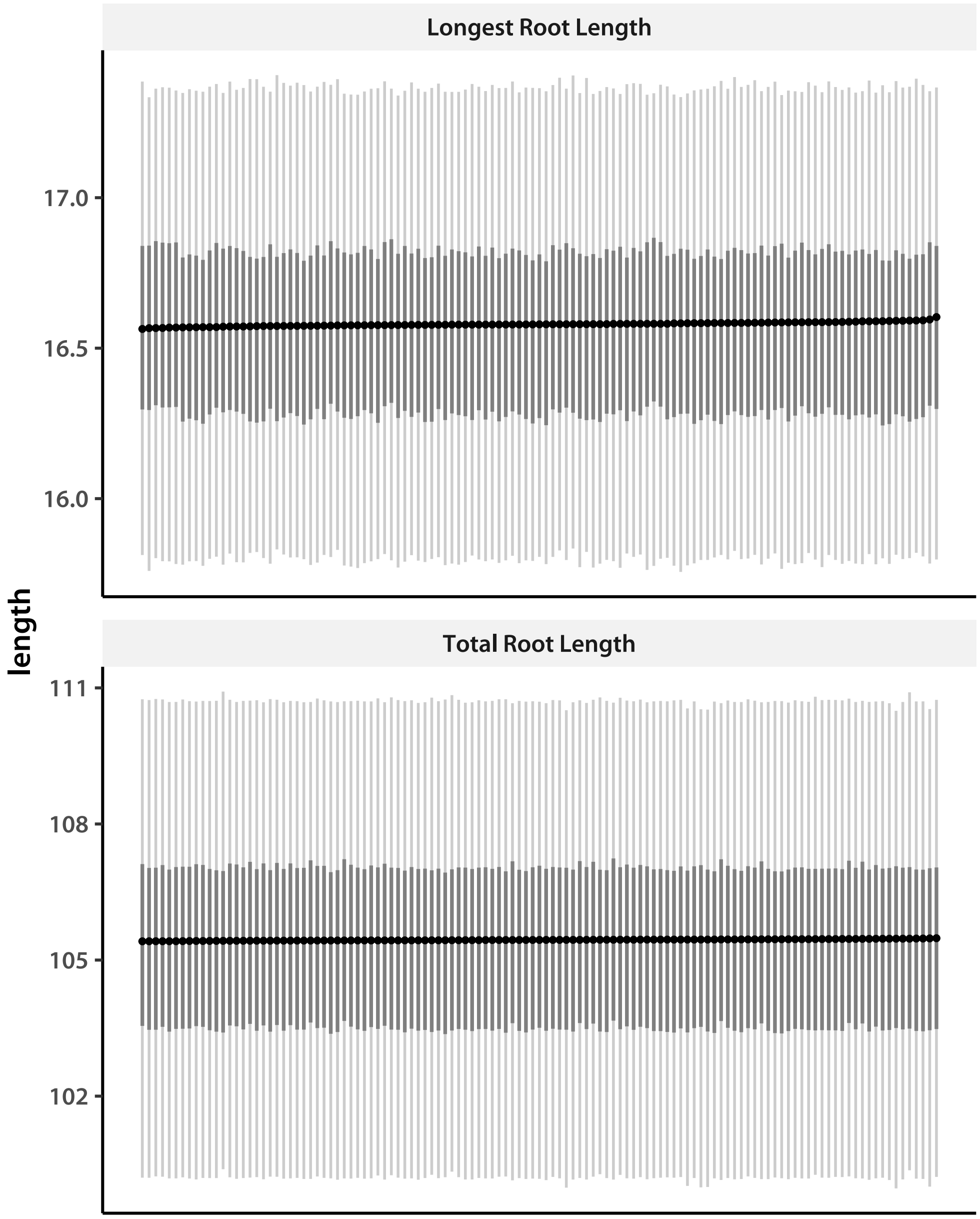


The correlations are high, but not as high as they were in the previous experiment. I calculate the $\log_{10}$(RRG) for each trait, and save point estimates.

```
> accNam  <- scan(file="accList.tsv", what = character())
> rrgMode <- log10(accMnExpt$mode/accMnCtrl$mode)
> modeMat <- cbind(matrix(accMnExpt$mode, ncol = 2, byrow = T),
+              matrix(accMnCtrl$mode, ncol = 2, byrow = T),
+              matrix(rrgMode, ncol = 2, byrow = T))
> cat(
+     apply(
```

```
+         rbind(c("HDRA_ID", paste(trtNam, "trt", sep = "_"),
+                 paste(trtNam, "ctrl", sep = "_"), paste(trtNam, "l10rrg", sep = "_")),
+             cbind(accNam, modeMat)), 1, paste, collapse = "\t"),
+     file = "accessionMeans.tsv", sep = "\n"
+     )
```

### 1 Heritability

I next estimate heritability, both broad-sense and marker-based narrow sense. Because I am using a Student-$t$ model for marker effects, I cannot use $\mathbf{\Sigma}_a$ directly, but have to calculate the variances from the GEBV estimates. In addition to broad-sense heritability, I am interested in the separate contribution of background effects. I first define the functions I need.

```
> makeVar <- cmpfun(function(vec){
+     apply(matrix(vec, ncol=d, byrow=T), 2, var)
+ })
> get.hsq <- cmpfun(function(vec){
+     vec[1:d]/rowSums(matrix(vec, nrow = d))
+ })
> get.Hsq <- cmpfun(function(vec){
+     (vec[1:d] + vec[(d+1):(2*d)])/rowSums(matrix(vec, nrow = d))
+ })
> get.bck <- cmpfun(function(vec){
+     (vec[(d+1):(2*d)])/rowSums(matrix(vec, nrow = d))
+ })
```

The function `get.hsq` calculates the marker heritability, which is a kind of narrow-sense heritability. It is

$$h^2 = \frac{\sigma^2_{\mathrm{GEBV}}}{\sigma^2_{\mathrm{GEBV}} + \sigma^2_s + \sigma^2_e}$$

for each treatment. The `get.Hsq` function calculates the broad-sense (among-accession) heritability:

$$H^2 = \frac{\sigma^2_{\mathrm{GEBV}} + \sigma^2_s}{\sigma^2_{\mathrm{GEBV}} + \sigma^2_s + \sigma^2_e}$$

The `get.bck` function calculates the fraction of total variance contributed by the background effects alone:

$$FVE_s = \frac{\sigma^2_s}{\sigma^2_{\mathrm{GEBV}} + \sigma^2_s + \sigma^2_e}$$

I read in covariance matrix chains, starting with the controls.

```
> diagInd <- diag(matrix(1:(d^2), ncol = d, byrow = T))
> chn1 <- matrix(.C("GSLmatLoad",
```

```
+     "chains/SgS_ctrl_IBSS_3_1000_1.gbin",
+     as.integer(chnLen), as.integer(d^2), out = double(chnLen*d^2))$out,
+         nrow = chnLen, byrow = T)[,diagInd]
> chn1 <- cbind(chn1,
+     matrix(.C("GSLmatLoad",
+     "chains/SgE_ctrl_IBSS_3_1000_1.gbin",
+     as.integer(chnLen), as.integer(d^2), out = double(chnLen*d^2))$out,
+         nrow = chnLen, byrow = T)[,diagInd])
> chn2 <- matrix(.C("GSLmatLoad",
+     "chains/SgS_ctrl_IBSS_3_1000_2.gbin",
+     as.integer(chnLen), as.integer(d^2), out = double(chnLen*d^2))$out,
+         nrow = chnLen, byrow = T)[,diagInd]
> chn2 <- cbind(chn2,
+     matrix(.C("GSLmatLoad",
+     "chains/SgE_ctrl_IBSS_3_1000_2.gbin",
+     as.integer(chnLen), as.integer(d^2), out = double(chnLen*d^2))$out,
+         nrow = chnLen, byrow = T)[,diagInd])
> chn3 <- matrix(.C("GSLmatLoad",
+     "chains/SgS_ctrl_IBSS_3_1000_3.gbin",
+     as.integer(chnLen), as.integer(d^2), out = double(chnLen*d^2))$out,
+         nrow = chnLen, byrow = T)[,diagInd]
> chn3 <- cbind(chn3,
+     matrix(.C("GSLmatLoad",
+     "chains/SgE_ctrl_IBSS_3_1000_3.gbin",
+     as.integer(chnLen), as.integer(d^2), out = double(chnLen*d^2))$out,
+         nrow = chnLen, byrow = T)[,diagInd])
> chn4 <- matrix(.C("GSLmatLoad",
+     "chains/SgS_ctrl_IBSS_3_1000_4.gbin",
+     as.integer(chnLen), as.integer(d^2), out = double(chnLen*d^2))$out,
+         nrow = chnLen, byrow = T)[,diagInd]
> chn4 <- cbind(chn4,
+     matrix(.C("GSLmatLoad",
+     "chains/SgE_ctrl_IBSS_3_1000_4.gbin",
+     as.integer(chnLen), as.integer(d^2), out = double(chnLen*d^2))$out,
+         nrow = chnLen, byrow = T)[,diagInd])
> chn5 <- matrix(.C("GSLmatLoad",
+     "chains/SgS_ctrl_IBSS_3_1000_5.gbin",
+     as.integer(chnLen), as.integer(d^2), out = double(chnLen*d^2))$out,
+         nrow = chnLen, byrow = T)[,diagInd]
> chn5 <- cbind(chn5,
+     matrix(.C("GSLmatLoad",
+     "chains/SgE_ctrl_IBSS_3_1000_5.gbin",
+     as.integer(chnLen), as.integer(d^2), out = double(chnLen*d^2))$out,
+         nrow = chnLen, byrow = T)[,diagInd])
> sigChnCtrl <- rbind(chn1, chn2, chn3, chn4, chn5)
```

```
> sigChnCtrl <- cbind(t(apply(chnGEBVctrl, 1, makeVar)), sigChnCtrl)
> chnhsqCtrl <- t(apply(sigChnCtrl, 1, get.hsq))
> chnHsqCtrl <- t(apply(sigChnCtrl, 1, get.Hsq))
> chnBckCtrl <- t(apply(sigChnCtrl, 1, get.bck))
```

Add the data from treated accessions.

```
> chn1 <- matrix(.C("GSLmatLoad",
+     "chains/SgS_expt_IBSS_3_1000_1.gbin",
+     as.integer(chnLen), as.integer(d^2), out = double(chnLen*d^2))$out,
+         nrow = chnLen, byrow = T)[,diagInd]
> chn1 <- cbind(chn1,
+     matrix(.C("GSLmatLoad",
+     "chains/SgE_expt_IBSS_3_1000_1.gbin",
+     as.integer(chnLen), as.integer(d^2), out = double(chnLen*d^2))$out,
+         nrow = chnLen, byrow = T)[,diagInd])
> chn2 <- matrix(.C("GSLmatLoad",
+     "chains/SgS_expt_IBSS_3_1000_2.gbin",
+     as.integer(chnLen), as.integer(d^2), out = double(chnLen*d^2))$out,
+         nrow = chnLen, byrow = T)[,diagInd]
> chn2 <- cbind(chn2,
+     matrix(.C("GSLmatLoad",
+     "chains/SgE_expt_IBSS_3_1000_2.gbin",
+     as.integer(chnLen), as.integer(d^2), out = double(chnLen*d^2))$out,
+         nrow = chnLen, byrow = T)[,diagInd])
> chn3 <- matrix(.C("GSLmatLoad",
+     "chains/SgS_expt_IBSS_3_1000_3.gbin",
+     as.integer(chnLen), as.integer(d^2), out = double(chnLen*d^2))$out,
+         nrow = chnLen, byrow = T)[,diagInd]
> chn3 <- cbind(chn3,
+     matrix(.C("GSLmatLoad",
+     "chains/SgE_expt_IBSS_3_1000_3.gbin",
+     as.integer(chnLen), as.integer(d^2), out = double(chnLen*d^2))$out,
+         nrow = chnLen, byrow = T)[,diagInd])
> chn4 <- matrix(.C("GSLmatLoad",
+     "chains/SgS_expt_IBSS_3_1000_4.gbin",
+     as.integer(chnLen), as.integer(d^2), out = double(chnLen*d^2))$out,
+         nrow = chnLen, byrow = T)[,diagInd]
> chn4 <- cbind(chn4,
+     matrix(.C("GSLmatLoad",
+     "chains/SgE_expt_IBSS_3_1000_4.gbin",
+     as.integer(chnLen), as.integer(d^2), out = double(chnLen*d^2))$out,
+         nrow = chnLen, byrow = T)[,diagInd])
> chn5 <- matrix(.C("GSLmatLoad",
+     "chains/SgS_expt_IBSS_3_1000_5.gbin",
+     as.integer(chnLen), as.integer(d^2), out = double(chnLen*d^2))$out,
```

```
+         nrow = chnLen, byrow = T)[,diagInd]
> chn5 <- cbind(chn5,
+     matrix(.C("GSLmatLoad",
+     "chains/SgE_expt_IBSS_3_1000_5.gbin",
+     as.integer(chnLen), as.integer(d^2), out = double(chnLen*d^2))$out,
+         nrow = chnLen, byrow = T)[,diagInd])
> sigChnExpt <- rbind(chn1, chn2, chn3, chn4, chn5)
> sigChnExpt <- cbind(t(apply(chnGEBVexpt, 1, makeVar)), sigChnExpt)
> chnhsqExpt <- t(apply(sigChnExpt, 1, get.hsq))
> chnHsqExpt <- t(apply(sigChnExpt, 1, get.Hsq))
> chnBckExpt <- t(apply(sigChnExpt, 1, get.bck))
> hsqHPD     <- as.data.frame(rbind(t(apply(chnhsqCtrl, 2, quantileLike)),
+                              t(apply(chnHsqCtrl, 2, quantileLike)),
+                              t(apply(chnhsqExpt, 2, quantileLike)),
+                              t(apply(chnHsqExpt, 2, quantileLike))))
> hsqHPD$trait        <- rep(rep(trtNam, times=2), times=d)
> hsqHPD$heritability <- rep(rep(c("GEBV", "Broad"), each=d), times=2)
> hsqHPD$treatment    <- rep(c("Control", "Treated"), each=2*d)
> fveHPD     <- as.data.frame(rbind(t(apply(chnhsqCtrl, 2, quantileLike)),
+                              t(apply(chnBckCtrl, 2, quantileLike)),
+                              t(apply(chnhsqExpt, 2, quantileLike)),
+                              t(apply(chnBckExpt, 2, quantileLike))))
> fveHPD$trait     <- rep(rep(trtNam, times=2), times=d)
> fveHPD$variance  <- factor(rep(rep(c("Marker", "Background"), each=d), times=2),
+                              levels=c("Marker", "Background"))
> fveHPD$treatment <- rep(c("Control", "Treated"), each=2*d)
```

Now plot both kinds of heritabilities on the same panel.

```
> pdfFlNam <- "heritabilityNEWqk.pdf"
> showtext_auto()
> ggplot(data=hsqHPD, aes(x=1:nrow(hsqHPD), y=mode, color=heritability)) +
+     geom_segment(aes(x=1:nrow(hsqHPD), y=lower95, xend=1:nrow(hsqHPD),
+         yend=upper95, color=heritability),
+         size=1.2) +
+     geom_segment(aes(x=1:nrow(hsqHPD), y=lower50, xend=1:nrow(hsqHPD),
+         yend=upper50, color=heritability),
+         size=1.75) +
+     geom_point(size=2) +
+     facet_wrap(~trait, ncol=2) +
+     theme_classic(base_size=18, base_family="myriad") +
+     theme(legend.title=element_blank(),
+         strip.background=element_rect(fill="grey95", linetype="blank")) +
+     scale_x_continuous(breaks=c(3,7), labels=c("Control", "Treated")) +
+     theme(panel.spacing.x=unit(4.0, "lines"), axis.ticks.x=element_blank()) +
+     ylim(c(0,1)) +
```

```
+     labs(y="heritability", x="treatment")
> ggsave(pdfFlNam, width=10, height=8, units="in", device="pdf", useDingbats=F)
> cat("\\includegraphics{", pdfFlNam, "}\n\n", sep="")
```

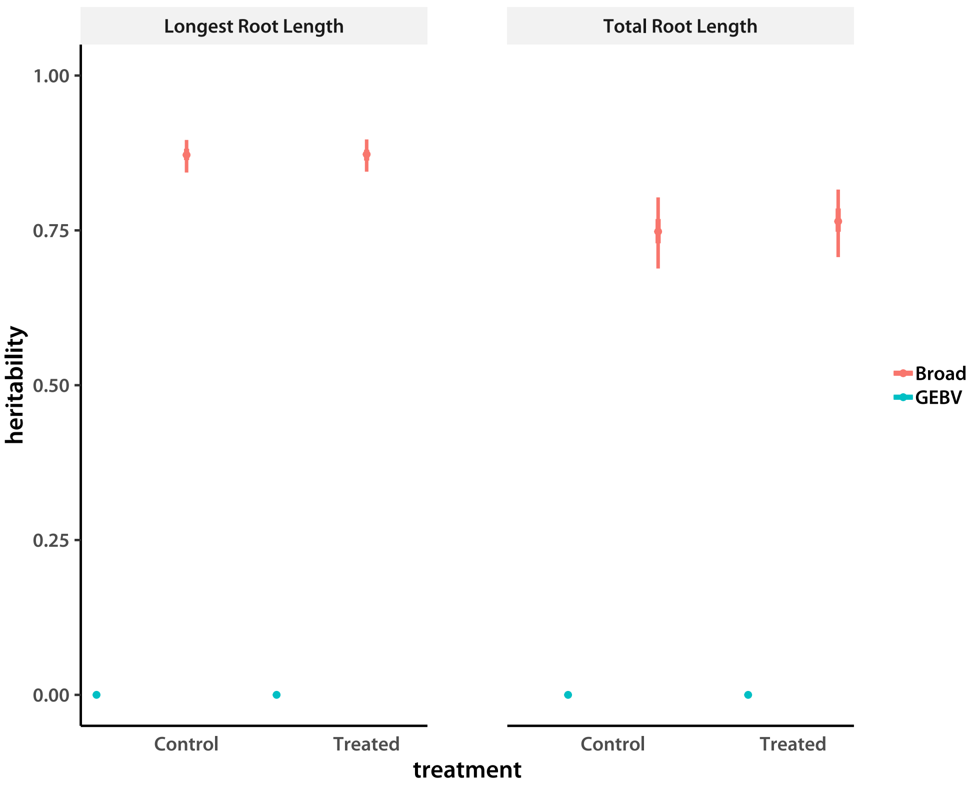


Finally plot fractions of variance explained on the same panel.

```
> pdfFlNam <- "fveNEWqk.pdf"
> showtext_auto()
> ggplot(data=fveHPD, aes(x=1:nrow(fveHPD), y=mode, color=variance)) +
+     geom_segment(aes(x=1:nrow(fveHPD), y=lower95, xend=1:nrow(fveHPD),
+         yend=upper95, color=variance),
+         size=1.2) +
+     geom_segment(aes(x=1:nrow(fveHPD), y=lower50, xend=1:nrow(fveHPD),
+         yend=upper50, color=variance),
+         size=1.75) +
+     geom_point(size=2) +
+     facet_wrap(~trait, ncol=2) +
+     theme_classic(base_size=18, base_family="myriad") +
+     theme(legend.title=element_blank(),
+         strip.background=element_rect(fill="grey95", linetype="blank")) +
+     scale_x_continuous(breaks=c(3,7), labels=c("Control", "Treated")) +
+     theme(panel.spacing.x=unit(4.0, "lines"), axis.ticks.x=element_blank()) +
+     ylim(c(0,1)) +
```

```
+     labs(y="fraction of variance", x="treatment")
> ggsave(pdfFlNam, width=10, height=8, units="in", device="pdf", useDingbats=F)
> cat("\\includegraphics{", pdfFlNam, "}\n\n", sep="")
```

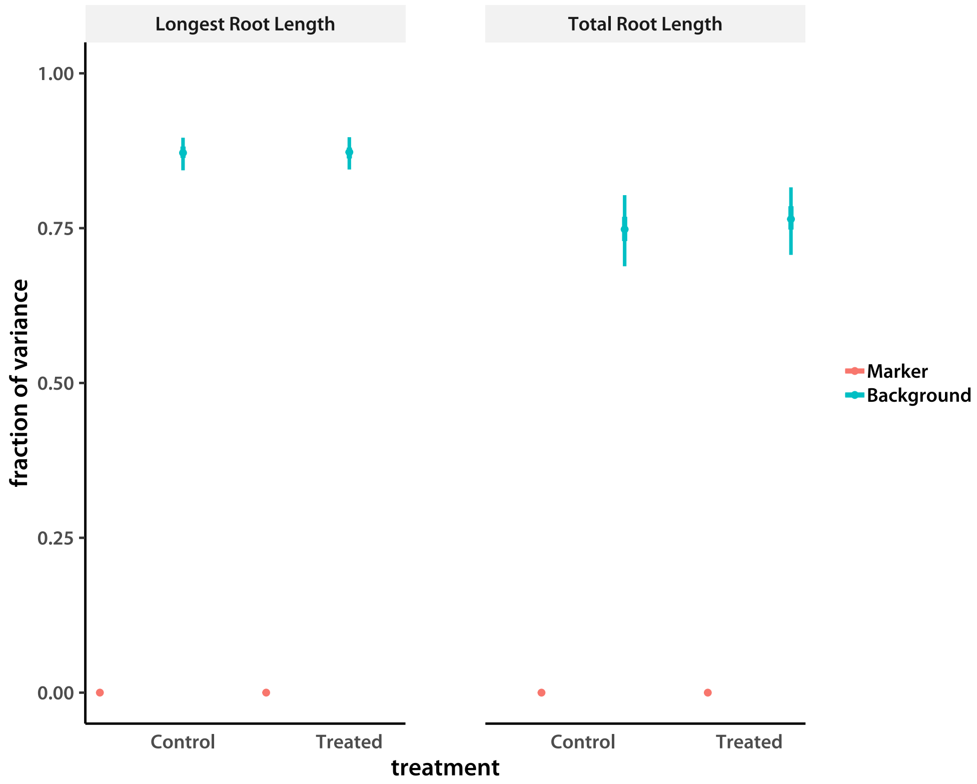
