## Supplemental methods and scripts for "Low additive genetic variation in a trait under selection in domesticated rice": phenoDataPrepFAM.pdf

### Preparing raw phenotype data for analysis

Tony Greenberg

December 21, 2018

```
R version 3.5.1 (2018-07-02)
Platform: x86_64-pc-linux-gnu (64-bit)
Running under: Pop!_OS 18.10

Matrix products: default
BLAS/LAPACK: /opt/intel/compilers_and_libraries_2019.1.144/linux/mkl/lib/intel64_lin/libmkl_gf_lp64.so

locale:
 [1] LC_CTYPE=en_US.UTF-8       LC_NUMERIC=C               LC_TIME=en_US.UTF-8
 [4] LC_COLLATE=en_US.UTF-8     LC_MONETARY=en_US.UTF-8    LC_MESSAGES=en_US.UTF-8
 [7] LC_PAPER=en_US.UTF-8       LC_NAME=C                  LC_ADDRESS=C
[10] LC_TELEPHONE=C             LC_MEASUREMENT=en_US.UTF-8 LC_IDENTIFICATION=C

attached base packages:
[1] compiler  stats     graphics  grDevices utils     datasets  methods   base

other attached packages:
[1] showtext_0.5-1 showtextdb_2.0 sysfonts_0.8   gridExtra_2.3  ggplot2_3.1.0

loaded via a namespace (and not attached):
 [1] Rcpp_1.0.0       crayon_1.3.4     withr_2.1.2      grid_3.5.1       plyr_1.8.4
 [6] gtable_0.2.0     scales_1.0.0     pillar_1.3.0     rlang_0.3.0.1    lazyeval_0.2.1
[11] tools_3.5.1      munsell_0.5.0    colorspace_1.3-2 tibble_1.4.2
```

In this document I detail the preparation of raw phenotype data from the initial tests of Al tolerance. I start by reading the phenotype data file. These data were provided by Lyza. The original file was modified to fix formatting and remove the caiapo control lines.

```
> AlTolRaw <- read.table("AlTolFixed.dat", sep = "\t", header = T)
> summary(AlTolRaw)
```

```
   Line_label     Genotype_ID        Tub       treatment          rep
 L85    : 180   NSFTV85 : 180          :1027   0uM  :5180   Min.   : 1.0
 L5     : 123   NSFTV152: 100   8      : 364   160uM:5174   1st Qu.: 3.0
 L152   : 100   NSFTV200: 100   6      : 361                Median : 6.0
 L200   : 100   NSFTV88 : 100   5      : 360                Mean   :10.4
 L88    : 100   NSFTV128:  99   2      : 358                3rd Qu.: 9.0
 L128   :  99   NSFTV174:  99   3      : 356                Max.   :96.0
 (Other):9652   (Other) :9676   (Other):7528
```

```
     Original_sample_ID   day_3_TRL           Experiment
 1_Tub1A_0uM_01:    1    Min.   :  0.29   2010_panelA :1824
 1_Tub1A_0uM_02:    1    1st Qu.: 30.41   2010_panelB :1688
 1_Tub1A_0uM_03:    1    Median : 55.66   2015        :1575
 1_Tub1A_0uM_04:    1    Mean   : 62.41   2008_screen3:1116
 1_Tub1A_0uM_05:    1    3rd Qu.: 88.26   2008_screen2:1054
 1_Tub1A_0uM_06:    1    Max.   :255.62   2008_screen4:1048
 (Other)       :10348                     (Other)     :2049
```

I next look for accessions that have been genotyped and retain only data for those. These accessions are in the first column of the HDRA meta-data file I read in the following segment.

```
> nsfFl  <- pipe("cut -f1 HDRA-G6-4-RDP1-RDP2-sativa-only_listPop.txt")
> nsfInd <- scan(nsfFl, what = character())
> close(nsfFl)
> nrow(AlTolRaw)

[1] 10354

> AlTolRaw <- AlTolRaw[as.character(AlTolRaw$Geno) %in% nsfInd,]
> nrow(AlTolRaw)

[1] 9015

> AlTolRaw$repID <- factor(paste(AlTolRaw$Genotype_ID, AlTolRaw$Experiment,
+     AlTolRaw$rep, sep = "."))
> AlTolRaw$Genotype_ID <- factor(as.character(AlTolRaw$Genotype_ID))
```

The phenotype table lists the control and Al treated plants as a factor. I would like to transform this into a matrix with treatment and control as "traits." The base function that does this takes advantage of the fact that the controls and treatments are listed continuously one after the other within each experiment. It also inserts `NA` as appropriate when the number of control and treatment plants is not equal.

```
> insertNA <- cmpfun(function(ind, v1, v2){
+     ln <- ind[3:4] - ind[1:2]
+     rbind( c(v1[ind[1]:ind[3]], rep(NA, times = max(ln) - ln[1])),
+         c(v2[ind[2]:ind[4]], rep(NA, times = max(ln) - ln[2])) )
+ })
```

Next I write the function that will transform the raw table into a matrix with the treatment levels as traits. As a by-product, I also generate the relevant factors.

```
> YmatAll    <- NULL
> lineFacAll <- NULL
> exprFacAll <- NULL
> buildY <- cmpfun(function(lnNam){
```

```
+     trtExpTab <- table(
+         as.character(AlTolRaw$Experiment[as.character(AlTolRaw$Genotype_ID) == lnNam]),
+         as.character(AlTolRaw$treatment[as.character(AlTolRaw$Genotype_ID) == lnNam]))
+     expNam <- NULL
+
+     if (sum(dim(trtExpTab)) == 2){
+         cat("WARNING: either treatment or control for line ", lnNam,
+             " is completely absent\n")
+         return(NULL)
+     }
+     if (0 %in% c(trtExpTab)){
+         non0ind <- apply(trtExpTab, 1, function(vec){!( 0 %in% vec )})
+         rwN     <- apply(matrix(trtExpTab[non0ind,], ncol = 2), 1, max)
+
+         expNam <- rownames(trtExpTab)[non0ind]
+         exprFacAll <<- c(exprFacAll, rep(rownames(trtExpTab)[non0ind], rwN))
+         lineFacAll <<- c(lineFacAll, rep(lnNam, sum(rwN)))
+
+         trtExpTab <- matrix(trtExpTab[non0ind,], ncol = 2)
+         trtExpTab <- matrix(apply(trtExpTab, 2, cumsum), ncol = 2)
+
+     } else {
+         rwN <- apply(trtExpTab, 1, max)
+
+         expNam <- rownames(trtExpTab)
+         exprFacAll <<- c(exprFacAll, rep(rownames(trtExpTab), rwN))
+         lineFacAll <<- c(lineFacAll, rep(lnNam, sum(rwN)))
+
+         trtExpTab <- matrix(apply(trtExpTab, 2, cumsum), ncol = 2)
+     }
+
+     ctrl <- AlTolRaw$day_3_TRL[ (AlTolRaw$treatment == "0uM")
+         & (as.character(AlTolRaw$Genotype_ID) == lnNam)
+         & (as.character(AlTolRaw$Experiment) %in% expNam) ]
+     trt  <- AlTolRaw$day_3_TRL[ (AlTolRaw$treatment == "160uM")
+         & (as.character(AlTolRaw$Genotype_ID) == lnNam)
+         & (as.character(AlTolRaw$Experiment) %in% expNam) ]
+
+     if (nrow(trtExpTab) == 1){
+         YmatAll <<- rbind( YmatAll,
+             cbind(c(ctrl, rep(NA, times = max(c(trtExpTab)) - c(trtExpTab)[1])),
+                 c(trt, rep(NA, times = max(c(trtExpTab)) - c(trtExpTab)[2]))) )
+     } else {
+         indTab  <- cbind( rbind(rep(1,2), trtExpTab[-nrow(trtExpTab),] + 1), trtExpTab )
+
```

```
+         YmatAll <<- rbind( YmatAll,
+             matrix(unlist(apply(indTab, 1, insertNA, ctrl, trt)),
+                 ncol = 2, byrow = T) )
+     }
+     return(NULL)
+ })
> trash <- sapply(levels(AlTolRaw$Genotype_ID), buildY)
```

```
WARNING: either treatment or control for line  NSFTV250  is completely absent
```

There is a warning about accession NSFTV250. Check the original data table.

```
> AlTolRaw[AlTolRaw$Geno == "NSFTV250",]
```

```
     Line_label Genotype_ID  Tub treatment rep Original_sample_ID day_3_TRL  Experiment
7522       L250    NSFTV250 Tub7       0uM   1    250_Tub7_0uM_01    120.48 2010_panelA
7523       L250    NSFTV250 Tub7       0uM   2    250_Tub7_0uM_02    129.00 2010_panelA
7524       L250    NSFTV250 Tub7       0uM   3    250_Tub7_0uM_03    121.48 2010_panelA
7525       L250    NSFTV250 Tub7       0uM   4    250_Tub7_0uM_04    145.33 2010_panelA
7526       L250    NSFTV250 Tub7       0uM   5    250_Tub7_0uM_05    136.86 2010_panelA
7527       L250    NSFTV250 Tub7       0uM   6    250_Tub7_0uM_06    157.24 2010_panelA
7528       L250    NSFTV250 Tub7       0uM   7    250_Tub7_0uM_07    148.15 2010_panelA
7529       L250    NSFTV250 Tub7       0uM   8    250_Tub7_0uM_08    184.68 2010_panelA
7530       L250    NSFTV250 Tub7       0uM   9    250_Tub7_0uM_09     89.66 2010_panelA
7531       L250    NSFTV250 Tub7       0uM  10    250_Tub7_0uM_10    163.52 2010_panelA
                        repID
7522  NSFTV250.2010_panelA.1
7523  NSFTV250.2010_panelA.2
7524  NSFTV250.2010_panelA.3
7525  NSFTV250.2010_panelA.4
7526  NSFTV250.2010_panelA.5
7527  NSFTV250.2010_panelA.6
7528  NSFTV250.2010_panelA.7
7529  NSFTV250.2010_panelA.8
7530  NSFTV250.2010_panelA.9
7531 NSFTV250.2010_panelA.10
```

Indeed, no treatment data.

In addition, manual examination revealed that the tub9 and tub10 in the 2010_panelB are most likely switched. Since these account for few data points, I will eliminate these tubs. However, one accession (NSFTV128) has data from other experiments, so I keep that.

```
> badGeno <- unique(as.character(AlTolRaw[(AlTolRaw$Exper == "2010_panelB")
+     & (AlTolRaw$Tub %in% c("Tub10", "Tub9")),][,2]))
> badGeno <- badGeno[!(badGeno %in% AlTolRaw$Geno[AlTolRaw$Exper != "2010_panelB"])]
> badGeno <- c(badGeno, "NSFTV250")
> badGeno
```

```
 [1] "NSFTV108" "NSFTV177" "NSFTV182" "NSFTV186" "NSFTV192" "NSFTV211" "NSFTV218"
 [8] "NSFTV232" "NSFTV236" "NSFTV249" "NSFTV300" "NSFTV302" "NSFTV307" "NSFTV387"
[15] "NSFTV250"
```

I rebuild the phenotype matrix.

```
> buildY <- cmpfun(function(lnNam){
+     trtExpTab <- NULL
+     if(lnNam == "NSFTV128"){ # this line has bad and good experiments
+         trtExpTab <- table(
+             as.character(AlTolRaw$Experiment[(as.character(AlTolRaw$Genotype_ID) == lnNam)
+                 & (as.character(AlTolRaw$Experiment) != "2010_panelB")]),
+             as.character(AlTolRaw$treatment[(as.character(AlTolRaw$Genotype_ID) == lnNam)
+                 & (as.character(AlTolRaw$Experiment) != "2010_panelB")]))
+     }
+     else {
+         trtExpTab <- table(
+             as.character(AlTolRaw$Experiment[as.character(AlTolRaw$Genotype_ID) == lnNam]),
+             as.character(AlTolRaw$treatment[as.character(AlTolRaw$Genotype_ID) == lnNam]))
+     }
+     expNam <- NULL
+
+     if (sum(dim(trtExpTab)) == 2){
+         cat("WARNING: either treatment or control for line ", lnNam,
+             " is completely absent\n")
+         return(NULL)
+     }
+     if (0 %in% c(trtExpTab)){
+         non0ind <- apply(trtExpTab, 1, function(vec){!( 0 %in% vec )})
+         rwN     <- apply(matrix(trtExpTab[non0ind,], ncol = 2), 1, max)
+
+         expNam <- rownames(trtExpTab)[non0ind]
+         exprFacAll <<- c(exprFacAll, rep(rownames(trtExpTab)[non0ind], rwN))
+         lineFacAll <<- c(lineFacAll, rep(lnNam, sum(rwN)))
+
+         trtExpTab <- matrix(trtExpTab[non0ind,], ncol = 2)
+         trtExpTab <- matrix(apply(trtExpTab, 2, cumsum), ncol = 2)
+
+     } else {
+         rwN <- apply(trtExpTab, 1, max)
+
+         expNam <- rownames(trtExpTab)
+         exprFacAll <<- c(exprFacAll, rep(rownames(trtExpTab), rwN))
+         lineFacAll <<- c(lineFacAll, rep(lnNam, sum(rwN)))
+
+         trtExpTab <- matrix(apply(trtExpTab, 2, cumsum), ncol = 2)
+     }
+
+     ctrl <- AlTolRaw$day_3_TRL[ (AlTolRaw$treatment == "0uM")
+         & (as.character(AlTolRaw$Genotype_ID) == lnNam)
```

```
+         & (as.character(AlTolRaw$Experiment) %in% expNam) ]
+     trt  <- AlTolRaw$day_3_TRL[ (AlTolRaw$treatment == "160uM")
+         & (as.character(AlTolRaw$Genotype_ID) == lnNam)
+         & (as.character(AlTolRaw$Experiment) %in% expNam) ]
+
+     if (nrow(trtExpTab) == 1){
+         YmatAll <<- rbind( YmatAll,
+             cbind(c(ctrl, rep(NA, times = max(c(trtExpTab)) - c(trtExpTab)[1])),
+                 c(trt, rep(NA, times = max(c(trtExpTab)) - c(trtExpTab)[2]))) )
+     } else {
+         indTab  <- cbind( rbind(rep(1,2), trtExpTab[-nrow(trtExpTab),] + 1), trtExpTab )
+
+         YmatAll <<- rbind( YmatAll,
+             matrix(unlist(apply(indTab, 1, insertNA, ctrl, trt)), ncol = 2, byrow = T) )
+     }
+     return(NULL)
+ })
> goodGeno <- levels(AlTolRaw$Genotype_ID)[!(levels(AlTolRaw$Genotype_ID) %in% badGeno)]
> YmatAll    <- NULL
> lineFacAll <- NULL
> exprFacAll <- NULL
> trash <- sapply(goodGeno, buildY)
> dim(YmatAll)

[1] 4428    2

> length(lineFacAll)

[1] 4428

> length(unique(lineFacAll))

[1] 361
```

Make the experiment factor levels a bit easier to fit on plots.

```
> exprFacAll <- sub("panel", "", sub("screen", "", exprFacAll))
```

I build a data frame with the phenotypic data, adding experiment, accession, and population information for plotting. I also create a version of the population index that is easier to see on plots.

```
> popInd <- matrix(scan(file="HDRA-G6-4-RDP1-RDP2-sativa-only_listPop.txt",
+                           what = character()), ncol=3, byrow=T)
> rownames(popInd) <- nsfInd
> popInd     <- popInd[lineFacAll,]
```

```
> displayPop <- popInd[,2]
> displayPop[grep("admixed", popInd[,2])]        <- "ADM"
> displayPop[popInd[,2] == "aus"]                <- "AUS"
> displayPop[popInd[,2] == "indica"]             <- "IND"
> displayPop[popInd[,2] == "aromatic"]           <- "ARO"
> displayPop[popInd[,2] == "temperate-japonica"] <- "TEJ"
> displayPop[popInd[,2] == "tropical-japonica"]  <- "TRJ"
> Yframe <- data.frame(TRSG      = c(YmatAll),
+                     treatment  = rep(c("Control", "Treated"), each=nrow(YmatAll)),
+                     experiment = rep(exprFacAll, 2),
+                     population = factor(rep(displayPop, 2),
+                             levels=c("IND", "AUS", "ARO", "TRJ", "TEJ", "ADM")),
+                     NSFTV.ID   = factor(rep(popInd[,1], 2)),
+                     HDRA.ID    = factor(rep(popInd[,3], 2))
+                     )
> exprFacAll <- rep(exprFacAll, 2)[!is.na(Yframe$TRSG)]
> Yframe     <- Yframe[!is.na(Yframe$TRSG),]
```

I plot overall histograms of the phenotype values.

```
> pdfFlNam <- "rawHist.pdf"
> showtext_auto()
> ggplot(data=Yframe, aes(x=TRSG, ..density.., fill=treatment)) +
+     geom_histogram(bins=200) +
+     theme_classic(base_size=18, base_family="myriad") +
+     xlab("total root system growth")
> ggsave(pdfFlNam, width=8, height=8, units="in", device="pdf", useDingbats=F)
> cat("\\includegraphics{", pdfFlNam, "}\n\n", sep="")
```

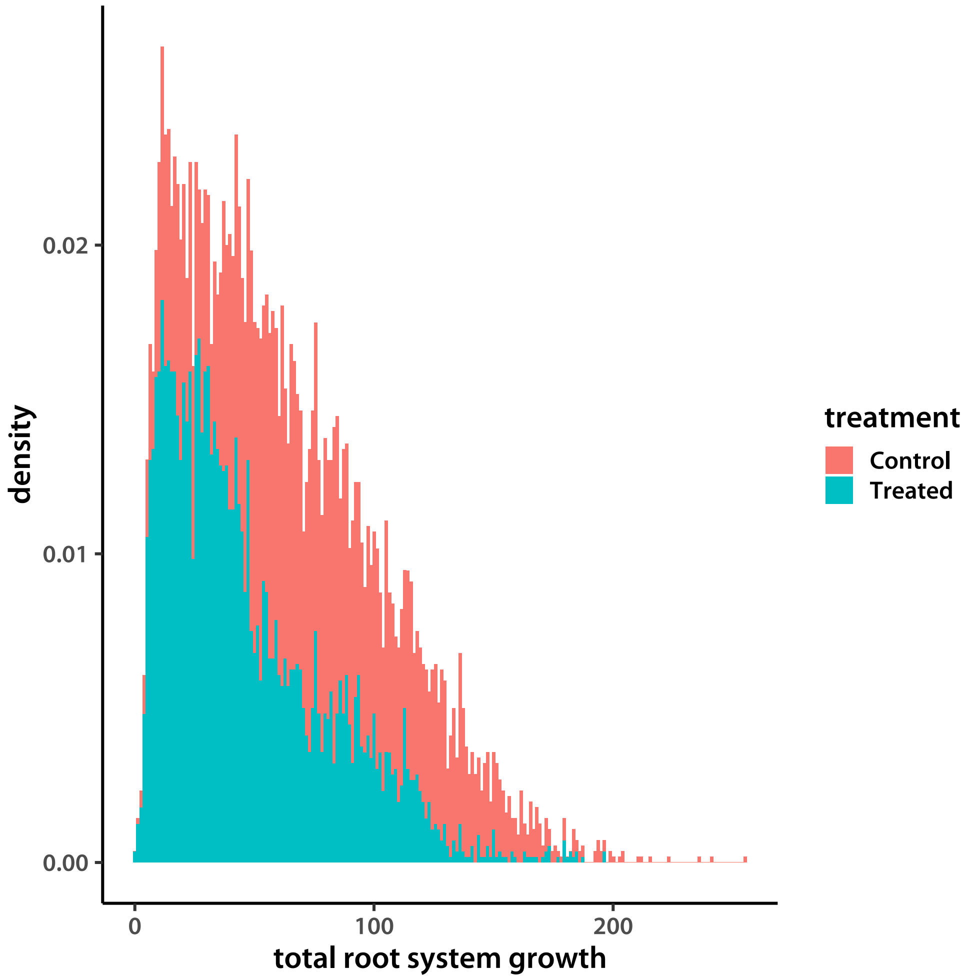


There are no outliers that suggest data entry errors. I next check the magnitude of the experiment effect, first creating a data frame to use for plotting.

```
> pdfFlNam <- "experimentVal.pdf"
> showtext_auto()
> ggplot(data=Yframe, aes(y=TRSG, x=experiment)) +
+     geom_boxplot(fill="grey80") +
+     theme_classic(base_size=18, base_family="myriad") +
+     facet_wrap(~treatment, scales="free_y", nrow=2) +
+     ylab("total root system growth") +
+     theme(axis.text.x=element_text(angle=-45))
> ggsave(pdfFlNam, width=8, height=10, units="in", device="pdf", useDingbats=F)
> cat("\\includegraphics{", pdfFlNam, "}\n\n", sep="")
```

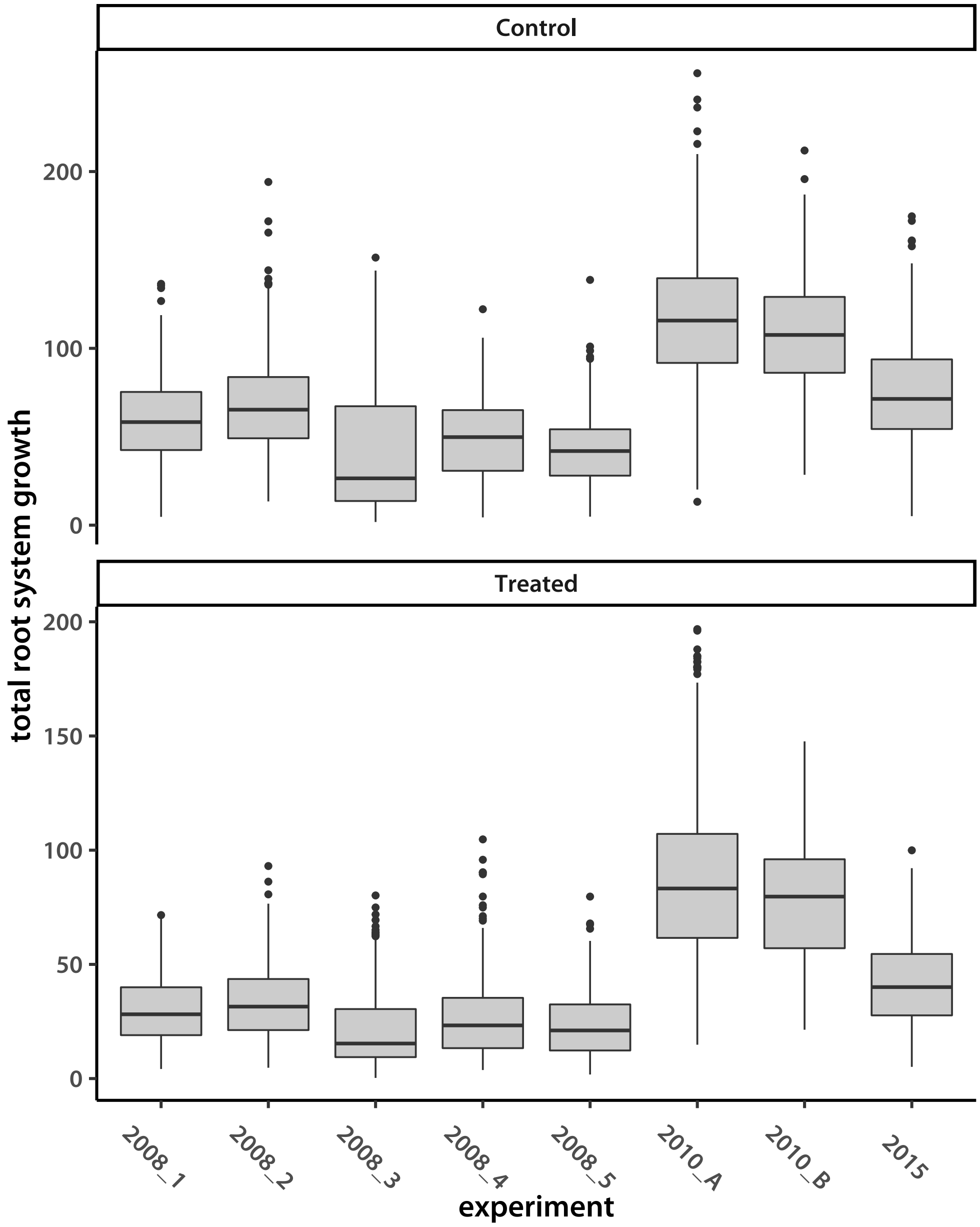


Now separate by population.

```
> pdfFlNam <- "experimentValPop.pdf"
> ggplot(data=Yframe, aes(y=TRSG, x=experiment, fill=population)) +
+     geom_boxplot() +
+     theme_classic(base_size=18, base_family="myriad") +
+     facet_wrap(~treatment, scales="free_y", nrow=2) +
+     ylab("total root system growth") +
+     theme(axis.text.x=element_text(angle=-45))
> showtext_auto()
```

```
> ggsave(pdfFlNam, width=8, height=10, units="in", device="pdf", useDingbats=F)
> cat("\\includegraphics{", pdfFlNam, "}\n\n", sep="")
```

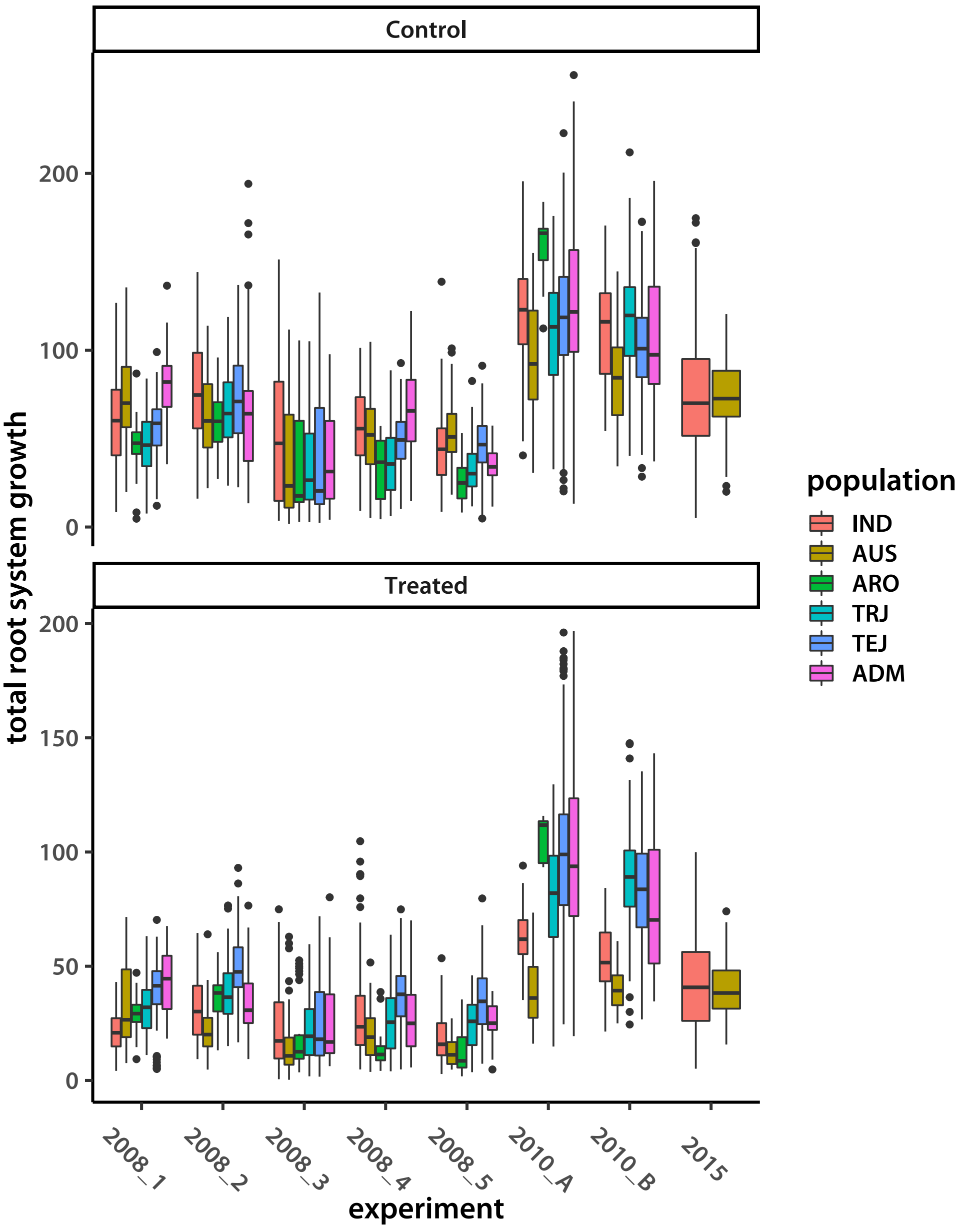


I also look at the number of samples by population in each experiment.

```
> pdfFlNam <- "experimentBar.pdf"
> ctVec <- tapply(1:nrow(Yframe), paste(Yframe$treatment,
+                                          Yframe$population,
+                                          Yframe$experiment, sep="."),
+                 length)
```

```
> ctID  <- matrix(unlist(strsplit(names(ctVec), "\\.")), ncol=3, byrow=T)
> ctFrame <- data.frame(count=ctVec,
+                     treatment=ctID[,1], population=ctID[,2], experiment=ctID[,3])
> showtext_auto()
> ggplot(data=ctFrame, aes(y=count, x=experiment, fill=population)) +
+     geom_bar(stat="identity") +
+     theme_classic(base_size=18, base_family="myriad") +
+     facet_wrap(~treatment, scales="free_y", nrow=2) +
+     theme(axis.text.x=element_text(angle=-45))
> ggsave(pdfFlNam, width=8, height=10, units="in", device="pdf", useDingbats=F)
> cat("\\includegraphics{", pdfFlNam, "}\n\n", sep="")
```

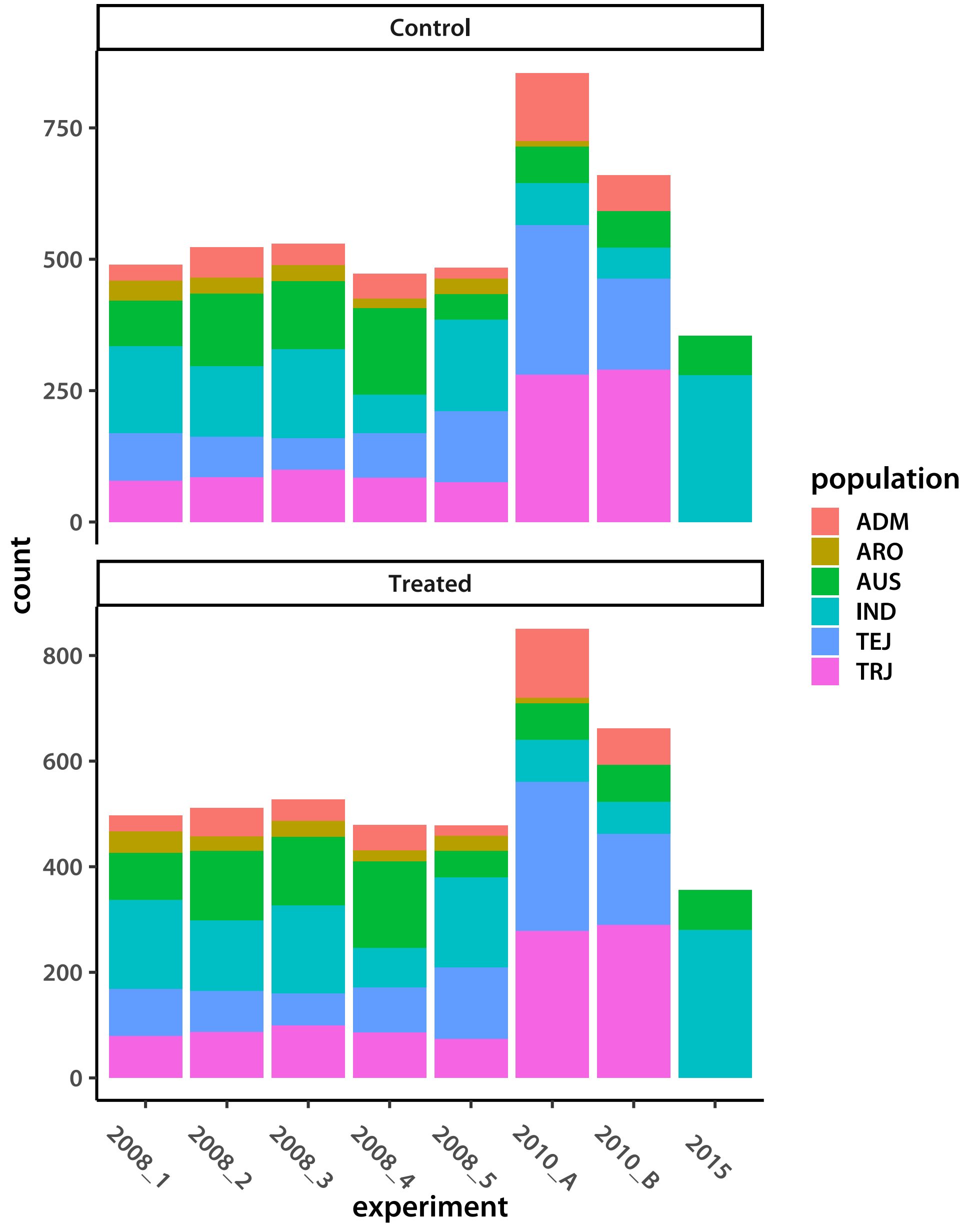


Population and experiment are somewhat confounded. In particular, 2015 is only *indica* and *aus*. Looking at the graph, it seems that distributions for these populations within experiment 2008_2 are the closest to 2015. I rename 2008_2 and 2015 as 2008_15.

```
> exprFacAll[exprFacAll == "2015"] <- "2008_15"
> exprFacAll[exprFacAll == "2008_2"] <- "2008_15"
> Yframe$experiment <- factor(exprFacAll)
```

Re-plot to see the effect of this arrangement.

```
> pdfFlNam <- "experimentValPooled.pdf"
> showtext_auto()
```

```
> ggplot(data=Yframe, aes(y=TRSG, x=experiment)) +
+     geom_boxplot(fill="grey80") +
+     theme_classic(base_size=18, base_family="myriad") +
+     facet_wrap(~treatment, scales="free_y", nrow=2) +
+     ylab("total root system growth") +
+     theme(axis.text.x=element_text(angle=-45))
> ggsave(pdfFlNam, width=8, height=10, units="in", device="pdf", useDingbats=F)
> cat("\\includegraphics{", pdfFlNam, "}\n\n", sep="")
```

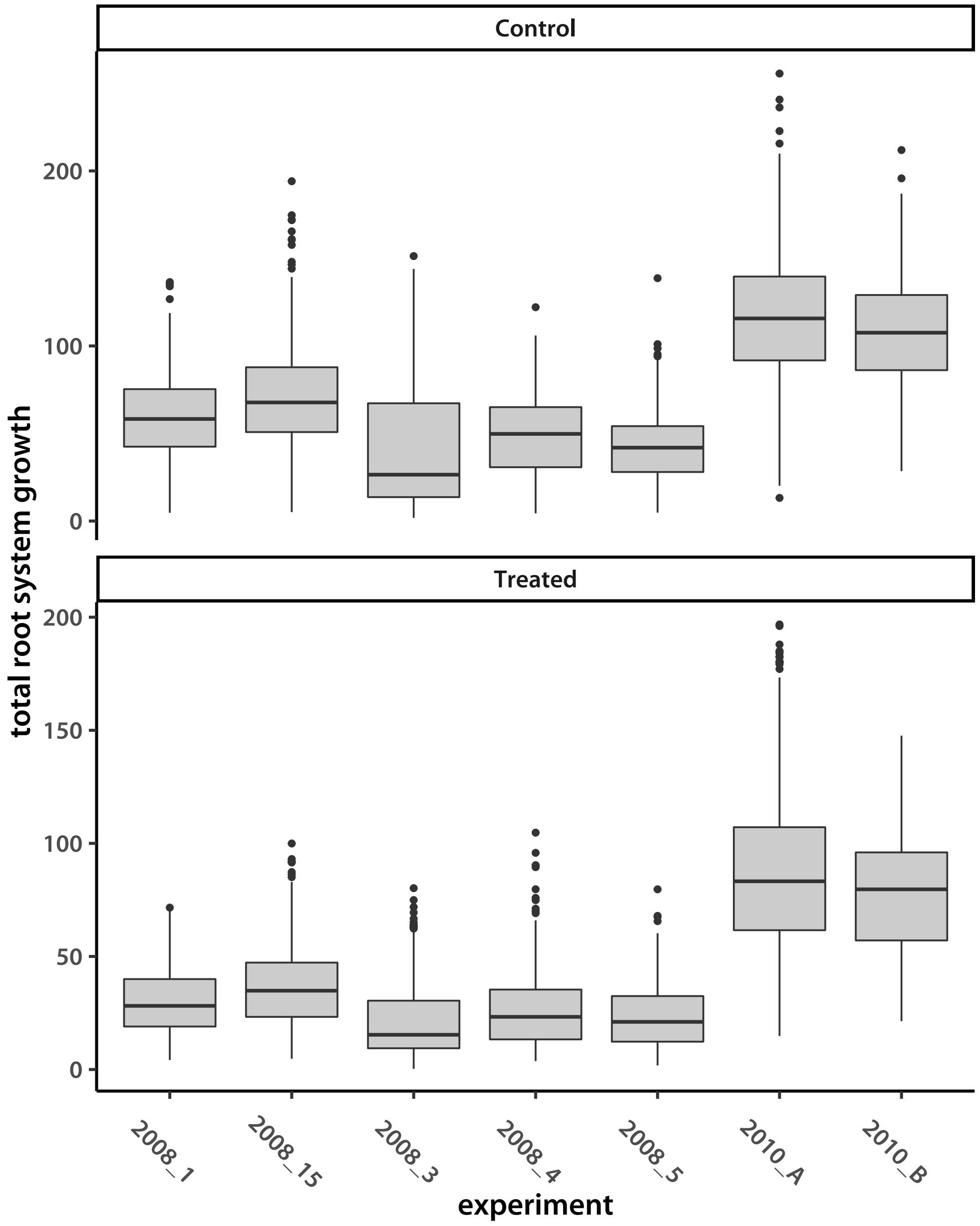


I save the raw data frame to a file.

```
> write.table(Yframe, file="famosoRAW.tsv", quote=F, sep="\t", row.names=F)
```

I prepare the data for output to files that will be used by the `MuGen` library-based code.

```
> matNAindALL <- apply(YmatAll, 2, function(vec){ifelse(is.na(vec), 1, 0)})
> signif(colSums(matNAindALL)/nrow(YmatAll),4)

[1] 0.01378 0.01513

> totNAindALL <- rowSums(matNAindALL)
> cat(unique(lineFacAll), file = "lineIDsAll.txt", sep = " ")
> blkIdx <- 1:2
```

I make a contrast matrix for the experiment factor. Since there are few levels, I will model with a vague prior (analogous to a fixed effect). The `2008_1` level is the base, simply because it is the first on the list.

```
> expMatAll <- sapply(unique(exprFacAll)[-1], function(val){as.integer(exprFacAll==val)})
> dim(expMatAll)

[1] 8728    6
```

Next I split the data into populations. First, generate the indexes for regular populations.

```
> IAind  <- popInd[,2] %in% c("admixed-indica", "aus", "indica")
> AUSind <- popInd[,2]  == "aus"
> JAPind <- popInd[,2] %in%
+     c("admixed-japonica", "temperate-japonica", "tropical-japonica", "aromatic")
> TRJind <- popInd[,2] == "tropical-japonica"
> sum(IAind)

[1] 2009

> sum(AUSind)

[1] 794

> sum(JAPind)

[1] 2349

> sum(TRJind)

[1] 1007
```

Next, I extract tropical *japonica* accessions that also have a significant admixture from *indica* and *aus*.

```
> admxFrame <- read.table(file="FastStructRes_K7_subpop_DW.csv", sep="\t", header=T)
> indSum    <- rowSums(admxFrame[,c("IND1", "IND2", "IND3", "AUS")])
> trjAdmNam <- as.character(admxFrame$ID)[(admxFrame$TRJ >= 0.3) & (indSum >= 0.1)
+             & (admxFrame$TEJ <= 0.1) & (admxFrame$ARO <= 0.1)]
> trjAdmNam <- trjAdmNam[grep("\\.0$", trjAdmNam)]
> arrFl     <- pipe("cut -f1,3 HDRA-G6-4-RDP1-RDP2-sativa-only_listPop.txt")
> arrInd    <- matrix(scan(arrFl, what = character()), ncol=2, byrow=T)
> close(arrFl)
> rownames(arrInd) <- arrInd[,1]
> trjAdmNam[trjAdmNam %in% arrInd[unique(lineFacAll),2]]
```

```
[1] "818143d8.0" "b0816082.0" "f25ad349.0" "5c592759.0" "642238d9.0" "c7301040.0"
```

```
> admxFrame[as.character(admxFrame$ID) %in%
+     trjAdmNam[trjAdmNam %in% arrInd[unique(lineFacAll),2]],]
```

```
            ID      TEJ      TRJ IND1     IND2      AUS      ARO     IND3
209 818143d8.0 0.000000 0.881691    0 0.000000 0.118307 0.000000 0.000000
245 b0816082.0 0.012675 0.804436    0 0.182887 0.000000 0.000000 0.000000
362 f25ad349.0 0.074808 0.815979    0 0.000000 0.000000 0.000000 0.109211
729 5c592759.0 0.061996 0.419183    0 0.518820 0.000000 0.000000 0.000000
797 642238d9.0 0.000000 0.530285    0 0.096401 0.000000 0.000000 0.373312
846 c7301040.0 0.000000 0.459639    0 0.000000 0.099597 0.008283 0.432479
              subpop  subpop.new
209 tropical-japonica         trj
245 tropical-japonica         trj
362 tropical-japonica         trj
729           admixed admixed-jap
797           admixed admixed-jap
846           admixed admixed-jap
```

There are six accessions I can use. Too few to analyze by themselves, but I will add them to the `TRJ` group to make `TRJA` and test that.

```
> TRJAind  <- TRJind
> trjaSubs <- trjAdmNam[trjAdmNam %in% arrInd[unique(lineFacAll),2]]
> TRJAind[names(TRJAind) %in% arrInd[arrInd[,2] %in% trjaSubs,1]] <- TRUE
```

Save the admixed accession names.

```
> cat(arrInd[arrInd[,2] %in% trjaSubs,1], file = "lineIDsADM.txt", sep = " ")
```

Make the subsets.

```
> YmatIA     <- YmatAll[IAind,]
> matNAindIA <- matNAindALL[IAind,]
> colSums(matNAindIA)
```

```
[1] 32 36
```

```
> totNAindIA <- totNAindALL[IAind]
> lineFacIA  <- factor(lineFacAll[IAind])
> expMatIA   <- expMatAll[IAind,]
> colSums(expMatIA)

 2010_B  2008_3  2008_5 2008_15  2008_4  2010_A
    549     492     397     865     436     770
```

```
> YmatAUS     <- YmatAll[AUSind,]
> matNAindAUS <- matNAindALL[AUSind,]
> colSums(matNAindAUS)

[1] 12 14
```

```
> totNAindAUS <- totNAindALL[AUSind]
> lineFacAUS  <- factor(lineFacAll[AUSind])
> expMatAUS   <- expMatAll[AUSind,]
> colSums(expMatAUS)

 2010_B  2008_3  2008_5 2008_15  2008_4  2010_A
    208     185      91     328     246     339
```

```
> YmatJAP     <- YmatAll[JAPind,]
> matNAindJAP <- matNAindALL[JAPind,]
> colSums(matNAindJAP)

[1] 28 30
```

```
> totNAindJAP <- totNAindALL[JAPind]
> lineFacJAP  <- factor(lineFacAll[JAPind])
> expMatJAP   <- expMatAll[JAPind,]
> colSums(expMatJAP)

 2010_B  2008_3  2008_5 2008_15  2008_4  2010_A
    736     554     547     872     504     895
```

```
> YmatTRJ     <- YmatAll[TRJind,]
> matNAindTRJ <- matNAindALL[TRJind,]
> colSums(matNAindTRJ)

[1] 11 11
```

```
> totNAindTRJ <- totNAindALL[TRJind]
> lineFacTRJ  <- factor(lineFacAll[TRJind])
> expMatTRJ   <- expMatAll[TRJind,]
> colSums(expMatTRJ)

 2010_B  2008_3  2008_5 2008_15  2008_4  2010_A
    332     199     262     322     222     445


> YmatTRJA     <- YmatAll[TRJAind,]
> matNAindTRJA <- matNAindALL[TRJAind,]
> colSums(matNAindTRJA)

[1] 12 11


> totNAindTRJA <- totNAindALL[TRJAind]
> lineFacTRJA  <- factor(lineFacAll[TRJAind])
> expMatTRJA   <- expMatAll[TRJAind,]
> colSums(expMatTRJA)

 2010_B  2008_3  2008_5 2008_15  2008_4  2010_A
    344     199     273     328     222     472


>
```

Finally, I save all the data to binary files for modeling. I have to first replace `NA` values in the data matrix. The value is not important because it will be ignored by the software I write.

```
> trash <- .C("GSLvecSaveInt", "phenoData/trtBlk.gbin",
+     as.integer(blkIdx), length(blkIdx))
> YmatAll[is.na(YmatAll)]   <- 0.01
> YmatIA[is.na(YmatIA)]     <- 0.01
> YmatAUS[is.na(YmatAUS)]   <- 0.01
> YmatJAP[is.na(YmatJAP)]   <- 0.01
> YmatTRJ[is.na(YmatTRJ)]   <- 0.01
> YmatTRJA[is.na(YmatTRJA)] <- 0.01
> lineFacAll  <- factor(lineFacAll)
> lineFacIA   <- factor(lineFacIA)
> lineFacAUS  <- factor(lineFacAUS)
> lineFacJAP  <- factor(lineFacJAP)
> lineFacTRJ  <- factor(lineFacTRJ)
> lineFacTRJA <- factor(lineFacTRJA)
> trash <- .C("GSLmatSave", "phenoData/YatAll.gbin",
+     as.double(YmatAll), nrow(YmatAll), ncol(YmatAll))
> trash <- .C("GSLvecSaveInt", "phenoData/matNAindAll.gbin",
+     as.integer(t(matNAindALL)), as.integer(prod(dim(matNAindALL))))
```

```
> trash <- .C("GSLvecSaveInt", "phenoData/totNAindAll.gbin",
+     as.integer(totNAindALL), length(totNAindALL))
> trash <- .C("GSLvecSaveInt", "phenoData/lnIndAll.gbin",
+     as.integer(lineFacAll), length(lineFacAll))
> trash <- .C("GSLmatSave", "phenoData/expMatAll.gbin",
+     as.double(expMatAll), nrow(expMatAll), ncol(expMatAll))
> trash <- .C("GSLmatSave", "phenoData/YatIA.gbin",
+     as.double(YmatIA), nrow(YmatIA), ncol(YmatIA))
> trash <- .C("GSLvecSaveInt", "phenoData/matNAindIA.gbin",
+     as.integer(t(matNAindIA)), as.integer(prod(dim(matNAindIA))))
> trash <- .C("GSLvecSaveInt", "phenoData/totNAindIA.gbin",
+     as.integer(totNAindIA), length(totNAindIA))
> trash <- .C("GSLvecSaveInt", "phenoData/lnIndIA.gbin",
+     as.integer(lineFacIA), length(lineFacIA))
> trash <- .C("GSLmatSave", "phenoData/expMatIA.gbin",
+     as.double(expMatIA), nrow(expMatIA), ncol(expMatIA))
> trash <- .C("GSLmatSave", "phenoData/YatAUS.gbin",
+     as.double(YmatAUS), nrow(YmatAUS), ncol(YmatAUS))
> trash <- .C("GSLvecSaveInt", "phenoData/matNAindAUS.gbin",
+     as.integer(t(matNAindAUS)), as.integer(prod(dim(matNAindAUS))))
> trash <- .C("GSLvecSaveInt", "phenoData/totNAindAUS.gbin",
+     as.integer(totNAindAUS), length(totNAindAUS))
> trash <- .C("GSLvecSaveInt", "phenoData/lnIndAUS.gbin",
+     as.integer(lineFacAUS), length(lineFacAUS))
> trash <- .C("GSLmatSave", "phenoData/expMatAUS.gbin",
+     as.double(expMatAUS), nrow(expMatAUS), ncol(expMatAUS))
> trash <- .C("GSLmatSave", "phenoData/YatJAP.gbin",
+     as.double(YmatJAP), nrow(YmatJAP), ncol(YmatJAP))
> trash <- .C("GSLvecSaveInt", "phenoData/matNAindJAP.gbin",
+     as.integer(t(matNAindJAP)), as.integer(prod(dim(matNAindJAP))))
> trash <- .C("GSLvecSaveInt", "phenoData/totNAindJAP.gbin",
+     as.integer(totNAindJAP), length(totNAindJAP))
> trash <- .C("GSLvecSaveInt", "phenoData/lnIndJAP.gbin",
+     as.integer(lineFacJAP), length(lineFacJAP))
> trash <- .C("GSLmatSave", "phenoData/expMatJAP.gbin",
+     as.double(expMatJAP), nrow(expMatJAP), ncol(expMatJAP))
> trash <- .C("GSLmatSave", "phenoData/YatTRJ.gbin",
+     as.double(YmatTRJ), nrow(YmatTRJ), ncol(YmatTRJ))
> trash <- .C("GSLvecSaveInt", "phenoData/matNAindTRJ.gbin",
+     as.integer(t(matNAindTRJ)), as.integer(prod(dim(matNAindTRJ))))
> trash <- .C("GSLvecSaveInt", "phenoData/totNAindTRJ.gbin",
+     as.integer(totNAindTRJ), length(totNAindTRJ))
> trash <- .C("GSLvecSaveInt", "phenoData/lnIndTRJ.gbin",
+     as.integer(lineFacTRJ), length(lineFacTRJ))
> trash <- .C("GSLmatSave", "phenoData/expMatTRJ.gbin",
```

```
+     as.double(expMatTRJ), nrow(expMatTRJ), ncol(expMatTRJ))
> trash <- .C("GSLmatSave", "phenoData/YatTRJA.gbin",
+     as.double(YmatTRJA), nrow(YmatTRJA), ncol(YmatTRJA))
> trash <- .C("GSLvecSaveInt", "phenoData/matNAindTRJA.gbin",
+     as.integer(t(matNAindTRJA)), as.integer(prod(dim(matNAindTRJA))))
> trash <- .C("GSLvecSaveInt", "phenoData/totNAindTRJA.gbin",
+     as.integer(totNAindTRJA), length(totNAindTRJA))
> trash <- .C("GSLvecSaveInt", "phenoData/lnIndTRJA.gbin",
+     as.integer(lineFacTRJA), length(lineFacTRJA))
> trash <- .C("GSLmatSave", "phenoData/expMatTRJA.gbin",
+     as.double(expMatTRJA), nrow(expMatTRJA), ncol(expMatTRJA))
>
```

Finally, I save the accession IDs and print all dimensions for easy reference.

```
> cat(levels(lineFacIA),   file = "lineIDsIA.txt",   sep = " ")
> cat(levels(lineFacAUS),  file = "lineIDsAUS.txt",  sep = " ")
> cat(levels(lineFacJAP),  file = "lineIDsJAP.txt",  sep = " ")
> cat(levels(lineFacTRJ),  file = "lineIDsTRJ.txt",  sep = " ")
> cat(levels(lineFacTRJA), file = "lineIDsTRJA.txt", sep = " ")
> cat(unique(exprFacAll)[-1], file = "expIDs.txt", sep = " ")
> dim(YmatAll)
```

```
[1] 4428    2
```

```
> dim(YmatIA)
```

```
[1] 2009    2
```

```
> dim(YmatAUS)
```

```
[1] 794   2
```

```
> dim(YmatJAP)
```

```
[1] 2349    2
```

```
> dim(YmatTRJ)
```

```
[1] 1007    2
```

```
> dim(YmatTRJA)
```

```
[1] 1037    2
```

```
> dim(expMatAll)
[1] 8728    6

> dim(expMatIA)
[1] 3988    6

> dim(expMatAUS)
[1] 1568    6

> dim(expMatJAP)
[1] 4600    6

> dim(expMatTRJ)
[1] 1965    6

> dim(expMatTRJA)
[1] 2025    6

> nlevels(lineFacAll)
[1] 361

> nlevels(lineFacIA)
[1] 141

> nlevels(lineFacAUS)
[1] 55

> nlevels(lineFacJAP)
[1] 213

> nlevels(lineFacTRJ)
[1] 92

> nlevels(lineFacTRJA)
[1] 95
```
