## Supplemental methods and scripts for "Low additive genetic variation in a trait under selection in domesticated rice": runGWA.pdf

### Running GWA using a REML mixed model

Tony Greenberg

May 2, 2019

```
R version 3.5.3 (2019-03-11)
Platform: x86_64-pc-linux-gnu (64-bit)
Running under: Arch Linux

Matrix products: default
BLAS/LAPACK: /opt/intel/compilers_and_libraries_2019.1.144/linux/mkl/lib/intel64_lin/libmkl_gf_lp64.so

locale:
 [1] LC_CTYPE=en_US.UTF-8       LC_NUMERIC=C               LC_TIME=en_US.UTF-8
 [4] LC_COLLATE=en_US.UTF-8     LC_MONETARY=en_US.UTF-8    LC_MESSAGES=en_US.UTF-8
 [7] LC_PAPER=en_US.UTF-8       LC_NAME=C                  LC_ADDRESS=C
[10] LC_TELEPHONE=C             LC_MEASUREMENT=en_US.UTF-8 LC_IDENTIFICATION=C

attached base packages:
[1] compiler  stats     graphics  grDevices utils     datasets  methods   base

other attached packages:
[1] GWAlikeMeth_1.0 showtext_0.6    showtextdb_2.0  sysfonts_0.8    ggplot2_3.1.0

loaded via a namespace (and not attached):
 [1] Rcpp_1.0.0       crayon_1.3.4     withr_2.1.2      grid_3.5.3       plyr_1.8.4
 [6] gtable_0.2.0     scales_1.0.0     pillar_1.3.1     rlang_0.3.1      lazyeval_0.2.1
[11] tools_3.5.3      munsell_0.5.0    pkgconfig_2.0.2  colorspace_1.4-0 tibble_2.0.1
```

#### 1 Famoso *et al.* data

I start by replicating the Famoso *et al.* results with aus. Read in the accession means (from my Bayesian modeling) and the relationship matrix.

```
> accAUS        <- read.table(file="accModeAUS.tsv", header=T, sep = "\t")
> nsf2hdra      <- matrix(scan(file="NSFTV2HDRA.tsv", what=character()),
+                               ncol=2, byrow=T)
> # cumulative chromosome lengths
> cumChrLen     <- scan(file="riceCumChromLen.txt", what=integer())
> cumChrLen     <- c(0, cumChrLen[-length(cumChrLen)])
> hdra          <- nsf2hdra[,1]
> names(hdra)   <- nsf2hdra[,2]
```

```
> accAUS$HDRAID <- hdra[as.character(accAUS$NSFTVID)]
> Nacc          <- nlevels(accAUS$NSFTVID)
> d             <- 3
> nPer          <- 15
> trtNam        <- c("Control", "Treated", "logRRG")
> K             <- matrix(scan(file="matAUS.rel", what=double()),
+                        ncol=Nacc, byrow=T)
> Knam          <- matrix(scan(file="matAUS.rel.id", what=character()),
+                        ncol=2, byrow=T)[,1]
> rownames(K)   <- Knam
> colnames(K)   <- Knam
> K             <- K[accAUS$HDRAID,accAUS$HDRAID]
```

I next read the SNP matrix prepared using `plink-1.9`. Each locus is represented by two columns, so I collapse them to get diploid allele counts. Missing data are marked by "9." I convert that to `NA`.

```
> snpFl   <- pipe("cut -d ' ' -f7- ausSNPs.ped")
> ausSNPs <- matrix(scan(snpFl, what = integer()), nrow=Nacc, byrow=T)
> close(snpFl)
> Nsnp    <- ncol(ausSNPs)/2
> ausSNPs <- ausSNPs[,(1:Nsnp)*2 - 1] + ausSNPs[,(1:Nsnp)*2]
> ausSNPs[ausSNPs == 18] <- NA
> genoNames <- matrix(scan(file="ausSNPs.nosex", what=character()),
+                      ncol=2, byrow=T)[,1]
> genoNames <- genoNames[genoNames %in% rownames(K)]
> rownames(ausSNPs) <- genoNames
> ausSNPs   <- ausSNPs[rownames(K),]
```

The "0" are the minor alleles. For computational purposes it is better if they are major alleles. I switch the 0 and 2.

```
> ausSNPs[ausSNPs == 2]  <- -2
> ausSNPs[ausSNPs == 0]  <-  2
> ausSNPs[ausSNPs == -2] <-  0
```

Now run the GWA.

```
> ausGWA <- gwa(Y=accAUS[,2:4], K=K, snps=ausSNPs, nPerm=nPer)
> ausGWA$hSq

[1] 4.799851e-01 7.124925e-02 5.374918e-09
```

Create a data frame of $-\log_{10}(p)$ for plotting. Start by importing genome positions and chromosome ID.

```
> posFl      <- pipe("cut -f1,4 ausSNPs.map")
> chrPos     <- matrix(scan(posFl, what=integer()), ncol=2, byrow=T)
```

```
> close(posFl)
> chrPos[,2] <- chrPos[,2] + cumChrLen[chrPos[,1]]
> lPvalFrame <- data.frame(lPval=array(ausGWA$lPval),
+                         trait=factor(rep(trtNam, each=Nsnp), levels=trtNam),
+                         chr=rep(chrPos[,1], d),
+                         pos=rep(chrPos[,2], d))
```

I calculate FDR cut-offs for each trait using the permutation $q$-values.

```
> trtFDR <- cmpfun(function(gwaNam, cutOff){
+     res   <- rep(NA, d)
+     qVmat <- eval(as.symbol(gwaNam))$qVal
+     nCut  <- apply(qVmat, 2, function(vec){sum(vec <= cutOff)})
+     if(sum(nCut == 0) == d) return(res)
+     non0  <- sapply(which(nCut != 0),
+         function(i, qm){min(eval(as.symbol(gwaNam))$lPval[qm[,i] <= cutOff,i])}, qVmat)
+     res[which(nCut != 0)] <- non0
+     return(res)
+ })
> fdrCut   <- 0.1
> lpCutVal <- trtFDR("ausGWA", fdrCut)
> lpCutVal
```

```
[1] 5.440368       NA 4.538376
```

Make a data frame for adding these values to plots.

```
> fdrFrame <- data.frame(trait=factor(trtNam), lp=lpCutVal)
```

I add indexes to color the plots. I want the chromosomes to switch between two shades of gray and "significant" ($FDR \leq 0.1$) points to be red.

```
> colIdx <- rep("C2", nrow(lPvalFrame))
> colIdx[as.logical(lPvalFrame$chr %% 2)] <- "C1"
> if(sum(!is.na(lpCutVal))){
+     lPvalFrame$colIdx <- ifelse(lPvalFrame$lPval >=
+                             rep(lpCutVal, each=Nsnp), "C3", colIdx)
+     lPvalFrame$colIdx[is.na(lPvalFrame$colIdx)] <- colIdx[is.na(lPvalFrame$colIdx)]
+ } else {
+     lPvalFrame$colIdx <- colIdx
+ }
> lPvalFrame$colIdx <- factor(lPvalFrame$colIdx)
```

There are too many points to plot. I thin out the uninformative SNPs ($FDR <= 0.1$).

```
> keepInd <- NULL
> if(sum(!is.na(lpCutVal))){
+     keepInd <- sort(c(which(lPvalFrame$lPval > min(lpCutVal, na.rm=T)),
```

```
+                 sample(which(lPvalFrame$lPval <= min(lpCutVal, na.rm=T)), d*5000,
+                     prob=lPvalFrame$lPval[lPvalFrame$lPval <= min(lpCutVal, na.rm=T)])))
+ } else {
+     keepInd <- sort(sample(1:length(lPvalFrame$lPval), d*5000,
+                         prob=lPvalFrame$lPval))
+ }
> lPvalFrame <- lPvalFrame[keepInd,]

> chrPos    <- tapply(lPvalFrame$pos, lPvalFrame$chr, median)
> nratFrame <- data.frame(trait=factor(trtNam),
+                         pos=c(NA, NA, 1660000+cumChrLen[2]),
+                         post=c(NA, NA, 25000000+cumChrLen[2]),
+                         lPval=c(NA, NA, fdrFrame$lp[3]+0.5),
+                         lab=c(NA, NA, "Nrat1"),
+                         col=rep("C4", 3))
> pdfFlNam  <- "ausGWAfamoso.pdf"
> showtext_auto()
> ggplot(data=lPvalFrame, aes(x=pos,y=lPval, color=colIdx)) +
+     scale_color_manual(values=c("C1"="grey40", "C2"="grey60", "C3"="red", "C4"="black")) +
+     geom_point(show.legend=F) +
+     facet_wrap(~trait, scales="free_y", nrow=d) +
+     scale_x_continuous(name="chromosomes", breaks=chrPos,
+                     labels=unique(lPvalFrame$chr)) +
+     theme_classic(base_size=18, base_family="myriad") +
+     theme(axis.ticks.x=element_blank(), axis.line.x=element_blank(),
+         strip.background=element_rect(fill="grey95", linetype="blank")) +
+     labs(y=expression(-log[10](p))) +
+     geom_vline(data=nratFrame, aes(xintercept=pos), color="grey85") +
+     geom_text(data=nratFrame, mapping=aes(x=post, y=lPval, color=col),
+                 label="Nrat1", size=6, show.legend=F, fontface="bold") +
+     if(sum(!is.na(lpCutVal))){geom_hline(data=fdrFrame, aes(yintercept=lp),
+                                     color="grey60")}
> ggsave(pdfFlNam, width=8, height=10, units="in", device="pdf", useDingbats=F)
> cat("\\includegraphics{", pdfFlNam, "}\n\n", sep="")
```

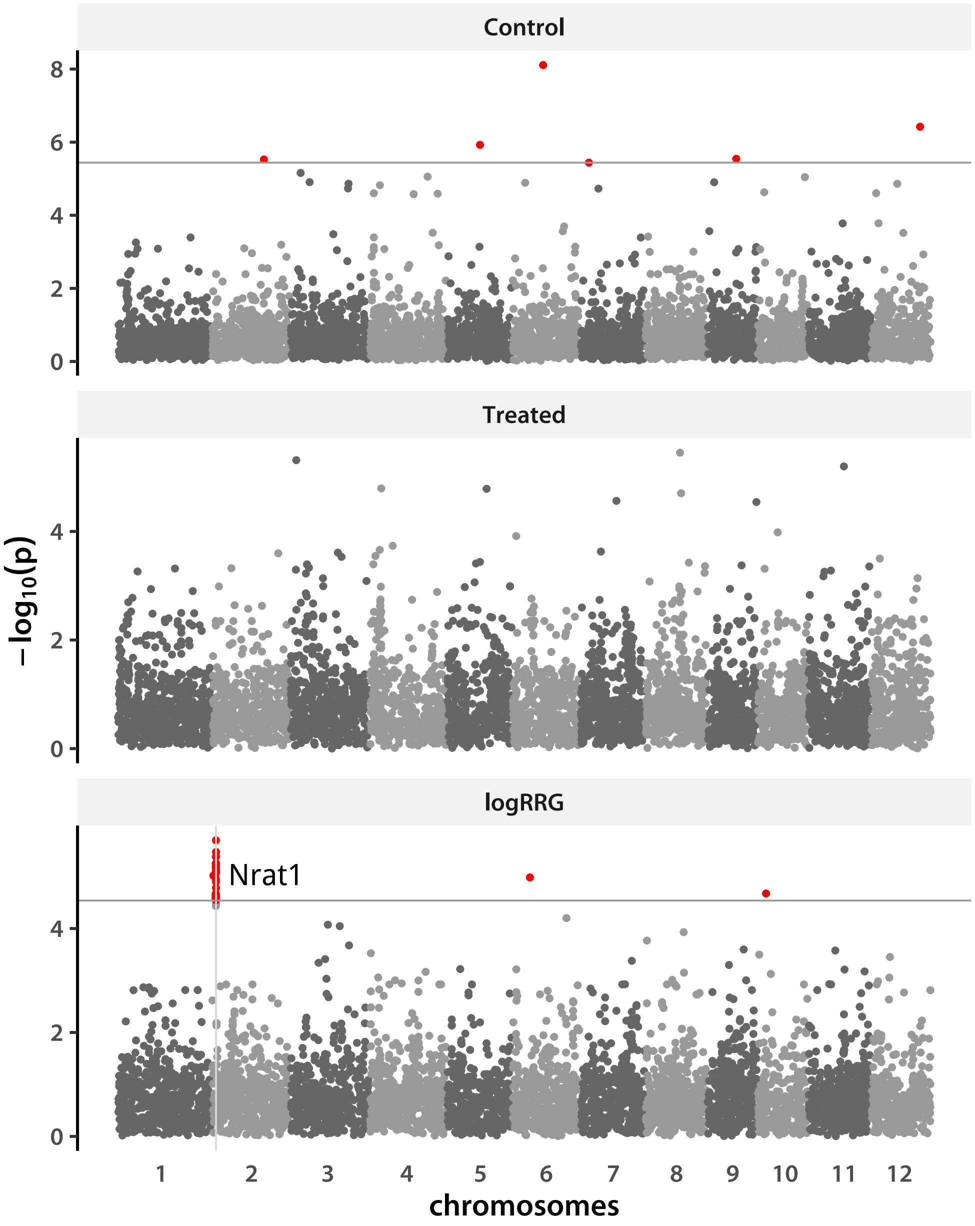


```
>
```

Now repeat the whole thing with tropical *japonica* data.

```
> accTRJ         <- read.table(file="accModeTRJ.tsv", header=T, sep = "\t")
> accTRJ$HDRAID <- hdra[as.character(accTRJ$NSFTVID)]
> Nacc           <- nlevels(accTRJ$NSFTVID)
> K              <- matrix(scan(file="matTRJ.rel", what=double()),
+                          ncol=Nacc, byrow=T)
> Knam           <- matrix(scan(file="matTRJ.rel.id", what=character()),
+                          ncol=2, byrow=T)[,1]
```

```
> rownames(K)   <- Knam
> colnames(K)   <- Knam
> K             <- K[accTRJ$HDRAID,accTRJ$HDRAID]
```

I next read the SNP matrix prepared using `plink-1.9`. Each locus is represented by two columns, so I collapse them to get diploid allele counts. Missing data are marked by "9." I convert that to `NA`.

```
> snpFl   <- pipe("cut -d ' ' -f7- trjSNPs.ped")
> trjSNPs <- matrix(scan(snpFl, what = integer()), nrow=Nacc, byrow=T)
> close(snpFl)
> Nsnp    <- ncol(trjSNPs)/2
> trjSNPs <- trjSNPs[,(1:Nsnp)*2 - 1] + trjSNPs[,(1:Nsnp)*2]
> trjSNPs[trjSNPs == 18] <- NA
> genoNames <- matrix(scan(file="trjSNPs.nosex", what=character()), ncol=2, byrow=T)[,1]
> genoNames <- genoNames[genoNames %in% rownames(K)]
> rownames(trjSNPs) <- genoNames
> trjSNPs   <- trjSNPs[rownames(K),]
```

The "0" are the minor alleles. For computational purposes it is better if they are major alleles. I switch the 0 and 2.

```
> trjSNPs[trjSNPs == 2]  <- -2
> trjSNPs[trjSNPs == 0]  <-  2
> trjSNPs[trjSNPs == -2] <-  0
```

Now run the GWA.

```
> trjGWA <- gwa(Y=accTRJ[,2:4], K=K, snps=trjSNPs, nPerm=nPer)
> trjGWA$hSq
```

```
[1] 0.7254627 0.6134491 0.1774834
```

Create a data frame of $-\log_{10}(p)$ for plotting. Start by importing genome positions and chromosome ID.

```
> posFl      <- pipe("cut -f1,4 trjSNPs.map")
> chrPos     <- matrix(scan(posFl, what=integer()), ncol=2, byrow=T)
> close(posFl)
> chrPos[,2] <- chrPos[,2] + cumChrLen[chrPos[,1]]
> lPvalFrame <- data.frame(lPval=array(trjGWA$lPval),
+                          trait=factor(rep(trtNam, each=Nsnp), levels=trtNam),
+                          chr=rep(chrPos[,1], d),
+                          pos=rep(chrPos[,2], d))
```

I calculate FDR cut-offs for each trait.

```
> lpCutVal <- trtFDR("trjGWA", fdrCut)
> lpCutVal
```

```
[1] NA NA NA
```

Make a data frame for adding these values to plots.

```
> fdrFrame <- data.frame(trait=factor(trtNam), lp=lpCutVal)
```

I add indexes to color the plots. I want the chromosomes to switch between two shades of gray and "significant" ($FDR \leq 0.1$) points to be red.

```
> colIdx <- rep("C2", nrow(lPvalFrame))
> colIdx[as.logical(lPvalFrame$chr %% 2)] <- "C1"
> if(sum(!is.na(lpCutVal))){
+     lPvalFrame$colIdx <- ifelse(lPvalFrame$lPval >=
+                                 rep(lpCutVal, each=Nsnp), "C3", colIdx)
+     lPvalFrame$colIdx[is.na(lPvalFrame$colIdx)] <- colIdx[is.na(lPvalFrame$colIdx)]
+ } else {
+     lPvalFrame$colIdx <- colIdx
+ }
> lPvalFrame$colIdx <- factor(lPvalFrame$colIdx)
```

There are too many points to plot. I thin out the uninformative SNPs.

```
> keepInd <- NULL
> if(sum(!is.na(lpCutVal))){
+     keepInd <- sort(c(which(lPvalFrame$lPval > min(lpCutVal, na.rm=T)),
+                 sample(which(lPvalFrame$lPval <= min(lpCutVal, na.rm=T)), d*5000,
+                     prob=lPvalFrame$lPval[lPvalFrame$lPval <= min(lpCutVal, na.rm=T)])))
+ } else {
+     keepInd <- sort(sample(1:length(lPvalFrame$lPval), d*5000,
+                         prob=lPvalFrame$lPval))
+ }
> lPvalFrame <- lPvalFrame[keepInd,]

> chrPos   <- tapply(lPvalFrame$pos, lPvalFrame$chr, median)
> pdfFlNam <- "trjGWAfamoso.pdf"
> showtext_auto()
> ggplot(data=lPvalFrame, aes(x=pos,y=lPval, color=colIdx)) +
+     scale_color_manual(values=c("C1"="grey40", "C2"="grey60", "C3"="red")) +
+     geom_point(show.legend=F) +
+     facet_wrap(~trait, scales="free_y", nrow=d) +
+     scale_x_continuous(name="chromosomes", breaks=chrPos,
+                     labels=unique(lPvalFrame$chr)) +
+     theme_classic(base_size=18, base_family="myriad") +
+     theme(axis.ticks.x=element_blank(), axis.line.x=element_blank(),
+         strip.background=element_rect(fill="grey95", linetype="blank")) +
+     labs(y=expression(-log[10](p))) +
+     if(sum(!is.na(lpCutVal))){geom_hline(data=fdrFrame,
```

```
+                                         aes(yintercept=lp), color="grey60")}
> ggsave(pdfFlNam, width=8, height=10, units="in", device="pdf", useDingbats=F)
> cat("\\includegraphics{", pdfFlNam, "}\n\n", sep="")
```

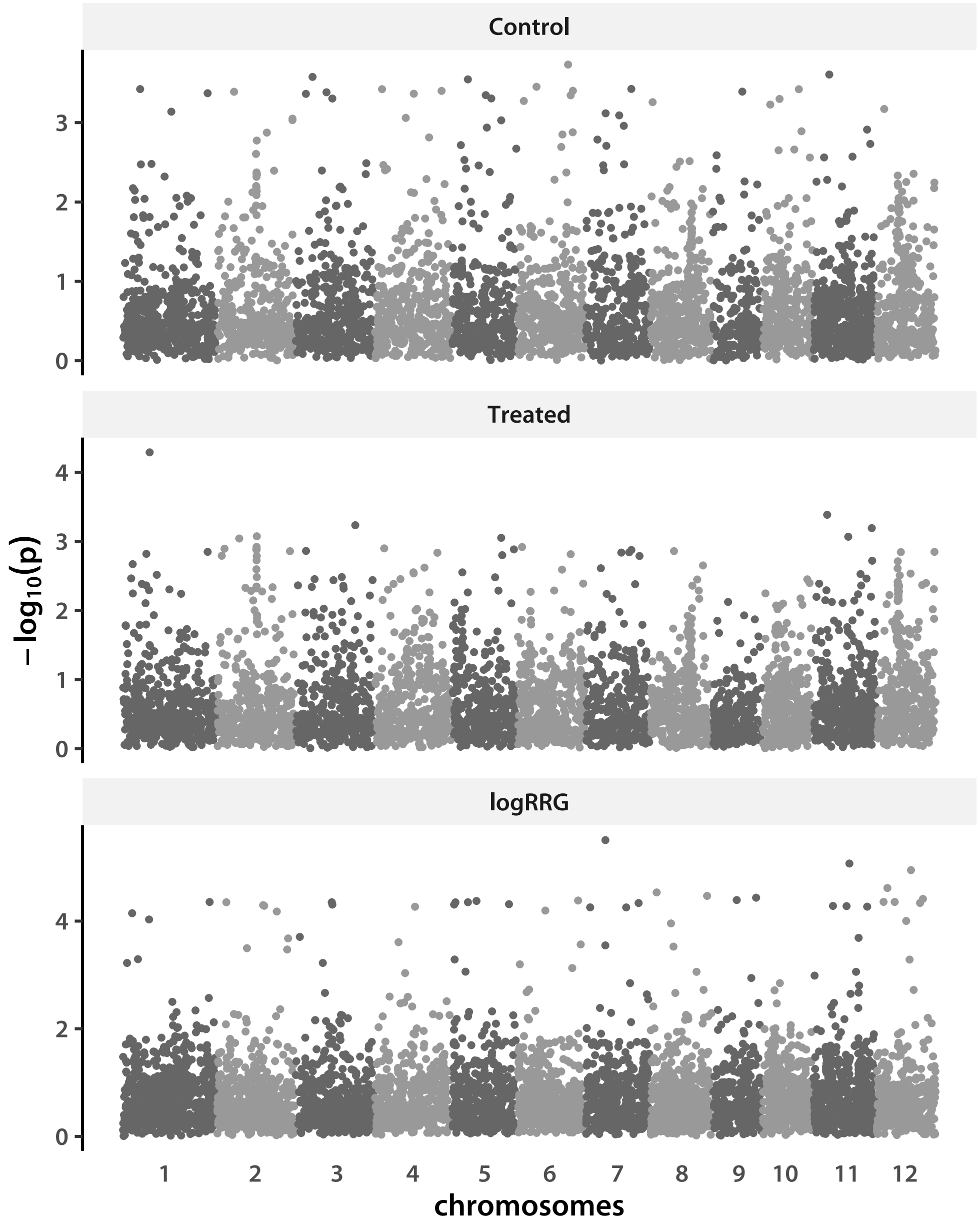


```
>
```

#### 2 Additional TRJ accessions

First remove previous SNP matrices and GWA objects to free memory.

```
> rm(list=c("ausSNPs", "trjSNPs", "ausGWA", "trjGWA"))
```

Read accession modes and establish constants.

```
> accADtrj    <- read.table(file="accModeADtrj.tsv", header=T, sep = "\t")
> Nacc        <- nlevels(accADtrj$acc_ID)
> d           <- 3
> K           <- matrix(scan(file="addDataTRJ.rel", what=double()),
+                       ncol=Nacc, byrow=T)
> Knam        <- matrix(scan(file="addDataTRJ.rel.id", what=character()),
+                       ncol=2, byrow=T)[,1]
> rownames(K) <- Knam
> colnames(K) <- Knam
> K           <- K[as.character(accADtrj$acc_ID), as.character(accADtrj$acc_ID)]
```

Read the SNP data.

```
> snpFl   <- pipe("cut -d ' ' -f7- addDataTRJ.ped")
> addSNPs <- matrix(scan(snpFl, what = integer()), nrow=Nacc, byrow=T)
> close(snpFl)
> Nsnp    <- ncol(addSNPs)/2
> addSNPs <- addSNPs[,(1:Nsnp)*2 - 1] + addSNPs[,(1:Nsnp)*2]
> addSNPs[addSNPs == 18] <- NA
> genoNames <- matrix(scan(file="addDataTRJ.nosex", what=character()),
+                       ncol=2, byrow=T)[,1]
> genoNames <- genoNames[genoNames %in% rownames(K)]
> rownames(addSNPs) <- genoNames
> addSNPs   <- addSNPs[rownames(K),]
```

Now run the GWA.

```
> addGWA <- gwa(Y=accADtrj[,2:4], K=K, snps=addSNPs, nPerm=nPer)
> addGWA$hSq
```

```
[1] 0.42531138 0.35863032 0.04608683
```

Create a data frame of $-\log_{10}(p)$ for plotting. Start by importing genome positions and chromosome ID.

```
> posFl      <- pipe("cut -f1,4 addDataTRJ.map")
> chrPos     <- matrix(scan(posFl, what=integer()), ncol=2, byrow=T)
> close(posFl)
> chrPos[,2] <- chrPos[,2] + cumChrLen[chrPos[,1]]
> lPvalFrame <- data.frame(lPval=array(addGWA$lPval),
+                          trait=factor(rep(trtNam, each=Nsnp), levels=trtNam),
+                          chr=rep(chrPos[,1], d),
+                          pos=rep(chrPos[,2], d))
```

I calculate FDR cut-offs for each trait.

```
> lpCutVal <- trtFDR("addGWA", fdrCut)
> lpCutVal
```

```
[1]       NA       NA 6.326049
```

Make a data frame for adding these values to plots.

```
> fdrFrame <- data.frame(trait=factor(trtNam), lp=lpCutVal)
```

I add indexes to color the plots. I want the chromosomes to switch between two shades of gray and "significant" ($FDR \leq 0.1$) points to be red.

```
> colIdx <- rep("C2", nrow(lPvalFrame))
> colIdx[as.logical(lPvalFrame$chr %% 2)] <- "C1"
> if(sum(!is.na(lpCutVal))){
+     lPvalFrame$colIdx <- ifelse(lPvalFrame$lPval >= rep(lpCutVal, each=Nsnp),
+                                 "C3", colIdx)
+     lPvalFrame$colIdx[is.na(lPvalFrame$colIdx)] <- colIdx[is.na(lPvalFrame$colIdx)]
+ } else {
+     lPvalFrame$colIdx <- colIdx
+ }
> lPvalFrame$colIdx <- factor(lPvalFrame$colIdx)
```

There are too many points to plot. I thin out the uninformative SNPs.

```
> keepInd <- NULL
> if(sum(!is.na(lpCutVal))){
+     keepInd <- sort(c(which(lPvalFrame$lPval > min(lpCutVal, na.rm=T)),
+                 sample(which(lPvalFrame$lPval <= min(lpCutVal, na.rm=T)), d*5000,
+                     prob=lPvalFrame$lPval[lPvalFrame$lPval <= min(lpCutVal, na.rm=T)])))
+ } else {
+     keepInd <- sort(sample(1:length(lPvalFrame$lPval), d*5000,
+                         prob=lPvalFrame$lPval))
+ }
> lPvalFrame <- lPvalFrame[keepInd,]
```

```
> chrPos   <- tapply(lPvalFrame$pos, lPvalFrame$chr, median)
> pdfFlNam <- "addGWAtrj.pdf"
> showtext_auto()
> ggplot(data=lPvalFrame, aes(x=pos,y=lPval, color=colIdx)) +
+     scale_color_manual(values=c("C1"="grey40", "C2"="grey60", "C3"="red")) +
+     geom_point(show.legend=F) +
+     facet_wrap(~trait, scales="free_y", nrow=d) +
+     scale_x_continuous(name="chromosomes", breaks=chrPos,
+                     labels=unique(lPvalFrame$chr)) +
+     theme_classic(base_size=18, base_family="myriad") +
+     theme(axis.ticks.x=element_blank(), axis.line.x=element_blank(),
+         strip.background=element_rect(fill="grey95", linetype="blank")) +
```

```
+     labs(y=expression(-log[10](p))) +
+     if(sum(!is.na(lpCutVal))){geom_hline(data=fdrFrame,
+                                    aes(yintercept=lp), color="grey60")}
> ggsave(pdfFlNam, width=8, height=10, units="in", device="pdf", useDingbats=F)
> cat("\\includegraphics{", pdfFlNam, "}\n\n", sep="")
```

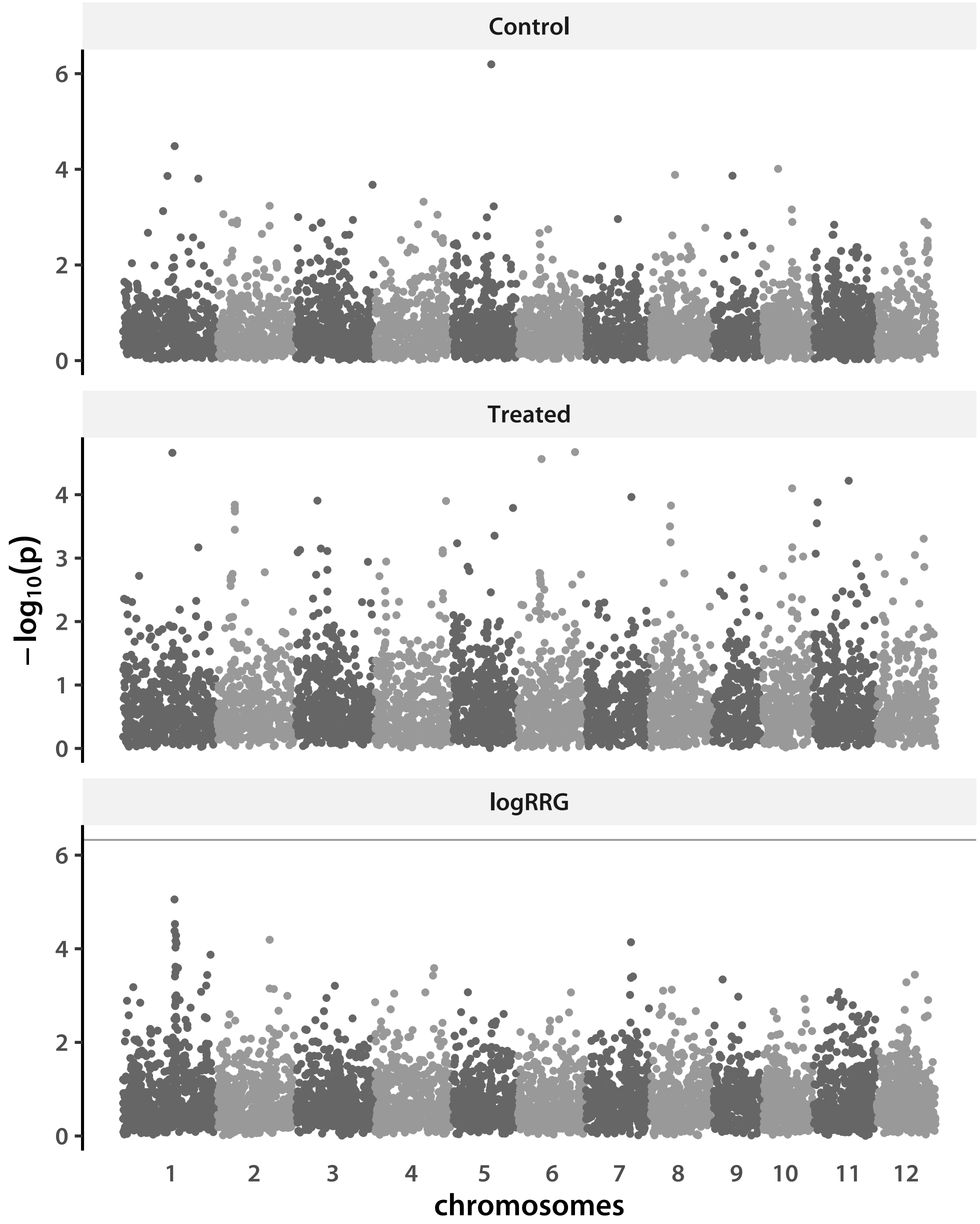


```
>
```

### 3 New phenotype data

Finally, I run GWA on the new phenotype data. I first read the phenotype file and set constants.

```
> accNDtrj     <- read.table(file="accModeNDtrj.tsv", header=T, sep = "\t")
> Nacc         <- nlevels(accNDtrj$HDRA_ID)
> d            <- 6
> trtNam       <- c("Longest Root Length (Treated)", "Total Root Length (Treated)",
+                    "Longest Root Length (Control)", "Total Root Length (Control)",
+                    "Longest Root Length (logRRG)", "Total Root Length (logRRG)")
> K            <- matrix(scan(file="newData.rel", what=double()),
+                        ncol=Nacc, byrow=T)
> Knam         <- matrix(scan(file="newData.rel.id", what=character()),
+                            ncol=2, byrow=T)[,1]
> rownames(K) <- Knam
> colnames(K) <- Knam
> K            <- K[as.character(accNDtrj$HDRA_ID), as.character(accNDtrj$HDRA_ID)]
```

Read the SNP data.

```
> snpFl   <- pipe("cut -d ' ' -f7- newData.ped")
> newSNPs <- matrix(scan(snpFl, what = integer()), nrow=Nacc, byrow=T)
> close(snpFl)
> Nsnp    <- ncol(newSNPs)/2
> newSNPs <- newSNPs[,(1:Nsnp)*2 - 1] + newSNPs[,(1:Nsnp)*2]
> newSNPs[newSNPs == 18] <- NA
> genoNames <- matrix(scan(file="newData.nosex", what=character()),
+                    ncol=2, byrow=T)[,1]
> genoNames <- genoNames[genoNames %in% rownames(K)]
> rownames(newSNPs) <- genoNames
> newSNPs   <- newSNPs[rownames(K),]
```

Now run the GWA.

```
> newGWA <- gwa(Y=accNDtrj[,2:7], K=K, snps=newSNPs, nPerm=nPer)
> newGWA$hSq

[1] 5.922939e-02 2.134305e-01 4.509077e-02 3.860885e-01 3.321888e-09 2.053023e-09
```

Create a data frame of $-\log_{10}(p)$ for plotting. Start by importing genome positions and chromosome ID.

```
> posFl      <- pipe("cut -f1,4 newData.map")
> chrPos     <- matrix(scan(posFl, what=integer()), ncol=2, byrow=T)
> close(posFl)
> chrPos[,2] <- chrPos[,2] + cumChrLen[chrPos[,1]]
> lPvalFrame <- data.frame(lPval=array(newGWA$lPval),
+                          trait=factor(rep(trtNam, each=Nsnp), levels=trtNam),
+                          chr=rep(chrPos[,1], d),
+                          pos=rep(chrPos[,2], d))
```

I calculate FDR cut-offs for each trait.

```
> lpCutVal <- trtFDR("newGWA", fdrCut)
> lpCutVal
```

```
[1] NA NA NA NA NA NA
```

Make a data frame for adding these values to plots.

```
> fdrFrame <- data.frame(trait=factor(trtNam), lp=lpCutVal)
```

I add indexes to color the plots. I want the chromosomes to switch between two shades of gray and "significant" ($FDR \leq 0.1$) points to be red.

```
> colIdx <- rep("C2", nrow(lPvalFrame))
> colIdx[as.logical(lPvalFrame$chr %% 2)] <- "C1"
> if(sum(!is.na(lpCutVal))){
+     lPvalFrame$colIdx <- ifelse(lPvalFrame$lPval >= rep(lpCutVal, each=Nsnp),
+                                 "C3", colIdx)
+     lPvalFrame$colIdx[is.na(lPvalFrame$colIdx)] <- colIdx[is.na(lPvalFrame$colIdx)]
+ } else {
+     lPvalFrame$colIdx <- colIdx
+ }
> lPvalFrame$colIdx <- factor(lPvalFrame$colIdx)
```

There are too many points to plot. I thin out the uninformative SNPs.

```
> keepInd <- NULL
> if(sum(!is.na(lpCutVal))){
+     keepInd <- sort(c(which(lPvalFrame$lPval > min(lpCutVal, na.rm=T)),
+                 sample(which(lPvalFrame$lPval <= min(lpCutVal, na.rm=T)), d*5000,
+                     prob=lPvalFrame$lPval[lPvalFrame$lPval <= min(lpCutVal, na.rm=T)])))
+ } else {
+     keepInd <- sort(sample(1:length(lPvalFrame$lPval), d*5000,
+                         prob=lPvalFrame$lPval))
+ }
> lPvalFrame <- lPvalFrame[keepInd,]
```

```
> chrPos   <- tapply(lPvalFrame$pos, lPvalFrame$chr, median)
> pdfFlNam <- "newGWAtrj.pdf"
> showtext_auto()
> ggplot(data=lPvalFrame, aes(x=pos,y=lPval, color=colIdx)) +
+     scale_color_manual(values=c("C1"="grey40", "C2"="grey60", "C3"="red")) +
+     geom_point(show.legend=F) +
+     facet_wrap(~trait, scales="free_y", ncol=2) +
+     scale_x_continuous(name="chromosomes", breaks=chrPos,
+                     labels=unique(lPvalFrame$chr)) +
+     theme_classic(base_size=18, base_family="myriad") +
```

```
+     theme(axis.ticks.x=element_blank(), axis.line.x=element_blank(),
+         strip.background=element_rect(fill="grey95", linetype="blank")) +
+     labs(y=expression(-log[10](p))) +
+     if(sum(!is.na(lpCutVal))){geom_hline(data=fdrFrame,
+                                     aes(yintercept=lp), color="grey60")}
> ggsave(pdfFlNam, width=12, height=10, units="in", device="pdf", useDingbats=F)
> cat("\\includegraphics{", pdfFlNam, "}\n\n", sep="")
```

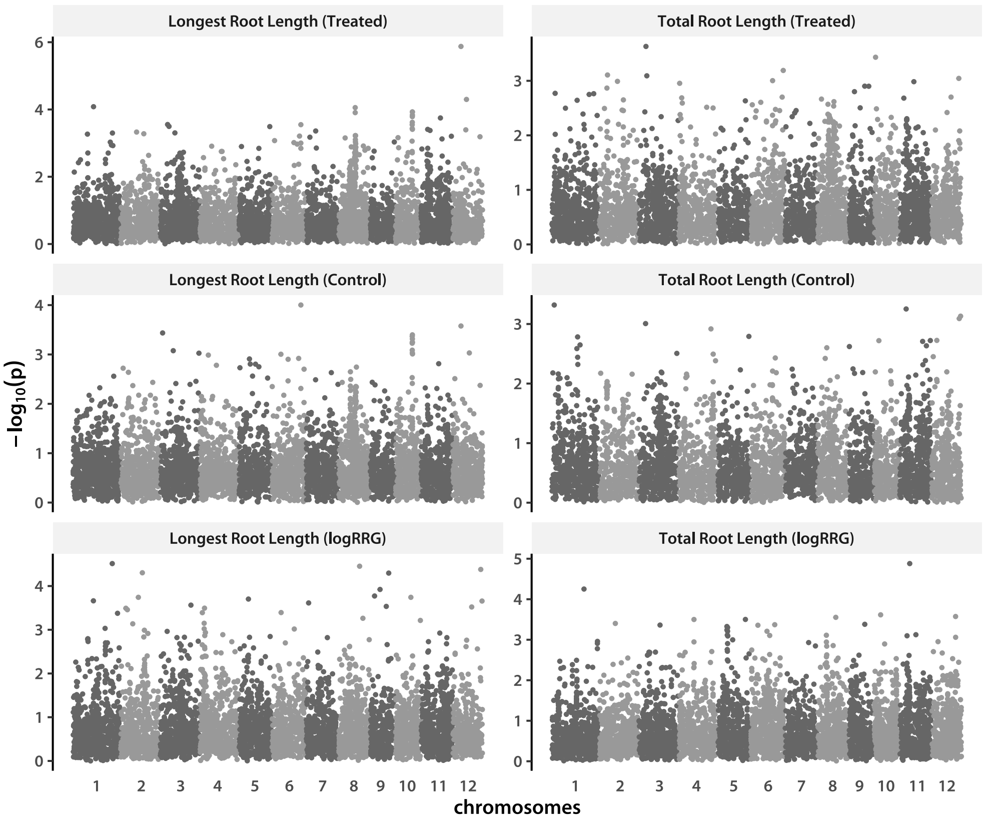


```
>
```
