## Supplemental methods and scripts for "Low additive genetic variation in a trait under selection in domesticated rice": dataPrepADtrj.Rnw

```
%@
\documentclass[fleqn,11pt]{article}
\usepackage{amsmath}
\usepackage{amssymb}
\usepackage{sectsty}
\usepackage{amsbsy}
\usepackage{fancyhdr}
\usepackage{color}
\usepackage{url}
\usepackage[indention=2em,labelsep=period,labelfont=bf]{caption}
\usepackage[normalsize,nooneline,sf,bf,FIGTOPCAP,tight]{subfigure}
\usepackage[noae]{Sweave} % noae required to sanitize single quotes
\usepackage{fancyvrb}

\textwidth = 6.5 in
\textheight = 8.5 in
\oddsidemargin = 0.0 in
\evensidemargin = 0.0 in
\topmargin = 0.0 in
%\voffset = - 0.4 in
\headheight = 15 pt
\headsep = 0.1 in
\parskip = 0.0 in
\parindent = 4 ex

\definecolor{grey}{rgb}{0.4,0.4,0.4}

\newcommand{\bcdot}{\boldsymbol{\cdot}}

\DefineVerbatimEnvironment{Sinput}{Verbatim}{tabsize=4}
\DefineVerbatimEnvironment{Scode}{Verbatim}{tabsize=4}
\DefineVerbatimEnvironment{Soutput}{Verbatim}{tabsize=4,formatcom=\color{grey},fontsize=\small}

\pagestyle{fancy}
\fancyhf{}
\rhead{Page \thepage}
\lhead{Data Prep}
\chead{Tony Greenberg}

\title{Preparing TRJ Al tolerance data for analysis}
\author{Tony Greenberg}

\begin{document}

\maketitle
\thispagestyle{empty}
<<echo=false>>=
rm(list = ls())
options(width=90)
version[c(1,13)]
@
In this document I prepare the new data set that includes additional measurements on the tropical \textit{japonica} accessions. Start by reading the phenotype data. The file has been modified from the original Excel spreadsheet to add ``.0'' to two accession names that did not have it.

\section{Phenotype data}
<<>>=
rawData <- read.table(file = "rawData.tsv", sep = "\t", header = T)
summary(rawData)
@
There are a few accessions that do not have IDs. These are not genotyped, so I am eliminating them.
<<>>=
dim(rawData)
rawData <- rawData[!is.na(rawData$assay_id),]
dim(rawData)
@
The control individuals are marked by ``0uM'' in the \verb|Original_sample_ID| column and the treated plants are ``160uM.'' I am making them into two separate traits. Here is the function that does that within each accession.
<<results=hide>>=
buildY <- cmpfun(function(accNam){
	accMat <- rawData[rawData$assay_id == accNam,]
	ctrlID <- grep("_0uM_", as.character(accMat$Original_sample_ID))
	ctrl   <- accMat$day_3_TRL[ctrlID]
	cExp   <- as.character(accMat$Experiment)[ctrlID]

	trtID  <- grep("_160uM_", as.character(accMat$Original_sample_ID))
	trt    <- accMat$day_3_TRL[trtID]
	tExp   <- as.character(accMat$Experiment)[trtID]
	res    <- matrix(NA, nrow = max(length(trt), length(ctrl)), ncol = 2)
	res[1:length(ctrl),1] <- ctrl
	res[1:length(trt),2]  <- trt

	acc  <- rep(accNam, nrow(res))
	if(length(cExp) >= length(tExp)) {
		expr <- cExp
	} else {
		expr <- tExp
	}

	Ymat  <<- rbind(Ymat, res)
	accID <<- c(accID, acc)
	expID <<- c(expID, expr)
	return(NULL)
})
@

Now run the function to build the trait matrix and the accession and experiment ID factors.

<<>>=
Ymat  <- NULL
accID <- NULL
expID <- NULL
trash <- sapply(levels(rawData$assay_id), buildY)
colnames(Ymat) <- c("CTRL", "TRT")
dim(Ymat)
length(unique(accID))
range(apply(Ymat, 1, function(vec){sum(is.na(vec))}))
@
Prepare the missing data matrix and save the phenotype and meta-data.
<<>>=
matNA <- apply(Ymat, 2, function(vec){ifelse(is.na(vec), 1, 0)})
colSums(matNA)
totNA <- rowSums(matNA)
trash   <- .C("GSLvecSaveInt", "phenoData/matNAind.gbin",
	as.integer(t(matNA)), as.integer(prod(dim(matNA))))
trash <- .C("GSLvecSaveInt", "phenoData/totNAind.gbin",
	as.integer(totNA), length(totNA))
Ymat[is.na(Ymat)] <- 0.0
trash <- .C("GSLmatSave", "phenoData/yMat.gbin",
	as.double(Ymat), nrow(Ymat), ncol(Ymat))
accID <- factor(accID)
trash <- .C("GSLvecSaveInt", "phenoData/accInd.gbin",
	as.integer(accID), length(accID))
accIDmat <- matrix(levels(accID), nrow = nlevels(accID), ncol = 2)
cat(apply(accIDmat, 1, paste, collapse = "\t"), file = "accID.tsv", sep = "\n")
@
Save the trait-block index for errors. In our case each of the two ``traits'' are in their own block so that the errors are not correlated (the data for treated and untreated come from different individuals).
<<>>=
blkIdx <- 1:2
trash  <- .C("GSLvecSaveInt", "phenoData/trtBlk.gbin",
	as.integer(blkIdx), length(blkIdx))
@

I will use a regression on contrasts to account for the experiment effects. To pick which level to set as the base, I plot effect distributions.
\begin{center}
<<fig=TRUE,width=8.0,height=6.0,echo=FALSE>>=
par(mfrow = c(1,2), mar = c(6.7, 3.0, 1.0, 0.1))
trash <- sapply(colnames(Ymat), function(val){boxplot(Ymat[,val]~expID, main = val, las = 2, pch = -0x00b7, col = "grey80", cex.axis = 0.9)})
@
\end{center}
\verb|2016-December| seems to be close to the overall mean, so I will use that. Furthermore, differences within a year are minor compared to among-year divergence. Trying to model each experiment resulted in shrinkage to zero and no correction. I will just estimate year effects, contrasting to 2016/2017 (those were done by the same person using the same reagents and there is no evident difference in mean).

<<>>=
nYr    <- length(unique(sapply(strsplit(expID, "-"), function(vec){vec[1]}))) - 2
Xexp   <- matrix(0, nrow = nrow(Ymat), ncol = nYr)
Xexp[grep("2008-", expID),1] <- 1
Xexp[grep("2010-", expID),2] <- 1
dim(Xexp)
trash <- .C("GSLmatSave", "phenoData/Xexp.gbin",
	as.double(Xexp), nrow(Xexp), ncol(Xexp))
@
I also want to construct an experiment predictor that corrects at the level of accessions (i.e. after accession mean estimation). I first test how many accessions were measured in more than one year.
<<>>=
multiExpAcc <- levels(accID)[
	sapply(
		tapply(
			sapply(strsplit(expID, "-"), function(vec){vec[1]}),
			accID, function(vec){unique(vec)}),
		length) > 1]
multiExpAcc
@
There are two accessions. I will keep only the data for the 2016/2017 for these.
<<>>=
dropInd <- which((rowSums(Xexp) == 1) & (accID %in% multiExpAcc))
YmatS   <- Ymat[-dropInd,]
accIDs  <- accID[-dropInd]
matNAs  <- matNA[-dropInd,]
totNAs  <- totNA[-dropInd]
trash   <- .C("GSLvecSaveInt", "phenoData/matNAindS.gbin",
	as.integer(t(matNAs)), as.integer(prod(dim(matNAs))))
trash <- .C("GSLvecSaveInt", "phenoData/totNAindS.gbin",
	as.integer(totNAs), length(totNAs))
trash <- .C("GSLmatSave", "phenoData/yMatS.gbin",
	as.double(YmatS), nrow(YmatS), ncol(YmatS))
trash <- .C("GSLvecSaveInt", "phenoData/accIndS.gbin",
	as.integer(accIDs), length(accIDs))
@
I determine the year ID for each accession and build the contrast matrix. I will include the intercept.
<<>>=
yrID <- tapply(sapply(strsplit(expID[-dropInd], "-"), function(vec){vec[1]}),
			accIDs, function(vec){unique(vec)[1]})
Xyr <- cbind(rep(1, nlevels(accIDs)),
	sapply(c("2008", "2010"), function(yr){yrID == yr}))
trash <- .C("GSLmatSave", "phenoData/Xyr.gbin",
	as.double(Xyr), nrow(Xyr), ncol(Xyr))
@

\section{Genotype data}
Next I prepare genotype data: the SNP table and the relationship matrix eigenvectors and -values. I first run \verb|plink| on the HDRA SNP master file to extract the accessions I need and trim out missing data and singleton SNPs. To do that, the MAF cut-off is $2/N_{acc} = 0.012$.
<<>>=
if(!file.exists("genoData/snpHDRAaltlTRJ.gbin")){
	plinkComm <- paste(
			"plink --noweb --silent",
			"--tfile HDRA-G6-4-RDP1-RDP2-sativa-only",
			"--keep accID.tsv --maf 0.012 --geno 0.3",
			"--recode12 --transpose --out alTolTRJ"
		)
	system(plinkComm)
}
@
Next I read in the genotype file and format the data.
<<>>=
if(!file.exists("genoData/snpHDRAaltlTRJ.gbin")){
	dimPipe <- pipe("wc -l alTolTRJ.tped")
	nSNP  <- as.integer(scan(dimPipe, what = character())[1])
	close(dimPipe)
	snpFl <- pipe("cut -d ' ' -f5- alTolTRJ.tped")
	SNPtab <- matrix(scan(snpFl, what = integer()), nrow = nSNP, byrow = T)
	close(snpFl)
}
@
Extract SNP ID, chromosome position and chromosome number from the \verb|.tped| file. This information is in the first four columns. I re-arrange the columns so that I have SNP ID then chromosome number then position. This is done only if the SNP data have not been saved yet.
<<>>=
if(!file.exists("genoData/snpHDRAaltlTRJ.gbin")){
	snpFl <- pipe("cut -d ' ' -f-4 alTolTRJ.tped")
	snpID <- matrix(scan(snpFl, what = character()), nrow = nSNP, byrow = T)
	close(snpFl)
	system("rm alTolTRJ.*")
	snpID <- snpID[,c(2, 1, 4)]
	cat(
		apply(rbind(c("SNP_ID", "CHROM", "POS"), snpID),
			1, paste, collapse = "\t"),
		file = "genoData/snpID.tsv", sep = "\n"
		)
	SNPtab[SNPtab == 0] <- NA
	SNPtab <- SNPtab - 1
	dim(SNPtab)
}
@

Convert the \verb|.tped| two rows per individual to one row per individual.
<<>>=
if(!file.exists("genoData/snpHDRAaltlTRJ.gbin")){
	SNPtab <- (SNPtab[,seq(from = 1, to = ncol(SNPtab) - 1, by = 2)] +
		SNPtab[,seq(from = 2, to = ncol(SNPtab), by = 2)])
	SNPtab <- t(SNPtab)
	dim(SNPtab)
}
@
Calculate the relationship matrix and its eigendecomposition unless that has been done before.
<<>>=
if(!file.exists("genoData/altlTRJevc.gbin") | !file.exists("genoData/altlTRJevl.gbin")){
	library(rrBLUP)
	SNPtab <- SNPtab - 1
	K      <- A.mat(SNPtab, impute.method="EM")
	Keig   <- eigen(K, symmetric = T)
	Keig$values

	eVal  <- Keig$val[Keig$val > 1e-8]
	eVec  <- Keig$vec[,Keig$val > 1e-8]
	trash <- .C("GSLvecSave", "genoData/altlTRJevl.gbin",
		as.double(eVal), length(eVal))
	trash <- .C("GSLmatSave", "genoData/altlTRJevc.gbin",
		as.double(eVec), nrow(eVec), ncol(eVec))

	SNPtab[is.na(SNPtab)] <- -9
	trash <- .C("GSLmatSave", "genoData/snpHDRAaltlTRJ.gbin",
		as.double(SNPtab), nrow(SNPtab), ncol(SNPtab))
	dim(SNPtab)
	dim(eVec)
}
@
Print dimensions for future reference.
<<>>=
dim(Ymat)
dim(YmatS)
nlevels(accID)
dim(Xexp)
dim(Xyr)
@

\end{document}
%<<>>=
```
