## Supplemental methods and scripts for "Low additive genetic variation in a trait under selection in domesticated rice": mcmcExamineADtrj.Rnw

```
\documentclass[fleqn,11pt]{article}
\usepackage{amsmath}
\usepackage{amssymb}
\usepackage{sectsty}
\usepackage{amsbsy}
\usepackage{fancyhdr}
\usepackage{color}
\usepackage{url}
\usepackage[indention=2em,labelsep=period,labelfont=bf]{caption}
\usepackage[normalsize,nooneline,sf,bf,FIGTOPCAP,tight]{subfigure}
\usepackage[noae]{Sweave} % noae required to sanitize single quotes
\usepackage{fancyvrb}

\textwidth = 6.5 in
\textheight = 8.5 in
\oddsidemargin = 0.0 in
\evensidemargin = 0.0 in
\topmargin = 0.0 in
%\voffset = - 0.4 in
\headheight = 15 pt
\headsep = 0.1 in
\parskip = 0.0 in
\parindent = 4 ex

\definecolor{grey}{rgb}{0.4,0.4,0.4}

\newcommand{\bcdot}{\boldsymbol{\cdot}}

\DefineVerbatimEnvironment{Sinput}{Verbatim}{tabsize=4}
\DefineVerbatimEnvironment{Scode}{Verbatim}{tabsize=4}
\DefineVerbatimEnvironment{Soutput}{Verbatim}{tabsize=4,formatcom=\color{grey},fontsize=\small}

\pagestyle{fancy}
\fancyhf{}
\rhead{Page \thepage}
\lhead{MCMC diagnostics}
\chead{Tony Greenberg}

\title{Examining MCMC chains for convergence and mixing}
\author{Tony Greenberg}

\begin{document}

\maketitle
\thispagestyle{empty}
<<echo=false>>=
dyn.load("GSLRIO.so")
options(width=90)
library(compiler)
sessionInfo()
@
Examining the Gibbs sampler chains generated for the Al tolerance experiment on TRJ. Starting with the bivariate model that generates accession means for further analyses relative root growth. 

Start by defining some useful functions.

I first define a function that calculates the Gelman-Rubin convergence statistic for multiple MCMC chains.
<<>>=
gelRub <- cmpfun(function(vrNam, nChn, chnLen){
	mat <- NULL
	for (i in 1:nChn){
		mat <- rbind(mat, eval(as.symbol(paste(vrNam, i, sep=""))))
	}
	chnFac <- factor(rep(1:nChn, each=chnLen))
	W <- colMeans(apply(mat, 2, tapply, chnFac, var))
	B <- apply(apply(mat, 2, tapply, chnFac, mean), 2, var)
	return( sqrt((((chnLen - 1)/chnLen)*W + B)/W) )
})

@
I am also interested in the quality of chain mixing. I use autocorrelations to test for that. The following function examines autocorrelations for each chain for a given variable and reports the chain with the highest mean value. The autocorrelations are calculated using default parameters of the \texttt{acf()} function from the \texttt{stats} R package.
<<>>=
eachAcf <- cmpfun(function(i, nChn, vrNam){
	tmp <- NULL
	for (j in 1:nChn){
		tmp <- cbind(tmp, 
				acf(eval(as.symbol(paste(vrNam, j, sep="")))[,i], plot = F)$acf
			)
	}
	return(tmp[,which.max(colMeans(tmp[-1,]))])
})

mcmcAcf <- cmpfun(function(vrNam, nChn){
	N <- ncol(eval(as.symbol(paste(vrNam, 1, sep="")))) # all chains should have the same dimensions, so take the first one
	t(sapply(1:N, eachAcf, nChn, vrNam))
})
@

\section{Raw data modeling}

The first task is to model raw data and extract accession means. I ran a 5000 iteration burn-in and 10 000 samples with, keeping every fifth value. I first look at the experiment effect.

<<>>=
trtNam <- c("CTRL", "TRT")
d      <- 2
Nyr    <- 3
yrDim  <- d*Nyr
nChn   <- 5
chnLen <- 2000

chn1 <- matrix(.C("GSLmatLoad", "chains/YR_3_1.gbin",
	as.integer(chnLen), as.integer(yrDim), out = double(chnLen*yrDim))$out,
		nrow = chnLen, byrow = T)
chn2 <- matrix(.C("GSLmatLoad", "chains/YR_3_2.gbin", 
	as.integer(chnLen), as.integer(yrDim), out = double(chnLen*yrDim))$out, 
		nrow = chnLen, byrow = T)
chn3 <- matrix(.C("GSLmatLoad", "chains/YR_3_3.gbin", 
	as.integer(chnLen), as.integer(yrDim), out = double(chnLen*yrDim))$out, 
		nrow = chnLen, byrow = T)
chn4 <- matrix(.C("GSLmatLoad", "chains/YR_3_4.gbin", 
	as.integer(chnLen), as.integer(yrDim), out = double(chnLen*yrDim))$out, 
		nrow = chnLen, byrow = T)
chn5 <- matrix(.C("GSLmatLoad", "chains/YR_3_5.gbin", 
	as.integer(chnLen), as.integer(yrDim), out = double(chnLen*yrDim))$out, 
		nrow = chnLen, byrow = T)

grMat    <- matrix(gelRub("chn",nChn,chnLen), ncol = d, byrow = T)
grMaxInd <- apply(grMat, 2, function(vec){which(vec == max(vec))[1]})
acMat    <- mcmcAcf("chn", 5)

@

Now I plot the Gelman-Rubin for each trait.

\begin{center}
<<fig=TRUE,width=8.0,height=6.0,echo=FALSE>>=
par(mar = c(5.2, 6, 2, 1), mgp = c(4,1,0))
plot(c(1,1,1,2,2,2), grMat, pch = -0x00b7, cex = 5, las = 1, ylab = "Gelman-Rubin R", xaxt = "n", xlab = "", main = "Year Effect")
axis(1, at = 1:d, trtNam, las = 1)
@
\end{center}
Values smaller than $\sim$1.2 indicate good convergence. I check by plotting an the chains with the worst G-R statistic for each trait.

<<>>=
plotLocEx <- cmpfun(function(trt.i, samp.j, N){
	i <- ( d*(samp.j-1) ) + trt.i
	plot( chn1[,i], type = "l", 
		ylim = range(c(chn1[,i], chn2[,i], chn3[,i], chn4[,i], chn5[,i])), 
		col = "orange", ylab = trtNam[trt.i], xaxt = "n", las = 1)
	lines(chn2[,i], col = "red")
	lines(chn3[,i], col = "blue")
	lines(chn4[,i], col = "green")
	lines(chn5[,i], col = "magenta")
})
@

\begin{center}
<<fig=TRUE,width=8.0,height=6.0,echo=FALSE>>=
par(mfrow = c(1,2), mar = c(0.2, 4, 0.5, 1), mgp = c(3,1,0))
for (trt in 1:d){
	plotLocEx(trt, grMaxInd[trt], Nyr)
}

@
\end{center}

Now I look at mixing. Lag is on the $x$ axis.
\begin{center}
<<fig=TRUE,width=8.0,height=6.0,useDingbats=F,echo=FALSE>>=
par(mfrow = c(1,2), mar = c(0.2, 4, 0.5, 1), mgp = c(3,1,0))
for (trt in 1:d){
	boxplot(acMat[(0:(Nyr-1)*d + trt),], pch = -0x00b7, outcex = 2, las = 1, ylab = trtNam[trt], col = "grey80", xaxt = "n")
}
@
\end{center}

Next I look at accession means.
<<>>=
Nacc   <- 169
accDim <- d*Nacc

chn1 <- matrix(.C("GSLmatLoad", "chains/LN_3_1.gbin", 
	as.integer(chnLen), as.integer(accDim), out = double(chnLen*accDim))$out, 
		nrow = chnLen, byrow = T)
chn2 <- matrix(.C("GSLmatLoad", "chains/LN_3_2.gbin", 
	as.integer(chnLen), as.integer(accDim), out = double(chnLen*accDim))$out, 
		nrow = chnLen, byrow = T)
chn3 <- matrix(.C("GSLmatLoad", "chains/LN_3_3.gbin", 
	as.integer(chnLen), as.integer(accDim), out = double(chnLen*accDim))$out, 
		nrow = chnLen, byrow = T)
chn4 <- matrix(.C("GSLmatLoad", "chains/LN_3_4.gbin", 
	as.integer(chnLen), as.integer(accDim), out = double(chnLen*accDim))$out, 
		nrow = chnLen, byrow = T)
chn5 <- matrix(.C("GSLmatLoad", "chains/LN_3_5.gbin", 
	as.integer(chnLen), as.integer(accDim), out = double(chnLen*accDim))$out, 
		nrow = chnLen, byrow = T)

grMat    <- matrix(gelRub("chn",nChn,chnLen), ncol = d, byrow = T)
grMaxInd <- apply(grMat, 2, function(vec){which(vec == max(vec))[1]})
acMat    <- mcmcAcf("chn", 5)

@

Now I plot the Gelman-Rubin for each trait.

\begin{center}
<<fig=TRUE,width=8.0,height=6.0,echo=FALSE>>=
par(mar = c(5.2, 6, 2, 1), mgp = c(4,1,0))
boxplot(grMat, pch = -0x00b7, outcex = 2, las = 1, ylab = "Gelman-Rubin R", xaxt = "n", col = "grey80", main = "Accession means")
axis(1, at = 1:d, trtNam, las = 1)
@
\end{center}
Values smaller than $\sim$1.2 indicate good convergence. I check by plotting an the chains with the worst G-R statistic for each trait.

\begin{center}
<<fig=TRUE,width=8.0,height=6.0,echo=FALSE>>=
par(mfrow = c(1,2), mar = c(0.2, 4, 0.5, 1), mgp = c(3,1,0))
for (trt in 1:d){
	plotLocEx(trt, grMaxInd[trt], Nacc)
}

@
\end{center}

Now I look at mixing. Lag is on the $x$ axis.
\begin{center}
<<fig=TRUE,width=8.0,height=6.0,useDingbats=F,echo=FALSE>>=
par(mfrow = c(1,2), mar = c(0.2, 4, 0.5, 1), mgp = c(3,1,0))
for (trt in 1:d){
	boxplot(acMat[(0:(Nacc-1)*d + trt),], pch = -0x00b7, outcex = 2, las = 1, ylab = trtNam[trt], col = "grey80", xaxt = "n")
}
@
\end{center}

Next I examine the GEBV chains.

<<>>=
chn1 <- matrix(.C("GSLmatLoad", "chains/BV_3_1.gbin", 
	as.integer(chnLen), as.integer(accDim), out = double(chnLen*accDim))$out, 
		nrow = chnLen, byrow = T)
chn2 <- matrix(.C("GSLmatLoad", "chains/BV_3_2.gbin", 
	as.integer(chnLen), as.integer(accDim), out = double(chnLen*accDim))$out, 
		nrow = chnLen, byrow = T)
chn3 <- matrix(.C("GSLmatLoad", "chains/BV_3_3.gbin", 
	as.integer(chnLen), as.integer(accDim), out = double(chnLen*accDim))$out, 
		nrow = chnLen, byrow = T)
chn4 <- matrix(.C("GSLmatLoad", "chains/BV_3_4.gbin", 
	as.integer(chnLen), as.integer(accDim), out = double(chnLen*accDim))$out, 
		nrow = chnLen, byrow = T)
chn5 <- matrix(.C("GSLmatLoad", "chains/BV_3_5.gbin", 
	as.integer(chnLen), as.integer(accDim), out = double(chnLen*accDim))$out, 
		nrow = chnLen, byrow = T)

grMat    <- matrix(gelRub("chn",nChn,chnLen), ncol = d, byrow = T)
grMaxInd <- apply(grMat, 2, function(vec){which(vec == max(vec))[1]})
acMat    <- mcmcAcf("chn", 5)
@

\begin{center}
<<fig=TRUE,width=8.0,height=6.0,echo=FALSE>>=
par(mar = c(5.2, 6, 2, 1), mgp = c(4,1,0))
boxplot(grMat, pch = -0x00b7, outcex = 2, las = 1, ylab = "Gelman-Rubin R", xaxt = "n", col = "grey80", main = "GEBV")
axis(1, at = 1:d, trtNam, las = 1)
@
\end{center}
Plot the worst chains.

\begin{center}
<<fig=TRUE,width=8.0,height=6.0,echo=FALSE>>=
par(mfrow = c(1,2), mar = c(0.2, 4, 0.5, 1), mgp = c(3,1,0))
for (trt in 1:d){
	plotLocEx(trt, grMaxInd[trt], Nacc)
}

@
\end{center}

Now I look at mixing.
\begin{center}
<<fig=TRUE,width=8.0,height=6.0,useDingbats=F,echo=FALSE>>=
par(mfrow = c(1,2), mar = c(0.2, 4, 0.5, 1), mgp = c(3,1,0))
for (trt in 1:d){
	boxplot(acMat[(0:(Nacc-1)*d + trt),], pch = -0x00b7, outcex = 2, las = 1, ylab = trtNam[trt], col = "grey80", xaxt = "n")
}
@
\end{center}

Next I look at covariance matrices. I create an index to isolate just the upper triangles.

<<>>=
upInd   <- matrix(1:(d^2), ncol = d, byrow = T)
upInd   <- upInd[row(upInd) <= col(upInd)]
diagInd <- diag(matrix(1:(d^2), ncol = d, byrow = T))
@
Read in covariance matrix chains. Error correlations are set to zero, I exclude them.

<<>>=
chn1 <- matrix(.C("GSLmatLoad", 
	"chains/SgE_3_1.gbin", 
	as.integer(chnLen), as.integer(d^2), out = double(chnLen*d^2))$out, 
		nrow = chnLen, byrow = T)[,diagInd]
chn1 <- cbind(chn1, 
	matrix(.C("GSLmatLoad", 
	"chains/SgS_3_1.gbin", 
	as.integer(chnLen), as.integer(d^2), out = double(chnLen*d^2))$out, 
		nrow = chnLen, byrow = T)[,upInd])
chn1 <- cbind(chn1, 
	matrix(.C("GSLmatLoad", 
	"chains/SgA_3_1.gbin", 
	as.integer(chnLen), as.integer(d^2), out = double(chnLen*d^2))$out, 
		nrow = chnLen, byrow = T)[,upInd])

chn2 <- matrix(.C("GSLmatLoad", 
	"chains/SgE_3_2.gbin", 
	as.integer(chnLen), as.integer(d^2), out = double(chnLen*d^2))$out, 
		nrow = chnLen, byrow = T)[,diagInd]
chn2 <- cbind(chn2, 
	matrix(.C("GSLmatLoad", 
	"chains/SgS_3_2.gbin", 
	as.integer(chnLen), as.integer(d^2), out = double(chnLen*d^2))$out, 
		nrow = chnLen, byrow = T)[,upInd])
chn2 <- cbind(chn2, 
	matrix(.C("GSLmatLoad", 
	"chains/SgA_3_2.gbin", 
	as.integer(chnLen), as.integer(d^2), out = double(chnLen*d^2))$out, 
		nrow = chnLen, byrow = T)[,upInd])

chn3 <- matrix(.C("GSLmatLoad", 
	"chains/SgE_3_3.gbin", 
	as.integer(chnLen), as.integer(d^2), out = double(chnLen*d^2))$out, 
		nrow = chnLen, byrow = T)[,diagInd]
chn3 <- cbind(chn3, 
	matrix(.C("GSLmatLoad", 
	"chains/SgS_3_3.gbin", 
	as.integer(chnLen), as.integer(d^2), out = double(chnLen*d^2))$out, 
		nrow = chnLen, byrow = T)[,upInd])
chn3 <- cbind(chn3, 
	matrix(.C("GSLmatLoad", 
	"chains/SgA_3_3.gbin", 
	as.integer(chnLen), as.integer(d^2), out = double(chnLen*d^2))$out, 
		nrow = chnLen, byrow = T)[,upInd])

chn4 <- matrix(.C("GSLmatLoad", 
	"chains/SgE_3_4.gbin", 
	as.integer(chnLen), as.integer(d^2), out = double(chnLen*d^2))$out, 
		nrow = chnLen, byrow = T)[,diagInd]
chn4 <- cbind(chn4, 
	matrix(.C("GSLmatLoad", 
	"chains/SgS_3_4.gbin", 
	as.integer(chnLen), as.integer(d^2), out = double(chnLen*d^2))$out, 
		nrow = chnLen, byrow = T)[,upInd])
chn4 <- cbind(chn4, 
	matrix(.C("GSLmatLoad", 
	"chains/SgA_3_4.gbin", 
	as.integer(chnLen), as.integer(d^2), out = double(chnLen*d^2))$out, 
		nrow = chnLen, byrow = T)[,upInd])

chn5 <- matrix(.C("GSLmatLoad", 
	"chains/SgE_3_5.gbin", 
	as.integer(chnLen), as.integer(d^2), out = double(chnLen*d^2))$out, 
		nrow = chnLen, byrow = T)[,diagInd]
chn5 <- cbind(chn5, 
	matrix(.C("GSLmatLoad", 
	"chains/SgS_3_5.gbin", 
	as.integer(chnLen), as.integer(d^2), out = double(chnLen*d^2))$out, 
		nrow = chnLen, byrow = T)[,upInd])
chn5 <- cbind(chn5, 
	matrix(.C("GSLmatLoad", 
	"chains/SgA_3_5.gbin", 
	as.integer(chnLen), as.integer(d^2), out = double(chnLen*d^2))$out, 
		nrow = chnLen, byrow = T)[,upInd])
acMat <- mcmcAcf("chn", 5)

@
I have to make an index that marks each covariance matrix elements.

<<>>=
cvrID <- c(rep("e", length(diagInd)), rep(c("s", "a"), each = length(upInd)))
cvrInd <- 1:ncol(chn1)
@

Now plot the diagnostics.

\begin{center}
<<fig=TRUE,width=8.0,height=6.0,echo=FALSE>>=
par(mar = c(5.2, 6, 2, 1), mgp = c(4,1,0))
plot(gelRub("chn",nChn,chnLen), pch = -0x00b7, cex = 3, las = 1, ylab = "Gelman-Rubin R", xaxt = "n", main = "Covariance matrices")
axis(1, at = c(1.5, 4, 7), c(expression(Sigma[e]), expression(Sigma[s]), expression(Sigma[a])), las = 1)
@
\end{center}
Now I look at mixing.
\begin{center}
<<fig=TRUE,width=8.0,height=8.0,useDingbats=F,echo=FALSE>>=
par(mfrow = c(3,1), mar = c(0.2, 4, 0.5, 1), mgp = c(3,1,0))
for (sgID in 1:3){
	boxplot(acMat[cvrInd[cvrID == c("e", "s", "a")[sgID]],], pch = -0x00b7, outcex = 2, las = 1, ylab = c(expression(Sigma[e]), expression(Sigma[s]), expression(Sigma[a]))[sgID], col = "grey80", xaxt = "n")
}
@
\end{center}

I plot the MCMC chains of variances (diagonal elements) for each covariance matrix. First, I get the diagonal index from the upper triangle index.
<<>>=
diagInd <- list(1:2, c(3,5), c(6,8))
@

First, I plot the error variances.

\begin{center}
<<fig=TRUE,width=8.0,height=6.0,echo=FALSE>>=

par(mfrow = c(2,1), mar = c(0.2, 4, 0.5, 1), mgp = c(3,1,0))
for (trt in 1:d){
	plot( chn1[,diagInd[[1]][trt]], type = "l", ylim = range(c(chn1[,diagInd[[1]][trt]], chn2[,diagInd[[1]][trt]], chn3[,diagInd[[1]][trt]], chn4[,diagInd[[1]][trt]], chn5[,diagInd[[1]][trt]])), col = "orange", ylab = trtNam[trt], xlab = "", xaxt = "n", las = 1)
	lines(chn2[,diagInd[[1]][trt]], col = "red")
	lines(chn3[,diagInd[[1]][trt]], col = "blue")
	lines(chn4[,diagInd[[1]][trt]], col = "green")
	lines(chn5[,diagInd[[1]][trt]], col = "magenta")
}
@
\end{center}

Next, non-additive genetic variances.

\begin{center}
<<fig=TRUE,width=8.0,height=6.0,echo=FALSE>>=

par(mfrow = c(2,1), mar = c(0.2, 4, 0.5, 1), mgp = c(3,1,0))
for (trt in 1:d){
	plot( chn1[,diagInd[[2]][trt]], type = "l", ylim = range(c(chn1[,diagInd[[2]][trt]], chn2[,diagInd[[2]][trt]], chn3[,diagInd[[2]][trt]], chn4[,diagInd[[2]][trt]], chn5[,diagInd[[2]][trt]])), col = "orange", ylab = trtNam[trt], xlab = "", xaxt = "n", las = 1)
	lines(chn2[,diagInd[[2]][trt]], col = "red")
	lines(chn3[,diagInd[[2]][trt]], col = "blue")
	lines(chn4[,diagInd[[2]][trt]], col = "green")
	lines(chn5[,diagInd[[2]][trt]], col = "magenta")
}
@
\end{center}
Finally, $\boldsymbol{\Sigma}_a$ which can be interpreted as the additive genetic covariance matrix in this case, where $\nu_g =1000$ and the model is essentially Gaussian.

\begin{center}
<<fig=TRUE,width=8.0,height=6.0,echo=FALSE>>=

par(mfrow = c(2,1), mar = c(0.2, 4, 0.5, 1), mgp = c(3,1,0))
for (trt in 1:d){
	plot( chn1[,diagInd[[3]][trt]], type = "l", ylim = range(c(chn1[,diagInd[[3]][trt]], chn2[,diagInd[[3]][trt]], chn3[,diagInd[[3]][trt]], chn4[,diagInd[[3]][trt]], chn5[,diagInd[[3]][trt]])), col = "orange", ylab = trtNam[trt], xlab = "", xaxt = "n", las = 1)
	lines(chn2[,diagInd[[3]][trt]], col = "red")
	lines(chn3[,diagInd[[3]][trt]], col = "blue")
	lines(chn4[,diagInd[[3]][trt]], col = "green")
	lines(chn5[,diagInd[[3]][trt]], col = "magenta")
}
@
\end{center}

It is OK to proceed with summarizing the MCMC output for further analyses.

\section{Population structure control for GWA}

Having extracted accession means from the model above, I collected three traits (the original two plus $\log_{10}\mathrm{RRG}$) and ran a model for genotypes with the accession means as responce. Here I check the MCMC output of this model.

I start with the genetic PC regression coefficients.
<<>>=
trtNam <- c("CTRL", "TRT", "RRG")
d      <- 3
Npc    <- 168
pcDim  <- d*Npc

chn1 <- matrix(.C("GSLmatLoad", "chainsRRG/GM_3_1.gbin", 
	as.integer(chnLen), as.integer(pcDim), out = double(chnLen*pcDim))$out, 
		nrow = chnLen, byrow = T)
chn2 <- matrix(.C("GSLmatLoad", "chainsRRG/GM_3_2.gbin", 
	as.integer(chnLen), as.integer(pcDim), out = double(chnLen*pcDim))$out, 
		nrow = chnLen, byrow = T)
chn3 <- matrix(.C("GSLmatLoad", "chainsRRG/GM_3_3.gbin", 
	as.integer(chnLen), as.integer(pcDim), out = double(chnLen*pcDim))$out, 
		nrow = chnLen, byrow = T)
chn4 <- matrix(.C("GSLmatLoad", "chainsRRG/GM_3_4.gbin", 
	as.integer(chnLen), as.integer(pcDim), out = double(chnLen*pcDim))$out, 
		nrow = chnLen, byrow = T)
chn5 <- matrix(.C("GSLmatLoad", "chainsRRG/GM_3_5.gbin", 
	as.integer(chnLen), as.integer(pcDim), out = double(chnLen*pcDim))$out, 
		nrow = chnLen, byrow = T)

grMat    <- matrix(gelRub("chn",nChn,chnLen), ncol = d, byrow = T)
grMaxInd <- apply(grMat, 2, function(vec){which(vec == max(vec))[1]})
acMat    <- mcmcAcf("chn", 5)

@
Plot the results.

\begin{center}
<<fig=TRUE,width=8.0,height=6.0,echo=FALSE>>=
par(mar = c(5.2, 6, 2, 1), mgp = c(4,1,0))
boxplot(grMat, pch = -0x00b7, outcex = 2, las = 1, ylab = "Gelman-Rubin R", xaxt = "n", col = "grey80", main = "PC coefficients")
axis(1, at = 1:d, trtNam, las = 1)
@
\end{center}
Plot the worst chains.

\begin{center}
<<fig=TRUE,width=8.0,height=8.0,echo=FALSE>>=
par(mfrow = c(2,2), mar = c(0.2, 4, 0.5, 1), mgp = c(3,1,0))
for (trt in 1:d){
	plotLocEx(trt, grMaxInd[trt], Npc)
}

@
\end{center}

Now I look at mixing.
\begin{center}
<<fig=TRUE,width=8.0,height=8.0,useDingbats=F,echo=FALSE>>=
par(mfrow = c(2,2), mar = c(0.2, 4, 0.5, 1), mgp = c(3,1,0))
for (trt in 1:d){
	boxplot(acMat[(0:(Npc-1)*d + trt),], pch = -0x00b7, outcex = 2, las = 1, ylab = trtNam[trt], col = "grey80", xaxt = "n")
}
@
\end{center}

Next, look at GEBVs.

<<>>=
accDim <- d*Nacc
chn1 <- matrix(.C("GSLmatLoad", "chainsRRG/BV_3_1.gbin", 
	as.integer(chnLen), as.integer(accDim), out = double(chnLen*accDim))$out, 
		nrow = chnLen, byrow = T)
chn2 <- matrix(.C("GSLmatLoad", "chainsRRG/BV_3_2.gbin", 
	as.integer(chnLen), as.integer(accDim), out = double(chnLen*accDim))$out, 
		nrow = chnLen, byrow = T)
chn3 <- matrix(.C("GSLmatLoad", "chainsRRG/BV_3_3.gbin", 
	as.integer(chnLen), as.integer(accDim), out = double(chnLen*accDim))$out, 
		nrow = chnLen, byrow = T)
chn4 <- matrix(.C("GSLmatLoad", "chainsRRG/BV_3_4.gbin", 
	as.integer(chnLen), as.integer(accDim), out = double(chnLen*accDim))$out, 
		nrow = chnLen, byrow = T)
chn5 <- matrix(.C("GSLmatLoad", "chainsRRG/BV_3_5.gbin", 
	as.integer(chnLen), as.integer(accDim), out = double(chnLen*accDim))$out, 
		nrow = chnLen, byrow = T)

grMat    <- matrix(gelRub("chn",nChn,chnLen), ncol = d, byrow = T)
grMaxInd <- apply(grMat, 2, function(vec){which(vec == max(vec))[1]})
acMat    <- mcmcAcf("chn", 5)

@
Plot the results.
\begin{center}
<<fig=TRUE,width=8.0,height=6.0,echo=FALSE>>=
par(mar = c(5.2, 6, 2, 1), mgp = c(4,1,0))
boxplot(grMat, pch = -0x00b7, outcex = 2, las = 1, ylab = "Gelman-Rubin R", xaxt = "n", col = "grey80", main = "GEBV")
axis(1, at = 1:d, trtNam, las = 1)
@
\end{center}
Plot the worst chains.

\begin{center}
<<fig=TRUE,width=8.0,height=8.0,echo=FALSE>>=
par(mfrow = c(2,2), mar = c(0.2, 4, 0.5, 1), mgp = c(3,1,0))
for (trt in 1:d){
	plotLocEx(trt, grMaxInd[trt], Nacc)
}

@
\end{center}

Now I look at mixing.
\begin{center}
<<fig=TRUE,width=8.0,height=8.0,useDingbats=F,echo=FALSE>>=
par(mfrow = c(2,2), mar = c(0.2, 4, 0.5, 1), mgp = c(3,1,0))
for (trt in 1:d){
	boxplot(acMat[(0:(Nacc-1)*d + trt),], pch = -0x00b7, outcex = 2, las = 1, ylab = trtNam[trt], col = "grey80", xaxt = "n")
}
@
\end{center}

Next I look at covariance matrices. I create an index to isolate just the upper triangles.

<<>>=
upInd   <- matrix(1:(d^2), ncol = d, byrow = T)
upInd   <- upInd[row(upInd) <= col(upInd)]
@
Read in covariance matrix chains. Error correlations are set to zero, I exclude them.

<<>>=
chn1 <- matrix(.C("GSLmatLoad", 
	"chainsRRG/SgS_3_1.gbin", 
	as.integer(chnLen), as.integer(d^2), out = double(chnLen*d^2))$out, 
		nrow = chnLen, byrow = T)[,upInd]
chn1 <- cbind(chn1, 
	matrix(.C("GSLmatLoad", 
	"chainsRRG/SgA_3_1.gbin", 
	as.integer(chnLen), as.integer(d^2), out = double(chnLen*d^2))$out, 
		nrow = chnLen, byrow = T)[,upInd])

chn2 <- matrix(.C("GSLmatLoad", 
	"chainsRRG/SgS_3_2.gbin", 
	as.integer(chnLen), as.integer(d^2), out = double(chnLen*d^2))$out, 
		nrow = chnLen, byrow = T)[,upInd]
chn2 <- cbind(chn2, 
	matrix(.C("GSLmatLoad", 
	"chainsRRG/SgA_3_2.gbin", 
	as.integer(chnLen), as.integer(d^2), out = double(chnLen*d^2))$out, 
		nrow = chnLen, byrow = T)[,upInd])

chn3 <- matrix(.C("GSLmatLoad", 
	"chainsRRG/SgS_3_3.gbin", 
	as.integer(chnLen), as.integer(d^2), out = double(chnLen*d^2))$out, 
		nrow = chnLen, byrow = T)[,upInd]
chn3 <- cbind(chn3, 
	matrix(.C("GSLmatLoad", 
	"chainsRRG/SgA_3_3.gbin", 
	as.integer(chnLen), as.integer(d^2), out = double(chnLen*d^2))$out, 
		nrow = chnLen, byrow = T)[,upInd])

chn4 <- matrix(.C("GSLmatLoad", 
	"chainsRRG/SgS_3_4.gbin", 
	as.integer(chnLen), as.integer(d^2), out = double(chnLen*d^2))$out, 
		nrow = chnLen, byrow = T)[,upInd]
chn4 <- cbind(chn4, 
	matrix(.C("GSLmatLoad", 
	"chainsRRG/SgA_3_4.gbin", 
	as.integer(chnLen), as.integer(d^2), out = double(chnLen*d^2))$out, 
		nrow = chnLen, byrow = T)[,upInd])

chn5 <- matrix(.C("GSLmatLoad", 
	"chainsRRG/SgS_3_5.gbin", 
	as.integer(chnLen), as.integer(d^2), out = double(chnLen*d^2))$out, 
		nrow = chnLen, byrow = T)[,upInd]
chn5 <- cbind(chn5, 
	matrix(.C("GSLmatLoad", 
	"chainsRRG/SgA_3_5.gbin", 
	as.integer(chnLen), as.integer(d^2), out = double(chnLen*d^2))$out, 
		nrow = chnLen, byrow = T)[,upInd])
acMat <- mcmcAcf("chn", 5)

@
Plot the results.

Gelman-Rubin:
\begin{center}
<<fig=TRUE,width=8.0,height=6.0,echo=FALSE>>=
par(mar = c(5.2, 6, 2, 1), mgp = c(4,1,0))
boxplot(matrix(gelRub("chn",nChn,chnLen), ncol = 2), pch = -0x00b7, outcex = 2, las = 1, ylab = "Gelman-Rubin R", xaxt = "n", col = "grey80", main = "Covariance matrices")
axis(1, at = 1:2, c(expression(Sigma[s]), expression(Sigma[a])), las = 1)
@
\end{center}
Now I look at mixing.
\begin{center}
<<fig=TRUE,width=8.0,height=6.0,useDingbats=F,echo=FALSE>>=
par(mfrow = c(2,1), mar = c(0.2, 4, 0.5, 1), mgp = c(3,1,0))
for (sgID in 1:2){
	boxplot(acMat[(0:(length(upInd)-1)*2 + sgID),], pch = -0x00b7, outcex = 2, las = 1, ylab = c(expression(Sigma[e]), expression(Sigma[a]))[sgID], col = "grey80", xaxt = "n")
}
@
\end{center}

<<>>=
tmpInd  <- matrix(1:(d^2), ncol = d, byrow = T)
tmpInd  <- tmpInd[row(tmpInd) < col(tmpInd)]
diagInd <- which(!(upInd %in% tmpInd))
rm(tmpInd)
@

First, I plot the error (SCA) variances.

\begin{center}
<<fig=TRUE,width=8.0,height=6.0,echo=FALSE>>=

par(mfrow = c(3,1), mar = c(0.2, 4, 0.5, 1), mgp = c(3,1,0))
for (trt in 1:d){
	plot( chn1[,diagInd[trt]], type = "l", ylim = range(c(chn1[,diagInd[trt]], chn2[,diagInd[trt]], chn3[,diagInd[trt]], chn4[,diagInd[trt]], chn5[,diagInd[trt]])), col = "orange", ylab = trtNam[trt], xlab = "", xaxt = "n", las = 1)
	lines(chn2[,diagInd[trt]], col = "red")
	lines(chn3[,diagInd[trt]], col = "blue")
	lines(chn4[,diagInd[trt]], col = "green")
	lines(chn5[,diagInd[trt]], col = "magenta")
}
@
\end{center}

Next, the $\sigma^2_a$.

\begin{center}
<<fig=TRUE,width=8.0,height=6.0,echo=FALSE>>=

par(mfrow = c(3,1), mar = c(0.2, 4, 0.5, 1), mgp = c(3,1,0))
for (trt in 1:d){
	plot( chn1[,diagInd[trt] + length(upInd)], type = "l", ylim = range(c(chn1[,diagInd[trt] + length(upInd)], chn2[,diagInd[trt] + length(upInd)], chn3[,diagInd[trt] + length(upInd)], chn4[,diagInd[trt] + length(upInd)], chn5[,diagInd[trt] + length(upInd)])), col = "orange", ylab = trtNam[trt], xlab = "", xaxt = "n", las = 1)
	lines(chn2[,diagInd[trt] + length(upInd)], col = "red")
	lines(chn3[,diagInd[trt] + length(upInd)], col = "blue")
	lines(chn4[,diagInd[trt] + length(upInd)], col = "green")
	lines(chn5[,diagInd[trt] + length(upInd)], col = "magenta")
}
@
\end{center}

Everything looks reasonable. Proceeding with MCMC summaries for use in GWA.

Finally, I try a model with essentially a Gaussian prior on the PC coefficients (analogous to a regular mixed model).
<<>>=
chn1 <- matrix(.C("GSLmatLoad", "chainsRRG/GM_1000_1.gbin", 
	as.integer(chnLen), as.integer(pcDim), out = double(chnLen*pcDim))$out, 
		nrow = chnLen, byrow = T)
chn2 <- matrix(.C("GSLmatLoad", "chainsRRG/GM_1000_2.gbin", 
	as.integer(chnLen), as.integer(pcDim), out = double(chnLen*pcDim))$out, 
		nrow = chnLen, byrow = T)
chn3 <- matrix(.C("GSLmatLoad", "chainsRRG/GM_1000_3.gbin", 
	as.integer(chnLen), as.integer(pcDim), out = double(chnLen*pcDim))$out, 
		nrow = chnLen, byrow = T)
chn4 <- matrix(.C("GSLmatLoad", "chainsRRG/GM_1000_4.gbin", 
	as.integer(chnLen), as.integer(pcDim), out = double(chnLen*pcDim))$out, 
		nrow = chnLen, byrow = T)
chn5 <- matrix(.C("GSLmatLoad", "chainsRRG/GM_1000_5.gbin", 
	as.integer(chnLen), as.integer(pcDim), out = double(chnLen*pcDim))$out, 
		nrow = chnLen, byrow = T)

grMat    <- matrix(gelRub("chn",nChn,chnLen), ncol = d, byrow = T)
grMaxInd <- apply(grMat, 2, function(vec){which(vec == max(vec))[1]})
acMat    <- mcmcAcf("chn", 5)

@

\begin{center}
<<fig=TRUE,width=8.0,height=6.0,echo=FALSE>>=
par(mar = c(5.2, 6, 2, 1), mgp = c(4,1,0))
boxplot(grMat, pch = -0x00b7, outcex = 2, las = 1, ylab = "Gelman-Rubin R", xaxt = "n", col = "grey80", main = "PC coefficients")
axis(1, at = 1:d, trtNam, las = 1)
@
\end{center}
Plot the worst chains.

\begin{center}
<<fig=TRUE,width=8.0,height=8.0,echo=FALSE>>=
par(mfrow = c(2,2), mar = c(0.2, 4, 0.5, 1), mgp = c(3,1,0))
for (trt in 1:d){
	plotLocEx(trt, grMaxInd[trt], Npc)
}

@
\end{center}

Now I look at mixing.
\begin{center}
<<fig=TRUE,width=8.0,height=8.0,useDingbats=F,echo=FALSE>>=
par(mfrow = c(2,2), mar = c(0.2, 4, 0.5, 1), mgp = c(3,1,0))
for (trt in 1:d){
	boxplot(acMat[(0:(Npc-1)*d + trt),], pch = -0x00b7, outcex = 2, las = 1, ylab = trtNam[trt], col = "grey80", xaxt = "n")
}
@
\end{center}

Next, look at GEBVs.
<<>>=
chn1 <- matrix(.C("GSLmatLoad", "chainsRRG/BV_1000_1.gbin", 
	as.integer(chnLen), as.integer(accDim), out = double(chnLen*accDim))$out, 
		nrow = chnLen, byrow = T)
chn2 <- matrix(.C("GSLmatLoad", "chainsRRG/BV_1000_2.gbin", 
	as.integer(chnLen), as.integer(accDim), out = double(chnLen*accDim))$out, 
		nrow = chnLen, byrow = T)
chn3 <- matrix(.C("GSLmatLoad", "chainsRRG/BV_1000_3.gbin", 
	as.integer(chnLen), as.integer(accDim), out = double(chnLen*accDim))$out, 
		nrow = chnLen, byrow = T)
chn4 <- matrix(.C("GSLmatLoad", "chainsRRG/BV_1000_4.gbin", 
	as.integer(chnLen), as.integer(accDim), out = double(chnLen*accDim))$out, 
		nrow = chnLen, byrow = T)
chn5 <- matrix(.C("GSLmatLoad", "chainsRRG/BV_1000_5.gbin", 
	as.integer(chnLen), as.integer(accDim), out = double(chnLen*accDim))$out, 
		nrow = chnLen, byrow = T)

grMat    <- matrix(gelRub("chn",nChn,chnLen), ncol = d, byrow = T)
grMaxInd <- apply(grMat, 2, function(vec){which(vec == max(vec))[1]})
acMat    <- mcmcAcf("chn", 5)
@
Plot the results.
\begin{center}
<<fig=TRUE,width=8.0,height=6.0,echo=FALSE>>=
par(mar = c(5.2, 6, 2, 1), mgp = c(4,1,0))
boxplot(grMat, pch = -0x00b7, outcex = 2, las = 1, ylab = "Gelman-Rubin R", xaxt = "n", col = "grey80", main = "GEBV")
axis(1, at = 1:d, trtNam, las = 1)
@
\end{center}
Plot the worst chains.

\begin{center}
<<fig=TRUE,width=8.0,height=8.0,echo=FALSE>>=
par(mfrow = c(2,2), mar = c(0.2, 4, 0.5, 1), mgp = c(3,1,0))
for (trt in 1:d){
	plotLocEx(trt, grMaxInd[trt], Nacc)
}

@
\end{center}

Now I look at mixing.
\begin{center}
<<fig=TRUE,width=8.0,height=8.0,useDingbats=F,echo=FALSE>>=
par(mfrow = c(2,2), mar = c(0.2, 4, 0.5, 1), mgp = c(3,1,0))
for (trt in 1:d){
	boxplot(acMat[(0:(Nacc-1)*d + trt),], pch = -0x00b7, outcex = 2, las = 1, ylab = trtNam[trt], col = "grey80", xaxt = "n")
}
@
\end{center}

Next I look at covariance matrices. I create an index to isolate just the upper triangles.
<<>>=
chn1 <- matrix(.C("GSLmatLoad", 
	"chainsRRG/SgS_1000_1.gbin", 
	as.integer(chnLen), as.integer(d^2), out = double(chnLen*d^2))$out, 
		nrow = chnLen, byrow = T)[,upInd]
chn1 <- cbind(chn1, 
	matrix(.C("GSLmatLoad", 
	"chainsRRG/SgA_1000_1.gbin", 
	as.integer(chnLen), as.integer(d^2), out = double(chnLen*d^2))$out, 
		nrow = chnLen, byrow = T)[,upInd])

chn2 <- matrix(.C("GSLmatLoad", 
	"chainsRRG/SgS_1000_2.gbin", 
	as.integer(chnLen), as.integer(d^2), out = double(chnLen*d^2))$out, 
		nrow = chnLen, byrow = T)[,upInd]
chn2 <- cbind(chn2, 
	matrix(.C("GSLmatLoad", 
	"chainsRRG/SgA_1000_2.gbin", 
	as.integer(chnLen), as.integer(d^2), out = double(chnLen*d^2))$out, 
		nrow = chnLen, byrow = T)[,upInd])

chn3 <- matrix(.C("GSLmatLoad", 
	"chainsRRG/SgS_1000_3.gbin", 
	as.integer(chnLen), as.integer(d^2), out = double(chnLen*d^2))$out, 
		nrow = chnLen, byrow = T)[,upInd]
chn3 <- cbind(chn3, 
	matrix(.C("GSLmatLoad", 
	"chainsRRG/SgA_1000_3.gbin", 
	as.integer(chnLen), as.integer(d^2), out = double(chnLen*d^2))$out, 
		nrow = chnLen, byrow = T)[,upInd])

chn4 <- matrix(.C("GSLmatLoad", 
	"chainsRRG/SgS_1000_4.gbin", 
	as.integer(chnLen), as.integer(d^2), out = double(chnLen*d^2))$out, 
		nrow = chnLen, byrow = T)[,upInd]
chn4 <- cbind(chn4, 
	matrix(.C("GSLmatLoad", 
	"chainsRRG/SgA_1000_4.gbin", 
	as.integer(chnLen), as.integer(d^2), out = double(chnLen*d^2))$out, 
		nrow = chnLen, byrow = T)[,upInd])

chn5 <- matrix(.C("GSLmatLoad", 
	"chainsRRG/SgS_1000_5.gbin", 
	as.integer(chnLen), as.integer(d^2), out = double(chnLen*d^2))$out, 
		nrow = chnLen, byrow = T)[,upInd]
chn5 <- cbind(chn5, 
	matrix(.C("GSLmatLoad", 
	"chainsRRG/SgA_1000_5.gbin", 
	as.integer(chnLen), as.integer(d^2), out = double(chnLen*d^2))$out, 
		nrow = chnLen, byrow = T)[,upInd])
acMat <- mcmcAcf("chn", 5)

@
Plot the results.

Gelman-Rubin:
\begin{center}
<<fig=TRUE,width=8.0,height=6.0,echo=FALSE>>=
par(mar = c(5.2, 6, 2, 1), mgp = c(4,1,0))
boxplot(matrix(gelRub("chn",nChn,chnLen), ncol = 2), pch = -0x00b7, outcex = 2, las = 1, ylab = "Gelman-Rubin R", xaxt = "n", col = "grey80", main = "Covariance matrices")
axis(1, at = 1:2, c(expression(Sigma[s]), expression(Sigma[a])), las = 1)
@
\end{center}
Now I look at mixing.
\begin{center}
<<fig=TRUE,width=8.0,height=6.0,useDingbats=F,echo=FALSE>>=
par(mfrow = c(2,1), mar = c(0.2, 4, 0.5, 1), mgp = c(3,1,0))
for (sgID in 1:2){
	boxplot(acMat[(0:(length(upInd)-1)*2 + sgID),], pch = -0x00b7, outcex = 2, las = 1, ylab = c(expression(Sigma[e]), expression(Sigma[a]))[sgID], col = "grey80", xaxt = "n")
}
@
\end{center}

\begin{center}
<<fig=TRUE,width=8.0,height=6.0,echo=FALSE>>=

par(mfrow = c(3,1), mar = c(0.2, 4, 0.5, 1), mgp = c(3,1,0))
for (trt in 1:d){
	plot( chn1[,diagInd[trt]], type = "l", ylim = range(c(chn1[,diagInd[trt]], chn2[,diagInd[trt]], chn3[,diagInd[trt]], chn4[,diagInd[trt]], chn5[,diagInd[trt]])), col = "orange", ylab = trtNam[trt], xlab = "", xaxt = "n", las = 1)
	lines(chn2[,diagInd[trt]], col = "red")
	lines(chn3[,diagInd[trt]], col = "blue")
	lines(chn4[,diagInd[trt]], col = "green")
	lines(chn5[,diagInd[trt]], col = "magenta")
}
@
\end{center}

Next, the $\sigma^2_a$.

\begin{center}
<<fig=TRUE,width=8.0,height=6.0,echo=FALSE>>=

par(mfrow = c(3,1), mar = c(0.2, 4, 0.5, 1), mgp = c(3,1,0))
for (trt in 1:d){
	plot( chn1[,diagInd[trt] + length(upInd)], type = "l", ylim = range(c(chn1[,diagInd[trt] + length(upInd)], chn2[,diagInd[trt] + length(upInd)], chn3[,diagInd[trt] + length(upInd)], chn4[,diagInd[trt] + length(upInd)], chn5[,diagInd[trt] + length(upInd)])), col = "orange", ylab = trtNam[trt], xlab = "", xaxt = "n", las = 1)
	lines(chn2[,diagInd[trt] + length(upInd)], col = "red")
	lines(chn3[,diagInd[trt] + length(upInd)], col = "blue")
	lines(chn4[,diagInd[trt] + length(upInd)], col = "green")
	lines(chn5[,diagInd[trt] + length(upInd)], col = "magenta")
}
@
\end{center}
Everything is ready to proceed.


\end{document}
%<<>>=
```
