## Supplemental methods and scripts for "Low additive genetic variation in a trait under selection in domesticated rice": mcmcExamineFAM.Rnw

```
%@
\documentclass[fleqn,11pt]{article}
\usepackage{amsmath}
\usepackage{amssymb}
\usepackage{sectsty}
\usepackage{amsbsy}
\usepackage{fancyhdr}
\usepackage{color}
\usepackage{url}
\usepackage[indention=2em,labelsep=period,labelfont=bf]{caption}
\usepackage[normalsize,nooneline,sf,bf,FIGTOPCAP,tight]{subfigure}
\usepackage[noae]{Sweave} % noae required to sanitize single quotes
\usepackage{fancyvrb}

\textwidth = 6.5 in
\textheight = 8.5 in
\oddsidemargin = 0.0 in
\evensidemargin = 0.0 in
\topmargin = 0.0 in
%\voffset = - 0.4 in
\headheight = 15 pt
\headsep = 0.1 in
\parskip = 0.0 in
\parindent = 4 ex

\definecolor{grey}{rgb}{0.4,0.4,0.4}

\newcommand{\bcdot}{\boldsymbol{\cdot}}

\DefineVerbatimEnvironment{Sinput}{Verbatim}{tabsize=4}
\DefineVerbatimEnvironment{Scode}{Verbatim}{tabsize=4}
\DefineVerbatimEnvironment{Soutput}{Verbatim}{tabsize=4,formatcom=\color{grey},fontsize=\small}

\pagestyle{fancy}
\fancyhf{}
\rhead{Page \thepage}
\lhead{MCMC Convergence}
\chead{Tony Greenberg}

\title{MCMC convergence analyses}
\author{Tony Greenberg}

\begin{document}

\maketitle
\thispagestyle{empty}
<<echo=false>>=
rm(list = ls())
options(width=90)
library(ggplot2)
library(gridExtra)
library(showtext)
font_add("myriad", regular="MYRIADPRO-SEMIBOLD.OTF", 
	bold="MYRIADPRO-BOLD.OTF", italic="MYRIADPRO-SEMIBOLDIT.OTF")
sessionInfo()
@
In this document I examine convergence of Markov chains generated for the Al tolerance project on the old multipopulation data set. I start by looking at the experiment ID regression coefficients, first in analyses using \textit{aus} accessions, then using tropical \textit{japonica} lines.

The chain files are in the binary format generated by saving GSL matrices. R functions to read these files are available from the \texttt{MuGen} GitHub page.

I first define a function that calculates the Gelman-Rubin convergence statistic for multiple MCMC chains.
<<>>=
gelRub <- cmpfun(function(vrNam, nChn, chnLen){
	mat <- NULL
	for (i in 1:nChn){
		mat <- rbind(mat, eval(as.symbol(paste(vrNam, i, sep=""))))
	}
	chnFac <- factor(rep(1:nChn, each=chnLen))
	W <- colMeans(apply(mat, 2, tapply, chnFac, var))
	B <- apply(apply(mat, 2, tapply, chnFac, mean), 2, var)
	return( sqrt((((chnLen - 1)/chnLen)*W + B)/W) )
})

@
I am also interested in the quality of chain mixing. I use autocorrelations to test for that. The following function examines autocorrelations for each chain for a given variable and reports the chain with the highest mean value. The autocorrelations are calculated using default parameters of the \texttt{acf()} function from the \texttt{stats} R package.
<<>>=
eachAcf <- cmpfun(function(i, nChn, vrNam){
	tmp <- NULL
	for (j in 1:nChn){
		tmp <- cbind(tmp, 
				acf(eval(as.symbol(paste(vrNam, j, sep="")))[,i], plot = F)$acf
			)
	}
	return(tmp[,which.max(colMeans(tmp[-1,]))])
})

mcmcAcf <- cmpfun(function(vrNam, nChn){
	 # all chains should have the same dimensions, so take the first one
	N <- ncol(eval(as.symbol(paste(vrNam, 1, sep=""))))
	t(sapply(1:N, eachAcf, nChn, vrNam))
})
@
Read the chains.
<<>>=
trtNam <- c("Control", "160uM_Al")
d      <- 2
Nexp   <- 6
expDim <- d*Nexp
nChn   <- 5
chnLen <- 2000

chn1 <- matrix(.C("GSLmatLoad", 
	"chains/EXPout_aus_3_1.gbin", 
	as.integer(chnLen), as.integer(expDim), out = double(chnLen*expDim))$out, 
		nrow = chnLen, byrow = T)
chn2 <- matrix(.C("GSLmatLoad", 
	"chains/EXPout_aus_3_2.gbin", 
	as.integer(chnLen), as.integer(expDim), out = double(chnLen*expDim))$out, 
		nrow = chnLen, byrow = T)
chn3 <- matrix(.C("GSLmatLoad", 
	"chains/EXPout_aus_3_3.gbin", 
	as.integer(chnLen), as.integer(expDim), out = double(chnLen*expDim))$out, 
		nrow = chnLen, byrow = T)
chn4 <- matrix(.C("GSLmatLoad", 
	"chains/EXPout_aus_3_4.gbin", 
	as.integer(chnLen), as.integer(expDim), out = double(chnLen*expDim))$out, 
		nrow = chnLen, byrow = T)
chn5 <- matrix(.C("GSLmatLoad", 
	"chains/EXPout_aus_3_5.gbin", 
	as.integer(chnLen), as.integer(expDim), out = double(chnLen*expDim))$out, 
		nrow = chnLen, byrow = T)

grMat    <- matrix(gelRub("chn", nChn, chnLen), ncol = d, byrow = T)
grFrame  <- data.frame(Gelman.Rubin=c(grMat), trait=rep(trtNam, each = Nexp))
grMaxInd <- apply(grMat, 2, function(vec){which(vec == max(vec))[1]})
acMat    <- mcmcAcf("chn", 5)
acFrame  <- data.frame(autocorrelation=c(acMat), trait=rep(trtNam, prod(dim(acMat))/2),
	lag=rep(0:33, each = nrow(acMat)))

@
Plot the \textit{aus} results.

\begin{center}
<<fig=TRUE,width=8.0,height=8.0>>=
showtext_auto()
ggplot(data=grFrame, 
		aes(x=trait, y=Gelman.Rubin)) +
	geom_boxplot(fill="grey80") + 
	theme_classic(base_size=18, base_family="myriad")

@
\end{center}

Values smaller than $\sim$1.2 indicate good convergence. I check by plotting an the chains with the worst G-R statistic for each trait.

<<>>=
plotLocEx <- cmpfun(function(trt.i, samp.j, N){
	i <- ( d*(samp.j-1) ) + trt.i
	plot( chn1[,i], type = "l", 
		ylim = range(c(chn1[,i], chn2[,i], chn3[,i], chn4[,i], chn5[,i])), 
		col = "orange", ylab = trtNam[trt.i], xaxt = "n", las = 1)
	lines(chn2[,i], col = "red")
	lines(chn3[,i], col = "blue")
	lines(chn4[,i], col = "green")
	lines(chn5[,i], col = "magenta")
})
@

\begin{center}
<<fig=TRUE,width=6.0,height=8.0,echo=FALSE>>=
par(mfrow = c(2,1), mar = c(0.2, 4, 0.5, 1), mgp = c(3,1,0))
for (trt in 1:d){
	plotLocEx(trt, grMaxInd[trt], Nexp)
}
@
\end{center}

Now I look at mixing. Lag is on the $x$ axis.
\begin{center}
<<fig=TRUE,width=8.0,height=10.0>>=
ggTrt1 <- ggplot(data=subset(acFrame, trait==trtNam[1]), 
		aes(x=lag, y=autocorrelation)) +
	geom_boxplot(fill="grey80", aes(group=cut_interval(lag, n=34))) + 
	theme_classic(base_size=18, base_family="myriad") +
	labs(title=trtNam[1])
ggTrt2 <- ggplot(data=subset(acFrame, trait==trtNam[2]), 
		aes(x=lag, y=autocorrelation)) +
	geom_boxplot(fill="grey80", aes(group=cut_interval(lag, n=34))) + 
	theme_classic(base_size=18, base_family="myriad") + 
	labs(title=trtNam[2])
showtext_auto()
grid.arrange(ggTrt1, ggTrt2, nrow = 2)

@
\end{center}
Now do the tropical \textit{japonica} results.

<<>>=

chn1 <- matrix(.C("GSLmatLoad", 
	"chains/EXPout_trj_3_1.gbin", 
	as.integer(chnLen), as.integer(expDim), out = double(chnLen*expDim))$out, 
		nrow = chnLen, byrow = T)
chn2 <- matrix(.C("GSLmatLoad", 
	"chains/EXPout_trj_3_2.gbin", 
	as.integer(chnLen), as.integer(expDim), out = double(chnLen*expDim))$out, 
		nrow = chnLen, byrow = T)
chn3 <- matrix(.C("GSLmatLoad", 
	"chains/EXPout_trj_3_3.gbin", 
	as.integer(chnLen), as.integer(expDim), out = double(chnLen*expDim))$out, 
		nrow = chnLen, byrow = T)
chn4 <- matrix(.C("GSLmatLoad", 
	"chains/EXPout_trj_3_4.gbin", 
	as.integer(chnLen), as.integer(expDim), out = double(chnLen*expDim))$out, 
		nrow = chnLen, byrow = T)
chn5 <- matrix(.C("GSLmatLoad", 
	"chains/EXPout_trj_3_5.gbin", 
	as.integer(chnLen), as.integer(expDim), out = double(chnLen*expDim))$out, 
		nrow = chnLen, byrow = T)

grMat    <- matrix(gelRub("chn", nChn, chnLen), ncol = d, byrow = T)
grFrame  <- data.frame(Gelman.Rubin=c(grMat), trait=rep(trtNam, each = Nexp))
grMaxInd <- apply(grMat, 2, function(vec){which(vec == max(vec))[1]})
acMat    <- mcmcAcf("chn", 5)
acFrame  <- data.frame(autocorrelation=c(acMat), trait=rep(trtNam, prod(dim(acMat))/2),
	lag=rep(0:33, each = nrow(acMat)))

@
\begin{center}
<<fig=TRUE,width=8.0,height=8.0>>=
showtext_auto()
ggplot(data=grFrame, 
		aes(x=trait, y=Gelman.Rubin)) +
	geom_boxplot(fill="grey80") + 
	theme_classic(base_size=18, base_family="myriad")

@
\end{center}

Values smaller than $\sim$1.2 indicate good convergence. I check by plotting an the chains with the worst G-R statistic for each trait.

\begin{center}
<<fig=TRUE,width=6.0,height=8.0,echo=FALSE>>=
par(mfrow = c(2,1), mar = c(0.2, 4, 0.5, 1), mgp = c(3,1,0))
for (trt in 1:d){
	plotLocEx(trt, grMaxInd[trt], Nexp)
}
@
\end{center}

Now I look at mixing. Lag is on the $x$ axis.
\begin{center}
<<fig=TRUE,width=8.0,height=10.0>>=
ggTrt1 <- ggplot(data=subset(acFrame, trait==trtNam[1]), 
		aes(x=lag, y=autocorrelation)) +
	geom_boxplot(fill="grey80", aes(group=cut_interval(lag, n=34))) + 
	theme_classic(base_size=18, base_family="myriad") +
	labs(title=trtNam[1])
ggTrt2 <- ggplot(data=subset(acFrame, trait==trtNam[2]), 
		aes(x=lag, y=autocorrelation)) +
	geom_boxplot(fill="grey80", aes(group=cut_interval(lag, n=34))) + 
	theme_classic(base_size=18, base_family="myriad") + 
	labs(title=trtNam[2])
showtext_auto()
grid.arrange(ggTrt1, ggTrt2, nrow = 2)

@
\end{center}
Next, tropical \textit{japonica} with admixture.
<<>>=

chn1 <- matrix(.C("GSLmatLoad", 
	"chains/EXPout_trja_3_1.gbin", 
	as.integer(chnLen), as.integer(expDim), out = double(chnLen*expDim))$out, 
		nrow = chnLen, byrow = T)
chn2 <- matrix(.C("GSLmatLoad", 
	"chains/EXPout_trja_3_2.gbin", 
	as.integer(chnLen), as.integer(expDim), out = double(chnLen*expDim))$out, 
		nrow = chnLen, byrow = T)
chn3 <- matrix(.C("GSLmatLoad", 
	"chains/EXPout_trja_3_3.gbin", 
	as.integer(chnLen), as.integer(expDim), out = double(chnLen*expDim))$out, 
		nrow = chnLen, byrow = T)
chn4 <- matrix(.C("GSLmatLoad", 
	"chains/EXPout_trja_3_4.gbin", 
	as.integer(chnLen), as.integer(expDim), out = double(chnLen*expDim))$out, 
		nrow = chnLen, byrow = T)
chn5 <- matrix(.C("GSLmatLoad", 
	"chains/EXPout_trja_3_5.gbin", 
	as.integer(chnLen), as.integer(expDim), out = double(chnLen*expDim))$out, 
		nrow = chnLen, byrow = T)

grMat    <- matrix(gelRub("chn", nChn, chnLen), ncol = d, byrow = T)
grFrame  <- data.frame(Gelman.Rubin=c(grMat), trait=rep(trtNam, each = Nexp))
grMaxInd <- apply(grMat, 2, function(vec){which(vec == max(vec))[1]})
acMat    <- mcmcAcf("chn", 5)
acFrame  <- data.frame(autocorrelation=c(acMat), trait=rep(trtNam, prod(dim(acMat))/2),
	lag=rep(0:33, each = nrow(acMat)))

@
\begin{center}
<<fig=TRUE,width=8.0,height=8.0>>=
showtext_auto()
ggplot(data=grFrame, 
		aes(x=trait, y=Gelman.Rubin)) +
	geom_boxplot(fill="grey80") + 
	theme_classic(base_size=18, base_family="myriad")

@
\end{center}

Values smaller than $\sim$1.2 indicate good convergence. I check by plotting an the chains with the worst G-R statistic for each trait.

\begin{center}
<<fig=TRUE,width=6.0,height=8.0,echo=FALSE>>=
par(mfrow = c(2,1), mar = c(0.2, 4, 0.5, 1), mgp = c(3,1,0))
for (trt in 1:d){
	plotLocEx(trt, grMaxInd[trt], Nexp)
}
@
\end{center}

Now I look at mixing. Lag is on the $x$ axis.
\begin{center}
<<fig=TRUE,width=8.0,height=10.0>>=
ggTrt1 <- ggplot(data=subset(acFrame, trait==trtNam[1]), 
		aes(x=lag, y=autocorrelation)) +
	geom_boxplot(fill="grey80", aes(group=cut_interval(lag, n=34))) + 
	theme_classic(base_size=18, base_family="myriad") +
	labs(title=trtNam[1])
ggTrt2 <- ggplot(data=subset(acFrame, trait==trtNam[2]), 
		aes(x=lag, y=autocorrelation)) +
	geom_boxplot(fill="grey80", aes(group=cut_interval(lag, n=34))) + 
	theme_classic(base_size=18, base_family="myriad") + 
	labs(title=trtNam[2])
showtext_auto()
grid.arrange(ggTrt1, ggTrt2, nrow = 2)

@
\end{center}

Everything looks good. I next look at the accession means, \textit{aus} first.

<<>>=
Nacc   <- 55
locDim <- d*Nacc

chn1 <- matrix(.C("GSLmatLoad", 
	"chains/LNout_aus_3_1.gbin", 
	as.integer(chnLen), as.integer(locDim), out = double(chnLen*locDim))$out, 
		nrow = chnLen, byrow = T)
chn2 <- matrix(.C("GSLmatLoad", 
	"chains/LNout_aus_3_2.gbin", 
	as.integer(chnLen), as.integer(locDim), out = double(chnLen*locDim))$out, 
		nrow = chnLen, byrow = T)
chn3 <- matrix(.C("GSLmatLoad", 
	"chains/LNout_aus_3_3.gbin", 
	as.integer(chnLen), as.integer(locDim), out = double(chnLen*locDim))$out, 
		nrow = chnLen, byrow = T)
chn4 <- matrix(.C("GSLmatLoad", 
	"chains/LNout_aus_3_4.gbin", 
	as.integer(chnLen), as.integer(locDim), out = double(chnLen*locDim))$out, 
		nrow = chnLen, byrow = T)
chn5 <- matrix(.C("GSLmatLoad", 
	"chains/LNout_aus_3_5.gbin", 
	as.integer(chnLen), as.integer(locDim), out = double(chnLen*locDim))$out, 
		nrow = chnLen, byrow = T)

grMat    <- matrix(gelRub("chn", nChn, chnLen), ncol = d, byrow = T)
grFrame  <- data.frame(Gelman.Rubin=c(grMat), trait=rep(trtNam, each = Nacc))
grMaxInd <- apply(grMat, 2, function(vec){which(vec == max(vec))[1]})
acMat    <- mcmcAcf("chn", 5)
acFrame  <- data.frame(autocorrelation=c(acMat), trait=rep(trtNam, prod(dim(acMat))/2),
	lag=rep(0:33, each = nrow(acMat)))

@
Plot the \textit{aus} results.

\begin{center}
<<fig=TRUE,width=8.0,height=8.0>>=
showtext_auto()
ggplot(data=grFrame, 
		aes(x=trait, y=Gelman.Rubin)) +
	geom_boxplot(fill="grey80") + 
	theme_classic(base_size=18, base_family="myriad")

@
\end{center}

Values smaller than $\sim$1.2 indicate good convergence. I check by plotting an the chains with the worst G-R statistic for each trait.

\begin{center}
<<fig=TRUE,width=6.0,height=8.0,echo=FALSE>>=
par(mfrow = c(2,1), mar = c(0.2, 4, 0.5, 1), mgp = c(3,1,0))
for (trt in 1:d){
	plotLocEx(trt, grMaxInd[trt], Nacc)
}
@
\end{center}

Now I look at mixing. Lag is on the $x$ axis.
\begin{center}
<<fig=TRUE,width=8.0,height=10.0>>=
ggTrt1 <- ggplot(data=subset(acFrame, trait==trtNam[1]), 
		aes(x=lag, y=autocorrelation)) +
	geom_boxplot(fill="grey80", aes(group=cut_interval(lag, n=34))) + 
	theme_classic(base_size=18, base_family="myriad") +
	labs(title=trtNam[1])
ggTrt2 <- ggplot(data=subset(acFrame, trait==trtNam[2]), 
		aes(x=lag, y=autocorrelation)) +
	geom_boxplot(fill="grey80", aes(group=cut_interval(lag, n=34))) + 
	theme_classic(base_size=18, base_family="myriad") + 
	labs(title=trtNam[2])
showtext_auto()
grid.arrange(ggTrt1, ggTrt2, nrow = 2)

@
\end{center}
Looks good. Now do the tropical \textit{japonica}.

<<>>=
Nacc   <- 92
locDim <- d*Nacc

chn1 <- matrix(.C("GSLmatLoad", 
	"chains/LNout_trj_3_1.gbin", 
	as.integer(chnLen), as.integer(locDim), out = double(chnLen*locDim))$out, 
		nrow = chnLen, byrow = T)
chn2 <- matrix(.C("GSLmatLoad", 
	"chains/LNout_trj_3_2.gbin", 
	as.integer(chnLen), as.integer(locDim), out = double(chnLen*locDim))$out, 
		nrow = chnLen, byrow = T)
chn3 <- matrix(.C("GSLmatLoad", 
	"chains/LNout_trj_3_3.gbin", 
	as.integer(chnLen), as.integer(locDim), out = double(chnLen*locDim))$out, 
		nrow = chnLen, byrow = T)
chn4 <- matrix(.C("GSLmatLoad", 
	"chains/LNout_trj_3_4.gbin", 
	as.integer(chnLen), as.integer(locDim), out = double(chnLen*locDim))$out, 
		nrow = chnLen, byrow = T)
chn5 <- matrix(.C("GSLmatLoad", 
	"chains/LNout_trj_3_5.gbin", 
	as.integer(chnLen), as.integer(locDim), out = double(chnLen*locDim))$out, 
		nrow = chnLen, byrow = T)

grMat    <- matrix(gelRub("chn", nChn, chnLen), ncol = d, byrow = T)
grFrame  <- data.frame(Gelman.Rubin=c(grMat), trait=rep(trtNam, each = Nacc))
grMaxInd <- apply(grMat, 2, function(vec){which(vec == max(vec))[1]})
acMat    <- mcmcAcf("chn", 5)
acFrame  <- data.frame(autocorrelation=c(acMat), trait=rep(trtNam, prod(dim(acMat))/2),
	lag=rep(0:33, each = nrow(acMat)))

@
Plot the \textit{aus} results.

\begin{center}
<<fig=TRUE,width=8.0,height=8.0>>=
showtext_auto()
ggplot(data=grFrame, 
		aes(x=trait, y=Gelman.Rubin)) +
	geom_boxplot(fill="grey80") + 
	theme_classic(base_size=18, base_family="myriad")

@
\end{center}

Values smaller than $\sim$1.2 indicate good convergence. I check by plotting an the chains with the worst G-R statistic for each trait.

\begin{center}
<<fig=TRUE,width=6.0,height=8.0,echo=FALSE>>=
par(mfrow = c(2,1), mar = c(0.2, 4, 0.5, 1), mgp = c(3,1,0))
for (trt in 1:d){
	plotLocEx(trt, grMaxInd[trt], Nacc)
}
@
\end{center}

Now I look at mixing. Lag is on the $x$ axis.
\begin{center}
<<fig=TRUE,width=8.0,height=10.0>>=
ggTrt1 <- ggplot(data=subset(acFrame, trait==trtNam[1]), 
		aes(x=lag, y=autocorrelation)) +
	geom_boxplot(fill="grey80", aes(group=cut_interval(lag, n=34))) + 
	theme_classic(base_size=18, base_family="myriad") +
	labs(title=trtNam[1])
ggTrt2 <- ggplot(data=subset(acFrame, trait==trtNam[2]), 
		aes(x=lag, y=autocorrelation)) +
	geom_boxplot(fill="grey80", aes(group=cut_interval(lag, n=34))) + 
	theme_classic(base_size=18, base_family="myriad") + 
	labs(title=trtNam[2])
showtext_auto()
grid.arrange(ggTrt1, ggTrt2, nrow = 2)

@
\end{center}

Next, admixed tropical \textit{japonica}.
<<>>=
Nacc   <- 95
locDim <- d*Nacc

chn1 <- matrix(.C("GSLmatLoad", 
	"chains/LNout_trja_3_1.gbin", 
	as.integer(chnLen), as.integer(locDim), out = double(chnLen*locDim))$out, 
		nrow = chnLen, byrow = T)
chn2 <- matrix(.C("GSLmatLoad", 
	"chains/LNout_trja_3_2.gbin", 
	as.integer(chnLen), as.integer(locDim), out = double(chnLen*locDim))$out, 
		nrow = chnLen, byrow = T)
chn3 <- matrix(.C("GSLmatLoad", 
	"chains/LNout_trja_3_3.gbin", 
	as.integer(chnLen), as.integer(locDim), out = double(chnLen*locDim))$out, 
		nrow = chnLen, byrow = T)
chn4 <- matrix(.C("GSLmatLoad", 
	"chains/LNout_trja_3_4.gbin", 
	as.integer(chnLen), as.integer(locDim), out = double(chnLen*locDim))$out, 
		nrow = chnLen, byrow = T)
chn5 <- matrix(.C("GSLmatLoad", 
	"chains/LNout_trja_3_5.gbin", 
	as.integer(chnLen), as.integer(locDim), out = double(chnLen*locDim))$out, 
		nrow = chnLen, byrow = T)

grMat    <- matrix(gelRub("chn", nChn, chnLen), ncol = d, byrow = T)
grFrame  <- data.frame(Gelman.Rubin=c(grMat), trait=rep(trtNam, each = Nacc))
grMaxInd <- apply(grMat, 2, function(vec){which(vec == max(vec))[1]})
acMat    <- mcmcAcf("chn", 5)
acFrame  <- data.frame(autocorrelation=c(acMat), trait=rep(trtNam, prod(dim(acMat))/2),
	lag=rep(0:33, each = nrow(acMat)))

@
Plot the \textit{aus} results.

\begin{center}
<<fig=TRUE,width=8.0,height=8.0>>=
showtext_auto()
ggplot(data=grFrame, 
		aes(x=trait, y=Gelman.Rubin)) +
	geom_boxplot(fill="grey80") + 
	theme_classic(base_size=18, base_family="myriad")

@
\end{center}

Values smaller than $\sim$1.2 indicate good convergence. I check by plotting an the chains with the worst G-R statistic for each trait.

\begin{center}
<<fig=TRUE,width=6.0,height=8.0,echo=FALSE>>=
par(mfrow = c(2,1), mar = c(0.2, 4, 0.5, 1), mgp = c(3,1,0))
for (trt in 1:d){
	plotLocEx(trt, grMaxInd[trt], Nacc)
}
@
\end{center}

Now I look at mixing. Lag is on the $x$ axis.
\begin{center}
<<fig=TRUE,width=8.0,height=10.0>>=
ggTrt1 <- ggplot(data=subset(acFrame, trait==trtNam[1]), 
		aes(x=lag, y=autocorrelation)) +
	geom_boxplot(fill="grey80", aes(group=cut_interval(lag, n=34))) + 
	theme_classic(base_size=18, base_family="myriad") +
	labs(title=trtNam[1])
ggTrt2 <- ggplot(data=subset(acFrame, trait==trtNam[2]), 
		aes(x=lag, y=autocorrelation)) +
	geom_boxplot(fill="grey80", aes(group=cut_interval(lag, n=34))) + 
	theme_classic(base_size=18, base_family="myriad") + 
	labs(title=trtNam[2])
showtext_auto()
grid.arrange(ggTrt1, ggTrt2, nrow = 2)

@
\end{center}

Everything looks good. Now examine GEBV chains, again starting with \textit{aus}.

<<>>=
Nacc   <- 55
locDim <- d*Nacc

chn1 <- matrix(.C("GSLmatLoad", 
	"chains/BVout_aus_3_1.gbin", 
	as.integer(chnLen), as.integer(locDim), out = double(chnLen*locDim))$out, 
		nrow = chnLen, byrow = T)
chn2 <- matrix(.C("GSLmatLoad", 
	"chains/BVout_aus_3_2.gbin", 
	as.integer(chnLen), as.integer(locDim), out = double(chnLen*locDim))$out, 
		nrow = chnLen, byrow = T)
chn3 <- matrix(.C("GSLmatLoad", 
	"chains/BVout_aus_3_3.gbin", 
	as.integer(chnLen), as.integer(locDim), out = double(chnLen*locDim))$out, 
		nrow = chnLen, byrow = T)
chn4 <- matrix(.C("GSLmatLoad", 
	"chains/BVout_aus_3_4.gbin", 
	as.integer(chnLen), as.integer(locDim), out = double(chnLen*locDim))$out, 
		nrow = chnLen, byrow = T)
chn5 <- matrix(.C("GSLmatLoad", 
	"chains/BVout_aus_3_5.gbin", 
	as.integer(chnLen), as.integer(locDim), out = double(chnLen*locDim))$out, 
		nrow = chnLen, byrow = T)

grMat    <- matrix(gelRub("chn", nChn, chnLen), ncol = d, byrow = T)
grFrame  <- data.frame(Gelman.Rubin=c(grMat), trait=rep(trtNam, each = Nacc))
grMaxInd <- apply(grMat, 2, function(vec){which(vec == max(vec))[1]})
acMat    <- mcmcAcf("chn", 5)
acFrame  <- data.frame(autocorrelation=c(acMat), trait=rep(trtNam, prod(dim(acMat))/2),
	lag=rep(0:33, each = nrow(acMat)))

@
Plot the \textit{aus} results.

\begin{center}
<<fig=TRUE,width=8.0,height=8.0>>=
showtext_auto()
ggplot(data=grFrame, 
		aes(x=trait, y=Gelman.Rubin)) +
	geom_boxplot(fill="grey80") + 
	theme_classic(base_size=18, base_family="myriad")

@
\end{center}

Values smaller than $\sim$1.2 indicate good convergence. I check by plotting an the chains with the worst G-R statistic for each trait.

\begin{center}
<<fig=TRUE,width=6.0,height=8.0,echo=FALSE>>=
par(mfrow = c(2,1), mar = c(0.2, 4, 0.5, 1), mgp = c(3,1,0))
for (trt in 1:d){
	plotLocEx(trt, grMaxInd[trt], Nacc)
}
@
\end{center}

Now I look at mixing. Lag is on the $x$ axis.
\begin{center}
<<fig=TRUE,width=8.0,height=10.0>>=
ggTrt1 <- ggplot(data=subset(acFrame, trait==trtNam[1]), 
		aes(x=lag, y=autocorrelation)) +
	geom_boxplot(fill="grey80", aes(group=cut_interval(lag, n=34))) + 
	theme_classic(base_size=18, base_family="myriad") +
	labs(title=trtNam[1])
ggTrt2 <- ggplot(data=subset(acFrame, trait==trtNam[2]), 
		aes(x=lag, y=autocorrelation)) +
	geom_boxplot(fill="grey80", aes(group=cut_interval(lag, n=34))) + 
	theme_classic(base_size=18, base_family="myriad") + 
	labs(title=trtNam[2])
showtext_auto()
grid.arrange(ggTrt1, ggTrt2, nrow = 2)

@
\end{center}
Looks good. Now do the tropical \textit{japonica}.

<<>>=
Nacc   <- 92
locDim <- d*Nacc

chn1 <- matrix(.C("GSLmatLoad", 
	"chains/BVout_trj_3_1.gbin", 
	as.integer(chnLen), as.integer(locDim), out = double(chnLen*locDim))$out, 
		nrow = chnLen, byrow = T)
chn2 <- matrix(.C("GSLmatLoad", 
	"chains/BVout_trj_3_2.gbin", 
	as.integer(chnLen), as.integer(locDim), out = double(chnLen*locDim))$out, 
		nrow = chnLen, byrow = T)
chn3 <- matrix(.C("GSLmatLoad", 
	"chains/BVout_trj_3_3.gbin", 
	as.integer(chnLen), as.integer(locDim), out = double(chnLen*locDim))$out, 
		nrow = chnLen, byrow = T)
chn4 <- matrix(.C("GSLmatLoad", 
	"chains/BVout_trj_3_4.gbin", 
	as.integer(chnLen), as.integer(locDim), out = double(chnLen*locDim))$out, 
		nrow = chnLen, byrow = T)
chn5 <- matrix(.C("GSLmatLoad", 
	"chains/BVout_trj_3_5.gbin", 
	as.integer(chnLen), as.integer(locDim), out = double(chnLen*locDim))$out, 
		nrow = chnLen, byrow = T)

grMat    <- matrix(gelRub("chn", nChn, chnLen), ncol = d, byrow = T)
grFrame  <- data.frame(Gelman.Rubin=c(grMat), trait=rep(trtNam, each = Nacc))
grMaxInd <- apply(grMat, 2, function(vec){which(vec == max(vec))[1]})
acMat    <- mcmcAcf("chn", 5)
acFrame  <- data.frame(autocorrelation=c(acMat), trait=rep(trtNam, prod(dim(acMat))/2),
	lag=rep(0:33, each = nrow(acMat)))

@
Plot the \textit{aus} results.

\begin{center}
<<fig=TRUE,width=8.0,height=8.0>>=
showtext_auto()
ggplot(data=grFrame, 
		aes(x=trait, y=Gelman.Rubin)) +
	geom_boxplot(fill="grey80") + 
	theme_classic(base_size=18, base_family="myriad")

@
\end{center}

Values smaller than $\sim$1.2 indicate good convergence. I check by plotting an the chains with the worst G-R statistic for each trait.

\begin{center}
<<fig=TRUE,width=6.0,height=8.0,echo=FALSE>>=
par(mfrow = c(2,1), mar = c(0.2, 4, 0.5, 1), mgp = c(3,1,0))
for (trt in 1:d){
	plotLocEx(trt, grMaxInd[trt], Nacc)
}
@
\end{center}

Now I look at mixing. Lag is on the $x$ axis.
\begin{center}
<<fig=TRUE,width=8.0,height=10.0>>=
ggTrt1 <- ggplot(data=subset(acFrame, trait==trtNam[1]), 
		aes(x=lag, y=autocorrelation)) +
	geom_boxplot(fill="grey80", aes(group=cut_interval(lag, n=34))) + 
	theme_classic(base_size=18, base_family="myriad") +
	labs(title=trtNam[1])
ggTrt2 <- ggplot(data=subset(acFrame, trait==trtNam[2]), 
		aes(x=lag, y=autocorrelation)) +
	geom_boxplot(fill="grey80", aes(group=cut_interval(lag, n=34))) + 
	theme_classic(base_size=18, base_family="myriad") + 
	labs(title=trtNam[2])
showtext_auto()
grid.arrange(ggTrt1, ggTrt2, nrow = 2)

@
\end{center}
Next, admixed tropical \textit{japonica}.

<<>>=
Nacc   <- 95
locDim <- d*Nacc

chn1 <- matrix(.C("GSLmatLoad", 
	"chains/BVout_trja_3_1.gbin", 
	as.integer(chnLen), as.integer(locDim), out = double(chnLen*locDim))$out, 
		nrow = chnLen, byrow = T)
chn2 <- matrix(.C("GSLmatLoad", 
	"chains/BVout_trja_3_2.gbin", 
	as.integer(chnLen), as.integer(locDim), out = double(chnLen*locDim))$out, 
		nrow = chnLen, byrow = T)
chn3 <- matrix(.C("GSLmatLoad", 
	"chains/BVout_trja_3_3.gbin", 
	as.integer(chnLen), as.integer(locDim), out = double(chnLen*locDim))$out, 
		nrow = chnLen, byrow = T)
chn4 <- matrix(.C("GSLmatLoad", 
	"chains/BVout_trja_3_4.gbin", 
	as.integer(chnLen), as.integer(locDim), out = double(chnLen*locDim))$out, 
		nrow = chnLen, byrow = T)
chn5 <- matrix(.C("GSLmatLoad", 
	"chains/BVout_trja_3_5.gbin", 
	as.integer(chnLen), as.integer(locDim), out = double(chnLen*locDim))$out, 
		nrow = chnLen, byrow = T)

grMat    <- matrix(gelRub("chn", nChn, chnLen), ncol = d, byrow = T)
grFrame  <- data.frame(Gelman.Rubin=c(grMat), trait=rep(trtNam, each = Nacc))
grMaxInd <- apply(grMat, 2, function(vec){which(vec == max(vec))[1]})
acMat    <- mcmcAcf("chn", 5)
acFrame  <- data.frame(autocorrelation=c(acMat), trait=rep(trtNam, prod(dim(acMat))/2),
	lag=rep(0:33, each = nrow(acMat)))

@
Plot the \textit{aus} results.

\begin{center}
<<fig=TRUE,width=8.0,height=8.0>>=
showtext_auto()
ggplot(data=grFrame, 
		aes(x=trait, y=Gelman.Rubin)) +
	geom_boxplot(fill="grey80") + 
	theme_classic(base_size=18, base_family="myriad")

@
\end{center}

Values smaller than $\sim$1.2 indicate good convergence. I check by plotting an the chains with the worst G-R statistic for each trait.

\begin{center}
<<fig=TRUE,width=6.0,height=8.0,echo=FALSE>>=
par(mfrow = c(2,1), mar = c(0.2, 4, 0.5, 1), mgp = c(3,1,0))
for (trt in 1:d){
	plotLocEx(trt, grMaxInd[trt], Nacc)
}
@
\end{center}

Now I look at mixing. Lag is on the $x$ axis.
\begin{center}
<<fig=TRUE,width=8.0,height=10.0>>=
ggTrt1 <- ggplot(data=subset(acFrame, trait==trtNam[1]), 
		aes(x=lag, y=autocorrelation)) +
	geom_boxplot(fill="grey80", aes(group=cut_interval(lag, n=34))) + 
	theme_classic(base_size=18, base_family="myriad") +
	labs(title=trtNam[1])
ggTrt2 <- ggplot(data=subset(acFrame, trait==trtNam[2]), 
		aes(x=lag, y=autocorrelation)) +
	geom_boxplot(fill="grey80", aes(group=cut_interval(lag, n=34))) + 
	theme_classic(base_size=18, base_family="myriad") + 
	labs(title=trtNam[2])
showtext_auto()
grid.arrange(ggTrt1, ggTrt2, nrow = 2)

@
\end{center}

Finally, I look at covariance matrices. I create an index to isolate just the upper triangles. I again start with \textit{aus}.

<<>>=
upInd   <- matrix(1:(d^2), ncol = d, byrow = T)
upInd   <- upInd[row(upInd) <= col(upInd)]
diagInd <- diag(matrix(1:(d^2), ncol = d, byrow = T))
@
Read in covariance matrix chains. Error correlations are set to zero, I exclude them.

<<>>=
chn1 <- matrix(.C("GSLmatLoad", 
	"chains/SgEout_aus_3_1.gbin", 
	as.integer(chnLen), as.integer(d^2), out = double(chnLen*d^2))$out, 
		nrow = chnLen, byrow = T)[,diagInd]
chn1 <- cbind(chn1, 
	matrix(.C("GSLmatLoad", 
	"chains/SgSout_aus_3_1.gbin", 
	as.integer(chnLen), as.integer(d^2), out = double(chnLen*d^2))$out, 
		nrow = chnLen, byrow = T)[,upInd])
chn1 <- cbind(chn1, 
	matrix(.C("GSLmatLoad", 
	"chains/SgAout_aus_3_1.gbin", 
	as.integer(chnLen), as.integer(d^2), out = double(chnLen*d^2))$out, 
		nrow = chnLen, byrow = T)[,upInd])

chn2 <- matrix(.C("GSLmatLoad", 
	"chains/SgEout_aus_3_2.gbin", 
	as.integer(chnLen), as.integer(d^2), out = double(chnLen*d^2))$out, 
		nrow = chnLen, byrow = T)[,diagInd]
chn2 <- cbind(chn2, 
	matrix(.C("GSLmatLoad", 
	"chains/SgSout_aus_3_2.gbin", 
	as.integer(chnLen), as.integer(d^2), out = double(chnLen*d^2))$out, 
		nrow = chnLen, byrow = T)[,upInd])
chn2 <- cbind(chn2, 
	matrix(.C("GSLmatLoad", 
	"chains/SgAout_aus_3_2.gbin", 
	as.integer(chnLen), as.integer(d^2), out = double(chnLen*d^2))$out, 
		nrow = chnLen, byrow = T)[,upInd])

chn3 <- matrix(.C("GSLmatLoad", 
	"chains/SgEout_aus_3_3.gbin", 
	as.integer(chnLen), as.integer(d^2), out = double(chnLen*d^2))$out, 
		nrow = chnLen, byrow = T)[,diagInd]
chn3 <- cbind(chn3, 
	matrix(.C("GSLmatLoad", 
	"chains/SgSout_aus_3_3.gbin", 
	as.integer(chnLen), as.integer(d^2), out = double(chnLen*d^2))$out, 
		nrow = chnLen, byrow = T)[,upInd])
chn3 <- cbind(chn3, 
	matrix(.C("GSLmatLoad", 
	"chains/SgAout_aus_3_3.gbin", 
	as.integer(chnLen), as.integer(d^2), out = double(chnLen*d^2))$out, 
		nrow = chnLen, byrow = T)[,upInd])

chn4 <- matrix(.C("GSLmatLoad", 
	"chains/SgEout_aus_3_4.gbin", 
	as.integer(chnLen), as.integer(d^2), out = double(chnLen*d^2))$out, 
		nrow = chnLen, byrow = T)[,diagInd]
chn4 <- cbind(chn4, 
	matrix(.C("GSLmatLoad", 
	"chains/SgSout_aus_3_4.gbin", 
	as.integer(chnLen), as.integer(d^2), out = double(chnLen*d^2))$out, 
		nrow = chnLen, byrow = T)[,upInd])
chn4 <- cbind(chn4, 
	matrix(.C("GSLmatLoad", 
	"chains/SgAout_aus_3_4.gbin", 
	as.integer(chnLen), as.integer(d^2), out = double(chnLen*d^2))$out, 
		nrow = chnLen, byrow = T)[,upInd])

chn5 <- matrix(.C("GSLmatLoad", 
	"chains/SgEout_aus_3_5.gbin", 
	as.integer(chnLen), as.integer(d^2), out = double(chnLen*d^2))$out, 
		nrow = chnLen, byrow = T)[,diagInd]
chn5 <- cbind(chn5, 
	matrix(.C("GSLmatLoad", 
	"chains/SgSout_aus_3_5.gbin", 
	as.integer(chnLen), as.integer(d^2), out = double(chnLen*d^2))$out, 
		nrow = chnLen, byrow = T)[,upInd])
chn5 <- cbind(chn5, 
	matrix(.C("GSLmatLoad", 
	"chains/SgAout_aus_3_5.gbin", 
	as.integer(chnLen), as.integer(d^2), out = double(chnLen*d^2))$out, 
		nrow = chnLen, byrow = T)[,upInd])
acMat   <- mcmcAcf("chn", 5)
acFrame <- data.frame(autocorrelation=c(acMat), 
	Sigma=rep(c(rep("SgE", length(diagInd)), 
	rep(c("SgS", "SgA"), each=length(upInd))), ncol(acMat)),
	lag=rep(0:33, each = nrow(acMat)))

gelRub("chn",nChn,chnLen)
@
\begin{center}
<<fig=TRUE,width=8.0,height=10.0>>=
ggSgE <- ggplot(data=subset(acFrame, Sigma=="SgE"), 
		aes(x=lag, y=autocorrelation)) +
	geom_point() + 
	theme_classic(base_size=18, base_family="myriad") +
	labs(title=expression(Sigma[e]))
ggSgS <- ggplot(data=subset(acFrame, Sigma=="SgS"), 
		aes(x=lag, y=autocorrelation)) +
	geom_point() + 
	theme_classic(base_size=18, base_family="myriad") + 
	labs(title=expression(Sigma[s]))
ggSgA <- ggplot(data=subset(acFrame, Sigma=="SgA"), 
		aes(x=lag, y=autocorrelation)) +
	geom_point() + 
	theme_classic(base_size=18, base_family="myriad") + 
	labs(title=expression(Sigma[a]))
showtext_auto()
grid.arrange(ggSgE, ggSgS, ggSgA, nrow = 3)

@
\end{center}
Plot variance chains. First, I get the diagonal index from the upper triangle index.
<<>>=
diagList <- list(1:d, c(3,5), c(6,8))
@

$\Sigma_e$:

\begin{center}
<<fig=TRUE,width=8.0,height=6.0,echo=FALSE>>=

par(mfrow = c(2,1), mar = c(0.2, 4, 0.5, 1), mgp = c(3,1,0))
for (trt in 1:d){
	plot( chn1[,diagList[[1]][trt]], type = "l", ylim = range(c(chn1[,diagList[[1]][trt]], chn2[,diagList[[1]][trt]], chn3[,diagList[[1]][trt]], chn4[,diagList[[1]][trt]], chn5[,diagList[[1]][trt]])), col = "orange", ylab = trtNam[trt], xlab = "", xaxt = "n", las = 1)
	lines(chn2[,diagList[[1]][trt]], col = "red")
	lines(chn3[,diagList[[1]][trt]], col = "blue")
	lines(chn4[,diagList[[1]][trt]], col = "green")
	lines(chn5[,diagList[[1]][trt]], col = "magenta")
}
@
\end{center}

Next, non-additive genetic variances.

\begin{center}
<<fig=TRUE,width=8.0,height=6.0,echo=FALSE>>=

par(mfrow = c(2,1), mar = c(0.2, 4, 0.5, 1), mgp = c(3,1,0))
for (trt in 1:d){
	plot( chn1[,diagList[[2]][trt]], type = "l", ylim = range(c(chn1[,diagList[[2]][trt]], chn2[,diagList[[2]][trt]], chn3[,diagList[[2]][trt]], chn4[,diagList[[2]][trt]], chn5[,diagList[[2]][trt]])), col = "orange", ylab = trtNam[trt], xlab = "", xaxt = "n", las = 1)
	lines(chn2[,diagList[[2]][trt]], col = "red")
	lines(chn3[,diagList[[2]][trt]], col = "blue")
	lines(chn4[,diagList[[2]][trt]], col = "green")
	lines(chn5[,diagList[[2]][trt]], col = "magenta")
}
@
\end{center}
Finally, $\boldsymbol{\Sigma}_a$ which can not be interpreted as the additive genetic covariance matrix in this case, where $\nu_g =3$ and the model is Student-$t$.

\begin{center}
<<fig=TRUE,width=8.0,height=6.0,echo=FALSE>>=

par(mfrow = c(2,1), mar = c(0.2, 4, 0.5, 1), mgp = c(3,1,0))
for (trt in 1:d){
	plot( chn1[,diagList[[3]][trt]], type = "l", ylim = range(c(chn1[,diagList[[3]][trt]], chn2[,diagList[[3]][trt]], chn3[,diagList[[3]][trt]], chn4[,diagList[[3]][trt]], chn5[,diagList[[3]][trt]])), col = "orange", ylab = trtNam[trt], xlab = "", xaxt = "n", las = 1)
	lines(chn2[,diagList[[3]][trt]], col = "red")
	lines(chn3[,diagList[[3]][trt]], col = "blue")
	lines(chn4[,diagList[[3]][trt]], col = "green")
	lines(chn5[,diagList[[3]][trt]], col = "magenta")
}
@
\end{center}
Everything looks good and I move on to the tropical \textit{japonica} covariances.
<<>>=
chn1 <- matrix(.C("GSLmatLoad", 
	"chains/SgEout_trj_3_1.gbin", 
	as.integer(chnLen), as.integer(d^2), out = double(chnLen*d^2))$out, 
		nrow = chnLen, byrow = T)[,diagInd]
chn1 <- cbind(chn1, 
	matrix(.C("GSLmatLoad", 
	"chains/SgSout_trj_3_1.gbin", 
	as.integer(chnLen), as.integer(d^2), out = double(chnLen*d^2))$out, 
		nrow = chnLen, byrow = T)[,upInd])
chn1 <- cbind(chn1, 
	matrix(.C("GSLmatLoad", 
	"chains/SgAout_trj_3_1.gbin", 
	as.integer(chnLen), as.integer(d^2), out = double(chnLen*d^2))$out, 
		nrow = chnLen, byrow = T)[,upInd])

chn2 <- matrix(.C("GSLmatLoad", 
	"chains/SgEout_trj_3_2.gbin", 
	as.integer(chnLen), as.integer(d^2), out = double(chnLen*d^2))$out, 
		nrow = chnLen, byrow = T)[,diagInd]
chn2 <- cbind(chn2, 
	matrix(.C("GSLmatLoad", 
	"chains/SgSout_trj_3_2.gbin", 
	as.integer(chnLen), as.integer(d^2), out = double(chnLen*d^2))$out, 
		nrow = chnLen, byrow = T)[,upInd])
chn2 <- cbind(chn2, 
	matrix(.C("GSLmatLoad", 
	"chains/SgAout_trj_3_2.gbin", 
	as.integer(chnLen), as.integer(d^2), out = double(chnLen*d^2))$out, 
		nrow = chnLen, byrow = T)[,upInd])

chn3 <- matrix(.C("GSLmatLoad", 
	"chains/SgEout_trj_3_3.gbin", 
	as.integer(chnLen), as.integer(d^2), out = double(chnLen*d^2))$out, 
		nrow = chnLen, byrow = T)[,diagInd]
chn3 <- cbind(chn3, 
	matrix(.C("GSLmatLoad", 
	"chains/SgSout_trj_3_3.gbin", 
	as.integer(chnLen), as.integer(d^2), out = double(chnLen*d^2))$out, 
		nrow = chnLen, byrow = T)[,upInd])
chn3 <- cbind(chn3, 
	matrix(.C("GSLmatLoad", 
	"chains/SgAout_trj_3_3.gbin", 
	as.integer(chnLen), as.integer(d^2), out = double(chnLen*d^2))$out, 
		nrow = chnLen, byrow = T)[,upInd])

chn4 <- matrix(.C("GSLmatLoad", 
	"chains/SgEout_trj_3_4.gbin", 
	as.integer(chnLen), as.integer(d^2), out = double(chnLen*d^2))$out, 
		nrow = chnLen, byrow = T)[,diagInd]
chn4 <- cbind(chn4, 
	matrix(.C("GSLmatLoad", 
	"chains/SgSout_trj_3_4.gbin", 
	as.integer(chnLen), as.integer(d^2), out = double(chnLen*d^2))$out, 
		nrow = chnLen, byrow = T)[,upInd])
chn4 <- cbind(chn4, 
	matrix(.C("GSLmatLoad", 
	"chains/SgAout_trj_3_4.gbin", 
	as.integer(chnLen), as.integer(d^2), out = double(chnLen*d^2))$out, 
		nrow = chnLen, byrow = T)[,upInd])

chn5 <- matrix(.C("GSLmatLoad", 
	"chains/SgEout_trj_3_5.gbin", 
	as.integer(chnLen), as.integer(d^2), out = double(chnLen*d^2))$out, 
		nrow = chnLen, byrow = T)[,diagInd]
chn5 <- cbind(chn5, 
	matrix(.C("GSLmatLoad", 
	"chains/SgSout_trj_3_5.gbin", 
	as.integer(chnLen), as.integer(d^2), out = double(chnLen*d^2))$out, 
		nrow = chnLen, byrow = T)[,upInd])
chn5 <- cbind(chn5, 
	matrix(.C("GSLmatLoad", 
	"chains/SgAout_trj_3_5.gbin", 
	as.integer(chnLen), as.integer(d^2), out = double(chnLen*d^2))$out, 
		nrow = chnLen, byrow = T)[,upInd])
acMat   <- mcmcAcf("chn", 5)
acFrame <- data.frame(autocorrelation=c(acMat), 
	Sigma=rep(c(rep("SgE", length(diagInd)), 
	rep(c("SgS", "SgA"), each=length(upInd))), ncol(acMat)),
	lag=rep(0:33, each = nrow(acMat)))

gelRub("chn",nChn,chnLen)
@
\begin{center}
<<fig=TRUE,width=8.0,height=10.0>>=
ggSgE <- ggplot(data=subset(acFrame, Sigma=="SgE"), 
		aes(x=lag, y=autocorrelation)) +
	geom_point() + 
	theme_classic(base_size=18, base_family="myriad") +
	labs(title=expression(Sigma[e]))
ggSgS <- ggplot(data=subset(acFrame, Sigma=="SgS"), 
		aes(x=lag, y=autocorrelation)) +
	geom_point() + 
	theme_classic(base_size=18, base_family="myriad") + 
	labs(title=expression(Sigma[s]))
ggSgA <- ggplot(data=subset(acFrame, Sigma=="SgA"), 
		aes(x=lag, y=autocorrelation)) +
	geom_point() + 
	theme_classic(base_size=18, base_family="myriad") + 
	labs(title=expression(Sigma[a]))
showtext_auto()
grid.arrange(ggSgE, ggSgS, ggSgA, nrow = 3)

@
\end{center}
Plot variance chains. First, I get the diagonal index from the upper triangle index.
<<>>=
diagList <- list(1:d, c(3,5), c(6,8))
@

$\Sigma_e$:

\begin{center}
<<fig=TRUE,width=8.0,height=6.0,echo=FALSE>>=

par(mfrow = c(2,1), mar = c(0.2, 4, 0.5, 1), mgp = c(3,1,0))
for (trt in 1:d){
	plot( chn1[,diagList[[1]][trt]], type = "l", ylim = range(c(chn1[,diagList[[1]][trt]], chn2[,diagList[[1]][trt]], chn3[,diagList[[1]][trt]], chn4[,diagList[[1]][trt]], chn5[,diagList[[1]][trt]])), col = "orange", ylab = trtNam[trt], xlab = "", xaxt = "n", las = 1)
	lines(chn2[,diagList[[1]][trt]], col = "red")
	lines(chn3[,diagList[[1]][trt]], col = "blue")
	lines(chn4[,diagList[[1]][trt]], col = "green")
	lines(chn5[,diagList[[1]][trt]], col = "magenta")
}
@
\end{center}

Next, non-additive genetic variances.

\begin{center}
<<fig=TRUE,width=8.0,height=6.0,echo=FALSE>>=

par(mfrow = c(2,1), mar = c(0.2, 4, 0.5, 1), mgp = c(3,1,0))
for (trt in 1:d){
	plot( chn1[,diagList[[2]][trt]], type = "l", ylim = range(c(chn1[,diagList[[2]][trt]], chn2[,diagList[[2]][trt]], chn3[,diagList[[2]][trt]], chn4[,diagList[[2]][trt]], chn5[,diagList[[2]][trt]])), col = "orange", ylab = trtNam[trt], xlab = "", xaxt = "n", las = 1)
	lines(chn2[,diagList[[2]][trt]], col = "red")
	lines(chn3[,diagList[[2]][trt]], col = "blue")
	lines(chn4[,diagList[[2]][trt]], col = "green")
	lines(chn5[,diagList[[2]][trt]], col = "magenta")
}
@
\end{center}
Finally, $\boldsymbol{\Sigma}_a$ which can not be interpreted as the additive genetic covariance matrix in this case, where $\nu_g =3$ and the model is Student-$t$.

\begin{center}
<<fig=TRUE,width=8.0,height=6.0,echo=FALSE>>=

par(mfrow = c(2,1), mar = c(0.2, 4, 0.5, 1), mgp = c(3,1,0))
for (trt in 1:d){
	plot( chn1[,diagList[[3]][trt]], type = "l", ylim = range(c(chn1[,diagList[[3]][trt]], chn2[,diagList[[3]][trt]], chn3[,diagList[[3]][trt]], chn4[,diagList[[3]][trt]], chn5[,diagList[[3]][trt]])), col = "orange", ylab = trtNam[trt], xlab = "", xaxt = "n", las = 1)
	lines(chn2[,diagList[[3]][trt]], col = "red")
	lines(chn3[,diagList[[3]][trt]], col = "blue")
	lines(chn4[,diagList[[3]][trt]], col = "green")
	lines(chn5[,diagList[[3]][trt]], col = "magenta")
}
@
\end{center}
Finally, admixed tropical \textit{japonica} covariances.
<<>>=
chn1 <- matrix(.C("GSLmatLoad", 
	"chains/SgEout_trja_3_1.gbin", 
	as.integer(chnLen), as.integer(d^2), out = double(chnLen*d^2))$out, 
		nrow = chnLen, byrow = T)[,diagInd]
chn1 <- cbind(chn1, 
	matrix(.C("GSLmatLoad", 
	"chains/SgSout_trja_3_1.gbin", 
	as.integer(chnLen), as.integer(d^2), out = double(chnLen*d^2))$out, 
		nrow = chnLen, byrow = T)[,upInd])
chn1 <- cbind(chn1, 
	matrix(.C("GSLmatLoad", 
	"chains/SgAout_trja_3_1.gbin", 
	as.integer(chnLen), as.integer(d^2), out = double(chnLen*d^2))$out, 
		nrow = chnLen, byrow = T)[,upInd])

chn2 <- matrix(.C("GSLmatLoad", 
	"chains/SgEout_trja_3_2.gbin", 
	as.integer(chnLen), as.integer(d^2), out = double(chnLen*d^2))$out, 
		nrow = chnLen, byrow = T)[,diagInd]
chn2 <- cbind(chn2, 
	matrix(.C("GSLmatLoad", 
	"chains/SgSout_trja_3_2.gbin", 
	as.integer(chnLen), as.integer(d^2), out = double(chnLen*d^2))$out, 
		nrow = chnLen, byrow = T)[,upInd])
chn2 <- cbind(chn2, 
	matrix(.C("GSLmatLoad", 
	"chains/SgAout_trja_3_2.gbin", 
	as.integer(chnLen), as.integer(d^2), out = double(chnLen*d^2))$out, 
		nrow = chnLen, byrow = T)[,upInd])

chn3 <- matrix(.C("GSLmatLoad", 
	"chains/SgEout_trja_3_3.gbin", 
	as.integer(chnLen), as.integer(d^2), out = double(chnLen*d^2))$out, 
		nrow = chnLen, byrow = T)[,diagInd]
chn3 <- cbind(chn3, 
	matrix(.C("GSLmatLoad", 
	"chains/SgSout_trja_3_3.gbin", 
	as.integer(chnLen), as.integer(d^2), out = double(chnLen*d^2))$out, 
		nrow = chnLen, byrow = T)[,upInd])
chn3 <- cbind(chn3, 
	matrix(.C("GSLmatLoad", 
	"chains/SgAout_trja_3_3.gbin", 
	as.integer(chnLen), as.integer(d^2), out = double(chnLen*d^2))$out, 
		nrow = chnLen, byrow = T)[,upInd])

chn4 <- matrix(.C("GSLmatLoad", 
	"chains/SgEout_trja_3_4.gbin", 
	as.integer(chnLen), as.integer(d^2), out = double(chnLen*d^2))$out, 
		nrow = chnLen, byrow = T)[,diagInd]
chn4 <- cbind(chn4, 
	matrix(.C("GSLmatLoad", 
	"chains/SgSout_trja_3_4.gbin", 
	as.integer(chnLen), as.integer(d^2), out = double(chnLen*d^2))$out, 
		nrow = chnLen, byrow = T)[,upInd])
chn4 <- cbind(chn4, 
	matrix(.C("GSLmatLoad", 
	"chains/SgAout_trja_3_4.gbin", 
	as.integer(chnLen), as.integer(d^2), out = double(chnLen*d^2))$out, 
		nrow = chnLen, byrow = T)[,upInd])

chn5 <- matrix(.C("GSLmatLoad", 
	"chains/SgEout_trja_3_5.gbin", 
	as.integer(chnLen), as.integer(d^2), out = double(chnLen*d^2))$out, 
		nrow = chnLen, byrow = T)[,diagInd]
chn5 <- cbind(chn5, 
	matrix(.C("GSLmatLoad", 
	"chains/SgSout_trja_3_5.gbin", 
	as.integer(chnLen), as.integer(d^2), out = double(chnLen*d^2))$out, 
		nrow = chnLen, byrow = T)[,upInd])
chn5 <- cbind(chn5, 
	matrix(.C("GSLmatLoad", 
	"chains/SgAout_trja_3_5.gbin", 
	as.integer(chnLen), as.integer(d^2), out = double(chnLen*d^2))$out, 
		nrow = chnLen, byrow = T)[,upInd])
acMat   <- mcmcAcf("chn", 5)
acFrame <- data.frame(autocorrelation=c(acMat), 
	Sigma=rep(c(rep("SgE", length(diagInd)), 
	rep(c("SgS", "SgA"), each=length(upInd))), ncol(acMat)),
	lag=rep(0:33, each = nrow(acMat)))

gelRub("chn",nChn,chnLen)
@
\begin{center}
<<fig=TRUE,width=8.0,height=10.0>>=
ggSgE <- ggplot(data=subset(acFrame, Sigma=="SgE"), 
		aes(x=lag, y=autocorrelation)) +
	geom_point() + 
	theme_classic(base_size=18, base_family="myriad") +
	labs(title=expression(Sigma[e]))
ggSgS <- ggplot(data=subset(acFrame, Sigma=="SgS"), 
		aes(x=lag, y=autocorrelation)) +
	geom_point() + 
	theme_classic(base_size=18, base_family="myriad") + 
	labs(title=expression(Sigma[s]))
ggSgA <- ggplot(data=subset(acFrame, Sigma=="SgA"), 
		aes(x=lag, y=autocorrelation)) +
	geom_point() + 
	theme_classic(base_size=18, base_family="myriad") + 
	labs(title=expression(Sigma[a]))
showtext_auto()
grid.arrange(ggSgE, ggSgS, ggSgA, nrow = 3)

@
\end{center}
Plot variance chains. First, I get the diagonal index from the upper triangle index.
<<>>=
diagList <- list(1:d, c(3,5), c(6,8))
@

$\Sigma_e$:

\begin{center}
<<fig=TRUE,width=8.0,height=6.0,echo=FALSE>>=

par(mfrow = c(2,1), mar = c(0.2, 4, 0.5, 1), mgp = c(3,1,0))
for (trt in 1:d){
	plot( chn1[,diagList[[1]][trt]], type = "l", ylim = range(c(chn1[,diagList[[1]][trt]], chn2[,diagList[[1]][trt]], chn3[,diagList[[1]][trt]], chn4[,diagList[[1]][trt]], chn5[,diagList[[1]][trt]])), col = "orange", ylab = trtNam[trt], xlab = "", xaxt = "n", las = 1)
	lines(chn2[,diagList[[1]][trt]], col = "red")
	lines(chn3[,diagList[[1]][trt]], col = "blue")
	lines(chn4[,diagList[[1]][trt]], col = "green")
	lines(chn5[,diagList[[1]][trt]], col = "magenta")
}
@
\end{center}

Next, non-additive genetic variances.

\begin{center}
<<fig=TRUE,width=8.0,height=6.0,echo=FALSE>>=

par(mfrow = c(2,1), mar = c(0.2, 4, 0.5, 1), mgp = c(3,1,0))
for (trt in 1:d){
	plot( chn1[,diagList[[2]][trt]], type = "l", ylim = range(c(chn1[,diagList[[2]][trt]], chn2[,diagList[[2]][trt]], chn3[,diagList[[2]][trt]], chn4[,diagList[[2]][trt]], chn5[,diagList[[2]][trt]])), col = "orange", ylab = trtNam[trt], xlab = "", xaxt = "n", las = 1)
	lines(chn2[,diagList[[2]][trt]], col = "red")
	lines(chn3[,diagList[[2]][trt]], col = "blue")
	lines(chn4[,diagList[[2]][trt]], col = "green")
	lines(chn5[,diagList[[2]][trt]], col = "magenta")
}
@
\end{center}
Finally, $\boldsymbol{\Sigma}_a$ which can not be interpreted as the additive genetic covariance matrix in this case, where $\nu_g =3$ and the model is Student-$t$.

\begin{center}
<<fig=TRUE,width=8.0,height=6.0,echo=FALSE>>=

par(mfrow = c(2,1), mar = c(0.2, 4, 0.5, 1), mgp = c(3,1,0))
for (trt in 1:d){
	plot( chn1[,diagList[[3]][trt]], type = "l", ylim = range(c(chn1[,diagList[[3]][trt]], chn2[,diagList[[3]][trt]], chn3[,diagList[[3]][trt]], chn4[,diagList[[3]][trt]], chn5[,diagList[[3]][trt]])), col = "orange", ylab = trtNam[trt], xlab = "", xaxt = "n", las = 1)
	lines(chn2[,diagList[[3]][trt]], col = "red")
	lines(chn3[,diagList[[3]][trt]], col = "blue")
	lines(chn4[,diagList[[3]][trt]], col = "green")
	lines(chn5[,diagList[[3]][trt]], col = "magenta")
}
@
\end{center}


\end{document}
%<<>>=
```
