## Supplemental methods and scripts for "Low additive genetic variation in a trait under selection in domesticated rice": mcmcResultsFAM.Rnw

```
\documentclass[fleqn,11pt]{article}
\usepackage{amsmath}
\usepackage{amssymb}
\usepackage{sectsty}
\usepackage{amsbsy}
\usepackage{fancyhdr}
\usepackage{color}
\usepackage{url}
\usepackage[indention=2em,labelsep=period,labelfont=bf]{caption}
\usepackage[normalsize,nooneline,sf,bf,FIGTOPCAP,tight]{subfigure}
\usepackage[noae]{Sweave} % noae required to sanitize single quotes
\usepackage{fancyvrb}

\textwidth = 6.5 in
\textheight = 8.5 in
\oddsidemargin = 0.0 in
\evensidemargin = 0.0 in
\topmargin = 0.0 in
%\voffset = - 0.4 in
\headheight = 15 pt
\headsep = 0.1 in
\parskip = 0.0 in
\parindent = 4 ex

\definecolor{grey}{rgb}{0.4,0.4,0.4}

\newcommand{\bcdot}{\boldsymbol{\cdot}}

\DefineVerbatimEnvironment{Sinput}{Verbatim}{tabsize=4}
\DefineVerbatimEnvironment{Scode}{Verbatim}{tabsize=4}
\DefineVerbatimEnvironment{Soutput}{Verbatim}{tabsize=4,formatcom=\color{grey},fontsize=\small}

\pagestyle{fancy}
\fancyhf{}
\rhead{Page \thepage}
\lhead{Old Phenotype Results}
\chead{Tony Greenberg}

\title{Markov chain results}
\author{Tony Greenberg}

\begin{document}

\maketitle
\thispagestyle{empty}
<<echo=false>>=
#rm(list = ls())
dyn.load("GSLRIO.so")
options(width=90)
library(ggplot2)
library(gridExtra)
library(showtext)
library(compiler)
font_add("myriad", regular="MYRIADPRO-SEMIBOLD.OTF",
	bold="MYRIADPRO-BOLD.OTF", italic="MYRIADPRO-SEMIBOLDIT.OTF")
sessionInfo()
@
In this document I summarize the results of modeling the old Al tolerance data using Bayesian hierarchical modeling.

\section{Model}
I used the following hierarchical model to account for experiments from a number of years.

\begin{align*}
	\boldsymbol{y}_{i\bcdot} &\sim t_{\nu_{e},d\left(\boldsymbol{\mu}^{acc}_{j[i]\bcdot} + \boldsymbol{x}_{i\bcdot}^{\textsc{exp}}\boldsymbol{B}^{\textsc{exp}}; \boldsymbol{\Sigma}_e \right)\\
\boldsymbol{\mu}^{acc}_{j\bcdot} &\sim \textrm{N}_d\left(\boldsymbol{\mu} + \boldsymbol{u}_{j\bcdot}\boldsymbol{\Gamma}; \boldsymbol{\Sigma}_s \right)\\
	\boldsymbol{\gamma}_{j\bcdot} &\sim t_{\nu_{g}, d}\left(\boldsymbol{0}_d; \boldsymbol{\Sigma}_a \right)\\
\boldsymbol{\Sigma}_e^{-1} &\sim \textrm{W}_{d, \nu_0}\left(\left[\sum_i\left(\boldsymbol{y}_{i\bcdot} - \boldsymbol{\mu}^{acc}_{j[i]\bcdot} - \boldsymbol{x}_{i\bcdot}^{\textsc{exp}}\boldsymbol{B}^{\textsc{exp}} \right)\left(\boldsymbol{y}_{i\bcdot} - \boldsymbol{\mu}^{acc}_{j[i]\bcdot} - \boldsymbol{x}_{i\bcdot}^{\textsc{exp}}\boldsymbol{B}^{\textsc{exp}} \right)^{\textsc{t}} \right]^{-1} \right)\\
\boldsymbol{\Sigma}_s^{-1} &\sim \textrm{W}_{d, \nu_0}\left(\left[\sum_j\left(\boldsymbol{\mu}^{acc}_{j\bcdot} - \boldsymbol{\mu} - \boldsymbol{u}_{j\bcdot}\boldsymbol{\Gamma} \right)\left(\boldsymbol{\mu}^{acc}_{j\bcdot} - \boldsymbol{\mu} - \boldsymbol{u}_{j\bcdot}\boldsymbol{\Gamma} \right)^{\textsc{t}} \right]^{-1} \right)\\
\boldsymbol{\Sigma}_a^{-1} &\sim \textrm{W}_{d, \nu_0}\left(\left[ \boldsymbol{\Gamma}^{\textsc{t}}\boldsymbol{\Gamma} \right]^{-1}\right),
\end{align*}
where bold lower-case symbols refer to row vectors and bold upper-case symbols are matrices. All vectors are of length $d$ and all matrices have $d$ columns, where $d$ is the number of traits ($d = 2$ in our case). $\boldsymbol{X}^{\textsc{exp}}$ is the matrix of experiment IDs and $\boldsymbol{B}^{\textsc{exp}}$ contains experimental effect coefficients. These are essentially years, but see the \verb|phenoDataPrep| documents for details. The $\boldsymbol{U}$ matrix contains principal components of the genetic relationship matrix, scaled by square roots of their eigenvalues. The $\boldsymbol{U\Gamma}$ matrix is the matrix of GEBVs (marker breeding values). The degrees of freedom parameter $\nu_g$ was set to 3, the smallest value that results in a distribution with all moments defined.  Notation of the $i\bcdot$ type refers to rows in matrices. Subscripts of the $j[i]$ type refer to a row $j$ in a upper level (in the model hierarchy) that corresponds to a replicate row $i$. Note also that the residual covariance matrix $\boldsymbol{\Sigma}_e^{-1}$ is diagonal, since the experiment and control phenotypes were measured on different plants and thus the residual correlation  is undefined. Location parameters without priors were modeled with very high-variance Gaussian priors.

I define functions that calculate summaries of Markov chains.

<<>>=
pmode <- cmpfun(function(vec){
	dst <- density(vec, adjust = 2)
	mxi <- which(dst$y == max(dst$y))
	if(length(mxi) > 1){
		warning("More than one mode in call to pmode(); picking randomly")
		mxi <- sample(mxi, 1)
	}
	dst$x[mxi]
})

HPDint <- cmpfun(function(vec, prob = 0.95){
	nsamp <- length(vec)
	if (nsamp <= 2) stop("vector must have length > 2")

	vals <- sort(vec)
	gap  <- max(1, min(nsamp - 1, round(nsamp * prob)))
	init <- 1:(nsamp - gap)
	mInd <- which.min(vals[init + gap] - vals[init])
	res  <- c(vals[mInd], vals[mInd + gap])
	names(res) <- c("lower", "upper")
	return(res)
})
@
Next, I write a function that is similar to \verb|quantile()|, but outputs the mode, lower and upper limits of the 95\% and 50\% HPD. The \verb|outr| parameter is the probability for the outer margins, default is 95\%.
<<>>=
quantileLike <- cmpfun(function(vec, outr = 0.95){
	if(outr <= 0.5) stop("Outer margin has to be >= 50%")
	md    <- pmode(vec)
	hpd95 <- HPDint(vec, outr)
	hpd50 <- HPDint(vec, 0.5)
	res   <- c(hpd95[1], hpd50[1], md, hpd50[2], hpd95[2])

	outNm <- paste(c("lower", "upper"), outr*100, sep = "")
	names(res) <- c(outNm[1], "lower50", "mode", "upper50", outNm[2])
	return(res)
})
@

\section{Experiment effect}
I start by checking the experiment effect. Start with the \textit{aus} results.

I start with total accession means.
<<>>=
trtNam <- c("Control", "Treated")
d      <- 2
Nexp   <- 6
expDim <- d*Nexp
nChn   <- 5
chnLen <- 2000

chn <- NULL
addSamp <- cmpfun(function(i, vrNam, popNam, ncl){
	inFlNam <- paste("chains/", vrNam, "out_", popNam, "_3_", i, ".gbin", sep = "")
	chn <<- rbind(chn,
				matrix(.C("GSLmatLoad",
					inFlNam, as.integer(chnLen), as.integer(ncl),
					out = double(chnLen*ncl))$out,
				nrow = chnLen, byrow = T)
		)
	return(NULL)
})

trash <- sapply(1:nChn, addSamp, "EXP", "aus", expDim)

expID               <- scan(file = "expIDs.txt", what=character())
expMnAUS            <- as.data.frame(t(apply(chn, 2, quantileLike)))
expMnAUS$trait      <- rep(trtNam, times = Nexp)
expMnAUS$experiment <- rep(expID, each=d)
@

\begin{center}
<<fig=TRUE,width=8.0,height=10.0>>=
ggTrt1 <- ggplot(data=subset(expMnAUS, trait==trtNam[1]), aes(x=experiment,y=mode)) +
	geom_segment(aes(x=experiment, y=lower95, xend=experiment, yend=upper95),
		color="grey80", size=0.75) +
	geom_segment(aes(x=experiment, y=lower50, xend=experiment, yend=upper50),
		color="grey50", size=1) +
	geom_point() +
	theme_classic(base_size=18, base_family="myriad") +
	theme(axis.title.x=element_blank(), axis.ticks.x=element_blank()) +
	labs(y=trtNam[1])
ggTrt2 <- ggplot(data=subset(expMnAUS, trait==trtNam[2]), aes(x=experiment,y=mode)) +
	geom_segment(aes(x=experiment, y=lower95, xend=experiment, yend=upper95),
		color="grey80", size=0.75) +
	geom_segment(aes(x=experiment, y=lower50, xend=experiment, yend=upper50),
		color="grey50", size=1) +
	geom_point() +
	theme_classic(base_size=18, base_family="myriad") +
	theme(axis.ticks.x=element_blank()) +
	labs(y=trtNam[2])
showtext_auto()
grid.arrange(ggTrt1, ggTrt2, nrow = 2)
@
\end{center}
The estimates look reasonable compared to plots of raw data. I look at the tropical \textit{japonica} results.
<<>>=
chn <- NULL
trash <- sapply(1:nChn, addSamp, "EXP", "trj", expDim)

expMnTRJ            <- as.data.frame(t(apply(chn, 2, quantileLike)))
expMnTRJ$trait      <- rep(trtNam, times = Nexp)
expMnTRJ$experiment <- rep(expID, each=d)
@

\begin{center}
<<fig=TRUE,width=8.0,height=10.0>>=
ggTrt1 <- ggplot(data=subset(expMnTRJ, trait==trtNam[1]), aes(x=experiment,y=mode)) +
	geom_segment(aes(x=experiment, y=lower95, xend=experiment, yend=upper95),
		color="grey80", size=0.75) +
	geom_segment(aes(x=experiment, y=lower50, xend=experiment, yend=upper50),
		color="grey50", size=1) +
	geom_point() +
	theme_classic(base_size=18, base_family="myriad") +
	theme(axis.title.x=element_blank(), axis.ticks.x=element_blank()) +
	labs(y=trtNam[1])
ggTrt2 <- ggplot(data=subset(expMnTRJ, trait==trtNam[2]), aes(x=experiment,y=mode)) +
	geom_segment(aes(x=experiment, y=lower95, xend=experiment, yend=upper95),
		color="grey80", size=0.75) +
	geom_segment(aes(x=experiment, y=lower50, xend=experiment, yend=upper50),
		color="grey50", size=1) +
	geom_point() +
	theme_classic(base_size=18, base_family="myriad") +
	theme(axis.ticks.x=element_blank()) +
	labs(y=trtNam[2])
showtext_auto()
grid.arrange(ggTrt1, ggTrt2, nrow = 2)
@
\end{center}
Results are very similar to \textit{aus}. Finally, I look at the admixed tropical \textit{japonica} results.
<<>>=
chn <- NULL
trash <- sapply(1:nChn, addSamp, "EXP", "trja", expDim)

expMnTRJA            <- as.data.frame(t(apply(chn, 2, quantileLike)))
expMnTRJA$trait      <- rep(trtNam, times = Nexp)
expMnTRJA$experiment <- rep(expID, each=d)
@

\begin{center}
<<fig=TRUE,width=8.0,height=10.0>>=
ggTrt1 <- ggplot(data=subset(expMnTRJA, trait==trtNam[1]), aes(x=experiment,y=mode)) +
	geom_segment(aes(x=experiment, y=lower95, xend=experiment, yend=upper95),
		color="grey80", size=0.75) +
	geom_segment(aes(x=experiment, y=lower50, xend=experiment, yend=upper50),
		color="grey50", size=1) +
	geom_point() +
	theme_classic(base_size=18, base_family="myriad") +
	theme(axis.title.x=element_blank(), axis.ticks.x=element_blank()) +
	labs(y=trtNam[1])
ggTrt2 <- ggplot(data=subset(expMnTRJA, trait==trtNam[2]), aes(x=experiment,y=mode)) +
	geom_segment(aes(x=experiment, y=lower95, xend=experiment, yend=upper95),
		color="grey80", size=0.75) +
	geom_segment(aes(x=experiment, y=lower50, xend=experiment, yend=upper50),
		color="grey50", size=1) +
	geom_point() +
	theme_classic(base_size=18, base_family="myriad") +
	theme(axis.ticks.x=element_blank()) +
	labs(y=trtNam[2])
showtext_auto()
grid.arrange(ggTrt1, ggTrt2, nrow = 2)
@
\end{center}

\section{Accession means}

Next, I look at accession means. Their variance reflects broad-sense heritability. Starting with \textit{aus}.

<<>>=
Nacc   <- 55
accDim <- d*Nacc

chn <- NULL
trash <- sapply(1:nChn, addSamp, "LN", "aus", accDim)

accMnAUS       <- as.data.frame(t(apply(chn, 2, quantileLike)))
accMnAUS$trait <- rep(trtNam, times = Nacc)
accMnAUSs      <- accMnAUS[order(accMnAUS[,6], accMnAUS[,3]),]
@
I calculate the relative root growth and check if all values are positive.
<<>>=
accNamAUS     <- scan(file="lineIDsAUS.txt", what=character())
colnames(chn) <- paste(rep(trtNam, Nacc), rep(accNamAUS, each=d), sep=".")
chnRRG        <- chn[,grep(trtNam[2], colnames(chn))]/chn[,grep(trtNam[1], colnames(chn))]
sum(chnRRG<=0)
@
They are. I calculate the $-\log_{10}(RRG)$ and save the modes of accession effect estimates for further analyses.
<<>>=
rrgModeAUS    <- apply(-log10(chnRRG), 2, pmode)
accModeTabAUS <- cbind(matrix(round(accMnAUS$mode, 4), ncol=d, byrow=T),
	round(rrgModeAUS, 4))
cat(c("NSFTVID\tControl\t160uM_Al\tneglgRRG",
		apply(cbind(accNamAUS, accModeTabAUS), 1, paste, collapse="\t")),
	file="accModeAUS.tsv",sep="\n"
	)
trash <- .C("GSLmatSave", "phenoData/accMdAUS.gbin",
	as.double(accModeTabAUS), nrow(accModeTabAUS), ncol(accModeTabAUS))
@
I plot sorted accession means.
\begin{center}
<<results=tex>>=
pdfFlNam <- "lineMeansFAMaus.pdf"
showtext_auto()
ggplot(data=accMnAUSs, aes(x=1:nrow(accMnAUSs),y=mode)) +
	geom_segment(aes(x=1:nrow(accMnAUSs), y=lower95, xend=1:nrow(accMnAUSs), yend=upper95),
		color="grey80", size=0.75) +
	geom_segment(aes(x=1:nrow(accMnAUSs), y=lower50, xend=1:nrow(accMnAUSs), yend=upper50),
		color="grey50", size=1) +
	geom_point() +
	facet_wrap(~trait, scales="free", nrow=2) +
	theme_classic(base_size=18, base_family="myriad") +
	theme(axis.title.x=element_blank(), axis.text.x=element_blank(),
		strip.background=element_rect(fill="grey95", linetype="blank"),
		axis.ticks.x=element_blank()) +
	labs(y="root length")
ggsave(pdfFlNam, width=8, height=10, units="in", device="pdf", useDingbats=F)
cat("\\includegraphics{", pdfFlNam, "}\n\n", sep="")
@
\end{center}
Seems like there are some particularly resistant accessions, which is expected when treatment has the desired stress effect. Now look at the tropical \textit{japonica}.

<<>>=
Nacc   <- 92
accDim <- d*Nacc

chn <- NULL
trash <- sapply(1:nChn, addSamp, "LN", "trj", accDim)

accMnTRJ       <- as.data.frame(t(apply(chn, 2, quantileLike)))
accMnTRJ$trait <- rep(trtNam, times = Nacc)
accMnTRJs      <- accMnTRJ[order(accMnTRJ[,6], accMnTRJ[,3]),]
@
I calculate the relative root growth and check if all values are positive.
<<>>=
accNamTRJ     <- scan(file="lineIDsTRJ.txt", what=character())
colnames(chn) <- paste(rep(trtNam, Nacc), rep(accNamTRJ, each=d), sep=".")
chnRRG        <- chn[,grep(trtNam[2], colnames(chn))]/chn[,grep(trtNam[1], colnames(chn))]
sum(chnRRG<=0)
@
There are negative values, due to correction for the experiment effect. I subtract the smallest value to make everything positive.
<<>>=
chnRRG <- (chn[,grep(trtNam[2], colnames(chn))]-min(chn)+1e-4)/
	(chn[,grep(trtNam[1], colnames(chn))]-min(chn)+1e-4)
sum(chnRRG<=0)
@

I calculate the $-\log_{10}(RRG)$ and save the modes of accession effect estimates for further analyses.
<<>>=
rrgModeTRJ    <- apply(-log10(chnRRG), 2, pmode)
accModeTabTRJ <- cbind(matrix(round(accMnTRJ$mode, 4), ncol=d, byrow=T),
	round(rrgModeTRJ, 4))
cat(c("NSFTVID\tControl\t160uM_Al\tneglgRRG",
		apply(cbind(accNamTRJ, accModeTabTRJ), 1, paste, collapse="\t")),
	file="accModeTRJ.tsv",sep="\n"
	)
trash <- .C("GSLmatSave", "phenoData/accMdTRJ.gbin",
	as.double(accModeTabTRJ), nrow(accModeTabTRJ), ncol(accModeTabTRJ))
@
I plot sorted accession means.
\begin{center}
<<results=tex>>=
pdfFlNam <- "lineMeansFAMtrj.pdf"
showtext_auto()
ggplot(data=accMnTRJs, aes(x=1:nrow(accMnTRJs),y=mode)) +
	geom_segment(aes(x=1:nrow(accMnTRJs), y=lower95, xend=1:nrow(accMnTRJs), yend=upper95),
		color="grey80", size=0.75) +
	geom_segment(aes(x=1:nrow(accMnTRJs), y=lower50, xend=1:nrow(accMnTRJs), yend=upper50),
		color="grey50", size=1) +
	geom_point() +
	facet_wrap(~trait, scales="free", nrow=2) +
	theme_classic(base_size=18, base_family="myriad") +
	theme(axis.title.x=element_blank(), axis.text.x=element_blank(),
		strip.background=element_rect(fill="grey95", linetype="blank"),
		axis.ticks.x=element_blank()) +
	labs(y="root length")
ggsave(pdfFlNam, width=8, height=10, units="in", device="pdf", useDingbats=F)
cat("\\includegraphics{", pdfFlNam, "}\n\n", sep="")
@
\end{center}
The distributions are very similar. Finally, look at the admixed tropical \textit{japonica}.

<<>>=
Nacc   <- 95
accDim <- d*Nacc

chn <- NULL
trash <- sapply(1:nChn, addSamp, "LN", "trja", accDim)

accNamTRJA      <- scan(file="lineIDsTRJA.txt", what=character())
accNamADM       <- scan(file="lineIDsADM.txt", what=character())
accMnTRJA       <- as.data.frame(t(apply(chn, 2, quantileLike)))
accMnTRJA$trait <- rep(trtNam, times = Nacc)
accMnTRJA$admix <- rep(ifelse(accNamTRJA %in% accNamADM, "admixed", "pure"), each=d)
accMnTRJAs      <- accMnTRJA[order(accMnTRJA[,6], accMnTRJA[,3]),]
@
Save the modes to a text file.
<<>>=
accModeTabTRJA <- matrix(round(accMnTRJA$mode, 4), ncol=d, byrow=T)
cat(c("NSFTVID\tControl\t160uM_Al",
		apply(cbind(accNamTRJA, accModeTabTRJA), 1, paste, collapse="\t")),
	file="accModeTRJA.tsv",sep="\n"
	)

@
I plot sorted accession means.
\begin{center}
<<results=tex>>=
pdfFlNam <- "lineMeansFAMtrja.pdf"
showtext_auto()
ggplot(data=accMnTRJAs, aes(x=1:nrow(accMnTRJAs),y=mode)) +
	geom_segment(aes(x=1:nrow(accMnTRJAs), y=lower95, xend=1:nrow(accMnTRJAs), yend=upper95),
		color="grey80", size=0.75) +
	geom_segment(aes(x=1:nrow(accMnTRJAs), y=lower50,
				xend=1:nrow(accMnTRJAs), yend=upper50, color=admix),size=1.5) +
	geom_point(size=1.5, color="grey60") +
	facet_wrap(~trait, scales="free", nrow=2) +
	theme_classic(base_size=18, base_family="myriad") +
	theme(axis.title.x=element_blank(), axis.text.x=element_blank(),
		strip.background=element_rect(fill="grey95", linetype="blank"),
		axis.ticks.x=element_blank(), legend.title=element_blank()) +
	labs(y="root length")
ggsave(pdfFlNam, width=8, height=10, units="in", device="pdf", useDingbats=F)
cat("\\includegraphics{", pdfFlNam, "}\n\n", sep="")
@
\end{center}
The admixed lines have smaller roots even in control, and even smaller in treatment (compared to the average tropical \textit{japonica}). I test this more formally by estimating mean rank of the admixed accessions, compared to the overall mean (which is $(1+N_{acc})/2$).
<<>>=
admInd <- accNamTRJA %in% accNamADM
mnRank <- cmpfun(function(vec, ind){
	colMeans(apply(matrix(vec, ncol=2, byrow=T), 2, rank)[ind,])
})
rankChn         <- t(apply(chn, 1, mnRank, admInd))
rankMnTRJ       <- as.data.frame(t(apply(rankChn, 2, quantileLike)))
rankMnTRJ$trait <- trtNam
rankMnTRJ
@
Plot the ranks.
\begin{center}
<<results=tex>>=
pdfFlNam <- "rankAdmLineMeans.pdf"
showtext_auto()
ggplot(data=rankMnTRJ, aes(x=1:d,y=mode)) +
	geom_segment(aes(x=1:d, y=lower95, xend=1:d, yend=upper95),
		color="grey80", size=1.25) +
	geom_segment(aes(x=1:d, y=lower50, xend=1:d, yend=upper50),
		color="grey50", size=2) +
	geom_point() +
	geom_hline(yintercept=(mean(1:Nacc)), color="grey50") +
	theme_classic(base_size=18, base_family="myriad") +
	theme(axis.title.x=element_blank(), axis.text.x=element_blank(),
		strip.background=element_rect(fill="grey95", linetype="blank"),
		axis.ticks.x=element_blank()) +
	labs(y="rank") + xlim(0.5, d+0.5) +
	annotate("text", label="Control", x=1, y=min(rankMnTRJ$lower95)-2) +
	annotate("text", label="Al treated", x=2, y=min(rankMnTRJ$lower95)-2)
ggsave(pdfFlNam, width=8, height=8, units="in", device="pdf", useDingbats=F)
cat("\\includegraphics{", pdfFlNam, "}\n\n", sep="")
@
\end{center}

\section{GEBV}
I next examine the distributions of genome (or SNP) estimated breeding values (GEBV).
<<>>=
Nacc   <- 55
accDim <- d*Nacc

chn        <- NULL
trash      <- sapply(1:nChn, addSamp, "BV", "aus", accDim)
gebvChnAUS <- chn

gebvMnAUS       <- as.data.frame(t(apply(chn, 2, quantileLike)))
gebvMnAUS$trait <- rep(trtNam, times = Nacc)
gebvMnAUSs      <- gebvMnAUS[order(gebvMnAUS[,6], gebvMnAUS[,3]),]
@
I plot sorted GEBVs.
\begin{center}
<<results=tex>>=
pdfFlNam <- "gebvFAMaus.pdf"
showtext_auto()
ggplot(data=gebvMnAUSs, aes(x=1:nrow(gebvMnAUSs),y=mode)) +
	geom_segment(aes(x=1:nrow(gebvMnAUSs), y=lower95, xend=1:nrow(gebvMnAUSs), yend=upper95),
		color="grey80", size=0.75) +
	geom_segment(aes(x=1:nrow(gebvMnAUSs), y=lower50, xend=1:nrow(gebvMnAUSs), yend=upper50),
		color="grey50", size=1) +
	geom_point() +
	facet_wrap(~trait, scales="free", nrow=2) +
	theme_classic(base_size=18, base_family="myriad") +
	theme(axis.title.x=element_blank(), axis.text.x=element_blank(),
		strip.background=element_rect(fill="grey95", linetype="blank"),
		axis.ticks.x=element_blank()) +
	labs(y="root length")
ggsave(pdfFlNam, width=8, height=10, units="in", device="pdf", useDingbats=F)
cat("\\includegraphics{", pdfFlNam, "}\n\n", sep="")
@
\end{center}
Again, seems like some breeding values stand out as high in the treated group. Now look at the tropical \textit{japonica}.

<<>>=
Nacc   <- 92
accDim <- d*Nacc

chn <- NULL
trash <- sapply(1:nChn, addSamp, "BV", "trj", accDim)
gebvChnTRJ <- chn

gebvMnTRJ       <- as.data.frame(t(apply(chn, 2, quantileLike)))
gebvMnTRJ$trait <- rep(trtNam, times = Nacc)
gebvMnTRJs      <- gebvMnTRJ[order(gebvMnTRJ[,6], gebvMnTRJ[,3]),]
@
I plot sorted accession means.
\begin{center}
<<results=tex>>=
pdfFlNam <- "gebvFAMtrj.pdf"
showtext_auto()
ggplot(data=gebvMnTRJs, aes(x=1:nrow(gebvMnTRJs),y=mode)) +
	geom_segment(aes(x=1:nrow(gebvMnTRJs), y=lower95, xend=1:nrow(gebvMnTRJs), yend=upper95),
		color="grey80", size=0.75) +
	geom_segment(aes(x=1:nrow(gebvMnTRJs), y=lower50, xend=1:nrow(gebvMnTRJs), yend=upper50),
		color="grey50", size=1) +
	geom_point() +
	facet_wrap(~trait, scales="free", nrow=2) +
	theme_classic(base_size=18, base_family="myriad") +
	theme(axis.title.x=element_blank(), axis.text.x=element_blank(),
		strip.background=element_rect(fill="grey95", linetype="blank"),
		axis.ticks.x=element_blank()) +
	labs(y="root length")
ggsave(pdfFlNam, width=8, height=10, units="in", device="pdf", useDingbats=F)
cat("\\includegraphics{", pdfFlNam, "}\n\n", sep="")
@
\end{center}

Finally, look at the admixed tropical \textit{japonica}.

<<>>=
Nacc   <- 95
accDim <- d*Nacc

chn <- NULL
trash <- sapply(1:nChn, addSamp, "BV", "trja", accDim)
gebvChnTRJA <- chn

gebvMnTRJA       <- as.data.frame(t(apply(chn, 2, quantileLike)))
gebvMnTRJA$trait <- rep(trtNam, times = Nacc)
gebvMnTRJA$admix <- rep(ifelse(accNamTRJA %in% accNamADM, "admixed", "pure"), each=d)
gebvMnTRJAs      <- gebvMnTRJA[order(gebvMnTRJA[,6], gebvMnTRJA[,3]),]
@
I plot sorted accession means.
\begin{center}
<<results=tex>>=
pdfFlNam <- "gebvFAMtrja.pdf"
showtext_auto()
ggplot(data=gebvMnTRJAs, aes(x=1:nrow(gebvMnTRJAs),y=mode,color=admix)) +
	geom_segment(aes(x=1:nrow(gebvMnTRJAs), y=lower95, xend=1:nrow(gebvMnTRJAs), yend=upper95),
		color="grey80", size=0.75) +
	geom_segment(aes(x=1:nrow(gebvMnTRJAs), y=lower50, xend=1:nrow(gebvMnTRJAs), yend=upper50),
		color="grey50", size=1) +
	geom_point() +
	facet_wrap(~trait, scales="free", nrow=2) +
	theme_classic(base_size=18, base_family="myriad") +
	theme(axis.title.x=element_blank(), axis.text.x=element_blank(),
		strip.background=element_rect(fill="grey95", linetype="blank"),
		axis.ticks.x=element_blank()) +
	labs(y=trtNam[1])
ggsave(pdfFlNam, width=8, height=10, units="in", device="pdf", useDingbats=F)
cat("\\includegraphics{", pdfFlNam, "}\n\n", sep="")
@
\end{center}
The effect is similar to the accession means, but seems more pronounced. Plot the ranks as before.
\begin{center}
<<results=tex>>=
pdfFlNam        <- "rankAdmGEBV.pdf"
rankChn         <- t(apply(chn, 1, mnRank, admInd))
rankBVTRJ       <- as.data.frame(t(apply(rankChn, 2, quantileLike)))
rankBVTRJ$trait <- trtNam

showtext_auto()
ggplot(data=rankBVTRJ, aes(x=1:d,y=mode)) +
	geom_segment(aes(x=1:d, y=lower95, xend=1:d, yend=upper95),
		color="grey80", size=1.25) +
	geom_segment(aes(x=1:d, y=lower50, xend=1:d, yend=upper50),
		color="grey50", size=2) +
	geom_point() +
	geom_hline(yintercept=(mean(1:Nacc)), color="grey50") +
	theme_classic(base_size=18, base_family="myriad") +
	theme(axis.title.x=element_blank(), axis.text.x=element_blank(),
		strip.background=element_rect(fill="grey95", linetype="blank"),
		axis.ticks.x=element_blank()) +
	labs(y="rank") + xlim(0.5, d+0.5) +
	annotate("text", label="Control", x=1, y=min(rankMnTRJ$lower95)-2) +
	annotate("text", label="Al treated", x=2, y=min(rankMnTRJ$lower95)-2)
ggsave(pdfFlNam, width=8, height=8, units="in", device="pdf", useDingbats=F)
cat("\\includegraphics{", pdfFlNam, "}\n\n", sep="")
@
\end{center}
Numerical values:
<<>>=
rankBVTRJ
@

\section{Heritabilities}
Finally, I directly compare heritabilities between control and treated, as well as between populations. The background genetic and residual variances are taken from the respective covariance matrices. However, because we used a Student-$t$ genetic model, the additive variance is estimated from the GEBV values rather $\boldsymbol{\Sigma}_a$.

I first define the functions I need.
<<>>=
makeVar <- cmpfun(function(vec){
	apply(matrix(vec, ncol=d, byrow=T), 2, var)
})
get.hsq <- cmpfun(function(vec){
	vec[1:d]/rowSums(matrix(vec, nrow = d))
})
@
The function \verb|get.hsq| calculates the marker heritability, which is a kind of narrow-sense heritability. It is
\begin{displaymath}
h^2 = \dfrac{\sigma^2_{\textsc{gebv}}}{\sigma^2_{\textsc{gebv}} + \sigma^2_s + \sigma^2_e}
\end{displaymath}
for each treatment.

I read in the covariance matrices and extract variances (diagonal elements).

<<>>=
diagInd <- diag(matrix(1:(d^2), ncol = d, byrow = T))

chn1 <- matrix(.C("GSLmatLoad",
	"chains/SgEout_aus_3_1.gbin",
	as.integer(chnLen), as.integer(d^2), out = double(chnLen*d^2))$out,
		nrow = chnLen, byrow = T)[,diagInd]
chn1 <- cbind(chn1,
	matrix(.C("GSLmatLoad",
	"chains/SgSout_aus_3_1.gbin",
	as.integer(chnLen), as.integer(d^2), out = double(chnLen*d^2))$out,
		nrow = chnLen, byrow = T)[,diagInd])

chn2 <- matrix(.C("GSLmatLoad",
	"chains/SgEout_aus_3_2.gbin",
	as.integer(chnLen), as.integer(d^2), out = double(chnLen*d^2))$out,
		nrow = chnLen, byrow = T)[,diagInd]
chn2 <- cbind(chn2,
	matrix(.C("GSLmatLoad",
	"chains/SgSout_aus_3_2.gbin",
	as.integer(chnLen), as.integer(d^2), out = double(chnLen*d^2))$out,
		nrow = chnLen, byrow = T)[,diagInd])

chn3 <- matrix(.C("GSLmatLoad",
	"chains/SgEout_aus_3_3.gbin",
	as.integer(chnLen), as.integer(d^2), out = double(chnLen*d^2))$out,
		nrow = chnLen, byrow = T)[,diagInd]
chn3 <- cbind(chn3,
	matrix(.C("GSLmatLoad",
	"chains/SgSout_aus_3_3.gbin",
	as.integer(chnLen), as.integer(d^2), out = double(chnLen*d^2))$out,
		nrow = chnLen, byrow = T)[,diagInd])

chn4 <- matrix(.C("GSLmatLoad",
	"chains/SgEout_aus_3_4.gbin",
	as.integer(chnLen), as.integer(d^2), out = double(chnLen*d^2))$out,
		nrow = chnLen, byrow = T)[,diagInd]
chn4 <- cbind(chn4,
	matrix(.C("GSLmatLoad",
	"chains/SgSout_aus_3_4.gbin",
	as.integer(chnLen), as.integer(d^2), out = double(chnLen*d^2))$out,
		nrow = chnLen, byrow = T)[,diagInd])

chn5 <- matrix(.C("GSLmatLoad",
	"chains/SgEout_aus_3_5.gbin",
	as.integer(chnLen), as.integer(d^2), out = double(chnLen*d^2))$out,
		nrow = chnLen, byrow = T)[,diagInd]
chn5 <- cbind(chn5,
	matrix(.C("GSLmatLoad",
	"chains/SgSout_aus_3_5.gbin",
	as.integer(chnLen), as.integer(d^2), out = double(chnLen*d^2))$out,
		nrow = chnLen, byrow = T)[,diagInd])

sigChnAUS <- rbind(chn1, chn2, chn3, chn4, chn5)
sigChnAUS <- cbind(t(apply(gebvChnAUS, 1, makeVar)), sigChnAUS)
chnhsqAUS <- t(apply(sigChnAUS, 1, get.hsq))

chn1 <- matrix(.C("GSLmatLoad",
	"chains/SgEout_trj_3_1.gbin",
	as.integer(chnLen), as.integer(d^2), out = double(chnLen*d^2))$out,
		nrow = chnLen, byrow = T)[,diagInd]
chn1 <- cbind(chn1,
	matrix(.C("GSLmatLoad",
	"chains/SgSout_trj_3_1.gbin",
	as.integer(chnLen), as.integer(d^2), out = double(chnLen*d^2))$out,
		nrow = chnLen, byrow = T)[,diagInd])

chn2 <- matrix(.C("GSLmatLoad",
	"chains/SgEout_trj_3_2.gbin",
	as.integer(chnLen), as.integer(d^2), out = double(chnLen*d^2))$out,
		nrow = chnLen, byrow = T)[,diagInd]
chn2 <- cbind(chn2,
	matrix(.C("GSLmatLoad",
	"chains/SgSout_trj_3_2.gbin",
	as.integer(chnLen), as.integer(d^2), out = double(chnLen*d^2))$out,
		nrow = chnLen, byrow = T)[,diagInd])

chn3 <- matrix(.C("GSLmatLoad",
	"chains/SgEout_trj_3_3.gbin",
	as.integer(chnLen), as.integer(d^2), out = double(chnLen*d^2))$out,
		nrow = chnLen, byrow = T)[,diagInd]
chn3 <- cbind(chn3,
	matrix(.C("GSLmatLoad",
	"chains/SgSout_trj_3_3.gbin",
	as.integer(chnLen), as.integer(d^2), out = double(chnLen*d^2))$out,
		nrow = chnLen, byrow = T)[,diagInd])

chn4 <- matrix(.C("GSLmatLoad",
	"chains/SgEout_trj_3_4.gbin",
	as.integer(chnLen), as.integer(d^2), out = double(chnLen*d^2))$out,
		nrow = chnLen, byrow = T)[,diagInd]
chn4 <- cbind(chn4,
	matrix(.C("GSLmatLoad",
	"chains/SgSout_trj_3_4.gbin",
	as.integer(chnLen), as.integer(d^2), out = double(chnLen*d^2))$out,
		nrow = chnLen, byrow = T)[,diagInd])

chn5 <- matrix(.C("GSLmatLoad",
	"chains/SgEout_trj_3_5.gbin",
	as.integer(chnLen), as.integer(d^2), out = double(chnLen*d^2))$out,
		nrow = chnLen, byrow = T)[,diagInd]
chn5 <- cbind(chn5,
	matrix(.C("GSLmatLoad",
	"chains/SgSout_trj_3_5.gbin",
	as.integer(chnLen), as.integer(d^2), out = double(chnLen*d^2))$out,
		nrow = chnLen, byrow = T)[,diagInd])

sigChnTRJ <- rbind(chn1, chn2, chn3, chn4, chn5)
sigChnTRJ <- cbind(t(apply(gebvChnTRJ, 1, makeVar)), sigChnTRJ)
chnhsqTRJ <- t(apply(sigChnTRJ, 1, get.hsq))

chn1 <- matrix(.C("GSLmatLoad",
	"chains/SgEout_trja_3_1.gbin",
	as.integer(chnLen), as.integer(d^2), out = double(chnLen*d^2))$out,
		nrow = chnLen, byrow = T)[,diagInd]
chn1 <- cbind(chn1,
	matrix(.C("GSLmatLoad",
	"chains/SgSout_trja_3_1.gbin",
	as.integer(chnLen), as.integer(d^2), out = double(chnLen*d^2))$out,
		nrow = chnLen, byrow = T)[,diagInd])

chn2 <- matrix(.C("GSLmatLoad",
	"chains/SgEout_trja_3_2.gbin",
	as.integer(chnLen), as.integer(d^2), out = double(chnLen*d^2))$out,
		nrow = chnLen, byrow = T)[,diagInd]
chn2 <- cbind(chn2,
	matrix(.C("GSLmatLoad",
	"chains/SgSout_trja_3_2.gbin",
	as.integer(chnLen), as.integer(d^2), out = double(chnLen*d^2))$out,
		nrow = chnLen, byrow = T)[,diagInd])

chn3 <- matrix(.C("GSLmatLoad",
	"chains/SgEout_trja_3_3.gbin",
	as.integer(chnLen), as.integer(d^2), out = double(chnLen*d^2))$out,
		nrow = chnLen, byrow = T)[,diagInd]
chn3 <- cbind(chn3,
	matrix(.C("GSLmatLoad",
	"chains/SgSout_trja_3_3.gbin",
	as.integer(chnLen), as.integer(d^2), out = double(chnLen*d^2))$out,
		nrow = chnLen, byrow = T)[,diagInd])

chn4 <- matrix(.C("GSLmatLoad",
	"chains/SgEout_trja_3_4.gbin",
	as.integer(chnLen), as.integer(d^2), out = double(chnLen*d^2))$out,
		nrow = chnLen, byrow = T)[,diagInd]
chn4 <- cbind(chn4,
	matrix(.C("GSLmatLoad",
	"chains/SgSout_trja_3_4.gbin",
	as.integer(chnLen), as.integer(d^2), out = double(chnLen*d^2))$out,
		nrow = chnLen, byrow = T)[,diagInd])

chn5 <- matrix(.C("GSLmatLoad",
	"chains/SgEout_trja_3_5.gbin",
	as.integer(chnLen), as.integer(d^2), out = double(chnLen*d^2))$out,
		nrow = chnLen, byrow = T)[,diagInd]
chn5 <- cbind(chn5,
	matrix(.C("GSLmatLoad",
	"chains/SgSout_trja_3_5.gbin",
	as.integer(chnLen), as.integer(d^2), out = double(chnLen*d^2))$out,
		nrow = chnLen, byrow = T)[,diagInd])

sigChnTRJA <- rbind(chn1, chn2, chn3, chn4, chn5)
sigChnTRJA <- cbind(t(apply(gebvChnTRJA, 1, makeVar)), sigChnTRJA)
chnhsqTRJA <- t(apply(sigChnTRJA, 1, get.hsq))

Npop   <- 3
hsqHPD <- as.data.frame(rbind(t(apply(chnhsqAUS, 2, quantileLike)),
	t(apply(chnhsqTRJ, 2, quantileLike)),
	t(apply(chnhsqTRJA, 2, quantileLike))))
hsqHPD$treatment  <- rep(trtNam, Npop)
hsqHPD$population <- rep(c("AUS", "TRJ", "TRJA"), each=d)

@
Plot the results.
\begin{center}
<<results=tex>>=
pdfFlNam <- "hSqFAMall.pdf"
showtext_auto()
ggplot(data=hsqHPD, aes(x=c(1,2,4,5,7,8), y=mode, color=population)) +
	geom_segment(aes(x=c(1,2,4,5,7,8), y=lower95, xend=c(1,2,4,5,7,8),
		yend=upper95, color=population),
		size=1.2) +
	geom_segment(aes(x=c(1,2,4,5,7,8), y=lower50, xend=c(1,2,4,5,7,8),
		yend=upper50, color=population),
		size=1.75) +
	geom_point(size=2) +
	theme_classic(base_size=18, base_family="myriad") +
	scale_x_continuous(breaks=c(1,2,4,5,7,8), labels=rep(trtNam, Npop)) +
	ylim(c(0,1)) +
	labs(y=expression(italic(h)^2), x="treatment")
ggsave(pdfFlNam, width=8, height=8, units="in", device="pdf", useDingbats=F)
cat("\\includegraphics{", pdfFlNam, "}\n\n", sep="")
@
\end{center}
Repeat, including only the pure accessions.

\begin{center}
<<results=tex>>=
pdfFlNam <- "hSqFAMpure.pdf"
showtext_auto()
ggplot(data=subset(hsqHPD, population != "TRJA"), aes(x=c(1,2,4,5), y=mode, color=population)) +
	geom_segment(aes(x=c(1,2,4,5), y=lower95, xend=c(1,2,4,5),
		yend=upper95, color=population),
		size=1.2) +
	geom_segment(aes(x=c(1,2,4,5), y=lower50, xend=c(1,2,4,5),
		yend=upper50, color=population),
		size=1.75) +
	geom_point(size=2) +
	theme_classic(base_size=18, base_family="myriad") +
	scale_x_continuous(breaks=c(1,2,4,5), labels=rep(trtNam, Npop-1)) +
	ylim(c(0,1)) +
	labs(y=expression(italic(h)^2), x="treatment")
ggsave(pdfFlNam, width=8, height=8, units="in", device="pdf", useDingbats=F)
cat("\\includegraphics{", pdfFlNam, "}\n\n", sep="")
@
\end{center}
Now I plot broad-sense (among-accession) heritability:
\begin{displaymath}
	H^2 = \dfrac{\sigma^2_{\textsc{gebv}} + \sigma^2_s}{\sigma^2_{\textsc{gebv}} + \sigma^2_s + \sigma^2_e}
\end{displaymath}
I am also interested in the contribution of the non-additive effects alone, excluding the marker influence:
\begin{displaymath}
	FVE_{s} = \dfrac{\sigma^2_s}{\sigma^2_{\textsc{gebv}} + \sigma^2_s + \sigma^2_e}
\end{displaymath}

<<>>=
get.Hsq <- cmpfun(function(vec){
	(vec[1:d] + vec[(2*d+1):(3*d)])/rowSums(matrix(vec, nrow = d))
})
get.fve <- cmpfun(function(vec){
	(vec[(2*d+1):(3*d)])/rowSums(matrix(vec, nrow = d))
})

chnHsqAUS  <- t(apply(sigChnAUS,  1, get.Hsq))
chnHsqTRJ  <- t(apply(sigChnTRJ,  1, get.Hsq))
chnHsqTRJA <- t(apply(sigChnTRJA, 1, get.Hsq))

chnFveAUS  <- t(apply(sigChnAUS,  1, get.fve))
chnFveTRJ  <- t(apply(sigChnTRJ,  1, get.fve))
chnFveTRJA <- t(apply(sigChnTRJA, 1, get.fve))

HsqHPD <- as.data.frame(rbind(t(apply(chnHsqAUS, 2, quantileLike)),
	t(apply(chnHsqTRJ, 2, quantileLike)),
	t(apply(chnHsqTRJA, 2, quantileLike))))
HsqHPD$treatment <- rep(trtNam, Npop)
HsqHPD$population <- rep(c("AUS", "TRJ", "TRJA"), each=d)

fveHPD <- as.data.frame(rbind(t(apply(chnhsqAUS, 2, quantileLike)),
	t(apply(chnhsqTRJ, 2, quantileLike)),
	t(apply(chnFveAUS, 2, quantileLike)),
	t(apply(chnFveTRJ, 2, quantileLike))))
fveHPD$treatment  <- rep(trtNam, 2*(Npop-1))
fveHPD$population <- rep(rep(c("AUS", "TRJ"), each=2), d)
fveHPD$kind       <- factor(rep(c("Marker", "Background"), each=d*(Npop-1)),
							levels=c("Marker", "Background"))
@

Plot the results.
\begin{center}
<<results=tex>>=
pdfFlNam <- "HsqFAMall.pdf"
showtext_auto()
ggplot(data=HsqHPD, aes(x=c(1,2,4,5,7,8), y=mode, color=population)) +
	geom_segment(aes(x=c(1,2,4,5,7,8), y=lower95, xend=c(1,2,4,5,7,8),
		yend=upper95, color=population),
		size=1.2) +
	geom_segment(aes(x=c(1,2,4,5,7,8), y=lower50, xend=c(1,2,4,5,7,8),
		yend=upper50, color=population),
		size=1.75) +
	geom_point(size=2) +
	theme_classic(base_size=18, base_family="myriad") +
	scale_x_continuous(breaks=c(1,2,4,5,7,8), labels=rep(trtNam, Npop)) +
	ylim(c(0,1)) +
	labs(y=expression(italic(H)^2), x="treatment")
ggsave(pdfFlNam, width=8, height=8, units="in", device="pdf", useDingbats=F)
cat("\\includegraphics{", pdfFlNam, "}\n\n", sep="")
@
\end{center}
Repeat, including only the pure accessions.

\begin{center}
<<results=tex>>=
pdfFlNam <- "HsqFAMpure.pdf"
showtext_auto()
ggplot(data=subset(HsqHPD, population != "TRJA"), aes(x=c(1,2,4,5), y=mode, color=population)) +
	geom_segment(aes(x=c(1,2,4,5), y=lower95, xend=c(1,2,4,5),
		yend=upper95, color=population),
		size=1.2) +
	geom_segment(aes(x=c(1,2,4,5), y=lower50, xend=c(1,2,4,5),
		yend=upper50, color=population),
		size=1.75) +
	geom_point(size=2) +
	theme_classic(base_size=18, base_family="myriad") +
	scale_x_continuous(breaks=c(1,2,4,5), labels=rep(trtNam, Npop-1)) +
	ylim(c(0,1)) +
	labs(y=expression(italic(H)^2), x="treatment")
ggsave(pdfFlNam, width=8, height=8, units="in", device="pdf", useDingbats=F)
cat("\\includegraphics{", pdfFlNam, "}\n\n", sep="")
@
\end{center}
I plot the fractions of variance explained side by side.
\begin{center}
<<results=tex>>=
pdfFlNam <- "fveFAMpure.pdf"
showtext_auto()
ggplot(data=fveHPD, aes(x=rep(1:4,2), y=mode, color=population)) +
	geom_segment(aes(x=rep(1:4,2), y=lower95, xend=rep(1:4,2),
		yend=upper95, color=population),
		size=1.2) +
	geom_segment(aes(x=rep(1:4,2), y=lower50, xend=rep(1:4,2),
		yend=upper50, color=population),
		size=1.75) +
	geom_point(size=2) +
	facet_wrap(~kind, scales="free_x", ncol=2) +
	theme_classic(base_size=18, base_family="myriad") +
	scale_x_continuous(breaks=rep(1:4,2), labels=rep(trtNam, d*(Npop-1))) +
	theme(axis.text.x=element_text(angle=45, hjust=1.0, vjust=1.0),
		strip.background=element_rect(fill="grey95", linetype="blank"),
		panel.spacing.x=unit(4.0, "lines")) +
	ylim(c(0,1)) +
	labs(y="fraction of variance", x="treatment")
ggsave(pdfFlNam, width=10, height=8, units="in", device="pdf", useDingbats=F)
cat("\\includegraphics{", pdfFlNam, "}\n\n", sep="")
@
\end{center}

Finally, put the plots together for direct comparison.
\begin{center}
<<fig=TRUE,width=12.0,height=6.0>>=
ggGEBV <- ggplot(data=hsqHPD, aes(x=c(1,2,4,5,7,8), y=mode, color=population)) +
	geom_segment(aes(x=c(1,2,4,5,7,8), y=lower95, xend=c(1,2,4,5,7,8),
		yend=upper95, color=population),
		size=1.2) +
	geom_segment(aes(x=c(1,2,4,5,7,8), y=lower50, xend=c(1,2,4,5,7,8),
		yend=upper50, color=population),
		size=1.75) +
	geom_point(size=2) +
	theme_classic(base_size=18, base_family="myriad") +
	scale_x_continuous(breaks=c(1,2,4,5,7,8), labels=rep(trtNam, Npop)) +
	theme(axis.text.x=element_text(angle=45, hjust=1.0, vjust=1.0)) +
	ylim(c(0,1)) +
	labs(y=expression(italic(h)^2), x="treatment")

ggHsq <- ggplot(data=HsqHPD, aes(x=c(1,2,4,5,7,8), y=mode, color=population)) +
	geom_segment(aes(x=c(1,2,4,5,7,8), y=lower95, xend=c(1,2,4,5,7,8),
		yend=upper95, color=population),
		size=1.2) +
	geom_segment(aes(x=c(1,2,4,5,7,8), y=lower50, xend=c(1,2,4,5,7,8),
		yend=upper50, color=population),
		size=1.75) +
	geom_point(size=2) +
	theme_classic(base_size=18, base_family="myriad") +
	scale_x_continuous(breaks=c(1,2,4,5,7,8), labels=rep(trtNam, Npop)) +
	theme(axis.text.x=element_text(angle=45, hjust=1.0, vjust=1.0)) +
	ylim(c(0,1)) +
	labs(y=expression(italic(H)^2), x="treatment")

showtext_auto()
grid.arrange(ggGEBV, ggHsq, ncol = 2)

@
\end{center}


\end{document}
```
