## Supplemental methods and scripts for "Low additive genetic variation in a trait under selection in domesticated rice": README.md

This directory contains the `Sweave()` R scripts that prepare the raw data for analysis, run convergence diagnostics on the MCMC output, and summarize and plot the results. They all require the GSLRIO.so file to run. Instructions to generate this library are in the _CppSource_ directory _README.md_ file. The raw phenotypic data files necessary to run these scripts are included in this directory. The genotype data must be generated from the master HDRA set available at the [Rice Diversity](ricediversity.org/data/index.cfm) web site.

The scripts use some Unix commands and assume a Unix-like file system, so will probably not work on Windows.

Once the data preparation scripts are run, the files they generate can be used to run the analysis programs in the _CppSource_ directory. The MCMC processing scripts rely on the results of these analyses.
