## Supplemental methods and scripts for "Low additive genetic variation in a trait under selection in domesticated rice": RFmixPlot.Rnw

```
\documentclass[fleqn,11pt]{article}
\usepackage{amsmath}
\usepackage{amssymb}
\usepackage{sectsty}
\usepackage{amsbsy}
\usepackage{fancyhdr}
\usepackage{color}
\usepackage{url}
\usepackage[indention=2em,labelsep=period,labelfont=bf]{caption}
\usepackage[normalsize,nooneline,sf,bf,FIGTOPCAP,tight]{subfigure}
\usepackage[noae]{Sweave} % noae required to sanitize single quotes
\usepackage{fancyvrb}

\textwidth = 6.5 in
\textheight = 8.5 in
\oddsidemargin = 0.0 in
\evensidemargin = 0.0 in
\topmargin = 0.0 in
\headheight = 15 pt
\headsep = 0.1 in
\parskip = 0.0 in
\parindent = 4 ex

\definecolor{grey}{rgb}{0.4,0.4,0.4}

\newcommand{\bcdot}{\boldsymbol{\cdot}}

\DefineVerbatimEnvironment{Sinput}{Verbatim}{tabsize=4}
\DefineVerbatimEnvironment{Scode}{Verbatim}{tabsize=4}
\DefineVerbatimEnvironment{Soutput}{Verbatim}{tabsize=4,formatcom=\color{grey},fontsize=\small}

\pagestyle{fancy}
\fancyhf{}
\rhead{Page \thepage}
\lhead{RFMix plots}
\chead{Tony Greenberg}

\title{Plotting RFMix results for the admix accessions}
\author{Tony Greenberg}

\begin{document}

\maketitle
\thispagestyle{empty}
<<echo=false>>=
options(width=90)
library(ggplot2)
library(showtext)
library(compiler)
font_add("myriad", regular="MYRIADPRO-SEMIBOLD.OTF",
	bold="MYRIADPRO-BOLD.OTF", italic="MYRIADPRO-SEMIBOLDIT.OTF")
sessionInfo()
@
In this document I plot the results of the RFMix run on the admixed tropical \textit{japonica} accessions. The output of RFMix is the ID of the population at each SNP position. I start by reading the RFMix output (I modified it by adding chromosome ID as the first column ahead of time).

<<>>=
nAcc       <- 6
rfmRes     <- matrix(scan(file="chrRFres.txt", what=integer()), ncol=nAcc+1, byrow=T)
chrPos     <- matrix(scan(file="chrPos.tsv", what=integer()), ncol=2, byrow=T)
cumChrLen  <- scan(file="riceCumChromLen.txt", what=integer())
chrLabPos  <- cumChrLen
cumChrLen  <- c(0, cumChrLen[-length(cumChrLen)])
chrLabPos  <- (cumChrLen + chrLabPos)/2
chrPos[,2] <- chrPos[,2] + cumChrLen[chrPos[,1]]
@
I next read the root length modes for the admixed accessions. I want to plot them in order of root length increase (under Al stress).
<<>>=
admAccID <- scan(file="admixedAccessions.txt", what=character())
rootMode <- read.table(file="accModeTRJA.tsv", header=T, sep="\t")
nsf2hdra <- matrix(scan(file="NSFTV2HDRA.tsv", what=character()), ncol=2, byrow=T)
hdraID   <- nsf2hdra[,1]
names(hdraID)   <- nsf2hdra[,2]
rootMode$HDRAID <- hdraID[as.character(rootMode$NSFTVID)]
admMode         <- rootMode[rootMode$HDRAID %in% admAccID,"X160uM_Al"]
names(admMode)  <- rootMode[rootMode$HDRAID %in% admAccID,"HDRAID"]
admMode         <- admMode[admAccID]
@
Now calculate lengths of population ID runs along chromosomes of each admixed accession.
<<>>=
runEnd  <- apply(rfmRes[,-1], 2, function(vec){cumsum(rle(vec)$lengths)})
runID   <- apply(rfmRes[,-1], 2, function(vec){rle(vec)$values})
accLen  <- sapply(runEnd, length)
cpEnd   <- sapply(runEnd, function(vec){chrPos[vec,2]})
cpStart <- sapply(cpEnd, function(vec){c(0, vec[-length(vec)])})
@
Build a data frame for plotting.
<<>>=
rfmFrame <- data.frame(start=unlist(cpStart),
						end=unlist(cpEnd),
						accession=rep(order(admMode), times=accLen),
						population=c("AUS", "IND", "TRJ")[unlist(runID)])
@
Plot the results.
\begin{center}
<<results=tex>>=
pdfFlNam  <- "RFMixRes.pdf"
showtext_auto()
ggplot(data=rfmFrame) +
	geom_segment(aes(x=start, y=accession,
			xend=end, yend=accession, color=population),size=4) +
	scale_color_manual(values=c("AUS"="red2", "IND"="green2", "TRJ"="grey85")) +
	geom_vline(xintercept=cumChrLen[-1], color="grey50") +
	theme_classic(base_size=18, base_family="myriad") +
	theme(axis.ticks=element_blank(), axis.line=element_blank(),
		axis.title.y=element_blank(), axis.text.y=element_blank()) +
	scale_x_continuous(name="chromosomes", breaks=chrLabPos, labels=1:12)
ggsave(pdfFlNam, width=8, height=2, units="in", device="pdf", useDingbats=F)
cat("\\includegraphics{", pdfFlNam, "}\n\n", sep="")
@
\end{center}

\end{document}
```
