## Supplemental methods and scripts for "Low additive genetic variation in a trait under selection in domesticated rice": README.md

This directory contains the C++ software, R (Sweave) scripts, PDFs generated using the R scripts, and the raw phenotypic data used in the study. Please refer to the _README_ files in included in the separate directories for further information. 

The R scripts require the `GSLRIO.so` library to read and save the binary files used by the C++ programs. The source code and compilation instructions for it are in the _software_ directory. The PDFs show the results we obtained running the R scripts on the data. Raw phenotype data needed to reproduce our analyses are included in that directory. The genotype data are available for download from the (Rice Diversity website)[ricediversity.org/data/index.cfm].

The intermediate files (such as accession means) have to be re-generated by running the scripts. The data preparation scripts must be run first, followed by analyses with the C++ programs in the _CppSource_ directory, then the MCMC examination scripts (that run convergence diagnostics), and finally MCMC results and plotting scripts to summarize and plot results.
